# Depth-Resolved Microbial Community Structure and Phosphorus Deficiency Across a 34-cm Mat–Sediment Core in Poza Roja 2, Cuatrociénegas

**DOI:** 10.64898/2026.10.04.756594

**Authors:** Antonio González-Sánchez, Georgina Ibarra-Arzave, Atzintli Paniagua-Vargas, Felipe García-Oliva, Gabriela Olmedo-Álvarez

## Abstract

Stratified benthic deposits offer crucial modern analogs for Precambrian biogeochemical dynamics, yet whether the vertical organization of deep sedimentary columns reflects depositional history, present-day geochemical conditions, or both remains poorly understood, particularly under extreme oligotrophy. Here, we present a depth-resolved exploratory survey of a continuous 34-cm sediment core partitioned into 11 discrete horizons (L1 to L11) from Poza Roja 2, an alkaline, sub-hypersaline pond in the Cuatro Ciénegas Basin, Mexico. Integrating physical stratigraphy, organic carbon and nutrient stoichiometry, and 16S rRNA gene metabarcoding, we evaluated down-core microbial community structure alongside acute phosphorus scarcity. Molar C:Po and N:Po ratios exceeded canonical Redfield proportions (106:16:1) in every layer, by up to 48.5-fold and 87.8-fold, respectively, both peaking in layer L4. Unconstrained ordination (PCoA) and hierarchical clustering grouped the layers into three compositional zones: upper (L2–L4), intermediate (L5–L6), and basal (L8–L11), with L1 and L7 occupying transitional positions. Archaea accounted for 31.0% of prokaryotic reads, ranging from 8.3% (L5) to 43.7% (L9), and were dominated by SM1-50 (Thermoplasmatota) and Halobacteriota. The phototrophic heliobacterium *Heliorestis* peaked in the intermediate zone (47.6% of bacterial reads in L6). Constrained Redundancy Analysis models lacked statistical significance (P ≥ 0.12), highlighting the exploratory nature of a single core (n = 11). This pilot characterization provides a baseline for understanding deep biogenic archives under severe nutrient limitation.

## Introduction

Stratified benthic ecosystems, such as microbial mats and shallow sediment columns, represent prime modern analogs for Precambrian environments, offering key insights into the biogeochemical conditions and microbial community structures that prevailed on the early Earth (Des Marais 2003; Nisbet and Sleep 2001). These ecosystems typically exhibit steep, millimetric physicochemical gradients (spanning light availability, redox potential, and nutrient stoichiometry) that drive sharp vertical zonation of specialized microbial functional guilds (Bolhuis *et al*. 2014; Stolz 2000). Depth-resolved characterization of these gradients, however, has been largely confined to the uppermost few centimeters of actively growing mats and associated surface sediments (Harris *et al*. 2013; Ladakis *et al*. 2006), leaving the vertical organization of thicker, decimeter-scale stratigraphic records comparatively unexplored. While upper photic zones are frequently dominated by oxygenic phototrophs and aerobic heterotrophs, deeper horizons transition into microoxic and anoxic regimes sustained by anaerobic respirations, fermentation, and methanogenesis (Canfield and Des Marais 1993; van Gemerden 1993). How these complex vertical communities self-organize across physical and chemical transitions remains a central question in microbial ecology and geobiology.

The Cuatro Ciénegas Basin (CCB), an oasis located in the Chihuahuan Desert of Coahuila, Mexico, is internationally recognized as a living archive of ancient microbial diversity and evolutionary processes (Souza *et al*. 2006, 2012, 2018). Due to its extreme oligotrophy, most notably an acute, chronic deficiency of phosphorus (P), and its unique geological history as an enclosed hydrological system fed by deep groundwater, CCB harbors exceptional levels of microbial endemism and novel phylogenetic lineages (Elser *et al*. 2005; Souza *et al*. 2008). Living organisms depend on more than 20 largely non-substitutable elements, and growth can in principle be constrained by whichever element is scarcest relative to metabolic demand (Hessen *et al*. 2013). Previous surveys across the basin’s aquatic systems have revealed distinctive microbial communities associated with extreme stoichiometric imbalances, where molar C:P and N:P ratios far exceed canonical Redfield proportions (106:16:1) (Bonilla-Rosso *et al*. 2012; Garcia-Ulloa *et al*. 2022; Peimbert *et al*. 2012; Redfield 1958).

To cope with this severe, systemic phosphorus scarcity, microbial communities in CCB have evolved specialized genomic, physiological, and microevolutionary strategies. Genomic sequencing of local isolates revealed that endemic bacteria can substitute membrane phospholipids with non-phosphorus sulfolipids via genes closely related to cyanobacterial *sqd1* and *sqdX*, apparently acquired by horizontal gene transfer; because membrane phospholipids account for approximately 30% of total cellular phosphate in most organisms, this substitution can release a substantial fraction of cellular phosphorus (Alcaraz *et al*. 2008). Furthermore, microbial communities across soil and sediment habitats in CCB actively scavenge alternative phosphorus sources across multiple oxidation states, heavily relying on extracellular DNA recycling via phosphodiesterases as well as the enzymatic degradation of phosphonates and phosphites (Tapia-Torres *et al*. 2016). Intraspecific microevolutionary analyses further demonstrate that local strains fine-tune phosphorus acquisition through gene gains, losses, and subtle regulatory adjustments of alkaline phosphatases and phosphonate transporters (Gómez-Lunar *et al*. 2016). However, while these metabolic adaptations have been characterized in laboratory isolates and surface habitats, how severe phosphorus limitation structures deep, continuous sedimentary columns remains unknown.

Most structural characterizations describe microbial mats as thin, millimeter- to centimeter-scale features (Des Marais 2003; Prieto-Barajas *et al*. 2018). Nevertheless, progressive biostabilization and sediment accretion can allow such communities to thicken substantially over time, eventually giving rise to deeper laminated or lithified stratigraphic records (Noffke and Awramik 2026). Within such thicker systems, it remains an open question how vertical structure reflects depositional history, the living community’s ongoing accommodation to present-day geochemical conditions, or both processes operating simultaneously.

In this study, we conducted a high-resolution, depth-resolved exploratory survey of a continuous 34-cm sediment core partitioned into 11 discrete stratigraphic layers (L1 to L11) from Poza Roja 2, an alkaline, sub-hypersaline pond in the Pozas Rojas complex of CCB. By integrating physical stratigraphy, organic carbon and nutrient stoichiometry, and high-throughput 16S rRNA gene metabarcoding, we aimed to map how domain-level community shifts track steep down-core biogeochemical gradients and persistent phosphorus scarcity. Rather than assuming strictly deterministic filtering, this pilot characterization provides a baseline for evaluating how Bacteria and Archaea co-occur alongside extreme Redfield-deviant stoichiometry across a deep, continuous sedimentary column.

## Materials and Methods

### Study Site and Core Sampling

The Cuatro Ciénegas Basin (CCB) is located in northern Mexico. The CCB climate is arid with an average annual temperature of 21.9 °C and an average annual rainfall of 253 mm (Montiel-González *et al*. 2021). Precipitation is highly variable between years but is concentrated in the summer months (Montiel-González *et al*. 2018). The Pozas Rojas system (PR) is within the Los Hundidos Reserve, located on the east side of the CCB (26°52’11"N, 102°01’12"W and 712 m.a.s.l.). This system consists of several small permanent and temporary ponds that exhibit a red-brown color. The PR ponds are fed by a subterranean river that emerges from several sinkholes (Rebollar *et al*. 2012). The water temperature fluctuates between 10 °C and 60 °C, with maximum daily fluctuations of 15 °C during winter and 45 °C in summer, as documented for the PR desiccation pond (Peimbert *et al*. 2012); given the shared biogeochemical and hydrological context of the PR complex, analogous thermal dynamics are expected across the system, including Pond 2. Some of these red ponds are characterized by extremely oligotrophic waters (Garcia-Ulloa *et al*. 2022; Peimbert *et al*. 2012; Vázquez-Rosas-Landa *et al*. 2020), with a high C:N:P ratio in the water column (15820:157:1; Peimbert *et al*. 2012). In September 2023, during one of the driest seasons for the CCB, a sediment core was collected from Poza Roja 2 (26°52’11"N, 102°01’12"W) ponds, using a 50-cm-long, 7.62-cm-diameter PVC tube, yielding a 34 cm sediment core for analysis. The sediment core was stored at 4 °C until subsequent analyses.

### Physical Core Description

The core from Pond 2 was processed at the Laboratorio de Estudios Paleoambientales y Paleoclimáticos de la Escuela Nacional de Estudios Superiores (ENES, UNAM) at the Morelia campus. Before opening the core, every centimeter was marked to better document its description. The top of the core was also identified and marked. The core was then cut longitudinally, and the sediment stratigraphy was described, considering texture, color, and sedimentary composition, following the protocol established by Schnurrenberger *et al*. (2003). Eleven layers, each with distinct laminations, were identified. Layers were distinguished by their predominant color. Additionally, samples were collected from each layer for biogeochemical and microbial analysis.

### Nutrient Analyses

The pH of deionized water (1:10 w/v) was measured with a digital potentiometer (Thermo Scientific Orion 3 Star Plus). A water sample was collected independently from Poza Roja 2 in June 2024 and submitted to Fertilab for physicochemical characterization. The analysis included major ions, salinity-related variables, nutrients, and trace elements. Sediment samples were dried for 72 h at 105 °C and then ground in an agate mortar for chemical analyses. Total Carbon (TC) and inorganic C (Ci) were determined by coulometric detection (Huffman 1977) using a total carbon analyzer (UIC model CM5012, Chicago, USA). Total organic C (TOC) was calculated as the difference between TC and Ci. Total Nitrogen (TN) and Total Phosphorus (TP) were determined after acid digestion; TN was measured by the Kjeldahl macro method (Bremner 1996), and TP by molybdate reduction with ascorbic acid (Murphy and Riley 1962). Both nutrients were measured by colorimetry in a Bran-Luebbe Auto Analyzer 3 (Norderstedt, Germany). Inorganic phosphorus (Pi) and organic phosphorus (Po) were determined using the method proposed by Tiessen and Moir (1993). Phosphorus was extracted from 3 g of sediment using 1 M HCl, and two aliquots were collected: one for the inorganic fraction (Pi) and another for the total digested phosphorus (DP). The Pi aliquots were acidified with 0.9 M H2SO4, while the DP aliquots were digested with 50% (NH4)2S2O8 (ammonium persulfate). Both extracts were measured by colorimetry using a Bran-Luebbe Auto Analyzer 3 (Norderstedt, Germany). Organic P (Po) was calculated as the difference between DP and Pi. Total phosphorus (PT) was calculated as the sum of the inorganic and organic fractions (PT = Pi + Po).

### DNA Extraction and 16S rRNA Gene Amplicon Sequencing

Total genomic DNA was extracted from 0.5 g of wet sediment from each distinct layer (L1–L11) using the DNeasy PowerSoil Kit (Qiagen, Germany). To overcome the structural complexity of the stratified sediment matrix and ensure efficient lysis of recalcitrant cells, a modified pretreatment protocol was implemented as follows: (i) Samples were prewashed with a 0.5 M EDTA solution, resuspended in sterile distilled water, and immediately frozen at −20 °C for 4–6 h; (ii) Samples were subsequently heated at 65 °C for 10 min and immediately refrozen at −20 °C for 2 h; (iii) After thawing at room temperature, enzymatic and chemical lysis was initiated according to the kit’s specifications, with the mechanical disruption phase extended to 15 min at maximum speed using a Vortex-Genie 2 mixer (Scientific Industries, USA). The remaining extraction steps were completed strictly adhering to the manufacturer’s instructions. DNA quality and yield were quantified using a Qubit 4 Fluorometer with the dsDNA BR Assay Kit (Thermo Fisher Scientific, USA), yielding final concentrations ranging between 8.8 and 149 ng/µL. DNA integrity was visualized by agarose gel electrophoresis. The V3–V4 hypervariable regions of the 16S rRNA gene were amplified using the primer pair 341F (5’-CCTACGGGNGGCWGCAG-3’) and 806R (5’-GGACTACNVGGGTWTCTAAT-3’) (Apprill *et al*. 2015; Wasimuddin *et al*. 2020). Subsequent library preparation and high-throughput paired-end sequencing (2 × 250 bp) were performed by Novogene Corporation utilizing the Illumina NovaSeq 6000 platform. The raw sequence datasets generated during this study were deposited in the NCBI Sequence Read Archive (SRA) under the accession number PRJNA1238309.

### Amplicon Sequence Variants and Taxonomic Classification

Demultiplexed paired-end reads provided by Novogene were processed using the Quantitative Insights Into Microbial Ecology 2 (QIIME2, v2024.10) pipeline (Bolyen *et al*. 2019). Quality filtering, trimming, denoising, and chimera removal were executed using the DADA2 algorithm via the q2-dada2 plugin to generate high-resolution Amplicon Sequence Variants (ASVs) (Callahan *et al*. 2016). Taxonomic assignment was performed using the q2-feature-classifier plugin with a Naive Bayes classifier trained against the GreenGenes2 database (v2024.09; McDonald *et al*. 2024), applying a confidence threshold of 0.9 for taxonomic assignment. ASVs classified as microbial dark matter (Unknown-Unknown and Known-Unknown; Escudeiro *et al*. 2022) were filtered out prior to downstream analyses.

### Microbial Community Ecology and Statistical Analyses

Downstream ecological analyses were performed in the R environment (v4.3) and Python (v3.11). Taxonomic profiles and relative abundances of Bacteria and Archaea lineages were characterized using QIIME2. Alpha diversity metrics and beta diversity distance matrices, including Bray–Curtis and Jaccard indices, were calculated using the microeco package. Canberra distance was specifically included to capture compositional dissimilarities driven by the rare biosphere, given the strong dominance of a few genera in relative abundance profiles. Unconstrained Principal Coordinate Analysis (PCoA) based on Bray–Curtis and Jaccard dissimilarity matrices, alongside hierarchical clustering using Canberra distances, served as the primary ordination tools to evaluate community architecture without imposing environmental constraints.

To identify persistent and localized micro-niches across the sediment core, core and unique taxa were determined from QIIME2 sequence count (feature abundance) data; the resulting distributions were visualized as flower plots and frequency histograms using the matplotlib library (v3.8.0) in Python (Hunter 2007). Layer-specific distribution of the top 20 dominant core genera was visualized using the ggplot2 library in R. To evaluate monotonic relationships between depth progression and specific ASV shifts, non-parametric Spearman rank correlations were computed, applying the Benjamini–Hochberg False Discovery Rate (FDR) correction (p < 0.05).

To explore potential co-linearity between community structure and down-core biogeochemical variables, Redundancy Analysis (RDA) was performed under two non-nested environmental models: Model A (absolute nutrient concentrations: pH, ln-transformed TOC, TN, and Po) and Model B (elemental stoichiometry: pH, ln C:N, and ln N:Po). Stoichiometric ratios were natural-log transformed rather than used in raw form to avoid the numerical asymmetry and spurious correlation inherent to untransformed compositional ratios (Pearson 1897). Genus-level count data were Centered Log-Ratio (CLR) transformed after adding a uniform pseudocount of +1 across the full community matrix (Aitchison 1986). Global model significance was assessed via a Freedman–Lane conditional permutation test (9,999 permutations) (Freedman and Lane 1983; ter Braak 1992), and overall adjusted R² values were calculated to account for predictor quantity relative to sample size (Peres-Neto *et al*. 2006). However, given the limited sample size of a single stratified core (n = 11 layers) and spatial autocorrelation between contiguous horizons, RDA models were interpreted strictly as exploratory indicators of co-occurrence rather than deterministic drivers.

### Phylogenetic Reconstruction and Comparative Analysis of Archaeal Lineages

To investigate the evolutionary relationships of the archaeal community, a maximum likelihood phylogenetic tree was constructed. The representative sequences of the archaeal ASVs were aligned and the phylogeny was inferred using IQ-TREE. The best-fit nucleotide substitution model, GTR+I+R9, was selected based on ModelFinder criteria. Branch supports were assessed using 1,000 ultrafast bootstrap replicates. *Escherichia coli* K12 was used as the outgroup to root the tree. The resulting tree was visualized and annotated in the Interactive Tree Of Life (iTOL) platform. Additionally, to contextualize the archaeal diversity of Poza Roja 2 within the broader Cuatro Ciénegas Basin, we performed a presence/absence comparative analysis of archaeal lineages against previously published data from the Archaean Domes (Medina-Chávez *et al*. 2023).

## Results

### Physical Stratigraphy and Biogeochemical Profiles Across the 34-cm Core

The 34-cm sediment core recovered from Poza Roja 2 in the Cuatro Ciénegas Basin (Figure 1) exhibited distinct physical laminations and color variations, allowing the identification of 11 discrete stratigraphic layers labeled sequentially from surface to depth (L1 to L11; Figure 2).

**Figure 1.**
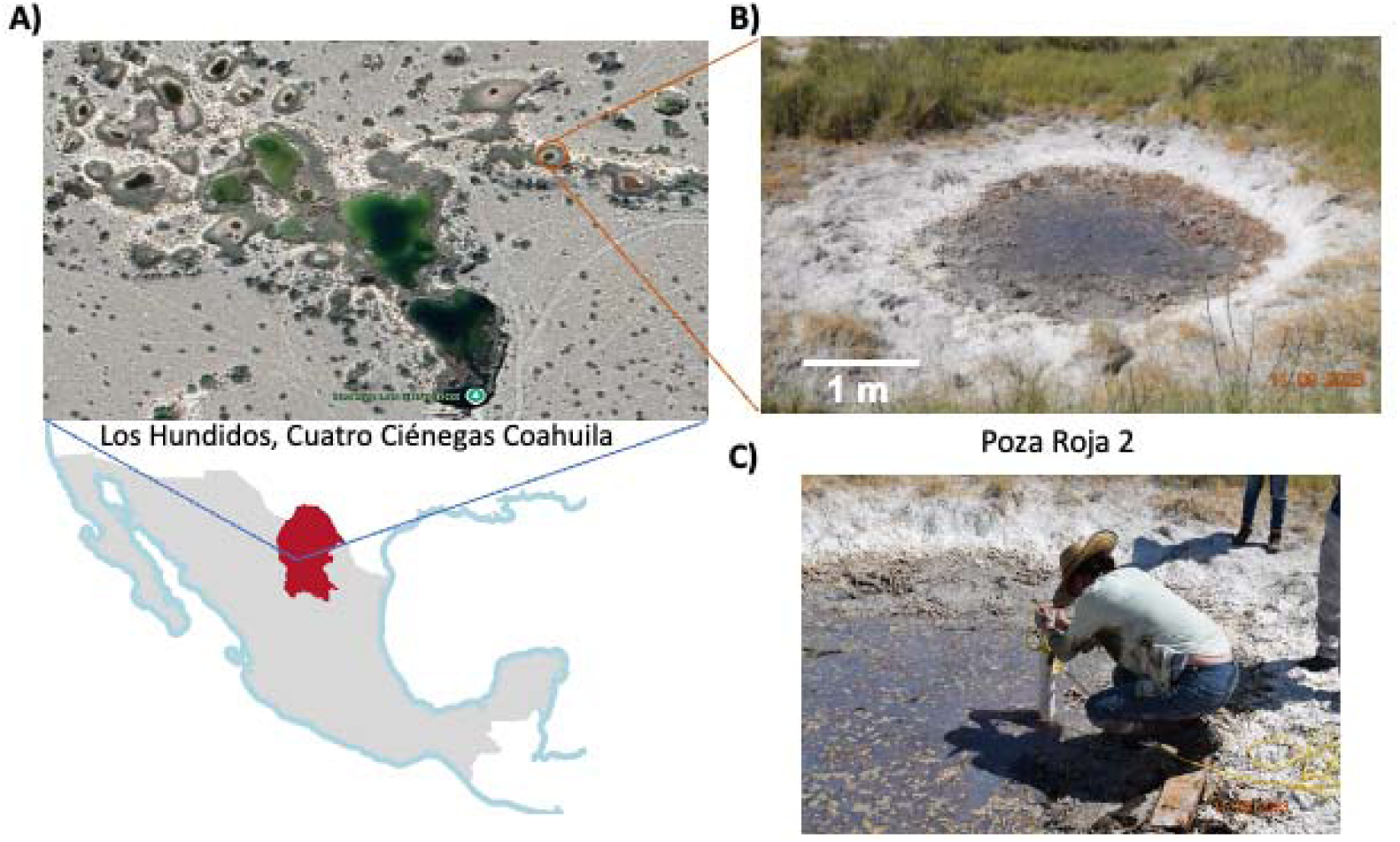
Study site and sediment sampling at Poza Roja 2, Cuatro Ciénegas Basin, Mexico. A) Geographic location of the Pozas Rojas hydrological system within Los Hundidos in the Cuatro Ciénegas Basin, including a satellite overview of the pond complex, which comprises multiple shallow ponds and a deeper lagoon. Satellite imagery was obtained from Google Earth (2026). B) Field photograph of Poza Roja 2. C) Sediment core collected from Poza Roja 2 using a PVC tube. The core was longitudinally sectioned into two halves before subsampling and downstream analyses.

**Figure 2.**
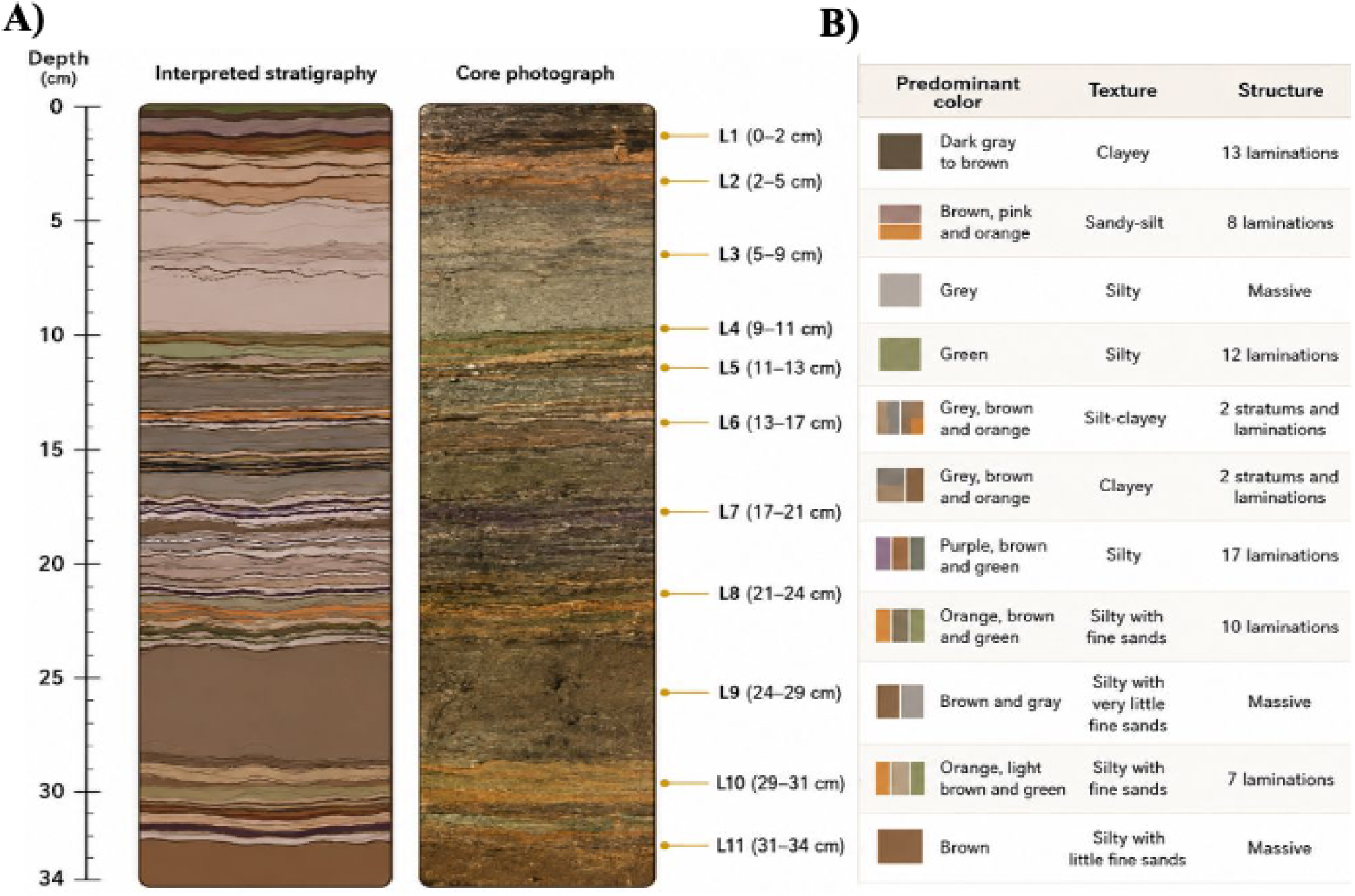
Physical stratification of the Poza Roja 2 sediment core. A) Interpreted stratigraphy and photograph of the 34-cm core, divided into 11 layers (L1–L11); layer boundaries and depth intervals are indicated. B) Stratigraphic characteristics of each layer, including predominant color, sediment texture, and internal structure (number of laminations or massive structure). The profile alternates between laminated and massive intervals and records pronounced vertical variation in color and texture.

The surface layer (L1, 0–2 cm) consisted of dark gray to brown clayey sediment with 13 laminations. Below it, the profile alternated between laminated and massive intervals with pronounced vertical variation in color and texture (Figure 2). Laminated layers included L2 (2–5 cm; brown, pink, and orange sandy silt), L4 (9–11 cm; green silt), L5 and L6 (11–17 cm; grey, brown, and orange silty clay to clay with two strata and laminations), L7 (17–21 cm; purple, brown, and green silt with 17 laminations, the only layer with purple laminations), L8 (21–24 cm; orange, brown, and green silt with fine sands), and L10 (29–31 cm; orange, light brown, and green silt with fine sands). Three layers lacked visible lamination and were classified as massive: L3 (5–9 cm; grey silt), L9 (24–29 cm; brown and grey silt), and L11 (31–34 cm; brown silt) (Figure 2).

Water column analysis confirmed sub-hypersaline, alkaline, and highly oligotrophic conditions in Poza Roja 2 (Table S1). Biogeochemical analysis across the sediment profile revealed marked, non-monotonic down-core variation (Figure 3; Table S2). Sediment pH ranged from 7.2 in surface layers to 8.4 in intermediate horizons, maintaining moderately alkaline conditions throughout the profile. Total organic carbon (TOC) ranged from 2.25% (L3) to 4.55% (L1) dry weight, with secondary maxima in L4 (3.98%) and L8 (4.11%). Total nitrogen (TN) ranged from 0.37% (L3) to 1.42% (L8). Organic phosphorus (Po) was low throughout the core, ranging from 0.002% (L4) to 0.016% (L1) dry weight (Figure 3; Table S2). Consequently, molar C:Po and N:Po ratios exceeded canonical Redfield proportions (106:16:1) in all 11 layers, by 4.6- to 48.5-fold and 4.3- to 87.8-fold, respectively, with the strongest imbalance in L4 (C:N:Po = 5,137:1,404:1), indicating severe phosphorus depletion throughout the sedimentary column (Figure 3; Table S2).

**Figure 3.**
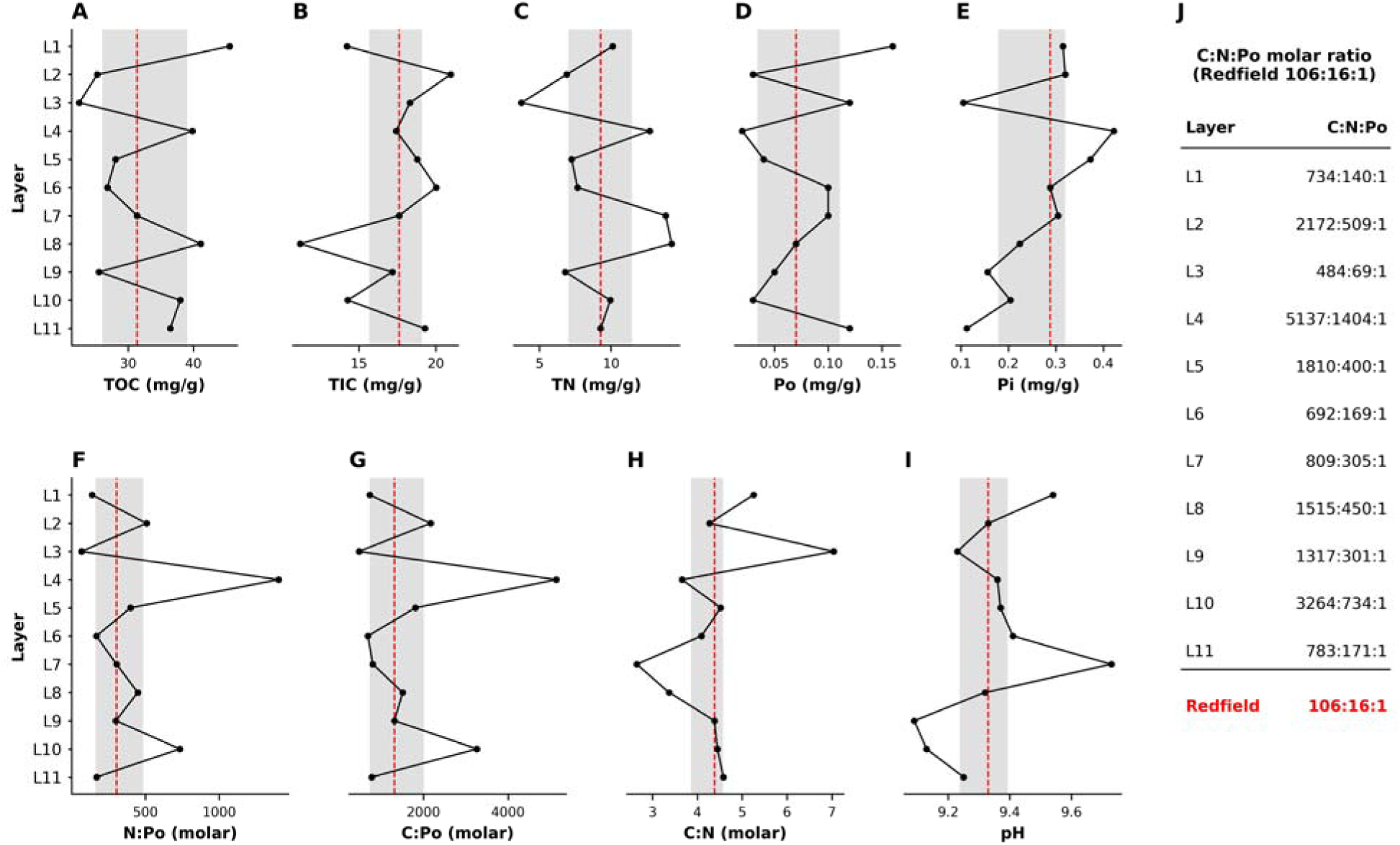
Depth profiles of nutrient concentrations, stoichiometric ratios, and pH across the Poza Roja 2 sediment core. A) Total organic carbon (TOC). B) Total inorganic carbon (TIC). C) Total nitrogen (TN). D) Organic phosphorus (Po). E) Inorganic phosphorus (Pi). F) N:Po molar ratio. G) C:Po molar ratio. H) C:N molar ratio. I) pH. Red dashed lines indicate the median, and shaded areas represent the interquartil range (Q1–Q3). J) Layer-specific C:N:Po molar ratios normalized to Po = 1; the canonical Redfield ratio (106:16:1) is shown as a reference.

### Domain-Level Microbial Composition and High Archaeal Abundance

High-throughput 16S rRNA gene amplicon sequencing yielded 742,204 reads across the 11 layers, resolved into 8,382 Amplicon Sequence Variants (ASVs; 7,139 bacterial and 1,243 archaeal). Taxonomic assignment evaluation demonstrated high confidence across known lineages alongside a substantial proportion of uncharacterized microbial dark matter (Figure 4A; Table S3). Archaea accounted for 31.0% of total prokaryotic reads, varying markedly among layers from 8.3% in L5 to 43.7% in L9 (Figure 4B). Alpha diversity metrics showed a bimodal profile, with a pronounced mid-depth decline in layer L7 (Figure 4C; Table S4).

**Figure 4.**
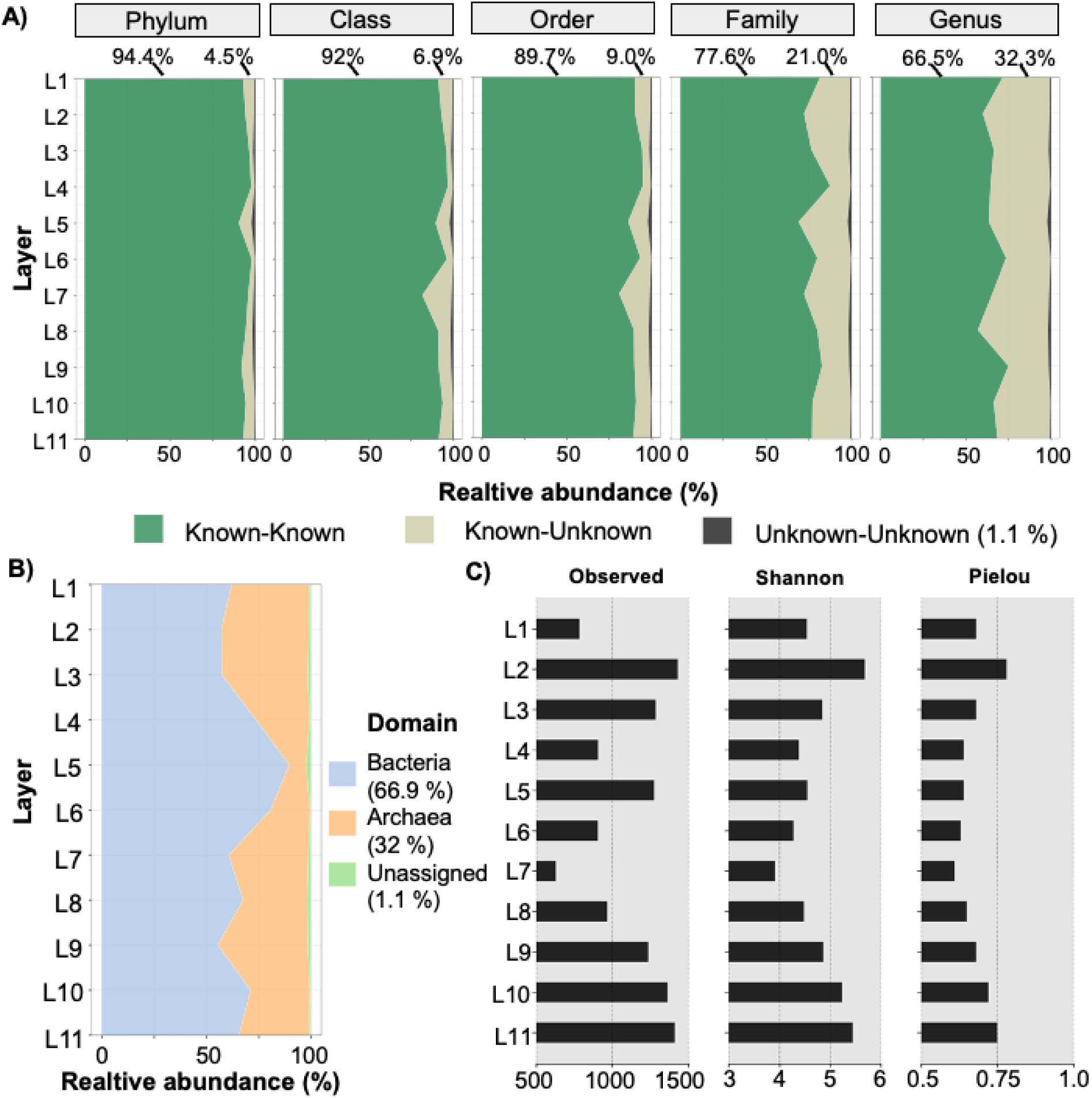
Taxonomic resolution, domain-level structure, and alpha-diversity dynamics of the prokaryotic community across the Poza Roja 2 sediment core. A) Relative abundance profiles of taxonomic classification confidence from phylum to genus. Known-Known denotes taxa resolved at the corresponding rank, Known-Unknown denotes sequences unresolved at that rank, and Unknown-Unknown denotes sequences lacking significant homology to current reference databases. Values above each panel indicate the global mean percentage in each category at that rank. B) Relative abundance of Bacteria, Archaea, and unassigned ASVs across layers L1–L11. C) Down-core profiles of observed ASV richness, Shannon diversity, and Pielou evenness, showing a bimodal diversity pattern and a pronounced mid-depth minimum centered on L7.

At the phylum level, bacterial reads were dominated by Firmicutes_B (20.3%), Proteobacteria (18.4%), Firmicutes_A (16.4%), Campylobacterota (8.8%), and Firmicutes_E (8.0%) (Greengenes2/GTDB nomenclature). Firmicutes_B was largely represented by the phototrophic heliobacterium *Heliorestis*, which showed a discontinuous vertical distribution: abundant in L1 (37.4% of bacterial reads), scarce in L2–L4 (0.8–8.7%), and peaking in L5–L6 (41.9% and 47.6%). Cyanobacteria remained a minor component throughout the core (≤2.3% of bacterial reads in every layer). Within the domain Archaea, Thermoplasmatota (58.4% of archaeal reads) and Halobacteriota (34.0%) dominated the profile. The candidate lineage SM1-50 (Thermoplasmatota; 55.8% of archaeal reads) and the halophilic genus *Halohasta* (10.5%) were detected across both upper and basal layers (Figure 4B; Table S3).

### Unconstrained Community Ordination and Vertical Zonation

To evaluate community architecture without imposing environmental constraints, Principal Coordinate Analysis (PCoA) based on Bray-Curtis and Jaccard dissimilarity matrices, alongside hierarchical clustering using Canberra distances, were performed as the primary ordination tools (Figure 5). PCoA based on Bray-Curtis distance explained 58.4% of total compositional variance along the first two axes (PCoA1 = 38.2%, PCoA2 = 20.2%), separating the 11 layers into three compositional zones (Figure 5B, C).

**Figure 5.**
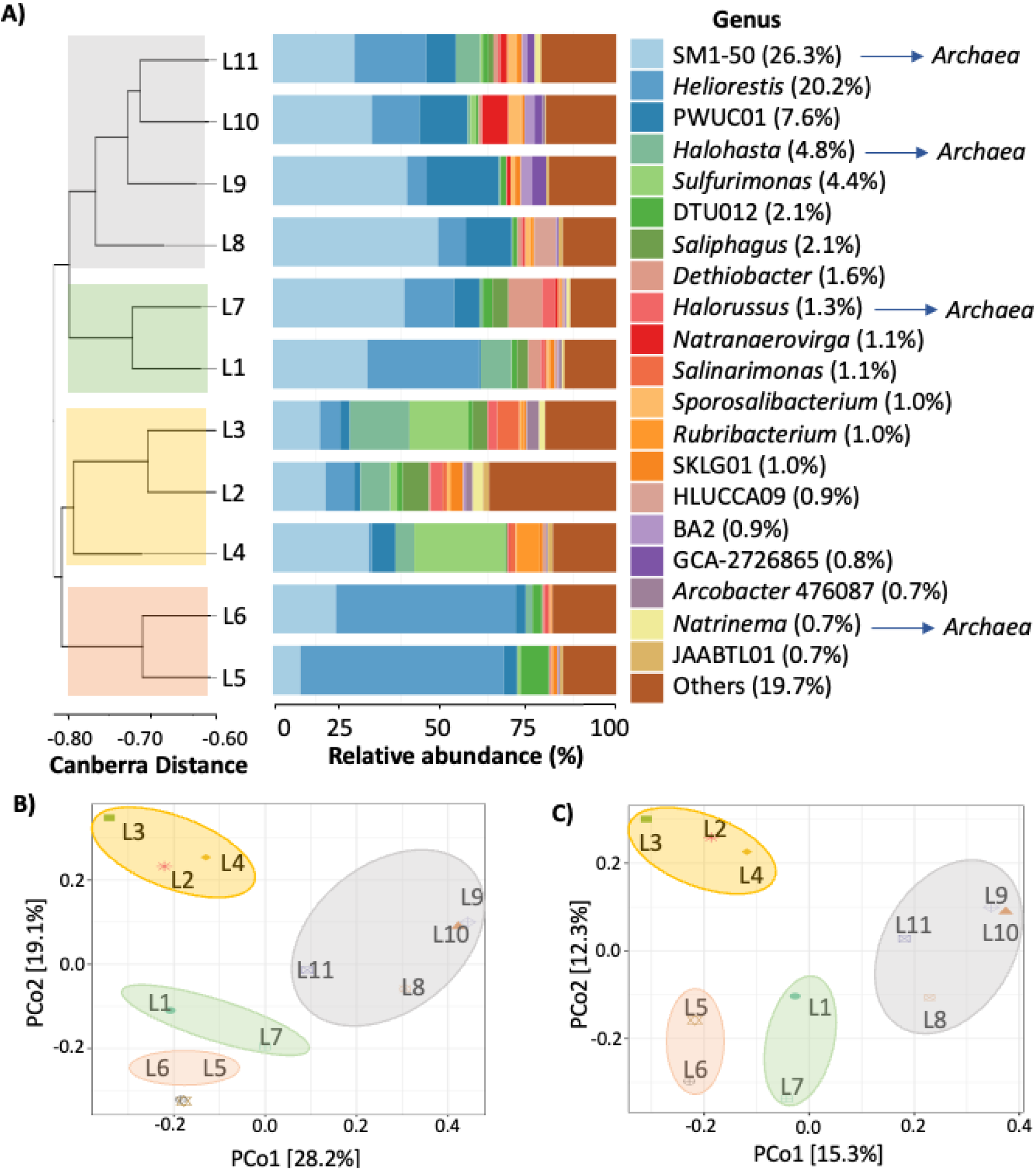
Taxonomic composition and compositional stratification of Known-Known genera along the sediment core. A) Relative abundance of the 20 most abundant bacterial and archaeal genera across layers L1–L11, with archaeal lineages identified in blue. Hierarchical clustering based on Canberra distance summarizes compositional similarity among layers. B) Principal coordinate analysis (PCoA) based on Bray–Curtis dissimilarity. C) PCoA based on Jaccard dissimilarity. Both ordinations support the separation of the sediment layers into distinct compositional zones.

The Upper Zone comprised layers L2 to L4 (2–11 cm). These layers combined high archaeal representation in L2–L3 (42.3% and 41.7%) with low *Heliorestis* abundance (0.8–8.7%), and included both the lowest TOC and TN values (L3) and the strongest phosphorus imbalance of the core (L4). The Intermediate Zone encompassed layers L5 and L6 (11–17 cm), where *Heliorestis* reached its maximum abundance (41.9% and 47.6% of bacterial reads) and archaeal representation its minimum (8.3% in L5). The Basal Zone spanned layers L8 to L11 (21–34 cm) and was dominated by Firmicutes_A (27.1% of bacterial reads), together with archaeal lineages including SM1-50 and Halobacteriota. Layers L1 (surface) and L7 occupied transitional coordinates along PCoA2 between these clusters. Hierarchical clustering based on Canberra distances was consistent with this three-zone partition (Figure 5A).

Detailed layer-by-layer distribution profiles of the 20 dominant core genera highlighted clear vertical turnover across these zones (Figure 6; Figure S1). The shared core microbiome comprised 45 persistent genera across the profile, whereas unique genera were enriched in specific micro-niches (Figure S2; Table S5, Table S6). Non-parametric Spearman rank correlation analysis identified specific bacterial taxa that increased significantly with depth, including PWUC01, *Natranaerovirga*, and DTU015 (FDR-adjusted P < 0.05; Figure S3; Table S7).

**Figure 6.**
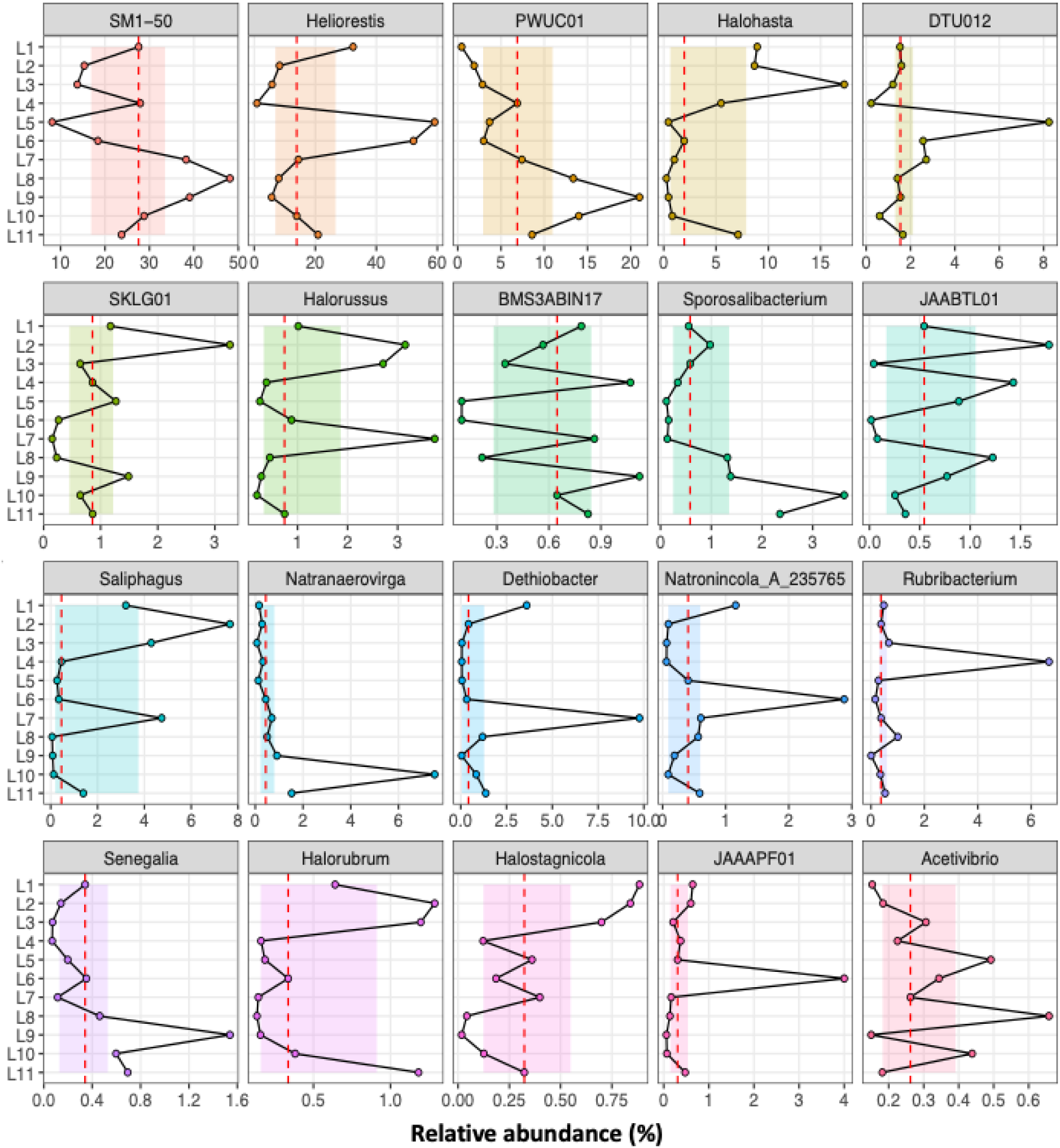
Layer-specific distributions of the 20 dominant core genera across the sediment column. Relative abundance profiles are shown for each genus across layers L1–L11. Red dashed lines indicate the median, and shaded areas represent the interquartile range (Q1–Q3), highlighting layer-specific enrichment relative to the community baseline.

### Phylogenetic Reconstruction of Archaea and Regional Comparative Analysis

Maximum likelihood phylogenetic reconstruction of archaeal 16S rRNA gene sequences resolved distinct monophyletic clades corresponding to Halobacteriota, Thermoplasmatota, Nanoarchaeota, and Methanobacteriota, with concentric rings illustrating taxonomic assignment and layer-specific occurrence from L1 to L11 (Figure 7).

**Figure 7.**
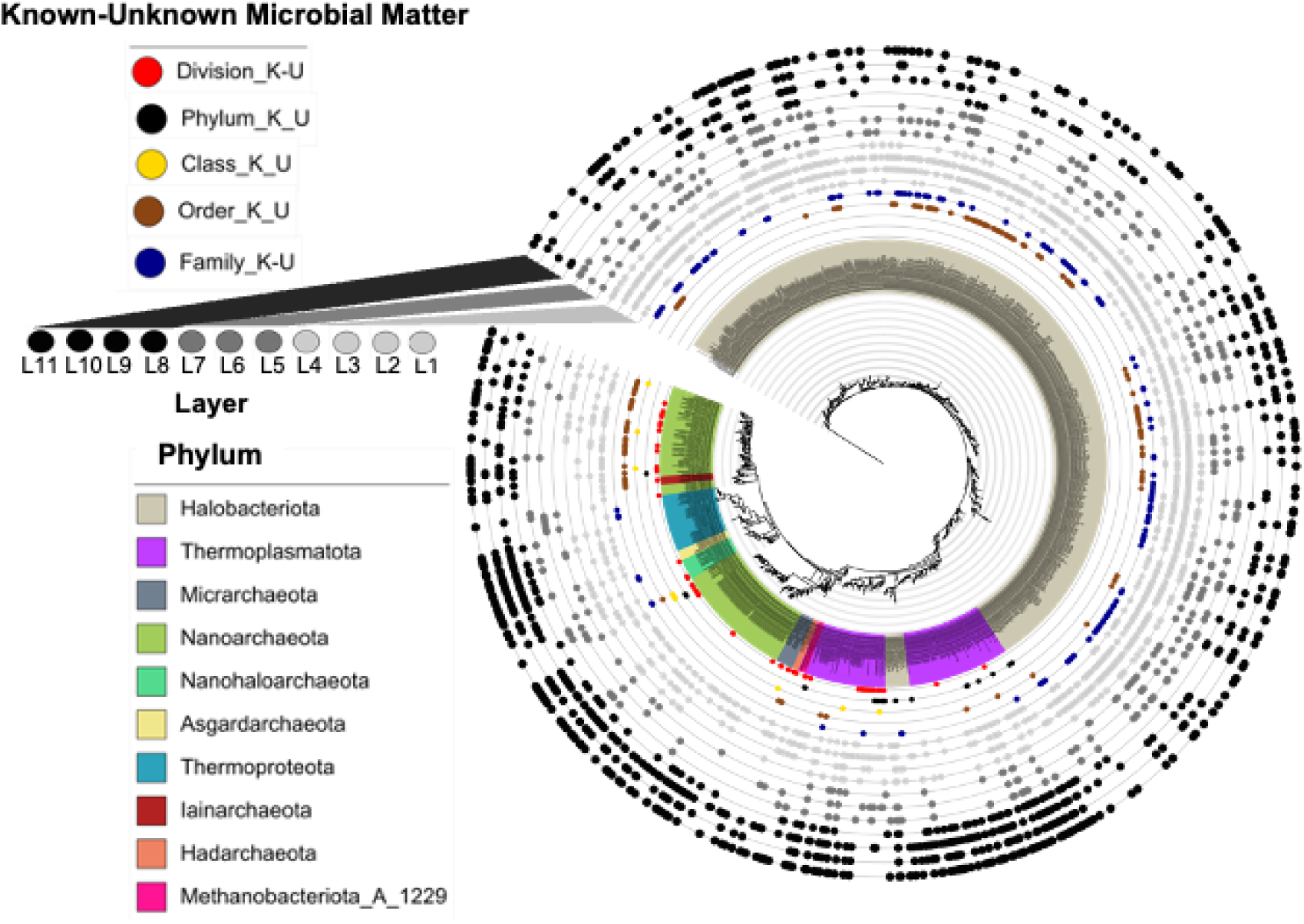
Maximum-likelihood phylogenetic tree reconstructed from archaeal 16S rRNA gene V3–V4 sequences (n = 1,243) recovered from the Poza Roja 2 sediment core. Tree tips are colored by archaeal phylum. Five inner annotation rings indicate the lowest taxonomic rank resolved for Known-Unknown ASVs: division (red), phylum (black), class (yellow), order (brown), and family (blue). Eleven outer rings show the presence of each ASV by sediment layer, from L1 (innermost, lightest gray; surface) to L11 (outermost, black; 31–34 cm), with grayscale intensity increasing with depth. The tree was rooted with *Escherichia coli* K-12 as the outgroup.

Comparative analysis between Poza Roja 2 and previously published datasets from the Archaean Domes of Cuatro Ciénegas revealed a broader phylum-level archaeal footprint in Poza Roja 2 (13 phyla versus 5 phyla in Archaean Domes; Figure 8; Table S10). Furthermore, genus-level overlap between both systems was nearly absent, reflecting distinct environmental selection pressures driven by local hydrochemistry and sediment depth (Figure 8; Table S9, Table S10). Literature synthesis of representative isolates further emphasized the unique metabolic capabilities of these resident archaeal and bacterial taxa (Table S11).

**Figure 8.**
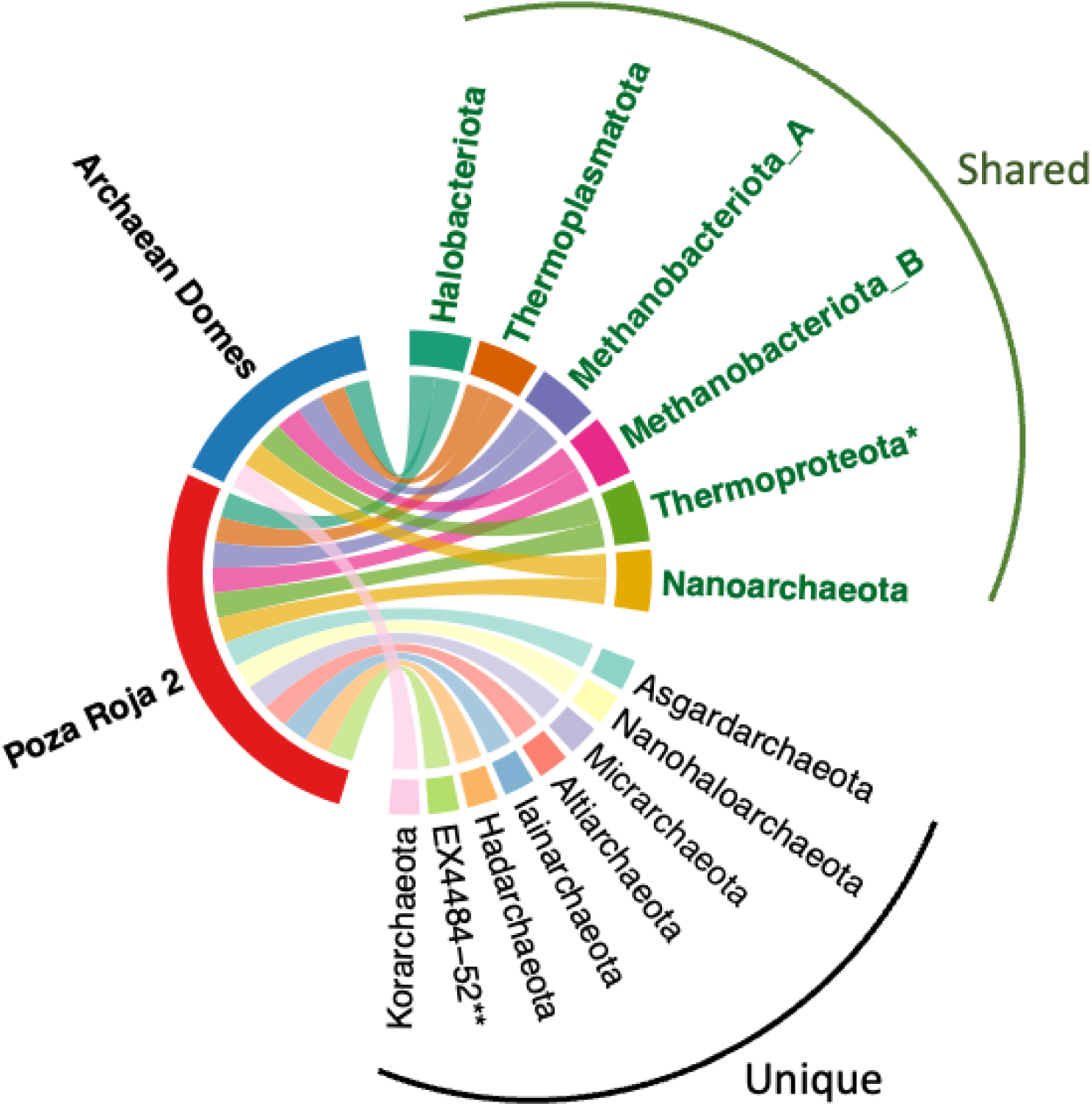
Comparison of archaeal phyla detected in Poza Roja 2 (13 phyla; Greengenes2/GTDB taxonomy) and Archaean Domes (five phyla; NCBI/SILVA taxonomy) (Medina-Chávez *et al*. 2023; Rodríguez-Cruz *et al*. 2024). Shared and system-specific phyla are indicated. *GTDB reclassified the former phylum Thaumarchaeota as the class Nitrososphaeria within Thermoproteota; the Thermoproteota ASVs in this study include the family Nitrososphaeraceae, the GTDB equivalent of the Thaumarchaeota reported for Archaean Domes. **EX4484-52 is a GTDB placeholder for a single-species lineage of uncertain phylogenetic placement; no established NCBI/SILVA equivalent is currently available.

### Exploratory Evaluation of Biogeochemical Co-Linearity

To exploratory evaluate potential co-variation between community composition and down-core biogeochemical parameters (pH, TOC, TN, Po, C:N, and N:Po), Redundancy Analysis (RDA) was conducted under two environmental models: Model A (absolute concentrations) and Model B (elemental stoichiometry) (Figure S4; Table S8). Global permutation tests revealed that neither constrained model achieved statistical significance (Model A: pseudo-F = 1.60, P = 0.120; Model B: pseudo-F = 1.55, P = 0.155; Table S8).

## Discussion

Depth-resolved characterizations of benthic microbial mats have traditionally focused on millimeter-scale photic horizons within upper surface layers (Des Marais 2003; Bolhuis *et al*. 2014). In contrast, the 34-cm continuous profile from Poza Roja 2 shows that compositional zonation and marked biogeochemical variation extend across decimeter scales in a thickened biogenic deposit. Unconstrained ordination (PCoA) and Canberra-based hierarchical clustering grouped the layers into three compositional zones: upper (L2–L4), intermediate (L5–L6), and basal (L8–L11), with L1 and L7 in transitional positions. This vertical organization co-occurs with phosphorus depletion throughout the core: molar C:Po and N:Po ratios exceeded canonical Redfield proportions (106:16:1) in all 11 layers, by up to 48.5-fold and 87.8-fold, respectively, both in L4 (C:N:Po = 5,137:1,404:1). Phosphorus scarcity is therefore not restricted to the surface but extends across the entire sedimentary column, consistent with the extreme stoichiometric imbalances previously reported for Pozas Rojas waters (Peimbert *et al*. 2012; Garcia-Ulloa *et al*. 2022).

This pattern bears directly on the question posed at the outset: whether the vertical structure of a thickened mat reflects depositional history, present-day conditions, or both. The physical stratigraphy, with its laminations, color bands, and massive intervals (Figure 2), most likely records depositional and post-depositional history. The microbial signal, however, does not simply follow depth from the surface: the phototrophic heliobacterium *Heliorestis* was abundant in L1 (37.4% of bacterial reads), nearly absent from L2–L4 (0.8–8.7%), and reached its maximum in L5–L6 (41.9–47.6%). This discontinuous distribution may reflect active populations responding to present-day conditions, DNA from dormant or inactive cells retained in the sediment (González-Sánchez *et al*. 2026), or both, a distinction that 16S rRNA gene amplicon data cannot resolve. Separating these two imprints will require activity-based approaches such as metatranscriptomics.

Understanding how diverse microbial communities maintain cellular integrity and turnover under such extreme, down-core phosphorus starvation requires examining established physiological adaptation frameworks within the Cuatro Ciénegas Basin. Although our 16S rRNA amplicon survey is descriptive and cannot measure metabolic fluxes directly, bacteria in this basin have been shown to overcome chronic phosphorus starvation by substituting membrane phospholipids with non-phosphorus sulfolipids via genes closely related to cyanobacterial *sqd1* and *sqdX*, apparently acquired by horizontal gene transfer, an important saving given that membrane phospholipids account for approximately 30% of total cellular phosphate in most organisms (Alcaraz *et al*. 2008). Furthermore, microbial communities across soil and sediment habitats in the valley actively exploit alternative phosphorus pools across multiple oxidation states, utilizing extracellular DNA recycling via phosphodiesterases alongside the enzymatic cleavage of phosphonates and phosphites (Tapia-Torres *et al*. 2016). Intraspecific comparative genomics also demonstrates that endemic strains fine-tune phosphorus acquisition through gene gains, losses, and subtle regulatory modifications of alkaline phosphatases (Gómez-Lunar *et al*. 2016). In Poza Roja 2, such strategies provide a plausible, though as yet untested, framework for how bacterial communities persist under severe, down-core phosphorus depletion.

A second notable feature of the Poza Roja 2 profile is the substantial representation of Archaea, which accounted for 31.0% of prokaryotic reads overall and ranged from 8.3% (L5) to 43.7% (L9). This layer-to-layer variation, obtained under an identical protocol, argues against a uniform primer-associated amplification bias. Archaeal reads were dominated by the candidate lineage SM1-50 (Thermoplasmatota) and by Halobacteriota, a phylum commonly associated with hypersaline settings (Jahnke *et al*. 2008), even though the overlying water was sub-hypersaline (5.0–8.1 g/L). Phosphorus scarcity, alongside salinity, is a plausible contributing factor to this archaeal representation, although the present data cannot establish causality. Recent work in the basin has likewise revealed substantial archaeal diversity (Medina-Chávez *et al*. 2023), with nearly half of recovered archaeal genomes unclassifiable even at the genus level (Rodríguez-Cruz *et al*. 2024).

Parallel to these dominant bacterial and archaeal lineages, the Poza Roja 2 sediment core harbors a substantial rare biosphere, comprising 39 of the 45 core genera, with relative abundances of 1% or less. This long tail of low-abundance microorganisms suggests niche partitioning along the down-core gradients documented here. In highly variable desert ecosystems subject to seasonal desiccation and hydrological shifts, low-abundance taxa can act as an ecological seed bank, remaining dormant or metabolically suppressed until environmental conditions favor their proliferation (Jousset *et al*. 2017).

Notwithstanding these ecological insights, a critical methodological consideration of this study is the constrained statistical power inherent to evaluating a single, highly stratified sediment core (n = 11 layers). Metagenomic time series from another CCB system indicate that a core community can account for more than 75% of total abundance despite seasonal turnover (Madrigal-Trejo *et al*. 2023), but the representativeness of a single core cannot be confirmed with the present dataset. Because contiguous sedimentary horizons exhibit spatial autocorrelation, constrained multivariate models such as RDA failed to achieve statistical significance (P > 0.12). Consequently, down-core biogeochemical profiles must be interpreted as co-occurring environmental conditions rather than proven deterministic drivers of community assembly. Future investigations incorporating multi-core spatial replication across the Pozas Rojas complex, paired with shotgun metagenomics, metatranscriptomics, and intact polar lipid (IPL) profiling, will be essential to directly quantify membrane lipid remodeling, verify phosphorus scavenging gene expression, and resolve the functional metabolic networks operating across these deep biogenic archives. This is particularly urgent given the ongoing hydrological stress and risk of ecosystem disassembly in this endangered desert oasis (González-Sánchez *et al*. 2026; Rodríguez-Cruz *et al*. 2025).

## Declarations

### Ethics Approval and Sampling Permits

Sample collection was conducted under official permits issued by the General Directorate of Wildlife (DGVS, Mexico): No. SPARN/DGVS/07643/23 and SPARN/DGVS/09800/24. Field access and sampling authorization for Los Hundidos reserve were granted by Genesis 4C and Grupo ECCO.

### Data Availability

The raw 16S rRNA gene amplicon sequencing datasets generated during the current study have been deposited in the NCBI Sequence Read Archive (SRA) under BioProject accession number PRJNA1238309. All physical, chemical, and taxonomic data matrices supporting the findings of this study are provided within the supplementary information files.

### Code Availability

Custom R and Python scripts used for bioinformatic processing, alpha and beta diversity calculations, ordination analyses, and data visualization are available from the corresponding authors upon reasonable request.

### Author Contributions

F.G.O. and G.O.A. conceived and designed the study. F.G.O., A.P.V., and G.I.A. conducted field sampling. G.I.A. performed the physical stratigraphy and lithological description of the sediment core. A.G.S. performed laboratory analyses, including DNA extraction and sequencing, and conducted bioinformatic and multivariate statistical analyses. G.O.A. and A.G.S. wrote the manuscript. All authors reviewed, revised, and approved the final manuscript.

## Acknowledgments

This study was supported by PAPIIT-DGAPA, UNAM (Grant number IN203526 to F. García-Oliva). We thank Rodrigo Velázquez-Durán for technical assistance in biogeochemical analyses at IIES, UNAM. We express our gratitude to Genesis 4C and Grupo ECCO for permission to sample the Red Ponds in Los Hundidos reserve. We also thank the members of our research team for insightful discussions, and Africa Islas-Robles for invaluable technical support from core subsampling to DNA isolation.

## Supplementary Figures

**Supplementary Figure 1.**
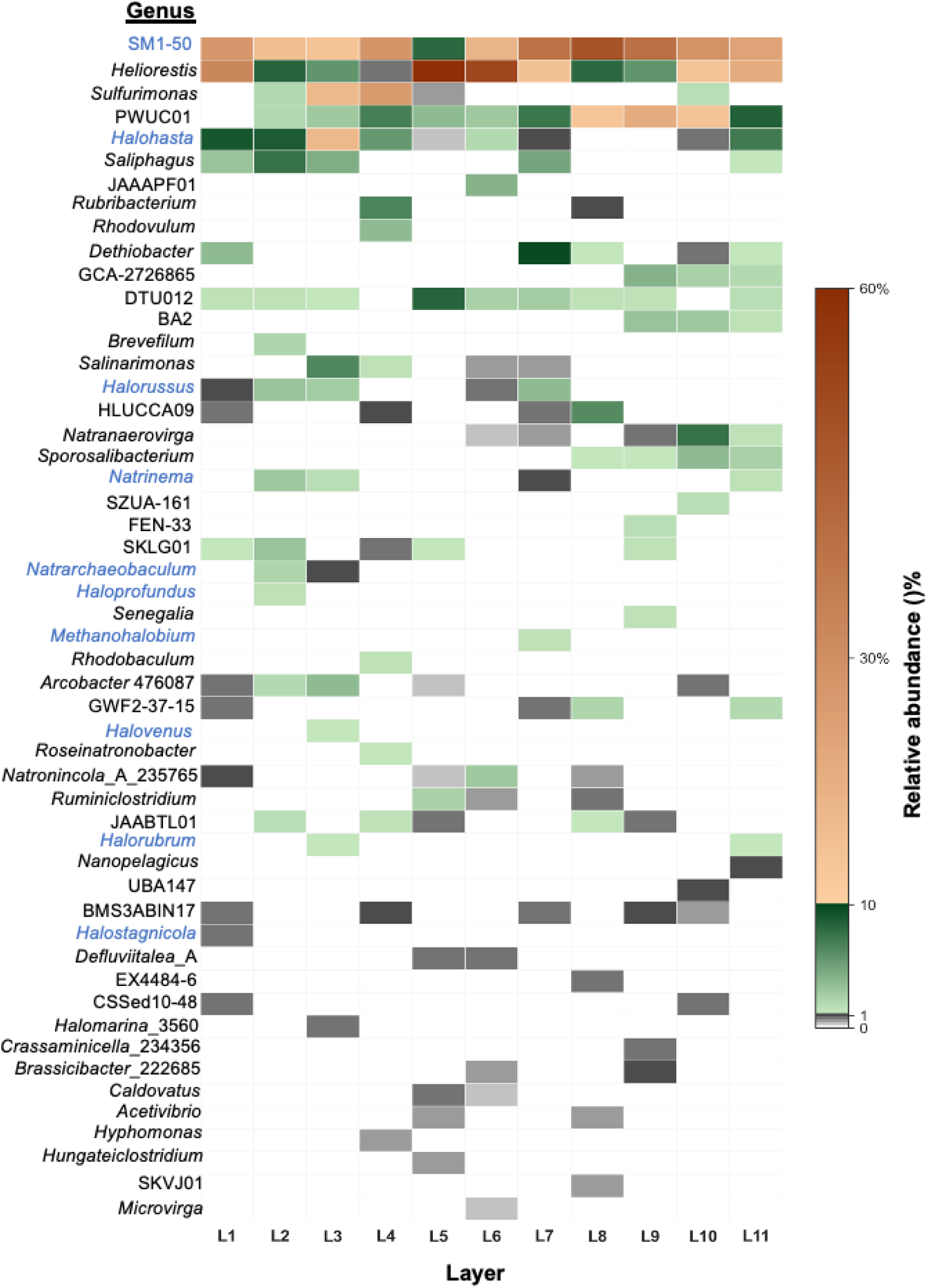
Relative abundance patterns of major microbial genera across sediment layers. The heatmap shows the relative abundance (%) of microbial genera across layers L1–L11. The piecewise color scale resolves three abundance ranges: gray for low-abundance or rare taxa (0–1%), green for intermediate abundance (1–10%), and orange-brown for highly abundant taxa (10–60%). Archaeal lineages are identified in blue.

**Supplementary Figure 2.**
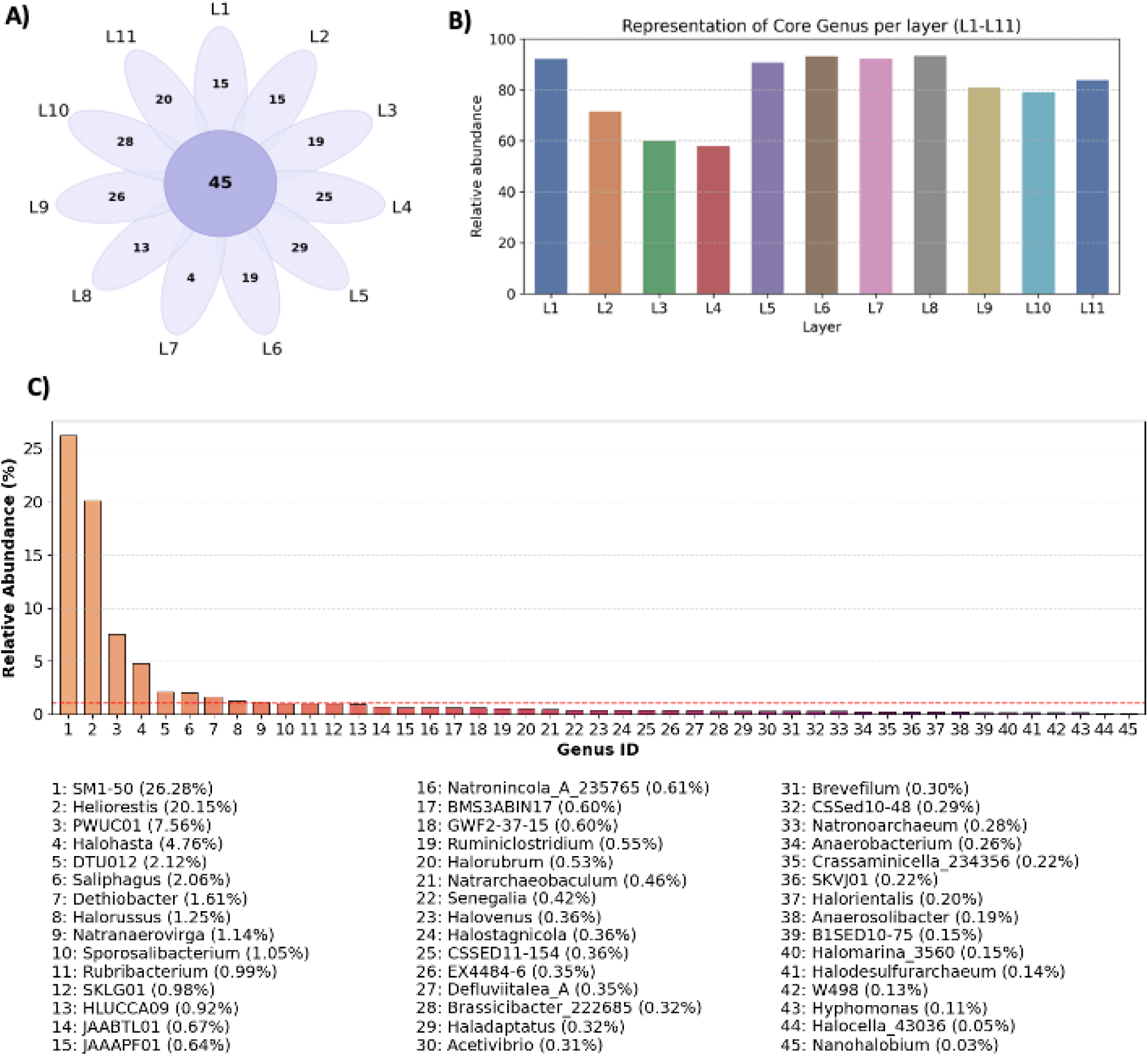
Distribution and abundance of core genera across stratified sediment layers. A) Flower diagram showing the number of genera unique to each layer in the petals and the number of genera shared across all layers in the center. B) Relative contribution of the shared core microbiome within each layer. C) Relative abundance of the 45 core genera across the sediment profile; the red dashed line marks 1% relative abundance.

**Supplementary Figure 3.**
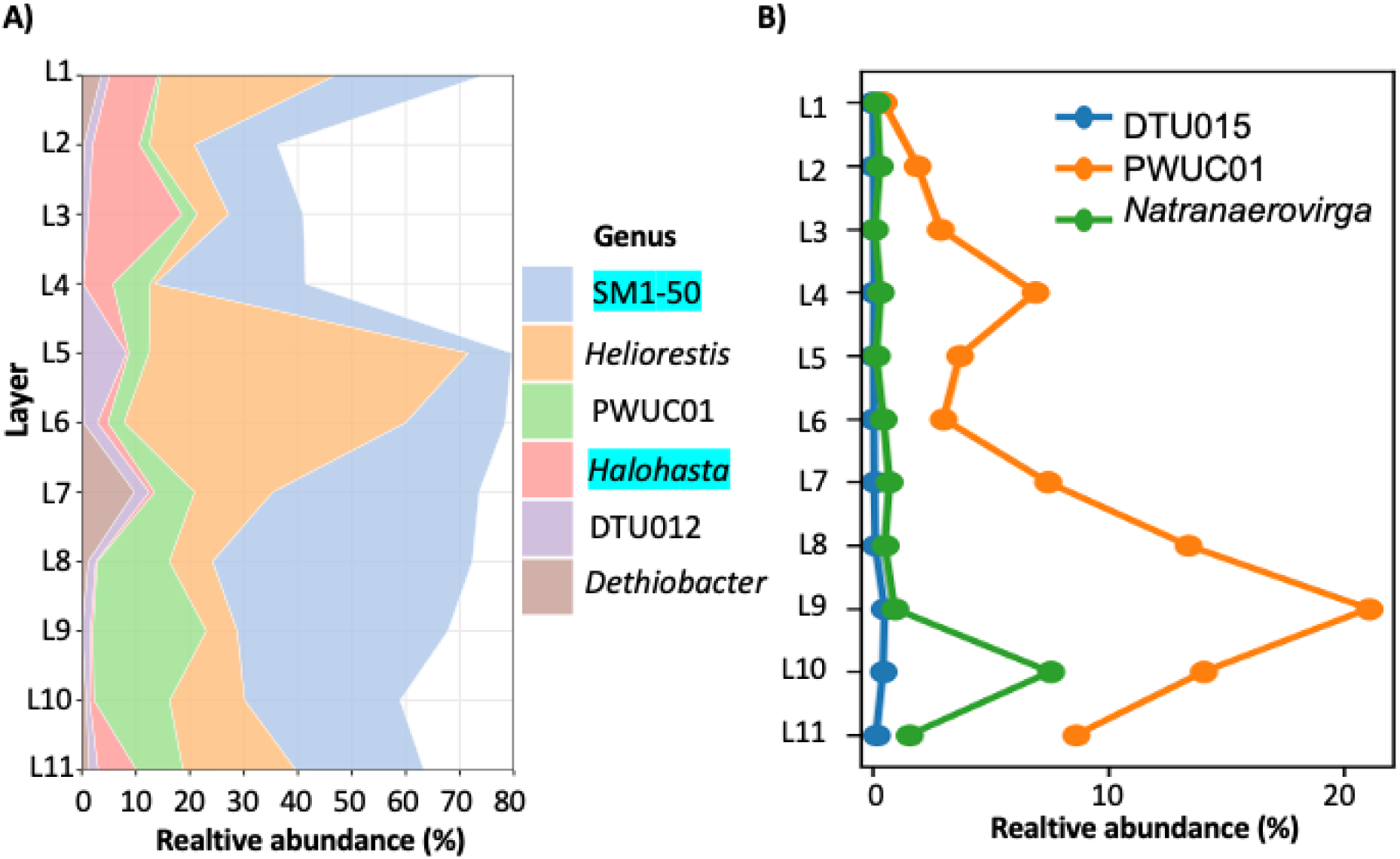
Distribution patterns of dominant core genera across sediment layers. A) Down-core dominance patterns of six core genera; asterisks denote archaeal lineages. B) Bacterial genera whose relative abundance increases significantly with depth (Spearman correlation, FDR ≤ 0.05).

**Supplementary Figure 4.**
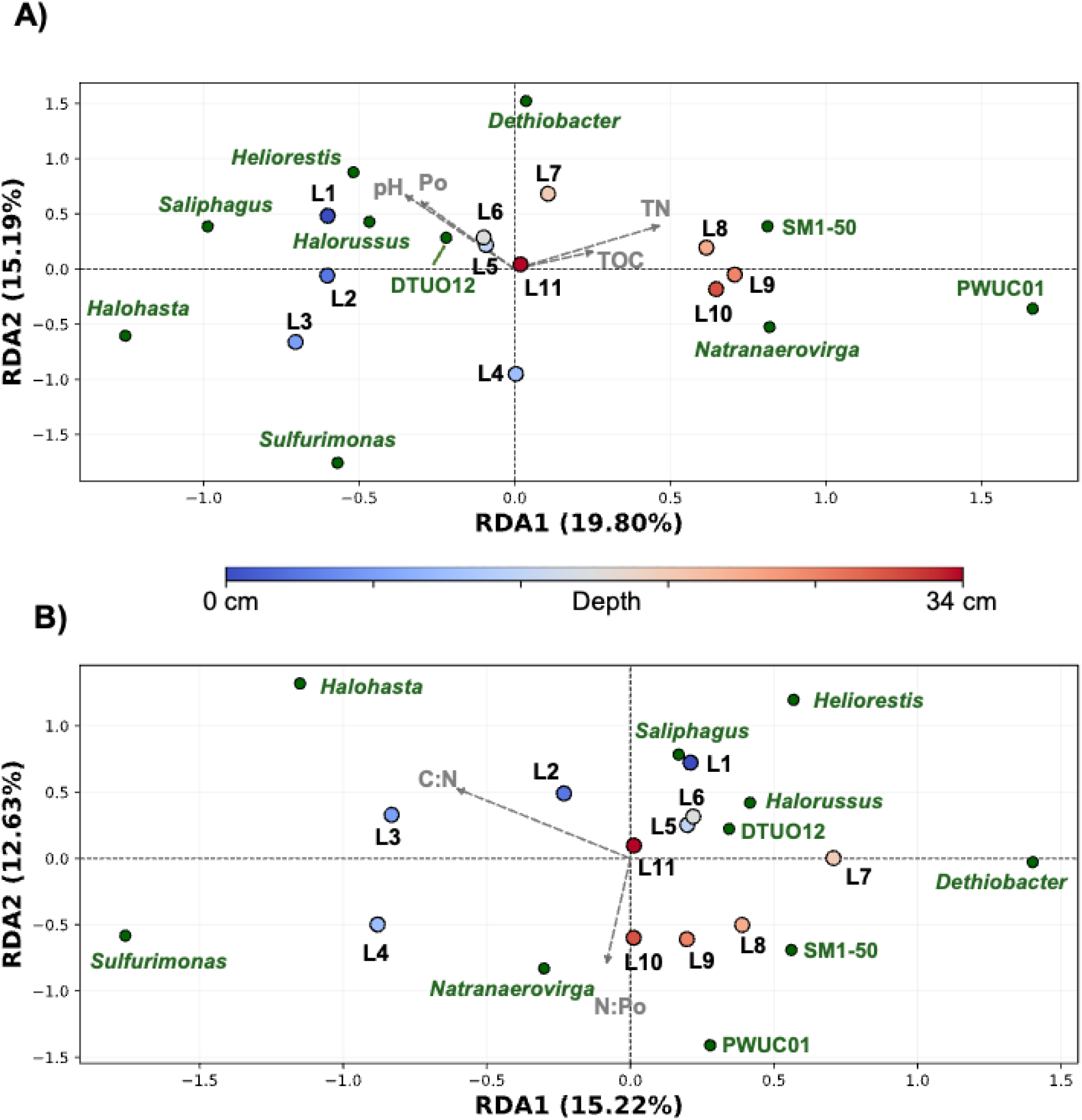
Redundancy analysis (RDA) of microbial community stratification and geochemical gradients across layers L1–L11. Biological community structure was assessed based on the centered log-ratio (CLR) transformed abundances of the top 10 most abundant microbial taxa. A) Model A (Absolute Nutrients): RDA constrained by pH and natural-log transformed absolute nutrient concentrations (TOC, TN, and Po). Model A accounted for 34.99% of the total compositional variance across the first two axes (RDA1 = 19.80%, RDA2 = 15.19%; R2 adjusted = 0.195, pseudo-F = 1.60, p = 0.12). Type III marginal permutation tests revealed that ln TN showed the strongest individual association with microbial turnover (F = 2.85, p = 0.059, FDR = 0.17), followed by pH (F = 2.36, p = 0.085, FDR = 0.17). B). Model B (Stoichiometry ratio C:N, C:Po, N:Po): RDA constrained by pH and log-transformed stoichiometric ratios (ln C:N, ln C:Po and ln N:Po). Model B explained 27.85% of total variance along the primary axes (RDA1 = 15.22%, RDA2 = 12.63%; overall model R2 adjusted = 0.099, pseudo-F = 1.55, p = 0.155). Marginal tests indicated a significant standalone effect of ln C:N (F = 2.9149, p = 0.0483, FDR = 0.1449). Vectors shown solely for exploratory illustration of trends (p > 0.05).

