## Supplementary Tables 1 to 11 for "Depth-Resolved Microbial Community Structure and Phosphorus Deficiency Across a 34-cm Mat–Sediment Core in Poza Roja 2, Cuatrociénegas"

**Supplementary Table 1. Physicochemical characterization of Poza Roja 2 water from Cuatro Ciénegas.**

| Parameter | Units | Result | Category / Remark (Fertilab) |
|---|---|---|---|
| **Salinity and sodicity** | | | |
| Electrical conductivity (EC) | dS/m | 11.6 | Very high |
| pH | — | 8.13 | Alkaline |
| Sodium Adsorption Ratio (SAR) | — | 13.5 | Very high |
| Adjusted SAR (SARadj) | — | 17.0 | Very high |
| Hardness | °f | 220.49 | Hard water (≈2,216 mg $CaCO_3$eq/L) |
| **Cations** | | | |
| Calcium (Ca) | mg/L (meq/L) | 555 (27.7) | — |
| Magnesium (Mg) | mg/L (meq/L) | 202 (16.6) | — |
| Sodium (Na) | mg/L (meq/L) | 1,463 (63.6) | — |
| Potassium (K) | mg/L (meq/L) | 114 (2.91)* | *Below the quantification limit (LC=145 mg/L); detected but not reliably quantifiable |
| Sum of cations | meq/L | 111 | — |
| **Anions** | | | |
| Sulfate (S-$SO_4$) | mg/L (meq/L) | 1,449 (90.4) | Very high |
| Bicarbonate ($HCO_3$) | mg/L (meq/L) | 230 (3.77) | Medium |
| Chloride (Cl) | mg/L (meq/L) | 984 (28.1) | Very high |
| Carbonate ($CO_3$) | mg/L (meq/L) | 41.7 (1.39) | Moderate |
| Nitrate (N-$NO_3$) | mg/L (meq/L) | 0.14 (0.01) | Very low |
| Sum of anions | meq/L | 124 | — |
| **Special determinations / micronutrients** | | | |
| Boron (B) | mg/L | 1.58 | Very high (potentially toxic) |
| Iron (Fe) | mg/L | 0.0210† | †Below the limit of detection (LD=0.062 mg/L); not reliably detected |
| Manganese (Mn) | mg/L | 0.0010† | †Below the LD (0.003 mg/L); not reliably detected |
| Copper (Cu) | mg/L | 0.1350 | Mod. high (>0.5 ppm would be toxic for crops) |
| Zinc (Zn) | mg/L | 0.0010† | †Below the LD (0.016 mg/L); not reliably detected |
| Arsenic (As) | mg/L | 0.0410* | *Between LD (0.009) and LC (0.050 mg/L); detected but not reliably quantifiable |
| **Heavy metals (subcontracted package, EPA 6010C 2007)** | | | |
| Arsenic (As) | mg/L | < 0.05 | Below quantification limit |
| **Other parameters** | | | |

| Parameter | Units | Result | Category / Remark (Fertilab) |
| --- | --- | --- | --- |
| Ammonium ($N\text{-}NH_4$) | mg/L | NA | Not analyzed |
| Phosphate ($P\text{-}PO_4$) | mg/L | < 0.091‡ | ‡Reported as "0" by Fertilab; below the limit of detection (LD=0.091 mg/L) — not a measured zero, but non-detect |
| Total solids | mg/L | NA | Not analyzed |
| Dissolved solids | mg/L | NA | Not analyzed |
| Nickel (Ni) | mg/L | NA | Not analyzed |
| Molybdenum (Mo) | mg/L | NA | Not analyzed |
| Fluoride | mg/L | NA | Not analyzed |

**Supplementary Table 2. Geochemical parameters and nutrient composition across sediment layers**

| Layer | TC % | TIC % | TOC % | TC (mg) | TIC (mg) | TOC (mg) | TN (mg/g) | TP (mg/g) | Po (mg/g) | Pi (mg/g) | C/Po | N/Po | C/N | pH |
|---|---|---|---|---|---|---|---|---|---|---|---|---|---|---|
| L1 | 5.98 | 1.42 | 4.55 | 59.76 | 14.21 | 45.55 | 10.11 | 0.48 | 0.16 | 0.32 | 285 | 63 | 5 | 9.54 |
| L2 | 4.62 | 2.10 | 2.53 | 46.23 | 20.96 | 25.27 | 6.90 | 0.35 | 0.03 | 0.32 | 842 | 230 | 4 | 9.33 |
| L3 | 4.08 | 1.83 | 2.25 | 40.82 | 18.31 | 22.50 | 3.73 | 0.23 | 0.12 | 0.11 | 188 | 31 | 6 | 9.23 |
| L4 | 5.73 | 1.74 | 3.98 | 57.26 | 17.42 | 39.84 | 12.70 | 0.44 | 0.02 | 0.42 | 1,992 | 635 | 3 | 9.36 |
| L5 | 4.69 | 1.88 | 2.81 | 46.86 | 18.79 | 28.08 | 7.24 | 0.41 | 0.04 | 0.37 | 702 | 181 | 4 | 9.37 |
| L6 | 4.69 | 2.00 | 2.68 | 46.86 | 20.03 | 26.83 | 7.65 | 0.39 | 0.10 | 0.29 | 268 | 77 | 4 | 9.41 |
| L7 | 4.90 | 1.76 | 3.14 | 48.97 | 17.61 | 31.36 | 13.81 | 0.40 | 0.10 | 0.30 | 314 | 138 | 2 | 9.73 |
| L8 | 5.23 | 1.12 | 4.11 | 52.30 | 11.18 | 41.12 | 14.24 | 0.29 | 0.07 | 0.22 | 587 | 203 | 3 | 9.32 |
| L9 | 4.27 | 1.72 | 2.55 | 42.70 | 17.16 | 25.54 | 6.80 | 0.21 | 0.05 | 0.16 | 511 | 136 | 4 | 9.09 |
| L10 | 5.22 | 1.43 | 3.80 | 52.23 | 14.26 | 37.98 | 9.95 | 0.23 | 0.03 | 0.20 | 1,266 | 332 | 4 | 9.13 |
| L11 | 5.57 | 1.93 | 3.64 | 55.73 | 19.29 | 36.44 | 9.27 | 0.23 | 0.12 | 0.11 | 304 | 77 | 4 | 9.25 |

**Supplementary Table 3. Genus-Level Taxonomic Classification and Relative Abundance of ASV's Using the GreenGenes2 v2 Database and Microbial Dark Matter Framework assignment.**

| Genus | L1 | L2 | L3 | L4 | L5 | L6 | L7 | L8 | L9 | L10 | L11 | Microbial_matter |
|---|---|---|---|---|---|---|---|---|---|---|---|---|
| c__ABY1 | 0.00E+00 | 0.00E+00 | 0.00E+00 | 0.00E+00 | 0.00E+00 | 0.00E+00 | 0.00E+00 | 0.00E+00 | 0.00E+00 | 0.00E+00 | 3.24E-05 | Known_Unknown |
| c__Acidimicrobiia_401430 | 0.00E+00 | 6.98E-04 | 0.00E+00 | 5.72E-05 | 1.91E-04 | 6.19E-04 | 1.07E-04 | 0.00E+00 | 1.19E-04 | 0.00E+00 | 3.57E-04 | Known_Unknown |
| c__Actinomycetia | 0.00E+00 | 1.04E-03 | 7.51E-04 | 8.01E-05 | 7.50E-04 | 3.04E-04 | 3.58E-05 | 0.00E+00 | 1.10E-03 | 1.62E-03 | 2.59E-04 | Known_Unknown |
| c__Alphaproteobacteria | 8.40E-03 | 1.87E-02 | 9.40E-03 | 1.35E-02 | 8.34E-03 | 2.16E-02 | 4.62E-03 | 8.28E-03 | 4.32E-03 | 9.16E-03 | 1.03E-02 | Known_Unknown |
| c__Anaerolineae | 3.97E-04 | 1.92E-03 | 0.00E+00 | 2.63E-04 | 4.09E-04 | 0.00E+00 | 0.00E+00 | 3.99E-04 | 2.74E-03 | 2.79E-03 | 1.31E-03 | Known_Unknown |
| c__Bacilli | 6.07E-04 | 1.45E-03 | 2.11E-03 | 2.75E-04 | 1.79E-02 | 2.70E-03 | 1.67E-03 | 5.54E-04 | 1.37E-03 | 4.39E-03 | 1.72E-03 | Known_Unknown |
| c__Bacteroidia | 0.00E+00 | 0.00E+00 | 2.35E-05 | 1.60E-04 | 8.19E-05 | 4.67E-05 | 7.16E-05 | 9.01E-05 | 0.00E+00 | 0.00E+00 | 0.00E+00 | Known_Unknown |
| c__Bradymonadia | 0.00E+00 | 0.00E+00 | 0.00E+00 | 0.00E+00 | 0.00E+00 | 0.00E+00 | 0.00E+00 | 0.00E+00 | 0.00E+00 | 7.29E-05 | 0.00E+00 | Known_Unknown |
| c__Chloroflexia | 0.00E+00 | 1.70E-04 | 0.00E+00 | 0.00E+00 | 0.00E+00 | 0.00E+00 | 0.00E+00 | 0.00E+00 | 5.93E-05 | 3.64E-05 | 0.00E+00 | Known_Unknown |
| c__Clostridia_258483 | 1.19E-03 | 2.18E-03 | 2.34E-03 | 2.39E-03 | 3.96E-03 | 3.28E-03 | 1.81E-03 | 4.64E-03 | 4.53E-03 | 7.45E-03 | 4.78E-03 | Known_Unknown |
| c__Clostridia_258486 | 9.33E-05 | 1.36E-04 | 0.00E+00 | 8.01E-05 | 4.91E-04 | 4.67E-05 | 1.07E-04 | 1.89E-03 | 6.97E-04 | 1.99E-03 | 2.74E-03 | Known_Unknown |
| c__Cyanobacteriia | 2.57E-04 | 6.81E-05 | 0.00E+00 | 5.15E-04 | 3.27E-04 | 0.00E+00 | 0.00E+00 | 0.00E+00 | 0.00E+00 | 0.00E+00 | 2.11E-04 | Known_Unknown |
| c__Dehalococcoidia | 1.40E-04 | 1.87E-04 | 1.88E-04 | 4.01E-04 | 5.18E-04 | 0.00E+00 | 0.00E+00 | 1.03E-04 | 2.97E-04 | 1.46E-04 | 2.11E-04 | Known_Unknown |
| c__Desulfobacteria | 0.00E+00 | 3.41E-05 | 0.00E+00 | 0.00E+00 | 0.00E+00 | 0.00E+00 | 0.00E+00 | 0.00E+00 | 1.48E-05 | 0.00E+00 | 0.00E+00 | Known_Unknown |
| c__Dethiobacteria | 1.17E-04 | 0.00E+00 | 0.00E+00 | 0.00E+00 | 0.00E+00 | 0.00E+00 | 0.00E+00 | 0.00E+00 | 0.00E+00 | 0.00E+00 | 0.00E+00 | Known_Unknown |
| c__Fusobacteriia | 0.00E+00 | 0.00E+00 | 0.00E+00 | 0.00E+00 | 0.00E+00 | 0.00E+00 | 0.00E+00 | 0.00E+00 | 0.00E+00 | 0.00E+00 | 6.48E-05 | Known_Unknown |
| c__Gammaproteobacteria | 1.87E-04 | 1.87E-04 | 9.39E-05 | 7.67E-04 | 4.09E-05 | 0.00E+00 | 0.00E+00 | 2.06E-04 | 3.71E-04 | 2.73E-04 | 1.46E-04 | Known_Unknown |
| c__Gemmatimonadetes | 0.00E+00 | 0.00E+00 | 0.00E+00 | 0.00E+00 | 0.00E+00 | 0.00E+00 | 5.37E-05 | 0.00E+00 | 0.00E+00 | 0.00E+00 | 0.00E+00 | Known_Unknown |
| c__Iainarchaeia | 0.00E+00 | 0.00E+00 | 0.00E+00 | 0.00E+00 | 0.00E+00 | 0.00E+00 | 0.00E+00 | 0.00E+00 | 4.45E-05 | 0.00E+00 | 0.00E+00 | Known_Unknown |
| c__Lentisphaeria | 0.00E+00 | 0.00E+00 | 0.00E+00 | 0.00E+00 | 0.00E+00 | 0.00E+00 | 0.00E+00 | 0.00E+00 | 0.00E+00 | 0.00E+00 | 8.10E-05 | Known_Unknown |
| c__Limnochordia | 0.00E+00 | 8.52E-05 | 2.93E-04 | 0.00E+00 | 1.09E-04 | 0.00E+00 | 0.00E+00 | 0.00E+00 | 0.00E+00 | 0.00E+00 | 0.00E+00 | Known_Unknown |
| c__Nanoarchaeia | 1.40E-04 | 8.52E-05 | 0.00E+00 | 1.49E-04 | 0.00E+00 | 0.00E+00 | 0.00E+00 | 0.00E+00 | 1.78E-04 | 1.82E-04 | 4.70E-04 | Known_Unknown |
| c__Paceibacteria | 0.00E+00 | 3.41E-05 | 0.00E+00 | 5.72E-05 | 0.00E+00 | 0.00E+00 | 0.00E+00 | 1.04E-03 | 5.19E-04 | 2.00E-04 | 1.13E-04 | Known_Unknown |
| c__Phycisphaerae | 0.00E+00 | 0.00E+00 | 0.00E+00 | 0.00E+00 | 0.00E+00 | 9.34E-05 | 0.00E+00 | 0.00E+00 | 0.00E+00 | 0.00E+00 | 0.00E+00 | Known_Unknown |
| c__Spirochaetia | 7.00E-05 | 6.81E-05 | 3.52E-05 | 5.72E-05 | 0.00E+00 | 2.34E-05 | 3.58E-05 | 0.00E+00 | 0.00E+00 | 1.82E-04 | 0.00E+00 | Known_Unknown |
| c__Thermoleophilia | 0.00E+00 | 0.00E+00 | 0.00E+00 | 3.43E-05 | 0.00E+00 | 0.00E+00 | 0.00E+00 | 0.00E+00 | 0.00E+00 | 0.00E+00 | 0.00E+00 | Known_Unknown |
| c__Thermoplasmata_1773 | 2.80E-04 | 2.21E-04 | 0.00E+00 | 0.00E+00 | 2.73E-05 | 0.00E+00 | 0.00E+00 | 1.03E-04 | 0.00E+00 | 1.27E-04 | 9.72E-05 | Known_Unknown |
| c__UBA5829 | 0.00E+00 | 0.00E+00 | 0.00E+00 | 0.00E+00 | 0.00E+00 | 0.00E+00 | 0.00E+00 | 0.00E+00 | 0.00E+00 | 9.11E-05 | 0.00E+00 | Known_Unknown |
| c__Vampirovibrionia | 0.00E+00 | 0.00E+00 | 0.00E+00 | 0.00E+00 | 0.00E+00 | 0.00E+00 | 0.00E+00 | 6.44E-05 | 0.00E+00 | 0.00E+00 | 0.00E+00 | Known_Unknown |
| d__Archaea | 3.76E-03 | 6.83E-03 | 1.46E-03 | 2.52E-04 | 4.09E-05 | 4.20E-04 | 3.04E-04 | 4.51E-04 | 1.99E-02 | 7.90E-03 | 5.01E-03 | Known_Unknown |
| d__Bacteria | 6.03E-02 | 4.20E-02 | 1.95E-02 | 1.79E-02 | 7.37E-02 | 1.73E-02 | 3.01E-02 | 3.79E-02 | 4.75E-02 | 4.36E-02 | 5.70E-02 | Known_Unknown |
| f__AB-539-J10 | 0.00E+00 | 0.00E+00 | 0.00E+00 | 0.00E+00 | 0.00E+00 | 0.00E+00 | 0.00E+00 | 1.80E-04 | 9.05E-04 | 6.56E-04 | 2.43E-04 | Known_Unknown |
| f__Acetivibrionaceae | 1.17E-04 | 1.06E-03 | 4.81E-04 | 0.00E+00 | 1.46E-03 | 1.28E-03 | 1.97E-04 | 0.00E+00 | 3.12E-04 | 7.29E-05 | 1.13E-04 | Known_Unknown |
| f__Acetivibrionaceae | 2.10E-04 | 3.07E-04 | 2.11E-04 | 2.06E-04 | 1.09E-04 | 1.75E-04 | 1.43E-04 | 2.70E-04 | 5.79E-04 | 1.13E-03 | 2.27E-04 | Known_Unknown |
| f__Acetobacteraceae | 0.00E+00 | 1.60E-03 | 3.76E-04 | 8.01E-05 | 3.18E-03 | 2.14E-03 | 0.00E+00 | 0.00E+00 | 5.79E-04 | 2.55E-04 | 0.00E+00 | Known_Unknown |
| f__Acidobacteriaceae | 0.00E+00 | 0.00E+00 | 0.00E+00 | 0.00E+00 | 0.00E+00 | 0.00E+00 | 0.00E+00 | 0.00E+00 | 0.00E+00 | 0.00E+00 | 3.24E-05 | Known_Unknown |
| f__Acutalibacteraceae | 0.00E+00 | 0.00E+00 | 0.00E+00 | 3.43E-05 | 0.00E+00 | 0.00E+00 | 0.00E+00 | 0.00E+00 | 0.00E+00 | 0.00E+00 | 0.00E+00 | Known_Unknown |
| f__Aminicenantaceae | 0.00E+00 | 0.00E+00 | 0.00E+00 | 0.00E+00 | 0.00E+00 | 0.00E+00 | 0.00E+00 | 0.00E+00 | 0.00E+00 | 0.00E+00 | 4.86E-05 | Known_Unknown |
| f__Aminicenantaceae | 0.00E+00 | 1.87E-04 | 0.00E+00 | 0.00E+00 | 0.00E+00 | 0.00E+00 | 0.00E+00 | 1.29E-05 | 8.90E-05 | 1.40E-03 | 5.19E-04 | Known_Unknown |
| f__Amorphaceae | 0.00E+00 | 8.52E-05 | 0.00E+00 | 2.98E-04 | 0.00E+00 | 0.00E+00 | 0.00E+00 | 0.00E+00 | 0.00E+00 | 0.00E+00 | 0.00E+00 | Known_Unknown |
| f__Amphibacillaceae | 7.00E-05 | 0.00E+00 | 9.39E-05 | 0.00E+00 | 3.41E-04 | 2.10E-04 | 5.37E-05 | 0.00E+00 | 1.48E-05 | 0.00E+00 | 3.24E-05 | Known_Unknown |
| f__Anaerolineaceae_409200 | 4.67E-05 | 2.21E-04 | 0.00E+00 | 0.00E+00 | 0.00E+00 | 0.00E+00 | 0.00E+00 | 7.73E-05 | 7.42E-05 | 0.00E+00 | 2.76E-04 | Known_Unknown |
| f__Anaerolineaceae_409201 | 0.00E+00 | 6.98E-04 | 0.00E+00 | 5.49E-04 | 2.59E-04 | 0.00E+00 | 6.27E-04 | 3.73E-04 | 1.34E-04 | 0.00E+00 | 6.48E-05 | Known_Unknown |
| f__Anaeromyxobacteraceae | 0.00E+00 | 0.00E+00 | 0.00E+00 | 0.00E+00 | 0.00E+00 | 0.00E+00 | 7.16E-05 | 0.00E+00 | 0.00E+00 | 0.00E+00 | 0.00E+00 | Known_Unknown |
| f__Anaerovoracaceae | 0.00E+00 | 0.00E+00 | 0.00E+00 | 0.00E+00 | 8.19E-05 | 0.00E+00 | 0.00E+00 | 0.00E+00 | 0.00E+00 | 0.00E+00 | 8.10E-05 | Known_Unknown |

| Genus | L1 | L2 | L3 | L4 | L5 | L6 | L7 | L8 | L9 | L10 | L11 | Microbial_matter |
|---|---|---|---|---|---|---|---|---|---|---|---|---|
| f__Arcobacteraceae | 0.00E+00 | 0.00E+00 | 3.52E-05 | 0.00E+00 | 0.00E+00 | 0.00E+00 | 0.00E+00 | 0.00E+00 | 0.00E+00 | 4.41E-03 | 2.98E-03 | Known_Unknown |
| f__Azospirillaceae_507917 | 1.40E-04 | 2.38E-04 | 9.39E-05 | 0.00E+00 | 9.55E-05 | 0.00E+00 | 0.00E+00 | 0.00E+00 | 0.00E+00 | 0.00E+00 | 0.00E+00 | Known_Unknown |
| f__Bacillaceae_H_294103 | 0.00E+00 | 0.00E+00 | 0.00E+00 | 0.00E+00 | 0.00E+00 | 0.00E+00 | 0.00E+00 | 0.00E+00 | 0.00E+00 | 0.00E+00 | 6.48E-05 | Known_Unknown |
| f__Beijerinckiaceae | 0.00E+00 | 2.45E-03 | 1.27E-03 | 0.00E+00 | 1.60E-03 | 2.30E-03 | 0.00E+00 | 0.00E+00 | 0.00E+00 | 0.00E+00 | 3.73E-04 | Known_Unknown |
| f__Borkfalkiaceae | 0.00E+00 | 0.00E+00 | 0.00E+00 | 8.01E-05 | 0.00E+00 | 3.50E-05 | 0.00E+00 | 0.00E+00 | 0.00E+00 | 0.00E+00 | 0.00E+00 | Known_Unknown |
| f__Bryobacteraceae | 0.00E+00 | 0.00E+00 | 0.00E+00 | 0.00E+00 | 0.00E+00 | 0.00E+00 | 0.00E+00 | 0.00E+00 | 4.45E-05 | 0.00E+00 | 0.00E+00 | Known_Unknown |
| f__C00003060 | 0.00E+00 | 0.00E+00 | 0.00E+00 | 0.00E+00 | 0.00E+00 | 0.00E+00 | 0.00E+00 | 0.00E+00 | 0.00E+00 | 1.82E-05 | 0.00E+00 | Known_Unknown |
| f__CAG-138 | 0.00E+00 | 0.00E+00 | 0.00E+00 | 0.00E+00 | 5.46E-05 | 0.00E+00 | 0.00E+00 | 0.00E+00 | 0.00E+00 | 0.00E+00 | 0.00E+00 | Known_Unknown |
| f__CAG-917 | 0.00E+00 | 0.00E+00 | 0.00E+00 | 0.00E+00 | 0.00E+00 | 0.00E+00 | 0.00E+00 | 0.00E+00 | 0.00E+00 | 0.00E+00 | 1.13E-04 | Known_Unknown |
| f__Caldilineaceae | 0.00E+00 | 5.11E-05 | 0.00E+00 | 1.60E-04 | 0.00E+00 | 0.00E+00 | 0.00E+00 | 1.16E-04 | 6.97E-04 | 0.00E+00 | 0.00E+00 | Known_Unknown |
| f__Caloramatoraceae | 0.00E+00 | 1.67E-03 | 8.33E-04 | 2.29E-04 | 1.91E-04 | 8.17E-05 | 0.00E+00 | 0.00E+00 | 0.00E+00 | 0.00E+00 | 1.30E-04 | Known_Unknown |
| f__Caulobacteraceae | 0.00E+00 | 0.00E+00 | 1.76E-04 | 4.58E-05 | 9.55E-05 | 9.34E-05 | 0.00E+00 | 3.86E-05 | 0.00E+00 | 0.00E+00 | 0.00E+00 | Known_Unknown |
| f__Cellulosilyticaceae | 0.00E+00 | 0.00E+00 | 0.00E+00 | 1.14E-05 | 0.00E+00 | 0.00E+00 | 0.00E+00 | 0.00E+00 | 1.48E-05 | 0.00E+00 | 0.00E+00 | Known_Unknown |
| f__CG2-30-64-16 | 9.33E-05 | 0.00E+00 | 0.00E+00 | 0.00E+00 | 0.00E+00 | 0.00E+00 | 0.00E+00 | 0.00E+00 | 4.45E-05 | 0.00E+00 | 0.00E+00 | Known_Unknown |
| f__CG2-30-64-17 | 0.00E+00 | 4.43E-04 | 0.00E+00 | 0.00E+00 | 0.00E+00 | 0.00E+00 | 0.00E+00 | 6.44E-04 | 3.56E-03 | 7.50E-03 | 1.49E-03 | Known_Unknown |
| f__Chitinispirillaceae | 0.00E+00 | 0.00E+00 | 0.00E+00 | 0.00E+00 | 0.00E+00 | 0.00E+00 | 0.00E+00 | 0.00E+00 | 0.00E+00 | 0.00E+00 | 6.48E-05 | Known_Unknown |
| f__Chlorobiaceae | 0.00E+00 | 0.00E+00 | 0.00E+00 | 3.43E-05 | 4.09E-05 | 0.00E+00 | 0.00E+00 | 0.00E+00 | 0.00E+00 | 0.00E+00 | 0.00E+00 | Known_Unknown |
| f__Chloroflexaceae | 0.00E+00 | 1.02E-04 | 0.00E+00 | 2.29E-04 | 1.91E-04 | 0.00E+00 | 4.12E-04 | 3.09E-04 | 1.90E-03 | 5.28E-04 | 4.05E-04 | Known_Unknown |
| f__Chroococcidiopsidaceae_29158 | 0.00E+00 | 0.00E+00 | 0.00E+00 | 0.00E+00 | 1.36E-04 | 4.67E-05 | 0.00E+00 | 0.00E+00 | 0.00E+00 | 0.00E+00 | 6.48E-05 | Known_Unknown |
| f__Clostridiaceae_222000 | 0.00E+00 | 5.94E-03 | 2.98E-03 | 1.16E-03 | 3.74E-03 | 2.05E-03 | 0.00E+00 | 3.99E-04 | 3.15E-03 | 8.74E-04 | 4.05E-04 | Known_Unknown |
| f__Clostridiaceae_222001 | 9.10E-04 | 3.27E-03 | 2.23E-03 | 7.33E-04 | 2.51E-03 | 2.49E-03 | 2.51E-04 | 9.53E-04 | 2.42E-03 | 1.46E-03 | 1.67E-03 | Known_Unknown |
| f__Cyanobiaceae | 1.17E-03 | 2.73E-04 | 7.28E-04 | 4.46E-04 | 1.64E-04 | 0.00E+00 | 0.00E+00 | 0.00E+00 | 0.00E+00 | 0.00E+00 | 2.27E-04 | Known_Unknown |
| f__Dermatophilaceae_390796 | 0.00E+00 | 3.41E-05 | 0.00E+00 | 0.00E+00 | 0.00E+00 | 0.00E+00 | 0.00E+00 | 0.00E+00 | 0.00E+00 | 0.00E+00 | 0.00E+00 | Known_Unknown |
| f__Desulfallaceae | 0.00E+00 | 0.00E+00 | 2.58E-04 | 0.00E+00 | 0.00E+00 | 0.00E+00 | 0.00E+00 | 9.01E-05 | 0.00E+00 | 0.00E+00 | 4.86E-05 | Known_Unknown |
| f__Desulfatiglandaceae | 7.00E-05 | 6.81E-05 | 1.17E-05 | 0.00E+00 | 4.09E-05 | 7.01E-05 | 0.00E+00 | 2.19E-04 | 2.08E-04 | 1.17E-03 | 3.24E-05 | Known_Unknown |
| f__Desulfohalotomaculaceae | 0.00E+00 | 2.73E-04 | 3.64E-04 | 0.00E+00 | 3.07E-03 | 2.04E-03 | 2.51E-04 | 8.76E-04 | 4.30E-04 | 0.00E+00 | 4.05E-04 | Known_Unknown |
| f__Desulfosarcinaceae | 0.00E+00 | 0.00E+00 | 0.00E+00 | 0.00E+00 | 0.00E+00 | 0.00E+00 | 0.00E+00 | 0.00E+00 | 1.48E-05 | 3.64E-05 | 0.00E+00 | Known_Unknown |
| f__Desulfovibrionaceae | 0.00E+00 | 0.00E+00 | 0.00E+00 | 0.00E+00 | 0.00E+00 | 0.00E+00 | 0.00E+00 | 0.00E+00 | 0.00E+00 | 1.27E-04 | 0.00E+00 | Known_Unknown |
| f__Desulfuromonadaceae_439376 | 0.00E+00 | 0.00E+00 | 0.00E+00 | 0.00E+00 | 0.00E+00 | 0.00E+00 | 0.00E+00 | 0.00E+00 | 0.00E+00 | 5.46E-05 | 0.00E+00 | Known_Unknown |
| f__Devosiaceae | 1.63E-04 | 7.32E-04 | 8.80E-04 | 1.37E-04 | 1.50E-04 | 1.05E-03 | 0.00E+00 | 0.00E+00 | 0.00E+00 | 0.00E+00 | 3.40E-04 | Known_Unknown |
| f__DSM-24629 | 0.00E+00 | 3.75E-04 | 1.88E-04 | 0.00E+00 | 2.59E-04 | 4.32E-04 | 5.37E-05 | 0.00E+00 | 0.00E+00 | 0.00E+00 | 0.00E+00 | Known_Unknown |
| f__DSM-24629 | 1.40E-04 | 0.00E+00 | 0.00E+00 | 1.60E-04 | 1.23E-04 | 1.09E-03 | 5.37E-05 | 1.03E-04 | 0.00E+00 | 1.82E-04 | 4.86E-05 | Known_Unknown |
| f__DSM-24630 | 0.00E+00 | 0.00E+00 | 0.00E+00 | 0.00E+00 | 0.00E+00 | 0.00E+00 | 0.00E+00 | 0.00E+00 | 1.44E-03 | 1.93E-03 | 0.00E+00 | Known_Unknown |
| f__EnvOPS12 | 0.00E+00 | 0.00E+00 | 0.00E+00 | 0.00E+00 | 0.00E+00 | 0.00E+00 | 0.00E+00 | 0.00E+00 | 0.00E+00 | 7.29E-05 | 0.00E+00 | Known_Unknown |
| f__EnvOPS13 | 0.00E+00 | 0.00E+00 | 0.00E+00 | 0.00E+00 | 0.00E+00 | 0.00E+00 | 0.00E+00 | 6.44E-05 | 1.90E-03 | 9.47E-04 | 1.78E-04 | Known_Unknown |
| f__Fastidiosipilaceae | 0.00E+00 | 0.00E+00 | 0.00E+00 | 0.00E+00 | 0.00E+00 | 3.50E-05 | 0.00E+00 | 0.00E+00 | 0.00E+00 | 0.00E+00 | 0.00E+00 | Known_Unknown |
| f__Fermentibacteraceae | 0.00E+00 | 0.00E+00 | 0.00E+00 | 0.00E+00 | 0.00E+00 | 0.00E+00 | 0.00E+00 | 0.00E+00 | 0.00E+00 | 0.00E+00 | 4.86E-05 | Known_Unknown |
| f__Gaiellaceae | 0.00E+00 | 0.00E+00 | 0.00E+00 | 0.00E+00 | 0.00E+00 | 0.00E+00 | 0.00E+00 | 0.00E+00 | 5.93E-05 | 0.00E+00 | 0.00E+00 | Known_Unknown |
| f__Geminicoccaceae | 0.00E+00 | 4.09E-04 | 0.00E+00 | 1.37E-04 | 1.11E-03 | 9.34E-04 | 1.25E-04 | 5.15E-05 | 4.01E-04 | 7.29E-05 | 1.62E-04 | Known_Unknown |
| f__Gemmataceae | 0.00E+00 | 0.00E+00 | 2.35E-05 | 0.00E+00 | 0.00E+00 | 0.00E+00 | 0.00E+00 | 0.00E+00 | 0.00E+00 | 0.00E+00 | 0.00E+00 | Known_Unknown |
| f__Geodermatophilaceae | 0.00E+00 | 3.41E-05 | 4.70E-05 | 0.00E+00 | 7.10E-04 | 7.01E-05 | 0.00E+00 | 0.00E+00 | 4.45E-05 | 1.27E-04 | 8.10E-05 | Known_Unknown |
| f__GWB1-42-6 | 0.00E+00 | 0.00E+00 | 0.00E+00 | 0.00E+00 | 0.00E+00 | 0.00E+00 | 0.00E+00 | 0.00E+00 | 0.00E+00 | 3.64E-05 | 0.00E+00 | Known_Unknown |
| f__Haladaptataceae | 0.00E+00 | 0.00E+00 | 1.17E-04 | 9.16E-05 | 0.00E+00 | 0.00E+00 | 0.00E+00 | 0.00E+00 | 0.00E+00 | 0.00E+00 | 0.00E+00 | Known_Unknown |
| f__Halanaerobiaceae_43120 | 1.87E-04 | 1.19E-04 | 2.35E-05 | 0.00E+00 | 5.46E-05 | 4.67E-05 | 1.07E-04 | 9.01E-05 | 1.48E-05 | 4.92E-04 | 5.19E-04 | Known_Unknown |
| f__Halieaceae | 2.33E-05 | 0.00E+00 | 0.00E+00 | 0.00E+00 | 0.00E+00 | 0.00E+00 | 0.00E+00 | 0.00E+00 | 0.00E+00 | 0.00E+00 | 0.00E+00 | Known_Unknown |
| f__Haloarculaceae_3992 | 2.33E-04 | 2.61E-03 | 5.14E-03 | 7.10E-04 | 1.50E-04 | 1.52E-04 | 1.97E-04 | 0.00E+00 | 0.00E+00 | 1.64E-04 | 3.40E-04 | Known_Unknown |
| f__Haloarculaceae_3993 | 0.00E+00 | 6.64E-04 | 1.58E-03 | 2.17E-04 | 0.00E+00 | 9.34E-05 | 0.00E+00 | 0.00E+00 | 0.00E+00 | 0.00E+00 | 0.00E+00 | Known_Unknown |

| Genus | L1 | L2 | L3 | L4 | L5 | L6 | L7 | L8 | L9 | L10 | L11 | Microbial_matter |
|---|---|---|---|---|---|---|---|---|---|---|---|---|
| f__Halobacillaceae | 4.67E-05 | 0.00E+00 | 2.35E-05 | 0.00E+00 | 9.55E-05 | 4.67E-05 | 0.00E+00 | 0.00E+00 | 0.00E+00 | 5.46E-05 | 0.00E+00 | Known_Unknown |
| f__Halobacteriaceae_4595 | 0.00E+00 | 1.48E-03 | 2.28E-03 | 0.00E+00 | 0.00E+00 | 0.00E+00 | 1.07E-04 | 0.00E+00 | 8.90E-05 | 4.01E-04 | 3.40E-04 | Known_Unknown |
| f__Haloferacaceae | 1.45E-03 | 9.76E-03 | 5.99E-03 | 2.17E-04 | 2.28E-03 | 1.67E-03 | 5.02E-04 | 2.70E-04 | 1.51E-03 | 1.68E-03 | 3.26E-03 | Known_Unknown |
| f__Halomonadaceae_641030 | 0.00E+00 | 2.21E-04 | 3.52E-05 | 0.00E+00 | 1.50E-04 | 0.00E+00 | 0.00E+00 | 0.00E+00 | 0.00E+00 | 2.00E-04 | 3.24E-05 | Known_Unknown |
| f__Heliobacteriaceae | 2.10E-04 | 5.45E-04 | 2.03E-03 | 2.17E-04 | 1.84E-03 | 8.17E-04 | 1.97E-04 | 4.12E-04 | 8.90E-05 | 1.91E-03 | 7.13E-04 | Known_Unknown |
| f__Hymenobacteraceae | 0.00E+00 | 0.00E+00 | 2.35E-05 | 0.00E+00 | 4.09E-05 | 0.00E+00 | 0.00E+00 | 0.00E+00 | 0.00E+00 | 0.00E+00 | 3.24E-05 | Known_Unknown |
| f__Hyphomicrobiaceae | 0.00E+00 | 0.00E+00 | 7.04E-05 | 1.03E-04 | 0.00E+00 | 0.00E+00 | 0.00E+00 | 0.00E+00 | 3.41E-04 | 3.10E-04 | 0.00E+00 | Known_Unknown |
| f__Hyphomonadaceae | 0.00E+00 | 0.00E+00 | 0.00E+00 | 3.89E-04 | 0.00E+00 | 0.00E+00 | 0.00E+00 | 3.99E-04 | 0.00E+00 | 0.00E+00 | 0.00E+00 | Known_Unknown |
| f__Ilumatobacteraceae | 0.00E+00 | 0.00E+00 | 0.00E+00 | 0.00E+00 | 0.00E+00 | 0.00E+00 | 0.00E+00 | 1.03E-04 | 0.00E+00 | 0.00E+00 | 4.86E-05 | Known_Unknown |
| f__Izemoplasmataceae | 0.00E+00 | 0.00E+00 | 0.00E+00 | 0.00E+00 | 0.00E+00 | 0.00E+00 | 0.00E+00 | 0.00E+00 | 0.00E+00 | 0.00E+00 | 1.62E-05 | Known_Unknown |
| f__Kiloniellaceae | 1.17E-04 | 1.36E-04 | 0.00E+00 | 0.00E+00 | 0.00E+00 | 0.00E+00 | 0.00E+00 | 2.45E-04 | 4.60E-04 | 2.73E-04 | 8.10E-05 | Known_Unknown |
| f__Kineosporiaceae | 0.00E+00 | 8.52E-05 | 0.00E+00 | 0.00E+00 | 4.09E-05 | 0.00E+00 | 0.00E+00 | 0.00E+00 | 0.00E+00 | 0.00E+00 | 0.00E+00 | Known_Unknown |
| f__Kosmotogaceae | 0.00E+00 | 8.52E-05 | 4.70E-05 | 0.00E+00 | 2.73E-05 | 0.00E+00 | 0.00E+00 | 0.00E+00 | 0.00E+00 | 0.00E+00 | 1.62E-05 | Known_Unknown |
| f__Lachnospiraceae | 3.73E-04 | 1.41E-03 | 1.51E-03 | 5.72E-05 | 1.34E-03 | 5.84E-04 | 0.00E+00 | 3.09E-04 | 5.19E-04 | 3.50E-03 | 6.97E-04 | Known_Unknown |
| f__LOWO2-01-FULL-3 | 0.00E+00 | 0.00E+00 | 0.00E+00 | 4.58E-05 | 0.00E+00 | 0.00E+00 | 0.00E+00 | 0.00E+00 | 3.12E-04 | 3.64E-05 | 0.00E+00 | Known_Unknown |
| f__Maricaulaceae | 0.00E+00 | 0.00E+00 | 0.00E+00 | 0.00E+00 | 0.00E+00 | 5.84E-05 | 0.00E+00 | 0.00E+00 | 0.00E+00 | 0.00E+00 | 0.00E+00 | Known_Unknown |
| f__Melioribacteraceae | 0.00E+00 | 0.00E+00 | 0.00E+00 | 0.00E+00 | 2.73E-05 | 0.00E+00 | 0.00E+00 | 0.00E+00 | 0.00E+00 | 0.00E+00 | 0.00E+00 | Known_Unknown |
| f__Methanomassiliicoccaceae | 4.67E-05 | 0.00E+00 | 0.00E+00 | 0.00E+00 | 0.00E+00 | 0.00E+00 | 0.00E+00 | 0.00E+00 | 0.00E+00 | 0.00E+00 | 0.00E+00 | Known_Unknown |
| f__Methanomethylophilaceae | 0.00E+00 | 0.00E+00 | 0.00E+00 | 0.00E+00 | 0.00E+00 | 0.00E+00 | 0.00E+00 | 0.00E+00 | 5.64E-04 | 1.64E-04 | 1.13E-04 | Known_Unknown |
| f__Methanosarcinaceae | 0.00E+00 | 0.00E+00 | 1.41E-04 | 1.49E-04 | 0.00E+00 | 0.00E+00 | 0.00E+00 | 0.00E+00 | 8.90E-05 | 0.00E+00 | 7.45E-04 | Known_Unknown |
| f__Methylophagaceae | 4.67E-05 | 0.00E+00 | 0.00E+00 | 0.00E+00 | 0.00E+00 | 0.00E+00 | 0.00E+00 | 0.00E+00 | 0.00E+00 | 0.00E+00 | 0.00E+00 | Known_Unknown |
| f__Micavibrionaceae | 2.57E-04 | 1.36E-04 | 0.00E+00 | 1.95E-04 | 0.00E+00 | 0.00E+00 | 6.09E-04 | 8.88E-04 | 0.00E+00 | 1.46E-04 | 3.73E-04 | Known_Unknown |
| f__Micrococcaceae | 0.00E+00 | 3.41E-05 | 4.70E-05 | 0.00E+00 | 3.14E-04 | 4.67E-05 | 0.00E+00 | 0.00E+00 | 0.00E+00 | 1.82E-04 | 1.62E-05 | Known_Unknown |
| f__Micromonosporaceae | 0.00E+00 | 0.00E+00 | 0.00E+00 | 0.00E+00 | 1.36E-05 | 3.50E-05 | 0.00E+00 | 0.00E+00 | 0.00E+00 | 0.00E+00 | 0.00E+00 | Known_Unknown |
| f__Mycobacteriaceae | 0.00E+00 | 0.00E+00 | 2.35E-05 | 0.00E+00 | 0.00E+00 | 0.00E+00 | 0.00E+00 | 0.00E+00 | 0.00E+00 | 0.00E+00 | 0.00E+00 | Known_Unknown |
| f__Myxococcaceae | 0.00E+00 | 5.11E-05 | 0.00E+00 | 0.00E+00 | 0.00E+00 | 0.00E+00 | 0.00E+00 | 0.00E+00 | 0.00E+00 | 0.00E+00 | 0.00E+00 | Known_Unknown |
| f__Natrialbaceae | 8.12E-03 | 3.16E-02 | 2.94E-02 | 5.93E-03 | 2.76E-03 | 6.30E-04 | 4.19E-03 | 5.92E-04 | 8.16E-04 | 7.29E-04 | 5.40E-03 | Known_Unknown |
| f__Natronincolaceae | 6.60E-03 | 3.63E-03 | 1.12E-03 | 8.81E-04 | 3.38E-03 | 6.90E-03 | 4.03E-03 | 7.97E-03 | 1.53E-02 | 1.39E-02 | 1.20E-02 | Known_Unknown |
| f__NBRC-103111 | 0.00E+00 | 0.00E+00 | 0.00E+00 | 0.00E+00 | 5.46E-05 | 3.50E-05 | 0.00E+00 | 0.00E+00 | 0.00E+00 | 0.00E+00 | 0.00E+00 | Known_Unknown |
| f__Nitrososphaeraceae | 0.00E+00 | 0.00E+00 | 0.00E+00 | 0.00E+00 | 2.05E-04 | 9.22E-04 | 0.00E+00 | 0.00E+00 | 1.34E-04 | 0.00E+00 | 0.00E+00 | Known_Unknown |
| f__Nocardioidaceae | 0.00E+00 | 5.11E-05 | 3.52E-05 | 0.00E+00 | 0.00E+00 | 0.00E+00 | 0.00E+00 | 0.00E+00 | 0.00E+00 | 0.00E+00 | 0.00E+00 | Known_Unknown |
| f__Oleiphilaceae_637066 | 0.00E+00 | 0.00E+00 | 5.87E-05 | 0.00E+00 | 0.00E+00 | 0.00E+00 | 0.00E+00 | 0.00E+00 | 0.00E+00 | 0.00E+00 | 0.00E+00 | Known_Unknown |
| f__Oscillospiraceae_88309 | 1.17E-04 | 0.00E+00 | 3.52E-05 | 0.00E+00 | 0.00E+00 | 0.00E+00 | 0.00E+00 | 0.00E+00 | 0.00E+00 | 0.00E+00 | 0.00E+00 | Known_Unknown |
| f__Paenibacillaceae_367444 | 0.00E+00 | 0.00E+00 | 0.00E+00 | 0.00E+00 | 1.09E-04 | 0.00E+00 | 0.00E+00 | 6.44E-05 | 0.00E+00 | 0.00E+00 | 0.00E+00 | Known_Unknown |
| f__Parvularculaceae | 0.00E+00 | 5.11E-05 | 3.52E-05 | 2.06E-04 | 1.23E-04 | 0.00E+00 | 0.00E+00 | 1.67E-04 | 0.00E+00 | 0.00E+00 | 0.00E+00 | Known_Unknown |
| f__Peptostreptococcaceae_256921 | 1.31E-03 | 2.21E-04 | 0.00E+00 | 0.00E+00 | 8.73E-04 | 4.09E-04 | 5.19E-04 | 4.40E-03 | 5.93E-05 | 2.19E-04 | 4.54E-04 | Known_Unknown |
| f__Phycisphaeraceae | 0.00E+00 | 3.41E-05 | 0.00E+00 | 0.00E+00 | 0.00E+00 | 0.00E+00 | 0.00E+00 | 0.00E+00 | 0.00E+00 | 0.00E+00 | 0.00E+00 | Known_Unknown |
| f__PJMF01 | 0.00E+00 | 0.00E+00 | 0.00E+00 | 0.00E+00 | 0.00E+00 | 0.00E+00 | 0.00E+00 | 0.00E+00 | 2.97E-05 | 0.00E+00 | 3.24E-05 | Known_Unknown |
| f__Planococcaceae | 0.00E+00 | 1.02E-04 | 9.39E-05 | 0.00E+00 | 1.23E-04 | 2.34E-05 | 0.00E+00 | 0.00E+00 | 0.00E+00 | 3.64E-05 | 0.00E+00 | Known_Unknown |
| f__Pleomorphomonadaceae | 0.00E+00 | 4.09E-04 | 0.00E+00 | 5.95E-04 | 0.00E+00 | 5.84E-05 | 0.00E+00 | 5.15E-04 | 0.00E+00 | 0.00E+00 | 1.94E-04 | Known_Unknown |
| f__Prolixibacteraceae | 0.00E+00 | 5.11E-05 | 0.00E+00 | 0.00E+00 | 0.00E+00 | 0.00E+00 | 0.00E+00 | 0.00E+00 | 0.00E+00 | 0.00E+00 | 6.48E-05 | Known_Unknown |
| f__Propionibacteriaceae | 0.00E+00 | 0.00E+00 | 0.00E+00 | 0.00E+00 | 6.82E-05 | 0.00E+00 | 0.00E+00 | 0.00E+00 | 0.00E+00 | 0.00E+00 | 0.00E+00 | Known_Unknown |
| f__Pseudonocardiaceae | 0.00E+00 | 0.00E+00 | 0.00E+00 | 0.00E+00 | 2.73E-05 | 0.00E+00 | 0.00E+00 | 0.00E+00 | 0.00E+00 | 0.00E+00 | 0.00E+00 | Known_Unknown |
| f__RBG-13-55-18 | 0.00E+00 | 0.00E+00 | 0.00E+00 | 0.00E+00 | 0.00E+00 | 0.00E+00 | 0.00E+00 | 0.00E+00 | 0.00E+00 | 5.46E-05 | 8.10E-05 | Known_Unknown |
| f__Rhizobiaceae_A_499470 | 0.00E+00 | 0.00E+00 | 0.00E+00 | 0.00E+00 | 4.09E-05 | 0.00E+00 | 0.00E+00 | 0.00E+00 | 0.00E+00 | 0.00E+00 | 0.00E+00 | Known_Unknown |
| f__Rhodobacteraceae | 6.16E-02 | 3.54E-02 | 3.29E-02 | 2.07E-01 | 9.88E-03 | 1.51E-02 | 4.42E-02 | 1.63E-01 | 9.51E-03 | 9.89E-03 | 2.82E-02 | Known_Unknown |
| f__Rhodospirillaceae | 0.00E+00 | 0.00E+00 | 4.70E-05 | 6.87E-05 | 0.00E+00 | 0.00E+00 | 0.00E+00 | 0.00E+00 | 0.00E+00 | 0.00E+00 | 0.00E+00 | Known_Unknown |

| Genus | L1 | L2 | L3 | L4 | L5 | L6 | L7 | L8 | L9 | L10 | L11 | Microbial_matter |
|---|---|---|---|---|---|---|---|---|---|---|---|---|
| f__Rhodovibrionaceae | 0.00E+00 | 1.36E-04 | 2.00E-04 | 5.72E-05 | 0.00E+00 | 0.00E+00 | 0.00E+00 | 0.00E+00 | 0.00E+00 | 0.00E+00 | 0.00E+00 | Known_Unknown |
| f__Roseiflexaceae | 0.00E+00 | 0.00E+00 | 0.00E+00 | 0.00E+00 | 0.00E+00 | 2.34E-05 | 0.00E+00 | 0.00E+00 | 0.00E+00 | 0.00E+00 | 0.00E+00 | Known_Unknown |
| f__Rubrobacteraceae | 4.67E-05 | 0.00E+00 | 0.00E+00 | 0.00E+00 | 1.77E-04 | 3.50E-05 | 0.00E+00 | 0.00E+00 | 0.00E+00 | 1.82E-05 | 0.00E+00 | Known_Unknown |
| f__Ruminococcaceae | 0.00E+00 | 0.00E+00 | 0.00E+00 | 0.00E+00 | 0.00E+00 | 0.00E+00 | 5.37E-05 | 3.22E-04 | 0.00E+00 | 3.64E-04 | 9.72E-05 | Known_Unknown |
| f__Saccharofermentanaceae | 2.10E-04 | 0.00E+00 | 2.00E-04 | 2.40E-04 | 4.50E-04 | 5.95E-04 | 5.37E-05 | 3.61E-04 | 1.93E-04 | 7.29E-05 | 9.72E-05 | Known_Unknown |
| f__Salisediminibacteriaceae | 0.00E+00 | 0.00E+00 | 5.87E-05 | 0.00E+00 | 8.19E-05 | 7.01E-05 | 7.16E-05 | 1.29E-05 | 4.45E-05 | 1.64E-04 | 8.10E-05 | Known_Unknown |
| f__Sedimentibacteraceae | 1.28E-03 | 3.92E-04 | 6.69E-04 | 5.95E-04 | 0.00E+00 | 0.00E+00 | 1.79E-04 | 2.45E-04 | 0.00E+00 | 1.15E-03 | 1.41E-03 | Known_Unknown |
| f__SLSP01 | 0.00E+00 | 0.00E+00 | 0.00E+00 | 0.00E+00 | 0.00E+00 | 2.34E-05 | 0.00E+00 | 0.00E+00 | 0.00E+00 | 0.00E+00 | 0.00E+00 | Known_Unknown |
| f__SLSP02 | 4.88E-03 | 1.28E-03 | 1.29E-04 | 0.00E+00 | 0.00E+00 | 0.00E+00 | 0.00E+00 | 1.80E-04 | 1.48E-04 | 1.09E-04 | 8.91E-04 | Known_Unknown |
| f__SM1A02 | 0.00E+00 | 0.00E+00 | 0.00E+00 | 5.72E-05 | 0.00E+00 | 0.00E+00 | 0.00E+00 | 3.86E-05 | 0.00E+00 | 0.00E+00 | 0.00E+00 | Known_Unknown |
| f__SM23-32 | 0.00E+00 | 0.00E+00 | 0.00E+00 | 0.00E+00 | 0.00E+00 | 0.00E+00 | 0.00E+00 | 0.00E+00 | 0.00E+00 | 0.00E+00 | 4.86E-05 | Known_Unknown |
| f__SM23-33 | 0.00E+00 | 0.00E+00 | 0.00E+00 | 0.00E+00 | 0.00E+00 | 0.00E+00 | 0.00E+00 | 2.58E-05 | 0.00E+00 | 0.00E+00 | 0.00E+00 | Known_Unknown |
| f__SM23-42 | 0.00E+00 | 0.00E+00 | 0.00E+00 | 0.00E+00 | 0.00E+00 | 0.00E+00 | 0.00E+00 | 0.00E+00 | 5.93E-05 | 0.00E+00 | 0.00E+00 | Known_Unknown |
| f__Solirubrobacteraceae_405341 | 0.00E+00 | 0.00E+00 | 0.00E+00 | 0.00E+00 | 5.46E-05 | 0.00E+00 | 0.00E+00 | 0.00E+00 | 0.00E+00 | 5.46E-05 | 0.00E+00 | Known_Unknown |
| f__Sphingomonadaceae | 1.35E-03 | 2.16E-03 | 1.98E-03 | 2.70E-03 | 1.57E-03 | 1.79E-03 | 1.61E-04 | 5.92E-04 | 2.82E-04 | 3.64E-05 | 3.40E-04 | Known_Unknown |
| f__Sporanaerobacteraceae | 0.00E+00 | 0.00E+00 | 0.00E+00 | 0.00E+00 | 0.00E+00 | 0.00E+00 | 0.00E+00 | 3.86E-05 | 0.00E+00 | 0.00E+00 | 0.00E+00 | Known_Unknown |
| f__Stappiaceae | 0.00E+00 | 0.00E+00 | 0.00E+00 | 5.72E-05 | 0.00E+00 | 0.00E+00 | 0.00E+00 | 0.00E+00 | 0.00E+00 | 0.00E+00 | 0.00E+00 | Known_Unknown |
| f__Streptomycetaceae_400641 | 0.00E+00 | 0.00E+00 | 0.00E+00 | 0.00E+00 | 0.00E+00 | 0.00E+00 | 0.00E+00 | 0.00E+00 | 1.48E-05 | 0.00E+00 | 0.00E+00 | Known_Unknown |
| f__Streptosporangiaceae | 0.00E+00 | 0.00E+00 | 4.70E-05 | 0.00E+00 | 0.00E+00 | 0.00E+00 | 0.00E+00 | 0.00E+00 | 0.00E+00 | 0.00E+00 | 0.00E+00 | Known_Unknown |
| f__Sulfurimonadaceae | 0.00E+00 | 0.00E+00 | 4.70E-05 | 4.58E-05 | 0.00E+00 | 0.00E+00 | 0.00E+00 | 0.00E+00 | 0.00E+00 | 0.00E+00 | 0.00E+00 | Known_Unknown |
| f__Sulfurovaceae | 7.00E-05 | 2.08E-03 | 0.00E+00 | 1.26E-04 | 0.00E+00 | 0.00E+00 | 0.00E+00 | 0.00E+00 | 1.34E-04 | 0.00E+00 | 0.00E+00 | Known_Unknown |
| f__SURF-3 | 0.00E+00 | 8.52E-05 | 1.06E-04 | 0.00E+00 | 1.91E-04 | 4.67E-05 | 1.79E-05 | 0.00E+00 | 8.90E-05 | 3.64E-04 | 4.86E-05 | Known_Unknown |
| f__Synergistaceae | 0.00E+00 | 0.00E+00 | 0.00E+00 | 0.00E+00 | 4.09E-05 | 4.67E-05 | 0.00E+00 | 0.00E+00 | 0.00E+00 | 0.00E+00 | 0.00E+00 | Known_Unknown |
| f__Syntrophomonadaceae_368540 | 5.60E-04 | 9.37E-04 | 3.87E-04 | 4.69E-04 | 1.50E-04 | 3.39E-04 | 6.99E-04 | 7.60E-04 | 2.97E-04 | 1.22E-03 | 7.94E-04 | Known_Unknown |
| f__TH1-2 | 0.00E+00 | 0.00E+00 | 0.00E+00 | 4.58E-05 | 0.00E+00 | 0.00E+00 | 0.00E+00 | 0.00E+00 | 0.00E+00 | 0.00E+00 | 3.24E-05 | Known_Unknown |
| f__Thermococcaceae | 0.00E+00 | 0.00E+00 | 0.00E+00 | 4.58E-05 | 0.00E+00 | 0.00E+00 | 0.00E+00 | 0.00E+00 | 0.00E+00 | 0.00E+00 | 0.00E+00 | Known_Unknown |
| f__Thermomicrobiaceae | 0.00E+00 | 0.00E+00 | 0.00E+00 | 0.00E+00 | 0.00E+00 | 0.00E+00 | 0.00E+00 | 0.00E+00 | 0.00E+00 | 9.11E-05 | 0.00E+00 | Known_Unknown |
| f__Thermotaleaceae | 1.17E-04 | 0.00E+00 | 1.53E-04 | 0.00E+00 | 7.50E-04 | 2.28E-03 | 5.02E-04 | 1.44E-03 | 1.13E-03 | 1.82E-05 | 4.86E-05 | Known_Unknown |
| f__Thorarchaeaceae | 0.00E+00 | 0.00E+00 | 0.00E+00 | 0.00E+00 | 0.00E+00 | 0.00E+00 | 0.00E+00 | 0.00E+00 | 1.10E-03 | 8.56E-04 | 2.76E-04 | Known_Unknown |
| f__Trueperaceae | 4.67E-05 | 3.92E-04 | 0.00E+00 | 0.00E+00 | 1.09E-04 | 0.00E+00 | 0.00E+00 | 0.00E+00 | 4.45E-05 | 0.00E+00 | 6.48E-05 | Known_Unknown |
| f__UBA10185 | 0.00E+00 | 0.00E+00 | 0.00E+00 | 2.40E-04 | 0.00E+00 | 0.00E+00 | 0.00E+00 | 0.00E+00 | 2.97E-05 | 0.00E+00 | 4.86E-05 | Known_Unknown |
| f__UBA10450 | 1.17E-04 | 6.81E-05 | 0.00E+00 | 2.98E-04 | 9.55E-05 | 0.00E+00 | 0.00E+00 | 2.58E-04 | 7.42E-05 | 0.00E+00 | 1.62E-04 | Known_Unknown |
| f__UBA12075 | 0.00E+00 | 0.00E+00 | 0.00E+00 | 2.29E-05 | 0.00E+00 | 0.00E+00 | 0.00E+00 | 5.15E-05 | 0.00E+00 | 5.46E-05 | 0.00E+00 | Known_Unknown |
| f__UBA1242 | 1.40E-04 | 0.00E+00 | 0.00E+00 | 5.72E-05 | 0.00E+00 | 1.63E-04 | 1.07E-04 | 2.45E-04 | 8.90E-05 | 8.74E-04 | 7.62E-04 | Known_Unknown |
| f__UBA1328 | 0.00E+00 | 1.36E-04 | 5.87E-05 | 5.72E-05 | 4.09E-05 | 0.00E+00 | 0.00E+00 | 0.00E+00 | 0.00E+00 | 0.00E+00 | 0.00E+00 | Known_Unknown |
| f__UBA2279 | 0.00E+00 | 0.00E+00 | 0.00E+00 | 0.00E+00 | 0.00E+00 | 0.00E+00 | 0.00E+00 | 0.00E+00 | 4.45E-05 | 0.00E+00 | 0.00E+00 | Known_Unknown |
| f__UBA2963 | 0.00E+00 | 0.00E+00 | 1.06E-04 | 3.43E-05 | 0.00E+00 | 0.00E+00 | 0.00E+00 | 0.00E+00 | 0.00E+00 | 0.00E+00 | 0.00E+00 | Known_Unknown |
| f__UBA4665 | 0.00E+00 | 0.00E+00 | 0.00E+00 | 5.72E-05 | 0.00E+00 | 0.00E+00 | 0.00E+00 | 0.00E+00 | 4.45E-05 | 0.00E+00 | 0.00E+00 | Known_Unknown |
| f__UBA4823 | 0.00E+00 | 0.00E+00 | 0.00E+00 | 0.00E+00 | 0.00E+00 | 0.00E+00 | 0.00E+00 | 0.00E+00 | 4.01E-04 | 0.00E+00 | 0.00E+00 | Known_Unknown |
| f__UBA4824 | 0.00E+00 | 2.04E-04 | 0.00E+00 | 1.95E-04 | 2.46E-04 | 0.00E+00 | 0.00E+00 | 1.42E-04 | 3.50E-03 | 8.92E-04 | 1.94E-04 | Known_Unknown |
| f__UBA5272 | 0.00E+00 | 1.70E-05 | 0.00E+00 | 0.00E+00 | 0.00E+00 | 0.00E+00 | 0.00E+00 | 0.00E+00 | 1.19E-04 | 0.00E+00 | 0.00E+00 | Known_Unknown |
| f__UBA5550 | 0.00E+00 | 0.00E+00 | 0.00E+00 | 0.00E+00 | 0.00E+00 | 0.00E+00 | 0.00E+00 | 0.00E+00 | 0.00E+00 | 0.00E+00 | 3.24E-05 | Known_Unknown |
| f__UBA8154 | 0.00E+00 | 0.00E+00 | 0.00E+00 | 0.00E+00 | 1.36E-05 | 0.00E+00 | 0.00E+00 | 5.15E-05 | 7.42E-05 | 0.00E+00 | 0.00E+00 | Known_Unknown |
| f__UBA8155 | 0.00E+00 | 3.75E-04 | 5.05E-04 | 1.73E-03 | 1.09E-03 | 1.21E-03 | 8.24E-04 | 2.83E-04 | 4.60E-04 | 2.55E-04 | 1.94E-04 | Known_Unknown |
| f__UBA9579 | 3.27E-04 | 5.11E-05 | 0.00E+00 | 4.58E-05 | 4.78E-04 | 0.00E+00 | 0.00E+00 | 1.03E-04 | 1.34E-04 | 0.00E+00 | 6.48E-05 | Known_Unknown |
| f__VadinHA17_874699 | 0.00E+00 | 0.00E+00 | 0.00E+00 | 0.00E+00 | 0.00E+00 | 0.00E+00 | 0.00E+00 | 0.00E+00 | 0.00E+00 | 1.82E-05 | 0.00E+00 | Known_Unknown |
| f__VadinHA17_877549 | 0.00E+00 | 0.00E+00 | 0.00E+00 | 0.00E+00 | 0.00E+00 | 0.00E+00 | 0.00E+00 | 0.00E+00 | 0.00E+00 | 3.64E-05 | 0.00E+00 | Known_Unknown |

| Genus | L1 | L2 | L3 | L4 | L5 | L6 | L7 | L8 | L9 | L10 | L11 | Microbial_matter |
|---|---|---|---|---|---|---|---|---|---|---|---|---|
| f__VENL01 | 7.47E-03 | 4.80E-03 | 2.22E-03 | 1.73E-03 | 7.50E-03 | 1.31E-02 | 8.36E-03 | 4.11E-02 | 1.94E-02 | 4.15E-02 | 1.39E-02 | Known_Unknown |
| f__VENL02 | 0.00E+00 | 0.00E+00 | 0.00E+00 | 0.00E+00 | 0.00E+00 | 0.00E+00 | 0.00E+00 | 0.00E+00 | 0.00E+00 | 2.64E-03 | 0.00E+00 | Known_Unknown |
| f__Xanthobacteraceae_503485 | 0.00E+00 | 0.00E+00 | 0.00E+00 | 5.72E-05 | 0.00E+00 | 0.00E+00 | 0.00E+00 | 0.00E+00 | 0.00E+00 | 0.00E+00 | 0.00E+00 | Known_Unknown |
| f__Xanthomonadaceae_616009 | 0.00E+00 | 0.00E+00 | 0.00E+00 | 0.00E+00 | 0.00E+00 | 0.00E+00 | 0.00E+00 | 0.00E+00 | 1.63E-04 | 5.46E-05 | 3.24E-05 | Known_Unknown |
| o__AB16 | 7.00E-05 | 0.00E+00 | 0.00E+00 | 0.00E+00 | 0.00E+00 | 0.00E+00 | 0.00E+00 | 0.00E+00 | 0.00E+00 | 3.64E-05 | 0.00E+00 | Known_Unknown |
| o__Acetivibrionales | 4.39E-03 | 8.09E-03 | 4.03E-03 | 4.74E-03 | 1.49E-02 | 7.27E-03 | 1.40E-02 | 2.11E-02 | 1.12E-02 | 4.98E-02 | 2.91E-02 | Known_Unknown |
| o__Acidimicrobiales | 1.63E-04 | 2.38E-04 | 0.00E+00 | 6.87E-05 | 5.46E-04 | 5.95E-04 | 0.00E+00 | 0.00E+00 | 1.78E-04 | 2.55E-04 | 0.00E+00 | Known_Unknown |
| o__Acidobacteriales | 0.00E+00 | 0.00E+00 | 0.00E+00 | 0.00E+00 | 0.00E+00 | 0.00E+00 | 0.00E+00 | 0.00E+00 | 4.45E-05 | 0.00E+00 | 0.00E+00 | Known_Unknown |
| o__Actinomycetales | 0.00E+00 | 0.00E+00 | 0.00E+00 | 0.00E+00 | 8.19E-05 | 0.00E+00 | 0.00E+00 | 0.00E+00 | 4.45E-05 | 0.00E+00 | 0.00E+00 | Known_Unknown |
| o__Aggregatilineales | 7.00E-05 | 0.00E+00 | 0.00E+00 | 0.00E+00 | 0.00E+00 | 0.00E+00 | 0.00E+00 | 0.00E+00 | 0.00E+00 | 0.00E+00 | 0.00E+00 | Known_Unknown |
| o__Aminicenantales | 7.00E-05 | 5.11E-05 | 0.00E+00 | 0.00E+00 | 0.00E+00 | 0.00E+00 | 0.00E+00 | 0.00E+00 | 2.37E-04 | 0.00E+00 | 0.00E+00 | Known_Unknown |
| o__Anaerolineales | 0.00E+00 | 0.00E+00 | 2.35E-05 | 3.89E-04 | 5.18E-04 | 0.00E+00 | 1.25E-04 | 0.00E+00 | 1.63E-04 | 5.46E-05 | 0.00E+00 | Known_Unknown |
| o__Azospirillales_507929 | 0.00E+00 | 5.28E-04 | 1.76E-04 | 2.29E-04 | 5.46E-05 | 5.84E-05 | 1.07E-04 | 1.03E-04 | 5.49E-04 | 3.10E-04 | 0.00E+00 | Known_Unknown |
| o__Azospirillales_507931 | 0.00E+00 | 6.47E-04 | 0.00E+00 | 0.00E+00 | 0.00E+00 | 0.00E+00 | 0.00E+00 | 2.96E-04 | 3.15E-03 | 6.48E-03 | 1.15E-03 | Known_Unknown |
| o__B13-G15 | 0.00E+00 | 5.11E-05 | 5.87E-05 | 0.00E+00 | 0.00E+00 | 0.00E+00 | 0.00E+00 | 0.00E+00 | 0.00E+00 | 7.29E-05 | 0.00E+00 | Known_Unknown |
| o__Bacillales_B_302584 | 0.00E+00 | 0.00E+00 | 1.17E-05 | 0.00E+00 | 1.36E-05 | 3.50E-05 | 0.00E+00 | 0.00E+00 | 4.45E-05 | 7.29E-05 | 4.86E-05 | Known_Unknown |
| o__Bacillales_D_310495 | 0.00E+00 | 0.00E+00 | 4.70E-05 | 0.00E+00 | 4.09E-05 | 3.50E-05 | 1.97E-04 | 2.58E-05 | 0.00E+00 | 1.09E-04 | 0.00E+00 | Known_Unknown |
| o__Bacillales_H | 1.63E-04 | 3.41E-05 | 1.41E-04 | 0.00E+00 | 7.91E-04 | 5.84E-04 | 3.04E-04 | 9.01E-05 | 1.48E-04 | 6.37E-04 | 1.78E-04 | Known_Unknown |
| o__Bacteroidales | 0.00E+00 | 0.00E+00 | 3.52E-05 | 3.43E-05 | 0.00E+00 | 0.00E+00 | 0.00E+00 | 5.15E-05 | 3.26E-04 | 1.31E-03 | 6.48E-05 | Known_Unknown |
| o__Bdellovibrionales | 4.67E-05 | 0.00E+00 | 0.00E+00 | 0.00E+00 | 0.00E+00 | 0.00E+00 | 0.00E+00 | 9.01E-05 | 0.00E+00 | 1.27E-04 | 0.00E+00 | Known_Unknown |
| o__Caldilineales | 0.00E+00 | 0.00E+00 | 0.00E+00 | 4.58E-05 | 0.00E+00 | 0.00E+00 | 0.00E+00 | 0.00E+00 | 0.00E+00 | 0.00E+00 | 0.00E+00 | Known_Unknown |
| o__Campylobacterales | 1.29E-02 | 3.63E-02 | 1.05E-01 | 5.93E-03 | 0.00E+00 | 4.67E-05 | 0.00E+00 | 0.00E+00 | 0.00E+00 | 1.88E-02 | 9.64E-03 | Known_Unknown |
| o__Christensenellales | 9.33E-05 | 0.00E+00 | 0.00E+00 | 3.32E-04 | 0.00E+00 | 7.01E-05 | 3.58E-04 | 1.16E-04 | 0.00E+00 | 1.82E-04 | 1.78E-04 | Known_Unknown |
| o__Clostridiales | 0.00E+00 | 4.09E-04 | 2.70E-04 | 0.00E+00 | 1.77E-04 | 0.00E+00 | 0.00E+00 | 0.00E+00 | 0.00E+00 | 3.64E-04 | 3.40E-04 | Known_Unknown |
| o__Cyanobacteriales | 9.33E-05 | 2.04E-04 | 4.81E-04 | 9.39E-04 | 8.19E-05 | 5.84E-05 | 0.00E+00 | 0.00E+00 | 1.48E-04 | 0.00E+00 | 1.46E-04 | Known_Unknown |
| o__Cytophagales | 0.00E+00 | 0.00E+00 | 0.00E+00 | 0.00E+00 | 0.00E+00 | 0.00E+00 | 0.00E+00 | 2.58E-05 | 0.00E+00 | 3.64E-05 | 6.48E-05 | Known_Unknown |
| o__Dehalococcoidales | 1.40E-03 | 1.55E-03 | 3.76E-04 | 2.86E-04 | 3.27E-04 | 8.17E-05 | 5.37E-05 | 4.89E-04 | 1.08E-03 | 3.81E-03 | 1.04E-03 | Known_Unknown |
| o__Deinococcales | 0.00E+00 | 0.00E+00 | 0.00E+00 | 0.00E+00 | 1.36E-05 | 0.00E+00 | 0.00E+00 | 0.00E+00 | 0.00E+00 | 0.00E+00 | 0.00E+00 | Known_Unknown |
| o__Desulfarculales | 0.00E+00 | 0.00E+00 | 0.00E+00 | 0.00E+00 | 0.00E+00 | 0.00E+00 | 0.00E+00 | 0.00E+00 | 0.00E+00 | 5.46E-05 | 0.00E+00 | Known_Unknown |
| o__Desulfatiglandales | 0.00E+00 | 0.00E+00 | 0.00E+00 | 0.00E+00 | 0.00E+00 | 0.00E+00 | 0.00E+00 | 0.00E+00 | 2.97E-05 | 9.11E-05 | 0.00E+00 | Known_Unknown |
| o__Desulfitobacteriales | 0.00E+00 | 6.81E-05 | 1.06E-04 | 0.00E+00 | 0.00E+00 | 0.00E+00 | 0.00E+00 | 7.73E-05 | 4.45E-05 | 2.00E-04 | 9.72E-05 | Known_Unknown |
| o__Desulfobacterales | 7.00E-05 | 3.41E-05 | 2.00E-04 | 9.16E-05 | 1.23E-04 | 0.00E+00 | 3.58E-05 | 1.29E-05 | 8.90E-05 | 9.11E-05 | 2.92E-04 | Known_Unknown |
| o__Desulfotomaculales_370525 | 0.00E+00 | 1.16E-03 | 1.80E-03 | 2.86E-04 | 1.23E-04 | 3.50E-05 | 0.00E+00 | 0.00E+00 | 0.00E+00 | 1.27E-04 | 1.62E-04 | Known_Unknown |
| o__Desulfotomaculales_370527 | 5.60E-04 | 7.65E-03 | 5.72E-03 | 2.34E-03 | 2.01E-03 | 3.97E-04 | 7.16E-05 | 0.00E+00 | 0.00E+00 | 0.00E+00 | 7.13E-04 | Known_Unknown |
| o__Desulfovibrionales | 2.10E-04 | 0.00E+00 | 1.06E-04 | 0.00E+00 | 0.00E+00 | 0.00E+00 | 0.00E+00 | 0.00E+00 | 0.00E+00 | 0.00E+00 | 1.62E-04 | Known_Unknown |
| o__Desulfuromonadales | 0.00E+00 | 0.00E+00 | 0.00E+00 | 0.00E+00 | 0.00E+00 | 0.00E+00 | 0.00E+00 | 0.00E+00 | 0.00E+00 | 1.82E-05 | 0.00E+00 | Known_Unknown |
| o__Dissulfuribacterales | 0.00E+00 | 0.00E+00 | 0.00E+00 | 0.00E+00 | 0.00E+00 | 0.00E+00 | 0.00E+00 | 0.00E+00 | 0.00E+00 | 1.09E-04 | 8.10E-05 | Known_Unknown |
| o__DSM-27196 | 0.00E+00 | 5.11E-05 | 0.00E+00 | 0.00E+00 | 5.46E-05 | 4.67E-05 | 0.00E+00 | 0.00E+00 | 0.00E+00 | 3.64E-05 | 3.24E-05 | Known_Unknown |
| o__DTU022 | 1.17E-04 | 3.92E-04 | 7.04E-05 | 0.00E+00 | 3.14E-04 | 0.00E+00 | 0.00E+00 | 0.00E+00 | 7.42E-05 | 1.09E-04 | 0.00E+00 | Known_Unknown |
| o__DTU024 | 6.37E-03 | 6.06E-02 | 6.10E-04 | 2.74E-03 | 3.23E-02 | 5.14E-04 | 3.53E-03 | 9.14E-03 | 7.14E-03 | 7.58E-03 | 7.52E-03 | Known_Unknown |
| o__Eubacteriales_258471 | 9.33E-05 | 0.00E+00 | 0.00E+00 | 5.72E-05 | 0.00E+00 | 0.00E+00 | 1.07E-04 | 0.00E+00 | 8.90E-05 | 0.00E+00 | 0.00E+00 | Known_Unknown |
| o__Eubacteriales_258473 | 0.00E+00 | 0.00E+00 | 0.00E+00 | 0.00E+00 | 0.00E+00 | 1.75E-04 | 1.43E-04 | 0.00E+00 | 0.00E+00 | 0.00E+00 | 0.00E+00 | Known_Unknown |
| o__Gaiellales | 0.00E+00 | 0.00E+00 | 0.00E+00 | 0.00E+00 | 0.00E+00 | 3.50E-05 | 0.00E+00 | 0.00E+00 | 0.00E+00 | 0.00E+00 | 0.00E+00 | Known_Unknown |
| o__Gastranaerophilales | 4.43E-04 | 1.70E-04 | 0.00E+00 | 1.72E-04 | 3.55E-04 | 0.00E+00 | 1.79E-05 | 1.16E-04 | 1.04E-04 | 0.00E+00 | 1.78E-04 | Known_Unknown |
| o__GWA2-44-7 | 0.00E+00 | 0.00E+00 | 0.00E+00 | 0.00E+00 | 0.00E+00 | 0.00E+00 | 0.00E+00 | 0.00E+00 | 0.00E+00 | 0.00E+00 | 3.24E-05 | Known_Unknown |
| o__Hadarchaeales | 2.57E-04 | 5.79E-04 | 0.00E+00 | 0.00E+00 | 1.09E-04 | 4.67E-05 | 0.00E+00 | 0.00E+00 | 2.97E-05 | 9.11E-05 | 1.78E-04 | Known_Unknown |
| o__Halanaerobiales | 5.60E-04 | 8.72E-03 | 1.11E-02 | 5.15E-04 | 1.04E-03 | 5.84E-04 | 7.16E-04 | 9.01E-05 | 7.57E-04 | 1.15E-03 | 1.73E-03 | Known_Unknown |

| Genus | L1 | L2 | L3 | L4 | L5 | L6 | L7 | L8 | L9 | L10 | L11 | Microbial_matter |
|---|---|---|---|---|---|---|---|---|---|---|---|---|
| o__Halobacteriales | 1.87E-03 | 1.45E-02 | 7.72E-03 | 2.37E-03 | 1.20E-03 | 6.65E-04 | 6.81E-04 | 1.93E-04 | 8.46E-04 | 6.01E-04 | 1.60E-03 | Known_Unknown |
| o__Iainarchaeales | 1.17E-04 | 0.00E+00 | 0.00E+00 | 0.00E+00 | 0.00E+00 | 0.00E+00 | 0.00E+00 | 1.55E-04 | 7.42E-05 | 2.19E-04 | 3.24E-04 | Known_Unknown |
| o__Lachnospirales | 2.57E-04 | 7.49E-04 | 2.23E-04 | 1.79E-03 | 1.94E-03 | 1.33E-03 | 5.02E-04 | 2.48E-03 | 4.44E-03 | 3.93E-03 | 2.04E-03 | Known_Unknown |
| o__Limnochordales_A | 0.00E+00 | 1.06E-03 | 6.34E-04 | 1.72E-04 | 1.36E-04 | 0.00E+00 | 5.37E-05 | 0.00E+00 | 1.19E-04 | 0.00E+00 | 1.46E-04 | Known_Unknown |
| o__Methanomassiliicoccales | 2.01E-03 | 3.24E-04 | 1.76E-04 | 0.00E+00 | 6.82E-05 | 1.11E-03 | 5.79E-03 | 5.01E-03 | 4.45E-04 | 1.22E-03 | 1.85E-03 | Known_Unknown |
| o__Mycobacteriales | 4.67E-05 | 3.41E-05 | 0.00E+00 | 0.00E+00 | 4.23E-04 | 7.01E-05 | 0.00E+00 | 0.00E+00 | 1.04E-04 | 0.00E+00 | 8.10E-05 | Known_Unknown |
| o__Myxococcales | 4.67E-05 | 0.00E+00 | 0.00E+00 | 0.00E+00 | 0.00E+00 | 0.00E+00 | 0.00E+00 | 3.86E-05 | 2.82E-04 | 5.46E-04 | 1.13E-04 | Known_Unknown |
| o__Nannocystales | 0.00E+00 | 0.00E+00 | 0.00E+00 | 0.00E+00 | 0.00E+00 | 0.00E+00 | 0.00E+00 | 2.58E-05 | 7.42E-05 | 0.00E+00 | 0.00E+00 | Known_Unknown |
| o__Oscillospirales | 0.00E+00 | 0.00E+00 | 0.00E+00 | 0.00E+00 | 0.00E+00 | 0.00E+00 | 0.00E+00 | 0.00E+00 | 4.45E-05 | 0.00E+00 | 0.00E+00 | Known_Unknown |
| o__Paenibacillales | 0.00E+00 | 1.70E-05 | 2.35E-05 | 4.58E-05 | 1.77E-04 | 1.17E-04 | 0.00E+00 | 1.80E-04 | 2.97E-05 | 2.91E-04 | 0.00E+00 | Known_Unknown |
| o__PCC-6307 | 1.87E-04 | 2.04E-04 | 8.22E-05 | 1.72E-04 | 0.00E+00 | 0.00E+00 | 0.00E+00 | 0.00E+00 | 0.00E+00 | 7.29E-05 | 8.10E-05 | Known_Unknown |
| o__Peptostreptococcales | 1.02E-02 | 7.03E-03 | 6.87E-03 | 6.32E-03 | 1.74E-02 | 3.57E-02 | 1.28E-02 | 1.83E-02 | 2.32E-02 | 2.30E-02 | 1.78E-02 | Known_Unknown |
| o__Phycisphaerales | 0.00E+00 | 6.81E-05 | 8.22E-05 | 5.72E-05 | 6.82E-05 | 0.00E+00 | 0.00E+00 | 5.15E-05 | 2.97E-05 | 1.27E-04 | 0.00E+00 | Known_Unknown |
| o__Pirellulales | 0.00E+00 | 0.00E+00 | 3.52E-05 | 0.00E+00 | 0.00E+00 | 0.00E+00 | 0.00E+00 | 0.00E+00 | 0.00E+00 | 0.00E+00 | 0.00E+00 | Known_Unknown |
| o__Propionibacteriales | 0.00E+00 | 0.00E+00 | 3.52E-05 | 0.00E+00 | 0.00E+00 | 0.00E+00 | 0.00E+00 | 0.00E+00 | 0.00E+00 | 0.00E+00 | 0.00E+00 | Known_Unknown |
| o__Proteinivoracales | 2.57E-04 | 1.36E-04 | 1.76E-04 | 0.00E+00 | 1.26E-03 | 2.90E-03 | 9.13E-04 | 6.95E-04 | 2.06E-03 | 5.46E-04 | 9.72E-04 | Known_Unknown |
| o__Rhizobiales_A_501582 | 1.10E-03 | 2.73E-04 | 4.58E-04 | 1.14E-03 | 0.00E+00 | 2.10E-04 | 5.02E-04 | 3.35E-04 | 1.78E-04 | 0.00E+00 | 4.21E-04 | Known_Unknown |
| o__Rhizobiales_A_504705 | 0.00E+00 | 0.00E+00 | 2.35E-05 | 0.00E+00 | 0.00E+00 | 0.00E+00 | 0.00E+00 | 0.00E+00 | 0.00E+00 | 0.00E+00 | 0.00E+00 | Known_Unknown |
| o__Rhizobiales_A_504721 | 0.00E+00 | 0.00E+00 | 0.00E+00 | 0.00E+00 | 0.00E+00 | 0.00E+00 | 0.00E+00 | 0.00E+00 | 0.00E+00 | 1.27E-04 | 2.27E-04 | Known_Unknown |
| o__Rhodospirillales_505895 | 0.00E+00 | 0.00E+00 | 0.00E+00 | 4.12E-04 | 0.00E+00 | 0.00E+00 | 0.00E+00 | 0.00E+00 | 0.00E+00 | 0.00E+00 | 0.00E+00 | Known_Unknown |
| o__Rickettsiales | 0.00E+00 | 0.00E+00 | 0.00E+00 | 3.43E-05 | 0.00E+00 | 0.00E+00 | 0.00E+00 | 0.00E+00 | 0.00E+00 | 0.00E+00 | 0.00E+00 | Known_Unknown |
| o__Saccharimonadales | 0.00E+00 | 1.02E-04 | 0.00E+00 | 5.72E-05 | 0.00E+00 | 0.00E+00 | 0.00E+00 | 0.00E+00 | 1.19E-04 | 0.00E+00 | 0.00E+00 | Known_Unknown |
| o__Saccharofermentanales | 4.67E-05 | 8.86E-04 | 9.39E-05 | 1.25E-03 | 9.14E-04 | 0.00E+00 | 0.00E+00 | 1.25E-03 | 2.08E-04 | 0.00E+00 | 1.30E-04 | Known_Unknown |
| o__SB-45 | 0.00E+00 | 3.41E-05 | 7.04E-05 | 0.00E+00 | 0.00E+00 | 0.00E+00 | 0.00E+00 | 0.00E+00 | 1.48E-04 | 0.00E+00 | 0.00E+00 | Known_Unknown |
| o__SG8-24 | 4.67E-05 | 0.00E+00 | 0.00E+00 | 0.00E+00 | 0.00E+00 | 0.00E+00 | 0.00E+00 | 0.00E+00 | 7.42E-05 | 1.27E-04 | 4.86E-05 | Known_Unknown |
| o__Solirubrobacterales | 9.33E-05 | 0.00E+00 | 2.35E-05 | 0.00E+00 | 4.09E-05 | 4.67E-05 | 0.00E+00 | 0.00E+00 | 0.00E+00 | 0.00E+00 | 0.00E+00 | Known_Unknown |
| o__Sphingomonadales | 0.00E+00 | 0.00E+00 | 1.29E-04 | 0.00E+00 | 0.00E+00 | 0.00E+00 | 0.00E+00 | 5.15E-05 | 0.00E+00 | 0.00E+00 | 0.00E+00 | Known_Unknown |
| o__SpSt-313 | 0.00E+00 | 0.00E+00 | 0.00E+00 | 2.29E-05 | 0.00E+00 | 0.00E+00 | 0.00E+00 | 0.00E+00 | 1.78E-04 | 0.00E+00 | 0.00E+00 | Known_Unknown |
| o__SpSt-315 | 1.63E-04 | 2.73E-04 | 0.00E+00 | 0.00E+00 | 8.19E-05 | 0.00E+00 | 7.16E-05 | 2.32E-04 | 3.56E-04 | 2.55E-04 | 1.30E-04 | Known_Unknown |
| o__Staphylococcales | 0.00E+00 | 0.00E+00 | 0.00E+00 | 2.29E-05 | 0.00E+00 | 0.00E+00 | 0.00E+00 | 0.00E+00 | 0.00E+00 | 3.64E-05 | 0.00E+00 | Known_Unknown |
| o__Synergistales | 1.46E-02 | 1.82E-02 | 2.62E-02 | 3.83E-02 | 8.56E-02 | 6.88E-02 | 4.04E-02 | 2.65E-02 | 2.64E-03 | 4.66E-03 | 2.94E-02 | Known_Unknown |
| o__Thermincolales | 0.00E+00 | 1.02E-04 | 0.00E+00 | 0.00E+00 | 0.00E+00 | 5.84E-05 | 0.00E+00 | 0.00E+00 | 0.00E+00 | 2.37E-04 | 1.78E-04 | Known_Unknown |
| o__Thermoactinomycetales_367866 | 0.00E+00 | 0.00E+00 | 0.00E+00 | 0.00E+00 | 5.46E-05 | 0.00E+00 | 1.61E-04 | 0.00E+00 | 0.00E+00 | 7.29E-05 | 0.00E+00 | Known_Unknown |
| o__Thermoanaerobaculales | 5.37E-04 | 5.45E-04 | 2.00E-04 | 0.00E+00 | 4.78E-03 | 2.15E-03 | 3.58E-04 | 1.42E-04 | 8.31E-04 | 5.46E-04 | 5.83E-04 | Known_Unknown |
| o__Thermomicrobiales | 2.71E-03 | 3.97E-03 | 3.31E-03 | 2.29E-04 | 1.15E-03 | 6.41E-03 | 2.11E-03 | 1.16E-04 | 1.02E-03 | 1.24E-03 | 5.33E-03 | Known_Unknown |
| o__Tissierellales | 7.14E-03 | 8.52E-04 | 4.58E-04 | 0.00E+00 | 1.17E-03 | 2.22E-03 | 1.18E-03 | 5.90E-03 | 6.29E-03 | 6.28E-03 | 2.98E-03 | Known_Unknown |
| o__UBA1369 | 0.00E+00 | 0.00E+00 | 0.00E+00 | 0.00E+00 | 0.00E+00 | 0.00E+00 | 0.00E+00 | 0.00E+00 | 3.12E-04 | 0.00E+00 | 0.00E+00 | Known_Unknown |
| o__UBA9983_A | 0.00E+00 | 0.00E+00 | 4.70E-05 | 0.00E+00 | 0.00E+00 | 0.00E+00 | 5.37E-05 | 1.55E-04 | 8.90E-05 | 0.00E+00 | 0.00E+00 | Known_Unknown |
| o__Woesearchaeales_581 | 1.37E-02 | 2.38E-03 | 3.52E-04 | 3.89E-04 | 0.00E+00 | 0.00E+00 | 1.07E-04 | 9.01E-05 | 3.56E-04 | 1.27E-04 | 3.97E-03 | Known_Unknown |
| o__Xenobiales | 0.00E+00 | 0.00E+00 | 0.00E+00 | 0.00E+00 | 0.00E+00 | 0.00E+00 | 8.96E-05 | 0.00E+00 | 0.00E+00 | 0.00E+00 | 0.00E+00 | Known_Unknown |
| p__Acidobacteriota | 0.00E+00 | 0.00E+00 | 0.00E+00 | 0.00E+00 | 0.00E+00 | 0.00E+00 | 0.00E+00 | 0.00E+00 | 4.45E-05 | 0.00E+00 | 0.00E+00 | Known_Unknown |
| p__Actinobacteriota | 4.67E-05 | 2.38E-04 | 0.00E+00 | 4.58E-05 | 2.73E-05 | 0.00E+00 | 0.00E+00 | 3.86E-05 | 2.08E-04 | 7.47E-04 | 3.08E-04 | Known_Unknown |
| p__Armatimonadota | 0.00E+00 | 0.00E+00 | 0.00E+00 | 0.00E+00 | 3.14E-04 | 1.40E-04 | 0.00E+00 | 0.00E+00 | 0.00E+00 | 0.00E+00 | 0.00E+00 | Known_Unknown |
| p__Chloroflexota | 1.96E-03 | 1.81E-03 | 8.22E-05 | 3.43E-05 | 6.28E-04 | 1.17E-04 | 7.70E-04 | 1.17E-03 | 5.64E-04 | 1.58E-03 | 1.26E-03 | Known_Unknown |
| p__Desulfobacterota_I | 2.33E-05 | 0.00E+00 | 0.00E+00 | 2.29E-04 | 0.00E+00 | 0.00E+00 | 0.00E+00 | 0.00E+00 | 4.45E-05 | 3.10E-04 | 4.86E-05 | Known_Unknown |
| p__Firmicutes_A | 0.00E+00 | 0.00E+00 | 0.00E+00 | 2.17E-04 | 5.32E-04 | 0.00E+00 | 0.00E+00 | 4.25E-04 | 0.00E+00 | 4.55E-04 | 1.94E-04 | Known_Unknown |
| p__Firmicutes_B_370522 | 0.00E+00 | 1.06E-03 | 7.16E-04 | 1.26E-04 | 1.36E-04 | 0.00E+00 | 0.00E+00 | 0.00E+00 | 0.00E+00 | 0.00E+00 | 0.00E+00 | Known_Unknown |

| Genus | L1 | L2 | L3 | L4 | L5 | L6 | L7 | L8 | L9 | L10 | L11 | Microbial_matter |
|---|---|---|---|---|---|---|---|---|---|---|---|---|
| p__Firmicutes_B_370539 | 0.00E+00 | 0.00E+00 | 0.00E+00 | 5.72E-05 | 0.00E+00 | 0.00E+00 | 0.00E+00 | 0.00E+00 | 0.00E+00 | 5.46E-05 | 0.00E+00 | Known_Unknown |
| p__Firmicutes_D | 1.36E-02 | 7.85E-03 | 2.23E-03 | 7.19E-03 | 2.81E-03 | 1.40E-02 | 1.37E-01 | 2.90E-02 | 1.08E-03 | 2.55E-03 | 1.11E-02 | Known_Unknown |
| p__Firmicutes_E | 0.00E+00 | 0.00E+00 | 8.57E-04 | 1.26E-03 | 7.23E-04 | 1.84E-03 | 2.17E-03 | 3.35E-04 | 1.19E-04 | 1.09E-04 | 6.48E-04 | Known_Unknown |
| p__Firmicutes_G | 1.40E-04 | 2.04E-04 | 5.17E-04 | 0.00E+00 | 2.21E-03 | 3.85E-04 | 0.00E+00 | 0.00E+00 | 0.00E+00 | 0.00E+00 | 9.72E-05 | Known_Unknown |
| p__Halobacteriota | 0.00E+00 | 0.00E+00 | 0.00E+00 | 0.00E+00 | 0.00E+00 | 0.00E+00 | 8.96E-05 | 0.00E+00 | 0.00E+00 | 0.00E+00 | 0.00E+00 | Known_Unknown |
| p__Myxococcota_A_473307 | 0.00E+00 | 0.00E+00 | 4.70E-05 | 0.00E+00 | 0.00E+00 | 0.00E+00 | 0.00E+00 | 0.00E+00 | 0.00E+00 | 0.00E+00 | 0.00E+00 | Known_Unknown |
| p__Nanoarchaeota | 4.67E-04 | 1.53E-04 | 5.87E-05 | 0.00E+00 | 0.00E+00 | 0.00E+00 | 0.00E+00 | 0.00E+00 | 1.10E-03 | 9.11E-05 | 2.59E-04 | Known_Unknown |
| p__Nitrospirota_A_437815 | 0.00E+00 | 0.00E+00 | 2.35E-05 | 0.00E+00 | 0.00E+00 | 0.00E+00 | 0.00E+00 | 0.00E+00 | 0.00E+00 | 0.00E+00 | 0.00E+00 | Known_Unknown |
| p__Patescibacteria | 2.57E-04 | 4.26E-04 | 9.39E-05 | 3.21E-04 | 2.46E-04 | 2.34E-05 | 7.16E-05 | 2.19E-04 | 5.64E-04 | 2.55E-04 | 4.21E-04 | Known_Unknown |
| p__Proteobacteria | 0.00E+00 | 2.73E-04 | 1.41E-04 | 0.00E+00 | 9.55E-05 | 0.00E+00 | 0.00E+00 | 0.00E+00 | 4.01E-04 | 8.38E-04 | 9.24E-04 | Known_Unknown |
| p__Thermoplasmatota | 0.00E+00 | 2.73E-04 | 1.88E-04 | 0.00E+00 | 4.09E-05 | 2.80E-04 | 3.58E-05 | 1.29E-04 | 1.78E-04 | 0.00E+00 | 8.10E-05 | Known_Unknown |
| p__Thermotogota | 1.63E-04 | 0.00E+00 | 0.00E+00 | 0.00E+00 | 0.00E+00 | 0.00E+00 | 0.00E+00 | 0.00E+00 | 0.00E+00 | 0.00E+00 | 0.00E+00 | Known_Unknown |
| Unassigned | 7.82E-03 | 8.81E-03 | 1.47E-02 | 5.98E-03 | 2.32E-02 | 3.58E-03 | 1.16E-02 | 1.80E-02 | 1.39E-02 | 4.41E-03 | 8.22E-03 | Unknown_Unknown |
| g__ZCTH02-B6 | 0.00E+00 | 8.86E-04 | 4.11E-04 | 3.78E-04 | 0.00E+00 | 9.34E-05 | 8.96E-05 | 0.00E+00 | 0.00E+00 | 0.00E+00 | 1.78E-04 | Known_Known |
| g__Xylanibacillus | 0.00E+00 | 0.00E+00 | 0.00E+00 | 0.00E+00 | 0.00E+00 | 1.17E-05 | 0.00E+00 | 0.00E+00 | 0.00E+00 | 0.00E+00 | 0.00E+00 | Known_Known |
| g__XYA2-FULL-43-10 | 0.00E+00 | 0.00E+00 | 0.00E+00 | 0.00E+00 | 0.00E+00 | 0.00E+00 | 0.00E+00 | 1.80E-04 | 7.42E-05 | 0.00E+00 | 0.00E+00 | Known_Known |
| g__XN24 | 0.00E+00 | 1.02E-04 | 0.00E+00 | 0.00E+00 | 0.00E+00 | 0.00E+00 | 0.00E+00 | 1.29E-04 | 0.00E+00 | 5.46E-05 | 0.00E+00 | Known_Known |
| g__Xenobium | 0.00E+00 | 0.00E+00 | 0.00E+00 | 0.00E+00 | 0.00E+00 | 0.00E+00 | 5.37E-05 | 0.00E+00 | 2.97E-05 | 0.00E+00 | 0.00E+00 | Known_Known |
| g__Wukongibacter | 2.33E-05 | 0.00E+00 | 0.00E+00 | 0.00E+00 | 0.00E+00 | 0.00E+00 | 0.00E+00 | 0.00E+00 | 0.00E+00 | 1.09E-04 | 0.00E+00 | Known_Known |
| g__WRJI01 | 0.00E+00 | 0.00E+00 | 0.00E+00 | 0.00E+00 | 0.00E+00 | 1.17E-05 | 0.00E+00 | 0.00E+00 | 0.00E+00 | 0.00E+00 | 0.00E+00 | Known_Known |
| g__WHTX01 | 0.00E+00 | 0.00E+00 | 0.00E+00 | 0.00E+00 | 0.00E+00 | 4.67E-05 | 0.00E+00 | 0.00E+00 | 1.48E-04 | 0.00E+00 | 0.00E+00 | Known_Known |
| g__WHTN01 | 0.00E+00 | 5.11E-05 | 3.52E-05 | 0.00E+00 | 0.00E+00 | 0.00E+00 | 0.00E+00 | 0.00E+00 | 0.00E+00 | 0.00E+00 | 0.00E+00 | Known_Known |
| g__WHTK01 | 0.00E+00 | 0.00E+00 | 0.00E+00 | 0.00E+00 | 0.00E+00 | 0.00E+00 | 0.00E+00 | 0.00E+00 | 7.42E-05 | 0.00E+00 | 0.00E+00 | Known_Known |
| g__WHTC01 | 0.00E+00 | 0.00E+00 | 3.52E-05 | 0.00E+00 | 8.19E-05 | 0.00E+00 | 0.00E+00 | 0.00E+00 | 1.48E-05 | 0.00E+00 | 0.00E+00 | Known_Known |
| g__W498 | 1.40E-04 | 9.88E-04 | 1.04E-03 | 7.44E-04 | 6.14E-04 | 1.32E-03 | 3.22E-04 | 4.76E-04 | 1.13E-03 | 1.60E-03 | 1.12E-03 | Known_Known |
| g__W-Firmicutes-14 | 0.00E+00 | 0.00E+00 | 9.39E-05 | 0.00E+00 | 0.00E+00 | 0.00E+00 | 0.00E+00 | 0.00E+00 | 0.00E+00 | 0.00E+00 | 0.00E+00 | Known_Known |
| g__VYCW01 | 0.00E+00 | 6.81E-05 | 0.00E+00 | 1.03E-04 | 0.00E+00 | 0.00E+00 | 0.00E+00 | 0.00E+00 | 0.00E+00 | 1.09E-04 | 0.00E+00 | Known_Known |
| g__Vulgatibacter | 7.00E-05 | 0.00E+00 | 8.22E-05 | 3.43E-05 | 0.00E+00 | 0.00E+00 | 0.00E+00 | 0.00E+00 | 0.00E+00 | 0.00E+00 | 0.00E+00 | Known_Known |
| g__Viridilinea | 0.00E+00 | 2.38E-04 | 0.00E+00 | 1.50E-03 | 4.09E-04 | 0.00E+00 | 0.00E+00 | 1.55E-04 | 3.10E-03 | 9.11E-05 | 1.78E-04 | Known_Known |
| g__VFBF01 | 0.00E+00 | 0.00E+00 | 0.00E+00 | 0.00E+00 | 8.19E-05 | 0.00E+00 | 0.00E+00 | 0.00E+00 | 0.00E+00 | 0.00E+00 | 0.00E+00 | Known_Known |
| g__Vermiphilus | 0.00E+00 | 0.00E+00 | 0.00E+00 | 0.00E+00 | 0.00E+00 | 0.00E+00 | 0.00E+00 | 0.00E+00 | 4.45E-05 | 0.00E+00 | 0.00E+00 | Known_Known |
| g__VENG01 | 1.63E-04 | 1.36E-04 | 0.00E+00 | 0.00E+00 | 1.91E-04 | 7.01E-05 | 1.07E-04 | 3.22E-04 | 0.00E+00 | 2.37E-04 | 0.00E+00 | Known_Known |
| g__VEMY01 | 0.00E+00 | 2.21E-04 | 0.00E+00 | 0.00E+00 | 0.00E+00 | 0.00E+00 | 1.00E-03 | 0.00E+00 | 0.00E+00 | 0.00E+00 | 0.00E+00 | Known_Known |
| g__Vampirovibrio | 0.00E+00 | 5.11E-05 | 0.00E+00 | 0.00E+00 | 0.00E+00 | 0.00E+00 | 0.00E+00 | 0.00E+00 | 5.93E-05 | 0.00E+00 | 0.00E+00 | Known_Known |
| g__Vallitalea_A | 0.00E+00 | 0.00E+00 | 0.00E+00 | 0.00E+00 | 0.00E+00 | 0.00E+00 | 0.00E+00 | 7.98E-04 | 4.15E-04 | 0.00E+00 | 3.57E-04 | Known_Known |
| g__UTCFX2 | 0.00E+00 | 0.00E+00 | 0.00E+00 | 0.00E+00 | 0.00E+00 | 0.00E+00 | 0.00E+00 | 0.00E+00 | 1.48E-04 | 0.00E+00 | 0.00E+00 | Known_Known |
| g__URHD0088 | 0.00E+00 | 0.00E+00 | 0.00E+00 | 0.00E+00 | 0.00E+00 | 0.00E+00 | 0.00E+00 | 0.00E+00 | 2.37E-04 | 2.73E-04 | 0.00E+00 | Known_Known |
| g__UNRQ01 | 0.00E+00 | 0.00E+00 | 0.00E+00 | 0.00E+00 | 0.00E+00 | 0.00E+00 | 0.00E+00 | 5.15E-05 | 2.97E-05 | 0.00E+00 | 0.00E+00 | Known_Known |
| g__UM-FILTER-47-13 | 0.00E+00 | 0.00E+00 | 3.52E-05 | 3.43E-04 | 5.46E-05 | 0.00E+00 | 0.00E+00 | 0.00E+00 | 1.04E-04 | 0.00E+00 | 6.48E-05 | Known_Known |
| g__UM-FILTER-43-11 | 1.17E-04 | 0.00E+00 | 0.00E+00 | 0.00E+00 | 0.00E+00 | 0.00E+00 | 0.00E+00 | 0.00E+00 | 5.93E-05 | 0.00E+00 | 0.00E+00 | Known_Known |
| g__UM-FILTER-41-35 | 0.00E+00 | 0.00E+00 | 0.00E+00 | 0.00E+00 | 0.00E+00 | 0.00E+00 | 0.00E+00 | 0.00E+00 | 0.00E+00 | 3.64E-05 | 0.00E+00 | Known_Known |
| g__UBA9971 | 0.00E+00 | 0.00E+00 | 0.00E+00 | 0.00E+00 | 0.00E+00 | 0.00E+00 | 0.00E+00 | 0.00E+00 | 0.00E+00 | 0.00E+00 | 3.24E-05 | Known_Known |
| g__UBA994 | 7.00E-05 | 1.70E-04 | 1.88E-04 | 1.83E-04 | 1.09E-04 | 1.17E-04 | 0.00E+00 | 6.44E-05 | 4.01E-04 | 3.46E-04 | 1.13E-04 | Known_Known |
| g__UBA9936 | 0.00E+00 | 0.00E+00 | 0.00E+00 | 0.00E+00 | 0.00E+00 | 0.00E+00 | 0.00E+00 | 3.86E-05 | 0.00E+00 | 0.00E+00 | 0.00E+00 | Known_Known |
| g__UBA9910 | 0.00E+00 | 0.00E+00 | 3.52E-05 | 0.00E+00 | 0.00E+00 | 0.00E+00 | 0.00E+00 | 0.00E+00 | 0.00E+00 | 0.00E+00 | 0.00E+00 | Known_Known |
| g__UBA9862 | 0.00E+00 | 0.00E+00 | 1.88E-04 | 9.16E-05 | 1.36E-04 | 0.00E+00 | 1.61E-04 | 1.29E-04 | 4.01E-04 | 6.92E-04 | 1.78E-04 | Known_Known |
| g__UBA9664 | 0.00E+00 | 0.00E+00 | 2.35E-05 | 2.86E-04 | 0.00E+00 | 0.00E+00 | 0.00E+00 | 1.16E-04 | 0.00E+00 | 0.00E+00 | 8.10E-05 | Known_Known |

| Genus | L1 | L2 | L3 | L4 | L5 | L6 | L7 | L8 | L9 | L10 | L11 | Microbial_matter |
|---|---|---|---|---|---|---|---|---|---|---|---|---|
| g__UBA9579 | 0.00E+00 | 0.00E+00 | 0.00E+00 | 0.00E+00 | 0.00E+00 | 0.00E+00 | 0.00E+00 | 0.00E+00 | 1.19E-04 | 9.11E-05 | 4.38E-04 | Known_Known |
| g__UBA95 | 0.00E+00 | 0.00E+00 | 0.00E+00 | 0.00E+00 | 0.00E+00 | 0.00E+00 | 0.00E+00 | 0.00E+00 | 5.34E-04 | 9.11E-05 | 6.48E-05 | Known_Known |
| g__UBA9296 | 0.00E+00 | 0.00E+00 | 0.00E+00 | 7.55E-04 | 0.00E+00 | 0.00E+00 | 0.00E+00 | 2.58E-05 | 0.00E+00 | 0.00E+00 | 8.10E-05 | Known_Known |
| g__UBA877 | 0.00E+00 | 3.07E-04 | 0.00E+00 | 0.00E+00 | 0.00E+00 | 0.00E+00 | 0.00E+00 | 0.00E+00 | 0.00E+00 | 0.00E+00 | 0.00E+00 | Known_Known |
| g__UBA876 | 0.00E+00 | 3.07E-04 | 0.00E+00 | 0.00E+00 | 0.00E+00 | 0.00E+00 | 0.00E+00 | 0.00E+00 | 0.00E+00 | 0.00E+00 | 0.00E+00 | Known_Known |
| g__UBA8592 | 0.00E+00 | 0.00E+00 | 0.00E+00 | 0.00E+00 | 0.00E+00 | 0.00E+00 | 0.00E+00 | 0.00E+00 | 4.15E-04 | 0.00E+00 | 0.00E+00 | Known_Known |
| g__UBA8473 | 0.00E+00 | 0.00E+00 | 0.00E+00 | 0.00E+00 | 2.73E-05 | 0.00E+00 | 0.00E+00 | 6.44E-05 | 2.52E-04 | 9.11E-04 | 8.10E-05 | Known_Known |
| g__UBA7879 | 0.00E+00 | 0.00E+00 | 0.00E+00 | 4.35E-04 | 0.00E+00 | 0.00E+00 | 0.00E+00 | 0.00E+00 | 0.00E+00 | 0.00E+00 | 1.30E-04 | Known_Known |
| g__UBA6170 | 0.00E+00 | 2.04E-04 | 0.00E+00 | 0.00E+00 | 0.00E+00 | 0.00E+00 | 0.00E+00 | 0.00E+00 | 2.97E-05 | 0.00E+00 | 0.00E+00 | Known_Known |
| g__UBA6092 | 0.00E+00 | 0.00E+00 | 0.00E+00 | 0.00E+00 | 0.00E+00 | 0.00E+00 | 0.00E+00 | 0.00E+00 | 7.42E-05 | 0.00E+00 | 0.00E+00 | Known_Known |
| g__UBA6055 | 1.63E-04 | 0.00E+00 | 0.00E+00 | 6.87E-05 | 0.00E+00 | 0.00E+00 | 0.00E+00 | 0.00E+00 | 0.00E+00 | 0.00E+00 | 6.48E-05 | Known_Known |
| g__UBA583 | 5.37E-04 | 0.00E+00 | 0.00E+00 | 0.00E+00 | 0.00E+00 | 0.00E+00 | 0.00E+00 | 0.00E+00 | 0.00E+00 | 0.00E+00 | 0.00E+00 | Known_Known |
| g__UBA5793 | 4.67E-05 | 0.00E+00 | 0.00E+00 | 2.29E-05 | 0.00E+00 | 0.00E+00 | 0.00E+00 | 0.00E+00 | 0.00E+00 | 0.00E+00 | 0.00E+00 | Known_Known |
| g__UBA5619 | 0.00E+00 | 0.00E+00 | 0.00E+00 | 0.00E+00 | 0.00E+00 | 0.00E+00 | 0.00E+00 | 3.86E-05 | 0.00E+00 | 0.00E+00 | 0.00E+00 | Known_Known |
| g__UBA5232 | 0.00E+00 | 0.00E+00 | 0.00E+00 | 0.00E+00 | 2.73E-05 | 0.00E+00 | 0.00E+00 | 0.00E+00 | 0.00E+00 | 0.00E+00 | 0.00E+00 | Known_Known |
| g__UBA4782 | 0.00E+00 | 3.07E-04 | 0.00E+00 | 0.00E+00 | 1.09E-04 | 0.00E+00 | 8.96E-05 | 1.03E-04 | 2.97E-04 | 4.19E-04 | 0.00E+00 | Known_Known |
| g__UBA4729 | 4.67E-05 | 0.00E+00 | 0.00E+00 | 0.00E+00 | 0.00E+00 | 0.00E+00 | 0.00E+00 | 0.00E+00 | 4.45E-05 | 0.00E+00 | 0.00E+00 | Known_Known |
| g__UBA4427 | 0.00E+00 | 3.41E-05 | 0.00E+00 | 0.00E+00 | 0.00E+00 | 0.00E+00 | 0.00E+00 | 0.00E+00 | 0.00E+00 | 0.00E+00 | 0.00E+00 | Known_Known |
| g__UBA3254 | 0.00E+00 | 1.70E-04 | 1.29E-04 | 0.00E+00 | 0.00E+00 | 0.00E+00 | 0.00E+00 | 1.29E-05 | 0.00E+00 | 0.00E+00 | 0.00E+00 | Known_Known |
| g__UBA2963 | 0.00E+00 | 8.52E-05 | 0.00E+00 | 0.00E+00 | 0.00E+00 | 0.00E+00 | 0.00E+00 | 0.00E+00 | 5.93E-05 | 0.00E+00 | 4.86E-05 | Known_Known |
| g__UBA2466 | 0.00E+00 | 0.00E+00 | 0.00E+00 | 0.00E+00 | 0.00E+00 | 0.00E+00 | 0.00E+00 | 0.00E+00 | 4.45E-05 | 0.00E+00 | 0.00E+00 | Known_Known |
| g__UBA2284 | 7.00E-05 | 0.00E+00 | 0.00E+00 | 0.00E+00 | 0.00E+00 | 0.00E+00 | 0.00E+00 | 0.00E+00 | 0.00E+00 | 0.00E+00 | 0.00E+00 | Known_Known |
| g__UBA2224 | 0.00E+00 | 0.00E+00 | 0.00E+00 | 0.00E+00 | 0.00E+00 | 0.00E+00 | 0.00E+00 | 5.15E-05 | 0.00E+00 | 0.00E+00 | 0.00E+00 | Known_Known |
| g__UBA2200 | 0.00E+00 | 5.11E-05 | 0.00E+00 | 0.00E+00 | 0.00E+00 | 0.00E+00 | 0.00E+00 | 0.00E+00 | 6.53E-04 | 2.19E-04 | 1.46E-04 | Known_Known |
| g__UBA2174 | 0.00E+00 | 0.00E+00 | 3.52E-05 | 8.01E-05 | 2.73E-05 | 0.00E+00 | 0.00E+00 | 0.00E+00 | 2.97E-05 | 0.00E+00 | 0.00E+00 | Known_Known |
| g__UBA2156 | 0.00E+00 | 0.00E+00 | 0.00E+00 | 0.00E+00 | 0.00E+00 | 0.00E+00 | 0.00E+00 | 0.00E+00 | 0.00E+00 | 5.46E-05 | 0.00E+00 | Known_Known |
| g__UBA2020 | 0.00E+00 | 0.00E+00 | 0.00E+00 | 4.58E-05 | 0.00E+00 | 0.00E+00 | 0.00E+00 | 0.00E+00 | 2.97E-05 | 0.00E+00 | 0.00E+00 | Known_Known |
| g__UBA147 | 4.67E-05 | 2.21E-04 | 0.00E+00 | 2.29E-05 | 0.00E+00 | 0.00E+00 | 0.00E+00 | 7.21E-04 | 4.90E-03 | 6.81E-03 | 1.83E-03 | Known_Known |
| g__UBA1369 | 0.00E+00 | 6.81E-05 | 0.00E+00 | 0.00E+00 | 0.00E+00 | 0.00E+00 | 0.00E+00 | 7.73E-05 | 2.97E-05 | 1.27E-04 | 0.00E+00 | Known_Known |
| g__UBA12474 | 0.00E+00 | 0.00E+00 | 0.00E+00 | 0.00E+00 | 0.00E+00 | 3.50E-05 | 0.00E+00 | 0.00E+00 | 0.00E+00 | 0.00E+00 | 0.00E+00 | Known_Known |
| g__UBA12044 | 1.17E-04 | 3.41E-05 | 0.00E+00 | 9.50E-04 | 0.00E+00 | 8.17E-05 | 5.37E-04 | 1.09E-03 | 3.12E-04 | 1.46E-04 | 3.08E-04 | Known_Known |
| g__UBA12014 | 0.00E+00 | 0.00E+00 | 0.00E+00 | 0.00E+00 | 2.73E-05 | 0.00E+00 | 0.00E+00 | 0.00E+00 | 0.00E+00 | 0.00E+00 | 0.00E+00 | Known_Known |
| g__UBA1162 | 1.17E-04 | 0.00E+00 | 0.00E+00 | 0.00E+00 | 0.00E+00 | 0.00E+00 | 0.00E+00 | 1.29E-04 | 3.41E-04 | 2.73E-04 | 6.48E-05 | Known_Known |
| g__UBA1140 | 0.00E+00 | 0.00E+00 | 0.00E+00 | 0.00E+00 | 0.00E+00 | 0.00E+00 | 0.00E+00 | 0.00E+00 | 0.00E+00 | 0.00E+00 | 6.48E-05 | Known_Known |
| g__UBA10424 | 0.00E+00 | 0.00E+00 | 0.00E+00 | 6.41E-04 | 0.00E+00 | 0.00E+00 | 0.00E+00 | 0.00E+00 | 0.00E+00 | 0.00E+00 | 0.00E+00 | Known_Known |
| g__Tumebacillus_A | 0.00E+00 | 0.00E+00 | 0.00E+00 | 0.00E+00 | 8.19E-05 | 0.00E+00 | 0.00E+00 | 0.00E+00 | 0.00E+00 | 5.46E-05 | 0.00E+00 | Known_Known |
| g__Truepera | 0.00E+00 | 4.94E-04 | 1.29E-04 | 0.00E+00 | 2.05E-04 | 1.05E-04 | 0.00E+00 | 0.00E+00 | 0.00E+00 | 3.64E-05 | 8.10E-05 | Known_Known |
| g__Tropicimonas_492701 | 0.00E+00 | 0.00E+00 | 0.00E+00 | 1.60E-04 | 0.00E+00 | 0.00E+00 | 0.00E+00 | 0.00E+00 | 0.00E+00 | 0.00E+00 | 0.00E+00 | Known_Known |
| g__TMED27 | 0.00E+00 | 0.00E+00 | 0.00E+00 | 1.03E-04 | 0.00E+00 | 0.00E+00 | 0.00E+00 | 0.00E+00 | 0.00E+00 | 0.00E+00 | 0.00E+00 | Known_Known |
| g__TMED2 | 0.00E+00 | 0.00E+00 | 0.00E+00 | 9.16E-05 | 0.00E+00 | 0.00E+00 | 0.00E+00 | 0.00E+00 | 0.00E+00 | 0.00E+00 | 0.00E+00 | Known_Known |
| g__Tindallia | 2.33E-05 | 3.41E-04 | 0.00E+00 | 0.00E+00 | 0.00E+00 | 0.00E+00 | 0.00E+00 | 2.06E-04 | 0.00E+00 | 0.00E+00 | 0.00E+00 | Known_Known |
| g__Thiomicrospira | 7.00E-05 | 1.19E-04 | 4.70E-05 | 0.00E+00 | 0.00E+00 | 0.00E+00 | 0.00E+00 | 0.00E+00 | 0.00E+00 | 1.46E-04 | 4.86E-05 | Known_Known |
| g__Thiolinea | 0.00E+00 | 0.00E+00 | 0.00E+00 | 0.00E+00 | 0.00E+00 | 0.00E+00 | 0.00E+00 | 0.00E+00 | 0.00E+00 | 0.00E+00 | 3.24E-05 | Known_Known |
| g__Thioflavicoccus | 0.00E+00 | 0.00E+00 | 0.00E+00 | 3.43E-05 | 0.00E+00 | 0.00E+00 | 0.00E+00 | 0.00E+00 | 0.00E+00 | 0.00E+00 | 0.00E+00 | Known_Known |
| g__Thermosulfidibacter | 0.00E+00 | 1.87E-04 | 0.00E+00 | 0.00E+00 | 0.00E+00 | 0.00E+00 | 0.00E+00 | 1.16E-04 | 1.34E-04 | 2.48E-03 | 1.13E-04 | Known_Known |
| g__Thalassospira | 0.00E+00 | 0.00E+00 | 0.00E+00 | 0.00E+00 | 0.00E+00 | 0.00E+00 | 0.00E+00 | 0.00E+00 | 0.00E+00 | 1.09E-04 | 9.72E-05 | Known_Known |
| g__Thalassobaculum_A_508031 | 0.00E+00 | 0.00E+00 | 0.00E+00 | 0.00E+00 | 0.00E+00 | 0.00E+00 | 0.00E+00 | 0.00E+00 | 0.00E+00 | 2.19E-04 | 0.00E+00 | Known_Known |

| Genus | L1 | L2 | L3 | L4 | L5 | L6 | L7 | L8 | L9 | L10 | L11 | Microbial_matter |
|---|---|---|---|---|---|---|---|---|---|---|---|---|
| g__Tepidiforma | 0.00E+00 | 0.00E+00 | 0.00E+00 | 0.00E+00 | 0.00E+00 | 0.00E+00 | 0.00E+00 | 3.86E-05 | 0.00E+00 | 0.00E+00 | 0.00E+00 | Known_Known |
| g__Tepidibaculum | 0.00E+00 | 0.00E+00 | 0.00E+00 | 0.00E+00 | 0.00E+00 | 0.00E+00 | 0.00E+00 | 0.00E+00 | 0.00E+00 | 0.00E+00 | 4.86E-05 | Known_Known |
| g__T3SED10-354 | 2.10E-04 | 6.81E-05 | 0.00E+00 | 0.00E+00 | 0.00E+00 | 0.00E+00 | 0.00E+00 | 0.00E+00 | 8.90E-05 | 5.46E-05 | 9.72E-05 | Known_Known |
| g__SZUA-442 | 2.33E-05 | 0.00E+00 | 0.00E+00 | 0.00E+00 | 0.00E+00 | 0.00E+00 | 0.00E+00 | 0.00E+00 | 0.00E+00 | 0.00E+00 | 0.00E+00 | Known_Known |
| g__SZUA-229 | 0.00E+00 | 0.00E+00 | 0.00E+00 | 0.00E+00 | 0.00E+00 | 0.00E+00 | 0.00E+00 | 0.00E+00 | 0.00E+00 | 0.00E+00 | 2.11E-04 | Known_Known |
| g__SZUA-161 | 1.17E-04 | 8.52E-05 | 0.00E+00 | 0.00E+00 | 0.00E+00 | 0.00E+00 | 0.00E+00 | 6.57E-04 | 3.03E-03 | 1.13E-02 | 3.26E-03 | Known_Known |
| g__SZUA-149 | 0.00E+00 | 0.00E+00 | 0.00E+00 | 0.00E+00 | 0.00E+00 | 0.00E+00 | 0.00E+00 | 0.00E+00 | 0.00E+00 | 7.29E-05 | 0.00E+00 | Known_Known |
| g__Syntrophobacter_452395 | 0.00E+00 | 0.00E+00 | 0.00E+00 | 0.00E+00 | 2.73E-05 | 0.00E+00 | 0.00E+00 | 0.00E+00 | 0.00E+00 | 0.00E+00 | 0.00E+00 | Known_Known |
| g__Symbiobacterium | 0.00E+00 | 5.11E-05 | 0.00E+00 | 0.00E+00 | 4.09E-05 | 0.00E+00 | 3.58E-05 | 0.00E+00 | 0.00E+00 | 0.00E+00 | 0.00E+00 | Known_Known |
| g__SYFI01 | 0.00E+00 | 0.00E+00 | 0.00E+00 | 0.00E+00 | 0.00E+00 | 0.00E+00 | 0.00E+00 | 0.00E+00 | 0.00E+00 | 1.82E-05 | 0.00E+00 | Known_Known |
| g__Sutcliffiella_A | 0.00E+00 | 0.00E+00 | 0.00E+00 | 0.00E+00 | 0.00E+00 | 0.00E+00 | 0.00E+00 | 0.00E+00 | 0.00E+00 | 0.00E+00 | 3.24E-05 | Known_Known |
| g__SURF-21 | 0.00E+00 | 0.00E+00 | 0.00E+00 | 0.00E+00 | 0.00E+00 | 0.00E+00 | 0.00E+00 | 0.00E+00 | 3.26E-04 | 5.28E-04 | 6.48E-05 | Known_Known |
| g__Sulfurovum | 0.00E+00 | 5.28E-04 | 8.22E-05 | 6.87E-05 | 0.00E+00 | 0.00E+00 | 0.00E+00 | 0.00E+00 | 0.00E+00 | 0.00E+00 | 0.00E+00 | Known_Known |
| g__Sulfurospirillum_A | 0.00E+00 | 0.00E+00 | 0.00E+00 | 6.30E-04 | 0.00E+00 | 0.00E+00 | 0.00E+00 | 0.00E+00 | 0.00E+00 | 0.00E+00 | 3.89E-04 | Known_Known |
| g__Sulfurimonas | 8.63E-04 | 1.20E-02 | 1.14E-01 | 1.70E-01 | 4.28E-03 | 0.00E+00 | 0.00E+00 | 3.86E-05 | 0.00E+00 | 1.10E-02 | 4.85E-03 | Known_Known |
| g__Stella | 0.00E+00 | 0.00E+00 | 0.00E+00 | 1.49E-04 | 0.00E+00 | 0.00E+00 | 0.00E+00 | 0.00E+00 | 0.00E+00 | 0.00E+00 | 0.00E+00 | Known_Known |
| g__Stappia | 2.10E-04 | 0.00E+00 | 0.00E+00 | 0.00E+00 | 0.00E+00 | 0.00E+00 | 0.00E+00 | 0.00E+00 | 0.00E+00 | 0.00E+00 | 0.00E+00 | Known_Known |
| g__SR-FBR-E99 | 0.00E+00 | 0.00E+00 | 2.35E-05 | 0.00E+00 | 0.00E+00 | 0.00E+00 | 0.00E+00 | 0.00E+00 | 0.00E+00 | 0.00E+00 | 0.00E+00 | Known_Known |
| g__SpSt-899 | 0.00E+00 | 0.00E+00 | 0.00E+00 | 0.00E+00 | 2.73E-05 | 0.00E+00 | 0.00E+00 | 3.99E-04 | 5.93E-05 | 1.09E-04 | 3.24E-05 | Known_Known |
| g__SpSt-77 | 1.12E-03 | 1.04E-03 | 0.00E+00 | 0.00E+00 | 0.00E+00 | 9.34E-05 | 3.40E-04 | 7.60E-04 | 4.01E-04 | 5.28E-04 | 4.86E-04 | Known_Known |
| g__Sporotomaculum | 4.67E-05 | 1.87E-04 | 1.41E-04 | 9.16E-05 | 2.73E-05 | 3.50E-05 | 0.00E+00 | 0.00E+00 | 0.00E+00 | 0.00E+00 | 0.00E+00 | Known_Known |
| g__Sporosalibacterium | 3.97E-03 | 5.88E-03 | 3.89E-03 | 2.19E-03 | 7.37E-04 | 1.17E-03 | 8.78E-04 | 7.53E-03 | 1.04E-02 | 2.41E-02 | 1.62E-02 | Known_Known |
| g__Sporobacter | 0.00E+00 | 0.00E+00 | 5.87E-05 | 0.00E+00 | 1.77E-04 | 0.00E+00 | 7.16E-05 | 0.00E+00 | 0.00E+00 | 0.00E+00 | 0.00E+00 | Known_Known |
| g__Sporacetigenium | 0.00E+00 | 0.00E+00 | 0.00E+00 | 0.00E+00 | 0.00E+00 | 7.01E-05 | 0.00E+00 | 0.00E+00 | 0.00E+00 | 0.00E+00 | 0.00E+00 | Known_Known |
| g__Spiro-10 | 0.00E+00 | 0.00E+00 | 3.52E-05 | 0.00E+00 | 0.00E+00 | 0.00E+00 | 0.00E+00 | 0.00E+00 | 0.00E+00 | 0.00E+00 | 0.00E+00 | Known_Known |
| g__Sphingomonas_L_486704 | 0.00E+00 | 2.55E-04 | 0.00E+00 | 0.00E+00 | 2.73E-05 | 0.00E+00 | 0.00E+00 | 0.00E+00 | 0.00E+00 | 0.00E+00 | 0.00E+00 | Known_Known |
| g__Sphingomicrobium_483265 | 9.33E-05 | 3.24E-04 | 2.00E-04 | 6.87E-05 | 5.05E-04 | 6.19E-04 | 1.43E-04 | 0.00E+00 | 0.00E+00 | 0.00E+00 | 0.00E+00 | Known_Known |
| g__Sphingoaurantiacus | 9.33E-05 | 0.00E+00 | 0.00E+00 | 0.00E+00 | 0.00E+00 | 0.00E+00 | 0.00E+00 | 0.00E+00 | 0.00E+00 | 0.00E+00 | 0.00E+00 | Known_Known |
| g__Sphaerobacter | 9.33E-05 | 1.36E-04 | 0.00E+00 | 0.00E+00 | 0.00E+00 | 4.67E-05 | 0.00E+00 | 0.00E+00 | 8.90E-05 | 0.00E+00 | 0.00E+00 | Known_Known |
| g__Solincola_A_402975 | 0.00E+00 | 0.00E+00 | 0.00E+00 | 0.00E+00 | 0.00E+00 | 0.00E+00 | 0.00E+00 | 0.00E+00 | 2.97E-05 | 0.00E+00 | 0.00E+00 | Known_Known |
| g__SM23-78 | 0.00E+00 | 0.00E+00 | 2.35E-05 | 3.43E-05 | 0.00E+00 | 0.00E+00 | 3.58E-05 | 0.00E+00 | 1.34E-04 | 0.00E+00 | 9.72E-05 | Known_Known |
| g__SM1-50 | 1.98E-01 | 9.25E-02 | 9.19E-02 | 1.80E-01 | 5.16E-02 | 1.36E-01 | 2.51E-01 | 2.75E-01 | 2.94E-01 | 1.91E-01 | 1.63E-01 | Known_Known |
| g__SLQV01 | 0.00E+00 | 0.00E+00 | 0.00E+00 | 0.00E+00 | 0.00E+00 | 0.00E+00 | 0.00E+00 | 0.00E+00 | 0.00E+00 | 5.46E-05 | 0.00E+00 | Known_Known |
| g__SLAK01 | 0.00E+00 | 6.81E-05 | 0.00E+00 | 0.00E+00 | 2.73E-05 | 0.00E+00 | 0.00E+00 | 0.00E+00 | 0.00E+00 | 0.00E+00 | 0.00E+00 | Known_Known |
| g__SKVJ01 | 7.70E-04 | 1.43E-03 | 8.33E-04 | 9.27E-04 | 3.41E-04 | 1.07E-03 | 4.30E-04 | 3.05E-03 | 1.78E-03 | 3.95E-03 | 1.38E-03 | Known_Known |
| g__SKLG01 | 8.40E-03 | 1.97E-02 | 4.26E-03 | 5.53E-03 | 8.05E-03 | 1.94E-03 | 9.85E-04 | 1.31E-03 | 1.12E-02 | 4.26E-03 | 5.90E-03 | Known_Known |
| g__SKKC01 | 7.00E-05 | 0.00E+00 | 1.06E-04 | 0.00E+00 | 2.73E-05 | 3.50E-05 | 0.00E+00 | 0.00E+00 | 0.00E+00 | 5.46E-05 | 0.00E+00 | Known_Known |
| g__Skermanella | 1.63E-04 | 6.18E-03 | 1.17E-03 | 0.00E+00 | 1.30E-03 | 2.16E-03 | 6.81E-04 | 3.09E-04 | 5.79E-04 | 8.01E-04 | 5.19E-04 | Known_Known |
| g__SIRW01 | 0.00E+00 | 1.02E-04 | 0.00E+00 | 0.00E+00 | 0.00E+00 | 3.50E-05 | 0.00E+00 | 3.86E-05 | 1.63E-04 | 5.28E-04 | 2.92E-04 | Known_Known |
| g__SIO2C1 | 7.00E-05 | 0.00E+00 | 0.00E+00 | 5.04E-04 | 0.00E+00 | 0.00E+00 | 0.00E+00 | 3.86E-05 | 0.00E+00 | 0.00E+00 | 0.00E+00 | Known_Known |
| g__SHYG01 | 0.00E+00 | 0.00E+00 | 0.00E+00 | 0.00E+00 | 0.00E+00 | 0.00E+00 | 0.00E+00 | 0.00E+00 | 0.00E+00 | 1.09E-04 | 0.00E+00 | Known_Known |
| g__SHVA01 | 0.00E+00 | 0.00E+00 | 0.00E+00 | 0.00E+00 | 0.00E+00 | 0.00E+00 | 0.00E+00 | 0.00E+00 | 0.00E+00 | 3.64E-05 | 0.00E+00 | Known_Known |
| g__SHUY01 | 0.00E+00 | 0.00E+00 | 0.00E+00 | 0.00E+00 | 0.00E+00 | 0.00E+00 | 0.00E+00 | 7.73E-05 | 1.04E-04 | 2.91E-04 | 6.48E-05 | Known_Known |
| g__Shivajiella | 0.00E+00 | 0.00E+00 | 0.00E+00 | 0.00E+00 | 1.36E-05 | 0.00E+00 | 0.00E+00 | 0.00E+00 | 0.00E+00 | 0.00E+00 | 0.00E+00 | Known_Known |
| g__Shimazuella | 0.00E+00 | 0.00E+00 | 0.00E+00 | 0.00E+00 | 0.00E+00 | 7.01E-05 | 0.00E+00 | 0.00E+00 | 0.00E+00 | 0.00E+00 | 0.00E+00 | Known_Known |
| g__SG8-5 | 0.00E+00 | 0.00E+00 | 0.00E+00 | 0.00E+00 | 0.00E+00 | 0.00E+00 | 0.00E+00 | 0.00E+00 | 1.19E-04 | 0.00E+00 | 0.00E+00 | Known_Known |
| g__SG-bin7 | 0.00E+00 | 5.11E-05 | 0.00E+00 | 5.72E-05 | 0.00E+00 | 0.00E+00 | 0.00E+00 | 0.00E+00 | 1.19E-04 | 0.00E+00 | 3.24E-05 | Known_Known |

| Genus | L1 | L2 | L3 | L4 | L5 | L6 | L7 | L8 | L9 | L10 | L11 | Microbial_matter |
|---|---|---|---|---|---|---|---|---|---|---|---|---|
| g__Senegalia | 2.45E-03 | 8.52E-04 | 4.81E-04 | 4.46E-04 | 1.26E-03 | 2.59E-03 | 7.52E-04 | 2.64E-03 | 1.16E-02 | 3.97E-03 | 4.76E-03 | Known_Known |
| g__Sediminibacillus_A | 0.00E+00 | 0.00E+00 | 0.00E+00 | 0.00E+00 | 4.09E-05 | 1.17E-05 | 0.00E+00 | 0.00E+00 | 4.45E-05 | 0.00E+00 | 0.00E+00 | Known_Known |
| g__SDRK01 | 0.00E+00 | 0.00E+00 | 0.00E+00 | 0.00E+00 | 2.73E-05 | 0.00E+00 | 0.00E+00 | 0.00E+00 | 0.00E+00 | 0.00E+00 | 0.00E+00 | Known_Known |
| g__SDNM01 | 4.67E-05 | 1.36E-04 | 9.39E-05 | 2.17E-04 | 2.73E-05 | 0.00E+00 | 0.00E+00 | 0.00E+00 | 3.12E-04 | 2.91E-04 | 9.72E-05 | Known_Known |
| g__SCUD01 | 0.00E+00 | 0.00E+00 | 0.00E+00 | 1.14E-05 | 0.00E+00 | 0.00E+00 | 0.00E+00 | 0.00E+00 | 0.00E+00 | 0.00E+00 | 0.00E+00 | Known_Known |
| g__SCSU01 | 0.00E+00 | 0.00E+00 | 0.00E+00 | 0.00E+00 | 0.00E+00 | 0.00E+00 | 0.00E+00 | 0.00E+00 | 0.00E+00 | 1.09E-04 | 9.72E-05 | Known_Known |
| g__SAR202-VII-2 | 0.00E+00 | 0.00E+00 | 0.00E+00 | 0.00E+00 | 4.09E-05 | 0.00E+00 | 0.00E+00 | 0.00E+00 | 0.00E+00 | 0.00E+00 | 0.00E+00 | Known_Known |
| g__Sandarakinorhabdus | 0.00E+00 | 0.00E+00 | 0.00E+00 | 4.58E-05 | 0.00E+00 | 0.00E+00 | 0.00E+00 | 0.00E+00 | 0.00E+00 | 0.00E+00 | 0.00E+00 | Known_Known |
| g__Sandaracinobacter | 0.00E+00 | 0.00E+00 | 0.00E+00 | 1.03E-04 | 0.00E+00 | 0.00E+00 | 0.00E+00 | 0.00E+00 | 0.00E+00 | 0.00E+00 | 0.00E+00 | Known_Known |
| g__Salirhabdus_286857 | 0.00E+00 | 0.00E+00 | 0.00E+00 | 0.00E+00 | 0.00E+00 | 0.00E+00 | 0.00E+00 | 0.00E+00 | 0.00E+00 | 5.46E-05 | 0.00E+00 | Known_Known |
| g__Saliphagus | 2.30E-02 | 4.61E-02 | 2.86E-02 | 2.96E-03 | 1.72E-03 | 2.53E-03 | 3.11E-02 | 3.61E-04 | 6.23E-04 | 8.20E-04 | 9.56E-03 | Known_Known |
| g__Salinivirga | 4.67E-05 | 5.11E-05 | 0.00E+00 | 0.00E+00 | 0.00E+00 | 0.00E+00 | 0.00E+00 | 0.00E+00 | 0.00E+00 | 0.00E+00 | 0.00E+00 | Known_Known |
| g__Salinigranum | 0.00E+00 | 1.29E-03 | 9.16E-04 | 1.95E-04 | 4.09E-05 | 4.67E-05 | 0.00E+00 | 0.00E+00 | 7.42E-05 | 0.00E+00 | 1.78E-04 | Known_Known |
| g__Salinibaculum | 0.00E+00 | 0.00E+00 | 2.35E-04 | 0.00E+00 | 0.00E+00 | 0.00E+00 | 0.00E+00 | 0.00E+00 | 0.00E+00 | 0.00E+00 | 0.00E+00 | Known_Known |
| g__Salinarimonas | 4.67E-03 | 7.49E-03 | 4.26E-02 | 1.02E-02 | 4.78E-04 | 4.31E-03 | 4.32E-03 | 9.01E-04 | 0.00E+00 | 7.65E-04 | 5.14E-03 | Known_Known |
| g__Salinarchaeum | 2.33E-04 | 5.62E-04 | 1.10E-03 | 6.87E-05 | 2.32E-04 | 1.17E-04 | 0.00E+00 | 0.00E+00 | 1.78E-04 | 0.00E+00 | 0.00E+00 | Known_Known |
| g__S5-A14a | 0.00E+00 | 0.00E+00 | 9.39E-05 | 0.00E+00 | 1.64E-04 | 0.00E+00 | 0.00E+00 | 0.00E+00 | 0.00E+00 | 5.46E-05 | 0.00E+00 | Known_Known |
| g__S20-B6 | 0.00E+00 | 1.07E-03 | 6.57E-04 | 0.00E+00 | 2.87E-04 | 7.24E-04 | 8.96E-05 | 0.00E+00 | 0.00E+00 | 1.46E-04 | 2.92E-04 | Known_Known |
| g__RZZM01 | 0.00E+00 | 0.00E+00 | 0.00E+00 | 5.15E-04 | 0.00E+00 | 0.00E+00 | 0.00E+00 | 5.15E-05 | 0.00E+00 | 0.00E+00 | 0.00E+00 | Known_Known |
| g__Ruminiclostridium | 8.63E-04 | 1.33E-03 | 4.58E-04 | 8.13E-04 | 1.53E-02 | 3.83E-03 | 1.45E-03 | 4.67E-03 | 1.96E-03 | 2.53E-03 | 6.06E-03 | Known_Known |
| g__RUG658 | 0.00E+00 | 0.00E+00 | 0.00E+00 | 0.00E+00 | 0.00E+00 | 0.00E+00 | 0.00E+00 | 0.00E+00 | 1.19E-04 | 1.27E-04 | 0.00E+00 | Known_Known |
| g__Rubrobacter_A_405431 | 0.00E+00 | 0.00E+00 | 0.00E+00 | 0.00E+00 | 0.00E+00 | 0.00E+00 | 0.00E+00 | 0.00E+00 | 4.45E-05 | 7.29E-05 | 0.00E+00 | Known_Known |
| g__Rubrimonas | 3.50E-04 | 2.55E-04 | 1.41E-04 | 9.39E-04 | 0.00E+00 | 0.00E+00 | 0.00E+00 | 1.93E-04 | 2.82E-04 | 0.00E+00 | 0.00E+00 | Known_Known |
| g__Rubribacterium | 3.43E-03 | 2.25E-03 | 4.46E-03 | 4.31E-02 | 1.80E-03 | 1.18E-03 | 2.45E-03 | 5.74E-03 | 5.93E-05 | 2.33E-03 | 3.63E-03 | Known_Known |
| g__Rubellimicrobium | 0.00E+00 | 5.06E-03 | 3.04E-03 | 1.37E-04 | 3.41E-04 | 1.06E-03 | 1.79E-04 | 0.00E+00 | 0.00E+00 | 0.00E+00 | 4.54E-04 | Known_Known |
| g__RSA9 | 0.00E+00 | 0.00E+00 | 0.00E+00 | 3.43E-05 | 0.00E+00 | 0.00E+00 | 0.00E+00 | 0.00E+00 | 0.00E+00 | 0.00E+00 | 0.00E+00 | Known_Known |
| g__Roseomonas_A_507058 | 0.00E+00 | 1.53E-04 | 1.29E-04 | 0.00E+00 | 0.00E+00 | 2.22E-04 | 0.00E+00 | 0.00E+00 | 0.00E+00 | 7.29E-05 | 0.00E+00 | Known_Known |
| g__Roseinatronobacter | 3.03E-04 | 0.00E+00 | 4.97E-03 | 8.88E-03 | 0.00E+00 | 3.39E-04 | 1.25E-04 | 0.00E+00 | 0.00E+00 | 0.00E+00 | 3.08E-04 | Known_Known |
| g__Roseicyclus | 0.00E+00 | 4.43E-04 | 0.00E+00 | 9.04E-04 | 0.00E+00 | 0.00E+00 | 1.43E-04 | 6.44E-05 | 0.00E+00 | 0.00E+00 | 0.00E+00 | Known_Known |
| g__Roseibium_497070 | 0.00E+00 | 0.00E+00 | 0.00E+00 | 2.98E-04 | 0.00E+00 | 0.00E+00 | 0.00E+00 | 2.96E-04 | 0.00E+00 | 0.00E+00 | 0.00E+00 | Known_Known |
| g__Romboutsia_B | 0.00E+00 | 0.00E+00 | 0.00E+00 | 0.00E+00 | 0.00E+00 | 0.00E+00 | 0.00E+00 | 0.00E+00 | 8.90E-05 | 0.00E+00 | 0.00E+00 | Known_Known |
| g__Robertmurraya_298194 | 0.00E+00 | 0.00E+00 | 0.00E+00 | 0.00E+00 | 1.09E-04 | 0.00E+00 | 0.00E+00 | 0.00E+00 | 0.00E+00 | 0.00E+00 | 0.00E+00 | Known_Known |
| g__Risungbinella | 0.00E+00 | 0.00E+00 | 0.00E+00 | 0.00E+00 | 0.00E+00 | 0.00E+00 | 0.00E+00 | 3.86E-05 | 0.00E+00 | 0.00E+00 | 0.00E+00 | Known_Known |
| g__Rhodovulum | 7.93E-04 | 6.47E-04 | 0.00E+00 | 2.38E-02 | 1.43E-03 | 1.28E-04 | 2.33E-04 | 1.64E-03 | 0.00E+00 | 2.37E-04 | 4.86E-04 | Known_Known |
| g__Rhodospira | 9.33E-05 | 0.00E+00 | 0.00E+00 | 2.63E-04 | 0.00E+00 | 0.00E+00 | 0.00E+00 | 5.15E-05 | 0.00E+00 | 0.00E+00 | 0.00E+00 | Known_Known |
| g__Rhodosalinus | 0.00E+00 | 3.24E-04 | 3.52E-03 | 4.35E-04 | 3.82E-04 | 1.31E-03 | 1.06E-03 | 0.00E+00 | 0.00E+00 | 0.00E+00 | 1.13E-04 | Known_Known |
| g__Rhodohalobacter | 0.00E+00 | 3.41E-05 | 0.00E+00 | 0.00E+00 | 0.00E+00 | 2.34E-05 | 0.00E+00 | 0.00E+00 | 2.97E-05 | 0.00E+00 | 0.00E+00 | Known_Known |
| g__Rhodobaculum | 0.00E+00 | 0.00E+00 | 3.39E-03 | 9.74E-03 | 0.00E+00 | 0.00E+00 | 0.00E+00 | 0.00E+00 | 0.00E+00 | 0.00E+00 | 0.00E+00 | Known_Known |
| g__Rhabdanaerobium | 0.00E+00 | 1.53E-04 | 0.00E+00 | 1.26E-04 | 0.00E+00 | 0.00E+00 | 0.00E+00 | 0.00E+00 | 0.00E+00 | 0.00E+00 | 0.00E+00 | Known_Known |
| g__Reyranella | 7.00E-05 | 0.00E+00 | 0.00E+00 | 0.00E+00 | 0.00E+00 | 0.00E+00 | 0.00E+00 | 0.00E+00 | 0.00E+00 | 0.00E+00 | 0.00E+00 | Known_Known |
| g__REEP01 | 7.00E-05 | 0.00E+00 | 0.00E+00 | 0.00E+00 | 0.00E+00 | 0.00E+00 | 0.00E+00 | 0.00E+00 | 0.00E+00 | 0.00E+00 | 0.00E+00 | Known_Known |
| g__REDI01 | 0.00E+00 | 4.94E-04 | 2.35E-04 | 1.14E-04 | 1.09E-04 | 4.67E-05 | 7.16E-05 | 3.48E-04 | 1.54E-03 | 2.50E-03 | 8.10E-04 | Known_Known |
| g__REDH01 | 0.00E+00 | 1.36E-04 | 0.00E+00 | 0.00E+00 | 0.00E+00 | 4.67E-05 | 0.00E+00 | 0.00E+00 | 1.19E-04 | 0.00E+00 | 0.00E+00 | Known_Known |
| g__RBG-16-45-19 | 0.00E+00 | 0.00E+00 | 0.00E+00 | 0.00E+00 | 0.00E+00 | 0.00E+00 | 0.00E+00 | 0.00E+00 | 0.00E+00 | 0.00E+00 | 3.24E-05 | Known_Known |
| g__RBG-13-50-9 | 0.00E+00 | 0.00E+00 | 0.00E+00 | 0.00E+00 | 0.00E+00 | 0.00E+00 | 0.00E+00 | 0.00E+00 | 0.00E+00 | 5.46E-05 | 0.00E+00 | Known_Known |
| g__RBC038 | 0.00E+00 | 1.02E-04 | 4.70E-05 | 0.00E+00 | 0.00E+00 | 5.84E-05 | 0.00E+00 | 0.00E+00 | 0.00E+00 | 0.00E+00 | 0.00E+00 | Known_Known |
| g__QPJE01 | 0.00E+00 | 0.00E+00 | 2.35E-05 | 0.00E+00 | 0.00E+00 | 0.00E+00 | 0.00E+00 | 0.00E+00 | 0.00E+00 | 0.00E+00 | 0.00E+00 | Known_Known |

| Genus | L1 | L2 | L3 | L4 | L5 | L6 | L7 | L8 | L9 | L10 | L11 | Microbial_matter |
|---|---|---|---|---|---|---|---|---|---|---|---|---|
| g__QALR01 | 0.00E+00 | 0.00E+00 | 0.00E+00 | 0.00E+00 | 0.00E+00 | 0.00E+00 | 0.00E+00 | 0.00E+00 | 0.00E+00 | 3.64E-05 | 0.00E+00 | Known_Known |
| g__PXFI01 | 0.00E+00 | 0.00E+00 | 0.00E+00 | 0.00E+00 | 0.00E+00 | 0.00E+00 | 0.00E+00 | 1.29E-05 | 0.00E+00 | 1.82E-04 | 0.00E+00 | Known_Known |
| g__PWVO01 | 2.57E-04 | 3.41E-05 | 0.00E+00 | 0.00E+00 | 2.73E-05 | 0.00E+00 | 0.00E+00 | 3.99E-04 | 3.26E-04 | 1.64E-04 | 4.86E-05 | Known_Known |
| g__PWVB01 | 2.80E-04 | 3.75E-04 | 0.00E+00 | 2.29E-05 | 5.73E-04 | 0.00E+00 | 0.00E+00 | 7.73E-05 | 1.63E-04 | 0.00E+00 | 0.00E+00 | Known_Known |
| g__PWUC01 | 3.24E-03 | 1.14E-02 | 1.91E-02 | 4.44E-02 | 2.33E-02 | 2.20E-02 | 4.86E-02 | 7.63E-02 | 1.58E-01 | 9.30E-02 | 5.90E-02 | Known_Known |
| g__PUND01 | 0.00E+00 | 0.00E+00 | 0.00E+00 | 0.00E+00 | 0.00E+00 | 0.00E+00 | 3.58E-05 | 0.00E+00 | 0.00E+00 | 0.00E+00 | 0.00E+00 | Known_Known |
| g__Pseudovibrio | 0.00E+00 | 0.00E+00 | 0.00E+00 | 6.87E-05 | 0.00E+00 | 0.00E+00 | 0.00E+00 | 3.22E-04 | 0.00E+00 | 7.29E-05 | 0.00E+00 | Known_Known |
| g__Pseudonocardia | 0.00E+00 | 0.00E+00 | 0.00E+00 | 0.00E+00 | 4.09E-05 | 2.34E-05 | 0.00E+00 | 0.00E+00 | 0.00E+00 | 0.00E+00 | 0.00E+00 | Known_Known |
| g__Pseudenhygromyxa | 0.00E+00 | 0.00E+00 | 0.00E+00 | 0.00E+00 | 0.00E+00 | 0.00E+00 | 0.00E+00 | 0.00E+00 | 1.19E-04 | 1.46E-04 | 0.00E+00 | Known_Known |
| g__Pseudalkalibacillus_286710 | 0.00E+00 | 0.00E+00 | 0.00E+00 | 0.00E+00 | 1.36E-05 | 3.50E-05 | 0.00E+00 | 0.00E+00 | 0.00E+00 | 0.00E+00 | 0.00E+00 | Known_Known |
| g__Proteinivorax | 0.00E+00 | 0.00E+00 | 0.00E+00 | 0.00E+00 | 4.50E-04 | 0.00E+00 | 5.37E-05 | 1.16E-04 | 2.37E-04 | 5.28E-04 | 2.92E-04 | Known_Known |
| g__Prosthecochloris | 1.17E-04 | 0.00E+00 | 0.00E+00 | 3.43E-05 | 1.64E-04 | 0.00E+00 | 0.00E+00 | 0.00E+00 | 0.00E+00 | 0.00E+00 | 0.00E+00 | Known_Known |
| g__Propylenella | 0.00E+00 | 0.00E+00 | 0.00E+00 | 0.00E+00 | 1.77E-04 | 0.00E+00 | 0.00E+00 | 0.00E+00 | 0.00E+00 | 0.00E+00 | 0.00E+00 | Known_Known |
| g__Pontibacter_909846 | 0.00E+00 | 0.00E+00 | 2.35E-05 | 0.00E+00 | 0.00E+00 | 0.00E+00 | 0.00E+00 | 0.00E+00 | 0.00E+00 | 0.00E+00 | 0.00E+00 | Known_Known |
| g__Pontibacter_909804 | 0.00E+00 | 5.11E-05 | 0.00E+00 | 0.00E+00 | 0.00E+00 | 1.17E-05 | 0.00E+00 | 0.00E+00 | 0.00E+00 | 0.00E+00 | 0.00E+00 | Known_Known |
| g__Polymorphobacter_A_486811 | 0.00E+00 | 0.00E+00 | 0.00E+00 | 8.01E-05 | 0.00E+00 | 0.00E+00 | 0.00E+00 | 0.00E+00 | 0.00E+00 | 0.00E+00 | 0.00E+00 | Known_Known |
| g__Phenylobacterium | 0.00E+00 | 0.00E+00 | 7.04E-05 | 0.00E+00 | 0.00E+00 | 0.00E+00 | 0.00E+00 | 0.00E+00 | 0.00E+00 | 0.00E+00 | 0.00E+00 | Known_Known |
| g__PH28-bin88 | 4.67E-05 | 0.00E+00 | 0.00E+00 | 0.00E+00 | 0.00E+00 | 0.00E+00 | 0.00E+00 | 0.00E+00 | 1.04E-04 | 2.55E-04 | 9.72E-05 | Known_Known |
| g__Petroclostridium | 1.17E-04 | 0.00E+00 | 0.00E+00 | 1.14E-05 | 5.46E-05 | 2.80E-04 | 4.66E-04 | 2.32E-04 | 2.52E-04 | 3.10E-04 | 1.78E-04 | Known_Known |
| g__Persicimonas | 0.00E+00 | 3.75E-04 | 8.22E-05 | 0.00E+00 | 0.00E+00 | 0.00E+00 | 0.00E+00 | 0.00E+00 | 0.00E+00 | 5.46E-05 | 0.00E+00 | Known_Known |
| g__Pelolinea | 0.00E+00 | 4.51E-03 | 0.00E+00 | 0.00E+00 | 1.23E-04 | 0.00E+00 | 1.97E-04 | 5.67E-04 | 1.44E-03 | 1.04E-03 | 8.75E-04 | Known_Known |
| g__Pelagibius | 9.33E-05 | 2.04E-04 | 0.00E+00 | 0.00E+00 | 0.00E+00 | 8.17E-05 | 0.00E+00 | 0.00E+00 | 1.19E-04 | 0.00E+00 | 0.00E+00 | Known_Known |
| g__Pelagibacterium | 0.00E+00 | 0.00E+00 | 0.00E+00 | 0.00E+00 | 0.00E+00 | 3.50E-05 | 0.00E+00 | 0.00E+00 | 0.00E+00 | 0.00E+00 | 0.00E+00 | Known_Known |
| g__PCC-7407 | 0.00E+00 | 0.00E+00 | 0.00E+00 | 0.00E+00 | 0.00E+00 | 1.17E-05 | 0.00E+00 | 0.00E+00 | 0.00E+00 | 0.00E+00 | 0.00E+00 | Known_Known |
| g__Pasteuria | 0.00E+00 | 0.00E+00 | 0.00E+00 | 0.00E+00 | 0.00E+00 | 0.00E+00 | 0.00E+00 | 0.00E+00 | 0.00E+00 | 4.19E-04 | 6.48E-05 | Known_Known |
| g__Parvularcula | 0.00E+00 | 0.00E+00 | 0.00E+00 | 1.37E-04 | 0.00E+00 | 9.34E-05 | 0.00E+00 | 2.70E-04 | 0.00E+00 | 0.00E+00 | 0.00E+00 | Known_Known |
| g__Paracoccus | 0.00E+00 | 3.92E-04 | 1.36E-03 | 9.39E-04 | 0.00E+00 | 1.52E-04 | 0.00E+00 | 0.00E+00 | 0.00E+00 | 0.00E+00 | 0.00E+00 | Known_Known |
| g__Paraclostridium | 3.97E-04 | 1.36E-04 | 0.00E+00 | 4.58E-05 | 0.00E+00 | 0.00E+00 | 0.00E+00 | 0.00E+00 | 1.34E-04 | 0.00E+00 | 3.73E-04 | Known_Known |
| g__Papillibacter | 0.00E+00 | 0.00E+00 | 0.00E+00 | 0.00E+00 | 8.19E-05 | 0.00E+00 | 0.00E+00 | 0.00E+00 | 0.00E+00 | 0.00E+00 | 0.00E+00 | Known_Known |
| g__Paludibaculum | 0.00E+00 | 1.19E-04 | 3.52E-05 | 1.27E-03 | 4.09E-05 | 0.00E+00 | 0.00E+00 | 3.86E-05 | 5.93E-05 | 0.00E+00 | 0.00E+00 | Known_Known |
| g__PALSA-986 | 3.97E-04 | 4.09E-04 | 1.53E-04 | 1.26E-04 | 0.00E+00 | 0.00E+00 | 8.60E-04 | 2.06E-04 | 2.85E-03 | 1.53E-03 | 1.57E-03 | Known_Known |
| g__PALSA-647 | 0.00E+00 | 0.00E+00 | 0.00E+00 | 0.00E+00 | 0.00E+00 | 0.00E+00 | 0.00E+00 | 0.00E+00 | 0.00E+00 | 3.64E-05 | 0.00E+00 | Known_Known |
| g__Palleronia_492160 | 0.00E+00 | 2.04E-04 | 0.00E+00 | 0.00E+00 | 0.00E+00 | 0.00E+00 | 0.00E+00 | 0.00E+00 | 0.00E+00 | 0.00E+00 | 0.00E+00 | Known_Known |
| g__Palleronia_492156 | 0.00E+00 | 6.98E-04 | 4.23E-04 | 0.00E+00 | 0.00E+00 | 2.34E-04 | 1.07E-04 | 1.29E-04 | 0.00E+00 | 0.00E+00 | 8.10E-05 | Known_Known |
| g__Paenibacillus_Z_364416 | 0.00E+00 | 0.00E+00 | 0.00E+00 | 0.00E+00 | 4.09E-05 | 2.34E-05 | 0.00E+00 | 0.00E+00 | 0.00E+00 | 0.00E+00 | 0.00E+00 | Known_Known |
| g__Paenibacillus_J_366884 | 0.00E+00 | 0.00E+00 | 0.00E+00 | 0.00E+00 | 2.73E-05 | 1.17E-05 | 0.00E+00 | 0.00E+00 | 0.00E+00 | 3.64E-05 | 0.00E+00 | Known_Known |
| g__Paenibacillus_D_363770 | 0.00E+00 | 0.00E+00 | 1.17E-05 | 0.00E+00 | 0.00E+00 | 0.00E+00 | 0.00E+00 | 0.00E+00 | 0.00E+00 | 0.00E+00 | 0.00E+00 | Known_Known |
| g__Paenibacillus_C | 0.00E+00 | 0.00E+00 | 0.00E+00 | 0.00E+00 | 0.00E+00 | 1.17E-05 | 0.00E+00 | 0.00E+00 | 0.00E+00 | 0.00E+00 | 0.00E+00 | Known_Known |
| g__Paenibacillus_AE | 2.33E-05 | 0.00E+00 | 7.04E-05 | 0.00E+00 | 8.19E-05 | 2.34E-05 | 0.00E+00 | 0.00E+00 | 0.00E+00 | 0.00E+00 | 0.00E+00 | Known_Known |
| g__OXYB2-FULL-41-6 | 0.00E+00 | 0.00E+00 | 0.00E+00 | 0.00E+00 | 0.00E+00 | 0.00E+00 | 0.00E+00 | 3.86E-05 | 1.04E-04 | 1.82E-04 | 0.00E+00 | Known_Known |
| g__Oxalophagus | 0.00E+00 | 0.00E+00 | 3.52E-05 | 0.00E+00 | 0.00E+00 | 4.67E-05 | 0.00E+00 | 0.00E+00 | 0.00E+00 | 2.19E-04 | 0.00E+00 | Known_Known |
| g__Oricola | 0.00E+00 | 1.02E-04 | 0.00E+00 | 0.00E+00 | 0.00E+00 | 0.00E+00 | 0.00E+00 | 3.73E-04 | 0.00E+00 | 2.00E-04 | 0.00E+00 | Known_Known |
| g__Orenia_42969 | 0.00E+00 | 0.00E+00 | 0.00E+00 | 0.00E+00 | 0.00E+00 | 0.00E+00 | 0.00E+00 | 0.00E+00 | 2.97E-05 | 0.00E+00 | 0.00E+00 | Known_Known |
| g__Oharaeibacter | 0.00E+00 | 0.00E+00 | 0.00E+00 | 1.95E-04 | 0.00E+00 | 0.00E+00 | 0.00E+00 | 0.00E+00 | 0.00E+00 | 2.00E-04 | 0.00E+00 | Known_Known |
| g__Oceanidesulfovibrio_446513 | 0.00E+00 | 0.00E+00 | 0.00E+00 | 0.00E+00 | 0.00E+00 | 0.00E+00 | 0.00E+00 | 0.00E+00 | 0.00E+00 | 0.00E+00 | 3.24E-05 | Known_Known |
| g__O2-02-FULL-42-17 | 7.00E-05 | 0.00E+00 | 0.00E+00 | 0.00E+00 | 0.00E+00 | 0.00E+00 | 0.00E+00 | 0.00E+00 | 0.00E+00 | 0.00E+00 | 0.00E+00 | Known_Known |
| g__Nodosilinea | 3.87E-03 | 8.52E-05 | 0.00E+00 | 0.00E+00 | 0.00E+00 | 0.00E+00 | 0.00E+00 | 2.58E-05 | 0.00E+00 | 0.00E+00 | 3.08E-04 | Known_Known |

| Genus | L1 | L2 | L3 | L4 | L5 | L6 | L7 | L8 | L9 | L10 | L11 | Microbial_matter |
|---|---|---|---|---|---|---|---|---|---|---|---|---|
| g__Nitrososphaera | 7.00E-05 | 0.00E+00 | 0.00E+00 | 0.00E+00 | 0.00E+00 | 3.97E-04 | 0.00E+00 | 0.00E+00 | 0.00E+00 | 0.00E+00 | 0.00E+00 | Known_Known |
| g__Nitrosocosmicus | 0.00E+00 | 1.36E-04 | 0.00E+00 | 0.00E+00 | 3.82E-04 | 2.60E-03 | 2.87E-04 | 0.00E+00 | 2.52E-04 | 0.00E+00 | 2.92E-04 | Known_Known |
| g__Nitrolancea | 1.87E-04 | 7.15E-04 | 3.64E-04 | 8.01E-05 | 4.09E-04 | 4.67E-05 | 0.00E+00 | 0.00E+00 | 3.41E-04 | 1.82E-04 | 1.46E-04 | Known_Known |
| g__Niameybacter | 0.00E+00 | 0.00E+00 | 0.00E+00 | 0.00E+00 | 0.00E+00 | 0.00E+00 | 0.00E+00 | 0.00E+00 | 0.00E+00 | 0.00E+00 | 1.30E-04 | Known_Known |
| g__Nesciobacter | 0.00E+00 | 0.00E+00 | 0.00E+00 | 2.29E-04 | 0.00E+00 | 0.00E+00 | 0.00E+00 | 0.00E+00 | 0.00E+00 | 0.00E+00 | 0.00E+00 | Known_Known |
| g__Natronorubrum_4312 | 0.00E+00 | 5.11E-05 | 0.00E+00 | 0.00E+00 | 2.05E-04 | 1.17E-04 | 0.00E+00 | 0.00E+00 | 0.00E+00 | 0.00E+00 | 0.00E+00 | Known_Known |
| g__Natronoplasma | 4.39E-03 | 9.71E-04 | 5.75E-04 | 0.00E+00 | 0.00E+00 | 1.28E-04 | 1.07E-03 | 7.98E-04 | 4.30E-04 | 0.00E+00 | 1.69E-03 | Known_Known |
| g__Natronomonas | 7.00E-05 | 4.16E-03 | 2.21E-03 | 1.14E-04 | 2.32E-04 | 0.00E+00 | 1.79E-04 | 0.00E+00 | 8.90E-05 | 2.55E-04 | 2.59E-04 | Known_Known |
| g__Natronobiforma | 0.00E+00 | 1.64E-03 | 2.89E-03 | 1.27E-03 | 8.19E-05 | 0.00E+00 | 0.00E+00 | 0.00E+00 | 8.90E-05 | 0.00E+00 | 2.11E-04 | Known_Known |
| g__Natronoarchaeum | 1.59E-03 | 7.87E-03 | 5.04E-03 | 9.50E-04 | 1.91E-04 | 2.10E-04 | 1.09E-03 | 7.73E-05 | 5.64E-04 | 1.06E-03 | 1.02E-03 | Known_Known |
| g__Natronincola_A_235765 | 8.31E-03 | 5.45E-04 | 4.23E-04 | 3.89E-04 | 2.57E-03 | 2.12E-02 | 3.98E-03 | 3.22E-03 | 1.41E-03 | 5.83E-04 | 4.04E-03 | Known_Known |
| g__Natronincola_A_235762 | 0.00E+00 | 2.21E-04 | 2.82E-04 | 0.00E+00 | 1.53E-03 | 2.69E-04 | 0.00E+00 | 0.00E+00 | 0.00E+00 | 0.00E+00 | 0.00E+00 | Known_Known |
| g__Natrinema | 2.38E-03 | 1.80E-02 | 1.10E-02 | 4.46E-04 | 9.55E-05 | 1.40E-04 | 6.63E-03 | 0.00E+00 | 0.00E+00 | 2.55E-04 | 1.06E-02 | Known_Known |
| g__Natrarchaeobaculum | 1.87E-03 | 1.28E-02 | 7.02E-03 | 1.64E-03 | 6.41E-04 | 7.94E-04 | 1.31E-03 | 3.86E-04 | 1.94E-03 | 1.53E-03 | 2.50E-03 | Known_Known |
| g__Natranaerovirga | 1.12E-03 | 1.72E-03 | 4.70E-04 | 2.07E-03 | 8.19E-04 | 3.26E-03 | 4.57E-03 | 2.85E-03 | 6.83E-03 | 4.99E-02 | 1.05E-02 | Known_Known |
| g__Natranaerobaculum | 0.00E+00 | 0.00E+00 | 7.04E-05 | 0.00E+00 | 0.00E+00 | 0.00E+00 | 0.00E+00 | 0.00E+00 | 0.00E+00 | 0.00E+00 | 0.00E+00 | Known_Known |
| g__Nanopelagicus | 0.00E+00 | 2.21E-04 | 4.70E-05 | 3.43E-05 | 1.23E-04 | 0.00E+00 | 3.58E-05 | 6.44E-04 | 1.97E-03 | 3.19E-03 | 7.13E-03 | Known_Known |
| g__Nanohalobium | 1.40E-04 | 6.13E-04 | 2.11E-04 | 2.52E-04 | 4.09E-05 | 1.87E-04 | 3.58E-04 | 2.58E-05 | 2.97E-05 | 3.64E-05 | 1.94E-04 | Known_Known |
| g__NAK82 | 4.67E-05 | 8.52E-05 | 0.00E+00 | 2.52E-04 | 0.00E+00 | 0.00E+00 | 0.00E+00 | 3.86E-05 | 0.00E+00 | 0.00E+00 | 0.00E+00 | Known_Known |
| g__MZ-XQ | 0.00E+00 | 0.00E+00 | 0.00E+00 | 0.00E+00 | 0.00E+00 | 0.00E+00 | 0.00E+00 | 0.00E+00 | 0.00E+00 | 1.82E-05 | 0.00E+00 | Known_Known |
| g__Mycobacterium | 0.00E+00 | 0.00E+00 | 0.00E+00 | 4.58E-05 | 9.55E-05 | 0.00E+00 | 0.00E+00 | 0.00E+00 | 0.00E+00 | 5.10E-04 | 4.86E-05 | Known_Known |
| g__MWDR01 | 0.00E+00 | 0.00E+00 | 0.00E+00 | 0.00E+00 | 0.00E+00 | 0.00E+00 | 0.00E+00 | 0.00E+00 | 2.97E-05 | 9.11E-05 | 4.86E-05 | Known_Known |
| g__MWBO01 | 0.00E+00 | 5.11E-05 | 0.00E+00 | 0.00E+00 | 0.00E+00 | 0.00E+00 | 0.00E+00 | 0.00E+00 | 0.00E+00 | 0.00E+00 | 0.00E+00 | Known_Known |
| g__MVZH01 | 0.00E+00 | 0.00E+00 | 2.35E-05 | 0.00E+00 | 0.00E+00 | 0.00E+00 | 5.37E-05 | 0.00E+00 | 0.00E+00 | 0.00E+00 | 0.00E+00 | Known_Known |
| g__Moorella | 7.00E-05 | 3.92E-04 | 4.34E-04 | 4.58E-05 | 9.69E-04 | 4.20E-04 | 1.61E-04 | 0.00E+00 | 1.04E-04 | 0.00E+00 | 6.48E-05 | Known_Known |
| g__Modestobacter | 0.00E+00 | 0.00E+00 | 0.00E+00 | 0.00E+00 | 4.09E-05 | 7.01E-05 | 0.00E+00 | 0.00E+00 | 2.97E-05 | 7.29E-05 | 1.62E-05 | Known_Known |
| g__Mobilitalea | 0.00E+00 | 0.00E+00 | 0.00E+00 | 1.05E-03 | 1.62E-03 | 8.87E-04 | 0.00E+00 | 1.16E-04 | 5.93E-04 | 0.00E+00 | 0.00E+00 | Known_Known |
| g__MnB-11 | 0.00E+00 | 6.81E-05 | 0.00E+00 | 0.00E+00 | 0.00E+00 | 0.00E+00 | 0.00E+00 | 0.00E+00 | 0.00E+00 | 0.00E+00 | 0.00E+00 | Known_Known |
| g__MLS-D | 0.00E+00 | 0.00E+00 | 0.00E+00 | 0.00E+00 | 6.82E-05 | 0.00E+00 | 0.00E+00 | 0.00E+00 | 0.00E+00 | 0.00E+00 | 0.00E+00 | Known_Known |
| g__Microvirga | 2.57E-04 | 8.65E-03 | 1.87E-03 | 0.00E+00 | 1.43E-03 | 2.78E-03 | 5.19E-04 | 6.44E-05 | 8.90E-05 | 9.11E-05 | 4.86E-04 | Known_Known |
| g__Micavibrio_B_508655 | 0.00E+00 | 0.00E+00 | 0.00E+00 | 6.87E-05 | 0.00E+00 | 0.00E+00 | 0.00E+00 | 0.00E+00 | 0.00E+00 | 3.64E-05 | 0.00E+00 | Known_Known |
| g__MGML01 | 1.63E-04 | 1.36E-04 | 0.00E+00 | 1.03E-04 | 4.09E-05 | 3.50E-05 | 1.07E-04 | 0.00E+00 | 2.97E-05 | 5.46E-05 | 9.72E-05 | Known_Known |
| g__Methyloceanibacter | 2.33E-04 | 1.70E-04 | 0.00E+00 | 4.01E-04 | 0.00E+00 | 1.52E-04 | 5.37E-05 | 6.31E-04 | 8.16E-04 | 1.38E-03 | 9.24E-04 | Known_Known |
| g__Methylobacterium | 0.00E+00 | 9.20E-04 | 0.00E+00 | 0.00E+00 | 8.05E-04 | 1.86E-03 | 1.97E-04 | 0.00E+00 | 0.00E+00 | 0.00E+00 | 3.89E-04 | Known_Known |
| g__Methanothrix_A | 0.00E+00 | 0.00E+00 | 0.00E+00 | 0.00E+00 | 0.00E+00 | 0.00E+00 | 0.00E+00 | 0.00E+00 | 8.46E-04 | 5.10E-04 | 2.59E-04 | Known_Known |
| g__Methanospirillum | 0.00E+00 | 0.00E+00 | 0.00E+00 | 0.00E+00 | 0.00E+00 | 0.00E+00 | 0.00E+00 | 0.00E+00 | 2.97E-05 | 0.00E+00 | 0.00E+00 | Known_Known |
| g__Methanolobus | 0.00E+00 | 6.30E-04 | 0.00E+00 | 0.00E+00 | 0.00E+00 | 0.00E+00 | 0.00E+00 | 0.00E+00 | 6.83E-04 | 3.28E-04 | 4.70E-04 | Known_Known |
| g__Methanohalophilus | 3.10E-03 | 2.23E-03 | 1.29E-04 | 6.64E-04 | 0.00E+00 | 1.87E-04 | 1.06E-03 | 8.37E-04 | 5.19E-04 | 2.73E-04 | 1.49E-03 | Known_Known |
| g__Methanohalobium | 4.67E-05 | 2.90E-04 | 7.63E-04 | 1.03E-04 | 0.00E+00 | 2.19E-03 | 1.00E-02 | 1.93E-04 | 0.00E+00 | 0.00E+00 | 5.02E-04 | Known_Known |
| g__Methanobrevibacter_A | 0.00E+00 | 0.00E+00 | 0.00E+00 | 0.00E+00 | 0.00E+00 | 3.50E-05 | 0.00E+00 | 0.00E+00 | 0.00E+00 | 3.64E-05 | 0.00E+00 | Known_Known |
| g__Methanobacterium_D_1054 | 0.00E+00 | 3.41E-05 | 0.00E+00 | 3.43E-05 | 0.00E+00 | 0.00E+00 | 0.00E+00 | 0.00E+00 | 5.93E-05 | 7.29E-05 | 6.48E-05 | Known_Known |
| g__Mesotoga | 1.17E-04 | 0.00E+00 | 0.00E+00 | 0.00E+00 | 2.73E-05 | 0.00E+00 | 0.00E+00 | 0.00E+00 | 0.00E+00 | 0.00E+00 | 1.46E-04 | Known_Known |
| g__MB-PLM-2 | 0.00E+00 | 0.00E+00 | 0.00E+00 | 0.00E+00 | 2.73E-05 | 0.00E+00 | 0.00E+00 | 3.86E-05 | 0.00E+00 | 0.00E+00 | 0.00E+00 | Known_Known |
| g__Marivirga | 7.00E-05 | 3.41E-05 | 1.17E-05 | 0.00E+00 | 0.00E+00 | 0.00E+00 | 0.00E+00 | 0.00E+00 | 0.00E+00 | 0.00E+00 | 4.86E-05 | Known_Known |
| g__Marispirochaeta | 0.00E+00 | 0.00E+00 | 0.00E+00 | 0.00E+00 | 2.73E-05 | 0.00E+00 | 0.00E+00 | 0.00E+00 | 0.00E+00 | 0.00E+00 | 0.00E+00 | Known_Known |
| g__Marinococcus | 0.00E+00 | 0.00E+00 | 2.35E-05 | 2.29E-05 | 0.00E+00 | 0.00E+00 | 0.00E+00 | 0.00E+00 | 0.00E+00 | 0.00E+00 | 1.62E-05 | Known_Known |
| g__Marinobacter_A_637054 | 9.33E-05 | 6.81E-05 | 8.22E-05 | 0.00E+00 | 0.00E+00 | 0.00E+00 | 0.00E+00 | 0.00E+00 | 0.00E+00 | 1.27E-04 | 4.86E-05 | Known_Known |

| Genus | L1 | L2 | L3 | L4 | L5 | L6 | L7 | L8 | L9 | L10 | L11 | Microbial_matter |
|---|---|---|---|---|---|---|---|---|---|---|---|---|
| g__Mariniphaga | 0.00E+00 | 0.00E+00 | 2.35E-05 | 0.00E+00 | 0.00E+00 | 0.00E+00 | 0.00E+00 | 0.00E+00 | 0.00E+00 | 2.19E-04 | 0.00E+00 | Known_Known |
| g__Marinilabilia | 0.00E+00 | 0.00E+00 | 0.00E+00 | 0.00E+00 | 0.00E+00 | 0.00E+00 | 0.00E+00 | 0.00E+00 | 0.00E+00 | 5.46E-05 | 0.00E+00 | Known_Known |
| g__Marinicaulis_488176 | 0.00E+00 | 0.00E+00 | 0.00E+00 | 0.00E+00 | 0.00E+00 | 0.00E+00 | 0.00E+00 | 2.19E-04 | 1.19E-04 | 0.00E+00 | 0.00E+00 | Known_Known |
| g__Marinicauda | 0.00E+00 | 3.07E-04 | 9.39E-05 | 1.03E-04 | 0.00E+00 | 0.00E+00 | 0.00E+00 | 0.00E+00 | 0.00E+00 | 2.37E-04 | 0.00E+00 | Known_Known |
| g__Maricaulis | 0.00E+00 | 0.00E+00 | 0.00E+00 | 1.03E-04 | 0.00E+00 | 0.00E+00 | 0.00E+00 | 0.00E+00 | 0.00E+00 | 0.00E+00 | 0.00E+00 | Known_Known |
| g__LZORAL124-64-63 | 0.00E+00 | 0.00E+00 | 0.00E+00 | 0.00E+00 | 4.09E-05 | 0.00E+00 | 0.00E+00 | 0.00E+00 | 0.00E+00 | 0.00E+00 | 0.00E+00 | Known_Known |
| g__Lutispora | 0.00E+00 | 1.35E-03 | 1.01E-03 | 5.04E-04 | 3.41E-04 | 0.00E+00 | 0.00E+00 | 0.00E+00 | 1.63E-04 | 0.00E+00 | 1.13E-04 | Known_Known |
| g__Lutibaculum | 0.00E+00 | 2.04E-04 | 0.00E+00 | 0.00E+00 | 0.00E+00 | 0.00E+00 | 0.00E+00 | 0.00E+00 | 0.00E+00 | 0.00E+00 | 0.00E+00 | Known_Known |
| g__Lujinxingia | 0.00E+00 | 0.00E+00 | 0.00E+00 | 0.00E+00 | 2.73E-05 | 4.67E-05 | 0.00E+00 | 0.00E+00 | 0.00E+00 | 4.37E-04 | 0.00E+00 | Known_Known |
| g__Litorilinea | 5.83E-04 | 3.75E-04 | 0.00E+00 | 1.37E-04 | 3.68E-04 | 2.34E-05 | 1.79E-04 | 1.55E-04 | 1.84E-03 | 4.55E-04 | 4.86E-04 | Known_Known |
| g__Limnospira | 0.00E+00 | 0.00E+00 | 0.00E+00 | 1.14E-05 | 0.00E+00 | 0.00E+00 | 0.00E+00 | 0.00E+00 | 0.00E+00 | 0.00E+00 | 0.00E+00 | Known_Known |
| g__Limnochorda | 0.00E+00 | 0.00E+00 | 2.35E-04 | 6.87E-05 | 1.09E-04 | 1.05E-04 | 0.00E+00 | 0.00E+00 | 1.93E-04 | 4.92E-04 | 3.40E-04 | Known_Known |
| g__Limivicinus | 0.00E+00 | 0.00E+00 | 0.00E+00 | 0.00E+00 | 0.00E+00 | 4.67E-05 | 0.00E+00 | 0.00E+00 | 0.00E+00 | 0.00E+00 | 0.00E+00 | Known_Known |
| g__Limiplasma | 0.00E+00 | 0.00E+00 | 9.39E-05 | 0.00E+00 | 2.87E-04 | 0.00E+00 | 0.00E+00 | 0.00E+00 | 0.00E+00 | 0.00E+00 | 0.00E+00 | Known_Known |
| g__Limibaculum | 0.00E+00 | 0.00E+00 | 0.00E+00 | 8.01E-05 | 0.00E+00 | 1.52E-04 | 0.00E+00 | 0.00E+00 | 0.00E+00 | 0.00E+00 | 0.00E+00 | Known_Known |
| g__Lenti-02 | 0.00E+00 | 0.00E+00 | 0.00E+00 | 0.00E+00 | 0.00E+00 | 0.00E+00 | 0.00E+00 | 0.00E+00 | 0.00E+00 | 0.00E+00 | 4.86E-05 | Known_Known |
| g__Legionella_B_628077 | 0.00E+00 | 0.00E+00 | 2.35E-05 | 0.00E+00 | 0.00E+00 | 0.00E+00 | 0.00E+00 | 0.00E+00 | 0.00E+00 | 0.00E+00 | 0.00E+00 | Known_Known |
| g__Latescibacter | 0.00E+00 | 0.00E+00 | 0.00E+00 | 0.00E+00 | 5.46E-05 | 0.00E+00 | 0.00E+00 | 0.00E+00 | 4.45E-05 | 0.00E+00 | 0.00E+00 | Known_Known |
| g__Lacrimispora | 2.10E-04 | 0.00E+00 | 1.29E-04 | 6.87E-05 | 5.59E-04 | 1.63E-04 | 3.04E-04 | 1.03E-04 | 0.00E+00 | 1.51E-03 | 5.19E-04 | Known_Known |
| g__Lachnoclostridium_A_184689 | 0.00E+00 | 0.00E+00 | 0.00E+00 | 3.43E-05 | 6.00E-04 | 1.23E-03 | 0.00E+00 | 0.00E+00 | 5.49E-04 | 0.00E+00 | 5.19E-04 | Known_Known |
| g__Kocuria | 0.00E+00 | 0.00E+00 | 0.00E+00 | 0.00E+00 | 4.09E-05 | 0.00E+00 | 0.00E+00 | 0.00E+00 | 0.00E+00 | 0.00E+00 | 0.00E+00 | Known_Known |
| g__Klenkia | 0.00E+00 | 3.41E-05 | 0.00E+00 | 0.00E+00 | 0.00E+00 | 0.00E+00 | 0.00E+00 | 0.00E+00 | 0.00E+00 | 0.00E+00 | 0.00E+00 | Known_Known |
| g__JC017 | 7.00E-05 | 0.00E+00 | 4.70E-05 | 0.00E+00 | 0.00E+00 | 0.00E+00 | 0.00E+00 | 0.00E+00 | 2.52E-04 | 3.64E-05 | 2.11E-04 | Known_Known |
| g__JACIWP01 | 0.00E+00 | 1.02E-04 | 0.00E+00 | 0.00E+00 | 0.00E+00 | 0.00E+00 | 0.00E+00 | 0.00E+00 | 0.00E+00 | 1.82E-04 | 0.00E+00 | Known_Known |
| g__JACAEJ01 | 0.00E+00 | 0.00E+00 | 0.00E+00 | 0.00E+00 | 0.00E+00 | 0.00E+00 | 0.00E+00 | 0.00E+00 | 8.90E-05 | 3.64E-05 | 0.00E+00 | Known_Known |
| g__JABFSM01 | 0.00E+00 | 0.00E+00 | 0.00E+00 | 0.00E+00 | 0.00E+00 | 8.17E-05 | 0.00E+00 | 0.00E+00 | 0.00E+00 | 0.00E+00 | 0.00E+00 | Known_Known |
| g__JAAZNL01 | 0.00E+00 | 0.00E+00 | 0.00E+00 | 8.01E-05 | 2.73E-05 | 0.00E+00 | 0.00E+00 | 0.00E+00 | 0.00E+00 | 0.00E+00 | 0.00E+00 | Known_Known |
| g__JAAYXY01 | 1.17E-04 | 0.00E+00 | 0.00E+00 | 0.00E+00 | 0.00E+00 | 0.00E+00 | 0.00E+00 | 0.00E+00 | 0.00E+00 | 0.00E+00 | 0.00E+00 | Known_Known |
| g__JAAYUB01 | 0.00E+00 | 0.00E+00 | 0.00E+00 | 0.00E+00 | 0.00E+00 | 0.00E+00 | 0.00E+00 | 0.00E+00 | 5.93E-05 | 0.00E+00 | 0.00E+00 | Known_Known |
| g__JAAYNZ01 | 0.00E+00 | 0.00E+00 | 0.00E+00 | 0.00E+00 | 0.00E+00 | 0.00E+00 | 0.00E+00 | 0.00E+00 | 1.04E-04 | 0.00E+00 | 0.00E+00 | Known_Known |
| g__JAAYDY01 | 0.00E+00 | 0.00E+00 | 0.00E+00 | 0.00E+00 | 0.00E+00 | 0.00E+00 | 0.00E+00 | 0.00E+00 | 2.97E-05 | 5.46E-05 | 0.00E+00 | Known_Known |
| g__JAAYAV01 | 0.00E+00 | 5.28E-04 | 0.00E+00 | 1.14E-04 | 4.37E-04 | 0.00E+00 | 0.00E+00 | 3.73E-04 | 2.77E-03 | 3.88E-03 | 1.30E-03 | Known_Known |
| g__JAAXQM01 | 2.33E-04 | 4.26E-04 | 5.87E-05 | 2.52E-04 | 1.64E-04 | 0.00E+00 | 0.00E+00 | 1.94E-03 | 3.56E-04 | 1.91E-03 | 4.05E-04 | Known_Known |
| g__JAAXHJ01 | 0.00E+00 | 0.00E+00 | 0.00E+00 | 0.00E+00 | 1.36E-05 | 0.00E+00 | 0.00E+00 | 0.00E+00 | 0.00E+00 | 9.11E-05 | 0.00E+00 | Known_Known |
| g__JAAUTF01 | 0.00E+00 | 0.00E+00 | 0.00E+00 | 0.00E+00 | 0.00E+00 | 0.00E+00 | 0.00E+00 | 5.15E-05 | 1.57E-03 | 1.46E-04 | 1.62E-04 | Known_Known |
| g__JAABVY01 | 3.50E-04 | 3.41E-05 | 0.00E+00 | 3.43E-05 | 0.00E+00 | 0.00E+00 | 7.16E-05 | 4.25E-04 | 0.00E+00 | 0.00E+00 | 9.72E-05 | Known_Known |
| g__JAABTL01 | 3.90E-03 | 1.07E-02 | 2.82E-04 | 9.23E-03 | 5.64E-03 | 1.28E-04 | 5.19E-04 | 6.99E-03 | 5.79E-03 | 1.69E-03 | 2.45E-03 | Known_Known |
| g__JAAAPF01 | 4.60E-03 | 3.61E-03 | 1.39E-03 | 2.40E-03 | 1.95E-03 | 2.95E-02 | 1.07E-03 | 7.73E-04 | 4.45E-04 | 4.55E-04 | 3.26E-03 | Known_Known |
| g__J25 | 0.00E+00 | 0.00E+00 | 0.00E+00 | 0.00E+00 | 0.00E+00 | 0.00E+00 | 0.00E+00 | 3.86E-05 | 0.00E+00 | 0.00E+00 | 0.00E+00 | Known_Known |
| g__J137 | 0.00E+00 | 0.00E+00 | 0.00E+00 | 4.58E-05 | 0.00E+00 | 0.00E+00 | 0.00E+00 | 0.00E+00 | 0.00E+00 | 0.00E+00 | 0.00E+00 | Known_Known |
| g__J124 | 4.67E-05 | 0.00E+00 | 0.00E+00 | 0.00E+00 | 0.00E+00 | 0.00E+00 | 0.00E+00 | 9.01E-05 | 1.63E-04 | 1.64E-04 | 1.13E-04 | Known_Known |
| g__J110 | 7.00E-05 | 0.00E+00 | 0.00E+00 | 0.00E+00 | 0.00E+00 | 0.00E+00 | 0.00E+00 | 0.00E+00 | 0.00E+00 | 0.00E+00 | 0.00E+00 | Known_Known |
| g__Isachenkonia | 4.67E-05 | 9.20E-04 | 1.03E-03 | 3.32E-04 | 0.00E+00 | 1.17E-04 | 1.61E-04 | 1.03E-04 | 7.42E-05 | 5.10E-04 | 1.13E-04 | Known_Known |
| g__Irregularibacter | 0.00E+00 | 0.00E+00 | 1.17E-04 | 0.00E+00 | 1.09E-04 | 7.01E-05 | 0.00E+00 | 1.93E-04 | 0.00E+00 | 0.00E+00 | 0.00E+00 | Known_Known |
| g__Inquilinus | 0.00E+00 | 0.00E+00 | 0.00E+00 | 0.00E+00 | 0.00E+00 | 0.00E+00 | 0.00E+00 | 0.00E+00 | 0.00E+00 | 2.19E-04 | 0.00E+00 | Known_Known |
| g__Indioceanicola | 0.00E+00 | 0.00E+00 | 1.29E-04 | 0.00E+00 | 0.00E+00 | 0.00E+00 | 0.00E+00 | 0.00E+00 | 1.19E-04 | 0.00E+00 | 0.00E+00 | Known_Known |
| g__IMCC26207 | 0.00E+00 | 1.53E-04 | 0.00E+00 | 1.14E-04 | 0.00E+00 | 0.00E+00 | 0.00E+00 | 1.16E-04 | 4.15E-04 | 0.00E+00 | 0.00E+00 | Known_Known |

| Genus | L1 | L2 | L3 | L4 | L5 | L6 | L7 | L8 | L9 | L10 | L11 | Microbial_matter |
|---|---|---|---|---|---|---|---|---|---|---|---|---|
| g__Ilumatobacter_A | 2.10E-03 | 6.47E-04 | 3.52E-05 | 3.66E-04 | 2.59E-04 | 0.00E+00 | 0.00E+00 | 5.15E-05 | 1.48E-04 | 0.00E+00 | 6.81E-04 | Known_Known |
| g__Idiomarina | 0.00E+00 | 0.00E+00 | 0.00E+00 | 3.43E-05 | 1.50E-04 | 0.00E+00 | 0.00E+00 | 0.00E+00 | 0.00E+00 | 0.00E+00 | 0.00E+00 | Known_Known |
| g__HyVt-337 | 0.00E+00 | 0.00E+00 | 0.00E+00 | 0.00E+00 | 0.00E+00 | 0.00E+00 | 0.00E+00 | 0.00E+00 | 0.00E+00 | 3.64E-05 | 0.00E+00 | Known_Known |
| g__HyVt-105 | 4.67E-05 | 0.00E+00 | 5.87E-05 | 4.58E-05 | 0.00E+00 | 0.00E+00 | 8.96E-05 | 0.00E+00 | 0.00E+00 | 3.64E-05 | 0.00E+00 | Known_Known |
| g__Hyphomonas | 8.63E-04 | 3.41E-04 | 7.51E-04 | 3.57E-03 | 6.82E-05 | 1.52E-04 | 2.33E-04 | 1.00E-03 | 7.42E-05 | 7.29E-05 | 3.40E-04 | Known_Known |
| g__Hyphomicrobium_A | 0.00E+00 | 3.92E-04 | 1.06E-04 | 3.32E-04 | 0.00E+00 | 0.00E+00 | 0.00E+00 | 1.56E-03 | 8.75E-04 | 8.20E-04 | 7.94E-04 | Known_Known |
| g__Hydrogenovibrio | 0.00E+00 | 0.00E+00 | 0.00E+00 | 1.14E-05 | 0.00E+00 | 0.00E+00 | 0.00E+00 | 0.00E+00 | 0.00E+00 | 4.55E-04 | 8.10E-05 | Known_Known |
| g__Hydrogenispora | 0.00E+00 | 1.87E-04 | 0.00E+00 | 0.00E+00 | 0.00E+00 | 0.00E+00 | 0.00E+00 | 0.00E+00 | 1.48E-04 | 0.00E+00 | 0.00E+00 | Known_Known |
| g__Hungateiclostridium | 0.00E+00 | 1.06E-03 | 3.99E-04 | 9.50E-04 | 3.45E-03 | 8.76E-04 | 4.84E-04 | 2.58E-04 | 1.82E-03 | 1.07E-03 | 5.19E-04 | Known_Known |
| g__HRBIN40 | 0.00E+00 | 6.81E-05 | 0.00E+00 | 4.58E-05 | 1.36E-04 | 0.00E+00 | 0.00E+00 | 0.00E+00 | 0.00E+00 | 0.00E+00 | 0.00E+00 | Known_Known |
| g__HRBIN30 | 0.00E+00 | 0.00E+00 | 0.00E+00 | 0.00E+00 | 0.00E+00 | 0.00E+00 | 0.00E+00 | 0.00E+00 | 4.45E-05 | 0.00E+00 | 0.00E+00 | Known_Known |
| g__HLUCCA09 | 6.58E-03 | 1.38E-03 | 1.80E-03 | 6.49E-03 | 2.87E-04 | 6.19E-04 | 5.03E-03 | 3.60E-02 | 1.75E-03 | 1.46E-04 | 1.77E-03 | Known_Known |
| g__Herbivorax | 0.00E+00 | 0.00E+00 | 0.00E+00 | 0.00E+00 | 1.36E-04 | 0.00E+00 | 0.00E+00 | 0.00E+00 | 0.00E+00 | 0.00E+00 | 0.00E+00 | Known_Known |
| g__Henriciella | 0.00E+00 | 0.00E+00 | 0.00E+00 | 0.00E+00 | 0.00E+00 | 2.34E-05 | 0.00E+00 | 3.86E-05 | 7.42E-05 | 0.00E+00 | 0.00E+00 | Known_Known |
| g__Heliorestis | 2.32E-01 | 5.00E-02 | 3.89E-02 | 5.64E-03 | 3.76E-01 | 3.85E-01 | 9.48E-02 | 4.58E-02 | 4.30E-02 | 9.24E-02 | 1.44E-01 | Known_Known |
| g__Heliomicrobium | 0.00E+00 | 0.00E+00 | 0.00E+00 | 0.00E+00 | 6.82E-05 | 0.00E+00 | 0.00E+00 | 0.00E+00 | 0.00E+00 | 0.00E+00 | 0.00E+00 | Known_Known |
| g__Halovivax | 7.23E-04 | 4.29E-03 | 2.27E-03 | 8.47E-04 | 0.00E+00 | 3.85E-04 | 1.56E-03 | 0.00E+00 | 1.93E-04 | 1.15E-03 | 5.67E-04 | Known_Known |
| g__Halovenus | 4.67E-04 | 6.52E-03 | 9.20E-03 | 1.79E-03 | 3.82E-04 | 1.00E-03 | 3.31E-03 | 1.67E-04 | 3.86E-04 | 5.83E-04 | 1.70E-03 | Known_Known |
| g__Halothece | 0.00E+00 | 0.00E+00 | 0.00E+00 | 0.00E+00 | 0.00E+00 | 0.00E+00 | 3.58E-05 | 0.00E+00 | 0.00E+00 | 0.00E+00 | 1.46E-04 | Known_Known |
| g__Halostella | 0.00E+00 | 1.02E-04 | 1.64E-04 | 0.00E+00 | 0.00E+00 | 0.00E+00 | 0.00E+00 | 0.00E+00 | 0.00E+00 | 0.00E+00 | 4.86E-05 | Known_Known |
| g__Halostagnicola | 6.35E-03 | 5.06E-03 | 4.66E-03 | 7.90E-04 | 2.29E-03 | 1.37E-03 | 2.61E-03 | 2.45E-04 | 1.34E-04 | 8.38E-04 | 2.22E-03 | Known_Known |
| g__Halosimplex | 0.00E+00 | 1.02E-03 | 9.04E-04 | 0.00E+00 | 0.00E+00 | 0.00E+00 | 0.00E+00 | 0.00E+00 | 0.00E+00 | 0.00E+00 | 1.62E-04 | Known_Known |
| g__Halorussus | 7.28E-03 | 1.90E-02 | 1.81E-02 | 2.44E-03 | 1.58E-03 | 6.49E-03 | 2.45E-02 | 2.56E-03 | 2.08E-03 | 1.26E-03 | 5.09E-03 | Known_Known |
| g__Halorubrum | 4.60E-03 | 7.75E-03 | 7.97E-03 | 1.03E-03 | 1.17E-03 | 2.48E-03 | 9.31E-04 | 7.60E-04 | 1.17E-03 | 2.53E-03 | 8.12E-03 | Known_Known |
| g__Halorientalis | 4.67E-04 | 4.99E-03 | 3.24E-03 | 5.27E-04 | 3.82E-04 | 6.89E-04 | 2.81E-03 | 1.03E-04 | 4.45E-05 | 2.00E-04 | 6.32E-04 | Known_Known |
| g__Halorhabdus | 0.00E+00 | 0.00E+00 | 2.00E-04 | 0.00E+00 | 0.00E+00 | 1.75E-04 | 1.61E-04 | 0.00E+00 | 0.00E+00 | 0.00E+00 | 0.00E+00 | Known_Known |
| g__Haloprofundus | 7.93E-04 | 9.28E-03 | 2.32E-03 | 1.72E-04 | 1.36E-04 | 8.17E-05 | 0.00E+00 | 7.73E-05 | 2.37E-04 | 7.29E-04 | 1.46E-03 | Known_Known |
| g__Haloplanus | 0.00E+00 | 0.00E+00 | 0.00E+00 | 0.00E+00 | 0.00E+00 | 0.00E+00 | 0.00E+00 | 0.00E+00 | 0.00E+00 | 0.00E+00 | 8.10E-05 | Known_Known |
| g__Halopiger_A | 8.87E-04 | 1.45E-03 | 1.95E-03 | 0.00E+00 | 1.02E-03 | 7.36E-04 | 1.27E-03 | 0.00E+00 | 2.97E-04 | 0.00E+00 | 0.00E+00 | Known_Known |
| g__Halopelagius | 0.00E+00 | 5.96E-04 | 1.41E-04 | 8.01E-05 | 0.00E+00 | 0.00E+00 | 0.00E+00 | 0.00E+00 | 0.00E+00 | 0.00E+00 | 4.86E-05 | Known_Known |
| g__Halonotius | 0.00E+00 | 0.00E+00 | 2.23E-04 | 0.00E+00 | 0.00E+00 | 0.00E+00 | 0.00E+00 | 0.00E+00 | 0.00E+00 | 0.00E+00 | 9.72E-05 | Known_Known |
| g__Halomonas_C_640989 | 0.00E+00 | 8.52E-05 | 1.17E-04 | 9.16E-05 | 0.00E+00 | 0.00E+00 | 0.00E+00 | 0.00E+00 | 0.00E+00 | 0.00E+00 | 0.00E+00 | Known_Known |
| g__Halomonas_B | 0.00E+00 | 0.00E+00 | 0.00E+00 | 0.00E+00 | 0.00E+00 | 0.00E+00 | 0.00E+00 | 0.00E+00 | 2.97E-05 | 1.27E-04 | 0.00E+00 | Known_Known |
| g__Halomicrobium_3849 | 5.37E-04 | 3.73E-03 | 3.18E-03 | 2.29E-04 | 1.21E-03 | 7.01E-04 | 1.97E-04 | 0.00E+00 | 5.93E-04 | 0.00E+00 | 6.00E-04 | Known_Known |
| g__Halomarina_3560 | 1.87E-04 | 2.30E-03 | 5.14E-03 | 3.21E-04 | 8.19E-05 | 4.67E-05 | 1.97E-04 | 1.93E-04 | 8.75E-04 | 3.46E-04 | 8.43E-04 | Known_Known |
| g__Halolamina | 0.00E+00 | 1.94E-03 | 1.67E-03 | 2.75E-04 | 1.64E-04 | 2.10E-04 | 1.25E-04 | 5.15E-05 | 0.00E+00 | 4.55E-04 | 3.73E-04 | Known_Known |
| g__Halolacustris | 2.33E-05 | 4.09E-04 | 1.17E-04 | 2.29E-05 | 1.77E-04 | 0.00E+00 | 8.96E-05 | 5.15E-05 | 7.42E-05 | 9.11E-05 | 4.86E-05 | Known_Known |
| g__Halolactibacillus | 0.00E+00 | 0.00E+00 | 1.17E-05 | 0.00E+00 | 1.36E-05 | 0.00E+00 | 0.00E+00 | 2.58E-05 | 0.00E+00 | 0.00E+00 | 0.00E+00 | Known_Known |
| g__Haloimpatiens | 0.00E+00 | 0.00E+00 | 0.00E+00 | 1.14E-04 | 0.00E+00 | 0.00E+00 | 0.00E+00 | 0.00E+00 | 0.00E+00 | 0.00E+00 | 0.00E+00 | Known_Known |
| g__Halohasta | 6.41E-02 | 5.22E-02 | 1.15E-01 | 3.54E-02 | 2.80E-03 | 1.44E-02 | 6.61E-03 | 1.45E-03 | 3.35E-03 | 5.43E-03 | 4.88E-02 | Known_Known |
| g__Halogeometricum | 0.00E+00 | 0.00E+00 | 0.00E+00 | 0.00E+00 | 0.00E+00 | 7.01E-05 | 0.00E+00 | 0.00E+00 | 0.00E+00 | 0.00E+00 | 0.00E+00 | Known_Known |
| g__Haloferax | 0.00E+00 | 2.55E-04 | 2.00E-04 | 0.00E+00 | 0.00E+00 | 0.00E+00 | 0.00E+00 | 0.00E+00 | 2.23E-04 | 1.27E-04 | 9.72E-05 | Known_Known |
| g__Halodesulfurarchaeum | 9.80E-04 | 1.16E-03 | 2.04E-03 | 3.66E-04 | 1.23E-04 | 5.95E-04 | 2.97E-03 | 5.41E-04 | 7.42E-05 | 1.64E-04 | 8.59E-04 | Known_Known |
| g__Halococcus | 0.00E+00 | 0.00E+00 | 0.00E+00 | 0.00E+00 | 8.19E-05 | 1.87E-04 | 0.00E+00 | 0.00E+00 | 8.90E-05 | 2.00E-04 | 1.30E-04 | Known_Known |
| g__Halocella_43036 | 7.00E-05 | 8.52E-05 | 5.87E-05 | 2.29E-05 | 6.96E-04 | 2.57E-04 | 4.84E-04 | 2.06E-04 | 3.41E-04 | 8.92E-04 | 3.40E-04 | Known_Known |
| g__Halobium | 2.33E-04 | 0.00E+00 | 7.04E-05 | 0.00E+00 | 0.00E+00 | 1.05E-04 | 7.16E-05 | 0.00E+00 | 0.00E+00 | 0.00E+00 | 0.00E+00 | Known_Known |
| g__Halobiforma | 2.33E-05 | 0.00E+00 | 0.00E+00 | 0.00E+00 | 0.00E+00 | 0.00E+00 | 0.00E+00 | 0.00E+00 | 0.00E+00 | 0.00E+00 | 0.00E+00 | Known_Known |

| Genus | L1 | L2 | L3 | L4 | L5 | L6 | L7 | L8 | L9 | L10 | L11 | Microbial_matter |
|---|---|---|---|---|---|---|---|---|---|---|---|---|
| g__Halobellus_3029 | 0.00E+00 | 3.41E-04 | 1.64E-04 | 0.00E+00 | 0.00E+00 | 0.00E+00 | 0.00E+00 | 0.00E+00 | 2.97E-05 | 0.00E+00 | 9.72E-05 | Known_Known |
| g__Halobellus_3026 | 0.00E+00 | 5.62E-04 | 2.23E-04 | 8.01E-05 | 1.23E-04 | 8.17E-05 | 1.43E-04 | 0.00E+00 | 0.00E+00 | 0.00E+00 | 0.00E+00 | Known_Known |
| g__Halobaculum | 0.00E+00 | 7.66E-04 | 6.22E-04 | 4.58E-05 | 1.77E-04 | 5.84E-05 | 0.00E+00 | 0.00E+00 | 0.00E+00 | 9.11E-05 | 4.05E-04 | Known_Known |
| g__Halobacterium | 0.00E+00 | 9.37E-04 | 1.07E-03 | 1.95E-04 | 0.00E+00 | 0.00E+00 | 0.00E+00 | 0.00E+00 | 1.04E-04 | 0.00E+00 | 1.30E-04 | Known_Known |
| g__Halobacillus_A_288222 | 2.33E-05 | 0.00E+00 | 2.35E-05 | 0.00E+00 | 1.36E-05 | 0.00E+00 | 0.00E+00 | 0.00E+00 | 0.00E+00 | 7.29E-05 | 0.00E+00 | Known_Known |
| g__Haloarchaeobius | 0.00E+00 | 0.00E+00 | 0.00E+00 | 0.00E+00 | 0.00E+00 | 0.00E+00 | 0.00E+00 | 0.00E+00 | 0.00E+00 | 0.00E+00 | 1.13E-04 | Known_Known |
| g__Haliangium_437647 | 0.00E+00 | 1.70E-05 | 0.00E+00 | 0.00E+00 | 0.00E+00 | 0.00E+00 | 0.00E+00 | 0.00E+00 | 0.00E+00 | 0.00E+00 | 0.00E+00 | Known_Known |
| g__Halapricum_3981 | 0.00E+00 | 0.00E+00 | 0.00E+00 | 0.00E+00 | 0.00E+00 | 8.17E-05 | 0.00E+00 | 0.00E+00 | 0.00E+00 | 0.00E+00 | 0.00E+00 | Known_Known |
| g__Halapricum_3815 | 0.00E+00 | 0.00E+00 | 0.00E+00 | 0.00E+00 | 0.00E+00 | 0.00E+00 | 0.00E+00 | 6.44E-05 | 0.00E+00 | 0.00E+00 | 0.00E+00 | Known_Known |
| g__Halanaerobium | 4.43E-04 | 1.02E-04 | 0.00E+00 | 0.00E+00 | 4.09E-05 | 2.34E-05 | 3.58E-05 | 3.86E-05 | 5.93E-05 | 9.11E-04 | 3.89E-04 | Known_Known |
| g__Halanaerobacter | 0.00E+00 | 0.00E+00 | 0.00E+00 | 0.00E+00 | 0.00E+00 | 0.00E+00 | 0.00E+00 | 3.86E-05 | 0.00E+00 | 0.00E+00 | 0.00E+00 | Known_Known |
| g__Halanaeroarchaeum | 3.50E-04 | 0.00E+00 | 0.00E+00 | 0.00E+00 | 0.00E+00 | 5.72E-04 | 7.70E-04 | 0.00E+00 | 0.00E+00 | 0.00E+00 | 0.00E+00 | Known_Known |
| g__Halalkalirubrum | 0.00E+00 | 0.00E+00 | 2.47E-04 | 0.00E+00 | 0.00E+00 | 0.00E+00 | 8.96E-05 | 0.00E+00 | 8.90E-05 | 0.00E+00 | 0.00E+00 | Known_Known |
| g__Halalkalicoccus | 1.24E-03 | 7.02E-03 | 4.80E-03 | 0.00E+00 | 1.11E-03 | 2.32E-03 | 1.41E-03 | 0.00E+00 | 3.71E-04 | 2.73E-04 | 7.13E-04 | Known_Known |
| g__Halalkaliarchaeum | 2.80E-04 | 3.58E-04 | 1.57E-03 | 1.26E-04 | 0.00E+00 | 0.00E+00 | 0.00E+00 | 0.00E+00 | 0.00E+00 | 0.00E+00 | 9.72E-05 | Known_Known |
| g__Haladaptatus | 1.38E-03 | 7.99E-03 | 1.94E-03 | 5.95E-04 | 6.28E-04 | 1.79E-03 | 1.04E-03 | 1.61E-03 | 1.45E-03 | 1.66E-03 | 2.40E-03 | Known_Known |
| g__GWF2-39-127 | 1.40E-04 | 0.00E+00 | 0.00E+00 | 0.00E+00 | 0.00E+00 | 0.00E+00 | 1.07E-04 | 5.15E-05 | 4.45E-05 | 0.00E+00 | 1.78E-04 | Known_Known |
| g__GWF2-37-15 | 6.44E-03 | 6.64E-04 | 1.53E-04 | 9.16E-04 | 1.04E-03 | 3.27E-04 | 5.36E-03 | 1.22E-02 | 8.31E-04 | 1.37E-03 | 1.33E-02 | Known_Known |
| g__GWE2-41-16 | 0.00E+00 | 0.00E+00 | 0.00E+00 | 0.00E+00 | 0.00E+00 | 0.00E+00 | 0.00E+00 | 0.00E+00 | 0.00E+00 | 3.64E-05 | 0.00E+00 | Known_Known |
| g__GWC2-73-18 | 0.00E+00 | 1.29E-03 | 1.88E-04 | 2.07E-03 | 8.19E-04 | 1.05E-04 | 5.37E-05 | 2.45E-04 | 3.56E-04 | 2.19E-04 | 5.83E-04 | Known_Known |
| g__GWC2-33-13 | 0.00E+00 | 0.00E+00 | 0.00E+00 | 2.52E-04 | 0.00E+00 | 0.00E+00 | 0.00E+00 | 0.00E+00 | 0.00E+00 | 5.46E-05 | 6.48E-05 | Known_Known |
| g__GWC1-49-16 | 0.00E+00 | 0.00E+00 | 0.00E+00 | 0.00E+00 | 0.00E+00 | 0.00E+00 | 0.00E+00 | 0.00E+00 | 1.34E-04 | 0.00E+00 | 0.00E+00 | Known_Known |
| g__GWB1-41-6 | 0.00E+00 | 0.00E+00 | 0.00E+00 | 0.00E+00 | 0.00E+00 | 0.00E+00 | 0.00E+00 | 2.58E-05 | 0.00E+00 | 0.00E+00 | 0.00E+00 | Known_Known |
| g__GWA2-35-9 | 0.00E+00 | 0.00E+00 | 0.00E+00 | 2.29E-05 | 0.00E+00 | 0.00E+00 | 0.00E+00 | 0.00E+00 | 0.00E+00 | 0.00E+00 | 0.00E+00 | Known_Known |
| g__Guyparkeria | 0.00E+00 | 6.81E-05 | 1.41E-04 | 0.00E+00 | 4.09E-05 | 0.00E+00 | 0.00E+00 | 0.00E+00 | 0.00E+00 | 2.00E-04 | 1.94E-04 | Known_Known |
| g__Gracilibacter | 0.00E+00 | 9.54E-04 | 5.28E-04 | 3.78E-04 | 4.23E-04 | 0.00E+00 | 0.00E+00 | 1.03E-04 | 0.00E+00 | 6.01E-04 | 8.43E-04 | Known_Known |
| g__Gottschalkia_A | 0.00E+00 | 0.00E+00 | 0.00E+00 | 0.00E+00 | 0.00E+00 | 0.00E+00 | 0.00E+00 | 0.00E+00 | 0.00E+00 | 3.64E-05 | 0.00E+00 | Known_Known |
| g__Glycocaulis | 0.00E+00 | 4.60E-04 | 1.88E-04 | 7.10E-04 | 0.00E+00 | 0.00E+00 | 0.00E+00 | 5.15E-05 | 0.00E+00 | 0.00E+00 | 1.30E-04 | Known_Known |
| g__Gloeocapsa | 1.87E-04 | 0.00E+00 | 0.00E+00 | 0.00E+00 | 0.00E+00 | 0.00E+00 | 0.00E+00 | 3.86E-05 | 0.00E+00 | 0.00E+00 | 1.13E-04 | Known_Known |
| g__Geothermomicrobium | 0.00E+00 | 0.00E+00 | 0.00E+00 | 0.00E+00 | 2.05E-04 | 4.67E-05 | 0.00E+00 | 0.00E+00 | 0.00E+00 | 0.00E+00 | 0.00E+00 | Known_Known |
| g__Geosporobacter_234399 | 0.00E+00 | 4.94E-04 | 4.70E-04 | 0.00E+00 | 4.09E-05 | 0.00E+00 | 0.00E+00 | 0.00E+00 | 4.60E-04 | 0.00E+00 | 0.00E+00 | Known_Known |
| g__Geodermatophilus_A | 0.00E+00 | 3.41E-05 | 2.35E-05 | 0.00E+00 | 8.46E-04 | 1.52E-04 | 8.96E-05 | 5.15E-05 | 1.48E-05 | 0.00E+00 | 1.30E-04 | Known_Known |
| g__Gemmiger_A_73129 | 0.00E+00 | 0.00E+00 | 0.00E+00 | 0.00E+00 | 0.00E+00 | 0.00E+00 | 0.00E+00 | 0.00E+00 | 0.00E+00 | 5.46E-05 | 0.00E+00 | Known_Known |
| g__Geminicoccus | 0.00E+00 | 1.19E-04 | 0.00E+00 | 0.00E+00 | 0.00E+00 | 0.00E+00 | 0.00E+00 | 0.00E+00 | 0.00E+00 | 0.00E+00 | 0.00E+00 | Known_Known |
| g__GCA-2770975 | 0.00E+00 | 1.70E-05 | 0.00E+00 | 2.29E-05 | 0.00E+00 | 0.00E+00 | 0.00E+00 | 2.58E-05 | 0.00E+00 | 0.00E+00 | 0.00E+00 | Known_Known |
| g__GCA-2746885 | 0.00E+00 | 0.00E+00 | 0.00E+00 | 2.29E-05 | 0.00E+00 | 0.00E+00 | 0.00E+00 | 7.73E-05 | 0.00E+00 | 0.00E+00 | 0.00E+00 | Known_Known |
| g__GCA-2733885 | 0.00E+00 | 0.00E+00 | 0.00E+00 | 0.00E+00 | 0.00E+00 | 0.00E+00 | 0.00E+00 | 0.00E+00 | 0.00E+00 | 8.74E-04 | 1.46E-04 | Known_Known |
| g__GCA-2726865 | 3.50E-04 | 5.28E-04 | 0.00E+00 | 2.52E-04 | 1.09E-04 | 0.00E+00 | 2.69E-04 | 1.89E-03 | 3.09E-02 | 1.57E-02 | 1.42E-02 | Known_Known |
| g__GCA-2699585 | 1.40E-04 | 0.00E+00 | 2.35E-05 | 2.75E-04 | 0.00E+00 | 0.00E+00 | 1.25E-04 | 5.15E-05 | 0.00E+00 | 0.00E+00 | 9.72E-05 | Known_Known |
| g__GCA-2699025 | 0.00E+00 | 0.00E+00 | 0.00E+00 | 0.00E+00 | 0.00E+00 | 0.00E+00 | 0.00E+00 | 0.00E+00 | 0.00E+00 | 5.46E-05 | 0.00E+00 | Known_Known |
| g__GCA-2401425 | 0.00E+00 | 0.00E+00 | 0.00E+00 | 0.00E+00 | 0.00E+00 | 0.00E+00 | 0.00E+00 | 0.00E+00 | 5.93E-05 | 0.00E+00 | 0.00E+00 | Known_Known |
| g__Ga0077555 | 0.00E+00 | 0.00E+00 | 0.00E+00 | 4.58E-05 | 1.23E-04 | 0.00E+00 | 0.00E+00 | 0.00E+00 | 0.00E+00 | 5.46E-05 | 0.00E+00 | Known_Known |
| g__Francisella_A | 0.00E+00 | 0.00E+00 | 0.00E+00 | 2.29E-05 | 0.00E+00 | 0.00E+00 | 0.00E+00 | 0.00E+00 | 0.00E+00 | 0.00E+00 | 0.00E+00 | Known_Known |
| g__Fodinicurvata | 0.00E+00 | 4.43E-04 | 3.52E-05 | 6.87E-05 | 0.00E+00 | 0.00E+00 | 0.00E+00 | 6.44E-05 | 0.00E+00 | 0.00E+00 | 0.00E+00 | Known_Known |
| g__FMMP01 | 1.17E-04 | 1.36E-04 | 0.00E+00 | 1.49E-04 | 5.59E-04 | 5.95E-04 | 0.00E+00 | 0.00E+00 | 0.00E+00 | 0.00E+00 | 0.00E+00 | Known_Known |
| g__Filobacillus | 0.00E+00 | 0.00E+00 | 0.00E+00 | 0.00E+00 | 1.36E-05 | 0.00E+00 | 0.00E+00 | 0.00E+00 | 0.00E+00 | 0.00E+00 | 0.00E+00 | Known_Known |
| g__Fictibacillus_C | 0.00E+00 | 0.00E+00 | 0.00E+00 | 0.00E+00 | 0.00E+00 | 7.01E-05 | 0.00E+00 | 0.00E+00 | 0.00E+00 | 0.00E+00 | 0.00E+00 | Known_Known |

| Genus | L1 | L2 | L3 | L4 | L5 | L6 | L7 | L8 | L9 | L10 | L11 | Microbial_matter |
|---|---|---|---|---|---|---|---|---|---|---|---|---|
| g__Ferruginivarius | 4.90E-04 | 7.66E-04 | 1.06E-04 | 1.25E-03 | 0.00E+00 | 0.00E+00 | 0.00E+00 | 0.00E+00 | 0.00E+00 | 1.46E-04 | 0.00E+00 | Known_Known |
| g__FEN-33 | 0.00E+00 | 5.79E-04 | 1.17E-04 | 1.03E-04 | 0.00E+00 | 0.00E+00 | 0.00E+00 | 1.53E-03 | 1.26E-02 | 3.81E-03 | 3.99E-03 | Known_Known |
| g__FEN-1185 | 0.00E+00 | 0.00E+00 | 5.87E-05 | 0.00E+00 | 0.00E+00 | 0.00E+00 | 0.00E+00 | 0.00E+00 | 4.45E-05 | 0.00E+00 | 0.00E+00 | Known_Known |
| g__F1-60-MAGs027 | 0.00E+00 | 0.00E+00 | 0.00E+00 | 0.00E+00 | 1.09E-04 | 8.17E-05 | 0.00E+00 | 0.00E+00 | 0.00E+00 | 0.00E+00 | 0.00E+00 | Known_Known |
| g__EX4484-6 | 1.05E-03 | 1.09E-03 | 3.87E-04 | 1.10E-03 | 1.47E-03 | 1.28E-03 | 2.02E-03 | 4.94E-03 | 3.96E-03 | 4.23E-03 | 3.44E-03 | Known_Known |
| g__EX4484-52 | 0.00E+00 | 6.81E-05 | 0.00E+00 | 0.00E+00 | 0.00E+00 | 0.00E+00 | 0.00E+00 | 0.00E+00 | 0.00E+00 | 0.00E+00 | 0.00E+00 | Known_Known |
| g__EX4484-2 | 9.33E-05 | 0.00E+00 | 0.00E+00 | 0.00E+00 | 0.00E+00 | 0.00E+00 | 0.00E+00 | 0.00E+00 | 0.00E+00 | 9.11E-05 | 0.00E+00 | Known_Known |
| g__Evtepia | 0.00E+00 | 0.00E+00 | 0.00E+00 | 0.00E+00 | 0.00E+00 | 0.00E+00 | 0.00E+00 | 0.00E+00 | 8.90E-05 | 0.00E+00 | 0.00E+00 | Known_Known |
| g__Erythrobacter_484280 | 0.00E+00 | 0.00E+00 | 7.63E-04 | 1.50E-03 | 0.00E+00 | 0.00E+00 | 0.00E+00 | 3.35E-04 | 0.00E+00 | 0.00E+00 | 1.13E-04 | Known_Known |
| g__Entotheonella | 0.00E+00 | 0.00E+00 | 0.00E+00 | 0.00E+00 | 1.77E-04 | 0.00E+00 | 0.00E+00 | 2.58E-05 | 5.93E-05 | 0.00E+00 | 0.00E+00 | Known_Known |
| g__Ensifer | 1.17E-04 | 5.96E-04 | 2.93E-04 | 9.50E-04 | 3.96E-04 | 6.30E-04 | 0.00E+00 | 0.00E+00 | 0.00E+00 | 0.00E+00 | 1.78E-04 | Known_Known |
| g__Enhygromyxa | 0.00E+00 | 0.00E+00 | 0.00E+00 | 0.00E+00 | 0.00E+00 | 0.00E+00 | 0.00E+00 | 6.44E-05 | 1.48E-05 | 0.00E+00 | 0.00E+00 | Known_Known |
| g__Endothiovibrio | 0.00E+00 | 0.00E+00 | 0.00E+00 | 0.00E+00 | 0.00E+00 | 0.00E+00 | 0.00E+00 | 0.00E+00 | 4.45E-05 | 0.00E+00 | 0.00E+00 | Known_Known |
| g__Elainella | 0.00E+00 | 5.11E-05 | 0.00E+00 | 0.00E+00 | 0.00E+00 | 0.00E+00 | 0.00E+00 | 0.00E+00 | 0.00E+00 | 0.00E+00 | 0.00E+00 | Known_Known |
| g__Ectothiorhodospira_607903 | 0.00E+00 | 0.00E+00 | 0.00E+00 | 1.14E-05 | 0.00E+00 | 0.00E+00 | 0.00E+00 | 0.00E+00 | 0.00E+00 | 0.00E+00 | 0.00E+00 | Known_Known |
| g__E85 | 0.00E+00 | 0.00E+00 | 3.52E-05 | 0.00E+00 | 0.00E+00 | 0.00E+00 | 0.00E+00 | 0.00E+00 | 0.00E+00 | 1.82E-05 | 0.00E+00 | Known_Known |
| g__E44-bin10 | 0.00E+00 | 0.00E+00 | 0.00E+00 | 0.00E+00 | 0.00E+00 | 0.00E+00 | 0.00E+00 | 9.01E-05 | 1.99E-03 | 2.73E-03 | 2.30E-03 | Known_Known |
| g__Dwaynesavagella | 0.00E+00 | 0.00E+00 | 0.00E+00 | 0.00E+00 | 0.00E+00 | 0.00E+00 | 0.00E+00 | 0.00E+00 | 0.00E+00 | 9.11E-05 | 0.00E+00 | Known_Known |
| g__DTU015 | 0.00E+00 | 1.02E-04 | 1.41E-04 | 2.63E-04 | 1.23E-04 | 2.34E-04 | 3.22E-04 | 5.54E-04 | 3.69E-03 | 2.77E-03 | 8.75E-04 | Known_Known |
| g__DTU012 | 1.09E-02 | 9.57E-03 | 8.04E-03 | 1.47E-03 | 5.23E-02 | 1.89E-02 | 1.78E-02 | 8.05E-03 | 1.16E-02 | 4.01E-03 | 1.14E-02 | Known_Known |
| g__DTJY01 | 0.00E+00 | 1.53E-04 | 0.00E+00 | 0.00E+00 | 0.00E+00 | 0.00E+00 | 0.00E+00 | 0.00E+00 | 0.00E+00 | 0.00E+00 | 0.00E+00 | Known_Known |
| g__DTIZ01 | 0.00E+00 | 0.00E+00 | 0.00E+00 | 0.00E+00 | 0.00E+00 | 0.00E+00 | 0.00E+00 | 0.00E+00 | 1.63E-04 | 0.00E+00 | 8.10E-05 | Known_Known |
| g__DTFT01 | 0.00E+00 | 0.00E+00 | 0.00E+00 | 0.00E+00 | 0.00E+00 | 0.00E+00 | 0.00E+00 | 0.00E+00 | 0.00E+00 | 3.10E-04 | 0.00E+00 | Known_Known |
| g__DSYW01 | 0.00E+00 | 0.00E+00 | 0.00E+00 | 0.00E+00 | 0.00E+00 | 0.00E+00 | 0.00E+00 | 0.00E+00 | 1.04E-04 | 0.00E+00 | 0.00E+00 | Known_Known |
| g__DSXH01 | 0.00E+00 | 0.00E+00 | 0.00E+00 | 0.00E+00 | 0.00E+00 | 0.00E+00 | 0.00E+00 | 0.00E+00 | 0.00E+00 | 0.00E+00 | 1.62E-05 | Known_Known |
| g__DSWT01 | 0.00E+00 | 0.00E+00 | 0.00E+00 | 1.26E-04 | 2.73E-05 | 0.00E+00 | 0.00E+00 | 3.86E-05 | 2.97E-04 | 0.00E+00 | 3.24E-05 | Known_Known |
| g__DSUL01 | 0.00E+00 | 0.00E+00 | 0.00E+00 | 0.00E+00 | 0.00E+00 | 0.00E+00 | 0.00E+00 | 1.03E-04 | 2.67E-04 | 0.00E+00 | 0.00E+00 | Known_Known |
| g__DSTP01 | 0.00E+00 | 0.00E+00 | 3.52E-05 | 0.00E+00 | 0.00E+00 | 0.00E+00 | 0.00E+00 | 0.00E+00 | 0.00E+00 | 0.00E+00 | 0.00E+00 | Known_Known |
| g__DSQQ01 | 0.00E+00 | 0.00E+00 | 0.00E+00 | 0.00E+00 | 0.00E+00 | 0.00E+00 | 0.00E+00 | 0.00E+00 | 0.00E+00 | 7.29E-05 | 8.10E-05 | Known_Known |
| g__DSQK01 | 0.00E+00 | 5.11E-05 | 0.00E+00 | 0.00E+00 | 0.00E+00 | 0.00E+00 | 0.00E+00 | 0.00E+00 | 5.49E-04 | 0.00E+00 | 0.00E+00 | Known_Known |
| g__DSGV01 | 0.00E+00 | 0.00E+00 | 0.00E+00 | 4.58E-05 | 0.00E+00 | 4.67E-05 | 0.00E+00 | 0.00E+00 | 0.00E+00 | 0.00E+00 | 3.24E-05 | Known_Known |
| g__DSDE01 | 0.00E+00 | 0.00E+00 | 0.00E+00 | 0.00E+00 | 0.00E+00 | 0.00E+00 | 0.00E+00 | 1.29E-05 | 0.00E+00 | 0.00E+00 | 0.00E+00 | Known_Known |
| g__DSBN01 | 0.00E+00 | 1.70E-05 | 0.00E+00 | 0.00E+00 | 0.00E+00 | 0.00E+00 | 0.00E+00 | 5.15E-05 | 1.48E-05 | 9.11E-05 | 0.00E+00 | Known_Known |
| g__DSBA01 | 0.00E+00 | 0.00E+00 | 0.00E+00 | 0.00E+00 | 0.00E+00 | 0.00E+00 | 0.00E+00 | 5.15E-05 | 1.19E-04 | 2.00E-04 | 0.00E+00 | Known_Known |
| g__DSAU01 | 0.00E+00 | 0.00E+00 | 0.00E+00 | 0.00E+00 | 0.00E+00 | 0.00E+00 | 0.00E+00 | 0.00E+00 | 0.00E+00 | 1.82E-05 | 0.00E+00 | Known_Known |
| g__DQWO01 | 0.00E+00 | 0.00E+00 | 0.00E+00 | 0.00E+00 | 0.00E+00 | 0.00E+00 | 0.00E+00 | 0.00E+00 | 0.00E+00 | 5.46E-05 | 0.00E+00 | Known_Known |
| g__Dissulfurispira | 7.00E-05 | 1.19E-04 | 0.00E+00 | 5.72E-05 | 0.00E+00 | 0.00E+00 | 0.00E+00 | 1.15E-03 | 2.42E-03 | 2.19E-04 | 1.46E-04 | Known_Known |
| g__Dissulfurirhabdus | 0.00E+00 | 3.41E-05 | 0.00E+00 | 0.00E+00 | 0.00E+00 | 0.00E+00 | 0.00E+00 | 0.00E+00 | 0.00E+00 | 0.00E+00 | 6.48E-05 | Known_Known |
| g__Dietzia | 0.00E+00 | 0.00E+00 | 0.00E+00 | 0.00E+00 | 0.00E+00 | 0.00E+00 | 0.00E+00 | 0.00E+00 | 0.00E+00 | 0.00E+00 | 1.62E-05 | Known_Known |
| g__Dichotomicrobium | 3.27E-04 | 0.00E+00 | 2.35E-04 | 0.00E+00 | 1.36E-04 | 4.32E-04 | 4.30E-04 | 6.44E-05 | 0.00E+00 | 0.00E+00 | 7.94E-04 | Known_Known |
| g__Devosia_A_502124 | 0.00E+00 | 3.58E-04 | 1.88E-04 | 0.00E+00 | 0.00E+00 | 0.00E+00 | 0.00E+00 | 0.00E+00 | 0.00E+00 | 0.00E+00 | 0.00E+00 | Known_Known |
| g__Devosia_A_501803 | 0.00E+00 | 4.60E-04 | 3.40E-04 | 0.00E+00 | 8.19E-05 | 3.39E-04 | 8.96E-05 | 0.00E+00 | 0.00E+00 | 0.00E+00 | 0.00E+00 | Known_Known |
| g__Dethiosulfatibacter | 0.00E+00 | 0.00E+00 | 0.00E+00 | 6.18E-04 | 0.00E+00 | 0.00E+00 | 0.00E+00 | 0.00E+00 | 0.00E+00 | 0.00E+00 | 0.00E+00 | Known_Known |
| g__Dethiobacter | 2.58E-02 | 2.55E-03 | 3.99E-04 | 3.55E-04 | 4.09E-04 | 2.38E-03 | 6.40E-02 | 6.82E-03 | 3.86E-04 | 5.59E-03 | 9.38E-03 | Known_Known |
| g__Desulfuribacillus_A | 0.00E+00 | 0.00E+00 | 1.17E-05 | 0.00E+00 | 0.00E+00 | 0.00E+00 | 0.00E+00 | 0.00E+00 | 0.00E+00 | 1.82E-05 | 0.00E+00 | Known_Known |
| g__Desulfovibrio_R_446519 | 0.00E+00 | 0.00E+00 | 0.00E+00 | 0.00E+00 | 0.00E+00 | 0.00E+00 | 0.00E+00 | 0.00E+00 | 0.00E+00 | 0.00E+00 | 8.10E-05 | Known_Known |
| g__Desulfovermiculus | 4.67E-04 | 2.04E-04 | 2.70E-04 | 4.58E-05 | 8.19E-05 | 0.00E+00 | 2.15E-04 | 5.28E-04 | 0.00E+00 | 2.91E-04 | 2.76E-04 | Known_Known |

| Genus | L1 | L2 | L3 | L4 | L5 | L6 | L7 | L8 | L9 | L10 | L11 | Microbial_matter |
|---|---|---|---|---|---|---|---|---|---|---|---|---|
| g__Desulfotomaculum | 2.80E-04 | 2.84E-03 | 2.76E-03 | 2.75E-04 | 1.34E-03 | 1.12E-03 | 0.00E+00 | 1.16E-04 | 2.69E-03 | 5.46E-05 | 1.09E-03 | Known_Known |
| g__Desulfosporosinus | 0.00E+00 | 0.00E+00 | 0.00E+00 | 0.00E+00 | 1.91E-04 | 0.00E+00 | 0.00E+00 | 0.00E+00 | 0.00E+00 | 0.00E+00 | 9.72E-05 | Known_Known |
| g__Desulfosarcina | 0.00E+00 | 0.00E+00 | 0.00E+00 | 0.00E+00 | 0.00E+00 | 0.00E+00 | 0.00E+00 | 0.00E+00 | 4.45E-05 | 0.00E+00 | 0.00E+00 | Known_Known |
| g__Desulfosalsimonas | 9.33E-05 | 5.11E-05 | 0.00E+00 | 0.00E+00 | 0.00E+00 | 2.34E-05 | 8.24E-04 | 2.06E-04 | 0.00E+00 | 0.00E+00 | 1.46E-04 | Known_Known |
| g__Desulforhopalus | 4.67E-05 | 0.00E+00 | 0.00E+00 | 0.00E+00 | 0.00E+00 | 0.00E+00 | 0.00E+00 | 0.00E+00 | 0.00E+00 | 0.00E+00 | 0.00E+00 | Known_Known |
| g__Desulfonispora | 0.00E+00 | 0.00E+00 | 0.00E+00 | 0.00E+00 | 0.00E+00 | 3.50E-05 | 0.00E+00 | 0.00E+00 | 0.00E+00 | 0.00E+00 | 0.00E+00 | Known_Known |
| g__Desulfonatronovibrio | 4.67E-05 | 0.00E+00 | 3.52E-05 | 0.00E+00 | 0.00E+00 | 3.50E-05 | 0.00E+00 | 6.44E-05 | 0.00E+00 | 0.00E+00 | 0.00E+00 | Known_Known |
| g__Desulfonatronobacter | 9.33E-05 | 1.70E-05 | 2.58E-04 | 6.87E-05 | 1.09E-04 | 0.00E+00 | 5.37E-05 | 1.55E-04 | 1.34E-04 | 4.74E-04 | 1.13E-04 | Known_Known |
| g__Desulfohalotomaculum | 0.00E+00 | 0.00E+00 | 0.00E+00 | 0.00E+00 | 0.00E+00 | 0.00E+00 | 0.00E+00 | 0.00E+00 | 0.00E+00 | 9.11E-05 | 0.00E+00 | Known_Known |
| g__Desulfocucumis | 0.00E+00 | 0.00E+00 | 1.88E-04 | 0.00E+00 | 0.00E+00 | 0.00E+00 | 0.00E+00 | 0.00E+00 | 0.00E+00 | 0.00E+00 | 0.00E+00 | Known_Known |
| g__Desulfitibacter | 0.00E+00 | 0.00E+00 | 0.00E+00 | 0.00E+00 | 4.09E-05 | 0.00E+00 | 0.00E+00 | 0.00E+00 | 0.00E+00 | 0.00E+00 | 0.00E+00 | Known_Known |
| g__Desulfatitalea | 0.00E+00 | 0.00E+00 | 0.00E+00 | 0.00E+00 | 0.00E+00 | 0.00E+00 | 0.00E+00 | 0.00E+00 | 0.00E+00 | 9.11E-05 | 6.48E-05 | Known_Known |
| g__Desulfatiglans | 7.00E-05 | 0.00E+00 | 9.39E-05 | 0.00E+00 | 1.36E-05 | 0.00E+00 | 1.25E-04 | 6.44E-05 | 0.00E+00 | 3.28E-04 | 0.00E+00 | Known_Known |
| g__Desnuesiella | 2.10E-04 | 2.06E-03 | 1.50E-03 | 8.36E-04 | 2.05E-03 | 1.63E-03 | 0.00E+00 | 1.16E-04 | 2.31E-03 | 1.09E-04 | 5.67E-04 | Known_Known |
| g__Deserticella | 0.00E+00 | 7.49E-04 | 1.53E-04 | 0.00E+00 | 0.00E+00 | 7.01E-05 | 0.00E+00 | 0.00E+00 | 0.00E+00 | 0.00E+00 | 0.00E+00 | Known_Known |
| g__Desertibacter | 0.00E+00 | 1.02E-03 | 1.88E-04 | 0.00E+00 | 3.14E-04 | 4.90E-04 | 0.00E+00 | 0.00E+00 | 0.00E+00 | 0.00E+00 | 1.62E-04 | Known_Known |
| g__Defluviitalea_A | 1.40E-03 | 8.52E-05 | 8.22E-04 | 1.03E-03 | 5.61E-03 | 6.43E-03 | 2.70E-03 | 3.03E-03 | 1.16E-03 | 1.04E-03 | 2.22E-03 | Known_Known |
| g__Defluviicoccus_A | 0.00E+00 | 0.00E+00 | 0.00E+00 | 1.64E-03 | 8.19E-05 | 0.00E+00 | 0.00E+00 | 0.00E+00 | 0.00E+00 | 0.00E+00 | 0.00E+00 | Known_Known |
| g__Cytobacillus_294142 | 0.00E+00 | 0.00E+00 | 0.00E+00 | 0.00E+00 | 0.00E+00 | 0.00E+00 | 5.37E-05 | 0.00E+00 | 0.00E+00 | 0.00E+00 | 0.00E+00 | Known_Known |
| g__CSSed11-175R1 | 0.00E+00 | 0.00E+00 | 0.00E+00 | 0.00E+00 | 0.00E+00 | 0.00E+00 | 0.00E+00 | 0.00E+00 | 0.00E+00 | 0.00E+00 | 4.86E-05 | Known_Known |
| g__CSSED11-154 | 5.60E-03 | 8.14E-03 | 1.04E-03 | 1.29E-03 | 6.55E-04 | 8.87E-04 | 2.38E-03 | 2.16E-03 | 1.37E-03 | 2.37E-04 | 2.24E-03 | Known_Known |
| g__CSSed10-48 | 5.93E-03 | 3.73E-03 | 3.64E-04 | 2.52E-04 | 1.91E-04 | 2.92E-04 | 1.61E-04 | 1.03E-03 | 9.20E-04 | 5.41E-03 | 2.66E-03 | Known_Known |
| g__CSP1-1 | 0.00E+00 | 0.00E+00 | 0.00E+00 | 0.00E+00 | 0.00E+00 | 0.00E+00 | 0.00E+00 | 0.00E+00 | 8.75E-04 | 0.00E+00 | 0.00E+00 | Known_Known |
| g__CSBr16-57R1 | 0.00E+00 | 0.00E+00 | 0.00E+00 | 0.00E+00 | 0.00E+00 | 0.00E+00 | 0.00E+00 | 0.00E+00 | 0.00E+00 | 0.00E+00 | 3.24E-05 | Known_Known |
| g__CSBR16-193 | 0.00E+00 | 5.11E-05 | 0.00E+00 | 0.00E+00 | 0.00E+00 | 0.00E+00 | 0.00E+00 | 0.00E+00 | 0.00E+00 | 0.00E+00 | 0.00E+00 | Known_Known |
| g__Croceicoccus_484963 | 0.00E+00 | 0.00E+00 | 0.00E+00 | 0.00E+00 | 0.00E+00 | 7.01E-05 | 0.00E+00 | 0.00E+00 | 0.00E+00 | 0.00E+00 | 0.00E+00 | Known_Known |
| g__Croceibacterium_484881 | 0.00E+00 | 3.41E-04 | 1.76E-04 | 0.00E+00 | 9.55E-05 | 6.19E-04 | 0.00E+00 | 0.00E+00 | 0.00E+00 | 5.46E-05 | 2.92E-04 | Known_Known |
| g__Craurococcus | 0.00E+00 | 2.21E-04 | 1.53E-04 | 0.00E+00 | 6.00E-04 | 5.84E-04 | 0.00E+00 | 7.73E-05 | 0.00E+00 | 0.00E+00 | 2.59E-04 | Known_Known |
| g__Crassaminicella_234356 | 1.17E-03 | 6.81E-05 | 2.58E-04 | 1.60E-04 | 1.60E-03 | 2.57E-03 | 1.00E-03 | 1.20E-03 | 5.77E-03 | 3.46E-04 | 2.56E-03 | Known_Known |
| g__CR-1 | 0.00E+00 | 0.00E+00 | 0.00E+00 | 0.00E+00 | 0.00E+00 | 0.00E+00 | 0.00E+00 | 3.86E-05 | 0.00E+00 | 0.00E+00 | 0.00E+00 | Known_Known |
| g__Congregibacter | 2.33E-05 | 0.00E+00 | 0.00E+00 | 2.29E-05 | 0.00E+00 | 0.00E+00 | 0.00E+00 | 0.00E+00 | 0.00E+00 | 0.00E+00 | 0.00E+00 | Known_Known |
| g__Conexibacter | 0.00E+00 | 0.00E+00 | 0.00E+00 | 0.00E+00 | 4.09E-05 | 0.00E+00 | 3.58E-05 | 0.00E+00 | 2.97E-05 | 0.00E+00 | 0.00E+00 | Known_Known |
| g__Cohnella | 0.00E+00 | 0.00E+00 | 0.00E+00 | 0.00E+00 | 1.36E-05 | 0.00E+00 | 0.00E+00 | 0.00E+00 | 0.00E+00 | 0.00E+00 | 0.00E+00 | Known_Known |
| g__Clostridium_T | 0.00E+00 | 0.00E+00 | 0.00E+00 | 4.58E-05 | 8.87E-04 | 0.00E+00 | 0.00E+00 | 3.86E-05 | 0.00E+00 | 0.00E+00 | 0.00E+00 | Known_Known |
| g__Clostridium_S_216331 | 0.00E+00 | 2.73E-04 | 1.06E-04 | 0.00E+00 | 0.00E+00 | 0.00E+00 | 0.00E+00 | 0.00E+00 | 8.90E-05 | 0.00E+00 | 0.00E+00 | Known_Known |
| g__Clostridium_J | 0.00E+00 | 0.00E+00 | 0.00E+00 | 0.00E+00 | 0.00E+00 | 0.00E+00 | 0.00E+00 | 2.32E-04 | 6.97E-04 | 0.00E+00 | 2.11E-04 | Known_Known |
| g__Clostridium_AR_212534 | 0.00E+00 | 0.00E+00 | 0.00E+00 | 5.72E-05 | 0.00E+00 | 0.00E+00 | 0.00E+00 | 0.00E+00 | 0.00E+00 | 0.00E+00 | 0.00E+00 | Known_Known |
| g__Clostridium_AR_193524 | 7.00E-05 | 0.00E+00 | 0.00E+00 | 0.00E+00 | 0.00E+00 | 0.00E+00 | 0.00E+00 | 0.00E+00 | 0.00E+00 | 0.00E+00 | 0.00E+00 | Known_Known |
| g__Clostridium_AM | 0.00E+00 | 1.19E-04 | 2.00E-04 | 0.00E+00 | 6.82E-05 | 0.00E+00 | 0.00E+00 | 0.00E+00 | 0.00E+00 | 0.00E+00 | 0.00E+00 | Known_Known |
| g__Clostridium_AF | 0.00E+00 | 0.00E+00 | 0.00E+00 | 0.00E+00 | 1.64E-04 | 0.00E+00 | 0.00E+00 | 0.00E+00 | 0.00E+00 | 0.00E+00 | 0.00E+00 | Known_Known |
| g__Clostridium_AD | 0.00E+00 | 0.00E+00 | 0.00E+00 | 0.00E+00 | 0.00E+00 | 0.00E+00 | 0.00E+00 | 0.00E+00 | 1.78E-04 | 0.00E+00 | 0.00E+00 | Known_Known |
| g__Clostridiisalibacter | 0.00E+00 | 0.00E+00 | 0.00E+00 | 0.00E+00 | 0.00E+00 | 0.00E+00 | 0.00E+00 | 0.00E+00 | 7.42E-05 | 0.00E+00 | 0.00E+00 | Known_Known |
| g__Chloroploca | 0.00E+00 | 2.73E-04 | 0.00E+00 | 5.84E-04 | 2.73E-05 | 0.00E+00 | 0.00E+00 | 1.03E-04 | 5.93E-04 | 7.10E-04 | 1.13E-04 | Known_Known |
| g__CG2-30-64-16 | 0.00E+00 | 0.00E+00 | 0.00E+00 | 4.58E-05 | 0.00E+00 | 0.00E+00 | 0.00E+00 | 2.96E-04 | 1.31E-03 | 1.49E-03 | 5.67E-04 | Known_Known |
| g__CG2-30-51-40 | 0.00E+00 | 5.11E-05 | 0.00E+00 | 0.00E+00 | 4.09E-05 | 0.00E+00 | 0.00E+00 | 3.86E-05 | 0.00E+00 | 9.11E-05 | 0.00E+00 | Known_Known |
| g__CG2-30-33-13 | 3.73E-04 | 2.21E-04 | 9.39E-05 | 0.00E+00 | 1.77E-04 | 0.00E+00 | 1.97E-04 | 6.44E-05 | 4.45E-05 | 7.29E-05 | 2.11E-04 | Known_Known |
| g__CG1-02-57-44 | 1.82E-03 | 0.00E+00 | 2.35E-05 | 6.87E-04 | 5.46E-05 | 1.98E-04 | 8.96E-04 | 1.55E-04 | 8.90E-05 | 1.46E-04 | 7.45E-04 | Known_Known |

| Genus | L1 | L2 | L3 | L4 | L5 | L6 | L7 | L8 | L9 | L10 | L11 | Microbial_matter |
|---|---|---|---|---|---|---|---|---|---|---|---|---|
| g__CF-167 | 0.00E+00 | 2.21E-04 | 0.00E+00 | 0.00E+00 | 2.32E-04 | 9.34E-05 | 0.00E+00 | 0.00E+00 | 0.00E+00 | 5.46E-05 | 8.10E-05 | Known_Known |
| g__CF-163 | 0.00E+00 | 0.00E+00 | 0.00E+00 | 0.00E+00 | 0.00E+00 | 0.00E+00 | 0.00E+00 | 0.00E+00 | 8.90E-05 | 1.09E-04 | 3.24E-05 | Known_Known |
| g__Cereibacter_A_494128 | 0.00E+00 | 0.00E+00 | 0.00E+00 | 0.00E+00 | 1.09E-04 | 0.00E+00 | 0.00E+00 | 0.00E+00 | 0.00E+00 | 0.00E+00 | 0.00E+00 | Known_Known |
| g__Cephalothrix | 0.00E+00 | 0.00E+00 | 0.00E+00 | 1.83E-04 | 8.19E-05 | 0.00E+00 | 5.37E-05 | 0.00E+00 | 0.00E+00 | 0.00E+00 | 0.00E+00 | Known_Known |
| g__Cellulomonas | 0.00E+00 | 0.00E+00 | 0.00E+00 | 0.00E+00 | 1.36E-05 | 0.00E+00 | 0.00E+00 | 0.00E+00 | 0.00E+00 | 0.00E+00 | 0.00E+00 | Known_Known |
| g__CCUG-7971 | 0.00E+00 | 2.04E-04 | 0.00E+00 | 0.00E+00 | 1.36E-04 | 2.69E-04 | 8.96E-05 | 0.00E+00 | 1.48E-04 | 1.82E-04 | 0.00E+00 | Known_Known |
| g__Caloranaerobacter | 2.33E-04 | 0.00E+00 | 0.00E+00 | 0.00E+00 | 0.00E+00 | 0.00E+00 | 4.66E-04 | 9.27E-04 | 4.45E-05 | 8.74E-04 | 1.94E-04 | Known_Known |
| g__Calidifontibacillus | 0.00E+00 | 0.00E+00 | 0.00E+00 | 0.00E+00 | 4.09E-05 | 0.00E+00 | 0.00E+00 | 0.00E+00 | 0.00E+00 | 0.00E+00 | 0.00E+00 | Known_Known |
| g__Caldovatus | 9.33E-05 | 4.43E-04 | 1.06E-04 | 0.00E+00 | 4.57E-03 | 3.43E-03 | 1.61E-04 | 0.00E+00 | 7.12E-04 | 4.55E-04 | 1.62E-04 | Known_Known |
| g__Caldicellulosiruptor | 0.00E+00 | 0.00E+00 | 0.00E+00 | 0.00E+00 | 0.00E+00 | 0.00E+00 | 7.16E-05 | 0.00E+00 | 0.00E+00 | 0.00E+00 | 0.00E+00 | Known_Known |
| g__Caenispirillum | 0.00E+00 | 0.00E+00 | 9.39E-05 | 0.00E+00 | 0.00E+00 | 0.00E+00 | 0.00E+00 | 0.00E+00 | 0.00E+00 | 0.00E+00 | 0.00E+00 | Known_Known |
| g__CADDYX01 | 1.40E-04 | 3.58E-04 | 2.35E-05 | 0.00E+00 | 0.00E+00 | 0.00E+00 | 0.00E+00 | 0.00E+00 | 2.97E-05 | 0.00E+00 | 0.00E+00 | Known_Known |
| g__CADDYV01 | 0.00E+00 | 0.00E+00 | 0.00E+00 | 0.00E+00 | 4.09E-05 | 0.00E+00 | 0.00E+00 | 0.00E+00 | 0.00E+00 | 0.00E+00 | 0.00E+00 | Known_Known |
| g__Caccocola | 0.00E+00 | 0.00E+00 | 0.00E+00 | 0.00E+00 | 0.00E+00 | 0.00E+00 | 0.00E+00 | 0.00E+00 | 0.00E+00 | 5.46E-05 | 0.00E+00 | Known_Known |
| g__CABMCB01 | 0.00E+00 | 0.00E+00 | 0.00E+00 | 8.01E-05 | 0.00E+00 | 0.00E+00 | 0.00E+00 | 0.00E+00 | 0.00E+00 | 0.00E+00 | 0.00E+00 | Known_Known |
| g__C00003060 | 0.00E+00 | 0.00E+00 | 0.00E+00 | 0.00E+00 | 2.73E-05 | 0.00E+00 | 0.00E+00 | 0.00E+00 | 0.00E+00 | 0.00E+00 | 0.00E+00 | Known_Known |
| g__BRH-c8a | 0.00E+00 | 1.96E-03 | 1.03E-03 | 4.69E-04 | 7.91E-04 | 4.09E-04 | 0.00E+00 | 1.03E-04 | 0.00E+00 | 2.19E-04 | 4.38E-04 | Known_Known |
| g__Brevibacillus_D | 0.00E+00 | 0.00E+00 | 0.00E+00 | 0.00E+00 | 0.00E+00 | 0.00E+00 | 0.00E+00 | 2.58E-05 | 0.00E+00 | 0.00E+00 | 0.00E+00 | Known_Known |
| g__Brevefilum | 8.17E-04 | 1.40E-02 | 2.82E-04 | 1.83E-04 | 1.91E-04 | 2.34E-04 | 1.54E-03 | 1.13E-03 | 7.86E-04 | 4.19E-04 | 6.48E-04 | Known_Known |
| g__Brassicibacter_222685 | 2.17E-03 | 3.58E-04 | 8.69E-04 | 3.43E-04 | 1.02E-03 | 3.46E-03 | 1.59E-03 | 2.64E-03 | 7.83E-03 | 6.01E-04 | 3.70E-03 | Known_Known |
| g__Brasilonema | 0.00E+00 | 2.73E-04 | 0.00E+00 | 4.24E-04 | 3.41E-04 | 0.00E+00 | 0.00E+00 | 0.00E+00 | 0.00E+00 | 0.00E+00 | 0.00E+00 | Known_Known |
| g__Bosea | 0.00E+00 | 0.00E+00 | 0.00E+00 | 0.00E+00 | 0.00E+00 | 1.17E-05 | 0.00E+00 | 0.00E+00 | 0.00E+00 | 0.00E+00 | 0.00E+00 | Known_Known |
| g__Bog-159 | 0.00E+00 | 1.19E-04 | 0.00E+00 | 0.00E+00 | 5.46E-05 | 4.67E-05 | 0.00E+00 | 0.00E+00 | 0.00E+00 | 0.00E+00 | 0.00E+00 | Known_Known |
| g__BN140002 | 0.00E+00 | 1.02E-04 | 8.22E-05 | 0.00E+00 | 0.00E+00 | 9.34E-05 | 0.00E+00 | 0.00E+00 | 0.00E+00 | 0.00E+00 | 0.00E+00 | Known_Known |
| g__BMS3ABIN17 | 5.65E-03 | 3.41E-03 | 2.31E-03 | 6.91E-03 | 6.14E-04 | 7.12E-04 | 5.66E-03 | 1.22E-03 | 8.44E-03 | 4.30E-03 | 5.67E-03 | Known_Known |
| g__Blastococcus | 0.00E+00 | 5.11E-05 | 1.17E-05 | 0.00E+00 | 6.82E-05 | 4.67E-05 | 0.00E+00 | 0.00E+00 | 0.00E+00 | 5.46E-05 | 4.86E-05 | Known_Known |
| g__Blastochloris | 0.00E+00 | 0.00E+00 | 0.00E+00 | 0.00E+00 | 1.23E-04 | 0.00E+00 | 0.00E+00 | 0.00E+00 | 0.00E+00 | 0.00E+00 | 0.00E+00 | Known_Known |
| g__Blastocatella | 0.00E+00 | 0.00E+00 | 2.35E-05 | 0.00E+00 | 0.00E+00 | 0.00E+00 | 0.00E+00 | 0.00E+00 | 0.00E+00 | 0.00E+00 | 0.00E+00 | Known_Known |
| g__Bin90 | 0.00E+00 | 0.00E+00 | 0.00E+00 | 0.00E+00 | 2.73E-05 | 0.00E+00 | 0.00E+00 | 0.00E+00 | 0.00E+00 | 0.00E+00 | 3.24E-05 | Known_Known |
| g__Bin65 | 0.00E+00 | 0.00E+00 | 0.00E+00 | 0.00E+00 | 0.00E+00 | 0.00E+00 | 0.00E+00 | 0.00E+00 | 8.90E-05 | 0.00E+00 | 0.00E+00 | Known_Known |
| g__Bin5 | 0.00E+00 | 5.11E-05 | 0.00E+00 | 0.00E+00 | 0.00E+00 | 0.00E+00 | 0.00E+00 | 0.00E+00 | 1.04E-04 | 0.00E+00 | 0.00E+00 | Known_Known |
| g__Bin22 | 0.00E+00 | 8.52E-05 | 0.00E+00 | 0.00E+00 | 1.23E-04 | 7.01E-05 | 0.00E+00 | 0.00E+00 | 0.00E+00 | 0.00E+00 | 0.00E+00 | Known_Known |
| g__Bin103 | 0.00E+00 | 0.00E+00 | 0.00E+00 | 0.00E+00 | 0.00E+00 | 0.00E+00 | 0.00E+00 | 0.00E+00 | 0.00E+00 | 1.09E-04 | 0.00E+00 | Known_Known |
| g__Bilamarchaeum | 0.00E+00 | 0.00E+00 | 0.00E+00 | 0.00E+00 | 0.00E+00 | 0.00E+00 | 0.00E+00 | 0.00E+00 | 0.00E+00 | 0.00E+00 | 3.24E-05 | Known_Known |
| g__BH258 | 0.00E+00 | 0.00E+00 | 0.00E+00 | 0.00E+00 | 2.73E-05 | 2.34E-05 | 0.00E+00 | 0.00E+00 | 0.00E+00 | 0.00E+00 | 0.00E+00 | Known_Known |
| g__Bauldia | 0.00E+00 | 0.00E+00 | 0.00E+00 | 6.87E-05 | 0.00E+00 | 0.00E+00 | 0.00E+00 | 0.00E+00 | 0.00E+00 | 5.46E-05 | 0.00E+00 | Known_Known |
| g__Baia | 0.00E+00 | 0.00E+00 | 0.00E+00 | 0.00E+00 | 0.00E+00 | 1.17E-05 | 0.00E+00 | 0.00E+00 | 0.00E+00 | 0.00E+00 | 0.00E+00 | Known_Known |
| g__Bacillus_288330 | 0.00E+00 | 0.00E+00 | 1.17E-05 | 0.00E+00 | 0.00E+00 | 0.00E+00 | 0.00E+00 | 0.00E+00 | 0.00E+00 | 0.00E+00 | 0.00E+00 | Known_Known |
| g__BA2 | 2.19E-03 | 3.46E-03 | 4.23E-04 | 7.55E-04 | 1.79E-03 | 0.00E+00 | 3.73E-03 | 2.55E-03 | 2.40E-02 | 1.87E-02 | 9.95E-03 | Known_Known |
| g__B9-G15 | 0.00E+00 | 0.00E+00 | 0.00E+00 | 0.00E+00 | 0.00E+00 | 0.00E+00 | 0.00E+00 | 0.00E+00 | 1.63E-04 | 5.46E-05 | 0.00E+00 | Known_Known |
| g__B3-Chlor | 0.00E+00 | 0.00E+00 | 0.00E+00 | 0.00E+00 | 0.00E+00 | 0.00E+00 | 0.00E+00 | 0.00E+00 | 1.63E-04 | 4.55E-04 | 2.11E-04 | Known_Known |
| g__B1SED10-75 | 3.27E-04 | 2.32E-03 | 1.17E-03 | 6.75E-04 | 1.05E-03 | 8.06E-04 | 9.13E-04 | 6.05E-04 | 1.01E-03 | 1.06E-03 | 8.10E-04 | Known_Known |
| g__B10-G9 | 0.00E+00 | 2.21E-04 | 0.00E+00 | 1.49E-04 | 6.82E-05 | 0.00E+00 | 0.00E+00 | 1.42E-04 | 1.71E-03 | 1.02E-03 | 2.59E-04 | Known_Known |
| g__Azospirillum | 0.00E+00 | 3.75E-04 | 0.00E+00 | 0.00E+00 | 0.00E+00 | 0.00E+00 | 0.00E+00 | 0.00E+00 | 0.00E+00 | 0.00E+00 | 0.00E+00 | Known_Known |
| g__Atribacter | 0.00E+00 | 1.02E-04 | 0.00E+00 | 0.00E+00 | 0.00E+00 | 0.00E+00 | 0.00E+00 | 0.00E+00 | 8.90E-05 | 1.09E-04 | 0.00E+00 | Known_Known |
| g__Asprobacter | 0.00E+00 | 0.00E+00 | 0.00E+00 | 0.00E+00 | 0.00E+00 | 0.00E+00 | 0.00E+00 | 0.00E+00 | 5.93E-05 | 0.00E+00 | 0.00E+00 | Known_Known |
| g__Ardenticatena | 0.00E+00 | 0.00E+00 | 0.00E+00 | 0.00E+00 | 0.00E+00 | 0.00E+00 | 0.00E+00 | 3.86E-05 | 8.90E-05 | 9.29E-04 | 0.00E+00 | Known_Known |

| Genus | L1 | L2 | L3 | L4 | L5 | L6 | L7 | L8 | L9 | L10 | L11 | Microbial_matter |
|---|---|---|---|---|---|---|---|---|---|---|---|---|
| g__Arcobacter_476087 | 5.65E-03 | 1.22E-02 | 2.34E-02 | 1.82E-03 | 2.65E-03 | 0.00E+00 | 0.00E+00 | 2.58E-05 | 0.00E+00 | 5.01E-03 | 1.07E-03 | Known_Known |
| g__Aphodomorpha | 0.00E+00 | 0.00E+00 | 0.00E+00 | 0.00E+00 | 5.46E-05 | 0.00E+00 | 0.00E+00 | 0.00E+00 | 0.00E+00 | 0.00E+00 | 0.00E+00 | Known_Known |
| g__Anoxymicrobium | 9.33E-05 | 1.36E-04 | 7.04E-05 | 0.00E+00 | 6.82E-05 | 0.00E+00 | 0.00E+00 | 3.86E-05 | 0.00E+00 | 0.00E+00 | 8.10E-05 | Known_Known |
| g__Anoxybacter | 0.00E+00 | 1.70E-05 | 1.17E-05 | 0.00E+00 | 1.36E-05 | 0.00E+00 | 0.00E+00 | 0.00E+00 | 0.00E+00 | 0.00E+00 | 0.00E+00 | Known_Known |
| g__Andreesenia | 0.00E+00 | 0.00E+00 | 0.00E+00 | 0.00E+00 | 0.00E+00 | 0.00E+00 | 0.00E+00 | 0.00E+00 | 5.93E-05 | 0.00E+00 | 0.00E+00 | Known_Known |
| g__Anaerovorax | 0.00E+00 | 0.00E+00 | 0.00E+00 | 1.26E-04 | 0.00E+00 | 0.00E+00 | 0.00E+00 | 0.00E+00 | 0.00E+00 | 0.00E+00 | 0.00E+00 | Known_Known |
| g__Anaerosolibacter | 5.37E-04 | 3.20E-03 | 2.45E-03 | 7.44E-04 | 2.29E-03 | 2.25E-03 | 1.97E-04 | 2.19E-04 | 9.50E-04 | 2.00E-04 | 7.45E-04 | Known_Known |
| g__Anaerosacchariphilus | 0.00E+00 | 1.36E-04 | 0.00E+00 | 0.00E+00 | 0.00E+00 | 0.00E+00 | 0.00E+00 | 3.86E-05 | 1.93E-04 | 0.00E+00 | 1.78E-04 | Known_Known |
| g__Anaerophaga | 0.00E+00 | 0.00E+00 | 3.52E-05 | 0.00E+00 | 0.00E+00 | 0.00E+00 | 0.00E+00 | 0.00E+00 | 0.00E+00 | 0.00E+00 | 0.00E+00 | Known_Known |
| g__Anaerocolumna | 1.87E-04 | 0.00E+00 | 0.00E+00 | 0.00E+00 | 2.46E-04 | 3.74E-04 | 1.61E-04 | 9.01E-05 | 0.00E+00 | 0.00E+00 | 6.48E-05 | Known_Known |
| g__Anaerobium | 0.00E+00 | 0.00E+00 | 0.00E+00 | 0.00E+00 | 4.64E-04 | 4.09E-04 | 0.00E+00 | 1.67E-04 | 0.00E+00 | 0.00E+00 | 0.00E+00 | Known_Known |
| g__Anaerobacterium | 1.17E-03 | 8.86E-04 | 1.10E-03 | 4.46E-04 | 2.31E-03 | 3.97E-04 | 1.86E-03 | 2.28E-03 | 5.79E-04 | 3.70E-03 | 4.23E-03 | Known_Known |
| g__Anaerobacillus | 0.00E+00 | 0.00E+00 | 3.52E-05 | 0.00E+00 | 0.00E+00 | 0.00E+00 | 0.00E+00 | 0.00E+00 | 0.00E+00 | 2.73E-04 | 0.00E+00 | Known_Known |
| g__Aminipila | 0.00E+00 | 0.00E+00 | 0.00E+00 | 0.00E+00 | 0.00E+00 | 4.67E-05 | 0.00E+00 | 0.00E+00 | 0.00E+00 | 3.64E-05 | 0.00E+00 | Known_Known |
| g__Aminicenans | 4.67E-05 | 2.38E-04 | 0.00E+00 | 0.00E+00 | 0.00E+00 | 0.00E+00 | 0.00E+00 | 1.16E-04 | 2.23E-04 | 1.42E-03 | 1.13E-04 | Known_Known |
| g__Alteraurantiacibacter | 0.00E+00 | 0.00E+00 | 2.47E-04 | 0.00E+00 | 1.50E-04 | 0.00E+00 | 1.25E-04 | 0.00E+00 | 0.00E+00 | 0.00E+00 | 0.00E+00 | Known_Known |
| g__Allosphingosinicella | 0.00E+00 | 3.92E-04 | 2.93E-04 | 0.00E+00 | 2.87E-04 | 3.04E-04 | 0.00E+00 | 0.00E+00 | 0.00E+00 | 0.00E+00 | 0.00E+00 | Known_Known |
| g__Alkalispirochaeta | 0.00E+00 | 0.00E+00 | 0.00E+00 | 0.00E+00 | 0.00E+00 | 0.00E+00 | 0.00E+00 | 3.86E-05 | 0.00E+00 | 0.00E+00 | 0.00E+00 | Known_Known |
| g__Alkaliphilus_C | 3.50E-04 | 5.11E-05 | 1.53E-04 | 0.00E+00 | 5.87E-04 | 9.81E-04 | 1.97E-04 | 5.02E-04 | 1.08E-03 | 5.28E-04 | 2.11E-04 | Known_Known |
| g__Alkaliphilus_A | 0.00E+00 | 1.36E-04 | 0.00E+00 | 0.00E+00 | 2.18E-04 | 5.72E-04 | 0.00E+00 | 0.00E+00 | 0.00E+00 | 0.00E+00 | 0.00E+00 | Known_Known |
| g__Alkalibacterium | 0.00E+00 | 0.00E+00 | 0.00E+00 | 0.00E+00 | 0.00E+00 | 0.00E+00 | 0.00E+00 | 0.00E+00 | 0.00E+00 | 0.00E+00 | 1.46E-04 | Known_Known |
| g__Alkalibacter | 0.00E+00 | 1.53E-04 | 0.00E+00 | 0.00E+00 | 1.09E-04 | 8.17E-05 | 0.00E+00 | 0.00E+00 | 0.00E+00 | 0.00E+00 | 0.00E+00 | Known_Known |
| g__Alienimonas | 7.00E-05 | 0.00E+00 | 0.00E+00 | 0.00E+00 | 0.00E+00 | 0.00E+00 | 0.00E+00 | 0.00E+00 | 0.00E+00 | 0.00E+00 | 0.00E+00 | Known_Known |
| g__Aidingibacillus | 0.00E+00 | 0.00E+00 | 0.00E+00 | 0.00E+00 | 0.00E+00 | 1.17E-05 | 0.00E+00 | 0.00E+00 | 0.00E+00 | 0.00E+00 | 0.00E+00 | Known_Known |
| g__Aggregatilinea | 0.00E+00 | 3.41E-05 | 0.00E+00 | 6.87E-05 | 0.00E+00 | 0.00E+00 | 0.00E+00 | 0.00E+00 | 2.97E-05 | 0.00E+00 | 0.00E+00 | Known_Known |
| g__Afifella | 0.00E+00 | 0.00E+00 | 0.00E+00 | 0.00E+00 | 0.00E+00 | 1.75E-04 | 0.00E+00 | 0.00E+00 | 0.00E+00 | 0.00E+00 | 0.00E+00 | Known_Known |
| g__Aegiribacteria | 0.00E+00 | 0.00E+00 | 0.00E+00 | 0.00E+00 | 4.09E-05 | 0.00E+00 | 0.00E+00 | 0.00E+00 | 0.00E+00 | 0.00E+00 | 0.00E+00 | Known_Known |
| g__Acuticoccus | 6.53E-04 | 0.00E+00 | 5.28E-04 | 1.50E-03 | 2.73E-05 | 4.79E-04 | 1.63E-03 | 7.73E-04 | 2.82E-04 | 3.82E-04 | 3.57E-04 | Known_Known |
| g__Actinopolyspora | 0.00E+00 | 0.00E+00 | 0.00E+00 | 0.00E+00 | 4.09E-05 | 4.67E-05 | 0.00E+00 | 0.00E+00 | 0.00E+00 | 0.00E+00 | 0.00E+00 | Known_Known |
| g__Actinomycetospora | 0.00E+00 | 0.00E+00 | 0.00E+00 | 0.00E+00 | 4.09E-05 | 0.00E+00 | 0.00E+00 | 0.00E+00 | 0.00E+00 | 0.00E+00 | 0.00E+00 | Known_Known |
| g__Actinomarinicola | 7.70E-04 | 2.90E-04 | 3.05E-04 | 1.26E-03 | 3.27E-04 | 1.63E-04 | 4.84E-04 | 1.80E-04 | 0.00E+00 | 0.00E+00 | 7.78E-04 | Known_Known |
| g__Acetivibrio | 1.10E-03 | 1.11E-03 | 2.04E-03 | 1.45E-03 | 3.12E-03 | 2.53E-03 | 1.72E-03 | 3.76E-03 | 1.13E-03 | 2.91E-03 | 1.25E-03 | Known_Known |
| g__Abyssisolibacter | 0.00E+00 | 0.00E+00 | 0.00E+00 | 0.00E+00 | 0.00E+00 | 0.00E+00 | 1.79E-05 | 0.00E+00 | 0.00E+00 | 0.00E+00 | 0.00E+00 | Known_Known |
| g__A07HB70 | 0.00E+00 | 0.00E+00 | 1.29E-04 | 0.00E+00 | 0.00E+00 | 0.00E+00 | 0.00E+00 | 0.00E+00 | 0.00E+00 | 0.00E+00 | 0.00E+00 | Known_Known |
| g__49-20 | 1.87E-04 | 7.32E-04 | 0.00E+00 | 1.26E-04 | 1.77E-04 | 0.00E+00 | 4.84E-04 | 3.22E-04 | 0.00E+00 | 0.00E+00 | 6.48E-05 | Known_Known |
| g__46-80 | 0.00E+00 | 0.00E+00 | 0.00E+00 | 0.00E+00 | 0.00E+00 | 0.00E+00 | 0.00E+00 | 0.00E+00 | 0.00E+00 | 7.29E-05 | 0.00E+00 | Known_Known |
| g__4572-78 | 0.00E+00 | 0.00E+00 | 0.00E+00 | 6.87E-05 | 0.00E+00 | 0.00E+00 | 0.00E+00 | 0.00E+00 | 0.00E+00 | 0.00E+00 | 0.00E+00 | Known_Known |
| g__20-14-all-45-10 | 0.00E+00 | 0.00E+00 | 0.00E+00 | 0.00E+00 | 0.00E+00 | 0.00E+00 | 0.00E+00 | 0.00E+00 | 0.00E+00 | 4.74E-04 | 1.62E-04 | Known_Known |
| g__2-12-FULL-45-10 | 0.00E+00 | 0.00E+00 | 2.35E-05 | 0.00E+00 | 0.00E+00 | 0.00E+00 | 0.00E+00 | 0.00E+00 | 0.00E+00 | 0.00E+00 | 0.00E+00 | Known_Known |
| g__2-01-FULL-41-26 | 1.40E-04 | 0.00E+00 | 0.00E+00 | 0.00E+00 | 0.00E+00 | 0.00E+00 | 5.37E-05 | 0.00E+00 | 0.00E+00 | 0.00E+00 | 0.00E+00 | Known_Known |
| g__1-14-0-10-56-10 | 0.00E+00 | 0.00E+00 | 0.00E+00 | 0.00E+00 | 0.00E+00 | 0.00E+00 | 0.00E+00 | 0.00E+00 | 0.00E+00 | 0.00E+00 | 3.24E-05 | Known_Known |
| g__1-14-0-10-45-29 | 3.03E-04 | 0.00E+00 | 0.00E+00 | 0.00E+00 | 0.00E+00 | 0.00E+00 | 0.00E+00 | 0.00E+00 | 0.00E+00 | 0.00E+00 | 0.00E+00 | Known_Known |
| g__1-14-0-10-44-13 | 0.00E+00 | 0.00E+00 | 0.00E+00 | 2.29E-05 | 0.00E+00 | 0.00E+00 | 0.00E+00 | 0.00E+00 | 0.00E+00 | 0.00E+00 | 0.00E+00 | Known_Known |

**Supplementary Table 4. Alpha diversity indices across sediment layers**

| Layer | Observed | Shannon | invSimpson | Pielou |
|---:|---:|---:|---:|---:|
| L1 | 784 | 4.54 | 26.13 | 0.68 |
| L2 | 1429 | 5.68 | 85.34 | 0.78 |
| L3 | 1285 | 4.84 | 26.93 | 0.68 |
| L4 | 905 | 4.38 | 19.03 | 0.64 |
| L5 | 1273 | 4.55 | 22.09 | 0.64 |
| L6 | 904 | 4.27 | 19.93 | 0.63 |
| L7 | 630 | 3.91 | 13.69 | 0.61 |
| L8 | 965 | 4.48 | 24.67 | 0.65 |
| L9 | 1237 | 4.86 | 25.22 | 0.68 |
| L10 | 1361 | 5.23 | 54.92 | 0.72 |
| L11 | 1411 | 5.44 | 62.59 | 0.75 |

**Supplementary Table 5. Relative abundance of core genera**

| Genus | L1 | L2 | L3 | L4 | L5 | L6 | L7 | L8 | L9 | L10 | L11 | mean |
|---|---|---|---|---|---|---|---|---|---|---|---|---|
| g__SM1-50 | 27.57 | 15.35 | 13.79 | 27.92 | 8.13 | 18.42 | 38.26 | 48.11 | 39.04 | 28.79 | 23.73 | 26.28 |
| g__Heliorestis | 32.38 | 8.30 | 5.84 | 0.87 | 59.12 | 52.16 | 14.45 | 8.01 | 5.72 | 13.91 | 20.90 | 20.15 |
| g__PWUC01 | 0.45 | 1.89 | 2.86 | 6.88 | 3.67 | 2.99 | 7.41 | 13.36 | 21.01 | 14.00 | 8.59 | 7.56 |
| g__Halohasta | 8.94 | 8.67 | 17.26 | 5.48 | 0.44 | 1.96 | 1.01 | 0.25 | 0.45 | 0.82 | 7.11 | 4.76 |
| g__DTU012 | 1.53 | 1.59 | 1.21 | 0.23 | 8.24 | 2.57 | 2.71 | 1.41 | 1.54 | 0.60 | 1.65 | 2.12 |
| g__Saliphagus | 3.21 | 7.65 | 4.29 | 0.46 | 0.27 | 0.34 | 4.74 | 0.06 | 0.08 | 0.12 | 1.39 | 2.06 |
| g__Dethiobacter | 3.60 | 0.42 | 0.06 | 0.05 | 0.06 | 0.32 | 9.76 | 1.19 | 0.05 | 0.84 | 1.37 | 1.61 |
| g__Halorussus | 1.01 | 3.15 | 2.71 | 0.38 | 0.25 | 0.88 | 3.74 | 0.45 | 0.28 | 0.19 | 0.74 | 1.25 |
| g__Natranaerovirga | 0.16 | 0.29 | 0.07 | 0.32 | 0.13 | 0.44 | 0.70 | 0.50 | 0.91 | 7.52 | 1.52 | 1.14 |
| g__Sporosalibacterium | 0.55 | 0.98 | 0.58 | 0.34 | 0.12 | 0.16 | 0.13 | 1.32 | 1.38 | 3.62 | 2.35 | 1.05 |
| g__Rubribacterium | 0.48 | 0.37 | 0.67 | 6.67 | 0.28 | 0.16 | 0.37 | 1.00 | 0.01 | 0.35 | 0.53 | 0.99 |
| g__SKLG01 | 1.17 | 3.26 | 0.64 | 0.86 | 1.27 | 0.26 | 0.15 | 0.23 | 1.49 | 0.64 | 0.86 | 0.98 |
| g__HLUCCA09 | 0.92 | 0.23 | 0.27 | 1.00 | 0.05 | 0.08 | 0.77 | 6.30 | 0.23 | 0.02 | 0.26 | 0.92 |
| g__JAABTL01 | 0.54 | 1.78 | 0.04 | 1.43 | 0.89 | 0.02 | 0.08 | 1.22 | 0.77 | 0.25 | 0.36 | 0.67 |
| g__JAAAPF01 | 0.64 | 0.60 | 0.21 | 0.37 | 0.31 | 4.00 | 0.16 | 0.14 | 0.06 | 0.07 | 0.47 | 0.64 |
| g__Natronincola_A_235765 | 1.16 | 0.09 | 0.06 | 0.06 | 0.40 | 2.88 | 0.61 | 0.56 | 0.19 | 0.09 | 0.59 | 0.61 |
| g__BMS3ABIN17 | 0.79 | 0.57 | 0.35 | 1.07 | 0.10 | 0.10 | 0.86 | 0.21 | 1.12 | 0.65 | 0.83 | 0.60 |
| g__GWF2-37-15 | 0.90 | 0.11 | 0.02 | 0.14 | 0.16 | 0.04 | 0.82 | 2.14 | 0.11 | 0.21 | 1.94 | 0.60 |
| g__Ruminiclostridium | 0.12 | 0.22 | 0.07 | 0.13 | 2.41 | 0.52 | 0.22 | 0.82 | 0.26 | 0.38 | 0.88 | 0.55 |
| g__Halorubrum | 0.64 | 1.29 | 1.20 | 0.16 | 0.18 | 0.34 | 0.14 | 0.13 | 0.16 | 0.38 | 1.18 | 0.53 |
| g__Natrarchaeobaculum | 0.26 | 2.12 | 1.05 | 0.25 | 0.10 | 0.11 | 0.20 | 0.07 | 0.26 | 0.23 | 0.36 | 0.46 |
| g__Senegalia | 0.34 | 0.14 | 0.07 | 0.07 | 0.20 | 0.35 | 0.11 | 0.46 | 1.54 | 0.60 | 0.69 | 0.42 |
| g__Halostagnicola | 0.88 | 0.84 | 0.70 | 0.12 | 0.36 | 0.19 | 0.40 | 0.04 | 0.02 | 0.13 | 0.32 | 0.36 |
| g__CSSED11-154 | 0.78 | 1.35 | 0.16 | 0.20 | 0.10 | 0.12 | 0.36 | 0.38 | 0.18 | 0.04 | 0.33 | 0.36 |
| g__Halovenus | 0.07 | 1.08 | 1.38 | 0.28 | 0.06 | 0.14 | 0.51 | 0.03 | 0.05 | 0.09 | 0.25 | 0.36 |
| g__Defluviitalea_A | 0.20 | 0.01 | 0.12 | 0.16 | 0.88 | 0.87 | 0.41 | 0.53 | 0.15 | 0.16 | 0.32 | 0.35 |
| g__EX4484-6 | 0.15 | 0.18 | 0.06 | 0.17 | 0.23 | 0.17 | 0.31 | 0.87 | 0.53 | 0.64 | 0.50 | 0.35 |
| g__Brassicibacter_222685 | 0.30 | 0.06 | 0.13 | 0.05 | 0.16 | 0.47 | 0.24 | 0.46 | 1.04 | 0.09 | 0.54 | 0.32 |
| g__Haladaptatus | 0.19 | 1.33 | 0.29 | 0.09 | 0.10 | 0.24 | 0.16 | 0.28 | 0.19 | 0.25 | 0.35 | 0.32 |
| g__Acetivibrio | 0.15 | 0.18 | 0.31 | 0.23 | 0.49 | 0.34 | 0.26 | 0.66 | 0.15 | 0.44 | 0.18 | 0.31 |
| g__Brevefilum | 0.11 | 2.32 | 0.04 | 0.03 | 0.03 | 0.03 | 0.23 | 0.20 | 0.10 | 0.06 | 0.09 | 0.30 |
| g__CSSed10-48 | 0.83 | 0.62 | 0.05 | 0.04 | 0.03 | 0.04 | 0.02 | 0.18 | 0.12 | 0.81 | 0.39 | 0.29 |
| g__Natronoarchaeum | 0.22 | 1.31 | 0.76 | 0.15 | 0.03 | 0.03 | 0.17 | 0.01 | 0.07 | 0.16 | 0.15 | 0.28 |
| g__Anaerobacterium | 0.16 | 0.15 | 0.17 | 0.07 | 0.36 | 0.05 | 0.28 | 0.40 | 0.08 | 0.56 | 0.62 | 0.26 |
| g__SKVJ01 | 0.11 | 0.24 | 0.13 | 0.14 | 0.05 | 0.15 | 0.07 | 0.53 | 0.24 | 0.59 | 0.20 | 0.22 |
| g__Crassaminicella_234356 | 0.16 | 0.01 | 0.04 | 0.02 | 0.25 | 0.35 | 0.15 | 0.21 | 0.77 | 0.05 | 0.37 | 0.22 |
| g__Halorientalis | 0.07 | 0.83 | 0.49 | 0.08 | 0.06 | 0.09 | 0.43 | 0.02 | 0.01 | 0.03 | 0.09 | 0.20 |

| Genus | L1 | L2 | L3 | L4 | L5 | L6 | L7 | L8 | L9 | L10 | L11 | mean |
|---|---|---|---|---|---|---|---|---|---|---|---|---|
| g__Anaerosolibacter | 0.07 | 0.53 | 0.37 | 0.12 | 0.36 | 0.31 | 0.03 | 0.04 | 0.13 | 0.03 | 0.11 | 0.19 |
| g__B1SED10-75 | 0.05 | 0.38 | 0.18 | 0.10 | 0.17 | 0.11 | 0.14 | 0.11 | 0.13 | 0.16 | 0.12 | 0.15 |
| g__Halomarina_3560 | 0.03 | 0.38 | 0.77 | 0.05 | 0.01 | 0.01 | 0.03 | 0.03 | 0.12 | 0.05 | 0.12 | 0.15 |
| g__Halodesulfurarchaeum | 0.14 | 0.19 | 0.31 | 0.06 | 0.02 | 0.08 | 0.45 | 0.09 | 0.01 | 0.02 | 0.13 | 0.14 |
| g__W498 | 0.02 | 0.16 | 0.16 | 0.12 | 0.10 | 0.18 | 0.05 | 0.08 | 0.15 | 0.24 | 0.16 | 0.13 |
| g__Hyphomonas | 0.12 | 0.06 | 0.11 | 0.55 | 0.01 | 0.02 | 0.04 | 0.18 | 0.01 | 0.01 | 0.05 | 0.11 |
| g__Halocella_43036 | 0.01 | 0.01 | 0.01 | 0.00 | 0.11 | 0.03 | 0.07 | 0.04 | 0.05 | 0.13 | 0.05 | 0.05 |
| g__Nanohalobium | 0.02 | 0.10 | 0.03 | 0.04 | 0.01 | 0.03 | 0.05 | 0.00 | 0.00 | 0.01 | 0.03 | 0.03 |

### Supplementary Table 6. Relative abundance of unique genera per layer

| Genus | L1 | L2 | L3 | L4 | L5 | L6 | L7 | L8 | L9 | L10 | L11 |
|---|---|---|---|---|---|---|---|---|---|---|---|
| 1-14-0-10-45-29 | 0.04227505 | 0 | 0 | 0 | 0 | 0 | 0 | 0 | 0 | 0 | 0 |
| Alienimonas | 0.00975578 | 0 | 0 | 0 | 0 | 0 | 0 | 0 | 0 | 0 | 0 |
| Clostridium_AR_193524 | 0.00975578 | 0 | 0 | 0 | 0 | 0 | 0 | 0 | 0 | 0 | 0 |
| Desulforhopalus | 0.00650385 | 0 | 0 | 0 | 0 | 0 | 0 | 0 | 0 | 0 | 0 |
| Halobiforma | 0.00325193 | 0 | 0 | 0 | 0 | 0 | 0 | 0 | 0 | 0 | 0 |
| J110 | 0.00975578 | 0 | 0 | 0 | 0 | 0 | 0 | 0 | 0 | 0 | 0 |
| JAAYXY01 | 0.01625963 | 0 | 0 | 0 | 0 | 0 | 0 | 0 | 0 | 0 | 0 |
| O2-02-FULL-42-17 | 0.00975578 | 0 | 0 | 0 | 0 | 0 | 0 | 0 | 0 | 0 | 0 |
| REEP01 | 0.00975578 | 0 | 0 | 0 | 0 | 0 | 0 | 0 | 0 | 0 | 0 |
| Reyranella | 0.00975578 | 0 | 0 | 0 | 0 | 0 | 0 | 0 | 0 | 0 | 0 |
| SZUA-442 | 0.00325193 | 0 | 0 | 0 | 0 | 0 | 0 | 0 | 0 | 0 | 0 |
| Sphingoaurantiacus | 0.01300771 | 0 | 0 | 0 | 0 | 0 | 0 | 0 | 0 | 0 | 0 |
| Stappia | 0.02926734 | 0 | 0 | 0 | 0 | 0 | 0 | 0 | 0 | 0 | 0 |
| UBA2284 | 0.00975578 | 0 | 0 | 0 | 0 | 0 | 0 | 0 | 0 | 0 | 0 |
| UBA583 | 0.07479432 | 0 | 0 | 0 | 0 | 0 | 0 | 0 | 0 | 0 | 0 |
| Azospirillum | 0 | 0.06222951 | 0 | 0 | 0 | 0 | 0 | 0 | 0 | 0 | 0 |
| CSBR16-193 | 0 | 0.00848584 | 0 | 0 | 0 | 0 | 0 | 0 | 0 | 0 | 0 |
| DTJY01 | 0 | 0.02545753 | 0 | 0 | 0 | 0 | 0 | 0 | 0 | 0 | 0 |
| EX4484-52 | 0 | 0.01131446 | 0 | 0 | 0 | 0 | 0 | 0 | 0 | 0 | 0 |
| Elainella | 0 | 0.00848584 | 0 | 0 | 0 | 0 | 0 | 0 | 0 | 0 | 0 |
| Geminicoccus | 0 | 0.0198003 | 0 | 0 | 0 | 0 | 0 | 0 | 0 | 0 | 0 |
| Haliangium_437647 | 0 | 0.00282861 | 0 | 0 | 0 | 0 | 0 | 0 | 0 | 0 | 0 |
| Klenkia | 0 | 0.00565723 | 0 | 0 | 0 | 0 | 0 | 0 | 0 | 0 | 0 |
| Lutibaculum | 0 | 0.03394337 | 0 | 0 | 0 | 0 | 0 | 0 | 0 | 0 | 0 |
| MWBO01 | 0 | 0.00848584 | 0 | 0 | 0 | 0 | 0 | 0 | 0 | 0 | 0 |
| MnB-11 | 0 | 0.01131446 | 0 | 0 | 0 | 0 | 0 | 0 | 0 | 0 | 0 |
| Palleronia_492160 | 0 | 0.03394337 | 0 | 0 | 0 | 0 | 0 | 0 | 0 | 0 | 0 |
| UBA4427 | 0 | 0.00565723 | 0 | 0 | 0 | 0 | 0 | 0 | 0 | 0 | 0 |
| UBA876 | 0 | 0.05091506 | 0 | 0 | 0 | 0 | 0 | 0 | 0 | 0 | 0 |
| UBA877 | 0 | 0.05091506 | 0 | 0 | 0 | 0 | 0 | 0 | 0 | 0 | 0 |
| 2-12-FULL-45-10 | 0 | 0 | 0.00352367 | 0 | 0 | 0 | 0 | 0 | 0 | 0 | 0 |
| A07HB70 | 0 | 0 | 0.01938019 | 0 | 0 | 0 | 0 | 0 | 0 | 0 | 0 |
| Anaerophaga | 0 | 0 | 0.00528551 | 0 | 0 | 0 | 0 | 0 | 0 | 0 | 0 |
| Bacillus_288330 | 0 | 0 | 0.00176184 | 0 | 0 | 0 | 0 | 0 | 0 | 0 | 0 |
| Blastocatella | 0 | 0 | 0.00352367 | 0 | 0 | 0 | 0 | 0 | 0 | 0 | 0 |
| Caenispirillum | 0 | 0 | 0.01409468 | 0 | 0 | 0 | 0 | 0 | 0 | 0 | 0 |
| DSTP01 | 0 | 0 | 0.00528551 | 0 | 0 | 0 | 0 | 0 | 0 | 0 | 0 |
| Desulfocucumis | 0 | 0 | 0.02818936 | 0 | 0 | 0 | 0 | 0 | 0 | 0 | 0 |
| Legionella_B_628077 | 0 | 0 | 0.00352367 | 0 | 0 | 0 | 0 | 0 | 0 | 0 | 0 |
| Natranaerobaculum | 0 | 0 | 0.01057101 | 0 | 0 | 0 | 0 | 0 | 0 | 0 | 0 |
| Paenibacillus_D_363770 | 0 | 0 | 0.00176184 | 0 | 0 | 0 | 0 | 0 | 0 | 0 | 0 |
| Phenylobacterium | 0 | 0 | 0.01057101 | 0 | 0 | 0 | 0 | 0 | 0 | 0 | 0 |
| Pontibacter_909846 | 0 | 0 | 0.00352367 | 0 | 0 | 0 | 0 | 0 | 0 | 0 | 0 |
| QPJE01 | 0 | 0 | 0.00352367 | 0 | 0 | 0 | 0 | 0 | 0 | 0 | 0 |
| SR-FBR-E99 | 0 | 0 | 0.00352367 | 0 | 0 | 0 | 0 | 0 | 0 | 0 | 0 |
| Salinibaculum | 0 | 0 | 0.0352367 | 0 | 0 | 0 | 0 | 0 | 0 | 0 | 0 |
| Spiro-10 | 0 | 0 | 0.00528551 | 0 | 0 | 0 | 0 | 0 | 0 | 0 | 0 |
| UBA9910 | 0 | 0 | 0.00528551 | 0 | 0 | 0 | 0 | 0 | 0 | 0 | 0 |
| W-Firmicutes-14 | 0 | 0 | 0.01409468 | 0 | 0 | 0 | 0 | 0 | 0 | 0 | 0 |

| Genus | L1 | L2 | L3 | L4 | L5 | L6 | L7 | L8 | L9 | L10 | L11 |
|---|---|---|---|---|---|---|---|---|---|---|---|
| 1-14-0-10-44-13 | 0 | 0 | 0 | 0.00354497 | 0 | 0 | 0 | 0 | 0 | 0 | 0 |
| 4572-78 | 0 | 0 | 0 | 0.0106349 | 0 | 0 | 0 | 0 | 0 | 0 | 0 |
| Anaerovorax | 0 | 0 | 0 | 0.01949732 | 0 | 0 | 0 | 0 | 0 | 0 | 0 |
| CABMCB01 | 0 | 0 | 0 | 0.01240739 | 0 | 0 | 0 | 0 | 0 | 0 | 0 |
| Clostridium_AR_212534 | 0 | 0 | 0 | 0.00886242 | 0 | 0 | 0 | 0 | 0 | 0 | 0 |
| Dethiosulfatibacter | 0 | 0 | 0 | 0.09571413 | 0 | 0 | 0 | 0 | 0 | 0 | 0 |
| Ectothiorhodospira_607903 | 0 | 0 | 0 | 0.00177248 | 0 | 0 | 0 | 0 | 0 | 0 | 0 |
| Francisella_A | 0 | 0 | 0 | 0.00354497 | 0 | 0 | 0 | 0 | 0 | 0 | 0 |
| GWA2-35-9 | 0 | 0 | 0 | 0.00354497 | 0 | 0 | 0 | 0 | 0 | 0 | 0 |
| Haloimpatiens | 0 | 0 | 0 | 0.01772484 | 0 | 0 | 0 | 0 | 0 | 0 | 0 |
| J137 | 0 | 0 | 0 | 0.00708994 | 0 | 0 | 0 | 0 | 0 | 0 | 0 |
| Limnospira | 0 | 0 | 0 | 0.00177248 | 0 | 0 | 0 | 0 | 0 | 0 | 0 |
| Maricaulis | 0 | 0 | 0 | 0.01595236 | 0 | 0 | 0 | 0 | 0 | 0 | 0 |
| Nesciobacter | 0 | 0 | 0 | 0.03544968 | 0 | 0 | 0 | 0 | 0 | 0 | 0 |
| Polymorphobacter_A_486811 | 0 | 0 | 0 | 0.01240739 | 0 | 0 | 0 | 0 | 0 | 0 | 0 |
| RSA9 | 0 | 0 | 0 | 0.00531745 | 0 | 0 | 0 | 0 | 0 | 0 | 0 |
| SCUD01 | 0 | 0 | 0 | 0.00177248 | 0 | 0 | 0 | 0 | 0 | 0 | 0 |
| Sandaracinobacter | 0 | 0 | 0 | 0.01595236 | 0 | 0 | 0 | 0 | 0 | 0 | 0 |
| Sandarakinorhabdus | 0 | 0 | 0 | 0.00708994 | 0 | 0 | 0 | 0 | 0 | 0 | 0 |
| Stella | 0 | 0 | 0 | 0.02304229 | 0 | 0 | 0 | 0 | 0 | 0 | 0 |
| TMED2 | 0 | 0 | 0 | 0.01417987 | 0 | 0 | 0 | 0 | 0 | 0 | 0 |
| TMED27 | 0 | 0 | 0 | 0.01595236 | 0 | 0 | 0 | 0 | 0 | 0 | 0 |
| Thioflavicoccus | 0 | 0 | 0 | 0.00531745 | 0 | 0 | 0 | 0 | 0 | 0 | 0 |
| Tropicimonas_492701 | 0 | 0 | 0 | 0.02481478 | 0 | 0 | 0 | 0 | 0 | 0 | 0 |
| UBA10424 | 0 | 0 | 0 | 0.0992591 | 0 | 0 | 0 | 0 | 0 | 0 | 0 |
| Actinomycetospora | 0 | 0 | 0 | 0 | 0.00644358 | 0 | 0 | 0 | 0 | 0 | 0 |
| Aegiribacteria | 0 | 0 | 0 | 0 | 0.00644358 | 0 | 0 | 0 | 0 | 0 | 0 |
| Aphodomorpha | 0 | 0 | 0 | 0 | 0.00859143 | 0 | 0 | 0 | 0 | 0 | 0 |
| Blastochloris | 0 | 0 | 0 | 0 | 0.01933073 | 0 | 0 | 0 | 0 | 0 | 0 |
| C00003060 | 0 | 0 | 0 | 0 | 0.00429572 | 0 | 0 | 0 | 0 | 0 | 0 |
| CADDYV01 | 0 | 0 | 0 | 0 | 0.00644358 | 0 | 0 | 0 | 0 | 0 | 0 |
| Calidifontibacillus | 0 | 0 | 0 | 0 | 0.00644358 | 0 | 0 | 0 | 0 | 0 | 0 |
| Cellulomonas | 0 | 0 | 0 | 0 | 0.00214786 | 0 | 0 | 0 | 0 | 0 | 0 |
| Cereibacter_A_494128 | 0 | 0 | 0 | 0 | 0.01718287 | 0 | 0 | 0 | 0 | 0 | 0 |
| Clostridium_AF | 0 | 0 | 0 | 0 | 0.0257743 | 0 | 0 | 0 | 0 | 0 | 0 |
| Cohnella | 0 | 0 | 0 | 0 | 0.00214786 | 0 | 0 | 0 | 0 | 0 | 0 |
| Desulfitibacter | 0 | 0 | 0 | 0 | 0.00644358 | 0 | 0 | 0 | 0 | 0 | 0 |
| Filobacillus | 0 | 0 | 0 | 0 | 0.00214786 | 0 | 0 | 0 | 0 | 0 | 0 |
| Heliomicrobium | 0 | 0 | 0 | 0 | 0.01073929 | 0 | 0 | 0 | 0 | 0 | 0 |
| Herbivorax | 0 | 0 | 0 | 0 | 0.02147859 | 0 | 0 | 0 | 0 | 0 | 0 |
| Kocuria | 0 | 0 | 0 | 0 | 0.00644358 | 0 | 0 | 0 | 0 | 0 | 0 |
| LZORAL124-64-63 | 0 | 0 | 0 | 0 | 0.00644358 | 0 | 0 | 0 | 0 | 0 | 0 |
| MLS-D | 0 | 0 | 0 | 0 | 0.01073929 | 0 | 0 | 0 | 0 | 0 | 0 |
| Marispirochaeta | 0 | 0 | 0 | 0 | 0.00429572 | 0 | 0 | 0 | 0 | 0 | 0 |
| Papillibacter | 0 | 0 | 0 | 0 | 0.01288715 | 0 | 0 | 0 | 0 | 0 | 0 |
| Propylenella | 0 | 0 | 0 | 0 | 0.02792216 | 0 | 0 | 0 | 0 | 0 | 0 |
| Robertmurraya_298194 | 0 | 0 | 0 | 0 | 0.01718287 | 0 | 0 | 0 | 0 | 0 | 0 |
| SAR202-VII-2 | 0 | 0 | 0 | 0 | 0.00644358 | 0 | 0 | 0 | 0 | 0 | 0 |
| SDRK01 | 0 | 0 | 0 | 0 | 0.00429572 | 0 | 0 | 0 | 0 | 0 | 0 |
| Shivajiella | 0 | 0 | 0 | 0 | 0.00214786 | 0 | 0 | 0 | 0 | 0 | 0 |

| Genus | L1 | L2 | L3 | L4 | L5 | L6 | L7 | L8 | L9 | L10 | L11 |
|---|---|---|---|---|---|---|---|---|---|---|---|
| Syntrophobacter_452395 | 0 | 0 | 0 | 0 | 0.00429572 | 0 | 0 | 0 | 0 | 0 | 0 |
| UBA12014 | 0 | 0 | 0 | 0 | 0.00429572 | 0 | 0 | 0 | 0 | 0 | 0 |
| UBA5232 | 0 | 0 | 0 | 0 | 0.00429572 | 0 | 0 | 0 | 0 | 0 | 0 |
| VFBF01 | 0 | 0 | 0 | 0 | 0.01288715 | 0 | 0 | 0 | 0 | 0 | 0 |
| Afifella | 0 | 0 | 0 | 0 | 0 | 0.02375335 | 0 | 0 | 0 | 0 | 0 |
| Aidingibacillus | 0 | 0 | 0 | 0 | 0 | 0.00158356 | 0 | 0 | 0 | 0 | 0 |
| Baia | 0 | 0 | 0 | 0 | 0 | 0.00158356 | 0 | 0 | 0 | 0 | 0 |
| Bosea | 0 | 0 | 0 | 0 | 0 | 0.00158356 | 0 | 0 | 0 | 0 | 0 |
| Croceicoccus_484963 | 0 | 0 | 0 | 0 | 0 | 0.00950134 | 0 | 0 | 0 | 0 | 0 |
| Desulfonispora | 0 | 0 | 0 | 0 | 0 | 0.00475067 | 0 | 0 | 0 | 0 | 0 |
| Fictibacillus_C | 0 | 0 | 0 | 0 | 0 | 0.00950134 | 0 | 0 | 0 | 0 | 0 |
| Halapricum_3981 | 0 | 0 | 0 | 0 | 0 | 0.01108489 | 0 | 0 | 0 | 0 | 0 |
| Halogeometricum | 0 | 0 | 0 | 0 | 0 | 0.00950134 | 0 | 0 | 0 | 0 | 0 |
| JABFSM01 | 0 | 0 | 0 | 0 | 0 | 0.01108489 | 0 | 0 | 0 | 0 | 0 |
| Limivicinus | 0 | 0 | 0 | 0 | 0 | 0.00633423 | 0 | 0 | 0 | 0 | 0 |
| PCC-7407 | 0 | 0 | 0 | 0 | 0 | 0.00158356 | 0 | 0 | 0 | 0 | 0 |
| Paenibacillus_C | 0 | 0 | 0 | 0 | 0 | 0.00158356 | 0 | 0 | 0 | 0 | 0 |
| Pelagibacterium | 0 | 0 | 0 | 0 | 0 | 0.00475067 | 0 | 0 | 0 | 0 | 0 |
| Shimazuella | 0 | 0 | 0 | 0 | 0 | 0.00950134 | 0 | 0 | 0 | 0 | 0 |
| Sporacetigenium | 0 | 0 | 0 | 0 | 0 | 0.00950134 | 0 | 0 | 0 | 0 | 0 |
| UBA12474 | 0 | 0 | 0 | 0 | 0 | 0.00475067 | 0 | 0 | 0 | 0 | 0 |
| WRJI01 | 0 | 0 | 0 | 0 | 0 | 0.00158356 | 0 | 0 | 0 | 0 | 0 |
| Xylanibacillus | 0 | 0 | 0 | 0 | 0 | 0.00158356 | 0 | 0 | 0 | 0 | 0 |
| Abyssisolibacter | 0 | 0 | 0 | 0 | 0 | 0 | 0.00272985 | 0 | 0 | 0 | 0 |
| Caldicellulosiruptor | 0 | 0 | 0 | 0 | 0 | 0 | 0.01091941 | 0 | 0 | 0 | 0 |
| Cytobacillus_294142 | 0 | 0 | 0 | 0 | 0 | 0 | 0.00818956 | 0 | 0 | 0 | 0 |
| PUND01 | 0 | 0 | 0 | 0 | 0 | 0 | 0.00545971 | 0 | 0 | 0 | 0 |
| Alkalispirochaeta | 0 | 0 | 0 | 0 | 0 | 0 | 0 | 0.00675934 | 0 | 0 | 0 |
| Brevibacillus_D | 0 | 0 | 0 | 0 | 0 | 0 | 0 | 0.00450623 | 0 | 0 | 0 |
| CR-1 | 0 | 0 | 0 | 0 | 0 | 0 | 0 | 0.00675934 | 0 | 0 | 0 |
| DSDE01 | 0 | 0 | 0 | 0 | 0 | 0 | 0 | 0.00225311 | 0 | 0 | 0 |
| GWB1-41-6 | 0 | 0 | 0 | 0 | 0 | 0 | 0 | 0.00450623 | 0 | 0 | 0 |
| Halanaerobacter | 0 | 0 | 0 | 0 | 0 | 0 | 0 | 0.00675934 | 0 | 0 | 0 |
| Halapricum_3815 | 0 | 0 | 0 | 0 | 0 | 0 | 0 | 0.01126557 | 0 | 0 | 0 |
| J25 | 0 | 0 | 0 | 0 | 0 | 0 | 0 | 0.00675934 | 0 | 0 | 0 |
| Risungbinella | 0 | 0 | 0 | 0 | 0 | 0 | 0 | 0.00675934 | 0 | 0 | 0 |
| Tepidiforma | 0 | 0 | 0 | 0 | 0 | 0 | 0 | 0.00675934 | 0 | 0 | 0 |
| UBA2224 | 0 | 0 | 0 | 0 | 0 | 0 | 0 | 0.00901246 | 0 | 0 | 0 |
| UBA5619 | 0 | 0 | 0 | 0 | 0 | 0 | 0 | 0.00675934 | 0 | 0 | 0 |
| UBA9936 | 0 | 0 | 0 | 0 | 0 | 0 | 0 | 0.00675934 | 0 | 0 | 0 |
| Andreesenia | 0 | 0 | 0 | 0 | 0 | 0 | 0 | 0 | 0.00789173 | 0 | 0 |
| Asprobacter | 0 | 0 | 0 | 0 | 0 | 0 | 0 | 0 | 0.00789173 | 0 | 0 |
| Bin65 | 0 | 0 | 0 | 0 | 0 | 0 | 0 | 0 | 0.01183759 | 0 | 0 |
| CSP1-1 | 0 | 0 | 0 | 0 | 0 | 0 | 0 | 0 | 0.11640295 | 0 | 0 |
| Clostridiisalibacter | 0 | 0 | 0 | 0 | 0 | 0 | 0 | 0 | 0.00986466 | 0 | 0 |
| Clostridium_AD | 0 | 0 | 0 | 0 | 0 | 0 | 0 | 0 | 0.02367518 | 0 | 0 |
| DSYW01 | 0 | 0 | 0 | 0 | 0 | 0 | 0 | 0 | 0.01381052 | 0 | 0 |
| Desulfosarcina | 0 | 0 | 0 | 0 | 0 | 0 | 0 | 0 | 0.00591879 | 0 | 0 |
| Endothiovibrio | 0 | 0 | 0 | 0 | 0 | 0 | 0 | 0 | 0.00591879 | 0 | 0 |
| Evtepia | 0 | 0 | 0 | 0 | 0 | 0 | 0 | 0 | 0.01183759 | 0 | 0 |

| Genus | L1 | L2 | L3 | L4 | L5 | L6 | L7 | L8 | L9 | L10 | L11 |
|---|---|---|---|---|---|---|---|---|---|---|---|
| GCA-2401425 | 0 | 0 | 0 | 0 | 0 | 0 | 0 | 0 | 0.00789173 | 0 | 0 |
| GWC1-49-16 | 0 | 0 | 0 | 0 | 0 | 0 | 0 | 0 | 0.01775638 | 0 | 0 |
| HRBIN30 | 0 | 0 | 0 | 0 | 0 | 0 | 0 | 0 | 0.00591879 | 0 | 0 |
| JAAYNZ01 | 0 | 0 | 0 | 0 | 0 | 0 | 0 | 0 | 0.01381052 | 0 | 0 |
| JAAYUB01 | 0 | 0 | 0 | 0 | 0 | 0 | 0 | 0 | 0.00789173 | 0 | 0 |
| Methanospirillum | 0 | 0 | 0 | 0 | 0 | 0 | 0 | 0 | 0.00394586 | 0 | 0 |
| Orenia_42969 | 0 | 0 | 0 | 0 | 0 | 0 | 0 | 0 | 0.00394586 | 0 | 0 |
| Romboutsia_B | 0 | 0 | 0 | 0 | 0 | 0 | 0 | 0 | 0.01183759 | 0 | 0 |
| SG8-5 | 0 | 0 | 0 | 0 | 0 | 0 | 0 | 0 | 0.01578345 | 0 | 0 |
| Solincola_A_402975 | 0 | 0 | 0 | 0 | 0 | 0 | 0 | 0 | 0.00394586 | 0 | 0 |
| UBA2466 | 0 | 0 | 0 | 0 | 0 | 0 | 0 | 0 | 0.00591879 | 0 | 0 |
| UBA6092 | 0 | 0 | 0 | 0 | 0 | 0 | 0 | 0 | 0.00986466 | 0 | 0 |
| UBA8592 | 0 | 0 | 0 | 0 | 0 | 0 | 0 | 0 | 0.05524208 | 0 | 0 |
| UTCFX2 | 0 | 0 | 0 | 0 | 0 | 0 | 0 | 0 | 0.01972931 | 0 | 0 |
| Vermiphilus | 0 | 0 | 0 | 0 | 0 | 0 | 0 | 0 | 0.00591879 | 0 | 0 |
| WHTK01 | 0 | 0 | 0 | 0 | 0 | 0 | 0 | 0 | 0.00986466 | 0 | 0 |
| 46-80 | 0 | 0 | 0 | 0 | 0 | 0 | 0 | 0 | 0 | 0.01096311 | 0 |
| Bin103 | 0 | 0 | 0 | 0 | 0 | 0 | 0 | 0 | 0 | 0.01644466 | 0 |
| Caccocola | 0 | 0 | 0 | 0 | 0 | 0 | 0 | 0 | 0 | 0.00822233 | 0 |
| DQWO01 | 0 | 0 | 0 | 0 | 0 | 0 | 0 | 0 | 0 | 0.00822233 | 0 |
| DSAU01 | 0 | 0 | 0 | 0 | 0 | 0 | 0 | 0 | 0 | 0.00274078 | 0 |
| DTFT01 | 0 | 0 | 0 | 0 | 0 | 0 | 0 | 0 | 0 | 0.04659321 | 0 |
| Desulfohalotomaculum | 0 | 0 | 0 | 0 | 0 | 0 | 0 | 0 | 0 | 0.01370389 | 0 |
| Dwaynesavagella | 0 | 0 | 0 | 0 | 0 | 0 | 0 | 0 | 0 | 0.01370389 | 0 |
| GCA-2699025 | 0 | 0 | 0 | 0 | 0 | 0 | 0 | 0 | 0 | 0.00822233 | 0 |
| GWE2-41-16 | 0 | 0 | 0 | 0 | 0 | 0 | 0 | 0 | 0 | 0.00548155 | 0 |
| Gemmiger_A_73129 | 0 | 0 | 0 | 0 | 0 | 0 | 0 | 0 | 0 | 0.00822233 | 0 |
| Gottschalkia_A | 0 | 0 | 0 | 0 | 0 | 0 | 0 | 0 | 0 | 0.00548155 | 0 |
| HyVt-337 | 0 | 0 | 0 | 0 | 0 | 0 | 0 | 0 | 0 | 0.00548155 | 0 |
| Inquilinus | 0 | 0 | 0 | 0 | 0 | 0 | 0 | 0 | 0 | 0.03288933 | 0 |
| MZ-XQ | 0 | 0 | 0 | 0 | 0 | 0 | 0 | 0 | 0 | 0.00274078 | 0 |
| Marinilabilia | 0 | 0 | 0 | 0 | 0 | 0 | 0 | 0 | 0 | 0.00822233 | 0 |
| PALSA-647 | 0 | 0 | 0 | 0 | 0 | 0 | 0 | 0 | 0 | 0.00548155 | 0 |
| QALR01 | 0 | 0 | 0 | 0 | 0 | 0 | 0 | 0 | 0 | 0.00548155 | 0 |
| RBG-13-50-9 | 0 | 0 | 0 | 0 | 0 | 0 | 0 | 0 | 0 | 0.00822233 | 0 |
| SHVA01 | 0 | 0 | 0 | 0 | 0 | 0 | 0 | 0 | 0 | 0.00548155 | 0 |
| SHYG01 | 0 | 0 | 0 | 0 | 0 | 0 | 0 | 0 | 0 | 0.01644466 | 0 |
| SLQV01 | 0 | 0 | 0 | 0 | 0 | 0 | 0 | 0 | 0 | 0.00822233 | 0 |
| SYFI01 | 0 | 0 | 0 | 0 | 0 | 0 | 0 | 0 | 0 | 0.00274078 | 0 |
| SZUA-149 | 0 | 0 | 0 | 0 | 0 | 0 | 0 | 0 | 0 | 0.01096311 | 0 |
| Salirhabdus_286857 | 0 | 0 | 0 | 0 | 0 | 0 | 0 | 0 | 0 | 0.00822233 | 0 |
| Thalassobaculum_A_508031 | 0 | 0 | 0 | 0 | 0 | 0 | 0 | 0 | 0 | 0.03288933 | 0 |
| UBA2156 | 0 | 0 | 0 | 0 | 0 | 0 | 0 | 0 | 0 | 0.00822233 | 0 |
| UM-FILTER-41-35 | 0 | 0 | 0 | 0 | 0 | 0 | 0 | 0 | 0 | 0.00548155 | 0 |
| 1-14-0-10-56-10 | 0 | 0 | 0 | 0 | 0 | 0 | 0 | 0 | 0 | 0 | 0.00471965 |
| Alkalibacterium | 0 | 0 | 0 | 0 | 0 | 0 | 0 | 0 | 0 | 0 | 0.02123844 |
| Bilamarchaeum | 0 | 0 | 0 | 0 | 0 | 0 | 0 | 0 | 0 | 0 | 0.00471965 |
| CSBr16-57R1 | 0 | 0 | 0 | 0 | 0 | 0 | 0 | 0 | 0 | 0 | 0.00471965 |
| CSSed11-175R1 | 0 | 0 | 0 | 0 | 0 | 0 | 0 | 0 | 0 | 0 | 0.00707948 |
| DSXH01 | 0 | 0 | 0 | 0 | 0 | 0 | 0 | 0 | 0 | 0 | 0.00235983 |

| Genus | L1 | L2 | L3 | L4 | L5 | L6 | L7 | L8 | L9 | L10 | L11 |
|---|---|---|---|---|---|---|---|---|---|---|---|
| Desulfovibrio_R_446519 | 0 | 0 | 0 | 0 | 0 | 0 | 0 | 0 | 0 | 0 | 0.01179913 |
| Dietzia | 0 | 0 | 0 | 0 | 0 | 0 | 0 | 0 | 0 | 0 | 0.00235983 |
| Haloarchaeobius | 0 | 0 | 0 | 0 | 0 | 0 | 0 | 0 | 0 | 0 | 0.01651878 |
| Haloplanus | 0 | 0 | 0 | 0 | 0 | 0 | 0 | 0 | 0 | 0 | 0.01179913 |
| Lenti-02 | 0 | 0 | 0 | 0 | 0 | 0 | 0 | 0 | 0 | 0 | 0.00707948 |
| Niameybacter | 0 | 0 | 0 | 0 | 0 | 0 | 0 | 0 | 0 | 0 | 0.01887861 |
| Oceanidesulfovibrio_446513 | 0 | 0 | 0 | 0 | 0 | 0 | 0 | 0 | 0 | 0 | 0.00471965 |
| RBG-16-45-19 | 0 | 0 | 0 | 0 | 0 | 0 | 0 | 0 | 0 | 0 | 0.00471965 |
| SZUA-229 | 0 | 0 | 0 | 0 | 0 | 0 | 0 | 0 | 0 | 0 | 0.03067774 |
| Sutcliffiella_A | 0 | 0 | 0 | 0 | 0 | 0 | 0 | 0 | 0 | 0 | 0.00471965 |
| Tepidibaculum | 0 | 0 | 0 | 0 | 0 | 0 | 0 | 0 | 0 | 0 | 0.00707948 |
| Thiolinea | 0 | 0 | 0 | 0 | 0 | 0 | 0 | 0 | 0 | 0 | 0.00471965 |
| UBA1140 | 0 | 0 | 0 | 0 | 0 | 0 | 0 | 0 | 0 | 0 | 0.00943931 |
| UBA9971 | 0 | 0 | 0 | 0 | 0 | 0 | 0 | 0 | 0 | 0 | 0.00471965 |

**Supplementary Table 7. Spearman rank correlations applying the Benjamini–Hochberg False Discovery Rate (FDR) correction**

| Genus | Spearman_rho | p_value | p_value_FDR |
|---|---|---|---|
| DTU015 | 0.918182 | 6.66E-05 | 0.03957 |
| PWUC01 | 0.9 | 0.0001600 | 0.04751 |
| Natranaerovirga | 0.881818 | 0.0003302 | 0.04903 |

**Supplementary Table 8A. Marginal permutation tests for environmental variables in the Redundancy Analysis (RDA)**

*Evaluated via 9999 Monte Carlo permutations (Freedman-Lane conditional test; n = 11)*

**Model A (absolute nutrient concentration) — Pseudo-F(global) = 1.60, R2adj = 0.195, p(global) = 0.1207, VIF_max = 4.70**

| Variable | Parameter description | pseudo-F | p-value | FDR (BH) |
|---|---|---|---|---|
| pH | Sediment alkalinity (pH) | 2.364 | 0.085 | 0.170 |
| ln_TOC | Total organic carbon (ln, mg/g) | 1.376 | 0.261 | 0.307 |
| ln_TN | Total nitrogen (ln, mg/g) | 2.851 | 0.059 | 0.170 |
| ln_Po | Organic phosphorus (ln, mg/g) | 1.217 | 0.331 | 0.307 |

**Model B (elemental stoichiometry) — Pseudo-F(global) = 1.55, R2adj = 0.099, p(global) = 0.1555, VIF_max = 1.49**

| Variable | Parameter description | pseudo-F | p-value | FDR (BH) |
|---|---|---|---|---|
| ln_C_N | TOC:TN molar ratio (ln) | 1.828 | 0.112 | 0.212 |
| ln_N_Po | TN:Po molar ratio (ln) | 1.51 | 0.212 | 0.212 |

Note. Model A and Model B are non-nested (do not share all predictors) and are therefore compared by effect size (adjusted R2), not by relative significance of their global tests. FDR = Benjamini-Hochberg false discovery rate, applied within each model separately.

**Supplementary Table 8B. Centered Log-Ratio (CLR)-transformed abundances of the driving microbial genera across the eleven stratified sediment layers (L1–L11).**

| Layer | SM1-50 | *Heliorestis* | PWUC01 | *Halohasta* | *Sulfurimonas* | DTU012 | *Saliphagus* | *Dethiobacter* | *Halorussus* | *Natranaerovirga* |
|---|---|---|---|---|---|---|---|---|---|---|
| L1 | 2.0077054 | 2.1633051 | -0.9730883 | 0.9519152 | -1.2584742 | -0.4164325 | 0.0928108 | 0.1814045 | -0.6465171 | -1.1962318 |
| L2 | 1.4338910 | 0.8696775 | -0.2990804 | 0.9086914 | -0.5718795 | -0.4086790 | 0.7972224 | -1.0096800 | 0.0627714 | -1.1056947 |
| L3 | 1.2624757 | 0.4913122 | -0.0808084 | 1.4732373 | 1.0894415 | -0.6384830 | 0.2343427 | -1.3732067 | -0.1204437 | -1.3638169 |
| L4 | 2.2424664 | -0.4961286 | 0.9422609 | 0.7466535 | 1.7705245 | -0.9150529 | -0.7436306 | -1.0732769 | -0.7999835 | -0.8444353 |
| L5 | 1.1614837 | 3.0462605 | 0.4910770 | -0.6854389 | -0.6924076 | 1.1734598 | -0.8110651 | -0.9918131 | -0.8269385 | -0.9278644 |
| L6 | 1.8394133 | 2.8464160 | 0.2569010 | -0.0417009 | -1.1268902 | 0.1456754 | -0.8342206 | -0.8492584 | -0.4956184 | -0.7622471 |
| L7 | 2.1357897 | 1.2031925 | 0.5950050 | -0.8362817 | -1.5344165 | -0.2233846 | 0.2130428 | 0.8414191 | 0.0216207 | -1.0037882 |
| L8 | 2.7740281 | 1.078300443 | 1.5444119 | -0.8968911 | -1.1200346 | -0.2404079 | -1.0617657 | -0.3361331 | -0.7484711 | -0.7145695 |
| L9 | 2.6101511 | 0.8253603 | 2.0117690 | -0.7081643 | -1.0797279 | -0.1475638 | -1.0027668 | -1.0309377 | -0.8328678 | -0.4326246 |
| L10 | 2.0137281 | 1.3215875 | 1.3276055 | -0.7816082 | -0.6385073 | -0.9104410 | -1.2671160 | -0.7706791 | -1.2064913 | 0.7619717 |
| L11 | 1.7138160 | 1.5922856 | 0.7665198 | 0.5988968 | -1.1021590 | -0.5196414 | -0.6229077 | -0.6313111 | -0.9403160 | -0.5699422 |

Values represent log-encoded ratios relative to the geometric mean of each respective layer.

### Supplementary Table 9. Presence (1) or absence (0) of amplicon sequence variants (ASVs) assigned to phylum and genus across sediment core layers (L1–L11), including genus representative sequences (Ref_seq)

| | Presence (1) or Absence (0) in each layer | | | | | | | | | | | |
|---|---|---|---|---|---|---|---|---|---|---|---|---|
| Phylum_Genus_ASV_number | L1 | L2 | L3 | L4 | L5 | L6 | L7 | L8 | L9 | L10 | L11 | Sequence |
| p__Thermoplasmatota_g__SM1-50_ASV_1 | 1 | 1 | 1 | 1 | 1 | 1 | 1 | 1 | 1 | 1 | 1 | CCTTCACACTGTGCGAAAGCGCGATGAGGGGATCCCAAGTGCTTGCTCTTAGGAGTAAGCTGTTTTCCTGTTTAAATCGCAGGAAGAGGAAGGGCTGGGTAAGACGGGTGCCAGCCGCCGCGGTAATACCTGCAGCTCAAGTGGTGACCGCTATTATTGGGCCTAAAACGTCCGTAGCCGGCTGAGTAAATGCCTGGGTAAATCGTGTCGCTTAACGATACGAATTCCGGGTAGACTGCTTAGCTTGAGACTGGGAGAAGCTAGAGGTACTCCTGGGGTAGAGGTGAAATTCTGTAATCCTGGGGGACCACCAGTGGCGAAAGCGTCTAGCTAGAACGGGTCTGACGGTGAGGGACGAAACCCTGGGGCG |
| p__Thermoplasmatota_g__SM1-50_ASV_7 | 0 | 1 | 1 | 1 | 1 | 1 | 1 | 0 | 0 | 0 | 1 | CCTTCACACTGTGCGAAAGCGCGATGAGGGAATCCCAAGTGCTTGCTCTTAGGAGTAAGCTGTTTTCCTGTTTAAATCGCAGGAAGAGGAAGGGCTGGGTAAGACGGGTGCCAGCCGCCGCGGTAATACCTGCAGCTCAAGTGGTGACCGCTATTATTGGGCCTAAAACGTCCGTAGCCGGCTGAGTAAATGCCTGGGTAAATCGTGTCGCTTAACGATACGAATTCCGGGTAGACTGCTTAGCTTGAGACTGGGAGAAGCTAGAGGTACTCCTGGGGTAGAGGTGAAATTCTGTAATCCTGGGGGACCACCAGTGGCGAAAGCGTCTAGCTAGAACGGGTCTGACGGTGAGGGACGAAACCCTGGGGCG |
| p__Thermoplasmatota_g__SM1-50_ASV_8 | 1 | 0 | 0 | 0 | 0 | 0 | 1 | 1 | 1 | 1 | 1 | CCTTCACACTGTGCGAAAGCGCGATGAGGGGATCCCAAGTGCTTGCTCTTAGGAGTAAGCTGTTTTCCTGTTTAAATCGCAGGAAGAGGAAGGGCTGGGTAAGACGGGTGCCAGCCGCCGCGGTAATACCTGCAGCTCAAGTGGTGACCGCTATTATTGGGCCTAAAACGTCCGTAGCCGGCTGAGTAAATGCCTGGGTAAATCGTGTCGCTTAACGATACGAATTCCGGGTAGACTGCTTAGCTTGAGACCGGGAGAAGCTAGAGGTACTCCTGGGGTAGAGGTGAAATTCTGTAATCCTGGGGGACCACCAGTGGCGAAAGCGTCTAGCTAGAACGGGTCTGACGGTGAGGGACGAAACCCTGGGGCG |
| p__Halobacteriota_g__Halohasta_ASV_9 | 1 | 1 | 1 | 1 | 1 | 1 | 1 | 1 | 0 | 1 | 1 | CCTTTACACTGCACGCAAGTGCGATAGGGGAATCCCAAGTGCGAGGGCATACTAGCCCTCGCTTTTGTGTACTGTAAGGAGGTACACGAATAAGAGCTGGGCAAGACCGGTGCCAGCCGCCGCGGTAATACCGGCAGCTCGAGTGATGTCCACTATTATTGGGCCTAAAGCGTCCGTAGCTGGCCGCGCAAGTCCGTCGGGAAATCCACCTGCCCAACAGGTGGGCGTCCGGCGGAAACTGCGTGGCTTGGAACCGGAAGGCTCAGAGGGTACGTCCGGGGTAGGAGTGAAATCCCGTAATCCCGGACGGACCACCGATGGCGAAAGCACTCTGAGAGGACGGATTCGACAGTGAGGGACGAAAGCTAGGGTCT |
| p__Thermoplasmatota_g__SM1-50_ASV_13 | 1 | 1 | 1 | 1 | 1 | 0 | 0 | 0 | 0 | 0 | 1 | CCTTCACACTGTGCGAAAGCGCGATGAGGGGATTCCAAGTGCTAACTCGAAAGAGTTGGCTGTTTATCTGTCTAAAACGCAGGTAGAGTAAGAGCTGGGTAAGACGGCTGCCAGCCGCCGCGGTAATAGCTGCAGCTCGAGTGGTGACCACTTTTATTGGGCCTAAAACGTCCGTAGCCGGTCTAGTAAATGCCTGGGTAAATCGTGCCGCTTAACGGTACGAATTCCGGGTAGACTGCTAGACTTGAGATCGGGAGAGGTTAGAGGTACTCCTTGGGTAGAGGTGAAATTCTGTAATCCTAGGGGGACCACCTGTGGCGAAAGCGTCTAACCAGAACGAGTCTGACGGTGAGGGACGAAGCCCTGGGTCA |
| p__Thermoplasmatota_g__SM1-50_ASV_16 | 1 | 1 | 1 | 0 | 0 | 0 | 1 | 1 | 1 | 1 | 1 | CCTTCACACTGTGCGAAAGCGCGATGAGGGGATCCCAAGTGCTTGCACGTAAGTGTAAGCTGTTTCTTTGTTTAAAATGCATAGAGAGTAAGGGCTGGGTAAGACGGGTGCCAGCCGCCGCGGTAATACCTGCAGCTCAAGTGGTGACCACTATTATTGGGCCTAAAACGTCCGTAGCCTGTTTGATAAATGCCTGGGTAAATCGTATAGCTCAACTATACGAATTCCGGGTAGACTGTCAAGCTTGAGACCGGGAGAAGCTAGAGGTACTCTTGGGGTAGAGGTGAAATTCTGTAATCCTGAGGGGACCACCAGTGGCGAAGGCGTCTAGCTAGAACGGGTCTGACGGTGAGGGACGAAGCCCTGGGGCG |
| p__Halobacteriota_g__Saliphagus_ASV_19 | 1 | 1 | 1 | 1 | 0 | 1 | 1 | 1 | 0 | 0 | 1 | CCTTTACACTGCACGACAGTGCGATAAGGGGACTCCAAGTGCGAGGGCATATCGTCCTCGCTTTTCACTACCGTAAGGTGGTAGTGGAATAAGGGCTGGGCAAGACCGGTGCCAGCCGCCGCGGTAATACCGGCAGCCCGAGTGATGACCGCTATTATTGGGCCTAAAGCGTCCGTAGCCGGCCACGCAAGTCCGTCGGGAAATCCGCGCGCCTAACGCGCGGGCGTCCGGCGGAAACTGCGTGGCTTGGGACCGGAAGACTCGAGGGGTACGTCCGGGGTAGGAGTGAAATCCTGTAATCCTGGACGGACCACCGGTGGCGAAAGCGCCTCGAGAGGACGGATCCGACGGTGAGGGACGAAAGCTAGGGTCA |
| p__Thermoplasmatota_g__SM1-50_ASV_20 | 0 | 0 | 0 | 1 | 1 | 0 | 0 | 1 | 1 | 1 | 1 | ACTTCGCACTGCGCGAAAGCGCGACGAGGGAATCCCAAGTGTATGCACGTAAGTGTATACTGTTTTTCTGTCTAAAAAGCAGAGAGAGTAAGAGCTGGGTAAGACGGGTGCCAGCCGCCGCGGTAATACCTGCAGCTCAAGTGGTGATCGTTATTATTGGGCCTAAAACGTCCGTAGCCGGTCTTGTAAATGCCTGGGTAAATCGTATGGCTTAACCATACGAATTCCGGGTAGACTGTTAGACTTGAGACCGGGAGAGGCTAGAGGTACTCCTGGGGTAGAGGTGAAATTCTGTAATCCTGGGGGACCACCAGTGGCGAAGGCGTCTAGCCAGAACGGGTCTGACGGTGAGGGACGAAGCCCTGGGGCG |
| p__Thermoplasmatota_g__SM1-50_ASV_24 | 0 | 0 | 0 | 0 | 0 | 0 | 0 | 1 | 1 | 1 | 1 | CCTTCACACTGTGCGAAAGCGCGATGAGGGGATCCCAAGTGCTTGCTCTTAGGAGTAAGCTGTTTTCCTGTTTAAATCGCAGGAAGAGGAAGGGCTGGGTAAGACGGGTGCCAGCCGCCGCGGTAATACCTGCAGCTCAAGTGGTGACCGCTATTATTGGGCCTAAAACGTCCGTAGCCGGCTGAGTAAATGCCTGGGTAAATCGTGTCGCTTAACGATACGAATTCCGGGTAGACTGCTTGGCTTGAGACCGGGAGAAGCTAGAGGTACTCCTGGGGTAGAGGTGAAATTCTGTAATCCTGGGGGACCACCAGTGGCGAAGGCGTCTAGCTAGAACGGGTCTGACGGTGAGGGACGAAACCCTGGGGCG |
| p__Halobacteriota_g__Halohasta_ASV_27 | 1 | 1 | 1 | 1 | 1 | 1 | 1 | 1 | 1 | 1 | 1 | CCTTTACACTGCACGCAAGTGCGATAGGGGAATCCCAAGTGCGAGGGCATACTAGCCCTCGCTTTTGTGTACTGTAAGGAGGTACACGAATAAGAGCTGGGCAAGACCGGTGCCAGCCGCCGCGGTAATACCGGCAGCTCGAGTGATGTCCACTATTATTGGGCCTAAAGCGTCCGTAGCTGGCCAGCCAAGTTCGTCGGGAAATCCACCTGCCCAACAGGTGGGCGTCCGGCGAAAACTGTCTGGCTTGGAACCGGAAGGCTCAGAGGGTACGTCCGGGGTAGGAGTGAAATCCTGTAATCCCGGACGGACCACCGATGGCGAAAGCACTCTGAGAGGACGGATTCGACAGTGAGGGACGAAAGCTAGGGTCT |
| p__Thermoplasmatota_g__SM1-50_ASV_31 | 1 | 1 | 1 | 1 | 0 | 1 | 1 | 1 | 1 | 1 | 1 | CCTTCACACTGTGCGAAAGCGCGATGAGGGGATCCCAAGTGCTTGCACGTAAGTGTAAGCTGTTTCTTTGTTTAAAACGCAGAGAGAGGAAGGGCTGGGTAAGACGGGTGCCAGCCGCCGCGGTAATACCTGCAGCTCGAGTGGTGACCGCTATTATTGGGCCTAAAACGTCCGTAGCCGGCTGAGTAAATGCTTGGGTAAATCGTGCCGCTTAACGGTACGAATTCCGGGTAAACTGCTCAGCTTGAGATCGGGAGAAGCTAGAGGTACTCCTGGGGTAGAGGTGAAATTCTGTAATCCTGGGGGACCACCAGTGGCGAAGGCGTCTAGCTAGAACGAGTCTGACGGTGAGGGACGAAACCCTGGGGCG |
| p__Thermoplasmatota_g__SM1-50_ASV_32 | 0 | 0 | 0 | 0 | 0 | 0 | 0 | 1 | 1 | 1 | 1 | CCTTCACAATGTGCGAAAGCACGATGAGGGGATCCCAAGTGCTTGCACGTAAGTGTAAGCTGTTTCTTTGTCTAAAACGCATAGAGAGTAAGGGCTGGGTAAGACGGGTGCCAGCCGCCGCGGTAATACCTGCAGCTCAAGTGGTGGCCACTATTATTGGGCCTAAAACGTTCGTAGCCGGCTTGATAAATGCCTGGGTAAATCGTATAGCTCAACTATACGAATTCCGGGTAGACTGTTAAGCTTGGGACCGGGAGAGGCTAGAGGTACTCCTGGGGTAGAGGTGAAATTCTGTAATCCTGGGGGACCACCAGTGGCGAAGGCGTCTAGCTAGAACGGGTCCGACGGTGAGGGACGAAGCCCTGGGGCG |
| p__Thermoproteota_g__GCA-2726865_ASV_35 | 0 | 1 | 0 | 0 | 0 | 0 | 0 | 1 | 1 | 1 | 1 | CCTCCTCAATGCACTAAAGTGTGAAGGGGTTACCTCGAGTGCTACCTGAAGAAGGTGGCTTTTCCTTGGTCTAAAACGCCTTGGGAATAAGAGGGGGGCAAGACTGGTGTCAGCCGCCGCGGTAACACCAGCCCCTCGAGTGGTGGGGATGATTATTTGGCCTAAAGCGTCCGTAGCCGGCCAGGTAAGTTTTCCGTCAAATCCAACGTCTCAAGCGTTGGGCTGCGGGAAATACTGCCAGGCTAGAGAGTGGGAGAGGTAAGCGGTATTCTGGGGTAGGGGCGAAATCCTTTGATCCCCGGAGGACCACCAGTGGCGAAGGCTGCTTACTAGAACACGTCTGACGGTCAGGGACGAAAGCTGGGGTAG |
| p__Halobacteriota_g__Halorussus_ASV_36 | 1 | 1 | 1 | 1 | 1 | 1 | 1 | 1 | 1 | 0 | 1 | ACTTTACACTGCACGACAGTGCGATAAGGGAACTCCGAGTGCGAGGGCATATAGTCCTCGCTTTTCTGCACCGTAAGGTGGTGCACGAATAAGAGCTGGGCAAGACCGGTGCCAGCCGCCGCGGTAATACCGGCAGCTCAAGTGATGGCCGCTATTATTGGGCCTAAAGCGTCCGTAGCCGGCCAGACAGGTCCGTCGGGAAATCCACGCGCTCAACGCGTGGGCGTCCGGCGGAAACCAGCTGGCTTGGGGCCGGAAGATCTGAGGGGTACGTCCGGGGTAGGAGTGAAATCCCGTAATCCTGGACGGACCACCGGTGGCGAAAGCGCCTCAGAAAAACGGACCCGACGGTGAGGGACGAAAGCTAGGGTCA |
| p__Thermoplasmatota_g__SM1-50_ASV_43 | 1 | 1 | 1 | 1 | 1 | 0 | 0 | 1 | 1 | 1 | 1 | CCTTCACAATGTGCGAAAGCGCGATGAGGGGATCCCAAGTGCTTGCACGTAAGTGTAAGCTGTTTCTTTGTTTAAAAAGCAGAGAGAGTAAGGGCTGGGTAAGACGGGTGCCAGCCGCCGCGGTAATACCTGCAGCTCAAGTGGTGGCCGCTATTATTGGGCCTAAAACGTCCGTAGCCTGCTGGGTAAATGCCTGGGTAAATCGTGTCGCTTAACGATACGAATTCCGGGTAGACTGCCTAGCTTGAGACCGGAAGAAGTTAGAGGTACTCCTGGGGTAGGGGTGAAATCCTGTAATCCTGGGGGACCACCAGTGGCGAAGGCGTCTAACTAGGACGGGTCTGACGGTGAGGGACGAAACCCTGGGGCG |
| p__Halobacteriota_f__Natrialbaceae_ASV_45 | 1 | 1 | 1 | 1 | 0 | 1 | 1 | 1 | 0 | 0 | 1 | CCTTTACACTGCACGCCAGTGCGATAAGGGGACTCCAAGTGCGAGGGCATACAGTCCTCGCTTTTGTGTACCGTAAGGTGGTACACGAATAAGGGCTGGGCAAGACCGGTGCCAGCCGCCGCGGTAATACCGGCAGCCCAAGTGATGACCGCTATTATTGGGCCTAAAGCGTCCGTAGCCGGCCGGGTAAGTCCGTCGGGAAATCCGCGCGCTCAACGCGCGGGCGTCCGGTGGAAACTGCCTGGCTTGGGACCGGAAGATCCAGAGGGTACGTCCGGGGTAGGAGTGAAATCCTGTAATCCTGGACGGACCACCGGTGGCGAAAGCGCTCTGGAAGGACGGATCCGACGGTGAGGGACGAAAGCTAGGGTCA |
| p__Halobacteriota_g__Natrinema_ASV_47 | 1 | 1 | 1 | 0 | 0 | 0 | 1 | 0 | 0 | 0 | 1 | CCTTTACACTGCACGCCAGTGCGATAGGGGGACTCCAAGTGCGAGGGCATACAGTCCTCGCTTTTCACGACCGTAAGGTGGTCGTGGAATAAGAGCTGGGCAAGACCGGTGCCAGCCGCCGCGGTAATACCGGCAGCTCGAGTGATGACCGCTTTTATTGGGCCTAAAGCGTCCGTAGCCGGCCGGGTAAGTCCATCGGGAAATCCGCGCGCTCAACGCGCGGGCGTCCGGTGGAAACTGCCTGGCTTGGGACCGGAAGATCCAGAGGGTACGTCTGGGGTAGGAGTGAAATCCCGTAATCCTGGACGGACCACCGGTGGCGAAAGCGCTCTGGAAAGACGGATCCGACGGTGAGGGACGAAAGCTCGGGTCA |
| p__Thermoplasmatota_g__SM1-50_ASV_48 | 1 | 1 | 1 | 0 | 0 | 0 | 0 | 0 | 0 | 0 | 1 | CCTTCACACTGTGCGAAAGCGCGATGAGGGGATCCCAAGTGCTTGCACGTAAGTGTAAGCTGTTTCTTTGTCTAAAACGCATAGAGAGTAAGGGCTGGGTAAGACGGGTGCCAGCCGCCGCGGTAATACCTGCAGCCCAAGTGGTGACCGCTTTTATTGGGCCTAAAACGTCCGTAGCCGGTCTAGTAAATGCCTGGGTAAATCGTGCCGCTTAACGGTACGAATTCCGGGTAGACTGCCAGACTTGAGACTGGGAGAGGCTAGAGGTACTCCTGGGGTAGAGGTGAAATTCTGTAATCCTGGGGGACGACCAGTGGCGAAAGCGTCTAGCTAGAACGGGTCTGACGGTGAGGGACGAAGCCCTGGGTAG |
| p__Halobacteriota_g__Halohasta_ASV_50 | 1 | 1 | 1 | 1 | 1 | 1 | 1 | 0 | 0 | 1 | 1 | CCTTTACACTGCACGCAAGTGCGATAGGGGAATCCCAAGTGCGAGGGCATACTAGCCCTCGCTTTTGTGTACTGTAAGGAGGTACACGAATAAGAGCTGGGCAAGACCGGTGCCAGCCGCCGCGGTAATACCGGCAGCTCGAGTGATGTCCAATATTATTGGGCCTAAAGCGTCCGTAGCTGGCCAGCCAAGTCCGTCGGGAAATCCACCTGCCCAACAGGTGGGCGTCCGGCGGAAACTGTCTGGCTTGGAACCGGAAGGCTCAGAGGGTACGTCCGGGGTAGGAGTGAAATCCCGTAATCCCGGACGGACCACCGATGGCGAAAGCACTCTGAGAAGACGGATTCGACAGTGAGGGACGAAAGCTAGGGTCT |
| p__Thermoplasmatota_g__SM1-50_ASV_52 | 1 | 0 | 0 | 1 | 0 | 0 | 0 | 1 | 1 | 1 | 1 | ACTTCACAATGCGCGAAAGCGCGATGAGGGGATCCCAAGTGCTTGCACGTAAGTGTAAGCTGTTTTTCTGTTTAAAAAGCAGAGAGAGTAAGGGCTGGGTAAGACGGGTGCCAGCCGCCGCGGTAATACCTGCAGCTCAAGTGGTGACCGCTATTATTGGGCCTAAAACGTCCGTAGCCGGCTAAGTAAATGCCTGGGTAAATCGTGCCGCTTAACGGTACGAATTCCGGGTAGACTGCTTAGCTTGGGACCGGAAGAAGTTAGAGGTACTCCTGGGGTAGGGGTGAAATCCTGTAATCCTGGGGGACCACCAGTGGCGAAGGCGTCTAACTAGGACGGGTCTGACGGTGAGGGACGAAGCCCTGGGGCG |
| p__Thermoplasmatota_g__SM1-50_ASV_62 | 0 | 0 | 0 | 0 | 0 | 0 | 0 | 1 | 1 | 1 | 1 | ACTTCGCACTGTGCGAAAGCACGACGAGGGAATCCCAAGTGCTGGCTCGTTAGAGTCAGCTGTTTCTTTGTCTAAAAAACAGAGAGAGTAAGGGCTGGGTAAGACGGGTGCCAGCCGCCGCGGTAATACCTGCAGCCCGAGTGGTGACTGTTATTATTGGGCCTAAAACGTCCGTAGCCGGTTCTGTAAATGCCTGGGTAAATCGTACGGCTCAACCGTACGAATTCCGGGTAGACTGCAGGGCTAGAGATCGGGAGAGGCTAGAGGTACTCTTGGGGTAGAGGTGAAATTCTGTAATCCTAGGGGGACCACCAGTGGCGAAGGCGTCTAGCTAGAACGAGTCTGACGGTCAGGGACGAAGCCCTGGGGCG |
| p__Thermoplasmatota_g__SM1-50_ASV_73 | 0 | 0 | 0 | 0 | 0 | 0 | 0 | 1 | 1 | 1 | 1 | CCTTCACACTGTGCGAAAGCGCGATGAGGGAATCCCAAGTGCTTGCTCGTAAGAGTAAGCTGTTTCTGTGTATAAAAAGCATAGAGAGTAAGGGCTGGGTAAGACGGGTGCCAGCCGCCGCGGTAATACCTGCAGCTCTAGTGGTGACCACTATTATTGGGCCTAAAACGTTCGTAGCCGGTTTGCTAAATGCCTGGGTAAATCGAGCCGCTTAACGGTTCGAATTCCGGGTAGACTGGCAAACTTGAGACCGGGAGAGGCTAGAGGTACTCCTGGGGTAGAGGTGAAATTCTGTAATCCTGGGGGACCACCAGTGGCGAAGGCGTCTAGCTAGAACGGGTCTGACGGTGAGGGACGAAGCCCTGGGGCG |
| p__Thermoplasmatota_g__FEN-33_ASV_77 | 0 | 1 | 0 | 0 | 0 | 0 | 0 | 1 | 1 | 1 | 1 | ACTTCGCAATGGGGGAAACCCTGACGAGGGGATTCCAAGTGCCAGCACACAGTGTTGGCTGTTCTTCTGTCCAAAACGCAGGAGAAGTAAGGGCCGGGTAAGACGGGTGCCAGCCGCCGCGGTAATACCCGCGGCCCGAGTGGTGGTCGATTTTATTGAGCCTAAAACGTCCGTAGCCTGCTTGGTAAATCCTTGGGTAAATCGGCCAGCTTAACTGTCCGAAGTCCGGGGAGACTGCCAGGCTTGGGACCGGGAGAGGTCAGAGGTACTTCTGGGGTAGGGGTAAAATCCTGTAATCCTAGAAGGACCACCGGTGGCGAAGGCGTCTGACTAGAACGGATTCGACGGTGAGGGACGAAGCCCTGGGGCG |
| p__Halobacteriota_g__Natrarchaeobaculum_ASV_79 | 1 | 1 | 1 | 1 | 1 | 1 | 1 | 1 | 1 | 1 | 1 | CCTTTACACTGCACGCGAGTGCGATAAGGGGACTCCAAGTGCCAGGGCATATCGTCCTGGCTTTTCACCACCGTAGGGAGGTGGTGGAATAAGGGCTGGGCAAGACCGGTGCCAGCCGCCGCGGTAATACCGGCAGCCCAAGTGATGACCGCTCTTATTGGGCCTAAAGCGTCCGTAGCTGGCCTCGCAAGTCCATCGGGAAATCCGCACGCCTAACGTGCGGGCGTCCGGTGGAAACTGCGTGGCTTGGGACCGGAAGACCCGAGGGGTACGTCCGGGGTAGGAGTGAAATCCTGTAATCCTGGACGGACCACCGGTGGCGAAAGCGCCTCGGGAGGACGGATCCGACGGTGAGGGACGAAAGCTAGGGTCA |
| p__Thermoproteota_g__BA2_ASV_82 | 1 | 1 | 0 | 1 | 1 | 0 | 1 | 1 | 1 | 1 | 1 | ACTCCGCAATGCACGAAAGTGTGACGGGGCTACCCCGAGTGCCATCCGCTGAGGATGGCTTTTCTTTGGTATAACTAGCCATGGGAATAAGGAGAGGGCAAGACTGGTGTCAGCCGCCGCGGTAATACCAGCTCTCCGAGTGGTAGGGATGATTATTGGGCTTAAAGCGTCCGTAGCCGGCTTAGCAAGTTTCCTGTTAAACCCAGCGACTCAATCGTTGGATCGCGGGAAATACTACTAGGCTAGGAGGCGGGAGAGGTCAACGGTACTTCTGGGGTAGGGGTGAAATCCTATAATCCCGGGAGGACCACCAGTGGCGAAGGCTGTTGACTAGAACGCGCTCGACGGTGAGGGACGAAAGCTGGGGGAG |

| Phylum_Genus_ASV_number | Presence (1) or Absence (0) in each layer L1 | L2 | L3 | L4 | L5 | L6 | L7 | L8 | L9 | L10 | L11 | Sequence |
|---|---|---|---|---|---|---|---|---|---|---|---|---|
| p__Thermoplasmatota_g__SM1-50_ASV_85 | 1 | 0 | 0 | 0 | 1 | 1 | 1 | 1 | 1 | 1 | 1 | CCTTCACACTGTGCGAAAGCGCGATGAGGGGATCCCAAGTGCTAACTCGAAAGAGTTGGCTGTTTATCTGTCTAAAACGCAGGTAGAGTAAGAGCTGGGTAAGACGGCTGCCAGCCGCCGCGGTAATAGCTGCAGCTCGAGTGGTGACCACTATTATTGGGCCTAAAACGTCCGTAGCCGGCCTAGTAAATGCCTGGGTAAATCGTGCCGCTTAACGGTACGAATTCCGGGTAGACTGCTAGACTTGAGATCGGGAGAGGTTAGAGGTACTCCTTGGGTAGAGGTGAAATTCTGTAATCCTAGGGGGACCACCTGTGGCGAAAGCGTCTAACCAGAACGAGTCTGACGGTGAGGGACGAAGCCCTGGGTCA |
| p__Thermoproteota_g__BA2_ASV_90 | 1 | 1 | 1 | 0 | 0 | 0 | 0 | 1 | 1 | 1 | 1 | ACTCCGCAATGCACGAAAGTGTGACGGGGCTACCCCGAGTGCCGTCCGCTGAGGATGGCTTTTCCCCGGTGTAACTAGCCGGGGGAATAAGGAGAGGGCAAGACTGGTGTCAGCCGCCGCGGTAATACCAGCTCTCCGAGTTGTAGGGATGATTATTGGGCTTAAAGCGTCCGTAGCCGGCTTAGCAAGTCTCCTGTTAAATTCAGCGACCTAATCGTTGAGCCGCGGGAGATACTACTAGGCTAGGAGGCGGGAGAGGTCGACGGTACTTTCGGGGTAGGGGTGAAATCCTATAATCCCGGGAGGACCACCAGTGGCGAAGGCTGTCGACTAGAACGCGCTCGACGGTGAGGGACGAAAGCTGGGGGAG |
| p__Thermoplasmatota_g__EX4484-6_ASV_93 | 0 | 0 | 1 | 1 | 1 | 1 | 1 | 1 | 1 | 1 | 1 | CCTTCGCAATGCGCGAAAGCGCGACGAGGGGATTCCAAGTGCATGCACTTAGTGTATGCTGTTTGCTAGTCTAAATCGCTAGCAGAGTAAGGGCTGGGTAAGACTGGTGCCAGCCGCCGCGGTAATACCAGCGGCCCTAGTGGTGACCGTTATTATTGGGCCTAAAGCGTTCGTAGCCGGTCCGGTAAATCTCTGGGTAAATCGATGGGCTTAACCCATCGAATTCTGGGGAGACTGCTGGACTTGAGACCGGGAGAGGTCGGAGGTACTACGGGGGTAGAGGTGAAATTCTGTAATCCCTGTGGGACCACCGGTGGCGAAAGCGTCCGACCAGAACGGGTCTGACGGTCAGGGACGAAGCCTTGGGGCG |
| p__Thermoplasmatota_g__SM1-50_ASV_94 | 1 | 0 | 0 | 0 | 0 | 0 | 0 | 1 | 1 | 1 | 1 | CCTTCGCAATGTGCGAAAGCACGACGAGGGAATCCCAAGTGTTTGCACGTAAGTGTAAACTGTTTTTCTGTCTAAAACACAGAGAGAGTAAGAGCTGGGTAAGACGGGTGCCAGCCGCCGCGGTAATACCTGCAGCTCAAGTGGTGGCCGTTATTATTGGGCCTAAAACGTTCGTAGCCGGTTTAGTAAATGCCTGGGTAAATCGTGTAGCTTAACTATACGAATTCCGGGTAGACTGCTAAACTTGAGACCGGGAGAAGTTAGAGGTACTCCTGGGGTAGAGGTGAAATTCTGTAATCCTAGGGGGACCACCAGTGGCGAAGGCGTCTAACTAGAACGGGTCTGACGGTGAGGGACGAAGCCCTGGGGCG |
| p__Thermoplasmatota_o__Methanomassiliicoccales_ASV_98 | 1 | 1 | 1 | 0 | 1 | 1 | 1 | 1 | 1 | 1 | 1 | ACTTCACACTGGGGGAAACCCCGATGAGGGGATTCCAAGTGCCAGCACACAGTGTTGGCTGTTCTCCTGTCTAAATCGCAGGAGGAGTAAGGGCCGGGTAAGACGGGTGCCAGCCGCCGCGGTAATACCCGCGGCCCGAGTGGTGGTCGATATTATTGAGCCTAAAACGTCCGTAGCCGGCCTTGTAAATCCTTGGGTAAATCGGCCAGCTTAACTGTCCGAAGTCCGGGGAGACTGCAAGGCTTGGGACCGGGAGAGGCCAGAGGTACTTCCAGGGTAGGGGTAAAATCCTGTAATCCTGGAAGGACCACCGGTGGCGAAGGCGTCTGGCTAGAACGGATCCGACGGTGAGGGACGAAGCCCTGGGGCG |
| p__Halobacteriota_g__Halohasta_ASV_100 | 0 | 0 | 1 | 1 | 0 | 0 | 0 | 0 | 1 | 0 | 0 | CCTTTACACTGCACGCAAGTGCGATAGGGGAATCCCAAGTGCGAGGGCATATTAGCCCTCGCTTTTGTGTACTGTAAGGAGGTACACGAATAAGAGCTGGGCAAGACCGGTGCCAGCCGCCGCGGTAATACCGGCAGCTCGAGTGATGTCCACTATTATTGGGCCTAAAGCGTCCGTAGCTGGCCGCGCAAGTCCGTCGGGAAATCCACCTGCCCAACAGGTGGGCGTCCGGCGGAAACTGCGTGGCTTGGAACCGGAAGGCTCAGAGGGTACGTCCGGGGTAGGAGTGAAATCCCGTAATCCCGGACGGACCACCGATGGCGAAAGCACTCTGAGAGGACGGATTCGACAGTGAGGGACGAAAGCTAGGGTCT |
| p__Halobacteriota_o__Halobacteriales_ASV_106 | 1 | 1 | 1 | 1 | 0 | 1 | 0 | 1 | 1 | 1 | 1 | CCTTTACACTGTACGAAAGTGCGATAAGGGGACCCCGAGTGCGAGGGCATACAGTCCTCGCTTTTGTGTACCGTACGATGGTACACGAATAAGAGCTGGGCAAGACCGGTGCCAGCCGCCGCGGTAATACCGGCAGCTCAAGTGATGGCCGTTATTATTGGGCCTAAAGCGTCCGTAGCCGGCCGAACAGGTCTGTTGGGAAATCCGCCAGCTTAACTGGCGGGCGTCCAGCGGAAACCAGTCGGCTTGGGGCCGGAAGACCTGAGGGGTACGTCCGGGGTAGGAGTGAAATCCTGTAATCCTGGACGGACCGTCGGTGGCGAAAGCGCCTCAGGAGGACGGACCCGACGGTGAGGGACGAAAGCTAGGGTCA |
| p__Nanoarchaeota_g__BMS3ABIN17_ASV_109 | 1 | 1 | 1 | 1 | 1 | 1 | 1 | 0 | 1 | 0 | 1 | CTTCTGCAATGCGCGAAAGCGTGACAGAGTGAGTCAGAGTGTCTGCCATTTGGTAGACTTTTGTGGAATGTAAAGAGTTCTACGAATAAGGACTGGGTAAGACTAGTGCCAGCCGCCGCGGTAATACTAGCGGTCCAAGTCGCAGCCACTTTTATTGGGTCTAAAACATCCGTAGCTTGCTTGATAAGTCTCTTGTGAAATCCTGTATCTCAAGTATAGGACTTGCAAGAGATACTGTCAGGCTAGAGACCGGGAGACGTAAGGAGTACGTTCGAAGTAGCGGTAAAATGTGTTGATCTCGAACGGACTAACAAGAGCGAAGGCACCTTACGAGAACGGATCTGACAGTAAGGGATGAAGGCCAGGGTCG |
| p__Halobacteriota_g__Halorubrum_ASV_114 | 1 | 1 | 1 | 1 | 0 | 0 | 1 | 1 | 1 | 1 | 1 | CCTTTACACTGCACGCCAGTGCGATAGGGGGATCCCAAGTGCACAGGCATAGTGCCTGTGCTTTTCGACACCCTAAGGCGGTGTCAGAATAAGGGCTGGGCAAGACCGGTGCCAGCCGCCGCGGTAATACCGGCAGCCCGAGTGATGGCCGATCTTATTGGGCCTAAAGCGTCCGTAGCTGGCCGCGCAAGTCCATCGGGAAATCCACCTGCTCAACAGGTGGCCGCCCGGTGGAAACTGCGTGGCTTGGGACCGGAAGGCGCGACGGGTACGTCCGGGGTAGGAGTGAAATCCCGTAATCCTGGACGGACCACCGATGGCGAAAGCACGTCGCGAGAACGGATCCGACAGTGAGGGACGAAAGCCAGGGTCT |
| p__Halobacteriota_g__Methanohalobium_ASV_118 | 0 | 1 | 1 | 1 | 0 | 1 | 1 | 1 | 0 | 0 | 1 | ACTTTACAATGCGGGAAACCGCGATAAGGGGACACTGAGTGCCAGCATATAATGCTGGCTGTCCGTATGTGTAAAATACATGCGTTAGCAAGGGCCGGGCAAGACCGGTGCCAGCCGCCGCGGTAACACCGGCGGCCCGAGTGGTGGTCGCTATTATTGGGTCTAAAGGGTCCGTAGCCGGTTTGATCAGTCCATTGGGAAATCTGGTGGCTTAACCGCCAGAATTCCAATGGATACTGTCAGACTTGGGACCGGGAGAGGTGAGAGGTACTACAGGGGTAGGAGTGAAATCTTGTAATCCCTGTAGGACCACCAGTTGCGAAGGCGTCTCACCAGAACGGGTCCGACGGTGAGGGACGAAAGCTGGGGGCA |
| p__Nanoarchaeota_o__Woesearchaeales_581_ASV_120 | 1 | 1 | 1 | 0 | 0 | 0 | 0 | 0 | 0 | 0 | 1 | CCTTTACAATGCACGAAAGTGTGATAAGGGAACCCCCAGTGCCAGTGGAAACGCTGGCTTTTTATGAGGGTAAGGTCCTCATAGAATAAGCAGTGGGTAAGACCGGTGCCAGCCGCCGCGGTAACACCGGCGCTGCTAGTGGGAGTCGCGATTATTGGGCCTAAAGCGTTCGTAGCCGGTTTGGTAAATTCTCTGTGAAATCCTTGGGCTTAACCTAAGGCGAGCAGGGGAGACTGCCAAACTAGAGATTGGGAAGCGTTGGAGGTATTTCAGGGGGAGCGGTAAAATGCTATAAGCCCTGAAAGACCACCTGTGGCGAAGGCGTCCAGCGGGAACAAATCTGACGGTGAGGGACGAAGGCCAGGGGAG |
| p__Halobacteriota_g__Halostagnicola_ASV_123 | 1 | 1 | 1 | 1 | 1 | 1 | 1 | 1 | 0 | 1 | 1 | CCTTTACACTGCACGCGAGTGCGATAAGGGGACTCCAAGTGCGAGGGCATATAGTCCTCGCTTTTTCCGACCGTAGGATGGTCGGCGAATAAGTGCTGGGCAAGACCGGTGCCAGCCGCCGCGGTAATACCGGCAGCACGAGTGATGACCGCTATTATTGGGCCTAAAGCGTCCGTAGCTGGCCACGCAAGTCCATCGGGAAATCTCTCCGCTCAACGGAGAGGCGTCCGGTGGAAACTGTGTGGCTTGGGACCGGAAGATCCAGAGGGTACGTCTGGGGTAGGAGTGAAATCCCGTAATCCGAGACGGACCACCGGTGGCGAAAGCGCTCTGGAAAGACGGATCCGACGGTGAGGGACGAAAGCTCGGGTCA |
| p__Halobacteriota_f__Natrialbaceae_ASV_126 | 1 | 1 | 1 | 0 | 0 | 0 | 1 | 0 | 0 | 0 | 1 | CCTTTACACTGCACGCCAGTGCGATAAGGGGACTCCAAGTGCGAGGGCATACAGTCCTCGCTTTTGTGTACCGTAAGGTGGTACACGAATAAGGGCTGGGCAAGACCGGTGCCAGCCGCCGCGGTAATACCGGCAGCCCGAGTGATGACCGCTATTATTGGGCCTAAAGCGTCCGTAGCCGGCCGGGTAAGTCCGTCGGGAAATCCGCGCGCTCAACGCGCGGGCGTCCGGTGGAAACTGCCTGGCTTGGGACCGGAAGATCCAGAGGGTACGTCCGGGGTAGGAGTGAAATCCTGTAATCCTGGACGGACCACCGGTGGCGAAAGCGCTCTGGAAGGACGGATCCGACGGTGAGGGACGAAAGCTAGGGTCA |
| p__Thermoplasmatota_g__SM1-50_ASV_127 | 0 | 0 | 0 | 0 | 0 | 0 | 0 | 1 | 0 | 0 | 0 | CCTTCACACTGTGCGAACGCGCGATGAGGGGATCCCAAGTGCTTGCACGTAAGTGTAAGCTGTTTCTTTGTTTAAAATGCATAGAGAGTAAGGGCTGGGTAAGACGGGTGCCAGCCGCCGCGGTAATACCTGCAGCTCAAGTGGTGACCACTATTATTGGGCCTAAAACGTCCGTAGCCTGTTTGATAAATGCCTGGGTAAATCGTATAGCTCAACTATACGAATTCCGGGTAGACTGTCAAGCTTGAGACCGGGAGAAGCTAGAGGTACTCTTGGGGTAGAGGTGAAATTCTGTAATCCTGAGGGGACCACCAGTGGCGAAGGCGTCTAGCTAGAACGGGTCTGACGGTGAGGGACGAAGCCCTGGGGCG |
| p__Halobacteriota_f__Natrialbaceae_ASV_135 | 1 | 1 | 1 | 1 | 0 | 0 | 0 | 0 | 1 | 1 | 1 | CCTTTACACTGCACGCCAGTGCGATAAGGGGACACCAAGTGCGAGGGCATATCGTCCTCGCTTTTCACCACCGTAAGGAGGTGGTAGAATAAGAGCTGGGCAAGACCGGTGCCAGCCGCCGCGGTAATACCGGCAGCTCGAGTGATGACCGCTATTATTGGGCCTAAAGCGTCCGTAGCTGGCCATACAAGTCCATCGGGAAATCTCCCTGCTTAACAGGGAGGCGTCCGGTAGAAACTGTGTGGCTTGGGACCGGAAGATCCAGAGGGTACGTCTGGGGTAGGAGTGAAATCCCGTAATCCTAGACGGACCACCGGTGGCGAAAGCGCTCTGGAAGGACGGATCCGACAGTGAGGGACGAAAGCTAGGGTCA |
| p__Thermoplasmatota_g__SM1-50_ASV_146 | 0 | 0 | 0 | 0 | 0 | 0 | 0 | 1 | 1 | 1 | 1 | CCTTCACACTGTGCGAAAGCGCGATGAGGGAATCCCAAGTGTATGCACGTAAGTGTATACTGTTTCTATGTCTAAAAAGCAGAGAGAGTAAGAGCTGGGTAAGACGGGTGCCAGCCGCCGCGGTAATACCTGCAGCTCAAGTGGTGGCCGCTATTATTGGGCCTAAAACGTTCGTAGCCGGTCTTTTAAATGCCTGGGTAAATCGAGCCGCTTAACGGTTCGAATTCCGGGTAGACTGGAAGACTTGAGACCGGGAGAGGCTAGAGGTACTCCTGGGGTAGAGGTGAAATTCTGTAATCCTGGGGGACCACCAGTGGCGAAGGCGTCTAGCTAGAACGGGTCTGACGGTGAGGGACGAAGCCCTGGGGCG |
| d__Archaea_d__Archaea_ASV_151 | 0 | 0 | 0 | 0 | 0 | 0 | 0 | 1 | 1 | 1 | 1 | CCTCCACAATGCGCGAAAGCGTGATGGGGGGAACCCGCGTGCTTGCGCAAAACGTAAGCTTTTCTGGAGTGTAAGTAGCTCTGGGAATAAGTGGTGGGCAAGACCGGTGCCAGCCGCCGCGGTAATACCGGCGCCACAAGTGATAGCCGCGTTTATTGGGCCTAAAGCATCCGTAGCCGGACAAGTAGGTTTCCAGTGAAATCTAACAGCTTAACTGTTAGGCGTGCTGGGAGCACCGCTTGTCTTGAGACCGGGAGGAGTTAAGAGAATTTCTGGGGGAGCGGTAAAATGTTGTAATCCCAGGAGGACTATCTGTGGCGAAGGCGCTTAACGAGAACGGGTCTGACGGTCAGGGATGAAAGCTGGGGGAG |
| p__Halobacteriota_f__Natrialbaceae_ASV_158 | 1 | 1 | 1 | 1 | 1 | 0 | 0 | 0 | 0 | 0 | 1 | CCTTTACACTGCACGCCAGTGCGATAAGGGGACTCCCAGTGCGAGGGCATATCGTCCTCGCTTTTGTGTACCGTAGGGAGGTACACGAATAAGGGCTGGGCAAGACCGGTGCCAGCCGCCGCGGTAATACCGGCAGCCCGAGTGATGACCGCTATTATTGGGCCTAAAGCGTCCGTAGCCGGCCACGCAAGTCCGTCGGGAAATCCGCACGCCCAACGTGCGGGCGTCCGGCGGAAACTGCCTGGCTTGGGACCGGAAGACCAGAGGGGTACGTCCGGGGTAGGAGTGAAATCCCGTAATCCTGGACGGACCACCGGTGGCGAAAGCGCCTCTGGAAGACGGATCCGACGGTGAGGGACGAAAGCTCGGGTCA |
| p__Halobacteriota_g__Methanohalophilus_ASV_159 | 1 | 1 | 1 | 1 | 0 | 1 | 1 | 1 | 1 | 1 | 1 | CCTTTACACTGCGGGAAACCGCGATAAGGGGACACCGAGTGCCAGCATACAATGCTGGCTGTCCACATGTGTAAATAGCATGTGATAGCAAGGGCCGGGCAAGACCGGTGCCAGCCGCCGCGGTAACACCGGCGGCCCGAGTGGTAATCACTATTATTGGGTCTAAAGGGTCCGTAGCCGGTTTAATCAGTCTTCCGGGAAATCTGACAGCTCAACTGTTAGGCTTCCGGGGGATACTGTTAGACTTGGGACCGGGAGAGGTAAGAGGTACTACAGGGGTAGGAGTGAAATCTTGTAATCCCTGTGGGACCACCAGTGGCGAAGGCGTCTTACCAGAACGGGTCCGACGGTGAGGGACGAAAGCTGGGGGCA |
| p__Thermoplasmatota_g__UBA147_ASV_161 | 0 | 0 | 0 | 0 | 0 | 0 | 0 | 0 | 1 | 1 | 1 | ACTTGTCAATGGGGGAAACCCCGAACAGGGGACTCCAAGTGCATGCGTTATACGTATGCTGTTCGGTTGCCTAAAAAGCAGCCGAAGGAAGGGCTGGGTAAGACAGGTGCCAGCCGCCGCGGTAATACCTGCAGCCCAAGTGGTGATCGTTATTATTGAGCCTAAAGCGTCCGTAGCCGGTCCGGTAAATCCTCGGGTAAATCGAGCAGCTTAACTGTTCGAACACCGGGGGGACTGCCGGACTTGGGACCGGGAGAGGTCAGAGGTACTTCAGGGGTAGGGGCTAAATCCTGTAATCCTTGAAGGACCACCAGTGGCGAAGGCGTCTGACTAGAACGGATCCGACGGTAAGGGACGAAACCCTGGGGCG |
| p__Thermoplasmatota_g__SM1-50_ASV_162 | 0 | 0 | 0 | 0 | 0 | 0 | 0 | 1 | 1 | 1 | 1 | ACTTCGCACTGTGCGAAAGCACGACGAGGGGATCCCAAGTGCTGGCTCGTAAGAGTCAGCTGTTTCTTTGTCTAAAAAACAGAGAGAGTAAGAGCTGGGTAAGACGGGTGCCAGCCGCCGCGGTAATACCTGCAGCTCAAGTGGTGGCCGCTATTATTGGGCCTAAAACGTCCGTAGCCGGTTCTATAAATGCCTGGGTAAATCGTACGGCTTAACCGTACGAATTCCGGGTAGACTGTTCAACTTGGGACCGGGAGAGGCTAGAGGTACTCCTGGGGTAGAGGTGAAATTCTGTAATCCTAGGGGGACCACCAGTGGCGAAGGCGTCTAGCTAGAACGGGTCTGACGGTGAGGGACGAAGCCCTGGGGAG |
| p__Halobacteriota_o__Halobacteriales_ASV_166 | 1 | 1 | 1 | 1 | 1 | 0 | 1 | 0 | 0 | 0 | 1 | CCTTTACACTGTACGAAAGTGCGATAAGGGGACCCCGAGTGCGAGGGCATACAGTCCTCGCTTTTGTGTACCGTACGATGGTACACGAATAAGAGCTGGGCAAGACCGGTGCCAGCCGCCGCGGTAATACCGGCAGCTCAAGTGATGGCCGTTATTATTGGGCCTAAAGCGTCCGTAGCCGGCCGAACAGGTCTGTTGGGAAATCCGCCAGCTTAACTGGCGGGCGTCCAGCAGAAACCAGTCGGCTTGGGGCCGGAAGACCTGAGGGGTACGTCCGGGGTAGGAGTGAAATCCTGTAATCCTGGACGGACCGTCGGTGGCGAAAGCGCCTCAGGAGGACGGACCCGACGGTGAGGGACGAAAGCTAGGGTCA |
| p__Halobacteriota_g__Halomarina_3560_ASV_169 | 1 | 1 | 1 | 1 | 1 | 0 | 0 | 0 | 0 | 0 | 0 | CCTTTACACTGCACGACAGTGCGATAGGGGGATTCCGAGTGCGAGGGCATACAGTCCTCGCTTTTGTGTACCGTAAGGTGGTACACGAATAAGAGCTGGGCAAGACCGGTGCCAGCCGCCGCGGTAATACCGGCAGCTCGAGTGATGGCCGATTTTATTGGGCCTAAAGCGTCCGTAGCCGGCCAGTCAGGTCCGTCGGGAAATCCACTCGCTCAACGAGTGGGCGTCCGGCGGAAACCGTCTGGCTTGGGACCGGGAGATCTGAGGGGTACGTCCGGGGTAGGAGTGAAATCCTGTAATCCTGGACGGACCATCGGTGGCGAAAGCGCCTCAGAAGAACGGATCCGACGGTGAGGGACGAAAGCTAGGGTCT |
| p__Thermoplasmatota_g__SM1-50_ASV_170 | 0 | 0 | 0 | 1 | 0 | 0 | 0 | 1 | 1 | 1 | 1 | ACTTCACAATGCGCGAAAGCGCGATGAGGGGATCCCAAGTGCTTGCACGTAAGTGTAAGCTGTTTTTCTGTCTAAAAAGCAGAGAGAGTAAGGGCTGGGTAAGACGGGTGCCAGCCGCCGCGGTAATACCTGCAGCTCAAGTGGTGGCCGCTATTATTGGGCCTAAAACGTCCGTAGCCGGCTGAGTAAATGCCTGGGTAAATCGTGCCGCTTAACGGTACGAATTCCGGGTAGACTGCTTAGCTTGGGACCGGAAGAAGTTAGAGGTACTCCTGGGGTAGGGGTGAAATCCTGTAATCCTGGGGGGACCACCAGTGGCGAAGGCGTCTAACTAGGACGGGTCTGACGGTGAGGGACGAAGCCCTGGGGCG |

| Phylum_Genus_ASV_number | Presence (1) or Absence (0) in each layer | | | | | | | | | | | |
|---|---|---|---|---|---|---|---|---|---|---|---|---|
| | L1 | L2 | L3 | L4 | L5 | L6 | L7 | L8 | L9 | L10 | L11 | Sequence |
| p__Halobacteriota_g__Halostagnicola_ASV_171 | 1 | 1 | 1 | 1 | 1 | 1 | 1 | 1 | 1 | 1 | 1 | CCTTTACACTGCACGCGAGTGCGATAAGGGGACTCCAAGTGCGAGGGCATATAGTCCTCGCTTTTCGTGACCGTAGGATGGTCACAGAATAAGTGCTGGGCAAGACCGGTGCCAGCCGCCGCGGTAATACCGGCAGCACGAGTGATGACCGCTATTATTGGGCCTAAAGCGTCCGTAGCTGGCCACGCAAGTCCATCGGGAAATCTCTCCGCTCAACGGAGAGGCGTCCGGTGGAAACTGTGTGGCTTGGGACCGGAAGATCCAGAGGGTACGTCTGGGGTAGGAGTGAAATCCCGTAATCCGAGACGGACCACCGGTGGCGAAAGCGCTCTGGAAAGACGGATCCGACGGTGAGGGACGAAAGCTCGGGTCA |
| p__Halobacteriota_g__Halohasta_ASV_177 | 1 | 1 | 1 | 1 | 0 | 0 | 0 | 0 | 0 | 0 | 0 | CCTTTACACTGCACGCAGGTGCGATAGGGGAATCCCAAGTGCGAGGGCATACTAGCCCTCGCTTTTGTGTACTGTAAGGAGGTACACGAATAAGAGCTGGGCAAGACCGGTGCCAGCCGCCGCGGTAATACCGGCAGCTCGAGTGATGTCCACTATTATTGGGCCTAAAGCGTCCGTAGCTGGCCGTACAAGTCCGTCGGGAAATCCACCTGCCCAACAGGTGGGCGTCCGGCGGAAACTGTATGGCTTGGAACCGGAAGGCTCAGAGGGTACGTCCGGGGTAGGAGTGAAATCCTGTAATCCCGGACGGACCACCGATGGCGAAAGCACTCTGAGAGGACGGATTCGACAGTGAGGGACGAAAGCTAGGGTCT |
| p__Halobacteriota_g__Halorientalis_ASV_179 | 1 | 1 | 1 | 1 | 1 | 1 | 1 | 1 | 0 | 1 | 1 | CCTTTACACTGCACGACAGTGCGATAGGGGGACTCCGAGTGCGAGGGCATATAGTCCTCGCTTTTCTGTACCGTAAGGTGGTACAGGAACAAGTGCTGGGCAAGACCGGTGCCAGCCGCCGCGGTAATACCGGCAGCACCAGTGATGGCCGCTTTTATTGGGCCTAAAGCGTTCGTAGCCGGCCATGCAAGTTCGTCGGGAAATCTGCCCGCTTAACGGGCAGGCGTCCGGCGAAAACTGCATGGCTTGGGGCCGGGAGATCCAAGGGGTACGTCCGGGGTAGGAGTGAAATCCCGTAATCCTGGACGGACCACCGATGGCGAAAGCACCTTGGAAACACGGACCCGACGGTGAGGAACGAAAGCTAGGGTCT |
| d__Archaea_d__Archaea_ASV_185 | 1 | 1 | 1 | 0 | 0 | 1 | 0 | 0 | 0 | 0 | 1 | CCTTCACAATGCTCGAAAGAGTGATGAGGGTACTCCGAGTGCCAGTGCTTAGCACTGGCTTTTCTGCAGCGTAAATAGTTGCAGGAATAAGGGCTGGGCAAGACCGGTGCCAGCCGCCGCGGTAACACCGGCAGCCCTAGTGGTGACCGCTTTTATTGGGCCTAAAGCGTTCGTAGCCGGGCCGGTAAGTCCTTGGGTAAATCCTGCGTCTCAAGCGCGGGAATTCTAAGGATACTACCGGCCTTGAGGCCGGGATAGGCGAGAGGTACTCCCGGGGTAGGGGTAAAATCCTATAATCCTGGGGGACCACCGATGGCGAAGGCATCTCGCTGGAACGGGCCTGACGGTGAGGGACGAAACCCAGGGGAG |
| p__Thermoproteota_g__GCA-2726865_ASV_189 | 0 | 0 | 0 | 1 | 1 | 0 | 0 | 1 | 1 | 1 | 1 | CCTCCTCAATGCACTAACGTGCGAAGGGGTTACCTCGAGTGCTACCTGAAGAAGGTGGCTTTTCCTTGGTCTAAAAAGCCTCGGGAATAAGAGGGGGGCAAGACTGGTGTCAGCCGCCGCGGTAACACCAGCCCCTCGAGTGGTGGGGATGATTATTTGGCCTAAAGCGTCCGTAGCCGGCCGGGTAAGTTTCCCGTTAAATCCAACGTCTCAAGCGTTGGACTGCGGGAAATACTGCCTGGCTAGAGAGTGGGAGAGGTCAGCGGTATTCTGGGGGTAGGGGCGAAATCCTTTGATCCCCGGAGGACCACCAGTGGCGAAGGCTGCTGACTAGAACACGTCTGACGGTCAGGGACGAAAGCTGGGGTAG |
| p__Thermoplasmatota_g__SM1-50_ASV_191 | 0 | 0 | 0 | 1 | 0 | 0 | 0 | 1 | 1 | 1 | 1 | ACTTCACAATGCGCGAAAGCGCGATGAGGGAATCCCAAGTGTATGCACGTAAGTGTATACTGTTTCTCTGTCTAAAAAGCAGAGAGAGTAAGGGCTGGGTAAGACGGGTGCCAGCCGCCGCGGTAATACCTGCAGCTCAAGTGGTGGCCGTTATTATTGGGCCTAAAACGTCCGTAGCCGGTCTATTAAATGCCTGGGTAAATCGTGCCGCTTAACGGTACGAATTCCGGGTAGACTGATAGACTTGAGATCGGGAGAGGCTAGAGGTACTCCTGGGGTAGAGGTGAAATTCTGTAATCCTGGGGGGACCACCAGTGGCGAAGGCGTCTAGCTAGAACGAGTCTGACGGTGAGGGACGAAGCCCTGGGGCG |
| p__Halobacteriota_g__Natrarchaeobaculum_ASV_193 | 1 | 1 | 1 | 1 | 0 | 1 | 1 | 0 | 0 | 1 | 1 | CCTTTACACTGCACGCCAGTGCGATAAGGGGACTCCATCTGCGAGGGCATATCGTCCTCGCTTTTTCCTACCGTAAGGTGGTAGGCGAATAAGGGCTGGGCAAGACCGGTGCCAGCCGCCGCGGTAATACCGGCAGCCCGAGTGATGACCGCTCTTATTGGGCCTAAAGCGTCCGTAGCTGGCCTCGCAAGTCCATCGGGAAATCCGCACGCTCAACGTGCGGGCGTCCGGTGGAAACTGCGTGGCTTGGGACCGGAAGACCCGAGGGGTACGTCCGGGGTAGGAGTGAAATCCCGTAATCCTGGACGGACCACCGGTGGCGAAAGCGCCTCGGGAGGACGGATCCGACGGTGAGGGACGAAAGCCAGGGTCA |
| d__Archaea_d__Archaea_ASV_194 | 0 | 0 | 0 | 0 | 0 | 0 | 0 | 0 | 1 | 1 | 1 | CTTCTGCAATGACCGTAAGGTTGACAGAGCGATTCCGAGTGCTCATGCAAAGCATGGGCTTTTGCCAAGTGTAAACAGCTTGGCGAATAAGGGGTGGGCAAGACCGGTGGCAGCAGCCGCGGTAATACCGGCGCCCCGAGTGGTGGTCATTATTATTGGGCCTAAAGCGTCCGTAGCTGGTTTGGTTAGTTTTTGGTGAAATCCATGCGCTCAACGTATGGGCGAGCCAGAAATACTACCAAGCTAGAGACCGGGAGAAGACAGGGGTATTCCTAGGGTAAGGGTAAAATCTTGTAATCCTAGGAGGACCACCGGTGGCGAAGGCGCCTGTCTAGAACGGGTCTGACAGTGAGGGACGAAAGCTGAGGGAG |
| p__Thermoproteota_g__BA2_ASV_199 | 0 | 0 | 0 | 0 | 0 | 0 | 0 | 1 | 1 | 1 | 1 | ACTCCGCAATGCACGAAAGTGTGACGGGGCTACCCCGAGTGCCATCCGCTGAGGATGGCTTTTCCTTGGTATAACTAGCCAGGGGAATAAGGAGAGGGCAAGACTGGTGTCAGCCGCCGCGGTAATACCAGCTCTCCGAGTGGTAAGGATGATTATTGGGCTTAAAGCGTCCGTAGCCGGCTTAGCAAGTCTCCTGTTAAACTCAGCGACCTAATCGTTGAATCGCGGGAGATACTATTAGGCTAGGAGGCGGGAGAGGTCGACGGTACTTCTGGGGTAGGGGTGAAATCCTATAATCCCGGGAGGACCACCAGTGGCGAAGGCTGTCGACTAGAACGCGCTCGACGGTGAGGGACGAAAGCTGGGGGAG |
| p__Halobacteriota_g__Natrinema_ASV_200 | 0 | 1 | 1 | 0 | 0 | 0 | 1 | 0 | 0 | 0 | 0 | CCTTTACACTGCACGCCAGTGCGATAAGGGGACTCCAAGTGCGAGGGCATACAGTCCTCGCTTTTGTGTACCGTAAGGTGGTACACGAATAAGGGCTGGGCAAGACCGGTGCCAGCCGCCGCGGTAATACCGGCAGCCCAAGTGATGACCGCTATTATTGGGCCTAAAGCGTCCGTAGCCGGCCAGGTAAGTCCGTCGGGAAATCCGCGCGCTCAACGCGCGGGCGTCCGGTGGAAACTGCCTGGCTTGGGACCGGAAGATCCAGAGGGTACGTCCGGGGTAGGAGTGAAATCCTGTAATCCTGGACGGACCACCGGTGGCGAAAGCGCTCTGGAAAGACGGATCCGACGGTGAGGGACGAAAGCTAGGGTCA |
| p__Halobacteriota_g__Halorussus_ASV_203 | 1 | 1 | 1 | 1 | 0 | 0 | 0 | 0 | 1 | 1 | 0 | ACTTTACACTGCACGACAGTGCGATAAGGGAACTCCGAGTGCGAGGGCATATAGTCCTCGCTTTTCTGCACCGTAAGGTGGTGCACGAATAAGAGCTGGGCAAGACCGGTGCCAGCCGCCGCGGTAATACCGGCAGCTCAAGTGATGGCCGCTATTATTGGGCCTAAAGCGTCCGTAGCCGGCCAGACAGGTCCGTCGGGAAATCCACGCGCTCAACGCGTGGGCGTCCGGCGGAAACCAGCTGGCTTGGGGCCGGAAGACCTGAGGGGTACGTCCGGGGTAGGAGTGAAATCCCGTAATCCTGGACGGACCACCGGTGGCGAAAGCGCCTCAGGAAAACGGACCCGACGGTGAGGGACGAAAGCTAGGGTCA |
| p__Halobacteriota_g__Natronoarchaeum_ASV_210 | 1 | 1 | 1 | 1 | 1 | 1 | 1 | 0 | 1 | 1 | 1 | CCTTTACACTGCACGCCAGTGCGATAAGGGGACCCCCAGTGCGAGGGCATACCGTCCTCGCTTTTCACCACCGTAAGGAGGTGGTGGAACAAGAGCTGGGCAAGACCGGTGCCAGCCGCCGCGGTAATACCGGCAGCTCGAGTGATGGCCGTTTTTATTGGGCCTAAAGCGTCCGTAGCCGGCCAGACAAGTTCGTCGGGAAATCCACGCGCCCAACGCGTGGGCGTCCGGCGAAACCTCTCTGGCTTGGGACCGGACGGCCCGAGGGGTACGGTCGGGGTAGGAGTGAAATCCTGTAATCCCGACCGGACCACCGGTGGCGAAAGCGCCTCGGGAGGACGGATCCGACGGTGAGGGACGAAAGCCAGGGTCA |
| p__Halobacteriota_g__Halohasta_ASV_217 | 0 | 0 | 1 | 1 | 0 | 1 | 1 | 1 | 1 | 1 | 1 | CCTTTACACTGCACGCAAGTGCGATAGGGGAATCCCAAGTGCGAGGGCATACTAGCCCTCGCTTTTGTGTACTGTAAGGAGGTACACGAATAAGAGCTGGGCAAGACCGGTGCCAGCCGCCGCGGTAATACCGGCAGCTCGAGTGATGTCCACTATTATTGGGCCTAAAGCGTCCGTAGCTGGCCGTACAAGTTCGTCGGGAAATCCACCTGCCCAACAGGTGGGCGTCCGGCGAAAACTGTATGGCTTGGAACCGGAAGGCTCAGAGGGTACGTCCGGGGTAGGAGTGAAATCCTGTAATCCCGGACGGACCACCGATGGCGAAAGCACTCTGAGAAGACGGATTCGACAGTGAGGGACGAAAGCTAGGGTCT |
| p__Thermoplasmatota_g__SM1-50_ASV_220 | 1 | 0 | 0 | 0 | 0 | 1 | 1 | 0 | 0 | 0 | 1 | ACTTCACACTGCGCGAAAGCGCGATGAGGGGATCCCAAGTGTATGCACGTAAGTGTATACTGTTTCTATGTTTAAAAAGCATTGAGAGTAAGGGCTGGGTAAGACGGGTGCCAGCCGCCGCGGTAATACCTGCAGCTCGAGTGGTGATCACTATTATTGGGCCTAAAACGTCCGTAGCCGGTCTAGTAAATGCCTGGGTAAATCGTACGGCTTAACCGTACGAATTCTGGGTAGACTGCTAAACTTGAGACCGGGAGAGGCTAGAGGTACTCCTGGGGTAGAGGTGAAATTCTGTAATCCTAGGGGGACCACCAGTGGCGAAGGCGTCTAGCTAGAACGGGTCTGACGGTGAGGGACGAAGCCCTGGGGCG |
| p__Halobacteriota_g__Halorubrum_ASV_225 | 1 | 1 | 1 | 1 | 0 | 1 | 0 | 0 | 0 | 0 | 1 | CCTTTACACTGCACGACAGTGCGATAGGGGGATCCCAAGTGCACAGGCATAGCGCCTGTGCTTTTCGGTACCGTAAGGTGGTACCAGAATAAGGGCTGGGCAAGACCGGTGCCAGCCGCCGCGGTAATACCGGCAGCCCAAGTGATGGCCGATCTTATTGGGCCTAAAGCGTCCGTAGCTGGCCACGCAAGTCCATCGGGAAATCCACCTGCTCAACAGGTGGGCGCCCGGTAGAAACTGCGTGGCTTGGGACCGGAAGGCGCGACGGGTACGTCCGGGGTAGGAGTGAAATCCCGTAATCCTGGACGGACCGCCGATGGCGAAAGCACGTCGCGAGAACGGATCCGACAGTGAGGGACGAAAGCCAGGGTCT |
| p__Halobacteriota_g__Halorubrum_ASV_226 | 1 | 1 | 1 | 1 | 1 | 0 | 1 | 1 | 0 | 0 | 1 | CCTTTACACTGCACGACAGTGCGATAAGGGAATCCCAAGTGCGTAGGCATAGAGCCTACGCTTTTGTCCACCGTAGGGAGGTGGACGAATAAGGGCTGGGCAAGACCGGTGCCAGCCGCCGCGGTAATACCGGCAGCCCGAGTGATGGCCGATCTTATTGGGCCTAAAGCGTCCGTAGCTGGCCGCACAAGTCCATCGGAAAATCCACCCGCTCAACGGGTGGGCGTCCGGTGGAAACTGTGTGGCTTGGGACCGGAAGGCGCGACGGGTACGTCCGGGGTAGGAGTGAAATCCCGTAATCCTGGACGGACCGCCGATGGCGAAAGCACGTCGCGAGAACGGATCCGACAGTGAGGGACGAAAGCTAGGGTCT |
| p__Halobacteriota_g__Halovenus_ASV_233 | 1 | 1 | 1 | 1 | 1 | 1 | 1 | 0 | 0 | 0 | 1 | CCTTTACACTGCACGACAGTGCGATAGGGGGACTCCGAGTGTGAGGGCATATAGTCCTCACTTTTGTACACCGTAAGGTGGTGTACGAATAAGAACTGGGCAAGACCGGTGCCAGCCGCCGCGGTAATACCGGCAGTTCGAGTGATGGCCGCTTTTATTGGGCCTAAAGCGTTCGTAGCTGGCCAGTCAAGTCTGTCGGGAAATCCAGTGGCTTAACCACTGGACGTCCGGCGGAAACTGATTGGCTTGGGACCGGGAGACTCGAGGAGTACGTTCGGGGTAGGAGTGAAATCCCATAATCCTGAACGGACTACCGATGGCGAAAGCACCTCGAGAGCACGGATCCGACAGTGAGGAACGAAAGCTGGGGTCT |
| d__Archaea_d__Archaea_ASV_241 | 0 | 0 | 0 | 0 | 0 | 0 | 0 | 0 | 1 | 1 | 1 | CCTCCACAATGCGCGAAAGCGTGATGGGGGGAACCCGCGTGCTTGCGCAAAACGTAAGCTTTTCTGGAGTGTAAGTAGCTCTGAGAATAAGTGGTGGGCAAGACCGGTGCCAGCCGCCGCGGTAATACCGGCGCCACAAGTGATAGCCGCGTTTATTGGGCCTAAAGCATCCGTAGCCGGACAAGTAGGTTTCCAGTGAAATCTAACAGCTTAACTGTTAGGCGTGCTGGGAGCACCGCTTGTCTTGAGACCGGGAGGAGTTAAGAGAATTTCTGGGGGAGCGGTAAAATGTTGTAATCCCAGGAGGACTATCTGTGGCGAAGGCGCTTAACGAGAACGGGTCTGACGGTCAGGGATGAAAGCTGGGGGAG |
| p__Halobacteriota_g__Halovivax_ASV_243 | 1 | 1 | 1 | 0 | 0 | 1 | 1 | 0 | 0 | 1 | 1 | CCTTTACACTGCACGCCAGTGCGATAGGGGGACTCCAAGTGCGAGGGCATACAGTCCTCGCTTTTCACGACCGTAAGGTGGTCGTGGAATAAGAGCTGGGCAAGACCGGTGCCAGCCGCCGCGGTAATACCGGCAGCTCGAGTGATGACCGCTTTTATTGGGCCTAAAGCGTCCGTAGCCGGCCGGGTAAGTCCATCGGGAAATCCGCACGCTCAACGTGCGGGCGTCCGGTGGAAACTGCCTGGCTTGGGACCGGAAGATCCAGAGGGTACGTCTGGGGTAGGAGTGAAATCCCGTAATCCTGGACGGACCACCGGTGGCGAAAGCGCTCTGGAAAGACGGATCCGACGGTGAGGGACGAAAGCTCGGGTCA |
| p__Halobacteriota_g__Haladaptatus_ASV_245 | 1 | 1 | 1 | 1 | 0 | 0 | 0 | 1 | 1 | 1 | 1 | ACTTTACACTGCACGACAGTGCGATAAGGGAACTCCAAGTGCGAGGGCATATAGTCCTCGCTTTTGTGTATCGTAAGGTGGTACACGAATAAGAGCTGGGCAAGACCGGTGCCAGCCGCCGCGGTAATACCGGCAGCTCAAGTGATGGCCAATTTTATTGGGCCTAAAGCGTCCGTAGCCGGCCGAACAGGTCCGTCGGGAAATCTGCTCGCTCAACGAGCAGGCGTCCGACGGAAACCAGTCGGCTTGGGACCGGAAGACCCGAGGGGTACGTCTGGGGTAGGAGTGAAATCCTGTAATCCTAGACGGACCGCCGGTGGCGAAAGCGCCTCGGGAAGACGGATCCGACGGTGAGGGACGAAAGCTAGGGTCA |
| p__Halobacteriota_g__Saliphagus_ASV_246 | 1 | 1 | 1 | 0 | 0 | 0 | 0 | 0 | 0 | 0 | 1 | CCTTTACACTGCACGCCAGTGCGATAAGGGGACTCCAGGTGCGAGGGCATACAGTCCTCGCTTTTGTGTACCGTAGGGAGGTACACGAATAAGGGCTGGGCAAGACCGGTGCCAGCCGCCGCGGTAATACCGGCAGCCCGAGTGATGACCGCTATTATTGGGCCTAAAGCGTCCGTAGCCGGCCTCGCAAGTCCGTCGGGAAATCCGCACGCCTAACGTGCGGGCGTCCGGCGGAAACTGCGTGGCTTGGGACCGGAAGACTCGAGGGGTACGTCCGGGGTAGGAGTGAAATCCCGTAATCCTGGACGGACCACCGGTGGCGAAAGCGCCTCGAGAGGACGGATCCGACGGTGAGGGACGAAAGCTAGGGTCA |
| p__Halobacteriota_g__Halorussus_ASV_248 | 0 | 1 | 1 | 1 | 0 | 1 | 1 | 0 | 0 | 0 | 1 | ACTTTACACTGCACGACAGTGCGATAAGGGGACTCCGAGTGCGAGGGCATATAGTCCTCGCTTTTGTACACCGTAAGGTGGTGTACGAATAAGGGCTGGGCAAGACCGGTGCCAGCCGCCGCGGTAATACCGGCAGCCCGAGTGATGGCCGCTATTATTGGGCCTAAAGCGTCCGTAGCTGGCCAGACAGGTCCGTCGGGAAATCCATCCGCTTAACGGATGGGCGTCCGGCGGAAACCAGCTGGCTTGGGGCCGGAAGATCTGAGGGGTACGTCCGGGGTAGGAGTGAAATCCCGTAATCCTGGACGGACCGCCGGTGGCGAAAGCGCCTCAGAAGGACGGACCCGACGGTGAGGGACGAAAGCTAGGGTCA |
| p__Halobacteriota_g__Haladaptatus_ASV_250 | 1 | 0 | 0 | 0 | 0 | 1 | 1 | 1 | 1 | 1 | 1 | ACTTTACACTGCACGTCAGTGCGATAAGGGAACTCCAAGTGCGAGGGCATATAGTCCTCGCTTTTGTGTATCGTAAGGTGGTACACGAATAAGAGCTGGGCAAGACCGGTGCCAGCCGCCGCGGTAATACCGGCAGCTCGAGTGATGGCCAATTTTATTGGGCCTAAAGCGTCCGTAGCCTGCCAGACAGGTCCGTCGGGAAATCTGCTCGCTCAACGAGCAGGCGTCCGGCGGAAACCCGCTGGCTTGGGGCCGGAAGACCCAAGGGGTACGTCTGGGGTAGGAGTGAAATCCTGTAATCCTAGACGGACCACCGATCGCGAAAGCACCTTGGGAGGACGGACCCGACGGTGAGGGACGAAAGCTAGGGTCA |
| p__Thermoproteota_g__GCA-2726865_ASV_258 | 0 | 0 | 0 | 0 | 0 | 0 | 0 | 1 | 1 | 1 | 1 | CCTCCTCAATACACTAACGTGTGAAGGGGTTACCTCGAGTGCTATCTGAAAAAGGTGGCTTTTCCTTGGTCTAAAAAGCCTTGGGAATAAGAGGGGGGCAAGACTGGTGTCAGCCGCCGCGGTAACACCAGCCCCTCGAGTGGTGGGGATGATTATTTGGCCTAAAGCGTCCGTAGCCGGCCGAGTAAGTTTTCCGTTAAATCCAACGTCTCAAGCGTTGGACTGCGGGAAATACTGTTCGGCTAGAGAGTGGGAGAGGTAAGCGGTATTCTGGGGGTAGGGGCGAAATCCTTTGATCCCCGGAGGACCACCAGTGGCGAAGGCTGCTTACTAGAACACGTCTGACGGTCAGGGACGAAAGCTGGGGTAG |
| p__Halobacteriota_g__Natronoarchaeum_ASV_259 | 1 | 1 | 1 | 1 | 0 | 0 | 1 | 1 | 1 | 1 | 1 | CCTTTACACTGCACGCCAGTGCGATAAGGGGACCCCCAGTGCGAGGGCATACCGTCCTCGCTTTTCACCACCGTAAGGAGGTGGTGGAACAAGAGCTGGGCAAGACCGGTGCCAGCCGCCGCGGTAATACCGGCAGCTCGAGTGATGGCCGTTTTTATTGGGCCTAAAGCGTCCGTAGCCGGCCAGACAAGTTCGTCGGGAAATCCACGTGCCCAACACGTGGGCGTCCGGCGAAACCTCTCTGGCTTGGGACCGGACGGCCCGAGGGGTACGGTCGGGGTAGGAGTGAAATCCTGTAATCCCGACCGGACCACCGGTGGCGAAAGCGCCTCGGGAGGACGGATCCGACGGTGAGGGACGAAAGCCAGGGTCA |

| | Presence (1) or Absence (0) in each layer | | | | | | | | | | | |
|---|---|---|---|---|---|---|---|---|---|---|---|---|
| **Phylum_Genus_ASV_number** | **L1** | **L2** | **L3** | **L4** | **L5** | **L6** | **L7** | **L8** | **L9** | **L10** | **L11** | **Sequence** |
| p__Halobacteriota_g__Saliphagus_ASV_262 | 1 | 1 | 1 | 0 | 0 | 0 | 0 | 0 | 1 | 0 | 1 | CCTTTACACTGCACGACAGTGCGATAAGGGGACTCCAAGTGCGAGGGCATATCGTCCTCGCTTTTCACTACCGTAAGGTGGTGGTGGAATAAGGGCTGGGCAAGACCGGTGCCAGCCGCCGCGGTAATACCGGCAGCCCGAGTGATGACCGCTATTATTGGGCCTAAAGCGTCCGTAGCCGGCCACGCAAGTCCGTCGGGAAATCCGCGCGCCTAACGCGCGGGCGTCCGGCGGAAACTGCGTGGCTTGGGACCGGAAGACTCGAGGGGTACGTCCGGGGTAGGAGTGAAATCCTGTAATCCTGGACGGACCACCGGTGGCGAAAGCGCCTCGAGAGGACGGATCCGACGGTGAGGGACGAAAGCTAGGGTCA |
| p__Thermoproteota_g__BA2_ASV_263 | 0 | 0 | 0 | 0 | 0 | 0 | 0 | 0 | 1 | 1 | 1 | ACTCCGCAATGCACGAAAGTGTGACGGGGCTACCCCGAGTGCCGTCCGCTGAGGATGGCTTTTCCCCGGTGTAACTAGCCGGGGAATAAGGAGAGGGCAAGACTGGTGTCAGCCGCCGCGGTAATACCAGCTCTCCGAGTTGTAGGGATGATTATTGGGCTTAAAGCGTCCGTAGCCGGCTTAGCTAGTCTCCTGTTAAATTCAGCGACCTAATCGTTGAGCCGCGGGAGATACTACTAGGCTAGGAGGCGGGAGAGGTCGACGGTACTTTCGGGGTAGGGGTGAAATCCTATAATCCCGGGAGGACCACCAGTGGCGAAGGCTGTCGACTAGAACGCGCTCGACGGTGAGGGACGAAAGCTGGGGGAG |
| p__Halobacteriota_g__Saliphagus_ASV_264 | 0 | 1 | 1 | 1 | 1 | 0 | 1 | 0 | 1 | 1 | 1 | CCTTTACACTGCACGACAGTGCGATAGGGGGACTCCAAGTGCGAGGGCATATCGTCCTCGCTTTTCACCACCGTAAGGTGGTGGTGGAATAAGTGCTGGGCAAGACCGGTGCCAGCCGCCGCGGTAATACCGGCAGCACAAGTGATGACCGCTATTATTGGGCCTAAAGCGTCCGTAGCTGGCCGATCAAGTCCGTCGGGAAATCCGCACGCCTAACGTGCGGGCGTCCGGTGGAAACTGTTCGGCTTGGGACCGGAAGACTCGAGGGGTACGTCTGGGGTAGGAGTGAAATCCTGTAATCCTGGACGGACCACCGGTGGCGAAAGCGCCTCGAGAGGACGGATCCGACGGTGAGGGACGAAAGCTAGGGTCA |
| p__Thermoplasmatota_g__SM1-50_ASV_269 | 0 | 0 | 0 | 0 | 0 | 0 | 1 | 0 | 0 | 0 | 0 | CCTTCACACTGTGCGAAAGCGCGATGAGGGGATCCCAAGTGCTAACTCGAAAGAGTTGGCTGTTTATCTGTCTAAAACGCAGGTAGAGTAAGAGCTGGGTAAGACGGCTGCCAGCCGCCGCGGTAATAGCTGCAGCTCGAGTGGTGACCACTATTATTGGGCCTAAAACGTCCGTAGCCGGTCTAGTAAATGCCTGGGTAAATCGTGCCGCTTAACGGTACGAATTCCGGGTAGACTGCTAGACTTGAGATCGGGAGAGGTTAGAGGTACTCCTTGGGTAGAGGTGAAATTCTGTAATCCTAGGGGGACCACCTGTGGCGAAAGCGTCTAACCAGAACGAGTCTGACGGTGAGGGACGAAGCCCTGGGTCA |
| p__Thermoplasmatota_g__SM1-50_ASV_270 | 0 | 0 | 0 | 0 | 0 | 0 | 0 | 1 | 1 | 1 | 1 | CCTTCACAATGTGCGAAAGCACGATGAGGGAATCCCAAGTGCTTGCACGTAAGTGTAAGCTGTTTCTTTGTCTAAAAAGCATTGAGAGTAAGGGCTGGGTAAGACGGGTGCCAGCCGCCGCGGTAATACCTGCAGCTCAAGTGGTGATCGCTATTATTGGGCCTAAAACGTTCGTAGCCGGTTTAGTAAATGCCTGGGTAAATCGTGCCGCTTAACGGTACGAATTCCGGGTAGACTGCTAAACTTGAGACCGGGAGAGGTTAGAGGTACTCCTGGGGTAGAGGTGAAATTCTGTAATCCTAGGGGGACCACCAGTGGCGAAGGCGTCTAACTAGAACGGGTCTGACGGTGAGGGACGAAGCCCTGGGGCG |
| p__Halobacteriota_g__Halohasta_ASV_272 | 1 | 1 | 1 | 1 | 0 | 0 | 1 | 1 | 1 | 1 | 1 | CCTTTACACTGCACGCACGTGCGATAGGGGAATCCCAAGTGCGAGGGCATACTAGCCCTCGCTTTTGTCTACCCTAAGGCGGTAGACGAATAAGAGCTGGGCAAGACCGGTGCCAGCCGCCGCGGTAATACCGGCAGCTCGAGTGATGTCCACTATTATTGGGCCTAAAGCGTCCGTAGCTGGCCGACCAAGTCCGTCGGGAAATCCACCTGCCCAACAGGTGGGCGTCCGGCGGAAACTGGCCGGCTTGGAACCGGAAGGCTCAGAGGGTACGTCCGGGGTAGGAGTGAAATCCCGTAATCCCGGACGGACCACCGATGGCGAAAGCACTCTGAGAGAACGGATTCGACAGTGAGGGACGAAAGCTAGGGTCT |
| p__Halobacteriota_g__Halopiger_A_ASV_274 | 1 | 1 | 1 | 0 | 1 | 1 | 1 | 0 | 1 | 0 | 0 | CCTTTACACTGCACGCCAGTGCGATAAGGGGACTCCATCTGCGAGGGCATATCGTCCTCGCTTTTCTCGACCGTAAGGTGGTCGAGGAATAAGGGCTGGGCAAGACCGGTGCCAGCCGCCGCGGTAATACCGGCAGCCCGAGTGATGACCGCTATTATTGGGCCTAAAGCGTCCGTAGCTGGCCAGGCAAGTCCATCGGGAAATCCGCGTGCCCAACACGCGGGCGTCCGGTGGAACCTGCCTGGCTTGGGACCGGAAGACCAGAGGGGTACGTCTGGGGTAGGAGTGAAATCCCGTAATCCTGGACGGACCACCGGTGGCGAAAGCGCCTCTGGAAGACGGATCCGACGGTGAGGGACGAAAGCTCGGGTCA |
| p__Halobacteriota_g__Haladaptatus_ASV_277 | 0 | 1 | 1 | 1 | 0 | 0 | 0 | 1 | 1 | 1 | 1 | ACTTTACACTGCACGACAGTGCGATAAGGGAACTCCAAGTGCGAGGGCATATAGTCCTCGCTTTTGTGTATCGTAAGGTGGTACACGAATAAGAGCTGGGCAAGACCGGTGCCAGCCGCCGCGGTAATACCGGCAGCTCAAGTGATGGCCAATTTTATTGGGCCTAAAGCGTCCGTAGCCTGCCAGACAGGTCCGTCGGGAAATCTGCTCGCTCAACGAGCAGGCGTCCGGCGGAAACCAGCTGGCTTGGGGCCGGAAGACCCAAGGGGTACGTCTGGGGTAGGAGTGAAATCCTGTAATCCTAGACGGACCACCGATCGCGAAAGCACCTTGGGAGGACGGACCCGACGGTGAGGGACGAAAGCTAGGGTCA |
| p__Halobacteriota_g__Halalkalicoccus_ASV_279 | 0 | 1 | 1 | 0 | 0 | 0 | 0 | 0 | 0 | 0 | 0 | CCTTTACACTGCACGAACGTGCGATAAGGGGACTCCAAGTGCGAGGGCATATAGTCCTCGCTTTTCCCTACCGTAAGGTGGTAGGTGAATAAGAGCTGGGCAAGACCGGTGCCAGCCGCCGCGGTAATACCGGCAGCTCGAGTGATGGCCAATATTATTGGGCCTAAAGCGTCCGTAGCTGGCCGAACAGGTCTGTCGGGAAATCCGCTCGCTCAACGAGCGGACGTCCGGCGGAAACCAGTCGGCTTGGGACCGGAAGACCCGAGGGGTACGTCCGGGGTAGGAGTGAAATCCTGTAATCCTGGACGGACCACCGGTGGCGAAAGCGCCTCGGGAGGACGGATCCGACAGTGAGGGACGAAAGCTTGGGTCA |
| p__Halobacteriota_g__Saliphagus_ASV_276 | 1 | 1 | 1 | 1 | 1 | 0 | 0 | 0 | 0 | 1 | 1 | CCTTTACACTGCACGAAAGTGCGATAAGGGGATCCCGAGTGCGAGGGCATACAGTCCTCGCTTTTCTGTACCGTAAGGTGGTACAGGAATAAGGGCTGGGCAAGACCGGTGCCAGCCGCCGCGGTAATACCGGCAGCTCAAGTGATGGCCGCTATTATTGGGCCTAAAGCGTCCGTAGCCGGCCGCACAAGTCCGTCGGGAAATCCACCCGCCTAACGGGTGGGCGTCCGGCGGAAACTGTGCGGCTTGGGACCGGAAGACTCGAGGGGTACGTCCGGGGTAGGAGTGAAATCCTGTAATCCTGGACGGACCGCCGGTGGCGAAAGCGCCTCGAGAGGACGGATCCGACGGTGAGGGACGAAAGCTAGGGTCT |
| p__Thermoplasmatota_g__EX4484-6_ASV_286 | 0 | 1 | 0 | 1 | 1 | 1 | 1 | 1 | 0 | 0 | 1 | ACTTCGCAATGCGCGCAAGCGCGACGAGGGGATTCCAAGTGCATGCACACAGTGTATGCTGTTCAATAGTCTAAAACGCTATTGGAGTAAGGGCTGGGTAAGACTGGTGCCAGCCGCCGCGGTAATACCAGCGGCCCTAGTGGTGATCGTTATTATTGGGCCTAAAGCGTCCGTAGCCGGCTCGGTAAATCTCTGGGTAAATCGTTGGGCTTAACTCAGCGAATTCTGGGGAGACTGCCGGGCTTGGGACCGGGAGAGGTCGGAGGTACTCCAGGGGTAGGGGTGAAATCCTGTAATCCTTGGGGGACCACCGGTGGCGAAAGCGTCCGGCCAGAACGGGTCCGACGGTAAGGGACGAAGCCCTGGGGCG |
| p__Halobacteriota_g__Halodesulfurarchaeum_ASV_288 | 1 | 1 | 1 | 1 | 0 | 0 | 1 | 1 | 0 | 0 | 1 | CCTTTACACTGCACGACAGTGCGATAAGGGGACTCCGAGTGCGAGGGCATAGAGCCCTCGCTTTTCTGCACCGTAAGGTGGTGCAGGAAGAAGGGCTGGGCAAGACCGGTGCCAGCCGCCGCGGTAATACCGGCAGCCCAAGTGATGGCCGATATTATTGGGCCTAAAGCGTCCGTAGCCGGCCGGACAAGTCCGTTGGGAAAGCCGCACGCTCAACGTGCGGGAGTCCAGCGGAAACTGTCCGGCTTGGGACCGGAAGACCCGAGGGGTACGTCCGGGGTAGGAGTGAAATCCTGTAATCCCGGACGGACCACCTGTGGCGAAAGCGCCTCGGGAAGACGGATCCGACGGTGAGGGACGAAAGCCAGGGTCT |
| p__Thermoplasmatota_g__UBA147_ASV_296 | 1 | 1 | 0 | 1 | 0 | 0 | 0 | 1 | 1 | 1 | 1 | ACTTGTCAATGGGGGCAACCCCGAACAGGGGATTCCAAGTGCATGCGTTTTACGTATGCTGTTCGGTTGTCTAAAAAACAGCCGAAGTAAGGGCTGGGTAAGACAGGTGCCAGCCGCCGCGGTAATACCTGCAGCCCAAGTGGTAATCGTTATTATTGAGCCTAAAGCGTTCGTAGCCGGTTTGGTAAACCCTCGGGTAAATCGAGCAGCTCAACTGTTCGAATTCCGGGGGGACTGCCAGACTTGGGACCGGGAGAGGTCAGAGGTACTCCGGGGGTAGGGGCTAAATCCTGTAATCCTTGGAGGACCACCAGTGGCGAAGGCGTCTGACTAGAACGGATCCGACGGTAAGGAACGAAACCCTGGGGCG |
| p__Halobacteriota_g__Halostagnicola_ASV_297 | 1 | 1 | 1 | 1 | 0 | 0 | 1 | 0 | 0 | 0 | 0 | CCTTTACACTGCACGCGAGTGCGATAAGGGGACTCCAAGTGCGAGGGCATATAGTCCTCGCTTTTTCCGACCGTAGGATGGTCGGCGAATAAGTGCTGGGCAAGACCGGTGCCAGCCGCCGCGGTAATACCGGCAGCACGAGTGATGACCGCTATTATTGGGCCTAAAGCGTCCGTAGCTGGCCACGCAAGTCCATCGGGAAATCTCCTCGCTCAACGGGGAGGCGTCCGGTGGAAACTGTGTGGCTTGGGACCGGAAGATCCAGAGGGTACGTCTGGGGTAGGAGTGAAATCCCGTAATCCGAGACGGACCACCGGTGGCGAAAGCGCTCTGGAAAGACGGATCCGACGGTGAGGGACGAAAGCTCGGGTCA |
| p__Halobacteriota_g__Halovenus_ASV_301 | 0 | 1 | 1 | 1 | 0 | 0 | 0 | 0 | 0 | 0 | 1 | CCTTTACACTGCACGACAGTGCGATAGGGGGACTCCGAGTGCGAGGGCATACAGTCCTCGCTTTTCTGTACCGTAAGGTGGTACAGGAACAAGTGCTGGGCAAGACCGGTGCCAGCCGCCGCGGTAATACCGGCAGCACAAGTGATGACCGATTTTATTGGGCCTAAAGCGTCCGTAGCTGGCCAGACAAGTCTGTTGGGAAATCCACGCGCCCAACGCGTGGGCGTCCAGCGGAAACTGTCTGGCTTGGGGCCGGAGGACTCGAGGGGTACGTCCGGGGTAGGAGTGAAATCCCGTAATCCTGGACGGACCACCGGTGGCGAAAGCGCCTCGAGAGGACGGACCCGACAGTGAGGGACGAAAGCCAGGGTCT |
| p__Halobacteriota_f__Haloferacaceae_ASV_302 | 1 | 1 | 1 | 0 | 0 | 0 | 0 | 0 | 1 | 1 | 1 | CCTTTACACTGCACGCAAGTGCGATAAGGGGACTCCGAGTGCGAGGGCATATAGTCCTCGCTTTTCTGTACCGTAGGGCGGTACAGGAACAAGAGCTGGGCAAGACCGGTGCCAGCCGCCGCGGTAATACCGGCAGCTCGAGTGATGACCGATTTTATTGGGCCTAAAGCGTCCGTAGCTGGCCAGACAAGTCCGTCGGGAAATCCATCTGCTCAACAGATGGGCGTCCGGCGGAAACTCTCTGGCTTGGGACCGGAAGGCTCGAAGGGTACGTCCGGGGTAGGAGTGAAATCCCGTAATCCTGGACGGACCACCGATGGCGAAAGCACTTCGAGAAGACGGATCCGACAGTGAGGGACGAAAGCTAGGGTCT |
| p__Halobacteriota_f__Haloferacaceae_ASV_304 | 0 | 1 | 1 | 1 | 0 | 1 | 1 | 0 | 0 | 1 | 1 | CCTTTACACTGCGCGCAAGCGCGATAAGGGGACCCCAAGTGCGAGGGCATACAGTCCTCGCTTTTCACATCCGTAGGGAGGATGTGGAATAAGGGCTGGGCAAGACCGGTGCCAGCCGCCGCGGTAATACCGGCAGCCCAAGTGATGGCCGCTCTTATTGGGCCTAAAGCGTCCGTAGCTGGCCACGTGTGTCCGTCGGGAAATCCACCAGCCCAACTGGTGGGCGGCCGGCGGAAACTGCGTGGCTTGGGACCGGAAGACCCACCGGGTACGTCCGGGGTAGGAGTGAAATCCCGTAATCCCGGACGGACCACCGATGGCGAAAGCACGGTGGGAAGACGGCTCCGACAGTGAGGGACGAAAGCTGGGGTCT |
| p__Halobacteriota_g__Halovenus_ASV_305 | 0 | 1 | 1 | 1 | 0 | 0 | 0 | 0 | 0 | 0 | 1 | CCTTTACACTGCACGCCAGTGCGATAGGGGGACTCCGAGTGCGAGGGCATATAGTCCTCGCTTTTCTGTACCGTAAGGTGGTACAGGAACAAGTGCTGGGCAAGACCGGTGCCAGCCGCCGCGGTAATACCGGCAGCACAAGTGATGGCCGATATTATTGGGCCTAAAGCGTCCGTAGCCGGCCGGACAAGTCTGTTGGGAAATCCACGCGCTTAACGCGTGGGCGTCCAGCGGAAACTGTCTGGCTTGGGGCCGGAAGACTCGAGGGGTACGTCCGGGGTAGGAGTGAAATCCCGTAATCCTGGACGGACCGCCGGTGGCGAAAGCGCCTCGAGAGGACGGACCCGACGGTGAGGGACGAAAGCCAGGGTCT |
| p__Halobacteriota_f__Haloferacaceae_ASV_307 | 1 | 1 | 1 | 0 | 1 | 1 | 0 | 0 | 1 | 1 | 1 | CCTTTACACTGCACGCCAGTGCGATAAGGGGACTCCGAGTGCGAGGGCATATCGTCCTCGCTTTTCTGTACCGTAGGGAGGTACAGGAACAAGAGCTGGGCAAGACCGGTGCCAGCCGCCGCGGTAATACCGGCAGCTCGAGTGATGGCCGATCTTATTGGGCCTAAAGCGTCCGTAGCCGGCCAGACAAGTCCGTCGGGAAATCCACCTGCTCAACAGGTGGGCGTCCGGCGGAAACTCTCTGGCTTGGGACCAGAAGACCCGAGGGGTACGTCCGGGGTAGGAGTGAAATCCCGTAATCCTGGACGGACCACCGATAGCGAAAGCACCTCGGGAAGATGGATCCGACGGTGAGGGACGAAAGCTGGGGTCT |
| p__Halobacteriota_g__Halohasta_ASV_310 | 0 | 1 | 1 | 1 | 0 | 0 | 0 | 0 | 0 | 0 | 1 | CCTTTACACTGCACGCAAGTGCGATAGGGGAATCCCAAGTGCGAGGGCATACTAGCCCTCGCTTTTGTTGACTGTAAGGTGGTCAACGAATAAGAGCTGGGCAAGACCGGTGCCAGCCGCCGCGGTAATACCGGCAGCTCGAGTGATGTCCACTATTATTGGGCCTAAAGCGTCCGTAGCTGGCCAGCCAAGTTCGTCGGGAAATCCACCTGCCCAACAGGTGGGCGTCCGGCGAAAACTGTCTGGCTTGGAACCGGAAGGCTCAGAGGGTACGTCCGGGGTAGGAGTGAAATCCCGTAATCCCGGACGGACCACCGATGGCGAAAGCACTCTGAGAGGACGGATTCGACAGTGAGGGACGAAAGCTAGGGTCT |
| p__Halobacteriota_g__Halovivax_ASV_311 | 1 | 1 | 1 | 1 | 0 | 1 | 1 | 0 | 0 | 0 | 1 | CCTTTACACTGCACGCGAGTGCGATAGGGGGACTCCAAGTGCGAGGGCATACAGTCCTCGCTTTTCACCACCGTAAGGAGGTGGTAGAATAAGTGCTGGGCAAGACCGGTGCCAGCCGCCGCGGTAATACCGGCAGCACGAGTGATGACCGCTATTATTGGGCCTAAAGCGTCCGTAGCCGGCCGGGCAAGTCCATCGGGAAATCCGCACGCCTAACGTGCGGGCGTCCGGTGGAAACTGTTCGGCTTGGGACCGGAAGATCCAGAGGGTACGTCTGGGGTAGGAGTGAAATCCTGTAATCCTGGACGGACCACCGGTGGCGAAAGCGCTCTGGAAAGACGGATCCGACGGTGAGGGACGAAAGCTTGGGTCA |
| p__Halobacteriota_g__Halohasta_ASV_313 | 0 | 1 | 1 | 1 | 0 | 0 | 0 | 0 | 0 | 0 | 0 | CCTTTACACTGCACGCAAGTGCGATAGGGGAATCCCAAGTGCGAGGGCATACTAGCCCTCGCTTTTGTGTACTGTAAGGAGGTACACGAATAAGAGCTGGGCAAGACCGGTGCCAGCCGCCGCGGTAATACCGGCAGCTCAAGTGATGTCCACTATTATTGGGCCTAAAGCGTCCGTAGCTGGCCGCGCAAGTCCGTCGGGAAATCCACCTGCCCAACAGGTGGGCGTCCGGCGGAAACTGCGTGGCTTGGAACCGGAAGGCTCAGAGGGTACGTCCGGGGTAGGAGTGAAATCCCGTAATCCCGGACGGACCACCGATGGCGAAAGCACTCTGAGAGGACGGATTCGACAGTGAGGGACGAAAGCTAGGGTCT |
| p__Halobacteriota_g__Halomicrobium_3849_ASV_316 | 0 | 1 | 1 | 0 | 1 | 1 | 1 | 0 | 0 | 0 | 1 | CCTTTACACTGCACGACAGTGCGATAGGGGGACTCCGAGTGCGAGGGCATATAGTCCTCGCTTTTCTGGATCGTAAGGAGATCCAGGAACAAGTGCTGGGCAAGACCGGTGCCAGCCGCCGCGGTAATACCGGCAGCACGAGTGATGGCCGCTCTTATTGGGCCTAAAGCGTCCGTAGCCGGCCAGTCAAGTCCGTTGGGAAATCGGCGCGCCTAACGCGTCGGCGTCCAGCGGAAACTGACTGGCTTGGGGCCGGAAGACCTGAGGGGTACGTCCGGGGTAGGAGTGAAATCCCGTAATCCTGGACGGACCACCGGTGGCGAAAGCGCCTCAGGAAGACGGACCCGACGGTGAGGGACGAAAGCTAGGGTCT |
| p__Halobacteriota_g__Halorientalis_ASV_321 | 0 | 1 | 1 | 1 | 0 | 0 | 0 | 0 | 0 | 0 | 0 | CCTTTACACTGCACGACAGTGCGATAGGGGGACTCCGAGTGCGAGGGCATACAGTCCTCGCTTTTCTGTACCGTAAGGTGGTACAGGAACAAGTGCTGGGCAAGACCGGTGCCAGCCGCCGCGGTAATACCGGCAGCACGAGTGATGGCCGCTCTTATTGGGCCTAAAGCGTCCGTAGCCGGCCACGCAAGTTCGTCGGGAAATCTGCCCGCTTAACGGGCAGGCGTCCGGCGAAAACTGCGTGGCTTGGGGCCGGGAGATCCAAGGGGTACGTCCTGGGTAGGAGTGAAATCCTGTAATCCTGGACGGACCACCGATGGCGAAAGCACCTTGGAAAAACGGACCCGACGGTGAGGGACGAAAGCTAGGGTCT |
| p__Halobacteriota_o__Halobacteriales_ASV_325 | 0 | 1 | 1 | 1 | 1 | 1 | 0 | 0 | 1 | 1 | 1 | CCTTTACACTGCACGACAGTGCGATAAGGGGACCCCGAATGCGAGGGCATACAGTCCTCGCTTTTGTGCACTGTAGGGAGGTGCACGAATAAGAGCTGGGCAAGACCGGTGCCAGCCGCCGCGGTAATACCGGCAGCTCGAGTGATGGCCGCTGTTATTGGGCCTAAAGCGTCCGTAGCTGGCCGCACAAGTCCGTCGGGAAATCCACCCGCCTAACGGGTGGGCGTCCGGCGGAAACTGTGCGGCTTGGGACCGGAAGACCCGAGGGGTACGTCCGGGGTAGGAGTGAAATCCTGTAATCCTGGACGGACCGCCGGTGGCGAAAGCGCCTCGGGAAGACGGATCCGACAGTGAGGGACGAAAGCTAGGGTCT |

| Phylum_Genus_ASV_ number | Presence (1) or Absence (0) in each layer L1 | L2 | L3 | L4 | L5 | L6 | L7 | L8 | L9 | L10 | L11 | Sequence |
|---|---|---|---|---|---|---|---|---|---|---|---|---|
| p__Halobacteriota_f__Haloarculaceae_3992_ASV_329 | 0 | 1 | 1 | 1 | 0 | 0 | 1 | 0 | 0 | 0 | 1 | CCTTTACACTGCACGCCAGTGCGATAGGGGGACTCCGAGTGCGAGGGCATATAGTCCTCGCTTTTGTACACCGTAAGGTGGTGTACGAATAAGAACTGGGCAAGACCGGTGCCAGCCGCCGCGGTAATACCGGCAGTTCGAGTGATGGCCGATATTATTGGGCCTAAAGCGTTCGTAGCTGGCCAGTCAAGTCTGTCGGGAAATCCAGCGGCTCAACCGCTGGGCGTCCGGCGGAAACTGATTGGCTTGGGACCGGGAGACCCGAAGAGTACGTCCGGGGTAGGAGTGAAATCCCGTAATCCTGGACGGACTACCGATGGCGAAAGCACTTCGGGAGCACGGATCCGACAGTGAGGAACGAAAGCTGGGGTCT |
| p__Halobacteriota_g__Halalkalicoccus_ASV_327 | 1 | 1 | 1 | 0 | 1 | 1 | 0 | 0 | 1 | 0 | 1 | CCTTTACACTGCACGAACGTGCGATAAGGGGACTCCAAGTGCGAGGGCATACAGTCCTCGCTTTTCCCTACCGTAAGGTGGTAGGTGAATAAGAGCTGGGCAAGACCGGTGCCAGCCGCCGCGGTAATACCGGCAGCTCGAGTGATGGCCAATATTATTGGGCCTAAAGCGTCCGTAGCTGGCCGAACAGGTCCGTCGGGAAATCCGCTCGCTCAACGAGCGGACGTCCGGCGGAAACCAGTCGGCTTGGGACCGGAAGACCCGAGGGGTACGGCCGGGGTAGGAGTGAAATCCTGTAATCCTGGCCGGACCACCGGTGGCGAAAGCGCCTCGGGAAGACGGATCCGACAGTGAGGGACGAAAGCTAGGGTCA |
| p__Halobacteriota_g__Halorussus_ASV_328 | 1 | 1 | 1 | 1 | 1 | 0 | 0 | 0 | 0 | 0 | 1 | CCTTTACACTGCACGAAAGTGCGATAGGGGGACCCCGAGTGCGAGGGCATACAGTCCTCGCTTTTGTGTACCGTAAGGAGGTACACGAATAAGGGCTGGGCAAGACCGGTGCCAGCCGCCGCGGTAATACCGGCAGCTCGAGTGATGGCCGCTGTTATTGGGCCTAAAGCGTCCGTAGCCGGCCGCACAAGTCCGTCGGGAAATCCACCCGCCTAACGGGTGGGCGTCCGGCGGAAACTGTGCGGCTTGGGGCCGGAAGATCCGAGGGGTACGTCCGGGGTAGGAGTGAAATCCCGTAATCCTGGACGGACCGCCGGTGGCGAAAGCGCCTCGGAAGGACGGACCCGACGGTGAGGGACGAAAGCTAGGGTCT |
| p__Thermoplasmatota_g__SM1-50_ASV_331 | 1 | 1 | 0 | 0 | 0 | 0 | 0 | 1 | 0 | 0 | 1 | ACTTCACACTGCGCGAAAGCGCGATGAGGGAATCCCAAGTGTATGCACGTAAGTGTATACTGTTTCTATGTTTAAAAAGCATTGAGAGTAAGGGCTGGGTAAGACGGGTGCCAGCCGCCGCGGTAATACCTGCAGCTCGAGTGGTGATCACTATTATTGGGCCTAAAACGTCCGTAGCCGGTCTAGTAAATGCCTGGGTAAATCGTACGGCTTAACCGTACGAATTCTGGGTAGACTGCTAAACTTGAGACCGGGAGAGGCTAGAGGTACTCCTGGGGTAGAGGTGAAATTCTGTAATCCTAGGGGGACCACCAGTGGCGAAGGCGTCTAGCTAGAACGGGTCTGACGGTGAGGGACGAAGCCCTGGGGCG |
| p__Thermoproteota_g__BA2_ASV_337 | 1 | 1 | 0 | 0 | 0 | 0 | 1 | 1 | 0 | 1 | 1 | ACTCCGCAATGCACGAAAGTGTGACGGGGCTACCCCGAGTGCCGTCCGCTGAGGATGGCTTTTCTTTGGTTTAACTAGCCAGGGGAATAAGGAGAGGGCAAGACTGGTGTCAGCCGCCGCGGTAATACCAGCTCTCCGAGTGGTAGGGATGATTATTGGGCTTAAAGCGTCCGTAGCCGGCTTAGCAAGTCTCCTGTTAAACTCAGCGACTCAATCGTTGAATCGCGGGAGATACTACTGGGCTAGGAGGCGGGAGAGGTCGACGGTACTTCTGGGGTAGGGGTGAAATCCTATAATCCCAGGAGGACCACCAGTGGCGAAGGCTGTCGACTAGAACGCGCTCGACGGTGAGGGACGAAAGCTGGGGGAG |
| p__Thermoplasmatota_g__Natronoplasma_ASV_341 | 1 | 1 | 0 | 0 | 0 | 0 | 0 | 0 | 0 | 0 | 1 | CCTTCACACTGTGGGAAACCACGATGAGGGGATTCCAAGTGCCAACACTTTGTGTTGGCTTTTCACCTATGTAAAAAGTAGGTGGAATAAGGGCCGGGTAAGACGGGTGCCAGCCGCCGCGGTAACACCCGCGGCCCAAGTAGTGGTCATTAATATTGAGCTTAAAGCGTCCGTAGCCGGTCTGGTAAATCTCTGGGTAAATCGAACGGCTCAACCGTTCGAATTCCAGAGAGACTGTCAGACTTGGGGCCGGGAGAGGTCAGAGGTACTCCTGGGGTAGAAGTAAAATTCTGTAATCCTAGGAGGACCACCAGTGGCGAAAGCGTCTGACTAGAACGGACTCGACGGTGAGGGACGAAGCCCTGGGGCG |
| p__Thermoproteota_g__Nitrososocosmicus_ASV_346 | 0 | 0 | 0 | 0 | 1 | 1 | 1 | 0 | 1 | 0 | 1 | CCTCTGCAATAGGCGAAAGCTTGACAGGGTTACTCTGAGTGATTTCCGTTAAGGAGATCTTTTGGCACCTCTAAAAATGGTGCAGAATAAGGGGTGGGCAAGTCTGGTGTCAGCCGCCGCGGTAATACCAGCACCCCGAGTGGTCGGGACGTTTATTGGGCCTAAAGCATCCGTAGCCGGTTCTACAAGTCTTCCGTTAAATCCACCTGCTCAACAGATGGGCTGCGGAAGATACTATGGAGCTAGGAGGCGGGAGAGGCAAGCGGTACTCGATGGGTAGGGGTAAAATCCGTTGATCCATTGAAGACCACCAGTGGCGAAGGCGGCTTGCCAGAACGCGCTCGACGGTGAGGGATGAAAGCTGGGGGAG |
| p__Halobacteriota_g__Haloprofundus_ASV_356 | 0 | 1 | 1 | 0 | 1 | 0 | 0 | 0 | 0 | 0 | 0 | CCTTTACACTGCACGCCAGTGCGACAAGGGGACCCCGAGTGCGAGGGCATACAGTCCTCGCTTTTCCCGACCGTAAGAAGGTCGGAGAAGAAGAGCTGGGCAAGACCGGTGCCAGCCGCCGCGGTAATACCGGCAGCTCGAGTGATGACCGCTGTTATTGGGCCTAAAGCGTCCGTAGCTGGCCGCGTAAGTCCATCGGGAAATCCATCCGCTCAACGGATGGGCGTCCGGTGGAAACTACACGGCTTGGAGCCGGAAGACCCAAGGGGTACGTCCGGGGTAGGAGTGAAATCCCGTAATCCTGGACGGACCGCCGATGGCGAAAGCACCTTGGGAAGACGGACTCGACAGTGAGGGACGAAAGCTAGGGTCT |
| p__Thermoplasmatota_g__SM1-50_ASV_357 | 0 | 0 | 0 | 1 | 0 | 0 | 0 | 0 | 1 | 0 | 0 | ACTTCACACTGCGCGAAAGCGCGATGAGGGAATCCCAAGTGTATGCACGTAAGTGTATACTGTTTTTATGTCTAAAAAGCATAGAGAGTAAGAGCTGGGTAAGACGGGTGCCAGCCGCCGCGGTAATACCTGCAGCTCAAGTGGTGGCCGCTATTATTGGGCCTAAAACGTCCGTAGCCGGTCTTGTAAATGCCTGGGTAAATCGTGTCGCTTAACGATACGAATTCCGGGTAGACTGTAAGACTTGAGACCGGGAGAGGCTAGAGGTACTCCTGGGGTAGAGGTGAAATTCTGTAATCCTAGGGGGACCACCAGTGGCGAAGGCGTCTAGCTAGAACGGGTCTGACGGTGAGGGACGAAGCCCTGGGGCG |
| p__Halobacteriota_g__Halorubrum_ASV_359 | 0 | 0 | 1 | 0 | 1 | 1 | 0 | 0 | 0 | 0 | 0 | CCTTTACACTGCACGACAGTGCGATAGGGGGATCCCAAGTGCACAGGCATAGCGCCTGTGCTTTTCGGTACCGTAAGGTGGTACCAGAATAAGGGCTGGGCAAGACCGGTGCCAGCCGCCGCGGTAATACCGGCAGCCCAAGTGATGGCCGATCTTATTGGGCCTAAAGCGTCCGTAGCTGGCCGCGCAAGTCCATCGGGAAATCCACCTGCTCAACAGGTGGGCGCCCGGTAGAAACTGCGTGGCTTGGGACCGGAAGGCGCGACGGGTACGTCCGGGGTAGGAGTGAAATCCCGTAATCCTGGACGGACCGCCGATGGCGAAAGCACGTCGCGAGAACGGATCCGACAGTGAGGGACGAAAGCCAGGGTCT |
| p__Halobacteriota_g__Haladaptatus_ASV_362 | 0 | 1 | 1 | 0 | 1 | 1 | 0 | 0 | 0 | 0 | 0 | ACTTTACACTGCACGCCAGTGCGATAAGGGAACTCCAAGTGCGAGGGCATATAGTCCTCGCTTTTGTGTATCGTAAGGTGGTACACGAATAAGAGCTGGGCAAGACCGGTGCCAGCCGCCGCGGTAATACCGGCAGCTCAAGTGATGGCCAATATTATTGGGCCTAAAGCGTCCGTAGCCGGCCGAACAGGTCCGTCGGGAAATCTGCTCGCTCAACGAGCAGGCGTCCGACGGAAACCAGTCGGCTTGGGACCGGAAGACCCGAGGGGTACGTCTGGGGTAGGAGTGAAATCCTGTAATCCTAGACGGACCGCCGGTGGCGAAAGCGCCTCGGGAAGACGGATCCGACGGTGAGGGACGAAAGCTAGGGTCA |
| p__Halobacteriota_g__Halalkalicoccus_ASV_360 | 1 | 1 | 1 | 0 | 1 | 1 | 0 | 0 | 0 | 0 | 0 | CCTTTACACTGCACGAACGTGCGATAAGGGGACTCCAAGTGCGAGGGCATACAGTCCTCGCTTTTCTGTACCGTAAGGTGGTACAGGAATAAGAGCTGGGCAAGACCGGTGCCAGCCGCCGCGGTAATACCGGCAGCTCGAGTGATGGCCAATATTATTGGGCCTAAAGCGTCCGTAGCTGGCCGAACAGGTCCGTCGGGAAATCCGCTCGCTCAACGAGCGGACGTCCGGCGGAAACCAGTCGGCTTGGGACCGGAAGACCCGAGGGGTACGGCCGGGGTAGGAGTGAAATCCTGTAATCCTGGCCGGACCACCGGTGGCGAAAGCGCCTCGGGAAGACGGATCCGACAGTGAGGGACGAAAGCTAGGGTCA |
| p__Halobacteriota_g__Haloprofundus_ASV_364 | 1 | 1 | 1 | 0 | 0 | 0 | 0 | 1 | 0 | 0 | 0 | CCTTTACACTGCACGACAGTGCGACAAGGGGACCCCGAGTGCGAGGGCATACAGTCCTCGCTTTTGTGAACCGTAAGGAGGTTCACGAATAAGAGCTGGGCAAGACCGGTGCCAGCCGCCGCGGTAATACCGGCAGCTCGAGTGATGACCGCTGTTATTGGGCCTAAAGCGTCCGTAGCTGGCCGCGTAAGTCCATCGGGAAATCCATCCGCTCAACGGATGGGCGTCCGGTGGAAACTACACGGCTTGGAGCCGGAAGACCCAAGGGGTACGTCCGGGGTAGGAGTGAAATCCCGTAATCCTGGACGGACCGCCGATGGCGAAAGCACCTTGGGAAAACGGACTCGACAGTGAGGGACGAAAGCTAGGGTCT |
| p__Halobacteriota_g__Halohasta_ASV_365 | 1 | 0 | 0 | 0 | 0 | 0 | 0 | 0 | 0 | 0 | 1 | CCTTTACACTGCACGCAAGTGCGATAGGGGAATCCCAAGTGCGAGGGCATACTAGCCCTCGCTTTTGTGTACTGTAAGGAGGTACACGAATAAGAGCTGGGCAAGACCGGTGCCAGCCGCCGCGGTAATACCGGCAGCTCGAGTGATGTCCACTATTATTGGGCCTAAAGCGTCCGTAGCTGGCCGCGCAAGTCCGTCGGGAAATCCACCTGCCCAACAGGTGGGCGTCCGGCGGAAACTGCGTGGCTTGGAACCGGAAGGCTCAGAGGGTACGTCCGGGGTAGGAGTGAAATCCCGTAATCCCGGACGGACCACCGATGGCGAAAGCACTCTGAGAGAACGGATTCGACAGTGAGGGACGAAAGCTAGGGTCT |
| p__Thermoproteota_g__BA2_ASV_371 | 0 | 1 | 0 | 0 | 0 | 0 | 0 | 1 | 1 | 1 | 1 | ACTCCGCAATGCACGAAAGTGTGACGGGGCTACCCCGAGTGCCATCCGCTGAGGATGGCTTTTCTTTGGTATAACTAGCCATTGGAATAAGGAGAGGGCAAGACTGGTGTCAGCCGCCGCGGTAATACCAGCTCTCCGAGTGGTAGGGATGATTATTGGGCTTAAAGCGTCCGTAGCCGGCTTAGCAAGTTTCCTGTTAAACTCAGCGACTCAATCGTTGGATCGCGGGAAATACTACTAGGCTAGGAGGCGGGAGAGGTCGACGGTACTTCTGGGGTAGGGGTGAAATCCTATAATCCCGGGAGGACCACCAGTGGCGAAGGCTGTCGACTAGAACGCGCTCGACGGTGAGGGACGAAAGCTGGGGGAG |
| p__Halobacteriota_f__Natrialbaceae_ASV_374 | 0 | 0 | 1 | 1 | 0 | 0 | 1 | 0 | 0 | 0 | 0 | CCTTTACACTGCACGCCAGTGCGATAAGGGGATACCAAGTGCGAGGGCATATCGTCCTCGCTTTTCACCACCGTAAGGAGGTGGTAGAATAAGAGCTGGGCAAGACCGGTGCCAGCCGCCGCGGTAATACCGGCAGCTCGAGTGATGACCGCTATTATTGGGCCTAAAGCGTCCGTAGCTGGCCATACAAGTCCATCGGGAAATCTCCCTGCTTAACAGGGAGGCGTCCGGTAGAAACTGTGTGGCTTGGGACCGGAAGATCCAGAGGGTACGTCTGGGGTAGGAGTGAAATCCCGTAATCCTAGACGGACCACCGGTGGCGAAAGCGCTCTGGAAGGACGGATCCGACAGTGAGGGACGAAAGCTAGGGTCA |
| p__Halobacteriota_g__Halalkalicoccus_ASV_373 | 0 | 1 | 1 | 0 | 0 | 0 | 1 | 0 | 0 | 0 | 0 | CCTTTACACTGCACGACAGTGCGATAAGGGGACTCCAAGTGCGAGGGCATACAGTCCTCGCTTTTCTGTACCGTAAGGTGGTACAGGAATAAGAGCTGGGCAAGACCGGTGCCAGCCGCCGCGGTAATACCGGCAGCTCGAGTGATGGCCAATATTATTGGGCCTAAAGCGTCCGTAGCCGGCCGAACAGGTCCGTCGGGAAATCCGCTCGCTCAACGAGCGGACGTCCGGCGGAAACCAGTCGGCTTGGGACCGGAAGACCCGAGGGGTACGGCCGGGGTAGGAGTGAAATCCCGTAATCCTGGCCGGACCACCGGTGGCGAAAGCGCCTCGGGAAGACGGATCCGACGGTGAGGGACGAAAGCTAGGGTCA |
| p__Thermoplasmatota_g__SM1-50_ASV_377 | 0 | 0 | 0 | 0 | 0 | 0 | 0 | 1 | 1 | 1 | 1 | ACTTCACACTGTGCGAAAGCGCGATGAGGGAATCCCAAGTGTATGCACGTAAGTGTATACTGTTTCTTTGTTTAAAATGCAGAGAGAGTAAGGGCTGGGTAAGACGGGTGCCAGCCGCCGCGGTAATACCTGCAGCTCAAGTGGTGACCACTATTATTGGGCCTAAAACGTTCGTAGCCGGTTTATTAAATGCCTGGGTAAATCGGACCGCTTAACGGTTCGAATTCTGGGTAGACTGATAGACTTGAGACCGGGAGAGGCTAGAGGTACTCCTGGGGTAGAGGTGAAATTCTGTAATCCTGGGGGACCACCAGTGGCGAAGGCGTCTAGCTAGAACGGGTCTGACGGTGAGGGACGAAGCCCTGGGGCG |
| p__Halobacteriota_g__Halohasta_ASV_383 | 0 | 1 | 1 | 1 | 0 | 0 | 0 | 0 | 0 | 0 | 0 | CCTTTACACTGCACGCAGGTGCGATAGGGGAATCCCAAGTGCGAGGGCATACTAGCCCTCGCTTTTGTGTACTGTAAGGAGGTACACGAATAAGAGCTGGGCAAGACCGGTGCCAGCCGCCGCGGTAATACCGGCAGCTCGAGTGATGTCCACTATTATTGGGCCTAAAGCGTCCGTAGCTGGCCGTACAAGTTCGTCGGGAAATCCACCTGCCCAACAGGTGGGCGTCCGGCGAAAACTGTATGGCTTGGAACCGGAAGGCTCAGAGGGTACGTCCGGGGTAGGAGTGAAATCCTGTAATCCCGGACGGACCACCGATGGCGAAAGCACTCTGAGAGGACGGATTCGACAGTGAGGGACGAAAGCTAGGGTCT |
| p__Thermoplasmatota_g__SM1-50_ASV_385 | 1 | 1 | 0 | 0 | 0 | 0 | 0 | 1 | 1 | 1 | 1 | TCTTCACAATGTGCGAAAGCACGATGAGGGGATCCCAAGTGCTGGCACGTAAGTGTCAGCTGTTTCTTAGTCTAAACAACTGAGAGAGTAAGGGCTGGGTAAGACGGGTGCCAGCCGCCGCGGTAATACCTGCAGCTCAAGTGGTGGCCGCTATTATTGGGCCTAAAACGTCCGTAGCCTGTTTAGTAAATGCTTGGGTAAATCGTGTAGCTTAACTATACGAATACCGAGTAGACTGCTAGACTTGGGACCGGGAGAGGCTAGAGGTACTCCGGGGGTAGAGGTGAAATTCTGTAATCCTCGGGGGACCACCAGTGGCGAAGGCGTCTAGCTAGAACGGGTCTGACGGTGAGGGACGAAGCCCTGGGGCG |
| p__Thermoplasmatota_g__SM1-50_ASV_386 | 0 | 0 | 0 | 0 | 0 | 0 | 0 | 1 | 1 | 1 | 1 | CCTTCACACTGTGCGAGAGCACGATGAGGGAATCCCAAGTGCTGGCACGTAAGTGTCAGCTGTTTCTTTGTCTAAAAAACAAAGAGAGTAAGGGCTGGGTAAGACGGGTGCCAGCCGCCGCGGTAATACCTGCAGCTCAAGTGGTGACCGCTATTATTGGGCCTAAAACGTCCGTAGCCTGTTTGGTAAATGCCTGGGTAAATCGTGTCGCTTAACGATACGAATTCCGGGTAGACTGCCAGGCTTGAGACCGGGAGAGGTTAGAGGTACTCCGGGGGTAGAGGTGAAATTCTGTAATCCTCGGGGGACCACCAGTGGCGAAGGCGTCTAACCAGAACGGGTCTGACGGTGAGGGACGAAGCCCTGGGGCG |
| p__Halobacteriota_f__Natrialbaceae_ASV_390 | 0 | 1 | 1 | 1 | 0 | 0 | 0 | 0 | 1 | 0 | 0 | CCTTTACACTGCACGACAGTGCGATAAGGGGACTCCAAGTGCGAGGGCATATCGTCCTCGCTTTTTCCTACCGTAAGGTGGTAGGCGAATAAGGGCTGGGCAAGACCGGTGCCAGCCGCCGCGGTAATACCGGCAGCCCAAGTGATGACCGCTATTATTGGGCCTAAAGCGTCCGTAGCCGGCCGAACAAGTCCGTCGGGAAATCCGCACGCCTAACGTGCGGGCGTCCGGCGGAAACTGTTCGGCTTGGGACCGGAAGATCCAGAGGGTACGTCTGGGGTAGGAGTGAAATCCCGTAATCCTGGACGGACCACCGGTGGCGAAAGCGCTCTGGAAGGACGGATCCGACGGTGAGGGACGAAAGCTAGGGTCA |
| p__Halobacteriota_g__Halodesulfurarchaeum_ASV_388 | 1 | 1 | 1 | 0 | 0 | 1 | 1 | 1 | 0 | 1 | 1 | CCTTTACACTGCACGACAGTGCGATAGGGGGACTCCGAGTGCGAGGGCATAGAGCCCTCGCTTTTCTGTACCGTAAGGTGGTACAGGAAGAAGGGCTGGGCAAGACCGGTGCCAGCCGCCGCGGTAATACCGGCAGCCCAAGTGATGGCCGATATTATTGGGCCTAAAGCGTCCGTAGCTGGCCGGACAAGTCCGTTGGGAAATCCGCACGCTTAACGTGCGGGCGTCCAGCGGAAACTGTCCGGCTTGGGACCGGAAGACCCGAGGGGTACGTCCGGGGTAGGAGTGAAATCCTGTAATCTCGGACGGACCACCTGTGGCGAAAGCGCCTCGGGAAGACGGATCCGACAGTGAGGGACGAAAGCCAGGGTCT |
| p__Halobacteriota_g__Halorubrum_ASV_389 | 0 | 1 | 1 | 0 | 0 | 0 | 0 | 1 | 1 | 1 | 1 | CCTTTACACTGCACGCCAGTGCGATAAGGGAATCCCAAGTGCGCAGGCATAGAGCCTGCGCTTTTGTCCACCGTAGGGAGGTGGACGAATAAGGGCTGGGCAAGACCGGTGCCAGCCGCCGCGGTAATACCGGCAGCCCGAGTGATGGCCGATCTTATTGGGCCTAAAGCGTCCGTAGCTGGCCGCGCAAGTCCATCGGAAAATCCACCCGCTCAACGGGTGGGCGTCCGGTGGAAACTGCGTGGCTTGGGACCGGAAGGCGCGACGGGTACGTCCGGGGTAGGAGTGAAATCCCGTAATCCTGGACGGACCGCCGATGGCGAAAGCACGTCGCGAGGACGGATCCGACAGTGAGGGACGAAAGCCAGGGTCT |

| Phylum_Genus_ASV_number | Presence (1) or Absence (0) in each layer | | | | | | | | | | | Sequence |
|---|---|---|---|---|---|---|---|---|---|---|---|---|
| | L1 | L2 | L3 | L4 | L5 | L6 | L7 | L8 | L9 | L10 | L11 | |
| p__Halobacteriota_f__Haloferacaceae_ASV_393 | 0 | 1 | 0 | 0 | 1 | 1 | 1 | 0 | 0 | 0 | 0 | CCTTTACACTGCACGACAGTGCGATAAGGGGACTCCGAGTGCGAGGGCATATCGTCCTCGCTTTTCTGGACCGTAGGGAGGTCCAGGAACAAGAGCTGGGCAAGACCGGTGCCAGCCGCCGCGGTAATACCGGCAGCTCGAGTGATGGCCGATCTTATTGGGCCTAAAGCGTCCGTAGCTGGCCAGACAAGTCCGTCGGGAAATCCACCTGCCCAACAGGTGGGCGTCCGGCGGAAACTCTCTGGCTTGGGACCGGAAGACCCGAGGGGTACGTCCGGGGTAGGAGTGAAATCCCGTAATCCTGGACGGACCGCCGATAGCGAAAGCACCTCGGGAAGACGGATCCGACAGTGAGGGACGAAAGCTGGGGTCT |
| p__Halobacteriota_f__Natrialbaceae_ASV_402 | 0 | 1 | 1 | 1 | 0 | 0 | 0 | 0 | 0 | 0 | 0 | CCTTTACACTGCACGCCAGTGCGATAAGGGGACTCCAGGTGCGAGGGCATATAGTCCTCGCTTTTCACCACCGTAAGGTGGTGGTGGAATAAGGGCTGGGCAAGACCGGTGCCAGCCGCCGCGGTAATACCGGCAGCCCAAGTGATGACCGCTATTATTGGGCCTAAAGCGTCCGTAGCTGGCCACACAAGTCCGTCGGGAAATCTCCTCGCCTAACGAGGAGGCGTCCGGTGGAAACTGTGTGGCTTGGGACCGGAAGATCCAGAGGGTACGTCCGGGGTAGGAGTGAAATCCCGTAATCCTGGACGGACCACCGGTGGCGAAAGCGCTCTGGAAGGACGGATCCGACGGTGAGGGACGAAAGCTCGGGTCA |
| p__Thermoproteota_g__GCA-2726865_ASV_409 | 0 | 0 | 0 | 0 | 0 | 0 | 0 | 0 | 1 | 1 | 1 | CCTCCTCAATGCACTAAAGTGTGAAGGGGTTACCTCGAGTGCTACCTGAAGAAGGTGGCTTTTCCTTGGTCTAAAACGCCTTGGGAATAAGAGGGGGGCAAGACTGGTGTCAGCCGCCGCGGTAACACCAGCCCCTCGAGTGGTGGGGATGATTATTTGGCCTAAAGCGTCCGTAGCCGGCCAGGTAAGTTTTCCGTCAAATCCAATGTCTCAAGCGTTGGGCTGCGGGAAATACTGCCAGGCTAGAGAGTGGGAGAGGTAAGCGGTATTCTGGGGGTAGGGGCGAAATCCTTTGATCCCCGGAGGACCACCAGTGGCGAAGGCTGCTTACTAGAACACGTCTGACGGTCAGGGACGAAAGCTGGGGTAG |
| p__Thermoplasmatota_g__SM1-50_ASV_408 | 1 | 1 | 0 | 0 | 0 | 0 | 0 | 0 | 0 | 0 | 1 | TCTTCACAATGTGCGAGAGCACGATGAGGGAATCCCAAGTGCTAACACGTTAGTGTTAGCTGTTTTCTAGTCTAAAAAACTAGGAGAGTAAGAGCTGGGTAAGACGGGTGCCAGCCGCCGCGGTAATACCTGCAGCTCAAGTGGTGGCCGCTATTATTGGGCCTAAAACGTCCGTAGCCGGTTTAGTAAATGCTTGGGTAAATTGTGTAGCTTAACTATACAAATACCGAGTAGACTGCTAAACTTGGGACCGGGAGAGGCTAGAGGTACTCCTGGGGTAGAGGTGAAATTCTGTAATCCTAGGGGGACCACCAGTGGCGAAGGCGTCTAGCTAGAACGGGTCTGACGGTGAGGGACGAAGCCCTGGGGCG |
| p__Halobacteriota_g__Haloprofundus_ASV_414 | 0 | 1 | 1 | 1 | 0 | 0 | 0 | 0 | 0 | 0 | 0 | CCTTTACACTGCACGACAGTGCGACAAGGGGACCCCGAGTGCGAGGGCATACAGTCCTCGCTTTTGTGAACCGTAAGGAGGTTCACGAATAAGAGCTGGGCAAGACCGGTGCCAGCCGCCGCGGTAATACCGGCAGCTCGAGTGATGACCGCTGTTATTGGGCCTAAAGCGTCCGTAGCTGGCCGCGTAAGTTCATCGGGAAATCCATCCGCTCAACGGATGGGCGTCCGGTGAAAACTACACGGCTTGGAGCCGGAAGACCCAAGGGGTACGTCCGGGGTAGGAGTGAAATCCCGTAATCCTGGACGGACCGCCGATGGCGAAAGCACCTTGGGAAAACGGACTCGACAGTGAGGGACGAAAGCTAGGGTCT |
| p__Halobacteriota_o__Halobacteriales_ASV_412 | 1 | 1 | 1 | 1 | 0 | 0 | 1 | 0 | 0 | 0 | 0 | CCTTTACACTGCACGACAGTGCGATAGGGGGACCCCGAGTGCGAGGGCATATCGTCCTCGCTTTTGTACACCGTAAGGTGGTGTACGAATAAGTGCTGGGCAAGACCGGTGCCAGCCGCCGCGGTAATACCGGCAGCACAAGTGATGACCGTTATTATTGGGCCTAAAGCGTCCGTAGCCGGCCAGACAAGTTCGTCGGGAAATCCACCCGCTCAACGGGTGGGCGTCCGGCGAAAACTGTCTGGCTTGGGGCCGGAAGACCCAGGGGGTACGTCCGGGGTAGGAGTGAAATCCCGTAATCCTGGACGGACCGCCGGTGGCGAAAGCGCCCTGGGAGGACGGACCCGACGGTCAGGGACGAAAGCTAGGGTCT |
| p__Halobacteriota_g__Halohasta_ASV_415 | 1 | 0 | 1 | 0 | 0 | 0 | 0 | 0 | 0 | 0 | 1 | CCTTTACACTGCACGCAAGTGCGATAGGGGAATCCCAAGTGCGAGGGCATACTAGCCCTCGCTTTTGTGTACTGTAAGGAGGTACACGAATAAGAGCTGGGCAAGACCGGTGCCAGCCGCCGCGGTAATACCGGCAGCTCGAGTGATGTCCAATATTATTGGGCCTAAAGCGTCCGTAGCTGGCCAGCCAAGTTCGTCGGGAAATCCACCTGCCCAACAGGTGGGCGTCCGGCGAAAACTGTCTGGCTTGGAACCGGAAGGCTCAGAGGGTACGTCCGGGGTAGGAGTGAAATCCCGTAATCCCGGACGGACCACCGATGGCGAAAGCACTCTGAGAAGACGGATTCGACAGTGAGGGACGAAAGCTAGGGTCT |
| p__Halobacteriota_g__Halorussus_ASV_417 | 0 | 1 | 1 | 1 | 0 | 0 | 0 | 0 | 0 | 0 | 0 | CCTTTACACTGCACGCAAGTGCGATAAGGGGATCCCGAGTGCGAGGGCATACAGTCCTCGCTTTTGTACACCGTAAGGTGGTGTACGAATAAGAGCTGGGCAAGACCGGTGCCAGCCGCCGCGGTAATACCGGCAGCTCGAGTGATGGCCGCTATTATTGGGCCTAAAGCGTCCGTAGCCGGCCGAACAGGTCCGTTGGGAAATCCATCTGCCTAACAGATGGGCGTCCAGCGGAAACCAGTCGGCTTGGGGCCGGAAGACCTGAGGGGTACGTCCGGGGTAGGAGTGAAATCCTGTAATCCTGGACGGACCGCCGGTGGCGAAAGCGCCTCAGGAGGACGGACCCGACGGTGAGGGACGAAAGCTAGGGTCA |
| p__Halobacteriota_g__Halalkalicoccus_ASV_420 | 0 | 0 | 1 | 0 | 0 | 1 | 1 | 0 | 0 | 0 | 0 | CCTTTACACTGCACGAACGTGCGATAAGGGGACTCCAAGTGCGAGGGCATACAGTCCTCGCTTTTCCCTACCGTAAGGTGGTAGGTGAATAAGAGCTGGGCAAGACCGGTGCCAGCCGCCGCGGTAATACCGGCAGCTCGAGTGATGGCCAATATTATTGGGCCTAAAGCGTCCGTAGCTGGCCGAACAGGTCCGTCGGGAAATCCGCTCGCTCAACGAGCGGACGTCCGGCGGAAACCAGTCGGCTTGGGACCGGAAGACCCGAGGGGTACGGCCGGGGTAGGAGTGAAATCCTGTAATCCTGGCCGGACCACCGGTGGCGAAAGCGCCTCGGGAGGACGGATCCGACAGTGAGGGACGAAAGCTTGGGTCA |
| p__Halobacteriota_g__Natronobiforma_ASV_419 | 0 | 1 | 1 | 1 | 0 | 0 | 0 | 0 | 1 | 0 | 1 | CCTTTACACTGCACGACAGTGCGATAAGGGGACTCCAAGTGCGAGGGCATACAGTCCTCGCTTTTTCCTACCGTAAGGTGGTAGGCGAATAAGGGCTGGGCAAGACCGGTGCCAGCCGCCGCGGTAATACCGGCAGCCCAAGTGATGACCGCTATTATTGGGCCTAAAGCGTCCGTAGCCGGCCACGCAAGTCCGTCGGGAAATCTGCTCGCCTAACGAGCAGGCGTCCGGCGGAAACTGCGTGGCTTGGGACCGGAAGATCTGAGGGGTACGTCTGGGGTAGGAGTGAAATCCTGTAATCCTGGACGGACCACCGGTGGCGAAAGCGCCTCAGAAGGACGGATCCGACGGTGAGGGACGAAAGCTAGGGTCA |
| p__Halobacteriota_g__Saliphagus_ASV_423 | 1 | 1 | 1 | 0 | 0 | 0 | 0 | 0 | 0 | 1 | 1 | CCTTTACACTGCACGACAGTGCGATAGGGGGACTCCAAGTGCGAGGGCATATAGTCCTCGCTTTTCACCACCGTAAGGTGGTGGTGGAATAAGTGCTGGGCAAGACCGGTGCCAGCCGCCGCGGTAATACCGGCAGCACGAGTGATGACCGCTATTATTGGGCCTAAAGCGTCCGTAGCCGGCCGAACAAGTCCGTCGGGAAATCTGCACGCCTAACGTGCAGGCGTCCGGCGGAAACTGTTCGGCTTGGGACCGGAAGATCCAGAGGGTACGTCCGGGGTAGGAGTGAAATCCTGTAATCCTGGACGGACCACCGGTGGCGAAAGCGCTCTGGAAGGACGGATCCGACGGTGAGGGACGAAAGCTAGGGTCA |
| d__Archaea_d__Archaea_ASV_438 | 0 | 0 | 0 | 0 | 0 | 0 | 0 | 0 | 1 | 1 | 1 | CTTCTGCAATGTCCGTAAGGATGACAGAGCAACTCCGAGTGCTCATACATTGTATGGGCTTTTGCCATGTGTAAAAAGCTTGGCGAATAAGGGGTGGGCAAGACCGGTGGCAGCAGCCGCGGTAATACCGGCGCCCCAAGTGGTGGTCGTTTTTATTGGGCCTAAAGCGTTCGTAGCCGGTTCAGTAAGTTCTTGGTGAAATCCTTGCGCTTAACGTAAGGGCGTGCCAGGAATACTGCTGGACTAGAGACTGGGAGAAGACAGAGGTATTCTTAGGGTAAGGGTAAAATCTAATAATCCTAAGAGGACCGCCGGTGGCGAAGGCGTCTGTCTAGAACAGGTCTGACGGTGAGGGACGAAAGCTGAGGGAT |
| p__Halobacteriota_f__Haloferacaceae_ASV_436 | 1 | 1 | 1 | 0 | 1 | 0 | 0 | 0 | 0 | 0 | 1 | CCTTTACACTGCACGCCAGTGCGATAAGGGGACTCCGAGTGCGAGGGCATATCGTCCTCGCTTTTCTCAACCGTAGGGAGGTTGAGGAACAAGAGCTGGGCAAGACCGGTGCCAGCCGCCGCGGTAATACCGGCAGCTCGAGTGATGGCCGATCTTATTGGGCCTAAAGCGTCCGTAGCCGGCCAGACAAGTCCGTCGGGAAATCCACCTGCTCAACAGGTGGGCGTCCGGCGGAAACTCTCTGGCTTGGGACCAGAAGACCCGAGGGGTACGTCCGGGGTAGGAGTGAAATCCCGTAATCCTGGACGGACCACCGATAGCGAAAGCACCTCGGGAAGATGGATCCGACGGTGAGGGACGAAAGCTGGGGTCT |
| p__Halobacteriota_g__Salinarchaeum_ASV_442 | 1 | 1 | 1 | 1 | 1 | 1 | 0 | 0 | 1 | 0 | 0 | CCTTTACACTGCACGAGAGTGCGATAGGGGGACCCCGAGTGCGAGGGCATACAGTCCTCGCTTTTCACCACTGTAAGGTGGTGGTGGAATAAGTGCTGGGCAAGACCGGTGCCAGCCGCCGCGGTAATACCGGCAGCACGAGTGATGGCCGATTTTATTGGGCCTAAAGCGTCCGTAGCCGGCCAGACAGGTCCATCGGGAAATCTGCACGCCCAACGTGCAGGCGTCCGGTGGAAACCAGCTGGCTTGGGGCCGGAAGATCCAGGGGGTACGTCCGGGGTAGGAGTGAAATCCCGTAATCCCGGACGGACCGCCGGTGGCGAAAGCGCCCTGGAAGGACGGACCCGACGGTGAGGGACGAAAGCTAGGGTCA |
| p__Halobacteriota_g__Natronomonas_ASV_446 | 0 | 1 | 1 | 1 | 0 | 0 | 0 | 0 | 0 | 1 | 1 | CCTTTACACTGCACGCCAGTGCGATAGGGGGACACCGAGTGCGAGGGCATAGAGCCCTCGCTTTTGTAGACCGTAAGGAGGTCTACGAATAAGGGCTGGGCAAGACCGGTGCCAGCCGCCGCGGTAACACCGGCAGCCCAAGTGATGGCCGATATTATTGGGCCTAAAGCGTCCGTAGCTGGCCGCGCAAGTTCGTCGGGAAATCCACTCGCTTAACGAGTGGGCGTCCGGCGAAAACTGTGCGGCTTGGGACCGGAAGATCCAAGGGGTACGTCCGGGGTAGGAGTGAAATCCCGTAATCCTGGACGGACCACCGATGGCGAAAGCACCTTGGAAGGACGGATCCGACAGTGAGGGACGAAAGCTAGGGTCT |
| p__Nanoarchaeota_g__BMS3ABIN17_ASV_458 | 0 | 0 | 0 | 0 | 0 | 0 | 0 | 0 | 1 | 1 | 1 | ATTCTGCAATGCGCGAAAGCGTGACAGATTGAGCCAGAGTGCTTCTCAAATGAGAAGCTTTTGAGCACTGTAAAAAGGTGCAAGAATAAGGACTGGGTAAGACGAGTGCCAGCCGCCGCGGTAATACTCGCGGTCCAAGTCGCAGCCATATTTATTGGGTCTAAAACATCCGTAGCTTGCTTGATAAGTTCCCTGTGAAATTTTATCTCTTAAAGGTAAAGCGTGCAGGAAATACTGTCAAGCTAGAGACTGGAAGACGTAGGAAGTACGATTAAAGTAGCGGTAAAATGTGTTGATCTTAATCGGACTCACAATAGCGAAGGCATCCTACGAGGACAGTTCTGACAGTCAGGGATGAAGGCTAGGGGCG |
| p__Halobacteriota_g__Halomarina_3560_ASV_465 | 0 | 1 | 0 | 0 | 0 | 1 | 1 | 0 | 1 | 1 | 1 | CCTTTACACTGCACGACAGTGCGATAGGGGGATTCCGAGTGCGAGGGCATACAGTCCTCGCTTTTGTGTACCTTAAGGTGGTACACGAATAAGAGCTGGGCAAGACCGGTGCCAGCCGCCGCGGTAATACCGGCAGCTCGAGTGATGGCCGATTTTATTGGGCCTAAAGCGTCCGTAGCCGGCCAGTCAGGTCCGTCGGGAAATCCACTCGCTCAACGAGTGGGCGTCCGGCGGAAACCGTCTGGCTTGGGACCGGGAGATCTGAGGGGTACGTCCGGGGTAGGAGTGAAATCCTGTAATCCTGGACGGACCATCGGTGGCGAAAGCGCCTCAGAAGAACGGATCCGACGGTGAGGGACGAAAGCTAGGGTCT |
| p__Halobacteriota_o__Halobacteriales_ASV_466 | 0 | 1 | 1 | 1 | 1 | 0 | 0 | 0 | 0 | 0 | 1 | CCTTTACACTGCACGACAGTGCGATAAGGGGACCCCGAGTGCGAGGGCATACAGTCCTCGCTTTTGTGTACCGTAGGGAGGTACACGAATAAGGGCTGGGCAAGACCGGTGCCAGCCGCCGCGGTAATACCGGCAGCTCAAGTGATGGCCGCTGTTATTGGGCCTAAAGCGTCCGTAGCCGGCCGCACAAGTCCGTCGGGAAATCCACCCGCCTAACGGGTGGGCGTCCGGCGGAAACTGTGCGGCTTGGGACCGGAAGACTCGAGGGGTACGTCCGGGGTAGGAGTGAAATCCTGTAATCCTGGACGGACCGCCGGTGGCGAAAGCGCCTCGAGAGGACGGATCCGACGGTGAGGGACGAAAGCTAGGGTCT |
| p__Halobacteriota_f__Haloferacaceae_ASV_469 | 1 | 1 | 1 | 0 | 0 | 0 | 0 | 1 | 0 | 0 | 1 | CCTTTACACTGCACGAGAGTGCGATAAGGGGACTCCGAGTGCGAGGGCATATCGTCCTCGCTTTTCTGTACCGTAGGGCGGTACAGGAACAAGAGCTGGGCAAGACCGGTGCCAGCCGCCGCGGTAATACCGGCAGCTCAAGTGATGGCCGCTCTTATTGGGCCTAAAGCGTCCGTAGCTGGCCCTGAAAGTCCGTCGGGAAATCCACGCGCTCAACGCGTGGGCGTCCGGCGGAAACTCCAGGGCTTGGGACCGGAAGGCTCGAGGGGTACGTCCGGGGTAGGAGTGAAATCCTGTAATCCCGGACGGACCACCGATGGCGAAAGCACCTCGAGAAGACGGATCCGACAGTGAGGGACGAAAGCTAGGGTCT |
| p__Halobacteriota_g__Saliphagus_ASV_477 | 0 | 1 | 1 | 0 | 1 | 1 | 0 | 0 | 0 | 0 | 0 | CCTTTACACTGCACGACAGTGCGATAGGGGGACTCCAAGTGCTAGGGCATATCGTCCTAGCTTTTCACCACCGTAAGGTGGTGGTGGAATAAGTGCTGGGCAAGACCGGTGCCAGCCGCCGCGGTAATACCGGCAGCACGAGTGATGACCGCTATTATTGGGCCTAAAGCGTCCGTAGCCGGCCGTATAAGTCCGTCGGGAAATCCGCACGCCTAACGTGCGGGCGTCCGGCGGAAACTGTTCGGCTTGGGACCGGAAGACTCGAGGGGTACGTCTGGGGTAGGAGTGAAATCCCGTAATCCTGGACGGACCACCGGTGGCGAAAGCGCCTCGAGAGGACGGATCCGACGGTGAGGGACGAAAGCTAGGGTCA |
| p__Nanoarchaeota_g__BMS3ABIN17_ASV_482 | 0 | 1 | 1 | 1 | 1 | 0 | 0 | 0 | 0 | 0 | 1 | ATTCTTCAATGCGCGTAAGCGTGAAAGATTGAGCCAGAGTGTTTTCCAATTGGAAAACTTTTGCCAAATGTAAAAAGTTTGGCGAATAAGGACTGGGTAAGACGAGTGCCAGCCGCCGCGGTAATACTCGCGGTCCAAGTCGCAGCCATTTTTATTGGGTCTAAAACATCCGTAGCTTGCTTAATAAGTTTCTTGTGAAATTTTATATCTCAAGTATAAAAAGTGCAAGAAGTACTGTTAAGCTAGAGACTGGAAGACGTAAGAAGTATGTTCGAAGTAGCGGTAAAATGTGTTAATCTCGGACAGACTAACAAGAGCGAAGGCATCTTACGAGGACAGTTCTGACAGTAAGGGATGAAAGCTAGGGGCG |
| p__Halobacteriota_f__Natrialbaceae_ASV_487 | 1 | 1 | 1 | 1 | 0 | 0 | 0 | 0 | 0 | 0 | 0 | CCTTTACACTGCACGACAGTGCGATAGGGGGACTCCAAGTGCGAGGGCATATCGTCCTCGCTTTTCACTACCGTAAGGTGGTAGTAGAATAAGGGCTGGGCAAGACCGGTGCCAGCCGCCGCGGTAATACCGGCAGCCCGAGTGATGACCGCTATTATTGGGCCTAAAGCGTCCGTAGCCGGCCTCGCAAGTCTGTCGGGAAATCCGCGCGCTCAACGCGCGGGCGTCCGGCGGAAACTGCGTGGCTTGGGACCGGAAGACTCAGGGGGTACGTCTTGGGTAGGAGTGAAATCCTGTAATCCTGGACGGACCACCGGTGGCGAAAGCGCCTTGAGAGAACGGATCCGACGGTGAGGGACGAAAGCTAGGGTCA |
| p__Halobacteriota_g__Halorussus_ASV_490 | 0 | 1 | 1 | 0 | 0 | 0 | 0 | 0 | 0 | 0 | 0 | CCTTTACACTGCACGAAAGTGCGATAGGGGGACCCCGAGTGCGAGGGCATACAGTCCTCGCTTTTGTGTACCGTAAGGAGGTACACGAATAAGGGCTGGGCAAGACCGGTGCCAGCCGCCGCGGTAATACCGGCAGCTCGAGTGATGGCCGCTGTTATTGGGCCTAAAGCGTCCGTAGCCGGCCGCACAAGTCCGTCGGGAAATCCACCCGCCTAACGGGTGGGCGTCCGGCGGAAACTGTGCGGCTTGGGGCCGGAAGATCCGAGGGGTACGTCCGGGGTAGGAGTGAAATCCTGTAATCCTGGACGGACCGCCGGTGGCGAAAGCGCCTCGGAAGGACGGACCCGACGGTGAGGGACGAAAGCTAGGGTCT |
| p__Thermoplasmatota_g__SM1-50_ASV_489 | 0 | 0 | 0 | 0 | 0 | 0 | 0 | 1 | 1 | 1 | 1 | CCTTCACACTGTGCGAAAGCGCGATGAGGGAATCCCAAGTGCTTGCACGTAAGTGTAAGCTGTTTCTCTGTTTAAAACGCAGGGGAAGAAAGGGCTGGGTAAGACGGGTGCCAGCCGCCGCGGTAATACCTGCAGCTCAAGTGGTGACCGCTATTATTGGGCCTAAAACGTTCGTAGCCGGCTGAATAAATGCTTGGGTAAATCGTGTAGCTCAACTATACGAATTCCGGGTAGACTGTTCAGCTTGAGACCGGGAGAAGCTAGAGGTACTCCTGGGGTAGAGGTGAAATTCTGTAATCCTGGGGGGACCACCAGTGGCGAAGGCGTCTAGCTAGAACGGGTCTGACGGTGAGGAACGAAACCCTGGGGCG |

| | Presence (1) or Absence (0) in each layer | | | | | | | | | | |
|---|---|---|---|---|---|---|---|---|---|---|---|
| Phylum_Genus_ASV_number | L1 | L2 | L3 | L4 | L5 | L6 | L7 | L8 | L9 | L10 | L11 | Sequence |
| p__Halobacteriota_g__Halolamina_ASV_498 | 0 | 1 | 1 | 1 | 0 | 0 | 0 | 0 | 0 | 0 | 0 | CCTTTACACTGCACGCCAGTGCGATAAGGGGACCCCGAGTGAAAGGGCATATCGCCCTTTCTTTTCTGAACCGTAGGGAGGTTGAGGAACAAGAGCTGGGCAAGACCGGTGCCAGCCGCCGCGGTAATACCGGCAGCTCGAGTGATGACCGATTTTATTGGGCCTAAAGCGTCCGTAGCTGGCCGCGCAAGTCTGTCGGGAAATCCACCCGCTTAACGGGTGGGCGTCCGGCGGAAACTGTGCGGCTTGGGACAGGAAGGCTCGAGGGGTACGTTCGGGGTAGGAGTGAAATCCCGTAATCCTGAACGGACCGCCGATGGCGAAAGCACCTCGAGAGGACTGATCCGACAGTGAGGGACGAAAGCTGGGGTCT |
| p__Halobacteriota_g__Natronoarchaeum_ASV_496 | 0 | 1 | 1 | 0 | 0 | 0 | 0 | 0 | 0 | 0 | 0 | CCTTTACACTGCACGCCAGTGCGATAAGGGGACCCCCAGTGCGAGGGCATACCGTCCTCGCTTTTCACCACCGTAAGGTGGTGGTGGAACAAGAGCTGGGCAAGACCGGTGCCAGCCGCCGCGGTAATACCGGCAGCTCGAGTGATGGCCGTTTTTATTGGGCCTAAAGCGTCCGTAGCCGGCCAGACAAGTTCGTCGGGAAATCCACGTGCCCAACACGTGGGCGTCCGGCGAAACCTCTCTGGCTTGGGACCGGACGGCCCGAGGGGTACGGTCGGGGTAGGAGTGAAATCCTGTAATCCCGACCGGACCACCGGTGGCGAAAGCGCCTCGGGAGGACGGATCCGACGGTGAGGGACGAAAGCCAGGGTCA |
| p__Halobacteriota_f__Natrialbaceae_ASV_507 | 0 | 1 | 1 | 1 | 0 | 0 | 0 | 0 | 0 | 1 | 1 | CCTTTACACTGCACGACAGTGCGATAGGGGGACTCCAAGTGCGAGGGCATATAGTCCTCGCTTTTCACCACCGTAAGGTGGTGGTGGAATAAGTGCTGGGCAAGACCGGTGCCAGCCGCCGCGGTAATACCGGCAGCACGAGTGATGACCGCTATTATTGGGCCTAAAGCGTCCGTAGCCGGCCGTGTAAGTCCATCGGGAAATCTCTCCGCCCAACGGAGAGGCGTCCGGTGGAAACTGCACGGCTTGGGACCGGAAGACCTGAGGGGTACGTCCGGGGTAGGAGTGAAATCCCGTAATCCTGGACGGACCACCTGTGGCGAAAGCGCCTCAGGAGAACGGATCCGACGGTGAGGGACGAAAGCTAGGGTCA |
| p__Halobacteriota_g__Halohasta_ASV_505 | 1 | 1 | 1 | 1 | 0 | 0 | 0 | 0 | 0 | 0 | 1 | CCTTTACACTGCACGCAAGTGCGATAGGGGAATCCCAAGTGCGAGGGCATACTAGCCCTCGCTTTTGTCTACCCTAAGGCGGTAGACGAATAAGAGCTGGGCAAGACCGGTGCCAGCCGCCGCGGTAATACCGGCAGCTCGAGTGATGTCCACTATTATTGGGCCTAAAGCGTCCGTAGCTGGCCGCGCAAGTCCGTCGGGAAATCCACCTGCCCAACAGGTGGGCGTCCGGCGGAAACTGCGTGGCTTGGAACCGGAAGGCTCAGAGGGTACGTCCGGGGTAGGAGTGAAATCCCGTAATCCCGGACGGACCACCGATGGCGAAAGCACTCTGAGAGGACGGATTCGACAGTGAGGGACGAAAGCTAGGGTCT |
| p__Halobacteriota_g__Halomicrobium_3849_ASV_509 | 0 | 1 | 1 | 1 | 0 | 0 | 0 | 0 | 0 | 0 | 0 | CCTTTACACTGCACGACAGTGCGATAAGGGGACTCCGAGTGCGAGGGCATATAGTCCTCGCTTTTCTGTACCGTAAGGTGGTACAGGAACAAGTGCTGGGCAAGACCGGTGCCAGCCGCCGCGGTAATACCGGCAGCACGAGTGATGGCCGCTCTTATTGGGCCTAAAGCGTCCGTAGCCGGCCAGACAAGTCCGTTGGGAAATCGACGCGCTCAACGCGTCGGCGTCCAGCGGAAACTGTCTGGCTTGGGGCCGGAAGACCTGAGGGGTACGTCCGGGGTAGGAGTGAAATCCCGTAATCCTGGACGGACCACCGGTGGCGAAAGCGCCTCAGGAAGACGGACCCGACGGTGAGGGACGAAAGCTAGGGTCT |
| p__Halobacteriota_g__Halorussus_ASV_511 | 0 | 1 | 1 | 1 | 0 | 0 | 0 | 0 | 0 | 0 | 0 | ACTTTACACTGCACGACAGTGCGATAAGGGGACTCCGAGTGCGAGGGCATATAGTCCTCGCTTTTGTACACCGTAAGGTGGTGTACGAATAAGGGCTGGGCAAGACCGGTGCCAGCCGCCGCGGTAATACCGGCAGCCCGAGTGATGGCCGCTATTATTGGGCCTAAAGCGTCCGTAGCCAGCCAGACAGGTCCGTCGGGAAATCCATCCGCTTAACGGATGGGCGTCCGGCGGAAACCAGCTGGCTTGGGGCCGGAAGACCTGAGGGGTACGTCCGGGGTAGGAGTGAAATCCCGTAATCCTGGACGGACCGCCGGTGGCGAAAGCGCCTCAGGAGGACGGACCCGACGGTGAGGGACGAAAGCTAGGGTCA |
| p__Nanoarchaeota_g__BMS3ABIN17_ASV_513 | 1 | 0 | 0 | 0 | 0 | 0 | 0 | 1 | 1 | 1 | 1 | CTTTTGCAATGCGCGAAAGCGTGACAAAGCAAGCCAGAGTGTTTCTTTTATTAAAGAAACTTTTGTCAGCTGCAAAAAGGCTGACGAATAAGGACTGGGTAAGACCGGTGCCAGCCGCCGCGGTAATACCGGCGGTCCAAGTCGCAGCCATCATTATTGGGTCTAAAACATCCGTAGCTTGTTCTGTAAGTCTCTTGTGAAATCTGACATCTCAAGTGTCAGGCGGGCAAGAGATACTGCAAGACTAGAGACCGGAAGACGTAACGAGTACAACTGAGGTAGTGGTAAAATACGTTAATCTTGGTTGGACTAACAATGGCGAAGGCACGTTACGAGGACGGATCTGACAGTGAGGGATGAAGGCTAGGGGCG |
| p__Halobacteriota_g__Halomicrobium_3849_ASV_512 | 1 | 1 | 1 | 0 | 0 | 0 | 0 | 0 | 0 | 0 | 1 | CCTTTACACTGCACGACAGTGCGATAAGGGGACTCCGAGTGCGAGGGCATATAGTCCTCGCTTTTCTGTACCGTAAGGAGGTACAGGAACAAGTGCTGGGCAAGACCGGTGCCAGCCGCCGCGGTAATACCGGCAGCACGAGTGATGGCCGCTATTATTGGGCCTAAAGCGTCCGTAGCCGGCCAGACAAGTCCGTTGGGAAATCGACGCGCTCAACGCGTCGGCGTCCAGCGGAAACTGTCTGGCTTGGGGCCGGAAGACCTGAGGGGTACGTCCGGGGTAGGAGTGAAATCCCGTAATCCTGGACGGACCACCGGTGGCGAAAGCGCCTCAGGAAGACGGACCCGACGGTGAGGGACGAAAGCTAGGGTCT |
| p__Halobacteriota_g__Saliphagus_ASV_518 | 0 | 1 | 1 | 0 | 0 | 0 | 0 | 0 | 0 | 0 | 0 | CCTTTACACTGCACGCCAGTGCGATAAGGGGACTCCAGGTGCGAGGGCATACAGTCCTCGCTTTTGTGTACCGTAGGGAGGTACACGAATAAGGGCTGGGCAAGACCGGTGCCAGCCGCCGCGGTAATACCGGCAGCCCGAGTGATGACCGCTATTATTGGGCCTAAAGCGTCCGTAGCCGGCCTCGCAAGTCCGTCGGGAAATCCGCACGCCCAACGTGCGGGCGTCCGGCGGAAACTGCGTGGCTTGGGACCGGAAGACTCGAGGGGTACGTCTGGGGTAGGAGTGAAATCCCGTAATCCTGGACGGACCACCGGTGGCGAAAGCGCCTCGAGAGGACGGATCCGACGGTGAGGGACGAAAGCTAGGGTCA |
| p__Halobacteriota_g__Natronobiforma_ASV_521 | 0 | 1 | 1 | 1 | 0 | 0 | 0 | 0 | 0 | 0 | 0 | CCTTTACACTGCACGCCAGTGCGATAAGGGGACACCAAGTGCGAGGGCATATCGTCCTCGCTTTTCACCACCGTAAGGTGGTGGTGGAATAAGAGCTGGGCAAGACCGGTGCCAGCCGCCGCGGTAATACCGGCAGCTCGAGTGATGACCGCTATTATTGGGCCTAAAGCGTCCGTAGCTGGCCACACAAGTCCATCGGGAAATCTCCCTGCCTAACAGGGAGGCGTCCGGTAGAAACTGTGTGGCTTGGGACCGGAAGATCCAGAGGGTACGTCTGGGGTAGGAGTGAAATCCTGTAATCCTGGACGGACCACCGGTGGCGAAAGCGCTCTGGAAGGACGGATCCGACAGTGAGGGACGAAAGCTAGGGTCA |
| p__Nanoarchaeota_g__BMS3ABIN17_ASV_526 | 1 | 1 | 1 | 0 | 0 | 0 | 0 | 0 | 1 | 1 | 1 | AATCTACAATGCGCGAAAGCGTGATAGTTTGAATCAGAGTGCTTTGCTTTGCAAAGCTTTTGTGAAATCTAAAAAGTTTCACGAATAAGGACTGGGTAAGACGAGTGCCAGCCGCCGCGGTAATACTCGCGGTCCAAGTCGTAGCCATTTTTATTGGGTCTAAAACATCCGTAGCTTGCTTGGTGAGTTCCTTGTGAAATCCTATGTCTCAAATATAGGGCGCGTGAGGAGTACTGCTAAGCTAGAGACTGGGAGACGTAAAGAGTACGTTCGAGGTAGCGGTAAAATGTGTTAATCTCGGACGGACTAACAATAGCGAAGGCACTTTACGAGAACAGTTCTGACAGTAAGGGATGAAGGCTAGGGGCG |
| p__Halobacteriota_g__Halovenus_ASV_529 | 0 | 1 | 1 | 1 | 0 | 0 | 0 | 0 | 0 | 0 | 0 | CCTTTACACTGCACGCCAGTGCGATAGGGGGATTCCGAGTGCGAGGGCATACAGTCCTCGCTTTTCTACACCGTAAGGAGGTGCAGGAACAAGTGCTGGGCAAGACCGGTGCCAGCCGCCGCGGTAATACCGGCAGCACGAGTGATGGCCGATCTTATTGGGCCTAAAGCGTCCGTAGCTGGCCGGACAAGTCCGTTGGGAAATCCACGTGCCCAACACGTGGGCGTCCAGCGGAAACTGTCCGGCTTGGGGCCGGAGGACTCGAGGGGTACGTCCGGGGTAGGAGTGAAATCCCGTAATCCTGGACGGACCACCGGTGGCGAAAGCGCCTCGAGAGGACGGACCCGACAGTGAGGGACGAAAGCCAGGGTCT |
| p__Thermoproteota_g__PALSA-986_ASV_533 | 0 | 1 | 0 | 1 | 0 | 0 | 0 | 0 | 1 | 1 | 1 | CCTTTGCAATACACGCAAGTGTGACAGGGCCATCCCGAGTGCCGACCGCTGAGGTTGGCTTTTACCCAGTTTAGAAAGCTGGGGGAATAAGGAGAGGGCAAGTCTGGTGTCAGCCGCCGCGGTAATACCAGCTCTCCGAGTGGTGTGGACGTTTATTGGGCCTAAAGCATCCGTAGCTGGCTGATCAAGTCCCCTGTTAAACCCACCGATTTAATCGTTGGCGTGCGGGGGAAACTGATCGGCTAGGGGACGAGAGAGGCAGACGGTATTCCCGGGGTAGGGGTGAAATCCTATAATCCCGAGAAGACCACCAGTGGCGAAGGCTGTCTGCTAGAACGCGCCCGACGGTGAGGGATGAAAGCTGGGGGAG |
| p__Halobacteriota_g__Halovenus_ASV_536 | 1 | 1 | 1 | 0 | 0 | 0 | 0 | 0 | 1 | 1 | 1 | CCTTTACACTGCACGACAGTGCGATAGGGGGACTCCGAGTGCGAGGGCATACAGTCCTCGCTTTTCTGTACCGTAAGGTGGTACAGGAACAAGTGCTGGGCAAGACCGGTGCCAGCCGCCGCGGTAATACCGGCAGCACAAGTGATGGCCGATATTATTGGGCCTAAAGCGTCCGTAGCTGGCCAGACAAGTCTGTTGGGAAATCCACGCGCCCAACGCGTGGGCGTCCAGCGGAAACTGTCTGGCTTGGGGCCGGAGGACTCGAGGGGTACGTCCGGGGTAGGAGTGAAATCCCGTAATCCTGGACGGACCACCGGTGGCGAAAGCGCCTCGAGAGGACGGACCCGACAGTGAGGGACGAAAGCCAGGGTCT |
| p__Halobacteriota_f__Natrialbaceae_ASV_550 | 0 | 0 | 1 | 1 | 0 | 0 | 0 | 0 | 0 | 1 | 1 | CCTTTACACTGCACGCCAGTGCGATAAGGGGACTCCATCTGCGAGGGCATATCGTCCTCGCTTTTCTCGACCGTAAGGTGGTCGAGGAATAAGGGCTGGGCAAGACCGGTGCCAGCCGCCGCGGTAATACCGGCAGCCCGAGTGATGACCGCTATTATTGGGCCTAAAGCGTCCGTAGCTGGCCAGGCAAGTCCATCGGGAAATCCGCGTGCCCAACACGCGGGCGTCCGGTGGAACCTGCCTGGCTTGGGACCGGAAGACCAGAGGGGTACGTCCGGGGTAGGAGTGAAATCCCGTAATCCTGGACGGACCACCGGTGGCGAAAGCGCCTCTGGAAGACGGATCCGACGGTGAGGGACGAAAGCTCGGGTCA |
| p__Halobacteriota_g__Halorussus_ASV_549 | 1 | 1 | 1 | 0 | 0 | 0 | 0 | 0 | 0 | 0 | 1 | CCTTTACACTGCACGAGAGTGCGATAAGGGGACCCCGAGTGCGAGGGCATACAGTCCTCGCTTTTGTACACCGTAAGGAGGTGTACGAATAAGAGCTGGGCAAGACCGGTGCCAGCCGCCGCGGTAATACCGGCAGCTCGAGTGATGGCCGCTATTATTGGGCCTAAAGCGTCCGTAGCCGGCCGAACAGGTCCGTTGGGAAATCCACCAGCCTAACTGGTGGGCGTCCAGCGGAAACCAGTCGGCTTGGGGCCGGAAGACCTGAGGGGTACGTCCGGGGTAGGAGTGAAATCCTGTAATCCTGGACGGACCGCCGGTGGCGAAAGCGCCTCAGGAGGACGGACCCGACGGTGAGGGACGAAAGCTAGGGTCA |
| p__Thermoplasmatota_g__SM1-50_ASV_555 | 1 | 1 | 0 | 0 | 0 | 0 | 0 | 0 | 0 | 0 | 1 | CCTTCGCACTGTGCGAAAGCGCGATGAGGGGATCCCAAGTGCTTGCACGTAAGTGTAAGCTGTTTCTATGTCTAAAACGCATTGAGAGTAAGGGCTGGGTAAGACGGGTGCCAGCCGCCGCGGTAATACCTGCAGCTCAAGTGGTGACCACTATTATTGGGCCTAAAACGTCCGTAGCCGGCTTGGTAAATGCCTGGGTAAATCGTATAGCTTAACTATACGAATTCCGGGTAGACTGTCAAGCTTGAGACTGGGAGAAGCTAGAAGTACTCCTGGGGTAGAGGTGAAATTCTGTAATCCTAGGGGGACTACCGGTGGCGAAAGCGTCTAGCTAGAACGGGTCTGACGGTGAGGGACGAAGCCCTGGGGCG |
| p__Halobacteriota_g__Natronoarchaeum_ASV_561 | 0 | 1 | 1 | 0 | 0 | 0 | 0 | 0 | 0 | 0 | 1 | CCTTTACACTGCACGACAGTGCGATAAGGGGACCCCCAGTGCGAGGGCATACCGTCCTCGCTTTTCAACACCGTAAGGAGGTGCTGGAACAAGAGCTGGGCAAGACCGGTGCCAGCCGCCGCGGTAATACCGGCAGCTCGAGTGATGGCCGTTTTTATTGGGCCTAAAGCGTCCGTAGCCGGCCAGACAAGTTCGTCGGGAAATCCACGTGCCCAACACGTGGGCGTCCGGCGAAACCTCTCTGGCTTGGGACCGGACGGCCCGAGGGGTACGGTCGGGGTAGGAGTGAAATCCTGTAATCCCGACCGGACCACCGGTGGCGAAAGCGCCTCGGGAGGACGGATCCGACGGTGAGGGACGAAAGCCAGGGTCA |
| p__Halobacteriota_g__Halohasta_ASV_565 | 1 | 1 | 0 | 0 | 0 | 0 | 0 | 0 | 0 | 0 | 1 | CCTTTACACTGCACGTAAGTGCGATAGGGGAATCCCAAGTGCGAGGGCATACTAGCCCTCGCTTTTGTTGACTGTAAGGTGGTCAACGAATAAGAGCTGGGCAAGACCGGTGCCAGCCGCCGCGGTAATACCGGCAGCTCGAGTGATGTCCAATATTATTGGGCCTAAAGCGTCCGTAGCTGGCCAGCCAAGTCCGTCGGGAAATCCACCTGCCCAACAGGTGGGCGTCCGGCGGAAACTGTCTGGCTTGGAACCGGAAGGCTCAGAGGGTACGTCCGGGGTAGGAGTGAAATCCCGTAATCCCGGACGGACCACCGATGGCGAAAGCACTCTGAGAAGACGGATTCGACAGTGAGGGACGAAAGCTAGGGTCT |
| p__Halobacteriota_g__Halorussus_ASV_567 | 1 | 1 | 0 | 0 | 0 | 0 | 0 | 0 | 0 | 0 | 0 | CCTTTACACTGTACGAAAGTGCGATAAGGGGACCCCGAGTGCGAGGGCATACAGTCCTCGCTTTTGTGTACCGTAAGGTGGTACACGAATAAGAGCTGGGCAAGACCGGTGCCAGCCGCCGCGGTAATACCGGCAGCTCGAGTGATGGCCGCTATTATTGGGCCTAAAGCGTCCGTAGCCGGCCGAACAGGTCCGTTGGGAAATCCGTCAGCCTAACTGACGGGCGTCCAGCGGAAACCAGTCGGCTTGGGGCCGGAAGACCTGAGGGGTACGTCCGGGGTAGGAGTGAAATCCTGTAATCCTGGACGGACCGCCGGTGGCGAAAGCGCCTCAGGAGGACGGACCCGACGGTGAGGGACGAAAGCTAGGGTCA |
| p__Halobacteriota_g__Salinigranum_ASV_571 | 0 | 1 | 1 | 1 | 1 | 1 | 0 | 0 | 1 | 0 | 0 | CCTTTACACTGCACGCGAGTGCGATAAGGGGACCCCGAGTGCGAGGGCATACTAGCCCTCGCTTTTGTGAACCGTACGAAGGTTCTCGAACAAGAGCTGGGCAAGACCGGTGCCAGCCGCCGCGGTAATACCGGCAGCTCGAGTGATGGCCGATCTTATTGGGCCTAAAGCGTCCGTAGCTGGCCGTGCAAGTCCGTCGGGAAATCCACCTGCTCAACAGGTGGGCGTCCGGCGGAAACTGCTCGGCTTGGGACCGGAAGGCTCGAGGGGTACGTCCGGGGTAGGAGTGAAATCCCGTAATCCTGGACGGACCACCGATGGCGAAAGCACCTCGAGAGGACGGCTCCGACAGTGAGGGACGAAAGCTGGGGTCT |
| p__Halobacteriota_f__Haloarculaceae_3993_ASV_584 | 0 | 1 | 1 | 1 | 0 | 1 | 0 | 0 | 0 | 0 | 0 | CCTTTACACTGCACGAAAGTGCGATAGGGGGACTCCGAGTGCGAGGGCATATAGTCCTCGCTTTTCTGTACCGTAAGGTGGTACAGGAACAAGTGCTGGGCAAGACCGGTGCCAGCCGCCGCGGTAATACCGGCAGCACGAGTGATGGCCGATATTATTGGGCCTAAAGCGTCCGTAGCTTGCTGTGTAAGTCCATTGGGAAATCCACGCGCTCAACGCGTGGGCGTCCGGTGGAAACTGCGCGGCTTGGGGCCGAGAGACTCGACGGGTACGTCCGGGGTAGGAGTGAAATCCCGTAATCCTGGACGGACCACCAATGGGGAAACCACGTCGAGAGACCGGACCCGACAGTGAGGGACGAAAGCCAGGGTCT |
| p__Halobacteriota_g__Haladaptatus_ASV_601 | 0 | 1 | 0 | 0 | 0 | 0 | 0 | 0 | 0 | 0 | 0 | ACTTTACACTGCACGCCAGTGCGATAAGGGAACTCCAAGTGCGAGGGCATATAGTCCTCGCTTTTGTGTATCGTAAGGTGGTACACGAATAAGAGCTGGGCAAGACCGGTGCCAGCCGCCGCGGTAATACCGGCAGCTCAAGTGATGGCCAATCTTATTGGGCCTAAAGCGTCCGTAGCCGGCCGAACAGGTCCGTCGGGAAATCTGCTCGCTCAACGAGCAGGCGTCCGACGGAAACCAGTCGGCTTGGGACCGGAAGACCCGAGGGGTACGTCTGGGGTAGGAGTGAAATCCTGTAATCCTAGACGGACCGCCGGTGGCGAAAGCGCCTCGGGAAGACGGATCCGACGGTGAGGGACGAAAGCTAGGGTCA |
| p__Nanoarchaeota_g__BMS3ABIN17_ASV_605 | 0 | 0 | 1 | 1 | 0 | 0 | 1 | 1 | 1 | 0 | 0 | CTTCTGCAATGCGCGAAAGCGTGACAGAGTGAGTCAGAGTGTCTGCCATTCGGTAGACTTTTGTGGAATGTAAAAAGTTTCACGAATAAGGACTGGGTAAGACTAGTGCCAGCCGCCGCGGTAATACTAGCGGTCCAAGTCGCAGCCACTATTATTGGGTCTAGAACATCCGTAGCTTGCTTAATAAGTCTCTTGTGAAATCCTGTGTCTCAAGTATAGGACTTGCAAGAGAAACTGTTAAGCTAGAGACCGGGAGACGTAAGGAGTACGTTTGAAGTAGCGGTAAAATGTGTTGATCTCAGACGGACTAACAAGAGCGAAGGCACCTTACGAGAACGGTTCTGACAGTAAGGGATGAAGGCCAGGGTCG |

| Phylum_Genus_ASV_number | Presence (1) or Absence (0) in each layer | | | | | | | | | | | |
|---|---|---|---|---|---|---|---|---|---|---|---|---|
| | L1 | L2 | L3 | L4 | L5 | L6 | L7 | L8 | L9 | L10 | L11 | Sequence |
| p__Asgardarchaeota_f__Thorarchaeaceae_ASV_611 | 0 | 0 | 0 | 0 | 0 | 0 | 0 | 0 | 1 | 1 | 1 | CCTCACCAATGCGCGAAAGCGTGAGTGGGCCAAGCCAAGTGATGGGAGATAATTCCATCTGTGACGAAACAGTAAGGATGTTCGTTAGAAAGGAGAGGGCAAGGCTGGTGCCAGCCGCCGCGGTAAAACCAGCTCTTCGAGTGGTGTCCATGATTATTGAGCTTAAAGCGTTCGTAGCCTGTTTGGTAGGTCCCTGCTCAAATCCGGTCGCTCAACGGTCGGACTGGTAGGGATACCGCCTTACTTGGGGACGGGAGACGGCGACGGTATTCCGTAGGTAAGGGTGAAATCTAATGATCTACGGAGGACCACCAGTGGCGAAGGCGGTCGTCGAGAACGTGCCCGACGGTGAGGGACGAAACCCAGGGGAG |
| p__Nanoarchaeota_g__BMS3ABIN17_ASV_608 | 1 | 1 | 0 | 0 | 0 | 0 | 0 | 0 | 0 | 0 | 1 | AATCTACAATGCGCGAAAGCGTGATAGTTTGAGCCAGAGTGCTTTCCTTTTGGAAAGCTTTTGCCAAATGTAAAAAGTTTGGCGAATAAGGACTGGGTAAGACTAGTGCCAGCCGCCGCGGTAATACTAGCGGTCCAAGTCGCAGCCATTTTTATTGGGTCTAAAACATCCGTAGCTTGCTAAATAAGTTTCTTGTGAAATCTTATATCTCAAGTATAAGGCGGGCAAGAAATACTATTTAGCTAGAGACTGGAAGACGTAAGAAGTACGTTTGAAGTAACGGTAAAATGTGTTAATCTCAGACGGACTTACAAGAGCGAAGGCATCTTACGAGGACAGTTCTGACAGTAAGGGATGAAAGCCAGGGGCG |
| p__Halobacteriota_f__Haloarculaceae_3992_ASV_617 | 0 | 1 | 1 | 1 | 0 | 0 | 0 | 0 | 0 | 0 | 1 | CCTTTACACTGCACGCCAGTGCGATAAGGGGACTCCGAGTGCGAGGGCATACAGTCCTCGCTTTTCTGTACCGTAAGGAGGTACAAGAATAAGGGCTGGGCAAGACCGGTGCCAGCCGCCGCGGTAATACCGGCAGCCCAAGTGATGGCCGATATTATTGGGCCTAAAGCGTCCGTAGCCGGCCGGACAAGTCCGTTGGGAAATCCACGCGCTCAACGCGTGGGCGTCCAGCGGAAACTGTCCGGCTTGGGGCCGGAGGACTCGAGGGGTACGTTCGGGGTAGGAGTGAAATCCTGTAATCCTGAACGGACCACCGGTGGCGAAAGCGCCTCGAGAGGACGGACCCGACGGTGAGGGACGAAAGCCAGGGTCT |
| d__Archaea_d__Archaea_ASV_620 | 0 | 0 | 0 | 0 | 0 | 0 | 0 | 0 | 1 | 1 | 1 | CCTTCGCAATGCAGGAAACTGCGACGAGGGGACTCCAAGTGCCTACGCTCAGCGTAGGCTGTTCATGAGCCTAGAAAGCTTTTGGAGTAAGGGCTGGGTAAGACGGGTGCCAGCCGCCGCGGTAACACCCGCAGCCCGAGTGGTGGCCGCTTCTATTGGGCCTAAAGCGTCCGTAGCCGGCCCGGTAAATGCCTGGGTAAATCGTGGGGCCTAACCTTACGAACACCGGGTAGACTGCCGAGCTAGGGACCGGAAGAGGGAGAGGGTACTTCAGGGGTAGGGGTGAAATCCTGTAATCCCTGGAGGACCACCCGTGGCGAAGGCGCTCTCCTAAGACGGGTCCGACGGTGAGGGACGAAAGCCAGGGGAG |
| p__Halobacteriota_g__Halorussus_ASV_622 | 0 | 1 | 1 | 0 | 0 | 0 | 0 | 0 | 0 | 0 | 0 | CCTTTACACTGTACGAAAGTGCGATAAGGGGACCCCGAGTGCGAGGGCATACAGTCCTCGCTTTTGTGTACCGTACGATGGTACACGAATAAGAGCTGGGCAAGACCGGTGCCAGCCGCCGCGGTAATACCGGCAGCTCAAGTGATGGCCGCTATTATTGGGCCTAAAGCGTCCGTAGCCGGCCGAACAGGTCTGTTGGGAAATCCGCCAGCTTAACTGGCGGGCGTCCAGCAGAAACCAGTCGGCTTGGGGCCGGAAGACCTGAGGGGTACGTCCGGGGTAGGAGTGAAATCCTGTAATCCTGGACGGACCGTCGGTGGCGAAAGCGCCTCAGGAGGACGGACCCGACGGTGAGGGACGAAAGCTAGGGTCA |
| p__Thermoplasmatota_g__Natronoplasma_ASV_623 | 0 | 0 | 0 | 0 | 0 | 0 | 1 | 1 | 1 | 0 | 0 | CCTTCACACTGTGGGAAACCACGATGAGGGGATTCCAAGTGCCAACACTAAGTGTTGGCTTTTCATCTGTGTAAAAAGCAGGTGGAATAAGGGCCGGGTAAGACGGGTGCCAGCCGCCGCGGTAACACCCGCGGCCCAAGTGGTGGTCATTATTATTGAGCTTAAAGCGTCCGTAGCCGGTCTGGTAAATCTCTGGGTAAATCGGACGGCTCAACCGTTCGAATACCAGGGAGACTGCTAGACTTGGGGCCGGGAGAGGTCAGAGGTACTCCTGGGGTAGAAGTAAAATTCTGTAATCCTAGGAGGACCACCAGTGGCGAAAGCGTCTGGCTAGAACGGACCCGACGGTGAGGGACGAAGCCCTGGGGCG |
| p__Halobacteriota_g__Methanothrix_A_ASV_626 | 0 | 0 | 0 | 0 | 0 | 0 | 0 | 0 | 1 | 1 | 1 | CCTTTACAATGCGGGAAACCGTGATAAGGGGACCTCGAGTGCTGATGCCCAGCATCGGCTGTCTTTATGCCTAAAAAGCATCTGTTAGCAAGGGCCGGGCAAGACCGGTGCCAGCCGCCGCGGTAACACCGGCGGCTCGAGTGGTAGCCAATATTATTGGGTCTAAAGCGTCCGTAGCCGGCTTGGCAAGTCTTCTGGGAAATCTGGCAGCTTAACTGTCAGGCGTCCAGGAGATACTGTTAGGCTTGGGACCGGGAGAGGTGAGAGGTACCTCGGGGGTAGGGGTGAAATCTTGTGATCCTTGAGGGACCACCAGTGGCGAAGGCGTCTCACTAGAACGGCTCCGACGGTCAGGGACGAAAGCTAGGGGCA |
| p__Halobacteriota_g__Natronomonas_ASV_630 | 0 | 1 | 1 | 0 | 0 | 0 | 1 | 0 | 0 | 0 | 0 | CCTTTACACTGCACGCCAGTGCGATAGGGGGACACCGAGTGCGAGGGCATAGAGCCCTCGCTTTTCTGTACCGTAAGGTGGTACAGGAACAAGGGCTGGGCAAGACCGGTGCCAGCCGCCGCGGTAACACCGGCAGCCCAAGTGATGGCCGATATTATTGGGCCTAAAGCGTCCGTAGCTGGCCGGGCAAGTCCGTCGGGAAATCCACGCGCTTAACGCGTGGGCGTCCGGCGGAAACTGTTCGGCTTGGGACCGGAAGATCCGAGGGGTACGTCCGGGGTAGGAGTGAAATCCCGTAATCCTGGACGGACCACCGATGGCGAAAGCACCTCGGAAGGACGGATCCGACAGTGAGGGACGAAAGCTAGGGTCT |
| p__Thermoplasmatota_g__EX4484-6_ASV_637 | 1 | 1 | 1 | 0 | 0 | 0 | 0 | 0 | 0 | 0 | 1 | ACTTCACAATGCGCGCAAGCGCGATGAGGGGATTCCAAGTGCATGCACTATGTGTATGCTGTTTACTAGTCTAAAAAGCTAGTAGAGTAAGGGCTGGGTAAGACTGGTGCCAGCCGCCGCGGTAATACCAGCGGCCCTAGTGGTGATCGTTGTTATTGGGCCTAAAGCGTCCGTAGCCGGCCCGGTAAATCTCTGGGTAAATCGTTGGGCTTAACCTAGCGAATTCTGGGGAGACTGCCGGGCTTGGGACCGGGAGAGGTTGGAGGTACTCCAGGGGTAGGGGTGAAATCCTGTAATCCTTGGGGGACCACCGGTGGCGAAAGCGTCCAACCAGAACGGGTCCGACGGTAAGGGACGAAGCCCTGGGGCG |
| p__Halobacteriota_f__Halobacteriaceae_4595_ASV_640 | 0 | 1 | 1 | 0 | 0 | 0 | 0 | 0 | 0 | 0 | 1 | CCTTTACACTGTACGCAAGTGCGATAAGGGGACCCCGAGTGCGAGGGCATACAGTCCTCGCTTTTGTGTACCGTAAGGAGGTACACGAATAAGGGCTGGGCAAGACCGGTGCCAGCCGCCGCGGTAATACCGGCAGCCCTAGTGATGGCCGCTATTATTGGGCCTAAAGCGTCCGTAGCCTGCCGGACAAGTCCGTTGGGAAATCCGTTCGCTTAACGAACGGGCGTCCAGCGGAAACTGTCCGGCTTGGGACCGGAAGACCCGAGGGGTACGTCCGGGGTAGGAGTGAAATCCTGTAATCCTGGACGGACCGCCGGTGGCGAAAGCGCCTCGGGAAGACGGATCCGACGGTGAGGGACGAAAGCCAGGGTCT |
| p__Nanoarchaeota_g__CG1-02-57-44_ASV_643 | 1 | 0 | 0 | 0 | 0 | 1 | 1 | 0 | 0 | 0 | 0 | CCTTTACAATGCACGCAAGTGTGATAAGGGGATCCCCAGTGCTCATGTACTGCATGGGCTTTTGCCAAGTGTAAATAGCTTGGCGAATAAGTGGTGGGTAAGACCGGTGCCAGCCGCCGCGGTAACACCGGCGCCACGAGTGGTAACCGCGTTTATTGGGCCTAAAGCGTCCGTAGCCGGCCTGATGAATCCTTTGTGAAATTGTCCAGCCTAACTGGGCAGCGCGCAAGGGACACTATCAGGCTTGAGACCGGGAGGAGTCAGAAGTATTCCAGGGGGAGCGGTAAAATGTTATAATCCCTGGAGGACTACCTGTGGCGAAAGCGTCTGACTAGAACGGATCTGACGGTGAGGGACGAAAGCCAGGGGAG |
| p__Halobacteriota_g__Natrinema_ASV_652 | 0 | 1 | 1 | 0 | 0 | 0 | 0 | 0 | 0 | 0 | 0 | CCTTTACACTGCACGCGAGTGCGATAAGGGGACTCCAAGTGCGAGGGCATATAGTCCTCGCTTTTCACCACCGTAAGGTGGTGGTGGAATAAGTGCTGGGCAAGACCGGTGCCAGCCGCCGCGGTAATACCGGCAGCACGAGTGATGACCGCTATTATTGGGCCTAAAGCGTCCGTAGCTGGCCGTGCAAGTCCATCGGGAAATCCGCGCGCTCAACGCGCGGGCGTCCGGTGGAAACTGTACGGCTTGGGACCGGAAGACCAGAGGGGTACGTCCGGGGTAGGAGTGAAATCCCGTAATCCTGGACGGACCACCGGTGGCGAAAGCGCCTCTGGAAGACGGATCCGACGGTGAGGGACGAAAGCTCGGGTCA |
| p__Halobacteriota_g__Saliphagus_ASV_653 | 0 | 0 | 1 | 0 | 0 | 0 | 0 | 0 | 0 | 0 | 0 | CCTTTACACTGCGCGACAGCGCGATAAGGGGACTCCAAGTGCGAGGGCATATCGTCCTCGCTTTTCACTATCGTAAGGAGATGGTAGAATAAGGGCTGGGCAAGACCGGTGCCAGCCGCCGCGGTAATACCGGCAGCCCGAGTGATGACCGCTATTATTGGGCCTAAAGCGTCCGTAGCCGGCCGAACAAGTCCGTCGGGAAATCCGCACGCCTAACGTGCGGGCGTCCGGCGGAAACTGTTCGGCTTGGGACCGGAAGATCCAGAGGGTACGTCCGGGGTAGGAGTGAAATCCCGTAATCCTGGACGGACCACCGGTGGCGAAAGCGCTCTGGAAGGACGGATCCGACGGTGAGGGACGAAAGCTAGGGTCA |
| p__Halobacteriota_g__Halomicrobium_3849_ASV_660 | 0 | 0 | 1 | 0 | 1 | 0 | 0 | 0 | 0 | 0 | 0 | CCTTTACACTGCACGACAGTGCGATAAGGGGACTCCGAGTGCGAGGGCATATAGTCCTCGCTTTTCTGTACCGTAAGGTGGTACAGGAACAAGTGCTGGGCAAGACCGGTGCCAGCCGCCGCGGTAATACCGGCAGCACGAGTGATGGCCGCTATTATTGGGCCTAAAGCGTCCGTAGCCGGCCAGACAAGTCCGTTGGGAAATCGACGCGCTCAACGCGTCGGCGTCCAGCGGAAACTGTCTGGCTTGGGGCCGGAAGACCTGAGGGGTACGTCCGGGGTAGGAGTGAAATCCCGTAATCCTGGACGGACCACCGGTGGCGAAAGCGCCTCAGGAAGACGGACCCGACGGTGAGGGACGAAAGCTAGGGTCT |
| p__Halobacteriota_f__Haloarculaceae_3992_ASV_667 | 1 | 1 | 1 | 0 | 1 | 0 | 0 | 0 | 0 | 0 | 0 | CCTTTACACTGCACGACAGTGCGATAGGGGGACTCCGAGTGCGAGGGCATATCGTCCTCGCTTTTCTCGACCGTAAGGTGGTCGAGGAATAAGGACTGGGCAAGACCGGTGCCAGCCGCCGCGGTAATACCGGCAGTCCGAGTGATGGCCGCTATTATTGGGCCTAAAGCGTCCGTAGCCGGCCGGACAAGTCCGTTGGGAAATCGACGCGCTCAACGCGTCGGCGTCCAGCGGAAACTGTCCGGCTTGGGGCCGAGAGACTCGACGGGTACGTCCGGGGTAGGAGTGAAATCCCGTAATCCTGGACGGACCACCGATGGCGAAAGCACGTCGAGAGATCGGACCCGACGGTGAGGGACGAAAGCTAGGGTCT |
| p__Halobacteriota_g__Saliphagus_ASV_670 | 0 | 1 | 1 | 1 | 0 | 1 | 1 | 0 | 0 | 0 | 0 | CCTTTACACTGCACGACAGTGCGATAGGGGGACTCCAAGTGCGAGGGCATATAGTCCTCGCTTTTGTGTACCGTAAGGAGGTACAGGAATAAGGGCTGGGCAAGACCGGTGCCAGCCGCCGCGGTAATACCGGCAGCCCGAGTGATGACCGCTATTATTGGGCCTAAAGCGTCCGTAGCCGGCCACACAAGTCCGTCGGGAAATCCGCACGCCCAACGTGCGGGCGTCCGGCTGAAACTGTGTGGCTTGGGACCGGAAGACTCGAGGGGTACGTCCGGGGTAGGAGTGAAATCCTGTAATCCTGGACGGACCGCCGGTGGCGAAAGCGCCTCGAGAGGACGGATCCGACGGTGAGGGACGAAAGCTAGGGTCA |
| p__Halobacteriota_f__Haloarculaceae_3993_ASV_679 | 0 | 1 | 1 | 0 | 0 | 0 | 0 | 0 | 0 | 0 | 0 | CCTTTACACTGCACGACAGTGCGATAGGGGGACTCCGAGTGCGAGGGCATATCGTCCTCGCTTTTCTCGACCGTAAGGTGGTCGAGGAACAAGTGCTGGGCAAGACCGGTGCCAGCCGCCGCGGTAATACCGGCAGCACGAGTGATGGCCAATATTATTGGGCCTAAAGCGTCCGTAGCCGGCCGGACAAGTCCGTTGGGAAATCGACGCGCTCAACGCGTCGGCGTCCAGCGGAAACTGTCCGGCTTGGGGCCGGAAGACCTGGGGGGTACGTCCGGGGTAGGAGTGAAATCCCGTAATCCTGGACGGACCACCGGTGGCGAAAGCGCCTCAGGAAGACGGACCCGACGGTGAGGGACGAAAGCTAGGGTCT |
| p__Halobacteriota_g__Halorubrum_ASV_681 | 0 | 0 | 1 | 1 | 0 | 0 | 0 | 0 | 0 | 0 | 0 | CCTTTACACTGCACGACAGTGCGATAGGGGGATCCCAAGTGCACAGGCATAGTGCCTGTGCTTTTCGACACCCTAAGGCGGTGTCAGAATAAGGGCTGGGCAAGACCGGTGCCAGCCGCCGCGGTAATACCGGCAGCCCAAGTGATGGCCGATCTTATTGGGCCTAAAGCGTCCGTAGCTGGCCGCGCAAGTCCATCGGGAAATCCACCTGCTCAACAGGTGGACGCCCGGTAGAAACTGCGTGGCTTGGGACCGGAAGGCGCGACGGGTACGTCCGGGGTAGGAGTGAAATCCCGTAATCCTGGACGGACCACCGATGGCGAAAGCACGTCGCGAGGACGGATCCGACAGTGAGGGACGAAAGCCAGGGTCT |
| p__Halobacteriota_g__Halorussus_ASV_682 | 0 | 0 | 1 | 1 | 0 | 0 | 0 | 0 | 0 | 1 | 0 | CCTTTACACTGCACGACAGTGCGATAAGGGGACTCCGAGTGCGAGGGCATATAGTCCTCGCTTTTGTACACCGTAAGGTGGTGTACGAATAAGGACTGGGCAAGACCGGTGCCAGCCGCCGCGGTAATACCGGCAGTCCGAGTGATGGCCGCTATTATTGGGCCTAAAGCGTCCGTAGCCGGCCAGACAGGTCCGTCGGGAAATCTGCTCGCTCAACGAGCAGGCGTCCGGCGGAAACCAGCTGGCTTGGGGCCGGAAGACTCGAGGGGTACGTCCGGGGTAGGAGTGAAATCCCGTAATCCTGGACGGACCACCGGTGGCGAAAGCGCCTCGAGAAGACGGACCCGACGGTGAGGGACGAAAGCTAGGGTCA |
| p__Halobacteriota_g__Halovenus_ASV_683 | 0 | 1 | 1 | 1 | 0 | 0 | 0 | 0 | 0 | 0 | 0 | CCTTTACACTGCACGACAGTGCGATAGGGGGACTCCGAGTGCGAGGGCATATAGTCCTCGCTTTTCTGTACCGTAAGGAGGTACAGGAACAAGTGCTGGGCAAGACCGGTGCCAGCCGCCGCGGTAATACCGGCAGCACAAGTGATGGCCGATCTTATTGGGCCTAAAGCGTTCGTAGCCGGCCGGACAAGTTCGTTGGGAAATCCACGCGCTTAACGCGTGGGCGTCCAGCGAAAACTGTCTGGCTTGGGGCCGGAAGACTCGAGGGGTACGTCCGGGGTAGGAGTGAAATCCCGTAATCCTGGACGGACCGCCGATGGCGAAAGCACCTCGAGAGGACGGACCCGACGGTGAGGGACGAAAGCCAGGGTCT |
| p__Thermoproteota_g__BA2_ASV_686 | 0 | 0 | 0 | 0 | 0 | 0 | 0 | 0 | 1 | 1 | 1 | ACTCCGCAATGCACGAAAGTGTGACGGGGCTACCCCGAGTGCCGTCCGCTGAGGATGGCTTTTCCCTGGTTTAACTAGCCAGGGGAATAAGGAGAGGGCAAGACTGGTGTCAGCCGCCGCGGTAATACCAGCTCTCCGAGTGGTAGGGATGATTATTGGGCTTAAAGCGTCCGTAGCCGGCTTAGCAAGTCTCCTGTTAAATTCAGCGACCTAATCGTTGAGCTGCGGGAGATACTACTAGGCTAGGAGGCGGGAGAGGTCGACGGTACTTCTGGGGTAGGGGTGAAATCCTATAATCCCGGGAGGACCACCAGTGGCGAAGGCTGTTGACTAGAACGCGCTCGACGGTGAGGGACGAAAGCTGGGGGAG |
| p__Halobacteriota_g__Halalkalicoccus_ASV_688 | 0 | 1 | 0 | 0 | 0 | 0 | 0 | 0 | 0 | 0 | 0 | CCTTTACACTGCACGAACGTGCGATAAGGGGACTCCAAGTGCGAGGGCATATAGTCCTCGCTTTTGCCTACCGTAAGGTGGTAGGCGAATAAGAGCTGGGCAAGACCGGTGCCAGCCGCCGCGGTAATACCGGCAGCTCGAGTGATGGCCAATATTATTGGGCCTAAAGCGTCCGTAGCTGGCCGAACAGGTCCGTCGGGAAATCCGCTCGCTCAACGAGCGGACGTCCGGCGGAAACCAGTCGGCTTGGGACCGGAAGACCCGAGGGGTACGTCCGGGGTAGGAGTGAAATCCTGTAATCCTGGACGGACCACCGGTGGCGAAAGCGCCTCGGGAGGACGGATCCGACAGTGAGGGACGAAAGCTTGGGTCA |
| p__Halobacteriota_g__Halorussus_ASV_689 | 0 | 1 | 1 | 0 | 0 | 1 | 0 | 0 | 0 | 0 | 0 | ACTTTACACTGCACGCGAGTGCGATAAGGGGACTCCGAGTGCGAGGGCATAGAGCCCTCGCTTTTCTACACCGTAAGGTGGTGTAGGAATAAGAGCTGGGCAAGACCGGTGCCAGCCGCCGCGGTAATACCGGCAGCTCGAGTGATGGCCGATTTTATTGGGCCTAAAGCGTCCGTAGCTTGCCAGACAGGTCTATCGGGAAATCTACCCGCTTAACGGGTAGGCGTCCGGTAGAAACCAGCTGGCTTGGGGCCGGAAGATCCAAGGGGTACGTCCGGGGTAGGAGTGAAATCCCGTAATCCTGGACGGACCACCGATGGCGAAAGCACCTTGGAAAGACGGACCCGACGGTGAGGGACGAAAGCTAGGGTCA |
| p__Halobacteriota_f__Natrialbaceae_ASV_690 | 0 | 1 | 1 | 0 | 0 | 0 | 0 | 0 | 0 | 0 | 0 | CCTTTACACTGCACGCCAGTGCGATAAGGGGACTCCAAGTGCCAGGGCATATCGTCCTGGCTTTTCACCACCGTAGGGTGGTGGTGGAATAAGGGCTGGGCAAGACCGGTGCCAGCCGCCGCGGTAATACCGGCAGCCCGAGTGATGGCCGCTCTTATTGGGCCTAAAGCGTCCGTAGCTGGCCTCGCAAGTCCGTCGGGAAATCCGCACGCTTAACGTGCGGGCGTCCGGTGGAAACTGCGTGGCTTGGGACCGGAAGACCCGAGGGGTACGTCCGGGGTAGGAGTGAAATCCTGTAATCCTGGACGGACCACCGGTGGCGAAAGCGCCTCGGGAGGACGGATCCGACGGTGAGGGACGAAAGCTAGGGTCA |

| | Presence (1) or Absence (0) in each layer | | | | | | | | | | |
|---|---|---|---|---|---|---|---|---|---|---|---|
| Phylum_Genus_ASV_number | L1 | L2 | L3 | L4 | L5 | L6 | L7 | L8 | L9 | L10 | L11 | Sequence |
| p__Halobacteriota_o__Halanaerobiales_ASV_687 | 0 | 1 | 1 | 0 | 0 | 0 | 0 | 0 | 0 | 0 | 1 | CCTTTACACTGCGCGCAAGCGCGATAAGGGGACCCCAAGTGCGAGGGCATACAGTCCTCGCTTTTCACGACCGTAAGGAGGTCGTCGAATAAGGGCTGGGCAAGACCGGTGCCAGCCGCCGCGGTAATACCGGCAGCCCAAGTGATGGCCGCTCTTATTGGGCCTAAAGCGTCCGTAGCTGGCCGTGCAAGTCCGTCGGGAAATCCACCAGCTCAACTGGTGGGCGGCCGACGGAACCTGCATGGCTTGGGACCGGAAGACCCACCGGGTACGTCCGGGGTAGGAGTGAAATCCCGTAATCCCGGACGGACCACCGATGGCGAAAGCACGGTGGGAGGACGGCTCCGACAGTGAGGGACGAAAGCTGGGGTCT |
| p__Halobacteriota_g__Halanaeroarchaeum_ASV_698 | 0 | 0 | 0 | 0 | 0 | 1 | 1 | 0 | 0 | 0 | 0 | CCTTTACACTGCACGACAGTGCGATAAGGGAACTCCGAGTGCGAGGGCATAGAGCCCTCGCTTTTGTACACCGTAAGGTGGTGTACGAATAAGGGCTGGGCAAGACCGGTGCCAGCCGCCGCGGTAATACCGGCAGCCCAAGTGATGGCCGATATTATTGGGCCTAAAGCGTCCGTAGCTGGCCGGACAAGTCCGTTGGGAAATCCGCGCGCTCAACGCGCGGGCGTCCAGCGGAAACTGTCCGGCTTGGGGCCGGAAGACCTGAGGGGTACGTCCGGGGTAGGAGTGAAATCCTGTAATCTCGGACGGACCGCCTGTGGCGAAAGCGCCTCAGGAAGACGGACCCGACAGTGAGGGACGAAAGCCAGGGTCT |
| p__Halobacteriota_g__Halovenus_ASV_702 | 0 | 1 | 1 | 1 | 0 | 0 | 1 | 0 | 0 | 0 | 0 | CCTTTACACTGCACGACAGTGCGATAGGGGGACTCCGAGTGCGAGGGCATAGAGCCCTCGCTTTTGTACACCGTAAGGTGGTGTACGAATAAGAACTGGGCAAGACCGGTGCCAGCCGCCGCGGTAATACCGGCAGTTCGAGTGATGGCCGATCTTATTGGGCCTAAAGCGTTCGTAGCTGGCCAGTCAAGTCTGTCGGGAAACCTGCCCGCTCAACGGGCAGTCGTCCGGCGGAAACTGATTGGCTTGGGACCGGGAGACTCGAGGAGTACGTCCGGGGTAGGAGTGAAATCCCGTAATCCTGGACGGACTACCGATGGCGAAAGCACCTCGAGAGAACGGATCCGACAGTGAGGGACGAAAGCTGGGGTCT |
| p__Thermoproteota_g__PALSA-986_ASV_701 | 0 | 0 | 0 | 0 | 0 | 0 | 0 | 0 | 1 | 1 | 1 | ACTTTGCAATAAGCGCAAGCTTGACAGGGCTATCCCGAGTGCCATCCGCTGAGGAAGGCTTTTACCCAGTCTAGAACGCTGGGGGAATAAGGAGAGGGCAAGTCTGGTGTCAGCCGCCGCGGTAATACCAGCTCTCCGAGTGGTGTGGATGTTTATTGGGCCTAAAGCATCCGTAGCTGGCTAGGTTAGTCCCCTGTTAAATCCACCGAATTAATCGTTGGATTGCGGGGGATACTGCTTGGCTAGGGGACGAGAGAGGCAGACGGTATTTTCGGGGTAGGGGTGAAATCCTATAATCCCGGGAAGACCACCAGTGGCGAAGGCTGTCTGCTAGAACGCGCCCGACGGTGAGGGATGAAAGCTGGGGGAG |
| p__Halobacteriota_g__Halalkaliarchaeum_ASV_705 | 0 | 0 | 1 | 0 | 0 | 0 | 0 | 0 | 0 | 0 | 0 | CCTTTACACTGCACGCCAGTGCGATAAGGGGACCCCGAGTGCGAGGGCATATCGTCCTCGCTTTTGTGTACCGTAGGGAGGTACACGAACAAGGGCTGGGCAAGACCGGTGCCAGCCGCCGCGGTAATACCGGCAGCCCGAGTGATGGCCACTGTTATTGGGCCTAAAGCGTCCGTAGCCGGCCTGGCAAGTCCGTCGGGAAATCCGCGCGCTCAACGCGCGGGCGTCCGGCGGAACCTGCCGGGCTCGGGACCGGAGGACCCAGCGGGTACGTCCGGGGTAGGAGTGAAATCCTGTAATCCCGGACGGACCGCCGATGGCGAAAGCACGCTGGGAAGACGGATCCGACGGTGAGGGACGAAAGCTAGGGTCT |
| p__Halobacteriota_f__Natrialbaceae_ASV_703 | 0 | 0 | 1 | 1 | 0 | 0 | 0 | 0 | 0 | 0 | 1 | CCTTTACACTGCACGCCAGTGCGATAAGGGGACTCCATCTGCGAGGGCATATCGTCCTCGCTTTTCTGCACCGTAAGGTGGTGCAGGAATAAGGGCTGGGCAAGACCGGTGCCAGCCGCCGCGGTAATACCGGCAGCCCGAGTGATGACCGCTATTATTGGGCCTAAAGCGTCCGTAGCTGGCCAGGCAAGTCCATCGGGAAATCCGCGTGCCCAACACGCGGGCGTCCGGTGGAACCTGCCTGGCTTGGGACCGGAAGACCAGAGGGGTACGTCCGGGGTAGGAGTGAAATCCCGTAATCCTGGACGGACCACCGGTGGCGAAAGCGCCTCTGGAAGACGGATCCGACGGTGAGGGACGAAAGCTCGGGTCA |
| p__Halobacteriota_g__Haloprofundus_ASV_708 | 0 | 0 | 0 | 0 | 0 | 1 | 0 | 0 | 0 | 1 | 1 | CCTTTACACTGCACGACAGTGCGACAAGGGGACCCCGAGTGCGAGGGCATACAGTCCTCGCTTTTGTGAACCGTAAGATGGTTCACGAATAAGAGCTGGGCAAGACCGGTGCCAGCCGCCGCGGTAATACCGGCAGCTCGAGTGATGACCGCTGTTATTGGGCCTAAAGCGTCCGTAGCTGGCCGCGTAAGTTCATCGGGAAATCCATCCGCTCAACGGATGGGCGTCCGGTGAAAACTACACGGCTTGGAGCCGGAAGACCCAAGGGGTACGTCCGGGGTAGGAGTGAAATCCTGTAATCCTGGACGGACCGCCGATGGCGAAAGCACCTTGGGAAAACGGACTCGACAGTGAGGGACGAAAGCTAGGGTCT |
| p__Halobacteriota_o__Halanaerobiales_ASV_710 | 0 | 0 | 1 | 0 | 0 | 0 | 0 | 0 | 0 | 0 | 0 | CCTTTACACTGCACGAAAGTGCGATAGGGGGACCCCGAGTGCGAGGGCATATCGTCCTCGCTTTTCTGCACCGTAAGGCGGTGCAGGAATAAGGGCTGGGCAAGACCGGTGCCAGCCGCCGCGGTAATACCGGCAGCCCGAGTGATGACCGCTATTATTGGGCCTAAAGCGTCCGTAGCCGGCCGGACAAGTCCGTCGGGAAATCCACCCGCTCAACGGGTGGGCGTCCGGCGGAAACTGTTCGGCTTGGGGCCGGAGGACCTGAGGGGTACGTCCGGGGTAGGAGTGAAATCCCGTAATCCTGGACGGACCACCGGTGGCGAAAGCGCCTCAGGAGGACGGACCCGACGGTCAGGGACGAAAGCTAGGGTCT |
| p__Halobacteriota_g__Halorussus_ASV_718 | 0 | 1 | 1 | 0 | 0 | 0 | 0 | 0 | 0 | 0 | 0 | CCTTTACACTGCACGACAGTGCGATAAGGGGACTCCGAGTGCGAGGGCATATAGTCCTCGCTTTTCTGCACCGTAAGGTGGTACAGGAATAAGGGCTGGGCAAGACCGGTGCCAGCCGCCGCGGTAATACCGGCAGCCCGAGTGATGGCCGCTATTATTGGGCCTAAAGCGTCCGTAGCCGGCCAGACAGGTCCGTCGGGAAATCCACGCGCTCAACGCGTGGGCGTCCGGCGGAAACCAGCTGGCTTGGGGCCGGAAGACCTGAGGGGTACGTCCGGGGTAGGAGTGAAATCCCGTAATCCTGGACGGACCACCGGTGGCGAAAGCGCCTCAGGAAGACGGACCCGACGGTGAGGGACGAAAGCTAGGGTCA |
| p__Halobacteriota_f__Natrialbaceae_ASV_720 | 0 | 0 | 1 | 0 | 0 | 0 | 0 | 0 | 0 | 0 | 0 | CCTTTACACTGCACGCCAGTGCGATAAGGGGACTCCAAGTGCGAGGGCATACAGTCCTCGCTTTTGTGTACCGTAAGGTGGTACACGAATAAGGGCTGGGCAAGACCGGTGCCAGCCGCCGCGGTAATACCGGCAGCCCGAGTGATGACCGCTATTATTGGGCCTAAAGCGTCCGTAGCCGGCCGGGTAAGTCCGTCGGGAAATCCGCGCGCTCACCGCGCGGGCGTCCGGTGGAAACTGCCTGGCTTGGGACCGGAAGATCCAGAGGGTACGTCCGGGGTAGGAGTGAAATCCTGTAATCCTGGACGGACCACCGGTGGCGAAAGCGCTCTGGAAAGACGGATCCGACGGTGAGGGACGAAAGCTAGGGTCA |
| p__Halobacteriota_g__Natronoarchaeum_ASV_727 | 0 | 1 | 1 | 0 | 1 | 0 | 0 | 0 | 0 | 0 | 0 | CCTTTACACTGCACGCCAGTGCGATAAGGGGATCCCCAGTGCGAGGGCATACAGTCCTCGCTTTTCACCACCGTACGATGGTGGTGGAACAAGAGCTGGGCAAGACCGGTGCCAGCCGCCGCGGTAATACCGGCAGCTCGAGTGATGGCCGTTCTTATTGGGCCTAAAGCGTCCGTAGCCGGCCACGCAAGTTCGTCGGGAAATCCACACGCTCAACGTGTGGGCGTCCGGCGAAACCTGCGTGGCTTGGGACCGGACGGCCCGAGGGGTACGGCCGGGGTAGGAGTGAAATCCTGTAATCCCGGCCGGACCACCGGTGGCGAAAGCGCCTCGGGAGGACGGATCCGACGGTGAGGGACGAAAGCCAGGGTCA |
| p__Halobacteriota_g__Halopiger_A_ASV_730 | 1 | 1 | 0 | 0 | 0 | 0 | 1 | 0 | 0 | 0 | 0 | CCTTTACACTGCACGCCAGTGCGATAAGGGGACTCCATCTGCGAGGGCATATCGTCCTCGCTTTTCTGCACCGTAAGGTGGTGCAGGAATAAGGGCTGGGCAAGACCGGTGCCAGCCGCCGCGGTAATACCGGCAGCCCGAGTGATGACCGCTATTATTGGGCCTAAAGCGTCCGTAGCTGGCCAGGCAAGTCCATCGGGAAATCCGCGTGCCCAACACGCGGGCGTCCGGTGGAACCTGCCTGGCTTGGGACCGGAAGACCAGAGGGGTACGTCTGGGGTAGGAGTGAAATCCCGTAATCCTGGACGGACCACCGGTGGCGAAAGCGCCTCTGGAAGACGGATCCGACGGTGAGGGACGAAAGCTCGGGTCA |
| p__Halobacteriota_g__Halovenus_ASV_735 | 0 | 1 | 1 | 0 | 0 | 1 | 0 | 0 | 0 | 0 | 0 | CCTTTACACTGCACGACAGTGCGATAGGGGGACTCCGAGTGCGAGGGCATACAGTCCTCGCTTTTGTACACCGTAAGGTGGTGTACGAATAAGAACTGGGCAAGACCGGTGCCAGCCGCCGCGGTAATACCGGCAGTTCGAGTGATGGCCGCTTTTATTGGGCCTAAAGCGTTCGTAGCTGGCCGAGCAAGTCTGTCGGGAAATCCAGAGGCTTAACTTCTGGACGTCCGGCGGAAACTGCTCGGCTTGGGACCGGGAGACTCGAGGAGTACGTCCGGGGTAGGAGTGAAATCCTGTAATCCTGGACGGACTACCGATGGCGAAAGCACCTCGAGAGCACGGATCCGACAGTGAGGAACGAAAGCTGGGGTCT |
| p__Halobacteriota_g__Halobacterium_ASV_744 | 0 | 1 | 1 | 1 | 0 | 0 | 0 | 0 | 0 | 0 | 1 | CCTTTACACTGTACGAAAGTGCGATAAGGGGACTCCGAGTGTGAAGGCATAGAGCCTTCACTTTTGTACACCGTAAGGTGGTGTACGAATAAGGACTGGGCAAGACCGGTGCCAGCCGCCGCGGTAATACCGGCAGTCCGAGTGATGGCCGATCTTATTGGGCCTAAAGCGTCCGTAGCTGGCTAGACAAGTCCGTTGGGAAATCTGCCCGCTTAACGGGCAGGCGTCCAGCGGAAACTGTCTAGCTTGGGACCGGAAGACCTGAGGGGTACGTCTGGGGTAGGAGTGAAATCCCGTAATCCTGGACGGACCGCCGGTGGCGAAAGCGCCTCAGGAGAACGGATCCGACAGTGAGGGACGAAAGCTAGGGTCT |
| p__Halobacteriota_g__Natrarchaeobaculum_ASV_741 | 1 | 1 | 1 | 1 | 0 | 0 | 0 | 0 | 0 | 0 | 0 | CCTTTACACTGCGCGACAGCGCGATAAGGGGACTCCATCTGCGAGGGCATATCGTCCTCGCTTTTTCCTACCGTAAGGTGGTAGGCGAATAAGGGCTGGGCAAGACCGGTGCCAGCCGCCGCGGTAATACCGGCAGCCCGAGTGATGACCGCTCTTATTGGGCCTAAAGCGTCCGTAGCTGGCCTCGCAAGTCCATCGGGAAATCCGCACGCCCAACGTGCGGGCGTCCGGTGGAAACTGCGTGGCTTGGGACCGGAAGACCTGAGGGGTACGTCTGGGGTAGGAGTGAAATCCCGTAATCCTGGACGGACCACCGGTGGCGAAAGCGCCTCAGGAGGACGGATCCGACGGTGAGGGACGAAAGCCAGGGTCA |
| p__Thermoproteota_g__PALSA-986_ASV_743 | 0 | 0 | 0 | 0 | 0 | 0 | 1 | 1 | 0 | 0 | 1 | ACTTTGCAATACACGAAAGTGTGACAGGGTCATCCCGAGTGCCGACCGCTGAGGTTGGCTTTTACCCAGTCTAGACAGCTGGGGGAATAAGGAGAGGGCAAGTCTGGTGTCAGCCGCCGCGGTAATACCAGCTCTCCGAGTGGTGTGGACGTTTATTGGGCCTAAAGCATCCGTAGCTTGCTGAGCAAGTCCCCTGTTAAACCCACCGATTTAATCGTTGGTGTGCGGGGGAAACTGTTCGGCTAGGGGACGAGAGAGGCAGACGGTATTCCCGGGGTAGGGGTGAAATCCTATAATCCCGAGAAGACCACCAGTGGCGAAGGCTGTCTGCTAGAACGCGCCCGACGGTGAGGGATGAAAGCTGGGGGAG |
| p__Halobacteriota_g__Natronobiforma_ASV_755 | 0 | 1 | 1 | 1 | 1 | 0 | 0 | 0 | 0 | 0 | 0 | CCTTTACACTGCACGCCAGTGCGATAAGGGGACACCAAGTGCGAGGGCATATCGTCCTCGCTTTTCACTACCGTAAGGTGGTAGTGGAATAAGAGCTGGGCAAGACCGGTGCCAGCCGCCGCGGTAATACCGGCAGCTCGAGTGATGACCGCTATTATTGGGCCTAAAGCGTCCGTAGCTGGCCACACAAGTCCATCGGGAAATCTCCCTGCCTAACAGGGAGGCGTCCGGTAGAAACTGTGTGGCTTGGGACCGGAAGATCCAGAGGGTACGTCTGGGGTAGGAGTGAAATCCTGTAATCCTGGACGGACCACCGGTGGCGAAAGCGCTCTGGAAGGACGGATCCGACAGTGAGGGACGAAAGCTAGGGTCA |
| p__Halobacteriota_o__Halanaerobiales_ASV_754 | 0 | 1 | 1 | 1 | 0 | 0 | 0 | 0 | 0 | 0 | 1 | CCTTTACACTGCGCGAAAGCGCGATAAGGGGACTCCAAGTGCGAGGGCATAAAGTCCTCGCTTTTCTACACCGTAAGGTGGTGTAGGAATAAGGGCTGGGCAAGACCGGTGCCAGCCGCCGCGGTAATACCGGCAGCCCAAGTGATGGCCGCTATTATTGGGCCTAAAGCGTCCGTAGCCGGCCGGGTAAGTCCGTCGGGAAATCCACACGCCTAACGTGTGGGCGTCCGGCGGAAACTGCTCGGCTTGGGACCGGGAGACCCGAGGGGTACGTCTGGGGTAGGAGTGAAATCCTGTAATCCTAGACGGACCGCCGGTGGCGAAAGCGCCTCGGGAGAACGGATCCGACGGTGAGGGACGAAAGCTAGGGTCA |
| d__Archaea_d__Archaea_ASV_756 | 1 | 1 | 0 | 0 | 0 | 0 | 0 | 0 | 0 | 1 | 1 | ACTTTACAATGCGCGAAAGCGTGATAAGGGGATCCCGAGTGTCACAGCCTAAGGTTGTGACTGTGGCGAAGTGTAAAGAGCTTCGCAAGAAAGGGGTGGGTAATACCGGTGCCAGCAGCCGCGGTAATCCCGGAGCCCCAAGTGGTGGCCAGTTTTATTGGGCCTAAAGCGTGCGTAGCCTGATTAGTAAGTTTTCGGTGAAATCGTCCAGCTTAACTGGACGGCGTGCCGAAAATACTGCTAGTCTAGGGATTGGGAGAGGTTAGGGGTATGCCTGGGGGAGGGGTGAAATCCTTTAATCCTAGGTAGACCATCGGTGGCGAAGGCGCCTAACTGGAACGAGTCCGACGGTGAGGCACGAAAGCTAGGGTAG |
| p__Thermoproteota_g__GCA-2726865_ASV_763 | 0 | 0 | 0 | 0 | 0 | 0 | 0 | 0 | 1 | 0 | 1 | CCTCCTCAATACACTAACGTGTGAAGGGGTTACCTCGAGTGCTACCTGAAGAAGGTGGCTTTTCCTTGGTCTAAAAAGCCTTGGGAATAAGAGGGGGGCAAGACTGGTGTCAGCCGCCGCGGTAACACCAGCCCCTCGAGTGGTAGGGATGATTATTTGGCCTAAAGCGTCCGTAGCCTGCCGAGTAAGTTTTCCGTTAAATCCAGCGTCTCAAGCGTTGGACTGCGGGAAATACTGCTTGGCTAGAGAGTGGGAGAGGTAAGCGGTATTCTGGGGGTAGGGGCGAAATCCTTTGATCCCCGGAGGACCACCAGTGGCGAAGGCTGCTTACTAGAACACGTCTGACGGTCAGGGACGAAAGCTGGGGTAG |
| p__Thermoproteota_f__Nitrososphaeraceae_ASV_765 | 0 | 0 | 0 | 0 | 1 | 1 | 0 | 0 | 1 | 0 | 0 | CCTCTGCAATAGGCGAAAGCTTGACAGGGTTACTCTGAGTGATGCCCGCTAAGGGTATCTTTTGGCACCTCTAAAAATGGTGCAGAATAAGGGGTGGGCAAGTCTGGTGTCAGCCGCCGCGGTAATACCAGCACCCCGAGTTGTCGGGACGATTATTGGGCCTAAAGCATCCGTAGCCTGTTCTGCAAGTCCTCCGTTAAATCCACCTGCTTAACGGATGGGCTGCGGGGGATACTGCAGAGCTAGGAGGCGGGAGAGGCAAACGGTACTCAGTGGGTAGGGGTAAAATCCTTTGATCTACTGAAGACCACCAGTGGCGAAGGCGGTTTGCCAGAACGCGCTCGACGGTGAGGGATGAAAGCTGGGGGAG |
| p__Halobacteriota_f__Natrialbaceae_ASV_776 | 1 | 0 | 1 | 0 | 0 | 0 | 0 | 0 | 0 | 0 | 0 | CCTTTACACTGCACGCCAGTGCGATAAGGGGACTCCCAGTGCGAGGGCATATCGTCCTCGCTTTTGTGTACCGTAGGGAGGTACACGAATAAGGGCTGGGCAAGACCGGTGCCAGCCGCCGCGGTAATACCGGCAGCCCAAGTGATGACCGCTATTATTGGGCCTAAAGCGTCCGTAGCCGGCCACGCAAGTCCGTCGGGAAATCCGCACGCCCAACGTGCGGGCGTCCGGCGGAAACTGCCTGGCTTGGGACCGGAAGACCAGAGGGGTACGTCCGGGGTAGGAGTGAAATCCCGTAATCCTGGACGGACCACCGGTGGCGAAAGCGCCTCTGGAAGACGGATCCGACGGTGAGGGACGAAAGCTCGGGTCA |
| p__Halobacteriota_g__Methanolobus_ASV_779 | 0 | 1 | 0 | 0 | 0 | 0 | 0 | 0 | 1 | 1 | 1 | ACTTTACAATGCGGGAAACCGCGATAAGGGGACACCGAGTGCCAGCACCATGTGTTGGCTGTCCGCATGTATAAATAACATGTGTTAGCAAGGGCCGGGCAAGACCGGTGCCAGCCGCCGCGGTAACACCGGCGGCCCGAGTGGTAGCCACTATTATTGGGTCTAAAGGGTCCGTAGCCGGTTTGATCAGTCTTCCGGGAAATCTGACAGCTTAACTGTTAGGCTTCTGGGGGATACTGTCAGACTTGGAACCGGGAGAGGTAAGAGGTACTACAGGGGTAGGAGTGAAATCTTGTAATCCCTGTGGGACCACCAGTGGCGAAGGCGTCTTACCAGAACGGGTTCGACGGTGAGGGACGAAAGCTGGGGGCA |
| p__Thermoplasmatota_g__SM1-50_ASV_784 | 0 | 0 | 0 | 0 | 0 | 0 | 0 | 0 | 1 | 0 | 0 | ACTTCGCACTGCGCGAAAGCGCGACGAGGGAATCCCAAGTGTATGCACGTAAGTGTATACTGTTTCTTTGTCTAAAAAGCAGAGAGAGTAAGAGCTGGGTAAGACGGGTGCCAGCCGCCGCGGTAATACCTGCAGCTCAAGTGGTGATCGTTATTATTGGGCCTAAAACGTCCGTAGCCGGTCTTGTAAATGCCTGGGTAAATTGTATGGCTTAACCATACGAATTCCGGGTAGACTGTTAGACTTGAGACCGGGAGAGGCTAGAGGTACTCCTGGGGTAGAGGTGAAATTCTGTAATCCTGGGGGACCACCAGTGGCGAAGGCGTCTAGCCAGAACGGGTCTGACGGTGAGGGACGAAGCCCTGGGGCG |

| Phylum_Genus_ASV_number | Presence (1) or Absence (0) in each layer | | | | | | | | | | | |
|---|---|---|---|---|---|---|---|---|---|---|---|---|
| | L1 | L2 | L3 | L4 | L5 | L6 | L7 | L8 | L9 | L10 | L11 | Sequence |
| p__Halobacteriota_g__Halorussus_ASV_788 | 0 | 1 | 1 | 0 | 0 | 1 | 0 | 0 | 0 | 0 | 0 | CCTTTACACTGCACGCCAGTGCGATAAGGGGATCCCGAGTGCGAGGGCATACAGTCCTCGCTTTTGTACACCGTAAGGTGGTGTACGAATAAGAGCTGGGCAAGACCGGTGCCAGCCGCCGCGGTAATACCGGCAGCTCGAGTGATGGCCGCTATTATTGGGCCTAAAGCGTCCGTAGCCGGCCGAACAGGTCCGTTGGGAAATCCATCTGCCTAACAGATGGGCGTCCAGCGGAAACCAGTCGGCTTGGGGCCGGAAGACCTGAGGGGTACGTCCGGGGTAGGAGTGAAATCCTGTAATCCTGGACGGACCGCCGGTGGCGAAAGCGCCTCAGGAGGACGGACCCGACGGTGAGGGACGAAAGCTAGGGTCA |
| p__Halobacteriota_g__Salinigranum_ASV_787 | 0 | 1 | 1 | 0 | 0 | 0 | 0 | 0 | 0 | 0 | 1 | CCTTTACACTGCGCGCAAGCGCGATAAGGGGACCCCAAGTGCGAGGGCATACAGTCCTCGCTTTTGTGTACCGTAAGGAGGTACACGAATAAGAGCTGGGCAAGACCGGTGCCAGCCGCCGCGGTAATACCGGCAGCTCGAGTGATGGCCGCTATTATTGGGCCTAAAGCGTCCGTAGCTGGCCACGCAAGTCCGTCGGGAAATCCACCAGCCCAACTGGTGGGCGGCCGACGGAACCTGCGTGGCTTGGGACCGGAAGACCCAACGGGTACGTCCGGGGTAGGAGTGAAATCCCGTAATCCCGGACGGACCACCGATGGCGAAAGCACGTTGGGAGGACGGCTCCGACAGTGAGGGACGAAAGCTGGGGTCT |
| p__Halobacteriota_g__Saliphagus_ASV_790 | 0 | 0 | 1 | 0 | 0 | 0 | 1 | 0 | 0 | 0 | 0 | CCTTTACACTGCACGACAGTGCGATAGGGGGACTCCAAGTGCTAGGGCATATCGTCCTAGCTTTTCACCACCGTAAGGTGGTGGTGGAATAAGTGCTGGGCAAGACCGGTGCCAGCCGCCGCGGTAATACCGGCAGCACAAGTGATGACCGCTATTATTGGGCCTAAAGCGTCCGTAGCCGGCCGTATAAGTCCGTCGGGAAATCCGCACGCCTAACGTGCGGGCGTCCGGCGGAAACTGTTCGGCTTGGGACCGGAAGACTCGAGGGGTACGTCTGGGGTAGGAGTGAAATCCCGTAATCCTGGACGGACCACCGGTGGCGAAAGCGCCTCGAGAGGACGGATCCGACGGTGAGGGACGAAAGCTAGGGTCA |
| p__Halobacteriota_g__Halorussus_ASV_801 | 0 | 0 | 1 | 1 | 0 | 0 | 0 | 0 | 0 | 0 | 0 | CCTTTACACTGTACGAAAGTGCGATAAGGGGACCCCGAGTGCGAGGGCATACAGTCCTCGCTTTTGTGTACCGTAAGATGGTACACGAATAAGAGCTGGGCAAGACCGGTGCCAGCCGCCGCGGTAATACCGGCAGCTCAAGTGATGGCCGCTATTATTGGGCCTAAAGCGTCCGTAGCCGGCCGAACAGGTCCGTTGGGAAATCCGCCAGCCTAACTGGCGGGCGTCCAGCGGAAACCAGTCGGCTTGGGGCCGGAAGACCTGAGGGGTACGTCCGGGGTAGGAGTGAAATCCTGTAATCCTGGACGGACCGCCGGTGGCGAAAGCGCCTCAGGAGGACGGACCCGACGGTGAGGGACGAAAGCTAGGGTCA |
| p__Halobacteriota_g__Halopiger_A_ASV_815 | 0 | 0 | 1 | 0 | 1 | 1 | 0 | 0 | 0 | 0 | 0 | CCTTTACACTGCACGCCAGTGCGATAAGGGGACTCCACCTGCGAGGGCATATCGTCCTCGCTTTTCTGCACCGTAAGGTGGTACAGGAATAAGGGCTGGGCAAGACCGGTGCCAGCCGCCGCGGTAATACCGGCAGCCCGAGTGATGACCGCTATTATTGGGCCTAAAGCGTCCGTAGCTGGCCAGGCAAGTCCATCGGGAAATCCGCGTGCCCAACACGCGGGCGTCCGGTGGAACCTGCCTGGCTTGGGACCGGAAGACCAGAGGGGTACGTCTGGGGTAGGAGTGAAATCCCGTAATCCTGGACGGACCACCGGTGGCGAAAGCGCCTCTGGAAGACGGATCCGACGGTGAGGGACGAAAGCTCGGGTCA |
| p__Thermoplasmatota_g__SM1-50_ASV_811 | 0 | 0 | 0 | 0 | 0 | 0 | 0 | 0 | 1 | 1 | 1 | ACTTCACACTGCGCGAAAGCGTGATGAGGGAATCCCAAGTGTATGCACGTAAGTGTATACTGTTTCTTTGTTTAAAAAGCAGAGAGAGTAAGGGCTGGGTAAGACGGGTGCCAGCCGCCGCGGTAATACCTGCAGCTCAAGTGGTGACCACTATTATTGGGCCTAAAACGTTCGTAGCCTGTCTTTTAAATGCCTGGGTAAATCGGACTGCTTAACAGTTCGAATTCCGGGTAGACTGGAAGACTTGAGACTGGGAGAGGCTAGAGGTACTCCTGGGGTAGAGGTGAAATTCTGTAATCCTAGGGGGACCACCAGTGGCGAAGGCGTCTAGCTAGAACAGGTCTGACGGTGAGGGACGAAGCCCTGGGGCG |
| p__Thermoproteota_g__GCA-2726865_ASV_817 | 1 | 1 | 0 | 0 | 0 | 0 | 1 | 1 | 1 | 0 | 1 | CCTCCTCAATGCACTAACGTGTGAAGGGGTTACCTCGAGTGCTACCTGAAGAAGGTGGCTTTTCCTTGGTCTAAAACGCCTTGGGAATAAGAGGGGGGCAAGACTGGTGTCAGCCGCCGCGGTAACACCAGCCCCTCGAGTGGTGGGGATGATTATTTGGCCTAGAGCGTCCGTAGCCTGTTAAGTAAGTTTCCCGTCAAATCCAATGTCTCAAGCGTTGGTCTGCGGGAAATACTGTTTAACTAGAGAGTGGGAGAGGTCAGCGGTATTCTGGGGGTAGGGGCGAAATCCTTTGATCCCCGGAGGACCACCAGTGGCGAAGGCTGCTGACTAGCACACGTCTGACGGTCAGGGACGAAAGCTGGGGTAG |
| p__Halobacteriota_g__Natrinema_ASV_823 | 0 | 0 | 1 | 0 | 0 | 0 | 0 | 0 | 0 | 0 | 0 | CCTTTACACTGCACGAGAGTGCGATAAGGGGACTCCGAGTGCGAGGGCATATAGCCCTCGCTTTTCACGACCGTAAGGTGGTCGTAGAATAAGTGCTGGGCAAGACCGGTGCCAGCCGCCGCGGTAATACCGGCAGCACGAGTGATGACCGCTATTATTGGGCCTAAAGCGTCCGTAGCTGGCCACGCAAGTCCATCGGGAAATCCGCGCGCTCAACGCGCGGGCGTCCGGTGGAAACTGCGTGGCTTGGGACCGGAAGACCAGAGGGGTACGTCCGGGGTAGGAGTGAAATCCCGTAATCCTGGACGGACCACCGGTGGCGAAAGCGCCTCTGGAAGACGGATCCGACGGTGAGGGACGAAAGCTCGGGTCA |
| p__Nanoarchaeota_p__Nanoarchaeota_ASV_819 | 0 | 0 | 0 | 0 | 0 | 0 | 0 | 0 | 1 | 1 | 0 | TCTCCACAATGCGCGAAAGCGTGATGGGGGAACTCCAAGTGCGCTACCACTGGTAGTGCTGTGGCGAAATGTACAAAGTTTCGCAAGAAAGGGGTGGGCAAGACAGGTGCCAGCAGCCGCGGTAATCCCTGCGCCTCAAGTCGTGACCATGCTTATTGGGCCTAAAGTATCCGTAGCCGGAGTATTAAGTTCTCTGTGAAATCGTACAGCTTAACTGTACGGCGTGCAGGGAATACTGTTATTCTAGGGACCGGGAGGTGTCAAGGGTATTTCTGGGGTAAGGGTTAAATCTTATAATCCCAGGAGGACCAACGGTGGCGAAGGCGCTTGACAAGAACGGGTCCGACGGTGAATGGATGAAAGCTGGGGGAG |
| p__Halobacteriota_o__Halanaerobiales_ASV_821 | 0 | 1 | 1 | 0 | 0 | 0 | 0 | 0 | 0 | 0 | 0 | CCTTTACACTGTACGAAAGTGCGATAAGGGGACCCCGAGTGCGAGGGCATACAGTCCTCGCTTTTGTGTACCGTAAGATGGTACACGAATAAGAGCTGGGCAAGACCGGTGCCAGCCGCCGCGGTAATACCGGCAGCTCAAGTGATGGCCGTTATTATTGGGCCTAAAGCGTCCGTAGCCGGCCGAACAGGTCCGTTGGGAAATCCGCCAGCCTAACTGGCGGGCGTCCAGCGGAAACCAGTTGGCTTGGGGCCGGAAGACCTGAGGGGTACGTCTGGGGTAGGAGTGAAATCCTGTAATCCTGGACGGACCGCCGGTGGCGAAAGCGCCTCAGGAGGACGGACCCGACGGTGAGGGACGAAAGCTAGGGTCA |
| p__Halobacteriota_g__Halodesulfurarchaeum_ASV_833 | 0 | 0 | 1 | 0 | 1 | 1 | 1 | 0 | 0 | 0 | 0 | CCTTTACACTGCACGACAGTGCGATAGGGGGACTCCGAGTGCGAGGGCATAGAGCCCTCGCTTTTCTGCACCGTAAGGTGGTGCAGGAAGAAGGGCTGGGCAAGACCGGTGCCAGCCGCCGCGGTAATACCGGCAGCCCAAGTGATGGCCGATATTATTGGGCCTAAAGCGTCCGTAGCTGGCCGGACAAGTCCGTTGGGAAATCCGCACGCTTAACGTGCGGGCGTCCAGCGGAAACTGTCCGGCTTGGGACCGGAAGACCCGAGGGGTACGTCCGGGGTAGGAGTGAAATCCTGTAATCTCGGACGGACCACCTGTGGCGAAAGCGCCTCGGGAAGACGGATCCGACAGTGAGGGACGAAAGCCAGGGTCT |
| p__Halobacteriota_g__Natrinema_ASV_832 | 0 | 0 | 1 | 0 | 0 | 0 | 0 | 0 | 0 | 0 | 0 | CCTTTACACTGCACGCGAGTGCGATAAGGGGACTCCAAGTGCGAGGGCATATCGTCCTCGCTTTTCGCGACCGTAGGAAGGTCGCAGAATAAGTGCTGGGCAAGACCGGTGCCAGCCGCCGCGGTAATACCGGCAGCACGAGTGATGACCGCTATTATTGGGCCTAAAGCGTCCGTAGCTGGCCGGCCAAGTCCATCGGGAAATCCGCGCGCTCAACGCGCGGGCGTCCGGTGGAAACTGGCTGGCTTGGGACCGGAAGATCCAGAGGGTACGTCCGGGGTAGGAGTGAAATCCCGTAATCCTGGACGGACCACCGGTGGCGAAAGCGCTCTGGAAAGACGGATCCGACGGTGAGGGACGAAAGCTCGGGTCA |
| p__Halobacteriota_g__Halodesulfurarchaeum_ASV_838 | 0 | 1 | 1 | 1 | 0 | 0 | 0 | 1 | 1 | 0 | 1 | CCTTTACACTGCACGACAGTGCGATAAGGGAACTCCGAGTGCGAGGGCATAGAGCCCTCGCTTTTGTACACCGTAAGGTGGTGTACGAATAAGGGCTGGGCAAGACCGGTGCCAGCCGCCGCGGTAATACCGGCAGCCCAAGTGATGGCCGATATTATTGGGCCTAAAGCGTCCGTAGCCGGCCGGACAAGTCCGTTGGGAAATCCGCACGCTCAACGTGCGGGCGTCCAGCGGAAACTGTTCGGCTTGGGACCGGAAGACCCGAGGGGTACGTCCGGGGTAGGAGTGAAATCCTGTAATCCCGGACGGACCGCCTGTGGCGAAAGCGCCTCGGGAAGACGGATCCGACGGTGAGGGACGAAAGCCAGGGTCT |
| p__Halobacteriota_g__Natrinema_ASV_840 | 0 | 1 | 1 | 0 | 0 | 0 | 0 | 0 | 0 | 0 | 0 | CCTTTACACTGCACGCGAGTGCGATAAGGGGACTCCGAGTGCGAGGGCATATAGTCCTCGCTTTTCACCACCGTAAGGAGGTGGTGGAATAAGTGCTGGGCAAGACCGGTGCCAGCCGCCGCGGTAATACCGGCAGCACGAGTGATGACCGCTATTATTGGGCCTAAAGCGTCCGTAGCTGGCCGTGCAAGTCCATCGGGAAATCCGCGCGCTCAACGCGCGGGCGTCCGGTGGAAACTGTACGGCTTGGGACCGGAAGACCAGAGGGGTACGTCCGGGGTAGGAGTGAAATCCCGTAATCCTGGACGGACCACCGGTGGCGAAAGCGCCTCTGGAAGACGGATCCGACGGTGAGGGACGAAAGCTCGGGTCA |
| p__Halobacteriota_o__Halanaerobiales_ASV_839 | 0 | 1 | 1 | 0 | 0 | 0 | 0 | 0 | 0 | 0 | 0 | CCTTTACACTGCACGACAGTGCGATAAGGGGACTCCGAGTGCGAGGGCATATAGTCCTCGCTTTTGTCGACCGTAAGGTGGTCGACGAATAAGGACTGGGCAAGACCGGTGCCAGCCGCCGCGGTAATACCGGCAGTCCGAGTGATGGTCGATATTATTGGGCCTAAAGCGTCCGTAGCTGGCCGAGCAAGTCTATCGGGAAATCCAGTAGCTCAACTACTGGACGTCCGGTGGAAACTGTTCGGCTTGGGACCGGAAGATCCAAGGGGTACGTCCGGGGTAGGAGTGAAATCCTGTAATCCTGGACGGACCACCGATGGCGAAAGCACCTTGGAAGCACGGATCCGACAGTGAGGGACGAAAGCTAGGGTCT |
| p__Nanoarchaeota_g__CG1-02-57-44_ASV_842 | 1 | 0 | 0 | 1 | 0 | 0 | 0 | 0 | 0 | 0 | 0 | CCTTCACAATTCACGAAAGTGTGATGAGGTAACTCCTAGTGCTCATGGAAACATGGGCTTTTCTTAAAGGTAGTGTCTTTAAGGAATAAGCAGTGGGTAAGATCGGTGCCAGCCGCCGCGGTAATACCGACACTGCCAGTGGCAACCGCATTTATTGGGCCTAAAGCGTCCGTAGCCGGCTTGGTAAGTCTTTTGTGAAATCATAGGACTTAATCTTATGGCGAGCAGAGGATACTGTCAAGCTAGGGACCGGGAGGCGTCAAAGGTATTCCTTGGGAAGCGGTAAAATGTGATAATCCTTGGAAGACCACCTGTGGTGAAAACGTTTGGCGAGAACGGGTCCGACGGTGAGGGTCGAAAGCCAGGGGAG |
| p__Halobacteriota_g__Halohasta_ASV_845 | 0 | 0 | 1 | 0 | 0 | 0 | 0 | 0 | 0 | 0 | 1 | CCTTTACACTGCACGCAAGTGCGATAGGGGAATCCCAAGTGCGAGGGCATACTAGCCCTCGCTTTTGTGTACTGTAAGGAGGTACACGAATAAGAGCTGGGCAAGACCGGTGCCAGCCGCCGCGGTAATACCGGCAGCTCGAGTGATGTCCACTATTATTGGGCCTAAAGCGTCCGTAGCTGGCCGTACAAGTCCGTCGGGAAATCCACCTGCCCAACAGGTGGGCGTCCGGCGGAAACTGTGCGGCTTGGAACCGGAAGGCTCAGAGGGTACGTCCGGGGTAGGAGTGAAATCCTGTAATCCCGGACGGACCACCGATGGCGAAAGCACTCTGAGAAGACGGATTCGACAGTGAGGGACGAAAGCTAGGGTCT |
| p__Halobacteriota_g__Halorussus_ASV_847 | 0 | 1 | 1 | 0 | 1 | 0 | 0 | 0 | 0 | 0 | 0 | CCTTTACACTGCACGACAGTGCGATAAGGGGACTCCGAGTGCGAGGGCATATAGTCCTCGCTTTTGTACACCGTAAGGTGGTGTACGAATAAGGACTGGGCAAGACCGGTGCCAGCCGCCGCGGTAATACCGGCAGTCCGAGTGATGGCCGCTATTATTGGGCCTAAAGCGTCCGTAGCCGGCCAGACAGGTCCGTCGGGAAATCCACGCGCTCAACGCGTGGGCGTCCGGCGGAAACCAGCTGGCTTGGGGCCGGAAGACTCGAGGGGTACGTCCGGGGTAGGAGTGAAATCCCGTAATCCTGGACGGACCACCGGTGGCGAAAGCGCCTCGAGAAGACGGACCCGACGGTGAGGGACGAAAGCTAGGGTCA |
| p__Halobacteriota_g__Saliphagus_ASV_846 | 0 | 1 | 1 | 0 | 0 | 0 | 0 | 0 | 0 | 0 | 0 | CCTTTACACTGCACGACAGTGCGATAAGGGGACTCCAAGTGCGAGGGCATATCGTCCTCGCTTTTCACTACCGTAAGGTGGTAGTGGAATAAGGGCTGGGCAAGACCGGTGCCAGCCGCCGCGGTAATACCGGCAGCCCGAGTGATGACCGCTATTATTGGGCCTAAAGCGTCCGTAGCCGGCCACGCAAGTCCGTCGGGAAATCCGCACGCCTAACGTGCGGGCGTCCGGCGGAAACTGCGTGGCTTGGGACCGGAAGACTCGAGGGGTACGTCCGGGGTAGGAGTGAAATCCTGTAATCCTGGACGGACCACCGGTGGCGAAAGCGCCTCGAGAGGACGGATCCGACGGTGAGGGACGAAAGCTAGGGTCA |
| p__Halobacteriota_o__Halanaerobiales_ASV_848 | 0 | 1 | 1 | 0 | 0 | 0 | 1 | 0 | 0 | 0 | 0 | CCTTTACACTGCACGACAGTGCGATAAGGGGACTCCGAGTGCGAGGGCATATAGTCCTCGCTTTTCTGTACCGTAGGGAGGTACAGGAACAAGTGCTGGGCAAGACCGGTGCCAGCCGCCGCGGTAATACCGGCAGCACGAGTGATGACCGATATTATTGGGCCTAAAGCGTCCGTAGCTGGCCAGACAAGTCCGTTGGGAAATCCACGCGCTCAACGCGTGGGCGTCCGGCGGAAACTGTTTGGCTTGGGGCCAGAAGACCTGGGGGGTACGTCCGGGGTAGGAGTGAAATCCTGTAATCCTGGACGGACCACCAATGGGGAAACCACCTCAGGAGGATGGACCCGACAGTGAGGGACGAAAGCTAGGGTCT |
| p__Nanoarchaeota_g__BMS3ABIN17_ASV_859 | 0 | 0 | 0 | 0 | 0 | 0 | 0 | 0 | 1 | 1 | 1 | CTTTTGCAATGCGCGAAAGCGTGACAAAGCAAGCCAGAGTGTTTCTTTTATTAAAGAAACTTTTGTCAGCTGTAAAAAGGCTGACGAATAAGGACTGGGTAAGACCGGTGCCAGCCGCCGCGGTAATACCGGCGGTCCAAGTCGCAGCCATCATTATTGGGTCTAAAACATCCGTAGCTTGTTTTGCAAGTCTCTTGTGAAATCTGACATCTCAAGTGTCAGGCGGGCAAGAGATACTGCAAGACTAGAGACCGGAAGACGTAACGAGTACGACCAAGGTAGTGGTAAAATACGTTAATCTTGGTTGGACTAACAATGGCGAAGGCACGTTACGAGGACGGATCTGACAGTGAGGGATGAAGGCCAGGGGCG |
| p__Halobacteriota_g__Halohasta_ASV_862 | 0 | 1 | 1 | 1 | 0 | 0 | 0 | 0 | 0 | 0 | 1 | CCTTTACACTGCACGCAAGTGCGATAGGGGAATCCCAAGTGCGAGGGCATACTAGCCCTCGCTTTTGTGTACTGTAAGGAGGTACACGAATAAGAGCTGGGCAAGACCGGTGCCAGCCGCCGCGGTAATACCGGCAGCTCGAGTGATGTCCACTATTATTGGGCCTAAAGCGTCCGTAGCTGGCCGTACAAGTTCGTCGGGAAATCCACCTGCCCAACAGGTGGGCGTCCGGCGAAAACTGTGCGGCTTGGAACCGGAAGGCTCAGAGGGTACGTCGAGGGTAGGAGTGAAATCCTGTAATCCTCGACGGACCACCGATGGCGAAAGCACTCTGAGAAGACGGATTCGACAGTGAGGGACGAAAGCTAGGGTCT |
| p__Halobacteriota_g__Halorussus_ASV_863 | 0 | 1 | 1 | 0 | 0 | 0 | 0 | 0 | 0 | 0 | 0 | CCTTTACACTGCACGAGAGTGCGATAAGGGGACCCCGAGTGCGAGGGCATACAGTCCTCGCTTTTGTACACCGTAAGGTGGTGTACGAATAAGAGCTGGGCAAGACCGGTGCCAGCCGCCGCGGTAATACCGGCAGCTCGAGTGATGGCCGCTATTATTGGGCCTAAAGCGTCCGTAGCCGGCCGAACAGGTCCGTTGGGAAATCCACCAGCCTAACTGGTGGGCGTCCAGCGGAAACCAGTCGGCTTGGGGCCGGAAGACCTGAGGGGTACGTCCGGGGTAGGAGTGAAATCCTGTAATCCTGGACGGACCGCCGGTGGCGAAAGCGCCTCAGGAGGACGGACCCGACGGTGAGGGACGAAAGCTAGGGTCA |
| p__Nanoarchaeota_g__PWVO01_ASV_866 | 0 | 0 | 0 | 0 | 0 | 0 | 0 | 1 | 1 | 1 | 0 | CCTTTACAATGCACGAAAGTGTGATAAGGGAATCCCGAGTGCCTTGCTTTCGCAAGGCTTTTCTTAAGAGTAAATATCTTAGGGAATAAGTGGTGGGAAAGACCGGTGCCAGCAGCCGCGGTAATACCGGCGCCACAAGTGGTGATCATTTTTATTGGGCCTAAAGCGTTCGTAGCCGGTTTGATAAATCTTCTGTGAAATTGTCAGGCTTAACCTGACAGCGTGCAGGAGACACTGTCAAACTTGGGACCGGGAGGAGTCAGAGGTATTCCTTGGGGAGCGGTAAAATGCTATAATCCTCGGAAGACTAACTGTGGCGAAGGCGTCTGACCAGAACGGATCCGACGGTGATTGGACGAAAGTCAGGGGAG |

| Phylum_Genus_ASV_number | Presence (1) or Absence (0) in each layer | | | | | | | | | | | Sequence |
|---|---|---|---|---|---|---|---|---|---|---|---|---|
| | L1 | L2 | L3 | L4 | L5 | L6 | L7 | L8 | L9 | L10 | L11 | |
| p__Halobacteriota_g__Halovenus_ASV_882 | 0 | 0 | 1 | 0 | 0 | 0 | 0 | 0 | 0 | 1 | 0 | CCTTTACACTGCACGACAGTGCGATAGGGGGATTCCGAGTGCGAGGGCATACAGTCCTCGCTTTTCTGTACCGTAAGGAGGTACAGGAACAAGTGCTGGGCAAGACCGGTGCCAGCCGCCGCGGTAATACCGGCAGCACGAGTGATGGCCGATCTTATTGGGCCTAAAGCGTCCGTAGCTCGCCGGACAAGTCCGTTGGGAAATCCACGTGCCCAACACGTGGGCGTCCAGCGGAAACTGTCCGGCTTGGGGCCGGAGGACTCGAGGGGTACGTCCGGGGTAGGAGTGAAATCCCGTAATCCTGGACGGACCACCGGTGGCGAAAGCGCCTCGAGAGGACGGACCCGACAGTGAGGGACGAAAGCCAGGGTCT |
| p__Thermoplasmatota_g__SM1-50_ASV_893 | 0 | 0 | 0 | 0 | 0 | 0 | 0 | 1 | 1 | 1 | 1 | ACTTCACACTGCGCGAAAGCGCGATGAGGGAATCCCAAGTGTATGCACGTAAGTGTATACTGTTTCTATGTCTAAAAAGCATGGAGAGTAAGGGCTGGGTAAGACGGGTGCCAGCCGCCGCGGTAATACCTGCAGCTCAAGTGGTGGCCGCTATTATTGGGCCTAAAACGTCCGTAGCCGGTCTTGTAAATGCCTGGGTAAATCGTGCCGCTTAACGGTACGAATTCCGGGTAGACTGTAAGACTTGAGACCGGGAGAGGCTAGAGGTACTCCAGGGGTAGAGGTGAAATTCTGTAATCCTTGGGGGACCACCAGTGGCGAAGGCGTCTAGCTAGAACGGGTCTGACGGTGAGGGACGAAGCCCTGGGGCG |
| p__Halobacteriota_f__Haloferacaceae_ASV_894 | 0 | 0 | 1 | 0 | 0 | 0 | 0 | 0 | 0 | 0 | 0 | CCTTTACACTGCACGCAAGTGCGATAGGGGAATCCCAAGTGCGAGGGCATACTAGCCCTCGCTTTTGTCTACCCTAAGGCGGTAGACGAATAAGAGCTGGGCAAGACCGGTGCCAGCCGCCGCGGTAATACCGGCAGCTCGAGTGATGTCCACTATTATTGGGCCTAAAGCGTCCGTAGCTGGCCGCGCAAGTCCGTCGGGAAATCCACCCGCCCAACGGGTGGGCGTCCGGCGGAAACTGCGTGGCTTGGAACCGGAAGGCTCAGAGGGTACGTCCGGGGTAGGAGTGAAATCCCGTAATCCCGGACGGACCACCGATGGCGAAAGCACTCTGAGAGGACGGATTCGACAGTGAGGGACGAAAGCTAGGGTCT |
| p__Thermoproteota_g__CSP1-1_ASV_908 | 0 | 0 | 0 | 0 | 0 | 0 | 0 | 0 | 1 | 0 | 0 | CCTTTGCAATGTGCGCAAGCACGACAAGGTTAATCCGAGTGATTCCTGCTAAAGGAATCTTTTGTTAGTCGTAAAATCACTAATGAATAAGGGGTGGGCAAGTTCTGGTGTCAGCCGCCGCGGTAAAACCAGCACCTCAAGTGGTCAGGAGGATTATTGGGCCTAAAGCATCCGTAGCCTGTTCGGTAAGTTTCCGGTTAAATCCATTAGCTCAACTAATGGGCTGCCGGAAATACTGTCGGACTAGGAAGTGGGAGAGGTAGACGGTACTCGGTAGGAAGGGGTAAAATCCTTTGATCTATTGATGACCACCTGTGGCGAAGGCGGTCTACCAGAACACGTTCGACGGTGAGGGATGAAAGCTGGGGGAG |
| p__Halobacteriota_g__Natrinema_ASV_899 | 0 | 0 | 0 | 0 | 0 | 0 | 1 | 0 | 0 | 1 | 1 | CCTTTACACTGCACGCGAGTGCGATAAGGGGACTCCGAGTGCGAGGGCATATAGTCCTCGCTTTTCACCACCGTAGGAAGGTGGTAGAATAAGTGCTGGGCAAGACCGGTGCCAGCCGCCGCGGTAATACCGGCAGCACGAGTGATGACCGCTATTATTGGGCCTAAAGCGTCCGTAGCTGGCCACGCAAGTCTATCGGGAAATCTGCGCGCTCAACGCGCAGGCGTCCGGTGGAAACTGCGTGGCTTGGGACCGGAAGATCCAGAGGGTACGTCCGGGGTAGGAGTGAAATCCCGTAATCCTGGACGGACCACCGGTGGCGAAAGCGCTCTGGAAAGACGGATCCGACGGTGAGGGACGAAAGCTCGGGTCA |
| p__Halobacteriota_g___ASV_912 | 0 | 1 | 1 | 0 | 0 | 0 | 0 | 0 | 0 | 1 | 0 | CCTTTACACTGCACGAGAGTGCGATAGGGGGATCCCAAGTGCGAGGGCATATTAGCCCTCGCTTTTGTGAACCGTAAGGCGGTTCACGAATAAGAGCTGGGCAAGACCGGTGCCAGCCGCCGCGGTAATACCGGCAGCTCGAGTGATGACCGATTTTATTGGGCCTAAAGCGTCCGTAGCTGGCCACACAAGTCCGTCGGGAAATCTACCTGCCCAACAGGTAGGCGTCCGGCGGAAACTGTGCGGCTTGGGACCGGAAGGCGTGAAGGGTACGTCCGGGGTAGGAGTGAAATCCTGTAATCCTGGACGGACCACCGATGGCGAAAGCACTTCACGAGGACGGATCCGACAGTGAGGGACGAAAGCTAGGGTCT |
| p__Halobacteriota_g__Natronoarchaeum_ASV_916 | 0 | 1 | 1 | 0 | 0 | 0 | 0 | 0 | 0 | 0 | 0 | CCTTTACACTGCACGCCAGTGCGATAAGGGGACCCCCAGTGCGAGGGCATACCGTCCTCGCTTTTCACCACCGTAAGGAGGTAGTGGAACAAGAGCTGGGCAAGACCGGTGCCAGCCGCCGCGGTAATACCGGCAGCTCGAGTGATGGCCGTTTTTATTGGGCCTAAAGCGTCCGTAGCCGGCCAGAAAAGTTCGTCGGGAAATCCACGTGCCCAACACGTGGGCGTCCGGCGAAACCTCTCTGGCTTGGGACCGGACGGCCCGAGGGGTACGGTCGGGGTAGGAGTGAAATCCTGTAATCCCGACCGGACCACCGGTGGCGAAAGCGCCTCGGGAGGACGGATCCGACGGTGAGGGACGAAAGCCAGGGTCA |
| p__Halobacteriota_g__Saliphagus_ASV_915 | 0 | 1 | 0 | 0 | 1 | 0 | 0 | 0 | 0 | 0 | 0 | CCTTTACACTGCACGCCAGTGCGATAAGGGGACTCCACGTGCGAGGGCATACAGTCCTCGCTTTTGTGTACCGTAGGAAGGTACCCGAATAAGTGCTGGGCAAGACCGGTGCCAGCCGCCGCGGTAATACCGGCAGCACGAGTGATGACCGCTATTATTGGGCCTAAAGCGTCCGTAGCCGGCCGGGCAAGTCCGTCGGGAAATCCGCACGCCTAACGTGCGGGCGTCCGGCGGAAACTGCCTGGCTTGGGACCGGAAGACTCGAGGGGTACGTCCGGGGTAGGAGTGAAATCCTGTAATCCTGGACGGACCACCGGTGGCGAAAGCGCCTCGAGAAGACGGATCCGACGGTGAGGGACGAAAGCTCGGGTCA |
| p__Thermoplasmatota_g__SM1-50_ASV_923 | 0 | 0 | 0 | 0 | 0 | 0 | 0 | 0 | 1 | 0 | 0 | ACTTCGCACTGTGCGAAAGCACGACGAGGGAATCCCAAGTGCTTGCACGTAAGTGTAAGCTGTTTTTATGTCTAAAACGCATAGAGAGTAAGGGCTGGGTAAGACGGGTGCCAGCCGCCGCGGTAATACCTGCAGCTCAAGTGGTGGCCGTTATTATTGGGCCTAAAACGTTCGTAGCCGGTTTATTAAATGCCTGGGTAAATCGGACCGCTTAACGGTTCGAATTCCGGGTAGACTGATAAACTTGAGACCGGGAGAGGTTAGAGGTACTCCTGGGGTAGAGGTGAAATTCTGTAATCCTAGGGGGACCACCAGTGGCGAAGGCGTCTAACTAGAACGGGTCTGACGGTGAGGGACGAAGCCCTGGGGCG |
| p__Halobacteriota_f__Halobacteriaceae_4595_ASV_917 | 0 | 0 | 1 | 0 | 0 | 0 | 0 | 0 | 0 | 0 | 0 | CCTTTACACTGCACGAAAGTGCGATAAGGGGACCCCGAGTGCGAGGGCATACAGTCCTCGCTTTTGTACACCGTAAGGTGGTGTACGAATAAGGGCTGGGCAAGACCGGTGCCAGCCGCCGCGGTAATACCGGCAGCCCAAGTGATGGCCGTTATTATTGGGCCTAAAGCGTCCGTAGCCGGCCGGGCAAGTCCGTTGGGAAATCCGCTCGCTTAACGAGCGGACGTCCAGCGGAAACTGTTCGGCTTGGGACCGGAAGACTCGAGGGGTACGTCCGGGGTAGGAGTGAAATCCTGTAATCCTGGACGGACCGCCGGTGGCGAAAGCGCCTCGAGAAGACGGATCCGACGGTGAGGGACGAAAGCTAGGGTCT |
| p__Halobacteriota_f__Methanosarcinaceae_ASV_913 | 0 | 0 | 1 | 0 | 0 | 0 | 0 | 0 | 0 | 0 | 1 | ACTTTACAATGCGGGAAACCGCGATAAGGGGACACCGAGTGCCAGCATATTATGCTGGCTGTCCACATGTGTAAACAGCATGTGTTAGCAAGGGCCGGGCAAGACCGGTGCCAGCCGCCGCGGTAACACCGGCGGCCCGAGTGGTAATCACTATTATTGGGTCTAAAGGGTCCGTAGCCGGTTTGATCAGTCTTTCGGGAAATCTGATGGCTTAACTATCAGGCTTCCGGGAGATACTGTCAGACTTGGGACCGGGAGAGGTAAGAGGTACTACAGGGGTAGGAGTGAAATCTTGTAATCCCTGTGGGACCACCGGTGGCGAAGGCGTCTTACCAGAACGGGTCCGACGGTGAGGGACGAAAGCTGGGGGCA |
| p__Halobacteriota_o__Halanaerobiales_ASV_918 | 0 | 0 | 1 | 0 | 1 | 1 | 1 | 0 | 1 | 0 | 0 | CCTTTACACTGCACGACAGTGCGATAGGGGGATCCCGAGTGCGAGGGCATACAGTCCTCGCTTTTGTACACCGTAAGGAGGTGTACGAATAAGGGCTGGGCAAGACCGGTGCCAGCCGCCGCGGTAATACCGGCAGCCCGAGTGATGGCCGATGTTATTGGGCCTAAAGCGTCCGTAGCCGGCCGAACAAGTTCGTCGGGAAATCCGCACGCTTAACGTGCGGGCGACCGGCGAAAACTGTTCGGCTTGGGGCCGGAAGATCCGAGGGGTACGTCCGGGGTAGGAGTGAAATCCCGTAATCCTGGACGGACCGCCGGTGGCGAAAGCGCCTCGGAAGGACGGACCCGACGGTGAGGGACGAAAGCTGGGATCT |
| p__Halobacteriota_g__Halorussus_ASV_934 | 0 | 1 | 1 | 0 | 0 | 0 | 0 | 0 | 0 | 0 | 0 | CCTTTACACTGCACGACAGTGCGATAAGGGGACTCCGAGTGCGAGGGCATATAGTCCTCGCTTTTCTGTACCGTAAGGTGGTACAGGAATAAGGACTGGGCAAGACCGGTGCCAGCCGCCGCGGTAATACCGGCAGTCCAAGTGATGGCCGCTATTATTGGGCCTAAAGCGTCCGTAGCCGGCCAGACAGGTCCGTCGGGAAATCCACGCGCTCAACGCGTGGGCGTCCGGCGGAAACCAGCTGGCTTGGGGCCGGAAGACTCGAGGGGTACGTCCGGGGTAGGAGTGAAATCCCGTAATCCTGGACGGACCACCGGTGGCGAAAGCGCCTCGAGAAGACGGACCCGACGGTGAGGGACGAAAGCTAGGGTCA |
| p__Halobacteriota_g__Natronoarchaeum_ASV_925 | 1 | 1 | 1 | 0 | 0 | 0 | 0 | 0 | 0 | 0 | 0 | CCTTTACACTGCACGCCAGTGCGATAAGGGGATCCCCAGTGCGAGGGCATACAGTCCTCGCTTTTCACCACCGTACGATGGTGGTGGAACAAGAGCTGGGCAAGACCGGTGCCAGCCGCCGCGGTAATACCGGCAGCTCGAGTGATGGCCGTTCTTATTGGGCCTAAAGCGTCCGTAGCCGGCCACGCAAGTTCGTCGGGAAATCCACGCGCTCAACGCGTGGGCGTCCGGCGAAACCTGCGTGGCTTGGGACCGGACGGCCCGAGGGGTACGGCCGGGGTAGGAGTGAAATCCTGTAATCCCGGCCGGACCACCGGTGGCGAAAGCGCCTCGGGAGGACGGATCCGACGGTGAGGGACGAAAGCCAGGGTCA |
| p__Thermoplasmatota_g__SM1-50_ASV_930 | 0 | 0 | 0 | 0 | 0 | 0 | 0 | 0 | 1 | 1 | 1 | CCTTCACAATGTGCGAAAGCGCGATGAGGGGATCCCAAGTGCTTGCACTTAAGTGTAAGCTGTTTCTTTGTTTAAAACGCAGAGAGAGGAAGGGCTGGGTAAGACGGGTGCCAGCCGCCGCGGTAATACCTGCAGCTCAAGTGGTGACCGCTATTATTGGGCCTAAAACGTTCGTAGCCGGCTGAGTAAATGCTTGGGTAAATCGTGCCGCTTAACGGTACGAATTCCGGGTAGACTGCTCAGCTTGAGACCGGGAGAAGCTAGAGGTACTCCTGGGGTAGAGGTGAAATTCTGTAATCCTGGGGGACCACCAGTGGCGAAGGCGTCTAGCTAGAACGGGTCTGACGGTGAGGGACGAAACCCTGGGGCG |
| p__Halobacteriota_f__Halobacteriaceae_4595_ASV_936 | 0 | 0 | 1 | 0 | 0 | 0 | 0 | 0 | 0 | 0 | 0 | CCTTTACACTGTACGCAAGTGCGATAAGGGGACCCCGAGTGCGAGGGCATACAGTCCTCGCTTTTGTGTACCGTAAGGAGGTACACGAATAAGGGCTGGGCAAGACCGGTGCCAGCCGCCGCGGTAATACCGGCAGCCCTAGTGATGGCCGCTATTATTGGGCCTAAAGCGTCCGTAGCCGGCCGGACAAGTCCGTTGGGAAATCCGTTCGCTTAACGAACGGGCGTCCAGCGGAAACTGTCCGGCTTGGGACCGGAAGACTCGAGGGGTACGTCCGGGGTAGGAGTGAAATCCTGTAATCCTGGACGGACCGCCGGTGGCGAAAGCGCCTCGAGAAGACGGATCCGACGGTGAGGGACGAAAGCCAGGGTCT |
| p__Nanoarchaeota_o__Woesearchaeales_581_ASV_924 | 1 | 1 | 0 | 0 | 0 | 0 | 0 | 0 | 0 | 0 | 1 | CCTTTACAATGCACGAAAGTGTGATAAGGGAACCCTAAGTGCTATAGTTCTGCTATGGCTTTTGCCGAAAGTAAATATTTCGGCGAATAAGCGGTGGGTAAGACTGGTGCCAGCCGCCGCGGTAACACCAGCGCCGCAAGTAGCAACCATGTTTACTGGGCCTAAAGCGTTCGTAGCCGGTTTGGTAAATCTTTTGTGAAATTGTTGGGCCTAACCTAACAGCATGCAGAGGAGACTGCCAGACTAGAGACCGGGAGGAGTTGGAAATATTCTGGGGGGAGCGGTAAAATGTGATAATCCCCAGAAGATTACCTGTGGCGAAAGCGTCCAACCAGAACGGTTCTGACGGTGAGGAACGAAAGCTAGGGGAG |
| p__Nanoarchaeota_g__BMS3ABIN17_ASV_945 | 0 | 0 | 0 | 0 | 0 | 0 | 0 | 0 | 1 | 1 | 1 | AATCTACAATGCGCGCAAGCGTGATAGTTTGAGCCAGAGTGTTTTTCAAATGAAAGACTTTTGTGGAATGTAAAAAGTTTCACGAATAAGGACTGGGTAAGACTAGTGCCAGCCGCCGCGGTAATACTAGCGGTCCAAGTCGCAGCCATTTTTATTGGGTCTAAAACATCCGTAGCTTGCTTAATAAGTCTTCTGTGAAATTTTATGTCTAAAGCATAGAGCGAGCAGAAGGTACTGTTGAGCTAGAGACTGGAAGACGTAAGAAGTACATTCAGAGTAGCGGTTAAATGTGTTAATCCTGGATGGACTAACAAGAGCGAAGGCATCTTACGAGGACAGTTCTGACAGTAAGGGATGAAAGCTAGGGGCG |
| p__Nanoarchaeota_g__BMS3ABIN17_ASV_951 | 0 | 0 | 0 | 0 | 0 | 0 | 0 | 1 | 1 | 0 | 0 | AATCTGCAATGCGCGAAAGCGTGACAGTTTGAGCCAGAGTGCTTCTCATTGAGAAGCTTTTGCCAAGTGTAAAAAGCCTGGCGAATAAGGACTGGGCAAGACTAGTGCCAGCCGCCGCGGTAATACTAGCGGTCCAAGTCGCAGCCACATTTATTGGGTCTAAAACATCTGTAGCTTGTTAGGTAAGTTTCTCGTGAAATCCCGGGTCTTCAGCTTGGGGCGTGCGAGAAGTACTGCTTAACTAGAGACTGGAAGGCGTAAGAAGTACGTTCAAAGTAGCGGTAAAATGTGTTAATCTTGGACGGACTTACAACAGCGAAGGCATCTTACGAGGACAGTTCTGACAGTAAAGGATGAAGGCTAGGGGCG |
| p__Halobacteriota_g__Halalkaliarchaeum_ASV_947 | 0 | 0 | 1 | 0 | 0 | 0 | 0 | 0 | 0 | 0 | 1 | CCTTTACACTGCACGCGAGTGCGATAAGGGGACCCCGAGTGCGAGGGCATATCGTCCTCGCTTTTGTGTACCGTAGGGAGGTACACGAACAAGGGCTGGGCAAGACCGGTGCCAGCCGCCGCGGTAATACCGGCAGCCCAAGTGATGGCCGCTGTTATTGGGCCTAAAGCGTCCGTAGCCGGCCTGGCAAGTCCGTCGGGAAATCCACGCGCCCAACGCGTGGGCGGCCGGCGGAACCTGCCGGGCTCGGGACCGGAGGACCCAGCGGGTACGTCCGGGGTAGGAGTGAAATCCTGTAATCCCGGACGGACCGCCGATGGCGAAAGCACGCTGGGAAGACGGATCCGACGGTGAGGGACGAAAGCTAGGGTCT |
| p__Halobacteriota_g__Halorussus_ASV_949 | 0 | 0 | 1 | 1 | 0 | 0 | 0 | 0 | 0 | 0 | 0 | CCTTTACACTGCACGAAAGTGCGATAAGGGGATCCCAAGTGCGAGGGCATACAGTCCTCGCTTTTCTGTACCGTAGGGAGGTACAAGAATAAGGGCTGGGCAAGACCGGTGCCAGCCGCCGCGGTAATACCGGCAGCCCAAGTGATGGCCGCTATTATTGGGCCTAAAGCGTCCGTAGCCGGCCGGATAAGTCCGTCGGGAAATCCACCCGCTTAACGGGTGGACGTCCGGCGGAAACTGTCCGGCTTGGGGTCGGAGGGCCTGAGGGGTACGTCCGGGGTAGGAGTGAAATCCTGTAATCCTGGTCGGACCACCGGTGGCGAAAGCGCCTCAGGAGGACGAACCCGACGGTGAGGGACGAAAGCTAGGGTCA |
| p__Thermoplasmatota_g__Natronoplasma_ASV_940 | 1 | 1 | 1 | 0 | 0 | 0 | 0 | 0 | 0 | 0 | 0 | CCTTCACACTGTGGGAAACCACGACGAGGGGATTCCAAGTGCCAACACCCAGTGTTGGCTTTTCATCTGTCTAAAACGCAGGCGGAATAAGGGCCGGGTAAGACTGGTGCCAGCCGCCGCGGTAACACCAGCGGCTCGAGTGGTGGCCATTATTATTGAGCTTAAAGCGTCCGTAGCCGGTCTGGTAAATCTCTGGGTAAATTGAACGGCTTAACCGTTCGAATTCCAGAGAGACTGCCAGACTTGGAGCCGGGAGAGGCTAGAGGTACTTTTGGGGTAGAAGTAAAATTCTGTAATCCTAAAAGGACCACCAGTGGCGAAGGCGTCTAGCTAGAACGGACTCGACGGTGAGGGACGAAGCCCTGGGTCG |
| p__Halobacteriota_g__Saliphagus_ASV_948 | 0 | 1 | 1 | 0 | 0 | 0 | 0 | 0 | 0 | 0 | 0 | CCTTTACACTGCACGACAGTGCGATAAGGGGACCCCGAGTGCGAGGGCATACAGTCCTCGCTTTTGTGTACCGTAGGGAGGTACACGAATAAGGGCTGGGCAAGACCGGTGCCAGCCGCCGCGGTAATACCGGCAGCTCAAGTGATGGCCGCTGTTATTGGGCCTAAAGCGTCCGTAGCCGGCCGCACAAGTCCGTCGGGAAATCCACCCGCCTAACGGGTGGGCGTCCGGCGGAAACTGTGCGGCTTGGGACCGGAAGACTCGAGGGGTACGTCCGGGGTAGGAGTGAAATCCCGTAATCCTGGACGGACCGCCGGTGGCGAAAGCGCCTCGAGAGGACGGATCCGACGGTGAGGGACGAAAGCTAGGGTCT |
| p__Halobacteriota_o__Halanaerobiales_ASV_950 | 0 | 0 | 1 | 0 | 0 | 1 | 0 | 0 | 1 | 0 | 0 | CCTTTACACTGTACGGAAGTGCGATAAGGGGACCCCGAGTGCGAGGGCATACAGTCCTCGCTTTTGTACACCGTAGGGAGGTGTACGAATAAGGGCTGGGCAAGACCGGTGCCAGCCGCCGCGGTAATACCGGCAGCCCAAGTGATGGCCGCTATTATTGGGCCTAAAGCGTCCGTAGCCGGCCGGACAAGTCCGTTGGGAAATCCGTTCGCTCAACGAACGGGCGTCCAGCGGAAACTGTCCGGCTTGGGACCGGAAGACCCGAGGGGTACGTCCGGGGTAGGAGTGAAATCCCGTAATCCTGGACGGACCGCCGGTGGCGAAAGCGCCTCGGGAAGACGGATCCGACGGTGAGGGACGAAAGCCAGGGTCT |

| Phylum_Genus_ASV_number | Presence (1) or Absence (0) in each layer | | | | | | | | | | | |
|---|---|---|---|---|---|---|---|---|---|---|---|---|
| | L1 | L2 | L3 | L4 | L5 | L6 | L7 | L8 | L9 | L10 | L11 | Sequence |
| p__Halobacteriota_g__Halorussus_ASV_958 | 0 | 1 | 0 | 0 | 1 | 0 | 0 | 0 | 0 | 0 | 0 | ACTTTACACTGCACGACAGTGCGATAAGGGGACTCCGAGTGCGAGGGCATATAGTCCTCGCTTTTGTACACCGTAGGGTGGTGTACGAATAAGGGCTGGGCAAGACCGGTGCCAGCCGCCGCGGTAATACCGGCAGCCCGAGTGATGGCCGCTATTATTGGGCCTAAAGCGTCCGTAGCCGGCCAGACAGGTCCGTCGGGAAATCCATCCGCTCAACGGATGGGCGTCCGGCGGAAACCAGCTGGCTTGGGGCCGGAAGATCCGAGGGGTACGTCCGGGGTAGGAGTGAAATCCCGTAATCCTGGACGGACCACCGGTGGCGAAAGCGCCTCGGAAAGACGGACCCGACGGTGAGGGACGAAAGCTAGGGTCA |
| p__Halobacteriota_g__Natrarchaeobaculum_ASV_959 | 0 | 1 | 1 | 0 | 0 | 0 | 1 | 0 | 0 | 0 | 0 | CCTTTACACTGCGCGACAGCGCGATAAGGGGACTCCACGTGCCAGGGCATATCGTCCTGGCTTTTGTGTACCGTAAGGTGGTACACGAATAAGGGCTGGGCAAGACCGGTGCCAGCCGCCGCGGTAATACCGGCAGCCCGAGTGATGACCGCTCTTATTGGGCCTAAAGCGTCCGTAGCTGGCCGCGCAAGTCCATCGGGAAATCCGCACGCCTAACGTGCGGGCGTCCGGTGGAAACTGCGCGGCTTGGGACCGGAAGACCCGAGGGGTACGTCTGGGGTAGGAGTGAAATCCCGTAATCCTGGACGGACCACCGGTGGCGAAAGCGCCTCGGGAGGACGGATCCGACGGTGAGGGACGAAAGCTAGGGTCA |
| p__Thermoplasmatota_g__Natronoplasma_ASV_956 | 0 | 0 | 1 | 0 | 0 | 1 | 1 | 0 | 0 | 0 | 1 | CCTTCACACTGTGGGAAACCACGATGAGGAGATTCCAAGTGCCAACACCCAGTGTTGGCTTTTCAGATGTCTAAAACGCATCTGGAATAAGGGCCGGGTAAGACGGGTGCCAGCCGCCGCGGTAACACCCGCGGCTCAAGTGGTGACCATTATTATTGAGCTTAAAGCGTTCGTAGCCGGTCTGGTAAATCTCTGGGTAAATCGAACGGCTCAACCGTTCGAATACCAGGGAGACTGCCAGACTTGGGACCGGGAGAGGCCAGAGGTACTCCTAGGGTAGAAGTAAAATTCTGTAATCCTAGGAGGACCACCAGTGGCGCAAGCGTCTGGCTAGAACGGATCCGACGGTGAGGGACGAAGCCCTGGGGCG |
| p__Halobacteriota_f__Haloferacaceae_ASV_957 | 0 | 1 | 0 | 0 | 0 | 0 | 0 | 0 | 0 | 0 | 0 | CCTTTACACTGCGCGCACGCGCGATAAGGGGACCCCAAGTGCGAGGGCATACAGTCCTCGCTTTTCACATCCGTAGGGAGGATGTGGAATAAGGGCTGGGCAAGACCGGTGCCAGCCGCCGCGGTAATACCGGCAGCCCAAGTGATGGCCGCTCTTATTGGGCCTAAAGCGTCCGTAGCTGGCCACGTGTGTCCGTCGGGAAATCCACCAGCCCAACTGGTGGGCGGCCGGCGGAAACTGCGTGGCTTGGGACCGGAAGACCCACCGGGTACGTCCGGGGTAGGAGTGAAATCCCGTAATCCCGGACGGACCACCGATGGCGAAAGCACGGTGGGAAGACGGCTCCGACAGTGAGGGACGAAAGCTGGGGTCT |
| p__Thermoplasmatota_f__Methanomethylophilaceae_ASV_967 | 0 | 0 | 0 | 0 | 0 | 0 | 0 | 0 | 1 | 1 | 1 | ACTTCACAATGGGGGAAACCCTGATGAGGGGATTCCAAGTGCCAGCACACAGTGTTGGCTGTTCTTCTGTCTAAATCGCAGGAGGAGTAAGGGCCGGGTAAGACGGGTGCCAGCCGCCGCGGTAATACCCGCGGCCCGAGTGGTGGTCGATTTTATTGAGCCTAAAACGTTCGTAGCCGGCCTGGTAAATCCTTGGGTAAATCGGCCAGCTTAACTGTCCGAATTCCGAGGAGACTGCCAGGCTTGGGACCGGGAGAGGTCAGAGGTACTTCTGGGGTAGGGGTAAAATCCTGTAATCCTGGAAGGACCACCGGTGGCGAAGGCGTCTGACTAGAACGGATCCGACGGTGAGGGACGAAGCCCTGGGGCG |
| p__Thermoproteota_g__BA2_ASV_976 | 0 | 0 | 0 | 0 | 0 | 0 | 0 | 0 | 1 | 0 | 0 | ACTCCGCAATGCACGAAAGTGTGACGGGGCTACCCCGAGTGCCATCCGCTGAGGATGGCTTTTCTTTGGTATAACTAGCCATGGGAATAAGGAGAGGGCAAGACTGGTGTCAGCCGCCGCGGTAATACCAGCTCTCCGAGTGGTAGGGATGATTATTGGGCTTAAAGCGTCCGTAGCCGGCTTAGCAAGTTTCCTGTTAAACCCAGCGACTCAATCGTTGGATCGCGAGAAATACTACTAGGCTAGGAGGCGGGAGAGGTCAACGGTACTTCTGGGGTAGGGGTGAAATCCTATAATCCCGGGAGGACCACCAGTGGCGAAGGCTGTTGACTAGAACGCGCTCGACGGTGAGGGACGAAAGCTGGGGGAG |
| p__Halobacteriota_g__Halorussus_ASV_971 | 0 | 1 | 1 | 0 | 0 | 0 | 0 | 0 | 0 | 0 | 0 | CCTTTACACTGCACGCAAGTGCGATAAGGGGATCCCGAGTGCGAGGGCATATAGTCCTCGCTTTTCTGGACTGTAGGGAGGTCCAGGAACAAGAGCTGGGCAAGACCGGTGCCAGCCGCCGCGGTAATACCGGCAGCTCAAGTGATGGCCGATATTATTGGGCCTAAAGCGTCCGTAGCCGGCCGGATAAGTCCGTCGGGAAATCCACCCGCTCAACGGGTGGGCGTCCGGCGGAAACTGTCTGGCTTGGGGCCGGAAGGCCTGAGGGGTACGTCCGGGGTAGGAGTGAAATCCTGTAATCCTGGTCGGACCACCGGTGGCGAAAGCGCCTCAGGAGGACGGACCCGACGGTGAGGGACGAAAGCTAGGGTCA |
| p__Halobacteriota_o__Halanaerobiales_ASV_980 | 0 | 1 | 1 | 0 | 0 | 0 | 0 | 0 | 0 | 0 | 0 | CCTTTACACTGCACGACAGTGCGATAAGGGGACTCCAAGTGCGAGGGCATATAGTCCTCGCTTTTGTGTACCGTAAGGTGGTACACGAATAAGGGCTGGGCAAGACCGGTGCCAGCCGCCGCGGTAATACCGGCAGCCCAAGTGATGACCGCTATTATTGGGCCTAAAGCGTCCGTAGCTGGCCGTACAAGTCCGTCGGGAAATCTACTCGCCTAACGAGTAGACGTCCGGCGGAAACTGTTCGGCTAGGGACCGGAAGACCTGAGGGGTACGTCTGGGGTAGGAGTGAAATCCCGTAATCCTGGACGGACCACCGGTGGCGAAAGCGCCTCAGGAAGACGGATCCGACAGTGAGGGACGAAAGCTCGGGTCA |
| p__Thermoproteota_g__GCA-2726865_ASV_988 | 0 | 0 | 0 | 0 | 0 | 0 | 0 | 0 | 1 | 1 | 1 | CCTCCTCAATGCACTAAAGTGTGAAGGGGTTACCTCGAGTGCTATCTGAAGAAGGTGGCTTTTCCATGGTCTAAAAAGCCTAGGGAATAAGAGGGGGGCAAGACTGGTGTCAGCCGCCGCGGTAACACCAGCCCCTCGAGTGGTGGGGATGATTATTTGGCCTAAAGCGTCCGTAGCCGGCCAGGCAAGTTTCCCGTTAAATCCAACGTCTCAAGCGTTGGACCGCGGGAAATACTGCCCGGCTAGAGAGTGGGAGAGGTAAGCGGTATTCTGGGGGTAGGGGCGAAATCCTTTGATCCCCGGAGGACCACCAGTGGCGAAGGCTGCTTACTAGAACACGTCTGACGGTCAGGGACGAAAGCTGGGGTAG |
| p__Thermoplasmatota_g__SM1-50_ASV_992 | 0 | 0 | 0 | 0 | 0 | 0 | 0 | 0 | 1 | 0 | 0 | ACTTCGCACTGTGCGAAAGCACGACGAGGGAATCCCAAGTGCTGGCTCGTTAGAGTCAGCTGTTTCTTTGTCTAAAAAACAGAGAGAGTAAGGGCTGGGTAAGACGGGTGCCAGCCGCCGCGGTAATACCTGCAGCCCGAGTGGTGACTGTTATTATTGGGCCTAAAACGTCCGTAGCCGGTTCTGTAAATGCCTGGGTAAATCGTACGGCTCAACCGTACGAATTCCGGGTAGACTGCAGGGCTAGAGATCGGGAGAAGCTAGAGGTACTCTTGGGGTAGAGGTGAAATTCTGTAATCCTAGGGGGACCACCAGTGGCGAAGGCGTCTAGCTAGAACGAGTCTGACGGTCAGGGACGAAGCCCTGGGGCG |
| p__Thermoproteota_g__BA2_ASV_996 | 0 | 0 | 0 | 0 | 0 | 0 | 0 | 0 | 1 | 1 | 0 | ACTCCGCAATGCACGAAAGTGTGACGGGGCTACCCCGAGTGCCATCCGCTGAGGATGGCTTTTCTTTGGTTTAATTCGCCAGGGGAATAAGGAGAGGGCAAGACTGGTGTCAGCCGCCGCGGTAATACCAGCTCTCCGAGTGGTAGGGATGATTATTGGGCTTAAAGCGTCCGTAGCCGGCTTAGCAAGTTTCCTGTTAAATTCAGCGACTTAATCGTTGAGCCGCGGGAAATACTACTAGGCTAGGAGGCGGGAGAGGTCGACGGTACTTCTGGGGTAGGGGTGAAATCCTATAATCCCGGGAGGACCACCAGTGGCGAAGGCTGTCGACTAGAACGCGCTCGACGGTGAGGGACGAAAGCTGGGGGAG |
| p__Halobacteriota_g__Natrinema_ASV_1002 | 0 | 0 | 1 | 0 | 0 | 0 | 0 | 0 | 0 | 0 | 0 | CCTTTACACTGCACGCGAGTGCGATAAGGGGACTCCAAGTGCGAGGGCATATAGTCCTCGCTTTTCACCACCGTAAGGTGGTGGTGGAATAAGTGCTGGGCAAGACCGGTGCCAGCCGCCGCGGTAATACCGGCAGCACGAGTGATGACCGCTATTATTGGGCCTAAAGCGTCCGTAGCTGGCCGCGCAAGTTCATCGGGAAATCCGCACGCTTAACGTGCGGGCGTCCGGTGGAAACTGTGTGGCTTGGGACCGGAAGACCAGAGGGGTACGTCCGGGGTAGGAGTGAAATCCCGTAATCCTGGACGGACCACCGGTGGCGAAAGCGCCTCTGGAAGACGGATCCGACGGTGAGGGACGAAAGCTCGGGTCA |
| p__Thermoproteota_g__BA2_ASV_1015 | 0 | 0 | 0 | 0 | 0 | 0 | 0 | 0 | 1 | 1 | 0 | ACTCCGCAATGCACGAAAGTGTGACGGGGCTACCCCGAGTGCCGTCCGCTGAGGATGGCTTTTCCTTGGTGTAACTAGCCGGGGGAATAAGGAGAGGGCAAGACTGGTGTCAGCCGCCGCGGTAATACCAGCTCTCCGAGTGGTAGGGATGTTTATTGGGCTTAAAGCGTCCGTAGCCGGCTTAGCAAGTCTCCTGTTAAATTCAGCGACCTAATCGTTGAGCCGCGGGAGATACTACTAGGCTAGGAGGCGGGAGAGGTCAACGGTACTTCCGGGGTAGGGGTGAAATCCTATAATCCCGGGAGGACCACCAGTGGCGAAGGCTGTTGACTAGAACGCGCTCGACGGTGAGGGACGAAAGCTGGGGGAG |
| p__Nanoarchaeota_g__BMS3ABIN17_ASV_1016 | 0 | 0 | 0 | 0 | 0 | 0 | 0 | 0 | 1 | 0 | 1 | AATCTACAATGCGCGCAAGCGTGATAGTTTGAGCCAGAGTGTTTTTCAAATGAAAGACTTTTGTGAAATGTAAAAAGTTTCGCGAATAAGGACTGGGTAAGACTAGTGCCAGCCGCCGCGGTAATACTAGCGGTCCAAGTCGCAGCCATTTTTATTGGGTCTAAAACATCCGTAGCTTGCTTAATAAGTCTTCTGTGAAATTTTATGTCTAAAGCATAGAGCGAGCAGAAGGTACTGTTGAGCTAGAGACTGGAAGACGTAAGAAGTACATTCAGAGTAGCGGTTAAATGTGTTAATCCTGGATGGACTAACAAGAGCGAAGGCATCTTACGAGGACAGTTCTGACAGTAAGGGATGAAAGCTAGGGGCG |
| p__Halobacteriota_g__Halorussus_ASV_1023 | 0 | 1 | 1 | 0 | 0 | 0 | 0 | 0 | 0 | 0 | 0 | CCTTTACACTGCACGACAGTGCGATAAGGGGACTCCGAGTGCGAGGGCATATAGTCCTCGCTTTTCTGTACCGTAAGGTGGTACAAGAATAAGGACTGGGCAAGACCGGTGCCAGCCGCCGCGGTAATACCGGCAGTCCGAGTGATGGCCGCTATTATTGGGCCTAAAGCGTCCGTAGCCGGCCAGACAGGTCCGTCGGGAAATCCACGCGCTCAACGCGTGGGCGTCCGGCGGAAACCAGCTGGCTTGGGGCCGGAAGACTCGAGGGGTACGTCCGGGGTAGGAGTGAAATCCCGTAATCCTGGACGGACCACCGGTGGCGAAAGCGCCTCGAGAAGACGGACCCGACGGTGAGGGACGAAAGCTAGGGTCA |
| p__Halobacteriota_g__Natronobiforma_ASV_1025 | 0 | 0 | 1 | 1 | 0 | 0 | 0 | 0 | 0 | 0 | 0 | CCTTTACACTGCACGCCAGTGCGATAAGGGGACACCAAGTGCGAGGGCATATCGTCCTCGCTTTTCACCACCGTAAGGTGGTGGTGGAATAAGAGCTGGGCAAGACCGGTGCCAGCCGCCGCGGTAATACCGGCAGCTCGAGTGATGACCGCTATTATTGGGCCTAAAGCGTCCGTAGCTGGCCACACAAGTCCATCGGGAAATCTCCCTGCCTAACAGGGAGGCGTCCGGTGGAAACTGTGTGGCTTGGGACCGGAAGATCCAGAGGGTACGTCTGGGGTAGGAGTGAAATCCTGTAATCCTGGACGGACCACCGGTGGCGAAAGCGCTCTGGAAGGACGGATCCGACAGTGAGGGACGAAAGCTAGGGTCA |
| p__Thermoproteota_g__PALSA-986_ASV_1010 | 1 | 1 | 1 | 1 | 0 | 0 | 0 | 0 | 0 | 0 | 0 | ACTTTGCAATACACGAAAGTGTGACAGGGTCATCCCGAGTGCCGACCGCTGAGGTTGGCTTTTACCCAGTCTAGACAGCTGGGGGAATAAGGAGAGGGCAAGTCTGGTGTCAGCCGCCGCGGTAATACCAGCTCTCCGAGTGGTGTGGACGTTTATTGGGCCTAAAGCATCCGTAGCTTGCTGGGCAAGTCCCCTGTTAAATCCACCGATTTAATCGTTGGTGTGCGGGGAAACTGTTCGGCTAGGGGACGAGAGAGGCAGACGGTATTCCCGGGGTAGGGGTGAAATCCTATAATCCCGAGAAGACCACCAGTGGCGAAGGCTGTCTGCTAGAACGCGCCCGACGGTGAGGGATGAAAGCTGGGGGAG |
| p__Halobacteriota_o__Halanaerobiales_ASV_1014 | 0 | 1 | 1 | 0 | 0 | 0 | 0 | 0 | 0 | 1 | 0 | CCTTTACACTGCACGTCAGTGCGATAGGGGGACCCCGAGTGCGAGGGCATACAGTCCTCGCTTTTATGCACCGTAAGGAGGTGCATGAATAAGTGCTGGGCAAGACCGGTGCCAGCCGCCGCGGTAATACCGGCAGCACAAGTGATGGCCGCTATTATTGGGCCTAAAGCGTCCGTAGCCGGCCGAACAAGTCCGTCGGGAAATCCATCCGCTCAACGGATGGGCGTCCGGCGGAAACTGTTCGGCTTGGGACAGGAAGACTCGAGGGGTACGTCCGGGGTAGGAGTGAAATCCCGTAATCCTGGACGGACCGCCGATGGCGAAAGCACCTCGAGAAGACTGATCCGACGGTGAGGGACGAAAGCTGGGGTCT |
| p__Nanoarchaeota_g__BMS3ABIN17_ASV_1029 | 1 | 0 | 0 | 0 | 0 | 0 | 0 | 1 | 1 | 0 | 0 | AATCTACAATGCGCGAAAGCGTGATAGTTTGAGTCAGAGTGCGTTTCTTTGAAATGCTTTTGTGAAATGTAAAAAGTTTCACGAATAAGGACTGGGTAAGACGAGTGCCAGCCGCCGCGGTAATACTCGCGGTCCAAGTCGCAGCCATTTTTATTGGGTCTAAAACATCCGTAGCTTGCTTTGTAAGTTCCTTGTGAAATCCCGTATCTCAAGTATGGGGCGGGCAAGGAGTACTGCTAAGCTAGAGACTGGGAGACGTAAAGAGTACGTTTAAGGTAGCGGTAAAATGTGTTAATCTTAAGCGGACTAACAATAGCGAAGGCACTTTACGAGAACAGTTCTGACAGTAAGGGATGAAGGCTAGGGGCG |
| p__Halobacteriota_g__Halorientalis_ASV_1040 | 0 | 0 | 1 | 0 | 1 | 0 | 0 | 0 | 0 | 0 | 0 | CCTTTACACTGCACGACAGTGCGATAGGGGGACTCCGAGTGCGAGGGCATACAGTCCTCGCTTTTCTGTACCGTAAGGTGGTACAGGAACAAGTGCTGGGCAAGACCGGTGCCAGCCGCCGCGGTAATACCGGCAGCACGAGTGATGGCCGCTATTATTGGGCCTAAAGCGTCCGTAGCCGGCCGAGCAAGTTCGTCGGGAAATCTGCCCGCTTAACGGGCAGGCGTCCGGCGAAAACTGTTCGGCTTGGGGCCGGGAGATCCAAGGGGTACGTCCGGGGTAGGAGTGAAATCCCGTAATCCTGGACGGACCACCGATGGCGAAAGCACCTTGGAAAAACGGACCCGACGGTGAGGGACGAAAGCTAGGGTCT |
| p__Halobacteriota_g__Halorussus_ASV_1037 | 0 | 1 | 0 | 0 | 0 | 1 | 0 | 0 | 0 | 0 | 0 | CCTTTACACTGCACGAAAGTGCGATAAGGGGATCCCGAGTGCGAGGGCATACAGTCCTCGCTTTTGTGTACCGTAAGGTGGTACACGAATAAGAGCTGGGCAAGACCGGTGCCAGCCGCCGCGGTAATACCGGCAGCTCGAGTGATGGCCGCTATTATTGGGCCTAAAGCGTCCGTAGCCGGCCGAACAGGTCCGTTGGGAAATCCATCTGCCTAACAGATGGGCGTCCAGCGGAAACCAGTCGGCTTGGGGCCGGAAGACCTGAGGGGTACGTCCGGGGTAGGAGTGAAATCCTGTAATCCTGGACGGACCGCCGGTGGCGAAAGCGCCTCAGGAGGACGGACCCGACGGTGAGGGACGAAAGCTAGGGTCA |
| p__Thermoplasmatota_g__Natronoplasma_ASV_1030 | 1 | 1 | 0 | 0 | 0 | 0 | 0 | 1 | 0 | 0 | 0 | CCTTCACACTGTGGGAAACCACGATGAGGGGATTCCAAGTGCCAACACTAAGTGTTGGCTTTTCATCTGTGTAAAAAGCAGGTGGAATAAGGGCCGGGTAAGACGGGTGCCAGCCGCCGCGGTAACACCCGCGGCCCAAGTGGTGGTCATTATTATTGAGCTTAAAGCGTCCGTAGCCGGTCTGGTAAATCTCTGGGTAAATCGGACGGCTCAACCGTTCGAATACCAGGGAGACTGCTAGACTTGGGGCCGGGAGAGGTCAGAGGTACTCCTGGGGTAGAAGTAAAATTCTGTAATCTTAGGAGGACCACCAGTGGCGAAAGCGTCTGGCTAGAACGGACCCGACGGTGAGGGACGAAGCCCTGGGGCG |
| p__Halobacteriota_g__Saliphagus_ASV_1038 | 0 | 1 | 0 | 0 | 0 | 0 | 1 | 0 | 0 | 0 | 0 | CCTTTACACTGCACGCCAGTGCGATAAGGGGATTCCAAGTGCGAGGGCATACAGTCCTCGCTTTTGTGTACCGTAGGAAGGTACACGAATAAGGGCTGGGCAAGACCGGTGCCAGCCGCCGCGGTAATACCGGCAGCCCAAGTGATGACCGATATTATTGGGCCTAAAGCGTCCGTAGCCGGCTACGCAAGTCCGTCGGGAAATCTGCTCGCTCAACGAGTAGGCGTCCGGCGGAAACTGCGTGGCTTGGGACCGGAAGACTCGAGGGGTACGTCCGGGGTAGGAGTGAAATCCCGTAATCCTGGACGGACCACCGGTGGCGAAAGCGCCTCGAGAGGACGGATCCGACGGTGAGGGACGAAAGCTAGGGTCA |
| p__Nanoarchaeota_g__BMS3ABIN17_ASV_1052 | 0 | 1 | 0 | 0 | 0 | 0 | 0 | 0 | 1 | 1 | 0 | TTTTTGCAATGCACGCAAGTGTGACAAAGCGAACCAGAGTGTTTCTCATTCAGAGAAACTTTTGTCAGCTGTAAAAAGGCTGACGAATAAGGACTGGGCAAGACCGGTGCCAGCCGCCGCGGTAATACCGGCGGTCCAAGTCGCAGCCACATTTATTGGGTCTAAAACATCCGTAGCTTGTTTTGCAAGTCTTTTGTGAAATCTTGTCTCTCAAGGACAAGGCGTGCAAGAGATACTGCATAACTAGAGACTGGGGGATGTAAAGAGTATGATTAAGGTAGTGGTAAAATACGTTGATCTTAATCAGACTAACAATTAGCGAAGGCACTTTACAACAACAGTTCTGACAGTGAGGGATGAAGGCTAGGGGCG |

| Phylum_Genus_ASV_ number | Presence (1) or Absence (0) in each layer L1 | L2 | L3 | L4 | L5 | L6 | L7 | L8 | L9 | L10 | L11 | Sequence |
|---|---|---|---|---|---|---|---|---|---|---|---|---|
| p__Halobacteriota_g__ Halobaculum_ASV_1053 | 0 | 0 | 1 | 0 | 0 | 0 | 0 | 0 | 0 | 0 | 1 | CCTTTACACTGCACGCCAGTGCGATAGGGGGACCCCGAGTGCGAGGGCATACAGTCCTCGCTTTTGCACACCGTAAGGAGGTGTGCGAATAAGTGCTGGGCAAGACCGGTGCCAGCCGCCGCGGTAATACCGGCAGCACAAGTGATGGCCGCTTTTATTGGGCCTAAAGCGTCCGTAGCCGGCCGAACAAGTCCGTCGGGAAATCCATCCGCTCAACGGATGGGCGTCCGGCGGAAACTGTTCGGCTTGGGACAGGAAGACCCGAGGGGTACGTCCGGGGTAGGAGTGAAATCCCGTAATCCTGGACGGACCGCCGATGGCGAAAGCACCTCGGGAAGACTGATCCGACGGTGAGGGACGAAAGCTGGGGTCT |
| p__Halobacteriota_g__ Halosimplex_ASV_1055 | 0 | 1 | 1 | 0 | 0 | 0 | 0 | 0 | 0 | 0 | 0 | CCTTTACACTGCACGCCAGTGCGATAGGGGGACTCCGAGTGCGAGGGCATACAGTCCTCGCTTTTCTGCACCGTAAGGTGGTGCAGGAACAAGTGCTGGGCAAGACCGGTGCCAGCCGCCGCGGTAATACCGGCAGCACGAGTGATGGCCGATATTATTGGGCCTAAAGCGTCCGTAGCCGGCCCGACAAGTCCGTTGGGAAATCCACGCGCCCAACGCGTGGGCGTCCAGCGGAAACTGTCGGGCTTGGGGCCGGAGGACTCGAGGGGTACGTCCGGGGTAGGAGTGAAATCCCGTAATCCTGGACGGACCACCGGTGGCGAAAGCGCCTCGAGAGGACGGACCCGACGGTGAGGGACGAAAGCCAGGGTCT |
| p__Halobacteriota_o__ Halanaerobiales_ASV_1058 | 0 | 1 | 1 | 0 | 0 | 0 | 0 | 0 | 0 | 0 | 0 | CCTTTACACTGCACGACAGTGCGATAAGGGGACCCCAGGTGCGAGGGCATATCGTCCTCGCTTTTCTGCACCGTAAGGAGGTGCAGGAATAAGGGCTGGGCAAGACCGGTGCCAGCCGCCGCGGTAATACCGGCAGCCCAAGTGATGGCCGCTATTATTGGGCCTAAAGCGTCCGTAGCCGGCCGGACAAGTCCGTCGGGAAATCCACCCGCTCAACGGGTGGGCGTCCGGCGGAAACTGTTCGGCTTGGGGCCGGAGGACCTGAGGGGTACGTCCGGGGTAGGAGTGAAATCCTGTAATCCTGGACGGACCACCGGTGGCGAAAGCGCCTCAGGAGGACGGACCCGACGGTCAGGGACGAAAGCTAGGGTCT |
| p__Thermoproteota_g__ BA2_ASV_1067 | 0 | 0 | 0 | 0 | 0 | 0 | 0 | 0 | 1 | 1 | 0 | ACTCCGCAATGCACGAAAGTGTGACGGGGCTACCCCGAGTGCCGTCCGCTGAGGATGGCTTTTCCTTGGTTTAACTAGCCAGGGGAATAAGGAGAGGGCAAGACTGGTGTCAGCCGCCGCGGTAATACCAGCTCTCCGAGTGGTAGGGATGATTATTGGGCTTAAAGCGTCCGTAGCCGGCTTAGCAAGTCTCCTGTTAAATTCAGCGACCTAATCGTTGAGCCGCGGGAGATACTACTAGGCTAGGAGGCGGGAGAGGTCGACGGTACTTCTGGGGTAGGGGTGAAATCCTATAATCCCGGGAGGACCACCAGTGGCGAAGGCTGTTGACTAGAACGCGCTCGACGGTGAGGGACGAAAGCTGGGGGAG |
| p__Thermoplasmatota_ g__SM1-50_ASV_1068 | 0 | 0 | 0 | 0 | 0 | 0 | 0 | 0 | 1 | 1 | 1 | CCTTCACACTGTGCGAAAGCACGATGAGGGGATCCCAAGTGCTGGCTCGTAAGAGTCAGCTGTTTCTCTGTCTAAAAAACAGAGAGAGTAAGGGCTGGGTAAGACGGGTGCCAGCCGCCGCGGTAATACCTGCAGCTCAAGTGGTGGCCGCTATTATTGGGCCTAAAACGTTCGTAGCCGGTTCAGTAAATGCCTGGGTAAATTGTATGGCTTAACCATACGAATTCCGGGTAGACTGCTCGACTTGAGACCGGGAGAGGCTAGAGGTACTCCTGGGGTAGAGGTGAAATTCTGTAATCCTAGGGGGACCACCAGTGGCGAAGGCGTCTAGCCAGAACGGGTCTGACGGTGAGGGACGAAGCCCTGGGGCG |
| p__Halobacteriota_g__ Halolamina_ASV_1083 | 0 | 1 | 1 | 0 | 0 | 0 | 0 | 0 | 0 | 0 | 0 | CCTTTACACTGCACGCCAGTGCGATAAGGGAACCCCGAGTGCGAGGGCATATCGCCCTCGCTTTTCTGGACCGTAGGGAGGTCCAGGAACAAGAGCTGGGCAAGACCGGTGCCAGCCGCCGCGGTAATACCGGCAGCTCGAGTGATGACCGATTTTATTGGGCCTAAAGCGTCCGTAGCTGGCCGTGCAAGTCCGTCGGGAAATCCACCCGCTTAACGGGTGGGCGTCCGGCGGAAACTGTGGGGCTTGGGACAGGAAGGCTCGAGGGGTACGTTCGGGGTAGGAGTGAAATCCCGTAATCCTGAACGGACCGCCGATGGCGAAAGCACCTCGAGAGGACTGATCCGACAGTGAGGGACGAAAGCTGGGGTCT |
| p__Halobacteriota_f__N atrialbaceae_ASV_1082 | 0 | 1 | 1 | 0 | 0 | 0 | 0 | 0 | 0 | 0 | 0 | CCTTTACACTGCACGACAGTGCGATAAGGGGACTCCAAGTGCGAGGGCATATAGTCCTCGCTTTTGTGTACCGTAAGGTGGTACACGAATAAGGGCTGGGCAAGACCGGTGCCAGCCGCCGCGGTAATACCGGCAGCCCGAGTGATGACCGCTATTATTGGGCCTAAAGCGTCCGTAGCTGGCCGATCAAGTCCGTCGGGAAATCCGCGCGCTTAACGTGCGGGCGTCCGGCAGAAACTGGTCGGCTTGGGACCGGAAGACCCGAGGGGTACGTCTGGGGTAGGAGTGAAATCCCGTAATCCTGGACGGACCGCCGGTGGCGAAAGCGCCTCGGGAAGACGGATCCGACGGTGAGGGACGAAAGCTAGGGTCA |
| p__Halobacteriota_g__ Halobellus_3026_ASV_1094 | 0 | 1 | 0 | 0 | 0 | 1 | 1 | 0 | 0 | 0 | 0 | CCTTTACACTGCACGAGAGTGCGATAAGGGGACTCCGAGTGCGAGGGCATATCGTCCTCGCTTTTCTGTACCGTAGGGCGGTACAGGAACAAGAGCTGGGCAAGACCGGTGCCAGCCGCCGCGGTAATACCGGCAGCTCAAGTGATGGCCGCTCTTATTGGGCCTAAAGCGTCCGTAGCTGGCCTCGAAAGTCCGTCGGGAAATCCACTGGCCTAACCAGTGGGCGTCCGGCGGAAACTCCGAGGCTTGGGACCGGAAGGCTCGAGGGGTACGTCCGGGGTAGGAGTGAAATCCTGTAATCCCGGACGGACCACCGATGGCGAAAGCACCTCGAGAGGACGGATCCGACAGTGAGGGACGAAAGCTAGGGTCT |
| p__Halobacteriota_g__ Halorubrum_ASV_1091 | 1 | 1 | 1 | 0 | 0 | 0 | 0 | 0 | 1 | 0 | 0 | CCTTTACACTGCACGACAGTGCGATAGGGGGATCCCAAGTGCACAGGCATAGCGCCTGTGCTTTTCGGCACCCTAAGGCGGTGCCAGAATAAGGGCTGGGCAAGACCGGTGCCAGCCGCCGCGGTAATACCGGCAGCCCAAGTGATGGCCGATCTTATTGGGCCTAAAGCGTCCGTAGCTGGCCACGCAAGTCCATCGGGAAATCCACCTGCTCAACAGGTGGGCGCCCGGTAGAAACTGCGTGGCTTGGGACCGGAAGGCGCGACGGGTACGTCCGGGGTAGGAGTGAAATCCCGTAATCCTGGACGGACCGCCGATGGCGAAAGCACGTCGCGAGAACGGATCCGACAGTGAGGGACGAAAGCCAGGGTCT |
| p__Halobacteriota_g__ Halorussus_ASV_1098 | 0 | 0 | 1 | 0 | 0 | 0 | 0 | 0 | 0 | 0 | 0 | ACTTTACACTGCACGACAGTGCGATAAGGGGACTCCGAGTGCGAGGGCATATAGTCCTCGCTTTTGTACACCGTAAGGTGGTGTACGAATAAGGGCTGGGCAAGACCGGTGCCAGCCGCCGCGGTAATACCGGCAGCCCGAGTGATGGCCGCTATTATTGGGCCTAAAGCGTCCGTAGCCAGCCAGACAGGTCCGTCGGGAAATCCATCCGCTCAACGGATGGGCGTCCGGCGGAAACCAGCTGGCTTGGGGCCGGAAGACCTGAGGGGTACGTCCGGGGTAGGAGTGAAATCCCGTAATCCTGGACGGACCGCCGGTGGCGAAAGCGCCTCAGGAGGACGGACCCGACGGTGAGGGACGAAAGCTAGGGTCA |
| p__Nanohaloarchaeota_ g__Nanohalobium_ASV _1090 | 1 | 1 | 1 | 0 | 0 | 0 | 0 | 1 | 0 | 0 | 0 | CCTTTACAATGCACGCAAGTGCGATAAGGGGACTCCGAGTGCGGTCTTCCTACACGGAAGGCCGCTTTTCTGTACTGTAAGAAGGTACCGGAACAAGGGGTGGGCAACACTGCTGCCAGCCGCCGCGGTAATAGCAGGACCCCAAGTGGTAGCCAGTTTTATTGGGCTTAAAGTGTTCGTAGCCGGCCATGTAAGTCCGCTGTGAAATCCAAGCGCTTAACGTTTGGAAGTGCAGCGGATACTGCATGGCTAGGGACCGGATGGCGTCGAGGCTATTCCAGGGGTAGGAGTGAAATCCAAAAATCCCTGGAGGAGCACCTGTGGCGAAAGCGCTCGACGAATACGGATCCGACGGTGAGGAACGAAAGCCAGGGGAG |
| p__Halobacteriota_g__S aliphagus_ASV_1089 | 1 | 1 | 0 | 0 | 1 | 1 | 0 | 0 | 0 | 1 | 0 | CCTTTACACTGCACGAAAGTGCGATAGGGGGACCCCGAGTGCGAGGGCATACAGTCCTCGCTTTTGTGTACCGTAAGGAGGTACACGAATAAGGGCTGGGCAAGACCGGTGCCAGCCGCCGCGGTAATACCGGCAGCTCGAGTGATGGCCGCTGTTATTGGGCCTAAAGCGTCCGTAGCCGGCCGCACAAGTCCGTCGGGAAATCCACCCGCCTAACGGGTGGGCGTCCGGCGGAAACTGTGCGGCTTGGGACCGGAAGACTCGAGGGGTACGTCCGGGGTAGGAGTGAAATCCTGTAATCCTGGACGGACCGCCGGTGGCGAAAGCGCCTCGAGAGGACGGATCCGACGGTGAGGGACGAAAGCTAGGGTCT |
| p__Thermoplasmatota_ g__SM1-50_ASV_1101 | 0 | 0 | 0 | 0 | 0 | 0 | 0 | 0 | 1 | 0 | 0 | ACTTCACAATGTGCGAAAGCGCGATGAGGGAATCCCAAGTGCTTGCACTTAGGTGTAAGCTGTTTCTCTGTCTAAAACGCAGAGAGAGTAAGGGCTGGGTAAGACGGGTGCCAGCCGCCGCGGTAATACCTGCAGCTCAAGTGGTGATCGTTATTATTGGGCCTAAAACGTTCGTAGCCGGCCTATTAAATGCCTGGGTAAATCGGACCGCTTAACGGTTCGAATTCCGGGTAGACTGATAGGCTTGAGACCGGGCGAGGCTAGAGGTACTCCTGGGGTAGAGGTGAAATTCTGTAATCCTAGGGGGACCACCAGTGGCGAAGGCGTCTAGCTAGAACGGGTCTGACGGTGAGGGACGAAGCCCTGGGGCG |
| p__Halobacteriota_g__ Natrarchaeobaculum_A SV_1115 | 0 | 1 | 1 | 0 | 0 | 0 | 0 | 0 | 0 | 0 | 0 | CCTTTACACTGCGCGACAGCGCGATAAGGGGACTCCACGTGCCAGGGCATATCGTCCTGGCTTTTGTGTACCGTAAGGTGGTACACGAATAAGGGCTGGGCAAGACCGGTGCCAGCCGCCGCGGTAATACCGGCAGCCCGAGTGATGACCGCTCTTATTGGGCCTAAAGCGTCCGTAGCTGGCCATCCAAGTCCATCGGGAAATCCGCACGCCCAACGTGCGGGCGTCCGGTGGAAACTGGATGGCTTGGGACCGGAAGACCCGAGGGGTACGTCTGGGGTAGGAGTGAAATCCCGTAATCCTGGACGGACCACCGGTGGCGAAAGCGCCTCGGGAGGACGGATCCGACGGTGAGGGACGAAAGCTAGGGTCA |
| p__Thermoplasmatota_c __Thermoplasmata_177 3_ASV_1104 | 1 | 1 | 0 | 0 | 1 | 0 | 0 | 1 | 0 | 1 | 1 | ACTCTACAATGTGGGAAACCACGATAGGGGAACTCCAAGTGCATGCACTTAGTGTATGCTGTTCTACTGCCTAAAAAGCAGTTGAAGTAAGGGCTGGGTAAGACGGGTGCCAGCCGCCGCGGTAATACCCGCAGCCCCAGTGGTGTTCAGCATTATTGAGCCTAAAGCGTCCGTAGCCGGTCCTGTAAATCTCTGGGTAAATCGAGCAGCTCAACTGTTCGAATACCGGAGAGACTGCAGGACTTGGGACTGGGAGAGGTGGAAGGTACTCCAGGGGTAGGGGTAAAATCCTGTAATCCTTGGGGGACCACCAGTGGCGAAGGCGTTTCACTAGAACAGATCCGACGGTGAGGGACGAAGCCCTGGGGCG |
| p__Nanoarchaeota_g__ BMS3ABIN17_ASV_1127 | 0 | 0 | 0 | 0 | 0 | 0 | 0 | 0 | 1 | 1 | 1 | ATTCTGCAATGCGCGCAAGCGTGACAGATTGAACCAGAGTGCTTTTCAATTTGAAAAGCTTTTGTGAAATGCAAAAAGTTTTACGAATAAGGACTGGGTAAGACTAGTGCCAGCCGCCGCGGTAATACTAGCGGTCCAAGTCGCAGCCACAATTATTGGGTCTAAAACATCCGTAGCTTGCTTGATAAGTTTCCTGTGAAATTCGGTTTCTTAAAAGCCGAGCTGGCGGGAAGTACTGTCAAGCTAGAGACTGGAAGACGTAAGAAGTATGTTCAAAGTAGTGGTAAAATACGTTAATCTTGAACAGACTACCAATAGCGAAGGCATCTTACGAGGACAGTTCTGACAGTAAGGGATGAAGGCCAGGGGCG |
| p__Halobacteriota_g__ Haladaptatus_ASV_1135 | 0 | 1 | 0 | 0 | 0 | 0 | 0 | 0 | 0 | 0 | 0 | ACTTTACACTGCACGTCAGTGCGATAAGGGAACTCCAAGTGCGAGGGCATATAGTCCTCGCTTTTGTGTATCGTAAGGTGGTACACGAATAAGAGCTGGGCAAGACCGGTGCCAGCCGCCGCGGTAATACCGGCAGCTCGAGTGATGGCCAATCTTATTGGGCCTAAAGCGTCCGTAGCCGGCCGAACAGGTCCGTCGGGAAATCTGCTCGCTCAACGAGCAGGCGTCCGACGGAAACCAGTCGGCTTGGGACCGGAAGACCCGAGGGGTACGTCTGGGGTAGGAGTGAAATCCTGTAATCCTAGACGGACCGCCGGTGGCGAAAGCGCCTCGGGAAGACGGATCCGACGGTGAGGGACGAAAGCTAGGGTCA |
| p__Halobacteriota_g__ Halorussus_ASV_1125 | 1 | 1 | 0 | 0 | 0 | 0 | 0 | 0 | 1 | 0 | 1 | CCTTTACACTGCACGCCAGTGCGATAAGGGGACCCCGAGTGCGAGGGCATACAGTCCTCGCTTTTGTACACCGTAAGGAGGTGTACGAATAAGGGCTGGGCAAGACCGGTGCCAGCCGCCGCGGTAATACCGGCAGCCCGAGTGATGGCCGCTGTTATTGGGCCTAAAGCGTCCGTAGCCGGCCGAACAAGTCTGTCGGGAAATCCATCCGCCTAACGGATGGGCGTCCGGCAGAAACTGTTCGGCTTGGGGCCGGAAGATCCGAGGGGTACGTCCGGGGTAGGAGTGAAATCCCGTAATCCTGGACGGACCGCCGGTGGCGAAAGCGCCTCGGAAGGACGGACCCGACGGTGAGGGACGAAAGCTGGGGTCT |
| p__Halobacteriota_g__ Halosimplex_ASV_1132 | 0 | 1 | 0 | 0 | 0 | 0 | 0 | 0 | 0 | 0 | 1 | CCTTTACACTGCACGACAGTGCGATAGGGGGACTCCGAGTGCGAGGGCATACAGTCCTCGCTTTTCTGTACCGTAAGGTGGTACAGGAACAAGTGCTGGGCAAGACCGGTGCCAGCCGCCGCGGTAATACCGGCAGCACGAGTGATGGCCGATATTATTGGGCCTAAAGCGTCCGTAGCCGGCCCGACAAGTCCGTTGGGAAATCCACGCGCCCAACGCGTGGGCGTCCAGCGGAAACTGTCGGGCTTGGGGCCGGAGGACTCGAGGGGTACGTCCGGGGTAGGAGTGAAATCCCGTAATCCTGGACGGACCACCGGTGGCGAAAGCGCCTCGAGAGGACGGACCCGACGGTGAGGGACGAAAGCCAGGGTCT |
| p__Thermoplasmatota_ g__SM1-50_ASV_1131 | 0 | 0 | 0 | 0 | 0 | 0 | 0 | 0 | 1 | 0 | 1 | ACTTCACACTGTGCGAAAGCACGATGAGGGAATCCCAAGTGCTGGTTCGTAAGAGTCAGCTGTTTCTTTGTCTAAAAAACAGAGAGAGTAAGAGCTGGGTAAGACGGGTGCCAGCCGCCGCGGTAATACCTGCAGCTCAAGTGGTGGCCGCTATTATTGGGCCTAAAACGTCCGTAGCCTGTTTAGTAAATGCCTGGGTAAATCGTGTAGCTTAACTATACGAATTCCGGGTAGACTGCTGAACTTGAGATCGGGAGAAGTTAGAGGTACTCCTGGGGTAGAGGTGAAATTTTGTAATCCTAGGGGGACCACCAGTGGCGAAGGCGTCTGACTAGAACGAGTCTGACGGTGAGGGACGAAGCCCTGGGGCG |
| p__Nanoarchaeota_p__ Nanoarchaeota_ASV_1124 | 1 | 1 | 1 | 0 | 0 | 0 | 0 | 0 | 0 | 0 | 1 | CCTCTACAATGCACGAAAGTGTGATAGGGGAACTCTGCGTGCTTTCGGATATTTCCGGAAGCTTTTCTGGAGTGTAGATAGCTTCAGGAATAAGTGGTGGGTAAGACCGGTGCCAGCCGCCGCGGTAATCCCGGCGCCACAAGCAGTGATCACGATTATTGGGCCTAAAGCATCCGTAGCCGGCTCATTAAGTTTCTAGTGAAATCCAACAGCTTAACTGTTGGGCGTGCTAGAAATACTAATGGGCTTGAGACCGGGAGGAGTCAGAGGTATTCTTTGGGGAGCGGTAAAATGTTGTAATCCAAGGAGGACCATCAGTGGCGAAGGCGTCTGACTAGAACGGGTCTGACGGTGAGGGATGAAAGCTAGGGGAG |
| p__Halobacteriota_o__ Halanaerobiales_ASV_1138 | 0 | 1 | 1 | 0 | 0 | 0 | 0 | 0 | 0 | 0 | 0 | CCTTTACACTGCACGCCAGTGCGATAAGGGGACCCCGAGTGCGAGGGCATACAGTCCTCGCTTTTGTGTACCGTAGGGAGGTACACGAATAAGGGCTGGGCAAGACCGGTGCCAGCCGCCGCGGTAATACCGGCAGCTCAAGTGATGGCCGCTATTATTGGGCCTAAAGCGTCCGTAGCCGGCCGCACAAGTCCGTCGGGAAATCCACCCGCTTAACGGGTGGGCGTCCGGTGGAAACTGTGCGGCTTGGGACCGGAAGACTCGAGGGGTACGTCCGGGGTAGGAGTGAAATCCTGTAATCCTGGACGGACCGCCGGTGGCGAAAGCGCCTCGAGAGAACGGATCCGACGGTGAGGGACGAAAGCTAGGGTCT |
| p__Nanoarchaeota_g__ BMS3ABIN17_ASV_1147 | 1 | 0 | 0 | 0 | 0 | 0 | 0 | 1 | 0 | 1 | 1 | CTTCTGCAATGCGCGAAAGCGTGACAGAGTGAACCAGAGTGTTCCTCAACAGAGGAACTTTTGTGAAATGTAAAAAGTTTCACGAATAAGGACTGGGTAAGACTAGTGCCAGCCGCCGCGGTAATACTAGCGGTCCAAGTCGCAGCCACTATTATTGGGTCTAAAACATCCGCAGCTTGTTTAGTAAGTTTCTTGTGAAATCTTATGTCTCAAGCATAAGGCGGGCAAGAAGTACTGCTAAACTAGAGACCGGGAGAAGTAAGGAGTACGTTCAGAGTAGCGGTAAAATGTGTTAATCCTGGACGGACTAACAATAGCGAAGGCACCTTACTAGAACGGTTCTGACAGTCAGGGATGAAGGCTAGGGGCG |

| | Presence (1) or Absence (0) in each layer | | | | | | | | | | | |
|---|---|---|---|---|---|---|---|---|---|---|---|---|
| Phylum_Genus_ASV_number | L1 | L2 | L3 | L4 | L5 | L6 | L7 | L8 | L9 | L10 | L11 | Sequence |
| p__Nanoarchaeota_g__BMS3ABIN17_ASV_1157 | 0 | 0 | 0 | 1 | 0 | 0 | 1 | 1 | 1 | 0 | 0 | CTTCTGCAATGCGCGAAAGCGTGACAGAGTGAGTCAGAGTGTCCGCCATTCGGTGGACTTTTGTGGAATGTAAAGTATTTCACGAATAAGGACTGGGTAAGACTAGTGCCAGCCGCCGCGGTAATACTAGCGGTCCGAGTTGCAGCCACTATTATTGGGTCTAAAACATCCGTAGCTTGCTTGATAAGTCTCTTGTGAAATCTTGTATCTCAAGTATAAGACTTGCAAGAGATACTGTTAGGCTAGAGACCGGGAGACGTAAGGAGTACGTTCGAAGTAGCGGTAAAATGTGTTGATCTCGGACGGACTAACGAGAGCGAAGGCACCTTACGAGAACGGTTCTGACAGTGAGGGATGAAGGCCAGGGTCG |
| p__Thermoproteota_g__PALSA-986_ASV_1149 | 0 | 0 | 0 | 0 | 0 | 0 | 0 | 0 | 1 | 1 | 1 | ACTTTGCAATACACGAAAGTGTGACAGGGTCATCCCGAGTGCCGACCGCTGAGGTTGGCTTTTAACCAGTTTAGAAAGCTGGTAGAATAAGGAGAGGGCAAGTCTGGTGTCAGCCGCCGCGGTAATACCAGCTCTCCGAGTGGTGTGGACGTTTATTGGGCCTAAAGCATCCGTAGCTTGCTGGGCAAGTCCCCTGTTAAACCCACCGATTTAATCGTTGGCTTGCGGGGGAAACTGCTCGGCTAGGGGACGAGAGAGGCAGACGGTATTTCCGGGGTAGGGGTGAAATCCTATAATCCCGGGAAGACCACCAGTGGCGAAGGCTGTCTGCTAGAACGAGCCCGACGGTGAGGGATGAAAGCTGGGGGAG |
| p__Nanoarchaeota_g__BMS3ABIN17_ASV_1161 | 1 | 1 | 1 | 0 | 0 | 0 | 0 | 0 | 0 | 0 | 1 | AATCTACAATGCGCGAAAGCGTGATAGTTTGAGTCAGAGTGTTTCTCATTACGAGAAACTTTTGCCAATTGTAAAAAGATTGGCGAATAAGGACTGGGTAAGACTGGTGCCAGCCGCCGCGGTAATACCAGCGGTCCAAGTCGCAGCCAACATTATTGGGTCTAAAACATCCGTAGCTTGTTTCGTAAGTCTCTTGTGAAATCCGGCTTCTTAAGAGTCGGTCGGGCAAGAGATACTGCGAGGCTAGAGACTGGGAGGCGTAAAGAGTACAGTTAAGGTAGCGGTTAAATGCGTTGATCTTAACTGGACTAACAATAGCGAAGGCGCTTTACGAGAACAGTTCTGACAGTGAGGGATGAAGGCCAGGGGCG |
| p__Halobacteriota_g__Natrinema_ASV_1173 | 0 | 0 | 1 | 0 | 0 | 0 | 0 | 0 | 0 | 0 | 0 | CCTTTACACTGCACGAGAGTGCGATAAGGGGACTCCGAGTGCGAGGGCATATAGTCCTCGCTTTTCTCGACCGTAAGGAGGTCGAGGAATAAGTGCTGGGCAAGACCGGTGCCAGCCGCCGCGGTAATACCGGCAGCACGAGTGATGACCGCTATTATTGGGCCTAAAGCGTCCGTAGCTGGCCACGCAAGTCCATCGGGAAATCCGCGCGCTCAACGCGCGGGCGTCCGGTGGAAACTGCGTGGCTTGGGACCGGAAGACCAGAGGGGTACGTCCGGGGTAGGAGTGAAATCCCGTAATCCTGGACGGACCACCGGTGGCGAAAGCGCCTCTGGAAGACGGATCCGACGGTGAGGGACGAAAGCTCGGGTCA |
| p__Halobacteriota_g__Saliphagus_ASV_1174 | 0 | 0 | 1 | 0 | 0 | 0 | 0 | 0 | 0 | 0 | 0 | CCTTTACACTGCACGCCAGTGCGATAAGGGGATTCCAAGTGCGAGGGCATACAGTCCTCGCTTTTGTGTACCGTAGGAAGGTACACGAATAAGGGCTGGGCAAGACCGGTGCCAGCCGCCGCGGTAATACCGGCAGCCCGAGTGATGACCGATATTATTGGGCCTAAAGCGTCCGTAGCCGGCTACGCAAGTCCGTCGGGAAATCTGCTCGCTCAACGAGTAGGCGTCCGGCGGAAACTGCGTGGCTTGGGACCGGAAGACTCGAGGGGTACGTCCGGGGTAGGAGTGAAATCCCGTAATCCTGGACGGACCACCGGTGGCGAAAGCGCCTCGAGAGGACGGATCCGACGGTGAGGGACGAAAGCTAGGGTCA |
| p__Thermoplasmatota_g__SM1-50_ASV_1177 | 0 | 0 | 0 | 1 | 1 | 0 | 1 | 1 | 0 | 0 | 0 | TCTTCACAATGTGCGAGAGCACGATGAGGGAATCCCAAGTGCTAGCACGTTAGTGTTAGCTGTTTTCTAGTCTAAAAAACTAGGAGAGTAAGAGCTGGGTAAGACGGGTGCCAGCCGCCGCGGTAATACCTGCAGCTCGAGTGGTGGCCGCTATTATTGGGCCTAAAACGTCCGTAGCCGGTTTAGTAAATGCTTGGGTAAATTGTGTAGCTTAACTATACAAATACCGAGTAGACTGCTAGACTTGGGACCGGGAGAGGCTAGAGGTACTCCTGGGGTAGAGGTGAAATTCTGTAATCCTAGGGGGACCACCAGTGGCGAAGGCGTCTAGCTAGAACGGGTCTGACGGTGAGGGACGAAGCCCTGGGGCG |
| d__Archaea_d__Archaea_ASV_1162 | 1 | 0 | 0 | 0 | 0 | 0 | 0 | 0 | 0 | 0 | 1 | TCTCTTCAATGCACGACAGTGTGAAAGGGGAATTCTACGTGCTCATGTTTTCATGGGCTTTTGCATAGTGTAAATAACTACGCGAATAAGGGGTGGGAAAGACCGGTGCCAGCAGCCGCGGTAACACCGGCGCCCCGAGTGGCAATCAGTATTATTGGGCCTAAAGCGTCCGTAGCCTGAAAAATAAATCTTTTGTGAAATCTATCTGCTTAACAGGTAGGATTGTGAAAGACACTGTTTTTCTAGGGACCGGGAGGGGTCAGAAGTATTATAGGGGGAGCGGTAAAATGTGATAATCCTTATAAGACTACCGATGGCGAAAGCATCTGACCAGAACGGATCCGACGGTGAGGGACGAAGGCCAGGGAAG |
| p__Hadarchaeota_o__Hadarchaeales_ASV_1180 | 1 | 1 | 0 | 0 | 0 | 0 | 0 | 0 | 0 | 0 | 1 | ATTCCGCAATGCGTGAAAGCGCGACGGAGTGACTCCTAGTGCCGGTGGTCTAGCGACCGTCGGCTTTTTTGGAACGTAAAAAGTTCCAAGAATAAGGGCTGGGTAAGACTGGTGCCAGCCGCCGCGGTAACACCAGCAGCTCAAGTGGTAGCCGCTTTTATTGGGCCTAAAGCGTCCGTAGCCGGACCTGCAAGTCCCTTGTGAAATCGGGCGGCTTAACCGTTCGGCTTGCAAGGGAAACTGTTGGTCTTGGGGCCGATAGGCGCCGAGGGTACTCCGGGGGTAGCGGTGAAATGCTATAATCCCTGGGGGACCGCCGGTGGCGAAAGCGCTCGACGAGGTCGGACCCGACGGTGAGGGACGAAGGCCAGGGGAG |
| p__Halobacteriota_f__Haloferacaceae_ASV_1183 | 0 | 0 | 0 | 0 | 0 | 0 | 0 | 0 | 1 | 0 | 1 | CCTTTACACTGCACGAGAGTGCGATAGGGGGATCCCAAGTGCGAGGGCATATTAGCCCTCGCTTTTGTGAACCGTAAGGCGGTTCACGAATAAGAGCTGGGCAAGACCGGTGCCAGCCGCCGCGGTAATACCGGCAGCTCGAGTGATGACCGATTTTATTGGGCCTAAAGCGTCCGTAGCTGGCCACACAAGTCCGTCGGGAAATCTACCTGCCCAACAGGTAGGCGTCCGGCGGAAACTGTGTGGCTTGGGACCGGAAGGCGTGAAGGGTACGTCCGGGGTAGGAGTGAAATCCTGTAATCCTGGACGGACCACCGATGGCGAAAGCACTTCACGAGGACGGATCCGACAGTGAGGGACGAAAGCTAGGGTCT |
| p__Halobacteriota_g__Halopelagius_ASV_1186 | 0 | 1 | 1 | 1 | 0 | 0 | 0 | 0 | 0 | 0 | 0 | CCTTTACACTGCACGCAAGTGCGATAAGGGGACTCCGAGTGCGAGGGCATATAGTCCTCGCTTTTGTTCACCGTAGGAAGGTGTACGAATAAGAGCTGGGCAAGACCGGTGCCAGCCGCCGCGGTAATACCGGCAGCTCGAGTGATGACCGATTTTATTGGGCCTAAAGCGTCCGTAGCTGGCCAGACAAGTCCGTCGGGAAATCCATCTGCTCAACAGATGGGCGTCCGGCGGAAACTCTCTGGCTTGGGACCGGAAGGCTCGAAGGGTACGTCTGGGGTAGGAGTGAAATCCTGTAATCCTGGACGGACCACCGATGGCGAAAGCACTTCGAGAAGACGGATCCGACAGTGAGGGACGAAAGCTAGGGTCT |
| p__Halobacteriota_g__Halorussus_ASV_1190 | 0 | 0 | 1 | 0 | 0 | 0 | 0 | 0 | 0 | 0 | 0 | CCTTTACACTGCACGAAAGTGCGATAAGGGGACCCCGAGTGCGAGGGCATACAGTCCTCGCTTTTGTGTACCGTAAGGTGGTACACGAATAAGAGCTGGGCAAGACCGGTGCCAGCCGCCGCGGTAATACCGGCAGCTCGAGTGATGGCCGCTATTATTGGGCCTAAAGCGTCCGTAGCCGGCCGAACAGGTCCGTTGGGAAATCCATCTGCCTAACAGATGGGCGTCCAGCGGAAACCAGTCGGCTTGGGGCCGGAAGACCTGAGGGGTACGTCCGGGGTAGGAGTGAAATCCTGTAATCCTGGACGGACCGCCGGTGGCGAAAGCGCCTCAGGAGGACGGACCCGACGGTGAGGGACGAAAGCTAGGGTCA |
| p__Halobacteriota_g__Natronomonas_ASV_1185 | 0 | 1 | 1 | 0 | 0 | 0 | 0 | 0 | 0 | 0 | 0 | CCTTTACACTGCACGCCAGTGCGATAGGGGGACACCGAGTGCGAGGGCATAGAGCCCTCGCTTTTCTGGACCGTAAGGTGGTCCAGGAACAAGGGCTGGGCAAGACCGGTGCCAGCCGCCGCGGTAACACCGGCAGCCCAAGTGATGGCCGATATTATTGGGCCTAAAGCGTCCGTAGCTGGCCGAGCAAGTCCGTCGGGAAATCCACTCGCTCAACGAGTGGGCGTCCGGCGGAAACTGTTCGGCTTGGGACCGGAAGATCCGAGGGGTACGTCCGGGGTAGGAGTGAAATCCCGTAATCCTGGACGGACCACCGATGGCGAAAGCACCTCGGAAGGACGGATCCGACAGTGAGGGACGAAAGCTAGGGTCT |
| p__Halobacteriota_f__Haloarculaceae_3992_ASV_1188 | 0 | 1 | 1 | 1 | 0 | 0 | 0 | 0 | 0 | 0 | 0 | CCTTTACACTGCACGACAGTGCGATAAGGGGACTCCGAGTGCGAGGGCATACAGTCCTCGCTTTTCTGTACCGTAGGGAGGTACAGGAACAAGTGCTGGGCAAGACCGGTGCCAGCCGCCGCGGTAATACCGGCAGCACGAGTGATGGCCGATATTATTGGGCCTAAAGCGTCCGTAGCTTGCCGGACAAGTCCGTTGGGAAATCCACGCGCTTAACGTGTGGGCGTCCAGCGGAAACTGTCCGGCTTGGGGCCGGAGGACTCGAGGGGTACGTCCGGGGTAGGAGTGAAATCCTGTAATCCTGGACGGACCGCCGGTGGCGAAAGCGCCTCGAGAAGACGGACCCGACAGTGAGGGACGAAAGCCAGGGTCT |
| p__Halobacteriota_o__Halanaerobiales_ASV_1187 | 0 | 1 | 1 | 0 | 0 | 0 | 0 | 0 | 0 | 0 | 0 | CCTTTACACTGCACGAGAGTGCGATAAGGGGATCCCGAGTGCGAGGGCATATCGTCCTCGCTTTTCTGTACCGTAGGGAGGTACAGGAATAAGTGCTGGGCAAGACCGGTGCCAGCCGCCGCGGTAATACCGGCAGCACAAGTGATGACCGTTATTATTGGGCCTAAAGCGTCCGTAGCCGGCCAGACAAGTTCGTTGGGAAATCCACCCGCTCAACGGGTGGGCGTCCAGCGAAAACTGTTTGGCTTGGGGCCGGAAGACCCAGGGGGTACGTCCGGGGTAGGAGTGAAATCCTGTAATCCTGGACGGACCGCCGGTGGCGAAAGCGCCCTGGGAGGACGGACCCGACGGTCAGGGACGAAAGCTAGGGTCT |
| p__Halobacteriota_g__Halomicrobium_3849_ASV_1204 | 0 | 1 | 0 | 0 | 0 | 0 | 0 | 0 | 1 | 0 | 0 | CCTTTACACTGCACGCCAGTGCGATAAGGGGACTCCGAGTGCGAGGGCATATAGTCCTCGCTTTTCTGTACCGTAGGGAGGTACAGGAACAAGTGCTGGGCAAGACCGGTGCCAGCCGCCGCGGTAATACCGGCAGCACAAGTGATGGCCGTTCTTATTGGGCCTAAAGCGTCCGTAGCCGGCCAGACAAGTCTGTTGGGAAATCGACGCGCTCAACGCGTCGGCGTCCAGCGGAAACTGTCTGGCTTGGGGCCGGAAGACCTGAGGGGTACGTCCGGGGTAGGAGTGAAATCCCGTAATCCTGGACGGACCACCGGTGGCGAAAGCGCCTCAGGAAGACGGACCCGACGGTGAGGGACGAAAGCTAGGGTCT |
| p__Halobacteriota_g__Natrinema_ASV_1209 | 0 | 0 | 1 | 0 | 0 | 0 | 0 | 0 | 0 | 0 | 0 | CCTTTACACTGCACGCGAGTGCGATAAGGGGACTCCAAGTGCGAGGGCATATAGTCCTCGCTTTTCACCACCGTAAGGTGGTGGTGGAATAAGTGCTGGGCAAGACCGGTGCCAGCCGCCGCGGTAATACCGGCAGCACGAGTGATGACCGCTATTATTGGGCCTAAAGCGTCCGTAGCTGGCCATCCAAGTTCATCGGGAAATCCGCACGCTTAACGTGCGGGCGTCCGGTGGAAACTGTGTGGCTTGGGACCGGAAGACCAGAGGGGTACGTCCGGGGTAGGAGTGAAATCCCGTAATCCTGGACGGACCACCGGTGGCGAAAGCGCCTCTGGAAGACGGATCCGACGGTGAGGGACGAAAGCTCGGGTCA |
| p__Halobacteriota_g__Natronomonas_ASV_1207 | 0 | 0 | 1 | 0 | 0 | 0 | 0 | 0 | 0 | 0 | 0 | CCTTTACACTGCACGCCAGTGCGATAAGGGGATACCGAGTGCGAGGGCATAGAGCCCTCGCTTTTCTGGACCGTAGGGAGGTCCACGAATAAGGGCTGGGCAAGACCGGTGCCAGCCGCCGCGGTAACACCGGCAGCCCAAGTGATGGCCGATATTATTGGGCCTAAAGCGTCCGTAGCTGGCCGAGCAAGTTCGTCGGGAAATCCACGCGCTTAACGCGTGGGCGTCCGGCGAAAACTGTTCGGCTTGGGACCGGAAGACCCGAGGGGTACGTCCGGGGTAGGAGTGAAATCCCGTAATCCTGGACGGACCACCGATGGCGAAAGCACCTCGGGAGGACGGATCCGACAGTGAGGGACGAAAGCTAGGGTCT |
| p__Halobacteriota_o__Halanaerobiales_ASV_1206 | 0 | 1 | 1 | 0 | 0 | 0 | 0 | 0 | 0 | 0 | 0 | CCTTTACACTGCACGAGAGTGCGATAAGGGGACTCCGAGTGCGAGGGCATACTAGCCCTCGCTTTTTCCGACCGTAAGGAGGTCGGCGAATAAGGGCTGGGCAAGACCGGTGCCAGCCGCCGCGGTAATACCGGCAGCCCAAGTGATGGCCGCTTTTATTGGGCCTAAAGCGTCCGTAGCTGGCCGACCAAGTCCGTCGGGAAATCCACTGGCTCAACCAGTGGGCGTCCGGTGGAAACTAGTCGGCTTGAGACCGGAAGACTCAGAGGGTACGTCTGGGGTAGGAGTGAAATCCTGTAATCCTGGACGGACCGCCGATGGCGAAAGCACTCTGAGAGGACGGATCTGACAGTGAGGGACGAAAGCTCGGGTCT |
| p__Halobacteriota_o__Halanaerobiales_ASV_1208 | 0 | 0 | 1 | 0 | 0 | 0 | 0 | 0 | 0 | 0 | 0 | CCTTTACACTGCACGAAAGTGCGATAAGGGGACCCCGAGTGCGAGGGCATACAGTCCTCGCTTTTGTGTACCGTAAGGTGGTACACGAATAAGAGCTGGGCAAGACCGGTGCCAGCCGCCGCGGTAATACCGGCAGCTCGAGTGATGGCCGCTATTATTGGGCCTAAAGCGTCCGTAGCCGGCCGAACAGGTCCGTTGGGAAATCCGCCAGCCTAACTGGCGGGCGTCCAGCGGAAACCAGTCGGCTTGGGACCGGAAGACCTGAGGGGTACGTCTGGGGTAGGAGTGAAATCCTGTAATCCTGGACGGACCGCCGGTGGCGAAAGCGCCTCAGGAGGACGGATCCGACGGTGAGGGACGAAAGCTAGGGTCA |
| d__Archaea_d__Archaea_ASV_1201 | 0 | 0 | 0 | 0 | 0 | 0 | 0 | 0 | 1 | 1 | 1 | TCTCCACAATGCACTAACGTGTGATGGGGGGACTCTGAGTGCTCATCCTTTGGGTGAGCTTTTGTGAAGTCTAAAAAGCTTCGCGAATAAGGGCTGGGTAAGATCGGTGCCAGCAGCCGCGGTAATCCCGGCGGCCCAAGTGGTGGCCGTTTTTACTGGGCTTGAAGCGTCCGTAGCTGGACTAGTCCGTCTTTGGTTAAATCCTCAAGCTTAACTTGGGGGCTGCTGAAGATACCACTAGTCTAGGGACCGGGAGAGGACGGAGGTATTACAAGGGTAGGGGTAAAATCCGTTAATCCTTGTAGGACCACCAGTAGCGAAGGCGTCCGTCTAGAACGGGTCCGACAGTGAGGGACGAGAGCCAGGGGAG |
| p__Nanoarchaeota_g__BMS3ABIN17_ASV_1222 | 1 | 1 | 0 | 0 | 0 | 0 | 0 | 1 | 1 | 0 | 1 | CTTCTACAATGCGCGTAAGCGTGATAGAGCAAGCCAGAGTGCTTTTCCGTATAGGGAAAGCTTTTGTGTGATGTAAAAAGTCACACGAATAAGGACTGGGCAAGACTGGTGCCAGCCGCCGCGGTAATACCAGCAGTCCAAGTCGCAGCCGCTTTTATTGGGTCTAAAACATCCGTAGCTTGCTTTGCAAGTCTCTTGTGAAATCTGGCATCTCAAGTGTCAGGCGTGCAGGAGATACTGCTTGGCTAGGGACCGGGAGACGTAAGGAGTACGCATAGGGTAGCGGTAAAATGCGTTGATCCAATGTGGACTAACGATAGCGAAGGCACCTTACGAGAACGGATCTGACAGTGAGGGATGAAGGCTAGGGGCG |
| p__Halobacteriota_g__Halopiger_A_ASV_1231 | 0 | 0 | 1 | 0 | 0 | 0 | 0 | 0 | 0 | 0 | 0 | CCTTTACACTGCACGCCAGTGCGATAAGGGGACTCCACCTGCGAGGGCATATCGTCCTCGCTTTTCTGCACCGTAAGGTGGTACAGGAATAAGGGCTGGGCAAGACCGGTGCCAGCCGCCGCGGTAATACCGGCAGCCCGAGTGATGACCGCTATTATTGGGCCTAAAGCGTCCGTAGCTGGCCAGGCAAGTCCATCGGGAAATCCGCGCGCCCAACGCGCGGGCGTCCGGTGGAACCTGCTTGGCTTGGGACCGGAAGACCAGAGGGGTACGTCTGGGGTAGGAGTGAAATCCCGTAATCCTGGACGGACCACCGGTGGCGAAAGCGCCTCTGGAAGACGGATCCGACGGTGAGGGACGAAAGCTCGGGTCA |
| p__Halobacteriota_g__Natronoarchaeum_ASV_1229 | 0 | 1 | 0 | 0 | 0 | 0 | 0 | 0 | 0 | 0 | 0 | CCTTTACACTGCACGCCAGTGCGATAAGGGGACCCCCAGTGCGAGGGCATACCGTCCTCGCTTTTCACCACCGTAAGGAGGTAGTGGAACAAGAGCTGGGCAAGACCGGTGCCAGCCGCCGCGGTAATACCGGCAGCTCGAGTGATGGCCGTTTTTATTGGGCCTAAAGCGTCCGTAGCCGGCCAGACAAGTTCGTCGGGAAATCCACGTGCCCAACACGTGGGCGTCCGGCGAAACCTCTCTGGCTTGGGACCGGACGGCCCGAGGGGTACGGTCGGGGTAGGAGTGAAATCCTGTAATCCCGACCGGACCACCGGTGGCGAAAGCGCCTCGGGAGGACGGATCCGACGGTGAGGGACGAAAGCCAGGGTCA |

| Phylum_Genus_ASV_ number | Presence (1) or Absence (0) in each layer L1 | L2 | L3 | L4 | L5 | L6 | L7 | L8 | L9 | L10 | L11 | Sequence |
|---|---|---|---|---|---|---|---|---|---|---|---|---|
| p__Halobacteriota_g__Natronomonas_ASV_1230 | 0 | 1 | 1 | 0 | 0 | 0 | 0 | 0 | 0 | 0 | 0 | CCTTTACACTGCACGCCAGTGCGATAGGGGGACACCGAGTGCGAGGGCATAGAGCCCTCGCTTTTGTAGACCGTAAGGAGGTCTACGAATAAGGGCTGGGCAAGACCGGTGCCAGCCGCCGCGGTAACACCGGCAGCCCAAGTGATGGCCGATATTATTGGGCCTAAAGCGTCCGTAGCTGGCCGCACAAGTCCGTCGGGAAATCCACTCGCTTAACGAGTGGGCGTCCGGCGGAAACTGTGCGGCTTGGGACCGGAAGATCCGAGGGGTACGTCCGGGGTAGGAGTGAAATCCTGTAATCCTGGACGGACCACCGATGGCGAAAGCACCTCGGAAGGACGGATCCGACAGTGAGGGACGAAAGCTAGGGTCT |
| p__Halobacteriota_f__Natrialbaceae_ASV_1233 | 0 | 0 | 1 | 0 | 0 | 0 | 0 | 0 | 0 | 0 | 0 | CCTTTACACTGCACGCCAGTGCGATAAGGGGACTCCATCTGCGAGGGCATATCGTCCTCGCTTTTCTCGACCGTAAGGTGGTCGAGGAATAAGGGCTGGGCAAGACCGGTGCCAGCCGCCGCGGTAATACCGGCAGCCCAAGTGATGACCGCTATTATTGGGCCTAAAGCGTCCGTAGCTGGCCAGGCAAGTCCATCGGGAAATCCGCGTGCCCAACACGCGGGCGTCCGGTGGAACCTGCCTGGCTTGGGACCGGAAGACCAGAGGGGTACGTCCGGGGTAGGAGTGAAATCCCGTAATCCTGGACGGACCACCGGTGGCGAAAGCGCCTCTGGAAGACGGATCCGACGGTGAGGGACGAAAGCTCGGGTCA |
| p__Nanoarchaeota_g__BMS3ABIN17_ASV_1243 | 0 | 0 | 0 | 0 | 0 | 0 | 0 | 0 | 1 | 1 | 0 | AATCTACAATGCGCGCAAGCGTGATAGTTTGAGCCAGAGTGCTTTTCTATTTGAAAAGCTTTTGCCAAATGTAAAAAGTTTGGCGAATAAGGACTGGGTAAGACGAGTGCCAGCCGCCGCGGTAATACTCGCGGTCCAAGTCGCAGCCACATTTATTGGGTCTAAAACATCCGTAGCTTGCTTTGTAAGTTTCTTGTGAAATCCTATGTCTCAAGCATGGGGCGGGCAAGAAGTACTGCAAAGCTAGAGACTGGAAGACGTAAGAAGTACGTTTGAAGTAGCGGTAAAATGCGTTAATCTCAGACGGACTCACAATAGCGAAGGCATCTTACGAGGACAGTTCTGACAGTGAGGGATGAAGGCTAGGGGCG |
| p__Nanoarchaeota_g__DSQK01_ASV_1257 | 0 | 0 | 0 | 0 | 0 | 0 | 0 | 0 | 1 | 0 | 0 | CCTCCACAATGCACGCAAGTGTGATGGGGGGATTCCGAGTGCTCAGGGAATACCTGGGCTTTTTATAAGGGTAAAAACCTTGTAGAATAAGTGGTGGGCAAGACTGGTGCCAGCCGCCGCGGTAACCCCAGCGCCACAAGTGGGAACCATGTTTATTGGGTCTAAAGCATCCGTAGCCGGTTCTGCAAATACCCTGTGAAATCGTCCCGCTCAACGGGGCGGCGTGCAGGGTAGACTACAGGACTAGGGACCAGGTGAGGTGGACGGTATTCCGTGGGGAGCGGTAAAATGTGATAATCCTCGGAGGACCACCAATGGCGAAGGCAGTTCACCAAAATGGATCCGACGGTGAGGGATGAAAGCCAGGGGAG |
| p__Halobacteriota_f__Halobacteriaceae_4595_ASV_1245 | 0 | 1 | 0 | 0 | 0 | 0 | 0 | 0 | 0 | 0 | 0 | CCTTTACACTGCACGAAAGTGCGATAAGGGGACCCCGAGTGCGAGGGCATACAGTCCTCGCTTTTGTACACCGTAAGGTGGTGTACGAATAAGGGCTGGGCAAGACCGGTGCCAGCCGCCGCGGTAATACCGGCAGCCCAAGTGATGGCCGTTATTATTGGGCCTAAAGCGTCCGTAGCCGGCCGGGCAAGTCCGTTGGGAAACCCGCTCGCTCAACGAGCGGACGTCCAGCGGAAACTGTTCGGCTTGGGACCGGAAGACTCGAGGGGTACGTCCGGGGTAGGAGTGAAATCCTGTAATCCTGGACGGACCGCCGGTGGCGAAAGCGCCTCGAGAAGACGGATCCGACGGTGAGGGACGAAAGCTAGGGTCT |
| p__Thermoplasmatota_g__EX4484-6_ASV_1269 | 0 | 0 | 0 | 0 | 0 | 0 | 0 | 0 | 1 | 1 | 1 | ACTTCGCAATGCGCGCAAGCGCGACGAGGGGATTCCAAGTGCATGCACTCAGTGTATGCTGTTTTCCAGTCTAAAAAGCTGGGAGAGTAAGGGCTGGGTAAGACTGGTGCCAGCCGCCGCGGTAATACCAGCGGCCCTAGTGGTGATCGTTATTATTGGGCCTAAAGCGTCCGTAGCCGGCTCGGTAAATCTCTGGGTAAATCGTTGGGCTTAACTCAACGAATTCTGGGGAGACTGCCGGGCTTGGGACCGGGAGAGGTCGGAGGTACTCCAGGGGTAGGGGTGAAATCCTGTAATCCTTGGGGGACCACCGGTGGCGAAGGCGTCCGACCAGAACGGGTCCGACGGTAAGGGACGAAGCCCTGGGGCG |
| p__Halobacteriota_g__Haloprofundus_ASV_1271 | 0 | 1 | 0 | 0 | 0 | 0 | 0 | 0 | 0 | 0 | 0 | CCTTTACACTGCACGACAGTGCGACAAGGGGACCCCGAGTGCGAGGGCATACAGTCCTCGCTTTTGTGTACCGTAAGGTGGTACACGAATAAGAGCTGGGCAAGACCGGTGCCAGCCGCCGCGGTAATACCGGCAGCTCGAGTGATGACCGCTATTATTGGGCCTAAAGCGTCCGTAGCTGGCCGTGTAAGTCCATCGGGAAATCCATCCGCTCAACGGATGGGCGTCCGGTGGAAACTGCACGGCTTGGAGCCGGAAGACCCAAGGGGTACGTCCGGGGTAGGAGTGAAATCCCGTAATCCTGGACGGACCGCCGATGGCGAAAGCACCTTGGGAAGACGGACTCGACAGTGAGGGACGAAAGCTAGGGTCT |
| p__Nanoarchaeota_g__BMS3ABIN17_ASV_1298 | 0 | 0 | 1 | 1 | 0 | 0 | 0 | 0 | 0 | 0 | 0 | AATCTACAATGCGCGAAAGCGTGATAGTTTGAACCAGAGTGTATCTCTATTGAGATGCTTTTGCCAAATGTAAAAAGTTTGGCGAATAAGGACTGGGTAAGACAGGTGCCAGCCGCCGCGGTAATACCTGCGGTCCAAGTCGCAGCCATCTTTATTGGGTCTAAAACATCCGTAGCTTGCTTTGTAAGTTTCTTGTGAAATTCTGCATCTCAAGTGCAGAGCGCGCAGGAAATACTGTAGAGCTAGAGACCGGAAGACGTAAGGAGTACGTCTGGAGTAGCGGTAAAATGTACTAATCTCAGACGGACTAACAATAGCGAAAGCACCTTACGAGGACGGTTCTGACAGTCAGGGATGAAGGCCAGGGGCG |
| p__Halobacteriota_g__Haloprofundus_ASV_1293 | 0 | 1 | 0 | 0 | 0 | 0 | 0 | 0 | 0 | 0 | 0 | CCTTTACACTGCACGACAGTGCGACAAGGGGACCCCGAGTGCGAGGGCATACAGTCCTCGCTTTTGTGAACCGTAAGGAGGTTCACGAATAAGAGCTGGGCAAGACCGGTGCCAGCCGCCGCGGTAATACCGGCAGCTCGAGTGATGACCGCTGTTATTGGGCCTAAAGCGTCCGTAGCTGGCCGCGTAAGTCCATCGGGAAATCCATCCGCTCAACGGATGGGCGTCCGGTGGAAACTACACGGCTTGGAGCCGGAAGACCCAAGGGGTACGTCCGGGGTAGGAGTGAAATCCTGTAATCCTGGACGGACCGCCGATGGCGAAAGCACCTTGGGAAAACGGACTCGACAGTGAGGGACGAAAGCTAGGGTCT |
| p__Halobacteriota_g__Saliphagus_ASV_1296 | 0 | 0 | 1 | 0 | 0 | 0 | 0 | 0 | 0 | 0 | 0 | CCTTTACACTGCACGACAGTGCGATAAGGGGACTCCAAGTGCGAGGGCATATAGTCCTCGCTTTTGTGTACCGTAAGGTGGTACACGAATAAGGGCTGGGCAAGACCGGTGCCAGCCGCCGCGGTAATACCGGCAGCCCGAGTGATGACCGCTATTATTGGGCCTAAAGCGTCCGTAGCTGGCCGATCAAGTCCGTCGGGAAATCCGCACGCTTAACGTGCGGGCGTCCGGCGGAAACTGTTCGGCTTGGGACCGGAAGACCCGAGGGGTACGTCTGGGGTAGGAGTGAAATCCCGTAATCCTGGACGGACCGCCGGTGGCGAAAGCGCCTCGGGAGGACGGATCCGACGGTGAGGGACGAAAGCTAGGGTCA |
| p__Thermoplasmatota_g__SM1-50_ASV_1306 | 0 | 0 | 0 | 0 | 0 | 0 | 0 | 0 | 1 | 0 | 0 | CCTTCGCACTGTGCGAAAGCGCGATGAGGGGATCCCAAGTGCTTGCTCTTAGGAGTAAGCTGTTTCTCTGTTTAAAACGCAGGGAGAGTAAGGGCTGGGTAAGACGGGTGCCAGCCGCCGCGGTAATACCTGCAGCTCAAGTGGTGACCGCTATTATTGGGCCTAAAACGTTCGTAGCCGGCTGAGTAAATGCCTGGGTAAATCGTGCCGCTTAACGGTACGAATTCCGGGTAGACTGCTTAGCTTGAGACCGGGAGAAGCTAGAGGTACTCCTGGGGTAGGGGTGAAATCCTGTAATCCTGGGGGGACCACCAGTGGCGAAGGCGTCTAGCTAGAACGGGTCTGACGGTGAGGAACGAAACCCTGGGGCG |
| p__Halobacteriota_f__Halobacteriaceae_4595_ASV_1290 | 0 | 0 | 0 | 0 | 0 | 0 | 0 | 0 | 1 | 1 | 1 | CCTTTACACTGCACGCAAGTGCGATAAGGGAACCCCAAGTGCGAGGGCATACAGTCCTCGCTTTTGTGTACCGTAAGGTGGTACACGAATAAGAGCTGGGCAAGACCGGTGCCAGCCGCCGCGGTAATACCGGCAGCTCGAGTGATGGCCGCTATTATTGGGCCTAAAGCGTCCGTAGCCTGCCAGACAGGTCCGTTGGGAAATCTGCTCGCTTAACGAGCAGGCGTCCAGCGGAAACCCGCTGGCTTGGGACCGGAAGACCTGAGGGGTACGTCTGGGGTAGGAGTGAAATCCTGTAATCCTGGACGGACCGCCGGTGGCGAAAGCGCCTCAGGAAGACGGATCCGACGGTGAGGGACGAAAGCTAGGGTCA |
| p__Halobacteriota_o__Halanaerobiales_ASV_1287 | 1 | 1 | 1 | 0 | 0 | 0 | 0 | 0 | 0 | 0 | 0 | CCTTTACACTGCACGACAGTGCGATAGGGGGACTCCGAGTGCGAGGGCATATAGTCCTCGCTTTTCTGTACCGTAAGGTGGTACAGGAACAAGTGCTGGGCAAGACCGGTGCCAGCCGCCGCGGTAATACCGGCAGCACGAGTGATGGCCGATCTTATTGGGCCTAAAGCGTCCGTAGCTTGCCGATCAAGTCTATCGGGAAATCGACTCGCTCAACGAGTCGGCGTCCGGTGGAAACTGTTCGGCTTGGGGCCGGAAGACTCAGCGGGTACGTCCGGGGTAGGAGTGAAATCCTGTAATCCTGGACGGACCACCGATGGCGAAAGCACGCTGAGAGCCCGGACCCGACAGTGAGGGACGAAAGCTAGGGTCT |
| d__Archaea_d__Archaea_ASV_1291 | 0 | 0 | 0 | 0 | 0 | 0 | 0 | 0 | 1 | 1 | 1 | CCTCCGCAATGCTCGAGAGAGTGACGGGGAGATTCCAAGTGCCTACGCACAGCGTAGGCTGTTCAGATGCCTACAACGCATCTGGAGGAAGGACTGGGTAAGACGGGTGCCAGCCGCCGCGGTAACACCCGCAGTTCAAGTGGCAGTCGATTTTATTGGGCCTAAAGCGTCCGTAGCCGGTCTGGCAAATGCCTGGGTAAATCGTCTGGCTCAACCAGACGAATTCCGGGTAGACTGCCAGACTTGGGACCGGTTGAGGGAGAGGGTACTCCAGGGGTAGGAGTGAAATCTTGTAATCCTTGGAGGACCACCAGTGGCGAAAGCGCTCTCCCAGGACGGGTCCGACGGTGAGGGACGAAAGCCAGGGTAG |
| d__Archaea_d__Archaea_ASV_1305 | 0 | 0 | 0 | 0 | 0 | 0 | 0 | 1 | 1 | 0 | 0 | CCTCTGCAATGCGCGAAAGCGTGACAGGGGAACTCCGAGTGGCGGAGACTAAGTCTCCGTCTTTTGCCAAGTGTAAATAGCTTGGAGAATAAGGGTGTGGGCAAGACTGGTGGCAGCCGCCGCGGTAACACCAGCGCCTCGAGTGGTACCCACGTATACTGGGCCTAAAGCGTCCGTAGCCGGCTTGAAAAGTCCTCTGTGAAATAGTGGGGCTTAACTCCACTGCGCGCAGGGGATACTTTCTGGCTTGGGAGCGGGGGAGGTAAGGGGTACACCTAGGGTAGGGGTAAAATCCGTTGAGCCTGGGTGGACCACCAGTGGCGAAGGCGCCTTACCAAAACGCGTCCGACGGTGAGGGACGAAAGCTGGGGGAG |
| p__Nanoarchaeota_c__Nanoarchaeia_ASV_1312 | 0 | 0 | 0 | 0 | 0 | 0 | 0 | 0 | 1 | 1 | 1 | CCTTTACAATGCGCGAAAGCGTGATAAGGGGATTCCAAGTGCCATAACTATGTTATGGCTGTGGCGAAGTGTAAAGTACTTCGCAAGAAAGGGGTGGGTAATACCGGTGCCAGCAGCCGCGGTAACCCCGGAGCCCCGAGTGGTGGCCATTTTTATTGGGCCTAAAGCGCCCGTAGCCGGGTTAGTAAGTTCTCGGTGAAATCGTTCAGCTTAACTGAACGAATTGCCTTGAATACTGCTAACCTAGGGATAGGGAGAGGTTAAGAGTATGCCTGGGGGAGGGGTAAAATCCTTTAATCCCAGGTGGACTATCAATGGCGAAGGCACTTAACTGGAACTAGTCCGACGGTGAGGGACGAAAGCTAGGGTAG |
| p__Halobacteriota_g__Halorussus_ASV_1307 | 0 | 0 | 0 | 0 | 0 | 0 | 0 | 0 | 0 | 1 | 0 | ACTTTACACTGCACGACAGTGCGATAAGGGAACTCCGAGTGCGAGGGCATATAGTCCTCGCTTTTCTGCACCGTAAGGTGGTGCACGAATAAGAGCTGGGCAAGACCGGTGCCAGCCGCCGCGGTAATACCGGCAGCTCAAGTGATGGCCGCTATTATTGGGCCTAAAGCGTCCGTAGCCAGCCAGACAGGTCCGTCGGGAAATCCACGCGCTCAACGCGTGGGCGTCCGGCGGAAACCAGCTGGCTTGGGGCCGGAAGATCTGAGGGGTACGTCCGGGGTAGGAGTGAAATCCCGTAATCCTGGACGGACCACCGGTGGCGAAAGCGCCTCAGAAAAACGGACCCGACGGTGAGGGACGAAAGCTAGGGTCA |
| p__Halobacteriota_g__Halorussus_ASV_1316 | 0 | 1 | 1 | 0 | 0 | 0 | 0 | 0 | 0 | 0 | 0 | CCTTTACACTGCACGAAAGTGCGATAAGGGGATCCCAAGTGCGAGGGCATACAGTCCTCGCTTTTCTGTACCGTAGGGAGGTACAAGAATAAGGGCTGGGCAAGACCGGTGCCAGCCGCCGCGGTAATACCGGCAGCCCAAGTGATGGCCGCTATTATTGGGCCTAAAGCGTCCGTAGCCGGCCGGATAAGTCCGTCGGGAAATCCACCCGCTTAACGGGTGGGCGTCCGGCGGAAACTGTCTGGCTTGGGGTCGGAGGGCCTGAGGGGTACGTCCGGGGTAGGAGTGAAATCCTGTAATCCTGGTCGGACCACCGGTGGCGAAAGCGCCTCAGGAGGACGAACCCGACGGTGAGGGACGAAAGCTAGGGTCA |
| p__Nanohaloarchaeota_g__Nanohalobium_ASV_1319 | 0 | 1 | 1 | 1 | 0 | 1 | 0 | 0 | 0 | 0 | 0 | CCTTCACAATGCACGAAAGTGCGATGGGGGGATTCCGAGTGTAGTCTTCCCACAAGGAAGGCTACTTTTCTGTACTGTAAGGAGGTACTGGAACAAGGGGTGGGCAATACTGCTGCCAGCCGCCGCGGTAATAGCAGAACCCCAAGTGGTAGTCGTTATTATTGGGCTTAAAGTGTTCGTAGCCGGCCGTGCAAGTCCGCTGTGAAATCCAAGCGCTTAACGTTTGGGCGTGCAGCGGAAACTGCACTGGCTAGGGACCGGACGGTGTCGAGGCTATTCCAGGAGTAGGGGTGAAATCCTAAAATTCCTGGAGGAGCGCCTGTGGCGGAAGCGCTCGACAAGTACGGATCCGACGGTGAGGAACGAAAGCCAGGGGAG |
| p__Thermoplasmatota_g__SM1-50_ASV_1309 | 0 | 0 | 0 | 0 | 0 | 0 | 0 | 0 | 0 | 1 | 1 | ACTTCACAATGCGCGAAAGCGCGATGAGGGAATCCCAAGTGCTTGCACGTAAGTGTAAGCTGTTTTTATGTGTAAATAGCATAGAGAGTAAGGGCTGGGTAAGACGGGTGCCAGCCGCCGCGGTAATACCTGCAGCTCAAGTGGTGACCACTATTATTGGGCCTAAAACGTTCGTAGCCGGTTTATTAAATGCCTGGGTAAATCGTACGGCTTAACCGTACGAATTCCGGGTAGACTGATAGACTTGAGACCGGGAGAAGTTAGAGGTACTCCTGGGGTAGAGGTGAAATTCTGTAATCCTGGGGGGACCACCAGTGGCGAAGGCGTCTAACTAGAACGGGTCTGACGGTGAGGGACGAAGCCCTGGGGCG |
| p__Halobacteriota_f__Haloferacaceae_ASV_1320 | 0 | 1 | 1 | 0 | 0 | 0 | 0 | 0 | 0 | 0 | 0 | CCTCAACACTGCACGCCAGTGCGATTGGGGGACTCCGAGTGCGAGGGCATACCGTCCTCGCTTTTCAGCACGGTAAGGACGTGTTGGAACAAGGGCTGGGCAAGACCGGTGCCAGCCGCCGCGGTAATACCGGCAGCCCGAGTGATGACCGATCTTATTGGGCCTAAAGCGTCCGTAGCCGGCCGGGCAAGTCCGTCGGGAAATCCGCGCGCTCAACGCGTGGGCGTCCGGCGGAACCTGTCCGGCTTGGGGCCGGAAGACCCAGGGGGTACGTCCGGGGTAGGAGTGAAATCCCGTAATCCCGGACGGACCGCCGATGGCGAAAGCACCCTGGGAGGACGGACCCGACGGTGAGGGACGAAAGCTGGGGTCT |
| p__Halobacteriota_o__Halanaerobiales_ASV_1321 | 0 | 0 | 1 | 0 | 0 | 0 | 0 | 0 | 0 | 0 | 0 | CCTTTACACTGCACGACAGTGCGATAGGGGGACTCCGAGTGCGAGGGCATACAGTCCTCGCTTTTTTGTACCGTAAGGAGGTACAAGAATAAGGGCTGGGCAAGACCGGTGCCAGCCGCCGCGGTAATACCGGCAGCCCAAGTGATGGCCGCTATTATTGGGCCTAAAGCGTCCGTAGCCGGCCAGTCAAGTCCGTTGGGAAATCCACGCGCCTAACGCGTGGGCGTCCAGCGGAAACTGACTGGCTTGGGGCCGGAAGACCTGAGGGGTACGTCCGGGGTAGGAGTGAAATCCCGTAATCCTGGACGGACCACCGGTGGCGAAAGCGCCTCAGGAAGACGGACCCGACGGTGAGGGACGAAAGCTAGGGTCT |
| p__Thermoproteota_g__GCA-2726865_ASV_1339 | 0 | 0 | 0 | 0 | 0 | 0 | 0 | 0 | 1 | 0 | 1 | CCTCCTCAATGCACTCACGTGCGAAGGGGTTACCTCGAGTGCTACCTGAAGAAGGTGGCTTTTCCTTGGTCTAAAAAGCCTTGGGAATAAGAGGGGGGCAAGACTGGTGTCAGCCGCCGCGGTAACACCAGCCCCTCGAGTGGTGGGGATGATTATTTGGCCTAAAGCGTCCGTAGCCGGCCGGGTAAGTCTCCCGTTAAATCCAGCGTCTCAAGCGTTGGACTGCGGGAGATACTGTCTGGCTAGAGAGTGGGAGAGGTGTACGGTATTCTGGGGGTAGGGGCGAAATCCTTTGATCCCCGGAGGACCACCAGTGGCGAAGGCTGTACACTAGAACACGTCTGACGGTCAGGGACGAAAGCTGGGGTAG |

| Phylum_Genus_ASV_number | Presence (1) or Absence (0) in each layer | | | | | | | | | | | |
|---|---|---|---|---|---|---|---|---|---|---|---|---|
| | L1 | L2 | L3 | L4 | L5 | L6 | L7 | L8 | L9 | L10 | L11 | Sequence |
| p__Halobacteriota_g__ Halobacterium_ASV_1344 | 0 | 1 | 1 | 0 | 0 | 0 | 0 | 0 | 0 | 0 | 0 | CCTTTACACTGTACGCAAGTGCGATAAGGGGACTCCGAGTGTGAAGGCATAGAGCCTTCACTTTTGTACACCGTAAGGTGGTGTACGAATAAGGGCTGGGCAAGACCGGTGCCAGCCGCCGCGGTAATACCGGCAGCCCGAGTGATGGCCGATATTATTGGGCCTAAAGCGTCCGTAGCTGGCCGGACAAGTCCGTTGGGAAATCTGCTCGCTTAACGAGCAGGCGTCCAGCGGAAACTGTTCGGCTTGGGACCGGAAGACCTGAGGGGTACGTCCGGGGTAGGAGTGAAATCCCGTAATCCTGGACGGACCACCGGTGGCGAAAGCGCCTCAGGAGGACGGATCCGACAGTGAGGGACGAAAGCTAGGGTCT |
| p__Halobacteriota_g__ Halovenus_ASV_1345 | 0 | 1 | 0 | 0 | 0 | 0 | 0 | 1 | 1 | 0 | 0 | CCTTTACACTGCACGACAGTGCGATAGGGGGACTCCGAGTGCGAGGGCATATAGTCCTCGCTTTTGTACACCGTAAGGTGGTGTACGAATAAGAACTGGGCAAGACCGGTGCCAGCCGCCGCGGTAATACCGGCAGTTCGAGTGATGGCCGCTTTTATTGGGCCTAAAGCGTTCGTAGCTGGCCAGTCAAGTCTGTCGGGAAATCCAGTGGCTTAACCACTGGACGTCCGGCGGAAACTGATTGGCTTGGGACCGGGAGACTCGAGGAGTACGTTCGGGGTAGGAGTGAAATCCCATAATCCTGAACGGACTACCGATGGCGAAAGCACCTCGAGAGCACGGATCCGACAGTGAGGAACGAAAGCTGGGGTCT |
| p__Thermoproteota_g__ PALSA-986_ASV_1335 | 0 | 0 | 0 | 0 | 0 | 0 | 0 | 0 | 0 | 1 | 1 | ACTTTGCAATACACGAAAGTGTGACAGGGCTATCCCGAGTGCCATCCGCTGAGGAAGGCTTTTACCCAGTATAGCAAGCTGGGAGAATAAGGAGAGGGCAAGACTGGTGTCAGCCGCCGCGGTAATACCAGCTCTCCGAGTGGTGTGGATGTTTATTGGGCCTAAAGCATCCGTAGCTGGCTGGGCAAGTCCCCTGTTAAATCCACTGATTTAATCATTGGTGTGCGGGGGATACTGCTCGGCTAGGGGACGGGAGAGGCAGACGGTATTTCCGGGGTAGGGGTGAAATCCTATAATCCCGGGAAGACCACCAGTGGCGAAGGCTGTCTGCTAGAACGCGCCCGACGGTGAGGGATGAAAGCTGGGGGAG |
| p__Halobacteriota_o__ Halanaerobiales_ASV_1343 | 0 | 1 | 1 | 0 | 0 | 0 | 0 | 0 | 0 | 0 | 0 | CCTTTACACTGCGCGAAAGCGCGATAAGGGGACTCCAAGTGCGAGGGCATAAAGTCCTCGCTTTTCTACACCGTAAGGTGGTATAGGAATAAGGGCTGGGCAAGACCGGTGCCAGCCGCCGCGGTAATACCGGCAGCCCAAGTGATGGCCGCTATTATTGGGCCTAAAGCGTCCGTAGCCTGCTGGGTAAGTCCGTCGGGAAATCCACGCGCCTAACGTGTGGGCGTCCGGCGGAAACTGCCCGGCTTGGGACCGGGAGACCCGAGGGGTACGTCTGGGGTAGGAGTGAAATCCTGTAATCCTGGACGGACCGCCGGTGGCGAAAGCGCCTCGGGAGAACGGATCCGACGGTGAGGGACGAAAGCTAGGGTCA |
| p__Halobacteriota_o__ Halanaerobiales_ASV_1347 | 0 | 0 | 1 | 0 | 0 | 0 | 0 | 0 | 0 | 0 | 0 | CCTTTACACTGCACGACAGTGCGATAGGGGGACTCCGAGTGCGAGGGCATACAGTCCTCGCTTTTCTGTACCGTAAGGTGGTACAAGAATAAGTGCTGGGCAAGACCGGTGCCAGCCGCCGCGGTAATACCGGCAGCACGAGTGATGGCCGATATTATTGGGCCTAAAGCGTCCGTAGCCGGCCAGCCAAGTCCGTTGGGAAATCCACACGCTTAACGTGTGGGCGTCCAGCGGAAACTGTCTGGCTTGGGACCGGAAGACCCGAGGGGTACGTCCGGGGTAGGAGTGAAATCCCGTAATCCTGGACGGACCACCGGTGGCGAAAGCGCCTCGGGAAGACGGATCCGACGGTGAGGGACGAAAGCCAGGGTCT |
| p__Nanoarchaeota_g__ BMS3ABIN17_ASV_1378 | 0 | 0 | 0 | 1 | 0 | 0 | 0 | 0 | 0 | 0 | 0 | AATCTACAATACGCGCAAGCGTGATAGTTTGAGTCAGAGTGCGTTTCAATTGAAATGCTTTTGCCAAATGTAAAAAGTTTGGCGAATAAGGACTGGGTAAGACGAGTGCCAGCCGCCGCGGTAATACTCGCGGTCCAAGTCGCAGCCATTTTTATTGGGTCTAAAACATCCGTAGCTTGCTCTGCAAGTCTCTTGTGAAATTCTATCTCTTAAGGGTAGAGCGGGCAAGAGGTACTACAAAGCTAGAGACTGGGAGACGTAAGAAGTACGTTCGAAGTAGCGGTAAAATGTGTTAATCTCGAGCGGACTACCAACAGCGAAGGCATCTTACGAGAACAGTTCTGACAGTAAGGGATGAAAGCTAGGGGCG |
| p__Nanoarchaeota_g__ BMS3ABIN17_ASV_1389 | 0 | 0 | 0 | 0 | 0 | 0 | 0 | 1 | 1 | 0 | 0 | ATTCTGCAATGCGCGAAAGCGTGACAGATTGAGCCAGAGTGCTTCTCAAATGAGAAGCTTTTGAGCACTGTAAAAAGGTGCAAGAATAAGGACTGGGTAAGACGAGTGCCAGCCGCCGCGGTAATACTCGCGGTCCAAGTCGCAGCCATATTTATTGGGTCTAAAACATCCGTAGCTTGCTTGATAAGTTCCCTGTGAAATTTTATCTCTTAAAGGTAAAGCGTGCAGGAAATACTGTCAAGCTAGAGACTGGAAGACGTAGGAAGTACGATTAAAGTAGCGGTAAAATGTGTTGATCTTAATCGGACTCACAATAGCGAAGGCATCCTACGAAGACAGTTCTGACAGTCAGGGATGAAGGCTAGGGGCG |
| p__Halobacteriota_g__ Haloferax_ASV_1371 | 0 | 1 | 1 | 0 | 0 | 0 | 0 | 0 | 0 | 0 | 0 | CCTTTACACTGTACGCAAGTGCGATAAGGGGACCCCAAGTGCGAGGGCATATAGTCCTCGCTTTTCCCGACTGTAAGGCGGTCGGGGAATAAGGGCTGGGCAAGACCGGTGCCAGCCGCCGCGGTAATACCGGCAGCCCAAGTGATGACCGATATTATTGGGCCTAAAGCGTCCGTAGCCGGCCACGAAGGTCCATCGGGAAATCCGCCAGCTTAACTGGCGGGCGTCCGGTGGAAACCACGTGGCTTGGGACCGGAAGGCTCGAGGGGTACGTCCGGGGTAGGAGTGAAATCCTGTAATCCTGGACGGACCACCGATGGCGAAAGCACCTCGAGAAGACGGATCCGACGGTGAGGGACGAAAGCTAGGGTCT |
| p__Halobacteriota_g__ Halorussus_ASV_1369 | 0 | 1 | 1 | 0 | 1 | 0 | 0 | 0 | 0 | 0 | 0 | CCTTTACACTGTACGCAAGTACGATAAGGGGACCCCGAGTGCGAGGGCATACAGTCCTCGCTTTTGTACACCGTAAGGAGGTGTACGAATAAGGGCTGGGCAAGACCGGTGCCAGCCGCCGCGGTAATACCGGCAGCCCGAGTGATGGCCGCTATTATTGGGCCTAAAGCGTCCGTAGCCGGCCGGACAAGTCTGTCGGGAAATCCATCCGCCTAACGGATGGGCGTCCGGCAGAAACTGTTCGGCTTGGGGCCGGAAGACCCGAGGGGTACGTCTGGGGTAGGAGTGAAATCCTGTAATCCTGGACGGACCGCCGGTGGCGAAAGCGCCTCGGGAGGACGGACCCGACGGTGAGGGACGAAAGCTAGGGTCT |
| p__Thermoproteota_g__ Nitrososcosmicus_ASV_1384 | 0 | 0 | 0 | 0 | 0 | 1 | 0 | 0 | 0 | 0 | 0 | CCTCTGCAATAGGCGAAAGCCTGACAGGGTTACTCTGAGTGATCTCCGTTAAGGAGGTCTTTTGGCATCTCTAAAAAGGGTGCAGAATAAGGGGTGGGCAAGTCTGGTGTCAGCCGCCGCGGTAATACCAGCACCCCGAGTGGTCGGGACGTTTATTGGGCCTAAAGCATCCGTAGCCGGTTCTGCAAGTCTTCCGTTAAATCCACCTGCTTAACAGATGGGCTGCGGAAGATACTACAGAGCTAGGAGGCGGGAGAGGCAAGCGGTACTCGATGGGTAGGGGTAAAATCCGTTGATCCATTGAAGACCACCAGTGGCGAAGGCGGCTTGCCAGAACGCGCTCGACGGTGAGGGATGAAAGCTGGGGGAG |
| p__Asgardarchaeota_g__SDNM01_ASV_1368 | 0 | 1 | 1 | 1 | 0 | 0 | 0 | 0 | 0 | 0 | 1 | CCTCCGCAATACACGAAAGTGTGACGGGGTTACCCAAAGTGTCCAACATTGGACTGTGGTAGGTGAGTAATGTCCCCTACTAGAAAGGAGAGGGCAAGGCTGGTGCCAGCCGCCGCGGTAAAACCAGCTCTTCAAGTGGTCGGGATAATTATTGGGCTTAAAGTGTCCGTAGCCGGATTAGTAAGTTCCTGGTAAAATCAGGTAGCTTAACTATCTGCATGCTAGGAATACTGCTATTCTAGAGGGCGGGAGAGGTCTGAGGTACTACAGGGGTAGGGGTGAAATCTTATAATCCTTGTAGGACCACCAGTGGCGAAGGCGTCAGACTGGAACGCGCCTGACGGTGAGGGACGAAAGCCAGGGGAG |
| p__Halobacteriota_f__Natrialbaceae_ASV_1359 | 1 | 0 | 0 | 0 | 0 | 0 | 0 | 0 | 0 | 0 | 0 | CCTTTACACTGCACGCCAGTGCGATAAGGGGACTCCCAGTGCGAGGGCATATCGTCCTCGCTTTTGTGTACCGTAGGGAGGTACACGAATAAGGGCTGGGCAAGACCGGTGCCAGCCGCCGCGGTAATACCGGCAGCCCAAGTGATGACCGCTATTATTGGGCCTAAAGCGTCCGTAGCTGGCCACGCAAGTCCGTCGGGAAATCCGCACGCCCAACGTGCGGGCGTCCGGCGGAAACTGCCTGGCTTGGGACCGGAAGACCAGAGGGGTACGTCCGGGGTAGGAGTGAAATCCCGTAATCCTGGACGGACCACCGGTGGCGAAAGCGCCTCTGGAAGACGGATCCGACGGTGAGGGACGAAAGCTCGGGTCA |
| p__Halobacteriota_g__ Halorussus_ASV_1410 | 0 | 1 | 1 | 0 | 0 | 0 | 0 | 0 | 0 | 0 | 0 | CCTTTACACTGCACGAAAGTGCGATAGGGGGACCCCGAGTGCGAGGGCATACAGTCCTCGCTTTTGTGTACCGTAAGGTGGTACACGAATAAGAGCTGGGCAAGACCGGTGCCAGCCGCCGCGGTAATACCGGCAGCTCGAGTGATGGCCGCTGTTATTGGGCCTAAAGCGTCCGTAGCCGGCCGAACAGGTCCGTTGGGAAATCCATCTGCCTAACAGATGGGCGTCCAGCGGAAACCAGTCGGCTTGGGGCCGGAAGACCTGAGGGGTACGTCCGGGGTAGGAGTGAAATCCTGTAATCCTGGACGGACCGCCGGTGGCGAAAGCGCCTCAGGAGGACGGACCCGACGGTGAGGGACGAAAGCTAGGGTCA |
| p__Halobacteriota_g__Saliphagus_ASV_1407 | 0 | 1 | 1 | 0 | 0 | 0 | 0 | 0 | 0 | 0 | 0 | CCTTTACACTGCACGACAGTGCGATAGGGGGACTCCAAGTGCGAGGGCATATAGTCCTCGCTTTTCACTACCGTAAGGTGGTAGTAGAATAAGTGCTGGGCAAGACCGGTGCCAGCCGCCGCGGTAATACCGGCAGCACGAGTGATGACCGCTATTATTGGGCCTAAAGCGTCCGTAGCCGGCCGAACAAGTCCGTCGGGAAATCCGCACGCCTAACGTGCGGGCGTCCGGCGGAAACTGTTCGGCTTGGGACCGGAAGGCTCGAGGGGTACGTCCGGGGTAGGAGTGAAATCCCGTAATCCTGGACGGACCACCGGTGGCGAAAGCGCCTCGAGAGGACGGATCCGACGGTGAGGGACGAAAGCTAGGGTCA |
| p__Halobacteriota_f__Natrialbaceae_ASV_1406 | 0 | 1 | 0 | 0 | 0 | 0 | 0 | 0 | 1 | 0 | 0 | CCTTTACACTGCACGACAGTGCGATAGGGGGACTCCGAGTGCGAGGGCATATAGTCCTCGCTTTTCACCACCGTAAGGTGGTGGTGGAATAAGTGCTGGGCAAGACCGGTGCCAGCCGCCGCGGTAATACCGGCAGCACGAGTGATGGCCGCTATTATTGGGCCTAAAGCGTCCGTAGCCGGCCGTGCAAGTCCATCGGGAAATCTCTCCGCCCAACGGAGAGGCGTCCGGTGGAAACTGTTCGGCTTGGGACCGGAAGACCTGAGGGGTACGTCCGGGGTAGGAGTGAAATCCCGTAATCCTGGACGGACCACCTGTGGCGAAAGCGCCTCAGGAGAACGGATCCGACGGTGAGGGACGAAAGCTAGGGTCA |
| p__Halobacteriota_o__ Halanaerobiales_ASV_1392 | 1 | 0 | 0 | 0 | 1 | 0 | 0 | 0 | 0 | 0 | 0 | CCTTTACACTGCACGCCAGTGCGATAGGGGGACCCCGAGTGCGAGGGCATACAGTCCTCGCTTTTATGCACCGTAAGGAGGTGCATGAATAAGTGCTGGGCAAGACCGGTGCCAGCCGCCGCGGTAATACCGGCAGCACAAGTGATGGCCGCTATTATTGGGCCTAAAGCGTCCGTAGCCGGCCGAACAAGTCCGTCGGGAAATCCATCCGCTCAACGGATGGGCGTCCGGCGGAAACTGTTCGGCTTGGGACAGGAAGACTCGAGGGGTACGTCCGGGGTAGGAGTGAAATCCCGTAATCCTGGACGGACCGCCGATGGCGAAAGCACCTCGAGAAGACTGATCCGACGGTGAGGGACGAAAGCTGGGGTCT |
| p__Halobacteriota_o__ Halanaerobiales_ASV_1404 | 0 | 1 | 0 | 0 | 0 | 0 | 0 | 0 | 0 | 0 | 0 | CCTTTACACTGCACGCCAGTGCGATAGGGGGACACCGAGTGCGAGGGCATAGAGCCCTCGCTTTTGTGTACCGTAAGGTGGTACACGAATAAGGGCTGGGCAAGACCGGTGCCAGCCGCCGCGGTAACACCGGCAGCCCAAGTGATGGCCGCTATTATTGGGCCTAAAGCGTCCGTAGCCGGCCGGGCAGGTCCGTCGGGAAATCCACTGGCTCAACCAGTGGGCGTCCGGCGGAAACCGCTCGGCTTGGGACCGGGAGACCAGAGGGGTACGTCCGGGGTAGGAGTGAAATCCCGTAATCCTGGACGGACCACCGGTGGCGAAAGCGCCTCTGGAGAACGGATCCGACGGTGAGGGACGAAAGCTAGGGTCT |
| d__Archaea_d__Archaea_ASV_1393 | 1 | 0 | 0 | 0 | 0 | 0 | 0 | 0 | 0 | 0 | 1 | AATCTACAATGCGCGCAAGCGTGATAGTTTGAGTCAGAGTGCGCTTCAATTTGGAGTGCTTTTGTGAAATGTAAAAAGTTTTACGAATAAGGACTGGGTAAGACGAGTGCCAGCCGCCGCGGTAATACTCGCGGTCCAAGTCGCAGCCATTTTTATTGGGTCTAAAACATCCGTAGCTTGCCATATTAAGTTCCTTGTGAAATCCTATATCTAAAATATAGGGCGGCAGGGAATACTAATTAGCTAGAGACTGGAAGACGTGATGAGTACGTTCAAAGTAGCGGTCAAATGTGTTAATCTTGGACGGACTAACAACAGCGAAGGCACATCACGAGGACAGTTCTGACAGTAAGGGATGAAAGCTGGGGGCG |
| p__Halobacteriota_g__ Haloprofundus_ASV_1436 | 0 | 1 | 0 | 0 | 0 | 0 | 0 | 0 | 0 | 0 | 0 | CCTTTACACTGCACGACAGTGCGACAAGGGGACCCCGAGTGCGAGGGCATACAGTCCTCGCTTTTGTCGACCGTAAGGTGGTCGACGAATAAGAGCTGGGCAAGACCGGTGCCAGCCGCCGCGGTAATACCGGCAGCTCGAGTGATGACCGCTATTATTGGGCCTAAAGCGTCCGTAGCTGGCCGTGTAAGTCCATCGGGAAATCCATCCGCTCAACGGATGGGCGTCCGGTGGAAACTGCACGGCTTGGAGCCGGAAGACCCAAGGGGTACGTCCGGGGTAGGAGTGAAATCCCGTAATCCTGGACGGACCGCCGATGGCGAAAGCACCTTGGGAAGACGGACTCGACAGTGAGGGACGAAAGCTAGGGTCT |
| p__Thermoproteota_g__ PALSA-986_ASV_1457 | 0 | 0 | 0 | 0 | 0 | 0 | 0 | 0 | 1 | 0 | 0 | ACTTTGCAATACACGAAAGTGTGACAGGGCTATCCCGAGTGCCATCCGCTGAGGAAGGCTTTTACCCAGTATAGCAAGCTGGGAGAATAAGGAGAGGGCAAGACTGGTGTCAGCCGCCGCGGTAATACCAGCTCTCCGAGTGGTGTGGATGTTTATTGGGCCTAAAGCATCCGTAGCTGGCTGGGCAAGTCCCCTGTTAAATCCACTGATTTAATCATTGGTGTGCGGGGGATACTGCTCGGCTAGGGGACGAGAGAGGCAGACGGTATTTCCGGGGTAGGGGTGAAATCCTATAATCCCGGGAAGACCACCAGTGGCGAAGGCTGTCTGCTAGAACGCGCCCGACGGTGAGGGATGAAAGCTGGGGGAG |
| p__Halobacteriota_g__Saliphagus_ASV_1444 | 0 | 1 | 1 | 0 | 1 | 0 | 0 | 0 | 0 | 0 | 0 | CCTTTACACTGCACGACAGTGCGATAAGGGGACCCCGAGTGCGAGGGCATACAGTCCTCGCTTTTCTGTACCGTAAGGAGGTACAGGAATAAGGGCTGGGCAAGACCGGTGCCAGCCGCCGCGGTAATACCGGCAGCTCAAGTGATGGCCGCTATTATTGGGCCTAAAGCGTCCGTAGCCGGCCGCACAAGTCCGTCGGGAAATCCACCCGCCTAACGGGTGGGCGTCCGGCGGAAACTGTGCGGCTTGGGACCGGAAGACTCGAGGGGTACGTCCGGGGTAGGAGTGAAATCCTGTAATCCTGGACGGACCGCCGGTGGCGAAAGCGCCTCGAGAGGACGGATCCGACGGTGAGGGACGAAAGCTAGGGTCT |
| p__Thermoplasmatota_g__SM1-50_ASV_1434 | 0 | 0 | 0 | 0 | 0 | 0 | 0 | 0 | 0 | 1 | 1 | CCTTCACACTGTGCGAAAGCGCGATGAGGGAATCCCAAGTGCTTGCACGTAAGTGTAAGCTGTTTCTTTGTTTAAAACGCAGAGAGAGGAAGGGCTGGGTAAGACGGGTGCCAGCCGCCGCGGTAATACCTGCAGCTCGAGTGGTGACCGCTATTATTGGGCCTAAAACGTCCGTAGCCTGCTGAGTAAATGCTTGGGTAAATCGTGCCGCTTAACGGTACGAATTCCGGGTACACTGCTTGGCTTGAGATCGGGAGAAGCTAGAGGTACTCCTGGGGTAGAGGTGAAATTCTGTAATCCTGGGGGGACCACCAGTGGCGAAGGCGTCTAGCTAGAACGAGTCTGACGGTGAGGGACGAAACCCTGGGGCG |
| p__Micrarchaeota_g__ UBA95_ASV_1458 | 0 | 0 | 0 | 0 | 0 | 0 | 0 | 0 | 1 | 0 | 0 | CTTCCGCAATGCGCGAAAGCGTGACGGGGGGAGTCCGAGTGCCATCCTAACGGATGGCTTTTCGTGAGACCAAATCTCTCACGGAATAAGTGGTGGGTAAATCTCGTGCCAGCCGCCGCGGTAATACGAGTAGCACAAGTGGTACCCGCGAATATTGAGCTTAAAGCGTTCGTAGCCGGCCAAGCCAGTTTCCTGTGAAATCTACCCGCTTAACGGGTAGGCGTGCAGGGAATACTACTTGGCTAGGGAACGGGGGAGGTCAGGAGTACTCTAGGGGGAGCGGTAAAATGCTGTAATCCTTGGAGGACTAACGATGGCGAAGGCACCTGACTAAAACGTGTCCGACGGTGAGGGACGAAGGCTAGGGGAG |
| p__Halobacteriota_f__Haloarculaceae_3992_ASV_1442 | 0 | 1 | 1 | 0 | 0 | 0 | 0 | 0 | 0 | 0 | 0 | CCTTTACACTGCACGCCAGTGCGATAAGGGGACTCCGAGTGCGAGGGCATATCGTCCTCGCTTTTCTGTACCGTAAGGTGGTACAGGAACAAGGGCTGGGCAAGACCGGTGCCAGCCGCCGCGGTAATACCGGCAGCCCGAGTAATGGCCGCTCTTATTGGGCCTAAAGCGTCCGTAGCCGGCCGGACAAGTCCGTTGGGAAATCCACGCGCTCAACGCGTGGGCGTCCAGCGGAAACTGTCCGGCTTGGGGCCGGAAGACCTGAGGGGTACGTCCGGGGTAGGAGTGAAATCCCGTAATCCTGGACGGACCACCGGTGGCGAAAGCGCCTCAGGAAGACGGACCCGACGGTGAGGGACGAAAGCTAGGGTCT |

| Phylum_Genus_ASV_number | L1 | L2 | L3 | L4 | L5 | L6 | L7 | L8 | L9 | L10 | L11 | Sequence |
|---|---|---|---|---|---|---|---|---|---|---|---|---|
| | Presence (1) or Absence (0) in each layer | | | | | | | | | | | |
| p__Halobacteriota_f__Natrialbaceae_ASV_1438 | 0 | 1 | 0 | 1 | 0 | 0 | 0 | 0 | 0 | 0 | 0 | CCTTTACACTGCGCGACAGCGCGATAAGGGGACTCCAAGTGCGAGGGCATATCGTCCTCGCTTTTCACTATCGTAAGGAGATGGTAGAATAAGGGCTGGGCAAGACCGGTGCCAGCCGCCGCGGTAATACCGGCAGCCCGAGTGATGACCGCTATTATTGGGCCTAAAGCGTCCGTAGCCGGCCGAACAAGTCCGTCGGGAAATCCGCACGCCTAACGTGCGGGCGTCCGGCGGAAACTGTTCGGCTTGGGACCGGAAGATCCAGAGGGTACGTCTGGGGTAGGAGTGAAATCCCGTAATCCTGGACGGACCACCGGTGGCGAAAGCGCTCTGGAAGGACGGATCCGACGGTGAGGGACGAAAGCTAGGGTCA |
| p__Halobacteriota_g__Halobaculum_ASV_1463 | 0 | 1 | 0 | 0 | 0 | 0 | 0 | 0 | 0 | 0 | 0 | CCTTTACACTGCACGCCAGTGCGATAAGGGGACTCCGAGTGCGAGGGCATATCGTCCTCGCTTTTCTGAACCGTAGGGAGGTTCAGGAACAAGAGCTGGGCAAGACCGGTGCCAGCCGCCGCGGTAATACCGGCAGCTCGAGTGATGGCCGATCTTATTGGGCCTAAAGCGTCCGTAGCCGGCCAGACAAGTCCGTCGGGAAATCCACCAGCTCAACTGGTGGGCGTCCGGCGGAAACTCTCTGGCTTGGGACCAGAAGACCCGAGAGGTACGTCCGGGGTAGGAGTGAAATCCCGTAATCCTGGACGGACCACCGATAGCGAAAGCATCTCGGGAGGATGGATCCGACGGTGAGGGACGAAAGCTGGGGTCT |
| p__Halobacteriota_g__Halobaculum_ASV_1466 | 0 | 1 | 1 | 1 | 0 | 1 | 0 | 0 | 0 | 0 | 0 | CCTTTACACTGCACGACAGTGCGATAGGGGGACTCCGAGTGCGGAGGCATAGTGCCTTCGCTTTTGTGCACCCTAAGGCGGTGCACGAATAAGAGCTGGGCAAGACCGGTGCCAGCCGCCGCGGTAATACCGGCAGCTCGAGTGATGGCCGATCTTATTGGGCCTAAAGCGTCCGTAGCCGGCCGGGCAAGTCCGTCGGGAAATCTACCAGCTCAACTGGTAGGCGTCCGGCGGAAACTGCTCGGCTTGGGACAGGAAGACCCGAGGGGTACGTCCGGGGTAGGAGTGAAATCCCGTAATCCTGGACGGACCACCGATGGCGAAAGCACCTCGGGAGGACTGATCCGACGGTCAGGGACGAAAGCTGGGGTCT |
| p__Halobacteriota_g__Halovenus_ASV_1473 | 0 | 0 | 1 | 0 | 0 | 0 | 0 | 0 | 0 | 0 | 0 | CCTTTACACTGCACGACAGTGCGATAGGGGGACTCCGAGTGCGAGGGCATAGAGCCCTCGCTTTTGTACACCGTAAGGTGGTGTACGAATAAGAACTGGGCAAGACCGGTGCCAGCCGCCGCGGTAATACCGGCAGTTCGAGTGATGGCCGATATTATTGGGCCTAAAGCGTTCGTAGCTGGCCGATCAAGTCTGTCGGGAAATCTGCTCGCCTAACGAGCAGGCGTCCGGCGGAAACTGGTCGGCTTGGGACCGGAAGACTCGAGGGGTACGTCCGGGGTAGGAGTGAAATCCCGTAATCCTGGACGGACCACCGATGGCGAAAGCACCTCGAGAGCACGGATCCGACAGTGAGGAACGAAAGCTGGGGTCT |
| p__Halobacteriota_g__Methanolobus_ASV_1461 | 0 | 0 | 0 | 0 | 0 | 0 | 0 | 0 | 1 | 1 | 0 | ACTTTACAATGCGGGAAACCGCGATAAGGGGACACTGAGTGCCAGCATACAATGTTGGCTGTCCGCATGTATAAATAGCATGTGTTAGCAAGGGCCGGGCAAGACCGGTGCCAGCCGCCGCGGTAACACCGGCGGCCCGAGTGGTGGCCACTATTATTGGGTCTAAAGGGTCCGTAGCCGGTTTGATCAGTCTTCCGGGAAATCTGACAGCTCAACTGTTAGGCTTCCGGGGGATACTGTCAGACTTGGGACCGGGAGAGGTAAGAGGTACTACAGGGGTAGGAGTGAAATCTTGTAATCCTTGTGGGACCACCAGTGGCGAAGGCGTCTTACCAGAACGGGTCCGACGGTGAGGGACGAAAGCTGGGGGCA |
| p__Halobacteriota_g__Saliphagus_ASV_1474 | 0 | 0 | 1 | 0 | 0 | 0 | 0 | 0 | 0 | 0 | 0 | CCTTTACACTGCACGACAGTGCGATAAGGGGACTCCAAGTGCGAGGGCATATCGTCCTCGCTTTTCACTACCGTAGGGAGGTAGTGGAATAAGGGCTGGGCAAGACCGGTGCCAGCCGCCGCGGTAATACCGGCAGCCCGAGTGATGACCGCTATTATTGGGCCTAAAGCGTCCGTAGCCGGCCACGCAAGTCCGTCGGGAAATCCGCACGCCTAACGTGCGGGCGTCCGGCGGAAACTGCGTGGCTTGGGACCGGAAGACTCGAGGGGTACGTCCGGGGTAGGAGTGAAATCCTGTAATCCTGGACGGACCACCGGTGGCGAAAGCGCCTCGAGAGGACGGATCCGACGGTGAGGGACGAAAGCTAGGGTCA |
| p__Asgardarchaeota_f__Thorarchaeaceae_ASV_1489 | 0 | 0 | 0 | 0 | 0 | 0 | 0 | 0 | 1 | 0 | 0 | CCTCACCAATGCGCGAAAGCGCGAGTGGGCCAAGCCGAGTGATGGGGGATAACTCCATCTGTGGCGGATCCGTAGGGTGGACCGCTAGAAAGGAGAGGGCAAGGCTGGTGCCAGCCGCCGCGGTAAAACCAGCTCTTCGAGTGGTGTCCATGATTATTGAGCTTAAAGCGTTCGTAGCCTGTTTAGTAGGTCCCTGCTTAAATCCGGTCGCTCAACGGTCGGCCTGGTAGGGATACCGCTTTACTTGGGGACGGGAGACGATGACGGTATTCCGTAGGTAAGGGTGAAATCTAATGATCTACGGAGGACCACCAGTGGCGAAGGCGGTCATCGAGAACGTGCCCGACGGTGAGGGACGAAACCCAGGGGAG |
| p__Halobacteriota_o__Halanaerobiales_ASV_1465 | 0 | 1 | 1 | 0 | 0 | 0 | 0 | 0 | 0 | 0 | 0 | CCTTTACACTGCACGCAAGTGCGATAGGGGGACCCCAAGTGCGAGGGCATACAGTCCTCGCTTTTCACAGCCGTAAGGAGGCTGTGGAATAAGGGCTGGGCAAGACCGGTGCCAGCCGCCGCGGTAATACCGGCAGCCCAAGTGATGGCCGCTGTTATTGGGCCTAAAGCGTCCGTAGCCGGCCGCGTGTGTCCATCGGGAAATCCACCCGCTCAACGGGTGGGCGGCCGATGGAAACTGCGCGGCTTGGGGCCGGAAGACCCACCGGGTACGTCCGGGGTAGGAGTGAAATCCCGTAATCCCGGACGGACCACCGATGGCGAAAGCACGGTGGGAAGACGGACCCGACGGTGAGGGACGAAAGCTGGGGTCT |
| d__Archaea_d__Archaea_ASV_1490 | 0 | 0 | 0 | 0 | 0 | 0 | 0 | 0 | 1 | 0 | 0 | ACTCTGCAATGACCGTAAGGTTGACAGGGGGACTCCAAGTGCTTATACACAGTATAGGCTTTTGCCACGTGTAAATAGCTTGGCGAATAAGGGGTGGGCAAGACCGGTGGCAGCAGCCGCGGTAATACCGGCGCCCCGAGTGGTGATCGTTTTTATTGGGCCTAAAGCGTTCGTAGCCGGTTTAGTAAGCTCTTGGTGAAATCCTTGCACTTAATGTAAGGGCGTGCCAGGAGAACTACTTAACTTGGGACCAGGAGAGGGCAAGAGTATTCCTAGGGTAGGGGTAAAATCCTATAATCCTAGGAGGACTACCTGTGGCGAAGGCGCTTGCCTAGAATGGGTCCGACGGTGAGGGACGAAAGCTGAGGGAT |
| p__Halobacteriota_g__Haloferax_ASV_1499 | 0 | 0 | 0 | 0 | 0 | 0 | 0 | 0 | 1 | 1 | 1 | CCTTTACACTGTACGCAAGTGCGATAAGGGGACCCCAAGTGCGAGGGCATATAGTCCTCGCTTTTCCCGACTGTAAGGCGGTCGGGGAATAAGGGCTGGGCAAGACCGGTGCCAGCCGCCGCGGTAATACCGGCAGCCCAAGTGATGACCGATATTATTGGGCCTAAAGCGTCCGTAGCCGGCCACGAAGGTCCATCGGGAAATCCGCCAGCTTAACTGGCGGGCGTCCGGTGGAAACCACGTGGCTTGGGACCGGAAGGCTCGAGGGGTACGTCCGGGGTAGGAGTGAAATCCCGTAATCCTGGACGGACCACCGATGGCGAAAGCACCTCGAGAAGACGGATCCGACGGTGAGGGACGAAAGCTAGGGTCT |
| p__Halobacteriota_g__Halohasta_ASV_1501 | 0 | 0 | 0 | 0 | 0 | 0 | 0 | 0 | 0 | 0 | 1 | CCTTTACACTGCACGCAAGTGCGATAGGGGAATCCCAAGTGCGAGGGCATACTAGCCCTCGCTTTTGTGTACTGTAAGGAGATACACGAATAAGAGCTGGGCAAGACCGGTGCCAGCCGCCGCGGTAATACCGGCAGCTCGAGTGATGTCCAATATTATTGGGCCTAAAGCGTCCGTAGCTGGCCAGCCAAGTCCGTCGGGAAATCCACCTGCCCAACAGGTGGGCGTCCGGCGGAAACTGTCTGGCTTGGAACCGGAAGGCTCAGAGGGTACGTCCGGGGTAGGAGTGAAATCCCGTAATCCCGGACGGACCACCGATGGCGAAAGCACTCTGAGAAGACGGATTCGACAGTGAGGGACGAAAGCTAGGGTCT |
| p__Halobacteriota_g__Halorussus_ASV_1506 | 0 | 1 | 0 | 0 | 0 | 0 | 0 | 0 | 0 | 0 | 0 | ACTTTACACTGCACGACAGTGCGATAAGGGGACTCCGAGTGCGAGGGCATATAGTCCTCGCTTTTGTACACCGTAGGGAGGTGTACGAATAAGGGCTGGGCAAGACCGGTGCCAGCCGCCGCGGTAATACCGGCAGCCCGAGTGATGGCCGCTATTATTGGGCCTAAAGCGTCCGTAGCCGGCCAGACAGGTCCGTCGGGAAATCCATCCGCTTAACGGATGGGCGTCCGGCGGAAACCAGCTGGCTTGGGGCCGGAAGACCTGAGGGGTACGTCCGGGGTAGGAGTGAAATCCCGTAATCCTGGACGGACCACCGGTGGCGAAAGCGCCTCAGGAAGACGGACCCGACGGTGAGGGACGAAAGCTAGGGTCA |
| p__Halobacteriota_g__Natrinema_ASV_1518 | 0 | 0 | 0 | 1 | 0 | 0 | 0 | 0 | 0 | 0 | 0 | CCTTTACACTGCACGCGAGTGCGATAAGGGGACTCCAAGTGCGAGGGCATATAGTCCTCGCTTTTCACCACCGTAAGGTGGTGGTGGAATAAGTGCTGGGCAAGACCGGTGCCAGCCGCCGCGGTAATACCGGCAGCACGAGTGATGACCGCTATTATTGGGCCTAAAGCGTCCGTAGCTGGCCATCCAAGTTCATCGGGAAATCCGCGCGCTTAACGCGCGGGCGTCCGGTGGAAACTGGATGGCTTGGGACCGGAAGACCAGAGGGGTACGTCCGGGGTAGGAGTGAAATCCCGTAATCCTGGACGGACCACCGGTGGCGAAAGCGCCTCTGGAAGACGGATCCGACGGTGAGGGACGAAAGCTCGGGTCA |
| p__Halobacteriota_g__Natronomonas_ASV_1509 | 0 | 1 | 0 | 0 | 0 | 0 | 0 | 0 | 0 | 0 | 0 | CCTTTACACTGCACGCCAGTGCGATAGGGGGACACCGAGTGCGAGGGCATAGAGCCCTCGCTTTTGTAGACCGTAAGGAGGTCTACGAATAAGGGCTGGGCAAGACCGGTGCCAGCCGCCGCGGTAACACCGGCAGCCCAAGTGATGGCCGATATTATTGGGCCTAAAGCGTCCGTAGCTGGCCGTTCAAGTTCGTCGGGAAATCCACTCGCTTAACGAGTGGGCGTCCGGCGAAAACTGTTCGGCTTGGGACCGGAAGATCCAAGGGGTACGTCCGGGGTAGGAGTGAAATCCTGTAATCCTGGACGGACCACCGATGGCGAAAGCACCTTGGAAGGACGGATCCGACAGTGAGGGACGAAAGCTAGGGTCT |
| p__Halobacteriota_g__Saliphagus_ASV_1513 | 0 | 0 | 1 | 1 | 0 | 0 | 0 | 0 | 0 | 0 | 0 | CCTTTACACTGCACGATAGTGCGATAGGGGGACTCCAAGTGCGAGGGCATATCGTCCTCGCTTTTGTGTACCGTAAGGTGGTACACGAATAAGGGCTGGGCAAGACCGGTGCCAGCCGCCGCGGTAATACCGGCAGCCCAAGTGATGACCGCTATTATTGGGCCTAAAGCGTCCGTAGCTGGCCGATCAAGTCCGTCGGGAAATCCGCACGCCTAACGTGCGGGCGTCCGGTGGAAACTGTTCGGCTTGGGACCGGAAGACTCGAGGGGTACGTCTGGGGTAGGAGTGAAATCCTGTAATCCTGGACGGACCACCGGTGGCGAAAGCGCCTCGAGAGGACGGATCCGACGGTGAGGGACGAAAGCTAGGGTCA |
| p__Halobacteriota_f__Natrialbaceae_ASV_1512 | 0 | 0 | 1 | 0 | 0 | 0 | 0 | 0 | 0 | 0 | 0 | CCTTTACACTGCACGCCAGTGCGATAAGGGGACTCCAGGTGCGAGGGCATATAGTCCTCGCTTTTCACCACCGTAAGGTGGTGGTGGAATAAGGGCTGGGCAAGACCGGTGCCAGCCGCCGCGGTAATACCGGCAGCCCGAGTGATGACCGCTATTATTGGGCCTAAAGCGTCCGTAGCTGGCCACACAAGTCCGTCGGGAAATCTCCTCGCCTAACGAGGAGGCGTCCGGTGGAAACTGTGTGGCTTGGGACCGGAAGATCCAGAGGGTACGTCCGGGGTAGGAGTGAAATCCCGTAATCCTGGACGGACCACCGGTGGCGAAAGCGCTCTGGAAGGACGGATCCGACGGTGAGGGACGAAAGCTCGGGTCA |
| p__Halobacteriota_f__Natrialbaceae_ASV_1514 | 0 | 0 | 1 | 0 | 0 | 0 | 0 | 0 | 1 | 0 | 0 | CCTTTACACTGCACGCCAGTGCGATAAGGGGACTCCAGGTGCGAGGGCATATAGTCCTCGCTTTTCTGTACCGTAAGGTGGTACAGGAATAAGGGCTGGGCAAGACCGGTGCCAGCCGCCGCGGTAATACCGGCAGCCCAAGTGATGACCGCTATTATTGGGCCTAAAGCGTCCGTAGCTGGCCACACAAGTCCGTCGGGAAATCTCCTCGCCTAACGGGGAGGCGTCCGGTGGAAACTGTGTGGCTTGGGACCGGAAGATCCAGAGGGTACGTCCGGGGTAGGAGTGAAATCCCGTAATCCTGGACGGACCACCGGTGGCGAAAGCGCTCTGGAAAGACGGATCCGACGGTGAGGGACGAAAGCTCGGGTCA |
| p__Halobacteriota_o__Halanaerobiales_ASV_1503 | 0 | 0 | 1 | 0 | 0 | 0 | 0 | 0 | 0 | 0 | 1 | CCTTTACACTGCACGCAAGTGCGATAAGGGGACCCCGAGTGCGAGGGCATACCGTCCTCGCTTTTGTCGACCGTAGGGAGGTCGACGAATAAGGGCTGGGCAAGACCGGTGCCAGCCGCCGCGGTAATACCGGCAGCCCAAGTGATGGCCGCTCTTATTGGGCCTAAAGCGTCCGTAGCTGGCCGCATTGGTCCGTCGGGAAATCCACTCGCTCAACGAGTGGGCGTCCGGCGGATACCATGCGGCTTGGGACCGGAAGACCCAGGGGGTACGTCCGGGGTAGGAGTGAAATCCTGTAATCCCGGACGGACCGCCGATGGCGAAAGCACTCTGGGAAGACGGATCCGACAGTGAGGGACGAAAGCTGGGGTCT |
| p__Halobacteriota_g__Halorussus_ASV_1546 | 0 | 1 | 1 | 0 | 0 | 1 | 1 | 0 | 0 | 0 | 0 | CCTTTACACTGCACGACAGTGCGATAAGGGGATCCCGAGTGCGAGGGCATACAGTCCTCGCTTTTCTGCACCGTAAGGTGGTGCAGGAATAAGAGCTGGGCAAGACCGGTGCCAGCCGCCGCGGTAATACCGGCAGCTCGAGTGATGGCCGCTATTATTGGGCCTAAAGCGTCCGTAGCCGGCCGAACAGGTCCGTCGGGAAATCTGCTCGCCTAACGAGCAGGCGTCCGGCGGAAACCAGTCGGCTTGGGGCCGGAGGACCTGAGGGGTACGTCCGGGGTAGGAGTGAAATCCCGTAATCCCGGACGGACCGCCGGTGGCGAAAGCGCCTCAGGAGGACGGACCCGACGGTGAGGGACGAAAGCTAGGGTCA |
| p__Halobacteriota_g__Halovenus_ASV_1543 | 0 | 1 | 1 | 0 | 0 | 0 | 0 | 0 | 0 | 0 | 0 | CCTTTACACTGCACGACAGTGCGATAGGGGGACTCCGAGTGCGAGGGCATATAGTCCTCGCTTTTCTGTACCGTAAGGTGGTACAGGAACAAGTGCTGGGCAAGACCGGTGCCAGCCGCCGCGGTAATACCGGCAGCACAAGTGATGGCCGATATTATTGGGCCTAAAGCGTTCGTAGCCGGCCGGACAAGTTCGTTGGGAAATCCACGCGCTCAACGCGTGGGCGTCCAGCGAATACTGTCTGGCTTGGGGCCGGAAGACTCAAGGGGTACGTCCGGGGTAGGAGTGAAATCCCGTAATCCTGGACGGACCGCCGATGGCGAAAGCACCTTGAGAGGACGGACCCGACGGTGAGGGACGAAAGCCAGGGTCT |
| p__Halobacteriota_g__Halovivax_ASV_1545 | 0 | 1 | 1 | 0 | 0 | 0 | 0 | 0 | 1 | 0 | 0 | CCTTTACACTGCACGCCAGTGCGATAAGGGGACTCCAAGTGCGAGGGCATATAGTCCTCGCTTTTTCCCACCGTAAGGTGGTGGGCGAATAAGGGCTGGGCAAGACCGGTGCCAGCCGCCGCGGTAATACCGGCAGCCCGAGTGATGACCGCTATTATTGGGCCTAAAGCGTCCGTAGCTGGCCGGGCAAGTCCGTCGGGAAATCCGCACGCCTAACGTGCGGGCGTCCGGTGGAAACTGCTCGGCTTGGGACCGGAAGATCCAGAGGGTACGTCTGGGGTAGGAGTGAAATCCCGTAATCCTGGACGGACCGCCGGTGGCGAAAGCGCTCTGGAAGGACGGATCCGACAGTGAGGGACGAAAGCTAGGGTCA |
| p__Thermoproteota_g__PALSA-986_ASV_1541 | 0 | 0 | 0 | 0 | 0 | 0 | 0 | 0 | 1 | 0 | 1 | ACTTTGCAATACACGCAAGTGTGACAGGGTCATCCCGAGTGCCGACCGCTGAGGTTGGCTTTTACCCAGTCTAGAAAGCTGGGGGAATAAGGAGAGGGCAAGTCTGGTGTCAGCCGCCGCGGTAATACCAGCTCTCCGAGTGGTGTGGACGTTTATTGGGCCTAAAGCATCCGTAGCTGGCTGAACAAGTCCCCTGTTAAACCCACCGATTTAATCGTTGGCTTGCGGGGGATACTGCTCGGCTAGGGGACGAGAGAGGCAGACGGTATTCTAGGGGTAGGGGTGAAATCCTATAATCCCTGGAAGACCACCAGTGGCGAAGGCTGTCTGCTAGAACGCGCCCGACGGTGAGGGATGAAAGCTGGGGGAG |
| p__Halobacteriota_g__Saliphagus_ASV_1542 | 0 | 1 | 0 | 0 | 0 | 0 | 0 | 0 | 0 | 0 | 0 | CCTTTACACTGCACGACAGTGCGATAGGGGGACTCCAAGTGCGAGGGCATATCGTCCTCGCTTTTGTGTACCGTAAGGTGGTACACGAATAAGGGCTGGGCAAGACCGGTGCCAGCCGCCGCGGTAATACCGGCAGCCCGAGTGATGACCGCTATTATTGGGCCTAAAGCGTCCGTAGCTGGCCGATCAAGTCCGTCGGGAAATCCGCACGCCTAACGTGCGGGCGTCCGGTGGAAACTGTTCGGCTTGGGACCGGAAGACTCGAGGGGTACGTCTGGGGTAGGAGTGAAATCCTGTAATCCTGGACGGACCACCGGTGGCGAAAGCGCCTCGAGAGGACGGATCCGACGGTGAGGGACGAAAGCTAGGGTCA |
| p__Halobacteriota_f__Haloarculaceae_3992_ASV_1547 | 0 | 0 | 1 | 0 | 0 | 0 | 0 | 0 | 0 | 0 | 0 | CCTTTACACTGCACGACAGTGCGATAGGGGGACTCCGAGTGCGAGGGCATATAGCCCTCGCTTTTCTGTACCGTAAGGTGGTACAGGAACAAGTGCTGGGCAAGACCGGTGCCAGCCGCCGCGGTAATACCGGCAGCACGAGTGATGGCCGATATTATTGGGCCTAAAGCGTCCGTAGCTTGCTGTGTTAGTCCGTTGGGAAATCGACGCGCTCAACGCGTCGGCGTCCAGCGGAAACTGCACGGCTTGGGGCCGAGAGACTCGACGGGTACGTCCGGGGTAGGAGTGAAATCCTGTAATCCTGGACGGACCACCAATGGGGAAACCACGTCGAGAGACCGGACCCGACAGTGAGGGACGAAAGCCAGGGTCT |

| Phylum_Genus_ASV_ number | Presence (1) or Absence (0) in each layer | | | | | | | | | | | |
| --- | --- | --- | --- | --- | --- | --- | --- | --- | --- | --- | --- | --- |
| | L1 | L2 | L3 | L4 | L5 | L6 | L7 | L8 | L9 | L10 | L11 | Sequence |
| p__Halobacteriota_f__Natrialbaceae_ASV_1548 | 0 | 0 | 1 | 0 | 0 | 0 | 0 | 0 | 0 | 0 | 0 | CCTTTACACTGCACGACAGTGCGATAAGGGGATTCCGAGTGCGAGGGCATACAGTCCTCGCTTTTGTGTACCGTAGGAAGGTACACGAATAAGGGCTGGGCAAGACCGGTGCCAGCCGCCGCGGTAATACCGGCAGCCCAAGTGATGACCGATATTATTGGGCCTAAAGCGTCCGTAGCCGGCTACGCAAGTCCGTCGGGAAATCTGCTCGCTCAACGAGTAGGCGTCCGGCGGAAACTGCGTGGCTTGGGACCGGAAGACTCGAGGGGTACGTCCGGGGTAGGAGTGAAATCCCGTAATCCTGGACGGACCACCGGTGGCGAAAGCGCCTCGAGAGGACGGATCCGACGGTGAGGGACGAAAGCTAGGGTCA |
| d__Archaea_d__Archaea_ASV_1556 | 0 | 0 | 0 | 0 | 0 | 0 | 0 | 0 | 1 | 0 | 0 | CCTCTACAATGCTGGAAACAGTGATAGGGGGACTCCGAGTGGCGTCGGCTAAGCTGGCGTCTTTTGCCAAGTGTAAATAGCTTGGAGAATAAGGGTGTGGGCAAGACTGGTGGCAGCCGCCGCGGTAACACCAGCGCCTCAAGTGGTACCCACGAATACTGGGCCTAGAGCGTTCGTAGCCGGCCAAGTAAGTCTTCTGTGAAATAGTGAGGCTTAACCTCACTGCGCGCAGGAGATACTGCCTGGCTTGGGAACGGGGGAGGTCAGGAGTACACCCGGGGTAGGGGTAAAATCCGTTGAGCCTGGGTGGACTACCAGTGGCGAAGGCGCCTGACCAAAACGTGTCCGACGGTGAGGAACGAAAGCTGGGGGAG |
| p__Halobacteriota_g__Halobellus_3029_ASV_1570 | 0 | 1 | 0 | 0 | 0 | 0 | 0 | 0 | 0 | 0 | 1 | CCTTTACACTGCACGCCAGTGCGATAAGGGGACTCCAAGTGCGAGGGCATATCGTCCTCGCTATCCTGTACTGTAGGGTAGTACAGTATCAAGAGCTGGGCAAGACCGGTGCCAGCCGCCGCGGTAATACCGGCAGCTCAAGTGATGGCCGATCTTATTGGGCCTAAAGCGTCCGTAGCTGGCCCTGCAAGTCCGTCGGGAAATCCATCAGCTCAACTGATGGGCGTCCGGCGGAAACTCCAGGGCTTGGGACCGGAAGGCTCGAGGGGTACGTCCGGGGTAGGAGTGAAATCCCGTAATCCCGAACGGACCACCGATGGCGAAAGCACCTCGAGAAGACGGCTCCGACAGTGAGGGACGAAAGCTAGGGTCT |
| p__Halobacteriota_g__Halohasta_ASV_1575 | 0 | 1 | 1 | 0 | 0 | 0 | 0 | 0 | 0 | 0 | 0 | CCTTTACACTGCACGCAGGTGCGATAGGGGAATCCCAAGTGCGAGGGCATACTAGCCCTCGCTTTTGTGTACTGTAAGGAGGTACACGAATAAGAGCTGGGCAAGACCGGTGCCAGCCGCCGCGGTAATACCGGCAGCTCGAGTGATGTCCACTATTATTGGGCCTAAAGCGTCCGTAGCTGGCCGTACAAGTTCGTCGGGAAATCCACCTGCCCAACAGGTGGGCGTCCGGCGAAAACTGTGCGGCTTGGAACCGGAAGGCTCAGAGGGTACGTCCGGGGTAGGAGTGAAATCCCGTAATCCCGGACGGACCACCGATGGCGAAAGCACTCTGAGAGGACGGATTCGACAGTGAGGGACGAAAGCTAGGGTCT |
| p__Halobacteriota_g__Halovivax_ASV_1579 | 0 | 1 | 1 | 1 | 0 | 0 | 0 | 0 | 0 | 0 | 0 | CCTTTACACTGCACGCGAGTGCGATAGGGGGACTCCAAGTGCGAGGGCATACAGTCCTCGCTTTTTCCTACCGTAAGGTGGTAGGCGAATAAGTGCTGGGCAAGACCGGTGCCAGCCGCCGCGGTAATACCGGCAGCACGAGTGATGACCGCTATTATTGGGCCTAAAGCGTCCGTAGCCGGCCGGGCAAGTCCATCGGGAAATCCGCACGCCTAACGTGCGGGCGTCCGGTGGAAACTGTTCGGCTTGGGACCGGAAGATCCAGAGGGTACGTCTGGGGTAGGAGTGAAATCCTGTAATCCTGGACGGACCACCGGTGGCGAAAGCGCTCTGGAAAGACGGATCCGACGGTGAGGGACGAAAGCTTGGGTCA |
| p__Halobacteriota_g__Halovivax_ASV_1587 | 0 | 0 | 0 | 1 | 0 | 0 | 0 | 0 | 0 | 0 | 0 | CCTTTACACTGCACGCCAGTGCGATAGGGGGACTCCAAGTGCGAGGGCATACAGTCCTCGCTTTTCACGACCGTAAGGTGGTCGTGGAATAAGAGCTGGGCAAGACCGGTGCCAGCCGCCGCGGTAATACCGGCAGCTCGAGTGATGACCGCTTTTATTGGGCCTAAAGCGTCCGTAGCCGGCCGGGTAAGTCCATCGGGAAATCCGCACGCTTAACGTGCGGGCGTCCGGTGGAAACTGCCTGGCTTGGGACCGGAAGATCCAGAGGGTACGTCTGGGGTAGGAGTGAAATCCCGTAATCCTGGACGGACCACCGGTGGCGAAAGCGCTCTGGAAAGACGGATCCGACGGTGAGGGACGAAAGCTCGGGTCA |
| p__Halobacteriota_g__Saliphagus_ASV_1581 | 0 | 0 | 1 | 0 | 0 | 0 | 0 | 0 | 0 | 0 | 0 | CCTTTACACTGCACGACAGTGCGATAGGGGGACCCCGAGTGCGAGGGCATACAGTCCTCGCTTTTGTGTACCGTAAGGAGGTACACGAATAAGAGCTGGGCAAGACCGGTGCCAGCCGCCGCGGTAATACCGGCAGCTCAAGTGATGGCCGCTGTTATTGGGCCTAAAGCGTCCGTAGCCGGCCGCACAAGTCCGTCGGGAAATCCACCCGCCTAACGGGTGGGCGTCCGGCGGAAACTGTGCGGCTTGGGACCGGAGGACTCGAGGGGTACGTCCGGGGTAGGAGTGAAATCCCGTAATCCTGGACGGACCGCCGGTGGCGAAAGCGCCTCGAGAGGACGGATCCGACGGTGAGGGACGAAAGCTAGGGTCT |
| p__Halobacteriota_g__Saliphagus_ASV_1583 | 0 | 0 | 1 | 1 | 0 | 0 | 0 | 0 | 1 | 0 | 0 | CCTTTACACTGCACGACAGTGCGATAGGGGGACTCCAAGTGCGAGGGCATATAGTCCTCGCTTTTCACCACCGTAAGGTGGTGGTGGAATAAGTGCTGGGCAAGACCGGTGCCAGCCGCCGCGGTAATACCGGCAGCACGAGTGATGACCGCTATTATTGGGCCTAAAGCGTCCGTAGCCGGCCGAACAAGTCCGTCGGGAAATCCGCACGCCTAACGTGCGGGCGTCCGGCGGAAACTGTTCGGCTTGGGACCGGAAGATCCAGAGGGTACGTCTGGGGTAGGAGTGAAATCCCGTAATCCTGGACGGACCACCGGTGGCGAAAGCGCTCTGGAAGGACGGATCCGACGGTGAGGGACGAAAGCTAGGGTCA |
| p__Halobacteriota_f__Haloarculaceae_3992_ASV_1578 | 0 | 1 | 1 | 0 | 0 | 0 | 0 | 0 | 0 | 0 | 0 | CCTTTACACTGCACGCCAGTGCGATAAGGGGACTCCGAGTGCGAGGGCATATCGTCCTCGCTTTTCTGTACCGTAAGGTGGTACAGGAACAAGTGCTGGGCAAGACCGGTGCCAGCCGCCGCGGTAATACCGGCAGCACGAGTGATGGCCAATCTTATTGGGCCTAAAGCGTCCGTAGCCGGCCGGACAAGTCCGTTGGGAAATCCACGCGCTCAACGCGTGGGCGTCCAGCGGAAACTGTCCGGCTTGGGGCCGGAAGACCTGAGGGGTACGTCCGGGGTAGGAGTGAAATCCCGTAATCCTGGACGGACCACCGGTGGCGAAAGCGCCTCAGGAAGACGGACCCGACGGTGAGGGACGAAAGCTAGGGTCT |
| p__Halobacteriota_f__Natrialbaceae_ASV_1574 | 0 | 1 | 1 | 0 | 0 | 0 | 0 | 0 | 0 | 0 | 0 | CCTTTACACTGCACGACAGTGCGATAGGGGGACTCCAAGTGCGAGGGCATATCGTCCTCGCTTTTCACTACCGTAAGGTGGTAGTAGAATAAGGGCTGGGCAAGACCGGTGCCAGCCGCCGCGGTAATACCGGCAGCCCAAGTGATGACCGCTATTATTGGGCCTAAAGCGTCCGTAGCCGGCCTCGCAAGTCCGTCGGGAAATCCGCGCGCTCAACGCGCGGGCGTCCGGCGGAAACTGCGTGGCTTGGGACCGGAAGACTCAGGGGGTACGTCCGGGGTAGGAGTGAAATCCTGTAATCCTGGACGGACCACCGGTGGCGAAAGCGCCCTGAGAGGACGGATCCGACGGTGAGGGACGAAAGCTAGGGTCA |
| p__Halobacteriota_o__Halanaerobiales_ASV_1571 | 0 | 1 | 0 | 0 | 0 | 0 | 0 | 0 | 0 | 0 | 0 | CCTTTACACTGCACGACAGTGCGATAGGGGGACTCCGAGTGCGAGGGCATATAGTCCTCGCTTTTCTGGACCGTAAGGAGGTCCAGGAACAAGTGCTGGGCAAGACCGGTGCCAGCCGCCGCGGTAATACCGGCAGCACGAGTGATGGCCGCTCTTATTGGGCCTAAAGCGTCCGTAGCCGGCCCGACAAGTCCGTTGGGAAATCCACGCGCTCAACGCGTGGGCGTCCAGCGGAAACTGTCGGGCTTGGGACCGGAAGACCCAGAGGGTACGTCCGGGGTAGGAGTGAAATCCCGTAATCCTGGACGGACCACCGGTGGCGAAAGCGCTCTGGGAAGACGGATCCGACGGTGAGGGACGAAAGCTAGGGTCT |
| p__Thermoproteota_g__BA2_ASV_1635 | 0 | 0 | 0 | 0 | 0 | 0 | 0 | 0 | 1 | 0 | 0 | ACTCCGCAATGCACGAAAGTGTGACGGGGCTACCCCGAGTGCTGTCCGCTGAGGATGGCTTTTCCTCGGTGTAACTAGCCGAGGGAATAAGGAGAGGGCAAGACTGGTGTCAGCCGCCGCGGTAATACCAGCTCTCCGAGTTGTAGGGATGATTATTGGGCTTAAAGCGTCCGTAGCCGGCTTAGCAAGTCTCCTGTTAAATTCAGCGACCTAATCGTTGAGCCGCGGGAGATACTACTAGGCTAGGAGGCGGGAGAGGTCGACGGTACTTTCGGGGTAGGGGTGAAATCCTATAATCCCGGGAGGACCACCAGTGGCGAAGGCTGTCGACTAGAACGCGCTCGACGGTGAGGGACGAAAGCTGGGGGAG |
| p__Nanoarchaeota_g__BMS3ABIN17_ASV_1596 | 1 | 1 | 0 | 0 | 0 | 1 | 1 | 0 | 0 | 0 | 0 | CTTTTACAATGCTCGAAAGAGTGATAGAGCAAGCCAGAGTGTCTCTCTTAGAGAGACTTTTGTCTGCTGTAAAAAGGCAGACGAATAAGGACTGGGTAAGACCGGTGCCAGCCGCCGCGGTAATACCGGCGGTCCAAGTCGCATTCACATTTATTGGGTCTAAAACATCCGTAGCTTGTTTTGTAAGTTTCTTGTGAAATCCTGTATCTCAAGTACAGGGCTGGCAAGAAATACTGCAAAACTAGAGACCGGAAGATGCAAGGAGTATGGTTAAGGTAGCGGTAAAATGTGTTAATCTTAACTAGACTAACAACGGCGAAGGCACCTTGCAAGTACGGATCTGACAGTGAGGGATGAAGGCTAGGGGCG |
| p__Halobacteriota_g__Halalkalicoccus_ASV_1595 | 1 | 1 | 1 | 0 | 1 | 0 | 0 | 0 | 0 | 0 | 0 | CCTTTACACTGCACGAACGTGCGATAAGGGGACTCCAAGTGCGAGGGCATATAGTCCTCGCTTTTGCCTACCGTAAGGTGGTAGGCGAATAAGAGCTGGGCAAGACCGGTGCCAGCCGCCGCGGTAATACCGGCAGCTCGAGTGATGGCCAATATTATTGGGCCTAAAGCGTCCGTAGCTGGCCGAACAGGTCCGTCGGGAAATCCGCTCGCTCAACGAGCGGACGTCCGGCGGAAACCAGTCGGCTTGGGACCGGAAGACCCGAGGGGTACGGCCGGGGTAGGAGTGAAATCCTGTAATCCTGGCCGGACCACCGGTGGCGAAAGCGCCTCGGGAAGACGGATCCGACAGTGAGGGACGAAAGCTAGGGTCA |
| p__Halobacteriota_g__Halohasta_ASV_1612 | 0 | 1 | 1 | 1 | 0 | 0 | 0 | 0 | 0 | 0 | 0 | CCTTTACACTGCACGGAAGTGCGATAGGGGAATCCCAAGTGCGAGGGCATACTAGCCCTCGCTTTTGTTGACTGTAAGGAGGTCAACGAATAAGAGCTGGGCAAGACCGGTGCCAGCCGCCGCGGTAATACCGGCAGCTCGAGTGATGTCCAATATTATTGGGCCTAAAGCGTCCGTAGCTGGCCAGCCAAGTTCGTCGGGAAATCCACCTGCCCAACAGGTGGGCGTCCGGCGAAAACTGTCTGGCTTGGAACCGGAAGGCTCAGAGGGTACGTCCGGGGTAGGAGTGAAATCCCGTAATCCCGGACGGACCACCGATGGCGAAAGCACTCTGAGAGGACGGATTCGACAGTGAGGGACGAAAGCTAGGGTCT |
| p__Halobacteriota_g__Halohasta_ASV_1620 | 0 | 0 | 1 | 0 | 0 | 0 | 0 | 0 | 0 | 0 | 0 | CCTTTACACTGCACGCAAGTGCGATAGGGGAATCCCAAGTGCGAGGGCATACTAGCCCTCGCTTTTGTTGACTGTAAGGAGGTCAACGAATAAGAGCTGGGCAAGACCGGTGCCAGCCGCCGCGGTAATACCGGCAGCTCGAGTGATGTCCAATATTATTGGGCCTAAAGCGTCCGTAGCTGGCCAGCCAAGTTCGTCGGGAAATCCACCTGCCCAACAGGTGGGCGTCCGGCGAAAACTGTCTGGCTTGGAACCGGAAGGCTCAGAGGGTACGTCCGGGGTAGGAGTGAAATCCCGTAATCCCGGACGGACCACCGATGGCGAAAGCACTCTGAGAAGACGGATTCGACAGTGAGGGACGAAAGCTAGGGTCT |
| p__Halobacteriota_g__Halolamina_ASV_1601 | 0 | 1 | 0 | 0 | 0 | 0 | 0 | 1 | 0 | 1 | 0 | CCTTTACACTGCACGCCAGTGCGATAAGGGGACCCCGAGTGAAAGGGCATATCGCCCTTTCTTTTCTGAACCGTAGGGAGGTTCAGGAACAAGAGCTGGGCAAGACCGGTGCCAGCCGCCGCGGTAATACCGGCAGCTCGAGTGATGACCGATTTTATTGGGCCTAAAGCGTCCGTAGCTGGCCTCGCAAGTCCATCGGGAAATCCACCCGCTTAACGGGTGGGCGTCCGGTGGAAACTGCGTGGCTTGGGACAGGAAGGCTCGAGGGGTACGTTCGGGGTAGGAGTGAAATCCTGTAATCCTGAACGGACCGCCGATGGCGAAAGCACCTCGAGAGGACTGATCCGACAGTGAGGGACGAAAGCTGGGGTCT |
| p__Halobacteriota_g__Halomicrobium_3849_ASV_1616 | 0 | 0 | 1 | 0 | 0 | 0 | 0 | 0 | 0 | 0 | 0 | CCTTTACACTGCACGACAGTGCGATAGGGGGACTCCGAGTGCGAGGGCATATAGTCCTCGCTTTTCTGGATCGTAAGGAGATCCAGGAACAAGTGCTGGGCAAGACCGGTGCCAGCCGCCGCGGTAATACCGGCAGCACGAGTGATGGCCGCTCTTATTGGGCCTAAAGCGTCCGTAGCCGGCCAGTCAAGTCCGTTGGGAAATCGGCGCGCCTAACGCGTCGGCGTCCAGCAGAAACTGACTGGCTTGGGGCCGGAAGACCTGAGGGGTACGTCCGGGGTAGGAGTGAAATCCTGTAATCCTGGACGGACCACCGGTGGCGAAAGCGCCTCAGGAAGACGGACCCGACGGTGAGGGACGAAAGCTAGGGTCT |
| p__Halobacteriota_g__Natrinema_ASV_1605 | 0 | 1 | 0 | 0 | 0 | 0 | 0 | 0 | 0 | 0 | 0 | CCTTTACACTGCACGCGAGTGCGATAAGGGGACTCCGAGTGCGAGGGCATATAGTCCTCGCTTTTCACCACCGTAAGGTGGTGGTGGAATAAGTGCTGGGCAAGACCGGTGCCAGCCGCCGCGGTAATACCGGCAGCACGAGTGATGACCGCTATTATTGGGCCTAAAGCGTCCGTAGCTGGCCACGCAAGTCTATCGGGAAATCTGCGCGCTTAACGCGCAGGCGTCCGGTGGAAACTGTGTGGCTTGGGACCGGAAGACCAGAGGGGTACGTCCGGGGTAGGAGTGAAATCCCGTAATCCTGGACGGACCACCGGTGGCGAAAGCGCCTCTGGAAGACGGATCCGACGGTGAGGGACGAAAGCTCGGGTCA |
| p__Halobacteriota_g__Natronorubrum_4312_ASV_1627 | 0 | 0 | 0 | 0 | 1 | 1 | 0 | 0 | 0 | 0 | 0 | CCTTTACACTGCACGCCAGTGCGATAAGGGGACTCCGAGTGCGAGGGCATATAGTCCTCGCTTTTCACCACCGTAAGGTGGTGGTGGAATAAGTGCTGGGCAAGACCGGTGCCAGCCGCCGCGGTAATACCGGCAGCACGAGTGATGACCGCTATTATTGGGCCTAAAGCGTCCGTAGCTGGCCACGCAAGTCTATCGGGAAATCTCTTCGCTTAACGAAGAGGCGTCCGGTGGAAACTGCGTGGCTTGGGACCGGAAGATCCAGAGGGTACGTCCGGGGTAGGAGTGAAATCCTGTAATCCTGGACGGACCGCCGGTGGCGAAAGCGCTCTGGAAAGACGGATCCGACGGTGAGGGACGAAAGCTCGGGTCA |
| p__Halobacteriota_f__Haloarculaceae_3992_ASV_1617 | 0 | 0 | 1 | 0 | 0 | 0 | 0 | 0 | 0 | 0 | 0 | CCTTTACACTGCACGACAGTGCGATAGGGGGACTCCGAGTGCGAGGGCATATAGTCCTCGCTTTTCTGTACCGTAAGGTGGTACAGGAACAAGTGCTGGGCAAGACCGGTGCCAGCCGCCGCGGTAATACCGGCAGCACGAGTGATGGCCGATCTTATTGGGCCTAAAGCGTCCGTAGCCGGCCGAGCAAGTCCGTTGGGAAATCCACGCGCTCAACGCGTGGGCGTCCAGCGGAAACTGTTCGGCTTGGGGCCGGAAGACCTGAGGGGTACGTCCGGGGTAGGAGTGAAATCCTGTAATCCTGGACGGACCACCGGTGGCGAAAGCGCCTCAGGAAGACGGACCCGACGGTGAGGGACGAAAGCTAGGGTCT |
| p__Halobacteriota_f__Haloferacaceae_ASV_1618 | 0 | 0 | 1 | 0 | 0 | 0 | 0 | 0 | 0 | 0 | 0 | CCTTTACACTGCACGCACGTGCGATAAGGGGACTCCGAGTGCGAGGGCATACAGTCCTCGCTTTTGAGAACCGTAAGGAGGTTCTCGAATAAGAGCTGGGCAAGACCGGTGCCAGCCGCCGCGGTAATACCGGCAGCTCGAGTGATGGCCGCTCTTATTGGGCCTAAAGCGTCCGTAGCCGGCCGAATTGGTCCGTCGGGAAATCCACTGGCTCAACCAGTGGGCGTCCGGCGGATACCGTTCGGCTTGGGACCGGAAGACTCGAGGGGTACGTCCGGGGTAGGAGTGAAATCCTGTAATCCTGGACGGACCACCGATGGCGAAAGCACCTCGAGAGGACGGATCCGACGGTGAGGGACGAAAGCTAGGGTCT |
| p__Halobacteriota_f__Natrialbaceae_ASV_1609 | 0 | 1 | 0 | 0 | 0 | 0 | 0 | 0 | 0 | 0 | 0 | ACTTTACACTGCACGCCAGTGCGATAAGGGAACTCCGAGTGCGAGGGCATATCGTCCTCGCTTTTCTGTACCGTAGGGAGGTACAGGAATAAGGGCTGGGCAAGACCGGTGCCAGCCGCCGCGGTAACACCGGCAGCCCGAGTGATGGCCGATATTATTGGGCCTAAAGCGTCCGTAGCCAGCCGAACAGGTCCGTCGGGAAATCTGCTCGCTCAACGAGCAGGCGTCCGGCGGAAACCAGTCGGCTTGGGATCGGAAGACCCGAGGGGTACGTCTGGGGTAGGAGTGAAATCCCGTAATCCTGGACGGACCACCGGTGGCGAAAGCGCCTCGGGAAGACGAATCCGACGGTGAGGGACGAAAGCTAGGGTCA |
| p__Halobacteriota_f__Natrialbaceae_ASV_1610 | 0 | 1 | 1 | 0 | 0 | 0 | 0 | 0 | 0 | 0 | 0 | CCTTTACACTGCGCGACAGCGCGATAAGGGGACTCCACGTGCCAGGGCATATCGTCCTGGCTTTTGTGTACCGTAAGGTGGTACACGAATAAGGGCTGGGCAAGACCGGTGCCAGCCGCCGCGGTAATACCGGCAGCCCGAGTGATGACCGCTCTTATTGGGCCTAAAGCGTCCGTAGCTGGCCATCCAAGTCCATCGGGAAATCCGCACGCCTAACGTGCGGGCGTCCGGTGGAAACTGGATGGCTTGGGACCGGAAGATCCAGAGGGTACGTCTGGGGTAGGAGTGAAATCCCGTAATCCTGGACGGACCACCGGTGGCGAAAGCGCTCTGGAAGGACGGATCCGACGGTGAGGGACGAAAGCTCGGGTCA |

| Phylum_Genus_ASV_number | L1 | L2 | L3 | L4 | L5 | L6 | L7 | L8 | L9 | L10 | L11 | Sequence |
|---|---|---|---|---|---|---|---|---|---|---|---|---|
| | Presence (1) or Absence (0) in each layer | | | | | | | | | | | |
| d__Archaea_d__Archaea_ASV_1597 | 1 | 0 | 0 | 0 | 0 | 0 | 1 | 1 | 0 | 0 | 1 | CCTTTACAATGCACGCAAGTGTGATAAGGGGATTCCAAGTGCTCATGCATAGCATGGGCTTTTTAGAAGAGTAAAAATCTTCTAGAATAAGTGGTGGGAAAGACTGGTGCCAGCAGCCGCGGTAACCCCAGCGCCACAAGTGGCAATCGCATTTATTGGGTCTAAAGCGTTCGTAGTTGGTTTGATCAATCTCTTGTGAAATCGTAGGGCCTAACTCTACGGCGCGCAGGAGATATTGTCAAACTAGGGACCGGGAGGAGTCAGAGGTATTCCTGAGGGAACGGTAAAATGTTATAATCTCAGGAAGACCACCTGTGGCAAAGGCGTCTGACTAGAACGGATCCGACGATGAGGAACGAAGGCCAGGGGAG |
| p__Halobacteriota_g__Halolamina_ASV_1662 | 0 | 0 | 1 | 0 | 1 | 0 | 0 | 0 | 0 | 0 | 0 | CCTTTACACTGCACGCCAGTGCGATAAGGGGACCCCGAGTGAAAGGGCATATCGCCCTTTCTTTTCTGAACCGTAGGGAGGTTCAGGAACAAGAGCTGGGCAAGACCGGTGCCAGCCGCCGCGGTAATACCGGCAGCTCGAGTGATGACCGATTTTATTGGGCCTAAAGCGTCCGTAGCTGGCCTCGCAAGTCCATCGGGAAATCCACCCGCTTAACGGGTGGGCGTCCGGTGGAAACTGTGTGGCTTGGGACAGGAAGGCTCGAGGGGTACGTTCGGGGTAGGAGTGAAATCCCGTAATCCTGAACGGACCGCCGATGGCGAAAGCACCTCGAGAGGACTGATCCGACAGTGAGGGACGAAAGCTGGGGTCT |
| p__Halobacteriota_g__Haloprofundus_ASV_1655 | 0 | 1 | 0 | 0 | 0 | 0 | 0 | 0 | 0 | 0 | 0 | CCTTTACACTGCACGACAGTGCGACAAGGGGACCCTGAGTGCGAGGGCATACAGTCCTCGCTTTTGTCGACCGTAAGAAGGTCGACGAATAAGAGCTGGGCAAGACCGGTGCCAGCCGCCGCGGTAATACCGGCAGCTCGAGTGATGACCGCTATTATTGGGCCTAAAGCGTCCGTAGCTGGCCGCGTAAGTCCATCGGGAAATCCATCCGCTCAACGGATGGGCGTCCGGTGGAAACTACACGGCTTGGAGCCGGAAGACCCAAGGGGTACGTCCGGGGTAGGAGTGAAATCCCGTAATCCTGGACGGACCGCCGATGGCGAAAGCACCTTGGGAAGACGGACTCGACAGTGAGGGACGAAAGCTAGGGTCT |
| p__Halobacteriota_g__Natronomonas_ASV_1660 | 0 | 0 | 1 | 0 | 0 | 0 | 0 | 0 | 0 | 0 | 0 | CCTTTACACTGCACGCCAGTGCGATAGGGGGACACCGAGTGCGAGGGCATAGAGCCCTCGCTTTTGTAGACCGTAAGGAGGTCTACGAATAAGGGCTGGGCAAGACCGGTGCCAGCCGCCGCGGTAACACCGGCAGCCCAAGTGATGGCCGATATTATTGGGCCTAAAGCGTCCGTAGCTGGCCGTTCAAGTTCGTCGGGAAATCCACTCGCTTAACGAGTGGGCGTCCGGCGAAAACTGTGCGGCTTGGGACCGGAAGATCCAAGGGGTACGTCCGGGGTAGGAGTGAAATCCTGTAATCCTGGACGGACCACCGATGGCGAAAGCACCTTGGAAGGACGGATCCGACAGTGAGGGACGAAAGCTAGGGTCT |
| p__Thermoproteota_g__Nitrososphaera_ASV_1671 | 0 | 0 | 0 | 0 | 0 | 1 | 0 | 0 | 0 | 0 | 0 | CCTCTGCAATAGGCGAAAGCTTGACAGGGTTACTCTGAGTGATGCCCGCTAAGGGTATCTTTTGGCACCTCTAAAAATGGTGCAGAATAAGGGGTGGGCAAGTCTGGTGTCAGCCGCCGCGGTAATACCAGCACCCCGAGTTGTCGGGACGATTATTGGGCCTAAAGCATCCGTAGCCTGTTCTGCAAGTCCTCCGTTAAATCCACCTGCTCAACGGATGGGCTGCGGAGGATACCGCAGAGCTAGGAGGCGGGAGAGGCAAACGGTACTCAGTGGGTAGGGGTAAAATCCATTGATCTACTGAAGACCACCAGTGGCGAAGGCGGTTTGCCAGAACGCGCTCGACGGTGAGGGATGAAAGCTGGGGGAG |
| p__Halobacteriota_g__Saliphagus_ASV_1658 | 0 | 1 | 1 | 0 | 0 | 0 | 0 | 0 | 0 | 0 | 0 | CCTTTACACTGCACGAGAGTGCGATAAGGGGACTCCGAGTGCGAGGGCATATCGTCCTCGCTTTTTCCTACCGTAAGGTGGTAGGCGAATAAGTGCTGGGCAAGACCGGTGCCAGCCGCCGCGGTAATACCGGCAGCACGAGTGATGACCGCTATTATTGGGCCTAAAGCGTCCGTAGCCGGCCTCGCAAGTCCGTCGGGAAATCCGCGCGCTCAACGCGCGGGCGTCCGGCGGAAACTGCGTGGCTTGGGACCGGAAGACTCAGGGGGTACGTCCGGGGTAGGAGTGAAATCCTGTAATCCTGGACGGACCACCGGTGGCGAAAGCGCCCTGAGAGGACGGATCCGACGGTGAGGGACGAAAGCTAGGGTCA |
| p__Thermoplasmatota_g__SM1-50_ASV_1640 | 1 | 1 | 0 | 0 | 0 | 0 | 0 | 0 | 0 | 1 | 0 | ACTTCACACTGCGCGAAAGCGCGATGAGGGGATCCCAAGTGCTTGCACGTAAGTGTAAGCTGTTTTTATGTTTAAAACGCATAGAGAGTAAGGGCTGGGTAAGACGGGTGCCAGCCGCCGCGGTAATACCTGCAGCTCAAGTGGTGGCCGCTATTATTGGGCCTAAAACGTCCGTAGCCGGTTTAGTAAATGCCTGGGTAAATCGTGCCGCTCAACGGTACGAATTCCGGGTAGACTGCTAAGCTTGAGACCGGGAGAGGCTAGAGGTACTCCTGGGGTAGAGGTGAAATTCTGTAATCCTAGGGGGACCACCAGTGGCGAAGGCGTCTAGCTAGAACGGGTCTGACGGTGAGGGACGAAGCCCTGGGGCG |
| p__Halobacteriota_o__Halanaerobiales_ASV_1657 | 0 | 1 | 1 | 0 | 0 | 0 | 0 | 0 | 0 | 0 | 0 | CCTTTACACTGCACGACAGTGCGATAAGGGGACTCCGAGTGCGAGGGCATACCGTCCTCGCTTTTCTGCACCGTAAGGTGGTGCAGGAATAAGAGCTGGGCAAGACCGGTGCCAGCCGCCGCGGTAATACCGGCAGCTCGAGTGATGGCCGTCATTATTGGGCCTAAAGCGTCCGTAGCCGGCCGGCCAAGTCCGTTGGGAAATCCACTCGCTTAACGAGTGGGCGGCCAGCGGAAACTGGCTGGCTTGGGACCGGAAGACCTGAGGGGTACGTCCGGGGTAGGAGTGAAATCCTGTAATCCTGGACGGACCGCCGGTGGCGAAAGCGCCTCAGGAAGACGGATCCGACGGTGAGGGACGAAAGCTAGGGTCT |
| p__Halobacteriota_o__Halanaerobiales_ASV_1659 | 0 | 0 | 1 | 0 | 0 | 0 | 0 | 0 | 0 | 0 | 0 | CCTTTACACTGCACGACAGTGCGATAAGGGGACCCCAAGTGCGAGGGCATATCGTCCTCGCTTTTCTGTACCGTAAGGAGGTGCAGGAATAAGGGCTGGGCAAGACCGGTGCCAGCCGCCGCGGTAATACCGGCAGCCCAAGTGATGGCCGCTATTATTGGGCCTAAAGCGTCCGTAGCCGGCCGGACAAGTCCGTCGGGAAATCCACCCGCTCAACGGGTGGGCGTCCGGCGGAAACTGTTCGGCTTGGGGCCGGAGGACCTGAGGGGTACGTCCGGGGTAGGAGTGAAATCCTGTAATCCTGGACGGACCACCGGTGGCGAAAGCGCCTCAGGAGGACGGACCCGACGGTCAGGGACGAAAGCTAGGGTCT |
| p__Halobacteriota_o__Halanaerobiales_ASV_1666 | 0 | 0 | 0 | 1 | 0 | 0 | 0 | 0 | 0 | 0 | 0 | CCTTTACACTGCACGACAGTGCGATAAGGGGACTCCGAGTGCGAGGGCATATAGTCCTCGCTTTTCTGTACCGTAGGGAGGTACAGGAACAAGTGCTGGGCAAGACCGGTGCCAGCCGCCGCGGTAATACCGGCAGCACGAGTGATGACCGATATTATTGGGCCTAAAGCGTCCGTAGCTGGCCAGACAAGTCCGTTGGGAAATCCACGCGCTCAACGCGTGGGCGTCCGGCGGAAACTGTTTGGCTTGGGGCCAGAAGACCTGAGGGGTACGTCCGGGGTAGGAGTGAAATCCTGTAATCCTGGACGGACCACCAATGGGGAAACCACCTCAGGAGGATGGACCCGACAGTGAGGGACGAAAGCTAGGGTCT |
| p__Nanoarchaeota_o__Woesearchaeales_581_ASV_1652 | 0 | 0 | 0 | 1 | 0 | 0 | 0 | 1 | 0 | 0 | 1 | CCTTTACAATGCACGAAAGTGTGATAAGGGGATCCCAAGTGCTCATGCACTGCATGGGCTTTTGCCAAGAGTAAATATCTTGGCGAATAAGTGGTGGGAAAGACTGGTGCCAGCAGCCGCGGTAACACCAGCGCCACGAGTGGCAGACATTATTATTGGGCCTAAAGCGTCCGTAGCCGGTTTAGTAAATCTTTTGTGAAATCAAGGGGCTTAACTCCTTGGCGTGCAGGAGAGACTGCTAGACTAGGAACCGGGAGGAGCCAGAAGTATTCCGGGAGTAGCGGTAAAATGCTATAATTCCCGGAGGACTACCTGTGGCGAAGGCGTCTGGCTAGAACGGATTCGACGGTGAGGGACGAAGGCCAGGGGAG |
| p__Nanoarchaeota_c__Nanoarchaeia_ASV_1678 | 1 | 1 | 0 | 0 | 0 | 0 | 0 | 0 | 0 | 0 | 1 | CCTTTACAATGCGCGAAAGCGTGATAAGGGAATCCCAAGTGCCATAGTAAATACTATGGCTGTGGCGAAGTGTAAAATACTTCGCAAGAAAGGGGTGGGTAATACCGGTGCCAGCAGCCGCGGTAATCCCGGAGCCTCAAGTGGTGACCATGTTTATTGGGCCTAAAGCGTTCGTAGCCTGATAAGTAAGTTCTCGGTGAAATCGTTCAGCTTAACTGAACGAATTGCCTTGAATACTGCTAATCTAGGGATTGGGAGAGGATAGAAGTATGCTCGGGGGAGGGGTAAAATCCTTTAATCCTGAGTAGACTGTCAATGGCGAAGGCATCTGTCTGGAACAAGTCCGACGGTGAGGGACGAAAGTTAGGGTAG |
| p__Iainarchaeota_o__Iainarchaeales_ASV_1690 | 0 | 0 | 0 | 0 | 0 | 0 | 0 | 0 | 0 | 1 | 1 | TCTCTACAATGCGCGAAAGCGTGATAGGGGGATTTCAAGTGCAAGCTTTGCTTGCTTTTGCATAATCCAAAAAGTTATGCGAATAAATGGAGGGTAAGATACGTGCCAGCCGCCGCGGTAATACGTACTCCATAAGTTGTGTCCACTAATATTGGGCCTAAAGCGTCTGTAGCCTGTCTGAAAAGTTTTTGGTGAAATCTGCAAGCCCAACTTGTAGGCGAGCCAAAAATACTCTTAGACTCGAGGCCGGGAGAAGTCAAAGGAATTCCTGAGGTAGCGGTGAAATGCTATGATCTTAGGAGGACCACCAGTAGCGAAGGCGTTTGACTAGAACGGACCTGACGGTGAGAGACGAAGGCTGGGGGAT |
| p__Nanoarchaeota_g__BMS3ABIN17_ASV_1686 | 0 | 0 | 0 | 0 | 0 | 0 | 0 | 0 | 1 | 1 | 1 | AATCTACAATGCGCGAAAGCGTGATAGTTTGAGTCAGAGTGCTATTCTTTTGAATAGCTTTTGCTAAATTTAAAAAGTTTGGCGAATAAGGACTGGGTAAGACAGGTGCCAGCCGCCGCGGTAATACCTGCGGTCCAAGTCGTAGTCACTTTTATTAGGTCTAAAACATCCGTAGCTTGCTTTGTAAGTCCCTTGTGAAATCCGGCATCTTAAGTGTTGGGCGGGCAAGGGATACTGCTTGGCTAGAGACTGGAAGACGCAAGAAGTATGTTCAGAGTAGCGGTCAAATGTGTTAATCCTGAGCAGACTAACAATAGCGAAGGCATCTTGCGAGGACAGTTCTGACAGTAAGGGATGAAGGCTAGGGGCG |
| p__Halobacteriota_g__Halobium_ASV_1676 | 1 | 0 | 0 | 0 | 0 | 1 | 1 | 0 | 0 | 0 | 0 | CCTTTACACTGCACGCACGTGCGATAAGGGGACCCCGAGTGCGAGGGCATACTAGCCCTCGCTTTTCGTAACCGTAAGGAGGTTACAGAATAAGAGCTGGGCAAGACCGGTGCCAGCCGCCGCGGTAATACCGGCAGCTCGAGTGATGACCGATTTTATTGGGCCTAAAGCGTCCGTAGCTGGCCTCGTCGGTCCGTCGGGAAATCCACCTGCCCAACAGGTGGGCGTCCGGCGGATACCACGGGGCTTGGGACCGGAAGACCCAGGGGGTACGTCCGGGGTAGGAGTGAAATCCCGTAATCCTGGACGGACCACCGATGGCGAAAGCACCCTGGGAGGACGGATCCGACAGTGAGGGACGAAAGCTCGGGTCT |
| p__Halobacteriota_g__Halohasta_ASV_1707 | 0 | 0 | 1 | 0 | 0 | 0 | 0 | 0 | 0 | 0 | 0 | CCTTTACACTGCACGGAAGTGCGATAGGGGAATCCCAAGTGCGAGGGCATACTAGCCCTCGCTTTTGTGTACCCTAAGGAGGTACACGAATAAGAGCTGGGCAAGACCGGTGCCAGCCGCCGCGGTAATACCGGCAGCTCGAGTGATGTCCACTATTATTGGGCCTAAAGCGTCCGTAGCTGGCCGCACAAGTCCGTCGGGAAATCCACCTGCCCAACAGGTGGGCGTCCGGCGGAAACTGTGTGGCTTGGAACCGGAAGGCTCAGAGGGTACGTCCGGGGTAGGAGTGAAATCCCGTAATCCCGGACGGACCACCGATGGCGAAAGCACTCTGAGAAGACGGATTCGACAGTGAGGGACGAAAGCTAGGGTCT |
| p__Halobacteriota_g__Halolamina_ASV_1694 | 0 | 0 | 0 | 0 | 0 | 0 | 0 | 0 | 0 | 0 | 1 | CCTTTACACTGCACGCCAGTGCGATAAGGGGACCCCGAGTGAAAGGGCATATCGCCCTTTCTTTTCTGAACCGTAGGGAGGTTGAGGAACAAGAGCTGGGCAAGACCGGTGCCAGCCGCCGCGGTAATACCGGCAGCTCGAGTGATGACCGATTTTATTGGGCCTAAAGCGTCCGTAGCTGGCCGCGCAAGTCTGTCGGGAAATCCACCCGCTCAACGGGTGGGCGTCCGGCGGAAACTGTGCGGCTTGGGACAGGAAGGCTCGAGGGGTACGTTCGGGGTAGGAGTGAAATCCCGTAATCCTGAACGGACCACCGATGGCGAAAGCACCTCGAGAGGACTGATCCGACAGTGAGGGACGAAAGCTGGGGTCT |
| p__Halobacteriota_g__Halolamina_ASV_1705 | 0 | 0 | 1 | 0 | 0 | 0 | 0 | 0 | 0 | 0 | 0 | CCTTTACACTGCACGCCAGTGCGATAAGGGGACCCCGAGTGAAAGGGCATATCGCCCTTTCTTTTCTGGACCGTAGGGAGGTCCAGGAACAAGAGCTGGGCAAGACCGGTGCCAGCCGCCGCGGTAATACCGGCAGCTCGAGTGATGACCGATCTTATTGGGCCTAAAGCGTCCGTAGCTGGCCGTGCAAGTCCGTCGGGAAATCCACCCGCTTAACGGGTGGGCGTCCGGCGGAAACTGCCCGGCTTGGGACAGGAAGGCTCGAGGGGTACGTTCGGGGTAGGAGTGAAATCCCGTAATCCTGAACGGACCGCCGATGGCGAAAGCACCTCGAGAGGACTGATCCGACAGTGAGGGACGAAAGCTGGGGTCT |
| p__Halobacteriota_g__Halorubrum_ASV_1700 | 0 | 1 | 1 | 0 | 0 | 0 | 0 | 0 | 0 | 0 | 0 | CCTTTACACTGCACGCCAGTGCGATAAGGGGATCCCAAGTGCGTAGGCATAGAGCCTACGCTTTTGTGAACCGTAGGGAGGTTCACGAATAAGGGCTGGGCAAGACCGGTGCCAGCCGCCGCGGTAATACCGGCAGCCCGAGTGATGGCCGATTTTATTGGGCCTAAAGCGTCCGTAGCTGGCCGCGCAAGTCCATTGGGAAATCTCCCCGCTCAACGGGGAGGCGCCCGGTGGAAACTGCGTGGCTTGGGACCGGAAGGCGCGACGGGTACGTCCGGGGTAGGAGTGAAATCCTGTAATCCTGGACGGACCGCCGATGGCGAAAGCACGTCGCGAGAACGGATCCGACAGTGAGGGACGAAAGCTAGGGTCT |
| p__Halobacteriota_g__Halorussus_ASV_1715 | 0 | 0 | 0 | 0 | 0 | 0 | 1 | 1 | 0 | 0 | 0 | CCTTTACACTGCACGACAGTGCGATAAGGGGACTCCGAGTGCGAGGGCATATAGTCCTCGCTTTTGTACACCGTAAGGTGGTGTACGAATAAGGACTGGGCAAGACCGGTGCCAGCCGCCGCGGTAATACCGGCAGTCCAAGTGATGGCCGCTATTATTGGGCCTAAAGCGTCCGTAGCCGGCCAGACAGGTCCGTCGGGAAATCTGCTCGCTCAACGAGCAGGCGTCCGGCGGAAACCAGCTGGCTTGGGGCCGGAAGACTCGAGGGGTACGTCCGGGGTAGGAGTGAAATCCCGTAATCCTGGACGGACCACCGGTGGCGAAAGCGCCTCGAGAAGACGGACCCGACGGTGAGGGACGAAAGCTAGGGTCA |
| p__Halobacteriota_g__Halovenus_ASV_1702 | 0 | 1 | 1 | 0 | 0 | 0 | 0 | 0 | 0 | 0 | 0 | CCTTTACACTGCACGACAGTGCGATAAGGGGACTCCGAGTGCGAGGGCATATAGTCCTCGCTTTTCTGTACCGTAAGGTGGTACAGGAACAAGTGCTGGGCAAGACCGGTGCCAGCCGCCGCGGTAATACCGGCAGCACGAGTGATGGCCGATTTTATTGGGCCTAAAGCGTCCGTAGCCGGCCAGTCAAGTTTGTTGGGAAATCCACGCGCTCAACGCGTGGGCGTCCAGCGAAAACTGTCTGGCTTGGGGCCGGAAGACTCGAGGGGTACGTCCGGGGTAGGAGTGAAATCCCGTAATCCTGGACGGACCGCCGGTGGCGAAAGCGCCTCGAGAGGACGGACCCGACGGTGAGGGACGAAAGCCAGGGTCT |
| p__Halobacteriota_g__Halovivax_ASV_1697 | 0 | 1 | 0 | 0 | 0 | 0 | 0 | 0 | 0 | 0 | 0 | CCTTTACACTGCACGACAGTGCGATAAGGGGACTCCAAGTGCGAGGGCATACAGTCCTCGCTTTTCACCACCGTAAGGAGGTGGTAGAATAAGGGCTGGGCAAGACCGGTGCCAGCCGCCGCGGTAATACCGGCAGCCCAAGTGATGACCGCTATTATTGGGCCTAAAGCGTCCGTAGCCGGCCAGGTAAGTCCGTCGGGAAATCCGCACGCCTAACGTGCGGGCGTCCGGTGGAAACTGCCTGGCTTGGGACCGGAAGATCCAGAGGGTACGTCCGGGGTAGGAGTGAAATCCTGTAATCCTGGACGGACCACCGGTGGCGAAAGCGCTCTGGAAGGACGGATCCGACGGTGAGGGACGAAAGCCAGGGTCA |
| p__Thermoproteota_g__Nitrosocosmicus_ASV_1713 | 0 | 0 | 0 | 0 | 0 | 1 | 0 | 0 | 0 | 0 | 0 | CCTCTGCAATAGGCGAAAGCTTGACAGGGTTACTCTGAGTGATTTCCGTTAAGGAGATCTTTTGGCACCTCTAAAAATGGTGCAGAATAAGGGGTGGGCAAGTCTGGTGTCAGCCGCCGCGGTAATACCAGCACCCCGAGTGGTCGGGACGTTTATTGGGCCTAAAGCATCCGTAGCCGGTTCTACAAGTCTTCCGTTAAATCCACCTGCTTAACAGATGGGCTGCGGAAGATACTATAGAGCTAGGAGGCGGGAGAGGCAAGCGGTACTCGATGGGTAGGGGTAAAATCCGTTGATCCATTGAAGACCACCAGTGGCGAAGGCGGCTTGCCAGAACGCGCTCGACGGTGAGGGATGAAAGCTGGGGGAG |
| p__Halobacteriota_g__Saliphagus_ASV_1706 | 0 | 0 | 1 | 0 | 0 | 0 | 0 | 0 | 0 | 0 | 0 | CCTTTACACTGCACGAAAGTGCGATAAGGGGATCCCGAGTGCGAGGGCATACAGTCCTCGCTTTTGTGTACCGTAAGGTGGTACACGAATAAGGGCTGGGCAAGACCGGTGCCAGCCGCCGCGGTAATACCGGCAGCTCAAGTGATGGCCGCTATTATTGGGCCTAAAGCGTCCGTAGCCGGCCGCACAAGTCCGTCGGGAAATCCACCCGCCTAACGGGTGGGCGTCCGGCGGAAACTGTGCGGCTTGGGACCGGAAGACTCGAGGGGTACGTCCGGGGTAGGAGTGAAATCCTGTAATCCTGGACGGACCGCCGGTGGCGAAAGCGCCTCGAGAGGACGGATCCGACGGTGAGGGACGAAAGCTAGGGTCT |

| Phylum_Genus_ASV_number | Presence (1) or Absence (0) in each layer | | | | | | | | | | | Sequence |
|---|---|---|---|---|---|---|---|---|---|---|---|---|
| | L1 | L2 | L3 | L4 | L5 | L6 | L7 | L8 | L9 | L10 | L11 | |
| p__Halobacteriota_o__Halanaerobiales_ASV_1691 | 0 | 1 | 1 | 0 | 0 | 0 | 0 | 0 | 0 | 1 | 1 | ACTTTACACTGCACGACAGTGCGATAAGGGAACTCCGAGTGCGAGGGCATATCGTCCTCGCTTTTCTGTACCGTAGGGAGGTACAGGAATAAGAGCTGGGCAAGACCGGTGCCAGCCGCCGCGGTAACACCGGCAGCTCGAGTGATGGCCGATATTATTGGGCCTAAAGCGTCCGTAGCCAGCTGAACAGGTCCGTCGGGAAATCTGCTCGCTCAACGAGCAGGCGTCCGGCGGAAACCGATCAGCTTGGGACCGAGAGGACCGAGGGGTACGTCCGGGGTAGGAGTGAAATCCCGTAATCCTGGACGGACCACCGATGGCGAAAGCACCTCGGTAGATCGGATCCGACGGTGAGGGACGAAAGCTAGGGTCA |
| p__Nanoarchaeota_g__BMS3ABIN17_ASV_1731 | 0 | 0 | 0 | 0 | 0 | 0 | 0 | 0 | 1 | 1 | 0 | CTTTTACAATGCGCGTAAGCGTGATAGAGCGAGCCAGAGTGCTTTTCCGTATTGGAAAAGCTTTTGTGAGATGCAAATAGTCTCACGAATAAGGACTGGGCAAGACTGGTGCCAGCCGCCGCGGTAATACCAGCGGTCCAAGTCGCAGCCAGTTTTATTGGGTCTAAAACATCCGTAGCTTGTTATGCAAGTCTCTTGTGAAATCTGGCATCTTAAGTGTCAGGCCTGCAGGAGATACTACATGGCTTGAGACCGGGAGACGTAAGGAGTACGGTTAGGGTAGCGGTAAAATGCGTTGATCCTTTCTGGACTAACGATAGCGAAGGCACCTTACGAGAACGGATCTGACAGTGAGGGATGAAGGCTAGGGGCG |
| p__Halobacteriota_g__Halohasta_ASV_1726 | 1 | 0 | 1 | 0 | 0 | 0 | 0 | 0 | 0 | 0 | 0 | CCTTTACACTGCACGCAAGTGCGATAGGGGAATCCCAAGTGCGAGGGCTAGTATGCCCTCGCTTTTGTGTACTGTAAGGAGGTACACGAATAAGAGCTGGGCAAGACCGGTGCCAGCCGCCGCGGTAATACCGGCAGCTCGAGTGATGTCCACTATTATTGGGCCTAAAGCGTCCGTAGCTGGCCGCGCAAGTCCGTCGGGAAATCCACCTGCCCAACAGGTGGGCGTCCGGCGGAAACTGCGTGGCTTGGAACCGGAAGGCTCAGAGGGTACGTCCGGGGTAGGAGTGAAATCCCGTAATCCCGGACGGACCACCGATGGCGAAAGCACTCTGAGAGGACGGATTCGACAGTGAGGGACGAAAGCTAGGGTCT |
| p__Halobacteriota_g__Halorientalis_ASV_1744 | 0 | 1 | 0 | 0 | 0 | 0 | 0 | 0 | 0 | 0 | 1 | CCTTTACACTGCACGACAGTGCGATAGGGGGACTCCGAGTGCGAGGGCATATAGTCCTCGCTTTTCTGTACCGTAAGGTGGTACAGGAACAAGTGCTGGGCAAGACCGGTGCCAGCCGCCGCGGTAATACCGGCAGCACGAGTGATGGCCGCTATTATTGGGCCTAAAGCGTCCGTAGCCGGCCAATCAAGTCCGTCGGGAAATCTGCCCGCTTAACGGGCAGGCGTCCGGCGGAAACTGATTGGCTTGGGGCCGGGAGATCCAAGGGGTACGTCCGGGGTAGGAGTGAAATCCTGTAATCCTGGACGGACCACCGATGGCGAAAGCACCTTGGAAAAACGGACCCGACGGTGAGGGACGAAAGCTAGGGTCT |
| p__Halobacteriota_g__Halorientalis_ASV_1756 | 0 | 0 | 1 | 0 | 0 | 0 | 0 | 0 | 0 | 0 | 0 | CCTTTACACTGCACGACAGTGCGATAGGGGGACTCCGAGTGCGAGGGCATACAGTCCTCGCTTTTCTGTACCGTAAGGTGGTACAGGAACAAGTGCTGGGCAAGACCGGTGCCAGCCGCCGCGGTAATACCGGCAGCACGAGTGATGGCCGCTATTATTGGGCCTAAAGCGTCCGTAGCCGGCCACGCAAGTCCGTCGGGAAATCTGCCCGCTTAACGGGCAGGCGTCCGGCGGAAACTGCGTGGCTTGGGGCCGGGAGATCCAAGGGGTACGTCCGGGGTAGGAGTGAAATCCTGTAATCCTGGACGGACCACCGATGGCGAAAGCACCTTGGAAAAACGGACCCGACGGTGAGGGACGAAAGCTAGGGTCT |
| p__Halobacteriota_g__Halorubrum_ASV_1727 | 1 | 0 | 1 | 0 | 0 | 0 | 0 | 0 | 0 | 0 | 0 | CCTTTACACTGCACGCAAGTGCGATAAGGGGACCCCGAGTGCGAGGGCATACCGTCCTCGCTTTTGTCGACCGTAGGGAGGTCGACGAATAAGGGCTGGGCAAGACCGGTGCCAGCCGCCGCGGTAATACCGGCAGCCCAAGTGATGGCCGCTCTTATTGGGCCTAAAGCGTCCGTAGCTGGCCGTGTAAGTTCATCGGGAAATCCACTCGCTCAACGGGTGGGCGTCCGGTGAAAACTGCACGGCTTGGGACCGGAAGACCCAGGGGGTACGTCCGGGGTAGGAGTGAAATCCCGTAATCCCGGACGGACCGCCGATGGCGAAAGCACTCTGGGAAGACGGCTCCGACAGTGAGGGACGAAAGCTGGGGTCT |
| p__Halobacteriota_g__Halorussus_ASV_1724 | 1 | 1 | 0 | 0 | 0 | 0 | 0 | 0 | 0 | 0 | 0 | CCTTTACACTGCACGACAGTGCGATAAGGGGACTCCGAGTGCGAGGGCATATAGTCCTCGCTTTTCTGTACCGTAAGGTGGTACAGGAATAAGGACTGGGCAAGACCGGTGCCAGCCGCCGCGGTAATACCGGCAGTCCGAGTGATGGCCGCTATTATTGGGCCTAAAGCGTCCGTAGCCGGCCAGACAGGTCCGTCGGGAAATCCACGCGCTCAACGCGTGGGCGTCCGGCGGAAACCAGCTGGCTTGGGGCCGGAAGACTCGAGGGGTACGTCCGGGGTAGGAGTGAAATCCTGTAATCCTGGACGGACCGCCGGTGGCGAAAGCGCCTCGAGAGGACGGACCCGACGGTGAGGGACGAAAGCTAGGGTCA |
| p__Halobacteriota_g__Natrinema_ASV_1752 | 0 | 1 | 0 | 0 | 0 | 0 | 0 | 0 | 0 | 0 | 0 | CCTTTACACTGCACGCGAGTGCGATAAGGGGACTCCAAGTGCGAGGGCATATAGTCCTCGCTTTTCACCACCGTAGGAAGGTGGTAGAATAAGTGCTGGGCAAGACCGGTGCCAGCCGCCGCGGTAATACCGGCAGCACGAGTGATGACCGCTATTATTGGGCCTAAAGCGTCCGTAGCTGGCCACGCAAGTCTATCGGGAAATCTGCGCGCTTAACGCGCAGGCGTCCGGTGGAAACTGTGTGGCTTGGGACCGGAAGACCAGAGGGGTACGTCCGGGGTAGGAGTGAAATCCCGTAATCCTGGACGGACCACCGGTGGCGAAAGCGCCTCTGGAAGACGGATCCGACGGTGAGGGACGAAAGCTCGGGTCA |
| p__Thermoplasmatota_g__SM1-50_ASV_1740 | 0 | 0 | 0 | 0 | 0 | 0 | 0 | 0 | 1 | 0 | 1 | CCTTCACACTGCGCGAAAGCGTGATGAGGGGATCCCAAGTGCTGGCACGTAAGTGTCAGCTGTTTCTCTGTCTAAAAAACAGAGAGAGTAAGGGCTGGGTAAGACGGGTGCCAGCCGCCGCGGTAATACCTGCAGCTCAAGTGGTGGCCGCTATTATTGGGCCTAAAACGTCCGTAGCCGGTCTTGTAAATGCCTGGGTAAATCGTGTCGCTTAACGATACGAATGCCGGGTAGACTGTAAGACTTGAGACCGGGAGAGGTTAGAGGTACTCCTGGGGTAGAGGTGAAATTCTGTAATCCTAGGGGGACCACCAGTGGCGAAGGCGTCTAACCAGAACGGGTCTGACGGTGAGGGACGAAGCCCTGGGGCG |
| p__Halobacteriota_f__Natrialbaceae_ASV_1755 | 0 | 1 | 1 | 0 | 0 | 1 | 0 | 0 | 1 | 0 | 0 | CCTTTACAATGCACGCAAGTGCGATAAGGGAACTCCAAGTGCGAGGGCATACCGTCCTCGCTTTTCTGCACTGTAAGGAGGTGCAGGAACAAGGGCTGGGCAAGACCGGTGCCAGCCGCCGCGGTAATACCGGCAGCCCAAGTGATGGCCAAGTTTATTGGGCCTAAAGCGTCCGTAGCCGGCCGGGCGGGTCCATCGGGAAATCCGCGCGCTCAACGTGTGGACGTCCGGTGGAAACTGTTCGGCTTGGGACCGGAGGACCCGAGGGGTACGTCCGGGGTAGGAGTGAAATCCTGTAATCCTGGACGGACCACCGGTGGCGAAAGCGCCTCGGGAAGACGGATCCGACGGTGAGGGACGAAAGCTAGGGTCT |
| p__Halobacteriota_o__Halanaerobiales_ASV_1757 | 0 | 0 | 1 | 0 | 0 | 0 | 0 | 0 | 0 | 0 | 0 | CCTTTACACTGCACGACAGTGCGATAGGGGGACTCCGAGTGCGAGGGCATATAGTCCTCGCTTTTCTGGACCGTAAGGAGGTCCAGGAACAAGTGCTGGGCAAGACCGGTGCCAGCCGCCGCGGTAATACCGGCAGCACGAGTGATGGCCAATCTTATTGGGCCTAAAGCGTCCGTAGCCGGCCCGACAAGTCCGTTGGGAAATCCACGCGCTCAACGCGTGGGCGTCCAGCGGAAACTGTCGGGCTTGGGACCGGAAGACCCAGAGGGTACGTCCGGGGTAGGAGTGAAATCCCGTAATCCTGGACGGACCACCGGTGGCGAAAGCGCTCTGGGAAGACGGATCCGACGGTGAGGGACGAAAGCTAGGGTCT |
| p__Nanoarchaeota_g__CG1-02-57-44_ASV_1774 | 1 | 0 | 0 | 0 | 0 | 0 | 0 | 0 | 0 | 0 | 0 | CCTTTACAATGCGCGAAAGCGTGATAAGGGAACCCTAAGTGCTCAACAAACGTTGGGCTTTTGCCAAGAGTAAATATCTTGGAGAATAAGTGGTGGGTAAGACTGGTGCCAGCCGCCGCGGTAACACCAGCGCCACGAGTGGGGATCGTGTTTATTGGGCCTAAAGCGTCCGTAGCCGGTCAAGTAAATCTTTTGTGAAATCTTCAGGCTCAACCTGTTGGCGCGCAGAGGACACTGCTTGACTAGGAACCGGGAGGAGTCAGAGGTATGGGGTGGGGAGCGGTAAAATGCGATAATCCACCTTAGACCACCTGTGGCGAAGGCGTCTGACCAGAACGGATTCGACGGTGAGGGACGAAAGCCAGGGGAG |
| p__Nanoarchaeota_g__CG1-02-57-44_ASV_1776 | 1 | 0 | 0 | 0 | 0 | 0 | 0 | 0 | 0 | 0 | 1 | CCTTTACAATGCACGCAAGTGTGATAAGGGGATCCCCAGTGCTCATGTACTGCATGGGCTTTTGCCAAGTGTAAATAGCTTGGCGAATAAGTGGTGGGTAAGACCGGTGCCAGCCGCCGCGGTAACACCGGCGCCACGAGTGGTAACCGCGTTTATTGGGCCTAAAGCGTCCGTAGCCGGCCTGATGAATCCTTTGTGAAATTGTCCGGCCTAACTGGGCAGCGCGCAAGGGACACTATCAGGCTCGAGACCGGGAGGAGTCAGAAGTATTCCAGGGGGAGCGGTAAAATGTTATAATCCCTGGAGGACTACCTGTGGCGAAAGCGTCTGACTAGAACGGATCTGACGGTGAGGGACGAAAGCCAGGGGAG |
| p__Halobacteriota_g__Halalkaliarchaeum_ASV_1793 | 0 | 1 | 0 | 0 | 0 | 0 | 0 | 0 | 0 | 0 | 0 | CCTTTACACTGCACGCGAGTGCGATAAGGGGACCCCGAGTGCGAGGGCATATCGTCCTCGCTTTTGTGTACCGTAGGGAGGTACACGAACAAGGGCTGGGCAAGACCGGTGCCAGCCGCCGCGGTAATACCGGCAGCCCAAGTGATGGCCGCTGTTATTGGGCCTAAAGCGTCCGTAGCCGGCCTGGCAAGTCCGTCGGGAAATCCACGCGCTCAACGCGTGGGCGGCCGGCGGAACCTGCCGGGCTCGGGACCGGAGGACCCAGCGGGTACGTCCGGGGTAGGAGTGAAATCCTGTAATCCCGGACGGACCGCCGATGGCGAAAGCACGCTGGGAAGACGGATCCGACGGTGAGGGACGAAAGCTAGGGTCT |
| p__Halobacteriota_g__Halalkalirubrum_ASV_1801 | 0 | 0 | 1 | 0 | 0 | 0 | 0 | 0 | 0 | 0 | 0 | CCTTTACACTGCACGCGAGTGCGATAGGGGGACCCCGAGTGCGAGGGCATACTAGCCCTCGCTTTTGTGAACCGTAAGGCGGTTTACGAACAAGAGCTGGGCAAGACCGGTGCCAGCCGCCGCGGTAATACCGGCAGCTCGAGTGATGACCGATTTTATTGGGCCTAAAGCGTCCGTAGCTGGCCACGAAAGTCCGTCGGGAAATCTCTCGGCTCAACCGAGAGGCGTCCGGCGGGACCTCCGTGGCTTGGAACCGGAAGGCTCAGAGGGTACGTCCAGGGTAGGAGTGAAATCCTGTAATCCTGGACGGACCACCGATGGCGAAAGCACTCTGAGAGGACGGATTCGACAGTGAGGGACGAAAGCTAGGGTCT |
| p__Halobacteriota_g__Halohasta_ASV_1808 | 0 | 0 | 1 | 0 | 0 | 0 | 0 | 0 | 0 | 0 | 0 | CCTTTACACTGCACGCAAGTGCGATAGGGGAATCCCAAGTGCGAGGGCATACTAGCCCTCGCTTTTGTGTACTGTAAGGAGGTACACGAATAAGAGCTGGGCAAGACCGGTGCCAGCCGCCGCGGTAATACCGGCAGCTCGAGTGATGTCCAATATTATTGGGCCTAAAGCGTCCGTAGCTGGCCAGCCAAGTCCGTCGGGAAATCCACCTGCCCAACAGGTGGGCGGCCGGCGGAAACTGTCTGGCTTGGAACCGGAAGGCTCAGAGGGTACGTCCGGGGTAGGAGTGAAATCCCGTAATCCCGGACGGACCACCGATGGCGAAAGCACTCTGAGAAGACGGATTCGACAGTGAGGGACGAAAGCTAGGGTCT |
| p__Halobacteriota_g__Halohasta_ASV_1820 | 0 | 0 | 0 | 0 | 0 | 1 | 0 | 0 | 0 | 0 | 0 | CCTTTACACTGCACGCAAGTGCGATAGGGGAATCCCAAGTGCGAGGGCATACTAGCCCTCGCTTTTGTTGACTGTAAGGTGGTCAACGAATAAGAGCTGGGCAAGACCGGTGCCAGCCGCCGCGGTAATACCGGCAGCTCGAGTGATGTCCAATATTATTGGGCCTAAAGCGTCCGTAGCTGGCCAGCCAAGTCCGTCGGGAAATCCACCTGCCCAACAGGTGGGCGTCCGGCGGAAACTGTCTGGCTTGGAACCGGAAGGCTCAGAGGGTACGTCCGGGGTAGGAGTGAAATCCCGTAATCCCGGACGGACCACCGATGGCGAAAGCACTCTGAGAAGACGGATTCGACAGTGAGGGACGAAAGCTAGGGTCT |
| p__Halobacteriota_g__Halolamina_ASV_1796 | 0 | 1 | 0 | 0 | 0 | 0 | 1 | 0 | 0 | 0 | 0 | CCTTTACACTGCACGCCAGTGCGATAAGGGAACCCCGAGTGCGAGGGCATATCGCCCTCGCTTTTCTGGACCGTAGGGAGGTCCAGGAACAAGAGCTGGGCAAGACCGGTGCCAGCCGCCGCGGTAATACCGGCAGCTCGAGTGATGACCGATTTTATTGGGCCTAAAGCGTCCGTAGCTGGCCGTGCAAGTCCGTCGGGAAATCCACCCGCTTAACGGGTGGGCGTCCGGCGGAAACTGTGCGGCTTGGGACAGGAAGGCTCGAGGGGTACGTTCGGGGTAGGAGTGAAATCCCGTAATCCTGAACGGACCGCCGATGGCGAAAGCACCTCGAGAGGACTGATCCGACAGTGAGGGACGAAAGCTGGGGTCT |
| p__Halobacteriota_g__Natrinema_ASV_1803 | 0 | 0 | 1 | 0 | 0 | 0 | 0 | 0 | 0 | 0 | 0 | CCTTTACACTGCACGCGAGTGCGATAAGGGGACTCCGAGTGCGAGGGCATATAGTCCTCGCTTTTCACCACCGTAAGGTGGTGGTGGAATAAGTGCTGGGCAAGACCGGTGCCAGCCGCCGCGGTAATACCGGCAGCACGAGTGATGACCGCTATTATTGGGCCTAAAGCGTCCGTAGCTGGCCACGCAAGTCTATCGGGAAATCTGCGCGCTTAACGCGCAGGCGTCCGGTGGAAACTGCGTGGCTTGGGACCGGAAGACCAGAGGGGTACGTCCGGGGTAGGAGTGAAATCCCGTAATCCTGGACGGACCACCGGTGGCGAAAGCGCCTCTGGAAGACGGATCCGACGGTGAGGGACGAAAGCTCGGGTCA |
| p__Thermoplasmatota_g__Natronoplasma_ASV_1807 | 0 | 0 | 1 | 0 | 0 | 0 | 0 | 0 | 0 | 0 | 0 | CCTTCACACTGTGGGAAACCACGATGAGGAGATTCCAAGTGCCAACACCCAGTGTTGGCTTTTCAGCTGTCTAAAACGCAGCTGGAATAAGGGCCGGGTAAGACGGGTGCCAGCCGCCGCGGTAACACCCGCGGCTCAAGTGGTGACCATTATTATTGAGCTTAAAGCGTTCGTAGCCGGTCTGGTAAATCTCTGGGTAAATCGAACGGCTCAACCGTTCGAATACCAGGGAGACTGCTAGACTTGGGACCGGGAGAGGCCAGAGGTACTTTTGGGGTAGAAGTAAAATTCTGTAATCCTAAAAGGACCACCAGTGGCGCAAGCGTCTGGCTAGAACGGATCCGACGGTGAGGGACGAAGCCCTGGGGCG |
| p__Asgardarchaeota_g__SDNM01_ASV_1826 | 0 | 0 | 0 | 0 | 0 | 0 | 0 | 0 | 1 | 0 | 0 | CCTCCACAATACACGAAAGTGTGATGGGGTTACCCAAAGTGTTCTATTAAGAACTGTGGTAGGTGAGTAATGTTCCCTACTAGAAAGGAGAGGGCAAGGCTGGTGCCAGCCGCCGCGGTAAAACCAGCTCTTCAAGTGGTCGGGATAATTATTGGGCTTAAAGTGTCCGTAGCCGGTTTAGTAAGTTCCTGGTAAAATCGGGTAGCTTAACTATCTGCATGCTAGGAATACTGCTATACTAGAGGGCGGGAGAGGTCTGAGGTACTACAGGGGTAGGGGTGAAATCTTATAATCCTTGTAGGACCACCAGTGGCGAAGGCGTCAGACTGGAACGCGCCTGACGGTGAGGGACGAAAGCCAGGGGAG |
| p__Thermoplasmatota_g__SM1-50_ASV_1824 | 0 | 0 | 0 | 0 | 0 | 0 | 0 | 0 | 1 | 0 | 0 | ACTTCACACTGCGCGAAAGCGCGATGAGGGAATCCCAAGTGTATGCACGTAAGTGTATACTGTTTTTATGTCTAAAAAGCATGGAGAGTAAGAGCTGGGTAAGACGGGTGCCAGCCGCCGCGGTAATACCTGCAGCTCGAGTGGTGATCACTATTATTGGGCCTAAAACGTCCGTAGCCGGTCTTTTAAATGCTTGGGTAAATCGTACGGCTTAACCGTACGAATTCCGGGTAGACTGTAAGACTTGAGACCGGGAGAAGTTAGAGGTACTCCTGGGGTAGAGGTGAAATTCTGTAATCCTAGGGGGACCACCAGTGGCGAAGGCGTCTAACTAGAACGGGTCTGACGGTGAGGGACGAAGCCCTGGGGCG |
| p__Halobacteriota_f__Haloferacaceae_ASV_1802 | 0 | 0 | 1 | 0 | 0 | 0 | 0 | 0 | 0 | 0 | 0 | CCTTTACACTGCGCGCACGCGCGATAAGGGGATCCCAAGTGCGAGGGCATACAGTCCTCGCTTTTCACATCCGTAGGGAGGATGTGGAATAAGGGCTGGGCAAGACCGGTGCCAGCCGCCGCGGTAATACCGGCAGCCCAAGTGATGGCCGCTCTTATTGGGCCTAAAGCGTCCGTAGCTGGCCACGTGTGTCCGTCGGGAAATCCACCAGCCCAACTGGTGGGCGGCCGGCGGAAACTGCGTGGCTTGGGACCGGAAGACCCACCGGGTACGTCCGGGGTAGGAGTGAAATCCCGTAATCCCGGACGGACCACCGATGGCGAAAGCACGGTGGGAAGACGGCTCCGACAGTGAGGGACGAAAGCTGGGGTCT |
| p__Halobacteriota_o__Halanaerobiales_ASV_1800 | 0 | 0 | 1 | 0 | 0 | 0 | 0 | 0 | 0 | 0 | 0 | CCTTTACACTGCACGCCCGTGCGATAAGGGGACTCCAAGTGCGAGGGCATACAGTCCTCGCTTTTCTCGACCGTAAGGTGGTCGAGGAACAAGAGCTGGGCAAGACCGGTGCCAGCCGCCGCGGTAATACCGGCAGCTCGAGTGATGGCCGATCTTATTGGGCCTAAAGCGTCCGTAGCCGGCCACGACGGTCCGTCGGGAAATCCATCCGCTCAACGGATGGGCGTCCGGCGGAAACCACGTGGCTTGGGGCCGGAAGGCTCAGGGGGTACGTCCGGGGTAGGAGTGAAATCCCGTAATCCTGGACGGACCACCGATGGCGAAAGCACCCTGAGAGAACGGACCCGACGGTGAGGGACGAAAGCTAGGGTCT |

| Phylum_Genus_ASV_number | Presence (1) or Absence (0) in each layer L1 | L2 | L3 | L4 | L5 | L6 | L7 | L8 | L9 | L10 | L11 | Sequence |
|---|---|---|---|---|---|---|---|---|---|---|---|---|
| d__Archaea_d__Archaea_ASV_1778 | 1 | 1 | 1 | 0 | 0 | 0 | 0 | 0 | 0 | 0 | 1 | CCTTTACAATGCACGGAAGTGCGATAAGGGGATCCCAAGTGCCAATGTTATGCATTGGCTTTTGCCAAGAGCAAAAATCTCGGCGAATAAGTGATGGGTAAGACTGGTGCCAGCCGCCGCGGTAACACCAGCGTCACGAGTGGCATCCACAATTATTGGGTCTAAAGTGTCCGTAGCTGGACATGTAAGTCCTCTGTGAAATTGTTAGGCTAAACCTAACGAACTGCGGAGGAGACTGCGTGTCTAGAGACCGGGAGGGGCTAGAGGTATTCTCGGGGAAGCGGTAAAATGCGATAATCCCGGGAAGACCACCTGTGGCGAAGGCGTCTAGCTAGAACGGGTCTGACAGTGAGGGACGAAGGCTAGGGGAG |
| p__Halobacteriota_g__Halopiger_A_ASV_1855 | 0 | 0 | 1 | 0 | 0 | 0 | 0 | 0 | 0 | 0 | 0 | CCTTTACACTGCACGCCAGTGCGATAAGGGGACTCCATCTGCGAGGGCATATCGTCCTCGCTTTTCTGCACCGTAAGGTGGTGCAGGAATAAGGGCTGGGCAAGACCGGTGCCAGCCGCCGCGGTAATACCGGCAGCCCGAGTGATGACCGCTATTATTGGGCCTAAAGCGTCCGTAGCTGGCCAGGCAAGTCCATCGGGAAATCCGCGTGCCCAACACGCGGGCGTCCGGTGGAACCTGCCTGGCTTGGGACCGGAAGACCTGAGGGGTACGTCCGGGGTAGGAGTGAAATCCCGTAATCCTGGACGGACCACCGGTGGCGAAAGCGCCTCAGGAAGACGGATCCGACGGTGAGGGACGAAAGCTCGGGTCA |
| p__Halobacteriota_g__Halorussus_ASV_1840 | 0 | 0 | 0 | 0 | 0 | 1 | 0 | 0 | 0 | 0 | 1 | CCTTTACCCTGCGCGAGAGCGCGATAAGGGGACCCCAAGTGCGAGGGCATATAGTCCTCGCTGTTCTGGACTGTAGGGAGGTTCAGAAGAAAGGGCTGGGCAAGACCGGTGCCAGCCGCCGCGGTAATACCGGCAGCCCGAGTGATGGCCGATTTTATTGGGCCTAAAGCGTCCGTAGCCGGCCAGACAGGTCCGTCGGGAAATCCACGCGCTTAACGCGTGGAGGTCCGGCGGAAACCAGCTGGCTTGGGGCCGGAAGACCCAGGGGGTACGTCTGGGGTAGGAGTGAAATCCTGTAATCCCGGACGGACCGCCGGTGGCGAAAGCGCCCTGGGAGGACGGACCCGACGGTGAGGGACGAAAGCCAAGGTCA |
| p__Halobacteriota_g__Halorussus_ASV_1843 | 0 | 1 | 0 | 0 | 0 | 0 | 0 | 0 | 0 | 0 | 0 | ACTTTACACTGCACGACAGTGCGATAAGGGGACTCCGAGTGCGAGGGCATATAGTCCTCGCTTTTGTACACCTTAAGGCGGTGTACGAATAAGGGCTGGGCAAGACCGGTGCCAGCCGCCGCGGTAATACCGGCAGCCCGAGTGATGGCCGCTATTATTGGGCCTAAAGCGTCCGTAGCCGGCCAGACAGGTCCGTCGGGAAATCCACGCGCTCAACGCGTGGGCGTCCGGCGGAAACCAGCTGGCTTGGGGCCGGAAGACCTGAGGGGTACGTCCGGGGTAGGAGTGAAATCCTGTAATCCTGGACGGACCGCCGGTGGCGAAAGCGCCTCAGGAAGACGGACCCGACGGTGAGGGACGAAAGCTAGGGTCA |
| p__Halobacteriota_g__Halorussus_ASV_1870 | 0 | 0 | 0 | 0 | 1 | 0 | 0 | 0 | 0 | 0 | 0 | CCTTTACACTGCACGCCAGTGCGATAAGGGGATCCCGAGTGCGAGGGCATACAGTCCTCGCTTTTGTACACCGTAAGGTGGTGTACGAATAAGAGCTGGGCAAGACCGGTGCCAGCCGCCGCGGTAATACCGGCAGCTCGAGTGATGGCCGCTATTATTGGGCCTAAAGCGTCCGTAGCCGGCCGAACAGGTCCGTTGGGAAATCCATCTGCCTAACAGATGGGCGTCCAGCGGAAACCAGTCGGCTTGGGGCCGGAAGACCTGAGGGGTACGTCCGGGGTAGGAGTGAAATCCTGTAATCCTGGACGGACCGCCGGTGGCGAAAGCGCCTCAGGAGGACGGACCCGACGGTGAGGGACGAAAGCTTGGGTCA |
| p__Halobacteriota_g__Halostella_ASV_1851 | 0 | 1 | 1 | 0 | 0 | 0 | 0 | 0 | 0 | 0 | 0 | CCTTTACACTGCACGCGAGTGCGATAAGGGGACCCCGAGTGCGAGGGCATATAGCCCTCGCTTTTGCACACCGTACTGAGGTGCGCGAATAAGGACTGGGCAAGACCGGTGCCAGCCGCCGCGGTAATACCGGCAGTCCAAGTGATGGCCGCTGTTATTGGGCCTAAAGCGTCCGTAGCCGGCCAGGCAAGTCCGTCGGGAAATCCACCTGCTCAACAGGTGGGCGTCCGGCGGAAACTGTCTGGCTAGGGACCGGAAGACCCGAGGGGTACGTCCAGGGTAGGAGTGAAATCCCGTAATCCTGGACGGACCGCCGGTGGCGAAAGCGCCTCGGGAGAACGGATCCGACGGTGAGGGACGAAAGCTTGGGTCA |
| p__Halobacteriota_g__Halovivax_ASV_1853 | 0 | 0 | 1 | 0 | 0 | 0 | 0 | 0 | 0 | 0 | 0 | CCTTTACACTGCACGAGAGTGCGATAGGGGGACTCCAAGTGCGAGGGCATATAGTCCTCGCTTTTCACTACCGTAAGGTGGTAGTAGAATAAGGGCTGGGCAAGACCGGTGCCAGCCGCCGCGGTAATACCGGCAGCTCGAGTGATGACCGCTATTATTGGGCCTAAAGCGTCCGTAGCCGGCCAGGCAAGTCTGTCGGGAAATCCGCACGCCTAACGTGCGGGCGTCCGGTGGAAACTGCTTGGCTTGGGACCGGAAGATCCAGAGGGTACGTCTGGGGTAGGAGTGAAATCCTGTAATCCTGGACGGACCACCGGTGGCGAAAGCGCTCTGGAAGGACGGATCCGACGGTGAGGGACGAAAGCTAGGGTCA |
| p__Halobacteriota_g__Methanolobus_ASV_1842 | 0 | 1 | 0 | 0 | 0 | 0 | 0 | 0 | 0 | 0 | 0 | ACTTTACAATGCGGGAAACCGCGATAAGGGGACACCGAGTGCCAGCATATTATGTTGGCTGTCCATATGTGTAAATAACATGTGTTAGCAAGGGCCGGGCAAGACCGGTGCCAGCCGCCGCGGTAACACCGGCGGCCCGAGTGGTAGCCACTATTATTGGGTCTAAAGGGTCCGTAGCCGGTTTGATCAGTCTTCCGGGAAATCTGACAGCTCAACTGTCAGGCTTCCGGGGGATACTGTCAGACTTGGGACCGGGAGAGGTAAGAGGTACTACAGGGGTAGGAGTGAAATCTTGTAATCCTTGTGGGACCACCAGTGGCGAAGGCGTCTTACCAGAACGGGTCCGACGGTGAGGGACGAAAGCTGGGGGCA |
| p__Halobacteriota_g__Natrarchaeobaculum_ASV_1869 | 0 | 0 | 0 | 0 | 1 | 0 | 0 | 0 | 0 | 0 | 0 | CCTTTACACTGCGCGACAGCGCGATAAGGGGACTCCATCTGCGAGGGCATATCGTCCTCGCTTTTCTCGACCGTAAGGTGGTCGAGGAATAAGGGCTGGGCAAGACCGGTGCCAGCCGCCGCGGTAATACCGGCAGCCCGAGTGATGACCGCTCTTATTGGGCCTAAAGCGTCCGTAGCTGGCCTCGCAAGTCCATCGGGAAATCCGCACGCTCAACGTGCGGGCGTCCGGTGGAAACTGCGTGGCTTGGGACCGGAAGACCCGAGGGGTACGTCTGGGGTAGGAGTGAAATCCCGTAATCCTGGACGGACCACCGGTGGCGAAAGCGCCTCGGGAGGACGGATCCGACGGTGAGGGACGAAAGCCAGGGTCA |
| p__Halobacteriota_g__Salinibaculum_ASV_1854 | 0 | 0 | 1 | 0 | 0 | 0 | 0 | 0 | 0 | 0 | 0 | CCTTTACACTGCACGAGAGTGCGATAGGGGGACTCCGAGTGCGAGGGCATACAGTCCTCGCTTTTCTGTACCGTAAGGTGGTACAGGAACAAGTGCTGGGCAAGACCGGTGCCAGCCGCCGCGGTAATACCGGCAGCACAAGTGATGGCCGATATTATTGGGCCTAAAGCGTCCGTAGCTGGCCGAGCAAGTCCGTTGGGAAATCGACGTGCCTAACACGTCGGCGTCCAGCGGAAACTGTTCGGCTTGGGGCCGGAAGACCCGAGGGGTACGTCCGGGGTAGGAGTGAAATCCCGTAATCCTGGACGGACCGCCGGTGGCGAAAGCGCCTCGGGAGGACGGACCCGACAGTGAGGGACGAAAGCCAGGGTCT |
| p__Halobacteriota_g__Saliphagus_ASV_1844 | 0 | 1 | 0 | 0 | 0 | 0 | 0 | 0 | 0 | 0 | 0 | CCTTTACACTGCACGACAGTGCGATAGGGGGACCCCGAGTGCGAGGGCATACAGTCCTCGCTTTTGTGTACCGTAAGGAGGTACACGAATAAGAGCTGGGCAAGACCGGTGCCAGCCGCCGCGGTAATACCGGCAGCTCAAGTGATGGCCGCTGTTATTGGGCCTAAAGCGTCCGTAGCCGGCCGCACAAGTCCGTCGGGAAATCCACCCGCCTAACGGGTGGGCGTCCGGCGGAAACTGTGCGGCTTGGGACCGGAAGACTCGAGGGGTACGTCCGGGGTAGGAGTGAAATCCCGTAATCCTGGACGGACCGCCGGTGGCGAAAGCGCCTCGAGAGGACGGATCCGACGGTGAGGGACGAAAGCTAGGGTCT |
| p__Halobacteriota_f__Haloferacaceae_ASV_1858 | 0 | 0 | 1 | 0 | 0 | 0 | 0 | 0 | 1 | 0 | 0 | CCTTTACACTGCACGCCAGTGCGATAGGGGGACCCCGAGTGCGAGGGCATATAGTCCTCGCTTTTGCACACCGTAAGGTGGTGCGCGAATAAGTGCTGGGCAAGACCGGTGCCAGCCGCCGCGGTAATACCGGCAGCACAAGTGATGGCCGCTATTATTGGGCCTAAAGCGTCCGTAGCCGGCCGAACAAGTCCGTCGGGAAATCCATCCGCTCAACGGATGGGCGTCCGGCGGAAACTGTTCGGCTTGGGACAGGAAGACTCGAGGGGTACGTCCGGGGTAGGAGTGAAATCCTGTAATCCTGGACGGACCGCCGATGGCGAAAGCACCTCGAGAAGACTGATCCGACGGTGAGGGACGAAAGCTGGGGTCT |
| p__Thermoproteota_f__Nitrososphaeraceae_ASV_1878 | 0 | 0 | 0 | 0 | 0 | 1 | 0 | 0 | 0 | 0 | 0 | CCTCTGCAATAGGCGAAAGCTTGACAGGGTTACTCTGAGTGATGCCCGCTAAGGGTATCTTTTGGCACCTCTAAAAATGGTGCAGAATAAGGGGTGGGCAAGTCTGGTGTCAGCCGCCGCGGTAATACCAGCACCCCGAGTTGTCGGGACGATTATTGGGCCTAAAGCATCCGTAGCCTGTTCTGCAAGTCCTCCGTTAAATCCACCCGCTTAACGGATGGGCTGCGGGGGATACTGCAGAGCTAGGAGGCGGGAGAGGCAAACGGTACTCAGTGGGTAGGGGTAAAATCCATTGATCTACTGAAGACCACCAGTGGCGAAGGCGGTTTGCCAGAACGCGCTCGACGGTGAGGGATGAAAGCTGGGGGAG |
| p__Nanoarchaeota_o__Woesearchaeales_581_ASV_1830 | 1 | 0 | 0 | 0 | 0 | 0 | 0 | 0 | 0 | 0 | 1 | CCTTTACAATGCACGAAAGTGTGATAAGGGAATCCCCAGTGCTCATGCACAGCATGGGCTTTTGCCAAGAGTAAAAACCTTGGCGAATAAGTGGTGGGTAAGACTGGTGCCAGCAGCCGCGGTAACACCAGCGCCACGAGTGGGAATCGCGATTATTGGGCCTAAAGCGTCCGTAGCCGGTTCAGTAAATCTTTAGTGAAATCGTTCTGCTTAACAGAACGGCGTGCTGAAGAGATTGCTGGACTAGGAACCGGGAGGAGCCAGAAGTATATTAGGGGGAGCGGTAAAATGCAATAATCCCTAATAGACTACCTGTGGCGAAGGCGTCTGGCTAGAACGGATTCGACGGTGAGGGACGAAAGCTAGGGGAG |
| p__Thermoplasmatota_g__FEN-33_ASV_1918 | 0 | 0 | 1 | 1 | 0 | 0 | 0 | 0 | 0 | 0 | 0 | ACTTCGCAATGGGGGAAACCCTGACGAGGGGATTCCAAGTGCCAGCACACAGTGTTGGCTGTTCTTCTGTCCAAAACGCAGGAGAAGTAAGGGCCGGGTAAGACGGGTGCCAGCCGCCGCGGTAATACCCGCGGCCCGAGTGGTGGTCGATTTTATTGAGCCTAAAACGTCCGTAGCCGGCCTGGTAAATCCTTGGGTAAATCGGCCAGCTTAACTGTCCGAAGTCCGGGGAGACTGCCAGGCTTGGGACCGGGAGAGGTCAGAGGTACTTCTGGGGTAGGGGTAAAATCCTGTAATCCTGGGAGGACCACCGGTGGCGAAGGCGTCTGACTAGAACGGATTCGACGGTGAGGGACGAAGCCCTGGGGCG |
| p__Halobacteriota_g__Halalkalicoccus_ASV_1934 | 0 | 0 | 0 | 0 | 0 | 1 | 0 | 0 | 0 | 0 | 0 | CCTTTACACTGCACGAACGTGCGATAAGGGGACTCCAAGTGCGAGGGCATATAGTCCTCGCTTTTGCCTACCGTAAGGTGGTAGGCGAATAAGAGCTGGGCAAGACCGGTGCCAGCCGCCGCGGTAATACCGGCAGCTCGAGTGATGGCCAATATTATTGGGCCTAAAGCGTCCGTAGCTGGCCGAACAGGTCCGTCGGGAAATCCGCTCGCTCAACGAGCGGACGTCCGGCGGAAACCAGTCGGCTTGGGACCGGAAGACCCGAGGGGTACGGCCGGGGTAGGAGTGAAATCCTGTAATCCTGGCCGGACCACCGGTGGCGAAAGCGCCTCGGGAGGACGGATCCGACAGTGAGGGACGAAAGCTTGGGTCA |
| p__Halobacteriota_g__Halobellus_3026_ASV_1914 | 0 | 0 | 1 | 0 | 0 | 0 | 0 | 0 | 0 | 0 | 0 | CCTTTACACTGCACGCCAGTGCGATAAGGGGACTCCAAGTGCGAGGGCATATCGTCCTCGCTATCCTGTACTGTAGGGTAGTACAGTATCAAGAGCTGGGCAAGACCGGTGCCAGCCGCCGCGGTAATACCGGCAGCTCAAGTGATGGCCGATCTTATTGGGCCTAAAGCGTCCGTAGCTGGCCCTGCAAGTCCGTCGGGAAATCCATCAGCCCAACTGATGGGCGGCCGGCGGAAACTCCAGGGCTTGGGACCGGAAGGCTCGAGGGGTACGTCCGGGGTAGGAGTGAAATCCCGTAATCCCGGACGGACCACCGATGGCGAAAGCACCTCGAGAAGACGGCTCCGACAGTGAGGGACGAAAGCTAGGGTCT |
| p__Halobacteriota_g__Halococcus_ASV_1892 | 0 | 0 | 0 | 0 | 0 | 0 | 0 | 0 | 0 | 1 | 1 | CCTTTACACTGCACGCCAGTGCGATAAGGGGATCCCGAGTGCGAGGGCATACAGTCCTCGCTTTTGTGTACCGTAAGGTGGTACTCGAATAAGGGCTGGGCAAGACCGGTGCCAGCCGCCGCGGTAATACCGGCAGCCCAAGTGATAGCCACTATTATTGGGCCTAAAGCGTCCGTAGCCTGCCGAACGGGTCCGTCGGGAAATCCACCCGCTCAACGGGTGGGCGTCCGGCGGAAACCAGTCGGCTTGGGACCGGGAGACCTGTGGGGTACGTCCGGGGTAGGAGTGAAATCCTGTAATCCTGGACGGACCACCGGTGGCGAAAGCGCCACAGGAGAACGGATCCGACGGTGAGGGACGAAAGCTAGGGTCT |
| p__Halobacteriota_g__Halonotius_ASV_1915 | 0 | 0 | 1 | 0 | 0 | 0 | 0 | 0 | 0 | 0 | 0 | CCTTTACACTGCACGGAAGTGCGATAGGGGAATCCCAAGTGCGAAGGCATACTAGCCTTCGCTTTTGTGTACTGTAAGGAGGTACACGAATAAGAGCTGGGCAAGACCGGTGCCAGCCGCCGCGGTAATACCGGCAGCTCGAGTGATGTCCCATATTATTGGGCCTAAAGCGTCCGTAGCTGGCCCGCGCAAGTCCGTCGGGAAATCCACCTGCCCAACAGGTGGGCGTCCGGCGGAAACTGCGTGGCTTGGAACCGGAAGGCTCAGAGAGTACGTCCGGGGTAGGAGTGAAATCCCGTAATCCCGGACGGACTACCGATGGCGAAAGCACTCTGAGAAGACGGATTCGACAGTGAGGGACGAAAGCTAGGGTCT |
| p__Halobacteriota_g__Haloprofundus_ASV_1899 | 0 | 1 | 0 | 0 | 0 | 0 | 0 | 0 | 0 | 0 | 0 | CCTTTACACTGCACGACAGTGCGACAAGGGGACCCCGAGTGCGAGGGCATACAGTCCTCGCTTTTGTGACCCGTAAGAAGGTTCACGAATAAGAGCTGGGCAAGACCGGTGCCAGCCGCCGCGGTAATACCGGCAGCTCGAGTGATGACCGATTTTATTGGGCCTAAAGCGTCCGTAGCTGGCCGTGTAAGTCCATCGGGAAATCCATCCGCTCAACGGATGGGCGTCCGGTGGAAACTGCGCGGCTTGGAGCCGGAAGACCCAAGGGGTACGTCTGGGGTAGGAGTGAAATCCCGTAATCCTGGACGGACCGCCGATGGCGAAAGCACCTTGGGAAGACGGACTCGACAGTGAGGGACGAAAGCTAGGGTCT |
| p__Halobacteriota_g__Halorussus_ASV_1907 | 0 | 0 | 1 | 0 | 0 | 0 | 0 | 0 | 0 | 0 | 0 | CCTTTACACTGCACGACAGTGCGATAAGGGGACTCCGAGTGCGAGGGCATATCGTCCTCGCTTTTGTACACCGTAAGGTGGTGTACGAATAAGGGCTGGGCAAGACCGGTGCCAGCCGCCGCGGTAATACCGGCAGCCCGAGTGATGGCCGCTATTATTGGGCCTAAAGCGTCCGTAGCCGGCCAGACAGGTCCGTCGGGAAATCCACGCGCTCAACGCGTGGGCGTCCGGCGGAAACCAGCTGGCTTGGGGCCGGAAGACCTGAGGGGTACGTCCGGGGTAGGAGTGAAATCCCGTAATCCTGGACGGACCACCGGTGGCGAAAGCGCCTCAGGAAGACGGACCCGACGGTGAGGGACGAAAGCTAGGGTCA |
| p__Halobacteriota_g__Halorussus_ASV_1908 | 0 | 0 | 1 | 0 | 0 | 0 | 0 | 0 | 0 | 0 | 0 | ACTTTACACTGCACGACAGTGCGATAAGGGGACTCCGAGTGCGAGGGCATATAGTCCTCGCTTTTGTACACCGTAGGGTGGTGTACGAATAAGGGCTGGGCAAGACCGGTGCCAGCCGCCGCGGTAATACCGGCAGCCCGAGTGATGGCCGCTATTATTGGGCCTAAAGCGTCCGTAGCCGGCCAGACAGGTCCGTCGGGAAATCCATCCGCTTAACGGATGGGCGTCCGGCGGAAACCAGCTGGCTTGGGGCCGGAAGATCCGAGGGGTACGTCCGGGGTAGGAGTGAAATCCCGTAATCCTGGACGGACCACCGGTGGCGAAAGCGCCTCGGAAAGACGGACCCGACGGTGAGGGACGAAAGCTAGGGTCA |
| p__Halobacteriota_g__Natronobiforma_ASV_1911 | 0 | 0 | 1 | 0 | 0 | 0 | 0 | 0 | 0 | 0 | 0 | CCTTTACACTGCACGAGAGTGCGATAGGGGGACTCCAAGTGCGAGGGCATATAGTCCTCGCTTTTGTGTACCGTAAGGTGGTACACGAATAAGGGCTGGGCAAGACCGGTGCCAGCCGCCGCGGTAATACCGGCAGCCCAAGTGATGGCCGCTATTATTGGGCCTAAAGCGTCCGTAGCCGGCCAAGCAAGTCCGTCGGGAAATCTGCTCGCCTAACGAGCAGGCGTCCGGCGGAAACTGTTTGGCTTGGGACCGGAGGACCTGGGGGGTACGTCTGGGGTAGGAGTGAAATCCCGTAATCCTGGACGGACCACCGGTGGCGAAAGCGCCTCAGGAAGACGGATCCGACGGTGAGGGACGAAAGCTAGGGTCA |
| p__Thermoplasmatota_g__Natronoplasma_ASV_1887 | 1 | 0 | 0 | 0 | 0 | 0 | 0 | 0 | 0 | 0 | 1 | CCTTCACAATGTGGGAAACCACGATGAGGAGATTCCAAGTGCCAACACCCAGTGTTGGCTTTTCAACTGTCTAAAACGCAGTTGGAATAAGGGCCGGGTAAGACTGGTGCCAGCCGCCGCGGTAACACCAGCGGCTCGAGTGGTGGCCTTTATTATTGAGCTTAAAGCGTCCGTAGCCGGTCTGGTAAATCTCTGGGTAAATCGAACAGCTTAACTGTTCGAATGCCAGAGAGACTGCTAGACTTGGGGCCGGGAGAGGCTAGAGGTACTCCTGGGGTAGAGGTAAAATTCCGTAATCTTAGGAGGACCACCAGTGGCGAAAGCGTCTAGCTAAAACGGACCCGACGGTGAGGGACGAAGCCCTGGGGCG |

| Phylum_Genus_ASV_number | Presence (1) or Absence (0) in each layer | | | | | | | | | | | |
|---|---|---|---|---|---|---|---|---|---|---|---|---|
| | L1 | L2 | L3 | L4 | L5 | L6 | L7 | L8 | L9 | L10 | L11 | Sequence |
| p__Nanoarchaeota_g__UBA583_ASV_1886 | 1 | 0 | 0 | 0 | 0 | 0 | 0 | 0 | 0 | 0 | 0 | CCTTTACAATGCACGAAAGTGTGATAAGGTGACCCCTCGTGCTTATCTAACGATAGGCTTTTTTTAGCTTTAGTGAAGCTAAAGAATAAGTGGTGGGCAAGACCGGTGCCAGCAGCCGCGGTAATACCGGCGCCACAAGTGGAAATCATGTTTACTGGGTCTAAAGTGTTCGTAGCTGGTTAAATAAGTTTCTTGTGAAAGACCTATTGCTTAACTTAGGAATTGCAGGAAATACTGTTTAACTTGAGACCGGAAGAGGTGCAGAGTATTGCTTGGGTAGCGGTAAAATGCTATAATCCTTGCAAGACTAACTAAGGCGAAGGCATTGCACTAGGACGGATCTGACAGTGAGGAACGAGAGCTGGGGGAG |
| p__Halobacteriota_f__Haloferacaceae_ASV_1901 | 0 | 1 | 0 | 0 | 0 | 0 | 0 | 0 | 1 | 0 | 0 | CCTTTACACTGCACGCAGGTGCGATAAGGGGACCCCGAGTGCGAGGGCATACAGTCCTCGCTTTTGAGAACCGTAAGGCGGTTCTCGAATAAGAGCTGGGCAAGACCGGTGCCAGCCGCCGCGGTAATACCGGCAGCTCGAGTGATGACCGATCTTATTGGGCCTAAAGCGTCCGTAGCTGGCCGCGTTGGTCCGTCGGGAAATCCAGCCGCTCAACGGCTGGGCGTCCGGCGGATACCGCGTGGCTTGGGACAGGAAGACTCAAGGGGTACGTCCGGGGTAGGAGTGAAATCCCGTAATCCTGGACGGACCGCCGATGGCGAAAGCACCTTGAGAGGACTGCTCCGACAGTGAGGGACGAAAGCTGGGGTCT |
| p__Halobacteriota_f__Haloferacaceae_ASV_1933 | 0 | 0 | 0 | 0 | 0 | 1 | 0 | 0 | 0 | 0 | 0 | CCTCAACACTGCACGCCAGTGCGATTGGGGGACTCCGAGTGCGAGGGCATACCGTCCTCGCTTTTCAGCACGGTAAGGACGTGTTGGAACAAGGGCTGGGCAAGACCGGTGCCAGCCGCCGCGGTAATACCGGCAGCCCGAGTGATGACCGATCTTATTGGGCCTAAAGCGTCCGTAGCCGGCCGGGCAAGTCCGTCGGGAAATCCGCGCGCTCAACGCGTGGGCGTCCGGCGGAACCTGTTCGGCTTGGGGCCGGAAGACCCAGGGGGTACGTCCGGGGTAGGAGTGAAATCCCGTAATCCCGGACGGACCGCCGATGGCGAAAGCACCCTGGGAGGACGGACCCGACGGTGAGGGACGAAAGCTGGGGTCT |
| p__Halobacteriota_f__Methanosarcinaceae_ASV_1925 | 0 | 0 | 0 | 1 | 0 | 0 | 0 | 0 | 1 | 0 | 0 | ACTTTACAATGCGGGAAACCGCGATAAGGGGACACCGAGTGCCAGCATATTATGTTGGCTGTCCACATGTGTAAACAGCATGTGTTAGCAAGGGCCGGGCAAGACCGGTGCCAGCCGCCGCGGTAACACCGGCGGCCCGAGTGGTAATCACTATTATTGGGTCTAAAGGGTCCGTAGCCGGTTTGATCAGTCTTTCGGGAAATCTGATGGCTTAACTATCAGGCTTCCGGGAGATACTGTCAGACTTGGGACCGGGAGAGGTAAGAGGTACTACAGGGGTAGGAGTGAAATCTTGTAATCCCTGTGGGACCACCGGTGGCGAAGGCGTCTTACCAGAACGGGTCCGACGGTGAGGGACGAAAGCTGGGGGCA |
| p__Halobacteriota_f__Natrialbaceae_ASV_1912 | 0 | 0 | 1 | 0 | 0 | 0 | 0 | 0 | 0 | 0 | 0 | CCTTTACACTGCGCGACAGCGCGATAAGGGGACTCCAAGTGCGAGGGCATATCGTCCTCGCTTTTCACCATCGTAAGGAGATGGTGGAATAAGGGCTGGGCAAGACCGGTGCCAGCCGCCGCGGTAATACCGGCAGCCCGAGTGATGACCGCTATTATTGGGCCTAAAGCGTCCGTAGCCGGCCGATCAAGTCCGTCGGGAAATCCGCACGCCTAACGTGCGGGCGTCCGGCGGAAACTGCTCGGCTTGGGACCGGAAGATCCAGAGGGTACGTCCGGGGTAGGAGTGAAATCCCGTAATCCTGGACGGACCACCGGTGGCGAAAGCGCTCTGGAAGGACGGATCCGACGGTGAGGGACGAAAGCTCGGGTCA |
| p__Halobacteriota_o__Halanaerobiales_ASV_1898 | 0 | 0 | 0 | 0 | 1 | 0 | 0 | 0 | 0 | 0 | 1 | CCTTTACACTGCACGACAGTGCGATAAGGGGACTCCGAGTGCGAGGGCATATAGTCCTCGCTTTTGTCGACCGTAAGGTGGTCGACGAATAAGGACTGGGCAAGACCGGTGCCAGCCGCCGCGGTAATACCGGCAGTCCGAGTGATGGTCGATATTATTGGGCCTAAAGCGTCCGTAGCTGGCCGAGCAAGTCTGTCGGGAAATCCAGTAGCTCAACTACTGGACGTCCGGTGGAAACTGTTCGGCTTGGGACCGGAAGATCCAAGGGGTACGTCCGGGGTAGGAGTGAAATCCTGTAATCCTGGACGGACCACCGATGGCGAAAGCACCTTGGAAGCACGGATCCGACAGTGAGGGACGAAAGCTAGGGTCT |
| p__Thermoplasmatota_p__Thermoplasmatota_ASV_1905 | 0 | 1 | 0 | 0 | 0 | 1 | 0 | 0 | 0 | 0 | 0 | ACTGCACAATGCTGGAAACAGCGATGCGGGGATTCCAAGTGCCGGCACTGAGTGTCGGCTGTTCCATTGTCTAAAAAACATAGGAAGTAAGGGCTGGGTAAGACGGGTGCCAGCCGCCGCGGTAATACCTGCAGCCCGAGTGGTGGTCGTTATTATTGGGCCTAAAACGTCCGTAGCCTGCTCGGTAAATGCCTGTGTAAATCGGACGGCTTAACCGTCCGAACTGCGGGTAGACTGCCGGGCTTGGGACCGGGAGAGGTGTGAAGTACTCCGGGGGTAGAGGTGAAATTCTGTAATCCTCGGGGGACTACCGGTGGCGAAGGCGTCACACTAGAACGGCTCCGACGGTGAGGGACGAAGCCCTGGGGCG |
| p__Nanoarchaeota_g__BMS3ABIN17_ASV_1947 | 1 | 1 | 1 | 0 | 0 | 0 | 0 | 0 | 0 | 0 | 0 | ATTCTGCAATGCGCGCAAGCGTGACAGAGCAAGTCAGAGTGCTTTACCATTAGGTAAAGCTTTTGCCAGATGTAAAAAGTCTGGCGAATAAGGACTGGGTAAGACTGGTGCCAGCCGCCGCGGTAATACCAGCAGTCCAAGTGGCAGCCGTTTTTGTTGGGTCTAAAACATCCGTAGCTTGTCACGTAAGTCTTTTGTGAAATCCGGCATCTCAAGTGTCGGGCGTGCAAAAGATACTGCGTGGCTAGAGACCGGGAGACGTAAGAAGTACGCAAGGGGTAGCGGTAAAATGCGTTGATCCTCTGTGGACTAACGATAGCGAAGGCATCTTACGAGAACGGGTCTGACAGTGAGGGATGAAGGCTAGGGGCG |
| p__Nanoarchaeota_g__CG1-02-57-44_ASV_1946 | 1 | 0 | 0 | 0 | 0 | 0 | 0 | 0 | 0 | 0 | 1 | CCTTTACAATGCACGTAAGTGTGATAAGGGAACCCTTAGTGCTTTGCTTATGCAAGGCTTTTGCCAATTTTAAAATGATTGGCGAATAAGTGGTGGGTAAGACGGGTGCCAGCCGCCGCGGTAACACCTGCGCCACAAGTGGCAACCGCTATTATTGGGCCTAAAGCGTCCGTAGCCTGTCTTGTAAATTCTTTGTGAAATCGTCATGCTTAACGTGACGGCGTGCAGGGAAGACTGCAAGGCTTGGAACCGGGAGGAGTTAGAGGAATGCCAGGGGGAGCGGTAAAATGCTATAATCCTTGGTAGACCACCTGTGGCGAAGGCGTCTAACTAGAACGGATTCGACGGTGAGGGACGAAAGCCAGGGGAG |
| p__Halobacteriota_g__Halobaculum_ASV_1959 | 0 | 0 | 0 | 0 | 1 | 0 | 0 | 0 | 0 | 0 | 1 | CCTTTACACTGCACGACAGTGCGATAAGGGGACCCCGAGTGCGAGGGCATACAGTCCTCGCTTTTCTGAACGGTAAGGCCGTTCAGGAACAAGAGCTGGGCAAGACCGGTGCCAGCCGCCGCGGTAATACCGGCAGCTCGAGTGATGACCGATCTTATTGGGCCTAAAGCGTCCGTAGCCGGCCGGGCAAGTCCGTCGGGAAATCCACCAGCTCAACTGGTGGGCGCCCGGCGGAAACTGCTCGGCTTGGGACAGGAAGACCCGAGGGGTACGTCCGGGGTAGGAGTGAAATCCCGTAATCCTGGACGGACCGCCGATGGCGAAAGCACCTCGGGAGGACTGATCCGACGGTGAGGGACGAAAGCTGGGGTCT |
| p__Halobacteriota_g__Halolamina_ASV_1988 | 0 | 0 | 0 | 0 | 0 | 1 | 0 | 0 | 0 | 0 | 0 | CCTTTACACTGCACGCCAGTGCGATAAGGGGACCCCGAGTGAAAGGGCATATCGCCCTTTCTTTTCTGAACCGTAGGGAGGTTCAGGAACAAGAGCTGGGCAAGACCGGTGCCAGCCGCCGCGGTAATACCGGCAGCTCGAGTGATGACCGATTTTATTGGGCCTAAAGCGTCCGTAGCTGGCCTCGCAAGTCCATCGGGAAATCCACCCGCTTAACGGGTGGGCGTCCGGTGGAAACTGTGTGGCTTGGGACAGGAAGGCTCGAGGGGTACGTTCGGGGTAGGAGTGAAATCCTGTAATCCTGAACGGACCGCCGATGGCGAAAGCACCTCGAGAGGACTGATCCGACAGTGAGGGACGAAAGCTGGGGTCT |
| p__Halobacteriota_g__Halomicrobium_3849_ASV_1989 | 0 | 0 | 0 | 0 | 0 | 1 | 0 | 0 | 1 | 0 | 0 | CCTTTACACTGCACGACAGTGCGATAAGGGGACTCCGAGTGCGAGGGCATATAGTCCTCGCTTTTCTGTACCGTAAGGTGGTACAGGAAGAAGTGCTGGGCAAGACCGGTGCCAGCCGCCGCGGTAATACCGGCAGCACGAGTGATGGCCGCTATTATTGGGCCTAAAGCGTCCGTAGCCGGCCGGACAAGTCCGTTGGGAAATCGACGCGCTCAACGCGTCGGCGTCCAGCGGAAACTGTCCGGCTTGGGGCCGGAAGACCTGAGGGGTACGTCCGGGGTAGGAGTGAAATCCTGTAATCCTGGACGGACCACCGGTGGCGAAAGCGCCTCAGGAAGACGGACCCGACGGTGAGGGACGAAAGCTAGGGTCT |
| p__Halobacteriota_g__Haloprofundus_ASV_1954 | 0 | 0 | 0 | 0 | 0 | 0 | 0 | 0 | 0 | 0 | 1 | CCTTTACACTGCACGACAGTGCGACAAGGGGACCCTGAGTGCGAGGGCATACAGTCCTCGCTTTTGTCGACCGTAAGAAGGTCGACGAATAAGAGCTGGGCAAGACCGGTGCCAGCCGCCGCGGTAATACCGGCAGCTCGAGTGATGACCGCTATTATTGGGCCTAAAGCGTCCGTAGCTGGCCGCGTAAGTCCATCGGGAAATCCATCCGCTCAACGGATGGGCGTCCGGTGGAAACTACACGGCTTGGAGCCGGAAGACCCAAGGGGTACGTCCGGGGTAGGAGTGAAATCCCGTAATCCTGGACGGACCACCGATGGCGAAAGCACCTTGGGAAGACGGACTCGACAGTGAGGGACGAAAGCTAGGGTCT |
| p__Halobacteriota_g__Halovenus_ASV_1964 | 0 | 1 | 0 | 0 | 0 | 0 | 0 | 0 | 0 | 0 | 0 | CCTTTACACTGCACGCCAGTGCGATAAGGGGACTCCGAGTGCGAGGGCATACAGTCCTCGCTTTTCTGCACCGTAAGGTGGTGCAGGAACAAGTGCTGGGCAAGACCGGTGCCAGCCGCCGCGGTAATACCGGCAGCACAAGTGATGGCCGATATTATTGGGCCTAAAGCGTCCGTAGCCGGCCGGACAAGTCTGTTGGGAAATCCACGTGCTTAACGCGTGGGCGTCCAGCGGAAACTGTCTGGCTTGGGGCCGGAAGACTCGAGGGGTACGTCCGGGGTAGGAGTGAAATCCCGTAATCCTGGACGGACCACCGGTGGCGAAAGCGCCTCGAGAGGACGGACCCGACGGTGAGGGACGAAAGCCAGGGTCT |
| p__Halobacteriota_g__Natronoarchaeum_ASV_1972 | 0 | 0 | 1 | 0 | 0 | 0 | 0 | 0 | 0 | 0 | 0 | CCTTTACACTGCACGCCAGTGCGATAAGGGGACCCCCAGTGCGAGGGCATACCGTCCTCGCTTTTCACCACCGTAAGGAGGTGCTGGAACAAGAGCTGGGCAAGACCGGTGCCAGCCGCCGCGGTAATACCGGCAGCTCGAGTGATGGCCGTTTTTATTGGGCCTAAAGCGTCCGTAGCCGGCCAGAAAAGTTCGTCGGGAAATCCACGTGCCCAACACGTGGGCGTCCGGCGAAACCTCTCTGGCTTGGGACCGGACGGCCCGAGGGGTACGGTCGGGGTAGGAGTGAAATCCTGTAATCCCGACCGGACCACCGGTGGCGAAAGCGCCTCGGGAGGACGGATCCGACGGTGAGGGACGAAAGCCAGGGTCA |
| p__Thermoproteota_g__PALSA-986_ASV_1999 | 0 | 0 | 0 | 0 | 0 | 0 | 0 | 0 | 1 | 0 | 0 | ACTTTGCAATACACGCAAGTGTGACAGGGCTATCCCGAGTGCCATCCGCTGAGGAAGGCTTTTACCCAGTCTAGAACGCTGGGGGAATAAGGAGAGGGCAAGTCTGGTGTCAGCCGCCGCGGTAATACCAGCTCTCCGAGTGGTGTGGATGTTTATTGGGCCTAAAGCATCCGTAGCTGGCTAAGTTAGTCCCTTGTTAAATCCACCGAATTAATCGTTGGATTGCGGGGGATACTGCTTGGCTAGGGGACGAGAGAGGCAGACGGTATTTTTGGGGTAGGGGTGAAATCCTATAATCCCAGGAAGACCACCAGTGGCGAAGGCTGTCTGCTAGAACGCGCCCGACGGTGAGGGATGAAAGCTGGGGGAG |
| p__Thermoplasmatota_g__SM1-50_ASV_1943 | 1 | 0 | 0 | 0 | 0 | 0 | 0 | 0 | 0 | 0 | 1 | ACTTCACACTGTGCGAAAGCGCGATGAGGGAATCCCAAGTGTATGCACGTAAGTGTATACTGTTTCTTTGTTTAAAATGCAGAGAGAGTAAGGGCTGGGTAAGACGGGTGCCAGCCGCCGCGGTAATACCTGCAGCTCAAGTGGTGACCACTATTATTGGGCCTAAAACGTTCGTAGCCGGTTTATTAAATGCCTGGGTAAATCGTGTCGCTTAACGATACGAATTCTGGGTAGACTGATAGACTTGGGACCGGGAGAGGCTAGAGGTACTCCTGGGGTAGAGGTGAAATTCTGTAATCCTGGGGGGACCACCAGTGGCGAAGGCGTCTAGCTAGAACGGGTCCGACGGTGAGGGACGAAGCCCTGGGGCG |
| p__Halobacteriota_f__Haloarculaceae_3992_ASV_1945 | 1 | 0 | 1 | 0 | 0 | 0 | 0 | 0 | 0 | 0 | 0 | CCTTTACACTGCACGCCAGTGCGATAGGGGGACTCCGAGTGCGAGGGCATACAGTCCTCGCTTTTCTGCACCGTAAGGAGGTGCAAGAATAAGGGCTGGGCAAGACCGGTGCCAGCCGCCGCGGTAATACCGGCAGCCCGAGTGATGGCCGATATTATTGGGCCTAAAGCGTCCGTAGCCGGCCGGACAAGTCCGTTGGGAAATCCACGCGCCCAACGCGTGGGCGTCCAGCGGAAACTGTCCGGCTTGGGGCCGGAGGACTCGAGGGGTACGTCCGGGGTAGGAGTGAAATCCCGTAATCCTGGACGGACCACCGGTGGCGAAAGCGCCTCGAGAGGACGGACCCGACGGTGAGGGACGAAAGCCAGGGTCT |
| p__Halobacteriota_f__Haloferacaceae_ASV_1956 | 0 | 0 | 0 | 0 | 0 | 0 | 0 | 0 | 0 | 0 | 1 | CCTTTACACTGCACGACAGTGCGATAAGGGGACTCCGAGTGCGAGGGCATATCGTCCTCGCTTTTCTGGACCGTAGGGAGGTCCAGGAACAAGAGCTGGGCAAGACCGGTGCCAGCCGCCGCGGTAATACCGGCAGCTCAAGTGATGGCCGATTTTATTGGGCCTAAAGCGTCCGTAGCTGGCCAGACAAGTCCGTCGGGAAATCCACCTGCCCAACAGGTGGGCGTCCGGCGGAAACTCTCTGGCTTGGGACCGGAAGACCCGAGGGGTACGTCCGGGGTAGGAGTGAAATCCCGTAATCCTGGACGGACCGCCGATAGCGAAAGCACCTCGGGAAGACGGATCCGACAGTGAGGGACGAAAGCTGGGGTCT |
| p__Halobacteriota_f__Haloferacaceae_ASV_1971 | 0 | 0 | 1 | 0 | 0 | 0 | 0 | 0 | 0 | 0 | 0 | CCTTTACACTGCACGCAAGTGCGATAAGGGGACTCCGAGTGCGAGGGCATACAGTCCTCGCTTTTGTGTACCGTAGGGCGGTACACGAATAAGAGCTGGGCAAGACCGGTGCCAGCCGCCGCGGTAATACCGGCAGCTCAAGTGATGGCCACTCTTATTGGGCCTAAAGCGTCCGTAGCTGGCCGAGAAAGTCCGTCGGGAAATCCACCTGCTCAACAGGTGGGCGTCCGGCGGAAACTTCTCGGCTTGGGACCGGAAGGCTCGAGGGGTACGTCTGGGGTAGGAGTGAAATCCTGTAATCCTGGACGGACCACCGATGGCGAAAGCACCTCGAGAAGACGGATCCGACAGTGAGGGACGAAAGCTAGGGTCT |
| p__Halobacteriota_f__Natrialbaceae_ASV_1953 | 0 | 1 | 0 | 0 | 0 | 0 | 0 | 0 | 0 | 1 | 0 | ACTTTACACTGCACGACAGTGCGATAAGGGAACTCCGAGTGCGAGGGCATATCGTCCTCGCTTTTCTGTACCGTAGGGAGGTACAGGAATAAGAGCTGGGCAAGACCGGTGCCAGCCGCCGCGGTAACACCGGCAGCTCGAGTGATGGCCGATATTATTGGGCCTAAAGCGTCCGTAGCCAGCTGAACAGGTCCGTCGGGAAATCTGCTCGCTCAACGAGCAGGCGTCCGGCGGAAACCAGTCAGCTTGGGACCGGAAGACCCGAGGGGTACGTCCGGGGTAGGAGTGAAATCCCGTAATCCTGGACGGACCACCGGTGGCGAAAGCGCCTCGGGAAGACGGATCCGACGGTGAGGGACGAAAGCTAGGGTCA |
| p__Halobacteriota_o__Halanaerobiales_ASV_1969 | 0 | 1 | 1 | 0 | 0 | 0 | 0 | 0 | 0 | 0 | 0 | CCTTTACACTGCACGAAAGTGCGATAGGGGGACTCCAAGTGCGAGGGCATACAGTCCTCGCTTTTGTGTACCGTAAGGTGGTACACGAATAAGAGCTGGGCAAGACCGGTGCCAGCCGCCGCGGTAATACCGGCAGCTCGAGTGATGACCGCTATTATTGGGCCTAAAGCGTCCGTAGCCGGCCGAACAGGTCCGTTGGGAAATCCACTGGCTCAACCAGTGGGCGTCCAGCGGAAACCAGTCGGCTTGGGGCCGGAAGACCTGAGGGGTACGTCTGGGGTAGGAGTGAAATCCTGTAATCCTGGACGGACCGCCGGTGGCGAAAGCGCCTCAGGAGGACGGACCCGACGGTGAGGGACGAAAGCTAGGGTCA |
| p__Halobacteriota_g__Halorhabdus_ASV_2026 | 0 | 0 | 1 | 0 | 0 | 0 | 0 | 0 | 0 | 0 | 0 | CCTTTACACTGCACGAAAGTGCGATAGGGGGACTCCGAGTGCGAGGGCATATAGTCCTCGCTTTTCTGTACCGTAAGGTGGTACAAGAATAAGGGCTGGGCAAGACCGGTGCCAGCCGCCGCGGTAATACCGGCAGCCCGAGTGATGGCCGCTATTATTGGGCCTAAAGCGTCCGTAGCCGGCCAGACAAGTCCGTTGGGAAATCGACGCGCTCAACGCGTCGGCGTCCGGCGGAAACTGTCTGGCTTGGGGCCGGAAGACTCGAGGGGTACGTCCGGGGTAGGAGTGAAATCCCGTAATCCTGGACGGACCACCGGTGGCGAAAGCGCCTCGAGAAAACGGACCCGACGGTGAGGGACGAAAGCTAGGGTCT |

| Phylum_Genus_ASV_number | Presence (1) or Absence (0) in each layer | | | | | | | | | | | |
|---|---|---|---|---|---|---|---|---|---|---|---|---|
| | L1 | L2 | L3 | L4 | L5 | L6 | L7 | L8 | L9 | L10 | L11 | Sequence |
| p__Halobacteriota_g__Halorubrum_ASV_2022 | 0 | 1 | 0 | 0 | 0 | 0 | 0 | 0 | 0 | 0 | 0 | CCTTTACACTGCACGACAGTGCGATAGGGGGATCCCAAGTGCACAGGCATAGTGCCTGTGCTTTTCGACACCCTAAGGCGGTGTCAGAATAAGGGCTGGGCAAGACCGGTGCCAGCCGCCGCGGTAATACCGGCAGCCCAAGTGATGGCCGAT<br>CTTATTGGGCCTAAAGCGTCCGTAGCTGGCCGCGCAAGTCCATCGGGAAATCCACCTGCTCAACAGGTGGACGCCCGGTAGAAACTGCGTGGCTTGGGACCGGAAGGCGCGACGGGTACGTCCGGGGTAGGAGTGAAATCCTGTAATCCTGGA<br>CGGACCACCGATGGCGAAAGCACGTCGCGAGGACGGATCCGACAGTGAGGGACGAAAGCCAGGGTCT |
| p__Halobacteriota_g__Halorussus_ASV_2003 | 1 | 0 | 0 | 0 | 0 | 0 | 0 | 0 | 0 | 0 | 0 | CCTTTACACTGCACGACAGTGCGATAAGGGGACTCCGAGTGCGAGGGCATATAGTCCTCGCTTTTGTACACCTTAAGGCGGTGTACGAATAAGGGCTGGGCAAGACCGGTGCCAGCCGCCGCGGTAATACCGGCAGCCCGAGTGATGGCCGCT<br>ATTATTGGGCCTAAAGCGTCCGTAGCCGGCCAGACAGGTCCGTCGGGAAATCCACGCGCTCAACGCGTGGGCGTCCGGCGGAAACCAGCTGGCTTGGGGCCGGAAGACCTGAGGGGTACGTCCGGGGTAGGAGTGAAATCCCGTAATCCTGGA<br>CGGACCACCGGTGGCGAAAGCGCCTCAGGAAAACGGACCCGACGGTGAGGGACGAAAGCTAGGGTCA |
| p__Halobacteriota_g__Halorussus_ASV_2063 | 0 | 0 | 0 | 0 | 0 | 0 | 0 | 0 | 1 | 0 | 0 | ACTTTACACTGCACGACAGTGCGATAAGGGGACTCCGAGTGCGAGGGCATATAGTCCTCGCTTTTCTGTACCGTAAGGTGGTACAGGAATAAGGACTGGGCAAGACCGGTGCCAGCCGCCGCGGTAATACCGGCAGTCCGAGTGATGGCCGCT<br>ATTATTGGGCCTAAAGCGTCCGTAGCCGGCCAGACAGGTCCGTCGGGAAATCCACGCGCTCAACGCGTGGGCGTCCGGCGGAAACCAGCTGGCTTGGGGCCGGAAGACTCGAGGGGTACGTCCGGGGTAGGAGTGAAATCCCGTAATCCTGGA<br>CGGACCACCGGTGGCGAAAGCGCCTCGAGAAGACGGACCCGACGGTGAGGGACGAAAGCTAGGGTCA |
| p__Halobacteriota_g__Halovenus_ASV_2017 | 0 | 0 | 0 | 0 | 0 | 0 | 0 | 0 | 0 | 0 | 1 | CCTTTACACTGCACGACAGTGCGATAGGGGGATTCCGAGTGCGAGGGCATACAGTCCTCGCTTTTCTGTACCGTAAGGAGGTACAGGAACAAGTGCTGGGCAAGACCGGTGCCAGCCGCCGCGGTAATACCGGCAGCACAAGTGATGGCCGAT<br>CTTATTGGGCCTAAAGCGTCCGTAGCTCGCCGGACAAGTCCGTTGGGAAATCCACGTGCCCAACACGTGGGCGTCCAGCGGAAACTGTCCGGCTTGGGGCCGGAGGACTCGAGGGGTACGTCCGGGGTAGGAGTGAAATCCCGTAATCCTGGA<br>CGGACCACCGGTGGCGAAAGCGCCTCGAGAGGACGGACCCGACAGTGAGGGACGAAAGCCAGGGTCT |
| p__Halobacteriota_g__Saliphagus_ASV_2023 | 0 | 1 | 0 | 0 | 0 | 0 | 0 | 0 | 0 | 0 | 0 | CCTTTACACTGCACGCGAGTGCGATAAGGGGACCCCGAGTGCGAGGGCATACAGTCCTCGCTTTTGTGTACCGTAAGGAGGTACACGAATAAGTGCTGGGCAAGACCGGTGCCAGCCGCCGCGGTAATACCGGCAGCACGAGTGATGACCGCT<br>ATTATTGGGCCTAAAGCGTCCGTAGCCGGCCAGACAGGTCCGTCGGGAAATCCGCACGCTCAACGTGCGGGCGTCCGGCGGAACCCAGCTGGCTAGGGACCGGAAGACCCGAGGGGTACGTCCGGGGTAGGAGTGAAATCCCGTAATCCTGGA<br>CGGACCGCCGGTGGCGAAAGCGCCTCGGGAGGACGGATCCGACGGTGAGGGACGAACGCTAGGGTCA |
| p__Halobacteriota_g__Saliphagus_ASV_2053 | 0 | 0 | 0 | 0 | 0 | 0 | 1 | 0 | 0 | 0 | 0 | CCTTTACACTGCACGAAAGTGCGATAAGGGGACCCCGAGTGCGAGGGCATACAGTCCTCGCTTTTGTGTACCGTAGGGAGGTACACGAATAAGGGCTGGGCAAGACCGGTGCCAGCCGCCGCGGTAATACCGGCAGCTCAAGTGATGGCCGCT<br>ATTATTGGGCCTAAAGCGTCCGTAGCCGGCCGCACAAGTCCGTCGGGAAATCCACCCGCCTAACGGGTGGGCGTCCGGCGGAAACTGTGCGGCTTGGGACCGGAAGACTCGAGGGGTACGTCCGGGGTAGGAGTGAAATCCTGTAATCCTGGA<br>CGGACCGCCGGTGGCGAAAGCGCCTCGAGAGGACGGATCCGACGGTGAGGGACGAAAGCTAGGGTCT |
| p__Halobacteriota_f__Haloferacaceae_ASV_2021 | 0 | 1 | 0 | 0 | 0 | 0 | 0 | 0 | 0 | 0 | 0 | CCTTTACACTGCACGCAAGTGCGATAAGGGGACCCCGAGTGCGAGGGCATACCGTCCTCGCTTTTGTCGACCGTAGGGAGGTCGACGAATAAGGGCTGGGCAAGACCGGTGCCAGCCGCCGCGGTAATACCGGCAGCCCAAGTGATGGCCGCT<br>CTTATTGGGCCTAAAGCGTCCGTAGCTGGCCGTGTAAGTTCATCGGGAAATCCACTCGCTCAACGAGTGGGCGTCCGGTGAAAACTGCACGGCTTGGGACCGGAAGACCCAGGGGGTACGTCCGGGGTAGGAGTGAAATCCTGTAATCCCGGA<br>CGGACCGCCGATGGCGAAAGCACTCTGGGAAGACGGATCCGACAGTGAGGGACGAAAGCTGGGGTCT |
| p__Halobacteriota_o__Halanaerobiales_ASV_2027 | 0 | 0 | 1 | 0 | 0 | 0 | 0 | 0 | 0 | 0 | 0 | CCTTTACACTGCACGAACGTGCGATAAGGGGACTCCGAGTGCGAGGGCATATCGTCCTCGCTTTTCTGTACCGTAAGGTGGTACAGGAAGAAGGGCTGGGCAAGACCGGTGCCAGCCGCCGCGGTAATACCGGCAGCCCGAGTGATGGCCGCT<br>ATTATTGGGCCTAAAGCGTCCGTAGCCGGCCGGGCAAGTCCGTTGGGAAATCCACCCGCTTAACGGGTGGGCGTCCAGCGGAAACTGCTCGGCTTGGGACCGGAAGACCCGAGGGGTACGTCTGGGGTAGGAGTGAAATCCTGTAATCCTGGA<br>CGGACCGCCGATGCCGAAAGGACCTCGGGAGGACGGATCCGACGGTGAGGGACGAAAGCTTGGATCT |
| p__Halobacteriota_f__Haloferacaceae_ASV_2129 | 0 | 0 | 0 | 0 | 0 | 0 | 0 | 1 | 0 | 0 | 0 | CCTTTACACTGCACGAGAGTGCGATAGGGGGATCCCAAGTGCGAGGGCATATTAGCCCTCGCTTTTGTGAACCGTAAGGCGGTTCACGAATAAGAGCTGGGCAAGACCGGTGCCAGCCGCCGCGGTAATACCGGCAGCTCGAGTGATGACCGA<br>TTTTATTGGGCCTAAAGCGTCCGTAGCTGGCCACACAAGTCCGTCGGGAAATCTACCTGCCCAACAGGTAGGCGTCCGGCGGAAACTGTGTGGCTTGGGACCGGAAGGCGTGAAGGGTACGTCCGGGGTAGGAGTGAAATCCTGTAATCCTGA<br>ACGGACCACCGATGGCGAAAGCACTTCACGAGGACGGATCCGACAGTGAGGGACGAAAGCTAGGGTCT |
| p__Nanoarchaeota_g__BMS3ABIN17_ASV_2136 | 0 | 0 | 0 | 0 | 0 | 0 | 0 | 0 | 1 | 0 | 0 | ATTCTGCAATGCGCGCAAGCGTGACAGATTGAACCAGAGTGCTTTTCAATTTGAAAAGCTTTTGTGAAATGCAAAAAGTTTTACGAATAAGGACTGGGTAAGACTAGTGCCAGCCGCCGCGGTAATACTAGCGGTCCAAGTCGCAGCCACAAT<br>TATTGGGTCTAAAACATCCGTAGCTTGCTTGATAAGTTTCCTGTGAAATTCGGCTTCTTAAAAGCCGAGCTGGCGGGAAGTACTGTCAAGCTAGAGACTGGAAGACGTAAGAAGTATGTTCAAAGTAGTGGTAAAATACGTTAATCTTGAACA<br>GACTACCAATAGCGAAGGCATCTTACGAGGACAGTTCTGACAGTAAGGGATGAAGGCCAGGGGCG |
| p__Halobacteriota_g__Halobaculum_ASV_2104 | 0 | 0 | 1 | 0 | 0 | 0 | 0 | 0 | 0 | 0 | 0 | CCTTTACACTGCACGACAGTGCGATAGGGGGACTCCGAGTGCGGAGGCATAGTGCCTTCGCTTTTGTGCACCCTAAGGCGGTGCACGAATAAGAGCTGGGCAAGACCGGTGCCAGCCGCCGCGGTAACACCGGCAGCTCGAGTGATGACCGAT<br>CTTATTGGGCCTAAAGCGTTCGTAGCCGGCCGGGCAAGTCCGTCGGGAAATCTACCAGCTCAACTGGTAGGCGTCCGGCGGAAACTGCTCGGCTTGGGACAGGGAGACTCGAAGGGTACGTCCGGGGTAGGAGTGAAATCCCGTAATCCTGGA<br>CGGACCACCGATGGCGAAAGCACTTCGAGAGAACTGATCCGACGGTCAGGAACGAAAGCTGGGGTCT |
| p__Halobacteriota_g__Halohasta_ASV_2096 | 0 | 0 | 1 | 0 | 0 | 0 | 0 | 0 | 0 | 0 | 0 | CCTTTACACTGCACGGAAGTGCGATAGGGGAATCCCAAGTGCGAGGGCATACTAGCCCTCGCTTTTGTTGACTGTAAGGAGGTCAACGAATAAGAGCTGGGCAAGACCGGTGCCAGCCGCCGCGGTAATACCGGCAGCTCGAGTGATGTCCAA<br>TATTATTGGGCCTAAAGCGTCCGTAGCTGGCCAGCCAAGTTCGTCGGGAAATCCACCTGCCCAACAGGTGGGCGTCCGGCGAAAACTGTCTGGCTTGGAACCGGAAGGCTCAGAGGGTACGTCCGGGGTAGGAGTGAAATCCTGTAATCCCGG<br>ACGGACCACCGATGGCGAAAGCACTCTGAGAAGACGGATTCGACAGTGAGGGACGAAAGCTAGGGTCT |
| p__Halobacteriota_g__Halohasta_ASV_2110 | 0 | 0 | 0 | 1 | 0 | 0 | 0 | 0 | 0 | 0 | 0 | CCTTTACACTGCACGGAAGTGCGATAGGGGAATCCCAAGTGCGAGGGCATACTAGCCCTCGCTTTTGTGTACCCTAAGGCGGTACACGAATAAGAGCTGGGCAAGACCGGTGCCAGCCGCCGCGGTAATACCGGCAGCTCGAGTGATGTCCAC<br>TATTATTGGGCCTAAAGCGTCCGTAGCTGGCCGCACAAGTCCGTCGGGAAATCCACCTGCCCAACAGGTGGGCGTCCGGCGGAAACTGTGTGGCTTGGAACCGGAAGGCTCAGAGGGTACGTCCGGGGTAGGAGTGAAATCCCGTAATCCCGG<br>ACGGACCACCGATGGCGAAAGCACTCTGAGAAGACGGATTCGACAGTGAGGGACGAAAGCTAGGGTCT |
| p__Halobacteriota_g__Halomicrobium_3849_ASV_2133 | 0 | 0 | 0 | 0 | 0 | 0 | 0 | 0 | 1 | 0 | 0 | CCTTTACACTGCACGACAGTGCGATAGGGGGACTCCGAGTGCGAGGGCATATAGTCCTCGCTTTTCTGGATCGTAAGGAGATCCAGGAACAAGTGCTGGGCAAGACCGGTGCCAGCCGCCGCGGTAATACCGGCAGCACGAGTGATGGCCGCT<br>CTTATTGGGCCTAAAGCGTCCGTAGCCGGCCAGTCAAGTCCGTTGGGAAATCGGCGCGCCTAACGCGTCGGCGTCCAGCGGAAACTGACTGGCTTGGGGCCGGAAGACCTGAGGAATACGTCCGGGGTAGGAGTGAAATCCCGTAATCCTGGA<br>CGGACCACCGGTGGCGAAAGCGCCTCAGGAAGACGGACCCGACGGTGAGGGACGAAAGCTAGGGTCT |
| p__Halobacteriota_g__Halorubrum_ASV_2076 | 0 | 0 | 0 | 0 | 0 | 0 | 0 | 0 | 1 | 1 | 0 | CCTTTACACTGCACGACAGTGCGATAAGGGAATCCCAAGTGCGTAGGCATAGAGCCTACGCTTTTGTGTACCCTAGGGCGGTACACGAATAAGGGCTGGGCAAGACCGGTGCCAGCCGCCGCGGTAATACCGGCAGCCCAAGTGATGGCCGAT<br>TTTATTGGGCCTAAAGCGTCCGTAGCTGGCCGCACAAGTCCATCGGAAAATCCACCCGCTCAACGGGTGGGCGTCCGGTGGAAACTGTGTGGCTTGGGACCGGAAGGCGCGACGGGTACGTCCGGGGTAGGAGTGAAATCCTGTAATCCTGGA<br>CGGACCACCGATGGCGAAAGCACGTCGCGAGAACGGATCCGACAGTGAGGGACGAAAGCTAGGGTCT |
| p__Halobacteriota_g__Halorubrum_ASV_2098 | 0 | 0 | 1 | 0 | 0 | 0 | 0 | 0 | 0 | 0 | 0 | CCTTTACACTGCACGCCAGTGCGATAAGGGGATCCCAAGTGCGTAGGCATAGAGCCTACGCTTTTGTCCACCGTAGGGAGGTGGACGAATAAGGGCTGGGCAAGACCGGTGCCAGCCGCCGCGGTAATACCGGCAGCCCAAGTGATGGCCGAT<br>CTTATTGGGCCTAAAGCGTCCGTAGCTGGCCGTACAAGTCCATCGGGAAATCCACCCGCTCAACGGGTGGCCGTCCGGTGGAAACTGTATGGCTTGGGACCGGAAGGCGCGACGGGTACGTCCGGGGTAGGAGTGAAATCCCGTAATCCTGGA<br>CGGACCGCCGATGGCGAAAGCACGTCGCGAGAACGGATCCGACAGTGAGGGACGAAAGCCAGGGTCT |
| p__Halobacteriota_g__Halovenus_ASV_2093 | 0 | 0 | 1 | 0 | 0 | 0 | 0 | 0 | 0 | 0 | 0 | CCTTTACACTGCACGCCAGTGCGATAGGGGGACTCCGAGTGCGAGGGCATACAGTCCTCGCTTTTCTGTACCGTAAGGTGGTACAGGAACAAGTGCTGGGCAAGACCGGTGCCAGCCGCCGCGGTAATACCGGCAGCACGAGTGATGGCCGAT<br>CTTATTGGGCCTAAAGCGTCCGTAGCCGGCCGGACAAGTCCGTTGGGAAATCCACGTGCCCAACACGTGGGCGTCCAGCGGAAACTGTCCGGCTTGGGGCCGGAAGACTCGAGGGGTACGTCCGGGGTAGGAGTGAAATCCTGTAATCCTGGA<br>CGGACCACCGGTGGCGAAAGCGCCTCGAGAAGACGGACCCGACGGTGAGGGACGAAAGCCAGGGTCT |
| p__Halobacteriota_g__Natrinema_ASV_2092 | 0 | 0 | 1 | 0 | 0 | 0 | 0 | 0 | 0 | 0 | 0 | CCTTTACACTGCACGAAAGTGCGATAAGGGGACTCCAAGTGCGAGGGCATATAGTCCTCGCTTTTCACCACCGTAAGGTGGTGGTGGAATAAGTGCTGGGCAAGACCGGTGCCAGCCGCCGCGGTAATACCGGCAGCACGAGTGATGACCGCT<br>ATTATTGGGCCTAAAGCGTCCGTAGCTGGCCGCGCAAGTTCATCGGGAAATCCGCACGCTTAACGTGCGGGCGTCCGGTGGAAACTGCTCGGCTTGGGACCGGAAGACCAGAGGGGTACGTCCGGGGTAGGAGTGAAATCCCGTAATCCTGGA<br>CGGACCACCGGTGGCGAAAGCGCCTCTGGAAGACGGATCCGACGGTGAGGGACGAAAGCTCGGGTCA |
| p__Halobacteriota_g__Natronomonas_ASV_2120 | 0 | 0 | 0 | 0 | 1 | 0 | 0 | 0 | 1 | 0 | 0 | CCTTTACACTGCACGCCAGTGCGATAGGGGGACACCGAGTGCGAGGGCATAGAGCCCTCGCTTTTCTGTACCGTAAGGTGGTACAGGAACAAGGGCTGGGCAAGACCGGTGCCAGCCGCCGCGGTAACACCGGCAGCCCGAGTGATGGCCGAT<br>ATTATTGGGCCTAAAGCGTCCGTAGCTGGCCGGGCAAGTCCGTCGGGAAATCCACGCGCTTAACGCGTGGGCGTCCGGCGGAAACTGTTCGGCTTGGGACCGGAAGATCCGAGGGGTACGTCCGGGGTAGGAGTGAAATCCCGTAATCCTGGA<br>CGGACCACCGATGGCGAAAGCACCTCGGAAGGACGGATCCGACAGTGAGGGACGAAAGCTAGGGTCT |
| p__Halobacteriota_g__Saliphagus_ASV_2094 | 0 | 0 | 1 | 0 | 0 | 0 | 0 | 0 | 0 | 0 | 0 | CCTTTACACTGCACGACAGTGCGATAAGGGGACTCCAAGTGCGAGGGCATATAGTCCTCGCTTTTGTGTACCGTAAGGTGGTACACGAATAAGGGCTGGGCAAGACCGGTGCCAGCCGCCGCGGTAATACCGGCAGCCCGAGTGATGACCGCT<br>ATTATTGGGCCTAAAGCGTCCGTAGCCGGCCGTACAAGTCCGTCGGGAAATCCGCACGCCTAACGTGCGGGCGTCCGGCGGAAACTGTTCGGCTTGGGACCGGAAGACTCGAGGGGTACGTCTGGGGTAGGAGTGAAATCCTGTAATCCTGGA<br>CGGACCACCGGTGGCGAAAGCGCCTCGAGAGGACGGATCCGACGGTGAGGGACGAAAGCTAGGGTCA |
| p__Halobacteriota_g__Saliphagus_ASV_2100 | 0 | 0 | 1 | 0 | 0 | 0 | 0 | 0 | 0 | 0 | 0 | CCTTTACACTGCACGACAGTGCGATAAGGGGACTCCAAGTGCCAGGGCATATCGTCCTGGCTTTTGTGTACCGTAAGGTGGTACACGAATAAGGGCTGGGCAAGACCGGTGCCAGCCGCCGCGGTAATACCGGCAGCCCGAGTGATGACCGCT<br>ATTATTGGGCCTAAAGCGTCCGTAGCCGGCCGTACAAGTCCGTCGGGAAATCCGCACGCCTAACGTGCGGGCGTCCGGCGGAAACTGTTCGGCTTGGGACCGGAAGACTCGAGGGGTACGTCTGGGGTAGGAGTGAAATCCTGTAATCCTGGA<br>CGGACCACCGGTGGCGAAAGCGCCTCGAGAGGACGGATCCGACGGTGAGGGACGAAAGCTAGGGTCA |
| p__Halobacteriota_g__Saliphagus_ASV_2103 | 0 | 0 | 1 | 0 | 0 | 0 | 0 | 0 | 0 | 0 | 0 | CCTTTACACTGCACGAAAGTGCGATAGGGGGATCCCGAGTGCGAGGGCATACAGTCCTCGCTTTTGTGTACCGTAAGGTGGTACACGAATAAGGGCTGGGCAAGACCGGTGCCAGCCGCCGCGGTAATACCGGCAGCTCAAGTGATGACCGTT<br>ATTATTGGGCCTAAAGCGTCCGTAGCCGGCCGCACAAGTCCGTCGGGAAATCCACCCGCCTAACGGGTGGGCGTCCGGCGGAAACTGTGCGGCTTGGGACCGGAAGACTCGAGGGGTACGTCCGGGGTAGGAGTGAAATCCTGTAATCCTGGA<br>CGGACCGCCGGTGGCGAAAGCGCCTCGAGAGGACGGATCCGACGGTGAGGGACGAAAGCTAGGGTCT |
| p__Halobacteriota_f__Haloarculaceae_3992_ASV_2099 | 0 | 0 | 1 | 0 | 0 | 0 | 0 | 0 | 0 | 0 | 0 | CCTTTACACTGCACGCCAGTGCGATAAGGGGACTCCGAGTGCGAGGGCATATAGTCCTCGCTTTTCTGGACCGTAAGGTGGTCCAGGAACAAGTGCTGGGCAAGACCGGTGCCAGCCGCCGCGGTAATACCGGCAGCACGAGTGATGGCCGCT<br>ATTATTGGGCCTAAAGCGTCCGTAGCCGGCCAGACAAGTCCGTTGGGAAATCCACGCGCTTAACGCGTGGGCGTCCAGCGGAAACTGTCTGGCTTGGGGCCGGAAGACCTGAGGGGTACGTCCGGGGTAGGAGTGAAATCCCGTAATCCTGGA<br>CGGACCACCGGTGGCGAAAGCGCCTCAGGAAGACGGACCCGACGGTGAGGGACGAAAGCTAGGGTCT |
| p__Halobacteriota_o__Halanaerobiales_ASV_2105 | 0 | 0 | 1 | 0 | 0 | 0 | 0 | 0 | 0 | 0 | 0 | CCTTTACACTGCACGCAAGTGCGATAAGGGGACCCCGAGTGCGAGGGCATACCGTCCTCGCTTTTGTCGACCGTAGGGAGGTCGACGAATAAGGGCTGGGCAAGACCGGTGCCAGCCGCCGCGGTAATACCGGCAGCCCAAGTGATGGCCGCT<br>CTTATTGGGCCTAAAGCGTCCGTAGCTGGCCGCATTGGTCCGTCGGGAAATCCACTCGCTCAACGAGTGGGCGTCCGGCGGATACCATGCGGCTTGGAACCGGAAGACCCAGGGGGTACGTCCGGGGTAGGAGTGAAATCCTGTAATCCCGGA<br>CGGACCGCCGATGGCGAAAGCACTCTGGGAAGACGGATTCGACAGTGAGGGACGAAAGCTGGGGTCT |

| | Presence (1) or Absence (0) in each layer | | | | | | | | | | | |
|---|---|---|---|---|---|---|---|---|---|---|---|---|
| **Phylum_Genus_ASV_number** | **L1** | **L2** | **L3** | **L4** | **L5** | **L6** | **L7** | **L8** | **L9** | **L10** | **L11** | **Sequence** |
| p__Nanoarchaeota_g__BMS3ABIN17_ASV_2139 | 1 | 0 | 0 | 0 | 0 | 0 | 0 | 0 | 0 | 0 | 1 | CTTTTACAATGCACGCAAGTGTGATATAGCGAGCCAAAGTGCTTTCTGCACAAGAAAGCTTTTCTTTGCTGTAAAAAGGCAGAGGAATAAGGACTGGGAAAGACGAGTGCCAGCCGCCGCGGTAATACTCGCAGTCCAAGTTGCAGCCAACATTATTGGGTCTAAAACATCCGTAGCTTGTCTGTTAAGTTTCTTGTGAAATTCCGTGTCTCAAGCATGGAGCTTGCAAGAAATACTGACAGACTAGAGACCGGTAGACGTAAGGAGTACGCCCGAGGTAGTGGTAAAATACGTTAATCTTGGGTGGACTAACGATTGGCGAAGGCACCTTACGAGGACGGATCTGACAGTGAGGGATGAAGGCCAGGGGCG |
| p__Halobacteriota_g__Halalkalicoccus_ASV_2143 | 0 | 0 | 0 | 0 | 0 | 0 | 0 | 0 | 0 | 1 | 0 | CCTTTACACTGCACGAACGTGCGATAAGGGGACTCCAAGTGCGAGGGCATACAGTCCTCGCTTTTCCCTACCGTAAGGTGGTAGGTGAATAAGAGCTGGGCAAGACCGGTGCCAGCCGCCGCGGTAATACCGGCAGCTCGAGTGATGGCCAATATTATTGGGCCTAAAGCGTCCGTAGCTGGCCGAACAGGTCCGTCGGGAAATCCGCTCGCTCAACGAGCGGACGTCCGGCGGAAACCAGTCGGCTTGGGACCGGAAGACCCGAGGGGTACGGCCGGGGTAGGAGTGAAATCCTGTAATCCTAGACGGACCATCGATCGCGAAAGCACCTTGGGAGGACGGACCCGACGGTGAGGGACGAAAGCTAGGGTCA |
| p__Halobacteriota_g__Halanaeroarchaeum_ASV_2138 | 1 | 0 | 0 | 0 | 0 | 0 | 0 | 0 | 0 | 0 | 0 | CCTTTACACTGCACGACAGTGCGATAAGGGAACTCCGAGTGCGAGGGCATAGAGCCCTCGCTTTTGTACACCGTAAGGTGGTGTACGAATAAGGGCTGGGCAAGACCGGTGCCAGCCGCCGCGGTAATACCGGCAGCCCAAGTGATGGCCGATATTATTGGGCCTAAAGCGTCCGTAGCTGGCCGAACAAGTCCGTTGGGAAATCCGCGCGCTCAACGCGCGGGCGTCCAGCGGAAACTGTCCGGCTTGGGGCCGGAAGACCTGAGGGGTACGTCCGGGGTAGGAGTGAAATCCTGTAATCTCGGACGGACCGCCTGTGGCGAAAGCGCCTCAGGAAGACGGACCCGACAGTGAGGGACGAAAGCCAGGGTCT |
| p__Halobacteriota_g__Halobacterium_ASV_2181 | 0 | 0 | 1 | 1 | 0 | 0 | 0 | 0 | 0 | 0 | 0 | CCTTTACACTGCACGACAGTGCGATAGGGGGACTCCGAGTGCGAGGGCGTATAGTCCTCGCTATTCTGAACCGTAAGGTGGTTCAGGAAAAGGGCTGGGCAAGACCGGTGCCAGCCGCCGCGGTAATACCGGCAGCCCGAGTGATGGCCAATATTATTGGGCCTAAAGCGTCCGTAGCCGGCCGGACAAGTCCGTTGGGAAATCTGCTCGCTTAACGAGCAGGCGTCCAGCGGAAACTGTTCGGCTTGGGGCCGGAGGACTCGAGGGGTACGTCCGGGGTAGGAGTGAAATCCTGTAATCCTGGACGGACCGCCGATGGCGAAAGCACCTCGAGAGGACGGACCCGACGGTGAGGGACGAAAGCTGGGGTCT |
| p__Halobacteriota_g__Halomarina_3560_ASV_2165 | 0 | 1 | 0 | 0 | 0 | 0 | 0 | 0 | 0 | 0 | 0 | CCTTTACACTGCACGACAGTGCGATAGGGGGATTCCGAGTGCGAGGGCATACAGTCCTCGCTTTTCTGTACCGTAAGGTGGTGCAGGAATAAGGGCTGGGCAAGACCGGTGCCAGCCGCCGCGGTAATACCGGCAGCTCGAGTGATGACCGATTTTATTGGGCCTAAAGCGTCCGTAGCCGGCCGGGTAGGTCCGTCGGGAAATCCACTCGCCTAACGAGTGGGCGTCCGGCGGATACCGCCTGGCTTGGGACCGGGAGACCAGAGGGGTACGTCCGGGGTAGGAGTGAAATCCTGTAATCCTGGACGGACCATCGGTGGCGAAAGCGCCTCTGGAGAACGGATCCGACGGTGAGGGACGAAAGCTAGGGTCT |
| p__Halobacteriota_g__Halomarina_3560_ASV_2206 | 0 | 0 | 0 | 0 | 0 | 0 | 0 | 1 | 0 | 0 | 0 | CCTTTACACTGCACGACAGTGCGATAGGGGGATTCCGAGTGCGAGGGCATACAGTCCTCGCTTTTGTGTACCGTAAGGTGGTACACGAATAAGAGCTGGGCAAGACCGGTGCCAGCCGCCGCGGTAATACCGGCAGCTCGAGTGATGGCCGATTTTATTGGGCCTAAAGCGTCCGTAGCCGGCCAGTCAGGTCCGTCGGGAAATCCACTCGCTCAACGAGTGGGCGTCCGGCGGAAACCGTCTGGCTTGGGACCGGGAGATCTGAGGGGTACGTCCGGGGTAGGAGTGAAATCCTGTAATCCTAGACGGACCATCGGTGGCGAAAGCGCCTCAGAAGAACGGATCCGACGGTGAGGGACGAAAGCTAGGGTCT |
| p__Halobacteriota_g__Halorussus_ASV_2178 | 0 | 0 | 1 | 0 | 0 | 0 | 0 | 0 | 0 | 0 | 0 | ACTTTACACTGCACGACAGTGCGATAAGGGGACTCCGAGTGCGAGGGCATATAGTCCTCGCTTTTGTACACCGTAAGGTGGTGTACGAATAAGGGCTGGGCAAGACCGGTGCCAGCCGCCGCGGTAATACCGGCAGCCCGAGTGATGGCCGCTGTTATTGGGCCTAAAGCGTCCGTAGCCGGCCAGACAGGTCCGTCGGGAAATCCACTCGCTCAACGAGTGGGCGTCCGGCGGAAACCAGCTGGCTTGGGGCCGGAAGATCCGAGGGGTACGTCCGGGGTAGGAGTGAAATCCCGTAATCCTGGACGGACCGCCGGTGGCGAAAGCGCCTCGGAAAGACGGACCCGACGGTGAGGGACGAAAGCTAGGGTCA |
| p__Halobacteriota_g__Halovenus_ASV_2182 | 0 | 0 | 1 | 0 | 0 | 0 | 1 | 0 | 0 | 0 | 0 | CCTTTACACTGCACGACAGTGCGATAGGGGGACTCCGAGTGCGAGGGCATATAGTCCTCGCTTTTCTGTACCGTAAGGTGGTACAGGAACAAGTGCTGGGCAAGACCGGTGCCAGCCGCCGCGGTAATACCGGCAGCACGAGTGATGGCCGATTTTATTGGGCCTAAAGCGTCCGTAGCCGGCCAGTCAAGTTTGTTGGGAAATCCACGTGCTCAACACGTGGGCGTCCAGCGAAAACTGTCTGGCTTGGGGCCGGAAGACTCGAGGGGTACGTCCGGGGTAGGAGTGAAATCCCGTAATCCTGGACGGACCGCCGGTGGCGAAAGCGCCTCGAGAGGACGGACCCGACGGTGAGGGACGAAAGCCAGGGTCT |
| p__Halobacteriota_g__Natronomonas_ASV_2176 | 0 | 0 | 1 | 0 | 0 | 0 | 0 | 0 | 0 | 0 | 0 | CCTTTACACTGCACGACAGTGCGATAAGGGGACTCCGAGTGCGAGGGCATATCGTCCTCGCTTTTGTCGACCGTAAGGTGGTCGACGAATAAGGGCTGGGCAAGACCGGTGCCAGCCGCCGCGGTAATACCGGCAGCCCGAGTGATGGTCGATATTATTGGGCCTAAAGCGTCCGTAGCTGGCCGAGCAAGTCCGTCGGGAAATCCAGTGGCTCAACCACTGGACGTCCGGTGGAAACTGTTCGGCTTGGGACCGGAAGACTCGAGGGGTACGTCCGGGGTAGGAGTGAAATCCCGTAATCCTGGACGGACCACCGATGGCGAAAGCACCTCGAGAGGACGGATCCGACAGTGAGGGACGAAAGCTAGGGTCT |
| p__Micrarchaeota_g__UBA95_ASV_2152 | 0 | 0 | 0 | 0 | 0 | 0 | 0 | 0 | 1 | 1 | 1 | TTTCCGCAATGCGCGAAAGCGTGACGGGGGCAGTCTGAGTGCCATCCTAACGGATGGCTTTTCTCGAGTTTTAATTGCTCGGGGAATAAGTGGTGGGTAAAACTCGTGCCAGCCGCCGCGGTAATACGAGTAGCACTAGTGGTATCCACGAATATTGAGCTTAAAGCGCCCGTAGCCGGCTCGGTCAGTTCCTTGTGAAATCTGTCGACTTAATCGACAGGCTTGCAGGGAATACTACCGGGCTCGGGAGTGGGGGAGGCAAGAGGTACTCTAGGGGGAGCGGTAAAATGCTTTAATCCTTAGAGGACCAACAGTGGCGAAGGCGTCTTGCCAAAACACGTCCGACGGTGAGGGACGAAGGCTAGGGGAG |
| p__Halobacteriota_f__Haloarculaceae_3992_ASV_2162 | 0 | 1 | 0 | 0 | 0 | 0 | 0 | 0 | 0 | 0 | 0 | CCTTTACACTGCACGACAGTGCGATAAGGGGACTCCGAGTGCGAGGGCATATAGTCCTCGCTTTTCTGTACCGTAGGGAGGTACAGGAACAAGTGCTGGGCAAGACCGGTGCCAGCCGCCGCGGTAATACCGGCAGCACGAGTGATGACCGATATTATTGGGCCTAAAGCGTCCGTAGCTGGCCGGGCAAGTCCGTTGGGAAATCCACGTGCTCAACGCGTGGGCGTCCGGCGGAAACTGTTCGGCTTGGGGCCGGAAGGCCTGAGGGGTACGTCCGGGGTAGGAGTGAAATCCCGTAATCCTGGACGGACCACCAATGGGGAAACCACCTCAGGAGGACGGACCCGACAGTGAGGGACGAAAGCTAGGGTCT |
| p__Halobacteriota_f__Halobacteriaceae_4595_ASV_2168 | 0 | 1 | 0 | 0 | 0 | 0 | 0 | 0 | 0 | 0 | 0 | CCTTTACACTGCACGACAGTGCGATAGGGGGACTCCGAGTGCGAGGGCATATAGTCCTCGCTTTTCTGGACCGTAAGGAGGTCCAGGAACAAGGGCTGGGCAAGACCGGTGCCAGCCGCCGCGGTAATACCGGCAGCCCAAGTGATGGCCAATATTATTGGGCCTAAAGCGTCCGTAGCTGGCCGGACAAGTCCGTTGGGAAATCCACGCGCTCAACGCGTGGGCGTCCAGCGGAAACTGTCCGGCTCGGGACCGGAAGACCCAGAGGGTACGTCCGGGGTAGGAGTGAAATCCCGTAATCCTGGACGGACCACCGGTGGCGAAAGCGCTCTGGGAGGACGGATCCGACAGTGAGGGACGAAAGCTAGGGTCT |
| p__Halobacteriota_f__Haloferacaceae_ASV_2164 | 0 | 1 | 0 | 0 | 0 | 0 | 0 | 0 | 0 | 0 | 0 | CCTTTACACTGCACGCAAGTGCGATAAGGGGACCCCGAGTGCGAGGGCATACCGTCCTCGCTTTTGTCAACCGTAGGGAGGTTGACGAATAAGGGCTGGGCAAGACCGGTGCCAGCCGCCGCGGTAATACCGGCAGCCCAAGTGATGGCCGCTCTTATTGGGCCTAAAGCGTCCGTAGCTGGCCGCATTGGTCCGTCGGGAAATCCACTCGCTCAACGAGTGGGCGTCCGGCGGATACCATGCGGCTTGGGACCGGAAGACCCAGGGGGTACGTCCGGGGTAGGAGTGAAATCCTGTAATCCCGGACGGACCGCCGATGGCGAAAGCACTCTGGGAAGACGGCTCCGACAGTGAGGGACGAAAGCTGGGGTCT |
| p__Halobacteriota_f__Haloferacaceae_ASV_2197 | 0 | 0 | 0 | 0 | 0 | 1 | 0 | 0 | 0 | 0 | 0 | CCTTTACACTGCACGCCAGTGCGATAGGGGGACCCCGAGTGCGAGGGCATACAGTCCTCGCTTTTGCACACCGTAAGGAGGTGTGCGAATAAGTGCTGGGCAAGACCGGTGCCAGCCGCCGCGGTAATACCGGCAGCACAAGTGATGGCCGCTTTTATTGGGCCTAAAGCGTCCGTAGCCGGCCGAACAAGTCCGTCGGGAAATCCATCCGCTCAACGGATGGGCGTCCGGCGGAAACTGTTCGGCTTGGGACAGGAAGACTCGAGGGGTACGTCCGGGGTAGGAGTGAAATCCTGTAATCCTGGACGGACCGCCGATGGCGAAAGCACCTCGAGAAGACTGATCCGACGGTGAGGGACGAAAGCTGGGGTCT |
| p__Halobacteriota_o__Halanaerobiales_ASV_2174 | 0 | 0 | 1 | 0 | 0 | 0 | 0 | 0 | 0 | 0 | 0 | CCTTTACACTGCACGCCAGTGCGATAGGGGGACCCCGAGTGCGAGGGCATACAGTCCTCGCTTTTGTACACCGTAAGGAGGTGTACGAATAAGTGCTGGGCAAGACCGGTGCCAGCCGCCGCGGTAATACCGGCAGCACAAGTGATGGCCGCTATTATTGGGCCTAAAGCGTCCGTAGCCGGCCGAACAAGTCCGTCGGGAAATCCATCCGCTCAACGGATGGGCGTCCGGCGGAAACTGTTCGGCTTGGGACAGGAAGACTCGAGGGGTACGTCCGGGGTAGGAGTGAAATCCCGTAATCCTGGACGGACCGCCGATGGCGAAAGCACCTCGAGAAGACTGATCCGACGGTGAGGGACGAAAGCTGGGGTCT |
| p__Nanoarchaeota_g__BMS3ABIN17_ASV_2215 | 1 | 0 | 0 | 0 | 0 | 0 | 0 | 0 | 0 | 0 | 1 | AATCTACAATGCGCGCAAGCGTGATAGTTTGAGCCAGAGTGCGTTTCAATTGAAATGCTTTTGCCGAATGTAAAAAGTTCGGCGAATAAGGACTGGGTAAGACTAGTGCCAGCCGCCGCGGTAATACTAGCGGTCCAAGTCGCAGCCATTTTTATTGGGTCTAAAACATCCGTAGCTTGCTTGATAAGTTTCTTGTGAAATATTATGTCTTAAGCATAATGCGTGCAAGAAGTACTGTTAAGCTAGAGACAGGGAGACGTGAAGAGTACGTTTAAAGTAGCGGTAAAATGCTTTAATCTTAAACGGACTAACAATAGCGAAGGCACTTCACGAGAACTGTTCTGACAGTCAGGGATGAAGGCTAGGGGCG |
| p__Halobacteriota_g__Halobacterium_ASV_2241 | 0 | 1 | 0 | 0 | 0 | 0 | 0 | 0 | 0 | 0 | 0 | CCTTTACACTGTACGCAAGTGCGATAGGGGGACTCCGAGTGCGAGGGCATAGAGCCCTCGCTTTTGTACACCGTAAGGTGGTGTACGAATAAGGGCTGGGCAAGACCGGTGCCAGCCGCCGCGGTAATACCGGCAGCCCGAGTGATGGCCGATATTATTGGGCCTAAAGCGTCCGTAGCTGGCCAGGCAAGTCCGTTGGGAAATCTGCTCGCTCAACGAGCAGGCGTCCAGCGGAAACTGCTTGGCTTGGGACCGGAGGACCTGAGGGGTACGTCCGGGGTAGGAGTGAAATCCTGTAATCCTGGACGGACCGCCGGTGGCGAAAGCGCCTCAGGAAGACGGATCCGACAGTGAGGGACGAAAGCTAGGGTCT |
| p__Halobacteriota_g__Halobellus_3029_ASV_2254 | 0 | 0 | 1 | 0 | 0 | 0 | 0 | 0 | 0 | 0 | 0 | CCTTTACACTGCACGCCAGTGCGATAAGGGGACTCCAAGTGCGAGGGCATATCGTCCTCGCTATCCTGTACTGTAGGGTAGTACAGTATCAAGAGCTGGGCAAGACCGGTGCCAGCCGCCGCGGTAATACCGGCAGCTCAAGTGATGGCCGATCTTATTGGGCCTAAAGCGTCCGTAGCTGGCCCTGCAAGTCCGTCGGGAAATCCATCAGCCCAACTGATGGGCGTCCGGCGGAAACTCCAGGGCTTGGGACCGGAAGGCTCGAGGGGTACGTCCGGGGTAGGAGTGAAATCCCGTAATCCCGAACGGACCACCGATGGCGAAAGCACCTCGAGAAGACGGCTCCGACAGTGAGGGACGAAAGCTAGGGTCT |
| p__Halobacteriota_g__Halohasta_ASV_2267 | 0 | 0 | 0 | 1 | 0 | 0 | 0 | 0 | 0 | 0 | 0 | CCTTTACACTGCACGCAAGTGCGATAGGGGAATCCCAAGTGCGAGGGCATACTAGCCCTCGCTTTTGTGTACTGTAAGGAGGTACACGAATAAGAGCTGGGCAAGACCGGTGCCAGCCGCCGCGGTAATACCGGCAGCTCGAGTGATGTCCAATTTTATTGGGCCTAAAGCGTCCGTAGCTGGCCAGCCAAGTCCGTCGGGAAATCCACCTGCCCAACAGGTGGGCGTCCGGCGGAAACTGTCTGGCTTGGAACCGGAAGGCTCAGAGGGTACGTCCGGGGTAGGAGTGAAATCCCGTAATCCCGGACGGACCACCGATGGCGAAAGCACTCTGAGAAGACGGATTCGACAGTGAGGGACGAAAGCTAGGGTCT |
| p__Halobacteriota_g__Halopelagius_ASV_2236 | 0 | 1 | 0 | 0 | 0 | 0 | 0 | 0 | 0 | 0 | 0 | CCTTTACACTGCACGCAAGTGCGATAAGGGGACTCCGAGTGCGAGGGCATATAGTCCTCGCTTTTCTGTACCGTAGGAAGGTACACGAATAAGAGCTGGGCAAGACCGGTGCCAGCCGCCGCGGTAATACCGGCAGCTCGAGTGATGACCGATTTTATTGGGCCTAAAGCGTCCGTAGCTGGCCAGACAAGTCCGTCGGGAAATCCATCTGCTCAACAGATGGGCGTCCGGCGGAAACTCTCTGGCTTGGGACCGGAAGGCTCGAAGGGTACGTCTGGGGTAGGAGTGAAATCCTGTAATCCTGGACGGACCACCGATGGCGAAAGCACTTCGAGAAGACGGATCCGACAGTGAGGGACGAAAGCTAGGGTCT |
| p__Halobacteriota_g__Haloprofundus_ASV_2224 | 0 | 0 | 0 | 0 | 0 | 0 | 0 | 0 | 0 | 0 | 1 | CCTTTACACTGCACGACAGTGCGACAAGGGGACCCCGAGTGCGAGGGCATACAGTCCTCGCTTTTGTGAACCGTAAGGAGGTTCACGAATAAGAGCTGGGCAAGACCGGTGCCAGCCGCCGCGGTAATACCGGCAGCTCGAGTGATGACCGCTGTTATTGGGCCTAAAGCGTCCGTAGCTGGCCGCGTAAGTCCATCGGGAAATCCATCCGCTCAACGGATGGGCGTCCGGTGGAAACTACACGGCTTGGAGCCGGAAGACCCAAGGGGTACGTCCGGGGTAGGAGTGAAATCCCGTAATCCTGGACGGACCACCGATGGCGAAAGCACCTTGGGAAGACGGACTCGACAGTGAGGGACGAAAGCTAGGGTCT |
| p__Halobacteriota_g__Halorubrum_ASV_2228 | 0 | 1 | 0 | 0 | 0 | 0 | 0 | 0 | 0 | 0 | 0 | CCTTTACACTGCACGCAAGTGCGATAAGGGGATCCCAAGTGCGAGGGCATACAGTCCTCGCTTTTCACACCCGTAAGGAGGTTGTGGAATAAGGGCTGGGCAAGACCGGTGCCAGCCGCCGCGGTAATACCGGCAGCCCAAGTGATGACCGCTATTATTGGGCCTAAAGCGTCCGTAGCTGGCCGCGCAAGTCCATCGGGAAATCCACCTGCCCAACAGGTGGGCGACCGGTGGAAACTGCGCGGCTTGGGACCGGAAGGCTCAGAGGGTACGTCCGGGGTAGGAGTGAAATCCCGTAATCCCGGACGGACCGCCGATGGCGAAAGCACTCTGAGAAGACGGATCCGACAGTGAGGGACGAAAGCTGGGGTCT |
| p__Halobacteriota_g__Halorubrum_ASV_2294 | 0 | 0 | 0 | 0 | 0 | 0 | 1 | 0 | 0 | 0 | 0 | CCTTTACACTGCACGACAGTGCGATAGGGGGATCCCAAGTGCACAGGCATAGCGCCTGTGCTTTTCGGTACCGTAAGGTGGTACCAGAATAAGGGCTGGGCAAGACCGGTGCCAGCCGCCGCGGTAATACCGGCAGCCCAAGTGATGGCCGATCTTATTGGGCCTAAAGCGTCCGTAGCTGGCCGCGCAAGTCCATCGGGAAATCCACCTGCTCAACAGGTGGCCGCCCGGTGGAAACTGCGTGGCTTGGGACCGGAAGGCGCGACGGGTACGTCCGGGGTAGGAGTGAAATCCCGTAATCCTGGACGGACCACCGATGGCGAAAGCACGTCGCGAGAACGGATCCGACAGTGAGGGACGAAAGCCAGGGTCT |

| Phylum_Genus_ASV_number | Presence (1) or Absence (0) in each layer L1 | L2 | L3 | L4 | L5 | L6 | L7 | L8 | L9 | L10 | L11 | Sequence |
|---|---|---|---|---|---|---|---|---|---|---|---|---|
| p__Halobacteriota_g__Halorussus_ASV_2223 | 0 | 1 | 0 | 0 | 0 | 0 | 0 | 0 | 0 | 1 | 0 | CCTTTACACTGTACGAAAGTACGATAGGGGGACCCCGAGTGCGAGGGCATACAGTCCTCGCTTTTGTGTACCGTAAGGTGGTACACGAATAAGAGCTGGGCAAGACCGGTGCCAGCCGCCGCGGTAATACCGGCAGCTCAAGTGATGGCCGATATTATTGGGCCTAAAGCGTCCGTAGCCGGCCGAACAGGTCCGTTGGGAAATCCATCTGCCTAACAGATGGGCGTCCAGCGGAAACCAGTCGGCTTGGGGCCGGAAGATCCGAGGGGTACGTCCGGGGTAGGAGTGAAATCCTGTAATCCTGGACGGACCGCCGGTGGCGAAAGCGCCTCGGAAGGACGGACCCGACGGTGAGGGACGAAAGCTAGGGTCA |
| p__Halobacteriota_g__Halorussus_ASV_2250 | 0 | 1 | 1 | 0 | 0 | 0 | 0 | 0 | 0 | 0 | 0 | CCTTTACACTGCGCGAAAGCGCGATAAGGGGACCCCGAGTGCGAGGGCATACAGTCCTCGCTTTTCCCCACCGTAAGGTGGTGGGGGAATAAGGGCTGGGCAAGACCGGTGCCAGCCGCCGCGGTAATACCGGCAGCCCGAGTGATGGCCGCTATTATTGGGCCTAAAGCGTCCGTAGCCGGCCGGACAAGTCCGTCGGGAAATCCAGTGGCCCAACCACTGGGCGACCGGCGGAAACTGTCCGGCTTGGGGCCGGAAGACCCGAGGGGTACGTCCGGGGTAGGAGTGAAATCCTGTAATCCTGGACGGACCGCCGGTGGCGAAAGCGCCTCGGGAGGACGGACCCGACGGTGAGGGACGAAAGCTGGGGTCT |
| p__Nanoarchaeota_g__PWVO01_ASV_2214 | 1 | 0 | 0 | 0 | 0 | 0 | 0 | 0 | 0 | 0 | 1 | CCTTTACAATGCACGAAAGTGTGATAAGGGAATCCCGAGTGCCTTGCTTTCGCAAGGCTTTTCTTAAGAGTAAATATCTTAGGGAATAAGTGGTGGGAAAGACCGGTGCCAGCAGCCGCGGTAATACCGGCGCCACAAGTGGTGATCATTTTTATTGGGCCTAAAGCGTTCGTAGCCGGTCTGATAAATCTTCTGTGAAATTGTCAGGCTTAACCTGACAGCGTGCAGAAGACACTATCAGACTTGGGACCGGGAGGAGTCAGAGGTATTCCCTGGGGAGCGGTAAAATGCTATAATCCTTGGAAGACTAACTGTGGCGAAGGCGTCTGACCAGAACGGATCCGACGGTGATTGGACGAAAGTCAGGGGAG |
| p__Halobacteriota_g__Saliphagus_ASV_2281 | 0 | 0 | 0 | 0 | 1 | 0 | 0 | 0 | 0 | 0 | 0 | CCTTTACACTGCACGACAGTGCGATAAGGGGACTCCAAGTGCGAGGGCATATCGTCCTCGCTTTTCACTACCGTAAGGTGGTAGTGGAATAAGGGCTGGGCAAGACCGGTGCCAGCCGCCGCGGTAATACCGGCAGCCCGAGTGATGACCGCTATTATTGGGCCTAAAGCGTCCGTAGCCGGCCTCGCAAGTCCGTCGGGAAATCCGCACGCCTAACGTGCGGGCGTCCGGCGGAAACTGCGTGGCTTGGGACCGGAAGACTCGAGGGGTACGTCCGGGGTAGGAGTGAAATCCTGTAATCCTGGACGGACCACCGGTGGCGAAAGCGCCTCGAGAGGACGGATCCGACGGTGAGGGACGAAAGCTAGGGTCA |
| p__Asgardarchaeota_g__SDNM01_ASV_2219 | 0 | 0 | 0 | 0 | 0 | 0 | 0 | 0 | 0 | 1 | 0 | CCTCCACAATACACGAAAGTGTGATGGGGTTACCCAAAGTGTTCTATTAAGAACTGTGGTAGGTGAGTAATGTTCCCTATTAGAAAGGAGAGGGCAAGGCTGGTGCCAGCCGCCGCGGTAAAACCAGCTCTTCAAGTGGTCGGGATAATTATTGGGCTTAAAGTGTCCGTAGCCGGTTTAGTAAGTTCCTGGTAAAATCGGGTAGCTTAACTATCTGCATGCTAGGAATACTGCTATACTAGAGGGCGGGAGAGGTCTGAGGTACTACAGGGGTAGGGGTGAAATCTTATAATCCTTGTAGGACCACCAGTGGCGAAGGCGTCAGACTGGAACGCGCCTGACGGTGAGGGACGAAAGCCAGGGGAG |
| p__Halobacteriota_f__Halobacteriaceae_4595_ASV_2240 | 0 | 1 | 0 | 0 | 0 | 0 | 0 | 0 | 0 | 0 | 0 | CCTTTACACTGCACGACAGTGCGATAGGGGGACTCCGAGTGCGAGGGCATACAGTCCTCGCTTTTCTGCACCGTAAGGAGGTGCAAGAATAAGGGCTGGGCAAGACCGGTGCCAGCCGCCGCGGTAATACCGGCAGCCCAAGTGATGGCCGCTATTATTGGGCCTAAAGCGTCCGTAGCCGGCCGGACAAGTCCGTTGGGAAATCTGCTCGCTCAACGAGCAGGCGTCCAGCGGAAACTGTCCGGCTTGGGGCCGGAAGACTCGAGGGGTACGTCCGGGGTAGGAGTGAAATCCTGTAATCCTGGACGGACCACCGGTGGCGAAAGCGCCTCGAGAAGACGGACCCGACGGTGAGGGACGAAAGCTAGGGTCT |
| p__Halobacteriota_f__Haloferacaceae_ASV_2242 | 0 | 1 | 0 | 0 | 0 | 0 | 0 | 0 | 0 | 0 | 0 | CCTTTACACTGCACGCACGTGCGATAAGGGGACTCCGAGTGCGAGGGCATACAGTCCTCGCTTTTGAGAACCGTAAGGAGGTTCTCGAATAAGAGCTGGGCAAGACCGGTGCCAGCCGCCGCGGTAATACCGGCAGCTCAAGTGATGGCCGCTCTTATTGGGCCTAAAGCGTCCGTAGCCGGCCGAATAGGTCCGTCGGGAAATCCACTGGCTCAACCAGTGGGCGTCCGGCGGATACCGTTCGGCTTGGGACCGGAAGACTCGAGGGGTACGTCCGGGGTAGGAGTGAAATCCTGTAATCCTGGACGGACCACCGATGGCGAAAGCACCTCGAGAGGACGGATCCGACGGTGAGGGACGAAAGCTAGGGTCT |
| p__Halobacteriota_f__Natrialbaceae_ASV_2217 | 1 | 1 | 1 | 0 | 0 | 0 | 0 | 0 | 0 | 0 | 1 | CCTTTACACTGCACGACAGTGCGATAAGGGGACTCCAAGTGCGAGGGCATATCGTCCTCGCTTTTTCCTACCGTAAGGTGGTAGGCGAATAAGGGCTGGGCAAGACCGGTGCCAGCCGCCGCGGTAATACCGGCAGCCCAAGTGATGACCGCTATTATTGGGCCTAAAGCGTCCGTAGCCGGCCGAACAAGTCCGTCGGGAAATCTGCACGCCTAACGTGCAGGCGTCCGGCGGAAACTGTTCGGCTTGGGACCGGAAGATCCAGAGGGTACGTCCGGGGTAGGAGTGAAATCCTGTAATCCTGGACGGACCACCGGTGGCGAAAGCGCTCTGGAAGGACGGATCCGACGGTGAGGGACGAAAGCTAGGGTCA |
| p__Halobacteriota_f__Natrialbaceae_ASV_2282 | 0 | 0 | 0 | 0 | 1 | 0 | 0 | 0 | 0 | 0 | 0 | CCTTTACACTGCACGCCAGTGCGATAAGGGGACTCCATCTGCGAGGGCATATAGTCCTCGCTTTTCTCGACCGTAAGGTGGTCGAGGAATAAGTGCTGGGCAAGACCGGTGCCAGCCGCCGCGGTAATACCGGCAGCACGAGTGATGACCGCTATTATTGGGCCTAAAGCGTCCGTAGCTGGCCAGGCAAGTCCATCGGGAAATCCGCGTGCTCAACGCGCGGGCGTCCGGTGGAAACTGCTTGGCTTGGGACCGGAAGACCTAGGGGGTACGTCTGGGGTAGGAGTGAAATCCCGTAATCCTGGACGGACCACCGGTGGCGAAAGCGCCCTAGGAAGACGGATCCGACGGTGAGGGACGAAAGCTCGGGTCA |
| p__Halobacteriota_o__Halanaerobiales_ASV_2232 | 0 | 1 | 0 | 0 | 0 | 0 | 0 | 0 | 0 | 0 | 0 | CCTTTACACTGTACGAAAGTGCGATAAGGGGACTCCAAGTGCGAGGGCATATAGTCCTCGCTTTTCTACACCGTAAGGTGGTGTAGGAATAAGGGCTGGGCAAGACCGGTGCCAGCCGCCGCGGTAATACCGGCAGCCCAAGTGATGGCCGCTATTATTGGGCCTAAAGCGTCCGTAGCCGGCCGAGTAAGTCCGTCGGGAAATCCACACGCCTAACGTGTGGGCGTCCGGCGGAAACTGCCCGGCTTGGGACCGGGAGACCCGAGGGGTACGTCTGGGGTAGGAGTGAAATCCTGTAATCCTGGACGGACCGCCGGTGGCGAAAGCGCCTCGGGAGAACGGATCCGACGGTGAGGGACGAAAGCTAGGGTCA |
| p__Micrarchaeota_g__1-14-0-10-45-29_ASV_2305 | 1 | 0 | 0 | 0 | 0 | 0 | 0 | 0 | 0 | 0 | 0 | CCTATACAATGCGCGAAAGCGTGATATGGGGAATCCGAGTGATGGACCACGCGTTCATCTTTTCTTAAGTGTAAATAACTTGAGGAATAAGAGCTGGGTAATATCGGTGCCAGCCGCCGCGGTAACACCGAAGGCTCGAGTGGTATCCACGAATATTGGGCCTAAAGCGCTCGTAGCTGGTCTAGTAAGTTCGCTGTGAAATCTTCAAGCTTAACTTGGAGGCGTGCAGCGGATACTGCTAAGACTTGAGAGTGGGAGAGGTCAGGAGTACTTGTGGGGTAGTGGTAAAATACTGTGATCCTACAAGGACTAGCAGTGGCGAAGGCGCCTGACCAGAACACGTCTGACAGTGAGGAGCGAAGGCTAGGAGAA |
| p__Nanoarchaeota_g__BMS3ABIN17_ASV_2307 | 1 | 0 | 0 | 1 | 0 | 0 | 0 | 0 | 0 | 0 | 0 | AATCTACAATGCGCGCAAGCGTGATAGTTTGAGCCAGAGTGTTTCTCATTGAGAAACTTTTGCCGAATGTAAAAAGTTCGGCGAATAAGGACTGGGTAAGACGGGTGCCAGCCGCCGCGGTAATACCCGCGGTCCAAGTCGCAGCCATATTTATTGGGTCTAAAACATCCGTAGCTTGTTAGGTAAGTTCCCTGTGAAATCCTATCTCTTAAAGGTGGGGCGTGCAAGGAATACTGCCTGACTAGAGACCGGAAGACGTAAGAAGTACGTTCGAAGTAGCGGTAAAATGTTTTAATCTCGGACGGACTCACAATAGCGAAGGCATCTTACGAGAACGGTTCTGACAGTAAGGGATGAAGGCCAGGGGCG |
| p__Nanoarchaeota_g__BMS3ABIN17_ASV_2309 | 1 | 0 | 0 | 0 | 0 | 0 | 0 | 0 | 0 | 0 | 1 | CTTTTACAATGCTCGAAAGAGTGATAGAGCGAGCCGAAGTGTTCCTCTTAGAGGAACTTTTGTCTACTCTAAATAGGTAGATGAATAAGGACTGGGCAAGACCGGTGCCAGCCGCCGCGGTAACACCGGCGGTCCGAGTCGCATCCACATTTATTGGGTCTAAAACATCCGTAGCTTGTTTTGCAAGTCTCTTGTGAAATCCAGCGTCTTAAGCGTTGGAATTGCAAGAGATACTGCATTACTAGAGACCGGAAGACGTAAGGAGTACGGTTAAGGTAGTGGTAAAATATGTTAATCTTAACTGGACTAACAACGGCGAAGGCACCTTACGAGTACGGATCTGACAGTGAGGGATGAAGGCTAGGGGCG |
| p__Halobacteriota_g__Haladaptatus_ASV_2387 | 0 | 0 | 0 | 0 | 0 | 0 | 1 | 0 | 0 | 0 | 0 | ACTTTACACTGCACGTCAGTGCGATAAGGGAACTCCAAGTGCGAGGGCATATAGTCCTCGCTTTTGTGTATCGTAAGGTGGTACACGAATAAGAGCTGGGCAAGACCGGTGCCAGCCGCCGCGGTAATACCGGCAGCTCAAGTGATGGCCAATTTTATTGGGCCTAAAGCGTCCGTAGCCGGCCGAACAGGTCCGTCGGGAAATCTGCTCGCTCAACGAGCAGGCGTCCGACGGAAACCAGTCGGCTTGGGACCGGAAGACCCGAGGGGTACGTCTGGGGTAGGAGTGAAATCCTGTAATCCTAGACGGACCGCCGGTGGCGAAAGCGCCTCGGGAAGACGGATCCGACGGTGAGGGACGAAAGCTAGGGTCA |
| p__Halobacteriota_g__Halolamina_ASV_2336 | 0 | 1 | 0 | 0 | 0 | 0 | 0 | 0 | 0 | 0 | 0 | CCTTTACACTGCACGCCAGTGCGATAAGGGGACCCCGAGTGAAAGGGCATATCGCCCTTTCTTTTCTGGACCGTAGGGAGGTCCAGGAACAAGAGCTGGGCAAGACCGGTGCCAGCCGCCGCGGTAATACCGGCAGCTCGAGTGATGACCGATCTTATTGGGCCTAAAGCGTCCGTAGCTGGCCGTGCAAGTCCGTCGGGAAATCCACCCGCTTAACGGGTGGGCGTCCGGCGGAAACTGTGCGGCTTGGGACAGGAAGGCTCGAGGGGTACGTTCGGGGTAGGAGTGAAATCCCGTAATCCTGAACGGACCGCCGATGGCGAAAGCACCTCGAGAGGACTGATCCGACAGTGAGGGACGAAAGCTGGGGTCT |
| p__Halobacteriota_g__Haloprofundus_ASV_2334 | 0 | 1 | 0 | 0 | 0 | 0 | 0 | 0 | 0 | 0 | 0 | CCTTTACACTGCACGCCAGTGCGATAAGGGGACCCCGAGTGCGAGGGCATACAGTCCTCGCTTTTCCCAACCGTAAGGTGGTTGGTGAAGAAGAGCTGGGCAAGACCGGTGCCAGCCGCCGCGGTAATACCGGCAGCTCGAGTGATGACCGATCTTATTGGGCCTAAAGCGTCCGTAGCTGGCCAGTCAAGTCCATCGGGAAATCCATCCGCTCAACGGATGGGCGTCCGGTGGAAACTGTCTGGCTTGGAGCCGGAAGACCCAAGGGGTACGTCCGGGGTAGGAGTGAAATCCCGTAATCCTGGACGGACCGCCGATGGCGAAAGCACCTTGGGAGGACGGACTCGACAGTGAGGGACGAAAGCTGGGGTCT |
| p__Halobacteriota_g__Haloprofundus_ASV_2401 | 0 | 0 | 0 | 0 | 0 | 0 | 0 | 0 | 1 | 0 | 0 | CCTTTACACTGCACGCCAGTGCGACAAGGGGACCCCGAATGCGAGGGCATACAGTCCTCGCTTTTCCCGACCGTAAGAAGGTCGGAGAAGAAGAGCTGGGCAAGACCGGTGCCAGCCGCCGCGGTAATACCGGCAGCTCGAGTGATGACCGCTGTTATTGGGCCTAAAGCGTCCGTAGCTGGCCGCGTAAGTCCATCGGGAAATCCATCCGCTCAACGGATGGGCGTCCGGTGGAAACTACACGGCTTGGAGCCGGAAGACCCAAGGGGTACGTCCGGGGTAGGAGTGAAATCCCGTAATCCTGGACGGACCGCCGATGGCGAAAGCACCTTGGGAAGACGGACTCGACAGTGAGGGACGAAAGCTAGGGTCT |
| p__Halobacteriota_g__Halorubrum_ASV_2353 | 0 | 0 | 1 | 0 | 0 | 0 | 0 | 0 | 0 | 0 | 0 | CCTTTACACTGCACGCCAGTGCGATAAGGGGACTCCGAGTGCGAGGGCATATCGCCCTCGCTTTTGTACACCGTAGGGAGGTGTACGAATAAGGGCTGGGCAAGACCGGTGCCAGCCGCCGCGGTAATACCGGCAGCCCGAGTGATGGCCGATCTTATTGGGCCTAAAGCGTCCGTAGCTCGCCGGGTAAGTCCGTCGGGAAATCCACCCGCTCAACGGGTGGGCGTCCGACGGAACCTGCCTGGCTTGGGACCGGAAGACCCAGCGGGTACGTCCTGGGTAGGAGTGAAATCCTGTAATCCTGGACGGACCACCGATGGCGAAAGCACGCTGGGAGGACGGATCCGACAGTGAGGGACGAAAGCTGGGGTCT |
| p__Halobacteriota_g__Halorussus_ASV_2402 | 0 | 0 | 0 | 0 | 0 | 0 | 0 | 0 | 1 | 0 | 0 | ACTTTACACTGCACGACAGTGCGATAAGGGGACTCCGAGTGCGAGGGCATATAGTCCTCGCTTTTGTACACCGTAAGGTGGTGTACGAATAAGGGCTGGGCAAGACCGGTGCCAGCCGCCGCGGTAATACCGGCAGCCCGAGTGATGGCCGCTATTATTGGGCCTAAAGCGTCCGTAGCTGGCCAGACAGGTCCGTCGGGAAATCCATCCGCTTAACGGATGGGCGTCCGGCGGAAACCAGCTGGCTTGGGGCCGGAAGATCTGAGGGGTACGTCCGGGGTAGGAGTGAAATCCTGTAATCCTGGACGGACCGCCGGTGGCGAAAGCGCCTCAGAAGGACGGACCCGACGGTGAGGGACGAAAGCTAGGGTCA |
| p__Halobacteriota_g__Halovenus_ASV_2352 | 0 | 0 | 1 | 0 | 0 | 0 | 0 | 0 | 0 | 0 | 0 | CCTTTACACTGCACGACAGTGCGATAGGGGGACTCCGAGTGCGAGGGCATACAGTCCTCGCTTTTGTACACCGTAAGGTGGTGTACGAATAAGAACTGGGCAAGACCGGTGCCAGCCGCCGCGGTAATACCGGCAGTTCGAGTGATGGCCGCTTTTATTGGGCCTAAAGCGTTCGTAGCTGGCCGAGCAAGTCTGTCGGGAAATCCAGTGGCTTAACCACTGGACGTCCGGCGGAAACTGCTCGGCTTGGGACCGGGAGACTCGAGGAGTACGTCCGGGGTAGGAGTGAAATCCTGTAATCCTGGACGGACTACCGATGGCGAAAGCACCTCGAGAGCACGGATCCGACAGTGAGGAACGAAAGCTGGGGTCT |
| p__Halobacteriota_g__Halovenus_ASV_2388 | 0 | 0 | 0 | 0 | 0 | 0 | 1 | 0 | 0 | 0 | 0 | CCTTTACACTGCACGCCAGTGCGATAGGGGGACTCCGAGTGCGAGGGCATATAGTCCTCGCTTTTCTGTACCGTAAGGTGGTACAGGAACAAGTGCTGGGCAAGACCGGTGCCAGCCGCCGCGGTAATACCGGCAGCACAAGTGATGGCCGATATTATTGGGCCTAAAGCGTCCGTAGCCGGCCGGACAAGTCTGTTGGGAAATCCACGCGCTTAACGCGTGGGCGTCCAGCGGAAACTGTCTGGCTTGGGGCCGGAAGACTCGAGGGGTACGTCCGGGGTAGGAGTGAAATCCCGTAATCCTGGACGGACCGCCGGTGGCGAAAGCGCCTCGAGAGGACGGACCCGACGATGAGGGACGAAAGCCAGGGTCT |
| p__Halobacteriota_g__Natrarchaeobaculum_ASV_2346 | 0 | 1 | 1 | 0 | 0 | 0 | 0 | 0 | 0 | 0 | 0 | CCTTTACACTGCGCGACAGCGCGATAAGGGGACTCCATCTGCGAGGGCATATCGTCCTCGCTTTTCTCGACCGTAAGGTGGTCGAGGAATAAGGGCTGGGCAAGACCGGTGCCAGCCGCCGCGGTAATACCGGCAGCCCGAGTGATGACCGCTCTTATTGGGCCTAAAGCGTCCGTAGCTGGCCTCGCAAGTCCATCGGGAAATCCGCACGCTCAACGTGCGGGCGTCCGGTGGAAACTGCGTGGCTTGGGACCGGAAGACCCGAGGGGTACGTCCGGGGTAGGAGTGAAATCCCGTAATCCTGGACGGACCACCGGTGGCGAAAGCGCCTCGGGAGGACGGATCCGACGGTGAGGGACGAAAGCCAGGGTCA |
| p__Halobacteriota_g__Natrarchaeobaculum_ASV_2371 | 0 | 0 | 0 | 0 | 1 | 0 | 0 | 0 | 0 | 0 | 0 | CCTTTACACTGCGCGACAGCGCGATAAGGGGACTCCATCTGCGAGGGCATATCGTCCTCGCTTTTTCCTACCGTAAGGTGGTAGGCGAATAAGGGCTGGGCAAGACCGGTGCCAGCCGCCGCGGTAATACCGGCAGCCCGAGTGATGACCGCTCTTATTGGGCCTAAAGCGTCCGTAGCTGGCCTCGCAAGTCCATCGGGAAATCCGCACGCTCAACGTGCGGGCGTCCGGTGGAAACTGCGTGGCTTGGGACCGGAAGACCCGAGGGGTACGTCTGGGGTAGGAGTGAAATCCCGTAATCCTGGACGGACCACCGGTGGCGAAAGCGCCTCGGGAGGACGGATCCGACGGTGAGGGACGAAAGCCAGGGTCA |
| p__Halobacteriota_g__Natrinema_ASV_2348 | 0 | 0 | 1 | 0 | 0 | 0 | 0 | 0 | 0 | 0 | 0 | CCTTTACACTGCACGAGAGTGCGATAAGGGGACTCCGAGTGCGAGGGCATATAGCCCTCGCTTTTCGCAACCGTAAGGTGGTTGCAGAATAAGTGCTGGGCAAGACCGGTGCCAGCCGCCGCGGTAATACCGGCAGCACGAGTGATGACCGCTATTATTGGGCCTAAAGCGTCCGTAGCTGGCCACGCAAGTCCATCGGGAAATCCGCGCGCTCAACGCGCGGGCGTCCGGTGGAAACTGCGTGGCTTGGGACCGGAAGACCAGAGGGGTACGTCCGGGGTAGGAGTGAAATCCCGTAATCCTGGACGGACCACCGGTGGCGAAAGCGCCTCTGGAAGACGGATCCGACGGTGAGGGACGAAAGCTCGGGTCA |

| | Presence (1) or Absence (0) in each layer | | | | | | | | | | | |
|---|---|---|---|---|---|---|---|---|---|---|---|---|
| **Phylum_Genus_ASV_number** | **L1** | **L2** | **L3** | **L4** | **L5** | **L6** | **L7** | **L8** | **L9** | **L10** | **L11** | **Sequence** |
| p__Thermoproteota_g__Nitrosocosmicus_ASV_2377 | 0 | 0 | 0 | 0 | 0 | 1 | 0 | 0 | 0 | 0 | 0 | CCTCTGCAATAGGCGAAAGCTTGACAGGGTTACTCTGAGTGATCTCCGATAAGGAGATCTTTTGGCATCTCTAAAAATGATGCAGAATAAGGGGTGGGCAAGTCTGGTGTCAGCCGCCGCGGTAATACCAGCACCCCGAGTGGTCGGGACGATTATTGGGCCTAAAGCATCCGTAGCCGGTTCTGCAAGTCTTCCGTTAAATCCAGCTGCTTAACAGATGGGCTGCGGGGGATACTACAGAGCTAGGAGGCGGGAGAGGCAAGCGGTACTCAGTGGGTAGGGGTAAAATCCGTTGATCCATTGAAGACCACCAGTGGCGAAGGCGGCTTGCCAGAACGCACTCGACGGTGAGGGATGAAAGCTGGGGGAG |
| p__Halobacteriota_g__Saliphagus_ASV_2341 | 0 | 1 | 0 | 0 | 0 | 0 | 0 | 0 | 0 | 0 | 0 | CCTTTACACTGCACGACAGTGCGATAAGGGGACTCCGAGTGCGAGGGCATATCGTCCTCGCTTTTCTGCACCGTAAGGTGGTGCAGGAATAAGTGCTGGGCAAGACCGGTGCCAGCCGCCGCGGTAATACCGGCAGCACGAGTGATGACCGCTATTATTGGGCCTAAAGCGTCCGTAGCTGGCCGATCAAGTCCGTCGGGAAATCCGCACGCCTAACGTGCGGGCGTCCGGCGGAAACTGTTCGGCTTGGGACCGGAAGACCCGAGGGGTACGTCCGGGGTAGGAGTGAAATCCTGTAATCCTGGACGGACCACCGGTGGCGAAAGCGCCTCGGGAGGACGGATCCGACGGTGAGGGACGAAAGCTAGGGTCA |
| p__Thermoplasmatota_g__SM1-50_ASV_2324 | 0 | 0 | 0 | 0 | 0 | 0 | 0 | 0 | 0 | 0 | 1 | CCTTCACACTGTGCGAAAGCGCGATGAGGGGATCCCAAGTGCTTGCACGTAAGTGTAAGCTGTTTCTTTGTCTAAAAAGCAGAGAGAGTAAGAGCTGGGTAAGACGGCTGCCAGCCGCCGCGGTAATAGCTGCAGCTCGAGTGGTGACCACTTTTATTGGGCCTAAAACGTCCGTAGCCGGTCTAGTAAATGCCTGGGTAAATCGTGCCGCTTAACGGTACGAATTCCGGGTAGACTGCTAGACTTGAGATCGGGAGAGGTTAGAGGTACTCCTTGGGTAGAGGTGAAATTCTGTAATCCTAGGGGGACCACCTGTGGCGAAAGCGTCTAACCAGAACGAGTCTGACGGTGAGGGACGAAGCCCTGGGTCA |
| p__Halobacteriota_f__Natrialbaceae_ASV_2327 | 0 | 1 | 0 | 0 | 0 | 0 | 0 | 0 | 0 | 0 | 0 | CCTTTACACTGCACGACAGTGCGATAAGGGGACTCCAAGTGCGAGGGCATATCGTCCTCGCTTTTCACCACCGTAAGGTGGTGGTGGAATAAGTGCTGGGCAAGACCGGTGCCAGCCGCCGCGGTAATACCGGCAGCACGAGTGATGACCGCTATTATTGGGCCTAAAGCGTCCGTAGCTGGCCGATCAAGTCCATCGGGAAATCCGCACGCCTAACGTGCGGGCGTCCGGTGGAAACTGTTCGGCTTGGGACCGGAAGACCCGAGGGGTACGTCTGGGGTAGGAGTGAAATCCTGTAATCCTGGACGGACCACCGGTGGCGAAAGCGCCTCGGGAGGACGGATCCGACGGTGAGGGACGAAAGCTAGGGTCA |
| p__Halobacteriota_f__Natrialbaceae_ASV_2351 | 0 | 0 | 1 | 0 | 0 | 0 | 0 | 0 | 0 | 0 | 0 | CCTTTACACTGCACGCCAGTGCGATAAGGGGACTCCAAGTGCGAGGGCATATAGTCCTCGCTTTTTCCTACCGTAAGATGGTAGGCGAATAAGGGCTGGGCAAGACCGGTGCCAGCCGCCGCGGTAATACCGGCAGCTCAAGTGATGACCGCTATTATTGGGCCTAAAGCGTCCGTAGCCGGCCAAACAAGTCCGTCGGGAAATCCGCACGCCTAACGTGTGGGCGTCCGGCGGAAACTGTTTGGCTTGGGACCGGAAGACTCGGGGGGTACGTCTGGGGTAGGAGTGAAATCCTGTAATCCTGGACGGACCACCGGTAGCGAAAGCGCCTCGAGAGAACGGATCCGACGGTGAGGGACGAAAGCTAGGGTCA |
| p__Halobacteriota_o__Halanaerobiales_ASV_2349 | 0 | 0 | 1 | 0 | 0 | 0 | 0 | 0 | 0 | 0 | 0 | CCTTTACACTGCACGAAAGTGCGATAAGGGGACCCCGAGTGCGAGGGCATACAGTCCTCGCTTTTGTGTACCGTACGATGGTACACGAATAAGAGCTGGGCAAGACCGGTGCCAGCCGCCGCGGTAATACCGGCAGCTCAAGTGATGGCCGTTATTATTGGGCCTAAAGCGTCCGTAGCCGGCCGAACAGGTCTGTTGGGAAATCCGCCAGCTTAACTGGTGGGCGTCCAGCAGAAACCCGTCGGCTTGGGGCCGGAAGACCTGAGGGGTACGTCCGGGGTAGGAGTGAAATCCTGTAATCCTGGACGGACCGTCGGTGGCGAAAGCGCCTCAGGAGGACGGACCCGACGGTGAGGGACGAAAGCTAGGGTCA |
| d__Archaea_d__Archaea_ASV_2396 | 0 | 0 | 0 | 0 | 0 | 0 | 0 | 0 | 1 | 0 | 0 | CTTCTGCAATGACCGCAAGGTTGACAGAGCGATTCCGAGTGCTCATGCAAAGCATGGGCTTTTGCCAAGTGTAAATAGCTTGGCGAATAAGGGGTGGGCAAGACCGGTGGCAGCAGCCGCGGTAATACCGGCGCCCCAAGTGGTGGTCATTATTATTGGGCCTAAAGCGTCCGTAGCTGGTTTGGTTAGTTTTTGGTGAAATCCATGCGCTCAACGTATGGGCGAGCCAGAAATACTACCAAGCTAGAGACCGGGAGAAGACAGGGGTATTCCTAGGGTAAGGGTAAAATCTTGTAATCCTAGGAGGACCACCGGTGGCGAAGGCGCCTGTCTAGAACGGGTCTGACAGTGAGGGACGAAAGCTGAGGGAG |
| p__Nanoarchaeota_g__CG1-02-57-44_ASV_2473 | 0 | 0 | 1 | 0 | 0 | 0 | 0 | 1 | 0 | 0 | 0 | CCTTTACCATGCACGAAAGTGTGATAAGGGGAACCCCAGTGCTCACGTACTGCGTGGGCTTTTGCCAAGTGTAAATAGCTTGGCGAATAAGTGGTGGGTAAGACTGGTGCCAGCCGCCGCGGTAACACCAGCGCCACGAGTGGTAACCGCGATTATTGGGCCTAAAGCGTCCGTAGCCGGCTTCGTAAATCCTTTGTGAAATCGTGCGGCTCAACCGTACGACGCGCAAGGGACACTGCGAGGCTCGAGACCGGGAGGAGTCAGAAGTATTCCAGAGGGAGCGGTAAAATGTTTTAATCTCTGGAGGACTACCTATGGCGAAGGCATCTGACCAGAACGGATCTGACGGTGAGGGACGAAAGCCAGGGGAG |
| p__Halobacteriota_g__Halalkaliarchaeum_ASV_2409 | 1 | 0 | 0 | 0 | 0 | 0 | 0 | 0 | 0 | 0 | 0 | CCTTTACACTGCACGCGAGTGCGATAAGGGGACCCCGAGTGCGAGGGCATATCGTCCTCGCTTTTGTGTACCGTAGGGAGGTACACGAACAAGGGCTGGGCAAGACCGGTGCCAGCCGCCGCGGTAATACCGGCAGCCCAAGTGATGGCCACTGTTATTGGGCCTAAAGCGTCCGTAGCCGGCCTGGCAAGTCCGTCGGGAAATCCACGCGCCCAACGCGTGGGCGGCCGGCGGAACCTGCCGGGCTCGGGACCGGAGGACCCAGCGGGTACGTCCGGGGTAGGAGTGAAATCCTGTAATCCCGGACGGACCGCCGATGGCGAAAGCACGCTGGGAAGACGGATCCGACGGTGAGGGACGAAAGCTAGGGTCT |
| p__Halobacteriota_g__Halobacterium_ASV_2459 | 0 | 0 | 1 | 0 | 0 | 0 | 0 | 0 | 0 | 0 | 0 | CCTTTACACTGTACGCAAGTGCGATAGGGGGACTCCGAGTGCGAGGGCATAGAGCCCTCGCTTTTGTACACCGTAAGGTGGTGTACGAATAAGGGCTGGGCAAGACCGGTGCCAGCCGCCGCGGTAATACCGGCAGCCCGAGTGATGGCCGATCTTATTGGGCCTAAAGCGTCCGTAGCTGGCCAGGCAAGTCCGTTGGGAAATCTGCTCGCTCAACGAGCAGGCGTCCAGCGGAAACTGCTTGGCTTGGGACCGGAGGACCTGAGGGGTACGTCCGGGGTAGGAGTGAAATCCTGTAATCCTGGACGGACCGCCGGTGGCGAAAGCGCCTCAGGAAGACGGATCCGACAGTGAGGGACGAAAGCTAGGGTCT |
| p__Halobacteriota_g__Halococcus_ASV_2493 | 0 | 0 | 0 | 0 | 1 | 1 | 0 | 0 | 0 | 0 | 0 | CCTTTACACTGCGCGACAGCGCGATAGGGGGACCCCAAGTGCGAGGGCATACAGTCCTCGCTTTTTCCGACCGTAAGGTGGTCGGCGAATAAGAGCTGGGCAAGACCGGTGCCAGCCGCCGCGGTAATACCGGCAGCTCGAGTGATGGCCGCTATTATTGGGCCTAAAGCGTCCGTAGCCGGCTAGACAGGTCCGTCGGGAAATCCACCCGCTCAACGGGTGGGCGTCCGGCGGAAACCAGTTAGCTTGGGACCGAGCGACCTGAGGGGTACGTCTGGGGTAGGAGTGAAATCCCGTAATCCCGGACGGACCATCGGTGGCGAAAGCGCCTCAGGAGAGCGGATCCGACGGTGAGGGACGAAAGCTAGGGTCT |
| p__Halobacteriota_g__Halodesulfurarchaeum_ASV_2464 | 0 | 0 | 1 | 0 | 0 | 0 | 0 | 0 | 0 | 0 | 0 | CCTTTACACTGCACGACAGTGCGATAGGGGGACTCCGAGTGCGAGGGCATAGAGCCCTCGCTTTTGTACACCGTAAGGTGGTGTACGAATAAGGGCTGGGCAAGACCGGTGCCAGCCGCCGCGGTAATACCGGCAGCCCAAGTGATGGCCGATATTATTGGGCCTAAAGCGTCCGTAGCTGGCCGGACAAGTCCGTTGGGAAATCCGCACGCTCAACGTGCGGGCGTCCAGCGGAAACTGTCCGGCTTGGGACCGGAAGACTCGAGGGGTACGTCCGGGGTAGGAGTGAAATCCCGTAATCTCGGACGGACCACCTGTGGCGAAAGCGCCTCGAGAAGACGGATCCGACAGTGAGGGACGAAAGCCAGGGTCT |
| p__Halobacteriota_g__Halohasta_ASV_2447 | 0 | 1 | 0 | 0 | 0 | 0 | 0 | 0 | 0 | 0 | 0 | CCTTTACACTGCACGCAAGTGCGATAAGGGGACCCCAAGTGCGAGGGCATACAGTCCTCGCTTTTCTACACCGTAAGGAGGTGTAGGAATAAGAGCTGGGCAAGACCGGTGCCAGCCGCCGCGGTAATACCGGCAGCTCGAGTGATGACCGATTTTATTGGGCCTAAAGCGTCCGTAGCTGGCCGCACAAGTCCGTCGGGAAATCCACCTGCCCAACAGGTGGGCGTCCGGCGGAAACTGTGTGGCTTGGAACCGGAAGGCTCAGAGGGTACGTCTGGGGTAGGAGTGAAATCCCGTAATCCTGGACGGACCACCGATGGCGAAAGCACTCTGAGAAGACGGATTCGACAGTGAGGGACGAAAGCTGGGGTCT |
| p__Halobacteriota_g__Halolamina_ASV_2417 | 0 | 0 | 0 | 0 | 0 | 0 | 0 | 0 | 0 | 1 | 0 | CCTTTACACTGCACGCCAGTGCGATAAGGGGACCCCGAGTGAAAGGGCATATCGCCCTTTCTTTTCTGAACCGTAGGGAGGTTCAGGAACAAGAGCTGGGCAAGACCGGTGCCAGCCGCCGCGGTAATACCGGCAGCTCAAGTGATGACCGATTTTATTGGGCCTAAAGCGTCCGTAGCTGGCCGCGCAAGTCTGTCGGGAAATCCACCCGCTTAACGGGTGGGCGTCCGGCGGAAACTGTGCGGCTTGGGACAGGAAGGCTCGAGGGGTACGTTCGGGGTAGGAGTGAAATCCCGTAATCCTGAACGGACCACCGATGGCGAAAGCACCTCGAGAGGACTGATCCGACAGTGAGGGACGAAAGCTGGGGTCT |
| p__Halobacteriota_g__Halorientalis_ASV_2451 | 0 | 1 | 0 | 0 | 0 | 0 | 0 | 0 | 0 | 0 | 0 | CCTTTACACTGCACGACAGTGCGATAAGGGGACTCCGAGTGCGAGGGCATACAGTCCTCGCTTTTCTGTACCGTAGGGAGGTACAGGAACAAGTGCTGGGCAAGACCGGTGCCAGCCGCCGCGGTAATACCGGCAGCACGAGTGATGGCCGCTATTATTGGGCCTAAAGCGTCCGTAGCCGGCCGAGCAAGTCCGTCGGGAAATCTACCCGCTTAACGGGTAGGCGTCCGGCGGAAACTGCTCGGCTTGGGGCCGGGAGATCCAAGGGGTACGTCCGGGGTAGGAGTGAAATCCCGTAATCCTGGACGGACCACCGGTGGCGAAAGCGCCTTGGAAAAACGGACCCGACGGTGAGGGACGAAAGCTAGGGTCT |
| p__Halobacteriota_g__Halorubrum_ASV_2497 | 0 | 0 | 0 | 0 | 0 | 1 | 0 | 0 | 0 | 0 | 0 | CCTTTACACTGCACGACAGTGCGATAGGGGGATCCCAAGTGCACAGGCATAGCGCCTGTGCTTTTCGGCACCGTAAGGTGGTGTCAGAATAAGGGCTGGGCAAGACCGGTGCCAGCCGCCGCGGTAATACCGGCAGCCCAAGTGATGGCCGATTTTATTGGGCCTAAAGCGTCCGTAGCTGGCCGCGCAAGTCCATCGGGAAATCCACCTGCTCAACAGGTGGGCGCCCGGTAGAAACTGCGTGGCTTGGGACCGGAAGGCGCGACGGGTACGTCCGGGGTAGGAGTGAAATCCCGTAATCCTGGACGGACCGCCGATGGCGAAAGCACGTCGCGAGAACGGATCCGACAGTGAGGGACGAAAGCCAGGGTCT |
| p__Halobacteriota_g__Halorussus_ASV_2483 | 0 | 0 | 0 | 1 | 0 | 0 | 0 | 0 | 0 | 0 | 0 | CCTTTACACTGCACGACAGTGCGATAAGGGGACTCCGAGTGCGAGGGCATATAGTCCTCGCTTTTCTGTACCGTAAGGTGGTACAGGAATAAGGACTGGGCAAGACCGGTGCCAGCCGCCGCGGTAATACCGGCAGTCCAAGTGATGGCCGCTATTATTGGGCCTAAAGCGTCCGTAGCCGGCCAGACAGGTCCGTCGGGAAATCCACGCGCTCAACGCGTGGGCGTCCGGCGGAAACCAGCTGGCTTGGGGCCGGAAGACCTGAGGGGTACGTCCGGGGTAGGAGTGAAATCCCGTAATCCTGGACGGACCACCGGTGGCGAAAGCGCCTCAGGAAAACGGACCCGACGGTGAGGGACGAAAGCTAGGGTCA |
| p__Halobacteriota_g__Halorussus_ASV_2492 | 0 | 0 | 0 | 0 | 1 | 0 | 0 | 0 | 0 | 0 | 0 | CCTTTACACTGCACGACAGTGCGATAAGGGGACTCCGAGTGCGAGGGCATATAGTCCTCGCTTTTCTGTACCGTAAGGTGGTACAGGAATAAGGGCTGGGCAAGACCGGTGCCAGCCGCCGCGGTAATACCGGCAGCCCGAGTGATGGCCGCTATTATTGGGCCTAAAGCGTCCGTAGCCGGCCAGACAGGTCCGTCGGGAAATCCACGCGCTCAACGCGTGGGCGTCCGGCGGAAACCAGCTGGCTTGGGGCCGGAAGACCTGAGGGGTACGTCCGGGGTAGGAGTGAAATCCTGTAATCCTGGACGGACCGCCGGTGGCGAAAGCGCCTCAGGAAGACGGACCCGACGGTGAGGGACGAAAGCTAGGGTCA |
| p__Halobacteriota_g__Natrinema_ASV_2495 | 0 | 0 | 0 | 0 | 0 | 1 | 0 | 0 | 0 | 0 | 0 | CCTTTACACTGCACGCCAGTGCGATAAGGGGACTCCGAGTGCGAGGGCATATAGTCCTCGCTTTTCACCACCGTAAGGTGGTGGTGGAATAAGTGCTGGGCAAGACCGGTGCCAGCCGCCGCGGTAATACCGGCAGCACGAGTGATGACCGCTATTATTGGGCCTAAAGCGTCCGTAGCTGGCCACGCAAGTCTATCGGGAAATCTGCGCGCTCAACGCGCAGGCGTCCGGTGGAAACTGCGTGGCTTGGGACCGGAAGATCCAGAGGGTACGTCCGGGGTAGGAGTGAAATCCCGTAATCCTGGACGGACCACCGGTGGCGAAAGCGCTCTGGAAAGACGGATCCGACGGTGAGGGACGAAAGCTCGGGTCA |
| p__Halobacteriota_g__Natronomonas_ASV_2467 | 0 | 0 | 1 | 0 | 0 | 0 | 0 | 0 | 0 | 0 | 0 | CCTTTACACTGCACGCCAGTGCGATAGGGGGACACCGAGTGCGAGGGCATAGAGCCCTCGCTTTTGTAGACCGTAAGGAGGTCTACGAATAAGGGCTGGGCAAGACCGGTGCCAGCCGCCGCGGTAACACCGGCAGCCCAAGTGATGGCCGATATTATTGGGCCTAAAGCGTCCGTAGCTGGCCGCACAAGTCCGTCGGGAAATCCACGCGCTTAACGCGTGGGCGTCCGGCGGAAACTGTGCGGCTTGGGACCGGAAGATCCGAGGGGTACGTCCGGGGTAGGAGTGAAATCCTGTAATCCTGGACGGACCACCGATGGCGAAAGCACCTCGGAAGGACGGATCCGACAGTGAGGGACGAAAGCTAGGGTCT |
| p__Halobacteriota_g__Saliphagus_ASV_2455 | 0 | 0 | 1 | 0 | 0 | 0 | 0 | 0 | 0 | 0 | 0 | CCTTTACACTGCACGCGAGTGCGATAAGGGGACCCCGAGTGCGAGGGCATACAGTCCTCGCTTTTGTGTACCGTAAGGAGGTACACGAATAAGTGCTGGGCAAGACCGGTGCCAGCCGCCGCGGTAATACCGGCAGCACGAGTGATGACCGCTGTTATTGGGCCTAAAGCGTCCGTAGCCGGCCAGACAGGTCCGTCGGGAAATCCGCACGCTCAACGTGCGGGCGTCCGGCGGAACCCAGCTGGCTAGGGACCGGAAGACCCGAGGGGTACGTCCGGGGTAGGAGTGAAATCCCGTAATCCTGGACGGACCGCCGGTGGCGAAAGCGCCTCGGGAAGACGGATCCGACGGTGAGGGACGAACGCTAGGGTCA |
| p__Halobacteriota_g__Saliphagus_ASV_2468 | 0 | 0 | 1 | 0 | 0 | 0 | 0 | 0 | 0 | 0 | 0 | CCTTTACACTGCACGAAAGTGCGATAAGGGGACCCCGAGTGCGAGGGCATACAGTCCTCGCTTTTGTGTACCGTAGGGAGGTACACGAATAAGGGCTGGGCAAGACCGGTGCCAGCCGCCGCGGTAATACCGGCAGCTCAAGTGATGGCCGCTGTTATTGGGCCTAAAGCGTCCGTAGCCGGCCGCGCAAGTCCGTCGGGAAATCCACCCGCCTAACGGGTGGGCGTCCGGCGGAAACTGCGCGGCTTGGGACCGGAAGACTCGAGGGGTACGTCCGGGGTAGGAGTGAAATCCCGTAATCCTGGACGGACCGCCGGTGGCGAAAGCGCCTCGAGAGGACGGATCCGACGGTGAGGGACGAAAGCTAGGGTCT |
| p__Thermoplasmatota_g__SM1-50_ASV_2521 | 0 | 0 | 0 | 0 | 0 | 0 | 0 | 0 | 1 | 0 | 0 | CCTTCACAATGTGCGAAAGCACGATGAGGGGATCCCAAGTGCTTGCACGTAAGTGTAAGCTGTTTCTTTGTCTAAAACGCATAGAGAGTAAGGGCTGGGTAAGACGGGTGCCAGCCGCCGCGGTAATACCTGCAGCTCAAGTGGTGGCCACTATTATTGGGCCTAAAACATTCATAGCCAGCTTGATAAATGCCTGGGTAAATCGTATAGCTCAACTATACGAATTCCGGGTAGACTGTTAAGCTTGGGACCGGGAGAGGCTAGAGGTACTCCTGGGGTAGAGGTGAAATTCTGTAATCCTGGGGGGACCACCAGTGGCGAAGGCGTCTAGCTAGAACGGGTCCGACGGTGAGGGACGAAGCCCTGGGGCG |
| p__Halobacteriota_f__Natrialbaceae_ASV_2462 | 0 | 0 | 1 | 0 | 0 | 0 | 0 | 0 | 0 | 0 | 0 | CCTTTACACTGCACGAGAGTGCGATAAGGGGACTCCAAGTGCGAGGGCATATAGTCCTCGCTTTTCACCACCGTAAGGAGGTGGTGGAATAAGTGCTGGGCAAGACCGGTGCCAGCCGCCGCGGTAATACCGGCAGCACGAGTGATGACCGCTATTATTGGGCCTAAAGCGTCCGTAGCTGGCCACGCAAGTCTATCGGGAAATCCGCGCGCCTAACGCGCGGGCGTCCGGTAGAAACTGCGTGGCTTGGGACCGGAAGATCCAGAGGGTACGTCTGGGGTAGGAGTGAAATCCCGTAATCCTGGACGGACCACCGGTGGCGAAAGCGCTCTGGAAGGACGGATCCGACGGTGAGGGACGAAAGCTTGGGTCA |

| Phylum_Genus_ASV_number | Presence (1) or Absence (0) in each layer | | | | | | | | | | | |
|---|---|---|---|---|---|---|---|---|---|---|---|---|
| | L1 | L2 | L3 | L4 | L5 | L6 | L7 | L8 | L9 | L10 | L11 | Sequence |
| p__Halobacteriota_o__Halanaerobiales_ASV_2440 | 0 | 1 | 0 | 0 | 0 | 0 | 0 | 0 | 0 | 0 | 1 | CCTTTACACTGCACGACAGTGCGATAAGGGGATTCCGAGTGCGAGGGCATACAGTCCTCGCTTTTGTACACCGTAGGGTGGTGTACGAATAAGGGCTGGGCAAGACCGGTGCCAGCCGCCGCGGTAACACCGGCAGCCCAAGTGATGGCCGCTATTATTGGGCCTAAAGCGTCCGTAGCTGGCCGGACAGGTCCGTTGGGAAATCCACTTGCTTAACGAGTGGGCGTCCAGCGGAAACCGTTCGGCTTGGGACCGGAAGACCCGAGGGGTACGTCTAGGGTAGGAGTGAAATCCTGTAATCCTGAACGGACCACCGGTGGCGAAAGCGCCTCGGGAGAACGGATCCGACAGTGAGGGACGAAAGCTAGGGTCT |
| p__Halobacteriota_o__Halanaerobiales_ASV_2487 | 0 | 0 | 0 | 0 | 1 | 0 | 0 | 0 | 0 | 0 | 0 | CCTTTACACTGCACGACAGTGCGATAGGGGGACTCCGAGTGCGAGGGCATATAGTCCTCGCTTTTCTGAACCGTAAGGAGGTTCAGGAACAAGTGCTGGGCAAGACCGGTGCCAGCCGCCGCGGTAATACCGGCAGCACGAGTGATGGCCAATCTTATTGGGCCTAAAGCGTCCGTAGCCGGCCCGACAAGTCCGTTGGGAAATCCACGCGCTCAACGCGTGGGCGTCCAGCGGAAACTGTCGGGCTTGGGACCGGAAGACCCAGAGGGTACGTCCGGGGTAGGAGTGAAATCCCGTAATCCTGGACGGACCACCGGTGGCGAAAGCGCTCTGGGAAGACGGATCCGACGGTGAGGGACGAAAGCTAGGGTCT |
| p__Halobacteriota_o__Halanaerobiales_ASV_2488 | 0 | 0 | 0 | 0 | 1 | 0 | 0 | 0 | 0 | 0 | 0 | CCTTTACACTGTACGGAAGTGCGATAAGGGGACCCCGAGTGCGAGGGCATACAGTCCTCGCTTTTGTACACCGTAGGGAGGTGTACGAATAAGGGCTGGGCAAGACCGGTGCCAGCCGCCGCGGTAATACCGGCAGCCCAAGTGATGGCCGCTATTATTGGGCCTAAAGCGTCCGTAGCCGGCCGGACAAGTCCGTTGGGAAATCCGTTCGCTTAACGAACGGGCGTCCAGCGGAAACTGTCCGGCTTGGGACCGGAAGACCCGAGGGGTACGTCCGGGGTAGGAGTGAAATCCCGTAATCCTGGACGGACCGCCGGTGGCGAAAGCGCCTCGGGAAGACGGATCCGACGGTGAGGGACGAAAGCCAGGGTCT |
| p__Nanoarchaeota_o__Woesearchaeales_581_ASV_2410 | 1 | 0 | 0 | 0 | 0 | 0 | 0 | 0 | 0 | 0 | 0 | CCTCAGCAATGCACGAAAGTGTGACTGGGGGATTCCGAGTGCTCATGTAACGCATGGGCTTTTGCCAAAGGTAATGTCTTTGGCGAATAAGTGGGGGGTAATACTGGTGCCAGCCGCCGCGGTAACACCAGAGCCACGAGTGGGGACCGCTATTATTGGGCCTAAAGCGTTCGTAGCCGGCCGGGTGAATCTCCTGTGAAATCGCAGCGCTCAACGTTGCGGCGCGCAGGAGACACTGCTCGGCTCGGGACCGGGAGGCGTATGAGGTATTCTGGGGGTAGCGGTAAAATGCGATAATCCTCAGAGGACCACCTGTGGCGAAGGCGTCATACGAGAACGGATCCGACGGTGAGGGACGAAAGCTAGGGGAG |
| d__Archaea_d__Archaea_ASV_2530 | 0 | 0 | 0 | 0 | 0 | 0 | 0 | 0 | 1 | 0 | 0 | CCTTTACAATGCGCGCAAGCGTGATAGGGGGATTTCGAGTGCACTCTTTGAGTGCTTTTGCCAAGCCCAAACAGCTTGGCGAATAAGTGGGGGGCAAGATACGTGCCAGTCGCCGCGGTAACACGTACTCCACAAGTGGTATCCATCTATATTGGGCCTAAAACGTCCGTAGCTCGCCTAAAAAGTCCTTGGTGAAAGCAGCTGGCTTAACCAGCTGAACTGCCAAGGAAACTGTTGGGCTAGAGGCCGGAGGAGGTCAAGGGAATTTCAGGGGTAGCGGTGAAATGCTATGATCCTTGAAAGACCACCAGTAGCGAAGGCGCTTGACCAATACGGTCCTGACAGTGAGGGACGAAAGCTGGGGGAG |
| p__Halobacteriota_g__A07HB70_ASV_2587 | 0 | 0 | 1 | 0 | 0 | 0 | 0 | 0 | 0 | 0 | 0 | CCTTTACACTGCACGAACGTGCGATAAGGGGACTCCAAGTGCGAGGGCATACAGTCCTCGCTTTTCTGTACCGTAGGGTGGTACAGGAACAAGTGCTGGGCAAGACCGGTGCCAGCCGCCGCGGTAATACCGGCAGCACGAGTGATGGCCGATCTTATTGGGCCTAAAGCGTCCGTAGCCGGTCGCGTCGGTCCGTCGGGAAATCCCTCGGCCCAACCGAGGGGCGTCCGGCGGAAACCACGCGACTTGGGGCCGGAAGATCCAACGGGTACGTCCGGGGTAGGAGTGAAATCCCGTAATCCTGGACGGACCGCCGATGGCGAAAGCACGTTGGAAAGACGGACCCGACGGTGAGGGACGAAAGCTCGGGTCT |
| p__Thermoproteota_g__BA2_ASV_2538 | 0 | 0 | 0 | 0 | 0 | 0 | 0 | 0 | 0 | 1 | 0 | ACTCCGCAATGCACGAAAGTGTGACGGGGCTACCCCGAGTGCCATCCGCTGAGGATGGCTTTTCTTTGGTATAACTAGCCATTGGAATAAGGAGAGGGCAAGACTGGTGTCAGCCGCCGCGGTAATACCAGCTCTCCGAGTGGTAGGGATGATTATTGGGCTTAAAGCGTCCGTAGCCGGCTTAGCAAGTTTCCTGTTAAACTCAGCGACTCAATCGTTGGATCGCGGGAAATACTACTAGGCTAGGAGGCGGGAGAGGTCAACGGTACTTCTGGGGTAGGGGTGAAATCCTATAATCCCGGGAGGACCACCAGTGGCGAAGGCTGTTGACTAGAACGCGCTCGACGGTGAGGGACGAAAGCTGGGGTAG |
| p__Nanoarchaeota_g__BMS3ABIN17_ASV_2548 | 0 | 0 | 0 | 0 | 0 | 0 | 0 | 0 | 1 | 1 | 0 | CTTCTGCAATACGCGCAAGCGTGACAGAGCAAGCCAGAGTGATGCTCTATAATTGAGCATCTTTTATATGATGCAAAAAGTCGTATGAATAAGGACTGGGAAAGACTGGTGCCAGCCGCCGCGGTAATACCAGCGGTCCAAGTCGCAGCCACTTTTATTGGGTCTAAAACATCCGTAGCTTGCCTCGTCAGTCTTTTGTGAAATCTGGCATCTTAAGTGTCGGGCGCGCAGGAGATACTGCAAGGCTGGAGACCGGGAGACGTAAGGAGTACGGTTTGGGTAGCGGTTAAATGCGTTGATCCTTTCCGGACTAACGATAGCGAAGGCACCTTACGAGAACGGATCTGACAGTGAGGGATGAAGGCTAGGGTCG |
| p__Nanoarchaeota_g__BMS3ABIN17_ASV_2662 | 0 | 0 | 0 | 0 | 0 | 0 | 0 | 1 | 1 | 0 | 0 | ATTTTACAATGCTCGCAAGAGTGATAGAATGAGTCAGAGTGTTTCTCCTAGAGAAACTTTTGTCTGTTGTAAAAAGACAGACGAATAAGGACTGGGCAAGACCGGTGCCAGCCGCCGCGGTAATACCGGCGGTCCAAGTCGTATCCACATTTATTGGGTCTAAAACATCCGTAGCTTGTTTTGCAAGTCTCTTGTGAAATCTGACATCTCAAGTGTCAGGCGCGCAAGAGATACTGCATTGCTAGAGACCGGAAGACGTAAGGAGTACGATTGAGGTAGCGGTAAAATGTTTTAATCTTAATCGGACTAACAATGGCGAAGGCACCTTACGAGTACGGATCTGACAGTCAGGGATGAAGGCTAGGGGCG |
| p__Halobacteriota_g__Halalkaliarchaeum_ASV_2612 | 0 | 0 | 0 | 1 | 0 | 0 | 0 | 0 | 0 | 0 | 0 | CCTTTACACTGCACGCGAGTGCGATAAGGGGACCCCGAGTGCGAGGGCATATCGTCCTCGCTTTTGTGTACCGTAGGGAGGTACACGAACAAGGGCTGGGCAAGACCGGTGCCAGCCGCCGCGGTAATACCGGCAGCCCGAGTGATGGCCACTGTTATTGGGCCTAAAGCGTCCGTAGCCGGCCTGGCAAGTCCGTCGGGAAATCCACGCGCCCAACGCGTGGGCGGCCGGCGGAACCTGCCGGGCTCGGGACCGGAGGACCCAGCGGGTACGTCCGGGGTAGGAGTGAAATCCTGTAATCCCGGACGGACCGCCGATGGCGAAAGCACGCTGGGAAGACGGATCCGACGGTGAGGGACGAAAGCTAGGGTCT |
| p__Halobacteriota_g__Halolamina_ASV_2565 | 0 | 1 | 0 | 0 | 0 | 0 | 0 | 0 | 0 | 0 | 0 | CCTTTACACTGCACGCCAGTGCGATAAGGGGACCCCGAGTGCGAGGGCATATCGTCCTCGCTTTTCTGGACCCTAAGGAGGTCCAGGAACAAGAGCTGGGCAAGACCGGTGCCAGCCGCCGCGGTAATACCGGCAGCTCGAGTGATGACCGATCTTATTGGGCCTAAAGCGTCCGTAGCTGGCCGCACAAGTCCATCGGGAAATCCACCCGCTTAACGGGTGGGCGTCCGGTGGAAACTGTGCGGCTTGGGACAGGAAGGCTCGAGGGGTACGTCCGGGGTAGGAGTGAAATCCCGTAATCCTGGACGGACCGCCGATGGCGAAAGCACCTCGAGAGGACTGCTCCGACAGTGAGGGACGAAAGCTGGGGTCT |
| p__Halobacteriota_g__Halomicrobium_3849_ASV_2640 | 0 | 0 | 0 | 0 | 0 | 1 | 0 | 0 | 0 | 0 | 0 | CCTTTACACTGCACGACAGTGCGATAGGGGGACTCCGAGTGCGAGGGCATATAGTCCTCGCTTTTCTGTACCGTAAGGTGGTACAGGAACAAGTGCTGGGCAAGACCGGTGCCAGCCGCCGCGGTAATACCGGCAGCACGAGTGATGGCCGATATTATTGGGCCTAAAGCGTCCGTAGCCGGCCAGACAAGTCCGTTGGGAAATCGACGCGCTCAACGCGTCGGCGTCCAGCGGAAACTGTCTGGCTTGGGGCCGGAAGACCTGAGGGGTACGTCCGGGGTAGGAGTGAAATCCCGTAATCCTGGACGGACCACCGGTGGCGAAAGCGCCTCAGGAAGACGGACCCGACGGTGAGGGACGAAAGCTAGGGTCT |
| p__Halobacteriota_g__Halorussus_ASV_2553 | 0 | 0 | 0 | 0 | 0 | 0 | 0 | 0 | 0 | 0 | 1 | CCTTTACACTGCACGAAAGTGCGATAAGGGGATCCCGAGTGCGAGGGCATACAGTCCTCGCTTTTGTACACCGTAAGGTGGTGTACGAATAAGAGCTGGGCAAGACCGGTGCCAGCCGCCGCGGTAATACCGGCAGCTCGAGTGATGGCCGCTATTATTGGGCCTAAAGCGTCCGTAGCCGGCCGAACAGGTCCGTTGGGAAATCCACCAGCCTAACTGGTGGGCGTCCAGCGGAAACCAGTCGGCTTGGGGCCGGAAGACCTGAGGGGTACGTCCGGGGTAGGAGTGAAATCCTGTAATCCTGGACGGACCGCCGGTGGCGAAAGCGCCTCAGGAGGACGGACCCGACGGTGAGGGACGAAAGCTAGGGTCA |
| p__Halobacteriota_g__Halorussus_ASV_2568 | 0 | 1 | 0 | 0 | 0 | 0 | 0 | 0 | 0 | 0 | 0 | CCTTTACACTGCACGACAGTGCGATAAGGGGACTCCGAGTGCGAGGGCATATAGTCCTCGCTTTTGTACACCGTAAGGTGGTGTACGAATAAGGACTGGGCAAGACCGGTGCCAGCCGCCGCGGTAATACCGGCAGTCCGAGTGATGGCCGCTATTATTGGGCCTAAAGCGTCCGTAGCCGGCCAGACAGGTCCGTCGGGAAATCTGCTCGCTCAACGAGCAGGCGTCCGGCGGAAACCAGCTGGCTTGGGGCCGGAAGACTCGAGGGGTACGTCCGGGGTAGGAGTGAAATCCCGTAATCCTGGACGGACCACCGGTGGCGAAAGCGCCTCGAGAGGACGGACCCGACGGTGAGGGACGAAAGCTAGGGTCA |
| p__Halobacteriota_g__Halorussus_ASV_2639 | 0 | 0 | 0 | 0 | 0 | 1 | 0 | 0 | 0 | 0 | 0 | CCTTTACACTGCACGACAGTGCGATAAGGGGACTCCGAGTGCGAGGGCATATAGTCCTCGCTTTTCTGTACCGTAAGGTGGTACAGGAATAAGGACTGGGCAAGACCGGTGCCAGCCGCCGCGGTAATACCGGCAGTCCGAGTGATGGCCGCTATTATTGGGCCTAAAGCGTCCGTAGCCGGCCAGACAGGTCCGTCGGGAAATCCACGCGCTCAACGCGTGGGCGTCCGGCGGAAACCAGCTGGCTTGGGGCCGGAAGACCTGAGGGGTACGTCCGGGGTAGGAGTGAAATCCCGTAATCCTGGACGGACCGCCGGTGGCGAAAGCGCCTCAGGAAGACGGACCCGACGGTGAGGGACGAAAGCTAGGGTCA |
| p__Halobacteriota_g__Halosimplex_ASV_2582 | 0 | 0 | 1 | 0 | 0 | 0 | 0 | 0 | 0 | 0 | 0 | CCTTTACACTGCACGCCAGTGCGATAGGGGGACTCCGAGTGCGAGGGCATACAGTCCTCGCTTTTCTGTACCGTAAGGAGGTACAGGAACAAGTGCTGGGCAAGACCGGTGCCAGCCGCCGCGGTAATACCGGCAGCACGAGTGATGGCCGATATTATTGGGCCTAAAGCGTCCGTAGCCGGCCGGACAAGTCCGTTGGGAAATCCACGCGCCCAACGCGTGGGCGTCCAGCGGAAACTGTCCGGCTTGGGGCCGGAGGACTCGAGGGGTACGTCCGGGGTAGGAGTGAAATCCCGTAATCCTGGACGGACCACCGGTGGCGAAAGCGCCTCGAGAGGACGGACCCGACGGTGAGGGACGAAAGCCAGGGTCT |
| p__Halobacteriota_g__Halosimplex_ASV_2583 | 0 | 0 | 1 | 0 | 0 | 0 | 0 | 0 | 0 | 0 | 0 | CCTTTACACTGCACGGAAGTGCGATAAGGGGACCCCAAGTGCGAGGGCATACAGTCCTCGCTTTTCCTGACTGTAGGGCGGTCAGGGAACAAGAGCTGGGCAAGACCGGTGCCAGCCGCCGCGGTAATACCGGCAGCTCGAGTGATGGCCGATATTATTGGGCCTAAAGCGTCCGTAGCCGGCCGGGCAAGTCCGTTGGGAAATCCATCAGCTCAACTGATGGGCGTCCAGCGGAAACTGTTCGGCTTGAGGCCGGAGGACCCGAGGGGTACGTCCGGGGTAGGAGTGAAATCCCGTAATCCCGGACGGACCGCCGGTGGCGAAAGCGCCTCGGGAAGACGGACCTGACGGTGAGGGACGAAAGCCAGGGTCT |
| p__Halobacteriota_g__Halosimplex_ASV_2591 | 0 | 0 | 1 | 0 | 0 | 0 | 0 | 0 | 0 | 0 | 0 | CCTTTACACTGCACGACAGTGCGATAAGGGGACTCCGAGTGCGAGGGCATATAGTCCTCGCTTTTGTCGACCGTAAGGTGGTCGACGAATAAGGACTGGGCAAGACCGGTGCCAGCCGCCGCGGTAATACCGGCAGTCCGAGTGATGGCCGATTTTATTGGGCCTAAAGCGTCCGTAGCTTGCCGAGCAAGTCCGTCGGGAAATCCAGTGGCTCAACCACTGGACGTCCGGCGGAAACTGTTCGGCTTGGGACCGGAAGACTCGAGGAGTACGTCCGGGGTAGGAGTGAAATCCCGTAATCCTGGACGGACTACCGATGGCGAAAGCACCTCGAGAGCACGGATCCGACAGTGAGGGACGAAAGCTAGGGTCT |
| p__Halobacteriota_g__Halostagnicola_ASV_2560 | 0 | 0 | 0 | 0 | 0 | 0 | 0 | 0 | 0 | 0 | 1 | CCTTTACACTGCACGCGAGTGCGATAAGGGGACTCCGAGTGCGAGGGCATATAGTCCTCGCTTTTCACCACCGTAGGAAGGTGGTAGAATAAGTGCTGGGCAAGACCGGTGCCAGCCGCCGCGGTAATACCGGCAGCACGAGTGATGACCGCTATTATTGGGCCTAAAGCGTCCGTAGCTGGCCACGCAAGTCTATCGGGAAATCTCTCCGCTCAACGGAGAGGCGTCCGGTGGAAACTGTGTGGCTTGGGACCGGAAGATCCAGAGGGTACGTCCGGGGTAGGAGTGAAATCCCGTAATCCTGGACGGACCACCGGTGGCGAAAGCGCTCTGGAAAGACGGATCCGACGGTGAGGGACGAAAGCTCGGGTCA |
| p__Halobacteriota_g__Halovenus_ASV_2628 | 0 | 0 | 0 | 0 | 1 | 0 | 0 | 0 | 0 | 0 | 0 | CCTTTACACTGCACGACAGTGCGATAGGGGGACTCCGAGTGCGAGGGCATACAGTCCTCGCTTTTCTGTACCGTAAGGTGGTACAGGAACAAGTGCTGGGCAAGACCGGTGCCAGCCGCCGCGGTAATACCGGCAGCACAAGTGATGACCGATTTCATTGGGCCTAAAGCGTCCGTAGCTGGCCAGACAAGTCTGTTGGGAAATCCACGCGCCCAACGCGTGGGCGTCCAGCGGAAACTGTCTGGCTTGGGGCCGGAGGACTCGAGGGGTACGTCCGGGGTAGGAGTGAAATCCCGTAATCCTGGACGGACCACCGGTGGCGAAAGCGCCTCGAGAGGACGGACCCGACAGTGAGGGACGAAAGCCAGGGTCT |
| p__Nanohaloarchaeota_g__Nanohalobium_ASV_2642 | 0 | 0 | 0 | 0 | 0 | 1 | 1 | 0 | 0 | 0 | 0 | CCTTCACAATGCACGCAAGTGCGATGGGGGGACTCCGAGTGTAGTCTCCTCACAAAGGAGGCTACTTTTCTGTACTGTAAGGAGGTACAGGAACAAGGGGTGGGCAAGACGGCTGCCAGCCGCCGCGGTAATAGCCGCACCCCAAGTGGTGACCAGTATTATTGGGCTTAAAGTGTTCGTAGCCGGCCATGTAAGTCCGCTGTGAAATCCAAGCGCTCAACGTTTGGAACTGCAGTGGATACTGCATGGCTAGGGACCGGACGGCGTCGAGGCTATTCCAGGGGTAGGAGTGAAATCCTAAAATCCCTGGAGGAGCACCTGTGGCGAAAGCGCTCGACGAGTACGGATCCGACGGTGAGGAACGAAAGCTAGGGGAG |
| p__Nanohaloarchaeota_g__Nanohalobium_ASV_2651 | 0 | 0 | 0 | 0 | 0 | 0 | 1 | 0 | 0 | 0 | 0 | CCTTTACAATGCACTAACGTGCGATAAGGGGACTCCGAGTGCAGTCTTCCCACAAGGAAGGCTGCTTTTCTGCACTGTAAGGAGGTGCCGGAACAAGGGGTGGGCAATACTGCTGCCAGCCGCCGCGGTAATAGCAGAACCCCAAGTGGTAGTCGTTATTATTGGGCTTAAAGTGTTCGTAGCCGGATACGCAAGTCCGCTGTGAAATCCACGCGCTCAACGTGTGGGCGTGCAGCGGATACTGCGTGTCTAGGGACCGGACGGCGTCGAGGCTATTCCAGGAGTAGGGGTGAAATCCTAAAATTCCTGGAGGAGCACCTGTGGCGAAAGCGCTCGACGAGTACGGATCCGACGGTGAGGAACGAAAGCCAGGGGAG |
| p__Halobacteriota_g__Natrinema_ASV_2609 | 0 | 0 | 0 | 1 | 0 | 0 | 0 | 0 | 0 | 0 | 0 | CCTTTACACTGCACGAAAGTGCGATAAGGGGACTCCAAGTGCGAGGGCATATAGTCCTCGCTTTTCACTACCGTAAGGTGGTAGTAGAATAAGTGCTGGGCAAGACCGGTGCCAGCCGCCGCGGTAATACCGGCAGCACGAGTGATGACCGCTCTTATTGGGCCTAAAGCGTCCGTAGCTGGCCACGCAAGTCTATCGGGAAATCCGCGCGCTTAACGCGCGGGCGTCCGGTGGAAACTGTGTGGCTTGGGACCGGAAGACCTGAGGGGTACGTCTGGGGTAGGAGTGAAATCCCGTAATCCTGGACGGACCACCGGTGACGAAAGCGCCTCAGGAAGACGGATCCGACGGTGAGGGACGAAAGCTCGGGTCA |

| Phylum_Genus_ASV_number | Presence (1) or Absence (0) in each layer | | | | | | | | | | | |
|---|---|---|---|---|---|---|---|---|---|---|---|---|
| | L1 | L2 | L3 | L4 | L5 | L6 | L7 | L8 | L9 | L10 | L11 | Sequence |
| p__Thermoplasmatota_g__SM1-50_ASV_2656 | 0 | 0 | 0 | 0 | 0 | 0 | 0 | 1 | 0 | 0 | 0 | CCTTCACACTGTGCGAAAGCGCGATGAGGGGATCCCAAGTGCTTGCTCTTAGGAGTAAGCTGTTTTCCTGTTTAAATCGCAGGAAGAGGAAGGGCTGGGTAAGACGGGTGCCAGCCGCCGCGGTAATACCTGCAGCTCAAGTGGTGACCGCTATTATTGGGCCTAAAACGTCCGTAGCCGGCTGAGTAAATGCCTGGGTAAATCGTGTCGCTTAACGATACGAATTCCGGGTAGACTGCTTAGCTTGAGACCGGGAGAAGCTAGAGGTACTCCTGGGGTAGAGGTGAAATTCTGTAATCCTGGGGGGACCACCAGTGGCGAAAGCGTGGGGAG |
| p__Nanoarchaeota_p__Nanoarchaeota_ASV_2671 | 0 | 0 | 0 | 0 | 0 | 0 | 0 | 0 | 1 | 0 | 0 | TCTCCGCAATGCGCGCAAGCGTGACGGGGGAACTCCAAGTGCCCTACCACTGGTAGGGCTGTGGCGAAGCGTAAATAACTTTGCAAGAAAGGGGTGGGCAAGACAGGTGCCAGCAGCCGCGGTAATCCCTGCGCCTCAAGTGGTGACCATTCTTATTGGGCCTAAAGTATCCGTAGCCGGACTATTAAGTCCCTTGTGAAATCGTGCGGCTTAACCGTACGGCGTGCAAGGGATACTGTTAGTCTTGGGACCGGGAGGGGTCAAGGGTATTCCTGGGGTAGCGGTTAAATGTTATAATCCCAGGAGGACCAACGGTGGCGAAGGCGCTTGACTAGAACGGGTCCGACGGTGAATGGATGAAAGCTGGGGGAG |
| p__Halobacteriota_f__Haloarculaceae_3992_ASV_2573 | 0 | 1 | 0 | 0 | 0 | 0 | 0 | 0 | 0 | 0 | 0 | CCTTTACACTGCACGACAGTGCGATAAGGGGACTCCGAGTGCGAGGGCATATAGTCCTCGCTTTTCTGTACCGTAAGGTGGTACAGGAAGAAGTGCTGGGCAAGACCGGTGCCAGCCGCCGCGGTAATACCGGCAGCACGAGTGATGGCCGCTATTATTGGGCCTAAAGCGTCCGTAGCCGGCCGGACAAGTCCGTTGGGAAATCGACGCGCTCAACGCGTCGGCGTCCAGCGGAAACTGTCCGGCTTGGGGCCGGAAGACCTGGGGGGTACGTCCGGGGTAGGAGTGAAATCCCGTAATCCTGGACGGACCACCGGTGGCGAAAGCGCCTCAGGAAGACGGACCCGACGGTGAGGGACGAAAGCTAGGGTCT |
| p__Halobacteriota_f__Haloarculaceae_3992_ASV_2593 | 0 | 0 | 1 | 0 | 0 | 0 | 0 | 0 | 0 | 0 | 0 | CCTTTACACTGCACGACAGTGCGATAAGGGGACTCCGAGTGCGAGGGCATATCGTCCTCGCTTTTCTGTACCGTAAGGTGGTACAGGAACAAGTGCTGGGCAAGACCGGTGCCAGCCGCCGCGGTAATACCGGCAGCACGAGTGATGGCCGCTATTATTGGGCCTAAAGCGTCCGTAGCCGGCCGGACAAGTCCGTTGGGAAATCCACGCGCCTAACGCGTGGGCGTCCAGCGGAAACTGTCCGGCTTGGGGCCGGAGGACCTGAGGGGTACGTCCGGGGTAGGAGTGAAATCCCGTAATCCTGGACGGACCACCGGTGGCGAAAGCGCCTCAGGAGGACGGACCCGACGGTGAGGGACGAAAGCTAGGGTCT |
| p__Halobacteriota_f__Halobacteriaceae_4595_ASV_2595 | 0 | 0 | 1 | 0 | 0 | 0 | 0 | 0 | 0 | 0 | 0 | CCTTTACACTGCACGACAGTGCGATAAGGGAACCCCAAGTGCGAGGGCATATCGTCCTCGCTTTTCTGCACCGTAAGGAGGTGCAGGAATAAGGGCTGGGCAAGACCGGTGCCAGCCGCCGCGGTAATACCGGCAGCCCAAGTGATGGCCGTTATTATTGGGCCTAAAGCGTCCGTAGCCGGCCGGACAAGTCCGTTGGGAAATCCACCCGCTCAACGGGTGGGCGTCCAGCGGAAACTGTTCGGCTTGGGGCCGGAGGACCTGAGGGGTACGTCCGGGGTAGGAGTGAAATCCTGTAATCCTGGACGGACCACCGGTGGCGAAAGCGCCTCAGGAGGACGGACCCGACGGTCAGGGACGAAAGCTAGGGTCT |
| p__Halobacteriota_f__Haloferacaceae_ASV_2574 | 0 | 1 | 0 | 0 | 0 | 0 | 0 | 0 | 0 | 0 | 0 | CCTTTACACTGCACGACAGTGCGATAAGGGGACCCCGAGTGCGAGGGCATACTAGCCCTCGCTTTTTTCGACCGTAAGGTGGTCGACGAAGAAGAGCTGGGCAAGACCGGTGCCAGCCGCCGCGGTAATACCGGCAGCTCGAGTGATGACCGATTTTATTGGGCCTAAAGCGTCCGTAGCTGGCCACGTAAGTCCATCGGGAAATCCACCCGCTCAACGGGTGGGCGTCCGGTGGAAACTACGTGGCTTGGGACCGAAAGACCCAGAGGGTACGTCCGGGGTAGGAGTGAAATCCCGTAATCCTGGACGGACCGCCGATGGCGAAAGCACTCTGGGAGGACGGATCCGACAGTGAGGGACGAAAGCTGGGGTCT |
| p__Halobacteriota_f__Natrialbaceae_ASV_2572 | 0 | 1 | 0 | 0 | 0 | 0 | 0 | 0 | 0 | 0 | 0 | CCTTTACACTGCACGCCAGTGCGATAAGGGGACTCCAAGTGCGAGGGCATACAGTCCTCGCTTTTCACGACCGTAAGGTGGTCGTGGAATAAGAGCTGGGCAAGACCGGTGCCAGCCGCCGCGGTAATACCGGCAGCTCGAGTGATGACCGCTATTATTGGGCCTAAAGCGTCCGTAGCCGGCCAGGTAAGTCCGTCGGGAAATCCGCGCGCCTAACGCGCGGGCGTCCGGTGGAAACTGCCTGGCTTGGGACCGGAAGATCCAGAGGGTACGTCCGGGGTAGGAGTGAAATCCCGTAATCCTGGACGGACCACCGGTGGCGAAAGCGCTCTGGAAAGACGGATCCGACGGTGAGGGACGAAAGCTCGGGTCA |
| p__Halobacteriota_f__Natrialbaceae_ASV_2596 | 0 | 0 | 1 | 0 | 0 | 0 | 0 | 0 | 0 | 0 | 0 | CCTTTACACTGCACGAGAGTGCGATAGGGGGACTCCAAGTGCGAGGGCATACAGTCCTCGCTTTTCTGCACCGTAAGGTGGTGCAGGAATAAGTGCTGGGCAAGACCGGTGCCAGCCGCCGCGGTAATACCGGCAGCACGAGTGATGACCGCTATTATTGGGCCTAAAGCGTCCGTAGCCGGCCGGGCAAGTCCATCGGGAAATCCGCACGCCTAACGTGTGGGCGTCCGGTGGAAACTGCTCGGCTTGGGACCGGAAGACCTGAGGGGTACGTCTGGGGTAGGAGTGAAATCCCGTAATCCTGGACGGACCACCGGTGGCGAAAGCGCCTCAGGAAGACGGATCCGACGGTGAGGGACGAAAGCTTGGGTCA |
| p__Halobacteriota_o__Halanaerobiales_ASV_2559 | 0 | 0 | 0 | 0 | 0 | 0 | 0 | 0 | 0 | 0 | 1 | CCTTTACACTGCACGCAAGTGCGATAAGGGGATCCCAAGTGCGAGGGCGTACAGTCCTCGCTTTTCTGTACCGTAGGGAGGTACAGGAATAAGAGCTGGGCAAGACCGGTGCCAGCCGCCGCGGTAATACCGGCAGCTCGAGTGATGGCCGATATTATTGGGCCTAAAGCGTCCGTAGCCGGCCGGATAAGTCCGTCGGGAAATCCACCCGCTCAACGGGTGGGCGTCCGGCGGAAACTGTCCGGCTTGGGGCCGGAAGGCCAGAGGGGTACGTCCGGGGTAGGAGTGAAATCCTGTAATCCTGGTCGGACCGCCGGTGGCGAAAGCGCCTCTGGAGGACGGACCCGACGGTGAGGGACGAAAGCTTGGGTCA |
| p__Halobacteriota_o__Halanaerobiales_ASV_2567 | 0 | 1 | 0 | 0 | 0 | 0 | 0 | 0 | 0 | 0 | 0 | CCTTTACACTGCGCGAAAGCGCGATAAGGGGACCCCGAGTGCGAGGGCATACAGTCCTCGCTTTTGTGTACCGTAAGGTGGTACACGAATAAGAGCTGGGCAAGACCGGTGCCAGCCGCCGCGGTAATACCGGCAGCTCGAGTGATGGCCGCTATTATTGGGCCTAAAGCGTCCGTAGCCGGCCGAACAGGTCCGTTGGGAAATCCGCCAGCCTAACTGGCGGGCGTCCAGCGGAAACCAGTCGGCTTGGGACCGGAAGACCTGAGGGGTACGTCTGGGGTAGGAGTGAAATCCTGTAATCCTGGACGGACCGCCGGTGGCGAAAGCGCCTCAGGAGGACGGATCCGACGGTGAGGGACGAAAGCTAGGGTCA |
| p__Halobacteriota_o__Halanaerobiales_ASV_2569 | 0 | 1 | 0 | 0 | 0 | 0 | 0 | 0 | 0 | 0 | 0 | CCTTTACACTGTACGAAAGTGCGATAAGGGGACCCCGAGTGCGAGGGCATACAGTCCTCGCTTTTGTGTACCGTAAGGTGGTACACGAATAAGAGCTGGGCAAGACCGGTGCCAGCCGCCGCGGTAATACCGGCAGCTCGAGTGATGGCCGCTATTATTGGGCCTAAAGCGTCCGTAGCCGGCCGAACAGGTCCGTTGGGAAATCCGCCAGCCTAACTGGCGGGCGTCCAGCGGAAACCAGTCGGCTTGGGACCGGAAGACCTGAGGGGTACGTCTGGGGTAGGAGTGAAATCCTGTAATCCTGGACGGACCGCCGGTGGCGAAAGCGCCTCAGGAGGACGGATCCGACGGTGAGGGACGAAAGCTAGGGTCA |
| p__Halobacteriota_o__Halanaerobiales_ASV_2638 | 0 | 0 | 0 | 0 | 0 | 1 | 0 | 0 | 0 | 0 | 0 | CCTTTACACTGCACGACAGTGCGATAGGGGGACTCCGAGTGCGAGGGCATATAGTCCTCGCTTTTCTGGACCGTAAGGAGGTCCAGGAACAAGTGCTGGGCAAGACCGGTGCCAGCCGCCGCGGTAATACCGGCAGCACGAGTGATGGCCAATCTTATTGGGCCTAAAGCGTCCGTAGCCGGCCGGACAAGTCCGTTGGGAAATCCACGCGCTCAACGCGTGGGCGTCCAGCGGAAACTGTCCGGCTTGGGACCGGAAGACCCAGAGGGTACGTCCGGGGTAGGAGTGAAATCCCGTAATCCTGGACGGACCACCGGTGGCGAAAGCGCTCTGGGAAGACGGATCCGACGGTGAGGGACGAAAGCTAGGGTCT |
| p__Halobacteriota_o__Halanaerobiales_ASV_2674 | 0 | 0 | 0 | 0 | 0 | 0 | 0 | 0 | 1 | 0 | 0 | CCTTTACACTGCGCGCAAGCGCGATAAGGGGACCCCAAGTGCGAGGGCATACAGTCCTCGCTTTTCACGACCGTAAGGAGGTCGTCGAATAAGGGCTGGGCAAGACCGGTGCCAGCCGCCGCGGTAATACCGGCAGCCCAAGTGATGGCCGCTCTTATTGGACCTAAAGCGTCCGTAGCTGGCCGTGCAAGTCCGTCGGGAAATCCACCAGCTCAACTGGTGGGCGGCCGACGGAACCTGCATGGCTTGGGACCGGAAGACCCACCGGGTACGTCCGGGGTAGGAGTGAAATCCCGTAATCCCGGACGGACCACCGATGGCGAAAGCACGGTGGGAGGACGGCTCCGACAGTGAGGGACGAAAGCTGGGGTCT |
| p__Nanoarchaeota_o__Woesearchaeales_581_ASV_2533 | 1 | 0 | 0 | 0 | 0 | 0 | 0 | 0 | 0 | 0 | 1 | CCTCCACAATGCACGAAAGTGTGATGGGGGGATCCTACGTGCTCATGTTTAACATGGGCTTTTGCCAAGAGTAGATACCTTGGCGAATAAGGGGTGGGCAAGACCGGTGCCAGCCGCCGCGGTAACACCGGCGCCCCAAGTGGCGATCGCGATTATTGGGCCTAAAGCGTCCGTAGCCGGATGAGTAAATCCGCTGTGAAATCGTTGCGCTTAACGTAACGGCGTGCAGTGGACACTGCTTATCTTGAGACCGGGAGGAGTCAGACGTATGCCGTGGGGACTGGTAAAATGGTATAATCCACGGTAGACGACCGATGGCGAAGGCATCTGACCAGAACGGATCTGACGGTGAGGGACGAAAGCCAGGGGAG |
| p__Thermoplasmatota_p__Thermoplasmatota_ASV_2576 | 0 | 1 | 0 | 0 | 0 | 1 | 0 | 0 | 0 | 0 | 0 | ACTTCGCAATGTGGGAAACCACGACGAGGGAACTCCAAGTGCCGGCGCAGAGCGTCGGCTGTTTCCAAGCCTAAAAAGCTTGGAGAGTAAGGGCTGGGTAAGACGGGTGCCAGCCGCCGCGGTAACACCTGCAGCTCAAGTGGTCGCCGCTATTATTGGGCCTAAAGCGTCCGTAGCCGGCCCGGTAAATGCCTGGGTAAATCGACTGGCTTAACCAGTCGAATTCCGGGTAGACTGCCGGGCTTGGGACCGGGAGAGTCACGAGGTACTCGGGGGGTAGAGGTGAAATTCTGTAATCCTCCGGGGACCACCGGTGGCGAAGGCGTCGTGATAGAACGGCTCCGACGGTGAGGGACGAAGCCCTGGGGCG |
| p__Iainarchaeota_o__Iainarchaeales_ASV_2679 | 1 | 0 | 0 | 0 | 0 | 0 | 0 | 1 | 0 | 0 | 0 | CCTTTACAATGCACGAAAGTGTGATAGGGGGATTTCGAGTGCTCTCTTTGAGAGCTTTTGCCAAACCCAAACCGTTTGGCGAATAAGTGGAGGGCAAGATACGTGCCAGCCGCCGCGGTAACACGTACTCCGCAAGTGGTATCCACTAATATTGGGCCTAAAACGTCTGTAGCCGGTTCAACAAGTCCTTGGTGAAATCATTGGGCTTAACTTAATGAATTGCCAAGGAAACTGCTGGACTCGAGGCCGGGAGACGCCAAGGGAATTTTTGGGGTAACGGTGAAATGTTATGATCCTGAAAAGACCACCAGTAGCGAAGGCGCTTGGCGAGCACGGGCCTGACGGTGAGAGACGAAAGCTGGGGGAG |
| p__Nanoarchaeota_g__BMS3ABIN17_ASV_2805 | 0 | 0 | 0 | 0 | 0 | 0 | 0 | 1 | 1 | 0 | 0 | ATTCTGCAATGCGCGCAAGCGTGACAGAGCAAGTCAGAGTGCTTTACCATTAGGTAAAGCTTTTGCCAGATGTAAAAAGTCTGGCGAATAAGGACTGGGTAAGACTGGTGCCAGCCGCCGCGGTAATACCAGCAGTCCAAGTGGCAGCCGCTTTTGTTGGGTCTAAAACATCCGTAGCTTGTCACGTAAGTCTTTTGTGAAATCCGGCATCTCAAGTGTCGGGCGTGCAAAAGATACTGCGTGGCTAGAGACCGGGAGACGTAAGAAGTACGCAAGGGGTAGCGGTAAAATGCGTTGATCCTCTGTGGACTAACGATAGCGAAGGCATCTTACGAGAACGGATCTGACAGTGAGGGATGAAGGCTAGGGGCG |
| p__Nanoarchaeota_g__BMS3ABIN17_ASV_2806 | 0 | 0 | 0 | 0 | 0 | 0 | 0 | 0 | 1 | 0 | 0 | CTTTTGCAATGCGCGTAAGCGTGACAAAGCAAGCCAGAGTGTTTCTCTATCAGAGAAACTTTTGTATGATGCAAAAAGTCATACGAATAAGGACTGGGCAAGACTGGTGCCAGCCGCCGCGGTAATCCCAGCGGTCCAAGTCGCAGCCACTTTTATTGGGTCTAAAACATCCGTAGCTTGTTCTGTAAGTCTCTTGTGAAATCTTACATCTTAAGTGTAAGGCGTGCGGGAGATACTGCAGGACTAGAGACCGGAAGACGCAAGGAGTACGCCTAAGGAAGTGGTAAAATACGTTAATCTTGGGTGGACTAACAATGGCGAAGGCACCTTGCGAGGACGGATCTGACAGTGAGGGATGAAGGCTAGGGGCG |
| p__Nanoarchaeota_g__BMS3ABIN17_ASV_2811 | 0 | 0 | 0 | 0 | 0 | 0 | 0 | 0 | 1 | 0 | 0 | CTTTTGCAATGCGCGAAAGCGTGACAGAGCGAGTCGAAGTGCTTCTCAATAGAGAAGCTTTTGCCAAATGTAAAAAGTTTGGCGAATAAGGACTGGGTAAGACGAGTGCCAGCCGCCGCGGTAATACTCGCGGTCCAAGTCGCAGCCATTTTTATTGGGTCTAAAACATCCGTAGCTTGCTTAGTAAGTTTCTTGTGAAATCCGGCTTCCTAAGAGTCGGGCGCGCAGGAAATACTACTAAGCTAGAGACTGGGAGACGTAAGGAGTACACTCGAAGTAGCGGTAAAATGTGTTAATCTCGGGTGGACTATCAACAGCGAAGGCATCTTACGAGAACAGTTCTGACAGTAAGGGATGAAAGCTAGGGGCG |
| p__Nanoarchaeota_g__BMS3ABIN17_ASV_2816 | 0 | 0 | 0 | 0 | 0 | 0 | 0 | 0 | 1 | 0 | 0 | TTTCTGCAATGCGCGAAAGCGTGACAGAGCGAGTCAGAGTGCTTTTCTATTTGAAAAGCTTTTGCTAAATGTAAAAAGTTTGGCGAATAAGGACTGGGTAAGACGAGTGCCAGCCGCCGCGGTAATACTCGCGGTCCAAGTCGCAGCCATTTTTATTGGGTCTAAAATATCCGTAGCTTGTTTGAAAAGTTTCTTGTGAAATCTCATCTCTCAAGGATGTGGCGAGCGAGAAATACTCTCAAACTAGAGACTGGAAGACGTAAGAAGTATGTTCGAAGTAGCGGTAAAATGTGTTAATCTCGGACAGACTAACAACAGCGAAGGCATCTTACGAGGACAGTTCTGACAGTTAAGGATGAAGGCTAGGGGCG |
| p__Thermoplasmatota_g__EX4484-6_ASV_2680 | 0 | 0 | 0 | 0 | 0 | 0 | 0 | 0 | 0 | 1 | 0 | ACTTCGCAATGCGCGCAAGCGCGACGAGGGGATTCCAAGTGCATGCACTCAGTGTATGCTGTTTGCTAGTTTAAAACGCTAGCAGAGTAAGGGCTGGGTAAGACTGGTGCCAGCCGCCGCGGTAATACCAGCGGCCCTAGTGGTGACCGTTATTATTGGGCCTAAAGCGTCCGTAGCCGGCCCGGTAAATCTCTGGGTAAATCGATGGGCTTAACCCATCGAATTCTGGGGAGACTGCTGGGCTTGGGACCGGGAGAGGTCGGAGGTACTACAGGGGTAGGGGTGAAATCCTGTAATCCCTGTGGGACCACCGGTGGCGAAGGCGTCCGACCAGAACGGGTCCGACGGTAAGGGACGAAGCCCTGGGGCG |
| p__Thermoplasmatota_g__EX4484-6_ASV_2733 | 0 | 1 | 0 | 1 | 0 | 0 | 0 | 0 | 0 | 0 | 0 | ACTTCGCAATGCGCGCAAGCGCGACGAGGGGATTCCAAGTGCATGCACTCAGTGTATGCTGTTTTCCAGTCTAAAAAGCTGGGAGAGTAAGGGCTGGGTAAGACTGGTGCCAGCCGCCGCGGTAATACCAGCGGCCCTAGTGGTGATCGTTATTATTGGGCCTAAAGCGTCCGTAGCCGGCTCGGTAAATCTCTGGGTAAATCGTTGGGCTTAACTCAGCGAATTCTGGGGAGACTGCCGGGCTTGGGACCGGGAGAGGTCGGAGGTACTCCAGGGGTAGGGGTGAAATCCTGTAATCCTTGGGGGACCACCGGTGGCGAAAGCGTCCGGCCAGAACGGGTCCGACGGTAAGGGACGAAGCCCTGGGGCG |

| Phylum_Genus_ASV_ number | Presence (1) or Absence (0) in each layer | | | | | | | | | | | |
|---|---|---|---|---|---|---|---|---|---|---|---|---|
| | L1 | L2 | L3 | L4 | L5 | L6 | L7 | L8 | L9 | L10 | L11 | Sequence |
| p__Thermoproteota_g__GCA-2726865_ASV_2682 | 0 | 0 | 0 | 0 | 0 | 0 | 0 | 0 | 0 | 1 | 0 | CCTCCTCAATACACTAACGTGTGAAAGGGTTACCTCGAGTGCTACCTGAAGAAGGTGGCTTTTCCTTGGTCTAAAAAGCCTTGGGAATAAGAGGGGGGCAAGACTGGTGTCAGCCGCCGCGGTAACACCAGCCCCTCGAGTGGTAGGGATGAT<br>TATTTGGCCTAAAGCGTCCGTAGCCTGCCGAGTAAGTTTTCCGTTAAATCCAGCGTCTCAAGCGTTGGACTGCGGGAAATACTGCTTGGCTAGAGAGTGGGAGAGGTAAGCGGTATTCTGGGGGTAGGGGCGAAATCCTTTGATCCCCGGAGG<br>ACCACCAGTGGCGAAGGCTGCTTACTAGAACACGTCTGACGGTCAGGGACGAAAGCTGGGGTAG |
| p__Halobacteriota_g__Haloprofundus_ASV_2710 | 0 | 0 | 0 | 0 | 0 | 0 | 0 | 0 | 0 | 0 | 1 | CCTTTACACTGCACGCCAGTGCGACAAGGGGACCCCGAGTGCGAGGGCATACAGTCCTCGCTTTTCCCGACCGTAAGAAGGTCGGAGAAGAAGAGCTGGGCAAGACCGGTGCCAGCCGCCGCGGTAATACCGGCAGCTCGAGTGATGACCGCT<br>GTTATTGGGCCTAAAGCGTCCGTAGCTGGCCGCGTAAGTCCATCGGGAAATCCATCCGCTCAACGGATGGGCGTCCGGTGGAAACTACACGGCTTGGAGCCGGAAGACCCAAGGGGTACGTCCGGGGTAGGAGTGAAATCCCGTAATCCTGGA<br>CGGACCACCGATGGCGAAAGCACCTTGGGAAGACGGACTCGACAGTGAGGGACGAAAGCTAGGGTCT |
| p__Halobacteriota_g__Halorhabdus_ASV_2787 | 0 | 0 | 0 | 0 | 0 | 1 | 0 | 0 | 0 | 0 | 0 | CCTTTACACTGCACGACAGTGCGATAGGGGGACTCCGAGTGCGAGGGCATATAGTCCTCGCTTTTCTGTACCGTAAGGAGGTAGAAGAATAAGGGCTGGGCAAGACCGGTGCCAGCCGCCGCGGTAATACCGGCAGCCCGAGTGATGGCCGCT<br>ATTATTGGGCCTAAAGCGTCCGTAGCCGGCCAGACAAGTCCGTTGGGAAATCCACGCGCTCAACGCGTGGACGTCCGGCGGAAACTGCCTGGCTTGGGGCCGGAAGACCTGAGGGGTACGTCCGGGGTAGGAGTGAAATCCCGTAATCCTGGA<br>CGGACCGCCGGTGGCGAAAGCGCCTCAGGAGGACGGACCCGACGGTGAGGGACGAAAGCTAGGGTCT |
| p__Halobacteriota_g__Halorubrum_ASV_2720 | 0 | 1 | 0 | 0 | 0 | 0 | 0 | 0 | 0 | 0 | 0 | CCTTTACACTGCACGCCAGTGCGATAAGGGAATCCCAAGTGCACAGGCATAGCGCCTGTGCTTTTGTACACCCTAGGGTGGTGTACGAATAAGGGCTGGGCAAGACCGGTGCCAGCCGCCGCGGTAATACCGGCAGCCCGAGTGATGGCCGAT<br>CTTATTGGGCCTAAAGCGTCCGTAGCTGGCCGCGCAAGTCCATCGGGAAATCCACCTGCTCAACAGGTGGGCGCCCGGTGGAAACTGCGTGGCTTGGGACCGGAAGGCGCGACGGGTACGTCCGGGGTAGGAGTGAAATCCTGTAATCCTGGA<br>CGGACCGCCGATGGCGAAAGCACGTCGCGAGGACGGATCCGACAGTGAGGGACGAAAGCCAGGGTCT |
| p__Halobacteriota_g__Halorussus_ASV_2741 | 0 | 0 | 1 | 0 | 0 | 0 | 0 | 0 | 0 | 0 | 0 | ACTTTACACTGCACGACAGTGCGATAAGGGAACTCCGAGTGCGAGGGCATATAGTCCTCGCTTTTGTACACCGTAGGGAGGTGTACGAATAAGGGCTGGGCAAGACCGGTGCCAGCCGCCGCGGTAATACCGGCAGCCCGAGTGATGGCCGCT<br>ATTATTGGGCCTAAAGCGTCCGTAGCCAGCCAGACAGGTCCGTCGGGAAATCCATCCGCTTAACGGATGGGCGTCCGGCGGAAACCAGCTGGCTTGGGGCCGGAAGACCTGAGGGGTACGTCCGGGGTAGGAGTGAAATCCCGTAATCCTGGA<br>CGGACCGCCGGTGGCGAAAGCGCCTCAGGAGGACGGACCCGACGGTGAGGGACGAAAGCTAGGGTCA |
| p__Halobacteriota_g__Halorussus_ASV_2807 | 0 | 0 | 0 | 0 | 0 | 0 | 0 | 0 | 1 | 0 | 0 | ACTTTACACTGCACGACAGTGCGATAAGGGAACTCCGAGTGCGAGGGCATATAGTCCTCGCTTTTGTACACCGTAAGGTGGTGTACGAATAAGAGCTGGGCAAGACCGGTGCCAGCCGCCGCGGTAATACCGGCAGCTCGAGTGATGGCCGCT<br>ATTATTGGGCCTAAAGCGTCCGTAGCCGGCCAGACAGGTCTGTCGGGAAATCCACTCGCTCAACGAGTGGGCGTCCGGCGGAAACCAGCTGGCTTGGGGCCGGAAGATCTGAGGGGTACGTCCGGGGTAGGAGTGAAATCCTGTAATCCTGGA<br>CGGACCGCCGGTGGCGAAAGCGCCTCAGAAAGACGGACCCGACGGTGAGGGACGAAAGCTAGGGTCA |
| p__Halobacteriota_g__Halosimplex_ASV_2753 | 0 | 0 | 1 | 0 | 0 | 0 | 0 | 0 | 0 | 0 | 0 | CCTTTACACTGCACGACAGTGCGATAGGGGGACTCCGAGTGCGAGGGCATATAGCCCTCGCTTTTCTGTACCGTAAGGTGGTACAGGAACAAGTGCTGGGCAAGACCGGTGCCAGCCGCCGCGGTAATACCGGCAGCACGAGTGATGGCCGAT<br>ATTATTGGGCCTAAAGCGTCCGTAGCCGGCCGAGCAAGTCCGTTGGGAAATCCACGCGCTCAACGCGTGGGCGTCCAGCGGAAACTGTTCGGCTTGGGGCCGGAAGACTCGAGGGGTACGTCCGGGGTAGGAGTGAAATCCCGTAATCCTGGA<br>CGGACCACCGGTGGCGAAAGCGCCTCGAGAAGACGGACCCGACGGTGAGGGACGAAAGCTTGGGTCT |
| p__Halobacteriota_g__Halovenus_ASV_2740 | 0 | 0 | 1 | 0 | 0 | 0 | 0 | 0 | 0 | 0 | 0 | CCTTTACACTGCACGACAGTGCGATAGGGGGACTCCGAGTGCGAGGGCATATAGTCCTCGCTTTTCTGTACCGTAAGGTGGTACAGGAACAAGTGCTGGGCAAGACCGGTGCCAGCCGCCGCGGTAATACCGGCAGCACAAGTGATGGCCGAT<br>ATTATTGGGCCTAAAGCGTTCGTAGCCGGCCGGACAAGTTCGTTGGGAAATCCACGCGCTCAACGCGTGGGCGTCCAGCGAATACTGTCTGGCTTGGGGCCGGAAGACTCGAGGGGTACGTCCGGGGTAGGAGTGAAATCCCGTAATCCTGGA<br>CGGACCACCGATGGCGAAAGCACCTCGAGAGGACGGACCCGACGGTGAGGGACGAAAGCCAGGGTCT |
| p__Halobacteriota_g__Halovenus_ASV_2750 | 0 | 0 | 1 | 0 | 0 | 0 | 0 | 0 | 0 | 0 | 0 | CCTTTACACTGCACGACAGTGCGATAGGGGGACTCCGAGTGCGAGGGCATACAGTCCTCGCTTTTGTACACCGTAAGGTGGTGTACGAATAAGAACTGGGCAAGACCGGTGCCAGCCGCCGCGGTAATACCGGCAGTTCGAGTGATGGCCGCT<br>TTTATTGGGCCTAAAGCGTTCGTAGCTGGCCGAGCAAGTCTGTCGGGAAATCCAGTGGCTTAACCACTGGACGTCCGGCGGAAACTGCTCGGCTTGGGACCGGGAGACTCGAGAAGTACGTCCGGGGTAGGAGTGAAATCCCGTAATCCTGGA<br>CGGACTACCGATGGCGAAAGCATCTCGAGAGCACGGATCCGACAGTGAGGAACGAAAGCTGGGGTCT |
| p__Halobacteriota_g__Halovenus_ASV_2754 | 0 | 0 | 1 | 0 | 0 | 0 | 0 | 0 | 0 | 0 | 0 | CCTTTACACTGCACGACAGTGCGATAGGGGGACTCCGAGTGCGAGGGCATATAGTCCTCGCTTTTCTGTACCGTAAGGTGGTACAGGAACAAGTGCTGGGCAAGACCGGTGCCAGCCGCCGCGGTAATACCGGCAGCACGAGTGATGGCCGAT<br>ATTATTGGGCCTAAAGCGTTCGTAGCCGGCCGGACAAGTTCGTTGGGAAATCCACGCGCTCAACGCGTGGGCGTCCAGCGAATACTGTCTGGCTTGGGGCCGGAAGACTCGAGGGGTACGTCCGGGGTAGGAGTGAAATCCCGTAATCCTGGA<br>CGGACCACCGATGGCGAAAGCACCTCGAGAGGACGGACCCGACGGTGAGGGACGAAAGCCAGGGTCT |
| p__Halobacteriota_g__Halovenus_ASV_2792 | 0 | 0 | 0 | 0 | 0 | 1 | 0 | 0 | 0 | 0 | 0 | CCTTTACACTGCACGACAGTGCGATAGGGGGACTCCGAGTGCGAGGGCATATAGTCCTCGCTTTTCTGTACCGTAAGGTGGTACAGGAACAAGTGCTGGGCAAGACCGGTGCCAGCCGCCGCGGTAATACCGGCAGCACAAGTGATGGCCGAT<br>TTTATTGGGCCTAAAGCGTTCGTAGCCGGCCGGACAAGTTCGTTGGGAAATCCACGCGCTTAACGCGTGGGCGTCCAGCGAATACTGTTCGGCTTGGGGCCGGAAGACTCGAGGGGTACGTCCGGGGTAGGAGTGAAATCCCGTAATCCTGGA<br>CGGACCACCGATGGCGAAAGCACCTCGAGAGGACGGACCCGACGGTGAGGGACGAAAGCCAGGGTCT |
| p__Thermoproteota_g__Nitrososphaera_ASV_2786 | 0 | 0 | 0 | 0 | 0 | 1 | 0 | 0 | 0 | 0 | 0 | CCTCTGCAATAGGCGAAAGCTTGACAGGGTTACTCTGAGTGATGCCCGCTAAGGGTATCTTTTGGCACCTCTAAAAATGGTGCAGAATAAGGGGTGGGCAAGTCTGGTGTCAGCCGCCGCGGTAATACCAGCACCCCGAGTTGTCGGGACGAT<br>TATTGGGCCTAAAGCATCCGTAGCCTGTTCTGCAAGTCCTCCGTTAAATCCACCTGCTCAACGGATGGGCTGCGGAGGAAACCGCAGAGCTAGGAGGCGGGAGAGGCAAACGGTACTCAGTGGGTAGGGGTAAAATCCATTGATCTACTGAAG<br>ACCACCAGTGGCGAAGGCGGTTTGCCAGAACGCGCTCGACGGTGAGGGATGAAAGCTGGGGGAG |
| p__Halobacteriota_g__Saliphagus_ASV_2790 | 0 | 0 | 0 | 0 | 0 | 1 | 0 | 0 | 0 | 0 | 0 | CCTTTACACTGCACGACAGTGCGATAAGGGGATTCCGAGTGCGAGGGCATACAGTCCTCGCTTTTGTGTACCGTAGGAAGGTACACGAATAAGGGCTGGGCAAGACCGGTGCCAGCCGCCGCGGTAATACCGGCAGCCCGAGTGATGACCGAT<br>ATTATTGGGCCTAAAGCGTCCGTAGCCGGCTACGCAAGTCCGTCGGGAAATCTGCTCGCTCAACGAGTAGGCGTCCGGCGGAAACTGCGTGGCTTGGGACCGGAAGACTCGAGGGGTACGTCCGGGGTAGGAGTGAAATCCCGTAATCCTGGA<br>CGGACCACCGGTGGCGAAAGCGCCTCGAGAGGACGGATCCGACGGTGAGGGACGAAAGCTAGGGTCA |
| p__Thermoplasmatota_g__SM1-50_ASV_2747 | 0 | 0 | 1 | 0 | 0 | 0 | 0 | 0 | 0 | 0 | 0 | CCTTTACACTGCACGAAAGTGCGATAGGGGGACCCCGAGTGCGAGGGCATACAGTCCTCGCTTTTGTGTACTGTAAGGAGGTACACGAATAAGAGCTGGGCAAGACCGGTGCCAGCCGCCGCGGTAATACCTGCAGCTCAAGTGGTGACCGCT<br>ATTATTGGGCCTAAAACGTCCGTAGCCGGCTGAGTAAATGCCTGGGTAAATCGTGTCGCTTAACGATACGAATTCCGGGTAGACTGCTTAGCTTGAGACTGGGAGAAGCTAGAGGTACTCCTGGGGTAGAGGTGAAATTCTGTAATCCTGGGG<br>GGACCACCAGTGGCGAAAGCGTCTAGCTAGAACGGGTCTGACGGTGAGGGACGAAACCCTGGGGCG |
| p__Halobacteriota_f__Haladaptataceae_ASV_2742 | 0 | 0 | 1 | 0 | 0 | 0 | 0 | 0 | 0 | 0 | 0 | CCTTTACACTGCACGAAAGTGCGATAAGGGGACCCCGAGTGCGAGGGCATACAGTCCTCGCTTTTGTGTACCGTAAGGAGGTACACGAATAAGAGCTGGGCAAGACCGGTGCCAGCCGCCGCGGTAATACCGGCAGCTCAAGTGATGGCCGCT<br>GTTATTGGGCCTAAAGCGTCCGTAGCCTGCCGAACAGGTCCGTTGGGAAATTCATCTGCTTAACAGATGGGCGTCCAGCGGAAACCAGTCGGCTTGGGGCCGGAAGACCCGAGGGGTACGTCCGGGGTAGGAGTGAAATCCTGTAATCCTGGA<br>CGGACCGCCGGTGGCGAAAGCGCCTCGGGAGGACGGACCCGACGGTGAGGGACGAAAGCTAGGGTCA |
| p__Halobacteriota_f__Haloferacaceae_ASV_2711 | 0 | 0 | 0 | 0 | 0 | 0 | 0 | 0 | 0 | 0 | 1 | CCTTTACACTGCACGCGAGTGCGATAAGGGGACTCCGAGTGCGAGGGCATATCGTCCTCGCTTTTCTGTACCGTAGGGCGGTACAGGAACAAGAGCTGGGCAAGACCGGTGCCAGCCGCCGCGGTAATACCGGCAGCTCAAGTGATGGCCGAT<br>CTTATTGGGCCTAAAGCGTCCGTAGCCGGCCCTGCAAGTCCGTCGGGAAATCCACGCGCTCAACGCGTGGGCGTCCGGCGGAAACTCCAGGGCTTGGGACCGGAAGGCTCGAAGGGTACGTCCGGGGTAGGAGTGAAATCCTGTAATCCCGGA<br>CGGACCACCGATGGCGAAAGCACTTCGAGAAGACGGATCCGACGGTGAGGGACGAAAGCCAGGGTCT |
| p__Halobacteriota_f__Haloferacaceae_ASV_2795 | 0 | 0 | 0 | 0 | 0 | 0 | 1 | 0 | 0 | 0 | 0 | CCTTTACACTGCACGCCAGTGCGATAGGGGGACCCCGAGTGCGAGGGCATACAGTCCTCGCTTTTGCACACCGTAAGGTGGTGCGCGAATAAGTGCTGGGCAAGACCGGTGCCAGCCGCCGCGGTAATACCGGCAGCACAAGTGATGGCCGCT<br>TTTATTGGGCCTAAAGCGTCCGTAGCCGGCCGAACAAGTCCGTCGGGAAATCCATCCGCTCAACGGATGGGCGTCCGGCGGAAACTGTTCGGCTTGGGACAGGAAGACTCGAGGGGTACGTCCGGGGTAGGAGTGAAATCCTGTAATCCTGGA<br>CGGACCGCCGATGGCGAAAGCACCTCGAGAAGACTGATCCGACGGTGAGGGACGAAAGCTGGGGTCT |
| p__Halobacteriota_f__Natrialbaceae_ASV_2808 | 0 | 0 | 0 | 0 | 0 | 0 | 0 | 0 | 1 | 0 | 0 | CCTTTACACTGCACGCCAGTGCGATAAGGGGACTCCATCTGCGAGGGCATATAGTCCTCGCTTTTCTCGACCGTAGGGTGGTCGAGGAATAAGTGCTGGGCAAGACCGGTGCCAGCCGCCGCGGTAATACCGGCAGCACGAGTGATGACCGCT<br>ATTATTGGGCCTAAAGCGTCCGTAGCTGGCCAGGCAAGTCCATCGGGAAATCCGCGTGCTCAACGCGCGGGCGTCCGGTGGAAACTGCTTGGCTTGGGACCGGAAGACCTAGGGGGTACGTCTGGGGTAGGAGTGAAATCCCGTAATCCTGGA<br>CGGACCACCGGTGGCGAAAGCGCCCTAGGAAGACGGATCCGACGGTGAGGGACGAAAGCTCGGGTCA |
| p__Nanoarchaeota_o__Woesearchaeales_581_ASV_2760 | 0 | 0 | 0 | 1 | 0 | 0 | 0 | 0 | 0 | 0 | 0 | CCTCAGCAATGCACGAGAGTGTGACTGGGGGATTCCGAGTGCTCATGTAACGCATGGGCTTTTGCCAAAGGTAAGTACTTTGGCGAATAAGTGGGGGGTAATACTGGTGCCAGCCGCCGCGGTAACACCAGAGCCACGAGTGGGGACCGCTAT<br>TATTGGGCCTAAAGCGTTCGTAGCCGGCCAGGTAAATCTCCTGTGAAATCGCAGCGCTCAACGTTGCGGCGTGCAGGAGAGACTGCCTGGCTTGGGACCGGGAGGCGTATGAGGTATTCTGGGGGTAGCGGTAAAATGCGATAATCCTCAGAG<br>GACCACCTGTGGCGAAGGCGTCATACGAGAACGGATCCGACGGTGAGGGACGAAAGCTAGGGGAG |
| p__Thermoplasmatota_p__Thermoplasmatota_ASV_2799 | 0 | 0 | 0 | 0 | 0 | 0 | 0 | 1 | 0 | 0 | 0 | CCTTCACACTGTGCGAAAGCGCGATGAGGGGATCCCAAGTGCTTGCTCTTAGGAGTAAGCTGTTTTCCTGTTTAAATCGCAGGAAGAGGAAGGGCTGGGTAAGACGGGTGCCAGCCGCCGCGGTAATACCTGCAGCTCAAGTGGTGACCGCTA<br>TTATTGGGCCTAAAACGTCCGTAGCCGGCTGAGTAAATGCCTGGGTAAATCGTGTCGCTTAACGATACGAATTCCGGGTAGACTGCTTAGCTTGAGACTGGGAGAAGCTAGAGGTACTCCTGGGGTAGAGGTGAAATTCTGTAATCCTGGGGG<br>GACCACCAGTGGCGAAGGCGACTTCCTGGACCAATACTGACGCTGAGGCGCGAAGGCGTGGGTAG |
| d__Archaea_d__Archaea_ASV_2709 | 0 | 0 | 0 | 0 | 0 | 0 | 0 | 0 | 0 | 0 | 1 | CCTTTACAATGCACGAAAGTGTGATAAGGGGATTCCAAGTGCTCATGCACAGCATGGGCTTTTCTTAAGAGTAGGTATCTTGAGGAATAAGTGGTGGGAAATACCGGTGCCAGCAGCCGCGGTAACCCCGGAGCCACAAGTGGCAATCACAAT<br>TATTGGGTCTAAAGCGTCCGTAGTCGGCCTGATCAATCTCTTGTGAAATCGTGGAGCCTAACTCCACGGCGTGCAGGAGATATTGTCAGGCTAGGGACCGGAAGGAGTGTGAGGTATTCCAAAGGGAGCGGTAAAATGTGATAATCTTTGGAA<br>GACCACCGGTGGCGAAGGCGTCACACTAATACGGATCCGACGATGAGGGACGAAGGCCAGGGGAG |
| d__Archaea_d__Archaea_ASV_2763 | 0 | 0 | 0 | 1 | 0 | 0 | 0 | 0 | 0 | 0 | 0 | AATCTACAATGCACGAAAGTGTGATAGTTTGAATCGGAGTGCGCTTCAATTTGAAGTGCTTTTCTGTACTGTAAAAAGGTATAGGAATAAGGACTGGGTAAGACGAGTGCCAGCCGCCGCGGTAATACTCGCGGTCCAAGTCGCAGCCATTTT<br>TATTGGGTCTAAAACATCCGTAGCTTGCCAGGTGAGTTTCTCGTGAAATCCCACATCTTAAGTGTGGGGCGGGCGGGAAGTACTGCCTGGCTAGAGACTGGAAGGCGTAACGAGTACGTTCAAAGTAGCGGTAAAATGTGTTGATCTTGGACG<br>GACTAACGATTAGCGAAGGCACGTTACGAGAACAGTTCTGACAGTAAGGGATGAAGGCCAGGGGCG |
| p__Nanoarchaeota_g__BMS3ABIN17_ASV_2827 | 1 | 0 | 0 | 0 | 0 | 0 | 0 | 0 | 0 | 0 | 0 | CTTCTACAATGCACGAAAGTGTGATAGAGTGAGCCAGAGTGCTTTTCAATGAAAAGCTTTTGCTAAGTGTAAAAAGCTTGGCGAATAAGGACTGGGTAAGACTAGTGCCAGCCGCCGCGGTAATACTAGCGGTCCAAGTCGCAGCCATTTTTA<br>TTGGGTCTAAAACATCCGTAGCTTGCTTAATAAGTTTCTTGTGAAATTTCATATCTCAAGTATGAAGCGTGCGGGAAATACTGTTAAGCTAGAGACTGGAAGACGTAACGAGTACGATCAAGGTAGCGGTAAAATGTGTTAATCTTGGCCGGA<br>CTAACAACAGCGAAGGCACGTTACGAGGACAGTTCTGACAGTAAGGGATGAAGGCTAGGGGCG |
| p__Nanoarchaeota_g__BMS3ABIN17_ASV_2835 | 1 | 0 | 0 | 0 | 0 | 0 | 0 | 1 | 1 | 0 | 0 | CATTTACAATGCACGAAAGTGTGATAATGCGAACCAAAGTGTTTTTCGTATAGAAAAACTTTTGTTCACTGTAAAAATGTGGACGAATAAGGACTGGGTAAGACTGGTGCCAGCCGCCGCGGTAATCCCAGCAGTCCAAGTGGCATCCAACAT<br>TATTGGGTCTAAAACATCCGTAGCTTGTTACTTAAGTCTCTTGTTAAATCCTATGTCTCAAGCATGGGTCCTGCAGGAGATACTGGGTAGCTAGAGATCGGTAGACGTAAGGAGTACGCCGGAGGTAGTGGTAAAATACGTTAATCTTCGGTG<br>GACTAACGATGAGCGAAGGCACCTTACGAGGACGAATCTGACAGTGAGGGATGAAGGCCAGGGGCG |

| | Presence (1) or Absence (0) in each layer | | | | | | | | | | |
|---|---|---|---|---|---|---|---|---|---|---|---|
| **Phylum_Genus_ASV_ number** | **L1** | **L2** | **L3** | **L4** | **L5** | **L6** | **L7** | **L8** | **L9** | **L10** | **L11** | **Sequence** |
| p__Nanoarchaeota_g__CG1-02-57-44_ASV_2869 | 0 | 0 | 0 | 0 | 0 | 0 | 0 | 0 | 0 | 0 | 1 | CCTTTACAATGCACGCAAGTGTGATAAGGGAATCCTTAGTGCTTTGCTTTTGCAAAGCTTTTGCCAAGTGTAAAAAGCTTGGCGAATAAGTGGTGGGTAAGACGGGTGCCAGCCGCCGCGGTAACACCTGCGCCACGAGTGGCAACCGGTTTTATTGGGTCTAAAGCGTCCGTAGCCGGCCTTGTAAATTCTTTGTGAAATCGTCATGCTTAACATGACGGCGTGCGAGGAAGACTGCAAGGCTTGGGACCGGGAAGAGTCAGAGGTATTTTAGGGGGAGCGGTAAAATGCTATAATCCCTAAAGGACCACCTGTGGCGAAAGCGTCTGACTGGAACGGATCCGACGGTGAGGGTCGAAAGCCAGGGGAG |
| p__Altiarchaeota_g__EX4484-2_ASV_2831 | 1 | 0 | 0 | 0 | 0 | 0 | 0 | 0 | 0 | 1 | 0 | CCTCTGCAATGCACGAAAGTGTGACAGGGGGACCCCAAGTGTCCAAGTACTGCTTGGACTTTTACAGAGTGTAAAAAGCTCTGGGAATAAGGGCTGGGTAAGACCGGTGCCAGCCTCCGCGGTAAAACCGGCAGCTCTAGTGGTAGTCGTTTTTATTGGGCTTAGAGCATCCGTAGCCGGCTTGGAAAGTCCTCTGTGAAATCCTACCGCTTAACGGTAAGACTTGCAAAGGATACTTCCAGGCTTGGGGCCGGAAGAAGCTAGGGGCATTTCTGGGGTAGCGGTGAAATGTTATAATCCTAGGAAGGTCACCAATGGCGAAGGCACCTAGCTAGGACGGGCCCGACGGTGAGGGATGTAAGCTAGGGGAG |
| p__Halobacteriota_g__Haladaptatus_ASV_2893 | 0 | 1 | 0 | 0 | 0 | 0 | 0 | 0 | 0 | 0 | 0 | ACTTTACACTGCACGACAGTGCGATAAGGGAACTCCAAGTGCGAGGGCATATAGTCCTCGCTTTTGTGTATCGTAAGGTGGTACACGAATAAGAGCTGGGCAAGACCGGTGCCAGCCGCCGCGGTAATACCGGCAGCTCAAGTGATGGCCGCTATTATTGGGCCTAAAGCGTCCGTAGCCTGCCAGACAGGTCCGTCGGGAAATCTGCTCGCTCAACGAGCAGGCGTCCGGCGGAAACCAGCTGGCTTGGGGCCGGAAGACCCAAGGGGTACGTCTGGGGTAGGAGTGAAATCCTGTAATCCTAGACGGACCACCGATCGCGAAAGCACCTTGGGAGGACGGACCCGACGGTGAGGGACGAAAGCTAGGGTCA |
| p__Halobacteriota_g__Haladaptatus_ASV_2907 | 0 | 1 | 0 | 0 | 0 | 0 | 0 | 0 | 1 | 0 | 0 | CCTTTACACTGCACGAAAGTGCGATAAGGGGACTCCAAGTGCGAGGGCATACAGTCCTCGCTTTTGTACACCGTAAGGAGGTGTACGAATAAGAGCTGGGCAAGACCGGTGCCAGCCGCCGCGGTAATACCGGCAGCTCAAGTGATGACCGCTATTATTGGGCCTAAAGCGTCCGTAGCCGGCCGAACAGGTCCGTTGGGAAATCTGCTCGCTTAACGAGCAGGCGTCCAGCGGAAACCAGTCGGCTTGGGACCGGAAGACCCGAGGGGTACGTCTGGGGTAGGAGTGAAATCCTGTAATCCTGGACGGACCGTCGGTGGCGAAAGCGCCTCGGGAGGACGGATCCGACGGTGAGGGACGAAAGCTAGGGTCA |
| p__Halobacteriota_g__Halomarina_3560_ASV_2854 | 0 | 0 | 0 | 0 | 0 | 0 | 0 | 0 | 0 | 1 | 1 | CCTTTACACTGCACGACAGTGCGATAGGGGGATTCCGAGTGCGAGGGCATACAGTCCTCGCTTTTGTGTACCTTAAGGTGGTACACGAATAAGGGCTGGGCAAGACCGGTGCCAGCCGCCGCGGTAATACCGGCAGCTCGAGTGATGACCGATATTATTGGGCCTAAAGCGTCCGTAGCCGGCCAGGTTGGTCCGTCGGGAAATCCACTCGCCTAACGAGTGGGCGTCCGGCGGATACCGCCTGGCTTGGGACCGGGAGACCAGAGGGGTACGTCTGGGGTAGGAGTGAAATCCTGTAATCCTAGACGGACCATCGGTGGCGAAAGCGCCTCTGGAGAACGGATCCGACGGTGAGGGACGAAAGCTAGGGTCT |
| p__Halobacteriota_g__Haloprofundus_ASV_2826 | 1 | 0 | 0 | 0 | 0 | 0 | 0 | 0 | 0 | 0 | 0 | CCTTTACACTGCACGACAGTGCGACAAGGGGACCCCGAGTGCGAGGGCATACAGTCCTCGCTTTTGTCGACCGTAAAGTGGTCGACGAATAAGAGCTGGGCAAGACCGGTGCCAGCCGCCGCGGTAATACCGGCAGCTCGAGTGATGACCGCTATTATTGGGCCTAAAGCGTCCGTAGCTGGCCGCGTAAGTCCATCGGGAAATCCATCCGCTCAACGGATGGGCGTCCGGTGGAAACTACACGGCTTGGAGCCGGAAGACCCAAGGGGTACGTCCGGGGTAGGAGTGAAATCCCGTAATCCTGGACGGACCGCCGATGGCGAAAGCACCTTGGGAAAACGGACTCGACAGTGAGGGACGAAAGCTAGGGTCT |
| p__Halobacteriota_g__Halorhabdus_ASV_2993 | 0 | 0 | 0 | 0 | 0 | 0 | 1 | 0 | 0 | 0 | 0 | CCTTTACACTGCACGACAGTGCGATAGGGGGACTCCGAGTGCGAGGGCATATAGTCCTCGCTTTTCTGTACCGTAAGGTGGTACAGGAATAAGGGCTGGGCAAGACCGGTGCCAGCCGCCGCGGTAATACCGGCAGCCCGAGTGATGGCCGCTATTATTGGGCCTAAAGCGTCCGTAGCCGGCCAGACAAGTCCGTTGGGAAATCCACGCGCTCAACGCGTGGACGTCCGGCGGAAACTGTCTGGCTTGGGGCCGGAAGATCTGAGGGGTACGTCCGGGGTAGGAGTGAAATCCCGTAATCCTGGACGGACCGCCGGTGGCGAAAGCGCCTCAGAAAGACGGACCCGACGGTGAGGGACGAAAGCTAGGGTCT |
| p__Halobacteriota_g__Halorientalis_ASV_2916 | 0 | 0 | 1 | 0 | 0 | 0 | 0 | 0 | 0 | 0 | 0 | CCTTTACACTGCACGACAGTGCGATAGGGGGACTCCGAGTGCGAGGGCATACAGTCCTCGCTTTTCTGTACCGTAAGGAGGTACAGGAACAAGTGCTGGGCAAGACCGGTGCCAGCCGCCGCGGTAATACCGGCAGCACAAGTGATGGCCGCTATTATTGGGCCTAAAGCGTCCGTAGCCGGCCACTCAAGTTCGTCGGGAAATCTGCCCGCTTAACGGCAGGCGTCCGGCGAAAACTGGGTGGCTTGGGGCCGGGAGATCCAAGGGGTACGTCCGGGGTAGGAGTGAAATCCCGTAATCCTGGACGGACCACCGGTGGCGAAAGCGCCTTGGAAAAACGGACCCGACGGTGAGGGACGAAAGCTAGGGTCT |
| p__Halobacteriota_g__Halorientalis_ASV_2990 | 0 | 0 | 0 | 0 | 0 | 1 | 0 | 0 | 0 | 0 | 0 | CCTTTACACTGCACGACAGTGCGATAAGGGGACTCCGAGTGCGAGGGCATACAGTCCTCGCTTTTCTGTACTGTAGGGAGGTACAGGAACAAGTGCTGGGCAAGACCGGTGCCAGCCGCCGCGGTAATACCGGCAGCACGAGTGATGGCCGATCTTATTGGGCCTAAAGCGTCCGTAGCCGGCCAATCAAGTCCGTCGGGAAATCTGCCCGCTTAACGGCAGGCGTCCGGCGGAAACTGATTGGCTTGGGGCCGGGAGATCCAAGGGGTACGTCCGGGGTAGGAGTGAAATCCCGTAATCCTGGACGGACCACCGATGGCGAAAGCACCTTGGAAAAACGGACCCGACGGTGAGGGACGAAAGCTAGGGTCT |
| p__Halobacteriota_g__Halorubrum_ASV_2912 | 0 | 0 | 1 | 0 | 0 | 0 | 0 | 0 | 0 | 0 | 0 | CCTTTACACTGCACGACAGTGCGATAAGGGGACTCCGAGTGCGAGGGCATATCGCCCTCGCTTTTGTACACCGTAGGGAGGTGTACGAATAAGCGCTGGGCAAGACCGGTGCCAGCCGCCGCGGTAATACCGGCAGCGCAAGTGATGGCCGATTTTATTGGGCCTAAAGCGTCCGTAGCTGGCCAGGCAAGTCCATCGGGAAATCCACCCGCTCAACGGGTGGGCGACCGGTGGAAACTGCCCGGCTCGGGACCGGAAGACCCAACGGGTACGTCCAGGGTAGGAGTGAAATCCCGTAATCCTGGACGGACCGCCGATGGCGAAAGCACGTTGGGAAGACGGATCCGACAGTGAGGGACGAAAGCTGGGGTCT |
| p__Halobacteriota_g__Halorubrum_ASV_2918 | 0 | 0 | 1 | 0 | 0 | 0 | 0 | 0 | 0 | 0 | 0 | CCTTTACACTGCACGCCAGTGCGATAAGGGGACTCCAAGTGCGAGGGCATATCGCCCTCGCTTTTCTACACCGTAGGGAGGTGTAGGAACAAGCGCTGGGCAAGACCGGTGCCAGCCGCCGCGGTAATACCGGCAGCGCAAGTGATGGCCGCTCTTATTGGGCCTAAAGCGTCCGTAGCTCGCCAGGCAAGTCCATCGGGAAATCCACCCGCTCAACGGGTGGGCGTCCGGTGGAAACTGTCTGGCTTGGGACCGAACGACCCAACGGGTACGTCCGGGGTAGGAGTGAAATCCCGTAATCCTGGACGGACCGCCGATGGCGAAAGCACGTTGGGAGGACGGATCCGACAGTGAGGGACGAAAGCTAGGGTCT |
| p__Halobacteriota_g__Halorussus_ASV_2887 | 0 | 1 | 0 | 0 | 0 | 0 | 0 | 0 | 0 | 0 | 0 | CCTTTACACTGCACGCTAGTGCGATAAGGGGACTCCGAGTGCGAGGGCATATAGTCCTCGCTTTTCTGTACCGTAAGGTGGTACAAGAATAAGGACTGGGCAAGACCGGTGCCAGCCGCCGCGGTAATACCGGCAGTCCGAGTGATGGCCGCTATTATTGGGCCTAAAGCGTCCGTAGCCGGCCAGACAGGTCCGTCGGGAAATCCACGCGCTCAACGCGTGGGCGTCCGGCGGAAACCAGCTGGCTTGGGGCCGGAAGACCTGAGGGGTACGTCCGGGGTAGGAGTGAAATCCCGTAATCCTGGACGGACCACCGGTGGCGAAAGCGCCTCAGGAAGACGGACCCGACGGTGAGGGACGAAAGCTAGGGTCA |
| p__Halobacteriota_g__Halorussus_ASV_2989 | 0 | 0 | 0 | 0 | 0 | 1 | 0 | 0 | 0 | 0 | 0 | CCTTTACACTGCACGCCAGTGCGATAAGGGGACCCCGAGTGCGAGGGCATACAGTCCTCGCTTTTGTACACCGTAAGGAGGTGTACGAATAAGGGCTGGGCAAGACCGGTGCCAGCCGCCGCGGTAATACCGGCAGCCCGAGTGATGGCCGCTGTTATTGGGCCTAAAGCGTCCGTAGCCGGCCGAACAAGTCTGTCGGGAAATCCATCTGCCTAACAGATGGGCGTCCGGCAGAAACTGTTCGGCTTGGGGCCGGAAGATCCGAGGGGTACGTCCGGGGTAGGAGTGAAATCCCGTAATCCTGGACGGACCGCCGGTGGCGAAAGCGCCTCGGAAGGACGGACCCGACGGTGAGGGACGAAAGCTGGGGTCT |
| p__Halobacteriota_g__Halosimplex_ASV_2930 | 0 | 0 | 1 | 0 | 0 | 0 | 0 | 0 | 0 | 0 | 0 | CCTTTACACTGCACGCCAGTGCGATAGGGGGACTCCGAGTGCGAGGGCATACAGTCCTCGCTTTTCTGTACCGTAAGGTGGTACAGGAACAAGTGCTGGGCAAGACCGGTGCCAGCCGCCGCGGTAATACCGGCAGCACGAGTGATGGCCGATCTTATTGGGCCTAAAGCGTCCGTAGCCGGCCGGACAAGTCCGTTGGGAAATCCACGCGCCCAACGCGTGGGCGTCCAGCGGAAACTGTCCGGCTTGGGGCCGGAAGACTCGAGGGGTACGTCCGGGGTAGGAGTGAAATCCCGTAATCCTGGACGGACCACCGGTGGCGAAAGCGCCTCGAGAAGACGGACCCGACGGTCAGGGACGAAAGCTAGGGTCT |
| p__Halobacteriota_g__Halovenus_ASV_2960 | 0 | 0 | 0 | 0 | 1 | 0 | 0 | 0 | 0 | 0 | 0 | CCTTTACACTGCACGACAGTGCGATAGGGGGACTCCGAGTGCGAGGGCATAGAGCCCTCGCTTTTGTACACCGTAAGGTGGTGTACGAATAAGAACTGGGCAAGACCGGTGCCAGCCGCCGCGGTAATACCGGCAGTTCGAGTGATGGCCGATATTATTGGGCCTAAAGCGTTCGTAGCTGGCCGATCAAGTCTGTCGGGAAATCTGCTCGCCTAACGAGCAGGCGTCCGGCGGAAACTGTTCGGCTTGGGACCGGAAGACTCGAGGAGTACGTCCGGGGTAGGAGTGAAATCCCGTAATCCTGGACGGACTACCGATGGCGAAAGCACCTCGAGAGCACGGATCCGACAGTGAGGAACGAAAGCTGGGGTCT |
| p__Halobacteriota_g__Halovivax_ASV_2894 | 0 | 1 | 0 | 0 | 0 | 0 | 0 | 0 | 0 | 0 | 0 | CCTTTACACTGCACGACAGTGCGATAAGGGGACTCCAAGTGCGAGGGCATATCGTCCTCGCTTTTTCCCACCCTAAGGAGGTGGGCGAATAAGTGCTGGGCAAGACCGGTGCCAGCCGCCGCGGTAATACCGGCAGCACGAGTGATGACCGCTATTATTGGGCCTAAAGCGTCCGTAGCCGGCCGACCAAGTCCATCGGGAAATCCGCACGCCTAACGTGCGGGCGTCCGGTGGAAACTGCTCGGCTTGGGACCGGAAGATCTGAGGGGTACGTCTGGGGTAGGAGTGAAATCCCGTAATCCTGGACGGACCGCCGGTGGCGAAAGCGCCTCAGAAGGACGGATCCGACGGTGAGGGACGAAAGCTAGGGTCA |
| p__Methanobacteriota_A_1229_g__Methanobacterium_D_1054_ASV_2855 | 0 | 1 | 0 | 1 | 0 | 0 | 0 | 0 | 0 | 1 | 0 | CCTCCGCAATGCACGAAAGTGCGACGGGGGGACCCCAAGTGCCACTCTTAACGGGGTGGCTTTTATGAAGTGTAAAAAGCTTCAAGAATAAGGGCTGGGCAAGACCGGTGCCAGCCGCCGCGGTAACACCGGCAGCTCAAGTGGTGGCCGTTTTTATTGGGCCTAAAGCGTTCGTAGCCGGCCTGATAAGTCTCTGGTGAAATCCCATAGCTTAACTGTGGGAATTGCTGGAGATACTATCAGGCTTGAGGTCGGGAGAGGTTAGGGGTACTCCCAGGGTAGGGGTGAAATCCTATAATCCTGGGAGGACCACCTGTGGCGAAGGCGCCTAACTGGAACGAACCTGACGGTGAGTAACGAAAGCCAGGGGCG |
| p__Halobacteriota_g__Natronomonas_ASV_2886 | 0 | 1 | 0 | 0 | 0 | 0 | 0 | 0 | 0 | 0 | 0 | CCTTTACACTGCACGCCAGTGCGATAAGGGGATACCGAGTGCGAGGGCATAGAGCCCTCGCTTTTCTGGACCGTAGGGAGGTCCACGAATAAGGGCTGGGCAAGACCGGTGCCAGCCGCCGCGGTAACACCGGCAGCCCAAGTGATGGCCGATATTATTGGGCCTAAAGCGTCCGTAGCTGGCCGAGCAAGTTCGTCGGGAAATCCACGCGCTTAACGCGTGGGCGTCCGGCGAAAACTGTTCGGCTTGGGACCGGAAGACCCGAGGGGTACGTCTGGGGTAGGAGTGAAATCCCGTAATCCTGGACGGACCACCGATGGCGAAAGCACCTCGGGAGGACGGATCCGACAGTGAGGGACGAAAGCTAGGGTCT |
| p__Thermoplasmatota_g__Natronoplasma_ASV_2832 | 1 | 0 | 0 | 0 | 0 | 0 | 0 | 0 | 0 | 0 | 1 | CCTTCACACTGTGGGAAACCACGACGAGGGGATTCCAAGTGCCAACACAATGTGTTGGCTTTTCATCTGTCTAAAACGCAGGTGGAATAAGAGCCGGGTAAGACTGGTGCCAGCCGCCGCGGTAACACCAGCGGCTCGAGTGGTGGCCTTTTTTATTGAGCTTAAAGCGTCCGTAGCCGGTCCAGTAAATCTCTGGGTAAATTGAACGGCTTAACCGTTCGAATTCCAGAGACACTGTTGGACTTGGGGCCGGGAGAGGCCAGAGGTACTCTCGGGGTAGAAGTAAAATTCTGTAATCCCGATAGGACCACCAGTGGCGAAGGCGTCTGGCTAGAACGGACTCGACGGTGAGGGACGAAGCCCTGGGGCG |
| p__Halobacteriota_g__Saliphagus_ASV_2891 | 0 | 1 | 0 | 0 | 0 | 0 | 0 | 0 | 0 | 0 | 0 | CCTTTACACTGCACGACAGTGCGATAAGGGGACTCCAAGTGCGAGGGCATACAGTCCTCGCTTTTCACTACCGTAAGGAGGTAGTGGAATAAGTGCTGGGCAAGACCGGTGCCAGCCGCCGCGGTAATACCGGCAGCACGAGTGATGACCGCTATTATTGGGCCTAAAGCGTCCGTAGCCGGCTTCGCAAGTCCGTCGGGAAATCCGCACGCTCAACGTGCGGGCGTCCGGCGGAAACTGCGTGGCTTGGGACCGGAAGACTCGAGGGGTACGTCCGGGGTAGGAGTGAAATCCCGTAATCCTGGACGGACCACCGGTGGCGAAAGCGCCTCGAGAGGACGGATCCGACGGTGAGGGACGAAAGCTAGGGTCA |
| p__Asgardarchaeota_g__SDNM01_ASV_2857 | 0 | 0 | 0 | 1 | 1 | 0 | 0 | 0 | 0 | 1 | 0 | ACTTCGCAATGCGCGCAAGCGTGACGAGGCTACCCGGAGTGTCTTCGCCTAAGGAAGACTGTGGCGTGTGCGTAAACTGCCGCGCTAGAAAGGAGAGGGCAAGGCTGGTGCCAGCCGCCGCGGTAAAACCAGCTCTTCAAGTCGTCGGGATAATTATTGGGCTTAAAGTGTTCGTAGCCGGTTAAGTAAGTTCTTGGTAAAACCTGGCAGCTCAACTGTCAGTATGCTAGGAATACTGCTTAACTAGAGGGCGGGAGAGGTCCGAGGTACTCCAGGGGTAGGGGTGAAATCCAATAATCCTTGGGGGACCACCAGTGGCGAAGGCGTCGGACTGGAACGCGCCTGACGGTGAGGGACGAAAGCCAGGGGAG |
| p__Thermoplasmatota_g__SM1-50_ASV_2871 | 0 | 0 | 0 | 0 | 0 | 0 | 0 | 0 | 0 | 0 | 1 | CCTTCACACTATGCGAAAGCGCGATGAGGGGATCCCAAGTGCTTGCTCTTAGGAGTAAGCTGTTTTCCTGTTTAAATCGCAGGAAGAGGAAGGGCTGGGTAAGACGGGTGCCAGCCGCCGCGGTAATACCTGCAGCCCGAGTGGTGACTGTTATTATTGGGCCTAAAACGTCCGTAGCCGGTTCTGTAAATGCCTGGGTAAATCGTACGGCTCAACCGTACGAATTCCGGGTAGACTGCAGGGCTAGAGATCGGGAGAGGCTAGAGGTACTCTTGGGGTAGAGGTGAAATTCTGTAATCCTAGGGGGACCACCAGTGGCGAAGGCGTCTAGCTAGAACGAGTCTGACGGTCAGGGACGAAGCCCTGGGGCG |
| p__Halobacteriota_f__Haloarculaceae_3992_ASV_2882 | 0 | 1 | 0 | 0 | 0 | 0 | 0 | 0 | 0 | 0 | 0 | CCTTTACACTGCGCGACAGCGCGATAGGGGGACTCCAAGTGCGAGGGCATATAGTCCTCGCTTTTGTACACCGTAAGGTGGTGTACGAATAAGAACTGGGCAAGACCGGTGCCAGCCGCCGCGGTAATACCGGCAGTTCGAGTGATGGCCGATTTTATTGGGCCTAAAGCGTTCGTAGCTGGCCAGTCAAGTCTGTCGGGAAACCTGCTCGCTCAACGAGCAGCCGTCCGGCGGAAACTGATTGGCTTGGGACCGGAAGACCTGAAGAGTACGTCCGGGGTAGGAGTGAAATCCTGTAATCCTGGACGGACTACCGATGGCGAAAGCACTTCAGGAGCACGGATCCGACAGTGAGGAACGAAAGCTGGGGTCT |

| | Presence (1) or Absence (0) in each layer | | | | | | | | | | |
|---|---|---|---|---|---|---|---|---|---|---|---|
| Phylum_Genus_ASV_number | L1 | L2 | L3 | L4 | L5 | L6 | L7 | L8 | L9 | L10 | L11 | Sequence |
| p__Halobacteriota_f__Haloarculaceae_3992_ASV_2984 | 0 | 0 | 0 | 0 | 0 | 1 | 0 | 0 | 0 | 0 | 0 | CCTTTACACTGCACGACAGTGCGATAAGGGGACTCCGAGTGCGAGGGCATACAGTCCTCGCTTTTCTGTACCGTAAGGTGGTACAGGAACAAGGGCTGGGCAAGACCGGTGCCAGCCGCCGCGGTAATACCGGCAGCCCGAGTGATGGCCGATATTATTGGGCCTAAAGCGTCCGTAGCCGGCCGGACAAGTCCGTTGGGAAATCCACGCGCCTAACGCGTGGGCGTCCAGCGGAAACTGTCCGGCTTGGGGCCGGAAGACTCGAGGGGTACGTCCGGGGTAGGAGTGAAATCCCGTAATCCTGGACGGACCGCCGGTGGCGAAAGCGCCTCGAGAGGACGGACCCGACGGTGAGGGACGAAAGCTAGGGTCT |
| p__Halobacteriota_f__Haloferacaceae_ASV_2906 | 0 | 1 | 0 | 0 | 0 | 0 | 0 | 0 | 0 | 0 | 0 | CCTTTACACTGCACGACAGTGCGATAAGGGGACTCCGAGTGCGAGGGCATATCGTCCTCGCTTTTCTCGACCGTAGGGAGGTCGAAGAATAAGAGCTGGGCAAGACCGGTGCCAGCCGCCGCGGTAATACCGGCAGCTCGAGTGATGGCCGATCTTATTGGGCCTAAAGCGTCCGTAGCCGGCCAGACAAGTCCGTCGGGAAATCCACCTACCCAACAGGTGGGCGTCCGGCGGAAACTCTCTGGCTTGGGACCAGACGACCCGAGAGGTACGTCCGGGGTAGGAGTGAAATCCCGTAATCCTGGACGGACCACCGATAGCGAAAGCATCTCGGGAGGATGGATCCGACGGTGAGGGACGAAAGCTGGGGTCT |
| p__Halobacteriota_f__Haloferacaceae_ASV_2910 | 0 | 0 | 1 | 0 | 0 | 0 | 0 | 0 | 0 | 0 | 0 | CCTTTACACTGCACGACAGTGCGATAAGGGGACTCCAAGTGCGAGGGCATACAGTCCTCGCTTTTCTGTACCGTAAGGAGGTACACGAATAAGGGCTGGGCAAGACCGGTGCCAGCCGCCGCGGTAATACCGGCAGCCCGAGTGATGACCGCTGTTATTGGGCCTAAAGCGTCCGTAGCCGGCCACGCAGGTCCATCGGGAAATCCATCCGCTCAACGGATGGGCGTCCGGTGGATACCACGTGGCTTGGGACCGAACGACTCGAGGGGTACGTCCGGGGTAGGAGTGAAATCCTGTAATCCTGGACGGACCGCCGATGGCGAAAGCACCTCGAGAGGACGGCTCCGACGGTGAGGGACGAAAGCTAGGGTCT |
| p__Halobacteriota_f__Haloferacaceae_ASV_2911 | 0 | 0 | 1 | 0 | 0 | 0 | 0 | 0 | 0 | 0 | 0 | CCTTTACACTGCACGCGAGTGCGATAAGGGGACTCCGAGTGCGAGGGCATATCGTCCTCGCTTTTCTGTACCGTAGGGCGGTACAGGAACAAGAGCTGGGCAAGACCGGTGCCAGCCGCCGCGGTAATACCGGCAGCTCGAGTGATGGCCGATCTTATTGGGCCTAAAGCGTCCGTAGCCGGCCCTGCAAGTCCGTCGGGAAATCCACGCGCTCAACGCGTGGGCGTCCGGCGGAAACTCCAGGGCTTGGGACCGGAAGGCTCGAAGGGTACGTCCGGGGTAGGAGTGAAATCCTGTAATCCCGGACGGACCACCGATGGCGAAAGCACTTCGAGAAGACGGATCCGACGGTGAGGGACGAAAGCCAGGGTCT |
| p__Halobacteriota_o__Halanaerobiales_ASV_2884 | 0 | 1 | 0 | 0 | 0 | 0 | 0 | 0 | 0 | 0 | 0 | CCTTTACACTGCACGCACGTGCGATAAGGGGACTCCAAGTGCGAGGGCATACAGTCCTCGCTTTTTGCGACCGTAAGGAGGTCGCAGAATAAGAGCTGGGCAAGACCGGTGCCAGCCGCCGCGGTAATACCGGCAGCTCGAGTGATGGCCGATCTTATTGGGCCTAAAGCGTCCGTAGCCGGCCACGAAGGTCCGTCGGGAAATCCATCCGCTCAACGGATGGGCGTCCGGCGGAAACCACGTGGCTTGGGGCCAGAAGGCTCAGGGGGTACGTCCGGGGTAGGAGTGAAATCCCGTAATCCTGGACGGACCACCGATGGCGAAAGCACCCTGAGAGAATGGACCCGACGGTGAGGGACGAAAGCTAGGGTCT |
| p__Halobacteriota_o__Halanaerobiales_ASV_2900 | 0 | 1 | 0 | 0 | 0 | 0 | 0 | 0 | 0 | 0 | 0 | CCTTTACACTGCACGACAGTGCGATAGGGGGACTCCGAGTGCGAGGGCATACAGTCCTCGCTTTTCTGTACCGTAAGGTGGTACAGGAACAAGTGCTGGGCAAGACCGGTGCCAGCCGCCGCGGTAATACCGGCAGCACGAGTGATGGCCGATATTATTGGGCCTAAAGCGTCCGTAGCTGGCCGCGCAAGTCCGTCGGGAAATCCAGGGGCTCAACCTCTGGGCGTCCGGCGGAAACTGTGCGGCTTGGGACCGGAAGACCCGAGGGGTACGTCCGGGGTAGGAGTGAAATCCTGTAATCCCGGACGGACCACCGATGGCGAAAGCACCTCGGGAGCACGGATCCGACAGTGAGGGACGAAAGCTGGGGTCT |
| p__Halobacteriota_o__Halanaerobiales_ASV_2919 | 0 | 0 | 1 | 0 | 0 | 0 | 0 | 0 | 0 | 0 | 0 | CCTTTACACTGCACGACAGTGCGATAGGGGGACTCCGAGTGCGAGGGCATACAGTCCTCGCTTTTCTGTACCGTAAGGAGGTACAGGAACAAGTGCTGGGCAAGACCGGTGCCAGCCGCCGCGGTAATACCGGCAGCACGAGTGATGGCCGCTATTATTGGGCCTAAAGCGTCCGTAGCCGGCCAGTCAAGTCCGTTGGGAAATCCACGCGCCTAACGCGTGGGCGTCCAGCGGAAACTGACTGGCTTGGGGCCGGAAGACCTGAGGGGTACGTCCGGGGTAGGAGTGAAATCCCGTAATCCTGGACGGACCACCGGTGGCGAAAGCGCCTCAGGAAGACGGACCCGACGGTGAGGGACGAAAGCTAGGGTCT |
| p__Halobacteriota_o__Halanaerobiales_ASV_2923 | 0 | 0 | 1 | 0 | 0 | 0 | 0 | 0 | 0 | 0 | 0 | CCTTTACACTGCACGCCAGTGCGATAAGGGGACTCCGAGTGCGAGGGCATATAGTCCTCGCTTTTCTGGACCGTAAGGTGGTCCAGGAACAAGTGCTGGGCAAGACCGGTGCCAGCCGCCGCGGTAATACCGGCAGCACGAGTGATGGTCGATATTATTGGGCCTAAAGCGTCCGTAGCTGGCCGAACAAGTCCGTCGGGAAATCCAGTGGCTCAACCACTGGGCGTCCGGTGGAAACTGTTCGGCTTGGGACCGGAAGACCCGAGGGGTACGTCTGGGGTAGGAGTGAAATCCCGTAATCCTGGACGGACCACCGATCGCGAAAGCACCTCGGGAAGACGGATCCGACAGTGAGGGACGAAAGCTGGGGTCT |
| p__Halobacteriota_o__Halanaerobiales_ASV_2924 | 0 | 0 | 1 | 0 | 0 | 0 | 0 | 0 | 0 | 0 | 0 | CCTTTACACTGCACGACAGTGCGATAGGGGGACCCCGAGTGCGAGGGCATACAGTCCTCGCTTTTTCACACCGTAAGGAGGTGTGCGAATAAGTGCTGGGCAAGACCGGTGCCAGCCGCCGCGGTAATACCGGCAGCACAAGTGATGGCCGCTGTTATTGGGCCTAAAGCGTCCGTAGCCGGCCGAACAAGTCCGTCGGGAAATCCATCCGCCTAACGGATGGGCGTCCGGCGGAAACTGTTCGGCTTGGGACAGGAAGACTCGAGGGGTACGTCCGGGGTAGGAGTGAAATCCCGTAATCCTGGACGGACCGCCGATGGCGAAAGCACCTCGAGAAGACTGATCCGACGGTGAGGGACGAAAGCTGGGGTCT |
| p__Halobacteriota_o__Halanaerobiales_ASV_2933 | 0 | 0 | 1 | 0 | 0 | 0 | 0 | 0 | 0 | 0 | 0 | CCTTTACACTGCACGACAGTGCGATAAGGGAACTCCGAGTGCGAGGGCATATAGTCCTCGCTTTTGTGTACCGTAAGGTGGTACACGAATAAGGGCTGGGCAAGACCGGTGCCAGCCGCCGCGGTAATACCGGCAGCCCGAGTGATGGCCGCTATTATTGGGCCTAAAGCGTCCGTAGCCGGCCGAACAGGTCCGTCGGGAAATCTGCTCGCTCAACGAGCAGGCGTCCGGCGGAAACCAGTCGGCTTGGGACCGGAAGACCTGAGGGGTACGTCCGGGGTAGGAGTGAAATCCCGTAATCCTGGACGGACCGCCGGTGGCGAAAGCGCCTCAGGAGGACGGATCCGACGGTGAGGGACGAAAGCTAGGGTCA |
| p__Nanoarchaeota_o__Woesearchaeales_581_ASV_2828 | 1 | 1 | 0 | 0 | 0 | 0 | 0 | 0 | 0 | 0 | 0 | CCTCTACAATGCACGCAAGTGTGATAGGGGGATCCCAAGTGCTCATGCTTTGCATGGGCTTTTGCCAAGAGTAAAAATCTTGGCGAATAAGTGGTGGGTAAGACTGGTGCCAGCCGCCGCGGTAACACCAGCGCCACGAGTGGCAATCGCATTTATTGGGCCTAAAGCGTCCGTAGCCGGTTCAGTAAATCTTTTGTGAAACTGTAAAGCTCAACTTTACAACATGCAGAAGACACTGCTGGACTTGAGACCGGAAGGGGTTGGAGGTATGTTAGGGGGAGCGGTAAAATGTTATAATCCCTGACAGACCACCTATGGCGAAGGCATCCAACCAGGACGGATCTGACGGTGAGGGACGAAAGCTAGGGGAG |
| p__Nanoarchaeota_o__Woesearchaeales_581_ASV_2833 | 1 | 1 | 0 | 0 | 0 | 0 | 0 | 0 | 0 | 0 | 0 | CCTTTACAATGCACGCAAGTGTGATAAGGGGATCCTAAGTGCTCGTGCACAGCACGGGCTTTTGCCAAGAGTAAATATCTTGGCGAATAAGTGGTGGGTAAGACTGGTGCCAGCCGCCGCGGTAACACCAGCGCCACGAGTGGCAACCACGTTTATTGGGCCTAAAGCGTCCGTAGCCGGTTTGGAAAATCTTTTGTGAAATCGACAGGCTTAACCTGTCGGCGTGCTTGAGACACTACCAAACTTGAGACCGGGAGGAGTCAGAGGTATTCCAGGGGGAGCGGTAAAATGTGATAATCCCTGGAGGACCATCTATGGCGAAGGCATCTGACCAGAACGGATCTGACGGTGAGGGACGAAGGCTAGGGGAG |
| p__Nanoarchaeota_g__BMS3ABIN17_ASV_3033 | 1 | 0 | 0 | 0 | 0 | 0 | 0 | 0 | 0 | 0 | 1 | ATTTTACAATGCTCGCAAGAGTGATAAAATAAACCAGAGTGTTCTTCTTAGAAGAACTTTTGTTTGTTGTAAAAAGACAGACGAATAAGGACTGGGTAAGATCGGTGCCAGCCGCCGCGGTAATACCGGCGGTCCGAGTCGCATCCACATTTATTGGGTCTAAAACATCCGTAGCTTGTTTTGTAAGTTTCTTGTTAAATCTGATATCTTAAGTATCAATCTGCAAGAAATACTGCATTGCTAGAGACCGGAAGATGCAGAGAGTACGATCAAGGTAGTGGTAAAATACGTTAAGCTTGATTGGACTAACAATGGCGAAGGCACTCTGCAAGTACGGATCTGACAGTGAGGGATGAAGGCTAGGGGCG |
| p__Nanoarchaeota_g__CG1-02-57-44_ASV_3041 | 0 | 0 | 0 | 0 | 0 | 0 | 0 | 0 | 0 | 1 | 0 | CCTCCACAATGCACGCAAGTGTGATGGGGGGATCCCACGTGCTCACGTTCTGCGTGGGCTTTTGCTGAATGTAAGGAGTTCAGCGAATAAGGGGTGGGTAAGACCGGTGCCAGCCGCCGCGGTAACACCGGCGCCCCGAGTGGCAGTCGCGATTATTGGGCCTAAAGCGTTCGTAGCCGGCCTCGTAAATCCCTTGTGAAATCGTCTCGCCTAACGTGACGGCGTGCAGGGGAGACTGCGGGGCTAGGGACCGGGAGGAGTCAGAGGTATGCCGTGGGGACCGGTAAAATGGGATAATCCGCGGTAGACCACCGATGGCGAAGGCATCTGACCAGAACGGATCCGACGGTGAGGGACGAAAGCCAGGGGAG |
| p__Halobacteriota_g__Haladaptatus_ASV_3047 | 0 | 1 | 0 | 0 | 0 | 0 | 0 | 0 | 1 | 1 | 1 | CCTTTACACTGCACGACAGTGCGATAAGGGGACTCCAAGTGCGAGGGCATACAGTCCTCGCTTTTCTGTACCGTAAGGTGGTACAGGAATAAGAGCTGGGCAAGACCGGTGCCAGCCGCCGCGGTAATACCGGCAGCTCGAGTGATGGCCAATATTATTGGGCCTAAAGCGTCCGTAGCCGGCCGAACAGGTCCGTCGGGAAATCTGCTCGCTCAACGAGCAGGCGTCCGACGGAAACCAGTCGGCTTGGGACCGGAAGACCCGAGGGGTACGTCTGGGGTAGGAGTGAAATCCTGTAATCCTAGACGGACCGCCGGTGGCGAAAGCGCCTCGGGAAGACGGATCCGACGGTGAGGGACGAAAGCTAGGGTCA |
| p__Halobacteriota_g__Halobacterium_ASV_3126 | 0 | 0 | 0 | 1 | 0 | 0 | 0 | 0 | 0 | 0 | 0 | CCTTTACACTGCACGAAAGTGCGATAGGGGGACCCCGAGTGCGAGGGCATACAGTCCTCGCTTTTGTACACCGTAAGGTGGTGTACGAATAAGGGCTGGGCAAGACCGGTGCCAGCCGCCGCGGTAATACCGGCAGCCCAAGTGATGGCCGTTATTATTGGGCCTAAAGCGTCCGTAGCCGGCCGGACAAGTCCGTTGGGAAATCCGCTCGCTCAACGAGCGGACGTCCAGCGGAAACTGTTCGGCTTGGGACCGGAAGACTCGAGGGGTACGTCCGGGGTAGGAGTGAAATCCTGTAATCCTGGACGGACCGCCGGTGGCGAAAGCGCCTCGAGAAGACGGATCCGACGGTGAGGGACGAAAGCTAGGGTCT |
| p__Halobacteriota_g__Halodesulfurarchaeum_ASV_3174 | 0 | 0 | 0 | 0 | 0 | 0 | 1 | 0 | 0 | 0 | 0 | CCTTTACACTGCACGACAGTGCGATAAGGGGATTCCGAGTGCGAGGGCATAGAGCCCTCGCTTTTCTGTACCGTAAGGAGGTAGAGGAAGAAGGGCTGGGCAAGACCGGTGCCAGCCGCCGCGGTAATACCGGCAGCCCAAGTGATGGCCGATATTATTGGGCCTAAAGCGTCCGTAGCCGGCCGGACAAGTCCGTTGGGAAATCCGCACGCTCAACGTGCGGGCGTCCAGCGGAAACTGTCCGGCTTGGGACCGGAGGACCCGAGGGGTACGTCCGGGGTAGGAGTGAAATCCTGTAATCCCGGACGGACCGCCTGTGGCGAAAGCGCCTCGGGAAGACGGATCCGACGGTGAGGGACGAAAGCCAGGGTCT |
| p__Halobacteriota_g__Halorussus_ASV_3085 | 0 | 1 | 0 | 0 | 0 | 0 | 0 | 0 | 0 | 0 | 0 | CCTTTACACTGCACGACAGTGCGATAAGGGGACTCCGAGTGCGAGGGCATATAGTCCTCGCTTTTCTGTACCGTAAGGTGGTACAGGAATAAGGACTGGGCAAGACCGGTGCCAGCCGCCGCGGTAATACCGGCAGTCCGAGTGATGGCCGCTATTATTGGGCCTAAAGCGTCCGTAGCCGGCCAGACAGGTCCGTCGGGAAATCTGCTCGCTCAACGAGCAGGCGTCCGGCGGAAACCAGCTGGCTTGGGGCCGGAAGACTCGAGGGGTACGTCCGGGGTAGGAGTGAAATCCTGTAATCCTGGACGGACCGCCGGTGGCGAAAGCGCCTCGAGAGGACGGACCCGACGGTGAGGGACGAAAGCTAGGGTCA |
| p__Halobacteriota_g__Halorussus_ASV_3090 | 0 | 1 | 0 | 0 | 0 | 0 | 0 | 0 | 0 | 0 | 0 | CCTTTACACTGCACGACAGTGCGATAGGGGGATCCCGAGTGCGAGGGCATACAGTCCTCGCTTTTCTGTACCGTAAGGTGGTACAGGAATAAGGGCTGGGCAAGACCGGTGCCAGCCGCCGCGGTAATACCGGCAGCCCGAGTGATGGCCGATGTTATTGGGCCTAAAGCGTCCGTAGCCGGCCGGACAAGTTCGTCGGGAAATCCGCACGCTTAACGTGCGGGCGACCGGCGAAAACTGTCCGGCTTGGGGCCGGAAGATCCGAGGGGTACGTCCGGGGTAGGAGTGAAATCCCGTAATCCTGGACGGACCGCCGGTGGCGAAAGCGCCTCGGAAAGACGGACCCGACGGTGAGGGACGAAAGCTGGGATCT |
| p__Thermoproteota_g__JACAEJ01_ASV_3048 | 0 | 0 | 0 | 0 | 0 | 0 | 0 | 0 | 1 | 1 | 0 | ACTTCGCAATGCGCGAAAGCGTGACGAGGCTATTCCGAGTGCCACCCGCTGAGGGAGGCTTTTACTAGATTTAAGAAATCTAGAGAATAAGGGGAGGGCAAGCCTGGTGTCAGCCGCCGCGGTAATACCAGCTCCCCGAGTGGTTGGGATGATTATTGGGCCTAAAGCATCCGTAGCCTGTTCTGTAGGTCCTCTGTTAAAACTAATGGCTTAACCGTTAGACTGCAGAGGATACCATAGGACTAGGAGGCGGGAGAGGCGGACGGTACTCCGAGTGTAGGGGTAAAATCCTTTGATCCTCGGAAGACCACCAGTGGCGAAGGCGGTTCGCCAGAACGCGCTCGACGGTGAGGGATGAAAGCTGGGGGAG |
| p__Halobacteriota_g__Natrinema_ASV_3031 | 1 | 0 | 0 | 0 | 0 | 0 | 0 | 0 | 0 | 0 | 0 | CCTTTACACTGCACGCGAGTGCGATAAGGGGACTCCGAGTGCGAGGGCATATAGTCCTCGCTTTTCACCACCGTAGGAAGGTGGTAGAATAAGTGCTGGGCAAGACCGGTGCCAGCCGCCGCGGTAATACCGGCAGCACGAGTGATGACCGCTTTTATTGGGCCTAAAGCGTCCGTAGCTGGCCACGCAAGTCTATCGGGAAATCTGCGCGCTCAACGCGCAGGCGTCCGGTGGAAACTGCGTGGCTTGGGACCGGAAGATCCAGAGGGTACGTCCGGGGTAGGAGTGAAATCCCGTAATCCTGGACGGACCACCGGTGGCGAAAGCGCTCTGGAAAGACGGATCCGACGGTGAGGGACGAAAGCTCGGGTCA |
| p__Thermoproteota_g__Nitrosocosmicus_ASV_3093 | 0 | 1 | 0 | 0 | 0 | 0 | 0 | 0 | 0 | 0 | 0 | CCTCTGCAATAGGCGAAAGCTTGACAGGGTTACTCTGAGTGATTTCCGTTAAGGAGATCTTTTGGCACCTCTAAAAATGGTGCAGAATAAGGGGTGGGCAAGTCTGGTGTCAGCCGCCGCGGTAATACCAGCACCCCGAGTGGTCGGGACGTTTATTGGGCCTAAAGCATCCGTAGCCGGTTCTACAAGTCTTCCGTTAAATCCACCTGCTCAACAGATGGGCTGCGGAAGATACTATGGATCTAGGAGGCGGGAGAGGCAAGCGGTACTCGATGGGTAGGGGTAAAATCCGTTGATCCATTGAAGACCACCAGTGGCGAAGGCGGCTTGCCAGAACGCGCTCGACGGTGAGGGATGAAAGCTGGGGGAG |
| p__Thermoplasmatota_g__SG8-5_ASV_3195 | 0 | 0 | 0 | 0 | 0 | 0 | 0 | 0 | 1 | 0 | 0 | ACTTGTCAATGGGGGAAACCCCGAACAGGGGAGTCCAAGTGCTTGCACTTTGTGTAAGCTGTTCGGTTGTCTAAAAAACAGCTGAAGGAAGGGCTGGGTAAGACAGGTGCCAGCCGCCGCGGTAATACCTGCAGCCCGAGTGGTGATCGATTTTATTGAGCCTAAAGCGTCCGTAGCCGGTCCAGTAAACCCGCGGGTAAATTGGGCAGCTCAACTGTCCAAATTCCGCGGGGACTGCTGGACTTGGGACCGGGAGAGGTCGGAGGTACTCCAGGGGTAGGGGCTAAATCCTGTAATCCTTGGGGGACCACCAGTGGCGAAGGCGTCCGACTAGAACGGATCCGACGGTAAGGGACGAAACCCTGGGTCG |

| Phylum_Genus_ASV_ number | Presence (1) or Absence (0) in each layer | | | | | | | | | | | |
|---|---|---|---|---|---|---|---|---|---|---|---|---|
| | L1 | L2 | L3 | L4 | L5 | L6 | L7 | L8 | L9 | L10 | L11 | Sequence |
| p__Halobacteriota_f__Haladaptataceae_ASV_3128 | 0 | 0 | 0 | 1 | 0 | 0 | 0 | 0 | 0 | 0 | 0 | CCTTTACACTGCACGACAGTGCGATAGGGGGACTCCGAGTGCGAGGGCATACAGTCCTCGCTTTTCTGTACCGTAAGGAGGTACAAGAATAAGGGCTGGGCAAGACCGGTGCCAGCCGCCGCGGTAATACCGGCAGCCCAAGTGATGGCCGCTATTATTGGGCCTAAAGCGTCCGTAGCCGGCCAGTCAAGTCTGTTGGGAAATCCACGCGCCTAACGCGTGGGCGTCCAGCGGAAACTGACTGGCTTGGGGCCGGAAGACCTGAGGGGTACGTCCGGGGTAGGAGTGAAATCCCGTAATCCTGGACGGACCACCGGTGGCGAAAGCGCCTCAGGAAGACGGACCCGACGGTGAGGGACGAAAGCTAGGGTCT |
| p__Halobacteriota_f__Haloarculaceae_3992_ASV_3098 | 0 | 0 | 1 | 0 | 0 | 0 | 0 | 0 | 0 | 0 | 0 | CCTTTACACTGCACGCCAGTGCGATAGGGGGACTCCGAGTGCGAGGGCATACAGTCCTCGCTTTTCTGTACCGTAAGGTGGTACAGGAACAAGTGCTGGGCAAGACCGGTGCCAGCCGCCGCGGTAATACCGGCAGCACGAGTGATGGCCGATCTTATTGGGCCTAAAGCGTCCGTAGCCGGCCGGACAAGTCCGTTGGGAAATCCACGCGCTCAACGCGTGGGCGTCCAGCGGAAACTGTCTGGCTTGGGGCCGGAAGACCCGAGGGGTACGTCCGGGGTAGGAGTGAAATCCTGTAATCCCGGACGGACCACCGGTGGCGAAAGCGCCTCGGGAAGACGGACCCGACGGTGAGGGACGAAAGCTAGGGTCT |
| p__Halobacteriota_f__Haloferacaceae_ASV_3161 | 0 | 0 | 0 | 0 | 0 | 1 | 0 | 0 | 0 | 0 | 0 | CCTTTACACTGCACGCCAGTGCGATAGGGGGACCCCGAGTGCGAGGGCATACAGTCCTCGCTTTTATGCACCGTAAGGAGGTGCATGAATAAGTGCTGGGCAAGACCGGTGCCAGCCGCCGCGGTAATACCGGCAGCACAAGTGATGGCCGCTATTATTGGGCCTAAAGCGTCCGTAGCCGGCCGAACAAGTCCGTCGGGAAATCCATCCGCTCAACGGATGGGCGTCCGGCGGAAACTGTTCGGCTTGGGACAGGAAGACTCGAGGGGTACGTCCGGGGTAGGAGTGAAATCCTGTAATCCTGGACGGACCGCCGATGGCGAAAGCACCTCGAGAAGACTGATCCGACGGTGAGGGACGAAAGCTGGGGTCT |
| p__Halobacteriota_f__Natrialbaceae_ASV_3091 | 0 | 1 | 0 | 0 | 0 | 0 | 0 | 0 | 0 | 0 | 0 | CCTTTACACTGCACGACAGTGCGATAGGGGGACTCCACGTGCGAGGGCATACAGTCCTCGCTTTTTCCTACCGTAAGGTGGTAGGCGAATAAGGGCTGGGCAAGACCGGTGCCAGCCGCCGCGGTAATACCGGCAGCCCAAGTGATGACCGCTATTATTGGGCCTAAAGCGTCCGTAGCCTGCCGAGAAAGTCCGTCGGGAAATCCGCACGCCCAACGTGCGGGCGTCCGGCGGAAACTACTCGGCTTGGGACCGGAGGACCCGAGGGGTACGTCTGGGGTAGGAGTGAAATCCCGTAATCCTGGACGGACCACCGGTGGCGAAAGCGCCTCGGGAAGACGGATCCGACGGTGAGGGACGAAAGCTCGGGTCA |
| p__Halobacteriota_f__Natrialbaceae_ASV_3099 | 0 | 0 | 1 | 0 | 0 | 0 | 0 | 0 | 0 | 0 | 0 | CCTTTACACTGCACGACAGTGCGATAGGGGGACTCCAAGTGCGAGGGCATATCGTCCTCGCTTTTCACTACCGTAAGGTGGTAGTAGAATAAGGGCTGGGCAAGACCGGTGCCAGCCGCCGCGGTAATACCGGCAGCCCGAGTGATGACCGCTATTATTGGGCCTAAAGCGTCCGTAGCTGGCCGATCAAGTCTATCGGGAAATCTGCGCGCTCAACGCGCAGGCGTCCGGTGGAAACTGTTCGGCTTGGGACCGGAAGACTCGAGGGGTACGTCCGGGGTAGGAGTGAAATCCCGTAATCCTGGACGGACCACCGGTGGCGAAAGCGCCTCGAGAGGACGGATCCGACGGTGAGGGACGAAAGCTTGGGTCA |
| p__Halobacteriota_f__Natrialbaceae_ASV_3204 | 0 | 0 | 0 | 0 | 0 | 0 | 0 | 0 | 1 | 0 | 0 | CCTTTACACTGCGCGACAGCGCGATAAGGGGACTCCAAGTGCGAGGGCATATCGTCCTCGCTTTTCACCATCGTAAGGAGATGGTGGAATAAGGGCTGGGCAAGACCGGTGCCAGCCGCCGCGGTAATACCGGCAGCCCAAGTGATGACCGCTATTATTGGGCCTAAAGCGTCCGTAGCCGGCCGATCAAGTCCGTCGGGAAATCCGCACGCCTAACGTGCGGGCGTCCGGCGGAAACTGCTCGGCTTGGGACCGGAAGATCCAGAGGGTACGTCCGGGGTAGGAGTGAAATCCCGTAATCCTGGACGGACCACCGGTGGCGAAAGCGCTCTGGAAGGACGGATCCGACGGTGAGGGACGAAAGCTCGGGTCA |
| p__Halobacteriota_o__Halanaerobiales_ASV_3064 | 0 | 0 | 0 | 0 | 0 | 0 | 1 | 0 | 0 | 0 | 1 | CCTTTACACTGCACGACAGTGCGATAGGGGGACTCCGAGTGCGAGGGCATACAGTCCTCGCTTTTGTCGACCGTAAGGAGGTCGACGAATAAGGGCTGGGCAAGACCGGTGCCAGCCGCCGCGGTAATACCGGCAGCCCGAGTAATGGCCGATCTTATTGGGCCTAAAGCGTCCGTAGCCGGCCCGACAAGTCCGTTGGGAAATCCACGCGCTCAACGCGTGGGCGTCCAGCGGAAACTGTCGGGCTTGGGACCGGAAGACCCAGAGGGTACGTCCGGGGTAGGAGTGAAATCCCGTAATCCTGGACGGACCACCGGTGGCGAAAGCGCTCTGGGAAGACGGATCCGACGGTGAGGGACGAAAGCTAGGGTCT |
| p__Halobacteriota_o__Halanaerobiales_ASV_3068 | 0 | 1 | 0 | 0 | 0 | 0 | 0 | 0 | 0 | 0 | 0 | CCTTTACACTGCACGCCAGTGCGATAAGGGGACCCCCAGTGCGAGGGCATACCGTCCTCGCTTTTCACCACCGTAAGGAGGTGGTGGAACAAGAGCTGGGCAAGACCGGTGCCAGCCGCCGCGGTAATACCGGCAGCTCGAGTGATGGCCGTTTTTATTGGGCCTAAAGCGTCCGTAGCCGGCCAGACAAGTTCGTCGGGAAATCCACGTGCCCAACGCGTGGGCGTCCGGCGAAACCTCTCTGGCTTGGGACCGGACGGCCCGAGGGGTACGGTCGGGGTAGGAGTGAAATCCCGTAATCCTGGACGGACCACCGGTGGCGAAAGCGCCTCGAGAAGACGGACCCGACGGTGAGGGACGAAAGCTAGGGTCT |
| p__Halobacteriota_o__Halanaerobiales_ASV_3083 | 0 | 1 | 0 | 0 | 0 | 0 | 0 | 0 | 0 | 0 | 0 | CCTTTACACTGCACGACAGTGCGATAAGGGGACTCCCAGTGCGAGGGCATATCGCCCTCGCTTTTGTACACCCTAAGGAGGTGCACGAATAAGGGCTGGGCAAGACCGGTGCCAGCCGCCGCGGTAATACCGGCAGCCCGAGTGATGGCCGCTATTATTGGGCCTAAAGCGTCCGTAGCCGGCCGGTCAAGTCCGTTGGGAAATCCACCCGCTTAACGGGTGGGCGTCCAGCGGAAACTGATCGGCTTGGGACCGAGTGACCCGAGGGGTACGTCCGGGGTAGGAGTGAAATCCCGTAATCCTGGACGGACCGCCGGTGGCGAAAGCGCCTCGGGAGAGCGGATCCGACGGTGAGGGACGAAAGCTTGGATCT |
| p__Halobacteriota_o__Halanaerobiales_ASV_3107 | 0 | 0 | 1 | 0 | 0 | 0 | 0 | 0 | 0 | 0 | 0 | CCTTTACACTGCACGAAAGTGCGATAAGGGGATCCCAAGTGCGAGGGCATACAGTCCTCGCTTTTCTGTACCGTAGGGAGGTACAAGAATAAGGGCTGGGCAAGACCGGTGCCAGCCGCCGCGGTAATACCGGCAGCCCAAGTGATGGCCGCTATTATTGGGCCTAAAGCGTCCGTAGCCGGCCGGATAAGTCCGTCGGGAAATCCACCCGCTCAACGGGTGGGCGTCCGGCGGAAACTGTCCGGCTTGGGGCCGGAAGGCCAGAGGGGTACGTCCGGGGTAGGAGTGAAATCCTGTAATCCTGGTCGGACCGCCGGTGGCGAAAGCGCCTCTGGAGGACGGACCCGACGGTGAGGGACGAAAGCTTGGGTCA |
| p__Halobacteriota_o__Halanaerobiales_ASV_3198 | 0 | 0 | 0 | 0 | 0 | 0 | 0 | 0 | 1 | 0 | 0 | CCTTTACACTGTACGAAAGTACGATAGGGGGACCCCGAGTGCGAGGGCATACAGTCCTCGCTTTTGTGTACCGTAAGGTGGTACACGAATAAGAGCTGGGCAAGACCGGTGCCAGCCGCCGCGGTAATACCGGCAGCTCAAGTGATGGCCGATATTATTGGGCCTAAAGCGTCCGTAGCCGGCCGAACAGGTCCGTTGGGAAATCCATCTGCCTAACAGATGGGCGTCCAGCGGAAACCAGTCGGCTTGGGGCCGGAAGACCTGAGGGGTACGTCCGGGGTAGGAGTGAAATCCTGTAATCCTGGACGGACCGCCGGTGGCGAAAGCGCCTCAGGAGGACGGACCCGACGGTGAGGGACGAAAGCTAGGGTCA |
| p__Nanoarchaeota_o__Woesearchaeales_581_ASV_3024 | 1 | 0 | 0 | 0 | 0 | 0 | 0 | 0 | 0 | 0 | 0 | CCTTTACACTGCACGAAAGTGTGATAAGGGGATCCTAAGTGCTCATGCTTTGCATGGGCTTTTGCCAAGAGTAAATATCTTGGCGAATAAGTGGTGGGTAAGACTGGTGCCAGCCGCCGCGGTAACACCAGCGCCACGAGTGGGGACCAGCATTATTGGGCCTAAAGCGTCCGTAGCAGGAAAGATAAATCTTCTGTGAAACTGTTAAGCTTAACTTAACAACGTGCAGAAGACACTATTTTTCTAGAGACCGGGAGGAGTCAGAGGTATTTCTGGGGGAGCGGTAAAATGCTATAATCCCAGGAGGACCACCGATGGCGAAGGCATCTGACTAGAACGGATCTGACTGTGAGGGACGAAGGCCAGGGGAG |
| d__Archaea_d__Archaea_ASV_3191 | 0 | 0 | 0 | 0 | 0 | 0 | 0 | 0 | 1 | 0 | 0 | CTTTCCCAATGCACGCAAGTGTGAGGAAGAGATCCGGAGTGCTTTCTCTTAGGAGGGAGCTTTTCTGAAATCTAAAGAGCTTCAGGAATAAGCAGTGGGTAAGACAGGTGCCAGCCGCCGCGGTAATACCTGCGCTGCAAGTGGTGACCACGTTTATTGGGCTTAAAGCGTTCGTAGCCGGTTTTGTAAGTCCCTGGTGAAATCGGTTGGTAAACTAACCGGCTTGCTGGGGATACTGCGAGACTTGGGACCGGGAGAGGTCAAGGGTATTCCTGGGGTAGGGGTAAAATCCTGTAATCCTTGGAGGACCACCTGTGGCGAAGGCGCTTGACTAGAACGGATCCGACGGTGAGGGACGAAAGCTAGGGGAG |
| p__Nanoarchaeota_g__BMS3ABIN17_ASV_3213 | 1 | 0 | 0 | 0 | 0 | 0 | 0 | 0 | 0 | 0 | 0 | AATCTACAATGCGCGAAAGCGTGATAGTTTGAGCCAGAGTGCTTTCCTTTTGGAAAGCTTTTGCCAAATGTAAAAAGTTTGGCGAATAAGGACTGGGTAAGACTAGTGCCAGCCGCCGCGGTAATACTAGCGGTCCAAGTCGCAGCCATTTTTATTGGGTCTAAAACATCCGTAGCTTGCTTGATAAGTCTCTTGTGAAATCCTGTATCTCAAGTATAGGACTTGCAAGAGATACTGTCAGGCTAGAGACCGGGAGACGTAAGGAGTACGTTCGAAGTAGCGGTAAAATGTGTTGATCTCGAACGGACTAACAAGAGCGAAGGCACCTTACGAGAACGGATCTGACAGTAAGGGATGAAGGCCAGGGTCG |
| p__Nanoarchaeota_g__BMS3ABIN17_ASV_3217 | 1 | 0 | 0 | 0 | 0 | 0 | 0 | 0 | 0 | 0 | 0 | AATCTACAATGCGCGCAAGCGTGATAGTTTGAGCCAGAGTGCTTCTCTATTGAGAAGCTTTTGTGGAATGTAAAAAGTTCTACGAATAAGGACTGGGTAAGACTAGTGCCAGCCGCCGCGGTAATACTAGCGGTCCAAGTCGCAGCCATCTTTATTGGGTCTAAAACATCCGTAGCTTGCTTGGCAAGTTCCTTGTGAAATCTCATGTCTCAAGCATGGGGCGGGCAAGGAGTACTACCAAGCTAGAGACTGGGAGACGTAAGAAGTACGTTCAGAGTAGCGGTAAAATGTTTTAATCCTGAGCGGACTACCAATAGCGAAGGCATCTTACGAGAACAGTTCTGACAGTCAGGGATGAAGGCTAGGGGCG |
| p__Nanoarchaeota_g__BMS3ABIN17_ASV_3225 | 1 | 0 | 0 | 1 | 0 | 0 | 0 | 0 | 0 | 0 | 0 | AATCTGCAATGCGCGAAAGCGTGACAGTTTGAGCCAGAGTGTTTGCCTATTGGTAAGCTTTTATGAAATGTAAAAAGTTTCATGAATAAGGACTGGGTAAGACTAGTGCCAGCCGCCGCGGTAATACTAGCGGTCCAAGTCGCAATCATTTTTATTTGGTCTAAAACATCCGTAGCTTGTTTAGTAAGTTTCTTGTGAAATCTTGTATCTCAAATACAAGGCGAGCAAGAAGTACTACTAAACTAGAGACTGGGAGACGTAAGAAGTATGTCTGAAGTAGCGGTAAAATGTGTTAATCTCAGACAGACTCACAATCAGCGAAGGCATCTTACGAGAACAGTTCTGACAGTAAGGGATGAAGGCTAGGGGCG |
| p__Nanoarchaeota_g__BMS3ABIN17_ASV_3228 | 0 | 0 | 0 | 0 | 0 | 0 | 0 | 0 | 0 | 1 | 0 | CTTCTGCAATGCGCGAAAGCGTGACAGAGTGAGTCAGAGTGTCTGCCATTCGGTAGACTTTTGTGGAATGTAAAAAGTTTCATGAATAAGGACTGGGTAAGACTAGTGCCAGCCGCCGCGGTAATACTAGCGGTCCAAGTCGCAGCCACTATTATTGGGTCTAGAACATCCGTAGCTTGCTTAATAAGTCTCTTGTGAAATCCTGTGTCTCAAGTATAGGACTTGCAAGAGAAACTGTTAAGCTAGAGACCGGGAGACGTAAGGAGTACGTTTGAAGTAGCGGTAAAATGTGTTGATCTCAGACGGACTAACAAGAGCGAAGGCACCTTACGAGAACGGTTCTGACAGTAAGGGATGAAGGCCAGGGTCG |
| p__Nanoarchaeota_g__BMS3ABIN17_ASV_3261 | 0 | 0 | 0 | 0 | 0 | 0 | 0 | 0 | 0 | 1 | 1 | TTTTTGCAATGTACGAAAGTATGACAAAGCGAGCCAGAGTGTTTTTCTATATGAAAAACTTTTGCTTGCTGTAAAAAGGCAGGCGAATAAGGACTGGGCAAGACCGGTGCCAGCCGCCGCGGTAATCCCGGCAGTCCAAGTCGCAGCCATCTTTATTGGGTCTAAAACATCCGTAGCTAGTAATAAAAGTCTCTTGTGAAATCTTAACTCTCAAGGTTAAGGCGTGCAAGAGATACTATACTACTCGAGACCGGAAGGCGCAAGAAGTACTATTAGGGTAGTGGTAAAATACGTTAATCCTAATTGGACTAACAATCGCGAAGGCATCTTGCGAGGACGGTTCTGACAGTGAGGGATGAAGGCTAGGGGCG |
| p__Nanoarchaeota_g__BMS3ABIN17_ASV_3270 | 0 | 0 | 0 | 0 | 0 | 0 | 0 | 0 | 0 | 0 | 1 | ATTCTGCAATGCGCGCAAGCGTGACAGATTGAGTCAGAGTGCGCTTCAATTTGGAGTGCTTTTGCCAAATGTAAAAAGTTTGGCGAATAAGGACTGGGTAAGACGAGTGCCAGCCGCCGCGGTAATACTCGCGGTCCAAGTCGCAGCCATTTTTATTGGGTCTAAAACATCCGTAGCTTGCTTGGTGAGTTCCTTGTGAAATTCTATATCTCAAATATGGAGCGTGCAGGGAGTACTGCCAAGCTAGAGACTGGAAGACGTAACGAGTACGTTCGGAGTAGCGGTAAAATGTGTTAATCCCGGACGGACTAACAATAGCGAAGGCACGTTACGAGGACAGTTCTGACAGTAAGGGATGAAGGCTAGGGGCG |
| p__Nanoarchaeota_g__BMS3ABIN17_ASV_3440 | 0 | 0 | 0 | 0 | 0 | 0 | 0 | 1 | 0 | 0 | 0 | CTTCTACAATGCGCGCAAGCGTGATAGAGTGAGCCAGAGTGCTTTCCAATTGGAAAGCTTTTGCCAAATGTAAATAGTTTGGCGAATAAGGACTGGGTAAGACTAGTGCCAGCCGCCGCGGTAATACTAGCGGTCCAAGTCGCAGCCATTTTTATTGGGTCTAAAACATCCGTAGCTTGCTTAGTAAGTCCCTTGTGAAATTCTATGTCTCAAACATAGAGCGCGCGGGGGGTACTGCTGAGCTAGAGACCGGGAGGCGTAGGAAGTACAATCAAAGTAACGGTTAAATGTGTTAATCTTGGTTGGACTCACAATCGCGAAGGCATCCTACGAGAACGGTTCTGATAGTCAGGGATGAAGGCTAGGGGCG |
| p__Nanoarchaeota_g__BMS3ABIN17_ASV_3460 | 0 | 0 | 0 | 0 | 0 | 0 | 0 | 0 | 1 | 0 | 0 | AATCTACAATGCGCGAAAGCGTGATAGTTTGAGCCAGAGTGCTTCTCATTGAGAAGCTTTTGTGGAATCTAAAAAGTTCTAAGAATAAGGACTGGGTAAGACTAGTGCCAGCCGCCGCGGTAATACTAGCGGTCCAAGTCGCAGCCATTTTTATTGGGTCTAAAACATCCGTAGCTTGCTTAATAAGTTTCTTGTGAAATCCTATATCTTAAATATAGGGCGGGCAAGAAGTACTGTTAAGCTAGAGACCGGAAGACGTGAGAAGTACGTTCAAAGTAGTGGTAAAATACGTTAATCTTGAACGGACTCACAATAGCGAAGGCATCTCACGAGGACGGTTCTGACAGTAAGGGATGAAAGCTAGGGGCG |
| p__Nanoarchaeota_g__BMS3ABIN17_ASV_3464 | 0 | 0 | 0 | 0 | 0 | 0 | 0 | 0 | 1 | 0 | 0 | ATTTTACAATGCTCGCAAGAGTGATAGAATAAACCAGAGTGTTTCTCCTAGAGAAACTTTTATCTGTTGTAAAAAGACAGATGAATAAGGACTGGGCAAGACCGGTGCCAGCCGCCGCGGTAATACCGGCGGTCCAAGTCGTATCCACATTTATTGGGTCTAAAACATTCGTAGCTTGTTTTGTAAGTTTCTTGTGAAATCTGATATCTCAAGTATCAGGCGTGCAAGAAATACTACATCACTAGAGACCGGAAGATGCAAAGAGTATGATTAAGGTAGCGGTAAAATGTGTTAATCTTAATCAGACTAACAATGGCGAAGGCACTTTGCAAGTACGGATCTGACAGTGAGGGATGAAGGCTAGGGGCG |

| Phylum_Genus_ASV_number | Presence (1) or Absence (0) in each layer L1 | L2 | L3 | L4 | L5 | L6 | L7 | L8 | L9 | L10 | L11 | Sequence |
|---|---|---|---|---|---|---|---|---|---|---|---|---|
| p__Nanoarchaeota_g__BMS3ABIN17_ASV_3471 | 0 | 0 | 0 | 0 | 0 | 0 | 0 | 0 | 1 | 0 | 0 | AATCTGCAATGCGCGAAAGCGTGACAGTTTGAGTCAGAGTGTTTCTCATTGAGAAACTTTTGCCAAGTGTAAAAAGTTTGGCGAATAAGGACTGGGTAAGACTAGTGCCAGCCGCCGCGGTAATACTAGCGGTCCAAGTCGCAGCCATATTTATTGGGTCTAAAACATCCGTAGCTTGTTTTGTAAGTTTTTTGTGAAATCCTGTTTCTTAAAGATAGGGCGCGCAGGAAGTACTGCAAAACTAGAGACTGGGAGACGTAAGAAGTACGTTCGAAGTAGTGGTAAAATGCTTTAATCTCGGACGGACTAACAATAGCGAAGGCATCTTACGAGAACAGTTCTGACAGTAAGGGATGAAGGCCAGGGGCG |
| p__Altiarchaeota_g__CABMCB01_ASV_3362 | 0 | 0 | 0 | 1 | 0 | 0 | 0 | 0 | 0 | 0 | 0 | TTTGCACAATGGACGAAAGTCTGATGCAGCGACGCCGCGTGAAGGATGACGGCCTTAGGGTTGTAAACTTCGATAAAGCACCGGCAAACTACGTGCCAGCAGCAGCGGTCATACGTAGGGTGCAAGCGTTGTCCGGATTTACTGGGCGTAAAGAGTTCTGTAGGCGGTTGCGTAAGTTGAAAGTTAAATCTCTCCGCTCAACGGGGAAACTGCTTTTAATACTGCGTGACTAGAGACATCTAGGGGACAGCGGAACTTATGGTGGAGCAGTGAAATGCGTTGATATCATAGGGAACACCAAGGGGGTAGCCAGCTGTCTGGGGATGTACTGACGCTGAGAGACGAAAGCGTGGGTAG |
| p__Nanoarchaeota_g__CG1-02-57-44_ASV_3224 | 1 | 0 | 0 | 0 | 0 | 0 | 0 | 0 | 1 | 0 | 0 | CCTTTACAATACACGCAAGTGTGATAAGGGGACCCAAAGTGCTTGCTTTCGGGCGAGCTTTTGTTAAGTGTAAAAATCTTGACGAATAAGTGATGGGAAAGACCGGTGCCAGCCGCCGCGGTAACACCGGCGTCACGAGTGGCGACCACGATTATTGGGCCTAAAGCGTCCGTAGCCGATTTGGTAAATTTCTTGTGAAAACATTAAGCTTAACTTAGTGAAGGGCAAGAAAAACTGCTAAATTAGGGACCGGGAGGAGGCAGAAGTATTCCTTGGGAAGCGGTAAAATGTTATAATCCTTGGAAGACTACCTGTGGCGAAGGCGTCTGCCTAGAACGGATCCGACGGTGAGGGACGAAAGCTAGGGGAG |
| p__Thermoplasmatota_g__EX4484-6_ASV_3386 | 0 | 0 | 0 | 0 | 1 | 0 | 0 | 0 | 0 | 0 | 0 | ACTTCGCAATGCGCGCAAGCGCGACGAGGGGATTCCAAGTGCATGCACTCAGTGTATGCTGTTTTCCAGTCTAAAAAGCTGGGAGAGTAAGGGCTGGGTAAGACTGGTGCCAGCCGCCGCGGTAATACCAGCGGCCCTAGTGGTGATCGTTATTATTGGGCCTAAAGCGTCCGTAGCCGGCTCGGTAAATCTCTGGGTAAATCGTTAGGCTTAACTCAACGAATTCTGGGGAGACTGCCGGGCTTGGGACCGGGAGAGGTCGGAGGTACTCCAGGGGTAGGGGTGAAATCCTGTAATCCTTGGGGGACCACCGGTGGCGAAGGCGTCCGACCAGAACGGGTCCGACGGTAAGGGACGAAGCCCTGGGGCG |
| p__Halobacteriota_g__Halapricum_3981_ASV_3409 | 0 | 0 | 0 | 0 | 0 | 1 | 0 | 0 | 0 | 0 | 0 | CCTTTACACTGCACGACAGTGCGATAGGGGGACTCCGAGTGCGAGGGCATATAGTCCTCGCTTTTCTGTACCGTAAGGTGGTACAGGAACAAGTGCTGGGCAAGACCGGTGCCAGCCGCCGCGGTAATACCGGCAGCACGAGTGATGGCCGATATTATTGGGCCTAAAGCGTCCGTAGCCGGCCAGACAAGTCCGTTGGGAAATCGGCGTGCCTAACACGTCGGCGTCCAGCGGAAACTGTTTGGCTTGGGGCCAGAAGATCCGAGGGGTACGTCCGGGGTAGGAGTGAAATCCCGTAATCCTGGACGGACCACCGGTGGCGAAAGCGCCTCGGAAAGATGGACCCGACGGTGAGGGACGAAAGCTGGGGTCT |
| p__Halobacteriota_g__Haloarchaeobius_ASV_3268 | 0 | 0 | 0 | 0 | 0 | 0 | 0 | 0 | 0 | 0 | 1 | ACTTTACACTGCACGACAGTGCGATAAGGGAACTCCAAGTGCGAGGGCATAGAGCCCTCGCTTTTCTGTACCGTAGGGTGGTACAGGAATAAGAGCTGGGCAAGACCGGTGCCAGCCGCCGCGGTAACACCGGCAGCTCGAGTGATGACCGATATTATTGGGCCTAAAGCGTCCGTAGCTGGCCGAGAGGGTCCGTCGGGAAATCTGCTCGCTCAACGAGCAGGCGTCCGGCGGAAACCAATCGGCTTGGGACCGGAAGACCTGAGGGGTACGTCTGGGGTAGGAGTGAAATCCCGTAATCCTGGACGGACCACCGGTGGCGAAAGCGCCTCAGGAAGACGGATCCGACAGTGAGGGACGAAAGCTGGGGTCA |
| p__Halobacteriota_g__Halobacterium_ASV_3447 | 0 | 0 | 0 | 0 | 0 | 0 | 0 | 0 | 1 | 0 | 0 | CCTTTACACTGCACGACAGTGCGATAGGGGGACTCCGAGTGTGAAGGCATAGAGCCTTCACTTTTGTACACCGTAAGGTGGTGTACGAATAAGGACTGGGCAAGACCGGTGCCAGCCGCCGCGGTAATACCGGCAGTCCGAGTGATGGCCGATCTTATTGGGCCTAAAGCGTCCGTAGCTGGCTAGACAAGTCCGTTGGGAAATCTGCCCGCTTAACGGGCAGGCGTCCAGCGGAAACTGTCTAGCTTGGGACCGGAAGACCTGAGGGGTACGTCTGGGGTAGGAGTGAAATCCCGTAATCCTGGACGGACCGCCGGTGGCGAAAGCGCCTCAGGAGAACGGATCCGACAGTGAGGGACGAAAGCTAGGGTCT |
| p__Halobacteriota_g__Halobellus_3026_ASV_3349 | 0 | 0 | 0 | 1 | 0 | 0 | 0 | 0 | 0 | 0 | 0 | CCTTTACACTGCACGAAAGTGCGATAAGGGGACTCCGAGTGCGAGGGCATATCGTCCTCGCTTTTCTGTACCGTAGGGCGGTACAGGAACAAGAGCTGGGCAAGACCGGTGCCAGCCGCCGCGGTAATACCGGCAGCTCAAGTGATGGCCGCTCTTATTGGGCCTAAAGCGTCCGTAGCTGGCCTCGCAAGTCCGTCGGGAAATCCACTGGCCCAACCAGTGGGCGTCCGGCGGAAACTCCGAGGCTTGGGACCGGAAGGCTCGAGGGGTACGTCCGGGGTAGGAGTGAAATCCTGTAATCCCGGACGGACCACCGATGGCGAAAGCACCTCGAGAGGACGGATCCGACAGTGAGGGACGAAAGCTAGGGTCT |
| p__Halobacteriota_g__Halohasta_ASV_3324 | 0 | 0 | 1 | 0 | 0 | 0 | 0 | 0 | 0 | 0 | 0 | CCTTTACACTGCACGACAGTGCGATAAGGGGACTCCGAGTGCGAGGGCATATAGTCCTCGCTTTTCTGCACCGTAAGGTGGTACAGGAATAAGGACTGGGCAAGACCGGTGCCAGCCGCCGCGGTAATACCGGCAGTCCGAGTGATGTCCACTATTATTGGGCCTAAAGCGTCCGTAGCTGGCCGCGCAAGTCCGTCGGGAAATCCACCTGCCCAACAGGTGGGCGTCCGGCGGAAACTGCGTGGCTTGGAACCGGAAGGCTCAGAGGGTACGTCCGGGGTAGGAGTGAAATCCCGTAATCCCGGACGGACCACCGATGGCGAAAGCACTCTGAGAGGACGGATTCGACAGTGAGGGACGAAAGCTAGGGTCT |
| p__Halobacteriota_g__Halomicrobium_3849_ASV_3458 | 0 | 0 | 0 | 0 | 0 | 0 | 0 | 0 | 1 | 0 | 0 | CCTTTACACTGCACGCCAGTGCGATAGGGGGACTCCGAGTGCGAGGGCATACAGTCCTCGCTTTTCTGTACCGTAAGGTGGTACAGGAACAAGTGCTGGGCAAGACCGGTGCCAGCCGCCGCGGTAATACCGGCAGCACGAGTGATGGCCGATCTTATTGGGCCTAAAGCGTCCGTAGCCGGCCGGACAAGTCCGTTGGGAAATCGACGCGCTCAACGCGTCGGCGTCCAGCGGAAACTGTCCGGCTTGGGGCCGGAAGACCTGAGGGGTACGTCCGGGGTAGGAGTGAAATCCTGTAATCCTGGACGGACCACCGGTGGCGAAAGCGCCTCAGGAAGACGGACCCGACGGTGAGGGACGAAAGCTAGGGTCT |
| p__Halobacteriota_g__Haloprofundus_ASV_3238 | 0 | 0 | 0 | 0 | 0 | 0 | 0 | 0 | 0 | 1 | 0 | ACTTTACGCTGCACGTCAGTGCGATAAGGGAACTCCAAGTGCGAGGGCATACAGTCCTCGCTTTTCCCGACCGTAAGAAGGTCGGAGAAGAAGAGCTGGGCAAGACCGGTGCCAGCCGCCGCGGTAATACCGGCAGCTCGAGTGATGACCGCTGTTATTGGGCCTAAAGCGTCCGTAGCTGGCCGCGTAAGTCCATCGGGAAATCCATCCGCTCAACGGATGGGCGTCCGGTGGAAACTACACGGCTTGGAGCCGGAAGACCCAAGGGGTACGTCCGGGGTAGGAGTGAAATCCCGTAATCCTGGACGGACCGCCGATGGCGAAAGCACCTTGGGAAGACGGACTCGACAGTGAGGGACGAAAGCTAGGGTCT |
| p__Halobacteriota_g__Haloprofundus_ASV_3311 | 0 | 1 | 0 | 0 | 0 | 0 | 0 | 0 | 0 | 0 | 0 | CCTTTACACTGCACGCAAGTGCGATAGGGGAATCCCAAGTGCGAGGGCATACTAGCCCTCGCTTTTGTGTACTGTAAGGAGGTACACGAATAAGAGCTGGGCAAGACCGGTGCCAGCCGCCGCGGTAATACCGGCAGCTCGAGTGATGACCGCTGTTATTGGACCTAAAGCGTCCGTAGCTGGCCGCGTAAGTTCATCGGGAAATCCATCCGCTCAACGGATGGGCGTCCGGTGAAAACTACACGGCTTGGAGCCGGAAGACCCAAGGGGTACGTCCGGGGTAGGAGTGAAATCCCGTAATCCTGGACGGACCGCCGATGGCGAAAGCACCTTGGGAAAACGGACTCGACAGTGAGGGACGAAAGCTAGGGTCT |
| p__Halobacteriota_g__Halorientalis_ASV_3332 | 0 | 0 | 1 | 0 | 0 | 0 | 0 | 0 | 0 | 0 | 0 | CCTTTACACTGCACGACAGTGCGATAGGGGGACTCCGAGTGCGAGGGCATACAGTCCTCGCTTTTCTGTACCGTAAGGTGGTACAGGAACAAGTGCTGGGCAAGACCGGTGCCAGCCGCCGCGGTAATACCGGCAGCACGAGTGATGGCCGCTATTATTGGGCCTAAAGCGTCCGTAGCCGGCCGAGCAAGTTCGTCGGGAAATCTGCCCGCTCAACGGGCAGGCGTCCGGCGAAAACTGTTCGGCTTGGGGCCGGGAGATCCAAGGGGTACGTCCGGGGTAGGAGTGAAATCCCGTAATCCTGGACGGACCACCGGTGGCGAAAGCGCTCTGGAAGGACGGATCCGACGGTGAGGGACGAAAGCTTGGGTCA |
| p__Halobacteriota_g__Halorubrum_ASV_3278 | 0 | 0 | 0 | 0 | 0 | 0 | 0 | 0 | 0 | 0 | 1 | CCTTTACACTGCACGCAAGTGCGATAAGGGGATCCCAAGTGCGAGGGCATACAGTCCTCGCTTTTCTCACCCGTAAGGAGGTTGAGGAATAAGGGCTGGGCAAGACCGGTGCCAGCCGCCGCGGTAATACCGGCAGCCCAAGTGATGACCGATATTATTGGGCCTAAAGCGTCCGTAGCTGGCTGCGCAAGTCCATCGGGAAATCCACCTGCCCAACAGGTGGGCGACCGGTGGAAACTGCGCGGCTTGGGACCGGAAGGCTCAGAGGGTACGTCCGGGGTAGGAGTGAAATCCCGTAATCCCGGACGGACCGCCGATGGCGAAAGCACTCTGAGAAGACGGATCCGACAGTGAGGGACGAAAGCTGGGGTCT |
| p__Halobacteriota_g__Halorussus_ASV_3267 | 0 | 0 | 0 | 0 | 0 | 0 | 0 | 0 | 0 | 0 | 1 | TCTTTACACTGCACGACAGTGCGATAAGGGGACTCCGAGTGCGAGGGCATATAGTCCTCGCTTTTCTGTACCGTAAGGTGGTACAGGAATAAGGGCTGGGCAAGACCGGTGCCAGCCGCCGCGGTAATACCGGCAGCCCGAGTGATGGCCGCTATTATTGGGCCTAAAGCGTCCGTAGCCGGCCAGACAGGTCCGTCGGGAAATCCACGCGCTCAACGCGTGGGCGTCCGGCGGAAACCAGCTGGCTTGGGGCCGGAAGATCTGAGGGGTACGTCCGGGGTAGGAGTGAAATCCCGTAATCCTGGACGGACCACCGGTGGCGAAAGCGCCTCAGAAAAACGGACCCGACGGTGAGGGACGAAAGCTAGGGTCA |
| p__Halobacteriota_g__Halorussus_ASV_3415 | 0 | 0 | 0 | 0 | 0 | 1 | 0 | 0 | 0 | 0 | 0 | CCTTTACACTGCACGACAGTGCGATAAGGGGACTCCGAGTGCGAGGGCATATAGTCCTCGCTTTTGTACACCTTAAGGCGGTGTACGAATAAGGGCTGGGCAAGACCGGTGCCAGCCGCCGCGGTAATACCGGCAGCCCGAGTGATGGCCGCTATTATTGGGCCTAAAGCGTCCGTAGCCGGCCAGACAGGTCCGTCGGGAAATCCACGCGCTCAACGCGTGGGCGTCCGGCGGAAACCAGCTGGCTTGGGGCCGGAAGACCTGAGGGGTACGTCCGGGGTAGGAGTGAAATCCCGTAATCCTGGACGGACCGCCGGTGGCGAAAGCGCCTCAGGAAGACGGACCCGACGGTGAGGGACGAAAGCTAGGGTCA |
| p__Halobacteriota_g__Halovenus_ASV_3233 | 0 | 0 | 0 | 0 | 0 | 0 | 0 | 0 | 0 | 1 | 0 | CCTTTACACTGCACGACAGTGCGATAGGGGGACTCCAAGTGCGAGGGCATATAGTCCTCGCTTTTGTACACCGTAAGGTGGTGTACGAATAAGAACTGGGCAAGACCGGTGCCAGCCGCCGCGGTAATACCGGCAGTTCGAGTGATGGCCGATTTTATTGGGCCTAAAGCGTTCGTAGCTGGCCAGTCAAGTCTGTCGGGAAATCCAGTGGCTTAACCACTGGACGTCCGGCGGAAACTGATTGGCTTGGGACCGGGAGACTCGAGGAGTACGTTCGGGGTAGGAGTGAAATCCCATAATCCTGAACGGACTACCGATGGCGAAAGCACCTCGAGAGCACGGATCCGACAGTGAGGAACGAAAGCTGGGGTCT |
| p__Halobacteriota_g__Halovenus_ASV_3352 | 0 | 0 | 0 | 1 | 0 | 0 | 0 | 0 | 0 | 0 | 0 | CCTTTACACTGCACGACAGTGCGATAGGGGGACTCCGAGTGCGAGGGCATACAGTCCTCGCTTTTGTACACCGTAAGGTGGTGTACGAATAAGAACTGGGCAAGACCGGTGCCAGCCGCCGCGGTAATACCGGCAGTTCGAGTGATGGCCGCTTTTATTGGGCCTAAAGCGTTCGTAGCCGGCCGAGCAAGTCCGTCGGGAAATCCAGTGGCTTAACCACTGGACGTCCGGCGGAAACTGCTCGGCTTGGGACCGGGAGACTCGAGAAGTACGTCCGGGGTAGGAGTGAAATCCCGTAATCCTGGACGGACTACCGATGGCGAAAGCATCTCGAGAGCACGGATCCGACGGTGAGGAACGAAAGCTGGGGTCT |
| p__Halobacteriota_g__Halovivax_ASV_3216 | 1 | 0 | 0 | 0 | 0 | 0 | 0 | 0 | 0 | 0 | 0 | CCTTTACACTGCGCGACAGCGCGATAAGGGGACTCCACGTGCCAGGGCATATCGTCCTGGCTTTTGTGTACCGTAAGGTGGTACACGAATAAGGGCTGGGCAAGACCGGTGCCAGCCGCCGCGGTAATACCGGCAGCCCGAGTGATGACCGCTCTTATTGGGCCTAAAGCGTCCGTAGCTGGCCATCCAAGTCCATCGGGAAATCCGCACGCCTAACGTGCGGGCGTCCGGTGGAAACTGGATGGCTTGGGACCGGAAGATCCAGAGGGTACGTCTGGGGTAGGAGTGAAATCCCGTAATCCTGGACGGACCACCGGTGGCGAAAGCGCTCTGGAAGGACGGATCCGACGGTGAGGGACGAAAGCTAGGGTCA |
| p__Halobacteriota_g__Methanolobus_ASV_3318 | 0 | 1 | 0 | 0 | 0 | 0 | 0 | 0 | 0 | 0 | 0 | ACTTTACAATGCGGGAAACCGCGATAAGGGGACACCGAGTGCCAGCATATTATGTTGGCTGTCCGCATGTATAAATAACATGTGTTAGCAAGGGCCGGGCAAGACCGGTGCCAGCCGCCGCGGTAACACCGGCGGCCCGAGTGGTAGCCACTTTTATTGGGTCTAAAGGGTCCGTAGCCGGTTTGATCAGTCTTCCGGGAAATCTGACAGCTTAACTGTTAGGCTTCTGGGGGATACTGTCAGACTTGGAACCGGGAGAGGTAAGAGGTACTACAGGGGTAGGAGTGAAATCTTGTAATCCCTGTGGGACCACCAGTGGCGAAGGCGTCTTACCAGAACGGGTTCGACGGTGAGGGACGAAAGCTGGGGGCA |
| p__Nanohaloarchaeota_g__Nanohalobium_ASV_3274 | 0 | 0 | 0 | 0 | 0 | 0 | 0 | 0 | 0 | 0 | 1 | CCTTTACAATGCACGCAAGTGCGATAAGGGGACTCCGAGTGCGGTCTTCCTACACGGAAGGCCGCTTTTCTGTACTGTAAGAAGGTACCGGAACAAGGGGTGGGCAATACTGCTGCCAGCCGCCGCGGTAATAGCAGAACCCCAAGTGGTAACCAGTTTTATTGGGCTTAAAGTGTTCGTAGCCGGCCATGTAAGTCCGCTGTGAAATCCAAGCGCTTAACGTTTGGAAGTGCAGCGGATACTGCATGGCTAGGAACCGGATGGCGTCGAGGCTATTCCAGGGGTAGGAGTGAAATCCAAAAATCCCTGGAGGAGCACCTGTGGCGAAAGCGCTCGACGAATACGGATTCGACGGTGAGGAACGAAAGCCAGGGGAG |
| p__Halobacteriota_g__Natronomonas_ASV_3389 | 0 | 0 | 0 | 0 | 1 | 0 | 0 | 0 | 0 | 0 | 0 | CCTTTACACTGCACGCCAGTGCGATAGGGGGACACCGAGTGCGAGGGCATAGAGCCCTCGCTTTTCTGTACCGTAAGGTGGTACAGGAACAAGGGCTGGGCAAGACCGGTGCCAGCCGCCGCGGTAACACCGGCAGCCCGAGTGATGGCCGATATTATTGGGCCTAAAGCGTCCGTAGCTGGCCGGGCAAGTCCGTCGGGAAATCCACTCGCTCAACGAGTGGGCGTCCGGCGGAAACTGTTCGGCTTGGGACCGGAAGATCCGAGGGGTACGTCCGGGGTAGGAGTGAAATCCCGTAATCCTGGACGGACCACCGATGGCGAAAGCACCTCGGAAGGACGGATCCGACAGTGAGGGACGAAAGCTAGGGTCT |
| p__Halobacteriota_g__Salinigranum_ASV_3353 | 0 | 0 | 0 | 1 | 0 | 0 | 0 | 0 | 0 | 0 | 0 | CCTTTACACTGCGCGCAAGCACGATAAGGGGACCCCAAGTGCGAGGGCATACAGTCCTCGCTTTTGTGTACCGTAAGGAGGTACACGAATAAGAGCTGGGCAAGACCGGTGCCAGCCGCCGCGGTAATACCGGCAGCTCGAGTGATGGCCGCTATTATTGGGCCTAAAGCGTCCGTAGCTGGCCACGCAAGTCCGTCGGGAAATCCACCAGCCCAACTGGTGGGCGGCCGACGGAACCTGCGTGGCTTGGGACCGGAAGACCCAACGGGTACGTCCGGGGTAGGAGTGAAATCCCGTAATCCCGGACGGACCACCGATGGCGAAAGCACGTTGGGAGGACGGCTCCGACAGTGAGGGACGAAAGCTGGGGTCT |

| Phylum_Genus_ASV_ number | Presence (1) or Absence (0) in each layer | | | | | | | | | | | |
|---|---|---|---|---|---|---|---|---|---|---|---|---|
| | L1 | L2 | L3 | L4 | L5 | L6 | L7 | L8 | L9 | L10 | L11 | Sequence |
| p__Thermoplasmatota_ g__SM1-50_ASV_3345 | 0 | 0 | 1 | 0 | 0 | 0 | 0 | 0 | 0 | 0 | 0 | CCTTTACACTGCACGCCAGTGCGATAAGGGGACACCAAGTGCGAGGGCATATCGTCCTCGCTTTTCACCACCGTAAGGAGGTGGTAGAATAAGAGCTGGGCAAGACCGGTGCCAGCTGCCGCGGTAATACCGGCAGCTCGAGTGATGACCGCTATTATTGGGCCTAAAACGTCCGTAGCCGGCTGAGTAAATGCCTGGGTAAATCGTGTCGCTTAACGATACGAATTCCGGGTAGACTGCTTAGCTTGAGACTGGGAGAAGCTAGAGGTACTCCTGGGGTAGAGGTGAAATTCTGTAATCCTGGGGGGACCACCAGTGGCGAAAGCGTCTAGCTAGAACGGGTCTGACGGTGAGGGACGAAACCCTGGGGCG |
| p__Thermoplasmatota_ g__SM1-50_ASV_3445 | 0 | 0 | 0 | 0 | 0 | 0 | 0 | 1 | 1 | 0 | 0 | ACTTCACACTGCGCGAAAGCGCGATGAGGGAATCCCAAGTGCTTGCACGTAAGTGTAAGCTGTTTTTCTGTTTAAAAAGCAGAGAGAGTAAGGGCTGGGTAAGACGGGTGCCAGCCGCCGCGGTAATACCTGCAGCTCAAGTGGTGGCCGCTATTATTGGGCCTAAAACGTTCGTAGCCTGTTTATTAAATGCCTAGGTAAATCGGGCCGCTTAACGGTTCGAATTCTGGGTAGACTGATAAACTCGAGACTGGGAGAGGCTAGAGGTACTCCTGGGGTAGAGGTGAAATTCTGTAATCCTAGGGGGACCACCAGTGGCGAAGGCGTCTAGCTAGAACGGGTCTGACGGTGAGGGACGAAGCCCTGGGGCG |
| p__Nanoarchaeota_p__ Nanoarchaeota_ASV_3272 | 0 | 0 | 0 | 0 | 0 | 0 | 0 | 0 | 0 | 0 | 1 | TCTCCACAATGCGCGAAAGCGTGATGGGGGAACTCCAAGTGCGCTACCACTGGTAGTGCTGTGGCGAAATGTACAAAGTTTCGCAAGAAAGGGGTGGGCAAGACAGGTGCCAGCAGCCGCGGTAATCCCTGCGCCTCAAGTCGTGACCATGCTTATTGGGCCTAAAGTATCCGTAGCCGGAGTATTAAGTTCTCTGTGAAATCGTACAGCTTAACTGTACGGCGTGCAGGGAATACTGTTATTCTAGGGACCGGGAGGTGTCAAGGGTATTTCTGGGGTAAGGGTTAAATCTTATAATCCCAGGAGGACCAACGGTGGCGAAGGCGCTTGACAAGAACGGGTCCGACGGTGAATGAATGAAAGCTGGGGGAG |
| p__Halobacteriota_f__H aloarculaceae_3992_AS V_3351 | 0 | 0 | 0 | 1 | 0 | 0 | 0 | 0 | 0 | 0 | 0 | CCTTTACACTGCACGACAGTGCGATAGGGGGACTCCGAGTGCGAGGGCATACAGTCCTCGCTTTTCTGTACCGTAAGGTGGTACAGGAATAAGTGCTGGGCAAGACCGGTGCCAGCCGCCGCGGTAATACCGGCAGCACGAGTGATGGCCGATATTATTGGGCCTAAAGCGTCCGTAGCCGGCCAGACAAGTCCGTTGGGAAATCGACGTGCCTAACACGTCGGCGTCCAGCGGAAACTGTCTGGCTCGGGGCCGGAAGACTCGAGGGGTACGTCCGGGGTAGGAGTGAAATCCCGTAATTCTGGATGGACCGCCGGTGGCGAAAGCGCCTCGAGAGGACGGACCCGACGGTGAGGGACGAAAGCTAGGGTCT |
| p__Halobacteriota_f__H aloferacaceae_ASV_3212 | 1 | 0 | 0 | 0 | 0 | 0 | 0 | 0 | 0 | 0 | 0 | CCTTTACACTGCACGCCAGTGCGATAAGGGGACCCCGAGTGCGAGGGCATACAGTCCTCGCTTTTGTGTACCGTAAGGTGGTACACGAATAAGGGCTGGGCAAGACCGGTGCCAGCCGCCGCGGTAATACCGGCAGCCCGAGTGATGGCCGCTCTTATTGGGCCTAAAGCGTCCGTAGCCGGCCAGACAGGTCCGTCGGGAAATCCACGCGCTCAACGCGTGGGCGTCCGGCGGAACCCCGCTGGCTAGGGACCGGACGACCCGAGGGGTACGTCTGGGGTAGGAGTGAAATCCTGTAATCCTGGACGGACCGCCGATGGCGAAAGCACGTCGCGAGAACGGATCCGACAGTGAGGGACGAAAGCCAGGGTCT |
| p__Halobacteriota_f__H aloferacaceae_ASV_3342 | 0 | 0 | 1 | 0 | 0 | 0 | 0 | 0 | 0 | 0 | 0 | CCTTTACACTGCACGACAGTGCGATAAGGGGACCCCGAGTGCGAGGGCATACAGTCCTCGCTTTTGTGTACCGTAGGGAGGTACACGAATAAGAGCTGGGCAAGACCGGTGCCAGCCGCCGCGGTAATACCGGCAGCTCAAGTGATGGCCGCTATTATTGGGCCTAAAGCGTCCGTAGCCGGCCGCACAAGTCCGTCGGGAAATCCACCCGCCTAACGGGTGGGCGTCCGGCGGAAACTGTGCGGCTTGGGACCGGAAGACTCGAGGGGTACGTCCGGGGTAGGAGTGAAATCCTGTAATCCCGGACGGACCACCGATGGCGAAAGCACTCTGAGAGGACGGATTCGACAGTGAGGGACGAAAGCTAGGGTCT |
| p__Halobacteriota_f__N atrialbaceae_ASV_3330 | 0 | 0 | 1 | 0 | 0 | 0 | 0 | 0 | 0 | 0 | 0 | CCTTTACACTGCGCGACAGCGCGATAAGGGGACTCCAAGTGCGAGGGCATATCGTCCTCGCTTTTCACTATCGTAAGGAGATGGTAGAATAAGGGCTGGGCAAGACCGGTGCCAGCCGCCGCGGTAATACCGGCAGCCCGAGTGATGACCGCTATTATTGGGCCTAAAGCGTCCGTAGCCGGCCGAACAAGTCCGTCGGGAAATCTGCACGCCTAACGTGCAGGCGTCCGGCGGAAACTGTTCGGCTTGGGACCGGAAGATCCAGAGGGTACGTCCGGGGTAGGAGTGAAATCCTGTAATCCCGGACGGACCACCGATGGCGAAAGCACTCTGAGAGGACGGATTCGACAGTGAGGGACGAAAGCTAGGGTCT |
| p__Thermoproteota_f__ Nitrososphaeraceae_AS V_3417 | 0 | 0 | 0 | 0 | 0 | 1 | 0 | 0 | 0 | 0 | 0 | ACTCTGCAATAAGCGAAAGCTTGACAGGGTTATTCTGAGTGATTTCCGCTAAGGAAATCTTTTGGCACCTCTAAAAATGGTGCAGAATAAGGGGTGGGCAAGCCTGGTGTCAGCCGCCGCGGTAATACCAGCACCCCGAGTGGTCGGGACGATTATTGGGCCTAAAGCATCCGTAGCCGGTTCTACAAGTCTTCCGTTAAATCCATCTGCTTAACAGATGGGCTGCGGAGGATACTATGGAGCTAGGAGGCGGGAGAGGCAAGCGGTACTTAGTGGGTAGGGGTAAAATCCGTTGATCCACTGAAGACCACCAGTGGCGAAGGCGGCTTGCCAGAACGCGCTCGACGGTGAGGGATGAAAGCTGGGGGAG |
| p__Halobacteriota_o__ Halanaerobiales_ASV_3235 | 0 | 0 | 0 | 0 | 0 | 0 | 0 | 0 | 0 | 1 | 0 | CCTTTACACTGCACGCAAGTGCGATAGGGGGACCCCAAGTGCGAGGGCATACAGTCCTCGCTTTTCACAGCCGTAAGGAGGCTGTAGAATAAGGGCTGGGCAAGACCGGTGCCAGCCGCCGCGGTAATACCGGCAGCCCAAGTGATGGCCGCTGTTATTGGGCCTAAAGCGTCCGTAGCCGGCCGCGTGTGTCCATCGGGAAATCCACCCGCTCAACGGGTGGGCGGCCGATGGAAACTGCGCGGCTTGGGGCCGGAAGACCCACCGGGTACGTCCGGGGTAGGAGTGAAATCCCGTAATCCCGGACGGACCACCGATGGCGAAAGCACGGTGGGAAGACGGACCCGACGGTGAGGGACGAAAGCTGGGGTCT |
| p__Halobacteriota_o__ Halanaerobiales_ASV_3281 | 0 | 0 | 0 | 0 | 0 | 0 | 0 | 0 | 0 | 0 | 1 | CCTTTACACTGCACGACAGTGCGATAAGGGGACCCCAAGTGCGAGGGCATATCGTCCTCGCTTTTCTGTACCGTAAGGAGGTGCAGGAATAAGGGCTGGGCAAGACCGGTGCCAGCCGCCGCGGTAATACCGGCAGCCCAAGTGATGGCCGCTATTATTGGGCCTAAAGCGTCCGTAGCCGGCCGGACAAGTCCGTCGGGAAATCCACCCGCTCAACGGGTGGGCGTCCGGCGGAAACTGTTCGGCTTGGGGCCGGAGGACCTGAGGGGTACGTCCGGGGTAGGAGTGAAATCCTGTAATCCTGGACGGACCACCGGTGGCGAAAGCGCTCTGGAAGGACGGATCCGACGGTGAGGGACGAAAGCTAGGGTCA |
| p__Halobacteriota_o__ Halanaerobiales_ASV_3329 | 0 | 0 | 1 | 0 | 0 | 0 | 0 | 0 | 0 | 0 | 0 | CCTTTACACTGCACGACAGTGCGATAAGGGGACTCCAAGTGCGAGGGCATATCGCCCTCGCTTTTGTACACCGTAAGGTGGTGCACGAATAAGGGCTGGGCAAGACCGGTGCCAGCCGCCGCGGTAATACCGGCAGCCCGAGTGATGGCCGCTATTATTGGGCCTAAAGCGTCCGTAGCCGGCCAGTCAAGTCCGTCGGGAAATCCACCCGCTTAACGGGTGGGCGTCCGGCGGAAACTGACTGGCTTGGGACCGGAAGACCCGGGGGGTACGTCTGGGGTAGGAGTGAAATCCTGTAATCCTGGACGGACCGCCGGTGGCGAAAGCGCCTCGGGAGAACGGATCCGACGGTGAGGGACGAAAGCTTGGATCT |
| p__Halobacteriota_o__ Halanaerobiales_ASV_3335 | 0 | 0 | 1 | 0 | 0 | 0 | 0 | 0 | 0 | 0 | 0 | CCTTTACACTGCACGACAGTGCGATAAGGGGACTCCGAGTGCGAGGGCATAACGTCCTCGCTTTTCTGGACCGTAAGGTGGTCCAGGAATAAGGGCTGGGCAAGACCGGTGCCAGCCGCCGCGGTAATACCGGCAGCCCGAGTGATGGCCGTCATTATTGGGCCTAAAGCGTCCGTAGCCGGCCGGGCAAGTCTGTTGGGAAATCCACCTGCTTAACGGGTGGGCGTCCAGCGGAAACTGCTCGGCTTGGGACCGGAAGACTCGAGGGGTACGTCTGGGGTAGGAGTGAAATCCTGTAATCCTGGACGGACCGCCGGTGGCGAAAGCGCCTCGAGAGGACGGATCCGACGGTGAGGGACGAAAGCTTGGATCT |
| p__Halobacteriota_o__ Halanaerobiales_ASV_3346 | 0 | 0 | 1 | 0 | 0 | 0 | 0 | 0 | 0 | 0 | 0 | CCTTTACACTGTACGAAAGTGCGATAAGGGGACCCCGAGTGCGAGGGCATACAGTCCTCGCTTTTGTGTACCGTAAGATGGTACACGAATAAGAGCTGGGCAAGACCGGTGCCAGCCGCCGCGGTAATACCGGCAGCTCAAGTGATGGCCGCTATTATTGGGCCTAAAGCGTCCGTAGCCGGCCGAACAGGTCCGTTGGGAAATCCACGCGCTCAACGCGTGGGCGTCCAGCGGAAACTGTCCGGCTTGGGACCGGAAGACCCAGAGGGTACGTCCGGGGTAGGAGTGAAATCCCGTAATCCTGGACGGACCACCGGTGGCGAAAGCGCTCTGAGAGGACGGATTCGACAGTGAGGGACGAAAGCTAGGGTCT |
| p__Nanoarchaeota_o__ Woesearchaeales_581_ ASV_3452 | 0 | 0 | 0 | 0 | 0 | 0 | 0 | 0 | 1 | 0 | 0 | CCTTTACAATGCACGAAAGTGTGATAAGGGGATCCTACGTGCTCATGCATTGCATGGGCTTTTGCCAAGAGTAAATATCTTGGAGAATAAGTGGTGGGTAAGACTGGTGCCAGCCGCCGCGGTAACACCAGCGCCACGAGTGGCAGTCACGATTATTGGGCCTAAAGCGTCCGTAGCCGGACGTGCACATCCCTTGTGAAATCACACGGCTTAACCGTGTGGCGTGCAGGGGACACGACACGTCTTGGGAGTGGGAGGAGTCAGAGGTATTCGAGGGGGACCGGTAAAATGGTATAATCCCTTGAGGACCACCGATGGCGAAGGCATCTGACCAGAACACGTCCGACGGTGAGGGACGAAAGCCAGGGGAG |
| p__Thermoplasmatota_ p__Thermoplasmatota_ ASV_3405 | 0 | 0 | 0 | 0 | 0 | 1 | 0 | 0 | 0 | 0 | 0 | ACTGCACAATGCTGGAAACAGCGATGCGGGAACTCCAAGTGCCGGCACAACGTGTCGGCTGTTCCATTGCCTAAAAAGCATAGGAAGTAAGGGCTGGGTAAGACGGGTGCCAGCCGCCGCGGTAATACCTGCAGCCCGAGTGGTGGTCGTTATTATTGGGCCTAAAACGTCCGTAGCCGGTTTGGTAAATGCCTGTGTAAATCGAACGGCTTAACCGTTCGAACTGCGGGTAGACTGCCAGACTTGGGACCGGGAGAGTCACGAAGTACTCCGGGGGTAGAGGTGAAATTCTGTAATCCTCGGGGGACTACCGGTGGCGAAGGCGTCGTGATAGAACGGCTCCGACGGTGAGGGACGAAGCCCTGGGGCG |
| p__Iainarchaeota_o__Iai narchaeales_ASV_3492 | 1 | 0 | 0 | 0 | 0 | 0 | 0 | 0 | 0 | 0 | 1 | CCTCTACAATGCACGAAAGTGTGATAGGGGGACTTCGAGTGAGCTCTCTGAGTTCTTTTGCCAAGCCTAAAAAGCTTGGCGAATAAGTGGGGGGTAATATACGTGCCAGCCGCCGCGGTAATACGTAACCCACAAGTGGTATCCACTAATATTGGGCCTAAAACGTCTGTAGCCTGTTTTGTAAGTCTTTTGTGAAATCAGCCAGCTCAACTGGTTGAATTGCAAAAGATACTGCACGACTCGAGGCCGGAAGAGGTAAAGGGAATTTCAGGGGTAACGGTGAAATGTTATGATCCCTGAAAGACCACCAGTAGCGAAGGCGCTTTACCAGGACGGTCCTGACGGTGAGAGACGAAAGCTGGGGGAG |
| p__Nanoarchaeota_g__ BMS3ABIN17_ASV_3496 | 0 | 0 | 0 | 0 | 0 | 0 | 0 | 0 | 0 | 1 | 0 | ATTTTACAATGCTCGCAAGAGTGATAAAATAAGCCAGAGTGTTTCTCTTAGAGAAACTTTTTTTTGATGTAAAAAGTCAGAAGAATAAGGACTGGGCAAGACCGGTGCCAGCCGCCGCGGTAATACCGGCGGTCCAAGTCGCATCCACAATTATTGGGTCTAAAACATTCGTAGCTTGTTTTACAAGTCTCTTGTGAAATCTGACATCTCAAGTGTCAGGCGTGCAAGAGATACTGTATTACTAGAGACCGGAAGATGTAAAGAGTATGGTTGAGGTAGCGGTAAAATGTGTTAATCTCAATTAGACTAACAATGGCGAAGGCACTTTACAAGTACGGATCTGACAGTGAGGAATGAAGGCTAGGGGCG |
| p__Nanoarchaeota_g__ BMS3ABIN17_ASV_3614 | 0 | 1 | 0 | 0 | 0 | 0 | 0 | 0 | 0 | 0 | 0 | TTTTTGCAATGCGCGAAAGCGTGACAGAGCGAACCAGAGTGCTTTTCCGTATAGGAAAAGCTTTTGTGAGATGTAAAAAGTCTCACGAATAAGGACTGGGCAAGACTGGTGCCAGCCGCCGCGGTAATACCAGCGGTCCAAGTCGCAACCACATTTATTGGGTCTAAAACATCCGTAGCTTGTTGTGTAAGTCCCTTGTGAAATCCGGCTTCTTAAGAGTCGAAGTGCAAGGGATACTGCATAGCTAGGGACCGGGAGACGTAAGGAGTACGGATAGGGTAACGGTTAAATGTGTTGATCCTATCTGGACTAACGATAGCGAAGGCACCTTACGAGAACGGATCTGACAGTGAGGGATGAAGGCTAGGGGCG |
| p__Nanoarchaeota_g__ BMS3ABIN17_ASV_3625 | 0 | 0 | 1 | 0 | 0 | 0 | 0 | 0 | 0 | 0 | 0 | AATCTACAATACGCGAAAGCGTGATAGTTTGAGTCAGAGTGCGTTTCTTTGAAATGCTTTTGTGAAATGCAAAAAGTTTCACGAATAAGGACTGGGTAAGACGAGTGCCAGCCGCCGCGGTAATACTCGCGGTCCAAGTCGCAGCCATTTTTATTGGGTCTAAAACATCCGTAGCTTGCTTTGTAAGTTCCTTGTGAAATCCTATATCTCAAATATAGGGCGGGCAAGGAGTACTGCTAAGCTAGAGACTGGGAGACGTAAAGAGTACGTTTAAGGTAGCGGTAAAATGTGTTAATCTTAAGCGGACTAACAATAGCGAAGGCACTTTACGAGAACAGTTCTGACAGTAAGGGATGAAGGCTAGGGGCG |
| p__Nanoarchaeota_g__ BMS3ABIN17_ASV_3757 | 0 | 0 | 0 | 0 | 0 | 0 | 1 | 0 | 1 | 0 | 0 | CTTTTGCAATGCACGAAAGTGTGACAAAGCGAACCAGAGTGCTTCTCTATTGAGAAGCTTTTGTAGAATGTAAAAAATTCTACGAATAAGGACTGGGTAAGACTAGTGCCAGCCGCCGCGGTAATACTAGCGGTCCAAGTCGCAACCATTGTTATTGGGTCTAAAACATCCGTAGCTTGTTTGATAAGTTTTCTGTAAAATCTTGTTTCTTAAGAACAAGGCGTGCAGGAAATACTGTCAAGCTAGAGACTGGAAGACGCAGAAAGTACGTCTGAGGTAGTGGTAAAATATTTTAATCTTGGATGGACTCACAACAGCGAAAGCATTCTGCGAGGACAGTTCTGACAGTGAGGGATGAAGGCTAGGGGCG |
| p__Nanoarchaeota_g__ BMS3ABIN17_ASV_3789 | 0 | 0 | 0 | 0 | 0 | 0 | 0 | 0 | 1 | 0 | 0 | CTTTTACAATGCTCGAAAGAGTGATAGAGCGAGCCGAAGTGTTCCTCTTAGAGGAACTTTTGTCTACTCTAAATAGGTAGACGAATAAGGACTGGGCAAGACCGGTGCCAGCCGCCGCGGTAACACCGGCGGTCCGAGTCGCATCCACATTTATTGGGTCTAAAACATCCGTAGCTTGTTCTGCAAGTCTCTTGTGAAATCTAGCGTCTCAAGCGTTGGAATTGCAAGAGATACTGTATTACTAGAGACCGGAAGACGTAAGGAGTACGGTTAAGGTAGTGGTAAAATATGTTAATCTTAACTGGACTAACAACGGCGAAGGCACCTTACGAGTACGGATCTGACAGTGAGGGATGAAGGCCAGGGGCG |
| p__Nanoarchaeota_g__ CG1-02-57- 44_ASV_3561 | 0 | 0 | 0 | 0 | 0 | 0 | 0 | 0 | 0 | 0 | 1 | CCTTTACAATGCGCGAAAGCGTGATAAGGGAACCCTAAGTGCTCAACAAACGTTGGGCTTTTGCCAAGAGTAAATATCTTGGAGAATAAGTGGTGGGTAAGACTGGTGCCAGCCGCCGCGGTAACACCAGCGCCACGAGTGGGAATCGTGTTTATTGGGCCTAAAGCGTCCGTAGCCGGTCAAGTAAATCTTTTGTGAAATCTTCAGGCTCAACCTGTTGGCGCGCAGAGGACACTGCTTGACTAGGAACCGGGAGGAGTCAGAGGTATGGGGTGGGGAGCGGTAAAATGCGATAATCCACCTTAGACCACCTGTGGCGAAGGCGTCTGACCAGAACGGATTCGACGGTGAGGGACGAAAGCCAGGGGAG |

| Phylum_Genus_ASV_number | Presence (1) or Absence (0) in each layer | | | | | | | | | | | |
|---|---|---|---|---|---|---|---|---|---|---|---|---|
| | L1 | L2 | L3 | L4 | L5 | L6 | L7 | L8 | L9 | L10 | L11 | Sequence |
| p__Nanoarchaeota_g__CG1-02-57-44_ASV_3721 | 0 | 0 | 0 | 0 | 1 | 0 | 0 | 1 | 0 | 0 | 0 | CCTTTACAATGCACGAAAGTGTGATAAGGGGATTCTGAGTGCTCATGCAAAGCATGGGCTTTTGCCAAGGGTAAATACCTTGGCGAATAAGTGGTGGGTAAGACTGGTGCCAGCCGCCGCGGTAACACCAGCGCCACAAGTGGGAACCAGATTTATTGGGCCTAAAGCGTCCGTAGCCGGTTTGGTAAATTTTTTGTGAAATTGTCGGGCTTAACCCGGCAACGTGCAGAAAAGACTGCCAGACTAGGGACCGGGAGGCGTGTGAGGTATTCCTTGGGGAGCGGTAAAATGTTATAATCCTTGGAAGACCACCTGTGGCGAAGGCGTCACACGAGAACGGATCCGACGGTGAGGGACGAAAGCCAGGGGAG |
| p__Halobacteriota_g__Haladaptatus_ASV_3656 | 0 | 0 | 1 | 0 | 0 | 0 | 0 | 0 | 0 | 0 | 0 | ACTTTACACTGCACGCCAGTGCGATAAGGGAACTCCAAGTGCGAGGGCATATAGTCCTCGCTTTTGTGTATCGTAAGGTGGTACACGAATAAGAGCTGGGCAAGACCGGTGCCAGCCGCCGCGGTAATACCGGCAGCTCAAGTGATGGCCAATATTATTGGGCCTAAAGCGTCCGTAGCCTGCCAGACAGGTCCGTCGGGAAATCTGCTCGCTCAACGAGCAGGCGTCCGGCGGAAACCAGCTGGCCTGGGGCCGGAAGACCCAAGGGGTACGTCTGGGGTAGGAGTGAAATTCTGTAATCCTAGACGGACCACCGATGGCGAAAGCACTCTGAGAGGACGGATTCGACAGTGAGGGACGAAAGCTAGGGTCT |
| p__Halobacteriota_g__Halalkalirubrum_ASV_3791 | 0 | 0 | 0 | 0 | 0 | 0 | 0 | 0 | 1 | 0 | 0 | CCTTTACACTGCACGCGAGTGCGATAGGGGGACCCCGAGTGCGAGGGCATACTAGCCCTCGCTTTTGTGAACCGTAAGGCGGTTCACGAACAAGAGCTGGGCAAGACCGGTGCCAGCCGCCGCGGTAATACCGGCAGCTCGAGTGATGACCGATTTTATTGGGCCTAAAGCGTCCGTAGCTGGCCACGAAAGTCCGTCGGGAAATCTCTCGGCTCAACCGAGAGGCGTCCGGCGGGACCTCCGTGGCTTGGAACCGGAAGGCTCAGAGGGTACGTCCAGGGTAGGAGTGAAATCCTGTAATCCTGGACGGACCACCGATGGCGAAAGCACTCTGAGAGGACGGATTCGACAGTGAGGGACGAAAGCTAGGGTCT |
| p__Halobacteriota_g__Halanaeroarchaeum_ASV_3743 | 0 | 0 | 0 | 0 | 0 | 1 | 0 | 0 | 0 | 0 | 0 | CCTTTACACTGCACGCCAGTGCGATAGGGGGACTCCGAGTGCGAGGGCATACAGTCCTCGCTTTTCTGTACCGTAAGGTGGTACAGGAACAAGTGCTGGGCAAGACCGGTGCCAGCCGCCGCGGTAATACCGGCAGCCCAAGTGATGGCCGATATTATTGGGCCTAAAGCGTCCGTAGCTGGCCGGACAAGTCCGTTGGGAAATCCGCGCGCTCAACGCGCGGGCGTCCAGCGGAAACTGTCCGGCTTGGGGCCGGAAGACCTGAGGGGTACGTCCGGGGTAGGAGTGAAATCCTGTAATCTCGGACGGACCGCCTGTGGCGAAAGCGCCTCAGGAAGACGGACCCGACAGTGAGGGACGAAAGCCAGGGTCT |
| p__Halobacteriota_g__Halobellus_3026_ASV_3692 | 0 | 0 | 0 | 0 | 1 | 0 | 0 | 0 | 0 | 0 | 0 | CCTTTACACTGCACGCACGTGCGATAAGGGGACTCCGAGTGCGAGGGCATATCGTCCTCGCTTTTCTGTACCGTAGGGCGGTACAGGAACAAGAGCTGGGCAAGACCGGTGCCAGCCGCCGCGGTAATACCGGCAGCTCAAGTGATGGCCGATCTTATTGGGCCTAAAGCGTCCGTAGCTGGCCTCGAAAGTCCGTCGGGAAATCCACACGCTCAACGTGTGGGCGTCCGGCGGAAACTCCGAGGCTTGGAACCGGAAGGCTCGAGAGGTACGTCCGGGGTAGGAGTGAAATCCTGTAATCCCGGACGGACCACCGATGGCGAAAGCATCTCGAGAAGACGGATTCGACAGTGAGGGACGAAAGCTAGGGTCT |
| p__Halobacteriota_g__Halobium_ASV_3626 | 0 | 0 | 1 | 0 | 0 | 0 | 0 | 0 | 0 | 0 | 0 | CCTTTACACTGCACGCAAGTGCGATAAGGGGACCCCGAGTGCGAGGGCATACTAGCCCTCGCTTTTCGTAACCGTAAGGAGGTTACAGAATAAGAGCTGGGCAAGACCGGTGCCAGCCGCCGCGGTAATACCGGCAGCTCGAGTGATGACCGATCTTATTGGGCCTAAAGCGTCCGTAGCTGGCCCCGTTGGTCCGTCGGGAAATCCACCTGCTCAACAGGTGGCCGTCCGGCGGATACCACGGGGCTTGGGACCGGAAGACCCGAGGGGTACGTCCGGGGTAGGAGTGAAATCCCGTAATCCTGGACGGACCACCGGTGGCGAAAGCGCTCTGGAAAGACGGATCCGACGGTGAGGGACGAAAGCTCGGGTCA |
| p__Halobacteriota_g__Halococcus_ASV_3741 | 0 | 0 | 0 | 0 | 0 | 1 | 0 | 0 | 0 | 0 | 0 | CCTTTACACTGCACGACAGTGTGATAAGGGGATCCCGAGTGCGAGGGCATACAGTCCTCGCTTTTCGTGACCGTAAGGTGGTCACAGAATAAGGGCTGGGCAAGACCGGTGCCAGCCGCCGCGGTAATACCGGCAGCCCGAGTGATAGCCACTATTATTGGGCCTAAAGCGTCCGTAGCCGGCCGAACAGGTCCGTCGGGAAATCCACCCGCTCAACGGGTGGGCGTCCGGCGGAAACCAGTCGGCTTGGGGCCGGAAGACCCGTGGGGTACGTCCGGGGTAGGAGTGAAATCCCGTAATCCCGGACGGACCGCCGGTGGCGAAAGCGCCACGGGAGGACGGACCCGACGGTGAGGGACGAAAGCTTGGGTCT |
| p__Halobacteriota_g__Halococcus_ASV_3806 | 0 | 0 | 0 | 0 | 0 | 0 | 0 | 0 | 1 | 0 | 0 | CCTTTACACTGCACGACAGTGCGATAAGGGGATCCCGAGTGCGAGGGCATACAGTCCTCGCTTTTCGTGACCGTAAGGTGGTCACAGAATAAGGGCTGGGCAAGACCGGTGCCAGCCGCCGCGGTAATACCGGCAGCCCGAGTGATAGCCACTATTATTGGGCCTAAAGCGTCCGTAGCCGGCCGAACAGGTCCGTCGGGAAATCCACCCGCTCAACGGGTGGGCGTCCGGCGGAAACCAGTCGGCTTGGGGCCGGAAGACCCGTGGGGTACGTCCGGGGTAGGAGTGAAATCCCGTAATCCCGGACGGACCGCCGGTGGCGAAAGCGCCACGGGAGGACGGACCCGACGGTGAGGGACGAAAGCTTGGGTCT |
| p__Halobacteriota_g__Halogeometricum_ASV_3722 | 0 | 0 | 0 | 0 | 0 | 1 | 0 | 0 | 0 | 0 | 0 | CCTTTACACTGCACGCAAGTGCGATAAGGGGACTCCGAGTGCGAGGGCATACAGTCCTCGCTTTTGTGTACCGTAGGGCGGTACACGAATAAGAGCTGGGCAAGACCGGTGCCAGCCGCCGCGGTAATACCGGCAGCTCAAGTGATGGCCAATCTTATTGGGCCTAAAGCGTCCGTAGCTGGTCGGGAAAGTTCGTCGGGAAATCCACTCGCTCAACGAGTGGGCGTCCGGCGAAAACTTCTCGACTTGGGACCGGAAGGCTCGAGGGGTACGTCCGGGGTAGGAGTGAAATCCTGTAATCCTGGACGGACCACCGATGGCGAAAGCACCTCGAGAAGACGGATCCGACAGTGAGGGACGAAAGCTAGGGTCT |
| p__Halobacteriota_g__Halohasta_ASV_3651 | 0 | 0 | 1 | 0 | 0 | 0 | 0 | 0 | 0 | 0 | 0 | CCTTTACACTGCACGAAAGTGCGATAGGGGAATCCCAAGTGCGAGGGCATATTAGCCCTCGCTTTTGTCGACTGTAAGGAGGTCGACGAATAAGAGCTGGGCAAGACCGGTGCCAGCCGCCGCGGTAATACCGGCAGCTCAAGTGATGTCCACTATTATTGGGCCTAAAGCGTCCGTAGCTGGCCGTACAAGTTCGTCGGGAAATCCACCTGCCCAACAGGTGGGCGTCCGGCGAAAACTGTGCGGCTTGGAACCGGAAGGCTCAGAGGGTACGTCGAGGGTAGGAGTGAAATCCTGTAATCCTCGACGGACCACCGATGGCGAAAGCACTCTGAGAAGACGGATTCGACAGTGAGGGACGAAAGCTAGGGTCT |
| p__Halobacteriota_g__Halolamina_ASV_3708 | 0 | 0 | 0 | 0 | 1 | 0 | 0 | 0 | 0 | 0 | 0 | CCTTTACACTGCACGCCAGTGCGATAAGGGAACCCCGAGTGCGAGGGCATATCGCCCTCGCTTTTCTGGACTGTAGGGAGGTCCAGGAACAAGAGCTGGGCAAGACCGGTGCCAGCCGCCGCGGTAATACCGGCAGCTCGAGTGATGACCGATTTTATTGGGCCTAAAGCGTCCGTAGCTGGCCGCGCAAGTCCGTCGGGAAATCCACCCGCTTAACGGGTGGGCGTCCGGCGGAAACTGCCCGGCTTGGGACAGGAAGGCTCGAGGGGTACGTTCGGGGTAGGAGTGAAATCCCGTAATCCCGGACGGACCACCGATGGCGAAAGCACTCTGAGAGGACGGATTCGACAGTGAGGGACGAAAGCTAGGGTCT |
| p__Halobacteriota_g__Halomicrobium_3849_ASV_3639 | 0 | 0 | 1 | 0 | 0 | 0 | 0 | 0 | 0 | 0 | 0 | CCTTTACACTGCACGACAGTGCGATAAGGGGACTCCGAGTGCGAGGGCATATAGTCCTCGCTTTTCTGTACCGTAAGGTGGTACAGGAACAAGTGCTGGGCAAGACCGGTGCCAGCCGCCGCGGTAATACCGGCAGCACGAGTGATGGCCGCTATTATTGGGCCTAAAGCGTCCGTAGCCGGCCAGACAAGTCCGTTGGGAAATCCACGCGCTCAACGCGTGGGCGTCCAGCGGAAACTGTCTGGCTTGGGGCCGGAAGACCTGAGGGGTACGTCCGGGGTAGGAGTGAAATCCTGTAATCCTGGACGGACCACCGGTGGCGAAAGCGCCTCAGGAGGACGGACCCGACGGTGAGGGACGAAAGCTAGGGTCT |
| p__Halobacteriota_g__Halomicrobium_3849_ASV_3723 | 0 | 0 | 0 | 0 | 0 | 1 | 0 | 0 | 0 | 0 | 0 | CCTTTACACTGCACGACAGTGCGATAGGGGGACTCCGAGTGCGAGGGCATATCGTCCTCGCTTTTCTCGACCGTAAGGTGGTCGAGGAACAAGTGCTGGGCAAGACCGGTGCCAGCCGCCGCGGTAATACCGGCAGCACGAGTGATGGCCAATATTATTGGGCCTAAAGCGTCCGTAGCCGGCCGGACAAGTCCGTTGGGAAATCGACGCGCTCAACGCGTCGGCGTCCAGCGGAAACTGTCCGGCTTGGGGCCGGAAGACCTGAGGGGTACGTCCGGGGTAGGAGTGAAATCCTGTAATCCTGGACGGACCACCGGTGGCGAAAGCGCCTCAGGAAGACGGACCCGACGGTGAGGGACGAAAGCTAGGGTCT |
| p__Halobacteriota_g__Halonotius_ASV_3548 | 0 | 0 | 0 | 0 | 0 | 0 | 0 | 0 | 0 | 0 | 1 | CCTTTACACTGCACGGAAGTGCGATAGGGGAATCCCAAGTGCGAGGGCATATTAGCCCTCGCTTTTGTTGACTCTAAGGTGGTCAACGAATAAGAGCTGGGCAAGACCGGTGCCAGCCGCCGCGGTAATACCGGCAGCTCGAGTGATGTCCCATATTATTGGGCCTAAAGCGTCCGTAGCTGGCCACGCAAGTCCATCGGGAAATCCACCCGCCCAACGGGTGGGCGTCCGGTGGAAACTGCGTGGCTTGGAACCGGAAGGCTCAGAGAGTACGTCGAGGGTAGGAGTGAAATCCCGTAATCCTCGACGGACTACCGATGGCGAAAGCACTCTGAGAAGACGGCTTCGACAGTGAGGGACGAAAGCTAGGGTCT |
| p__Halobacteriota_g__Halorubrum_ASV_3711 | 0 | 0 | 0 | 0 | 1 | 0 | 0 | 0 | 0 | 0 | 0 | CCTTTACACTGCACGCCAGTGCGATAAGGGGACTCCAAGTGCGAGGGCATATCGCCCTCGCTTTTCTGTACCGTAGGGAGGTACAGGAACAAGCGCTGGGCAAGACCGGTGCCAGCCGCCGCGGTAATACCGGCAGCGCAAGTGATGGCCGATCTTATTGGGCCTAAAGCGTCCGTAGCTGGCCAGGCAAGTCCATCGGGAAATCCACCCGCTCAACGGGTGGGCGCCCGGTGGAAACTGCCCGGCTCGGGACCGGAAGACCCAACGGGTACGTCCAGGGTAGGAGTGAAATCCCGTAATCCTGGACGGACCGCCGATGGCGAAAGCACGTTGGGAAGACGGATCCGACAGTGAGGGACGAAAGCTGGGGTCT |
| p__Halobacteriota_g__Halorubrum_ASV_3785 | 0 | 0 | 0 | 0 | 0 | 0 | 0 | 0 | 1 | 0 | 0 | CCTTTACACTGCACGACAGTGCGATAAGGGAATCCCAAGTGCGTAGGCATAGAGCCTACGCTTTTGTCCACCGTAGGGAGGTGGACGAATAAGGGCTGGGCAAGACCGGTGCCAGCCGCCGCGGTAATACCGGCAGCCCGAGTGATGGCCGATCTTATTGGGCCTAAAGCGTCCGTAGCTGGCCGCACAAGTCCATCGGAAAATCCACCCGCTCAACGGGTGGGCGTCCGGTGGAAACTGTGTGGCTTGGGACCGGAAGGCGCGACGGGTACGTCCGGGGTAGGAGTGAAATCCTGTAATCCTGGACGGACCACCGATGGCGAAAGCACGTCGCGAGAACGGATCCGACAGTGAGGGACGAAAGCTAGGGTCT |
| p__Halobacteriota_g__Halorussus_ASV_3486 | 1 | 0 | 0 | 0 | 0 | 0 | 0 | 0 | 0 | 0 | 0 | CCTTTACACTGCACGACAGTGCGATAAGGGGACTCCGAGTGCGAGGGCATATAGTCCTCGCTTTTGTACACCTTAAGGCGGTGTACGAATAAGGGCTGGGCAAGACCGGTGCCAGCCGCCGCGGTAATACCGGCAGCCCGAGTGATGGCCGCTATTATTGGGCCTAAAGCGTCCGTAGCCGGCCAGACAGGTCCGTCGGGAAATCTGCTCGCTCAACGAGCAGGCGTCCGGCGGAAACCAGCTGGCTTGGGGCCGGAAGACTCGAGGGGTACGTCCGGGGTAGGAGTGAAATCCCGTAATCCTGGACGGACCACCGGTGGCGAAAGCGCCTCGAGAAGACGGACCCGACGGTGAGGGACGAAAGCTAGGGTCA |
| p__Halobacteriota_g__Halorussus_ASV_3550 | 0 | 0 | 0 | 0 | 0 | 0 | 0 | 0 | 0 | 0 | 1 | TCTTTACACTGCACGACAGTGCGATAAGGGGACTCCGAGTGCGAGGGCATATAGTCCTCGCTTTTCTGTACCGTAAGGTGGTACAGGAATAAGGGCTGGGCAAGACCGGTGCCAGCCGCCGCGGTAATACCGGCAGCCCGAGTGATGGCCGCTATTATTGGGCCTAAAGCGTCCGTAGCCGGCCAGACAGGTCCGTCGGGAAATCCACGCGCTCAACGCGTGGGCGTCCGGCGGAAACCAGCTGGCTTGGGGCCGGAAGACTCGAGGGGTACGTCCGGGGTAGGAGTGAAATCCCGTAATCCTGGACGGACCACCGGTGGCGAAAGCGCCTCGAGAAGACGGACCCGACGGTGAGGGACGAAAGCTAGGGTCA |
| p__Halobacteriota_g__Halorussus_ASV_3729 | 0 | 0 | 0 | 0 | 0 | 1 | 0 | 0 | 0 | 0 | 0 | CCTTTACACTGCACGACAGTGCGATAAGGGGACTCCGAGTGCGAGGGCATATAGTCCTCGCTTTTCTGTACCGTAAGGTGGTACAGGAATAAGGACTGGGCAAGACCGGTGCCAGCCGCCGCGGTAATACCGGCAGTCCGAGTGATGGCCGCTATTATTGGGCCTAAAGCGTCCGTAGCCGGCCAGACAGGTCCGTCGGGAAATCTGCTCGCTCAACGAGCAGGCGTCCGGCGGAAACCAGCTGGCTTGGGGCCGGAAGACTCGAGGGGTACGTCCGGGGTAGGAGTGAAATCCTGTAATCCTGGACGGACCGCCGGTGGCGAAAGCGCCTCGAGGAGGACGGACCCGACGATGAGGGACGAAAGCTAGGGTCA |
| p__Halobacteriota_g__Halorussus_ASV_3773 | 0 | 0 | 0 | 0 | 0 | 0 | 0 | 1 | 0 | 0 | 0 | ACTTTACACTGCACGACAGTGCGATAAGGGAACTCCGAGTGCGAGGGCATATAGTCCTCGCTTTTGTACACCGTAAGATGGTGTACGAATAAGAGCTGGGCAAGACCGGTGCCAGCCGCCGCGGTAATACCGGCAGCTCGAGTGATGGCCGCTATTATTGGGCCTAAAGCGTCCGTAGCCGGCCAGACAGGTCTGTCGGGAAATCCATCCGCTTAACGGATGGTCGTCCGGCGGAAACCAGCTGGCTTGGGACCGGAAGACCTGGGGGGTACGTCTGGGGTAGGAGTGAAATCCTGTAATCCTGGACGGACCGCCGGTGGCGAAAGCGCCTCAGGAAGACGGACCCGACGGTGAGGGACGAAAGCTAGGGTCA |
| p__Halobacteriota_g__Halorussus_ASV_3786 | 0 | 0 | 0 | 0 | 0 | 0 | 0 | 0 | 1 | 0 | 0 | CCTTTACACTGCACGCAAGTGCGATAAGGGGATCCCGAGTGCGAGGGCATATAGTCCTCGCTTTTCTGGACTGTAGGGAGGTCCAGGAACAAGAGCTGGGCAAGACCGGTGCCAGCCGCCGCGGTAATACCGGCAGCTCAAGTGATGGCCGATATTATTGGGCCTAAAGCGTCCGTAGCCGGCCGGATAAGTCCGTCGGGAAATCCACCCGCTCAATGGGTGGGCGTCCGGCGGAAACTGTCCGGCTTGGGGCCGGAGGGCCTGAGGGGTACGTCCGGGGTAGGAGTGAAATCCTGTAATCCCGGTCGGACCACCGGTGGCGAAAGCGCCTCAGGAGGACGGACCCGACGGTGAGGGACGAAAGCTAGGGTCA |
| p__Halobacteriota_g__Methanohalophilus_ASV_3475 | 1 | 0 | 0 | 0 | 0 | 0 | 0 | 0 | 0 | 0 | 0 | CCTTTACACTGCGGGAAACCGCGATAAGGGGACACCGAGTGCCAGCATACAATGCTGGCTGTCCACATGTGTAAATAGCATGTGATAGCAAGGGCCGGGCAAGACCGGTGCCAGCCGCCGCGGTAACACCGGCGGCCCGAGTGGTAATCACTATTATTGGGTCTAAAGGGTCCGTAGCCGGTTTAATCAGTCTTCCGGGAAATCTGACAGCTCAACTGTTAGGCTTCCGGGGGATACTGTTAGACTTGGGACCGGGAGAGGTAAGAGGTACTACAGGGGTAGGAGTGAAATCTTGTAATCCCTGTGGGACCACCAGTGGCGAAAGCACCTCGGAAGGACGGATCCGACAGTGAGGGACGAAAGCTAGGGTCT |
| p__Halobacteriota_g__Natrinema_ASV_3755 | 0 | 0 | 0 | 0 | 0 | 0 | 1 | 0 | 0 | 0 | 0 | CCTTTACACTGCACGACAGTGCGATAAGGGGACTCCAAGTGCGAAGGCATATCGTCCTCGCTTTTCACTACCGTAAGGTGGTAGTGGAATAAGAGCTGGGCAAGACCGGTGCCAGCCGCCGCGGTAATACCGGCAGCTCGAGTGATGACCGCTTTTATTGGGCCTAAAGCGTCCGTAGCCGGCCGGGTAAGTCCATCGGGAAATCCGCGCGCTCAACGCGCGGGCGTCCGGTGGAAACTGCCTGGCTTGGGACCGGAAGATCCAGAGGGTACGTCTGGGGTAGGAGTGAAATCCCGTAATCCTGGACGGACCACCGGTGGCGAAAGCGCTCTGGAAAGACGGATCCGACGGTGAGGGACGAAAGCTCGGGTCA |
| p__Halobacteriota_g__Natronobiforma_ASV_3644 | 0 | 0 | 1 | 0 | 0 | 0 | 0 | 0 | 0 | 0 | 0 | CCTTTACACTGCACGACAGTGCGATAAGGGGACTCCAAGTGCGAGGGCATATCGTCCTCGCTTTTTCCTACCGTAAGGTGGTAGGCGAATAAGGGCTGGGCAAGACCGGTGCCAGCCGCCGCGGTAATACCGGCAGCCCAAGTGATGACCGCTATTATTGGGCCTAAAGCGTCCGTAGCCGGCCGAACAAGTCCGTCGGGAAATCTGCTCGCCTAACGAGCAGGCGTCCGGCGGAAACTGTTCGGCTTGGGACCGGAAGATCCAGAGGGTACGTCTGGGGTAGGAGTGAAATCCTGTAATCCTGGACGGACCACCGGTGGCGAAAGCGCTCTGGAAGGACGGATCCGACGGTGAGGGACGAAAGCTAGGGTCA |

| | Presence (1) or Absence (0) in each layer | | | | | | | | | | | |
|---|---|---|---|---|---|---|---|---|---|---|---|---|
| **Phylum_Genus_ASV_number** | **L1** | **L2** | **L3** | **L4** | **L5** | **L6** | **L7** | **L8** | **L9** | **L10** | **L11** | **Sequence** |
| p__Thermoproteota_g__PALSA-986_ASV_3802 | 0 | 0 | 0 | 0 | 0 | 0 | 0 | 0 | 1 | 0 | 0 | CCTTTGCAATACACGAAAGTGTGACAGGGCTATCCCGAGTGCCATCCGCTGAGGAAGGCTTTTACCCAGTCTAGAAAGCTGGGGGAATAAGGAGAGGGCAAGTCTGGTGTCAGCCGCCGCGGTAATACCAGCTCTTCGAGTGGTGTGGATGTTTATTGGGCCTAAAGCATCCGTAGCTGGCTAGGTAAGTCCTTTGTTAAATCCACCGAATTAATCGTTGGATCGCGGGGGATACTGCTTGGCTAGGGGACGAGAGAGGCAGACGGTATTTTCGGGGTAGGGGTGAAATCCTATAATCCCGGGAAGACCACCAGTGGCGAAGGCTGTCTGCTAGAACGCGCCCGACGGTGAGGGATGAAAGCTGGGGGAG |
| p__Halobacteriota_g__Saliphagus_ASV_3702 | 0 | 0 | 0 | 0 | 1 | 0 | 0 | 0 | 0 | 0 | 0 | CCTTTACACTGCACGACAGTGCGATAGGGGGACTCCAAGTGCGAGGGCATATCGTCCTCGCTTTTGTGTACCGTAAGGTGGTACACGAATAAGGGCTGGGCAAGACCGGTGCCAGCCGCCGCGGTAATACCGGCAGCCCAAGTGATGACCGCTATTATTGGGCCTAAAGCGTCCGTAGCTGGCCGATCAAGTCCGTCGGGAAATCCGCACGCCTAACGTGCGGGCGTCCGGTGGAAACTGTTCGGCTTGGGACCGGAAGACTCGAGGGGTACGTCTGGGGTAGGAGTGAAATCCTGTAATCCTGGACGGACCACCGGTGGCGAAAGCGCCTCGAGAGGACGGATCCGACGGTGAGGGACGAAAGCTAGGGTCA |
| p__Thermoplasmatota_g__SM1-50_ASV_3765 | 0 | 0 | 0 | 0 | 0 | 0 | 0 | 1 | 0 | 0 | 0 | CCTTCACACTGTGCGAAAGCACGATGAGGGGATCCCAAGTGCTGGCTCGTAAGAGTCAGCTGTTTCTTTGTCTAAAAAACAGAGAGAGTAAGAGCTGGGTAAGACGGGTGCCAGCCGCCGCGGTAATACCTGCAGCTCAAGTGGTGGCCGCTATTATTGGGCCTAAAACGTTCGTAGCCTGTTTAGTAAATGCTTGGGTAAATTGTGTCGCTTAACGATACGAATACCGGGTAGACTGCTAAACTTGGGACCGGGAGAGGCTAGAGGTACTCCTGGGGTAGAGGTGAAATTCTGTAATCCTGGGGGACCACCAGTGGCGAAGGCGTCTAGCCAGAACGGGTCTGACGGTGAGGGACGAAGCCCTGGGGCG |
| p__Nanoarchaeota_g__SM23-78_ASV_3783 | 0 | 0 | 0 | 0 | 0 | 0 | 0 | 0 | 1 | 0 | 0 | CCTTTACAATGCACGAAAGTGTGATAAGAGGATCCCAAGTGCTTAACATTGTTAGGCTTTTGCCAAGTTTAGAACGCTTGGCGAATAAGGGGTGGGTAAGACTGGTGCCAGCCGCCGCGGTAACACCAGCGCCCCGAGTGGCAATCGCGATTATTGGGTCTAAAGCGTCCGTAGCAGGTTCTGTAAATCCCTTGTGAAATCGGCAAGCTTAACTTGTCGGCGTGCAGAGGACACTGCAGGACTCGAGACCGGGAGAGGTCAAAAGTATTTCCTGGGTACCGGTAAAATGGGTTAATCCAGGGAGGACTACCTATGGCGAAGGCATTTGACCAGAACGGATCTGACTGTGAGGGACGAAAGCTAGGGGAG |
| p__Thermoplasmatota_g__UBA147_ASV_3797 | 0 | 0 | 0 | 0 | 0 | 0 | 0 | 0 | 1 | 0 | 0 | ACTCTACAATGTGGGAAACCACGATAGGGGAACTCCAAGTGCATGCACTTAGTGTATGCTGTTCTACTGCCTAAAAAGCAGTTGAAGTAAGGGCTGGGTAAGACGGGTGCCAGCCGCCGCGGTAATACCTGCAGCCCCAGTGGTGTTCAGCATTATTGAGCCTAAAGCGTCCGTAGCCGGTCCTGTAAATCTCTGGGTAAATCGAGCAGCTCAACTGTTCGAATACCGGAGAGACTGCAGGACTTGGGACTGGGAGAGGTGGAAGGTACTCCAGGGGTAGGGGTAAAATCCTGTAATCCTTGGGGGACCACCAGTGGCGAAGGCGTTTCACTAGAACAGATCCGACGGTGAGGGACGAAGCCCTGGGGCG |
| p__Halobacteriota_f__Haloarculaceae_3992_ASV_3497 | 0 | 0 | 0 | 0 | 0 | 0 | 0 | 0 | 0 | 1 | 0 | CCTTTACACTGCACGACAGTGCGATAGGGGGACTCCAAGTGCGAGGGCATATAGTCCTCGCTTTTGTACACCGTAAGGTGGTGTACGAATAAGAACTGGGCAAGACCGGTGCCAGCCGCCGCGGTAATACCGGCAGTTCGAGTGATGGCCGATTTTATTGGGCCTAAAGCGTTCGTAGCTGGCCAGTCAAGTCTGTCGGGAAACCTGCTCGCTCAACGGGCAGCCGTCCGGCGGAAACTGATTGGCTTGGGACCGGAAGACCTGAAGAGTACGTCCGGGGTAGGAGTGAAATCCTGTAATCCTGGACGGACTACCGATGGCGAAAGCACTTCAGGAGCACGGATCCGACAGTGAGGAACGAAAGCTGGGGTCT |
| p__Halobacteriota_f__Haloarculaceae_3992_ASV_3586 | 0 | 1 | 0 | 0 | 0 | 0 | 0 | 0 | 0 | 0 | 0 | CCTTTACACTGCACGACAGTGCGATAGGGGGACTCCGAGTGCGAGGGCATAGAGCCCTCGCTTTTGTACACCGTAAGGTGGTGTACGAATAAGAACTGGGCAAGACCGGTGCCAGCCGCCGCGGTAATACCGGCAGTTCGAGTGATGGCCGATATTATTGGGCCTAAAGCGTTCGTAGCTGGCCGACCAAGTCTGTCGGGAAATCTGCTCGCCTAACGAGCAGGCGTCCGGCGGAAACTGGTCGGCTTGGGACCGGAAGACTCGAGGGGTACGTCCGGGGTAGGAGTGAAATCCTGTAATCCTGGACGGACCACCGATGGCGAAAGCACCTCGAGAGCACGGATCCGACAGTGAGGAACGAAAGCTGGGGTCT |
| p__Halobacteriota_f__Haloarculaceae_3992_ASV_3635 | 0 | 0 | 1 | 0 | 0 | 0 | 0 | 0 | 0 | 0 | 0 | CCTTTACACTGCACGACAGTGCGATAGGGGGACTCCGAGTGCGAGGGCATATAGTCCTCGCTTTTCTGAACCGTAAGGTGGTTCAGGAACAAGGACTGGGCAAGACCGGTGCCAGCCGCCGCGGTAATACCGGCAGTCCGAGTGATGGCCGATTTTATTGGGCCTAAAGCGTCCGTAGCCTGCCCTGTAAGTCCATTGGGAAATCGACCAGCTCAACTGGTCGGCGTCCAGTGGAAACTACAGGGCTTGGGGCCGAGAGACTCGACGGGTACGTCTGGGGTAGGAGTGAAATCCTGTAATCCTGGACGGACCACCGGTGGCGAAAGCGCCTCGGGAGGACGGATCCGACGGTGAGGGACGAAAGCTAGGGTCA |
| p__Halobacteriota_f__Halobacteriaceae_4595_ASV_3746 | 0 | 0 | 0 | 0 | 0 | 0 | 1 | 0 | 0 | 0 | 0 | CCTTTACACTGTACGAAAGTGCGATAAGGGGACCCCGAGTGCGAGGGCATACAGTCCTCGCTTTTGTGTACCGTAAGGAGGTACACGAATAAGGGCTGGGCAAGACCGGTGCCAGCCGCCGCGGTAATACCGGCAGCCCTAGTGATGGCCGCTATTATTGGGCCTAAAGCGTCCGTAGCCTGCCGGACAAGTCCGTTGGGAAATCCGTTCGCTTAACGAACGGGCGTCCAGCGGAAACTGTCCGGCTTGGGACCGGAAGACCCGAGGGGTACGTCCGGGGTAGGAGTGAAATCCTGTAATCCTGGACGGACCGCCGGTGGCGAAAGCGCCTCGGGAAGACGGATCCGACGGTGAGGGACGAAAGCCAGGGTCT |
| p__Halobacteriota_f__Haloferacaceae_ASV_3646 | 0 | 0 | 1 | 0 | 0 | 0 | 0 | 0 | 0 | 0 | 0 | CCTTTACACTGCACGACAGTGCGATAGGGGGACCCCGAGTGCGAGGGCATACAGTCCTCGCTTTTGCACACCGTAAGGAGGTGTGCGAATAAGTGCTGGGCAAGACCGGTGCCAGCCGCCGCGGTAATACCGGCAGCACAAGTGATGGCCGCTATTATTGGGCCTAAAGCGTCCGTAGCCGGCCGAACAAGTCCGTCGGGAAATCCATCCGCTCAACGGATGGGCGTCCGGCGGAAACTGTTCGGCTTGGGACAGGAAGACTCGAGGGGTACGTCCGGGGTAGGAGTGAAATCCTGTAATCCTGGACGGACCGCCGATGGCGAAAGCACCTCGAGAAGACTGATCCGACGGTGAGGGACGAAAGCTGGGGTCT |
| p__Halobacteriota_f__Haloferacaceae_ASV_3707 | 0 | 0 | 0 | 0 | 1 | 0 | 0 | 0 | 0 | 0 | 0 | CCTTTACACTGCACGCAAGTGCGATAGGGGAATCCCAAGTGCGAGGGCATACTAGCCCTCGCTTTTGTGTACTGTAAGGAGGTACACGAATAAGAGCTGGGCAAGACCGGTGCCAGCCGCCGCGGTAATACCGGCAGCTCGAGTGATGTCCACTATTATTGGGCCTAAAGCGTCCGTAGCTGGCCGCGCAAGTCCGTCGGGAAATCCACCTGCCCAACAGGTGGGCGTCCGGCGGAAACTGCGTGGCTTGGAACCGGAAGGCTCAGAGGGTACGTCCGGGGTAGGAGTGAAATCCCGTAATCCTGGACGGACCACTGGTGGCGAAAGCGCCTCGAGAGGACGGACCCGACAGTGAGGGACGAAAGCCAGGGTCT |
| p__Halobacteriota_f__Natrialbaceae_ASV_3629 | 0 | 0 | 1 | 0 | 0 | 0 | 0 | 0 | 0 | 0 | 0 | CCTTTACACTGCACGCCAGTGCGATAAGGGGACTCCAAGTGCGAGGGCATACAGTCCTCGCTTTTCACGACCGTAAGGTGGTCGTGGAATAAGAGCTGGGCAAGACCGGTGCCAGCCGCCGCGGTAATACCGGCAGCTCGAGTGATGACCGCTATTATTGGGCCTAAAGCGTCCGTAGCCGGCCGGGTAAGTCCGTCGGGAAATCCGCGCGCTCAACGCGCGGGCGTCCGGTGGAAACTGCCTGGCTTGGGACCGGAAGATCCAGAGGGTACGTCCGGGGTAGGAGTGAAATCCTGTAATCCTGGACGGACCACCGGTGGCGAAAGCGCTCTGGAAAGACGGATCCGACGGTGAGGGACGAAAGCTAGGGTCA |
| p__Halobacteriota_o__Halanaerobiales_ASV_3563 | 0 | 0 | 0 | 0 | 0 | 0 | 0 | 0 | 0 | 0 | 1 | CCTTTACACTGCACGACAGTGCGATAAGGGGACTCCGAGTGCGAGGGCATATAGTCCTCGCTTTTCTGTACCGTAGGGAGGTACAGGAACAAGTGCTGGGCAAGACCGGTGCCAGCCGCCGCGGTAATACCGGCAGCACGAGTGATGACCGATATTATTGGGCCTAAAGCGTCCGTAGCTGGCCAGACAAGTCCGTTGGGAAATCCACGCGCTCAACGCGTGGGCGTCCGGCGGAAACTGTTTGGCTTGGGGCCAGAAGACCTGAGGGGTACGTCCGGGGTAGGAGTGAAATCCCGTAATCCTGGACGGACCACCGATGGGGAAACCACCTCAGGAGGATGGACCCGACAGTGAGGGACGAAAGCTAGGGTCT |
| p__Halobacteriota_o__Halanaerobiales_ASV_3592 | 0 | 1 | 0 | 0 | 0 | 0 | 0 | 0 | 0 | 0 | 0 | CCTTTACACTGCACGCCAGTGCGATAGGGGGACCCCGAGTGCGAGGGCATACAGTCCTCGCTTTTGTGTACCGTAAGGAGGTACACGAATAAGTGCTGGGCAAGACCGGTGCCAGCCGCCGCGGTAATACCGGCAGCACAAGTGATGGCCGCTATTATTGGGCCTAAAGCGTCCGTAGCCGGCCGAACAAGTCCGTCGGGAAATCCATCCGCTCAACGGATGGGCGTCCGGCGGAAACTGTTCGGCTTGGGACAGGAAGACTCGAGGGGTACGTCCGGGGTAGGAGTGAAATCCCGTAATCCTGGACGGACCGCCGATGGCGAAAGCACCTCGAGAAGACTGATCCGACGGTGAGGGACGAAAGCTGGGGTCT |
| p__Halobacteriota_o__Halanaerobiales_ASV_3599 | 0 | 1 | 0 | 0 | 0 | 0 | 0 | 0 | 0 | 0 | 0 | CCTTTACACTGCACGAAAGTGCGATAAGGGGATCCCAAGTGCGAGGGCATACAGTCCTCGCTTTTCTGTACCGTAGGGAGGTACAAGAATAAGGGCTGGGCAAGACCGGTGCCAGCCGCCGCGGTAATACCGGCAGCCCAAGTGATGGCCGCTATTATTGGGCCTAAAGCGTCCGTAGCCGGCCGGATAAGTCCGTCGGGAAATCCACCCGCTTAACGGGTGGACGTCCGGCGGAAACTGTCCGGCTTGGGGTCGGAGGGCCTGAGGGGTACGTCCGGGGTAGGAGTGAAATCCCGTAATCCCGGACGGACCACCGATGGCGAAAGCACTCTGAGAAGACGGATTCGACAGTGAGGGACGAAAGCTAGGGTCT |
| p__Halobacteriota_o__Halanaerobiales_ASV_3617 | 0 | 1 | 0 | 0 | 0 | 0 | 0 | 0 | 0 | 0 | 0 | CCTTTACACTGCACGACAGTGCGATAAGGGGACTCCAAGTGCGAGGGCATATCGCCCTCGCTTTTGTACACCGTAAGGTGGTGCACGAATAAGGGCTGGGCAAGACCGGTGCCAGCCGCCGCGGTAATACCGGCAGCCCGAGTGATGGCCGCTATTATTGGGCCTAAAGCGTCCGTAGCCGGCCGGTCAAGTCCGTCGGGAAATCCACCCGCTTAACGGGTGGGCGTCCGGCGGAAACTGATCGGCTTGGGACCGGAAGATCCGAGGGGTACGTCCGGGGTAGGAGTGAAATCCTGTAATCCTGGACGGACCGCCGGTGGCGAAAGCGCCTCGGAAGGACGGATCCGACGGTGAGGGACGAAAGCTTGGATCT |
| p__Halobacteriota_o__Halanaerobiales_ASV_3627 | 0 | 0 | 1 | 0 | 0 | 0 | 0 | 0 | 0 | 0 | 0 | CCTTTACACTGCACGACAGTGCGATAGGGGGATCCCGAGTGCGAGGGCATACAGTCCTCGCTTTTCTGCACCGTAAGGTGGTGCAGGAATAAGTGCTGGGCAAGACCGGTGCCAGCCGCCGCGGTAATACCGGCAGCACAAGTGATGGCCGCTTTTATTGGGCCTAAAGCGTCCGTAGCCGGCCGAACAAGTCCGTCGGGAAATCCATCCGCCTAACGGATGGGCGTCCGGCGGAAACTGTTCGGCTTGGGACAGGAAGACTCGAGGGGTACGTCCGGGGTAGGAGTGAAATCCCGTAATCCTGGACGGACCGCCGATGGCGAAAGCACCTCGAGAAGACTGATCCGACGGTGAGGGACGAAAGCTGGGGTCT |
| p__Halobacteriota_o__Halanaerobiales_ASV_3631 | 0 | 0 | 1 | 0 | 0 | 0 | 0 | 0 | 0 | 0 | 0 | CCTTTACACTGCACGACAGTGCGATAGGGGGACTCCGAGTGCGAGGGCATACAGTCCTCGCTTTTGTCGACCGTAAGGAGGTCGACGAATAAGGGCTGGGCAAGACCGGTGCCAGCCGCCGCGGTAATACCGGCAGCCCAAGTAATGGCCGATCTTATTGGGCCTAAAGCGTCCGTAGCCGGCCCGACAAGTCCGTTGGGAAATCCACGCGCTCAACGCGTGGGCGTCCAGCGGAAACTGTCGGGCTTGGGACCGGAAGACCCAGAGGGTACGTCCGGGGTAGGAGTGAAATCCCGTAATCCTGGACGGACCACCGGTGGCGAAAGCGCTCTGGGAAGACGGATCCGACGGTGAGGGACGAAAGCTAGGGTCT |
| p__Halobacteriota_o__Halanaerobiales_ASV_3751 | 0 | 0 | 0 | 0 | 0 | 0 | 1 | 0 | 0 | 0 | 0 | CCTTTACACTGCACGACAGTGCGATAAGGGGACCCCGAATGCGAGGGCATACAGTCCTCGCTTTTGTGCACTGTAGGGAGGTGCACGAATAAGAGCTGGGCAAGACCGGTGCCAGCCGCCGCGGTAATACCGGCAGCTCGAGTGATGGCCGCTGTTATTGGGCCTAAAGCGTCCGTAGCTGGCCGCACAAGTCCGTCGGGAAATCCACCCGCCTAACGGGTGGGCGTCCGGCGGAAACTGTGCGGCTTGGGACCGGAAGACCCGAGGGGTACGTCCGGGGTAGGAGTGAAATCCCGTAATCCTAGACGGACCGCCGGTGGCGAAAGCGCCTCGGGAAGACGGATCCGACAGTGAGGGACGAAAGCTAGGGTCT |
| p__Halobacteriota_o__Halanaerobiales_ASV_3792 | 0 | 0 | 0 | 0 | 0 | 0 | 0 | 0 | 1 | 0 | 0 | CCTTTACACTGCACGCAAGTGCGATAAGGGGACCCCGAGTGCGAGGGCATACCGTCCTCGCTTTTGTCGACCGTAGGGAGGTTGACGAATAAGGGCTGGGCAAGACCGGTGCCAGCCGCCGCGGTAATACCGGCAGCCCAAGTGATGGCCGCTCTTATTGGGCCTAAAGCGTCCGTAGCTGGCCGCATTGGTCCGTCGGGAAATCCACTCGCTCAACGAGTGGGCGTCCGGCGGATACCATGCGGCTTGGGACCGGAAGACCCAGGGGGTACGTCCGGGGTAGGAGTGAAATCCTGTAATCCCGGACGGACCGCCGATGGCGAAAGCACTCTGGGAAGACGGATCCGACAGTGAGGGACGAAAGCTGGGGTCT |
| p__Nanoarchaeota_o__Woesearchaeales_581_ASV_3590 | 0 | 1 | 0 | 0 | 0 | 0 | 0 | 0 | 0 | 0 | 0 | CCTTTACAATGCACGCAAGTGTGATAAGGGGATCCTCAGTGCTCATGCATTGCATGGGCTTTTGCCAAACGTAAATAGTTTGGCGAATAAGTGGTGGGTAAGACTGGTGCCAGCCGCCGCGGTAACACCAGCGCCACAAGTGGCAACCACGTTTATTGGGCCTAAAGCGTCCGTAGCCGGTGAGGTAAATCTCTTGTGAAATCGTCATGCCTAACGTGACGGCGCGCTCGAGACACTGCCTTGCTCGAGACCGGGAGGAGTCAGAGGTATTCCAGGGGGAGCGGTAAAATGTGATAATCCCTGGAGGATCACCTATGGCGAAGGCATCTGACCAGAACGGATCTGACGGTGAGGGACGAAGGCTAGGGGAG |
| p__Nanoarchaeota_o__Woesearchaeales_581_ASV_3670 | 0 | 0 | 0 | 1 | 0 | 0 | 0 | 0 | 0 | 0 | 0 | CCTTTACAATGCACGCAAGTGTGATAAGGGGATCCTAAGTGCTTATACAATGTATAGGCTTTTGTCAAGAGTAAATATCTTGGCGAATAAGTGGTGGGTAAGACTGGTGCCAGCCGCCGCGGTAACACCAGCGCCACGAGTGGCAACCGCGTTTATTGGGCCTAAAGCGTCCGTAGCCGGTTCGGCAAATCTCTTGTGAAATCGTCAGGCTTAACTTGACGGCGCGCTTGAGAGACTACCGAACTTGAGACCGGGAGGAGTCAGAGGTATTCCAGGGGGAGCGGTAAAATGTGATAATCCCTGGAGGATCACCTATGGCGAAGGCATCTGACCAGAACGGATCTGACGGTGAGGGACGAAGGCTAGGGGAG |

| Phylum_Genus_ASV_ number | Presence (1) or Absence (0) in each layer | | | | | | | | | | | |
|---|---|---|---|---|---|---|---|---|---|---|---|---|
| | L1 | L2 | L3 | L4 | L5 | L6 | L7 | L8 | L9 | L10 | L11 | Sequence |
| p__Thermoplasmatota_ p__Thermoplasmatota_ ASV_3813 | 0 | 0 | 0 | 0 | 0 | 0 | 0 | 0 | 1 | 0 | 0 | ACTGCGCAATGCGAGCAATCGCGACGCGGGAACGCCAAGTGCCGGCACAGAGTGTCGGCTGTTTCCAAGCCTAAAAAGCTCGGAGAGTAAGGGCTGGGTAAGACGGGTGCCAGCCGCCGCGGTAATACCTGCAGCCCAAGTGGTGGTCGCTAT<br>TATTGGGCCTAAAGCGTCCGTAGCCGGTTTGGTAAATGCCTGTGTAAATCGAACGGCTTAACCGTTCGACTTGCGGGTAGACTGCCAGACTAGGGACCGGGAGAGGCACGAAGTACTCCGGGGGTAGAGGTGAAATTCTGTAATCCTCGGGGG<br>ACTACCGGTGGCGAAGGCGTCGTGCTAGAACGGCTCCGACGGTGAGGGACGAAGCCCTGGGGCG |
| d__Archaea_d__Archae a_ASV_3483 | 1 | 0 | 0 | 0 | 0 | 0 | 0 | 0 | 0 | 0 | 0 | ATTTCGCAATGCGTGAAAGCGTGACGGAGCGACTCCTAGTGTCGGTGGTCTAGCGACCGTCGACTTTTTCGGAACGTAAAAAGTTCTGAGAATAAGGGCTGGGTAAGACTGGTGCCAGCCGCCGCGGTAACACCAGCAGCTCGAGTGGTGGCC<br>ACTTTTATTGGGCCTAAAGCGTCCGTAGCCGGACTAGCAAGTCCTTTGTGAAATCGGGCGGCTTAACCGTTCGGCTTGCAAGGGAAACTGTTAGTCTTGGGGCCGACAGGCGCCGAGGGTACTCCGGGGGTAGCGGTGAAATGCTATAATCCC<br>TGGGGGACCGCCGGTGGCGAAAGCGCTCAGCGAGGTCGGACCCGACGGTGAGGGACGAAAGCCAGGGGAG |
| d__Archaea_d__Archae a_ASV_3606 | 0 | 1 | 0 | 0 | 0 | 0 | 0 | 0 | 0 | 0 | 0 | ACTGCACAATGCGGGCAACCGCGATGCGGGGACTCCAAGTGCCGGCACAGAGTGTCGGCTGTTTCCAAGCCTAAAAAGCTCGGAGAGTAAGGGCTGGGTAAGACGGGTGCCAGCCGCCGCGGTAATACCTGCAGCCCAAGTGGTGGCCGCTAT<br>TATTGGGCCTAAAGCGTCCGTAGCCGGACCAGTAAATGCCTGTGTAAATTGTACCGCTTAACGGTGCGAACCGCGGGTAGACTGCTGGTCTTGGGACCGGGAGAGGCGCGAAGTACTCCGGGGGTAGAGGTGAAATTCTGTAATCCTCGGGGG<br>ACTACCGGTGGCGAAGGCGTCGCGCTAGAACGGCTCCGACGGTGAGGGACGAAGCCCTGGGGCG |
| d__Archaea_d__Archae a_ASV_3734 | 0 | 0 | 0 | 0 | 0 | 1 | 0 | 0 | 0 | 0 | 0 | ACTGCACAATGCGGGAAACCGCGATGCGGGGACTCCAAGTGCCAGCACAGAGTGTTGGCTGTTTCTGAGTCTAAAAAACTCAGAGAGTAAGGGCTGGGTAAGACGGGTGCCAGCCGCCGCGGTAATACCTGCAGCCCAAGTGGTGGTCGCTAT<br>TATTGGGCCTAAAGCGTCCGTAGCCGGATCAGTAAATGCCTGTGTAAATTGTACCGCTTAACGGTGCGAACTGCGGGTAGACTGCTGGTCTTGGGACCGGGAGAGGTGTGGAGTACTCCGGGGGTAGAGGTGAAATTCTGTAATCCTCGGGGG<br>ACTACCGGTGGCGAAGGCGCCACACTAGAACGGCTCCGACGGTGAGGGACGAAGCCCTGGGGCG |
| p__Nanoarchaeota_c__ Nanoarchaeia_ASV_41 17 | 0 | 0 | 0 | 1 | 0 | 0 | 0 | 0 | 0 | 0 | 0 | ACTTTACAATGCACGAAAGTGTGATAGGGGGATCCTAAGTGATATAGGTTATGCTTATATCTGTGGAGGAGTGTAAATTACTCTTCAAGAAAGGGGTGGGCAATATCGGTGCCAGCAGCCGCGGTAATCCCGGAGCCCCAAGTGGTGACCATG<br>TTTATTGGGCCTAAAGCGCCCGTAGCCGGATTAGTAAGTTCTCGGTGAAATTGTTTGGCTTAACTAAACGAATTGCCTTGAATACTGCTAATCTAGGGATTGGGAGAGGTCAAGAGTATGCCTGGGGGAGGGGTAAAATCCTTTAATCCTAGG<br>TAGACTACCAATGGCAAAGGCACTTGACTAGAACAAGTCCGACGGTGAGGGACGAAAGTTAGGGTAG |
| p__Nanoarchaeota_c__ Nanoarchaeia_ASV_41 23 | 0 | 0 | 0 | 1 | 0 | 0 | 0 | 0 | 0 | 0 | 0 | ACTTTACAATGCGCGAAAGCGTGATAAGGGGATCCCAAGTGTCCCAGTTTTACTGGGACTGTGGCGAAGTGTAAAGCGCTTCGCAAGAAAGGGGTGGGTAATACCGGTGCCAGCAGCCGCGGTAATCCCGGAGCCTCAAGTGGTGACCATGTT<br>TATTGGGCCTAAAGCGCTCGTAGCCGGATTAGTAAGTTCTCGGTGAAATTGTTTAGCTTAACTAAACGAATTGCCTTGAATACTGCTAGTCTAGGGATAGGGAGAGGTCAGGAGTATGCTTGGGGGAGGGGTAAAATCCTTTAATCCTAAGTA<br>GACTATCAATGGCGAAGGCACCTGACTAGAACTAGTCCGACGGTGAGGGACGAAAGTTAGGGTAG |
| p__Iainarchaeota_o__Iai narchaeales_ASV_4313 | 0 | 0 | 0 | 0 | 0 | 0 | 0 | 0 | 1 | 0 | 0 | CCTTTACAATGCACGAAAGTGTGATAGGGGGATTTCGAGTGCGATCTTTGATCGCTTTTGCCAAACCTAAATAGTTTGGCGAATAAGTGGGGGGCAAGATACGTGCCAGCCGCCGCGGTAACACGTACTCCGCAAGTGGTATCCACTAATATT<br>GGGCCTAAAACGTCTGTAGCCGGCTCAACAAGTCCTTGGTGAAATCATTGCGCTCAACGTAATGAATTGCCAAGGAAACTGCTGGGCTCGAGGCCGGGAGACGCCAGGGGAATTTCCGGGGTAGCGGTGAAATGCTATGATCCTGGAAGGACC<br>ACCGGTAGCGAAGGCGCCTGGCGAGAACGGGCCTGACGGTGAGAGACGAAAGCTGGGGGAG |
| p__Thermoproteota_g__ BA2_ASV_4148 | 0 | 0 | 0 | 0 | 1 | 0 | 0 | 0 | 0 | 0 | 0 | CCTTTACACTGTACGAAAGTGCGATAAGGGGACCCCGAGTGCGAGGGCATACAGTCCTCGCTTTTGTGTACCGTAAGATGGTACACGAATAAGAGCTGGGCAAGACCGGTGCCAGCCGCCGCGGTAATACCAGCTCTCCGAGTGGTAGGGATG<br>ATTATTGGGCTTAAAGCGTCCGTAGCCGGCTTAGCAAGTTTCCTGTTAAACCCAGCGACTCAATCGTTGGATCGCGGGAAATACTACTAGGCTAGGAGGCGGGAGAGGTCAACGGTACTTCTGGGGTAGGGGTGAAATCCTATAATCCCGGGA<br>GGACCACCAGTGGCGAAGGCTGTTGACTAGAACGCGCTCGACGGTGAGGGACGAAAGCTGGGGGAG |
| p__Nanoarchaeota_g__ BMS3ABIN17_ASV_3 826 | 1 | 0 | 0 | 0 | 0 | 0 | 0 | 0 | 0 | 0 | 0 | ATTCTGCAATGCGCGAAAGCGTGACAGATTGAGTCGGAGTGCTTCTCTATTGAGAAGCTTTTGCCAAATGTAAAAAGTTTGGCGAATAAGGACTGGGTAAGACTAGTGCCAGCCGCCGCGGTAATACTAGCGGTCCAAGTCGCAGCCATTTTT<br>ATTGGGTCTAAAACATCCGTAGCTTGCTTAGTAAGTTTCTTGTGAAATCTGGCTTTTTAAGAGTCAGGCGTGCAGGAAATACTACTAAGCTAGAGACCGGGAGACGTAAGAAGTACGCCCAAAGTAGCGGTAAAATGTGTTAATCTTGGGTGG<br>ACTATCAACAGCGAAGGCATCTTACGAGAACGGTTCTGACAGTAAGGGATGAAAGCTAGGGGCG |
| p__Nanoarchaeota_g__ BMS3ABIN17_ASV_3 841 | 1 | 0 | 0 | 0 | 0 | 0 | 0 | 0 | 0 | 0 | 0 | CTTTTGCAATGCGCGAAAGCGTGACAGAGTGAGTCGAAGTGCTTCTCACATGAGAAGCTTTTGCCAAATGTAAAAAGTTTGGCGAATAAGGACTGGGTAAGACTAGTGCCAGCCGCCGCGGTAATACTAGCGGTCCAAGTCGCAGCCATTTTT<br>ATTGGGTCTAAAACATCCGTAGCTTGCTTGGTAAGTTCCTTGTGAAATCTGGCTTCCTAAAAGTCGGGCGTGCAGGGAATACTGCCAAGCTAGAGACTGGGAGACGTAAAGAGTACGTCTGAAGTAGCGGTAAAATGTATTAATCTCAGGCGG<br>ACTATCAACAGCGAAGGCATTTTACGAGAACAGTTCTGACAGTGAGGGATGAAAGCCAGGGGCG |
| p__Nanoarchaeota_g__ BMS3ABIN17_ASV_3 843 | 1 | 0 | 0 | 0 | 0 | 0 | 0 | 0 | 0 | 0 | 0 | CTTTTACAATGCTCGAAAGAGTGATAGAGCGAGCCGGAGTGTTTCTCTTTGAGAAACTTTTGTCTGCTGTAAATAGGCAGACGAATAAGGACTGGGCAAGACCGGTGCCAGCCGCCGCGGTAACACCGGCGGTCCGAGTCGCATCCACATTTA<br>TTGGGTCTAAAACATTCGTAGCTTGTTTTACAAGTCTCTTGTGAAATCCTATGTCTCAAGCATAGGAATTGCAAGAGATACTGTATTGCTAGAGACCGGAAGACGCAAGGAGTACGGTTAAGGTAGCGGTAAAATGTGTTAATCTTAACTGGA<br>CTAACAACGGCGAAGGCACCTTGCGAGTACGGATCTGACAGTGAGGGATGAAGGCCAGGGGCG |
| p__Nanoarchaeota_g__ BMS3ABIN17_ASV_3 991 | 0 | 1 | 0 | 0 | 0 | 0 | 0 | 0 | 0 | 0 | 0 | AATCTACAATGCGCGAAAGCGTGATAGTTTGAGTCAGAGTGCGTTTCTTTGAAATGCTTTTGTGAAATGTAAAAAGTTTCACGAATAAGGACTGGGTAAGACGAGTGCCAGCCGCCGCGGTAATACTCGCGGTCCAAGTCGCAGCCATTTTTA<br>TTGGGTCTAAAACATCCGTAGCTTGCTTTGTAAGTTCCTTGTGAAATCCCGTATCTCAAGTATGGGGCGGGCAAGGAGTACTGCCAAGCTAGAGACTGGGAGACGTAAAGAGTACGTTTAAGGTAGCGGTAAAATGTGTTAATCTTAAGCGGA<br>CTAACAATAGCGAAGGCACTTTACGAGAACAGTTCTGACAGTAAGGGATGAAGGCTAGGGGCG |
| p__Nanoarchaeota_g__ BMS3ABIN17_ASV_4 290 | 0 | 0 | 0 | 0 | 0 | 0 | 0 | 1 | 1 | 0 | 0 | CTTTTGCAATGCGCGCAAGCGTGACAAAGCGAATCAGAGTGCTTTTCAATTTTGAAAAGCTTTTATGCGCTGTAAAAAGGCGCAGGAATAAGGACTGGGTAAGACGAGTGCCAGCCGCCGCGGTAATACTCGCGGTCCAAGTCGCAGCCAATT<br>TTATTGGGTCTAAAACATCCGTAGCTTGCTTGGTAAGTTTCTTGTGAAATCCTGTCTCCTAAGGGCAGGGCGAGCAGGAAGTACTGTCAAGCTAGAGACTGGAAGACGTGAGAAGTACGTTCAAAGTAACGGTAAAATGTGTTAATCTTGAAC<br>GGACTCACAATAGCGAAGGCATCTCACGAGGACAGTTCTGACAGTAAGGGATGAAAGCTAGGGGCG |
| p__Nanoarchaeota_g__ CG1-02-57- 44_ASV_3955 | 0 | 0 | 0 | 0 | 0 | 0 | 0 | 0 | 0 | 0 | 1 | CCTTTACCATGTACGAAAGTGTGATAAGGGGATCCCCAGTGCTCATGTAATGCATGGGCTTTTGCCAAGCGTAAATAACTTGGCGAATAAGTGGTGGGTAAGACTGGTGCCAGCCGCCGCGGTAACACCAGCGCCACGAGTGGTAACCGCGAT<br>TATTGGGCCTAAAGCGTCCGTAGCCGGCCTCGTAAATCCCTTGTGAAATTGTGCGGCTCAACCGAACAACGCGCAAGGGACACTGCAAGGCTCGAGACCGGGAGGAGTCAGAAGTATTCCAGAGGGAGCGGTAAAATGTTATAATCTCTGGAG<br>GACTACCTGTGGCGAAGGCGTCTGACCAGAACGGATCTGACGGTGAGGGACGAAAGCCAGGGGAG |
| p__Thermoplasmatota_ g__EX4484- 6_ASV_4321 | 0 | 0 | 0 | 0 | 0 | 0 | 0 | 0 | 1 | 0 | 0 | CCTTCGCACTGTGCGAAAGCGCGATGAGGGGATCCCAAGTGCTTGCTCTTAGGAGTAAGCTGTTTTCCTGTTTAAATCGCAGGAAGAGGAAGGGCTGGGTAAGACTGGTGCCAGCCGCCGCGGTAATACCAGCGGCCCTAGTGGTGACCGTTA<br>TTATTGGGCCTAAAGCGTTCGTAGCCGGTCCGGTAAATCTCTGGGTAAATCGATGGGCTTAACCCATCGAATTCTGGGGAGACTGCTGGACTTGAGACCGGGAGAGGTCGGAGGTACTACGGGGGTAGAGGTGAAATTCTGTAATCCCTGTGG<br>GACCACCGGTGGCGAAAGCGTCCGACCAGAACGGGTCTGACGGTCAGGGACGAAGCCTTGGGGCG |
| p__Halobacteriota_g__ Halalkalirubrum_ASV_ 4241 | 0 | 0 | 0 | 0 | 0 | 0 | 1 | 0 | 0 | 0 | 0 | CCTTTACACTGCACGCGAGTGCGATAGGGGGACCCCGAGTGCGAGGGCATACTAGCCCTCGCTTTTGTGAACCGTAAGGCGGTTCACGAACAAGAGCTGGGCAAGACCGGTGCCAGCCGCCGCGGTAATACCGGCAGCTCGAGTGATGACCGA<br>TTTTATTGGGCCTAAAGCGTCCGTAGCTGGCCACGAAAGCCCGTCGGGAAATCTCTCGGCTCAACCGAGAGGCGTCCGGCGGAACCTCCGTGGCTTGGAACCGGAAGGCTCAGAGGGTACGTCCAGGGTAGGAGTGAAATCCTGTAATCCTGG<br>ACGGACCACCGATGGCGAAAGCACTCTGAGAGGACGGATTCGACAGTGAGGGACGAAAGCTAGGGTCT |
| p__Halobacteriota_g__ Halapricum_3815_ASV _4255 | 0 | 0 | 0 | 0 | 0 | 0 | 0 | 1 | 0 | 0 | 0 | CCTTTACACTGCACGACAGTGCGATAGGGGGACTCCGAGTGCGAGGGCATATAGTCCTCGCTTTTGTACACCGTAAGGTGGTGTACGAATAAGAACTGGGCAAGACCGGTGCCAGCCGCCGCGGTAATACCGGCAGTTCGAGTGATGGCCGCT<br>TTTATTGGGCCTAAAGCGTTCGTAGCTGGCCAGTCAAGTCTGTCGGGAAATCCAGCGGCTCAACCGCTGGGCGTCCGGCGGAAACTGATTGGCTTGGGACCGGGAGACCCGAAGAGTACGTCCGGGGTAGGAGTGAAATCCCGTAATCCTGGA<br>CGGACTACCGATGGCGAAAGCACTTCGGGAGCACGGATCCGACAGTGAGGAACGAAAGCTGGGGTCT |
| p__Halobacteriota_g__ Halobaculum_ASV_386 7 | 0 | 0 | 0 | 0 | 0 | 0 | 0 | 0 | 0 | 1 | 0 | CCTTTACACTGCACGACAGTGCGATAGGGGGACTCCGAGTGCGAGGGCATATCGTCCTCGCTTTTCTGTACCGTAAGGTGGTACAGGAAGAAGGGCTGGGCAAGACCGGTGCCAGCCGCCGCGGTAATACCGGCAGCCCGAGTGATGGCCGTT<br>ATTATTGGGCCTAAAGCGTCCGTAGCCGGCCGGGCAAGTCCGTTGGGAAATCCACCCGCTTAACGGGTGGGCGTCCGGCGGAAACTGCTCGGCTTGGGACCGGAAGACCCGAGGGGTACGTCCGGGGTAGGAGTGAAATCCTGTAATCCTGGA<br>CGGACCGCCGATGCCGAAAGGACCTCGGGAGGACGGATCCGACGGTGAGGGACGAAAGCTTGGATCT |
| p__Halobacteriota_g__ Halobaculum_ASV_400 9 | 0 | 1 | 0 | 0 | 0 | 0 | 0 | 0 | 0 | 0 | 0 | CCTTTACACTGCACGCGAGTGCGATAGGGGGACCCCGAGTGCGAGGGCATATAGTCCTCGCTTTTCTGGACGGTAAGGCCGTCCAGGAACAAGAGCTGGGCAAGACCGGTGCCAGCCGCCGCGGTAATACCGGCAGCTCGAGTGATGACCGAT<br>TTTATTGGGCCTAAAGCGTCCGTAGCCGGCCGGGCAAGTCCGTCGGGAAATCTACCAGCTCAACTGGTAGGCGTCCGGCGGAAACTGCTCGGCTTGGGACAGGAAGACCCGAGGGGTACGTCCGGGGTAGGAGTGAAATCCCGTAATCCTGGA<br>CGGACCACCGATGGCGAAAGCACCTCGGGAGGACTGATCCGACGGTCAGGGACGAAAGCTGGGGTCT |
| p__Halobacteriota_g__ Halohasta_ASV_3836 | 1 | 0 | 0 | 0 | 0 | 0 | 0 | 0 | 0 | 0 | 0 | CCTTTACACTGCGCGACAGCGCGATAAGGGGACTCCACGTGCCAGGGCATATCGTCCTGGCTTTTGTGTACTGTAAGGAGATACACGAATAAGAGCTGGGCAAGACCGGTGCCAGCCGCCGCGGTAATACCGGCAGCTCGAGTGATGTCCAAT<br>ATTATTGGGCCTAAAGCGTCCGTAGCTGGCCAGCCAAGTCCGTCGGGAAATCCACCTGCCCAACAGGTGGGCGTCCGGCGGAAACTGTCTGGCTTGGAACCGGAAGGCTCAGAGGGTACGTCCGGGGTAGGAGTGAAATCCCGTAATCCCGGA<br>CGGACCACCGATGGCGAAAGCACTCTGAGAAGACGGATTCGACAGTGAGGGACGAAAGCTAGGGTCT |
| p__Halobacteriota_g__ Halohasta_ASV_4084 | 0 | 0 | 1 | 0 | 0 | 0 | 0 | 0 | 0 | 0 | 0 | TCTTCACAATGCACGAAAGTGCGATGGGGGGACTCCGAGTGCGTGATTCCCACAAGGAATCACGCTTTTCTGCACTGTAAGGAGGTACACGAATAAGAGCTGGGCAAGACCGGTGCCAGCCGCCGCGGTAATACCGGCAGCTCGAGTGATGTC<br>CACTATTATTGGGCCTAAAGCGTCCGTAGCTGGCCGCGCAAGTCCGTCGGGAAATCCACCTGCCCAACAGGTGGGCGTCCGGCGGAAACTGCGTGGCTTGGAACCGGAAGGCTCAGAGGGTACGTCCGGGGTAGGAGTGAAATCCCGTAATCC<br>CGGACGGACCACCGATGGCGAAAGCACTCTGAGAGGACGGATTCGACAGTGAGGGACGAAAGCTAGGGTCT |
| p__Halobacteriota_g__ Haloplanus_ASV_3931 | 0 | 0 | 0 | 0 | 0 | 0 | 0 | 0 | 0 | 0 | 1 | CCTTTACACTGCACGCAAGTGCGATAAGGGGACCCCGAGTGCGAGGGCATACAGTCCTCGCTTTTCACCACCGTAGGGAGGTGGTGGAATAAGAGCTGGGCAAGACCGGTGCCAGCCGCCGCGGTAATACCGGCAGCTCGAGTGATGGCCGAT<br>TTTATTGGGCCTAAAGCGTCCGTAGCCGGCCGAGCAAGTCCATCGGGAAATCCGCCAGCCTAACTGGCGGGCGTCCGGTGGAAACTGCACGGCTCGGGACCGGAAGACCCGAAGGGTACGTCTTGGGTAGGAGTGAAATCCCGTAATCCTGGA<br>CGGACCACCGATGGCGAAAGCACTTCGGGAAGACGGATCCGACGGTGAGGGACGAAAGCCAGGGTCT |
| p__Halobacteriota_g__ Haloprofundus_ASV_3 973 | 0 | 0 | 0 | 0 | 0 | 0 | 0 | 0 | 0 | 0 | 1 | CCTTTACACTGCACGACAGTGCGACAAGGGGACCCTGAATGCGAGGGCATACAGTCCTCGCTTTTCCCGACCGTAAGGAGGTCGGAGAATAAGAGCTGGGCAAGACCGGTGCCAGCCGCCGCGGTAATACCGGCAGCTCGAGTGATGACCGCT<br>GTTATTGGGCCTAAAGCGTCCGTAGCTGGCCGCGTAAGTCCATCGGGAAATCCATCCGCTCAACGGATGGGCGTCCGGTGGAAACTACACGGCTTGGAGCCGGAAGACCCAAGGGGTACGTCCGGGGTAGGAGTGAAATCCCGTAATCCTGGA<br>CGGACCACCGATGGCGAAAGCACCTTGGGAAGACGGACTCGACAGTGAGGGACGAAAGCTAGGGTCT |

| Phylum_Genus_ASV_number | Presence (1) or Absence (0) in each layer | | | | | | | | | | | Sequence |
|---|---|---|---|---|---|---|---|---|---|---|---|---|
| | L1 | L2 | L3 | L4 | L5 | L6 | L7 | L8 | L9 | L10 | L11 | |
| p__Halobacteriota_g__Halorhabdus_ASV_4202 | 0 | 0 | 0 | 0 | 0 | 1 | 0 | 0 | 0 | 0 | 0 | CCTTTACACTGCACGACAGTGCGATAGGGGGACTCCGAGTGCGAGGGCATATAGTCCTCGCTTTTCTGTACCGTAAGGAGGTAGAAGAATAAGGGCTGGGCAAGACCGGTGCCAGCCGCCGCGGTAATACCGGCAGCCCGAGTGATGGCCGCTATTATTGGGCCTAAAGCGTCCGTAGCCGGCCAGACAAGTCCGTTGGGAAATCGACGCGCTCAACGCGTCGGCGTCCGGCGGAAACTGTCTGGCTTGGGGCCGGAAGACTCGAGGGGTACGTCCGGGGTAGGAGTGAAATCCCGTAATCCTGGACGGACCACCGGTGGCGAAAGCGCCTCGAGAAAACGGACCCGACGGTGAGGGACGAAAGCTAGGGTCT |
| p__Halobacteriota_g__Halorubrum_ASV_3934 | 0 | 0 | 0 | 0 | 0 | 0 | 0 | 0 | 0 | 0 | 1 | CCTTTACACTGCACGACAGTGCGATAAGGGAATCCCAAGTGCGTAGGCATAGAGCCTACGCTTTTGTGTACCCTAGGGCGGTACACGAATAAGGGCTGGGCAAGACCGGTGCCAGCCGCCGCGGTAATACCGGCAGCCCAAGTGATGGCCGATTTTATTGGGCCTAAAGCGTCCGTAGCTGGCCGCACAAGTCCATCGGAAAATCCACCCGCTCAACGGGTGGGCGTCCGGTGGAAACTGTGTGGCTTGGGACCGGAAGGCGCGACGGGTACGTCCGGGGTAGGAGTGAAATCCCGTAATCCTGGACGGACCGCCGATGGCGAAAGCACGTCGCGAGAACGGATCCGACAGTGAGGGACGAAAGCTAGGGTCT |
| p__Halobacteriota_g__Halorubrum_ASV_3994 | 0 | 1 | 0 | 0 | 0 | 0 | 0 | 0 | 0 | 0 | 0 | CCTTTACACTGCACGCCAGTGCGATAAGGGAATCCCAAGTGCGTAGGCATAGAGCCTACGCTTTTGTGTACCCTAGGGCGGTACACGAATAAGGGCTGGGCAAGACCGGTGCCAGCCGCCGCGGTAATACCGGCAGCCCAAGTGATGGCCGATTTTATTGGGCCTAAAGCGTCCGTAGCTGGCCGCACAAGTCCATCGGAAAATCCACCCGCTCAACGGGTGGGCGTCCGGTGGAAACTGTGTGGCTTGGGACCGGAAGGCGCGACGGGTACGTCCGGGGTAGGAGTGAAATCCCGTAATCCTGGACGGACCGCCGATGGCGAAAGCACGTCGCGAGAACGGATCCGACAGTGAGGGACGAAAGCTAGGGTCT |
| p__Halobacteriota_g__Halorubrum_ASV_4108 | 0 | 0 | 0 | 1 | 0 | 0 | 0 | 0 | 0 | 0 | 0 | CCTTTACACTGCACGACAGTGCGATAAGGGGACTCCGAGTGCGAGGGCATATCGCCCTCGCTTTTGTACACCGTAGGGAGGTGTACGAATAAGCGCTGGGCAAGACCGGTGCCAGCCGCCGCGGTAATACCGGCAGCGCAAGTGATGGCCGATTTTATTGGGCCTAAAGCGTCCGTAGCTGGCCGGGCAAGTCCATCGGGAAATCCACCCGCTCAACGGGTGGGCGACCGGTGGAAACTGCCCGGCTCGGGACCGGAAGACCCAACGGGTACGTCCAGGGTAGGAGTGAAATCCCGTAATCCTGGACGGACCGCCGATGGCGAAAGCACGTTGGGAAGACGGATCCGACAGTGAGGGACGAAAGCTGGGGTCT |
| p__Halobacteriota_g__Halorussus_ASV_4004 | 0 | 1 | 0 | 0 | 0 | 0 | 0 | 0 | 0 | 0 | 0 | CCTTTACACTGCACGCCAGTGCGATAAGGGGACCCCGAGTGCGAGGGCATACCGTCCTCGCTTTTCACGACCGTAAGGTGGTCGTGGAATAAGGGCTGGGCAAGACCGGTGCCAGCCGCCGCGGTAATACCGGCAGCCCGAGTAATGGCCGCTATTATTGGGCCTAAAGCGTCCGTAGCCGGCCGGACAGGTCCGTCGGGAAATCCACGCGCCTAACGCGTGGGCGTCCGGCGGAAACCAGCCGGCTTGGGGCCGGAAGACCCAGGGGGTACGTCCGGGGTAGGAGTGAAATCCTGTAATCCCGGACGGACCGCCGGTGGCGAAAGCGCCCTGGGAGGACGGACCCGACGGTGAGGGACGAAAGCTAGGGTCA |
| p__Halobacteriota_g__Halorussus_ASV_4039 | 0 | 0 | 1 | 0 | 0 | 0 | 0 | 0 | 0 | 0 | 0 | CCTTTACACTGCACGACAGTGCGATAGGGGGACTCCGAGTGCGAGGGCATACAGTCCTCGCTTTTCTGTACCGTAAGGAGGTACACGAATAAGGGCTGGGCAAGACCGGTGCCAGCCGCCGCGGTAATACCGGCAGCCCGAGTGATGGCCGCTATTATTGGGCCTAAAGCGTCCGTAGCCGGCCGGGCAAGTCCGTTGGGAAATCCACGCGCCTAACGCGTGGGCGTCCAGCGGAAACTGCCCGGCTTGGGGCCGGAAGACCTGAGGGGTACGTCCGGGGTAGGAGTGAAATCCCGTAATCCTGGACGGACCACCGGTGGCGAAAGCGCCTCAGGAAGACGGACCCGACGGTGAGGGACGAAAGCTAGGGTCT |
| p__Halobacteriota_g__Halorussus_ASV_4340 | 0 | 0 | 0 | 0 | 0 | 0 | 0 | 0 | 1 | 0 | 0 | CCTTTACACTGCACGAAAGTGCGATAAGGGGATCCCAAGTGCGAGGGCATACAGTCCTCGCTTTTCTGTACCGTAGGGAGGTACAAGAATAAGGGCTGGGCAAGACCGGTGCCAGCCGCCGCGGTAATACCGGCAGCCCAAGTGATGGCCGCTATTATTGGGCCTAAAGCGTCCGTAGCCGGCCGGATAAGTCCGTCGGGAAATCCACCCGCTCAATGGGTGGGCGTCCGGCGGAAACTGTCCGGCTTGGGGCCGGAGGGCCTGAGGGGTACGTCCGGGGTAGGAGTGAAATCCTGTAATCCCGGTCGGACCACCGGTGGCGAAAGCGCCTCAGGAGGACGGACCCGACGGTGAGGGACGAAAGCTAGGGTCA |
| p__Nanohaloarchaeota_g__Nanohalobium_ASV_3960 | 0 | 0 | 0 | 0 | 0 | 0 | 0 | 0 | 0 | 0 | 1 | CCTTCACAATGCACGACAGTGCGATGGGGGGATTCCGAGTGTGGTCTTCCCACAAGGAAGGCTACTTTTCTGTACTGTAAGGAGGTACTGGAAAAAGGGGTGGGCAATACTGCTGCCAGCCGCCGCGGTAATAGCAGAACCCCAAGTGGTAGTCGTTATTATTGGGCTTAAAGTGTTCGTAGCCGGCCGTGCAAGTCCGCTGTGAAATCCAAGCGCTTAACGTTTGGGCGTGCAGCGGAAACTGCACTGGCTAGGGACCGGACGGTGTCGAGGCTATTCCAGGAGTAGGGGTGAAATCCTAAAATTCCTGGAGGAGCGCCTGTGGCGGAAGCGCTCGACAAGTACGGATCCGACGGTGAGGAACGAAAGCCAGGGGAG |
| p__Nanohaloarchaeota_g__Nanohalobium_ASV_4130 | 0 | 0 | 0 | 1 | 0 | 0 | 0 | 0 | 0 | 0 | 0 | CCTTCACAATGCACGCAAGTGCGATGGGGGGACTCCGAGTGTAGTCTCCTCACAAAGGAGGCTACTTTTCTGTACTGTAAGGAGGTACAGGAACAAGGGGTGGGCAAGACGGCTGCCAGCCGCCGCGGTAATAGCCGCACCCCAAGTGGTGACCAGTATTATTGGGCTTAAAGTGTTCGTAGCCGGCCATGTAAGTCCGCTGTGAAATCCAAGCGCTTAACGTTTGGAAGTGCAGCGGATACTGCATGGCTAGGGACCGGATGGCGTCGAGGCTATTCCAGGGGTAGGAGTGAAATCCAAAAATCCCTGGAGGAGCACCTGTGGCGAAAGCGCTCGACGAATACGGATCCGACGGTGAGGAACGAAAGCCAGGGGAG |
| p__Halobacteriota_g__Natrarchaeobaculum_ASV_4060 | 0 | 0 | 1 | 0 | 0 | 0 | 0 | 0 | 0 | 0 | 0 | CCTTTACACTGCACGCCAGTGCGATAAGGGGACACCAAGTGCGAGGGCATATCGTCCTCGCTTTTCACCACCGTAAGGAGGTGGTAGAATAAGGGCTGGGCAAGACCGGTGCCAGCCGCCGCGGTAATACCGGCAGCCCGAGTGATGACCGCTCTTATTGGGCCTAAAGCGTCCGTAGCTGGCCTCGCAAGTCCATCGGGAAATCCGCACGCTCAACGTGCGGGCGTCCGGTGGAAACTGCGTGGCTTGGGACCGGAAGACCCGAGGGGTACGTCCGGGGTAGGAGTGAAATCCTGTAATCCTGGACGGACCGTCGGTGGCGAAAGCGCCTCAGGAGGACGGACCCGACGGTGAGGGACGAAAGCTAGGGTCA |
| p__Halobacteriota_g__Natrinema_ASV_4166 | 0 | 0 | 0 | 0 | 1 | 0 | 0 | 0 | 0 | 0 | 0 | CCTTTACACTGCACGCCAGTGCGATAAGGGGACTCCGAGTGCGAGGGCATATAGTCCTCGCTTTTCACCACCGTAAGGTGGTGGTGGAATAAGTGCTGGGCAAGACCGGTGCCAGCCGCCGCGGTAATACCGGCAGCACGAGTGATGACCGCTATTATTGGGCCTAAAGCGTCCGTAGCTGGCCACGCAAGTCTATCGGGAAATCTGCGCGCTTAACGCGCAGGCGTCCGGTGGAAACTGCGTGGCTTGGGACCGGAAGACCAGAGGGGTACGTCCGGGGTAGGAGTGAAATCCCGTAATCCTGGACGGACCGCCGATAGCGAAAGCACCTCGGGAAGACGGATCCGACAGTGAGGGACGAAAGCTGGGGTCA |
| p__Halobacteriota_g__Natronomonas_ASV_4037 | 0 | 0 | 1 | 0 | 0 | 0 | 0 | 0 | 0 | 0 | 0 | CCTTTACACTGCACGCCAGTGCGATAGGGGGACACCGAGTGCGAGGGCATAGAGCCCTCGCTTTTCTGGACCGTAAGGAGGTCCCGGAACAAGGGCTGGGCAAGACCGGTGCCAGCCGCCGCGGTAACACCGGCAGCCCAAGTGATGGCCGATATTATTGGGCCTAAAGCGTCCGTAGCTGGCCGAGCAAGTCCGTCGGGAAATCCACTCGCTCAACGAGTGGGCGTCCGGCGGAAACTGTTCGGCTTGGGACCGGAAGATCCGAGGGGTACGTCCGGGGTAGGAGTGAAATCCCGTAATCCTGGACGGACCACCGATGGCGAAAGCACCTCGGAAGGACGGATCCGACAGTGAGGGACGAAAGCTAGGGTCT |
| p__Halobacteriota_g__Salinarchaeum_ASV_4054 | 0 | 0 | 1 | 0 | 0 | 0 | 0 | 0 | 0 | 0 | 0 | CCTTTACACTGCACGAGAGTGCGATAGGGGGACCCCGAGTGCGAGGGCATACAGTCCTCGCTTTTCACCACTGTAAGGTGGTGGTGGAATAAGTGCTGGGCAAGACCGGTGCCAGCCGCCGCGGTAATACCGGCAGCACGAGTGATGGCCGATTTTATTGGGCCTAAAGCGTCCGTAGCCGGCCAGACAGGTCCATCGGGAAATCTGCACGCCCAACGTGCAGGCGTCCGGTGGAAACCAGTTGGCTTGGGGCCGGAAGATCCAGGGGGTACGTCCGGGGTAGGAGTGAAATCCTGTAATCCTGGACGGACCACCGGTGGCGAAAGCGCCTTGAGAGAACGGATCCGACGGTGAGGGACGAAAGCTAGGGTCA |
| p__Halobacteriota_g__Saliphagus_ASV_4033 | 0 | 1 | 0 | 0 | 0 | 0 | 0 | 0 | 0 | 0 | 0 | CCTTTACACTGCACGCGAGTGCGATAAGGGGACTCCAAGTGCGAGGGCATATAGCCCTCGCTTTTCACCACCGTAAGGTGGTGGTGGAATAAGTGCTGGGCAAGACCGGTGCCAGCCGCCGCGGTAATACCGGCAGCTCAAGTGATGGCCAATCTTATTGGGCCTAAAGCGTCCGTAGCCGGCCTCGCAAGTCCGTCGGGAAATCCGCACGCCTAACGTGCGGGCGTCCGGCGGAAACTGCGTGGCTTGGGACCGGAAGACTCGAGGGGTACGTCCGGGGTAGGAGTGAAATCCCGTAATCCTGGACGGACCACCGGTGGCGAAAGCGCCTCGAGAGGACGGATCCGACGGTGAGGGACGAAAGCTAGGGTCA |
| p__Halobacteriota_g__Saliphagus_ASV_4089 | 0 | 0 | 1 | 0 | 0 | 0 | 0 | 0 | 0 | 0 | 0 | CCTTTACACTGCACGCGAGTGCGATAAGGGGACTCCGAGTGCGAGGGCATATAGTCCTCGCTTTTCACCACCGTAAGGAGGTGGTGGAATAAGTGCTGGGCAAGACCGGTGCCAGCCGCCGCGGTAATACCGGCAGCACAAGTGATGGCCGCTATTATTGGGCCTAAAGCGTCCGTAGCCGGCCACGCAAGTCCGTCGGGAAATCCGCGCGCCTAACGCGCGGGCGTCCGGCGGAAACTGCGTGGCTTGGGACCGGAAGACTCGAGGGGTACGTCCGGGGTAGGAGTGAAATCCTGTAATCCTGGACGGACCACCGGTGGCGAAAGCGCCTCGAGAGGACGGATCCGACGGTGAGGGACGAAAGCTAGGGTCA |
| p__Halobacteriota_g__Saliphagus_ASV_4159 | 0 | 0 | 0 | 0 | 1 | 0 | 0 | 0 | 0 | 0 | 0 | CCTTTACACTGTACGCGAGTGCGATAAGGGGACCCCGAGTGCGAGGGCATACAGTCCTCGCTTTTGTGTACCGTAAGGAGGTACACGAATAAGTGCTGGGCAAGACCGGTGCCAGCCGCCGCGGTAATACCGGCAGCACGAGTGATGACCGCTATTATTGGGCCTAAAGCGTCCGTAGCCGGCCAGACAGGTCCGTCGGGAAATCCGCACGCTCAACGTGCGGGCGTCCGGCGGAACCCAGCTGGCTAGGGACCGGAAGACCCGAGGGGTACGTCCGGGGTAGGAGTGAAATCCCGTAATCCTGGACGGACCGCCGGTGGCGAAAGCGCCTCGGGAGGACGGATCCGACGGTGAGGGACGAACGCTAGGGTCA |
| p__Thermoplasmatota_g__SM1-50_ASV_3963 | 0 | 0 | 0 | 0 | 0 | 0 | 0 | 0 | 0 | 0 | 1 | CCTTCACACTGTGCGAAAGCGCGATGAGGGGATCCCAAGTGCTTGCACGTAAGTGTAAGCTGTTTCTTTGTTTAAAATGCATAGAGAGTAAGGGCTGGGTAAGACGGGTGCCAGCCGCCGCGGTAATACCTGCAGCTCAAGTGGTGACCACTATTATTGGGCCTAAAACGTCCGTAGCCTGTTTGATAAATGCCTGGGTAAATCGTATAGCTCAACTATACGAATTCCGGGTAGACTGCAGGGCTAGAGATCGGGAGAAGCTAGAGGTACTCTTGGGGTAGAGGTGAAATTCTGTAATCCTAGGGGGACCACCAGTGGCGAAGGCGTCTAGCTATAACGAGTCTGACGGTCAGGGACGAAGCCCTGGGGCG |
| p__Thermoplasmatota_g__SM1-50_ASV_3989 | 0 | 1 | 0 | 0 | 0 | 0 | 0 | 0 | 0 | 0 | 0 | ACTTCACACTGTGCGAAAGCACGATGAGGGAATCCCAAGTGCTTGCACGTAAGTGTAAGCTGTTTTTCTGTCTAAAACGCAGAGAGAGTAAGGGCTGGGTAAGACGGGTGCCAGCCGCCGCGGTAATACCTGCAGCTCAAGTGGTGGCCACTATTATTGGGCCTAAAACGTCCGTAGCCGGTTTGTTAAATGCCTGGGTAAATCGGACCGCTTAACGGTTCGAATTCCGGGTAGACTGACAAACTTGAGACCGGGAGAAGTTAGAGGTACTCCTAGGGTAGAGGTGAAATTCTGTAATCCTGGGGGACCACCAGTGGCGAAAGCGTCTAACTAGAACGGGTCTGACGGTGAGGGACGAAGCCCTGGGGCG |
| p__Thermoplasmatota_g__SM1-50_ASV_4316 | 0 | 0 | 0 | 0 | 0 | 0 | 0 | 0 | 1 | 0 | 0 | CCTTCACACTGTGCGAAAGCGCGATGAGGGGATCCCAAGTGCTTGCACGTAAGTGTAAGCTGTTTCTTTGTTTAAAACGCAGAGAGAGGAAGGGCTGGGTAAGACGGGTGCCAGCCGCCGCGGTAATACCTGCAGCTCGAGTGGTGACCGCTATTATTGGGCCTAAAACGTCCGTAGCCGGCTGAGTAAATGCTTGGGTAAATCGTGCCGCTTAACGGTACGAATTCCGGGTAAACTGCTCAGCTTGAGATCGGGAGAAGCTAGAGGTACTCCTGGGGTAGAGGTGAAATTCTGTAATCCTCGGGGGACCACCAGTGGCGAAGGCATCTAACCAGAACGGGTCTGACGGTGAGGGACGAAGCCCTGGGGCG |
| p__Thermoplasmatota_g__SM1-50_ASV_4319 | 0 | 0 | 0 | 0 | 0 | 0 | 0 | 0 | 1 | 0 | 0 | CCTTCACACTGTGCGAGAGCACGATGAGGGAATCCCAAGTGCTGGCACGTAAGTGTCAGCTGTTTCTTTGTCTAAAAAACAAAGAGAGTAAGGGCTGGGTAAGACGGGTGCCAGCCGCCGCGGTAATACCTGCAGCTCAAGTGGTGACCGCTATTATTGGGCCTAAAACGTCCGTAGCCTGTTTGGTAAATGCCTGGGTAAATCGTGTAGCTTAACTATACGAATTCCGGGTAGACTGCCAGGCTTGAGACCGGGAGAGGTCAGAGGTACTCCTGGGGTAGAGGTGAAATTCTGTAATCCTAGGGGGACCACCAGTGGCGAAGGCGTCTGACTAGAACGGGTCTGACGGTGAGGGACGAAGCCCTGGGGCG |
| p__Thermoplasmatota_g__SM1-50_ASV_4338 | 0 | 0 | 0 | 0 | 0 | 0 | 0 | 0 | 1 | 0 | 0 | ACTTCACAATGCGCGAAAGCGCGATGAGGGGATCCCAAGTGCTTGCACGTAAGTGTAAGCTGTTTTTCTGTCTAAAAAGCAGAGAGAGTAAGGGCTGGGTAAGACGGGTGCCAGCCGCCACAATAATACCTACAACTCAAATAGTGGCCGCTATTATTGGGCCTAAAACGTCCGTAGCCGGCTGAGTAAATGCCTGGGTAAATCGTGCCGCTTAACGGTACGAATTCCGGGTAGACTGCTTAGCTTGGGACCGGAAGAAGTTAGAGGTACTCCTGGGGTAGGGGTGAAATCCTGTAATCCTGGGGGGACCACCAGTGGCGAAGGCGTCTAACTAGGACGGGTCTGACGGTGAGGGACGAAGCCCTGGGGCG |
| p__Halobacteriota_f__Haloferacaceae_ASV_4087 | 0 | 0 | 1 | 0 | 0 | 0 | 0 | 0 | 0 | 0 | 0 | CCTTTACACTGCACGGAAGTGCGATAGGGGGACCCCGAGTGCGAGGGCATACAGTCCTCGCTTTTGTGTACCGTAAGGAGGTACACGAATAAGTGCTGGGCAAGACCGGTGCCAGCCGCCGCGGTAATACCGGCAGCACAAGTGATGGCCGCTATTATTGGGCCTAAAGCGTCCGTAGCCGGCCGAACAAGTCCGTCGGGAAATCCATCCGCTCAACGGATGGGCGTCCGGCGGAAACTGTTCGGCTTGGGACAGGAAGACTCGAGGGGTACGTCCGGGGTAGGAGTGAAATCCTGTAATCCTGGACGGACCGCCGATGGCGAAAGCACCTCGAGAAGACTGATCCGACGGTGAGGGACGAAAGCTGGGGTCT |
| p__Halobacteriota_f__Haloferacaceae_ASV_4150 | 0 | 0 | 0 | 0 | 1 | 0 | 0 | 0 | 0 | 0 | 0 | CCTCAACACTGCACGACAGTGCGATTGGGGGACTCCGAGTGCGAGGGCATACCGTCCTCGCTTTTCAGCACGGTAAGGACGTGTTGGAACAAGGGCTGGGCAAGACCGGTGCCAGCCGCCGCGGTAATACCGGCAGCCCGAGTGATGACCGATCTTATTGGGCCTAAAGCGTCCGTAGCCGGCCGGGCAAGTCCGTCGGGAAATCCGCGCGCTCAACGCGTGGGCGTCCGGCGGAACCTGTTCGGCTTGGGGCCGGAAGACCCAGGGGGTACGTCCGGGGTAGGAGTGAAATCCCGTAATCCCGGACGGACCGCCGATGGCGAAAGCACCCTGGGAGGACGGACCCGACGGTGAGGGACGAAAGCTGGGGTCT |
| p__Halobacteriota_f__Haloferacaceae_ASV_4327 | 0 | 0 | 0 | 0 | 0 | 0 | 0 | 0 | 1 | 0 | 0 | CCTTTACACTGCACGAGAGTGCGATAGGGGGATCCCAAGTGCGAGGGCATATTAGCCCTCGCTTTTGTGAACCGTAAGGCGGTTCACGAATAAGAGCTGGGCAAGACCGGTGCCAGCCGCCGCGGTAATACCGGCAGCTCGAGTGATGACCGATTTTATTGGGCCTAAAGCGTCCGTAGCTGGCCACACAAGTCCGTCGGGAAATCTACCTGCCCAACAGGTAGGCGTCCGGCGGAAACTGTGCGGCTTGGGACCGGAAGGCGTGAAGGGTACGTCCGGGGTAGGAGTGAAATCCTGTAATCCTGGGGGGACCACCAGTGGCGAAAGCGTCTAGCTAGAACGGGTCTGACGGTGAGGGACGAAACCCTGGGGCG |

| Phylum_Genus_ASV_number | L1 | L2 | L3 | L4 | L5 | L6 | L7 | L8 | L9 | L10 | L11 | Sequence |
|---|---|---|---|---|---|---|---|---|---|---|---|---|
| | Presence (1) or Absence (0) in each layer | | | | | | | | | | | |
| p__Halobacteriota_f__Natrialbaceae_ASV_3972 | 0 | 0 | 0 | 0 | 0 | 0 | 0 | 0 | 0 | 0 | 1 | CCTTTACACTGCACGCCAGTGCGATAAGGGGACTCCAGGTGCGAGGGCATACAGTCCTCGCTTTTGTGTACCGTAGGGAGGTACACGAATAAGGGCTGGGCAAGACCGGTGCCAGCCGCCGCGGTAATACCGGCAGCCCGAGTGATGACCGCTATTATTGGGCCTAAAGCGTCCGTAGCCGGCCTCGCAAGTCCGTCGGGAAATCCGCACGCCCAACGTGCGGGCGTCCGGCGGAAACTGCGTGGCTTGGGACCGGAAGACTCGAGGGGTACGTCTGGGGTAGGAGTGAAATCCTATAATCCCGGGAGGACCACCAGTGGCGAAGGCTGTCGACTAGAACGCGCTCGACGGTGAGGGACGAAAGCTGGGGGAG |
| p__Halobacteriota_f__Natrialbaceae_ASV_4006 | 0 | 1 | 0 | 0 | 0 | 0 | 0 | 0 | 0 | 0 | 0 | CCTTTACAATGCGCGAAAGCGCGATAAGGGGAGTCCAAGTGGTCGTTCCTCACAAAGGAGCGACCTTTTGCCAAGTGTAAGGAGCTTGGTGAATAAGGGGTGGGAAAGACGGCTGCCAGCCGCCGCGGTAATAGCCGCACCCCAAGTGGTGACCACGATTATTGGGCTTAAAGAGTCCGTAGCCGGCCACGCAAGTCTGCTGTGAAATCTGGCTGCTTAACAGTCAGGAGGGCAGCAGAAACTCCGTGGCTTGGGACCGGACGGCGTTAGGGCTATTCCAGGAGTAGGAGTGAAATCCTGAAATTCCTGGAGGAGCACCGGTGGCGAAAGCGCCTAACGAGGACGGATCCGACGGTGAGGGACGAAAGCCAGGGGAG |
| p__Halobacteriota_f__Natrialbaceae_ASV_4070 | 0 | 0 | 1 | 0 | 0 | 0 | 0 | 0 | 0 | 0 | 0 | CCTTTACACTGCACGCCAGTGCGATAGGGGGACTCCGAGTGCGAGGGCATATAGTCCTCGCTTTTGTACACCGTAAGGTGGTGTACGAATAAGAACTGGGCAAGACCGGTGCCAGCCGCCGCGGTAATACCGGCAGTTCGAGTGATGGCCGATATTATTGGGCCTAAAGCGTTCGTAGCTGGCCAGTCAAGTCTGTCGGGAAATCCAGCGACTCAACCGCTGGGCGTCCGGCGGAAACTGATTGGCTTGGGACCGGAAGATCCAGAGGGTACGTCTGGGGTAGGAGTGAAATCCCGTAATCCTAGACGGACCACCGGTGGCGAAAGCGCTCTGGAAGGACGGATCCGACAGTGAGGGACGAAAGCTAGGGTCA |
| p__Halobacteriota_f__Natrialbaceae_ASV_4228 | 0 | 0 | 0 | 0 | 0 | 0 | 1 | 0 | 0 | 0 | 0 | CCTTTACACTGCACGCCAGTGCGATAAGGGGACTCCATCTGCGAGGGCATATAGTCCTCGCTTTTCTCGACCGTAAGGTGGTCGAGGAATAAGTGCTGGGCAAGACCGGTGCCAGCCGCCGCGGTAATACCGGCAGCACGAGTGATGACCGCTATTATTGGGCCTAAAGCGTCCGTAGCTGGCCAGGCAAGTCCATCGGGAAATCCGCGTGCCCAACACGCGGGCGTCCGGTGGAACCTGCCTGGCTTGGGACCGGAAGACCAGAGGGGTACGTCTGGGGTAGGAGTGAAATCCCGTAATCCTGGACGGACCACCGGTGGCGAAAGCGCCTCTGGAAGACGGATCCGACGGTGAGGGACGAAAGCTCGGGTCA |
| p__Asgardarchaeota_f__Thorarchaeaceae_ASV_4322 | 0 | 0 | 0 | 0 | 0 | 0 | 0 | 0 | 1 | 0 | 0 | CCTCACCAATGCGCGAAAGCGTGAGTGGGCCAAGCCAAGTGACGGGGGATAACTCCGTCTGTGATGGATCCGTAAGGAGGACCATTAGAAAGGAGAGGGCAAGGCTGGTGCCAGCCGCCGCGGTAAAACCAGCTCTTCGAGTGGTGTCCATGATTATTGAGCTTAAAGCGTTCGTAGCCGGTTTGGTAGGTCCCTGCTTAAATCCGGCCGCCTAACGGTCGGTCTGGTAGGGATACCGCCTTACTTGGGGACGGGAGACGATGACGGTATTCCGCAGGTAGGGGTGAAATCCTATGATCTGCGGAGGACCACCAGTGGCGAAGGCGGTCATCGAGAACGTGCCCGACGGTGAGGAACGAAACCCAGGGTAG |
| p__Halobacteriota_o__Halanaerobiales_ASV_3858 | 0 | 0 | 0 | 0 | 0 | 0 | 0 | 0 | 0 | 1 | 0 | CCTTTACACTGCACGCCAGTGCGATAAGGGGACCCCGAGTGAAAGGGCATATCGCCCTTTCTTTTCTGAACCGTAGGGAGGTTCAGGAACAAGAGCTGGGCAAGACCGGTGCCAGCCGCCGCGGTAATACCGGCAGCTCGAGTGATGACCGATTTTATTGGGCCTAAAGCGTCCGTAGCTGGCCTCGCAAGTCCATCGGGAAATCCGCCAGCTTAACTGGCGGGCGTCCAGCAGAAACCAGTCGGCTTGGGGCCGGAAGACCTGAGGGGTACGTCCGGGGTAGGAGTGAAATCCTGTAATCCTGGACGGACCGTCGGTGGCGAAAGCGCCTCAGGAGGACGGACCCGACGGTGAGGGACGAAAGCTAGGGTCA |
| p__Halobacteriota_o__Halanaerobiales_ASV_3860 | 0 | 0 | 0 | 0 | 0 | 0 | 0 | 0 | 0 | 1 | 0 | CCTTCACACTGTGCGAAAGCGCGATGAGGGGATCCCAAGTGCTTGCTCTTAGGAGTAAGCTGTTTTCCTGTGTAAATCGCAGGAAGAGGAAGGGCTGGGTAAGACGGGTGCCAGCCGCCGCGGTAATACCGGCAGCTCAAGTGATGGCCGTTATTATTGGGCCTAAAGCGTCCGTAGCCGGCCGAACAGGTCTGTTGGGAAATCCGCCAGCTTAACTGGCGGGCGTCCAGCGGAAACCAGTCGGCTTGGGGCCGGAAGACCTGAGGGGTACGTCCGGGGTAGGAGTGAAATCCTGTAATCCTGGACGGACCGTCGGTGGCGAAAGCGCCTCAGGAGGACGGACCCGACGGTGAGGGACGAAAGCTAGGGTCA |
| p__Halobacteriota_o__Halanaerobiales_ASV_3933 | 0 | 0 | 0 | 0 | 0 | 0 | 0 | 0 | 0 | 0 | 1 | CCTTTACACTGCACGACAGTGCGATAGGGGGACTCCGAGTGCGAGGGCATACAGTCCTCGCTTTTCTGTACCGTAAGGTGGTACAGGAACAAGTGCTGGGCAAGACCGGTGCCAGCCGCCGCGGTAATACCGGCAGCACGAGTGATGGCCGATATTATTGGGCCTAAAGCGTCCGTAGCCGGCCCGACAAGTCCGTTGGGAAATCCACGCGCTCAACGCGTGGGCGTCCAGCGGAAACTGTCGGGCTTGGGACCGGAAGACCCAGAGGGTACGTCCGGGGTAGGAGTGAAATCCCGTAATCCTGGACGGACCACCGGTGGCGAAAGCGCTCTGGGAAGACGGATCCGACGGTGAGGGACGAAAGCTAGGGTCT |
| p__Halobacteriota_o__Halanaerobiales_ASV_4026 | 0 | 1 | 0 | 0 | 0 | 0 | 0 | 0 | 0 | 0 | 0 | ACTTTACACTGCACGTCAGTGCGATAAGGGGACTCCGAGTGCGAGGGCATATAGCCCTCGCTTTTGTCGACCGTAAGGTGGTCGACGAACAAGTGCTGGGCAAGACCGGTGCCAGCCGCCGCGGTAATACCGGCAGCACGAGTGATGGTCGATTTTATTGGGCCTAAAGCGTCCGTAGCTGGCCGAGCAAGTCTGTCGGGAAATCCAGTGGCTCAACCACTGGGCGTCCGGTGGAAACTGTTCGGCTTGGGACCGGAAGACCCGAGGGGTACGTCTGGGGTAGGAGCGAAATCCTATAATCCTGGACGGACCACCGATGGCGAAAGCACCTCGGGAGCACGGATCCGACAGTGAGGGACGAAAGCTAGGGTCT |
| p__Halobacteriota_o__Halanaerobiales_ASV_4036 | 0 | 0 | 1 | 0 | 0 | 0 | 0 | 0 | 0 | 0 | 0 | CCTTTACACTGCACGACAGTGCGATAAGGGGATCCCGAGTGCGAGGGCATATCGTCCTCGCTTTTGTGTACCGTAAGGTGGTACACGAATAAGTGCTGGGCAAGACCGGTGCCAGCCGCCGCGGTAATACCGGCAGCACGAGTGATGACCGTTGTTATTGGGCCTAAAGCGTCCGTAGCCGGCCAGACAAGTTCGTCGGGAAATCCACCCGCTCAACGGGTGGGCGTCCGGCGAAAACTGTCTGGCTTGGGGCCGGAAGACCCAGGGGGTACGTCCGGGGTAGGAGTGAAATCCCGTAATCCTGGACGGACCGCCGGTGGCGAAAGCGCCCTGGGAGGACGGACCCGACGGTCAGGGACGAAAGCTAGGGTCT |
| p__Halobacteriota_o__Halanaerobiales_ASV_4043 | 0 | 0 | 1 | 0 | 0 | 0 | 0 | 0 | 0 | 0 | 0 | CCTTTACACTGCACGCAAGTGCGATAAGGGGATCCCGAGTGCGAGGGCATATAGTCCTCGCTTTTCTGGACTGTAGGGAGGTCCAGGAACAAGAGCTGGGCAAGACCGGTGCCAGCCGCCGCGGTAATACCGGCAGCTCAAGTGATGGCCGATATTATTGGGCCTAAAGCGTCCGTAGCCGGCCGGATAAGTCCGTCGGGAAATCCACCCGCTCAACGGGTGGGCGTCCGGCGGAAACTGTCCGGCTTGGGGCCGGAAGGCCTGAGGGGTACGTCCGGGGTAGGAGTGAAATCCTGTAATCCTGGTCGGACCACCGATGGCGAAAGCACTCTGAGAGAACGGATTCGACAGTGAGGGACGAAAGCTAGGGTCT |
| p__Halobacteriota_o__Halanaerobiales_ASV_4079 | 0 | 0 | 1 | 0 | 0 | 0 | 0 | 0 | 0 | 0 | 0 | CCTTTACACTGCACGACAGTGCGATAGGGGGATTCCGAGTGCGAGGGCATACAGTCCTCGCTTTTGTGTACCGTAAGGTGGTACACGAATAAGAGCTGGGCAAGACCGGTGCCAGCCGCCGCGGTAATACCGGCAGCACAAGTGATGGCCGCTATTATTGGGCCTAAAGCGTCCGTAGCCGGCCGAACAAGTCCGTCGGGAAATCCATCCGCCTAACGGATGGGCGTCCGGCGGAAACTGTTCGGCTTGGGACAGGAAGACTCGAGGGGTACGTCCGGGGTAGGAGTGAAATCCCGTAATCCTGGACGGACCGCCGATGGCGAAAGCACCTCGAGAAGACTGATCCGACGGTGAGGGACGAAAGCTGGGGTCT |
| p__Halobacteriota_o__Halanaerobiales_ASV_4320 | 0 | 0 | 0 | 0 | 0 | 0 | 0 | 0 | 1 | 0 | 0 | CCTTTACACTGTACGCAAGTGCGATAAGGGGACCCCGAGTGCGAGGGCATACAGTCCTCGCTTTTGTGTACCGTAAGGAGGTACACGAATAAGGGCTGGGCAAGACCGGTGCCAGCCGCCGCGGTAATACCGGCAGCCCTAGTGATGGCCGCTATTATTGGGCCTAAAGCGTCCGTAGCCGGCCGGACAAGTCCGTTGGGAAATCCGTTCGCTCAACGAACGGGCGTCCAGCGGAAACTGTCCGGCTTGGGACCGGAAGACCCGAGGGGTACGTCCGGGGTAGGAGTGAAATCCCGTAATCCTGGACGGACCGCCGGTGGCGAAAGCGCCTCGGGAAGACGGATCCGACGGTGAGGGACGAAAGCCAGGGTCT |
| p__Halobacteriota_p__Halobacteriota_ASV_4226 | 0 | 0 | 0 | 0 | 0 | 0 | 1 | 0 | 0 | 0 | 0 | ACTTTACAATGCGGGAAACCGCGATAAGGGGACACTGAGTGCCAGCATATAATGCTGGCTGTCCGTATGTGTAAAATACATGCGTTAGCAAGGGCCGGGCAAGACCGGTGCCAGCCGCCGCGGTAACACCGGCGGCCCGAGTGGTGGTCGCTATTATTGGGTCTAAAGGGTCCGTAGCCGGTTTGATCAGTCCATTGGGAAATCTGGTGGCTTAACCGCCAGAATTCCAATGGATACTGTCAGACTTGGGACCGGAAGACTCGAGGGGTACGTCCGGGGTAGGAGTGAAATCCTGTAATCCTGGACGGACCACAGGTGGCGAAAGCGCCTCGAGAGGACGGATCCGACGGTGAGGGACGAAAGCTAGGGTCA |
| p__Thermoplasmatota_p__Thermoplasmatota_ASV_4042 | 0 | 0 | 1 | 0 | 0 | 0 | 0 | 0 | 0 | 0 | 0 | CCTTCACACTGTGCGAAAGCGCGATGAGGGGATTCCAAGTGCTAACTCGAAAGAGTTGGCTGTTTATCTGTCTAAAACGCAGGTAGAGTAAGAGCTGGGTAAGACGGCTGCCAGCCGCCGCGGTAATAGCTGCAGCTCGAGTGGTGACCACTTTTATTGGGCCTAAAACGTCCGTAGCCGGTCTAGTAAATGCCTGGGTAAATCGTGCCGCTTAACGGTACGAATTCCGGGTAGACTGCTAGACTTGAGATCGGGAGAGGTTAGAGGTACTCCTTGGGTAGAGGTGAAATTCTGTAATCCTAGGGGGACCACCTGTGGCGAAAGCGTGGGGAG |
| p__Thermoplasmatota_p__Thermoplasmatota_ASV_4059 | 0 | 0 | 1 | 0 | 0 | 0 | 0 | 0 | 0 | 0 | 0 | CCTTTACACTGCACGACAGTGCGATAGGGGGACTCCAAGTGCTAGGGCATATCGTCCTAGCTTTTCACCACCGTAAGGTGGTGGTGGAATAAGTGCTGGGCAAGACCGGTGCCAGCCGCCGCGGTAATACCGGCAGCACGAGTGATGACCGCTCTTATTGGGCCTAAAACGTCCGTAGCCGGCTGAGTAAATGCCTGGGTAAATCGTGTCGCTTAACGATACGAATTCCGGGTAGACTGCTTAGCTTGAGACTGGGAGAAGCTAGAGGTACTCCTGGGGTAGAGGTGAAATTCTGTAATCCTGGGGGGACCACCAGTGGCGAAAGCGTCTAGCTAGAACGGGTCTGACGGTGAGGGACGAAACCCTGGGGCG |
| d__Archaea_d__Archaea_ASV_4010 | 0 | 1 | 0 | 0 | 0 | 0 | 0 | 0 | 0 | 0 | 0 | ACTGCACAATGCGGGAAACCGCGATGCGGGAACTCCAAGTGCCAGCACAGAGTGTTGGCTGTTCCTTAGCCTAAACAGCTTTGGGAGTAAGGGCTGGGTAAGACGGGTGCCAGCCGCCGCGGTAATACCTGCAGCCCGAGTGGTGGCCGCTATTATTGGGCCTAAAGCGTCCGTAGCCGGCTCGGTAAATGCCTGTGTAAATTGTACCGCTTAACGGTGCGAACTGCGGGTAGACTGCCGGGCTTGGGACCGGGAGAGGCGCGGGGTACTCCAGGGGTAGAGGTGAAATTCTGTAATCCTTGGGGGACCACCGGTGGCGAAGGCGCCGCGCTGGAACGGGTCCGACGGTGAGGGACGAAGCCCTGGGGCG |
| d__Archaea_d__Archaea_ASV_4022 | 0 | 1 | 0 | 0 | 0 | 0 | 0 | 0 | 0 | 0 | 0 | ACTGCACAATGTGGGAAACCACGATGCGGGGACTCCAAGTGCCAGCACAGAGTGTTGGCTGTTTCCAAGTCTAAAAAACTCGGAGAGTAAGGGCTGGGTAAGACGGGTGCCAGCCGCCGCGGTAATACCTGCAGCCCGAGTGGTGGCCGCTATTATTGGGCCTAAAGCGTCCGTAGCCGGTCCGGTAAATGCCTGTGTAAATTGTACGGCTTAACCGTGCGAACTGCGGGTAGACTGCCGGACTTGGGACCGGGAGAGTCACGAAGTACTCCGGGGGTAGAGGTGAAATTCTGTAATCCTCGGGGGACTACCGGTGGCGAAGGCGTCGTGATAGAACGGCTCCAACGGTGAGGGACGAAGCCCTGGGGCG |
| d__Archaea_d__Archaea_ASV_4064 | 0 | 0 | 1 | 0 | 0 | 0 | 0 | 0 | 0 | 0 | 0 | CCTTCACAATGCACGCAAGTGCGATGGGGGGACTCCGAGTGTGGTCTTCCCACAAGGAAGGCCACTTTTCTGCACTGTAAGAAGGTGCCGGAACAAGGGGTGGGCAAGACTGCTGCCAGCCGCCGCGGTAATAGCAGCATCCCAAGTGGTGGCCAGTATTATTGGGCTTAAAGCGTTCGTAGCCGGACTTGTAAGTCCGCTGTGAAATCCACTAGCTTAACTAGTGGGCGTGCAGCGGATACTGCAGGTCTAGGGACCGGATGGCGTCGGGGCTATTCTCAGGGTAGGAGTGAAATCCTAAAATCCTGAGAGGAGCACCTGTGGCGAAAGCGCCCGACGAATACGGATCCGACGGTGAGGAACGAAAGCTGGGGGAG |
| d__Archaea_d__Archaea_ASV_4110 | 0 | 0 | 0 | 1 | 0 | 0 | 0 | 0 | 0 | 0 | 0 | CCTTTACAATGCACGAAAGTGTGATAAGGGGATCCTAAGTGCTCATGCAAATGTATGGGCTTTTGCCAAGAGTAAATATCTTGGAGAATAAGGGGTGGGTAAGACTGGTGCCAGCCGCCGCGGTAACACCAGCGCCTCAAGTGGCAACCACGTTTATTGGGCCTAAAGCGTCCGTAGCCGGTTAGGTAAATCCTTTGTGAAATCGTCAGGCTCAACCTGACGGCTTGCTTAGGAGACTGCCTAACTTGAGACCGGGAGGGGTCAAAGGTATTCCATGGGGAGCGGTAAAATGTGATAATCCATGGAGGATCGCCTATGGCGAAGGCATTTGACTAGAACGGATCTGACGGTCAGGGACGAAGGCCAGGGGAG |
| d__Archaea_d__Archaea_ASV_4233 | 0 | 0 | 0 | 0 | 0 | 0 | 1 | 0 | 0 | 0 | 0 | ACTGCACAATGCGGGAAACCGCGATGCGGGAACTCCAAGTGCCAGCACAGAGTGTTGGCTGTTCCTTAGCCTAAAAAGCTCTGGGAGTAAGGGCTGGGTAAGACGGGTGCCAGCCGCCGCGGTAATACCTGCAGCCCGAGTGGTGGCCGCTATTATTGGGCCTAAAGCGTCCGTAGCCTGCTCGGTAAATGCCTGTGTAAATTGTACCGCTTAACGGTGCGAAGTGCGGGTAGACTGCCGGGCTTGGGACCGGGAGAGGCGCGAAGTACTCCGGGGGTAGAGGTGAAATTCTGTAATCCTCGGGGGACTACCGGTGGCGAAGGCGTCGCGCTGGAACGGGTCCGACGGTGAGGGACGAAGCCCTGGGGCG |
| d__Archaea_d__Archaea_ASV_4337 | 0 | 0 | 0 | 0 | 0 | 0 | 0 | 0 | 1 | 0 | 0 | CCTTCACACTGTGCGAAAGCGCGATGAGGGAATCCCAAGTGCTTGCTCGTAAGAGTAAGCTGTTTCTGTGTATAAAAAGCATAGAGAGTAAGGGCTGGGTAAGACGGGTGCCAGCCGCCGCGGTAATACCTGCAGCTCTAGTGGTGACCACTATTATTGGGCCTAAAACGTTCGTAGCCGGTTTGCTAAATGCCTGGGTAAATCGAGCCGCTTAACGGTTCGAATTCCGGGTAGACTGGCAAACTTGAGACCGGGAGAGGCTAGAGGTACTCCTGGGGTAGAGGTGAAATCCTATAATCCCGGGAGGACCACCAGTGGCGAAGGCTGTTGACTAGAATGCGCTCGACGGTGAGGGACGAAAGCTGGGGGAG |
| p__Hadarchaeota_o__Hadarchaeales_ASV_4600 | 0 | 1 | 0 | 0 | 0 | 0 | 0 | 0 | 0 | 0 | 0 | CCTTCACACTGTGCGAAAGCGCGATGAGGGAATCCCAAGTGCTTGCTCTTAGGAGTAAGCTGTTTTTCTGTTTAAATCGCAGGAAGAGGAAGGGCTGGGTAAGACGGGTGCCAGCCGCCGCGGTAATACCTGCAGCTCAAGTGGTGACCGCTATTATTGGGCCTAAAACGTCCGTAGCCGGACCTGCAAGTCCCTTGTGAAATCGGGCGGCTTAACCGTTCGGCTTGCAAGGGAAACTGTTGGTCTTGGGGCCGATAGGCGCCGAGGGTACTCCGGGGGTAGCGGTGAAATGCTATAATCCCTGGGGGACCGCCGGTGGCGAAAGCGCTCGACGAGGTCGGACCCGACGGTGAGGGACGAAGGCCAGGGGAG |

| Phylum_Genus_ASV_number | Presence (1) or Absence (0) in each layer L1 | L2 | L3 | L4 | L5 | L6 | L7 | L8 | L9 | L10 | L11 | Sequence |
|---|---|---|---|---|---|---|---|---|---|---|---|---|
| p__Hadarchaeota_o__Hadarchaeales_ASV_4648 | 0 | 1 | 0 | 0 | 0 | 0 | 0 | 0 | 0 | 0 | 0 | CCTTCACACTGTGCGAAAGCGCGATGAGGGGATCCCAAGTGCTTGCACGTAAGTGTAAGCTGTTTCTTTGTCTAAAACGCATAGAGAGTAAGGGCTGGGTAAGACGGGTGCCAGCCGCCGCGGTAATACCTGCAGCCCAAGTGGTGACCGCTTTTATTGGGCCTAAAGCGTCCGTAGCCGGACTAGCAAGTCCTTTGTGAAATCGGGCGGCTTAACCGTTCGGCTTGCAAGGGAAACTGTTAGTCTTGGGGCCGACAGGCGCCGAGGGTACTCCGGGGGTAGCGGTGAAATGCTATAATCCCTGGGGGACCGCCGGTGGCGAAAGCGCTCAGCGAGGTCGGACCCGACGGTGAGGGACGAAAGCCAGGGGAG |
| p__Iainarchaeota_o__Iainarchaeales_ASV_5057 | 0 | 0 | 0 | 0 | 0 | 0 | 0 | 1 | 0 | 0 | 0 | CCTTTACACTGCACGAAAGTGTGATAAGGGGATTTCGAGTGCGCTCACTGAGCGCTTTTGCCAAATGTAAATAGTTTGGCGAATAAAGGGGGGGCAATATACGTGCCAGCCGCCGCGGTAATACGTAACCCCTAAGTGGTTTCCACTTGTATTGGGCCTAAAACACCTGTAGCCTGTTCAATAAGTCTTTGGTGAAATCAACCAGCTCAACTGGTTGAATTGCTAAAGATACTGTTGTGCTTGGGGAAGGAAGACGTGAAGAGAATTTTTGGGGTAACGGTGAAATGTGATGATCCCAAAAGGACTACCAGTAGCGAAGGCGCTTCACGAGAACTATCCCGACGGTGAGAGATGAAAGCTGGGGGAG |
| p__Halobacteriota_f__Natrialbaceae_ASV_4358 | 1 | 0 | 0 | 0 | 0 | 0 | 0 | 0 | 0 | 0 | 0 | CCTTTACACTGCGCGAAAGCGCGATAAGGGGACCCCGAGTGCGAGGGCGTACAGTCCTCGCTTTTGTACACCGTAAGGAGGTGTACGAATAAGGGCTGGGCAAGACCGGTGCCAGCCGCCGCGGTAATACCGGCAGCCCGAGTGATGACCGCTATTATTGGGCCTAAAGCGTCCGTAGCCGGCCACGCAAGTCCGTCGGGAAATCCGCACGCCCAACGTGCGGGCGTCCGGCGGAAACTGCCTGGCTTGGGACCGGAAGACCAGAGGGGTACGTCCGGGGTAGGAGTGAAATCCCGTAATCCTGGACGGACCACCGGTGGCGAAAGCGCCTCTGGAAGACGGATCCGACGGTGAGGGACGAAAGCTCGGGTCA |
| p__Nanoarchaeota_g__BMS3ABIN17_ASV_4497 | 0 | 0 | 0 | 0 | 0 | 0 | 0 | 0 | 0 | 1 | 0 | AATCTACAATGCACGCAAGTGTGATAGTTTGAGCCAGAGTGTTTCTCACTGAGAAACTTTTATGAAATGCAAAAAGTTTTATGAATAAGGACTGGGTAAGACGAGTGCCAGCCGCCGCGATAATACTCGCGGTCCAAGTCGCAGCCATTTTTATTGGGTCTAAAACATCCGTAGCTTGCCTGGTAAGTCCTTTGTGAAATCCTGTTCCTTAAGTTCAGGGCGCGCAGGGGGTACTGCCATGCTAGAGACCGGAAGACGTAACGAGTACGCTTAAAGTAATGGTAAAATATGTTAATCTTAAGCGGACTTACAACAGCGAAGGCACGTTACGAGGACGGTTCTGACAGTAAGGGATGAAGGCTAGGGGCG |
| p__Nanoarchaeota_g__BMS3ABIN17_ASV_4515 | 0 | 0 | 0 | 0 | 0 | 0 | 0 | 0 | 0 | 0 | 1 | ATTCTGCAATGTACGCAAGTATGACAGAGCAAGCCAGAGTGCGTTCCTATAATTGGAACGCTTTTATGTAATGTAAAAAGTTACATGAATAAGGACTGGGTAAGACTGGTGCCAGCCGCCGCGGTAATACCAGCGGTCCAAGTGGCAGCCGCGTTTATTGGGTCTAAAACATCCGTAGCTCGCCAAGTCAGCCTCTTGTGAAATCCGGCATCTCAAGTGTCGGGCGTGCAAGAGGTACTGCTTAGCTAGAGACCGGGAGACGTAAGGAGTACGGTTAGGGTAGCGGTAAAATGCGTTGATCCTTACTGGACTAACGATAGCGAAGGCACCTTACGAGAACGGATCTGACAGTGAGGGATGAAGGCTAGGGTCG |
| p__Nanoarchaeota_g__BMS3ABIN17_ASV_4520 | 0 | 0 | 0 | 0 | 0 | 0 | 0 | 0 | 0 | 0 | 1 | AATCTACAATGCGCGAAAGCGTGATAGTTTGAGCCAGAGTGTTTTACAATCGTAAAACTTTTGCCAAATGTAAAAAGTTTGGCGAATAAGGACTGGGTAAGACAGGTGCCAGCCGCCGCGGTAATACCTGCGGTCCAAGTCGCAGCCATTTTTATTGGGTCTAAAACATCCGTAGCTTGCTTTGTAAGTTCTCTGTGAAATCCTATATCTTAAATATGGGGCGTGCAGGGAATACTGCTTAGCTAGAGACCGGGAGACGTAAAGAGTATGTTCAAAGTAGCGGTAAAATGTTTTAATCTTGAGCAGACTAACAATAGCGAAGGCACTTTACGAGAACGGTTCTGACAGTCAGGGATGAAGGCTAGGGGCG |
| p__Nanoarchaeota_g__BMS3ABIN17_ASV_4529 | 0 | 0 | 0 | 0 | 0 | 0 | 0 | 0 | 0 | 0 | 1 | AATCTACAATGCGCGAAAGCGTGATAGTTTGAGTCAGAGTGCGTTTCTTTGAAATGCTTTTGTGAAATGTAAAAAGTTTCACGAATAAGGACTGGGTAAGACTAGTGCCAGCCGCCGCGGTAATACTAGCGGTCCAAGTCGCAGCCATTTTTATTGGGTCTAAAACATCCGTAGCTTGCTTGGTGAGTTCCTTGTGAAATCCTACATCTCAAGTGTAGGGCACGCAAGGAGTACTGCCAAGCTAGAGACTGGGAGACGTAAAGAGTACGTTCGAGGTAGCGGTAAAATGTGTTAATCTCGGACGGACTAACAATAGCGAAGGCACTTTACGAGAACAGTTCTGACAGTAAGGGATGAAGGCTAGGGGCG |
| p__Nanoarchaeota_g__BMS3ABIN17_ASV_4565 | 0 | 0 | 0 | 0 | 0 | 0 | 0 | 0 | 0 | 0 | 1 | CTTTTACAATGCACTAAAGTGTGATAGAGTGAGTCAGAGTGTTTTCCAATTGGAAAACTTTTGCCAAATGTAAAAAGTTTGGCGAATAAGGACTGGGTAAGACGAGTGCCAGCCGCCGCGGTAATACTCGCGGTCCAAGTCGCAGCCATTTTTATTGGGTCTAAAACATCCGTAGCTTGCTATATAAGTTTCCTGTGAAATCTTATCTCTTAAAGATAGGGCGTGCAGGAAATACTATTTAGCTAGAGACTGGGAGACGTAAGGAGTACGTTTGGAGTAGCGGTAAAATGTGTTAATCCCAAACGGACTAACAATAGCGAAGGCACCTTACGAGAACAGTTCTGACAGTAAGGGATGAAGGCTAGGGGCG |
| p__Nanoarchaeota_g__BMS3ABIN17_ASV_4630 | 0 | 1 | 0 | 0 | 0 | 0 | 0 | 0 | 0 | 0 | 0 | AATCTACAATGCGCGAAAGCGTGATAGTTTGAGCCAGAGTGTTTCTCAATGAGAGACTTTTGTGGAATGTAAAAAGTTCTACGAATAAGGACTGGGTAAGACGAGTGCCAGCCGCCGCGGTAATACTCGCGGTCCAAGTCGCAGCCATTTTTATTGGGTCTAAAACATCCGTAGCTTGCTTTGTAAGTCCTTTGTGAAATTTGATATCTTAAGTATTGAGCGTGCAAAGGGTACTGCGAAGCTAGAGACTGGTAGACGTAACGAGTATGTTCAGAGTAGCGGTAAAATGTGTTAATCCTGGACAGACTAACAATAGCGAAGGCACGTTACGAGGACAGTTCTGACAGTAAGGGATGAAGGCTAGGGGCG |
| p__Nanoarchaeota_g__BMS3ABIN17_ASV_4945 | 0 | 0 | 0 | 0 | 0 | 1 | 0 | 0 | 0 | 0 | 0 | AATCTACAATGCGCGAAAGCGTGATAGTTTGAGTCAGAGTGCGTTTCTTTGAAATGCTTTTGTGAAATGTAAAAAGTTTCACGAATAAGGACTGGGTAAGACGAGTGCCAGCCGCCGCGGTAATACTCGCGGTCCAAGTCGCAGCCATTTTTATTGGGTCTAAAACATCCGTAGCTTGCTTTGTAAGTTCCTTGTGAAATCCCGTATCTCAAGTATGGGGCGGGCAAGGAGTACTGCTAAGCTAGAGACTGGGAGACGTAAAGAGTACGTTTAAGGTAGCGGTAAAATGTGTTAATCTTAAGCGGACTAACAATAGCGAAGACACTTTACGAGAACAGTTCTGACAGTAAGGGATGAAGGCTAGGGGCG |
| p__Nanoarchaeota_g__BMS3ABIN17_ASV_5102 | 0 | 0 | 0 | 0 | 0 | 0 | 0 | 0 | 1 | 0 | 0 | AATCTACAATGCGCGAAAGCGTGATAGTTTGAGTCAGAGTGCGTTTCTTTGAAATGCTTTTGTGAAATGTAAAAAGTTTCACGAATAAGGACTGGGTAAGACGAGTGCCAGCCGCCGCGGTAATACTCGCGGTCCAAGTCGCAGCCATTTTTATTGGGTCTAAAACATCCGTAGCTTGCTTTGTAAGTTCCTTGTGAAATCCTATATCTCAAATATAGGGCGGGCAAGGAGTACTGCTAAGCTAGAGACTGGGAGACGTAAAGAGTACGTTTAAGGTAGCGGTAAAATGTGTTAATCTTAAGCGGACTAACAATAGCGAAGGCACTTTACGAGAACAGTTCTGACAGTAAGGGATGAAGGCTAGGGGCG |
| p__Nanoarchaeota_g__BMS3ABIN17_ASV_5103 | 0 | 0 | 0 | 0 | 0 | 0 | 0 | 0 | 1 | 0 | 0 | CTTCTGCAATGTACGAAAGTATGACAGAGCAAGCCAGAGTGATTCTCTATAGCTGAGAATCTTTTGTGTGATGTAAAAAGTCGCACGAATAAGGACTGGGCAAGACTGGTGCCAGCCGCCGCGGTAATCCCAGCGGTCCAAGTCGCAGCCGCTTTTATTGGGTCTAAAACATCCGTAGCTTGTTTCGTTAGTCTCGTGTGAAATCGGGCTTCTCAAGGGTCCGGCGGGCACGAGATACTGCGTTGCTAGGGACCGGGAGACGTAAGGAGTACACATCGGGTAGCGGTAAAATGCGTTGATCCAATGTGGACTAACGATAGCGAAGGCACCTTACGAGAACGGATCCGACAGTGAGGGATGAAGGCTAGGGGCG |
| p__Nanoarchaeota_g__BMS3ABIN17_ASV_5106 | 0 | 0 | 0 | 0 | 0 | 0 | 0 | 0 | 1 | 0 | 0 | CTTCTTCAATGCGCGCAAGCGTGAAAGAGTGAACCAGAGTGCTTTCCAATTTGGAAAGCTTTTGCCAAATGTAAAAAGTTTGGCGAATAAGGACTGGGTAAGACGAGTGCCAGCCGCCGCGGTAATACTCGCGGTCCAAGTCGCAGCCATATTTATTAGGTCTAAAACATCCGTAGCTTGTTAGAAAAGTTCTCTGTGAAATCTTATCTCTTAAAGATAAGGCGGGCAGGGAATACTTTCAAACTAGAGACTGGAAGACGTAAGAAGTATGGTCAAAGTAGTGGTAAAATTCTTTAATCTTGACCAGACTAACAACAGCGAAGGCATCTTACGAGGACAGTTCTGACAGTAAAGGATGAAGGCTAGGGGCG |
| p__Nanoarchaeota_g__BMS3ABIN17_ASV_5112 | 0 | 0 | 0 | 0 | 0 | 0 | 0 | 0 | 1 | 0 | 0 | CTTTTACAATGCACGAAAGTGTGATAGAGCGAGCCAAAGTGCTTTCCGCACGGAAAGCTTTTCTTCGCTGTAAAAAGGCGGAGGAATAAGGACTGGGTAAGACTGGTGCCAGCCGCCGCGGTAATCCCAGCAGTCCAAGTTGCAGCCAACATTATTGGGTCTAAAACATCCGTAGCTTGCTAGGTAAGTTTCTTGTGAAATCCTACTTCTCAAGGGTAGGGCGTGCAAGAAATACTATTTAGCTAGAGACCGGTAGATGTAAGGAGTATGTTCAAGGTAGTGGTAAAATATGTTAATCTTGAGCAGACTAACAATCAGCGAAGGCACCTTACAAGGACGGATCTGACAGTGAGGGATGAAGGCCAGGGGCG |
| p__Nanoarchaeota_g__BMS3ABIN17_ASV_5118 | 0 | 0 | 0 | 0 | 0 | 0 | 0 | 0 | 1 | 0 | 0 | AATCTACAATGCGCGCAAGCGTGATAGTTTGAATCAGAGTGCTTCTCAATGAGAAGCTTTTGTGAAATCTAAAAAGTTTTACGAATAAGGACTGGGTAAGACGAGTGCCAGCCGCCGCGGTAATACTCGCGGTCCAAGTCGCAGCCATATTTATTGGGTCTAAAACATCCGTAGCTTGCTTAATAAGTTCCTTGTGAAATTCTATGTCTCAAGCATAGGAAGTGCAAGGAGTACTGTTAAGCTAGAGACTGGAAGACGTAACGAGTACGTTCAGAGTAGCGGTTAAATGCGTTAATCCTGAATGGACTAACACGAGCGAAGGCACGTTACGAGGACAGTTCTGACAGTAAGGGATGAAAGCCGGGGTCG |
| p__Nanoarchaeota_g__BMS3ABIN17_ASV_5129 | 0 | 0 | 0 | 0 | 0 | 0 | 0 | 0 | 1 | 0 | 0 | ATTTTACAATGCACGAAAGTGTGATAGAATCAGCCAGAGTGTTTCTCTTTGAGAAACTTTTGCCAAATGTAAAAAGTTTGGCGAATAAGGACTGGGCAAGACCGGTGCCAGCCGCCGCGGTAATACCGGCGGTCCAAGTCGCATCCACATTTATTGGGTCTAAAACATTCGTAGCTTGTTTTGTAAGTCTCTTGTGAAATCCCACATCTCAAGTGTGGGGCGTGCAAGAGATACTACATTGCTAGAGACCGGAAGACGTAAGGAGTACATTTGAGGTAGTGGTAAAATACGTTAATCTTAAATGGACTAACAATGGCGAAGGCACCTTACGAGTACGGATCTGACAGTGAGGGATGAAGGCCAGGGGCG |
| p__EX4484-52_g__EX4484-52_ASV_4653 | 0 | 1 | 0 | 0 | 0 | 0 | 0 | 0 | 0 | 0 | 0 | CCTCCGCAATGCACGCAAGTGTGACGGGGGGATTCCAAGTGTTATTGTTTTACAATAACTTTTCTGAAGTGTAAATAGCTTCAGGAATAAGAGCTGGGTAAGACGGGTGCCAGCCGCCGCGGTAATACCCGCAGCTCAAGTGGTCTGGACTTTTATTGGGTCTAAAGCGTTCGTAGCTGGATTTTTAAGTTCTCTGTGAAATCTAACAGCTCAACTGTTAGGCGTGCAGAGGATACTATTAATCTTGAAATCGGGATGGGCCAAGGGTACTCTCGGGGTAGGGGTGAAATCCAATAAGCCTGAGGGGACCACCTGTGGCGAAGGCGCTTGGCTAGAACGAGTTTGACAGTGAGGGACGAAAGCTAGGGGAG |
| p__Thermoplasmatota_g__EX4484-6_ASV_4597 | 0 | 1 | 0 | 0 | 0 | 0 | 0 | 0 | 0 | 0 | 0 | ACTTCACAATGCGAGAAATCGCGATGAGGGAACTCCAAGTGCCTATGCATAGCATAGGCTGTTCAGGTGTCTAAAAAACATCTGGAGTAAGGGCTGGGAAAGACTGGTGCCAGCCGCCGCGGTAATACCAGCAGCTCGAGTGGTGGCCGCTTTTATTGGGCCTAAAGCGTCCGTAGCCGGTCCGGTAAATCCTTGGGTAAATCGTCGGGCTCAACTCGACGAATTCCGAAGAGACTGCCGGACTTGGGACTGGGAGAGGTCGGAGGTACTCTAGGGGTAGGGGTGAAATCCTGTAATCCTTAGAGGACCACCAGTGGCGAAGGCGTCCGACTAGAACGGGTCCGACGGTAAGGGACGAAGCCGTGGGTCG |
| p__Halobacteriota_g__Haladaptatus_ASV_4682 | 0 | 0 | 1 | 0 | 0 | 0 | 0 | 0 | 0 | 0 | 0 | ACTTTACACTGCACGCCAGTGCGATAAGGGAACTCCAAGTGCGAGGGCATATAGTCCTCGCTTTTGTGTATCGTAAGGTGGTACACGAATAAGAGCTGGGCAAGACCGGTGCCAGCCGCCGCGGTAATACCGGCAGCTCAAGTGATGGCCAATATTATTGGGCCTAAAGCGTCCGTAGCCTGCCAGACAGGTCCGTCGGGAAATCTGCTCGCTCAACGAGCAGGCGTCCGGCGGAAACCAGCTGGCTTGGGGCCGGAAGACCCAAGGGGTACGTCTGGGGTAGGAGTGAAATCCTGTAATCCTGGACGGACCACCAATGGCGAAAGCACGTCGAGAGACCGGACCCGACGGTGAGGGACAAAAGCCAGGGTCT |
| p__Halobacteriota_g__Halalkalicoccus_ASV_4999 | 0 | 0 | 0 | 0 | 0 | 0 | 1 | 0 | 0 | 0 | 0 | CCTTTACACTGCACGACAGTGCGATAGGGGGACTCCGAGTGCGAGGGCATACAGTCCTCGCTTTTGTACACCGTAAGGTGGTGTACGAATAAGAGCTGGGCAAGACCGGTGCCAGCCGCCGCGGTAACACCGGCAGCTCGAGTGATGGCCGATTTTACTGGGCCTAAAGCGTCCGTAGCTGGCCGCACAAGTCCATCGGGAAATCCAGGCGCTCAACGTCTGGCCGTCCGGTGGAAACTGTGCGGCTTGGGACCGGAAGACCCGAGGGGTACGTCCGGGGTAGGAGTGAAATCCTGTAATCCTGGACGGACCACCGGTGGCGAAAGCGCCTCGGGAAGACGGATCCGACAGTGAGGGACGAAAGCTGGGGTCT |
| p__Halobacteriota_g__Halococcus_ASV_4955 | 0 | 0 | 0 | 0 | 0 | 1 | 0 | 0 | 0 | 0 | 0 | CCTTTACACTGCACGCCAGTGCGATAAGGGGATCCCGAGTGCGAGGGCATACAGTCCTCGCTTTTGTGTACCGTAAGGTGGTACTCGAATAAGGGCTGGGCAAGACCGGTGCCAGCCGCCGCGGTAATACCGGCAGCCCAAGTGATAGCCACTATTATTGGGCCTAAAGCGTCCGTAGCCTGCCGAACGGGTCCGTCGGGAAATCCACCCGCTCAACGGGTGGGCGTCCGGCGGAAACCAGTCGGCTTGGGACCGGGAGACCTGTGGGGTACGTCCGGGGTAGGAGTGAAATCCTGTAATCCTGGACGGACCGCCGGTGGCGAAAGCGCCACAGGAGAACGGATCCGACGGTGAGGGACGAAAGCTAGGGTCT |
| p__Halobacteriota_g__Halohasta_ASV_4710 | 0 | 0 | 1 | 0 | 0 | 0 | 0 | 0 | 0 | 0 | 0 | CCTTTACACTGCACGACAGTGCGATAAGGGAACTCCGAGTGCGAGAGCATATAGTCCTCGCTTTTCTGCACCGTAAGGTGGTGCACGAATAAGAGCTGGGCAAGACCGGTGCCAGCCGCCGCGGTAATACCGGCAGCTCAAGTGATGTCCACTATTATTGGGCCTAAAGCGTCCGTAGCTGGCCGCGCAAGTCCGTCGGGAAATCCACCTGCCCAACAGGTGGGCGTCCGGCGGAAACTGCGTGGCTTGGAACCGGAAGGCTCAGAGGGTACGTCCGGGGTAGGAGTGAAATCCCGTAATCCCGGACGGACCACCGATGGCGAAAGCACTCTGAGAGGACGGATTCGACAGTGAGGGACGAAAGCTAGGGTCT |
| p__Halobacteriota_g__Halohasta_ASV_4728 | 0 | 0 | 1 | 0 | 0 | 0 | 0 | 0 | 0 | 0 | 0 | CCTTTACACTGCACGCAAGTGCGATAGGGGAATCCCAAGTGCGAGGGCATACTAGCCCTCGCTTTTGTGTACTGTAAGGAGGTACACGAATAAGAGCTGGGCAAGACCGGTGCCAGCCGCCGCGGTAATACCGGCAGCTCGAGTGATGTCCACTATTATTGGGCCTAAAGCGTCCGTAGCTGGCCGCGCAAGTCCGTCGGGAAATCCACCCGCTCAACGGGTGGGCGTCCGGCGAAAACTGTCTGGCTTGGGGCCGGAAGACCCAGGGGGTACGTCCGGGGTAGGAGTGAAATCCCGTAATCCCGGACGGACCACCGATGGCGAAAGCACTCTGAGAGGACGGATTCGACAGTGAGGGACGAAAGCTAGGGTCT |

| Phylum_Genus_ASV_ number | Presence (1) or Absence (0) in each layer | | | | | | | | | | | |
|---|---|---|---|---|---|---|---|---|---|---|---|---|
| | L1 | L2 | L3 | L4 | L5 | L6 | L7 | L8 | L9 | L10 | L11 | Sequence |
| p__Halobacteriota_g__Halohasta_ASV_5115 | 0 | 0 | 0 | 0 | 0 | 0 | 0 | 0 | 1 | 0 | 0 | CCTTTACACTGCACGACAGTGCGATAGGGGGACTCCGAGTGCGAGGGCATACAGTCCTCGCTTTTCTGTACCGTAAGGAGGTACAGGAACAAGTGCTGGGCAAGACCGGTGCCAGCCGCCGCGGTAATACCGGCAGCACGAGTGATGGCCGATATTATTGGGCCTAAAGCGTCCGTAGCTGGCCGTACAAGTTCGTCGGGAAATCCACCTGCCCAACAGGTGGGCGTCCGGCGAAAACTGTATGGCTTGGAACCGGAAGGCTCAGAGGGTACGTCCGGGGTAGGAGTGAAATCCTGTAATCCCGGACGGACCACCGATGGCGAAAGCACTCTGAGAAGACGGATTCGACAGTGAGGGACGAAAGCTAGGGTCT |
| p__Halobacteriota_g__Halomicrobium_3849_ASV_4628 | 0 | 1 | 0 | 0 | 0 | 0 | 0 | 0 | 0 | 0 | 0 | CCTTCACACTGTGCGAAAGCGCGATGAGGGGATTCCAAGTGCTAACTCGAAAGAGTTGGCTGTTTATCTGTCTAAATCGCAGGAAGAGGAAGGGCTGGGTAAGACGGGTGCCAGCCGCCGCGGTAATACCGGCAGCACGAGTGATGGCCGCTCTTATTGGGCCTAAAGCGTCCGTAGCCGGCCAGTCAAGTCCGTTGGGAAATCGGCGCGCCTAACGCGTCGGCGTCCAGCGGAAACTGACTGGCTTGGGGCCGGAAGACCTGAGGGGTACGTCCGGGGTAGGAGTGAAATCCCGTAATCCTGGACGGACCACCGGTGGCGAAAGCGCCTCAGGAAGACGGACCCGACGGTGAGGGACGAAAGCTAGGGTCT |
| p__Halobacteriota_g__Halorubrum_ASV_4619 | 0 | 1 | 0 | 0 | 0 | 0 | 0 | 0 | 0 | 0 | 0 | CCTTTACACTGCACGCCAGTGCGATAAGGGGACTCCGAGTGCGAGGGCATATCGCCCTCGCTTTTGTACACCGTAGGGAGGTGTACGAATAAGGGCTGGGCAAGACCGGTGCCAGCCGCCGCGGTAATACCGGCAGCCCGAGTGATGGCCGATCTTATTGGGCCTAAAGCGTCCGTAGCTCACCGGGTAAGTCCGTCGGGAAATCCACCCGCTCAACGGGTGGGCGTCCGACGGAACCTGCCTGGCTTGGGACCGGAAGACCCAGCGGGTACGTCCTGGGTAGGAGTGAAATCCTGTAATCCTGGACGGACCACCGATGGCGAAAGCACGCTGGGAGGACGGATCCGACAGTGAGGGACGAAAGCTGGGGTCT |
| p__Halobacteriota_g__Halorubrum_ASV_4693 | 0 | 0 | 1 | 0 | 0 | 0 | 0 | 0 | 0 | 0 | 0 | CCTTTACACTGCACGACAGTGCGATAAGGGGACTCCAAGTGCGAGGGCATATCGTCCTCGCTTTTCACTACCGTAAGGTGGCGGTGGAATAAGGGCTGGGCAAGACCGGTGCCAGCCGCCGCGGTAATACCGGCAGCCCGAGTGATGGCCGATCTTATTGGGCCTAAAGCGTCCGTAGCTGGCCGCACAAGTCCATCGGAAAATCCACCCGCTCAACGGGTGGGCGTCCGGTGGAAACTGTGTGGCTTGGGACCGGAAGGCGCGACGGGTACGTCCGGGGTAGGAGTGAAATCCCGTAATCCTGGACGGACCGCCGATGGCGAAAGCACGTCGCGAGAACGGATCCGACAGTGAGGGACGAAAGCTAGGGTCT |
| p__Halobacteriota_g__Halorubrum_ASV_4880 | 0 | 0 | 0 | 0 | 1 | 0 | 0 | 0 | 0 | 0 | 0 | CCTTTACACTGCACGACAGTGCGATAGGGGGATCCCAAGTGCACAGGCATAGCGCCTGTGCTTTTCGGTACCGTAAGGTGGTACCAGAATAAGGGCTGGGCAAGACCGGTGCCAGCCGCCGCGGTAATACCGGCAGCCCAAGTGATGGCCGATCTTATTGGGCCTAAAGCGTCCGTAGCTGGCCACGCAAGTCCATCGGGAAATCCACCTGCTCAACAGGTGGGCGCCCGGTAGAAACTGCGTGGCTTGGGACCGGAAGGCGCGACGGGTACGTCCGGGGTAGGAGTGAAATCCCGTAATCCTGGCCGGACCACCGGTGGCGAAAGCGCCTCGGGAAGACGGATCCGACGGTGAGGGACGAAAGCTAGGGTCA |
| p__Halobacteriota_g__Halorussus_ASV_4482 | 0 | 0 | 0 | 0 | 0 | 0 | 0 | 0 | 0 | 1 | 0 | ACTTTACACTGCACGACAGTGCGATAAGGGAACTCCGAGTGCGAGGGCATATAGTCCTCGCTTTTCTGCACCGTAAGGTGGTGCACGAATAAGAGCTGGGCAAGACCGGTGCCAGCCGCCGCGGTAATACCGGCAGCTCAAGTGATGGCCGCTATTATTGGGCCTAAAGCGTCCGTAGCCGGCCAGACAGGTCCGTCGGGAAATCCAGTGGCTTAACCACTGGGCGTCCGGCGGAAACCAGCTGGCTTGGGGCCGGAAGATCTGAGGGGTACGTCCGGGGTAGAAGTGAAATCCCGTAATCCTGGACGGACCGCCGGTGGCGAAAGCGCCTCAGAAGGACGGACCCGACGGTGAGGGACGAAAGCTAGGGTCA |
| p__Halobacteriota_g__Halosimplex_ASV_4603 | 0 | 1 | 0 | 0 | 0 | 0 | 0 | 0 | 0 | 0 | 0 | CCTTTACACTGCACGACAGTGCGATAGGGGAACTCCGAGTGCGAGGGCATAGAGCCCTCGCTTTTGTAGACCCTAAGGCGGTCTACGAACAAGTGCTGGGCAAGACCGGTGCCAGCCGCCGCGGTAATACCGGCAGCACGAGTGATGGCCGATCTTATTGGGCCTAAAGCGTCCGTAGCTGGCCGTGCAAGTCCGTCGGGAAATCCAGTGGCTCAACCTCTGGGCTTCCGGCGGAAACTGCTCGGCTTGGGACCGGAAGACTCGAGGGGTACGTTCGGGGTAGGAGTGAAATCCCGTAATCCTGAACGGACCACCGATGGCGAAAGCACCTCGAGAGCACGGATCCGACAGTGAGGGACGAAAGCTAGGGTCT |
| p__Halobacteriota_g__Halostagnicola_ASV_4376 | 1 | 0 | 0 | 0 | 0 | 0 | 0 | 0 | 0 | 0 | 0 | ATTCCGCAATGCGTGAAAGCGCGACGGAGTGACTCCTAGTGCCGGTGGTCTAGCGACCGTCGGCTTTTTTGGAACGTAAAAAGTTCCAAGAATAAGAGCTGGGCAAGACCGGTGCCAGCCGCCGCGGTAATACCGGCAGCACGAGTGATGACCGCTATTATTGGGCCTAAAGCGTCCGTAGCTGGCCACGCAAGTCCATCGGGAAATCTCTCCGCTCAACGGAGAGGCGTCCGGTGGAAACTGTGTGGCTTGGGACCGGAAGATCCAGAGGGTACGTCTGGGGTAGGAGTGAAATCCCGTAATCCGAGACGGACCACCGGTGGCGAAAGCGCTCTGGAAAGACGGATCCGACGGTGAGGGACGAAAGCTCGGGTCA |
| p__Halobacteriota_g__Halostagnicola_ASV_4504 | 0 | 0 | 0 | 0 | 0 | 0 | 0 | 0 | 0 | 0 | 1 | CCTTTACACTGCACGCGAGTGCGATAAGGGGACTCCAAGTGCGAGGGCATATAGTCCTCGCTTTTTCCGACCGTAGGATGGTCGGCGAATAAGTGCTGGGCAAGACCGGTGCCAGCCGCCGCGGTAATACCGGCAGCACGAGTGATGACCGCTATTATTGGGCCTAAAGCGTCCGTAGCTGGCCACGCAAGTCCATCGGGAAATCTCTCCGCTCAACGGAGAGGCGTCCGGTGGAAACTGTGTGGCTTGGGACCGGAAGATCCAGAGGGTACGTCTGGGGTAGGAGTGAAATCCCGTAATCCTGGACGGACCGCCGATGGCGAAAGCACCTTGGGAAGACGGACTCGACAGTGAGGGACGAAAGCTAGGGTCT |
| p__Halobacteriota_g__Halovenus_ASV_4690 | 0 | 0 | 1 | 0 | 0 | 0 | 0 | 0 | 0 | 0 | 0 | CCTTTACACTGCACGCCAGTGCGATAAGGGGACTCCGAGTGCGAGGGCATACAGTCCTCGCTTTTCTGCACCGTAAGGTGGTGCAGGAACAAGTGCTGGGCAAGACCGGTGCCAGCCGCCGCGGTAATACCGGCAGCACAAGTGATGGCCGATATTATTGGGCCTAAAGCGTCCGTAGCCGGCCGGACAAGTCTGTTGGGAAATCCACGCGCTTAACGCGTGGGCGTCCAGCGGAAACTGTCTGGCTTGGGGCCGGAAGACTCGAGGGGTACGTCCGGGGTAGGAGTGAAATCCCGTAATCCTGGACGGACCGCCGGTGGCGAAAGCGCCTCGAGAGGACGGACCCGACGGTGAGGGACGAAAGCCAGGGTCT |
| p__Halobacteriota_g__Halovenus_ASV_4734 | 0 | 0 | 1 | 0 | 0 | 0 | 0 | 0 | 0 | 0 | 0 | CCTTTACACTGCACGCCAGTGCGATAAGGGGACACCGAGTGCGAGGGCATAGAGCCCTCGCTTTTCTGGACCGTAGGGAGGTCCAGGAACAAGGGCTGGGCAAGACCGGTGCCAGCCGCCGCGGTAATACCGGCAGCACAAGTGATGGCCGATATTATTGGGCCTAAAGCATCCGTAGCTGGCCAGACAAGTCTGTTGGGAAATCCACGCGCCCAACGCGTGGGCGTCCAGCGGAAACTGTCTGGCTTGGGGCCGGAGGACTCGAGGGGTACGTCCGGGGTAGGAGTGAAATCCCGTAATCCTGGACGGACCACCGGTGGCGAAAGCGCCTCGAGAGGACGGACCCGACAGTGAGGGACGAAAGCCAGGGTCT |
| p__Methanobacteriota_A_1229_g__Methanobacterium_D_1054_ASV_4569 | 0 | 0 | 0 | 0 | 0 | 0 | 0 | 0 | 0 | 0 | 1 | CCTCCGCAATGCACGAAAGTGCGACGGGGGGACCCCAAGTGCCACTCTTAACGGGGTGGCTTTTATGAAGTGTAAAAAGCTTTAAGAATAAGGGCTGGGCAAGACCGGTGCCAGCCGCCGCGGTAACACCGGCAGCTCAAGTGGTGGCCGTTTTTATTGGGTCTAAAGCGTTCGTAGCCGGCCTGATAAGTCTCTGGTGAAATCCCATAGCTTAACTGTGGGAATTGCTGGAGATACTATCAGGCTTGAGATCGGGAGAGGTTAGGGGTACTCCCAGGGTAGGGGTGAAATCCTATAATCCTGGGAGGACCACCTGTGGCGAAGGCGCCTAACTGGAACGAATCTGACGGTGAGTAACGAAAGCCAGGGGCG |
| p__Methanobacteriota_A_1229_g__Methanobacterium_D_1054_ASV_5157 | 0 | 0 | 0 | 0 | 0 | 0 | 0 | 0 | 1 | 0 | 0 | CCTCCGCAATGCACGAAAGTGCGACGGGGGGACCCCAAGTGCCACTCTTAACGGGGTGGCTTTTATGAAGTGTAAAAAGCTTCAAGAATAAGGGCTGGGCAAGACCGGTGCCAGCCGCCGCGGTAACACCGGCAGCTCAAGTGGTGGCCGTTTTTATTGGGCCTAAAGCGTTCGTAGCCGGCCTGATAAGTCTCTGGTGAAATCCCATAGCTTAACTGTGGGAATTGCTGGAGATACTATCAGGCTTGAGGTCGGGAGAGGTTAGGGGTACTCCCAGGGTAGGGGTGAAATCCTATAATTCTGGGAGGACCACCTGTGGCGAAGGCGCCTAACTGGAACGAACCTGACGGTGAGTAACGAAAGCCAGGGGCG |
| p__Halobacteriota_g__Natronobiforma_ASV_4521 | 0 | 0 | 0 | 0 | 0 | 0 | 0 | 0 | 0 | 0 | 1 | CCTTTACACTGCACGCCAGTGCGATAAGGGGACTCCAAGTGCGAGGGCATATAGTCCTCGCTTTTCACCACCGTAAGGTGGTGGTGGAATAAGTGCTGGGCAAGACCGGTGCCAGCCGCCGCGGTAATACCGGCAGCACAAGTGATGACCGCTATTATTGGGCCTAAAGCGTCCGTAGCCGGCCACGCAAGTCCATCGGGAAATCTGCTCGCCTAACGAGCAGGCGTCCGACGGAAACTGTGTGGCTTGGGACCGGAAGATCCAGAGGGTACGTCTGGGGTAGGAGTGAAATCCCGTAATCCTGGACGGACCACCGGTGGCGAAAGCGCTCTGGAAGGACGGATCCGACGGTGAGGGACGAAAGCTAGGGTCA |
| p__Thermoplasmatota_g__Natronoplasma_ASV_4373 | 1 | 0 | 0 | 0 | 0 | 0 | 0 | 0 | 0 | 0 | 0 | CCTTCACACTGTGGGAAACCACGATGAGGGGATTCCAAGTGCCAACACTTTGTGTTGGCTTTTCACCTATGTAAAAAGTAGGTGGAATAAGGGCCGGGTAAGACGGGTGCCAGCCGCCGCGGTAACACCCGCGGCCCAAGTAGTGGTCATTAATATTGAGCTTAAAGCGTCCGTAGCCGGTCTGGTAAATCTCTGGGTAAATCGGACGGCTCAACCGTTCGAATACCAGGGAGACTGCTAGACTTGGGGCCGGGAGAGGTCAGAGGTACTCCTGGGGTAGAAGTAAAATTCTGTAATCCTAGGGGGACCACCTGTGGCGAAAGCGTCTAACCAGAACGAGTCTGACGGTGAGGGACGAAGCCCTGGGTCA |
| p__Halobacteriota_g__Salinarchaeum_ASV_4661 | 0 | 1 | 0 | 0 | 0 | 0 | 0 | 0 | 0 | 0 | 0 | CCTTTACACTGCACGAAAGTGCGATAAGGGGACCCCGAGTGCGAGGGCATACAGTCCTCGCTTTTGTGTACCGTACGATGGTACACGAATAAGAGCTGGGCAAGACCGGTGCCAGCCGCCGCGGTAATACCGGCAGCACAAGTGATGGCCGATTTTATTGGGCCTAAAGCGTCCGTAGCCGGCCAGACAGGTCCATCGGGAAATCTGCACGCCCAACGTGCAGGCGTCCGGTGGAAACCAGCTGGCTTGGGGCCGGAAGATCCAGGGGGTACGTCCGGGGTAGGAGTGAAATCCCGTAATCCCGGACGGACCGCCGGTGGCGAAAGCGCCCTGGAAGGACGGACCCGACGGTGAGGGACGAAAGCTAGGGTCA |
| p__Halobacteriota_g__Salinarchaeum_ASV_4853 | 0 | 0 | 0 | 0 | 1 | 0 | 0 | 0 | 0 | 0 | 0 | CCTTTACACTGCACGAGAGTGCGATAGGGGGACCCCGAGTGCGAGGGCATACAGTCCTCGCTTTTCTGCACCGTAAGGTGGTGCAGGAACAAGTGCTGGGCAAGACCGGTGCCAGCCGCCGCGGTAATACCGGCAGCACGAGTGATGGCCGATCTTATTGGGCCTAAAGCGTCCGTAGCCGGCCAGACGGGTCCGTCGGGAAATCTGCACGCCCAACGTGCAGGCGTCCGGCGGAAACCAGCTGGCTCGGGGCCGGAAGACCCAGGGGGTACGTCCGGGGTAGGAGTGAAATCCCGTAATCCTGGACGGACCGCCGGTGGCGAAAGCGCCCTGGGAGGACGGACCCGACGGTGAGGGACGAAAGCTAGGGTCA |
| p__Halobacteriota_g__Saliphagus_ASV_4697 | 0 | 0 | 1 | 0 | 0 | 0 | 0 | 0 | 0 | 0 | 0 | CCTTTACACTGCACGACAGTGCGATAAGGGGACTCCAAGTGCGAGGGCATACAGTCCTCGCTTTTCACTACCGTAAGGTGGTAGTGGAATAAGTGCTGGGCAAGACCGGTGCCAGCCGCCGCGGTAATACCGGCAGCACGAGTGATGACCGCTATTATTGGGCCTAAAGCGTCCGTAGCCGGCCTCGCAAGTCCGTCGGGAAATCCGCACGCCTAACGTGCGGGCGTCCGGCGGAAACTGCGTGGCTTGGGACCGGAAGACTCGAGGGGTACGTCCGGGGTAGGAGTGAAATCCCGTAATCCTGGACGGACCACCGGTGGCGAAAGCGCCTCGAGAGGACGGATCCGACGGTGAGGGACGAAAGCTAGGGTCA |
| p__Halobacteriota_g__Saliphagus_ASV_4834 | 0 | 0 | 0 | 1 | 0 | 0 | 0 | 0 | 0 | 0 | 0 | CCTTTACACTGCACGCCAGTGCGATAAGGGGACTCCCAGTGCGAGGGCATATCGTCCTCGCTTTTGTGTACCGTAGGGAGGTACACGAATAAGGGCTGGGCAAGACCGGTGCCAGCCGCCGCGGTAATACCGGCAGCCCGAGTGATGACCGCTATTATTGGGCCTAAAGCGTCCGTAGCCGGCCACGCAAGTCCGTCGGGAAATCCGCACGCCCAACGTGCGGGCGTCCGGCGGAAACTGCCTGGCTTGGTACCGGAAGACCAGAGGGGTACGTCCGGGGTAGGAGTGAAATCCCGTAATCCCGGACGGACCACCGATGGCGAAAGCACTCTGAGAGGACGGATTCGACAGTGAGGGACGAAAGCTAGGGTCT |
| p__Halobacteriota_g__Saliphagus_ASV_5014 | 0 | 0 | 0 | 0 | 0 | 0 | 1 | 0 | 0 | 0 | 0 | CCTTTACACTGCACGCAAGTGCGATAGGGGAATCCCAAGTGCGAGGGCATATTAGCCCTCGCTTTTGTGTACTGTAAGGAGGTACACGAATAAGAGCTGGGCAAGACCGGTGCCAGCCGCCGCGGTAATACCGGCAGCCCGAGTGATGACCGCTATTATTGGGCCTAAAGCGTCCGTAGCCGGCCACGCAAGTCCGTCGGGAAATCCGCGCGCCTAACGCGCGGGCGTCCGGCGGAAACTGCGTGGCTTGGGACCGGAAGACTCGAGGGGTACGTCCGGGGTAGGAGTGAAATCCTGTAATCCTGGACGGACCACCGGTGGCGAAAGCGCCTCGAGAGGACGGATCCGACGGTGAGGGACGAAAGCTAGGGTCA |
| p__Thermoplasmatota_g__SM1-50_ASV_4369 | 1 | 0 | 0 | 0 | 0 | 0 | 0 | 0 | 0 | 0 | 0 | CCTTCACACTGTGCGAAAGCGCGATGAGGGGATTCCAAGTGCTAACTCGAAAGAGTTGGCTGTTTATCTGTCTAAAACGCAGGTAGAGTAAGAGCTGGGTAAGACGGCTGCCAGCCGCCGCGGTAATAGCTGCAGCTCGAGTGGTGACCACTTTTATTGGGCCTAAAACGTCCGTAGCCGGTCTAGTAAATGCCTGGGTAAATCGTGCCGCTTAACGGTACGAATTCCGGGTAGACTGCTAGACTTGAGATCGGGAGAGGTTAGAGGTACTCCTTGGGTAGAGGTGAAATTCTGTAATCCCTGAAGGACCACCGGTGGCGAAGGCGCTCTCCCAGAACGGGTCCGACGGTGAGGGACGAAAGCCAGGGGAG |
| p__Thermoplasmatota_g__SM1-50_ASV_4379 | 1 | 0 | 0 | 0 | 0 | 0 | 0 | 0 | 0 | 0 | 0 | CCTTCACACTGTGCGAAAGCGCGATGAGGGGATCCCAAGTGCTTGCACGTAAGTGTAAGCTGTTTCTTTGTTTAAAACGCAGAGAGAGGAAGGGCTGGGTAAGACGGGTGCCAGCCGCCGCGGTAATACCTGCAGCTCGAGTGGTGACCGCTATTATTGGGCCTAAAACGTCCGTAGCCGGCTGAGTAAATGCTTGGGTAAATCGTGCCGCTTAACGGTACGAATTCCGGGTAAACTGCTCAGCTTGAGATCGGGAGAAGCTAGAGGTACTCCTGGGGTAGAGGTGAAATTCTGTAATCCTGGGGGGACCACCAGTGGCGAAAGCGTGGGGAG |
| p__Thermoplasmatota_g__SM1-50_ASV_4391 | 0 | 0 | 0 | 0 | 0 | 0 | 0 | 0 | 0 | 1 | 0 | ACTTCGCACTGCGCGAAAGCGCGACGAGGGAATCCCAAGTGTATGCACGTAAGTGTATACTGTTTTTCTGTCTAAAAAGCAGAGAGAGTAAGAGCTGGGTAAGACGGGTGCCAGCCGCCGCGGTAATACCTGCAGCTCAAGTGGTGATCGTTATTATTGGGCCTAAAACGTCCGTAGCCGGTCTTGTAAATGCCTGGGTAAATCGTATGGCTTAACCATACGAATTCCGGGTAGACTGCAGGGCTAGAGATCGGGAGAGGCTAGAGGTACTCCTGGGGTAGAGGTGAAATTCTGTAATCCTAGGGGGACCACCAGTGGCGAAGGCGTCTAACTAGAACGGGTCTGACGGTGAGGGACGAAGCCCTGGGGCG |
| p__Thermoplasmatota_g__SM1-50_ASV_4406 | 0 | 0 | 0 | 0 | 0 | 0 | 0 | 0 | 0 | 1 | 0 | ACTTCGCACTGTGCGCAAGCACGACGAGGGAATCCCAAGTGCCAGCACGTTAGTGTTGGCTGTTTTTCTGTCTAAAAAGCAGAGAGAGTAAGGGCTGGGTAAGACGGGTGCCAGCCGCCGCGGTAATACCTGCAGCCCAAGTGGTGGCCGCTATTATTGGGCCTAAAACGTCCGTAGCCGGTCTATTAAATGCCTGGGTAAATTGTGTGGCTCAACCATACGAATTCCGGGTAGACTGGTAGACTTGAGATCGGGAGAGGCTAGAGGTACTCCTGGGGTAGAGGTGAAATTCTGTAATCCTAGGGGGACCACCAGTGGCGAAGGCGTCTAGCTAGAACGAGTCTGACGGTGAGGGGCGAAGCCCTGGGGCG |

| Phylum_Genus_ASV_number | Presence (1) or Absence (0) in each layer L1 | L2 | L3 | L4 | L5 | L6 | L7 | L8 | L9 | L10 | L11 | Sequence |
|---|---|---|---|---|---|---|---|---|---|---|---|---|
| p__Thermoplasmatota_g__SM1-50_ASV_4564 | 0 | 0 | 0 | 0 | 0 | 0 | 0 | 0 | 0 | 0 | 1 | CCTTCACACTGTGCGAAAGCGCGATGAGGGGATCCCAAGTGCTAACTCGAAAGAGTTGGCTGTTTATCTGTCTAAAACGCAGGTAGAGTAAGAGCTGGGTAAGACGGCTGCCAGCCGCCGCGGTAATACCTGCAGCTCAAGTGGTGGCCGTTATTATTGGGCCTAAAACGTTCGTAGCCGGTTTATTAAATGCCTGGGTAAATCGGACCGCTTAACGGTTCGAATTCCGGGTAGACTGATAAACTTGAGACCGGGAGAGGTTAGAGGTACTCCTGGGGTAGAGGTGAAATTCTGTAATCCTAGGGGGACCACCAGTGGCGAAGGCGTCTAACTAGAACGGGTCTGACGGTGAGGGACGAAGCCCTGGGGCG |
| p__Thermoplasmatota_g__SM1-50_ASV_4581 | 0 | 0 | 0 | 0 | 0 | 0 | 0 | 0 | 0 | 0 | 1 | CCTTTACAATGCACGAAAGTGTGATAAGGGGATTCTAAGTGCTCATGCAAAGCATGGGCTTTTGCCAAGGGTAAATACCTTGGCGAATAAGGGGTGGGCAAGACTGGTGCCAGCCGCCGCGGTAATACCTGCAGCCCGAGTGGTGACTGTTATTATTGGGCCTAAAACGTCCGTAGCCGGTTCTGTAAATGCCTGGGTAAATCGTACGGCTCAACCGTACGAATTCCGGGTAGACTGCAGGGCTAGAGATCGGGAGAGGCTAGAGGTACTCTTGGGGTAGAGGTGAAATTCTGTAATCCTAGGGGGACCACCAGTGGCGAAGGCGTCTAGCTAGAACGAGTCTGACGGTCAGGGACGAAGCCCTGGGGCG |
| p__Thermoplasmatota_g__SM1-50_ASV_4592 | 0 | 0 | 0 | 0 | 0 | 0 | 0 | 0 | 1 | 0 | 1 | ACTTCGCACTGTGCGCAAGCACGACGAGGGAATCCCAAGTGCCAGCACGTTAGTGTTGGCTGTTTTTCTGTCTAAAAAGCAGAGAGAGTAAGGGCTGGGTAAGACGGGTGCCAGCCGCCGCGGTAATACCTGCAGCCCAAGTGGTGGCCGCTATTATTGGGCCTAAAACGTCCGTAGCCGGTCTATTAAATGCCTGGGTAAATCGTGCCGCTTAACGGTACGAATTCCGGGTAGACTGATAGACTTGAGATCGGGAGAGGCTAGAGGTACTCCTGGGGTAGAGGTGAAATTCTGTAATCCTGGGGGGACCACCAGTGGCGAAGGCGTCTAGCTAGAACGAGTCTGACGGTGAGGGACGAAGCCCTGGGGCG |
| p__Thermoplasmatota_g__SM1-50_ASV_5120 | 0 | 0 | 0 | 0 | 0 | 0 | 0 | 0 | 1 | 0 | 0 | CCTTCACACTGTGCGAAAGCGCGATGAGGGGATCCCAAGTGCTGGCTCGTTAGAGTCAGCTGTTTCTTTGTCTAAAAAACAGAGAGAGTAAGGGCTGGGTAAGACGGGTGCCAGCCGCCGCGGTAATACCTGCAGCTCAAGTGGTGGCCGCTATTATTGGGCCTAAAACGTTCGTAGCCTGTTTATTAAATGCCTAGGTAAATCGGGCCGCTTAACGGTTCGAATTCTGGGTAGACTGATAAACTCGAGACTGGGAGAGGCTAGAGGTACTCCTGGGGTAGAGGTGAAATTCTGTAATCCTAGGGGGACCACCAGTGGCGAAGGCGTCTAGCTAGAACGGGTCTGACGGTGAGGGACGAAGCCCTGGGGCG |
| p__Thermoplasmatota_g__SM1-50_ASV_5156 | 0 | 0 | 0 | 0 | 0 | 0 | 0 | 0 | 1 | 0 | 0 | CTTTTACAATGCACGAAAGTGTGATAGAGCGAGCCAGAGTGTTTTTCTGTATAAGAAAAACTTTTGCCAAATGTAAAAAATTTGGCGAATAAGGACTGGGCAAGACCGGTGCCAGCCGCCGCGGTAATCCCGGCAGTCCAAGTTGTAGCCACATTTATTGGGTCTAAAACGTTCGTAGCCGGTTTAGTAAATGCCTGGGTAAATCGTGTAGCTTAACTATACGAATTCCGGGTAGACTGCTAAACTTGAGACCGGGAGAAGTTAGAGGTACTCCTGGGGTAGAGGTGAAATTCTGTAATCCTAGGGGGACCACCAGTGGCGAAGGCGTCTAACTAGAACGGGTCTGACGGTGAGGGACGAAGCCCTGGGGCG |
| p__Thermoplasmatota_g__SM1-50_ASV_5158 | 0 | 0 | 0 | 0 | 0 | 0 | 0 | 0 | 1 | 0 | 0 | TCTTCACAATGTGCGAGAGCACGATGAGGGAATCCCAAGTGCTAGCACGTTAGTGTTAGCTGTTTTCTAGTCTAAAAAACTAGGAGAGTAAGAGCTGGGTAAGACGGGTGCCAGCCGCCGCGGTAATACCTGCAGCTCAAGTGGTGATCGTTATTATTGGGCCTAAAACGTCCGTAGCCGGTCTTGTAAATGCCTGGGTAAATCGTATGGCTTAACCATACGAATTCCGGGTAGACTGTTAGACTTGAGACCGGGAGAGGCTAGAGGTACTCCTGGGGTAGAGGTGAAATTCTGTAATCCTGGGGGGACCACCAGTGGCGAAGGCGTCTAGCCAGAACGGGTCTGACGGTGAGGGACGAAGCCCTGGGGCG |
| p__Nanoarchaeota_g__SM23-78_ASV_4559 | 0 | 0 | 0 | 0 | 0 | 0 | 0 | 0 | 0 | 0 | 1 | CCTTTACAATGCACGAAAGTGTGATAAGGGAATCCTGAGTGCTCATGCTTTGCATGGGCTTTTGTTAAGAGTAAATATCTTAGCGAATAAGTGGTGGGTAAGACTGGTGCCAGCCGCCGCGGTAACACCAGCGCCACAAGTGGGGACCAGCATTATTGGGCCTAAAGCGTCCGTAGCAGGAAAGATAAATCTTCCGTGAAACTGTTCAGCTTAACTGAACAACGTGCAGAAGACACTATTTTTCTAGAGACCGGGAGGAGTCAGAGGTATTTCTGGGGGAGCGGTAAAATGCTATAATCCCAGGAGGACCACCGATGGCGAAGGCATCTGACTAGAACGGATCTGACTGTGAGGGACGAAGGCCAGGGGAG |
| p__Nanoarchaeota_g__SM23-78_ASV_4593 | 0 | 0 | 0 | 0 | 0 | 0 | 1 | 0 | 0 | 0 | 1 | CCTTTACAATGCACGCAAGTGTGATAAGGGGATCCCAAGTGCTCAAGCATTGCTTGGGCTTTTGCCATGTGTAAATAGCATGGCGAATAAGTGGTGGGAAAGACTGGTGCCAGCCGCCGCGGTAACACCAGCGCCACGAGTGGCAATCGCGTTTATTGGGCCTAAAGCGTCCGTAGCAGGTTTCGTAAATCTCTTGTGAAATTGCCGGGCTCAATCCGGCAGACTGCAGGAGACACTGCGAGACTTGAGACCGGGAGGAGTTAGAGGTATTCTTTAGGGAGCGGTAAAATGCTATAATCTAAGGAGGACCACCTGTGGCGAAGGCGTCTAACTAGAACGGATCTGACTGTGAGGGACGAAAGCTAGGGGAG |
| p__Nanoarchaeota_g__UBA583_ASV_4356 | 1 | 0 | 0 | 0 | 0 | 0 | 0 | 0 | 0 | 0 | 0 | CCTCTACAATGCACGCAAGTGTGATAGGGTAACCCCTCGTGCTTATCGAAAGATAGGCTTTTGTTGAGTTTAGTGCACTCATCGAATAAGTGGTGGGCAAGACCGGTGCCAGCAGCCGCGGTAATACCGGCGCCACAAGTGGAGATCATGATTACTGGGTCTAAAGTGTTCGTAGCTGGCTTTGAAAGTTTCTTGTGAAAGCTCCGTGGCTTAACTCGGAAATTGCAAGAAATACTCCAAAGCTTGAGACCGGAAGAGGTAAGAGGTATTGATTGGGTAGCGGTAAAATGCTATAATCCTTTCAAGACCAACTAAGGCGAAGGCATCTTACTTGGACGGATCTGACAGTGAGGAACGAGAGCTGGGGGAG |
| p__Halobacteriota_f__Haloarculaceae_3992_ASV_4933 | 0 | 0 | 0 | 0 | 0 | 1 | 0 | 0 | 0 | 0 | 0 | CCTTTACACTGCACGCCAGTGCGATAGGGGGACTCCGAGTGCGAGGGCATACAGTCCTCGCTTTTCTCGACCGTAAGAAGGTCGAAGAATAAGGGCTGGGCAAGACCGGTGCCAGCCGCCGCGGTAATACCGGCAGCCCAAGTGATGGCCGATATTATTGGGCCTAAAGCGTCCGTAGCCGGCCGGACAAGTCCGTTGGGAAATCCACGCGCCTAACGCGTGGGCGTCCAGCGGAAACTGTCCGGCTTGGGGCCGGAAGACTCGAGGGGTACGTCCGGGGTAGGAGTGAAATCCCGTAATCCTGGACGGACCGCCGGTGGCGAAAGCGCCTCGAGAGGACGGACCCGACGGTGAGGGACGAAAGCTAGGGTCT |
| p__Halobacteriota_f__Haloarculaceae_3992_ASV_4992 | 0 | 0 | 0 | 0 | 0 | 0 | 1 | 0 | 0 | 0 | 0 | CCTTTACACTGCACGCCAGTGCGATAAGGGGACTCCGAGTGCGAGGGCATATCGTCCTCGCTTTTCTGTACCGTAAGGTGGTACAGGAACAAGGGCTGGGCAAGACCGGTGCCAGCCGCCGCGGTAATACCGGCAGCCCGAGTAATGGCCGCTCTTATTGGGCCTAAAGCGTCCGTAGCCGGCCGGACAAGTCCGTTGGGAAATCCACGCGCTCAACGCGTGGGCGTCCAGCGGAAACTGTCTGGCTTGGGGCCGGAAGACCTGGGGGGTACGTCCGGGGTAGGAGTGAAATCCTGTAATCCTGGACGGACCACCGATGGCGAAAGCACCTCAGGAAGACGGACCCGACGGTGAGGGACGAAAGCTAGGGTCT |
| p__Halobacteriota_f__Haloferacaceae_ASV_4656 | 0 | 1 | 0 | 0 | 0 | 0 | 0 | 0 | 0 | 0 | 0 | CCTTTACACTGCACGCCAGTGCGATAAGGGGACCCCGAGTGCGAGGGCATACAGTCCTCGCTTTTGTGTACCGTAGGGAGGTACACGAATAAGGGCTGGGCAAGACCGGTGCCAGCCGCCGCGGTAATACCGGCAGCTCAAGTGATGGCCGCTATTATTGGGCCTAAAGCGTCCGTAGCCGGCCGCACAAGTCCGTCGGGAAATCCACCCGCCTAACGGGTGGGCGTCCGGCGGAAACTGTGCGGCTTGGGACCGGAAGACTCGAGGGGTACGTCCGGGGTAGGAGTGAAATCCCGTAATCCCGGACGGACCACCGATGGCGAAAGCACGGTGGGAAGACGGCTCCGACAGTGAGGGACGAAAGCTGGGGTCT |
| p__Halobacteriota_f__Natrialbaceae_ASV_4677 | 0 | 1 | 1 | 0 | 0 | 0 | 0 | 0 | 0 | 0 | 0 | CCTTTACACTGCACGCCAGTGCGATAAGGGGACTCCAAGTGCGAGGGCATACAGTCCTCGCTTTTTCCTACCGTAAGGTGGTAGGCGAATAAGGGCTGGGCAAGACCGGTGCCAGCCGCCGCGGTAATACCGGCAGCCCAAGTGATGACCGCTATTATTGGGCCTAAAGCGTCCGTAGCCGGCCGGGTAAGTCCGTCGGGAAATCCGCGCGCTCAACGCGCGGGCGTCCGGTGGAAACTGCCTGGCTTGGGACCGGAAGATCCAGAGGGTACGTCCGGGGTAGGAGTGAAATCCTGTAATCCTGGACGGACCACCGGTGGCGAAAGCGCTCTGGAAGGACGGATCCGACGGTGAGGGACGAAAGCTAGGGTCA |
| p__Halobacteriota_f__Natrialbaceae_ASV_4683 | 0 | 0 | 1 | 0 | 0 | 0 | 0 | 0 | 0 | 0 | 0 | CCTTTACAATGCGCGAAAGCGCGATAAGGGGAGTCCAAGTGGTCGTTCCTCACAAAGGAGCGACCTTTTGCCAAGTGTAAGGAGCTTGGTGAATAAGGGGTGGGAAAGACGGCTGCCAGCCGCCGCGGTAATAGCCGCACCCCAAGTGGTGACCACGATTATTGGGCTTAAAGAGTCCGTAGCCGGCCACGCAAGTCTGCTGTGAAATCTGGCTGCTTAACAGTCAGGAGGGCAGCAGAAACTCCGTGGCTTGGGACCGGACGGCGTTAGGGCTATTCCAGGAGTAGGAGTGAAATCCTGAAATTCCTGGAGGAGCACCGGTGGCGAAAGCGCCTAACGAGGACGGATCCGACGGTGAGGGACGAAAGCTAGGGTCT |
| p__Halobacteriota_f__Natrialbaceae_ASV_4711 | 0 | 0 | 1 | 0 | 0 | 0 | 0 | 0 | 0 | 0 | 0 | CCTTTACACTGCACGCCAGTGCGATAGGGGGACTCCAAGTGCCAGGGCATATCGTCCTGGCTTTTCACGACCGTAAGGTGGTAGTGGAATAAGGGCTGGGCAAGACCGGTGCCAGCCGCCGCGGTAATACCGGCAGCCCGAGTGATGGCCGCTCTTATTGGGCCTAAAGCGTCCGTAGCTGGCCTCGCAAGTCCGTCGGGAAATCCGCACGCTTAACGTGCGGGCGTCCGGTGGAAACTGCGTGGCTTGGGACCGGAAGACCCGAGGGGTACGTCCGGGGTAGGAGTGAAATCCTGTAATCCTGGACGGACCACCGGTGGCGAAAGCGCCTCGGGAGGACGGATCCGACGGTGAGGGACGAAAGCTAGGGTCA |
| p__Halobacteriota_f__Natrialbaceae_ASV_4722 | 0 | 0 | 1 | 0 | 0 | 0 | 0 | 0 | 0 | 0 | 0 | CCTTTACACTGCACGACAGTGCGATAGGGGGATTCCGAGTGCGAGGGCATACAGTCCTCGCTTTTCTGTACCGTAAGGAGGTACAGGAACAAGTGCTGGGCAAGACCGGTGCCAGCCGCCGCGGTAATACCGGCAGCACGAGTGATGGCCGATCTTATTGGGCCTAAAGCGTCCGTAGCTCGCCGGACAAGTCCGTTGGGAAATCCACGTGCCCAACACGTGGGCGTCCAGCGGAAACTGTCTGGCTTGGGACCGGAAGATCCAGAGGGTATGTCCGGGGTAGGAGTGAAATCCTGTAATCCTGGACGGACCACCGGTGGCGAAAGCGCTCTGGAAAGACGGATCCGACGGTGAGGGACGAAAGCTAGGGTCA |
| p__Halobacteriota_f__Natrialbaceae_ASV_4757 | 0 | 0 | 1 | 0 | 0 | 0 | 0 | 0 | 0 | 0 | 0 | CCTTTACACTGCACGACAGTGCGATAGGGGGACTCCGAGTGCGAGGGCATATAGTCCTCGCTTTTCTGGACCGTAAGGAGGTCCAGGAATAAGGGCTGGGCAAGACCGGTGCCAGCCGCCGCGGTAATACCGGCAGCCCGAGTGATGGCCGCTATTATTGGGCCTAAAGCGTCCGTAGCCGGCCGGGTAAGTCCGTCGGGAAATCCGCGCGCTCAACGCGCGGGCGTCCGGTGGAAACTGCCTGGCTTGGGACCGGAAGATCCAGAGGGTACGTCCGGGGTAGGAGTGAAATCCTGTAATCCTGGACGGACCACCGGTGGCGAAAGCGCTCTGGAAGGACGGATCCGACGGTGAGGGACGAAAGCTAGGGTCA |
| p__Halobacteriota_f__Natrialbaceae_ASV_4764 | 0 | 0 | 1 | 0 | 0 | 0 | 0 | 0 | 0 | 0 | 0 | CCTTTACACTGCACGACAGTGCGATAGGGGGACTCCAAGTGCGAGGGCATATAGTCCTCGCTTTTGTGTACCGTAAGGAGGTACAGGAATAAGGGCTGGGCAAGACCGGTGCCAGCCGCCGCGGTAATACCGGCAGCCCGAGTGATGACCGCTATTATTGGGCCTAAAGCGTCCGTAGCCGGCCACACAAGTCCGTCGGGAAATCCGCACGCCCAACGTGCGGGCGTCCGGCTGAAACTGTGTGGCTTGGGACCGGAGGACCTGGGGGGTACGTCTGGGGTAGGAGTGAAATCCCGTAATCCTGGACGGACCACCGGTGGCGAAAGCGCCTCAGGAAGACGGATCCGACGGTGAGGGACGAAACCCTGGGGCG |
| p__Halobacteriota_f__Natrialbaceae_ASV_4766 | 0 | 0 | 1 | 0 | 0 | 0 | 0 | 0 | 0 | 0 | 0 | CCTTTACACTGCACGCACGTGCGATAAGGGGACACCGAGTGCGAGGGCATACAGTCCTCGCTTTTGTACACCGTAAGGAGGTGTACGAATAAGGGCTGGGCAAGACCGGTGCCAGCCGCCGCGGTAATACCGGCAGCCCGAGTGATGGCCGCTATTATTGGGCCTAAAGCGTCCGTAGCCGGCCGGACAAGTCCATTGGGAAATCCACGCGCCTAACGTGTGGGCGACCGGTGGAAACTGTCCGGCTTGGGACCGGAAGACCCGAGGGGTACGTCCGGGGTAGGAGTGAAATCCCGTAATCCTGGACGGACCGCCGGTGGCGAAAGCGCCTCGGGAGGACGGATCCGACGGTGAGGGACGAAAGCTAGGGTCT |
| p__Halobacteriota_f__Natrialbaceae_ASV_4833 | 0 | 0 | 0 | 1 | 0 | 0 | 0 | 0 | 0 | 0 | 0 | CCTTTACACTGCACGAGAGTGCGATAAGGGGACTCCAAGTGCGAGGGCATACAGTCCTCGCTTTTCACGACCGTAAGGTGGTCGTGGAATAAGGGCTGGGCAAGACCGGTGCCAGCCGCCGCGGTAATACCGGCAGCCCAAGTGATGGCCGCTATTATTGGGCCTAAAGCGTCCGTAGCCGGCCGAACAAGTCCGTCGGGAAATCCGCACGCCTAACGTGCGGGCGTCCGGCGGAAACTGTTCGGCTTGGGACCGGAAGATCCAGAGGGTACGTCTGGGGTAGGAGTGAAATCCCGTAATCCTGGACGGACCACCGGTGGCGAAAGCGCTCTGGAAGGACGGATCCGACGGTGAGGGACGAAAGCTAGGGTCA |
| p__Halobacteriota_f__Natrialbaceae_ASV_4988 | 0 | 0 | 0 | 0 | 0 | 0 | 1 | 0 | 0 | 0 | 0 | CCTTCACACTGTGCGAAAGCGCGATGAGGGGATCCCAAGTGCTTGCTCTTAGGAGTAAGCTGTTTTCCTGTTTAAATCGCAGGAAGAGGAAGGGCTGGGTAAGACGGGTGCCAGCCGCCGCGGTAATACCTGCAGCTCAAGTGGTGACCGCTATTATTGGGCCTAAAGCGTCCGTAGCCGGCCACGCAAGTCCGTCGGGAAATCCGCGCGCCTAACGCGCGGGCGTCCGGCGGAAACTGCGTGGCTTGGGACCGGAAGACTCGAGGGGTACGTCCGGGGTAGGAGTGAAATCCTGTAATCCTGGACGGACCACCGGTGGCGAAGGCGGCTCTCTATTCCGATACTGACGCCGAAGTGCGAAAGCCAGGGGAG |
| p__Halobacteriota_f__Natrialbaceae_ASV_5172 | 0 | 0 | 0 | 0 | 0 | 0 | 0 | 0 | 1 | 0 | 0 | CCTTTACACTGCACGCCAGTGCGATAAGGGGACTCCAACTGCCAGGGCATATCGTCCTGGCTTTTGTTTACCGTAAGGAGGTAAACGAATAAGTGCTGGGCAAGACCGGTGCCAGCCGCCGCGGTAATACCGGCAGCACAAGTGATGACCGCTATTATTGGGCCTAAAGCGTCCGTAGCTGGCCGGACAAGTCCGTCGGGAAATCTCTCCGCCTAACGGAGAGGCGTCCGGTGGAAACTGTTCGGCTTGGGACCGGAAGATCCAGAGGGTACGTCCGGGGTAGGAGTGAAATCCTGTAATCCTGGACGGACCACCGGTGGCGAAAGCGCTCTGGAAGGACGGATCCGACGGTGAGGGACGAAAGCTAGGGTCA |
| p__Methanobacteriota_B_f__Thermococcaceae_ASV_4829 | 0 | 0 | 0 | 1 | 0 | 0 | 0 | 0 | 0 | 0 | 0 | ACTGCACAATGCGGGAAACCGCGATGCGGGAACTCCAAGTGCCAGCACAGAGTGTTGGCTGTTCCTTAGCCTAAAAAGCTCTGGGAGTAAGGGCTGGGTAGACGGGTGCCAGCCGCCGCGGTAATACCTGCAGCCCGAGTGGTGGCCGCTATTATTGGGCCTAAAGCGTCCGTAGCCTGCTCGGTAAATGCCTGTGTAAATTGTACCGCTTAACGGTGCGAAGTGCGGGTAGACTGCCGGGCTTGGGACCGGGAGAGGCGCGAAGTACTCCGGGGGTAGAGGTGAAATTCTGTAATCCTGGGGGGACCACCAGTGGCGAAAGCGTCTAGCTAGAACGGGTCTGACGGTGAGGGACGAAACCCTGGGGCG |
| p__Halobacteriota_o__Halanaerobiales_ASV_4401 | 0 | 0 | 0 | 0 | 0 | 0 | 0 | 0 | 0 | 1 | 0 | ACTTCACAATGCGCGAAAGCGCGATGAGGGGATCCCAAGTGCTTGCACGTAAGTGTAAGCTGTTTTTCTGTTTAAAAAGCAGAGAGAGTAAGGGCTGGGTAAGACGGGTGCCAGCCGCCGCGGTAATACCTGCAGCTCAAGTGGTGACCGCTATTATTGGGCCTAAAACGTCCGTAGCCGGCTAAGTAAATGCCTGGGTAAATCGTGCCGCTTAACGGTACGAATTCCGGGTAGACTGCTTAGCTTGGGACCGGAAGGCTCGAGGGGTACGTCCGGGGTAGGAGTGAAATCCTGTAATCCTGGACGGACCACCGATGGCGAAAGCGTCTAGCTAGAACGGGTCTGACGGTGAGGGACGAAACCCTGGGGCG |
| p__Halobacteriota_o__Halanaerobiales_ASV_4599 | 0 | 1 | 0 | 0 | 0 | 0 | 0 | 0 | 0 | 0 | 0 | CCTTTACACTGCACGACAGTGCGATAGGGGGACCCCGAGTGCGAGGGCATACAGTCCTCGCTTTTGCACACCGTAAGGTGGTGCGCGAATAAGTGCTGGGCAAGACCGGTGCCAGCCGCCGCGGTAATACCGGCAGCACAAGTGATGGCCAATATTATTGGGCCTAAAGCGTCCGTAGCCGGCCGAACAAGTCCGTCGGGAAATCCATCCGCCTAACGGATGGGCGTCCGGCGGAAACTGTTCGGCTTGGGACAGGAAGACTCGAGGGGTACGTCCGGGGTAGGAGTGAAATCCCGTAATCCTGGACGGACCGCCGATGGCGAAAGCACCTCGAGAAGACTGATCCGACGGTGAGGGACGAAAGCTGGGGTCT |

| | Presence (1) or Absence (0) in each layer | | | | | | | | | | | |
|---|---|---|---|---|---|---|---|---|---|---|---|---|
| **Phylum_Genus_ASV_number** | **L1** | **L2** | **L3** | **L4** | **L5** | **L6** | **L7** | **L8** | **L9** | **L10** | **L11** | **Sequence** |
| p__Halobacteriota_o__Halanaerobiales_ASV_4658 | 0 | 1 | 0 | 0 | 0 | 0 | 0 | 0 | 0 | 0 | 0 | CCTTTACACTGCACGGAAGTGCGATAAGGGGACCCCAAGTGCGAGGGCATACAGTCCTCGCTTTTCCTGACTGTAGGGCGGTCAGGGAACAAGAGCTGGGCAAGACCGGTGCCAGCCGCCGCGGTAATACCGGCAGCTCGAGTGATGGCCGATATTATTGGGCCTAAAGCGTCCGTAGCCGGCCGGTCAAGTCCGTTGGGAAATCCACCAGCTCAACTGGTGGGCGTCCAGCGGAAACTGATCGGCTTGAGGCCGGAGGACCCGAGGGGTACGTCCGGGGTAGGAGTGAAATCCTGTAATCCTGGACGGACCACCGGTGGCGAAAGCGCCTCGAGAGGACGGATCCGACGGTGAGGGACGAAAGCTAGGGTCA |
| p__Halobacteriota_o__Halanaerobiales_ASV_4691 | 0 | 0 | 1 | 0 | 0 | 0 | 0 | 0 | 0 | 0 | 0 | CCTTTACACTGCACGACAGTGCGATAAGGGGACTCCAAGTGCGAGGGCATATAGTCCTCGCTTTTGTGTACCGTAAGGTGGTACACGAATAAGGGCTGGGCAAGACCGGTGCCAGCCGCCGCGGTAATACCGGCAGCCCGAGTGATGACCGCTATTATTGGGCCTAAAGCGTCCGTAGCTGGCCGATCAAGTCCGTCGGGAAATCCGCGCGCCCAACGCGCGGGCGTCCGGCGGAAACTGTTCGGCTTGGGACCGGAAGACCCGAGGGGTACGTCCAGGGTAGGAGTGAAATCCTGTAATCCTGGACGGACCGCCGGTGGCGAAAGCGCCTCGGGAGGACGGATCCGACGGTGAGGGACGAAAGCTAGGGTCA |
| p__Halobacteriota_o__Halanaerobiales_ASV_4730 | 0 | 0 | 1 | 0 | 0 | 0 | 0 | 0 | 0 | 0 | 0 | CCTTTACACTGCACGACAGTGCGATAAGGGGACTCCGAGTGCGAGGGCATAGAGCCCTCGCTTTTCTGCACCGTAAGGTGGTGCAGGAAGAAGGGCTGGGCAAGACCGGTGCCAGCCGCCGCGGTAATACCGGCAGCCCAAGTGATGGCCGATATTATTGGGCCTAAAGCGTCCGTAGCCGGCCGGACAAGTCCGTTGGGAAATCCGTTCGCTTAACGAACGGGCGTCCAGCGGAAACTGTCCGGCTTGGGACCGGAAGACCCGAGGGGTACGTCCGGGGTAGGAGTGAAATCCCGTAATCCCGGACGGACCACCGATGGCGAAAGCACTCTGAGAAGACGGATTCGACAGTGAGGGACGAAAGCTAGGGTCT |
| p__Halobacteriota_o__Halanaerobiales_ASV_4781 | 0 | 0 | 1 | 0 | 0 | 0 | 0 | 0 | 0 | 0 | 0 | CCTTTACACTGTGCGCAAGCGCGATAAGGGGACTCCGAGTGCGAGGGCATATCGTCCTCGCTTTTCTGGATCGTAAGGTGGTACAGGAATAAGGGCTGGGCAAGACCGGTGCCAGCCGCCGCGGTAATACCGGCAGCCCAAGTGATGGCCGTTCTTATTGGGCCTAAAGCGTCCGTAGCTGGCCGGGCAAGTCTGTTGGGAAATCCACTCGCTTAACGAGTGGGCGTCCAGCGGAAACTGCTCGGCTTGGGACCGGAAGACCCGAGGGGTACGTCCGGGGTAGGAGTGAAATCCTGTAATCCTGGACGGACCACCGATGGCGAAAGCACCTTGGAAACACGGACCCGACGGTGAGGAACGAAAGCTAGGGTCT |
| p__Halobacteriota_o__Halanaerobiales_ASV_4785 | 0 | 0 | 1 | 0 | 0 | 0 | 0 | 0 | 0 | 0 | 0 | CCTTTACACTGCACGACAGTGCGATAGGGGGACTCCGAGTGCGAGGGCATATAGTCCTCGCTTTTCTGTACCGTAAGGAGGTCGTGGAATAAGAGCTGGGCGAGACCGGTGCCAGCCGCCGCGGTAATACCGGCAGCTCGAGTGATGACCGCTTTTATTGGGCCTAAAGCGTCCGTAGCCGGCCGAACAGGTCCGTTGGGAAATCCGCCAGCCTAACTGGCGGGCGTCCAGCGGAAACCAGTCGGCTTGGGGCCGGAAGACCTGAGGGGTACGTCCGGGGTAGGAGTGAAATCCTGTAATCCTGGACGGACCGCCGGTGGCGAAAGCGCCTCAGGAGGACGGACCCGACGGTGAGGGACGAAAGCTAGGGTCA |
| p__Halobacteriota_o__Halanaerobiales_ASV_4793 | 0 | 0 | 0 | 1 | 0 | 0 | 0 | 0 | 0 | 0 | 0 | CCTTTACACTGCACGCGAGTGCGATAAGGGGACTCCAAGTGCGAGGGCATACAGTCCTCGCTTTTCTCGACCGTAAGGTGGTCGACGAATAAGGGCTGGGCAAGACCGGTGCCAGCCGCCGCGGTAATACCGGCAGCCCGAGTGATGACCGCTATTATTGGGCCTAAAGCGTCCGTAGCCGGCCAGGCAAGTCCGTCGGGAAATCCATCCGCTCAACGGATGGACGTCCGGCGGAACCTCCCTGGCTTGGGACCGGAAGACCCGACGGGTACGTCCGGGGTAGGAGTGAAATCCCGTAATCCTGGACGGACCACCGATGGCGAAAGCACGTCGGGAGGACGGCTCCGACGGTGAGGGACGAAAGCTAGGGTCT |
| p__Halobacteriota_o__Halanaerobiales_ASV_4796 | 0 | 0 | 0 | 1 | 0 | 0 | 0 | 0 | 0 | 0 | 0 | CCTTTACACTGCACGACAGTGCGATAGGGGGACTCCAAGTGCGAGGGCATATAGTCCTCGCTTTTCACCACCGTAAGGTGGTGGTGGAATAAGTGCTGGGCAAGACCGGTGCCAGCCGCCGCGGTAATACCGGCAGCACGAGTGATGACCGCTATTATTGGGCCTAAAGCGTCCGTAGCCGGCCGAACAAGTCCGTCGGGAAATCCATCCGCTCAACGGATGGGCGTCCGGCGGAAACTGTTCGGCTTGGGACAGGAAGACTCGAGGGGTACGTCCGGGGTAGGAGTGAAATCCTGTAATCCTGGACGGACCGCCGATGGCGAAAGCACCTCGAGAAGACTGATCCGACGGTGAGGGACGAAAGCTGGGGTCT |
| p__Halobacteriota_o__Halanaerobiales_ASV_4808 | 0 | 0 | 0 | 1 | 0 | 0 | 0 | 0 | 0 | 0 | 0 | CCTTTACAATGTGCGCAAGCACGATAAGGGGATCCCAAGCGCGAGGGCATATCGTCCTCGCTTTTCTGGACCGTAAGGTGGTCCAGGAACAAGTGCTGGGCAAGACCGGTGCCAGCCGCCGCGGTAATACCGGCAGCACAAGTGATGGCCAATATTATTGGGCCTAAAGCGTCCGTAGCCGGCCGGATAAGTTCGTCGGGAAATCTACCCGCTTAACGGGTGGGCGTCCGGCGGAAACTGTTCGGCTTGGGGCCGGAAGGCCAGAGGGGTACGTCCGGGGTAGGAGTGAAATCCTGTAATCCCGGGCGGACCGCCGGTGGCGAAAGCGCCTCTGGAGGACGGACCCGACGGTGAGGGACGAAAGCTAGGGTCT |
| p__Halobacteriota_o__Halanaerobiales_ASV_4990 | 0 | 0 | 0 | 0 | 0 | 0 | 1 | 0 | 0 | 0 | 0 | CCTTTACACTGTGCGAAAGCGCGATAAGGGAACTCCCAGTGCGAGGGCATATCGTCCTCGCTTTTCTGTACCGTAAGGTGGTACAGGAACAAGGGCTGGGCAAGACCGGTGCCAGCCGCCGCGGTAATACCGGCAGCCCGAGTGATGGCCGCTATTATTGGGCCTAAAGCGTCCGTAGCCGGCCGGGCAAGTCCGTTGGGAAATCCACCCGCTTAACGGGTGGGCGACCGGCGGAAACTGCCTGGCTTGGGACCGGGAGATCCGAGGGGTACGTCCGGGGTAGGAGTGAAATCCTGTAATCCTGGACGGACCGCCGGTGGCGAAAGCGCCTCGGAAGAACGGATCCGACGGTGAGGGACGAAAGCCAGGGTCT |
| p__Thermoplasmatota_o__Methanomassiliicoccales_ASV_5096 | 0 | 0 | 0 | 0 | 0 | 0 | 0 | 0 | 1 | 0 | 0 | ACTTCGCAATGGGGGAAACCCTGACGAGGGGATTCCAAGTGCCAGCACACAGTGTTGGCTGTTCTTCTGTCCAAAACGCAGGAGAAGTAAGGGCCGGGTAAGACGGGTGCCAGCCGCCGCGGTAATACCCGCGGCCCGAGTGGTGGTCGATTTTATTGAGCCTAAAACGTCCGTAGCCTGCTTGGTAAATCCTTGGGTAAATCGGCCAGCTTAACTGTCCGAAGTCCGGGGAGACTGCCAGGCTTGGGACCGGGAGAGGTCAGAGGTACTTCTGGGGTAGGGGTAAAATCCTGTAATCCTGGACGGACTACCGATGGCGAAAGCACCTTGGAAGCACGGATCCGACAGTGAGGGACGAAAGCTGGGGTCT |
| p__Nanoarchaeota_o__Woesearchaeales_581_ASV_4500 | 0 | 0 | 0 | 0 | 0 | 0 | 0 | 0 | 1 | 1 | 0 | CCTTTACAATGCACGAAAGTGTGATAAGGTAACCCCAAGTGCCAGTACTTTGTATTGGCTTTTGCAAAGAGTAAATATCTTGGCGAATAAGTGGTGGGTAAGACTGGTGCCAGCCGCCGCGGTAACACCAGCGCCACGAGTGGCGTCCGCTTTTATTGGGCCTAAAGTGTCCGTAGCCGGTTTAGACAGTCTTCTGTGAAAGTGGTAAGCTTAACTTATCAAATTGCAGGAGATCCTACTAAACTAGAGACCGGGAGGAGTTAGAGGTATTCTGGGGGAAGCGGTAAAATGCGATAATCCTCGGAGGACCACCTGTGGCGAAGGCGTCTAACCAGAACGGGTCTGACGGTGAGGGACGAAGGCTAGGGGAG |
| p__Nanoarchaeota_o__Woesearchaeales_581_ASV_5005 | 0 | 0 | 0 | 0 | 0 | 0 | 1 | 0 | 0 | 0 | 0 | CCTTTACAATGCACGCAAGTGTGATAAGGGAATCCTAAGTGCTCATGCAATGCATGGACTTTTGCCAAGAGTAAATATCTTGGCGAATAAGTGGTGGGAAAGACCGGTGCCAGCCGCCGCGGTAACACCGGCGCCACGAGTGGCAACCGCGTTTATTGGGCCTAAAGCGTCCGTAGCCTGTGTGGTAAATCTCATGTGAAATCGTCAAGCTCAACTTGACGGCGAGCTTGGGACACTGCCACGCTCGAGACCGGGAGGGGTCAAAGGTATTCCAAGGGGAGCGGTAAAATGTGATAATCCTTGGAGGATCGCCTATGGCGAAGGCATTTGACTAGAACGGATCTGACGGTGAGGGACGAAGGCCAGGGGAG |
| p__Thermoplasmatota_c__Thermoplasmata_1773_ASV_4596 | 0 | 1 | 0 | 0 | 0 | 0 | 0 | 0 | 0 | 0 | 0 | ACTGCGCAATGTGAGAAATCACGACGCGGGAACACCAAGTGCCAGCACAACGTGTTGGCTGTTTTCTTGTCTAAAAAACTAGGAGAGTAAGGGCTGGGTAAGACGGGTGCCAGCCGCCGCGGTAATACCTGCAGCCCAAGTGGTGGTCGTTATTATTGGGCCTAAAGCGTCCGTAGCCGGTTTGGTAAATGCCTGTGTAAATCGAACGGCTTAACCGTTCGAACTGCGGGTAGACTGCCAGACTAGGGACCGGGAGAGGTGCGAAGTACTCCGGGGGTAGAGGTGAAATTCTGTAATCCTCGGGGGACTACCGGTGGCGAAGGCGTCGCACTAGAACGGCTCCGACGGTGAGGGACGAAGCCCTGGGGCG |
| p__Thermoplasmatota_p__Thermoplasmatota_ASV_4705 | 0 | 0 | 1 | 0 | 0 | 0 | 0 | 0 | 0 | 0 | 0 | CCTTCACACTGTGCGAAAGCGCGATGAGGGGATTCCAAGTGCTAACTCGAAAGAGTTGGCTGTTTATCTGTCTAAAACGCAGGTAGAGTAAGAGCTGGGTAAGACGGCTGCCAGCCGCCGCGGTAATAGCTGCAGCTCGAGTGGTGACCACTTTTATTGGGCCTAAAACGTCCGTAGCCGGTCTAGTAAATGCCTGGGTAAATCGTGCCGCTTAACGGTACGAATTCCGGGTAGACTGCTAGACTTGAGATCGGGAGAGGTTAGAGGTACTCCTTGGGTAGAGGTGAAATTCTGTAATCCTCGGGGGACTACCGGTGGCGAAGGCGTCGTGATAGAACGGCTCCGACGGTGAGGGACGAAGCCCTGGGGCG |
| d__Archaea_d__Archaea_ASV_4348 | 1 | 0 | 0 | 0 | 0 | 0 | 0 | 0 | 0 | 0 | 0 | CCTCCGCAATGCTCGAGAGAGTGACGGGGAGATTCCAAGTGCCTACGCACAGCGTAGGCTGTTCAGATGCCTACAACGCATCTGGAGGAAGGACTGGGTAAGACGGGTGCCAGCCGCCGCGGTAACACCCGCAGTTCAAGTGGCAGTCGATTTTATTGGGCCTAAAGCGTCCGTAGCCGGTCTGGCAAATGCCTGGGTAAATCGTCTGGCTCAACCAGACGAATTCCGGGTAGACTGCCAGACTTGGGACCGGTTGAGGGAGAGGGTACTCCAGGGGTAGGAGTGAAATCCTGTAATCCTTGGAGGACCACCAGTGGCGAAAGCGCTCTCCCAGGACGGGTCCGACGGTGAGGGACGAAAGCCAGGGTAG |
| d__Archaea_d__Archaea_ASV_4513 | 0 | 0 | 0 | 0 | 0 | 0 | 0 | 0 | 0 | 0 | 1 | ACTTCACAATGCGCGAAAGCGCGATGAGGGGATCCCAAGTGCTTGCACGTAAGTGTAAGCTGTTTTTCTGTTTAAAAAGCAGAGAGAGTAAGGGCTGGGTAAGACGGGTGCCAGCCGCCGCGGTAATACCTGCAGCTCAAGTGGTGACCGCTATTATTGGGCCTAAAACGTCCGTAGCCGGCTAAGTAAATGCCTGGGTAAATCGTGCCGCTTAACGGTACGAATTCCGGGTAGACTGCTTAGCTTGGGACCGGAAGAAGTTAGAGGTACTCCTGGGGTAGGGGTGAAATCCTGTAATCCCGAACGGACCACCGATGGCGAAAGCACTCTGAGAAGACGGATTCGACAGTGAGGGACGAAAGCTAGGGTCT |
| d__Archaea_d__Archaea_ASV_4594 | 0 | 1 | 0 | 0 | 0 | 0 | 0 | 0 | 0 | 0 | 0 | ACTTCGCAATGCTGGAAACAGCGACGAGGTGACTCCAAGTGCCGGCACAGAGTGTCGGCTGTTTCCAAGCCTAAAAAGCTTGGAGAGTAAGGGCTGGGTAAGACGGGTGCCAGCCGCCGCGGTAATACCTGCAGCCCAAGTGGTGGTCGCTATTATTGGGCCTAAAGCGTCCGTAGCCGGACCAGTAAATGCCTGTGTAAATTGCACGGCTCAACCGTGCGAACTGCGGGTAGACTGCTGGTCTTGGGACCGGGAGAGGTGCGAGGTACTCGGGGGGTAGAGGTGAAATTCTGTAATCCTCCGGGGACCACCGGTGGCGAAGGCGTCGCACTAGAACGGCTCCGACGGTGAGGGACGAAGCCCTGGGGCG |
| d__Archaea_d__Archaea_ASV_4608 | 0 | 1 | 0 | 0 | 0 | 0 | 0 | 0 | 0 | 0 | 0 | CCTGCGCAATGTGCGAAAGCGCGACGCGGGAACTTCAAGTGCCGTTGTTACACTAGTGATAACGGCTGTTCTTGTGTCTAAAAAACATTTGGAGTAAGGGCTGGGTAAGACCGGTGCCAGCCGCCGCGGTAATACCGGCAGCCCAAGTGGTGACCAACATTATTGGGTCTAAAACGTCCGTAGCCGGCCTGGTAAATCCCTCGTTAAATCGAACGGCTTAACCGTTCGGCCTCGTGGGAGACTGTCAGGCTTGGGACCGAGAGAGGTCAGAGGTACTCCAGGGGTAGGGGTGAAATCCTGTAATCCTTGGGGGACCACCAGTGGCGAAAGCGTCTGACTTGAGCGGGTCCGACGGTAAGGGACGAAAGCCAGGGTCG |
| d__Archaea_d__Archaea_ASV_4618 | 0 | 1 | 0 | 0 | 0 | 0 | 0 | 0 | 0 | 0 | 0 | CCTCCACAATGCACGAAAGTGTGATGGGGGGATCCTCCGTGCTCATGTTATGCATGGGCTTTTGCCAAGACTAGACCCCTTGGCGAATAAGGGGTGGGCAAGACCGGTGCCAGCCGCCGCGGTAACACCGGCGCCCCAAGTGGCAATCACGTTTATTGGGCCTAAAGCGTCCGTAGCCGGACTGATAAATCCGCTGTGAAATCGTCGTGCTCAACGCGACGGCGTGCAGTGGACACTATCAGTCTCGAGACCGGGAGGAGCAAGAGGTATGCCGTGGGGACTGGTAAAATGGTATAATCCGCGGTAGACCACCGATGGCGAAGGCATCTTGCCAGAACGGATCTGACGGTGAGGGACGAAAGCCAGGGGAG |
| d__Archaea_d__Archaea_ASV_4643 | 0 | 1 | 0 | 0 | 0 | 0 | 0 | 0 | 0 | 0 | 0 | CCTTCGCAATGCTCGAAAGAGTGACGAGGGGATTCCAAGTGCCTACGCTCAGCGTAGGCTGTTCAGATGCCTAAAAAGCATCTGGAGGAAGGACTGGGTAAGACGGGTGCCAGCCGCCGCGGTAATACCCGCAGTCCGAGTGGTAGTCGATTTTATTGGGCCTAAAGCGTCCGTAGCCGGTCTGGAAAATGCCTGGGTAAATCGTCTGGCTCAACCAAACGAATTCCGGGTAGACTACCAGACTTGGGACCGGTTGAGGGAGAGGGTACTTCAGGGGTAGGAGTGAAATCCTGTAATCCCTGAAGGACCACCGGTGGCGAAGGCGCTCTCCCAGAACGGGTCCGACGGTGAGGGACGAAAGCCAGGGGAG |
| d__Archaea_d__Archaea_ASV_4971 | 0 | 0 | 0 | 0 | 0 | 1 | 0 | 0 | 0 | 0 | 0 | TCTTCACAATGCACGAAAGTGCGATGGGGGGACTCCGAGTGCGTTCTTCCTACAAGGAAGGGCGCTTTTCTGCACTGTAAGAAGGTGCCGGAACAAGGGGCGGGCAACACTGCTGCCAGCCGCCGCGGTAATAGCAGGACCCCAAGTGGTGGTCGCTATTATTGGGCTTAAAGCGTTCGTAGCCGGCCGTGAAAGTCTGCTGTGAAATCCGCGTGCTTAACACGTGGGCGTGCAGCGGAAACTTCACGGCTAGGGACCGGATGGCGTTGGGGCTATTCCCGGGGTAGGAGTGAAATCCTAAAATCCCGGGAGGAGCACCTGTGGCGAAAGCGCCCAACGAATACGGATCCGACGGTGAGGAACGAAAGCCAGGGGAG |
| d__Archaea_d__Archaea_ASV_5119 | 0 | 0 | 0 | 0 | 0 | 0 | 0 | 0 | 1 | 0 | 0 | ACTGCACAATGCTGGAAACAGCGATGCGGGAACTCCAAGTGCCGGCACACAGTGTCGGCTGTTCCATTGTCTAAAAAACATAGGAAGTAAGGGCTGGGTAAGACGGGTGCCAGCCGCCGCGGTAATACCTGCAGCCCGAGTGGTGGCCGTTATTATTGGGCCTAAAGCGTCCGTAGCCGGATTGGTAAATGCCTGTGTAAATCGAACGGCTCAACCGTTCGAACTGCGGGTAGACTGCCAGTCTTGGGACCGGGAGAGTCACGAAGTACTCCGGGGGTAGAGGTGAAATTCTGTAATCCTCGGGGGACTACCGGTGGCGAAGGCGTCGTGATAGAACGGCTCCGACGGTGAGGGACGAAGCCCTGGGGCG |
| p__Iainarchaeota_c__Iainarchaeia_ASV_6414 | 0 | 0 | 0 | 0 | 0 | 0 | 0 | 0 | 1 | 0 | 0 | CCTTTGCAATGCGCGAAAGCGTGACAAGGGAACTCCAAGTGATCCAACGCAAGTTGGATCTTTTCTCAACTCTAGAAAGGTTGGGGAATAAGGACTGGGCAAGACCCGTGCCAGCCGCCACGGTAATACGGGCAGTTCGAGTGATATCCGCGTATACTGGGCTTAAAGCGTCCGTAGCCGGTTTGGCAAGTTTCTGGTGAAATCCCGTCGCTTAACGGCGGGAAGTGCTGGAAATACTACCAGGCTTGGAGGCGGGAGAGGATAGAGGTACGTTAGGGGTAGCGGTTAAATGCAATAATCCTTAACGGACCACCGGTAGCGAAGGCGTCTATCTAGAACGCGCTCGACGGTGAGGGACGAAGGCCAGGGGAG |

| | Presence (1) or Absence (0) in each layer | | | | | | | | | | | |
|---|---|---|---|---|---|---|---|---|---|---|---|---|
| Phylum_Genus_ASV_number | L1 | L2 | L3 | L4 | L5 | L6 | L7 | L8 | L9 | L10 | L11 | Sequence |
| p__Iainarchaeota_o__Iainarchaeales_ASV_5321 | 0 | 0 | 0 | 0 | 0 | 0 | 0 | 0 | 0 | 1 | 0 | CCTTTACAATGCGCGAAAGCGTGATAGGGGGATTTCGAGTGCAAGCTTTGCTTGCTTTTGCCAAGCCCAAATCGCTTGGCGAATAAGTGGGGGGCAAGATACGTGCCAGCCGCCGCGGTAACACGTACTCCACAAGTGGTATCCATCTATATTGGGCCTAAAACGTCCGTAGCTCGTCCAAAAAGTCCTTGGTGAAAGCAGCTGGCTTAACTAGCTGAACTGCCAAGGAAACTGTTGGGCTAGAGGCCGGAAGAGGTCAAGGGAATTTCAGGGGTAGCGGTGAAATGTTATGATCCTTGAAAGACCACCAGTAGCGAAGGCGCTTGACCAGGACGGTCCTGACAGTGAGGGACGAAAGCTGGGGGAG |
| p__Iainarchaeota_o__Iainarchaeales_ASV_5466 | 0 | 0 | 0 | 0 | 0 | 0 | 0 | 0 | 0 | 0 | 1 | ACTCCGCAATGCACGAAAGTGTGATAGGGGGATTTCGAGTGCTCTCTTTGAGAGCTTTTGCCAAACCCAAACCGTTTGGCGAATAAGTGGAGGGCAAGATACGTGCCAGCCGCCGCGGTAACACGTACTCCGCAAGTGGTATCCACTAATATTGGGCCTAAAACGTCTGTAGCCGGTTCAACAAGTCCTTGGTGAAATCATTGGGCTTAACTTAATGAATTGCCAAGGAAACTGCTGGACTCGAGGCCGGGAGACGCCAAGGGAATTTTTGGGGTAACGGTGAAATGTTATGATCCTGAAAAGACCACCAGTAGCGAAGGCGCTTGGCGAGCACGGGCCTGACGGTGAGAGACGAAAGCTGGGGGAG |
| p__Thermoproteota_g__BA2_ASV_6511 | 0 | 0 | 0 | 0 | 0 | 0 | 0 | 0 | 1 | 0 | 0 | CCTTCACAATGCACGGAAGTGCGATGGGGGGACTCCGAGTGCATTCTTCTCACAAAGAAGGATGCTTTTCTGAACTGTAAGGAGGTTCAGGAAGAAGGAGTGGGCAAGACTGCTGCCAGCCGCCGCGGTAATAGCAGCACTCCAAGTGGTAGTCGTTATTATTGGGCTTAAAGTGTTCGTAGCCGGCCGTGCAAGTCCGCTGTGAAATCCAGGCGCTTAACGTCTGGGCGTGCAGCGGAAACTGCACTGGCTAGGGACCGGACGGTGTCGAGGCTATTCCCGGGGTAGGAGTGAAATCCTATAATCCCGGGAGGACCACCAGTGGCGAAGGCTGTCGACTAGAACGCGCTCGACGGTGAGGGACGAAAGCTGGGGGAG |
| p__Nanoarchaeota_g__BMS3ABIN17_ASV_5184 | 1 | 0 | 0 | 0 | 0 | 0 | 0 | 0 | 0 | 0 | 0 | ATTTTACAATGCTCGAAAGAGTGATCGAGTAAGCCAGAGTGTTTCTCTTTGAGAAACTTTTGTCTGCTGTAAAAAGGCAGACGAATAAGGACTGGGCAAGACCGGTGCCAGCCGCCGCGGTAATACCGGCGGTCCGAGTCGCATCCACATTTATTGGGTCTAAAACATCCGTAGCTTGTTTTGTAAGTCTCTTGTGAAATCTAATATCTCAAGTATTAGTCGTGCAAGAGATACTGCATTGCTAGAGACCGGAAGACGTAAGGAGTACGGTTAAGGTAGCGGTAAAATGTGTTAATCTTAACTGGACTAACAATGGCGAAGGCACCTTACGAGTACGGATCTGACAGTGAGGGATGAAGGCTAGGGGCG |
| p__Nanoarchaeota_g__BMS3ABIN17_ASV_5223 | 1 | 0 | 0 | 0 | 0 | 0 | 0 | 0 | 0 | 0 | 0 | CTTTTACAATGCACGCAAGTGTGATAAAGCGAATCAGAGTGTTTTCCAATTGGAAAACTTTTGCCAAATGTAAAAAGTTTGGCGAATAAGGACTGGGTAAGACGAGTGCCAGCCGCCGCGGTAATACTCGCGGTCCAAGTCGCAGCCATTTTTATTGGGTCTAAAACATCCGTAGCTTGCTAAATAAGTTTTCTGTGAAATCCTATTTCTTAAAGATGGGGCTTGCAGAAAATACTGTTCAGCTAGAGACTGGGAGGCGTAAGGAGTACGTTCGAAGTAGCGGTAAAATGTGTTAATCTCGGACGGACTAACAACAGCGAAGGCACCTTACGAGAACAGTTCTGACAGTGAGGGATGAAGGCTAGGGGCG |
| p__Nanoarchaeota_g__BMS3ABIN17_ASV_5266 | 0 | 0 | 0 | 0 | 0 | 0 | 0 | 0 | 0 | 1 | 0 | ATTCTGCAATGCGCGCAAGCGTGACAGATTGAACCAGAGTGCTTTTCAATTTGAAAAGCTTTTGTGAAATGCAAAAAGTTTTACGAATAAGGACTGGGTAAGACTAGTGCCAGCCGCCGCGGTAATACTAGCGGTCCAAGTCGCAGCCACAATTATTGGGTCTAAAACATCCGTAGCTTGCTTGATAAGTTTCCTGTGAAATTCGGTTTCTTAAAAGCCGAGCTGGCGGGAAGTACTGTCAAGCTAGAGACTGGAAGACGTAAGAAGTATGTTCAAAGTAGTGGTAAAATGGGTTGATCTTGAGCAGACTAACGATAGCGAAGGCACCTTACGAGAACAGATCTGACAGTGAGAGATGAAGGCCAGGGTCG |
| p__Nanoarchaeota_g__BMS3ABIN17_ASV_5729 | 0 | 1 | 0 | 0 | 0 | 0 | 0 | 0 | 0 | 0 | 0 | AATCTGCAATGCGCGCAAGCGTGACAGTTTGAACCAGAGTGCTTCTCTATTGAGAAGCTTTTGTAGAATGTAAAAAGTTCTACGAATAAGGACTGGGTAAGACTAGTGCCAGCCGCCGCGGTAATACTAGCGGTCCAAGTCGCAGCCATATTTATTGGGTCTAAAACATCCGTAGCTTGCTTGGCAAGTTCCTTGTGAAATCCTGTGTCTTAAGCATAGGGCGTGCAGGGAGTACTACCAAGCTAGAGACTGGGAGACGTGAGAAGTACGTTCGGAGTAGCGGTAAAATGTTTTAATCCTGAGCGGACTAACAATAGCGAAGGCATCTCACGAGAACAGTTCTGACAGTCAGGGATGAAGGCTAGGGGCG |
| p__Nanoarchaeota_g__BMS3ABIN17_ASV_5818 | 0 | 0 | 1 | 0 | 0 | 0 | 0 | 0 | 0 | 0 | 0 | CTTTTACAATGCACTAAAGTGTGATAGAGTGAGTCAGAGTGTTTTCCAATTGGAAAACTTTTGCCAAATGTAAAAAGTTTGGCGAATAAGGACTGGGTAAGACGAGTGCCAGCCGCCGCGGTAATACTCGCGGTCCAAGTCGCAGCCATTTTTATTGGGTCTAAAACATCCGTAGCTTGCTATCTAAGTTTCCTGTGAAATCTTATCTCTTAAAGATAGGGCGTGCAGGAAATACTATTTAGCTAGAGACTGGGAGACGTAAGGAGTACGTTTGGAGTAGCGGTAAAATGTGTTAATCCCAAACGGACTAACAATAGCGAAGGCACCTTACGAGAACAGTTCTGACAGTAAGGGATGAAGGCTAGGGGCG |
| p__Nanoarchaeota_g__BMS3ABIN17_ASV_6325 | 0 | 0 | 0 | 0 | 0 | 0 | 0 | 1 | 0 | 0 | 0 | CTTTTACAATGCACGAAAGTGTGATAGAGCGAGCCAAAGTGCTTAACTTTTTGTTAAGCTTTTGTCAGTTGTAAACATACTGACGAATAAGGACTGGGAAAGACCGGTGCCAGCCGCCGCGGTAATACCGGCAGTTCTAGTTGCAACCAACATTATTGGGTCTAAAACATCCGTAGCTTGTTTAGTAAGTTTCTTGTGAAATCTCGTGTCTCAAGCACGAGGCGTGCAGGAAATACTACTAGGCTAGAGACCGGTAGACGTAAGAAGTACGCTGGAGGTAGTGGTAAAATACGTTAATCTTCAGTGGACTCACAATTAGCGAAGGCATCTTACGAGGACGGATCTGACAGTGAGGGATGAAGGCCAGGGGCG |
| p__Nanoarchaeota_g__BMS3ABIN17_ASV_6403 | 0 | 0 | 0 | 0 | 0 | 0 | 0 | 0 | 1 | 0 | 0 | ATTCTGCAATGCTCGAAAGAGTGACAGAATGAGTCAGAGTGTTTGATTTTATCAAACTTTTATATATTGCAAAAAGATATATGAATAAGGACTGGGCAAGACCGGTGCCAGCCGCCGCGGTAATCCCGGCGGTCCAAGTCGCATCCACAATTATTGGGTCTAAAACATTCGTAGCTTGTTTTGCAAGTTTCTTGTGAAATCTGGTATCTTAAGTTCCAGGCGTGCAAGAAGTACTGCAATACTAGAGACCGGAAGACGCAAAGAGTATGTTTGAGGTAGCGGTAAAATGTGTTAATCTCAAATAGACTAACAATGGCGAAGGCACTTTGCGAGCACGGATCTGACAGTGAGGAATGAAGGCTAGGGGCG |
| p__Nanoarchaeota_g__BMS3ABIN17_ASV_6447 | 0 | 0 | 0 | 0 | 0 | 0 | 0 | 0 | 1 | 0 | 0 | ATTTTACAATGCTCGAAAGAGTGATAAAGCAAGCCAGAGTGTTTCTCTTAGAGAAACTTTTTTCTGCTGTAAAAATGCAGAAGAATAAGGACTGGGCAAGACCGGTGCCAGCCGCCGCGGTAATACCGGCGGTCCAAGTCGTATCCACTTTTATTGGGTCTAAAACATCCGTAGCTTGTTTTGTAAGTCTCTTGTGAAATTTTATATCTCAAATATAAAGCTTGCAAGAGATACTACATTGCTAGAGACCGGAAGACGTAACGAGTACAATAGAGGTAGCGGTAAAATGTGTTAATCTTTATTGGACTAACAATGGCGAAGGCACGTTACGAGTACGGATCTGACAGTGAGGGATGAAGGCTAGGGGCG |
| p__Nanoarchaeota_g__BMS3ABIN17_ASV_6529 | 0 | 0 | 0 | 0 | 0 | 0 | 0 | 0 | 1 | 0 | 0 | CTTTTGCAATGCGCGCAAGCGTGACAAAGCAAGCCAGAGTGTTTCTCTTATTTGAGAAACTTTTGTCAGCTGTAAAAAGGCTGGCGAATAAGGACTGGGCAAGACCGGTGCCAGCCGCCGCGGTAATACCGGCGGTCCAAGTCGCAGCCACAATTATTGGGTCTAAAACATCCGTAGCTTGTTATGCAAGTCTCCTGTGAAATCTGACATCTCAAGTGTCAGGCGTGCGAGAGATACTGCATGGCTAGAGACCGGAAGACGTAGCGAGTACGGCCAAGGTAGTGGTAGAATACGTTAATCTTGGTTGGACTAACAATGGCGAAGGCACGCTACGAGGACGGATCTGACAGTGAGGGATGAAGGCCAGGGGCG |
| p__Nanoarchaeota_g__BMS3ABIN17_ASV_6557 | 0 | 0 | 0 | 0 | 0 | 0 | 0 | 0 | 1 | 0 | 0 | CTTTTGCAATGCGCGAAAGCGTGACAAAGCAAGCCAGAGTGTTTCTTTTATTAAAGAAACTTTTGTCAGCTGTAAAAAGTCTGGCGAATAAGGACTGGGTAAGACTGGTGCCAGCCGCCGCGGTAATACCAGCGGTCCAAGTGGCAGCCGCGTTTATTGGGTCTAAAACATCCGTAGCTTGCTTCGTAAGTCTCATGTGAAATCTGGCCTCTCAAGGGTCAGGCGGGCATGAGATACTGCGTTGCTAGAGACCGGGAGGCGTAAGGAGTACAGTTAGGGTAGCGGTAAAATGCGTTGATCCTTACTGGACTAACGATTAGCGAAGGCACCTTACGAGAACGGATCTGACAGTGAGGGATGAAGGCTAGGGTCG |
| p__Nanoarchaeota_g__CG1-02-57-44_ASV_5483 | 0 | 0 | 0 | 0 | 0 | 0 | 0 | 0 | 0 | 0 | 1 | CCTTTACAATGCACGAAAGTGTGATAAGGGGATTCTAAGTGCTCATGCAAAGCATGGGCTTTTGCCAAGGGTAAATACCTTGGCGAATAAGGGGTGGGCAAGACTGGTGCCAGCCGCCGCGGTAACACCAGCGCCCCAAGTGGGAATCGGATTTATTGGGCCTAAAGCGTCCGTAGCCGGATTGTTAAATCCTTTGTGAAATTGTCAAGCCCAACTTGACAACGCGCAGAGGAGACTGGCAGTCTAGGGACCGGGAGGCGTGTGAGGTATTCCTTGGGGAGCGGTAAAATGTTATAATCCTTGGAAGACCACCTGTGGCGAAGGCGTCACACGAGAACGGATCCGACGGTGAGGGACGAAAGCCAGGGGAG |
| p__Nanoarchaeota_g__CG1-02-57-44_ASV_6212 | 0 | 0 | 0 | 0 | 0 | 1 | 0 | 0 | 0 | 0 | 0 | CCTTTACAATGCACGAAAGTGTGATAAGGGGATCCCAAGTGCTCATGCTCAGCATGGGCTTTTGCCAAGAGTAAATATCTTGGCGAATAAGTGGTGGGAAAGACTGGTGCCAGCAGCCGCGGTAACACCAGCGCCACGAGTGGCAGACGTTATTATTGGGCCTAAAGCGTCCGTAGCCGGTCTGGTAAATCTTTTGTGAAATCAAGAAGCTTAACTTCTTGGCGTGCAGAAGAGACTGCCAGGCTAGGAACCGGGAGGAGCCAGAAGTATTCTGGGGGTAGCGGTAAAATGCTATAATCCCCAGAGGACTACCTGTGGCGAAGGCGTCTGGCTAGAACGGATTCGACGGTGAGGGACGAAGGCCAGGGGAG |
| p__Nanoarchaeota_g__DSQK01_ASV_5704 | 0 | 1 | 0 | 0 | 0 | 0 | 0 | 0 | 0 | 0 | 0 | CCTCCACAATGCACGCAAGTGTGATGGGGGGATTCCGAGTGCTCAGGGATAACCTGGGCTTTTTACAAGGGTAAAGACCTTGTAGAATAAGTGGTGGGCAAGACTGGTGCCAGCCGCCGCGGTAACCCCAGCGCCACAAGTGGGAACCACGTTTATTGGGTCTAAAGCATCCGTAGCCGGTCTGGCAAATGCCCTGTGAAATCGTCCCGCTTAACGGAACGGCGTGCAGGGTAGACTACCAGACTAGGGACCGGGAGAAGCAAACGGTATTCCTGAGGGAGCGGTAAAATGTGATAATCTCAGGAGGACCACCGATGGCGAAGGCAGTTTGCTATAACGGATCCGACGGTGAGGGATGAAAGCTAGGGGAG |
| p__Thermoplasmatota_g__EX4484-6_ASV_5205 | 1 | 0 | 0 | 0 | 0 | 0 | 0 | 0 | 0 | 0 | 0 | ACTTCGCAATGCGGGCAACCGCGACGAGGGAACTCCCAGTGCTTGCGCAAAGCGTAAGCTGTTCCCATGCCTAAAAAGCATGGAGAGAAAGGGCTGGGTAAGACTGGTGCCAGCCGCCGCGGTAATACCAGCAGCCCTAGTGGTGTCCGCTATTATTGGGCCTAAAGCGTCCGTAGCCGGTCCGGTAAATCCTTGGGCAAATCGGGTCGCTTAACGATCCGAATTCCGGGGAGACTGCCGGACTTGGGACCGGGAGAGGTCGGAGGTACTTTGGGGGTAGGGGTGAAATCCTGTAATCCTCAAAGGACCACCAGTGGCGAAGGCGTCCGACTAGAACGGATCCGACGGTAAGGGACGAAGCCCTGGGGCA |
| p__Thermoplasmatota_g__EX4484-6_ASV_6293 | 0 | 0 | 0 | 0 | 0 | 0 | 0 | 1 | 0 | 0 | 0 | CCTTCACACTGTGCGAAAGCGCGATGAGGGGATCCCAAGTGCTTGCTCTTAGGAGTAAGCTGTTTTCCTGTTTAAATCGCAGGAAGAGGAAGGGCTGGGTAAGACGGGTGCCAGCCGCCGCGGTAATACCTGCAGCTCAAGTGGTGACCGCTATTATTGGGCCTAAAACGTCCGTAGCCGGCTGAGTAAATGCCTGGGTAAATCGTGTCGCTTAACGATACGAATTCCGGGTAGACTGCTTAGCTTGAGATCGGGAGAAGCTAGAGGTACTCCTGGGGTAGGGGTGAAATCCTGTAATCCTTGGGGGACCACCGGTGGCGAAGGCGTCCGACCAGAACGGGTCCGACGGTAAGGGACGAAGCCCTGGGGCG |
| p__Thermoplasmatota_g__EX4484-6_ASV_6561 | 0 | 0 | 0 | 0 | 0 | 0 | 0 | 0 | 1 | 0 | 0 | CCTTCGCAATGCGCGAAAGCGCGACGAGGGGATTCCAAGTGCATGCACTTAGTGTATGCTGTTTGCTAGTCTAAATCGCTAGCAGAGTAAGGGCTGGGTAAGACTGGTGCCAGCCGCCGCGGTAATACCAGCGGCCCTAGTGGTGACCGTTATTATTGGGCCTAAAGCGTTCGTAGCCGGTCCGGTAAATCTCTGGGTAAATCGATGGGCTTAACCCATCGAATTCTGGGGAGACTGCTGGACTTGAGACCGGGAGAGGCTAGAGGTACTCCTGGGGTAGAGGTGAAATTCGTAGATATCGGGAAGAACACCGGTGGCGAAGGCGGCCTTCTGGCCCGATACTGACGCTGAGACGCGAAAGCGTGGGGAG |
| p__Halobacteriota_g__Halobacterium_ASV_5666 | 0 | 1 | 0 | 0 | 0 | 0 | 0 | 0 | 0 | 0 | 0 | CCTTTACACTGCACGACAGTGCGATAGGGGGACTCCGAGTGCGAGGGCATATAGTCCTCGCTTTTCTGGACCGTAAGGAGGTCCAGGAACAAGGGCTGGGCAAGACCGGTGCCAGCCGCCGCGGTAATACCGGCAGCCCAAGTGATGGCCAATATTATTGGGCCTAAAGCGTCCGTAGCTGGCCGGACAAGTCCGTTGGGAAATCTGCTCGCTTAACGAGCAGGCGTCCAGCGGAAACTGTTCGGCTTGGGACCGGAGGACCCGAGGGGTACGTCCGGGGTAGGAGTGAAATCCTGTAATCCTGAGGGGACCACCAGTGGCGAAGGCGTCTAGCTAGAACGGGTCTGACGGTGAGGGACGAAGCCCTGGGGCG |
| p__Halobacteriota_g__Halobacterium_ASV_5809 | 0 | 0 | 1 | 0 | 0 | 0 | 0 | 0 | 0 | 0 | 0 | CCTTTACACTGCACGACAGTGCGATAGGGGGACTCCGAGTGCGAGGGCATACAGTCCTCGCTTTTCTGGACCGTAAGGAGATCCAGGAACAAGTGCTGGGCAAGACCGGTGCCAGCCGCCGCGGTAATACCGGCAGCACGAGTGATGGCCGATATTATTGGGCCTAAAGCGTCCGTAGCTGGCCGGACAAGTCCGTTGGGAAATCCACGCGCTCAACGCGTGGGCGTCCAGCGGAAACTGTCCGGCTTGGGACCGGAAGACCCAGAGGGTACGTCCGGGGTAGGAGTGAAATCCCGTAATCCTGGACGGACCACCGGTGGCGAAAGCGCTCTGGGAAGACGGATCCGACAGTGAGGGACGAAAGCTAGGGTCT |
| p__Halobacteriota_g__Halobellus_3026_ASV_6136 | 0 | 0 | 0 | 0 | 1 | 0 | 0 | 0 | 0 | 0 | 0 | CCTTTACACTGCACGCCAGTGCGATAAGGGGACTCCAAGTGCGAGGGCATATCGTCCTCGCTATCCTGTACTGTAGGGTAGTACAGTATCAAGAGCTGGGCAAGACCGGTGCCAGCCGCCGCGGTAATACCGGCAGCTCAAGTGATGGCCGATCTTATTGGGCCTAAAGCGTCCGTAGCTGGCCCTGCAAGTCCGTCGGGAAATCCATCAGCCCAACTGATGGGCGGCCGGCGGAAACTCCAGGGCTTGGGACCGGAAGGCTCGAGGGGTACGTCCGGGGTAGGAGTGAAATCCCGTAATCCCGGACGGACCACCGATGGCGAAAGCACCTCGAGAAGACGGCTCCGACAGTGAGGGACGAAAGCTAGGGGAG |

| | Presence (1) or Absence (0) in each layer | | | | | | | | | | | |
|---|---|---|---|---|---|---|---|---|---|---|---|---|
| Phylum_Genus_ASV_number | L1 | L2 | L3 | L4 | L5 | L6 | L7 | L8 | L9 | L10 | L11 | Sequence |
| p__Halobacteriota_g__Halobellus_3026_ASV_6253 | 0 | 0 | 0 | 0 | 0 | 0 | 1 | 0 | 0 | 0 | 0 | CCTTTACACTGCACGCAAGTGCGATAAGGGGACTCCGAGTGCGAGGGCATATCGTCCTCGCTTTTGTGTACCGTAGGGTGGTACACGAACAAGAGCTGGGCAAGACCGGTGCCAGCCGCCGCGGTAATACCGGCAGCTCGAGTGATGGCCGATCTTATTGGGCCTAAAGCGTCCGTAGCTGGCCTCGCAAGTCCGTCGGGAAATCCACGCGCCCAACGCGTGGGCGTCCGGCGGAAACTCCGAGGCTTGGGACCGGAAGGCTCGAGGGGTACGTCCGGGGTAGGAGTGAAATCCCGTAATCCCGGACGGACCACCGATGGCGAAAGCACCTCGAGAAGACGGATCCGACAGTGAGGGACGAAAGCTTGGGTCT |
| p__Halobacteriota_g__Halohasta_ASV_5232 | 1 | 0 | 0 | 0 | 0 | 0 | 0 | 0 | 0 | 0 | 0 | CCTTTACACTGCACGCCAGTGCGATAGGGGGATCCCAAGTGCACAGGCATAGTGCCTGTGCTTTTCGACACCCTAAGGCGATGTCAGAATAAGAGCTGGGCAAGACCGGTGCCAGCCGCTGCGGTAATACCGGCAGCTCGAGTGATGTCCACTATTATTGGGCCTAAAGCGTCCGTAGCTGGCCGCGCAAGTCCGTCGGGAAATCCACCTGCCCAACAGGTGGGCGTCCGGCGGAAACTGCGTGGCTTGGAACCGGAAGGCTCAGAGGGTACGTCCGGGGTAGGAGTGAAATCCCGTAATCCCGGACGGACCACCGATGGCGAAAGCACTCTGAGAGGACGGATTCGACAGTGAGGGACGAAAGCTAGGGTCT |
| p__Halobacteriota_g__Halohasta_ASV_5253 | 1 | 0 | 0 | 0 | 0 | 0 | 0 | 0 | 0 | 0 | 0 | CCTTCACACTGTGGGAAACCACGACGAGGGGATTCCAAGTGCCAACACAATGTGTTGGCTTTTCATCTGTCTAAAACGCAGGTGGAATAAGAGCCGGGTAAGACTGGTGCCAGCCGCCGCGGTAACACCAGCGGCTCGAGTGGTGGCCTTTTTATTGAGCTTAAAGCGTCCGTAGCTGGCCAGCCAAGTCCGTCGGGAAATCCACCTGCCCAACAGGTGGGCGTCCGGCGGAAACTGTCTGGCTTGGAACCGGAAGGCTCAGAGGGTACGTCCGGGGTAGGAGTGAAATCCCGTAATCCCGGACGGACCACCGATGGCGAAAGCACTCTGAGAAGACGGATTCGACAGTGAGGGACGAAAGCTAGGGTCT |
| p__Halobacteriota_g__Halohasta_ASV_5849 | 0 | 0 | 1 | 0 | 0 | 0 | 0 | 0 | 0 | 0 | 0 | CCTTTACACTGCACGCCAGTGCGATAAGGGGACACCAAGTGCGAGGGCATATCGTCCTCGCTTTTCACCACCGTAAGGAGGTACACGAATAAGAGCTGGGCAAGACCGGTGCCAGCCGCCGCGGTAATACCGGCAGCTCGAGTGATGTCCACTATTATTGGGCCTAAAGCGTCCGTAGCTGGCCAGCCAAGTCCGTCGGGAAATCCACCTGCCCAACAGGTGGGCGTCCGGCGGAAACTGTCTGGCTTGGAACCGGAAGGCTCAGAGGGTACGTCCGGGGTAGGAGTGAAATCCCGTAATCCCGGACGGACCACCGATGGCGAAAGCACTCTGAGAAGACGGATTCGACAGTGAGGGACGAAAGCTAGGGTCT |
| p__Halobacteriota_g__Halohasta_ASV_6365 | 0 | 0 | 0 | 0 | 0 | 0 | 0 | 1 | 0 | 0 | 0 | CCTTTACACTGCACGCCAGTGCGATAGGGGGACACCGAGTGCGAGGGCATAGAGCCCTCGCTTTTCTGGACCGTAAGGTGGTCCCGGAACAAGGGCTGGGCAAGACCGGTGCCAGCCGCCGCGGTAACACCGGCAGCCCAAGTGATGGCCGATATTATTGGGCCTAAAGCGTCCGTAGCTGGCCGTACAAGTTCGTCGGGAAATCCACCTGCCCAACAGGTGGGCGTCCGGCGAAAACTGTATGGCTTGGAACCGGAAGGCTCAGAGGGTACGTCCGGGGTAGGAGTGAAATCCTGTAATCCCGGACGGACCACCGATGGCGAAAGCACTCTGAGAAGACGGATTCGACAGTGAGGGACGAAAGCTAGGGTCT |
| p__Halobacteriota_g__Halolamina_ASV_5303 | 0 | 0 | 0 | 0 | 0 | 0 | 0 | 0 | 0 | 1 | 0 | CCTTTACACTGTACGAAAGTGCGATAAGGGGACCCCGAGTGCGAGGGCATACAGTCCTCGCTTTTGTGTACCGTAAGGAGGTACACGAATAAGGGCTGGGCAAGACCGGTGCCAGCCGCCGCGGTAATACCGGCAGCCCTAGTGATGGCCGCTATTATTGGGCCTAAAGCGTCCGTAGCCTGCCGGACAAGTCCGTTGGGAAATCCGTTCGCTTAACGAACGGGCGTCCAGCGGAAACTGTCCGGCTTGGGACCGGAAGACCCGAGGGGTACGTTCGGGGTAGGAGTGAAATCCCGTAATCCTGAACGGACCGCCGATGGCGAAAGCACCTCGAGAGGACTGCTCCGACAGTGAGGGACGAAAGCTGGGGTCT |
| p__Halobacteriota_g__Halomicrobium_3849_ASV_6080 | 0 | 0 | 0 | 0 | 1 | 0 | 0 | 0 | 0 | 0 | 0 | CCTTTACACTGCACGCCAGTGCGATAAGGGGACTCCGAGTGCGAGGGCATATAGCCCTCGCTTTTGTCGACCGTAAGGTGGTCGACGAACAAGTGCTGGGCAAGACCGGTGCCAGCCGCCGCGGTAATACCGGCAGCACGAGTGATGGCCGCTCTTATTGGGCCTAAAGCGTCCGTAGCCGGCCAGACAAGTCCGTTGGGAAATCGACGCGCTCAACGCGTCGGCGTCCAGCGGAAACTGTCTGGCTTGGGGCCGGAAGACCTGAGGGGTACGTCCGGGGTAGGAGTGAAATCCCGTAATCCTGGACGGACCACCGGTGGCGAAAGCGCCTCAGGAAGACGGACCCGACGGTGAGGGACGAAAGCTAGGGTCT |
| p__Halobacteriota_g__Halopelagius_ASV_5539 | 0 | 0 | 0 | 0 | 0 | 0 | 0 | 0 | 0 | 0 | 1 | CCTTTACACTGCACGCAAGTGCGATAAGGGGACTCCGAGTGCGAGGGCATATAGTCCTCGCTTTTCTGTACCGTAGGAAGGTACAGGAATAAGGGCTGGGCAAGACCGGTGCCAGCCGCCGCGGTAATACCGGCAGCCCGAGTGATGACCGATATTATTGGGCCTAAAGCGTCCGTAGCTGGCCAAGCAAGTCTGTCGGGAAATCCATCCGCTCAACGGATGGGCGTCCGGCGGAAACTCCTTGGCTTGGGACCGGAAGGCTCGAGGGGTACGTCTGGGGTAGGAGTGAAATCCTGTAATCCTGGACGGACCACCGATGGCGAAAGCACCTCGAGAAGACGGATCCGACAGTGAGGGACGAAAGCTAGGGTCT |
| p__Halobacteriota_g__Haloprofundus_ASV_6408 | 0 | 0 | 0 | 0 | 0 | 0 | 0 | 0 | 1 | 0 | 0 | CCTTTACACTGCACGACAGTGCGACAAGGGGACCCCGAGTGCGAGGGCATACAGTCCTCGCTTTTGTGAACCGTAAGATGGTTCACGAATAAGAGCTGGGCAAGACCGGTGCCAGCCGCCGCGGTAATACCGGCAGCTCGAGTGATGACCGCTGTTATTGGGCCTAAAGCGTCCGTAGCTGGCCGCGTAAGTTCATCGGGAAATCCATCCGCTCAACGGATGGGCGTCCGGTGAAAACTACACGGCTTGGAGCCGGAAGACCCAAGGGGTACGTCCGGGGTAGGAGTGAAATCCTGTAATCCTGGGGGGACCACCAGTGGCGAAAGCGTCTAGCTAGAACGGGTCTGACGGTGAGGGACGAAACCCTGGGGCG |
| p__Halobacteriota_g__Halorientalis_ASV_5879 | 0 | 0 | 1 | 0 | 0 | 0 | 0 | 0 | 0 | 0 | 0 | ACTTTACAATGCGGGAAACCGCGATAAGGGGACACTGAGTGCCAGCATATAATGCTGGCTGTCCGTATGTGTAAAATACATGCGTTAGCAAGGGCCGGGCAAGACCGGTGCCAGCCGCCGCGGTAACACCGGCGGCCCGAGTGGTGGTCGCTATTATTGGGTCTAAAGGGTCCGTAGCCGGTTTGATCAGTCCATTGGGAAATCTGGTGGCTTAACCGCCAGAATTCCAATGGATACTGATTGGCTTGGGGCCGGGAGATCCAAGGGGTACGTCCGGGGTAGGAGTGAAATCCCGTAATCCTGGACGGACCACCGATGGCGAAAGCACCTTGGAAAAATGGACCCGACGGTGAGGGACGAAAGCTAGGGTCT |
| p__Halobacteriota_g__Halorientalis_ASV_6498 | 0 | 0 | 0 | 0 | 0 | 0 | 0 | 0 | 1 | 0 | 0 | ACTTCGCACTGTGCGAAAGCACGACGAGGGAATCCCAAGTGCTGGCTCGTTAGAGTCAGCTGTTTCTTTGTCTAAAAAACAGAGAGAGTAAGGGCTGGGTAAGACGGGTGCCAGCCGCCGCGGTAATACCTGCAGCCCGAGTGGTGACTGTTATTATTGGGCCTAAAACGTCCGTAGCCGGTTCTGTAAATGCCTGGGTAAATCGTACGGCTCAACCGTACGAATTCCGGGTAGACTGCAGGGCTAGAGATCGGGAGAGGCTAGAGGTACTCTTGGGGTAGAGGTGAAATTCTGTAATCCTGGACGGACCACCGATGGCGAAAGCACCTTGGAAAAACGGACCCGACGGTGAGGAACGAAAGCTAGGGTCT |
| p__Halobacteriota_g__Halorubrum_ASV_5779 | 0 | 0 | 1 | 0 | 0 | 0 | 0 | 0 | 0 | 0 | 0 | CCTTTACACTGCACGCCAGTGCGATAAGGGAATCCCAAGTGCACAGGCATAGCGCCTGTGCTTTTGTACACCCTAGGGTGGTGTACGAATAAGGGCTGGGCAAGACCGGTGCCAGCCGCCGCGGTAATACCGGCAGCCCGAGTGATGGCCGATCTTATTGGGCCTAAAGCGTCCGTAGCTGGCCGCGCAAGTCCATCGGGAAATCCACCTGCTCAACAGGTGGGCGCCCGGTAGAAACTGCGTGGCTTGGGACCGGAAGGCGCGACGGGTACGTCCGGGGTAGGAGTGAAATCCCGTAATCCTGGACGGACCGCCGATGGCGAAAGCACGTCGCGAGAACGGATCCGACAGTGAGGGACGAAAGCCAGGGTCT |
| p__Halobacteriota_g__Halorubrum_ASV_6107 | 0 | 0 | 0 | 0 | 1 | 0 | 0 | 0 | 0 | 0 | 0 | ACTTTACAATGCGGGAAACCGCGATAAGGGGACACCGAGTGCCAGCATATTATGTTGGCTGTCCGCATGTATAAATAACATGTGTTAGCAAGGGCCGGGCAAGACCGGTGCCAGCCGCCGCGGTAATACCGGCAGCCCAAGTGATGGCCGATCTTATTGGGCCTAAAGCGTCCGTAGCTGGCCGCGCAAGTCCATCGGGAAATCCACCTGCTCAACAGGTGGGCGCCCGGTAGAAACTGCGTGGCTTGGGACCGGAAGGCGCGACGGGTACGTCCGGGGTAGGAGTGAAATCCCGTAATCCTGGACGGACCGCCGATGGCGAAAGCACGTCGCGAGAACGGATCCGACAGTGAGGGACGAAAGCCAGGGTCT |
| p__Halobacteriota_g__Halorussus_ASV_5196 | 1 | 0 | 0 | 0 | 0 | 0 | 0 | 0 | 0 | 0 | 0 | CCTTTACAATGTGCGAAAGCACGATAAGGGGATCCCAAGTGCGAGGGCATATCGTCCTCGCTTTTCTGGACTGTAAGGTGGTCCAGGAACAAGTGCTGGGCAAGACCGGTGCCAGCCGCCGCGGTAATACCGGCAGCACAAGTGATGGCCAATATTATTGGGCCTAAAGCGTCCGTAGCCGGCCGGATAAGTCCGTCGGGAAATCTACCCGCTTAACGGGTGGGCGTCCGGCGGAAACTGTTCGGCTTGGGGCCGGAAGGCCAGAGGGGTACGTCCGGGGTAGGAGTGAAATCCTGTAATCCCGGGCGGACCGCCGGTGGCGAAAGCGCCTCTGGAGGACGGACCCGACGGTGAGGGACGAAAGCTAGGGTCT |
| p__Halobacteriota_g__Halorussus_ASV_5645 | 0 | 1 | 0 | 0 | 0 | 0 | 0 | 0 | 0 | 0 | 0 | CCTTTACACTGCACGACAGTGCGATAAGGGAACTCCGAGTGCGAGGGCATATAGTCCTCGCTTTTGTGTACCGTAAGGTGGTACACGAATAAGGGCTGGGCAAGACCGGTGCCAGCCGCCGCGGTAATACCGGCAGCCCGAGTGATGGCCGCTATTATTGGGCCTAAAGCGTCCGTAGCCGGCCGAACAGGTCCGTCGGGAAATCTGCTCGCCTAACGAGCAGGCGTCCGGCGGAAACCAGTCGGCTTGGGGCCGGAGGACCTGAGGGGTACGTCCGGGGTAGGAGTGAAATCCCGTAATCCCGGACGGACCGCCGGTGGCGAAAGCGCCTCAGGAGGACGGACCCGACGGTGAGGGACGAAAGCTAGGGTCA |
| p__Halobacteriota_g__Halorussus_ASV_5774 | 0 | 0 | 1 | 0 | 0 | 0 | 0 | 0 | 0 | 0 | 0 | CCTTTACACTGTACGCCAGTGCGATAGGGGGACTCCAAGTGCGAGGGCATACAGTCCTCGCTTTTCACGACCGTAAGGTGGTCGTGGAATAAGAGCTGGGCAAGACCGGTGCCAGCCGCCGCGGTAATACCGGCAGCTCAAGTGATGGCCGTTATTATTGGGCCTAAAGCGTCCGTAGCCAGCCAGACAGGTCCGTCGGGAAATCCATCCGCTCAACGGATGGGCGTCCGGCGGAAACCAGCTGGCTTGGGGCCGGAAGACCTGAGGGGTACGTCCGGGGTAGGAGTGAAATCCCGTAATCCTGGACGGACCGCCGGTGGCGAAAGCGCCTCAGGAGGACGGACCCGACGGTGAGGGACGAAAGCTAGGGTCA |
| p__Halobacteriota_g__Halorussus_ASV_6188 | 0 | 0 | 0 | 0 | 0 | 1 | 0 | 0 | 0 | 0 | 0 | ACTTTACACTGCACGACAGTGCGATAAGGGGACTCCGAGTGCGAGGGCATATAGTCCTCGCTTTTGTACACCGTAAGGTGGTGTACGAATAAGGGCTGGGCAAGACCGGTGCCAGCCGCCGCGGTAATACCGGCAGCCCGAGTGATGGCCGCTATTATTGGGCCTAAAGCGTCCGTAGCTGGCCAGACAGGTCCGTCGGGAAATCCATCCGCTTAACGGATGGGCGTCCGGCGGAAACCAGCTGGCTTGGGGCCGGAAGACCTGAGGGGTACGTCCGGGGTAGAGGTGAAATTCTGTAATCCTGGGGGGACCACCAGTGGCGAAAGCGTCTAGCTAGAACGGGTCTGACGGTGAGGGACGAAACCCTGGGGCG |
| p__Halobacteriota_g__Halorussus_ASV_6277 | 0 | 0 | 0 | 0 | 0 | 0 | 0 | 1 | 0 | 0 | 0 | CCTTTACACTGCGGGAAACCGCGATAAGGGGACACCGAGTGCCAGCATACAATGCTGGCTGTCCACATGTGTAAATAGCATGTGATAGCAAGGGCCGGGCAAGACCGGTGCCAGCCGCCGCGGTAACACCGGCGGCCCGAGTGGTAATCACTATTATTGGGTCTAAAGGGTCCGTAGCCGGTTTAATCAGTCTTCCGGGAAATCTGACAGCTCAACTGTTAGGCTTCCGGCGGAAACCAGCTGGCTTGGGGCCGGAAGACCTGAGGGGTACGTCCAGGGTAGGAGTGAAATCCCGTAATCCTGGACGGACCACCGGTGGCGAAAGCGCCTCAGGAAAACGGACCCGACGGTGAGGGACGAAAGCTAGGGTCA |
| p__Halobacteriota_g__Halostagnicola_ASV_5864 | 0 | 0 | 1 | 0 | 0 | 0 | 0 | 0 | 0 | 0 | 0 | CCTTTACACTGCACGAAAGTGCGATAAGGGGATCCCAAGTGCGAGGGCGTACAGTCCTCGCTTTTCTGTACTGTAGGGAGGTACAGGAATAAGAGCTGGGCAAGACCGGTGCCAGCCGCCGCGGTAATACCGGCAGCTCGAGTGATGGCCGCTATTATTGGGCCTAAAGCGTCCGTAGCTGGCCACGCAAGTCCATCGGGAAATCTCTCCGCTCAACGGAGAGGCGTCCGGTGGAAACTGTGTGGCTTGGGACCGGAAGATCCAGAGGGTACGTCTGGGGTAGGAGTGAAATCCCGTAATCCGAGACGGACCACCGGTGGCGAAAGCGCTCTGGAAAGACGGATCCGACGGTGAGGGACGAAAGCTCGGGTCA |
| p__Halobacteriota_g__Halostella_ASV_5616 | 0 | 0 | 0 | 0 | 0 | 0 | 0 | 0 | 0 | 0 | 1 | CCTTTACACTGCACGCGAGTGCGATAAGGGGACCCCGAGTGCGAGGGCATATAGCCCTCGCTTTTGCACACCGTACTGAGGTGCGCGAATAAGGACTGGGCAAGACCGGTGCCAGCCGCCGCGGTAATACCGGCAGTCCAAGTGATGGTCGCTGTTATTGGGCCTAAAGCGTCCGTAGCCGGCCAGGCAAGTCCGTCGGGAAATCCACCTGCTCAACAGGTGGGCGTCCGGCGGAAACTGTCTGGCTAGGGACCGGAAGACCCGAGGGGTACGTCCAGGGTAGGAGTGAAATCCCGTAATCCTGGACGGACCGCCGGTGGCGAAAGCGCCTCGGGAGAACGGATCCGACGGTGAGGGACGAAAGCTTGGGTCA |
| p__Halobacteriota_g__Halovenus_ASV_5832 | 0 | 0 | 1 | 0 | 0 | 0 | 0 | 0 | 0 | 0 | 0 | CCTTTACACTGCACGACAGTGCGATAGGGGGACTCCGAGTGCGAGGGCATATAGTCCTCGCTTTTCTGTACCGTAAGGTGGTACAGGAACAAGTGCTGGGCAAGACCGGTGCCAGCCGCCGCGGTAATACCGGCAGCACGAGTGATGGCCGATATTATTGGGCCTAAAGCGTTCGTAGCCGGCCGGACAAGTTCGTTGGGAAATCCACGCGCTCAACGCGTGGGCGTCCAGCGAAAACTGTTCGGCTTGGGGCCGGAAGACTCGAGGGGTACGTCCGGGGTAGGAGTGAAATCCCGTAATCCTGGGGGGACCACCAGTGGCGAAAGCGTCTAGCTAGAACGGGTCTGACGGTGAGGGACGAAACCCTGGGGCG |
| p__Nanoarchaeota_g__J110_ASV_5245 | 1 | 0 | 0 | 0 | 0 | 0 | 0 | 0 | 0 | 0 | 0 | CCTTTACAATGCACGCAAGTGTGATAAGGGGATCCTAAGTGCTCATGCAATGCATGGGCTTTTGCCAAGAGTAAATATCTTGGCGAATAAGTGGTGGGTAAGACCGGTGCCAGCCGCCGCGGTAACACCGGCGCCACAAGTGGCAACCGCGTTTATTGGGCCTAAAGCGTCCGTAGCCGGCTTGGTAAATCTCATGTGAAATCGTCAGGCTCAACCTGACGGCTTGCTTGAGACACTGCCTCGCTAGAGACCGGGAGGGGTCAAAGGTATTCCGTGGGGAGCGGTAAAATGTGATAATCCTCGGAAGATCGCCTATGGCGAAGGCATTTGACTAGAACGGATCTGACGGTCAGGGACGAAGACCAGGGGAG |
| p__Methanobacteriota_A_1229_g__Methanobrevibacter_A_ASV_6153 | 0 | 0 | 0 | 0 | 0 | 1 | 0 | 0 | 0 | 0 | 0 | CCTCCGCAATGTGAGAAATCGCGACGGGGGGATCCCAAGTGCCATTCTTAACGGGATGGCTTTTCATTAGTGTAAAGAGCTTTTGGAATAAGAGCTGGGCAAGACCGGTGCCAGCCGCCGCGGTAACACCGGCAGCTCTAGTGGTAACAATTTTTATTGGGCCTAAAGCGTCCGTAGCCGGTTTAATAAGTCTCTGGTGAAATCCTGTAGCTTAACTGTGGGAATTGCTGGAGATACTATTAGACTTGAGACCGGGAGAGGTTAGAGGTACTCCCAGGGTAGAGGTGAAATTCTGTAATCCTGGGAGGACCGCCTGTTGCGAAGGCGTCTAACTGGAACGGTTCTGACGGTGAGGGACGAAAGTTAGGGGCG |
| p__Nanohaloarchaeota_g__Nanohalobium_ASV_5905 | 0 | 0 | 0 | 1 | 0 | 0 | 0 | 0 | 0 | 0 | 0 | CCTTCACAATGCACGAAAGTGCGATGGGGGGACTCCGAGTGCGTTCTTCTCACAAAGAAGGGCGCTTTTCTGCACTGTAAGGAGGTGCCGGAAAAAGGGGTGGGCAATACTGCTGCCAGCCGCCGCGGTAATAGCAGAGCCCCAAGTGGTAGTCGTTATTATTGGGTTTAAAGTGTTCGTAGCCGGCCGTGCAAGTCCGCTGTGAAATCCAAGCGCTTAACGTTTGGGCGTGCAGCGGAAACTGCACTGGCTAGGGACCGGACGGTGTCGAGGCTATTCCAGGAGTAGGGGTGAAATCCTAAAATTCCTGGAGGAGCGCCTGTGGCGGAAGCGCTCGACAAGTACGGATCCGACGGTGAGGAACGAAAGCCAGGGGAG |

| | Presence (1) or Absence (0) in each layer | | | | | | | | | | |
|---|---|---|---|---|---|---|---|---|---|---|---|
| Phylum_Genus_ASV_number | L1 | L2 | L3 | L4 | L5 | L6 | L7 | L8 | L9 | L10 | L11 | Sequence |
| p__Nanohaloarchaeota_g__Nanohalobium_ASV_6114 | 0 | 0 | 0 | 0 | 1 | 0 | 0 | 0 | 0 | 0 | 0 | CCTTCACAATGCACGACAGTGCGATGGGGGACTCCGAGTGTAGTCTTCCCACAAGGAAGGCTACTTTTCTGTACTGTAAGGAGGTACCGGAACAAGGGGTGGGCAATACTGCTGCCAGCCGCCGCGGTAATAGCAGAACCCCAAGTGGTAGTCGTTATTATTGGGCTTAAAGTGTTCGTAGCCGGCCGTGCAAGTCCGCTGTGAAATCCAGGTGCTTAACATCTGGGCGTGCAGCGGAAACTGCACTGGCTAGGGACCGGACGGTGTCGAGGCTATTCCAGGAGTAGGGGTGAAATCCTAAAATTCCTGGAGGAGCGCCTGTGGCGGAAGCGCTCGACAAGTACGGATCCGACGGTGAGGAACGAAAGCCAGGGGAG |
| p__Nanohaloarchaeota_g__Nanohalobium_ASV_6163 | 0 | 0 | 0 | 0 | 0 | 1 | 0 | 0 | 0 | 0 | 0 | CCTTCACAATGCACGAAAGTGCGATGGGGGACTCCGAGTGCGTTCTCCTTACAAAGGAGGGCGCTTTTCTGCACTTTAAGACGGTGCAGGAACAAGGGGTGGGTAAGACTGCTGCCAGCCGCCGCGGTAATAGCAGCACCCCAAGTGGTGGCCGATATTATTGGGCTTAAAGCGTTCGTAGCCGGCTGTGTAAGTCCGCTGTGAAATCCACCTGCTTAACAGGTGGGCGTGCAGCGGATACTGCATGGCTAGAGGCCGGACGGCGTCGAGGCTATTCCAGGGGTAGGAGTGAAATCCTAAAATCCCTGGAGGAGCACCTGTGGCGAAAGCGCTCGACGAGTACGGACCTGACGGTGAGGAACGAAAGCCAGGGGAG |
| p__Nanohaloarchaeota_g__Nanohalobium_ASV_6184 | 0 | 0 | 0 | 0 | 0 | 1 | 0 | 0 | 0 | 0 | 0 | CCTTCACAATGCACGCAAGTGCGATGGGGGACTCCGAGTGTGGTCTTCCTACAAGGAAGGCCACTTTTCTGCACTGTAAGAAGGTGCCGGAACAAGGGGTGGGCAAGACTGCTGCCAGCCGCCGCGGTAATAGCAGCATCCCAAGTGGTGGCCAGTATTATTGGGCTTAAAGCGTTCGTAGCCGGATCTGTAAGTCCGCTGTGAAATCCACACGCTTAACGTGTGGGCGTGCAGCGGATACTGCAGGTCTAGGGACCGGACGGCGTCAGGGCTATTCCCAGGGTAGGAGTGAAATCCTAAAATCCTGGGAGGAGCACCTGTGGCGAAAGCGCCTGACGAGTACGGATCCGACGGTGAGGAACGAAAGCCGGGGGAG |
| p__Nanohaloarchaeota_g__Nanohalobium_ASV_6224 | 0 | 0 | 0 | 0 | 0 | 0 | 1 | 0 | 0 | 0 | 0 | CCTTTACAATGCACGCAAGTGCGATAAGGGGACTCCGAGTGCGGTCTTCCTACACGGAAGGCCGCTTTTCTGTACTGTAAGAAGGTACCGGAACAAGGGGTGGGCAACACTGCTGCCAGCCGCCGCGGTAATAGCAGGACCCCAAGTGGTAGCCAGTTTTATTGGGCTTAAAGTGTTCGTAGCCGGCCATGTAAGTCCGCTGTGAAATCCAAGCGCTCAACGTTTGGAACTGCAGTGGATACTGCATGGCTAGGGACCGGACGGCGTCGAGGCTATTCCAGGGGTAGGAGTGAAATCCTAAAATCCCTGGAGGAGCACCTGTGGCGAAAGCGCTCGACGAGTACGGATCCGACGGTGAGGAACGAAAGCTAGGGGAG |
| p__Halobacteriota_g__Natronomonas_ASV_5187 | 1 | 0 | 0 | 0 | 0 | 0 | 0 | 0 | 0 | 0 | 0 | CCTTTACACTGCACGCCAGTGCGATAGGGGGACTCCGAGTGCGAGGGCATATAGTCCTCGCTTTTGTACACCGTAAGGTGGTGTACGAATAAGAACTGGGCAAGACCGGTGCCAGCCGCCGCGGTAATACCGGCAGTTCGAGTGATGGCCGATATTATTGGGCCTAAAGCGTTCGTAGCTGGCCAGTCAAGTCTGTCGGGAAATCCAGCGGCTCAACCGCTGGGCGTCCGGCGGAAACTGATTGGCTTGGGACCGGGAGACCCGAAGAGTACGTCCGGGGTAGGAGTGAAATCCCGTAATCCTGGACGGACCACCGATGGCGAAAGCACCTCGGAAGGACGGATCCGACAGTGAGGGACGAAAGCTAGGGTCT |
| p__Halobacteriota_g__Natronomonas_ASV_5740 | 0 | 1 | 0 | 0 | 0 | 0 | 0 | 0 | 0 | 0 | 0 | CCTTTACACTGCACGCCAGTGCGATAGGGGGACACCGAGTGCGAGGGCATAGAGCCCTCGCTTTTGTAGACCGTAAGGAGGTCTACGAATAAGGGCTGGGCAAGACCGGTGCCAGCCGCCGCGGTAACACCGGCAGCCCAAGTGATGGCCGATATTATTGGGCCTAAAGCGTCCGTAGCTGGCCGCGCAAGTTCGTCGGGAAATCCACTCGCTTAACGAGTGGGCGTCCGGCGAAAACTGTGCGGCTTGGGACCGGAAGATCCAAGGGGTACGTCCGGGGTAGGAGTGAAATCCTGTAATCCTGGACGGACCACCGGTGGCGAAAGCGCTCTGGAAGGACGGATCCGACGGTGAAAAACAAAAGCTCGGGTCA |
| p__Thermoplasmatota_g__Natronoplasma_ASV_5599 | 0 | 0 | 0 | 0 | 0 | 0 | 0 | 0 | 0 | 0 | 1 | CCTTCACACTGTGGGAAACCACGATGAGGGGATTCCAAGTGCCAACACTAAGTGTTGGCTTTTCATCTGTGTAAAAAGCAGGTGGAATAAGGGCCGGGTAAGACGGGTGCCAGCCGCCGCGGTAACACCCGCGGCCCAAGTGGTGGTCATTATTATTGAGCTTAAAGCGTCCGTAGCCGGTCTGGTAAATCTCTGGGTAAATCGAACGGCTCAACCGTTCGAATTCCAGAGAGACTGTCAGACTTGGGGCCGGGAGAGGTCAGAGGTACTCCTGGGGTAGAAGTAAAATTCTGTAATCCTAGGAGGACCACCAGTGGCGAAAGCGTCTGACTAGAACGGACTCGACGGTGAGGGACGAAGCCCTGGGGCG |
| p__Thermoplasmatota_g__Natronoplasma_ASV_5667 | 0 | 1 | 0 | 0 | 0 | 0 | 0 | 0 | 0 | 0 | 0 | CCTTCACACTGTGGGAAACCACGACGAGGGGATTCCAAGTGCCAACACCCAGTGTTGGCTTTTCATCTGTCTAAAACGCAGGCGGAATAAGGGCCGGGTAAGACTGGTGCCAGCCGCCGCGGTAACACCAGCGGCTCGAGTGGTGGCCATTATTATTGAGCTTAAAGCGTCCGTAGCCGGTCTGGTAAATCTCTGGGTAAATTGAACGGCTTAACCATTCGAATTCCGGGTAGACTGCTAGACTTGAGATCGGGAGAGGTTAGAGGTACTCCTTGGGTAGAGGTGAAATTCTGTAATCCTAGGGGGACCACCTGTGGCGAAAGCGTCTAACCAGAACGAGTCTGACGGTGAGGGACGAAGCCCTGGGTCA |
| p__Halobacteriota_g__Natronorubrum_4312_ASV_5635 | 0 | 1 | 0 | 0 | 0 | 0 | 0 | 0 | 0 | 0 | 0 | CCTTTACACTGCACGCGAGTGCGATAAGGGGACTCCGAGTGCGAGGGCATATAGTCCTCGCTTTTCACCACCGTAAGGTGGTGGTGGAATAAGTGCTGGGCAAGACCGGTGCCAGCCGCCGCGGTAATACCGGCAGCACGAGTGATGACCGCTATTATTGGGCCTAAAGCGTCCGTAGCTGGCCACGCAAGTCTATCGGGAAATCTCTTCGCTTAACGAAGAGGCGTCCGGTGGAAACTGCGTGGCTTGGGACCGGAAGATCCAGAGGGTACGTCCGGGGTAGGAGTGAAATCCTGTAATCCTGGACGGACCGCCGGTGGCGAAAGCGCTCTGGAAAGACGGATCCGACGGTGAGGGACGAAAGCTCGGGTCA |
| p__Thermoproteota_g__Nitrososphaera_ASV_5225 | 1 | 0 | 0 | 0 | 0 | 0 | 0 | 0 | 0 | 0 | 0 | CCTCTGCAATAGGCGAAAGCCTGACAGGGTTACTCTGAGTGATGCCCGTTAAGGGTATCTTTTGGCACCTCTAAAAATGGTGCAGAATAAGGGGTGGGCAAGTCTGGTGTCAGCCGCCGCGGTAATACCAGCACCCCGAGTTGTCGGGACGATTATTGGGCCTAAAGCATCCGTAGCCTGTTCTGCAAGTCCTCCGTTAAATCCACCTGCTCAACGGATGGGCTGCGGAGGATACCGCAGAGCTAGGAGGCGGGAGAGGCAAACGGTACTCAGTGGGTAGGGGTAAAATCCATCGATCTACTGAAGACCACCAGTGGCGAAGGCGGTTTGCCAGAACGCGCTCGACGGTGAGGGATGAAAGCTGGGGGAG |
| p__Halobacteriota_g__Saliphagus_ASV_5770 | 0 | 0 | 1 | 0 | 0 | 0 | 0 | 0 | 0 | 0 | 0 | CCTTTACACTGCACGACAGTGCGATAGGGGGACTCCAAGTGCGAGGGCATATAGTCCTCGCTTTTCTCAACCGTAAGGTGGTTGAGGAATAAGTGCTGGGCAAGACCGGTGCCAGCCGCCGCGGTAATACCGGCAGCACGAGTGATGACCGCTATTATTGGGCCTAAAGCGTCCGTAGCCGGCCGTCCAAGTCCGTCGGGAAATCCGCACGCTTAACGTGCGGGCGTCCGGCGGAAACTGTTCGGCTTGGGACCGGAAGACTCGAGGGGTACGTCTGGGGTAGGAGTGAAATCCCGTAATCCTGGACGGACCACCGGTGGCGAAAGCGCCTCGAGAGAACGGATCCGACGGTGAGGGACGAAAGCTAGGGTCA |
| p__Halobacteriota_g__Saliphagus_ASV_5868 | 0 | 0 | 1 | 0 | 0 | 0 | 0 | 0 | 0 | 0 | 0 | CCTTTACACTGCACGACAGTGCGATAAGGGGACCCCCAGTGCGAGGGCATACCGTCCTCGCTTTTCAACACCGTAAGGAGGTGCTGGAACAAGAGCTGGGCAAGACCGGTGCCAGCCGCCGCGGTAATACCGGCAGCCCAAGTGATGACCGCTATTATTGGGCCTAAAGCGTCCGTAGCCGGCCTCGCAAGTCCGTCGGGAAATCCGCACGCCTAACGTGCGGGCGTCCGGCGGAAACTGCGTGGCTTGGGACCGGAAGACTCGAGGGGTACGTCCGGGGTAGGAGTGAAATCCCGTAATCCTGGACGGACCACCGGTGGCGAAAGCGCCTCGAGAGGACGGATCCGACGGTGAGGGACGAAAGCTAGGGTCA |
| p__Thermoplasmatota_g__SM1-50_ASV_5268 | 0 | 0 | 0 | 0 | 0 | 0 | 0 | 0 | 0 | 1 | 0 | CCTTCACACTGTGCGAAAGCGCGATGAGGGGATTCCAAGTGCTAACTCGAAAGAGTTGGCTGTTTCTCTGTCTAAAAAGCAGAGAGAGTAAGGGCTGGGTAAGACGGGTGCCAGCCGCCGCGGTAATACCTGCAGCTCAAGTGGTGACCGCTATTATTGGGCCTAAAACGTCCGTAGCCGGCTGAGTAAATGCCTGGGTAAATCGTGTCGCTTAACGATACGAATTCCGGGTAGACTGCTTAGCTTGAGACTGGGAGAAGCTAGAGGTACTCCTGGGGTAGAGGTGAAATTCTGTAATCCTGGGGGGACCACCAGTGGCGAAAGCGTCTAGCTAGAACGGGTCTGACGGTGAGGGACGAAACCCTGGGGCG |
| p__Thermoplasmatota_g__SM1-50_ASV_5461 | 0 | 0 | 0 | 0 | 0 | 0 | 0 | 0 | 0 | 0 | 1 | ACTTCACACTGTGCGAAAGCACGATGAGGGAATCCCAAGTGCCAGCACGTAAGTGTTGGCTGTTTCTTTGTCTAAAAAACAGAGAGAGTAAGGGCTGGGTAAGACGGGTGCCAGCCGCCGCGGTAATACCTGCAGCCCGAGTGGTGACTGTTATTATTGGGCCTAAAACGTCCGTAGCCGGTTCTGTAAATGCCTGGGTAAATCGTACGGCTCAACCGTACGAATTCCGGGTAGACTGCAGGGCTAGAGATCGGGAGAGGCTAGAGGTACTCTTGGGGTAGAGGTGAAATTCTGTAATCCTAGGGGGACCACCAGTGGCGAAGGCGTCTAGCTAGAACGAGTCTGACGGTCAGGGACGAAGCCCTGGGGCG |
| p__Thermoplasmatota_g__SM1-50_ASV_5595 | 0 | 0 | 0 | 0 | 0 | 0 | 0 | 0 | 0 | 0 | 1 | CCTTTACACTGCACGCAGGTGCGATAGGGGAATCCCAAGTGCGAGGGCATACTAGCCCTCGCTTTTGTGTACTGTAAGGAGGTACACGAATAAGAGCTGGGTAAGACGGCTGCCAGCCGCCGCGGTAATAGCTGCAGCTCGAGTGGTGACCACTTTTATTGGGCCTAAAACGTCCGTAGCCGGTCTAGTAAATGCCTGGGTAAATCGTGCCGCTTAACGGTACGAATTCCGGGTAGACTGCTAGACTTGAGATCGGGAGAGGTTAGAGGTACTCCTTGGGTAGAGGTGAAATTCTGTAATCCTAGGGGGACCACCTGTGGCGAAAGCGTCTAACCAGAACGAGTCTGACGGTGAGGGACGAAGCCCTGGGTCA |
| p__Thermoplasmatota_g__SM1-50_ASV_5890 | 0 | 0 | 1 | 0 | 0 | 0 | 0 | 0 | 0 | 0 | 0 | CCTTTACACTGCACGACAGTGCGATAAGGGGACACCGAGTGCGAGGGCATATCGTCCTCGCTTTTCACCACCGTAAGGTGGTGGTGGAATAAGTGCTGGGCAAGACCGGTGCCAGCCGCCGCGGTAATACCGGCAGCTCGAGTGATGTCCACTATTATTGGGCCTAAAACGTCCGTAGCCGGCTGAGTAAATGCCTGGGTAAATCGTGTCGCTTAACGATACGAATTCCGGGTAGACTGCTAGACTTGAGATCGGGAGAGGTTAGAGGTACTCCTTGGGTAGAGGTGAAATTCTGTAATCCTAGGGGGACCACCTGTGGCGAAAGCGTCTAACCAGAACGAGTCTGACGGTGAGGGACGAAGCCCTGGGTCA |
| p__Thermoplasmatota_g__SM1-50_ASV_6295 | 0 | 0 | 0 | 0 | 0 | 0 | 0 | 1 | 0 | 0 | 0 | CCTTTACACTGCACGCCAGTGCGATAAGGGAATCCCAAGTGCGTAGGCATAGAGCCTACGCTTTTGTGTACCCTAGGGCGGTACACGAATAAGGGCTGGGTAAGACGGGTGCCAGCCGCCGCGGTAATACCTGCAGCTCAAGTGGTGACCGCTATTATTGGGCCTAAAACGTCCGTAGCCGGCTGAGTAAATGCCTGGGTAAATCGTGTCGCTTAACGATACGAATTCCGGGTAGACTGCTTAGCTTGAGACTGGGAGAAGCTAGAGGTACTCCTGGGGTAGAGGTGAAATTCTGTAATCCTGGGGGGACCACCAGTGGCGAAAGCGTCTAGCTAGAACGGGTCTGACGGTGAGGGACGAAACCCTGGGGCG |
| p__Thermoplasmatota_g__SM1-50_ASV_6466 | 0 | 0 | 0 | 0 | 0 | 0 | 0 | 0 | 1 | 0 | 0 | CCTCTGCAATAGGCGAAAGCCTGACAGGGTTACTCTGAGTGATCTCCGTTAAGGAGGTCTTTTGGCATCTCTAAAAAGGGTGCAGAATAAGAGCTGGGTAAGACGGGTGCCAGCCGCCGCGGTAATACCTGCAGCTCAAGTGGTGGCCGCTATTATTGGGCCTAAAACGTTCGTAGCCGGTCTTTTAAATGCCTGGGTAAATCGAGCCGCTTAACGGTTCGAATTCCGGGTAGACTGGAAGACTTGAGACCGGGAGAGGCTAGAGGTACTCCTGGGGTAGAGGTGAAATTCTGTAATCCTGGGGGGACCACCAGTGGCGAAGGCGTCTAGCTAGAACGGGTCTGACGGTGAGGGACGAAGCCCTGGGGCG |
| p__Thermoplasmatota_g__SM1-50_ASV_6509 | 0 | 0 | 0 | 0 | 0 | 0 | 0 | 0 | 1 | 0 | 0 | CCTCCGCAATGCGAGCAATCGCGACGGGGGGACCCCAAGTGCCACTCTTAACGGGGTGGCTTTTATGAAGTGTAAAAAGCTTCAAGAATAAGGGCTGGGCAAGACCGGTGCCAGCCGCCGCGGTAATACCTGCAGCTCAAGTGGTGGCCGCTATTATTGGGCCTAAAACGTCCGTAGCCGGTTCTATAAATGCCTGGGTAAATCGTACGGCTTAACCGTACGAATTCCGGGTAGACTGTTCAACTTGGGACCGGGAGAGGCTAGAGGTACTCCTGGGGTAGAGGTGAAATTCTGTAATCCTAGGGGGACCACCAGTGGCGAAGGCGTCTAGCTAGAACGGGTCTGACGGTGAGGGACGAAGCCCTGGGGAG |
| p__Thermoplasmatota_g__SM1-50_ASV_6540 | 0 | 0 | 0 | 0 | 0 | 0 | 0 | 0 | 1 | 0 | 0 | ACTTCACACTGTGCGAAAGCACGATGAGGGAATCCCAAGTGCTTGCTCGCAAGAGTAAGCTGTTTCTATGTTTAAAATGCATTGAGAGTAAGGGCTGGGTAAGACGGGTGCCAGCCGCCGTGGTAATACCTGCAGCTCGAGTGGTGACCGCTATTATTGGGCCTAAAACGTCCGTAGCCGGCTGAGTAAATGCCTGGGTAAATCGTGTCGCTTAACGATACGAATTCCGGGTAGACTGCTTAGCTTGAGACCGGGAGAAGCTAGAGGTACTCCTGGGGTAGAGGTGAAATTCTGTAATCCTGGGGGGACCACCAGTGGCGAAAGCGTCTAGCTAGAACGGGTCTGACGGTGAGGGACGAAACCCTGGGGCG |
| p__Nanoarchaeota_g__SM23-78_ASV_5925 | 0 | 0 | 0 | 1 | 0 | 0 | 0 | 0 | 0 | 0 | 0 | CCTTTACAATGCACGCAAGTGTGATAAGGGGATCCCAAGTGCTCAAGCATTGCTTGGGCTTTTGCCATGTGTAAATAGCATGGCGAATAAGTGGTGGGAAAGACTGGTGCCAGCCGCCGCGGTAACACCAGCGCCACGAGTGGCAATCACGTTTATTGGGCCTAAAGCGTCCGTAGCAGGTTTCGTAAATCTCTTGTGAAATTGCCAAGCTCAACTTGGCAAAATGCAGGAGACACTGCGAGACTTGAGACCGGGAGGAGTTAGAAGTATTCCTTAGGGAGCGGTAAAATGCTATAATCTAAGGAGGACTACCTGTGGCGAAGGCGTCTAACTAGAACGGATCTGACTGTGAGGGACGAAAGCTAGGGGAG |
| p__Nanoarchaeota_g__SM23-78_ASV_6527 | 0 | 0 | 0 | 0 | 0 | 0 | 0 | 0 | 1 | 0 | 0 | CCTCTACAATGCACGCAAGTGTGATAGGGGGATCCCAAGTGCTCAAGCACTGCTTGGGCTTTTGCTTAGTGTAAATAGCTAGGCGAATAAGTGGTGGGAAAGACTGGTGCCAGCCGCCGCGGTAACACCAGCGCCACGAGTGGCAATCGCGTTTATTGGGCCTAAAGCGTCCGTAGCAGGTTTGTTGAATTTCTTGTGAAATTGCCAGGCTTAACTTGGCAGTGTGCAGGGAACACTGACAAACTTGGGACCGGGAAGGGTCAGAGGTATTCCTGAGGGAGCGGTAAAATGCTATAATCTCAGGAGGACCACCTGTGGCGAAGGCGTCTGACTAGAACGGATCCGACTGTGAGGGACGAAAGCCAGGGGAG |
| p__Halobacteriota_f__Haloarculaceae_3992_ASV_5397 | 0 | 0 | 0 | 0 | 0 | 0 | 0 | 0 | 0 | 1 | 0 | CCTTTACACTGCACGACAGTGCGATAGGGGGACTCCGAGTGCGAGGGCATATAGTCCTCGCTTTTCTGAACCGTAAGGTGGTTCAGGAACAAGGACTGGGCAAGACCGGTGCCAGCCGCCGCGGTAATACCGGCAGTCCGAGTGATGGCCGATTTTATTGGGCCTAAAGCGTCCGTAGCCTGCCCTGTAAGTCCATTGGGAAATCGACCAGCTCAACTGGTCGGCGTCCAGTGGAAACTACAGGGCTTGGGGCCGAGAGACTCGACGGGTACGTCCGGGGTAGGAGTGAAATCCTGTAATCCTGGACGGACCACCAATGGCGAAAGCACGTCGAGAGACCGGACCCGACGGTGAGGGACGAAAGCTAGGGTCT |
| p__Halobacteriota_f__Haloarculaceae_3992_ASV_5521 | 0 | 0 | 0 | 0 | 0 | 0 | 0 | 0 | 0 | 0 | 1 | CCTTTACACTGCACGACAGTGCGATAGGGGGACTCCGAGTGCGAGGGCATATAGTCCTCGCTTTTCTGAACCGTAAGGTGGTTCAGGAACAAGGACTGGGCAAGACCGGTGCCAGCCGCCGCGGTAATACCGGCAGTCCGAGTGATGGCCGATTTTATTGGGCCTAAAGCGTTCGTAGCCTGCCCTGTAAGTCCATTGGGAAATCGACCAGCTCAACTGGTCGGCGTCCAGTGGAAACTACAGAGCTTGGGGCCGAGAGACTCGACGGGTACGTCCGGGGTAGGAGTGAAATCCTGTAATCCTGGACGGACCACCAATGGCGAAAGCACGTCGAGAGACCGGACCCGACGGTGAGGGACGAAAGCTAGGGTCT |

| | Presence (1) or Absence (0) in each layer | | | | | | | | | | |
|---|---|---|---|---|---|---|---|---|---|---|---|
| Phylum_Genus_ASV_number | L1 | L2 | L3 | L4 | L5 | L6 | L7 | L8 | L9 | L10 | L11 | Sequence |
| p__Halobacteriota_f__Haloarculaceae_3992_ASV_5808 | 0 | 0 | 1 | 0 | 0 | 0 | 0 | 0 | 0 | 0 | 0 | CCTTTACACTGCACGAAAGTGCGATAGGGGGACTCCGAGTGCGAGGGCATATAGTCCTCGCTTTTCTGTACCGTAAGGTGGTACAGGAACAAGTGCTGGGCAAGACCGGTGCCAGCCGCCGCGGTAATACCGGCAGCACGAGTGATGGCCGATATTATTGGGCCTAAAGCGTCCGTAGCTTGCTGTGTAAGTCCATTGGGAAATCCACGCGCTCAACGCGTGGGCGTCCGGTGGAAACTGCACGGCTTGGGGCCGAGAGACTCGACGGGTACGTCCGGGGTAGGAGTGAAATCCCGTAATCCTGGACGGACCACCGATGGGGAAACCACGTCGAGAGACCGGACCCGACAGTGAGGGACGAAAGCCAGGGTCT |
| p__Halobacteriota_f__Halobacteriaceae_4595_ASV_5712 | 0 | 1 | 0 | 0 | 0 | 0 | 0 | 0 | 0 | 0 | 0 | CCTTTACACTGTACGAAAGTGCGATAAGGGGACCCCGAGTGCGAGGGCATACAGTCCTCGCTTTTGTGTACCGTACGATGGTACACGAATAAGAGCTGGGCAAGACCGGTGCCAGCCGCCGCGGTAATACCGGCAGCTCGAGTGATGGCCGCTATTATTGGGCCTAAAGCGTCCGTAGCCGGCCGAACAGGTCCGTTGGGAAATCCGCCAGCTTAACTGGCGGGCGTCCAGCGGAAACCAGTCGGCTTGGGGCCGGAAGACCTGAGGGGTACGTCCGGGGTAGGAGTGAAATCCTGTAATCCTGGACGGACCACCGATGGCGAAAGCACTTCACGAGGACGGATCCGACAGTGAGGGACGAAAGCTAGGGTCT |
| p__Halobacteriota_f__Haloferacaceae_ASV_6064 | 0 | 0 | 0 | 0 | 1 | 0 | 0 | 0 | 0 | 0 | 0 | CCTTTACACTGCGCGCACGCGCGATAAGGGGACCCCAAGTGCGAGGGCATACAGTCCTCGCTTTTCACATCCGTAGGGAGGATGTGGAATAAGGGCTGGGCAAGACCGGTGCCAGCCGCTGCGGTAATACCGGCAGCCCAAGTGATGGCCGCTCTTATTGGGCCTAAAGCGTCCGTAGCTGGCCACGTGTGTCCGTCGGGAAATCCACCAGCCCAACTGGTGGGCGGCCGGCGGAAACTGCGTGGCTTGGGACCGGAAGACCCACCGGGTACGTCCGGGGTAGGAGTGAAATCCCGTAATCCCGGACGGACCACCGATGGCGAAAGCACGGTGGGAAGACGGCTCCGACAGTGAGGGACGAAAGCTGGGGTCT |
| p__Halobacteriota_f__Natrialbaceae_ASV_5570 | 0 | 0 | 0 | 0 | 0 | 0 | 0 | 0 | 0 | 0 | 1 | CCTCCACAATGCACGCAAGTGTGATGGGGGGATCCCACGTGCTCACGTTCTGCGTGGGCTTTTGCTGAATGTAAGGAGTTCAGCGAATAAGGGCTGGGCAAGACCGGTGCCAGCCGCCGCGGTAATACCGGCAGCCCAAGTGATGACCGATATTATTGGGCCTAAAGCGTCCGTAGCCGGCTACGCAAGTCCGTCGGGAAATCTGCTCGCTCAACGAGTAGGCGTCCGGCGGAAACTGCGTGGCTTGGGACCGGAAGACTCGAGGGGTACGTCCGGGGTAGGAGTGAAATCCCGTAATCCTGGACGGACCACCGGTGGCGAAAGCGCCTCGAGAGGACGGATCCGACGGTGAGGGACGAAAGCTAGGGTCA |
| p__Halobacteriota_f__Natrialbaceae_ASV_6046 | 0 | 0 | 0 | 0 | 1 | 0 | 0 | 0 | 0 | 0 | 0 | CCTTTACACTGCACGCCAGTGCGATAAGGGGATTCCGAGTGCGAGGGCATACAGTCCTCGCTTTTGTGTACCGTAGGGAGGTACACGAATAAGGGCTGGGCAAGACCGGTGCCAGCCGCCGCGGTAATACCGGCAGCCCGAGTGATGGCCGATATTATTGGGCCTAAAGCGTCCGTAGCCGGCCGGGCAGGTCCATCGGGAAATCCACGCGCCTAACGCGTGGGCGTCCGGTGGATACCGTCCGGCTTGGGACCGGGAGACCCGAGGGGTACGTCTGGGTTAGGAGTGAAATCCCGTAATCCTGGACGGACCACCGGTGGCGAAAGCGCCTCTGGAAGACGGATCCGACGGTGAGGGACGAAAGCTCGGGTCA |
| p__Halobacteriota_f__Natrialbaceae_ASV_6111 | 0 | 0 | 0 | 0 | 1 | 0 | 0 | 0 | 0 | 0 | 0 | ACTTCGCAATGCGCGCAAGCGCGACGAGGGGATTCCAAGTGCATGCACACAGTGTATGCTGTTCAATAGTCTAAAACGCTATTGGAGTAAGGGCTGGGTAAGACTGGTGCCAGCCGCCGCGGTAATACCAGCGGCCCTAGTGGTGATCGTTATTATTGGGCCTAAAGCGTCCGTAGCCGGCCGAACAGGTCTGTTGGGAAATCCGCCAGCTTAACTGGCGGGCGTCCAGCAGAAACCAGTCGGCTTGGGGCCGGAAGACCTGAGGGGTACGTCTGGGGTAGGAGTGAAATCCCGTAATCCGAGACGGACCACCGGTGGCGAAAGCGCTCTGGAAAGACGGATCCGACGGTGAGGGACGAAAGCTCGGGTCA |
| p__Halobacteriota_f__Natrialbaceae_ASV_6388 | 0 | 0 | 0 | 0 | 0 | 0 | 0 | 1 | 0 | 0 | 0 | CCTTTACACTGCACGACAGTGCGATAAGGGGACTCCAAGTGCGAGGGCATATCGTCCTCGCTTTTCACTACCGTAAGGTGGTAGTGGAATAAGGGCTGGGCAAGACCGGTGCCAGCCGCCGCGGTAATACCGGCAGCCCGAGTGATGACCGCTATTATTGGGCCTAAAGCGTCCGTAGCCGGCCACGCAAGTCCGTCGGGAAATCCGTGCGCCTAACGCGCGGGCGTCCGGCGAAAACTGCGTGGCTTGGGACCGGAAGACTCGAGGGGTACGTCCGGGGTAGGAGTGAAATCCTGTAATCCTGGGGGGACCACCAGTGGCGAAAGCGTCTAGCTAGAACGGGTCTGACGGTGAGGGACGAAACCCTGGGGCG |
| p__Halobacteriota_o__Halanaerobiales_ASV_5210 | 1 | 0 | 0 | 0 | 0 | 0 | 0 | 0 | 0 | 0 | 0 | CCTTTACACTGTACGAAAGTGCGATAAGGGGACCCCGAGTGCGAGGGCATACAGTCCTCGCTTTTGTGTACCGTAAGGTGGTACACGAATAAGAGCTGGGCAAGACCGGTGCCAGCCGCCGCGGTAATACCGGCAGCTCGAGTGATGGCCGCTATTATTGGGCCTAAAGCGTCCGTAGCCGGCCGAACAGGTCCGTTGGGAAATCTGCTCGCTTAACGAGCAGGCGTCCAGCGGAAACCAGTCGGCTTGGGGCCGGAAGACCTGAGGGGTACGTCTGGGGTAGGAGTGAAATCCTGTAATCCTGGACGGACCATCGGTGGCGAAAGCGCCTCAGGAGGACGGACCCGACGGTGAGGGACGAAAGCTAGGGTCA |
| p__Halobacteriota_o__Halanaerobiales_ASV_5275 | 0 | 0 | 0 | 0 | 0 | 0 | 0 | 0 | 0 | 1 | 0 | CCTTTACACTGTACGAAAGTGCGATAAGGGGACCCCGAGTGCGAGGGCATACAGTCCTCGCTTTTGTGTACCGTACGATGGTACACGAATAAGAGCTGGGCAAGACCGGTGCCAGCCGCCGCGGTAATACCGGCAGCTCAAGTGATGGCCGTTATTATTGGGCCTAAAGCGTCCGTAGCCGGCCGAACAGGTCTGTTGGGAAATCCGCCAGTTTAACTGGCGGGCGTCCAGCAGAAACCAGTCGGCTTGGGGCCGGAAGACCTGAGGGGTACGTCCGGGGTAGGAGTGAAATCCTGTAATCCTGGGGGGACCACCAGTGGCGAAGGCGTCTAGCTAGAACGGGTCTGACGGTGAGGGACGAAACCCTGGGGCG |
| p__Halobacteriota_o__Halanaerobiales_ASV_5488 | 0 | 0 | 0 | 0 | 0 | 0 | 0 | 0 | 0 | 0 | 1 | CCTTTACACTGCACGAAAGTGCGATAAGGGGATCCCGAGTGCGAGGGCATACAGTCCTCGCTTTTGTACACCGTAAGGTGGTGTACGAATAAGAGCTGGGCAAGACCGGTGCCAGCCGCCGCGGTAATACCGGCAGCTCGAGTGATGGCCGCTATTATTGGGCCTAAAGCGTCCGTAGCCGGCCGAACAGGTCCGTTGGGAAATCCATCTGCCTAACAGATGGGCGTCCAGCGGAAACCAGTCGGCTTGGGGCCGGAAGACCTGAGGGGTACGTCTGGGGTAGGAGTGAAATCCTGTAATCCCGGACGGACCACCGATGGCGAAAGCACTCTGAGAGGACGGATTCGACAGTGAGGGACGAAAGCTAGGGTCT |
| p__Halobacteriota_o__Halanaerobiales_ASV_5509 | 0 | 0 | 0 | 0 | 0 | 0 | 0 | 0 | 0 | 0 | 1 | ACTTTACAATGCACGCAAGTGTGATAGGGGGATTCCGAGTGCTCAGGTATCGCCTGGGCTTTTGCCAAAGGTAAGGACTTTGGCGAATAAGTGGTGGGCAAGACTGGTGCCAGCCGCCGCGGTAACCCCAGCGCCACAAGTGGGGACCAACCTTATTGGGTCTAAAGCATCCGTAGCCGGTTGAGTAAATGCCCTGTGAAATCGTTGCGCTTAACGTAACGGCGTGCGGGGTAGACTGCTCGACTAGGGACCGGGAGAGGCAGGCAGTATTCTTGGGGGAGCGGTTAAATGCTATAATCCTGGACGGACCACCGGTGGCGAAAGCGCCTCGAGAGGACGGATCCGACGGTGAGGGACGAAAGCTAGGGTCA |
| p__Halobacteriota_o__Halanaerobiales_ASV_5598 | 0 | 0 | 0 | 0 | 0 | 0 | 0 | 0 | 0 | 0 | 1 | CCTTTACACTGCACGCCAGTGCGATAAGGGGACCCCGAGTGAAAGGGCATATCGCCCTTTCTTTTCTGAACCGTAGGGAGGTTGAGGAACAAGAGCTGGGCAAGACCGGTGCCAGCCGCCGCGGTAATACCGGCAGCTCGAGTGATGACCGATTTTATTGGGCCTAAAGCGTCCGTAGCTGGCCGCGCAAGTCTGTCGGGAAATCCACCCGCTTAACGGGTGGGCGTCCGGCGGAAACCAGCTGGCTTGGGGCCGGAAGACCTGAGGGGTACGTCCGGGGTAGGAGTGAAATCCCGTAATCCTGGACGGACCACCGGTGGCGAAAGCGCCTCAGGAAAACGGACCCGACGGTGAGGGACGAAAGCTAGGGTCA |
| p__Halobacteriota_o__Halanaerobiales_ASV_5763 | 0 | 1 | 0 | 0 | 0 | 0 | 0 | 0 | 0 | 0 | 0 | CCTTTACACTGCACGCAAGTGCGATAGGGGGACTCCCAGTGCGAGGGCATATCGTCCTCGCTTTTCTGCACCGTAAGGTGGTGCAGGAAGAAGGGCTGGGCAAGACCGGTGCCAGCCGCCGCGGTAATACCGGCAGCCCGAGTGATGGCCGTTATTATTGGGCCTAAAGCGTCCGTAGCCGGCCGGGCAAGTCCGTTGGGAAATCCACTCGCTTAACGAGTGGGCGTCCGGCGGAAACTGCTCGGCTTGGGACCGGAGGACCCGAGGGGTACGTCCAGGGTAGGAGTGAAATCCCGTAATCCTGGACGGACCGCCGGTGGCGAAAGCGCCTCGGGAGGACGGATCCGACGGTGAGGGACGAAAGCTAGGGTCT |
| p__Halobacteriota_o__Halanaerobiales_ASV_5783 | 0 | 0 | 1 | 0 | 0 | 0 | 0 | 0 | 0 | 0 | 0 | CCTTTACACTGCACGACAGTGCGATAGGGGGACCCCCAGTGCGAGGGCATATCGTCCTCGCTTTTCTGTACCGTAAGGTGGTACAGGAACAAGGGCTGGGCAAGACCGGTGCCAGCCGCCGCGGTAATACCGGCAGCCCAAGTGATGGCCGCTATTATTGGGCCTAAAGCGTCCGTAGCCGGCCGGGCAAGTCCGTTGGGAAATCCACCCGCTTAACGGGTGGGCGTCCAGCGGAAACTGCTCGGCTTGGGACCGGAAGACCCGAGGGGTACGTCTGGGGTAGGAGTGAAATCCTGTAATCCTGGACGGACCGCCGATGCCGAAAGGACCTCGGGAGGACGGATCCGACGGTGAGGGACGAAAGCTTGGATCT |
| p__Halobacteriota_o__Halanaerobiales_ASV_5840 | 0 | 0 | 1 | 0 | 0 | 0 | 0 | 0 | 0 | 0 | 0 | CCTTTACACTGCACGACAGTGCGATAAGGGGACCCCAGGTGCGAGGGCATATCGTCCTCGCTTTTCTGCACCGTAAGGAGGTGCAGGAATAAGGGCTGGGCAAGACCGGTGCCAGCCGCCGCGGTAATACCGGCAGCCCAAGTGATGGCCGTTATTATTGGGCCTAAAGCGTCCGTAGCCGGCCGGGCAAGTCCGTCGGGAAATCCACCCGCTCAACGGGTGGGCGTCCGGCGGAAACTGTTCGGCTTGGGGCCGGAGGACCTGAGGGGTACGTCCGGGGTAGGAGTGAAATCCTGTAATCCTGGGGGGACCACCAGTGGCGAAAGCGTCTAGCTAGAACGGGTCTGACGGTGAGGGACGAAACCCTGGGGAG |
| p__Halobacteriota_o__Halanaerobiales_ASV_5863 | 0 | 0 | 1 | 0 | 0 | 0 | 0 | 0 | 0 | 0 | 0 | CCTTTACACTGCACGACAGTGCGATAAGGGGACTCCAAGTGCGAGGGCATATCGTCCTCGCTTTTCACTACCGTAAGGTGGTATAGGAATAAGGGCTGGGCAAGACCGGTGCCAGCCGCCGCGGTAATACCGGCAGCCCAAGTGATGGCCGCTATTATTGGGTCTAAAGCGTCCGTAGCCTGCTGGGTAAGTCCGTCGGGAAATCCACGCGCCTAACGTGTGGGCGTCCGGCGGAAACTGCCCGGCTTGGGACCGGGAGACCCGAGGGGTACGTCTGGGGTAGGAGTGAAATCCTGTAATCCTGGACGGACCGCCGGTGGCGAAAGCGCCTCGGGAGAACGGATCCGACGGTGAGGGACGAAAGCTAGGGTCA |
| p__Halobacteriota_o__Halanaerobiales_ASV_5876 | 0 | 0 | 1 | 0 | 0 | 0 | 0 | 0 | 0 | 0 | 0 | CCTTTACACTGTACGAAAGTGCGATAAGGGGACCCCGAGTGCGAGGGCATACAGTCCTCGCTTTTGTGTACCGTACGATGGTACACGAATAAGAGCTGGGCAAGACCGGTGCCAGCCGCCGCGGTAATACCGGCAGCTCAAGTGATGGCCGTTATTATTGGGCCTAAAGCGTCCGTAGCCGGCCGAACAGGTCTGTTGGGAAATCCGCCAGCTTAACTGGCGGGCGTCCAGCGGAAACCAGTCGGCTTGGGGCCGGAAGACCTGAGAGGACGGATTCGACAGTGAGGGACGAAAGCTAGGGTCT |
| p__Halobacteriota_o__Halanaerobiales_ASV_5888 | 0 | 0 | 1 | 0 | 0 | 0 | 0 | 0 | 0 | 0 | 0 | CCTTTACACTGCACGAAAGTGCGATAAGGGGACCCCGAGTGCGAGGGCATACAGTCCTCGCTTTTGTGTACCGTAAGGTGGTACACGAATAAGAGCTGGGCAAGACCGGTGCCAGCCGCCGCGGTAATACCGGCAGCTCGAGTGATGGCCGCTATTATTGGGCCTAAAGCGTCCGTAGCCGGCCGAACAGGTCCGTTGGGAAATCCATCTGCCTAACAGATGGGCGTCCAGCGGAAACCAGTCGGCTTGGGGCCGGAAGACCTGAGGGGTATGTCCGGGGTAGGAGTGAAATCCTGTAATCCTAGGGGGACCACCTGTGGCGAAAGCGTCTAACCAGAACGAGTCTGACGGTGAGGGACGAAGCCCTGGGTCA |
| p__Halobacteriota_o__Halanaerobiales_ASV_5895 | 0 | 0 | 1 | 0 | 0 | 0 | 0 | 0 | 0 | 0 | 0 | CCTTTACACTGCACGACAGTGCGATAGGGGGACTCCGAGTGCGAGGGCATATCGTCCTCGCTTTTCTGTACCGTAAGGTGGTACAGGAAGAAGGGCTGGGCAAGACCGGTGCCAGCCGCCGCGGTAATACCGGCAGCCCGAGTGATGGCCAATATTATTGGGCCTAAAGCGTCCGTAGCCGGCCAGTCAGGTCCGTCGGGAAATCCACTCGCTCAACGAGTGGGCGTCCGGCGGAAACCGTCTGGCTTGGGACCGGGAGATCTGAGGGGTACGTCCGGGGTAGGAGTGAAATCCTGTAATCCTGGACGGACCATCGGTGGCGAAAGCGCCTCAGAAGAACGGATCCGACGGTGAGGGACGAAAGCTAGGGTCT |
| p__Halobacteriota_o__Halanaerobiales_ASV_5944 | 0 | 0 | 0 | 1 | 0 | 0 | 0 | 0 | 0 | 0 | 0 | CCTTTACACTGCACGCCAGTGCGATAAGGGGACTCCGAGTGCGAGGGCATATAGTCCTCGCTTTTCTGGACCGTAAGGTGGTCCAGGAACAAGTGCTGGGCAAGACCGGTGCCAGCCGCCGCGGTAATACCGGCAGCACGAGTGATGGTCGATATTATTGGGCCTAAAGCGTCCGTAGCTGGCCGAACAAGTCCGTCGGGAAATCCAGTGGCTCAACCACTGGGCGTCCGGTGGAAACTGTTCGGCTTGGGACCGGAAGACCCGAGGGGTACGTCTGGGGTAGGAGTGAAATCCCGTAATCCTGGACGGACCACCGATCGCGAAAGCACCTCGGGAGGACGGATCCGACAGTGAGGGACGAAAGCTGGGGTCT |
| p__Halobacteriota_o__Halanaerobiales_ASV_6115 | 0 | 0 | 0 | 0 | 1 | 0 | 0 | 0 | 0 | 0 | 0 | CCTTTACAATGCACACACGTGCGATAAGGGGAATCCAAGTGGTCGTTCCATACAACGGAGCGACCTTTTGCCAAGTGTAAGGAGCTGGGTGAATAAGGGGTGGGAAAGACGGCTGCCAGCCGCCGCGGTAATAGCCGCACCCCAAGTGGTGGCCACGATTATTGGGCTTAAAGAGTCCGTAGCCGGCCACGCAAGTCCACTGTGAAATCTGGCTGCTTAACAGTCAGGAGCGCAGCGGAAACTCCGTGGCTTGGGGCCGGAAGGCGTCAGGGCTATTCCTGGAGTAGGAGTGAAATCCTGAAATTCCAGGAGGAGCACCGGTGGCGAAAGCGCCTGACGATGACGGACCCGACGGTGAGGGACGAAAGCCAGGGGAG |
| p__Halobacteriota_o__Halanaerobiales_ASV_6238 | 0 | 0 | 0 | 0 | 0 | 0 | 1 | 0 | 0 | 0 | 0 | CCTTTACACTGCACGGAAGTGCGATAAGGGGACTCCGAGTGCGAGGGCATACAACGCCCTCGCTTTTCTGCACCGTAGGGAGGTGCAGGAACAAGGACTGGGCAAGACCGGTGCCAGCCGCCGCGGTAATACCGGCAGTCCGAGTGATGGCCGATCTTATTGGGCCTAAAGCGTTCGTAGCTGGCCGCGCAAGTCCATCGGGAAATCCAGGGGCTCAACCTCTGGGCGTCCGGTGGAAACTGTGCGGCTTGGGACCGGAAGACCCGAGGGGTACGTCCGGGGTAGGAGTGAAATCCTGTAATCCCGGACGGACCACCGATGGCGAAAGCACCTCGGGAGCACGGATCCGACAGTGAGGAACGAAAGCTGGGGTCT |

| | Presence (1) or Absence (0) in each layer | | | | | | | | | | |
|---|---|---|---|---|---|---|---|---|---|---|---|
| Phylum_Genus_ASV_number | L1 | L2 | L3 | L4 | L5 | L6 | L7 | L8 | L9 | L10 | L11 | Sequence |
| p__Halobacteriota_o__Halanaerobiales_ASV_6239 | 0 | 0 | 0 | 0 | 0 | 0 | 1 | 0 | 0 | 0 | 0 | CCTTTACACTGCACGACAGTGCGATAAGGGGACTCCGAGTGCGAGGGCATATCGTCCTCGCTTTTGTGTACCGTAGGGTGGTACACGAATAAGGGCTGGGCAAGACCGGTGCCAGCCGCCGCGGTAATACCGGCAGCCCGAGTGATGGCCGATTTTATTGGGCCTAAAGCGTCCGTAGCTGGCCGAGAAAGTCCGTCGGGAAATCCACAGGCTCAACCTGTGGGCGTCCGGCGGAAACTGTTCGGCTTGGGACCGGAAGGCCCGAGGGGTACGTCCGGGGTAGGAGTGAAATCCTGTAATCCCGGACGGACCACCGATGGCGAAAGCACCTCGGGAAGACGGATCCGACAGTGAGGGACGAAAGCTGGGGTCT |
| p__Halobacteriota_o__Halanaerobiales_ASV_6313 | 0 | 0 | 0 | 0 | 0 | 0 | 0 | 1 | 0 | 0 | 0 | CCTTTACAATGCGCGCAAGCGTGATAAGGGAATTCTATGTGCTTTCGTATTGCGAAAGCTTTTTTAGAATGTAAAAAGTTCTAAGAATAAGTGGTGGGCAAGACCGGTGCCAGCCGCCGCGGTAACACCGGCGCCACAAGTGGTGACCACATTTATTGGGCTTAAAGCATCCGTAGCTGGATATTCAAGTTTCTAGTGAAATCCTGTAATCCTGGACGGACCGTCGGTGGCGAAAGCGCCTCAGGAGGACGGACCCGACGGTGAGGGACGAAAGCTAGGGTCA |
| p__Halobacteriota_o__Halanaerobiales_ASV_6444 | 0 | 0 | 0 | 0 | 0 | 0 | 0 | 0 | 1 | 0 | 0 | CCTTTACACTGCACGCAAGTGCGATAAGGGGATCCCGAGTGCGAGGGCATATAGTCCTCGCTTTTCTGGACTGTAGGGAGGTCCAGGAACAAGAGCTGGGCAAGACCGGTGCCAGCCGCCGCGGTAATACCGGCAGCTCAAGTGATGGCCGATATTATTGGGCCTAAAGCGTCCGTAGCCGGCCGGATAAGTCCGTCGGGAAATCCACCCGCTCAACGGGTGGGCGTCCGGCGGAAACTGTCCGGCTTGGGGCCGGAAGGCCTGAGGGGTACGTCCGGGGTAGGAGTGAAATCCTGTAATCCTGGACGGACCGCCGATGGCGAAAGCACCTTGGGAAAACGGACTCGACAGTGAGGGACGAAAGCTAGGGTCT |
| p__Halobacteriota_o__Halanaerobiales_ASV_6538 | 0 | 0 | 0 | 0 | 0 | 0 | 0 | 0 | 1 | 0 | 0 | CCTTTACAATGTGCGCAAGCACGATAAGGGGATCCCAAGTGCGAGGGCATATCGTCCTCGCTTTTCTGGACCGTAAGGTGGTCCAGGAACAAGTGCTGGGCAAGACCGGTGCCAGCCGCCGCGGTAATACCGGCAGCACAAGTGATGGCCAATATTATTGGGCCTAAAGCGTCCGTAGCCGGCCGGATAAGTCCGTCGGGAAATCTACCCGCTTAACGGGTGGGCGTCCGGCGGAAACTGTTCGGCTTGGGGCCGGAAGGCCAGAGGGGTACGTCCGGGGTAGGAGTGAAATCCTGTAATCCCGGGCGGACCGCCGGTGGCGAAAGCGCCTCTGGAGGACGGACCCGACGGTGAGGGACGAAAGCTAGGGTCT |
| p__Nanoarchaeota_o__Woesearchaeales_581_ASV_5264 | 0 | 0 | 0 | 0 | 0 | 0 | 0 | 0 | 0 | 1 | 0 | CCTCAGCAATGCACGAAAGTGTGACTGGGGGATTCCGAGTGCTCATGTAACGCATGGGCTTTTGCCAAAGGTAATGTCTTTGGCGAATAAGTGGGGGGTAATACTGGTGCCAGCCGCCGCGGTAACACCAGAGCCACGAGTGGGGACCGCTATTATTGGGCCTAAAGCGTTCGTAGCCGGCCGGGTAAATCTCCTGTGAAATCGCAGCGCTCAACGTTGCGGCGTGCAGGAGAGACTGCTCGGCTTGGGACCGGGAGGCGTATGAGGTATTCTGGGGGTAGCGGTAAAATGCGATAATCCTCAGAGGACCACCTGTGGCGAAGGCGTCATACGAGAACGGATCCGACGGTGAGGGACGAAAGCTAGGGGAG |
| p__Nanoarchaeota_o__Woesearchaeales_581_ASV_5919 | 0 | 0 | 0 | 1 | 0 | 0 | 0 | 0 | 0 | 0 | 0 | CCTTTACAATGCACGAAAGTGTGATAAGGGAATTCCTAGTGCTTATACTAAGTATAGGCTTTTGCCAAAGGCAAAAACTTTGGCGAATAAGTGGTGGGTAAGACTGGTGCCAGCCGCCGCGGTAATACCAGCGCCACGAGTGGGGATCATATTTATTGGGCCTAAAGCGTCCGTAGCTGGTTGGATAAATCTCTTGTGAAATCGTGGAGCTTAACTTCACGGCATGCAGAAGACACTATTCGACTAGGAACCGGGAGAAGTAAGAGGTATTCTGGGGGGAGCGGTAAAATGCTATAATCCTCGGAGGACCACCGATAGCGAAGGCATCTTACTAGAACGGATTCGACAGTGAGGGACGAAAGCCAGGGGAG |
| p__Nanoarchaeota_o__Woesearchaeales_581_ASV_6393 | 0 | 0 | 0 | 0 | 0 | 0 | 0 | 1 | 0 | 0 | 0 | CCTTTACAATGCACGAAAGTGTGATAAGGGGATCCGAAGTGCCTATACATTGTATAGGCTTTTGCCAAAGGTAAAATCTTTGGCGAATAAGGGGTGGGCAAGACTGGTGCCAGCCGCCGCGGTAACACCAGCGCCTCGAGTGGCAACCGTGTTTATTGGGCCTAAAGCGTTCGTAGCCGGTTCAGTAAATCTTCTGTGAAATTGTGCAGCTTAACTGTACAGCGTGCAGGAGACACTGCTGAACTAGAGACCGGGAGGAGCTGAGAGTATTCTTGGGGGAGCGGTAAAATGTTGTGATCCCTTGAAGACTACCTGTGGCGAAGGCGCTCAGCTAGAACGGATCTGACGGTGAGGGACGAAAGCCAGGGGAG |
| p__Nanoarchaeota_o__Woesearchaeales_581_ASV_6428 | 0 | 0 | 0 | 0 | 0 | 0 | 0 | 0 | 1 | 0 | 0 | CCTCTACAATGCACGAAAGTGTGATAGGGGAATCCGAAGTGCTCATACATTGTATGGGCTTTTGCCAAAGCTAAATCCTTTGGCGAATAAGTGGTGGGTAAGACTGGTGCCAGCCGCCGCGGTAACACCAGCGCCACAAGTGGCAACCGCGTTTATTGGGCCTAAAGCGTTCGTAGCCGGCCGCGTAAATCTTCTGTGAAATTGTTTCGCTCAACGTAACAGTGTGCAGGAGACACTGCGTGGCTTGAGACCGGGAGGAGCTAGAGGTATTCTTGGGGGAGCGGTAAAATGTGATAATCCCTTGAAGACCACCTGTGGCGAAGGCGTCTAGCTAGAACGGATCTGACGGTGAGGGACGAAAGCCAGGGGAG |
| p__Nanoarchaeota_o__Woesearchaeales_581_ASV_6461 | 0 | 0 | 0 | 0 | 0 | 0 | 0 | 0 | 1 | 0 | 0 | CCTCTACACTGCACGAAAGTGTGATAGGGGGATCCTAAGTGCTCATGCTTTGCATGGGCTTTTGTCAAGAGTAAATATCTTGGCGAATAAGTGGTGGGTAAGACTGGTGCCAGCCGCCGCGGTAACACCAGCGCCACAAGTGGTGACCAGCATTATTGGGCCTAAAGCGTCCGTAGCAGGAAAAACAAATCTTTTGTGAAACTGTTCAGCTCAACTGAACAACTGGCAGAAGACACTATTTTTCTAGAGACCGGGAGGAGTCAGAGGTATTTTTGGGGGAGCGGTAAAATGCTATAATCCCAAGAGGACCACCGATGGCGAAGGCATCTGACTAGAACGGATCTGACTGTGAGGGACGAAGGCCAGGGGAG |
| p__Nanoarchaeota_o__Woesearchaeales_581_ASV_6472 | 0 | 0 | 0 | 0 | 0 | 0 | 0 | 0 | 1 | 0 | 0 | CCTTTACAATGCACGAAAGTGTGATAAGGGAATCCCAAGTGCTTATGCATTGCATAGGCTTTTGTCAAGAGTAAATACCTTGGCGAATAAGTGGTGGGCAAGACTGGTGCCAGCCGCCGCGGTAACACCAGCGCCGCAAGTGGCAACCCGCATTACTGGGTCTAAAGAGTCCGTAGCCGGTTTGTTAAATCTTTTGTGAAATCGTTGTTGTTAACATAACGGCGTGCAAAAGAGACTGGCAAACTAGAAACCGGGAGGAGTCAGAGGTATTTTCAGGGGAGCGGTAAAATGCTATAATCCTGAAAGGACCACCTGTGGCAAAGGCGTCTGACTAGAACGGATTTGACGGTGAGGGACGAAAGCCAGGGGAG |
| p__Nanoarchaeota_c__Nanoarchaeia_ASV_5605 | 0 | 0 | 0 | 0 | 0 | 0 | 0 | 0 | 0 | 0 | 1 | ACTCCACAATGCACGAAAGTGTGATGGGGGGATTCCGAGTGCTCACGGACTTCCGTGGGCTTTTTACGAGGGTAAATACCTCGTAGAATAAGTGGTGGGCAAGACTGGTGCCAGCCGCCGCGGTAACCCCAGCGCCACAAGTGGGAACCAGTATTATTGGGTCTAAAGCATCCGTAGCCGGTTAAGCAAATGCCTTGTGAAATCGTCCAGCCTAACTGAACGGCGAGCAAGGTAGACTACTTAACTTGGGACCAGGAGAGGCTAGCGGTATTCCGAGGGGAGCGGTAAAATGTGATAATCCTCGGAGGACCACCAATGGCGAAGGCAGCTAGCTAGAATGGATCCGACGGTGAGGGATGAAAGCCAGGGGAG |
| p__Nanoarchaeota_c__Nanoarchaeia_ASV_5949 | 0 | 0 | 0 | 1 | 0 | 0 | 0 | 0 | 0 | 0 | 0 | CCTTTACAATGCACGAAAGTGTGATAAGGGGATCCAAAGTGCTTATACATTGTATAGGCTTTTGCCAAAGTTAAATCCTTTGGCGAATAAGTGGTGGGCAAGACTGGTGCCAGCCGCCGCGGTAACACCAGCGCCACGAGTGGCAACCGTGATTATTGGGCCTAAAGCGTTCGTAGCCGGTTTAATAAATCTTTTGTGAAATCGTGTGGCTTAACTATACGACGTGCAAGAGACACTGTTAAACTTGAGACCGGGAGGAGCTAGGGGTATTCTTGGGGGAGCGGTAAAATGTTATAATCCCTTGAAGACCACCTGTGGCGAAGGCGCCTAGCTAGAACGGATCTGACGGTGAGGGACGAAAGCCAGGGGAG |
| p__Thermoplasmatota_p__Thermoplasmatota_ASV_5564 | 0 | 0 | 0 | 0 | 0 | 0 | 0 | 0 | 0 | 0 | 1 | CTTCTGCAATGCGCGCAAGCGTGACAGAGCGAGCCAGAGTGCTTTTCCATATAGGAAAAGCTTTTGTGTGATGTAAAAAGTCACACGAATAAGGACTGGGTAAGATTGGTGCCAGCCGCCGCGGTAATACCAACAGTCCAAGTTGCAGCCACAATTATTGGGTCTAAAACGTCCGTAGCCGGCTGAGTAAATGCCTGGGTAAATCGTGTCGCTTAACGATACGAATTCCGGGTAGACTGCTTAGCTTGAGACTGGGAGAAGCTAGAGGTACTCCTGGGGTAGAGGTGAAATTCTGTAATCCTGGGGGGACCACCAGTGGCGAAAGCGTCTAGCTAGAACGGGTCTGACGGTGAGGGACGAAACCCTGGGGCG |
| p__Thermoplasmatota_p__Thermoplasmatota_ASV_6009 | 0 | 0 | 0 | 0 | 1 | 0 | 0 | 0 | 0 | 0 | 0 | ACTCTGCAATGCTGGAAACAGCGACAGGGGAACACCCAGTGCTTGCACTTTGTGCAAGCTGTTTCCATGCCTAAAAAGCATGGAGAGTAAGGGCTGGGTAAGACGGGTGCCAGCCGCCGCGGTAATACCCGCAGCCCGAGTGGTGCTCGTTATTATTGGGCCTAAAGCGTCCGTAACCGGCCTCCTAAATGCCTGGGGAAATCGTCGTGCTCAACGCGACGAAGGCCCGGGTAGACTGGGAGGCTTGAGACCGGGAGAGGCCAGAGGTACTCCAGGGGTAGAGGTGAAATTCTGTAATCCTTGGGGGACCACCGGTGGCGAAGGCGTCTGGCTAGAACGGATCTGACGGCGAGGGACGAAGCCCTGGGGCG |
| p__Thermoplasmatota_p__Thermoplasmatota_ASV_6186 | 0 | 0 | 0 | 0 | 0 | 1 | 0 | 0 | 0 | 0 | 0 | ACTGCACAATGTGGGCAACCACGATGCGGTGACTCCAAGTGCCGGCACAGAGTGTCGGCTGTTTCCAAGCCTAAAAAGCTTGGAGAGTAAGGGCTGGGTAAGACGGGTGCCAGCCGCCGCGGTAATACCTGCAGCCCAAGTGGTGGTCGTTATTATTGGGCCTAAAGCGTCCGTAGCCGGCCGGGTAAATGCCTGTGTAAATCGAACGACTCAACCGTTCGAATTGCGGGTAGACTGCCTGGCTTGGGACTGGGAGAGCCGAGGGGTACTCCAGGGGTAGAGGTGAAATTCTGTAATCCTTGGGGGACCACCGGTGGCGAAGGCGCCTCGGTTGAACAGCTCCGACGGTGAGGGACGAAGCCCTGGGGCG |
| p__Thermoplasmatota_p__Thermoplasmatota_ASV_6526 | 0 | 0 | 0 | 0 | 0 | 0 | 0 | 0 | 1 | 0 | 0 | ACTTGTCAATGTGGGAAACCACGAACGGGGTATTCCAAGTGCAAGTACAATGTACTTGCTGTTCCCGAACCTAAAAAGTTCGAGAAGTAAGGGCTGGGTAAGACGGGTGCCAGCCGCCGCGGTAATACCCGCAGCCCAAGTGGTGGTCATTATTATTGAGCCTAAAACGTCCGTAGCCGGCTGAGTAAATGCCTGGGTAAATCGTGTCGCTTAACGATACGAATTCCGGGTAGACTGCTTAGCTTGAGACCGGGAGAAGCTAGAGGTACTCCTGGGGTAGAGGTGAAATTCTGTAATCCTGGGGGGACCACCAGTGGCGAAAGCGTCTAGCTAGAACGGGTCTGACGGTGAGGGACGAAACCCTGGGGCG |
| p__Thermoplasmatota_p__Thermoplasmatota_ASV_6544 | 0 | 0 | 0 | 0 | 0 | 0 | 0 | 0 | 1 | 0 | 0 | CCTTTACACTGCACGCCAGTGCGATAAGGGAACCCCGAGTGCGAGGGCATATCGCCCTCGCTTTTCTGGACCGTAGGGAGGTCCAGGAACAAGAGCTGGGCAAGACCGGTGCCAGCCGCCGCGGTAATACCGGCAGCTCGAGTGATGACCGATTTTATTGGGCCTAAAACGTCCGTAGCCGGCTGAGTAAATGCCTGGGTAAATCGTGTCGCTTAACGATACGAATTCCGGGTAGACTGCTTAGCTTGAGACCGGGAGAAGCTAGAGGTACTCCTGGGGTAGAGGTGAAATTCTGTAATCCTGGGGGGACCACCAGTGGCGAAAGCGTCTAGCTAGAACGGGTCTGACGGTGAGGGACGAAACCCTGGGGCG |
| d__Archaea_d__Archaea_ASV_5242 | 1 | 0 | 0 | 0 | 0 | 0 | 0 | 0 | 0 | 0 | 0 | ACTGCGCAATGTGAGAAATCACGACGCGGGAACACCAAGTGCCGGCACAGAGTGTCGGCTGTTTTCTAGCCTAAAAAGCTAGGAGAGTAAGGGCTGGGTAAGACGGGTGCCAGCCGCCGCGGTAATACCTGCAGCCCAAGTGGTGGCCGTTATTATTGGGCCTAAAACGTCCGTAGCCGGCTCGGTAAATGCCTGTGTAAATCGAACGGCTTAACCGTTCGAATGGCGGGTAGACTGCCGGGCTTGGGACCGGGAGAGTCACGAAGTACTCCGGGGGTAGAGGTGAAATTCTGTAATCCTCGGGGGACTACCGGTGGCGAAGGCGTCGTGATAGAACGGCTCCGACGGTGAGGGACGAAGCCCTGGGGCG |
| d__Archaea_d__Archaea_ASV_5503 | 0 | 0 | 0 | 0 | 0 | 0 | 0 | 0 | 0 | 0 | 1 | CTTCTGCAATGCGCGCAAGCGTGACAGAGCGACTCCGAGTGCTTATGTACAACATAGGCTTTTGCAAAGCGTAAATAACTTGGCGAATAAGGGGTGGGCAAGACCGGTGGCAGCAGCCGCGGTAACACCGGCGCCCCAAGTGGTGATCGCGATTATTGGGCCTAAAGCGTTCGTAGCCGGCTTAGCAAGTTCTTGGTGAAATCCTTGCGTATAACGTAAGGGCGTGCCAGGAATACTACTAAGCTAGGGACTGGGAGACGACAGAGGTATTCCTAGGGTAGGGGTAGAATCCTGTAATCCTGGGAGGACCACCGGTGGCGAAGGCGTCTGTCGAGAACAGGTCCGACGGTGAGGGACGAGAGCTAGGGGAG |
| d__Archaea_d__Archaea_ASV_5628 | 0 | 1 | 0 | 0 | 0 | 0 | 0 | 0 | 0 | 0 | 0 | CCTTTACACTGCACGACAGTGCGATAAGGGGACTCCAAGTGCGAGGGCATATCGTCCTCGCTTTTCACTACCGTAAGGTGGTAGTGGAATAAGGGCCGGGTAAGACTGGTGCCAGCCGCCGCGGTAACACCAGCGGCCCAAGTGGTAGTCGCTTTTATTGGGCCTAAAGCGTCCGTAGCCGGATTGACAAGTCCTTTGTGAAATCGGACGGCTTAACCGTTCGGCTTGCAGAGGAAACTGTCGATCTTGGGACCGGTAAGCGCTAAGGGTACTCCGGGGGTAGCGGTGAAATGCTATAATCCCTGGGGGACCGCCAGTGGCGAAAGCGCTTAGCGAGGCCGGATCCGACGGTGAGGGACGAAGGCTAGGGGAG |
| d__Archaea_d__Archaea_ASV_5748 | 0 | 1 | 0 | 0 | 0 | 0 | 0 | 0 | 0 | 0 | 0 | AATCTACAATACGCGAAAGCGTGATAGTTTGAGTCAGAGTGCGTTTCTTTGAAATGCTTTTGTGAAATGCAAAAAGTTTCACGAATAAGGGCTGGGTAAGACGGGTGCCAGCCGCCGCGGTAATACCTGCAGCCCAAGTGGTGACCGCTTTTATTGGGCCTAAAACGTCCGTAGCCGGTCTAGTAAATGCCTGGGTAAATCGTGCCGCTTAACGGTACGAATTCCGGGTAGACTGCCAGACTTGAGACTGGGAGAGGCTAGAGGTACTCCTGGGGTAGGGGTAAAATCCTATAATCCTGGGGGGACCACCGATGGCGAAGGCATCTCGCTGGAACGGGCCTGACGGTGAGGGACGAAACCCAGGGGAG |
| d__Archaea_d__Archaea_ASV_5759 | 0 | 1 | 0 | 0 | 0 | 0 | 0 | 0 | 0 | 0 | 0 | CCTTTACACTGCACGAAAGTGCGATAAGGGGATCCCGAGTGCGAGGGCATACAGTCCTCGCTTTTGTGTACCGTAAGGTGGTACACGAATAAGAGCTGGGCAAGACCGGTGCCAGCCGCCGCGGTAATACCGGCAGCTCGAGTGATGGCCGTTATTATTGGGCCTAAAGCGTCCGTAGCCGGTTTAGTAAATCTTTTGTGAAATCAAGGGGCTTAACTCCTTGGCGTGCAGGAGAGACTGCTAGACTAGGAACCGGGAGGAGCCAGAAGTATTCCGGGAGTAGCGGTAAAATGCTATAATTCCCGGAGGACTACCTGTGGCGAAGGCGTCTGGCTAGAACGGATTCGACGGTGAGGGACGAAGGCCAGGGGAG |
| d__Archaea_d__Archaea_ASV_5842 | 0 | 0 | 1 | 0 | 0 | 0 | 0 | 0 | 0 | 0 | 0 | ACTTCGCAATGCTGGAAACAGCGACGAGGTGATTCCAAGTGCCGGCACAGAGTGTCGGCTGTTTCTTAGCCTAAAAAGCTTTGAGAGTAAGGGCTGGGTAAGACGGGTGCCAGCCGCCGCGGTAATACCTGCAGCCCAAGTGGTGGTCGTTTTTATTGGGCCTAAAGCGTCCGTAGCCGGACCAGTAAATGCCTGTGTAAATTGTACGGCTTAACCGTGCGAACTGCGGGTAGACTGCTGGTCTTGGGACCGGGAGAGCCAACAAGTACTCCGGGGGTAGAGGTGAAATTCTGTAATCCTAGGGGGACCACCTGTGGCGAAAGCGTCTAACCAGAACGAGTCTGACGGTGAGGGACGAAGCCCTGGGTCA |

| Phylum_Genus_ASV_number | Presence (1) or Absence (0) in each layer L1 | L2 | L3 | L4 | L5 | L6 | L7 | L8 | L9 | L10 | L11 | Sequence |
|---|---|---|---|---|---|---|---|---|---|---|---|---|
| d__Archaea_d__Archaea_ASV_5886 | 0 | 0 | 1 | 0 | 0 | 0 | 0 | 0 | 0 | 0 | 0 | ACTGTACAATGCGGGAAACCGCGATACGGGGACTCCAAGTGCCAGCACAGAGTGTTGGCTGTTTCTGAGCCTAAAAAGCTCAGAGAGTAAGGGCTGGGTAAGACGGGTGCCAGCCGCCGCGGTAATACCTGCAGCTCAAGTGGTGGCCGCTATTATTGGGCCTAAAGCGTCCGTAGCCGGATCAGTAAATGCCTGTGTAAATTGTACCGCTCAACGGTGCGAACTGCGGGTAGACTGCTGGTCTTGGGACCGGGAGAGTCACGAAGTACTCCGGGGGTAGAGGTGAAATTCTGTAATCCTGGGGGGACCACCAGTGGCGAAGGCGTCTAGCTAGAACGAGTCTGACGGTGAGGGACGAAACCCTGGGGCG |
| d__Archaea_d__Archaea_ASV_5897 | 0 | 0 | 0 | 1 | 0 | 0 | 0 | 0 | 0 | 0 | 0 | CCTTTACAATGTACGAAAGTGCGATAAGGGGATCCTTAGTGCTCAAGAGATTTTCTTTTGGGCTTTTGCTAAATGTAAAAAGTTTAGTGAATAAGTGGTGGGCAAGACCGGTGCCAGCAGCCGCGGTAATACCGGCGCCACGAGTGGCAATCTTGTTTATTGGGCCTAAAGCGTTTGTAGCATGCTTTGTAAGTTCTTTGTGAAATCTTTTATGTCGAGTAAAAGTTGTGTATAGAATACTGCATCGCTAGGAACCGAGAGAGGTAAGAGGTATTTCTGGGGGAGCGGTAAAATGTTATAATCCCAGGAAGACGACCTGTGGCGAAGGCGTCTTACCAGAACGGTTTTGACTGTGAGAAACGAAAGCTGGGGGAG |
| d__Archaea_d__Archaea_ASV_6131 | 0 | 0 | 0 | 0 | 1 | 0 | 0 | 0 | 0 | 0 | 0 | ACTTCGCAATGCGCGCAAGCGCGACGAGGGGATTCCAAGTGCATGCACACAGTGTATGCTGTTCAATAGTCTAAAACGCTATTGGAGTAAGGGCTGGGTAAGACTGGTGCCAGCCGCCGCGGTAATACCAGCGGCCCAAGTCGCAGCCATTATTATTGGGTCTAAAACATCCGTAGCTTGCTTGACAAGTCTCTTGTGAAATCTGGCATCTAAAATGTCGGGCGCGCAAGAGATACTGTCTCGCTAGAGATCGGGAGACGTGATGAGTACGTAGGAGGTAGCGGTGAAATGTGTTAATCTCCGACGGACTAACAATAGCGAAGGCACATCACGAGAACGGTTCTGACAGTCAGGGATGAAGGCTAGGGGCG |
| d__Archaea_d__Archaea_ASV_6427 | 0 | 0 | 0 | 0 | 0 | 0 | 0 | 0 | 1 | 0 | 0 | CTTCTGCAATGACCGTAAGGTTGACAGAGCGATCCCGAGTGCTCATGCAAAGCATGGGCTTTTGCCAAGTGTAAGTAGCTCTGGGAATAAGTGGTGGGCAAGACCGGTGCCAGCCGCCGCGGTAATACCGGCGCCACAAGTGATAGCCGCGTTTATTGGGCCTAAAGCATCCGTAGCCGGACAAGTAGGTTTCCAGTGAAATCTAACAGCTTAACTGTTAGGCGTGCTGGGAGCACCGCTTGTCTTGAGACCGGGAGGAGTTAAGAGAATTTCTGGGGGAGCGGTAAAATGTTGTAATCCCAGGAGGACTATCTGTGGCGAAGGCGCTTAACGAGAACGGGTCTGACGGTCAGGGATGAAAGCTGGGGGAG |
| d__Archaea_d__Archaea_ASV_6460 | 0 | 0 | 0 | 0 | 0 | 0 | 0 | 0 | 1 | 0 | 0 | CCTCTGCAATGTACGAAAGTGTGACAGGGGAATTCTACGTGCAAGCTATGCTTGCTTTTCTCAAGTTTAAAACACTTGGGGAATAAGGGCTGGGCAAGACCCGTGCCAGCCGCCGCGGTAATACGGGCAGCCCTAGTGGCGTTCGCGAATATTGGGCCTAAAGCGTACGTAGCCTGTTAAACAAGTTCCTGGTGAAAGCATGCGACTTAATCGTGTGAGTGGCCAGGAAAACTGTTTGACTTGGGACCGGAAAGGGTCGGAGGTATTCCAGGGGTAGGGGTAAAATCCTGTAATCCTTGGAGGACCATCGGTGGCGAAGGCGTCCGACTGGGACGGGTCCGACGGTGAGGTACGAAAGCTAGGGGAG |
| p__Nanoarchaeota_g__1-14-0-10-44-13_ASV_7246 | 0 | 0 | 0 | 1 | 0 | 0 | 0 | 0 | 0 | 0 | 0 | CCTTTACACTGCACGAAAGTGTGATAAGGGAACTCCCAGTGCTTGTGTTCTACACTGGCTTTTGCCAATAGTAAATATATTGGCGAATAAGCGGTGGGTAAGACTGGTGCCAGCCGCCGCGGTAATACCAGCGCCGCAAGTGGCAACCGTGTTTATTGGGCCTAAAGCGTTCGTAGCCGGTTTGGTAAATTTCTTGTGAAATTGATAAGCTTAACTTATCAGCGTGCATGAAAGACTGCTAAACTAGGGACCGGGAAGGGTTAGAAGTATGCCCGGGGGAGCGGTAAAATGTTATAATCCCAGGTAGACTACCTGTGGCGAAGGCGTCTAACTAGAACGGATCCGACGGTGAGGAACGAAAGCTAGGGGAG |
| p__Micrarchaeota_g__Bilamarchaeum_ASV_6914 | 0 | 0 | 0 | 0 | 0 | 0 | 0 | 0 | 0 | 0 | 1 | ACACCAGCGCCCCAAGTAGTATCCACGAATATTGGGCCTAGAGCGTCCGTAGCCGGTCAAGTTAGTCTTCTGTGAAATGTTGGGGCTCAACCTTAACGCGCGCAGGAGATACTGCTAGACTAGGGAACGGAGGAGGACAGGAGTACACCCGGGGTAGGGGTAAAAACCTGTGGATTTAATTTGTAATCCGCAGGATTACAATCCGTTGAGCCTGGGTGGACTACCAGTGGCGAAGGCGCCTGTCTAAGACGTATCCGACGGTGAGGGACGAAAGCTGGGGGAG |
| p__Nanoarchaeota_g__BMS3ABIN17_ASV_6564 | 1 | 0 | 0 | 0 | 0 | 0 | 0 | 0 | 0 | 0 | 0 | AATCTACAATGCGCGCAAGCGTGATAGTTTGAGCCAGAGTGCTTCTCAACAGAGAAGCTTTTGCCAAGTGTAAAAAGCCTGGCGAATAAGGACTGGGTAAGACTAGTGCCAGCCGCCGCGGTAATACTAGCGGTCCAAGTCGCAGCCATTATTATTGGGTCTAAAACATCCGTAGCTTGCCTGGCAAGTTCTTTGTGAAATCTCATGTCTCAAACATGGGGCGGGCAAAGAGTACTACTAGGCTAGAGACAGGGAGAAGTAAGAAGTACGTTCGGAGTAGCGGTAAAATGTGTTAATCCTGGACGGACTCACAATAGCGAAGGCATCTTACTAGAACTGTTCTGACAGTAAGGGATGAAGGCTAGGGGCG |
| p__Nanoarchaeota_g__BMS3ABIN17_ASV_6643 | 0 | 0 | 0 | 0 | 0 | 0 | 0 | 0 | 0 | 1 | 0 | CCTTTACACTGCACGCCAGTGCGATAGGGGGACACCGAGTGCGAGGGCATAGAGCCCTCGCTTTTGTGTACCGTAAGGTGGTACACGAATAAGGACTGGGTAAGACCGGTGCCAGCCGCCGCGGTAATACCGGCGGTCCAAGTCGCAGCCATCATTATTGGGTCTAAAACATCCGTAGCTTGTTCTGTAAGTCTCTTGTGAAATCTGACATCTCAAGTGTCAGGCGGGCAAGAGATACTGCAAGACTAGAGACCGGAAGACGTAACGAGTACAACTGAGGTAGTGGTAAAATACGTTAATCTTGGTTGGACTAACAATGGCGAAGGCACGTTACGAGGACGGATCTGACAGTGAGGGATGAAGGCTAGGGGCG |
| p__Nanoarchaeota_g__BMS3ABIN17_ASV_6654 | 0 | 0 | 0 | 0 | 0 | 0 | 0 | 0 | 0 | 1 | 0 | CTTTTGCAATACGCGCAAGCGTGACAAAGTGAGCCAGAGTGTTTTTCAAATGAAAAGCTTTTGCCAAATCTAAAAAGTTTGGCGAATAAGGACTGGGTAAGACGAGTGCCAGCCGCCGCGGTAATACTCGCGGTCCAAGTCGCAGCCACATTTATTGGGTCTAAAACATCCGTAGCTTGCTTTGTAAGTTTCTTGTGAAATCTCACATCTTAAGTGTGGGGCGAGCAAGAAATACTGCAAAGCTAGAGACTGGAAGACGTAGGAAGTACGTTCAAAGTAGTGGTAAAATATTTTAATCTTGGACGGACTCACAACAGCGAAGGCATCCTACGAGGACAGTTCTGACAGTAAGGGATGAAGGCCAGGGGCG |
| p__Nanoarchaeota_g__BMS3ABIN17_ASV_6658 | 0 | 0 | 0 | 0 | 0 | 0 | 0 | 0 | 0 | 1 | 0 | CCTCCACAATGCGCGAAAGCGTGATGGGGGGAACCCGCGTGCTTGCGCAAAACGTAAGCTTTTCTGGAGTGTAAAAAGACAGATGAATAAGGACTGGGTAAGACCGGTGCCAGCCGCCGCGGTAATACCGGCGGTCCAAGTCGCATCCACAATTATTGGGTCTAAAACATCCGTAGCTTGTTTTGTAAGTCTCTTGTGAAATTCCATATCTCAAGTATGGAACTTGGAAGAGATACTACATGACTAGAGACCGGAAGACGCAAAGAGTATGATTGAGGTAGTGGTAAAATATGTTAATCTCAATTAGACTAACAATAGCGAAGGCACTTTGCGAGTACGGTTCTGACAGTGAGGGATGAAGGCTAGGGGCG |
| p__Nanoarchaeota_g__BMS3ABIN17_ASV_6673 | 0 | 0 | 0 | 0 | 0 | 0 | 0 | 0 | 0 | 1 | 0 | CTTTTGCAATGCACGAAAGTGTGACAAAGCGAGCCAAAGTGCTTTCTGCACAAGAAAGCTTTTCTCCTCTTTAAAAAGGTGGGGGAATAAGGACTGGGCAAGACCGGTGCCAGCCGCCGCGGTAATCCCGGCAGTCCAAGTTGCAGCCAACTTTATTGGGTCTAAAACATCCGTAGCTTGTTAAATAAGTTCCTTGTGAAATCTTACATCTCAAGGGTAAGGCGAGCAAGGAATACTGTTTAGCTAGAGACCGGTAGATGTAAGGAGTACACCTGAGGTAGTGGTAAAATACGTTAATCTTGGGCGGACTAACAATTGGCGAAGGCACCTTACAAGGACGGATCTGACAGTGAGGGATGAAGGCCAGGGGCG |
| p__Nanoarchaeota_g__BMS3ABIN17_ASV_6850 | 0 | 0 | 0 | 0 | 0 | 0 | 0 | 0 | 0 | 0 | 1 | AATCTACAATGCACGAAAGTGTGATAGTTTGAGCCAGAGTGCTTCTCATTGAGAAGCTTTTGCTAAATGTAAAAAGTTTGGCGAATAAGGACTGGGTAAGACAGGTGCCAGCCGCCGCGGTAATACCTGCGGTCCAAGTCGCAGCCATTATTATTGGGTCTAAAACATCCGTAGCTTGCCTGACAAGTTCCTTGTGAAATCCTATGTCTCAAGCATAGGGCTTGCAAGGAGTACTACCTGGCTAGAGACTGGGAGACGTAAGAAGTACGTTCAGAGTAGCGGTAAAATGTTTTAATCCTGAACGGACTCACAATAGCGAAGGCATCTTACGAGAACAGTTCTGACAGTAAGGGATGAAGGCTAGGGGCG |
| p__Nanoarchaeota_g__BMS3ABIN17_ASV_6968 | 0 | 0 | 0 | 0 | 0 | 0 | 0 | 0 | 0 | 0 | 1 | ATTCTGCAATGCGCGCAAGCGTGACAGATTGAACCAGAGTGCTTTTCAATTTGAAAAGCTTTTGTGAAATGCAAAAAGTTTTACGAATAAGGACTGGGTAAGACTAGTGCCAGCCGCCGCGGTAATACTAGCGGTCCAAGTCGCAGCCACAATTATTGGGTCTAAAACATCCGTAGCTTGCTTGATAAGTTTCCTGTGAAATCTTGTTTCTTAAGAACAAGGCGTGCAGGAAATACTGTCAAGCTAGAGACTGGAAGACGCAGAAAGTACGTCTGAGGTAGTGGTAAAATATTTTAATCTTGGATGGACTCACAACAGCGAAAGCATTCTGCGAGGACAGTTCTGACAGTGAGGGATGAAGGCTAGGGGCG |
| p__Nanoarchaeota_g__BMS3ABIN17_ASV_6992 | 0 | 1 | 0 | 0 | 0 | 0 | 0 | 0 | 0 | 0 | 0 | CTTCTGCAATGCGCGAAAGCGTGACAGAGTGAATCAGAGTGCTTTTCAATTTGAAAAGCTTTTGCTTAGTGTAAAAAGCTAGGCGAATAAGGACTGGGTAAGACGAGTGCCAGCCGCCGCGGTAATACTCGCGGTCCAAGTCGCAGCCATTTTTATTGGGTCTAAAACATCCGTAGCTTGCTTAATAAGTTCTTTGTGAAATCCTATATCTCAAGTATAGGGTGTGCAGAGAATACTATTGAGCTAGAGACTGGAAGACGTGATGAGTACAATCAAGGTAGCGGTAAAATGTTTTAATCTTGGTTGGACTAACAAGAGCGAAGGCACATCACGAGGACAGTTCTGACAGTAAGGGATGAAGGCCAGGGGCG |
| p__Nanoarchaeota_g__BMS3ABIN17_ASV_7075 | 0 | 1 | 0 | 0 | 0 | 0 | 0 | 0 | 0 | 0 | 0 | ATTCTGCAATGCGCGCAAGCGTGACAGAGCAAGTCAGAGTGCTTTACCATTAGGTAAAGCTTTTGCCAGATGTAAAAAGTCTGGCGAATAAGGACTGGGTAAGACTGGTGCCAGCCGCCGCGGTAATACCAGCAGTCCAAGTGGCAGCCGTTTTTGTTGGGTCTAAAACATCCGTAGCTTGTCACGTAAGTCTTTTGTGAAATCCTATATCTTAAATATGGGGCGTGCAGGGAATACTGCTTAGCTAGAGACCGGGAGACGTAAAGAGTATGTTCAAAGTAGCGGTAAAATGTTTTAATCTTGAGCAGACTAACAATAGCGAAGGCACTTTACGAGAACGGTTCTGACAGTCAGGGATGAAGGCTAGGGGCG |
| p__Nanoarchaeota_g__BMS3ABIN17_ASV_7268 | 0 | 0 | 0 | 0 | 1 | 0 | 0 | 0 | 0 | 0 | 0 | AATCTACAATACGCGAAAGCGTGATAGTTTGAGTCAGAGTGCGTTTCTTTGAAATGCTTTTGTGAAATGCAAAAAGTTTCACGAATAAGGACTGGGTAAGACGAGTGCCAGCCGCCGCGGTAATACTCGCGGTCCAAGTCGCAGCCATTTTTATTGGGTCTAAAACATCCGTAGCTTGCTTTGTAAGTTCCTTGTGAAATCCTATATCTCAAATATAGGGCGGGCAAGGAGTACTGCTAAGCTAGAGACTGGGAGACGTAAAGAGTACGTTTAAGGTAGCGGTAAAATGTGTTAATCTTAAGCGGACTAACAAGAGCGAAGGCATCTTACGAGGACAGTTCTGACAGTAAGGGATGAAAGCTAGGGGCG |
| p__Nanoarchaeota_g__BMS3ABIN17_ASV_7595 | 0 | 0 | 0 | 0 | 0 | 0 | 0 | 1 | 0 | 0 | 0 | CTTTTGCAATGCACGAAAGTGTGACAAAGCGAGCCAAAGTGTATTTCTTTTTGAAATACTTTTCTTCACCGTAAAAACGTGGAGGAATAAGGACTGGGAGAGACCGGTGCCAGCCGCCGCGGTAATACCGGCAGTCCAAGTTGCATCCCATTATTGGGTCTAAAACATCCGTAGCTTGCCGTTTAAGTCTCTTGTGAAATCTGGCGTCTCAAGCGCCAGGCTTGCGGGAGATACTGGATGGCTAGAGATCGGTAGACGTAAGGAGTACGCTGGAGGTAGTGGTTAAATACGTTAATCTTCGGTGGACTAACAATTTAGCGAAGGCACCTTACGAGGACGAATCTGACAGTGAGGGATGAAGGCCAGGGGCG |
| p__Nanoarchaeota_g__CG1-02-57-44_ASV_7715 | 0 | 0 | 0 | 0 | 0 | 0 | 0 | 0 | 1 | 0 | 0 | CCTTTACAATGCACGAAAGTGTGATAAGGGGATCCCAAGTGCTTATGCATTGCATAGGCTTTTGCCAAGAGTAAATTCCTTGGCGAATAAGTGGTGGGAAAGACTGGTGCCAGCCGCCGCGGTAACACCAGCGCCACGAGTGGGAGTCGCGATTATTGGGCCTAAAGCGTCCGTAGCCGGTTCAGCAAATCTTTTGTGAAATCGTTGGGCTCAACCCAACGGCGCGCAGAGGACACTACTGAACTAGGGACCGGGAGGAGTTGGAGGTACTTGTGAGGGAGCGGTAAAATGCTATAATCTCATGAGGACCACCTGTGGCGAAGGCGTCCAACTAGAACGGATCCGACGGTGAGGGACGAAAGCTGGGGGAG |
| p__Thermoplasmatota_g__EX4484-6_ASV_7195 | 0 | 0 | 1 | 0 | 0 | 0 | 0 | 0 | 0 | 0 | 0 | ACTTCGCAATGCGCGCAAGCGCGACGAGGGGATTCCAAGTGCATGCACACAGTGTATGCTGTTCAATAGTCTAAAACGCTATTGGAGTAAGGGCTGGGTAAGACTGGTGCCAGCCGCCGCGGTAATACCAGCGGCCCTAGTGGTGATCGTTATTATTGGGCCTAAAGCGTCCGTAGCCGGCTCGGTAAATCTCTGGGTAAATCGTTGGGCTTAACTCAGCGAATTCTGGGGAGACTGCCGGGCTTGGGACCGGGAGAGGTCGGAGGTACTCCAGGGGTAGGGGTGAAATCCTGTAATCCTGGACGGACCACCGATGGCGAAAGCACCTCGGAAGGACGGATCCGACAGTGAGGGACGAAAGCTAGGGTCT |
| p__Thermoplasmatota_g__FEN-33_ASV_6851 | 0 | 0 | 0 | 0 | 0 | 0 | 0 | 0 | 0 | 0 | 1 | ACTTCGCAATGGGGGAAACCCTGACGAGGGGATTCCAAGTGCCAGCACACAGTGTTGGCTGTTCTTCTGTCCAAAACGCAGGAGAAGTAAGGGCCGGGTAAGACGGGTGCCAGCCGCCGCGGTAATACCCGCGGCCCGAGTGGTGGTCGATTTTATTGAGCCTAAAACGTCCGTAGCCTGCTTGGTAAATCCTTGGGTAAATCGGCCAGCTTAACTGTCCGAAGTCCGGGGAGACTGCCAGGCTTGGGACCGGGAGAGGTCAGAGGTACTTCTGGGGTAGGGGTAAAATACGTAGATATTAGGAAGAATACCAATGGCGAAAGCAGTTTTCTATCTTAAACCTGACGCTGAGGTACGAAAGCTAGGGTAG |
| p__Thermoproteota_g__GCA-2726865_ASV_6958 | 0 | 0 | 0 | 0 | 0 | 0 | 0 | 0 | 0 | 0 | 1 | CCTTTACACTGCACGCAAGTGCGATAAGGGGACCCCGAGTGCGAGGGCATACCGTCCTCGCTTTTGTCAACCTTAGGGAGGTTGACGAATAAGGAGAGGGCAAGACTGGTGTCAGCCGCCGCGGTAACACCAGCCCCTCGAGTGGTGGGGATGATTATTTGGCCTAAAGCGTCCGTAGCCGGCCGGGTAAGTTTCCCGTTAAATCCAACGTCTCAAGCGTTGGACTGCGGGAAATACTGCCTGGCTAGAGAGTGGGAGAGGTCAGCGGTATTCTGGGGGTAGGGGCGAAATCCTTTGATCCCCGGAGGACCACCAGTGGCGAAGGCTGCTGACTAGAACACGTCTGACGGTCAGGGACGAAAGCTGGGGTAG |
| p__Thermoproteota_g__GCA-2726865_ASV_7633 | 0 | 0 | 0 | 0 | 0 | 0 | 0 | 1 | 0 | 0 | 0 | ATTGGGCAAGACTGGTGTCAGCCGTCGCGGTAACACCAGCCCCTCGAGTGGTGGGGATGATTATTTGGCCTAAAGCGTCCGTAGCCGGCCAGGTAAGTTTTCCGTCAAATCCAACGTCTCAAGCGTTGGGCTGCGGGAAATACTGCCAGGCTAGAGAGTGGGAGAGGTAAGCGGTATTCTGGGGGTAGGGGCGAAATCCTTTGATCCCCGGAGGACCACCAGTGGCGAAGGCTGCTTACTAGAACACGTCTGACGGTCAGGGACGAAAGCTGGGGTAG |

| | Presence (1) or Absence (0) in each layer | | | | | | | | | | |
|---|---|---|---|---|---|---|---|---|---|---|---|
| Phylum_Genus_ASV_number | L1 | L2 | L3 | L4 | L5 | L6 | L7 | L8 | L9 | L10 | L11 | Sequence |
| p__Thermoproteota_g__GCA-2726865_ASV_7671 | 0 | 0 | 0 | 0 | 0 | 0 | 0 | 0 | 1 | 0 | 0 | CCTTCACAATGTGCGAAAGCACGATGAGGGGATCCCAAGTGCTTGCACGTAAGTGTAAGCTGTTTCTTTGTCTAAAACGCATAGAGAGTAAGGGCTGGGTAAGACGGGTGCCAGCCGCCGCGGTAATACCAGCCCCTCGAGTGGTGGGGATGATTATTTGGCCTAAAGCGTCCGTAGCCGGCCAGGTAAGTTTTCCGTCAAATCCAACGTCTCAAGCGTTGGGCTGCGGGAAATACTGCCAGGCTAGAGAGTGGGAGAGGTAAGCGGTATTCTGGGGGTAGGGGCGAAATCCTTTGATCCCTGTGGGACCACCGGTGGCGAAAGCGTCCGACCAGAACGGGTCTGACGGTCAGGGACGAAGCCTTGGGGCG |
| p__Halobacteriota_g__Haladaptatus_ASV_7050 | 0 | 1 | 0 | 0 | 0 | 0 | 0 | 0 | 0 | 0 | 0 | CCTTCACACTGTGGGAAACCACGACGAGGGGATTCCAAGTGCCAACACCCAGTGTTGGCTTTTCATCTGTCTAAAACGCAGGCGGAATAAGAGCTGGGCAAGACCGGTGCCAGCCGCCGCGGTAATACCGGCAGCTCGAGTGATGGCCAATTTTATTGGGCCTAAAGCGTCCGTAGCCTGCCAGACAGGTCCGTCGGGAAATCTGCTCGCTCAACGAGCAGGCGTCCGGCGGAAACCAGCTGGCTTGGGGCCGGAAGACCCAAGGGGTACGTCTGGGGTAGGAGTGAAATCCTGTAATCCTAGACGGACCACCGATCGCGAAAGCACCTTGGGAGGACGGACCCGACGGTGAGGGACGAAAGCTAGGGTCA |
| p__Halobacteriota_g__Haladaptatus_ASV_7444 | 0 | 0 | 0 | 0 | 0 | 1 | 0 | 0 | 0 | 0 | 0 | CCTTTACACTGCACGACAGTGCGATAAGGGGACTCCAAGTGCGAGGGCATACAGTCCTCGCTTTTCTGTACCGTAAGGTGGTACAGGAATAAGAGCTGGGCAAGACCGGTGCCAGCCGCCGCGGTAATACCGGCAGCTCGAGTGATGGCCAATATTATTGGGCCTAAAGCGTCCGTAGCCGGCCGAACAGGTCCGTCGGGAAATCTGCTCGCTCAACGAGCAGGCGTCCGACGGAAACCAGTCGGCTTGGGACCGGAAGACCCGAGGGGTACGTCTGGGGTAGGAGTGAAATCCTGTAATCCTGGGGGGACCACCAGTGGCGAAAGCGTCTAGCTAGAACGGGTCTGACGGTGAGGGACGAAACCCTGGGGCG |
| p__Halobacteriota_g__Halobacterium_ASV_7014 | 0 | 1 | 0 | 0 | 0 | 0 | 0 | 0 | 0 | 0 | 0 | CCTTTACACTGCACGACAGTGCGATAAGGGAACTCCGAGTGCGAGGGCATAGAGCCCTCGCTTTTGTACACCGTAAGGTGGTGTACGAATAAGGGCTGGGCAAGACCGGTGCCAGCCGCCGCGGTAATACCGGCAGCCCAAGTGATGGCCGATATTATTGGGCCTAAAGCGTCCGTAGCCGGCCGGACAAGTCCGTTGGGAAATCCGCACGCTCAACGTGCGGGCGTCCAGCGGAAACTGTTCGGCTTGGGACTGGAAGACCCGAGGGGTACGTCCGGGGTAGGAGTGAAATCCTGTAATCCTTGGGGGACCACCGGTGGCGAAAGCGTCCGGCCAGAACGGGTCCGACGGTAAGGGACGAAGCCCTGGGGCG |
| p__Halobacteriota_g__Halobacterium_ASV_7100 | 0 | 0 | 1 | 0 | 0 | 0 | 0 | 0 | 0 | 0 | 0 | CCTTTACACTGCACGACAGTGCGATAGGGGGACTCCGAGTGCGAGGGCGTATAGTCCTCGCTATTCTGAACCGTAAGGTGGTTCAGGAAAAGGGCTGGGCAAGACCGGTGCCAGCCGCCGCGGTAATACCGGCAGCCCGAGTGATGGCCAATATTATTGGGCCTAAAGCGTCCGTAGCCGGCCGGACAAGTCCGTTGGGAAATCTGCTCGCTCAACGAGCAGGCGTCCAGCGGAAACTGTCCGGCTTGGGGCCGGAAGACTCGAGGGGTACGTCCGGGGTAGGAGTGAAATCCTGTAATCCTGGACGGACCACCGGTGGCGAAAGCGCCTCGAGAAGACGGACCCGACGGTGAGGGACGAAAGCTAGGGTCT |
| p__Halobacteriota_g__Halobellus_3029_ASV_7693 | 0 | 0 | 0 | 0 | 0 | 0 | 0 | 0 | 1 | 0 | 0 | CCTCCACAATGCGCGAAAGCGTGATGGGGGGAACCCGCGTGCTTGCGCAAAACGTAAGCTTTTCTGGAGTGTAAGTAGCTCTGGGAATAAGTGGTGGGCAAGACCGGTGCCAGCCGCCGCGGTAATACCGGCAGCTCAAGTGATGGCCGATCTTATTGGGCCTAAAGCGTCCGTAGCTGGCCCTGCAAGTCCGTCGGGAAATCCATCAGCTCAACTGATGGGCGTCCGGCGGAAACTCCAGGGCTTGGGACCGGAAGGCTCGAGGGGTACGTCCGGGGTAGGAGTGAAATCCCGTAATCCCGAACGGACCACCGATGGCGAAAGCACCTCGAGAAGACGGCTCCGACAGTGAGGGACGAAAGCTAGGGTCT |
| p__Halobacteriota_g__Halodesulfurarchaeum_ASV_7492 | 0 | 0 | 0 | 0 | 0 | 0 | 1 | 0 | 0 | 0 | 0 | CCTTTACACTGCACGAAAGTGCGATAGGGGGACCCCGAGTGCGAGGGCATACAGTCCTCGCTTTTGTACACCGTAAGGTGGTGTACGAATAAGGGCTGGGCAAGACCGGTGCCAGCCGCCGCGGTAATACCGGCAGCCCAAGTGATGGCCGTTATTATTGGGCCTAAAGCGTCCGTAGCCGGCCGGACAAGTCCGTTGGGAAATCCGCACGCTCAACGTGCGGGCGTCCAGCGGAAACTGTCCGGCTTGGGACCGGAGGACCCGAGGGGTACGTCCGGGGTAGGAGTGAAATCCTGTAATCCCGGACGGACCGCCTGTGGCGAAAGCGCCTCGGGAAGACGGATCCGACGGTGAGGGACGAAAGCCAGGGTCT |
| p__Halobacteriota_g__Halohasta_ASV_7084 | 0 | 1 | 0 | 0 | 0 | 0 | 0 | 0 | 0 | 0 | 0 | TCTTCACAATGCACGAAAGTGCGATGGGGGAACTCCGAGTGCGTCTTTCCCACAAGGAAGGACGCTTTTCTGCACTCTAAGAAGGTGCAGGAACAAGGGGCGGGCAAGACTGCTGCCAGCCGCCGCGGTAATAGCAGCACCCCAAGTGGTGGTCAGTATTATTGGGCTTAAAGCGTCCGTAGCTGGCCGCGCAAGTCCGTCGGGAAATCCACCTGCCCAACAGGTGGGCGTCCGGCGGAAACTGCGTGGCTTGGAACCGGAAGGCTCAGAGGGTACGTCCGGGGTAGGAGTGAAATCCCGTAATCCCGGACGGACCACCGATGGCGAAAGCACTCTGAGAGGACGGATTCGACAGTGAGGGACGAAAGCTAGGGTCT |
| p__Halobacteriota_g__Halopiger_A_ASV_7153 | 0 | 0 | 1 | 0 | 0 | 0 | 0 | 0 | 0 | 0 | 0 | CCTTTACAATGCACGCAAGTGCGATAAGGGGACTCCGAGTGCGGTCTTCCTACACGGAAGGCCGCTTTTCTGCACTGTAAGAAGGTGCCGGAACAAGGGGTGGGCAATACTGCTGCCAGCCGCCGCGGTAATACCGGCAGCCCGAGTGATGACCGCTATTATTGGGCCTAAAGCGTCCGTAGCTGGCCAGGCAAGTCCATCGGGAAATCCGCGTGCCCAACACGCGGGCGTCCGGTGGAACCTGCCTGGCTTGGGACCGGAAGACCAGAGGGGTACGTCCGGGGTAGGAGTGAAATCCCGTAATCCTGGACGGACCGCCGGTGGCGAAAGCGCCTCGAGAGGACGGATCCGACGGTGAGGGACGAAAGCTAGGGTCT |
| p__Halobacteriota_g__Haloprofundus_ASV_7179 | 0 | 0 | 1 | 0 | 0 | 0 | 0 | 0 | 0 | 0 | 0 | CCTTTACACTGCACGCCAGTGCGACAAGGGGACCCCGAGTGCGAGGGCATACAGTCCTCGCTTTTCCCGACCGTAAGAAGGTCGGAGAAGAAGAGCTGGGCAAGACCGGTGCCAGCCGCCGCGGTAATACCGGCAGCTCGAGTGATGACCGCTGTTATTGGGCCTAAAGCGTCCGTAGCTGGCCGCGTAAGTCCATCGGGAAATCCATCCGCTCAACGGATGGGCGTCCGGTGGAAACTACACGGCTTGGAGCCGGAAGACCCAAGGGGTACGTCCGGGGTAGGAGTGAAATCCCGTAATCCTGGACGGACCACCGATGGCGAAAGCGTGGGGAG |
| p__Halobacteriota_g__Halorientalis_ASV_7172 | 0 | 0 | 1 | 0 | 0 | 0 | 0 | 0 | 0 | 0 | 0 | CCTTTACAATGCACGAAAGTGTGATAAGGGGATCCCAAGTGCTCATGCACTGCATGGGCTTTTGCCAAGAGTAAATATCTTGGCGAATAAGTGGTGGGAAAGACTGGTGCCAGCAGCCGCGGTAACACCAGCGCCACGAGTGGCAGACATTATTATTGGGCCTAAAGCGTCCGTAGCCGGCCACGCAAGTCCGTCGGGAAATCTGCCCGCTTAACGGGCAGGCGTCCGGCGGAAACTGCGTGGCTTGGGGCCGGGAGATCCAAGGGGTACGTCCGGGGTAGGAGTGAAATCCTGTAATCCTGGACGGACCACCGATGGCGAAAGCACCTTGGAAAAACGGACCCGACGGTGAGGGACGAAAGCTAGGGTCT |
| p__Halobacteriota_g__Halorubrum_ASV_6847 | 0 | 0 | 0 | 0 | 0 | 0 | 0 | 0 | 0 | 0 | 1 | CCTTTACAATGCACGAAAGTGTGATAAGGGAACCCCCAGTGCCAGTGGAAACGCTGGCTTTTTATGAGGGTAAGGTCCTCATAGAATAAGCAGTGGGTAAGACCGGTGCCAGCCGCCGCGGTAACACCGGCGCTGCTAGTGGGAGTCGCGATTATTGGGCCTAAAGCGTTCGTAGCCGGTTTGGTAAATTCTCTGTGAAATCCCGTAATCCTGGACGGACCACCGATGGCGAAAGCACGTCGCGAGAACGGATCCGACAGTGAGGGACGAAAGCCAGGGTCT |
| p__Halobacteriota_g__Halorussus_ASV_6885 | 0 | 0 | 0 | 0 | 0 | 0 | 0 | 0 | 0 | 0 | 1 | TCTTTACACTGCACGACAGTGCGATAAGGGGACTCCGAGTGCGAGGGCATATAGTCCTCGCTTTTCTGTACCGTAAGGTGGTACAGGAATAAGGGCTGGGCAAGACCGGTGCCAGCCGCCGCGGTAATACCGGCAGCCCGAGTGATGGCCGCTATTATTGGGCCTAAAGCGTCCGTAGCCGGCCAGACAGGTCCGTCGGGAAATCCACGCGCTCAACGCGTGGGCGTCCGGCGGAAACCAGCTGGCTTGGGGCCGGAAGACCTGAGGGGTAGAGGTGAAATTCTGTAATCCTGAGGGGACCACCAGTGGCGAAGGCGTCTAGCTAGAACGGGTCTGACGGTGAGGGACGAAGCCCTGGGGCG |
| p__Halobacteriota_g__Halorussus_ASV_6919 | 0 | 0 | 0 | 0 | 0 | 0 | 0 | 0 | 0 | 0 | 1 | ACTTTACACTGCACGACAGTGCGATAAGGGAACTCCGAGTGCGAGGGCATATAGTCCTCGCTTTTCTGCACCGTAAGGTGGTGCACGAATAAGAGCTGGGCAAGACCGGTGCCAGCCGCCGCGGTAATACCGGCAGCTCAAGTGATGGCCGCTATTATTGGGCCTAAAGCGTCCGTAGCCGGCCAGACAGGTCCGTCGGGAAATCCAGTGGCTTAACCACTGGGCGTCCGGCGGAAACCAGCTGGCTTGGGGCCGGAAGATCTGAGGGGTACGTCCGGGGTAGGAGTGAAATCCCGTAATCCTGGACGGACCGCCGGTGGCGAAAGCGCCTCAGAAGGACGGACCCGACGGTGAGGGACGAAAGCTAGGGTCA |
| p__Halobacteriota_g__Halorussus_ASV_7048 | 0 | 1 | 0 | 0 | 0 | 0 | 0 | 0 | 0 | 0 | 0 | ACTTTACACTGCACGACAGTGCGATAAGGGAACTCCGAGTGCGAGGGCATATAGTCCTCGCTTTTCTGCACCGTAAGGTGGTGCACGAATAAGAGCTGGGCAAGACCGGTGCCAGCCGCCGCGGTAATACCGGCAGCTCAAGTGATGGCCGCTATTATTGGGCCTAAAGCGTCCGTAGCCGGCCAGACAGGTCCGTCGGGAAATCCAGTGGCTTAACCACTGGGCGTCCGGCGGAAACCAGCTGGCTTGGGGCCGGAAGACCTGAGGGGTACGTCCGGGGTAGGAGTGAAATCCCGTAATCCTGGACGGACCGCCGGTGGCGAAAGCGCCTCAGGAAGACGGACCCGACGGTGAGGGACGAAAGCTAGGGTCA |
| p__Halobacteriota_g__Halorussus_ASV_7068 | 0 | 1 | 0 | 0 | 0 | 0 | 0 | 0 | 0 | 0 | 0 | ACTTTACACTGCACGACAGTGCGATAAGGGAACTCCGAGTGCGAGGGCATATAGTCCTCGCTTTTCTGCACCGTAAGGTGGTGCACGAATAAGAGCTGGGCAAGACCGGTGCCAGCCGCCGCGGTAATACCGGCAGCTCAAGTGATGGCCGCTATTATTGGGCCTAAAGCGTCCGTAGCCGGCCAGACAGGTCCGTCGGGAAATCCACTCGCTCAACGAGTGGGCGTCCGGCGGAAACCAGCTGGCTTGGGGCCGGAAGATCCGAGGGGTACGTCCGGGGTAGGAGTGAAATCCCGTAATCCTGGACGGACCGCCGGTGGCGAAAGCGCCTCGGAAAGACGGACCCGACGGTGAGGGACGAAAGCTAGGGTCA |
| p__Halobacteriota_g__Halorussus_ASV_7528 | 0 | 0 | 0 | 0 | 0 | 0 | 1 | 0 | 0 | 0 | 0 | ACTTTACACTGCACGACAGTGCGATAAGGGAACTCCGAGTGCGAGGGCATATAGTCCTCGCTTTTCTGCACCGTAAGGTGGTGCACGAATAAGAGCTGGGCAAGACCGGTGCCAGCCGCCGCGGTAATACCGGCAGCTCAAGTGATGGCCGCTATTATTGGGCCTAAAGCGTCCGTAGCCGGCCAGACAGGTCCGTCGGGAAATCCACGCGCTCAACGCGTGGGCGTCCGGCGGAAACCAGCTGGCTTGGGGCCGGAAGATCTGAGGGGTACGTCCGGGGTAGGAGTGAAATCCGTAGATATCTGGAGGAACACCAGAGGCGAAGGCGGCCACCTGGACCGAAACTGATGCTGAGGTGCGAAAGCGTGGGGAG |
| p__Halobacteriota_g__Halorussus_ASV_7530 | 0 | 0 | 0 | 0 | 0 | 0 | 1 | 0 | 0 | 0 | 0 | ACTTTACACTGCACGACAGTGCGATAAGGGAACTCCGAGTGCGAGGGCATATAGTCCTCGCTTTTCTGCACCGTAAGGTGGTGCACGAATAAGAGCTGGGCAAGACCGGTGCCAGCCGCCGCGGTAATACCGGCAGCTCAAGTGATGGCCGCTATTATTGGGCCTAAAGCGTCCGTAGCCGGCCAGACAGGTCCGTCGGGAAATCCACGCGCTCAACGCGTGGGCGTCCGGCGGAAACCAGCTGGCTTGGGGCCGGAAGATCTGAGGGGACGGATTCGACAGTGAGGGACGAAAGCTAGGGTCT |
| p__Halobacteriota_g__Halostagnicola_ASV_7214 | 0 | 0 | 0 | 1 | 0 | 0 | 0 | 0 | 0 | 0 | 0 | CCTTTACACTGCACGAGAGTGCGATAAGGGGACTCCGAGTGCGAGGGCATATAGCCCTCGCTTTTCACGACCGTAAGGTGGTCGTAGAATAAGTGCTGGGCAAGACCGGTGCCAGCCGCCGCGGTAATACCGGCAGCACGAGTGATGACCGCTATTATTGGGCCTAAAGCGTCCGTAGCTGGCCACGCAAGTCCATCGGGAAATCTCTCCGCTCAACGGAGAGGCGTCCGGTGGAAACTGTGTGGCTTGGGACCGGAAGATCCAGAGGGTACGTCTGGGGTAGGAGTGAAATCCCGTAATCCGAGACGGACCACCGGTGGCGAAAGCGCTCTGGAAAGACGGATCCGACGGTGAGGGACGAAAGCTCGGGTCA |
| p__Halobacteriota_g__Halovenus_ASV_7498 | 0 | 0 | 0 | 0 | 0 | 0 | 1 | 0 | 0 | 0 | 0 | ATTGCGCAATGGGGGAAACCCTGACGCAGCAACGCCGCGTGAAGGAAGAAGGCCTTCGGGTTGTAAACTTTTTTGATTGGGGAAGAAAAAAATGACGGTATCCAAGGAGGAAGCCCCGGCTAACTACGTGCCAGCAGCCGCGGTAATACCGGCAGTTCGAGTGATGGCCGCTTTTATTGGGCCTAAAGCGTTCGTAGCTGGCCAGTCAAGTCTGTCGGGAAATCCAGTGGCTTAACCACTGGACGTCCGGCGGAAACTGATTGGCTTGGGACCGGGAGACTCGAGGAGTACGTTCGGGGTAGGAGTGAAATCCCATAATCCTGAACGGACTACCGATGGCGAAAGCACCTCGAGAGCACGGATCCGACAGTGAGGAACGAAAGCTGGGGTCT |
| p__Methanobacteriota_A_1229_g__Methanobrevibacter_A_ASV_6645 | 0 | 0 | 0 | 0 | 0 | 0 | 0 | 0 | 0 | 1 | 0 | CCTCCGCAATGTGAGAAATCGCGACGGGGGGATCCCAAGTGCCATTCTTAACGGGATGGCTTTTCATTAGTGTAAAGAGCTTTTGGAATAAGGGCTGGGCAAGACCGGTGCCAGCCGCCGCGGTAACACCGGCAGCTCTAGTGGTAGCAGTTTTTATTGGGCCTAAAGCGTCCGTAGCCGGTTTAATAAGTCTCTGGTGAAATCCTGTAGCTTAACTGTGGGAATTGCTGGAGATACTATTAGACTTGAGACCGGGAGAGGTTAGAGGTACTCCCAGGGTAGAGGTGAAATTCTGTAATCCTGGGAGGACCGCCTGTTGCGAAGGCGTCTAACTGGAACGGTTCTGACGGTGAGGGACGAAAGTTAGGGGCG |
| p__Halobacteriota_g__Methanohalobium_ASV_6580 | 1 | 0 | 0 | 0 | 0 | 0 | 0 | 0 | 0 | 0 | 0 | CCGCGATAAGGGGACACTGAGTGCCAGCATATAATGCTGGCTGTCCGTATGTGTAAAATACATGCGTTAGCAAGGGCCGGGCAAGACCGGTGCCAGCCGCCGCGGTAACACCGGCGGCCCGAGTGGTGGTCGCTATTATTGGGTCTAAAGGGTCCGTAGCCGGTTTGATCAGTCCATTGGGAAATCTGGTGGCTTAACCGCCAGAATTCCAATGGATACTGTCAGACTTGGGACCGGGAGAGGTGAGAGGTACTACAGGGGTAGGAGTGAAATCTTGTAATCCCTGTAGGACCACCAGTTGCGAAGGCGTCTCACCAGAACGGGTCCGACGGTGAGGGACGAAAGCTGGGGGCA |
| p__Halobacteriota_g__Methanohalophilus_ASV_7021 | 0 | 1 | 0 | 0 | 0 | 0 | 0 | 0 | 0 | 0 | 0 | CCTTTACACTGCGGGAAACCGCGATAAGGGGACACCGAGTGCCAGCATACAATGCTGGCTGTCCACATGTGTAAATAGCATGTGATAGCAAGGGCCGGGCAAGACCGGTGCCAGCCGCCGCGGTAACACCGGCGGCCCGAGTGGTAATCACTATTATTGGGTCTAAAGGGTCCGTAGCCGGTTTAATCAGTCTTCCGGGAAATCTGACAGCTCAACTGTTAGGCTTCCGGGGGATACTGTTAGACTTGGGACCGGGAGAGGTAAGAGGTACTACAGGGGTAGGAGTGAAATTCGTAGATATTCGGAGGAACACCAGTGGCGAAGGCGGCTCACTGGTCCATTACTGACGCTGAGGTGCGAAAGCGTGGGGAG |
| p__Halobacteriota_g__Methanohalophilus_ASV_7493 | 0 | 0 | 0 | 0 | 0 | 0 | 1 | 0 | 0 | 0 | 0 | CCTTTACACTGCGGGAAACCGCGATAAGGGGACACCGAGTGCCAGCATACAATGCTGGCTGTCCACATGTGTAAATAGCATGTGATAGCAAGGGCCGGGCAAGACCGGTGCCAGCCGCCGCGGTAACACCGGCGGCCCGAGTGGTAATCACTATTATTGGGTCTAAAGGGTCCGTAGCCGGTTTAATCAGTCTTCCGGGAAATCTGACAGCTCAACTGTTAGGCTTCCGGGGGATACTGTTAGACTTGGGACCGGGAGAGGTAAGAGGTACTACAGGGGTAGGAGTGAAATCCCGTAATCCCGGACGGACCGCCGGTGGCGAAAGCGCCCTGGAAGGACGGACCCGACGGTGAGGGACGAAAGCTAGGGTCA |

| Phylum_Genus_ASV_number | Presence (1) or Absence (0) in each layer | | | | | | | | | | | Sequence |
|---|---|---|---|---|---|---|---|---|---|---|---|---|
| | L1 | L2 | L3 | L4 | L5 | L6 | L7 | L8 | L9 | L10 | L11 | |
| p__Halobacteriota_g__Methanospirillum_ASV_7729 | 0 | 0 | 0 | 0 | 0 | 0 | 0 | 0 | 1 | 0 | 0 | ACTTCACACTGGGGGCAACCCCGATGAGGGAATTCCTAGTGCCTGCGCAATTGCGTTTGGCTTTTCTTTAGTGTAGATAACTGGTGAAGAAAGGGCCGGGCAAGACCGGTGCCAGCCGCCGCGGTAATACCGGCGGCTCGAGTGGTGGCCGCTATTACTGGGCTTAAAGGGTCCGTAGCTTGATAACTGGGTCCCTTGAGAAATCCGCTGGCTTAACTTGCGGGCGTTCAGGGGAAACTTGTTTTCTTGGGACCGGGAGAGGTGAGGGGTACTGCCGGGGTAGGAGTGAAATCCTGTAATCCCGGTGGGACCACCTATGGCGAAGGCATCTCACCAGAACGGGTCCGACAGTGAGGGACGAAAGCTGGGGGAG |
| p__Nanohaloarchaeota_g__Nanohalobium_ASV_6644 | 0 | 0 | 0 | 0 | 0 | 0 | 0 | 0 | 0 | 1 | 0 | CCTTCACAATGCACGACAGTGCGATGGGGGGACTCCGAGTGTAGTCTTCCCACAAGGAAGGCTACTTTTCTGTACTGTAAGGAGGTACCGGAACAAGGGGTGGGCAATACTGCTGCCAGCCGCCGCGGTAATAGCAGAACCCCGAGTGGTAGTCGTTATTATTGGGCTTAAAGTGTTCGTAGCCGGCCGTGCAAGTCCGCTGTGAAATCCAAGCGCTTAACGTTTGGGCGTGCAGCGGATACTGCACTGGCTAGGGACCGGACGGTGTCGAGGCTATTCCAGGAGTAGGGGTGAAATCCTAAAATTCCTGGAGGAGCGCCTGTGGCGGAAGCGCTCGACAAGTACGGATCCGACGGTGAGGAACGAAAGCCAGGGGAG |
| p__Nanohaloarchaeota_g__Nanohalobium_ASV_7212 | 0 | 0 | 0 | 1 | 0 | 0 | 0 | 0 | 0 | 0 | 0 | CCTTCACAATGCACGAAAGTGCGATGGGGGGACTCCGAGTGCGTTCTTCTCACAAAGAAGGGCGCTTTTCTGCACTGTAAGGAGGTGCCGGAAAAAGGGGTGGGCAATACTGCTGCCAGCCGCCGCGGTAATAGCAGAGCCCCAAGTGGTAGTCGTTATTATTGGGTTTAAAGTGTTCGTAGCCGGCCGTGCAAGTCCGCTGTGAAATCCAGGCGCTTAACGTCTGGGCGTGCAGCGGAAACTGCACTGGCTAGGGACCGGACGGTGTCGAGGCTATTCCCGGGGTAGGAGTGAAATCCTAAAATCCCGGGAGGAGCGCCTGTGGCGTAAGCGCTCGACAAGGACGGATCCGACGGTGAGGGACGAAAGCCAGGGGAG |
| p__Nanohaloarchaeota_g__Nanohalobium_ASV_7674 | 0 | 0 | 0 | 0 | 0 | 0 | 0 | 0 | 1 | 0 | 0 | CCTTCACAATGCACGACAGTGCGATGGGGGGATTCCGAGTGTGGTCTTCCCACAAGGAAGGCTACTTTTCTGTACTGTAAGGAGGTACTGGAAAAAGGGGTGGGCAATACTGCTGCCAGCCGCCGCGGTAATAGCAGAACCCCAAGTGGTAGTCGTTATTATTGGGCTTAAAGTGTTCGTAGCCGGCCGTGCAAGTCCGCTGTGAAATCCAGGCGCTTAACGTCTGGGCGTGCAGCGGATACTGCACTGGCTAGGGACCGGACGGTGTCGAGGCTATTCCAGGAGTAGGGGTGAAATCCTAAAATTCCTGGAGGAGCGCCTGTGGCGGAAGCGCTCGACAAGTACGGATCCGACGGTGAGGAACGAAAGCCAGGGGAG |
| p__Halobacteriota_g__Natrinema_ASV_7353 | 0 | 0 | 0 | 0 | 1 | 0 | 0 | 0 | 0 | 0 | 0 | CCTTTACACTGCACGCGAGTGCGATAAGGGGACTCCGAGTGCGAGGGCATATAGTCCTCGCTTTTCACCACCGTAGGAAGGTGGTAGAATAAGTGCTGGGCAAGACCGGTGCCAGCCGCCGCGGTAATACCGGCAGCACGAGTGATGACCGCTATTATTGGGCCTAAAGCGTCCGTAGCTGGCCACGCAAGTCTATCGGGAAATCTGCGCGCTTAACGCGCAGGCGTCCGGTGGAAACTGCGTGGCTTGGGACCGGAAGACCAGAGGGGTACGTCCGGGGTAGGAGTGAAATCCCGTAATCCTGGACGGACCGCCGATAGCGAAAGCACCTCGGGAAGACGGATCCGACAGTGAGGGACGAAAGCTGGGGTCA |
| p__Thermoplasmatota_g__Natronoplasma_ASV_7085 | 0 | 1 | 0 | 0 | 0 | 0 | 0 | 0 | 0 | 0 | 0 | ACTTTACAATGCACGAAAGTGCGATAAGGGGAATCCAAGTGGTAGTTCTACACAAAGGAGCTACCTTTTGCCAAGTGTAAGGAGCTTGGTGAATAAGGGGTGGGCAAGACGGCTGCCAGCCGCCGCGGTAATAGCCGCACCCCAAGTGGTGACCACGATTATTGGGCTTAAAGCGTCCGTAGCCGGCTGAGTAAATGCCTGGGTAAATCGTGTCGCTTAACGATACGAATTCCGGGTAGACTGCTTAGCTTGAGACTGGGAGAAGCTAGAGGTACTCCTGGGGTAGAGGTGAAATTCTGTAATCCTGGGGGGACCACCAGTGGCGAAAGCGTCTAGCTAGAACGGGTCTGACGGTGAGGGACGAAACCCTGGGGCG |
| p__Nanoarchaeota_g__PWVO01_ASV_7025 | 0 | 1 | 0 | 0 | 0 | 0 | 0 | 0 | 0 | 0 | 0 | CCTTTACAATGCACGAAAGTGTGATAAGGGAATCCCAAGTGCCTTGCTTTCGCAAGGCTTTTCTTAAGAGTAAATATCTTAGGGAATAAGTGGTGGGAAAGACCGGTGCCAGCAGCCGCGGTAATACCGGCGCCACAAGTGGTGATCATTTTTATTGGGCCTAAAGCGTTCGTAGCCGGTTTGATAAATCTTCTGTGAAATTGTCAGGCTTAACCTGACAGCTTGCAGAAGACACTATTAAACTAGGGACCGGGAGGAGTCAGAGGTATTCTTGGGGGAGCGGTAAAATGCTATAATCCCAGGAAGACTAACAGTGGCGAAGGCGTCTGACCAGAACGGATCCGACGGTGATTGGACGAAAGTCAGGGGAG |
| p__Nanoarchaeota_g__PWVO01_ASV_7426 | 0 | 0 | 0 | 0 | 1 | 0 | 0 | 0 | 0 | 0 | 0 | CCTTTACAATGCACGAAAGTGTGATAAGGGAATCCCGAGTGCCTTGCTTTCGCAAGGCTTTTCTTAAGAGTAAATATCTTAGGGAATAAGTGGTGGGAAAGACCGGTGCCAGCAGCCGCGGTAATACCGGCGCCACAAGTGGTGATCATTTTTATTGGGCCTAAAGCGTTCGTAGCCGGTTTGATAAATCTTCTGTGAAATTGTCAGGCTTAACCTGACAGCGTGCAGGAGACACTGTCAAACTTGGGACCGGGAGGAGTCAGAGGTATTCCTTGGGGAGCGGTAAAATGCTATAATCCTCGGAAGACTAACTGTGGCGAAGGCGTCTGACCAGAACGGGTCTGACGGTGAGGGATGAAAGCTAGGGGAG |
| p__Halobacteriota_g__Saliphagus_ASV_6983 | 0 | 1 | 0 | 0 | 0 | 0 | 0 | 0 | 0 | 0 | 0 | CCTTTACACTGCACGCAAGTGCGATAAGGGGACCCCGAGTGCGAGGGCATACAGTCCTCGCTTTTGACACCCGTAAGGAGGTTGTCGAATAAGGGCTGGGCAAGACCGGTGCCAGCCGCCGCGGTAATACCGGCAGCCCAAGTGATGGCCGATCTTATTGGGCCTAAAGCGTCCGTAGCCGGCCACGCAAGTCCGTCGGGAAATCCGCGCGCCTAACGCGCGGGCGTCCGGCGGAAACTGCGTGGCTTGGGACCGGAAGACTCGAGGGGTACGTCCGGGGTAGGAGTGAAATCCTGTAATCCTGGACGGACCACCGGTGGCGAAAGCGCCTCGAGAGGACGGATCCGACGGTGAGGGACGAAAGCTAGGGTCA |
| p__Halobacteriota_g__Saliphagus_ASV_7005 | 0 | 1 | 0 | 0 | 0 | 0 | 0 | 0 | 0 | 0 | 0 | CCTTTACACTGCACGAGAGTGCGATAAGGGGACTCCAAGTGCGAGGGCATATCGTCCTCGCTTTTCACTACCGTAAGGAGGTAGTAGAATAAGTGCTGGGCAAGACCGGTGCCAGCCGCCGCGGTAATACCGGCAGCACGAGTGATGACCGCTATTATTGGGCCTAAAGCGTCCGTAGCCGGCCTCGCAAGTCCGTCGGGAAATCCGCGCGCTCAACGCGCGGGCGTCCGGCGGAAACTGCGTGGCTTGGGACCGGAAGACTCAGGGGGTACGTCCGGGGTAGGAGTGAAATCCTGTAATCCTGGACGGACCACCGGTGGCGAAAGCGCCCTGAGAGGACGGATCCGACGGTGAGGGACGAAAGCTAGGGTCA |
| p__Halobacteriota_g__Saliphagus_ASV_7127 | 0 | 0 | 1 | 0 | 0 | 0 | 0 | 0 | 0 | 0 | 0 | CCTTTACACTGCACGCCAGTGCGATAGGGGGACCCCGAGTGCGAGGGCATACAGTCCTCGCTTTTGTACACCGTAAGGAGGTGTACGAATAAGTGCTGGGCAAGACCGGTGCCAGCCGCCGCGGTAATACCGGCAGCACAAGTGATGGCCGCTATTATTGGGCCTAAAGCGTCCGTAGCCGGCCGAACAAGTCCGTCGGGAAATCCATCCGCCTAACGGATGGGCGTCCGGCGGAAACTGTTCGGCTTGGGACAGGAAGACTCGAGGGGTACGTCCGGGGTAGGAGTGAAATCCCGTAATCCCGGACGGACCACCGATGGCGAAAGCACTCTGAGAGGACGGATTCGACAGTGAGGGACAAAAGCTAGGGTCT |
| p__Asgardarchaeota_g__SDNM01_ASV_6626 | 1 | 0 | 0 | 0 | 0 | 0 | 0 | 0 | 0 | 0 | 0 | CCTCCGCAATACACGAAAGTGTGACGGGGTTACCCAAAGTGTCCATCATTGGACTGTGGTAGGTGAGTAATGTCCCCTACTAGAAAGGAGAGGGCAAGGCTGGTGCCAGCCGCCGCGGTAAAACCAGCTCTTCAAGTGGTCGGGATAATTATTGGGCTTAAAGCGTCCGTAGCCGGTTTAGTAAGTTCCTGGTAAAATCGGGTAGCTTAACTATCTGTATGCTAGGAATACTGCTATACTAGAGGGCGGGAGAGGTCTGAGGTACTACAGGGGTAGGGGTGAAATCTTATAATCCTTGTAGGACCACCAGTGGCGAAGGCGTCAGACTGGAACGCGCCTGACGGTGAGGGACGAAAGCCAGGGGAG |
| p__Asgardarchaeota_g__SDNM01_ASV_6938 | 0 | 0 | 0 | 0 | 0 | 0 | 0 | 0 | 0 | 0 | 1 | ACTTCGCAATGCGCGCAAGCGTGACGAGGCTACCCGGAGTGTCTTCGCCTAAGGAAGACTGTGGCGTGTGCGTAAACTGTCGCGCTAGAAAGGAGAGGGCAAGGCTGGTGCCAGCCGCCGCGGTAAAACCAGCTCTTCAAGTCGTCGGGATAATTATTGGGCTTAAAGTGTTCGTAGCCGGTTAAGTAAGTTCTTGGTAAAACCTGGCAGCTCAACTGTCAGTATGCTAGGAATACTGCTTAACTAGAGGGCGGGAGAGGTCCGAGGTACTCCAGGGGTAGGGGTGAAATCCAATAATCCTTGGGGGACCACCAGTGGCGAAGGCGTCGGACTGGAACGCGCCTGACGGTGAGGGACGAAAGCCAGGGGAG |
| p__Asgardarchaeota_g__SDNM01_ASV_7226 | 0 | 0 | 0 | 1 | 0 | 0 | 0 | 0 | 0 | 0 | 0 | CCTTTACAATGCACGAAAGTGTGATAAGGTTACCCAAAGTGGCACAAAAATGGTTGTGTCTTTTACGAGGTGTAAAATACCTTGTGAATAAGGAGAGGGCAAGGCTGGTGCCAGCCGCCGCGGTAAAACCAGCTCTTCAAGTGGTCGGGAATTTTATTGGGCTTAAAGTGTCCGTAGCCTGTTTGATAAGTTCCTGGTTAAATCTAGTGGCTTAACCATTAATTAGCTAGGAATACTGTCTTACTAGAGGGTGGGAAAGGTCTGAGGTACTCCAGGGGTAGCGGTGAAATGCGATAATCCTTGGGGGACCACCAGTGGCGAAGGCGTCAGACTGGAACACGCCTGACGGTGAGGGACGAAAGCTAGGGGAG |
| p__Thermoplasmatota_g__SM1-50_ASV_7604 | 0 | 0 | 0 | 0 | 0 | 0 | 0 | 1 | 0 | 0 | 0 | ACTTTACAATGCGGGAAACCGCGATAAGGGGACACTGAGTGCCAGCATACAATGTTGGCTGTCCGCATGTATAAATAGCATGTGTTAGCAAGGGCCGGGCAAGACCGGTGCCAGCCGCCGCGGTAACACCGGCGGCCCGAGTGGTGGCCACTATTATTGGGCCTAAAACGTCCGTAGCCTGTTTGATAAATGCCTGGGTAAATCGTATAGCTCAACTATACGAATTCCGGGTAGACTGTCAAGCTTGAGACCGGGAGAAGCTAGAGGTACTCTTGGGGTAGAGGTGAAATTCTGTAATCCTGAGGGGACCACCAGTGGCGAAGGCGTCTAGCTAGAACGGGTCTGACGGTGAGGGACGAAGCCCTGGGGCG |
| p__Thermoplasmatota_g__SM1-50_ASV_7613 | 0 | 0 | 0 | 0 | 0 | 0 | 0 | 1 | 0 | 0 | 0 | CCTTTACAATGCACGAAAGTGTGATAAGGGGATCCTCAGTGCTTATGCTTTGCATAGGCTTTTGCCAAATGTAAATAGTTTGGCGAATAAGTGGTGGGTAAGACTGGTGCCAGCCGCCGCGGTAACACCAGCGCCACGAGTGGTAGCCGCTATTATTGGGCCTAAAACGTCCGTAGCCTGTTTGATAAATGCCTGGGTAAATCGTATAGCTCAACTATACGAATTCCGGGTAGACTGTCAAGCTTGAGACCGGGAGAAGCTAGAGGTACTCTTGGGGTAGAGGTGAAATTCTGTAATCCTGAGGGGACCACCAGTGGCGAAGGCGTCTAGCTAGAACGGGTCTGACGGTGAGGGACGAAGCCCTGGGGCG |
| p__Thermoplasmatota_g__SM1-50_ASV_7698 | 0 | 0 | 0 | 0 | 0 | 0 | 0 | 0 | 1 | 0 | 0 | ACTTCACACTGCGCGAAAGCGCGATGAGGGGATCCCAAGTGTATGCACGTAAGTGTATACTGTTTCTATGTATAAAAAGCATAGAGAGTAAGGGCTGGGTAAGACGGGTGCCAGCCGCCGCGGTAATACCTGCAGCTCGAGTGGTGATCACTATTATTGGGCCTAAAACGTCCGTAGCCGGTCTTGTAAATGCCTGGGTAAATCGTGTCGCTTAACGATACGAATTCCGGGTAGACTGTAAGACTTGAGACCGGGAGAGGCTAGAGGTACTCCTGGGGTAGAGGTGAAATTCTGTAATCCTAGGGGGACCACCAGTGGCGAAGGCGTCTAGCTAGAACGGGTCTGACGGTGAGGGACGAAGCCCTGGGGCG |
| p__Thermoplasmatota_g__SM1-50_ASV_7734 | 0 | 0 | 0 | 0 | 0 | 0 | 0 | 0 | 1 | 0 | 0 | ACTTCACACTGTGCGAAAGCACGATGAGGGAATCCCAAGTGCCAGCACGTAAGTGTTGGCTGTTTCTTTGTCTAAAAAACAAAGAGAGTAAGGGCTGGGTAAGACGGGTGCCAGCCGCCGCGGTAATACCTGCAGCTCAAGTGGTGGCCGTTATTATTGGGCCTAAAACGTCCGTAGCCGGTCTATTAAATGCCTGGGTAAATCGTGCCGCTTAACGGTACGAATTCCGGGTAGACTGATAGACTTGAGATCGGGAGAGGCTAGAGGTACTCCTGGGGTAGAGGTGAAATTCTGTAATCCTGGGGGGACCACCAGTGGCGAAGGCGTCTAGCTAGAACGAGTCTGACGGTGAGGGACGAAGCCCTGGGGCG |
| p__Thermoplasmatota_g__SM1-50_ASV_7747 | 0 | 0 | 0 | 0 | 0 | 0 | 0 | 0 | 1 | 0 | 0 | ACTTCACACTGCGCGAAAGCGCGATGAGGGGATCCCAAGTGTATGCACGTAAGTGTATACTGTTTCTATGTATAAAAAGCATAGAGAGTAAGGGCTGGGTAAGACGGGTGCCAGCCGCCGCGGTAATACCTGCAGCTCGAGTGGTGATCACTATTATTGGGCCTAAAACGTCCGTAGCCGGTCTTGTAAATGCCTGGGTAAATCGTATGGCTTAACCATACGAATTCCGGGTAGACTGTTAGACTTGAGACCGGGAGAGGCTAGAGGTACTCCTGGGGTAGAGGTGAAATTCTGTAATCCTGGGGGGACCACCAGTGGCGAAAGCGTCTAGCTAGAACGGGTCTGACGGTGAGGGACGAAACCCTGGGGCG |
| p__Nanoarchaeota_g__SM23-78_ASV_7197 | 0 | 0 | 1 | 0 | 0 | 0 | 0 | 0 | 0 | 0 | 0 | CCTTTACAATGCACGAAAGTGTGATAAGGGAATCCCAAGTGCTCAAGCATAGCTTGGGCTTTTGCCATATGTAAATAGTATGGCGAATAAGTGGTGGGAAATACCGGTGCCAGCCGCCGCGGTAACACCGGAGCCACAAGTGGCGATCGCGATTATTGGGCCTAAAGCGTCCGTAGCAGGTCATGTAAATCTCTTGTGAAATTGCTAAGCTCAACTTAGCAGAGTGCAGGAGACACTGCATGACTCGAGACCGGGAGGGGTCAGAAGTATTCCTGAGGGAGCGGTAAAATGCTATAATCTCAGGAGGACTACCTGTGGCGAAAGCGTCTGACTAGAACGGGTCTGACTGTGAGGGACGAAAGCTAGGGGAG |
| p__Thermoplasmatota_g__UBA147_ASV_6917 | 0 | 0 | 0 | 0 | 0 | 0 | 0 | 0 | 0 | 0 | 1 | ACTTGTCAATGGGGGCAACCCCGAACAGGGGATTCCAAGTGCATGCGTTTTACGTATGCTGTTCGGTTGTCTAAAAAACAGCCGAAGTAAGGGCTGGGTAAGACAGGTGCCAGCCGCCGCGGTAATACCTGCAGCCCAAGTGGTAATCGTTATTATTGAGCCTAAAGCGTTCGTAGCCGGTTTGGTAAACCCTCGGGTAAATCGAGCAGCTCAACTGTTCTAATTCCGGGGGGACTGCCAGACTTGGGACCGGGAGAGGTCAGAGGTACTCCGGGGGTAGGGGCTAAATCCTGTAATCCCGGACGGACCACCGATGGCGAAAGCACGGTGGGAAGACGGCTCCGACAGTGAGGGACGAAAGCTGGGGTCT |
| p__Halobacteriota_f__Halobacteriaceae_4595_ASV_6809 | 0 | 0 | 0 | 0 | 0 | 0 | 0 | 0 | 0 | 0 | 1 | CCTTTACACTGCACGACAGTGCGATAGGGGGACTCCGAGTGCGAGGGCATAGAGCCCTCGCTTTTCTGTACCGTAAGGTGGTACAGGAAGAAGGGCTGGGCAAGACCGGTGCCAGCCGCCGCGGTAATACCGGCAGCCCAAGTGATGGCCGATATTATTGGGCCTAAAGCGTCCGTAGCTGGCCGGACAAGTCCGTTGGGAAATCCACGCGCTCAACGCGTGGGCGTCCAGCGGAAACTGTCCGGCTCGGGACCGGAAGACCCAGAGGGTACGTCCGGGGTAGGAGTGAAATCCCGTAATCCTGGACGGACCACCGGTGGCGAAAGCGCTCTGGGAGGACGGATCCGACAGTGAGGGACGAAAGCTAGGGTCT |
| p__Halobacteriota_f__Haloferacaceae_ASV_7341 | 0 | 0 | 0 | 0 | 1 | 0 | 0 | 0 | 0 | 0 | 0 | CTGGGCAAGACCGGTGCCAGCCGCCGCGGTAATACCGGCAGCTCGAGTGATGGCCGATCTTATTGGGCCTAAAGCGTCCGTAGCTGGCCAGACAAGTCCGTCGGGAAATCCACCTGCCCAACAGGTGGGCGTCCGGCGGAAACTCTCTGGCTTGGGACCGGAAGACCCGAGGGGTACGTCCGGGGTAGGAGTGAAATCCCGTAATCCTGGACGGACCGCCGATAGCGAAAGCACCTCGGGAAGACGGATCCGACAGTGAGGGACGAAAGCTGGGGTCT |
| p__Thermoplasmatota_f__Methanomassiliicoccaceae_ASV_6567 | 1 | 0 | 0 | 0 | 0 | 0 | 0 | 0 | 0 | 0 | 0 | CCTTCACACTGTGCGAAAGCGCGATGAGGGGATCCCAAGTGCTTGCACGTAAGTGTAAGCTGTTTCTTTGTTTAAAATGCATAGAGAGTAAGGGCTGGGTAAGACGGGTGCCAGCCGCCGCGGTAATACCTGCAGCTCAAGTGGTGACCACTATTATTGGGCCTAAAACGTCCGTAGCCTGTTTGATAAATGCCTGGGTAAATCGTATAGCTCAACTATACGAATTCCGGGGAGACTGCAAGGCTTGGGACCGGGAGAGGCCAGAGGTACTTCCAGGGTAGGAGTAAAATCCTGTAATCCTGGAAGGACCACCGGTGGCGAAGGCGTCTGGCTAGAACGGATCCGACGGTGAGGGACGAAGCCCTGGGGCG |

| Phylum_Genus_ASV_ number | Presence (1) or Absence (0) in each layer L1 | L2 | L3 | L4 | L5 | L6 | L7 | L8 | L9 | L10 | L11 | Sequence |
|---|---|---|---|---|---|---|---|---|---|---|---|---|
| p__Halobacteriota_f__Natrialbaceae_ASV_6838 | 0 | 0 | 0 | 0 | 0 | 0 | 0 | 0 | 0 | 0 | 1 | ACTTCACACTGTGTGAAAGCGCGATGAGGGAATCCCAAGTGTATGCACGTAAGTGTATACTGTTTCTTTGTTTAAAATGCAGAGAGAGTAAGAGCTGGGCAAGACCGGTGCCAGCCGCCGCGGTAATACCGGCAGCTCGAGTGATGACCGCTATTATTGGGCCTAAAGCGTCCGTAGCTGGCCATACAAGTCCATCGGGAAATCTCCCTGCTTAACAGGGAGGCGTCCGGTAGAAACTGTGTGGCTTGGGACCGGAAGATCCAGAGGGTACGTCTGGGGTAGGAGTGAAATCCCGTAATCCTAGACGGACCACCGGTGGCGAAAGCGCTCTGGAAGGACGGATCCGACAGTGAGGGACGAAAGCTAGGGTCA |
| p__Halobacteriota_f__Natrialbaceae_ASV_7042 | 0 | 1 | 0 | 0 | 0 | 0 | 0 | 0 | 0 | 0 | 0 | CCTTTACACTGCACGAGAGTGCGATAAGGGGACTCCAAGTGCGAGGGCATATCGTCCTCGCTTTTCACTACCGTAAGGAGGTAGTAGAATAAGTGCTGGGCAAGACCGGTGCCAGCCGCCGCGGTAATACCGGCAGCACGAGTGATGACCGCTATTATTGGGCCTAAAGCGTCCGTAGCCGGCCTCGCAAGTCCGTCGGGAAATCCGCGCGCTTAACGCGCGGGCGTCCGGCGGAAACTGCGTGGCTTGGGACCGGAAGACTCAGGGGGTACGTCTGGGGTAGGAGTGAAATCCTGTAATCCTGGACGGAACACCGGTGGCGAAGGCGGGTCTCTGGGCCATAACTGACGCTGAGGAGCGAAAGCGTGGGGAG |
| p__Halobacteriota_f__Natrialbaceae_ASV_7156 | 0 | 0 | 1 | 0 | 0 | 0 | 0 | 0 | 0 | 0 | 0 | CCTTTACACTGCACGCCAGTGCGATAAGGGGACTCCAAGTGCGAGGGCATACAGTCCTCGCTTTTTCCTACCGTAAGGTGGTAGGCGAATAAGGGCTGGGCAAGACCGGTGCCAGCCGCCGCGGTAATACCGGCAGCCCAAGTGATGACCGCTATTATTGGGCCTAAAGCGTCCGTAGCCGGCCGGGTAAGTCCGTCGGGAAATCCGCGCGCTCAACGCGCGGGCGTCCGGTGGAAACTGCCTGGCTTGGGACCGGAAGATCCAGAGGGTACGTCCGGGGTAGGAGTGAAATCCCGTAATCCCGGACGGACCACCGATGGCGAAAGCACTCTGAGAGGACGGATTCGACAGTGAGGGACGAAAGCTAGGGTCT |
| p__Halobacteriota_f__Natrialbaceae_ASV_7415 | 0 | 0 | 0 | 0 | 1 | 0 | 0 | 0 | 0 | 0 | 0 | CCTTTACACTGCACGACAGTGCGACAAGGGGACCCCGAGTGCGAGGGCATACAGTCCTCGCTTTTGTCGACCGTAAGGTGGTCGACGAATAAGAGCTGGGCAAGACCGGTGCCAGCCGCCGCGGTAATACCGGCAGCTCGAGTGATGACCGCTATTATTGGGCCTAAAGCGTCCGTAGCTGGCCGTGTAAGTCCATCGGGAAATCCATCCGCTCAACGGATGGGCGTCCGGTGGAAACTGCACGGCTTGGAGCCGGAAGACCCAAGGGGTACGTCCGGGGTAGGAGTGAAATCCCGTAATCCTGGGGGGACCACCAGTGGCGAAAGCGTCTAGCTAGAACGGGTCTGACGGTGAGGGACGAAACCCTGGGGCG |
| p__Halobacteriota_o__Halobacteriales_ASV_6693 | 0 | 0 | 0 | 0 | 0 | 0 | 0 | 0 | 0 | 1 | 0 | CCTTCACACTGTGCGAGAGCACGATGAGGGAATCCCAAGTGCTGGCACGTAAGTGTCAGCTGTTTCTTTGTCTAAAAAACAAAGAGAGTAAGGGCTGGGTAAGACGGGTGCCAGCCGCCGCGGTAATACCTGCAGCTCAAGTGGTGACCGCTATTATTGGGCCTAAAACGTCCGTAGCCTGTTTGGTAAATGCCTGGGTAAATCGTGTCGCTTAACGATACGAATTCCGGGTAGACTGCCAGGCTTGAGACCGGGAGAGGTTAGAGGTACTCCGGGGTAGGAGTGAAATCCTGTAATCCTGGACGGACCACCGGTGGCGAAAGCGCCTCAGGAGGACGGACCCGACGGTCAGGGACGAAAGCTAGGGTCT |
| p__Halobacteriota_o__Halobacteriales_ASV_6890 | 0 | 0 | 0 | 0 | 0 | 0 | 0 | 0 | 0 | 0 | 1 | CCTTTACACTGCACGCCAGTGCGATAAGGGGACTCCAAGTGCGAGGGCATACAGTCCTCGCTTTTGTGTACCGTAAGGTGGTACACGAATAAGGGCTGGGCAAGACCGGTGCCAGCCGCCGCGGTAATACCGGCAGCCCGAGTGATGACCGCTATTATTGGGCCTAAAGCGTCCGTAGCCGGCCGGGTAAGTCCGTCGGGAAATCCACCCGCTCAACGGGTGGGCGTCCGGCGGAAACTGCTCGGCTTGGGGCCGGAAGACCTGGAGGGTACGTCCGGGGTAGGAGTGAAATCCTGTAATCCTGGACGGACCGCCGGTGGCGAAAGCGCTCCAGGAGGACGGACCCGACGGTCAGGGACGAAAGCTAGGGTCT |
| p__Halobacteriota_o__Halobacteriales_ASV_6987 | 0 | 1 | 0 | 0 | 0 | 0 | 0 | 0 | 0 | 0 | 0 | CCTTTACACTGCACGCCAGTGCGATAAGGGGACTCCAAGTGCGAGGGCATACAGTCCTCGCTTTTTCCTACCGTAAGGTGGTAGGCGAATAAGGGCTGGGCAAGACCGGTGCCAGCCGCCGCGGTAATACCGGCAGCCCAAGTGATGACCGCTATTATTGGGCCTAAAGCGTCCGTAGCCGGCCGGGTAAGTCCGTCGGGAAATCCGCGCGCTCAACGCGCGGGCGTCCGGTGGAAACTGCCTGGCTTGGGACCGGAAGATCCAGAGGGTACGTCCGGGGTAGGAGTGAAATCCTGTAATCCCGGACGGACCACCGATGGCGAAAGCACTCTGAGAGGACGGATTCGACAGTGAGGGACGAAAGCTAGGGTCT |
| p__Halobacteriota_o__Halobacteriales_ASV_7251 | 0 | 0 | 0 | 0 | 1 | 0 | 0 | 0 | 0 | 0 | 0 | CCTTTACACTGCACGACAGTGCGATAGGGGGACTCCGAGTGCGAGGGCATACAGTCCTCGCTTTTCTGTACCGTAAGGAGGTACAGGAACAAGTGCTGGGCAAGACCGGTGCCAGCCGCCGCGGTAATACCGGCAGCACGAGTGATGGCCGATATTATTGGGCCTAAAGCGTCCGTAGCCGGCCGGACAAGTCCGTTGGGAAATCCGTTCGCTTAACGAACGGGCGTCCAGCGGAAACTGTCCGGCTTGGGACCGGAAGACCCGAGGGGTACGTCCGGGGTAGGAGTGAAATCCCGTAATCCTGGACGGACCGCCGGTGGCGAAAGCGCCTCGGGAAGACGGATCCGACGGTGAGGGACGAAAGCCAGGGTCT |
| p__Halobacteriota_o__Halobacteriales_ASV_7337 | 0 | 0 | 0 | 0 | 1 | 0 | 0 | 0 | 0 | 0 | 0 | GCATACAGTCCTCGCTTTTATGCACCGTAAGGAGGTGCATGAATAAGTGCTGGGCAAGACCGGTGCCAGCCGCCGCGGTAATACCGGCAGCACAAGTGATGGCCGCTATTATTGGGCCTAAAGCGTCCGTAGCCGGCCGAACAAGTCCGTCGGGAAATCCATCCGCTCAACGGATGGGCGTCCGGCGGAAACTGTTCGGCTTGGGACAGGAAGACTCGAGGGGTACGTCCGGGGTAGGAGTGAAATCCCGTAATCCTGGACGGACCGCCGATGGCGAAAGCACCTCGAGAAGACTGATCCGACGGTGAGGGACGAAAGCTGGGGTCT |
| p__Nanoarchaeota_o__Woesearchaeales_579_ASV_6583 | 1 | 0 | 0 | 0 | 0 | 0 | 0 | 0 | 0 | 0 | 0 | CCTCTACAATGCACGAAAGTGTGATAGGGGGACTCTCAGTGCTCAAGTTCTACTTGGGCTTTTGCCAAGAGTAAGTATCTTGGAGAATAAGTGGTGGGCAAGACCGGTGCCAGCCGCCGCGGTAATACCGGCGCCACAAGTGGCATTCGCGATTATTGGGCCTAAAGCGTCCGTAGCCGGTTTGGTAAATCCTCTGTGAAATTGTACAGCTTAACTGTACGGCGTGCAGGGGACACTGCTAGACTTGAGATCGGGAGGAGCTAGAAGTATGCTAGGGGGACCGGTAAAATGGGATAATCCCTGGTAGACTACCGATGGCGAAGGCATCTAGCCAGAACGAATCTGACGGTGAGGGACGAAAGCTAGGGGAG |
| p__Nanoarchaeota_o__Woesearchaeales_579_ASV_6632 | 1 | 0 | 0 | 0 | 0 | 0 | 0 | 0 | 0 | 0 | 0 | CCTTTACAATGCACGAAAGTGTGATAAGGGGATTCCAAGTGCTCAAGCAATGCTTGGGCTTTTGCCAAGAGTAAATACCTTGGCGAATAAGTGGTGGGCAAGACTGGTGCCAGCAGCCGCGGTAACACCAGCGCCACGAGTGGGAACCAGAATTATTGGGCCTAAAGCGTTCGTAGCCGGCTGTGTAAATCTTTTGTGAAATCGCCAGGCTCAACTTGGCGGAATGCAGAAGAGACTGCACAGCTAGGGACCGGGAGGGGTCAGGAGTATTTTTTGGGGAGCGGTAAAATGCTATAATCCAAAAAAGACTACCTGTGGCGAAGGCGCCTGACCAGAACGGATCCGACGGTGAGGAACGAAAGCTAGGGGAG |
| p__Nanoarchaeota_o__Woesearchaeales_579_ASV_6649 | 0 | 0 | 0 | 0 | 0 | 0 | 0 | 0 | 0 | 1 | 0 | CCTTTACAATGCACGAAAGTGTGATAAGGGAATCCTAAGTGCTCATGTACTACATGGGCTTTTGCTAAGAGTAAGTATCTTGGCGAATAAGTGGTGGGTAAGACTGGTGCCAGCCGCCGCGGTAACACCAGCGCCACGAGTGGGGGCCACGTTTATTGGGCCTAAAGCGTCCGTAGCCGGTCTGTTAAATCTTTTGTGAAATCGTAAGGCTTAACCTTACGGCTTGCAAGAGACACTGGCAGACTAGGAACCGGGAAGGGCAAGAGGTATTCTAGGGGTAGCGGTAAAATGTTATAATCCCTAGAGGACCACCTGTGGCGAAGGCGTCTTGCTAGAACGGATTCGACGGTGAGGGACGAAAGCCAGGGGAG |
| p__Nanoarchaeota_o__Woesearchaeales_579_ASV_6834 | 0 | 0 | 0 | 0 | 0 | 0 | 0 | 0 | 0 | 0 | 1 | CCTTCACACTGTGTGAAAGCGCGATGAGGGGATCCCAAGTGCTTGCTCTTAGGAGTAAGCTGTTTTCCTGTTTAAATCGCAGGAAGAGGAAGGGCTGGATAAGACGGGTGCCAGCCGCCGCGGTAACACCAGCGCCGCAAGTAGCAACCATGTTTACTGGGCCTAAAGCGTTCGTAGCCGGTTTGGTAAATCTTTTGTGAAATTGTTGGGCCTAACCTAACAGCATGCAGAGGAGACTGCCAGACTAGAGACCGGGAGGAGTTGGAAATATTCTGGGGGGAGCGGTAAAATGTGATAATCCCCAGAAGATTACCTGTGGCGAAAGCGTCCAACCAGAACGGTTCTGACGGTGAGGAACGAAAGCTAGGGGAG |
| p__Nanoarchaeota_o__Woesearchaeales_579_ASV_7495 | 0 | 0 | 0 | 0 | 0 | 0 | 1 | 0 | 0 | 0 | 0 | CCTTTACAATGCACTAACGTGTGATAAGGGGACTCTAAGTGCTCATACAACGTATGGGCTTTTCTAAAGAGTAAATATCTTTAGGAATAAGTGGTGGGAAAGACTGGTGCCAGCCGCCGCGGTAACACCAGCGCCACGAGTGGGAACCATCATTATTGGGCCTAAAGCGTCCGTAGCCGGTTTGATAAATTTCTTGTGAAATTATCCAGCTTAACTGGATAGCGTGCAGGAAAGACTGTCAGACTAGGGACCGGGAGGAGCGTGAGGTATTCCTTGGGGAGCGGTAAAATGTTATAATCCTTGGAAGACCACCTGTGGCGAAGGCGTCACGCTAGAACGGATCCGACGGTGAGGGACGAAAGCTAGGGTAG |
| p__Nanoarchaeota_o__Woesearchaeales_579_ASV_7663 | 0 | 0 | 0 | 0 | 0 | 0 | 0 | 0 | 1 | 0 | 0 | CCTTTACAATGCACGAAAGTGTGATAGGGGGACTCAAAGTGCTCATGTTAAACATGGGCTTTTGCCAAGTGCAAACAGCTTGGCGAATAAGGGGTGGGTAAGACTGGTGCCAGCCGCCGCGGTAACACCAGCGCCTCAAGTGGGAGCCGATATTATTGGGCCTAAAGCGTCCGTAGCAGGTTCTGTAAATTACTTGTGAAATCGCACAGCCTAACTGTGCGGATTGCGGGTAACACTGCAGAACTTGAGACCGGGAGACGTTAGAGGTATTCAAGGGGGAGCGGTAAAATGCGAGAATCCCTTGAGGACCACCTGTTGCGAAGGCGTCTAACAATAACGGGTCTGACTGTGAGGGACGAAAGCTAGGGGAG |
| p__Nanoarchaeota_o__Woesearchaeales_579_ASV_7667 | 0 | 0 | 0 | 0 | 0 | 0 | 0 | 0 | 1 | 0 | 0 | CCTCTACAATGCACGAAAGTGTGATAGGGGGATTCTCAGTGCTCAAGTTATGCTTGGGCTTTTGCTTAGAGTAAGTATCTATGCGAATAAGTGGTGGGCAAGACCGGTGCCAGCCGCCGCGGTAATACCGGCGCCACGAGTGGCAATCACGATTATTGGGCCTAAAGCGTCCGTAGCCGGCTTAGTAAATCTTTTGTGAAATCGTGCAGCTCAACTGTACGGCGTGCAGAAGACACTGCTGAGCTAGAGACAGGGAGGAGCTAGAAGTATGCTAAGGGGACCGGTAAAATGGGATAATCCTTAGTAGACTACCGATGGCGAAGGCATCTAGCCAGAACTGATCTGACGGTGAGGGACGAAAGCCAGGGGAG |
| p__Nanoarchaeota_o__Woesearchaeales_579_ASV_7720 | 0 | 0 | 0 | 0 | 0 | 0 | 0 | 0 | 1 | 0 | 0 | CCTCTACAATGCACGAAAGTGTGATAGGGGGATCCCAAGTGCTTACACACTGTGTAGGCTTTTGCCAAGAGTAAATACCTTGGCGAATAAGTGGTGGGTAAGACTGGTGCCAGCAGCCGCGGTAACACCAGCGCCACGAGTGGCATACACATTTATTGGGCCTAAAGCGTCCGTAGCTGGTCTGGTAAATCTTTTGTGAAATCATAGGGCTTAACTCTATGGCGTGCAGAAGACACTGCCAGACTTGGAACCGGGAGGAGTCAGAGGAATTCCGGGGGGAGCGGTAAAATGCTATAATCCCCGGAGGACCACCTGTGGCGAAGGCGTCTGACTAGAACGGATTCGACAGTGAGGGACGAAAGCCAGGGGAG |
| p__Nanoarchaeota_c__Nanoarchaeia_ASV_6871 | 0 | 0 | 0 | 0 | 0 | 0 | 0 | 0 | 0 | 0 | 1 | CCTCTTCAATGCACGACAGTGTGAAAGGGGAATCCTACGTGCTTATGTACAACATAGGCTTTTGCATAGTGTAAATAGCTATGCGAATAAGGGGTGGGAAAGACCGGTGCCAGCAGCCGCGGTAACACCGGCGCCCCAAGTGGCAATCATAATTACTGGGCCTAAAGCATCCGTAGCCGGATCAGTAAATTTCCTGTGAAATCTTTCAAGTAAACTGGAAGGCGTGCAGGAAAGACTGCTGATCTAGAGACCGGGAGGAGTTAGAGGTATTCTGAGGGGAGCGGTAAAATGTGATAATCCTCGGAAGACCACCGATGGCGAAGGCATCTAACTAGAACGGATCTGACGGTGAGGGATGAAGGCTAGGGGAG |
| p__Thermoplasmatota_p__Thermoplasmatota_ASV_6949 | 0 | 0 | 0 | 0 | 0 | 0 | 0 | 0 | 0 | 0 | 1 | CCTTCACACTGTGCGAAAGCGCGATGAGGGGATCCCAAGTGCTTGCACGTAAGTGTAAGCTGTTTCTTTGTTTAAAACGCAGAGAGAGGAAGGGCTGGGTAAGACGGGTGCCAGCCGCCGCGGTAATACCTGCAGCTCGAGTGGTGACCGCTATTATTGGGCCTAAAACGTCCGTAGCCGGCTGAGTAAATGCTTGGGTAAATCGTGCCGCTTAACGGTACGAATTCCGGGTAAACTGCTCAGCTTGAGATCGGGAGAAGCTAGAGGTACTCCTGGGGTAGGGGCTAAATCCTGTAATCCTTGAAGGACCACCAGAGGCGAAGGCGGTTCTCTAAACTACTCCTGACACTGAGGTACGAAAGCTAGGGGAG |
| p__Thermoplasmatota_p__Thermoplasmatota_ASV_7114 | 0 | 0 | 1 | 0 | 0 | 0 | 0 | 0 | 0 | 0 | 0 | CTTCTACAATGCGCGTAAGCGTGATAGAGCAAGCCAGAGTGCTTTTCCGTATAGGGAAAGCTTTTGTGTGATGTAAAAAGTCACACGAATAAGGACTGGGCAAGACTGGTGCCAGCCGCCGCGGTAATACCAGCAGTCCAAGTCGCAGCCGCTTTTATTGGGCCTAAAACGTCCGTAGCCGGTCTAGTAAATGCCTGGGTAAATCGTGCCGCTTAACGGTACGAATTCCGGGTAGACTGCTAGACTTGAGATCGGGAGAGGTTAGAGGTACTCCTTGGGTAGAGGTGAAATTCTGTAATCCTAGGGGGACCACCTGTGGCGAAAGCGTCTAACCAGAACGAGTCTGACGGTGAGGGACGAAGCCCTGGGTCA |
| p__Thermoplasmatota_p__Thermoplasmatota_ASV_7518 | 0 | 0 | 0 | 0 | 0 | 0 | 1 | 0 | 0 | 0 | 0 | ACTGCACAATGCTGGAAACAGCGATGCGGGAACTCCAAGTGCCGGCACAACGTGTCGGCTGTTCCATTGCCTAAAAAGCATAGGAAGTAAGGGCTGGGTAAGACGGGTGCCAGCCGCCGCGGTAATACCTGCAGCCCGAGTGGTGATCGTTATTATTGGGCCTAAAACGTCCGTAGCCGGTTCGGTAGATGCCTGTGTAAATCGAACAGCTTAACTGTTCGAACTGCGGGTAGACCGCCGGACTTGGGACCGGGAGAGTCACGAAGTACTCCGGGGGTAGAGGTGAAATTCTGTAATCCTCGGGGGACTACCGGTGGCGAAGGCGTCGTGATAGAACGGCTCCGACGGTGAGGGACGAAGCCCTGGGGCG |
| d__Archaea_d__Archaea_ASV_6582 | 1 | 0 | 0 | 0 | 0 | 0 | 0 | 0 | 0 | 0 | 0 | CCTCTACAATGCACGCAAGTGTGATAGGGGAACCCAAAGTGCTTACACTTTGTGTAGGCTTTTGCCAAGTGTAAATAACTTGGCGAATAAGTGGTGGGAAAGACTGGTGCCAGCAGCCGCGGTAACACCAGCGCCACGAGTGGCAATCATTTTTATTGGGCCTAAAGCGTCCGTAGCTGGTTCTGTAAGTCTTTTGTGAAATCGTTTGGCTCAACCAAACGGCGTGCAAGAGATACTGCAGTTCTAGGGACCGGGAGGAGCATAGAGTATTCCAGGGGGAACGGTAAAATGTAATAATCCCTGGAGGACTACCTGTGGCGAAGGCGCTATGCTAGAACGGATCCGACAGTGAGGGACGAAGGCCAGGGGAG |
| d__Archaea_d__Archaea_ASV_6812 | 0 | 0 | 0 | 0 | 0 | 0 | 0 | 0 | 0 | 0 | 1 | ATTCCGCAATGCGCGCAAGCGCGACGGAGTGACTCCCAGTGCCGATGGTCTAACGACCGTCGGCTTTTCTGAAGTGTAAACGGCTCCAGGTAATAAGGGCCGGGTAAGACTGGTGCCAGCCGCCGCGGTAACACCAGCGGCCCAAGTGGTAGTCGCTTTTATTGGGCCTAAAGCGTCCGTAGCCGGATTGACAAGTCCTTTGTGAAATCGGACGGCTTAACCGTTCGGCTTGCAGAGGAAACTGTCGATCTTGGGACCGGTAAGCGCTAAGGGTACTCCGGGGTAGCGGTGAAATGCTATAATCCCTGGGGGACCGCCAGTGGCGAAAGCGCTTAGCGAGGCCGGATCCGACGGTGAGGGACGAAGGCTAGGGGAG |

| Phylum_Genus_ASV_number | Presence (1) or Absence (0) in each layer L1 | L2 | L3 | L4 | L5 | L6 | L7 | L8 | L9 | L10 | L11 | Sequence |
|---|---|---|---|---|---|---|---|---|---|---|---|---|
| d__Archaea_d__Archaea_ASV_7040 | 0 | 1 | 0 | 0 | 0 | 0 | 0 | 0 | 0 | 0 | 0 | CCTTCACAATGCACGCAAGTGCGATGGGGGACTCCGAGTGTGGTCTTCCCACAAGGAAGGCCACTTTTCTGCACTGTAAGAAGGTACAGGAACAAGGGGTGGGCAAGACTGCTGCCAGCCGCCGCGGTAATAGCAGCATCCCAAGTGGTGGCCAGTATTATTGGGCTTAAAGCGTTCGTAGCCGGACTTGTAAGTCCGCTGTGAAATCCACTAGCTTAACTAGTGGGCATGCAGCGGATACTGCAGGTCTAGGGACCGGATGGCGTCAGGGCTATTCCCAGGGTAGGAGTGAAATCCTAAAATCCTGGGAGGAGCACCTGTGGCGAAAGCGCCTGACGAATACGGATCCGACGGTGAGGAACGAAAGCCGGGGGAG |
| d__Archaea_d__Archaea_ASV_7148 | 0 | 0 | 1 | 0 | 0 | 0 | 0 | 0 | 0 | 0 | 0 | CCTTCACAATGCTCGAAAGAGTGATGAGGGTACTCCGAGTGCCAGTGCTTAGCACTGGCTTTTCTGCAGCGTAAATAGTTGCAGGAATAAGGGCTGGGCAAGACCGGTGCCAGCCGCCGCGGTAACACCGGCAGCCCTAGTGGTGACCGCTTTTATTGGGCCTAAAGCGTTCGTAGCCGGGCCGGTAAGTCCTTGGGTAAATCCTGCGTCTCAAGCGCGGGAATTCTAAGGATACTACGGTGAGGGACGAAAGCTCGGGTCA |
| d__Archaea_d__Archaea_ASV_7220 | 0 | 0 | 0 | 1 | 0 | 0 | 0 | 0 | 0 | 0 | 0 | ACTTCGCAATGTGGGAAACCACGACGAGGGAACTCCAAGTGCCGGCGCAGAGCGTCGGCTGTTTCCAAGCCTAAAAAGCTTGGAGAGTAAGGGCTGGGTAAGACGGGTGCCAGCCGCCGCGGTAACACCTGCAGCTCAAGTGGTCGCCGCTATTATTGGGCCTAAAGCGTCCGTAGCCGGCCCGGTAAATGCCTGGGTAAATCGACTGGCTTAACCAGTCGAATTCCGGGTAGACTGCCGGGCTTGGGACCGGGAGAGTCACGAAGTACTCCGGGGGTAGAGGTGAAATTCTGTAATCCTGGGGGACCACCAGTGGCGAAAGCGTCTAGCTAGAACGGGTCTGACGGTGAGGGACGAAACCCTGGGGCG |
| d__Archaea_d__Archaea_ASV_7228 | 0 | 0 | 0 | 1 | 0 | 0 | 0 | 0 | 0 | 0 | 0 | CCTTTACAATGCACGAAAGTGTGATAAGGGAATCCCGAGTGCTAATGCAAAACATTGGCTTTTATGAAGGGTAGATACCTTTATGAATAAGGGGTGGGTAAGACCGGTGCCAGCCGCCGCGGTAATACCGGCGCCCCGAGTAGCACTCGCGTTTATTGGGCCTAAAGCGTCCGTAGCTGGTTTGGTAAATCGCCTGTGAAATCTCTTGGCTCAACTAAGAGACGTGCAGGTGACACTACTAGGCTTGGAGTCGGGAGGTGCTAGAAGTATTCTAAGGGGAGCGGTAAAATGCTATAATCCTTGGAGGACTGCCGGTGGCGAAGGCGTCTAGCAAGAACGAGCTCGACAGTGAGGGACGAAAGCTGGGGGAG |
| d__Archaea_d__Archaea_ASV_7547 | 0 | 0 | 0 | 0 | 0 | 0 | 1 | 0 | 0 | 0 | 0 | CCTTTACAATGCACGAAAGTGTGATAAGGCAACCCCTCGTGCTCACCGAGAGGTGGGCTTTTGTCATGTGTAGGGAGCATGACGAATAAGTGGTGGGCAAGACCGGTGCCAGCAGCCGCGGTAATACCGGCGCCACAAGTGGAGGCCGTGTTTATTGGGTCTAAAGCGTCCGTAGCCGGCTTTCCAAGTCTCTTGTCAAAGTTCTGAGCTCAACTCAGGATTAGCAAGGGAAACTGGAAGGCTTGGGACCGGGAGAGGCACAGGGTATTGTAAGGGTAGCGGTAAAATGCTATAATCCTTACAAGACCAACGAGTGCGAAGGCACTGTGCTGGAACGGATCCGACGGTCAGGGACGAAAGCCAGGGGAG |
| d__Archaea_d__Archaea_ASV_7557 | 0 | 0 | 0 | 0 | 0 | 0 | 0 | 1 | 0 | 0 | 0 | CCTTTACAATGCACGAAAGTGTGATAAGGGAATCCCAAGTGCCATAGTTTTACTATGGCTGTGGAAAAGCGTAAAATACTTTTCAAGAAAGGGGTGGGTAATACCGGTGCCAGCAGCCGCGGTAACCCCGGAGCCTCAAGTGGTGGCCGTGTTTATTGGGCCTAAAGCGCCCGTAGCCGGATTAATAAGTTCTCGGTGAAATCGTCTAGCTTAACTAAACGAATTGCCTTGAATACTATTAGTCTAGGGATAGGGAGAGGTCAGTAGTATGCCTGGGGGAGGGGTAAAATCCTTTAATCCTGGGTAGACTATCAATGGCGAAGGCAACTGACTGGAACTAATCCGACGGTGAGGGACGAAAGTTAGGGTAG |
| d__Archaea_d__Archaea_ASV_7634 | 0 | 0 | 0 | 0 | 0 | 0 | 0 | 1 | 0 | 0 | 0 | CCTTTACAATGCACGAAAGTGTGATAGGGGGATTCCAAGTGCTCATGCATTGCATGGGCTTTTCTCAAGAGTACAAATCTTGGGGAATAAGTGGTGGGAAAGACCGGTGCCAGCAGCCGCGGTAACCCCGGCGCCACAAGTGGCAATCATCATTATTGGGTCTAAAGCGTCCGTAGTCGGTCTGATCAATCTTTTGTGAAATCGTGGGGCTTAACTTCACGGCGCGCGGAAGATATTGTCAGACTAGGGACCGGGAGGAGTGTGAGGTATTGGCAAGGGAGCGGTAAAATGTGATAATCTTGCCAAGACCACCGGTGGCGAAGGCGTCACACCAGAACGGATCCGACGATCAGGGACGAAGGCCAGGGGAG |
| d__Archaea_d__Archaea_ASV_7640 | 0 | 0 | 0 | 0 | 0 | 0 | 0 | 1 | 0 | 0 | 0 | CCTCTACAATGCACGCAAGTGTGATAGGGGAACCCTAAGTGCCTTGTTTAACAAGGCTTTTGCCAAATGTAAATAGTTTGGCGAATAAGTGATGGGCAAGACTGGTGCCAGCAGCCGCGGTAACACCAGCGTCACAAGTGGGAACCGCAATTATTGGGCCTAAAGCGTCCGTAGCCGGGTTGTTAAATCTTTTGTGAAATTGTTAGGCTTAACTTAACAAATTGCAAGAGACACTGACAATCTTGGGACCGGGAGAAGTCAAAGGTATGGATGGGGGAGCGGTAAAATGTTATAATCCCTTCTAGACCACCAGTGGCAAAGGCGTTTGACTAGAACGGATCCGACGGTGAGGGACGAAAGCTAGGGGAG |
| d__Archaea_d__Archaea_ASV_7748 | 0 | 0 | 0 | 0 | 0 | 0 | 0 | 0 | 1 | 0 | 0 | ACTTTACAATGCACGAAAGTGTGATAAGGAAACTCCGAGTGCTCTTGGGAAACCAAGAGCTTTTGCCAAGCGTAGATAACTTGGCGAATAAGGACTGGGTAATACCCGTGCCAGCCGCCGCGGTAATACGGGAAGTCCAAGTGGTGTCCGCGTATATTGGGCTTAAAGCGTCCGTAGCTGGTCTGGAAAGTCTTTGGTGAAAGCCCGCGGCTCAACCGTGGGAAGTGCTAGGGAAACTTTCAGACTTGGAGGCGGGAGAGGTCAGAGGTACTTTTGGGGTAGCGGTAGAATGTGTTGATCCCGAAAGGACCACCGGTAGCGAAGGCGTCTGACTAGAACGTGCTCGACAGTGAGGGACGAAGGCCAGGGGAG |
| p__Nanoarchaeota_g__BMS3ABIN17_ASV_8437 | 0 | 0 | 0 | 0 | 0 | 0 | 0 | 0 | 1 | 0 | 0 | TTTTTACAATGCACGCAAGTGTGATAAAGCGAGCCAAAGTGCTTTCCGTACAGGAAAGCTTTTCTTCACTCTAAAAATGTGGAGGAATAAGGAGTGGGTAAGATCGGTGCCAGCCGCCGCGGTAATACCGACAGTTCAAGTAGCATCCTTTTTTATTGGGTCTAAAACATCCGTAGCTTGTAATTTAAGTCTCTTGTGAAATCCTATGTCTCAAGCATAGGGCTTGCAAGAGATACTGGATTACTAGAGATCGGTAGACGTAAGGAGTACGCCGGAGGTAGTGGTTAAATACGTTAATCTTCGGTGGACTAACAATCAGCGAAGGCACCTTACGAGGACGAATCTGACAGTGAGGGATGAAGGCCAGGGGCG |
| p__Halobacteriota_g__Halobiforma_ASV_7773 | 1 | 0 | 0 | 0 | 0 | 0 | 0 | 0 | 0 | 0 | 0 | CCTTTACACTGCACGCCAGTGCGATAAGGGGACCCCGAGTGCGAGGGCATACAGTCCTCGCTTTTGTGTACCGTAAGGTGGTACACGAATAAGGGCTGGGCAAGACCGGTGCCAGCCGCCGCGGTAATACCGGCAGCCCGAGTGATGGCCGCTCTTATTGGGCCTAAAGCGTCCGTAGCCGGCCAGACAGGTCCGTCGGGAAATCCGCACGCCCAACGTGCGGGCGTCCGGCGGAACCCAGCTGGCTTGGGACCGGAAGACCCGAGGGGTACGTCCGGGGTAGGAGTGAAATCCCGTAATCCTGGCCGGACCACCGGTGGCGAAAGCGCCTCGGGAGGACGGATCCGACGGTGAGGGACGAAAGCTCGGGTCA |
| p__Halobacteriota_g__Halorussus_ASV_8054 | 0 | 0 | 1 | 0 | 0 | 0 | 0 | 0 | 0 | 0 | 0 | CCTTTACACTGCACGAAAGTGCGATAAGGGGATCCCAAGTGCGAGGGCATACAGTCCTCGCTTTTCTGTACCGTAGGGAGGTACAAGAATAAGGGCTGGGCAAGACCGGTGCCAGCCGCCGCGGTAATACCGGCAGCCCAAGTGATGGCCGCTATTATTGGGCCTAAAGCGTCCGTAGCCGGCCGGATAAGTCCGTCGGGAAATCCACCTGCTCAACAGGTGGGCGTCCGGCGGAAACTGTCTGGCTTGGGGCCGGAAGATCTGAGGGGTACGTCCGGGGTAGGAGTGAAATCCTGTAATCCTGGACGGACCGCCGGTGGCGAAAGCGCCTCAGGAGGACGGACCCGACGGTGAGGGACGAAAGCTAGGGTCA |
| p__Halobacteriota_g__Halorussus_ASV_8435 | 0 | 0 | 0 | 0 | 0 | 0 | 0 | 0 | 1 | 0 | 0 | ACTTTACACTGCACGACAGTGCGATAAGGGAACTCCGAGTGCGAGGGCATATAGTCCTCGCTTTTCTGCACCGTAAGGTGGTGCACGAATAAGAGCTGGGCAAGACCGGTGCCAGCCGCCGCGGTAATACCGGCAGCTCAAGTGATGGCCGCTATTATTGGGCCTAAAGCGTCCGTAGCCGGCCAGACAGGTCCGTCGGGAAATCCATCCGCTTAACGGATGGGCGTCCGGCGGAAACCAGCTGGCTTGGGGCCGGAAGATCCGAGGGGTACGTCCGGGGTAGGAGTGAAATCCCGTAATCCTGGGGGGACCACCAGTGGCGAAGGCGTCTAGCTAGAACGGGTCCGACGGTGAGGGACGAAGCCCTGGGGCG |
| p__Halobacteriota_g__Methanolobus_ASV_7943 | 0 | 0 | 0 | 0 | 0 | 0 | 0 | 0 | 0 | 0 | 1 | ACTTTACAATGCGGGAAACCGCGATAAGGGGACACCGAGTGCCAGCATATTATGTTGGCTGTCCATATGTGTAAATAACATGTGTTAGCAAGGGCCGGGCAAGACCGGTGCCAGCCGCCGCGGTAACACCGGCGGCCCGAGTGGTAGCCACTATTATTGGGTCTAAAGGGTCCGTAGCCGGTTTGATCAGTCTTCCGGGAAATCTGACAGCTTAACTGTTAGGCTTCTGGGGGATACTGTCAGACTTGGAACCGGGAGAGGTAAGAGGTACTACAGGGGTAGGAGTGAAATCCCGTAATCCTGGACGGACCACCGGTGGCGAAAGCGCCTCAGAAAAACGGACCCGACGGTGAGGGACGAAAGCTAGGGTCA |
| p__Thermoplasmatota_g__SM1-50_ASV_8369 | 0 | 0 | 0 | 0 | 0 | 0 | 0 | 1 | 0 | 0 | 0 | CCTTCACACTGTGCGAAAGCGCGATGAGGGGATCCCAAGTGCTTGCACGTAAGTGTAAGCTGTTTCTTTGTTTAAAATGCATAGAGAGTAAGGGCTGGGTAAGACGGGTGCCAGCCGCCGCGGTAATACCTGCAGCTCAAGTGGTGATCGTTATTATTGGGCCTAAAACGTCCGTAGCCGGTCTTGTAAATGCCTGGGTAAATCGTAACGCTTAACGTTACGAATTCCGGGTAGACTGTAAGACTTGAGACCGGGAGAGGCTAGAGGTACTCCTGGGGTAGAGGTGAAATTCTGTAATCCTAGGGGGACCACCAGTGGCGAAGGCGTCTAGCTAGAACGGGTCTGACGGTGAGGGACGAAGCCCTGGGGCG |
| p__Halobacteriota_f__Haloarculaceae_3992_ASV_8058 | 0 | 0 | 1 | 0 | 0 | 0 | 0 | 0 | 0 | 0 | 0 | CCTTTACACTGCACGACAGTGCGATAAGGGGACTCCGAGTGCGAGGGCATATAGTCCTCGCTTTTCTGTACCGTAAGGAGGTACAGGAACAAGTGCTGGGCAAGACCGGTGCCAGCCGCCGCGGTAATACCGGCAGCACGAGTGATGGCCGCTATTATTGGGCCTAAAGCGTCCGTAGCCGGCCAGACAAGTCCGTTGGGAAATCCACGCGCCCAACGCGTGGGCGTCCAGCGGAAACTGTCGGGCTTGGGGCCGGAGGACTCGAGGGGTACGTCCGGGGTAGGAGTGAAATCCCGTAATCCTGGACGGACCACCGATGGCGAAAGCACTCTGAGAGGACGGATTCGACAGTGAGGGACGAAAGCTAGGGTCT |
| p__Halobacteriota_f__Haloarculaceae_3992_ASV_8068 | 0 | 0 | 1 | 0 | 0 | 0 | 0 | 0 | 0 | 0 | 0 | CCTTTACACTGCACGACAGTGCGATAAGGGGACTCCGAGTGCGAGGGCATATAGTCCTCGCTTTTCTGTACCGTAAGGTGGTACAGGAACAAGTGCTGGGCAAGACCGGTGCCAGCCGCCGCGGTAATACCGGCAGCACGAGTGATGGCCGCTATTATTGGGCCTAAAGCGTCCGTAGCCGGCCAGACAAGTCCGTTGGGAAATCCACGCGCCCAACGCGTGGGCGTCCAGCGGAAACTGTCGGGCTTGGGGCCGGAGGACTCGAGGGGTACGTCCGGGGTAGGAGTGAAATCCCGTAATCCTGGACGGACCACCGATGGCGAAAGCACTCTGAGAGGACGGATTCGACAGTGAGGGACGAAAGCTAGGGTCT |
| p__Halobacteriota_f__Haloarculaceae_3992_ASV_8073 | 0 | 0 | 1 | 0 | 0 | 0 | 0 | 0 | 0 | 0 | 0 | CCTTTACACTGCACGACAGTGCGATAAGGGGACTCCGAGTGCGAGGGCATATAGTCCTCGCTTTTCTGTACCGTAAGGTGGTACAGGAACAAGTGCTGGGCAAGACCGGTGCCAGCCGCCGCGGTAATACCGGCAGCACGAGTGATGGCCGCTCTTATTGGGCCTAAAGCGTCCGTAGCCGGCCAGACAAGTCCGTTGGGAAATCCACGCGCCCAACGCGTGGGCGTCCAGCGGAAACTGTCGGGCTTGGGGCCGGAGGACTCGAGGGGTACGTCCGGGGTAGGAGTGAAATCCCGTAATCCTGGACGGACCACCGATGGCGAAAGCACTCTGAGAGGACGGATTCGACAGTGAGGGACGAAAGCTAGGGTCT |
| p__Halobacteriota_o__Halobacteriales_ASV_8053 | 0 | 0 | 1 | 0 | 0 | 0 | 0 | 0 | 0 | 0 | 0 | CCTTTACACTGCACGAACGTGCGATAAGGGGACTCCGAGTGCGAGGGCATATCGTCCTCGCTTTTCTGTACCGTAAGGTGGTACAGGAAGAAGGGCTGGGCAAGACCGGTGCCAGCCGCCGCGGTAATACCGGCAGCCCGAGTGATGGCCGCTATTATTGGGCCTAAAGCGTCCGTAGCCGGCCGGGCAAGTCCGTTGGGAAATCCACCCGCTTAACGGGTGGGCGACCGGCGGAAACTGCCTGGCTTGGGACCGGGAGATCCGAGGGGTACGTCCGGGGTAGGAGTGAAATCCTGTAATCCTGGACGGACCGCCGGTGGCGAAAGCGCCTCGGAAGAACGGATCCGACGGTGAGGGACGAAAGCCAGGGTCT |
| Escherichia_coli_K12_16S_rRNA_rrs | | | | | | Out group | | | | | | ACCTACGGGAGGCAGCAGTGGGGAATATTGCACAATGGGCGCAAGCCTGATGCAGCCATGCCGCGTGTATGAAGAAGGCCTTCGGGTTGTAAAGTACTTTCAGCGGGGAGGAAGGGAGTAAAGTTAATACCTTTGCTCATTGACGTTACCCCGCAGAAGAAGCACCGGCTAACTCCGTGCCAGCAGCCGCGGTAATACGGAGGGTGCAAGCGTTAATCGGAATTACTGGGCGTAAAGCGCACGCAGGCGGTTTGTTAAGTCAGATGTGAAATCCCCGGGCTCAACCTGGGAACTGCATCTGATACTGGCAAGCTTGAGTCTCGTAGAGGGGGGTAGAATTCCAGGTGTAGCGGTGAAATGCGTAGAGATCTGGAGGAATACCGGTGGCGAAGGCGGCCCCCTGGACGAAGACTGACGCTCAGGTGCGAAAGCGTGGGGAGCAAACAGGATTAGATACCCTGGTAGTC |
| Escherichia_coli_strain_Nissle_1917_16S_rRNA | | | | | | | | | | | | CCTACGGGAGGCAGCAGTGGGGAATATTGCACAATGGGCGCAAGCCTGATGCAGCCATGCCGCGTGTATGAAGAAGGCCTTCGGGTTGTAAAGTACTTTCAGCGGGGAGGAAGGGAGTAAAGTTAATACCTTTGCTCATTGACGTTACCCCGCAGAAGAAGCACCGGCTAACTCCGTGCCAGCAGCCGCGGTAATACGGAGGGTGCAAGCGTTAATCGGAATTACTGGGCGTAAAGCGCACGCAGGCGGTTTGTTAAGTCAGATGTGAAATCCCCGGGCTCAACCTGGGAACTGCATCTGATACTGGCAAGCTTGAGTCTCGTAGAGGGGGGTAGAATTCCAGGTGTAGCGGTGAAATGCGTAGAGATCTGGAGGAATACCGGTGGCGAAGGCGGCCCCCTGGACGAAGACTGACGCTCAGGTGCGAAAGCGTGGGGAGCAAACAGGATTAGATACCCTGGTAGTC |
| Escherichia_coli_O157_H7_strain_EDL933_16S_rRNA | | | | | | | | | | | | CCTACGGGAGGCAGCAGTGGGGAATATTGCACAATGGGCGCAAGCCTGATGCAGCCATGCCGCGTGTATGAAGAAGGCCTTCGGGTTGTAAAGTACTTTCAGCGGGGAGGAAGGGAGTAAAGTTAATACCTTTGCTCATTGACGTTACCCCGCAGAAGAAGCACCGGCTAACTCCGTGCCAGCAGCCGCGGTAATACGGAGGGTGCAAGCGTTAATCGGAATTACTGGGCGTAAAGCGCACGCAGGCGGTTTGTTAAGTCAGATGTGAAATCCCCGGGCTCAACCTGGGAACTGCATCTGATACTGGCAAGCTTGAGTCTCGTAGAGGGGGGTAGAATTCCAGGTGTAGCGGTGAAATGCGTAGAGATCTGGAGGAATACCGGTGGCGAAGGCGGCCCCCTGGACGAAGACTGACGCTCAGGTGCGAAAGCGTGGGGAGCAAACAGGATTAGATACCCTGGTAGTC |
| Escherichia_coli_strain_CFT073_16S_rRNA | | | | | | | | | | | | CCTACGGGAGGCAGCAGTGGGGAATATTGCACAATGGGCGCAAGCCTGATGCAGCCATGCCGCGTGTATGAAGAAGGCCTTCGGGTTGTAAAGTACTTTCAGCGGGGAGGAAGGGAGTAAAGTTAATACCTTTGCTCATTGACGTTACCCCGCAGAAGAAGCACCGGCTAACTCCGTGCCAGCAGCCGCGGTAATACGGAGGGTGCAAGCGTTAATCGGAATTACTGGGCGTAAAGCGCACGCAGGCGGTTTGTTAAGTCAGATGTGAAATCCCCGGGCTCAACCTGGGAACTGCATCTGATACTGGCAAGCTTGAGTCTCGTAGAGGGGGGTAGAATTCCAGGTGTAGCGGTGAAATGCGTAGAGATCTGGAGGAATACCGGTGGCGAAGGCGGCCCCCTGGACGAAGACTGACGCTCAGGTGCGAAAGCGTGGGGAGCAAACAGGATTAGATACCCTGGTAGTC |
| Escherichia_coli_strain_W3110_16S_rRNA | | | | | | | | | | | | CCTACGGGAGGCAGCAGTGGGGAATATTGCACAATGGGCGCAAGCCTGATGCAGCCATGCCGCGTGTATGAAGAAGGCCTTCGGGTTGTAAAGTACTTTCAGCGGGGAGGAAGGGAGTAAAGTTAATACCTTTGCTCATTGACGTTACCCCGCAGAAGAAGCACCGGCTAACTCCGTGCCAGCAGCCGCGGTAATACGGAGGGTGCAAGCGTTAATCGGAATTACTGGGCGTAAAGCGCACGCAGGCGGTTTGTTAAGTCAGATGTGAAATCCCCGGGCTCAACCTGGGAACTGCATCTGATACTGGCAAGCTTGAGTCTCGTAGAGGGGGGTAGAATTCCAGGTGTAGCGGTGAAATGCGTAGAGATCTGGAGGAATACCGGTGGCGAAGGCGGCCCCCTGGACGAAGACTGACGCTCAGGTGCGAAAGCGTGGGGAGCAAACAGGATTAGATACCCTGGTAGTC |

| | Presence (1) or Absence (0) in each layer | | | | | | | | | | | |
|---|---|---|---|---|---|---|---|---|---|---|---|---|
| Phylum_Genus_ASV_number | L1 | L2 | L3 | L4 | L5 | L6 | L7 | L8 | L9 | L10 | L11 | Sequence |
| g__Haladaptatus_AB663345_Ref_seq | | | | | | Reference Sequences | | | | | | ACGGGGCGCAGCAGGCGCGAAAACTTTACACTGCACGACAGTGCGATAAGGGAACTCCAAGTGCGAGGGCATATAGTCCTCGCTTTTGTGTATCGTAAGGTGGTACACGAATAAGAGCTGGGCAAGACCGGTGCCAGCCGCCGCGGTAATACCGGCAGCTCGAGTGATGGCCAATTTTATTGGGCCTAAAGCGTCCGTAGCCTGCCAGACAGGTCCGTCGGGAAATCTGCTCGCTCAACGAGCAGGCGTCCGGCGGAAACCAGCTGGCTTGGGGCCGGAAGACCCAAGGGGTACGTCTGGGGTAGGAGTGAAATCCTGTAATCCTAGACGGACCACCGATCGCGAAAGCACCTTGGGAGGACGGACCCGACGGTGAGGGACGAAAGCTAGGGTCACGAACCGGATTAGATACCCGGGTAGTCC |
| g__Halalkaliarchaeum_KY612370_Ref_seq | | | | | | | | | | | | ACGGGGCGCAGCAGGCGCGAAACCTTTACACTGCACGCCAGTGCGATAAGGGGACCCCGAGTGCGAGGGCATATCGTCCTCGCTTTTGTACACCGTAGGGAGGTGTACGAACAAGGGCTGGGCAAGACCGGTGCCAGCCGCCGCGGTAATACCGGCAGCCCGAGTGATGGCCGATTTTATTGGGCCTAAAGCGTCCGTAGCCGGCCGAGCAAGTCCGTCGGGAAATCCACGCGCTCAACGCGTGGGCGTCCGGCGGAACCTGCTCGGCTTGGGACCGGAAGACCCAGCGGGTACGTCCGGGGTAGGAGTGAAATCCTGTAATCCCGAACGGACCGCCGATGGCGAAAGCACGCTGGGAAGACGGATCCGACGGTGAGGGACGAAAGCTAGGGTCTCGAACCGGATTAGATACCCGGGTAGTCC |
| g__Halalkalicoccus_CP002062_Ref_seq | | | | | | | | | | | | ACGGGGCGCAGCAGGCGCGAAACCTTTACACTGCACGAACGTGCGATAAGGGGACTCCAAGTGCGAGGGCATACAGTCCTCGCTTTTCCCTACCGTAAGGTGGTAGGTGAATAAGAGCTGGGCAAGACCGGTGCCAGCCGCCGCGGTAATACCGGCAGCTCGAGTGATGGCCAATATTATTGGGCCTAAAGCGTCCGTAGCTGGCCGAACAGGTCTGTCGGGAAATCCGCTCGCTCAACGAGCGGACGTCCGGCGGAAACCAGTTGGCTTGGGACCGGAAGACCCGAGGGGTACGTCCGGGGTAGGAGTGAAATCCTGTAATCCTGGACGGACCGCCGGTGGCGAAAGCGCCTCGGGAGAACGGATCCGACAGTGAGGGACGAAAGCTTGGGTCACGAACCGGATTAGATACCCGGGTAGTCC |
| g__Halanaeroarchaeum_KM875608_Ref_seq | | | | | | | | | | | | ACGGGGCGCAGCAGGCGCGAAACCTTTACACTGCACGACAGTGCGATAAGGGAACTCCGAGTGCGAGGGCATAGAGCCCTCGCTTTTGTACACCGTAAGGTGGTGCACGAATAAGGGCTGGGCAAGACCGGTGCCAGCCGCCGCGGTAATACCGGCAGCCCAAGTGATGGCCGATATTATTGGGCCTAAAGCGTCCGTAGCTGGCCGGACAAGTCCGTTGGGAAATCCGCGTGCTTAACGCGCGGGCGTCCAGCGGAAACTGTCCGGCTTGGGGCCGGAAGACCTGAGGGGTACGTCCGGGGTAGGAGTGAAATCCCGTAATCTCGGACGGACCGCCTGTGGCGAAAGCGCCTCAGGAAGACGGACCCGACAGTGAGGGACGAAAGCCAGGGTCTCGAACCGGATTAGATACCCGGGTAGTCC |
| g__Halapricum_3815_CP064788r2_Ref_seq | | | | | | | | | | | | ACGGGGCGCAGCAGGCGCGAAACCTTTACACTGCACGACAGTGCGATAGGGGGACTCCGAGTGCGAGGGCATAGCGCCCTCGCTTTTGCGCACCGTAAGGTGGTGCGTGAATAAGGACTGGGCAAGACCGGTGCCAGCCGCCGCGGTAATACCGGCAGTCCGAGTGATGGTCGATCTTATTGGGCCTAAAGCGTTCGTAGCTGGCCACGCAAGTCCGTCGGGAAATCTACCCGCCTAACGGGTAGGCGTCCGGCGGAAACTGCGTGGCTTGGGACCGGGAGACCCGAGGAGTACGTCCGGGGTAGGAGTGAAATCCCGTAATCCCGGACGGACTACCGATGGCGAAAGCACCTCGGGAGCACGGATCCGACAGTGAGGAACGAAAGCTGGGGTCTCGAACCGGATTAGATACCCGGGTAGTCC |
| g__Halapricum_3981_KF314042_Ref_seq | | | | | | | | | | | | ACGGGGCGCAGCAGGCGCGAAACCTTTACACTGCACGACAGTGCGATAAGGGGACTCCGAGTGCGAGGGCATATAGTCCTCGCTTTTCTGTACCGTAGGGAGGTACAGGAACAAGTGCTGGGCAAGACCGGTGCCAGCCGCCGCGGTAATACCGGCAGCACGAGTGATGGCCGATATTATTGGGCCTAAAGCGTCCGTAGCCGGCCAGACAAGTCCGTTGGGAAATCGGCGTGCCTAACACGTCGGCGTCCAGCGGAAACTGTTTGGCTTGGGGCCAGAAGATCCGAGGGGTACGTCCGGGGTAGGAGTGAAATCCCGTAATCCTGGACGGACCACCGGTGGCGAAAGCGCCTCGGAAAGATGGACCCGACGGTGAGGGACGAAAGCTGGGGTCTCGAACCGGATTAGATACCCGGGTAGTCC |
| g__Haloarchaeobius_GU951428_Ref_seq | | | | | | | | | | | | ACGGGGCGCAGCAGGCGCGAAAACTTTACACTGCACGACAGTGCGATAAGGGAACTCCAAGTGCGAGGGCATAGAGCCCTCGCTTTTCTGTACCGTAGGGTGGTACAGGAATAAGAGCTGGGCAAGACCGGTGCCAGCCGCCGCGGTAACACCGGCAGCTCGAGTGATGACCGATATTATTGGGCCTAAAGCGTCCGTAGCTGGCCGAGAGGGTCCGTCGGGAAATCTGCTCGCTCAACGAGCAGGCGTCCGGCGGAAACCAATCGGCTTGGGACCGGAAGACCTGAGGGGTACGTCTGGGGTAGGAGTGAAATCCCGTAATCCTGGACGGACCACCGGTGGCGAAAGCGCCTCAGGAAGACGGATCCGACAGTGAGGGACGAAAGCTGGGGTCACGAACCGGATTAGATACCCGGGTAGTCC |
| g__Halobacterium_KC914879_Ref_seq | | | | | | | | | | | | ACGGGGCGCAGCAGGCGCGAAACCTTTACACTGTACGCAAGTGCGATAAGGGGACTCCGAGTGTGAAGGCATAGAGCCTTCACTTTTGTACACCCTAAGGTGGTGCACGAATAAGGACTGGGCAAGACCGGTGCCAGCCGCCGCGGTAATACCGGCAGTCCGAGTGATGGCCGATCTTATTGGGCCTAAAGCGTCCGTAGCTGGCTAGACAAGTCCGTTGGGAAATCTGCCCGCTTAACGGGCAGGCGTCCAGCGGAAACTGTCTAGCTTGGGACCGGAAGACCTGAGGGGTACGTCTGGGGTAGGAGTGAAATCCTGTAATCCTGGACGGACCGCCGGTGGCGAAAGCGCCTCAGGAGAACGGATCCGACAGTGAGGGACGAAAGCTAGGGTCTCGAACCGGATTAGATACCCGGGTAGTCC |
| g__Halobaculum_HM159610_Ref_seq | | | | | | | | | | | | ACGGGGCGCAGCAGGCGCGAAACCTTTACACTGCACGACAGTGCGATAGGGGGACTCCGAGTGCGGAGGCATAGTGCCTTCGCTTTTGTGCACCCTAAGGCGGTGCACGAATAAGAGCTGGGCAAGACCGGTGCCAGCCGCCGCGGTAATACCGGCAGCTCAAGTGATGGCCGATTTTATTGGGCCTAAAGCGTCCGTAGCCGGCCGGGCAAGTTCGTCGGGAAATCTACCAGCTCAACTGGTAGGCGTCCGGCGAAAACTGCACGGCTTGGGACAGGGAGACTCGAAGGGTACGTCCGGGGTAGGAGTGAAATCCTGTAATCCTGGACGGACCACCGATGGCGAAAGCACTTCGAGAGAACTGATTCGACGGTCAGGGACGAAAGCTGGGGTCTCGAACCGGATTAGATACCCGGGTAGTCC |
| g__Halobellus_3026_JQ237122_Ref_seq | | | | | | | | | | | | ACGGGGCGCAGCAGGCGCGAAACCTTTACACTGCACGCACGTGCGATAAGGGGACTCCGAGTGCGAGGGCATATCGTCCTCGCTTTTGTGTACCGTAGGGCGGTACACGAACAAGAGCTGGGCAAGACCGGTGCCAGCCGCCGCGGTAATACCGGCAGCTCAAGTGATGGCCGATCTTATTGGGCCTAAAGCGTCCGTAGCTGGCCTCGAAAGTCCGTCGGGAAATCCACGCGCCCAACGCGTGGGCGTCCGGCGGAAACTCCGAGGCTTGGGACCGGAAGGCTCGAGGGGTACGTCCGGGGTAGGAGTGAAATCCCGTAATCCCGGACGGACCACCGATGGCGAAAGCACCTCGAGAAAACGGATCCGACAGTGAGGGACGAAAGCTAGGGTCTCGAACCGGATTAGATACCCGGGTAGTCC |
| g__Halobellus_3029_GQ282620_Ref_seq | | | | | | | | | | | | ACGGGGCGCAGCAGGCGCGAAACCTTTACACTGCGCGCAAGTGCGATAAGGGGACTCCGAGTGCGAGGGCATATCGTCCTCGCTTTTCTGTACCGTAGGGCGGTACAGGAACAAGGACTGGGCAAGACCGGTGCCAGCCGCCGCGGTAATACCGGCAGTCCAAGTGATGGCCAATCTTATTGGGCCTAAAGCGTCCGTAGCCGGCCTCGAAAGTCCGTCGGGAAATCCACTGGCTCAACCAGTGGGTGTCCGGCGGAAACTTCGAGGCTTGGGACCGGAAGGCTCGAGGGGTACGTCCGGGGTAGGAGTGAAATCCTGTAATCCCGGACGGACCACCGATGGCGAAAGCACCTCGAGAAGACGGATCCGACGGTGAGGGACGAAAGCTAGGGTCTCGAACCGGATTAGATACCCGGGTAGTCC |
| g__Halobiforma_AB663434_Ref_seq | | | | | | | | | | | | ACGGGGCGCAGCAGGCGCGAAACCTTTACACTGCACGCCAGTGCGATAAGGGGACTCCAAGTGCGAGGGCATATAGTCCTCGCTTTTCTGGACCGTAAGGTGGTCCAGGAATAAGTGCTGGGCAAGACCGGTGCCAGCCGCCGCGGTAATACCGGCAGCACGAGTGATGGCCGCTCTTATTGGGCCTAAAGCGTCCGTAGCTGGCCGATCAAGTCCGTCGGGAAATCCGCACGCCCAACGTGCGGGCGTCCGGTGGAAACTGATCGGCTTGGGACCGGAAGACCAGAGGGGTACGTCCGGGGTAGGAGTGAAATCCCGTAATCCTGGACGGACCACCGGTGGCGAAAGCGCCTCTGGAAGACGGATCCGACGGTGAGGGACGAAAGCTCGGGTCACGAACCGGATTAGATACCCGGGTAGTCC |
| g__Halobium_LC109314_Ref_seq | | | | | | | | | | | | ACGGGGCGCAGCAGGCGCGAAACCTTTACACTGCACGCACGTGCGATAAGGGGACCCCGAGTGCGAGGGCATACTAGCCCTCGCTTTTCGTAACCGTAAGGAGGTTACAGAATAAGAGCTGGGCAAGACCGGTGCCAGCCGCCGCGGTAATACCGGCAGCTCGAGTGATGACCGATCTTATTGGGCCTAAAGCGTCCGTAGCTGGCCTCGACGGTCCGTCGGGAAATCCACCTGCTCAACAGGTGGGCGTCCGGCGGATACCACGGGGCTTGGGACCGGAAGACCCGAGGGGTACGTCCGGGGTAGGAGTGAAATCCCGTAATCCTGGACGGACCACCGATGGCGAAAGCACCTCGGGAGGACGGATCCGACAGTGAGGGACGAAAGCTCGGGTCTCGAACCGGATTAGATACCCGGGTAGTCC |
| g__Halococcus_AJ420376_Ref_seq | | | | | | | | | | | | ACGGGGCGCAGCAGGCGCGAAACCTTTACACTGCACGCCAGTGCGATAAGGGGACCCCGAGTGCGAGGGCATACAGTCCTCGCTTTTCGTGACCGTAAGAAGGTCTCAGAATAAGAGCTGGGCAAGACCGGTGCCAGCCGCCGCGGTAATACCGGCAGCTCGAGTGATAGCCACTATTATTGGGCCTAAAGCGTCCGTAGCCGGCCGAACAGGTCCGTCGGGAAATCCACCCGCTCAACGGGTGGACGTCCGGCGGAAACCAGTCGGCTTGGGGCCGGGAGACCAGAGAGGTACGTCCGGGGTAGGAGTGAAATCCTGTAATCCTGGACGGACCACCGGTAGCGAAAGCGTCTCTGGAGAACGGACCCGACGGTGAGGGACGAAAGCTTGGGTCTCGAACCGGATTAGATACCCGGGTAGTCC |
| g__Halodesulfurarchaeum_CP016804_Ref_seq | | | | | | | | | | | | ACGGGGCGCAGCAGGCGCGAAACCTTTACACTGCACGACAGTGCGATAGGGGGACTCCGAGTGCGAGGGCATAGAGCCCTCGCTTTTCTGTACCGTAAGGTGGTACAGGAAGAAGGGCTGGGCAAGACCGGTGCCAGCCGCCGCGGTAATACCGGCAGCCCAAGTGATGGCCGATATTATTGGGCCTAAAGCGTCCGTAGCTGGCCGGACAAGTCCGTTGGGAAATCCGCACGCTTAACGTGCGGGCGTCCAGCGGAAACTGTCCGGCTTGGGACCGGAAGACCCGAGGGGTACGTCCGGGGTAGGAGTGAAATCCTGTAATCTCGGACGGACCACCTGTGGCGAAAGCGCCTCGGGAAGACGGATCCGACAGTGAGGGACGAAAGCCAGGGTCTCGAACCGGATTAGATACCCGGGTAGTCC |
| g__Haloferax_AB037474_Ref_seq | | | | | | | | | | | | ACGGGGCGCAGCAGGCGCGAAACCTTTACACTGCACGCAAGTGCGATAAGGGGACCCCAAGTGCGAGGGCATATAGTCCTCGCTTTTCACGACTGTAAGGCGGTCGTGGAATAAGAGCTGGGCAAGAGCGGTGCCAGCCGCCGCGGTAATACCGGCAGCTCAAGTGATGACCGATATTATTGGGCCTAAAGCGTCCGTAGCCGGCCACGAAGGTTCATCGGGAAATCCGCCAGCTCAACTGGCGGGCGTCCGGTGAAAACCACGTGGCTTGGGACCGGAAGGCTCGAGGGGTACGTCCGGGGTAGGAGTGAAATCCCGTAATCCTGGACGGACCACCGATGGCGAAAGCACCTCGAGAAGACGGATCCGACGGTGAGGGACGAAAGCTAGGGTCTCGAACCGGATTAGATACCCGGGTAGTCC |
| g__Halogeometricum_AF002984_Ref_seq | | | | | | | | | | | | ACGGGGCGCAGCAGGCGCGAAACCTTTACACTGCACGCAAGTGCGATAAGGGGACTCCGAGTGCGAGGGCATATAGTCCTCGCTTTTGAGAACCGTAAGGCGGTTCTCGAATAAGAGCTGGGCAAGACCGGTGCCAGCCGCCGCGGTAATACCGGCAGCTCAAGTGATGGCCAATCTTATTGGGCCTAAAGCGTCCGTAGCTGGCCACGAAAGTCCGTCGGGAAATCCATCTGCCCAACAGATGGGCGTCCGGCGGAAACTTCGTGGCTTGGGACCGGAAGGCTCGAGGGGTACGTCCGGGGTAGGAGTGAAATCCCGTAATCCTGGACGGACCGCCGATGGCGAAAGCACCTCGAGAAGACGGATCCGACAGTGAGGGACGAAAGCTAGGGTCTCGAACCGGATTAGATACCCGGGTAGTCC |
| g__Halohasta_JF421969_Ref_seq | | | | | | | | | | | | ACGGGGCGCAGCAGGCGCGAAACCTTTACACTGCACGGAAGTGCGATAGGGGAATCCCAAGTGCGAAGGCATACTAGCCTTCGCTTTTGTTGACTGTAAGGAGGTCAACGAATAAGAGCTGGGCAAGACCGGTGCCAGCCGCCGCGGTAATACCGGCAGCTCGAGTGATGTCCAATATTATTGGGCCTAAAGCGTCCGTAGCTGGCCAGCCAAGTTCGTCGGGAAATCCACCTGCCCAACAGGTGGGCGTCCGGCGAAAACTGTCTGGCTTGGAACCGGAAGGCTCAGAGGGTACGTCCGGGGTAGGAGTGAAATCCTGTAATCCCGGACGGACCACCGATGGCGAAAGCACTCTGAGAAGACGGATTCGACAGTGAGGGACGAAAGCTAGGGTCTCGAACCGGATTAGATACCCGGGTAGTCC |
| g__Halolamina_KJ689297_Ref_seq | | | | | | | | | | | | ACGGGGCGCAGCAGGCGCGAAACCTTTACACTGCACGCCAGTGCGATAAGGGGACCCCGAGTGAAAGGGCATATCGCCCTTTCTTTTCTGAACCGTAGGGAGGTTCAGGAACAAGAGCTGGGCAAGACCGGTGCCAGCCGCCGCGGTAATACCGGCAGCTCAAGTGATGACCGATTTTATTGGGCCTAAAGCGTCCGTAGCTGGCCGCGCAAGTTCGTCGGGAAATCCACCTGCTTAACGGGTGGGCGTCCGGCGAAAACTGCACGGCTTGGGACAGGAAGGCTCGAGGGGTACGTTCGGGGTAGGAGTGAAATCCCGTAATCCTGAACGGACCACCGATGGCGAAAGCACCTCGAGAGGACTGATCCGACAGTGAGGGACGAAAGCTGGGGTCTCGAACCGGATTAGATACCCGGGTAGTCC |
| g__Halomarina_3560_KC918830_Ref_seq | | | | | | | | | | | | ACGGGGCGCAGCAGGCGCGAAACCTTTACACTGCACGACAGTGCGATAGGGGGATTCCGAGTGCGAGGGCATACAGTCCTCGCTTTTGTGTACCTTAAGGTGGTACACGAATAAGGGCTGGGCAAGACCGGTGCCAGCCGCCGCGGTAATACCGGCAGCTCAAGTGATGACCGATATTATTGGGCCTAAAGCGTCCGTAGCCGGCCAGGTAGGTCCGTCGGGAAATCCACTCGCCTAACGAGTGGGCGTCCGGCGGATACCGCCTGGCTTGGGACCGGGAGACCAGAGGGGTACGTCTGGGGTAGGAGTGAAATCCTGTAATCCTAGACGGACCATCGGTGGCGAAAGCGCCTCTGGAGAACGGATCCGACGGTGAGGGACGAAAGCTAGGGTCTCGAACCGGATTAGATACCCGGGTAGTCC |
| g__Halomicrobium_3849_HM063952_Ref_seq | | | | | | | | | | | | ACGGGGCGCAGCAGGCGCGAAACCTTTACACTGCACGACAGTGCGATAAGGGGACTCCGAGTGCGAGGGCATATAGTCCTCGCTTTTCTGTACCGTAAGGTGGTACAGGAACAAGTGCTGGGCAAGACCGGTGCCAGCCGCCGCGGTAATACCGGCAGCACGAGTGATGGCCGCTCTTATTGGGCCTAAAGCGTCCGTAGCCGGCCAGACAAGTCCGTTGGGAAATCGACGCGCTCAACGCGTCGACGTCCAGCGGAAACTGTCTGGCTTGGGGCCGGAAGACCTGAGGGGTACGTCCGGGGTAGGAGTGAAATCCTGTAATCCTGGACGGACCACCGGTGGCGAAAGCGCCTCAGGAAGACGGACCCGACGGTGAGGGACGAAAGCTAGGGTCTCGAACCGGATTAGATACCCGGGTAGTCC |
| g__Halonotius_AB663393_Ref_seq | | | | | | | | | | | | ACGGGGCGCAGCAGGCGCGAAACCTTTACACTGCACGAAAGTGCGATAGGGGAATCCCGAGTGCGAGGGCATATTAGCCCTCGCTTTTGTTGACTCTAAGGCGGTCAACGAATAAGAGCTGGGCAAGACCGGTGCCAGCCGCCGCGGTAATACCGGCAGCTCAAGTGATGTCCCATATTATTGGGCCTAAAGCGTCCGTAGCTGGCCGACCAAGTCTATCGGGAAATCCACCTGCCCAACAGGTGGGCGTCCGGTAGAAACTGGCCGGCTTGGAACCGGAAGGCTCAGAGAGTACGTCCGGGGTAGGAGTGAAATCCCGTAATCCCGGACGGACTACCGATGGCGAAAGCACTCTGAGAAGACGGCTTCGACAGTGAGGGACGAAAGCTCGGGTCTCAAACCGGATTAGATACCCGGGTAGTCC |
| g__Halopelagius_JQ996497_Ref_seq | | | | | | | | | | | | ACGGGGCGCAGCAGGCGCGAAACCTTTACACTGCACGCAAGTGCGATAAGGGGACTCCGAGTGCGAGGGCATATAGTCCTCGCTTTTCTGTACCGTAGGAAGGTACAGGAATAAGGGCTGGGCAAGACCGGTGCCAGCCGCCGCGGTAATACCGGCAGCCCGAGTGATGACCGATTTTATTGGGCCTAAAGCGTCCGTAGCTGGCCAAGCAAGTCTGTCGGGAAATCCATCCGCTCAACGGATGGGCGTCCGGCGGAAACTCCTTGGCTTGGGACCGGAAGGCTCGAGGGGTACGTCTGGGGTAGGAGTGAAATCCTGTAATCCTGGACGGACCACCGATGGCGAAAGCACCTCGAGAAGACGGATCCGACAGTGAGGGACGAAAGCTAGGGTCTCGAACCGGATTAGATACCCGGGTAGTCC |
| g__Halopiger_A_AB663437_Ref_seq | | | | | | | | | | | | ACGGGGCGCAGCAGGCGCGAAACCTTTACACTGCACGCCAGTGCGATAAGGGGACTCCATCTGCGAGGGCATATCGTCCTCGCTTTTCTGTACCGTAAGGTGGTACAGGAATAAGGGCTGGGCAAGACCGGTGCCAGCCGCCGCGGTAATACCGGCAGCCCGAGTGATGACCGCTATTATTGGGCCTAAAGCGTCCGTAGCTGGCCAGGCAAGTCCATCGGGAAATCCGCGTGCCCAACACGCGGGCGTCCGGTGGAACCTGCTTGGCTTGGGACCGGAAGACCTGAGGGGTACGTCTGGGGTAGGAGTGAAATCCCGTAATCCTGGACGGACCACCGGTGGCGAAAGCGCCTCAGGAAGACGGATCCGACGGTGAGGGACGAAAGCTCGGGTCACGAACCGGATTAGATACCCGGGTAGTCC |
| g__Haloplanus_AB663396_Ref_seq | | | | | | | | | | | | ACGGGGCGCAGCAGGCGCGAAACCTTTACACTGCACGCAAGTGCGATAAGGGGACCCCGAGTGCGAGGGCATACAGTCCTCGCTTTTCACTACCGTAAGGTGGTAGTGGAATAAGAGCTGGGCAAGACCGGTGCCAGCCGCCGCGGTAATACCGGCAGCTCGAGTGATGGCCGATCTTATTGGGCCTAAAGCGTCCGTAGCCGGCCACACAAGTCCATCGGGAAATCCGCCAGCCCAACTGGCGGGCGTCCGGTGGAAACTGTGTGGCTCGGGATCGGAAGACCCAACGGGTACGTCTTGGGTAGGAGTGAAATCCTGTAATCCTGGACGGACCACCGATAGCGAAAGCACGTTGGGAAGACGAATCCGACGGTGAGGGACGAAAGCCAGGGTCTCGAACCGGATTAGATACCCGGGTAGTCC |
| g__Haloprofundus_KJ999759_Ref_seq | | | | | | | | | | | | ACGGGGCGCAGCAGGCGCGAAACCTTTACACTGCACGACAGTGCGACAAGGGGACCCCGAGTGCGAGGGCATACAGTCCTCGCTTTTGTGAACCGTAAGGTGGTTCACGAATAAGAGCTGGGCAAGACCGGTGCCAGCCGCCGCGGTAATACCGGCAGCTCGAGTGATGACCGATTTTATTGGGCCTAAAGCGTCCGTAGCTGGCCGCGTAAGTTCATCGGGAAATCCATCCGCTCAACGGATGGGCGTCCGGTGAAAACTACACGGCTTGGAGCCGGAAGACCCAAGGGGTACGTCCGGGGTAGGAGTGAAATCCCGTAATCCTGGACGGACCGCCGATGGCGAAAGCACCTTGGGAAGACGGACTCGACAGTGAGGGACGAAAGCTAGGGTCTCGAACCGGATTAGATACCCGGGTAGTCC |

| | Presence (1) or Absence (0) in each layer | | | | | | | | | | | |
|---|---|---|---|---|---|---|---|---|---|---|---|---|
| Phylum_Genus_ASV_number | L1 | L2 | L3 | L4 | L5 | L6 | L7 | L8 | L9 | L10 | L11 | Sequence |
| g__Halorhabdus_AB663399_Ref_seq | | | | | | | | | | | | ACGGGGCGCAGCAGGCGCGAAACCTTTACACTGCACGACAGTGCGATAGGGGGACTCCGAGTGCGAGGGCATATAGTCCTCGCTTTTCTGTACCGTAAGGTGGTACAGGAATAAGGGCTGGGCAAGACCGGTGCCAGCCGCCGCGGTAATACCGGCAGCCCGAGTGATGGCCGCTATTATTGGGCCTAAAGCGTCCGTAGCCGGCCAGACAAGTCCGTTGGGAAATCCACGCGCTTAACGCGTGGACGTCCGGCGGAAACTGTCTGGCTTGGGGCCGGAAGATCTGAGGGGTACGTCCGGGGTAGGAGTGAAATCCCGTAATCCTGGACGGACCGCCGGTGGCGAAAGCGCCTCAGAAAGACGGACCCGACGGTGAGGGACGAAAGCTAGGGTCTCGAACCGGATTAGATACCCGGGTAGTCC |
| g__Halorientalis_KF313134_Ref_seq | | | | | | | | | | | | ACGGGGCGCAGCAGGCGCGAAACCTTTACACTGCACGACAGTGCGATAAGGGGACTCCGAGTGCGAGGGCATACAGTCCTCGCTTTTCTGTACCGTAAGGTGGTACAGGAACAAGTGCTGGGCAAGACCGGTGCCAGCCGCCGCGGTAATACCGGCAGCACGAGTGATGGCCGCTATTATTGGGCCTAAAGCGTCCGTAGCCGGCCGAGCAAGTTCGTTGGGAAATCTGCCCGCTTAACGGGCAGGCGTCCGGCGAAAACTGTTCGGCTTGGGCCGGGAGATCTGAGGGGTACGTCCGGGGTAGGAGTGAAATCCCGTAATCCTGGACGGACCACCGGTGGCGAAAGCGCCTCAGAAAAACGGACCCGACGGTGAGGGACGAAAGCTAGGGTCTCGAACCGGATTAGATACCCGGGTAGTCC |
| g__Halorubrum_AB663424_Ref_seq | | | | | | | | | | | | ACGGGGCGCAGCAGGCGCGAAACCTTTACACTGCACGACAGTGCGATAGGGGGATCCCAAGTGCACAGGCATAGCGCCTGTGCTTTTCGGTACCGTAAGGTGGTACCAGAATAAGGGCTGGGCAAGACCGGTGCCAGCCGCCGCGGTAATACCGGCAGCCCAAGTGATGGCCGATCTTATTGGGCCTAAAGCGTCCGTAGCTGGCCGCGCAAGTCCATCGGGAAATCCACCTGCTCAACAGGTGGGCGCCCGGTAGAAACTGCGTGGCTTGGGACCGGAAGGCGCGACGGGTACGTCCGGGGTAGGAGTGAAATCCCGTAATCCTGGACGGACCGCCGATGGCGAAAGCACGTCGCGAGAACGGATCCGACAGTGAGGGACGAAAGCCAGGGTCTCGAACCGGATTAGATACCCGGGTAGTCC |
| g__Halorussus_KJ689292_Ref_seq | | | | | | | | | | | | ACGGGGCGCAGCAGGCGCGAAAACTTTACACTGCACGCGAGTGCGATAAGGGGACTCCGAGTGCGAGGGCATAGAGCCCTCGCTTTTCTGTACCGTAAGGTGGTACAGGAATAAGAACTGGGCAAGACCGGTGCCAGCCGCCGCGGTAATACCGGCAGTTCGAGTGATGGCCGATTTTATTGGGCCTAAAGCGTCCGTAGCTTGCTAGACAGGTTCATCGGGAAATCTACCCGCCTAACGGGTAGGCGTCCGGTGAAAACCAGCTAGCTTGGGGCCGGAAGATCCAAGGGGTACGTCCGGGGTAGGAGTGAAATCCCGTAATCCTGGACGGACCACCGATGGCGAAAGCACCTTGGAAAGACGGACCCGACGGTGAGGGACGAAAGCTAGGGTCACGAACCGGATTAGATACCCGGGTAGTCC |
| g__Halosimplex_AB072815_Ref_seq | | | | | | | | | | | | ACGGGGCGCAGCAGGCGCGAAACCTTTACACTGCACGACAGTGCGATAGGGGAACTCCGAGTGCGAGGGCATAGAGCCCTCGCTTTTCCACACCGTAAGGTGGTGCGGGAACAAGTGCTGGGCAAGACCGGTGCCAGCCGCCGCGGTAATACCGGCAGCACAAGTGATGGCCGATTTTATTGGGCCTAAAGCGTCCGTAGCTGGCCGTGCAAGTCTGTCGGGAAATCCAGTGGCTCAACCTCTGGGCTTCCGGCGGAAACTGCTCGGCTTGGGACCGGAAGACTCGAGGGGTACGTTCGGGGTAGGAGTGAAATCCTGTAATCCTGAACGGACCACCGATGGCGAAAGCACCTCGAGAGCACGGATCCGACAGTGAGGGACGAAAGCTAGGGTCTCGAACCGGATTAGATACCCGGGTAGTCC |
| g__Halostagnicola_AB663439_Ref_seq | | | | | | | | | | | | ACGGGGCGCAGCAGGCGCGAAACCTTTACACTGCACGAGAGTGCGATAAGGGGACTCCAAGTGCGAGGGCATATAGTCCTCGCTTTTTCCGACCGTAGGATGGTCGGCGAATAAGTGCTGGGCAAGACCGGTGCCAGCCGCCGCGGTAATACCGGCAGCACAAGTGATGACCGCTATTATTGGGCCTAAAGCGTCCGTAGCTGGCCACGCAAGTCCATCGGGAAATCTACTCGCTCAACGAGTAGGCGTCCGGTGGAAACTGCATGGCTTGGGACCGGAAGATCCAGAGGGTACGTCTGGGGTAGGAGTGAAATCCTGTAATCCGAGACGGACCACCGGTGGCGAAAGCGCTCTGGAAAGACGGATCCGACGGTGAGGGACGAAAGCTCGGGTCACGAACCGGATTAGATACCCGGGTAGTCC |
| g__Halostella_KM068159_Ref_seq | | | | | | | | | | | | ACGGGGCGCAGCAGGCGCGAAACCTTTACACTGCACGCGAGTGCGATAAGGGGACCCCGAGTGCGAGGGCATATAGCCCTCGCTTTTGCACACCGTACTGAGGTGCGCGAATAAGGACTGGGCAAGACCGGTGCCAGCCGCCGCGGTAATACCGGCAGTCCAAGTGATGGCCGCTGTTATTGGGCCTAAAGCGTCCGTAGCCGGCCAGGCAAGTCCATCGGGAAATCCACCCGCTCAACGGGTGGGCGTCCGGTGGAAACTGTCTGGCTAGGGACCGGAAGACCCGAGGGGTACGTCCAGGGTAGGAGTGAAATCCCGTAATCCTGGACGGACCGCCGGTGGCGAAAGCGCCTCGGGAAGACGGATCCGACGGTGAGGGACGAAAGCTTGGGTCACGAACCGGATTAGATACCCGGGTAGTCC |
| g__Halovenus_HM159605_Ref_seq | | | | | | | | | | | | ACGGGGCGCAGCAGGCGCGAAACCTTTACACTGCACGACAGTGCGATAGGGGGACTCCGAGTGCGAGGGTATACAGTCCTCGCTTTTCTGTACCGTAAGGTGGTACAGGAACAAGTGCTGGGCAAGACCGGTGCCAGCCGCCGCGGTAATACCGGCAGCACGAGTGATGGCCGATCTTATTGGGCCTAAAGCGTCCGTAGCTGGCCAGTCAAGTCTGTTGGGAAATCCACGTGCCCAACGCGTGGGCGTCCAGCGGAAACTGTCTGGCTTGGAGCCGGAGGACTCGAGGGGTACGTCCGGGGTAGGAGTGAAATCCCGTAATCCTGGACGGACCACCGGTGGCGAAAGCGCCTCGAGAGGACGGACTCGACAGTGAGGGACGAAAGCCAAGGTCTCGAACCGGATTAGATACCCGGGTAGTCT |
| g__Halovivax_AOIQ01000016_Ref_seq | | | | | | | | | | | | ACGGGGCGCAGCAGGCGCGAAACCTTTACACTGCACGCGAGTGCGATAGGGGGACTCCAAGTGCGAGGGCATACAGTCCTCGCTTTTCACCACCGTAAGGAGGTGGTAGAATAAGTGCTGGGCAAGACCGGTGCCAGCCGCCGCGGTAATACCGGCAGCACGAGTGATGACCGCTATTATTGGGCCTAAAGCGTCCGTAGCCGGCCGGGCAAGTCCATCGGGAAATCCGCACGCCTAACGTGCGGGCGTCCGGTGGAAACTGCTTGGCTTGGGACCGGAAGATCCAGAGGGTACGTCTGGGGTAGGAGTGAAATCCTGTAATCCTGGACGGACCACCGGTGGCGAAAGCGCTCTGGAAAGACGGATCCGACGGTGAGGGACGAAAGCTTGGGTCACGAACCGGATTAGATACCCGGGTAGTCC |
| g__Methanobacterium_D_1054_DQ517520_Ref_seq | | | | | | | | | | | | ACGGGGCGCAGCAGGCGCGAAACCTCCGCAATGCACGAAAGTGCGACGGGGGGACCCCAAGTGCCGCTCTTAACGGGGTGGCTTTTTTTGAGTGTAAAAAGCTTTGAGAATAAGAGCTGGGCAAGACCGGTGCCAGCCGCCGCGGTAACACCGGCAGCTCAAGTGGTGGCCATTTTTATTGGGCCTAAAGCGTTCGTAGCCGGCTTGATAAGTCTCTGGTGAAATCCCATAGCTTAACTGTGGGAATTGCTGGAGATACTATTAGGCTTGAGGCCGGGAGAGGTTAGGGGTACTCCCAGGGTAGGGGTGAAATCCTATAATCCTGGGAGGACCACCTGTGGCGAAGGCGCCTAACTGGAACGGACCTGACGGTGAGTAACGAAAGCCAGGGGCGCGAACCGGATTAGATACCCGGGTAGTCC |
| g__Methanobrevibacter_A_U55237_Ref_seq | | | | | | | | | | | | ACGGGGCGCAGCAGGCGCGAAACCTCCGCAATGTGAGAAATCGCGACGGGGGGATCCCAAGTGCCATTCTTAACGGGATGGCTTTTCATTAGTGTAAAGAGCTTTTGGAATAAGAGCTGGGCAAGACCGTTGCCAACCGCCGCGGTAACACCGTCAGCTCTAGTGGTAGCAGTTTTTATTGGGCCTAAAGCGTCCGTAGCCGGTTTATTAAGTCTCTGGTGAAATCCTGTAGCTTAACTGTGGGAATTGCTGGAGATACTAGTAGACTTGAGATCGGGAGAGGTTAGAGGTACTCCCAGGGTAGAGGTGAAATTCTGTAATCCTGGGAGGACCGCCTGTGGCGAAGGCGTCTAGCTGGAACGATTCTGCCGGTGAGGGACGAAAGCTAGGGGCGCGAACCGGATTAGATACCCGGGTAGTCC |
| g__Methanohalobium_FR733675_Ref_seq | | | | | | | | | | | | ACGGGGCGCAGCAGGCGCGAAACCTTTACACTGCGGGAAACCGCGATAAGGGGACACTGAGTGCCAGCATATAATGCTGGCTGTCCGTATGTGTAAAATACATGCGTTAGCAAGGGCCGGGCAAGACCGGTGCCAGCCGCCGCGGTAACACCGGCGGCCCGAGTGGTGATCGCTATTATTGGGTCTAAAGGGTCCGTAGCCGGTTTGATCAGTCCATTGGGAAATCTGGTGGCTTAACCGTCAGAATTCCAATGGATACTGTCAGACTTGGGACCGGGAGAGGTAAGAGGTACTACAGGGGTAGGAGTGAAATCTTGTAATCCCTGTAGGACCACCAGTTGCGAAGGCGTCTTACCAGAACGGGTCCGACGGTGAGGGACGAAAGCTGGGGGCACGAACCGGATTAGATACCCGGGTAGTCC |
| g__Methanohalophilus_FR733671_Ref_seq | | | | | | | | | | | | ACGGGGCGCAGCAGGCGCGAAACCTTTACACTGCGGGAAACCGCGATAAGGGGACACCGAGTGCCAGCATACAATGCTGGCTGTCCACATGTGTAAATAGCATGTGATAGCAAGGGCCGGGCAAGACCGGTGCCAGCCGCCGCGGTAACACCGGCGGCCCGAGTGGTAATCACTATTATTGGGTCTAAAGGGTCCGTAGCCGGTTTAATCAGTCTTCCGGGAAATCTGACAGCTCAACTGTTAGGCTTCCGGGGGATACTGTTAGACTTGGGACCGGGAGAGGTAAGAGGTACTACAGGGGTAGGAGTGAAATCTTGTAATCCCTGTGGGACCACCAGTGGCGAAGGCGTCTTACCAGAACGGGTCCGACGGTGAGGGACGAAAGCTGGGGGCACGAACCGGATTAGATACCCGGGTAGTCC |
| g__Methanolobus_FR733684_Ref_seq | | | | | | | | | | | | ACGGGGCGCAGCAGGCGCGAAATCTTTACAATGCGGGGAACCGCGATAAGGGGATACTGAGTGCCAGCATATTATGTTGGCTGTCCACCTGTATAAACAGCAGGTGTTAGCAAGGGCCGGGCAAGACCGGTGCCAGCCGCCGCGGTAACACCGGCGGCCCGAGTGGTGGCCACTATTATTGGGTCTAAAGGGTCCGTAGCCTGTTCGATCAGTCTTCCGGGAAATCTGACAGCTCAACTGTTAGGCTTCCGGGGGATACTGTCGGACTTGGGACCGGGAGAGGTAAGAGGTACTACAGGGGTAGGAGTGAAATCTTGTAATCCTTGTGGGACCACCAGTGGCGAAGGCGTCTTACCAGAACGGGTCCGACGGTGAGGGACGAAAGCTGGGGGCACGAACCGGATTAGATACCCGGGTAGTCC |
| g__Methanospirillum_CP000254_Ref_seq | | | | | | | | | | | | ACGGGGCGCAGCAGGCGCGAAAACTTTACCATGCGGGCAACCGTGATAAGGAAACCCCGAGTGCCAGCACAGGCTGGCTGTCCACCAGTGTAAATAACTGGTGAAGAAAGGGCCGGGCAAGACCGGTGCCAGCCGCCGCGGTAATACCGGCGGCTCGAGTGGTGGCCGCTATTACTGGGCTTAAAGGGTCCGTAGCTGGATATACAAGTCCCTTGAGAAATCCGCCGGCTTAACCGGTGGGCGTTCAGGGGAAACTGTATTTCTAGGGACCGGGAGAGGTGAGAGGTACTGCCGGGGTAGGAGTGAAATCCTGTAATCCCGGTGGGACCACCTATGGCGAAGGCATCTCACCAGAACGGGTCCGACAGTGAGGGACGAAAGCTGGGGGAGCAAACCGGATTAGATACCCGGGTAGTCC |
| g__Methanothrix_A_AY817738_Ref_seq | | | | | | | | | | | | ACGGGGTGCAGCAGGCGCGAAAACTTTACAATGCGGGAAACCGTGATAAGGGAATCTCGAGTGCCAGCATATAATGTTGGCTGTCCAGATGCCTAAAAAGCATCTGTTAGCAAGGGCCGGGCAAGACCGGTGCCAGCCGCCGCGGTAACACCGGCGGCCCGAGTGGTAACCGCTTTTATTGGGTCTAAAGGGTCTGTAGCCGGCCAAGTAAGTCCCTTGGGAAATCTGGCAGCTTAACTGTCAGGCTGCTAGGGGATACTGCTGGGCTTGGGACCGGGAGAGGTGAGAGGTACCTTGGGGGTAGGGGTGAAATCTTGTGATCCTCGGGGGACCACCAGTGGCGAAGGCGTCTCACCAGAACGGGTCCGACGGTAAGGGACGAAAGCTAGGGGCACGAACCGGATTAGATACCCGGGTAGTCC |
| g__Natrarchaeobaculum_KY612364_Ref_seq | | | | | | | | | | | | ACGGGGCGCAGCAGGCGCGAAACCTTTACACTGCACGCGAGTGCGATAAGGGGACTCCAAGTGCGAGGGCATATCGTCCTCGCTTTTTCCTACCGTAAGGTGGTAGGCGAATAAGGGCTGGGCAAGACCGGTGCCAGCCGCCGCGGTAATACCGGCAGCCCAAGTGATGACCGCTATTATTGGGCCTAAAGCGTCCGTAGCTGGCCAGGCAAGTCCGTCGGGAAATCCGCGCGCTCAACGCGCGGGCGTCCGGTGGAAACTGCGTGGCTTGGGACCGGAAGACCTGAGGGGTACGTCTGGGGTAGGAGTGAAATCCCGTAATCCTGGACGGACCACCGGTGGCGAAAGCGCCTCAGGAAGACGGATCCGACGGTGAGGGACGAAAGCTCGGGTCACGAACCGGATTAGATACCCGGGTAGTCC |
| g__Natrinema_AB663444_Ref_seq | | | | | | | | | | | | ACGGGGCGCAGCAGGCGCGAAACCTTTACACTGCACGAAAGTGCGATAAGGGGACTCCAAGTGCGAGGGCATATAGTCCTCGCTTTTTGCGACCGTAAGGTGGTCGCGGAATAAGTGCTGGGCAAGACCGGTGCCAGCCGCCGCGGTAATACCGGCAGCACGAGTGATGACCGCTATTATTGGGCCTAAAGCGTCCGTAGCTGGCCGTGCAAGTCCATCGGGAAATCCGCGCGCTTAACGCGCGGGCGTCCGGTGGAAACTGCATGGCTTGGGACCGGAAGACCAGAGGGGTACGTCCGGGGTAGGAGTGAAATCCCGTAATCCTGGACGGACCACCGGTGGCGAAAGCGCCTCTGGAAGACGGATCCGACGGTGAGGGACGAAAGCTCGGGTCACGAACCGGATTAGATACCCGGGTAGTCC |
| g__Natronoarchaeum_AB457580_Ref_seq | | | | | | | | | | | | ACGGGGCGCAGCAGGCGCGAAACCTTTACACTGCACGACAGTGCGATAAGGGGATCCCCAGTGCGAGGGCATACAGTCCTCGCTTTTCACCACCGTAAGGTGGTGGTGGAACAAGAGCTGGGCAAGACCGGTGCCAGCCGCCGCGGTAATACCGGCAGCTCGAGTGATGGCCGTTCTTATTGGGCCTAAAGCGTCCGTAGCCGGCCACATAAGTTCGTCGGGAAATCCACATGCTCAACATGTGGGCGTCCGGCGAAACCTGTGTGGCTTGGGACCGGACGGCCCAGAGGGTACGGCCGGGGTAGGAGTGAAATCCTGTAATCCCGGCCGGACCACCGGTGGCGAAAGCGCTCTGGGAGGACGGATCCGACGGTGAGGGACGAAAGCCAGGGTCACGAACCGGATTAGATACCCGGGTAGTCC |
| g__Natronobiforma_KT247980_Ref_seq | | | | | | | | | | | | ACGGGGCGCAGCAGGCGCGAAACCTTTACACTGCGCGACAGCGCGATAAGGGGACTCCAAGTGCGAGGGCGTATAGTCCTCGCTTTTAACTACCGTAAGGAGGTAGTAGAATAAGGGCTGGGCAAGACCGGTGCCAGCCGCCGCGGTAATACCGGCAGCCCGAGTGATGACCGCTATTATTGGGCCTAAAGCGTCCGTAGCTGGCCACACAAGTCCGTCGGGAAATCTGCTCGCCTAACGAGCAGGCGTCCGGCGGAAACTGTGTGGCTTGGGACCGGAAGATCCAGAGGGTACGTCTGGGGTAGGAGTGAAATCCTGTAATCCTGGACGGACCACCGGTGGCGAAAGCGCTCTGGAAGGACGGATCCGACAGTGAGGGACGAAAGCTAGGGTCACGAACCGGTATTAGATACCCGGGTAGTCC |
| g__Natronomonas_KX898188_Ref_seq | | | | | | | | | | | | ACGGGGCGCAGCAGGCGCGAAACCTTTACACTGCACGACAGTGCGATAGGGGGACACCGAGTGCGAGGGCATAGAGCCCTCGCTTTTCTGGACCTTAAGGTGGTCCAGGAACAAGGGCTGGGCAAGACCGGTGCCAGCCGCCGCGGTAACACCGGCAGCCCAAGTGATGGCCGATATTATTGGGCCTAAAGCGTCCGTAGCTGGCCGGGCAAGTCCGTCGGGAAATCCACGCGCTTAACGCGTGGGCGTCCGGCGGAAACTGTTCGGCTTGGGACCGGAAGATCCGAGGGGTACGTCCGGGGTAGGAGTGAAATCCCGTAATCCTGGACGGACCACCGATGGCGAAAGCACCTCGGAAGGACGGATCCGACAGTGAGGGACGAAAGCTAGGGTCTCGAACCGGATTAGATACCCGGGTAGTCC |
| g__Natronorubrum_4312_AB935414_Ref_seq | | | | | | | | | | | | ACGGGGCGCAGCAGGCGCGAAACCTTTACACTGCACGCCAGTGCGATAAGGGGACTCCGAGTGCGAGGGCATATAGTCCTCGCTTTTCACCACCGTAGGAAGGTAGTGGAATAAGTGCTGGGCAAGACCGGTGCCAGCCGCCGCGGTAATACCGGCAGCACGAGTGATGACCGCTATTATTGGGCCTAAAGCGTCCGTAGCTGGCCGCGCAAGTCTATCGGGAAATCTGCTCGCCTAACGAGCAGGCGTCCGGTGGAAACTGTGTGGCTTGGGACCGGAAGACCAGAGGGGTACGTCTGGGGTAGGAGTGAAATCCCGTAATCCTGGACGGACCACCGGTGGCGAAAGCGCCTCTGGAGGACGGATCCGACGGTGAGGGACGAAAGCTCGGGTCACGAACCGGATTAGATACCCGGGTAGTCC |
| g__Nitrososphaera_CP007536_Ref_seq | | | | | | | | | | | | ATGGGGCGCAGCAGGCGCGAAACCTCTGCAATAGGCGAAAGCTTGACAGGGTTACTCTGAGTGACGTCCGTTAAGGACGTCTTTTGGCACCTCTAAAAATGGTGCAGAATAAGGGGTGGGCAAGTCTGGTGTCAGCCGCCGCGGTAATACCAGCACCCCGAGTGGTCGGGACGATTATTGGGCCTAAAGCATCCGTAGCCGGTTCTGCAAGTCTTCCGTTAAATCCACCTGCTCAACAGATGGGCTGCGGAGGATACTACAGGGCTAGGAGGCGGGAGAGGCAAGCGGTACTCAGTGGTAGGGGTAAAATCCTCTGATCCACTGAAGACCACCAGTGGCGAAGGCGGCTTGCCAGAACGCGCTCGACGGTGAGGGATGAAAGCTGGGGGAGCAAACCGGATTAGATACCCGGGTAGTCC |
| g__Salinarchaeum_HM159601_Ref_seq | | | | | | | | | | | | ACGGGGCGCAGCAGGCGCGAAACCTTTACACTGCACGCGAGTGCGATAGGGGGACCCCGAGTGCGAGGGCATACAGTCCTCGCTTTTCACCACTGTAAGGTGGTGGTGGAATAAGTGCTGGGCAAGACCGGTGCCAGCCGCCGCGGTAATACCGGCAGCACGAGTGATGGCCGATCTTATTGGGCCTAAAGCGTCCGTAGCCGGCCAGACAGGTCCATCGGGAAATCTGCACGCCCAACGTGCAGGCGTCCGGTGGAAACCAGTTGGCTTGGGGCCGGAAGATCCAGGGGGTACGTCCGGGGTAGGAGTGAAATCCCGTAATCCCGGACGGACCACCGGTGGCGAAAGCGCCCTGGAAGGACGGACCCGACGGTGAGGGACGAAAGCTAAGGTCACGAACCGGATTAGATACCCGGGTAGTCT |
| g__Salinibaculum_KT261638_Ref_seq | | | | | | | | | | | | ACGGGGCGCAGCAGGCGCGAAACCTTTACACTGCACGAGAGTGCGATAGGGAGACTCCGAGTGCGAGGGCATACAGTCCTCGCTTTTCTGTACCGTAAGGTGGTACAGGAACAAGTGCTGGGCAAGACCGGTGCCAGCCGCCGCGGTAATACCGGCAGCACGAGTGATGGCCGATATTATTGGGCCTAAAGCGTCCGTAGCTGGCCGAGCAAGTCCGTTGGGAAATCGACGCGCCTAACGCGTCGGCGTCCAGCGGAAACTGTTCGGCTTGGGGCCGGAAGACCCGAGGGGTACGTCCGGGGTAGGAGTGAAATCCCGTAATCTTGGACGGACCGCCGGTGGCGAAAGCGCCTCGGGAGGACGGACCCGACAGTGAGGGACGAAAGCCAGGGTCTCGAACCGGATTAGATACCCGGGTAGTCC |
| g__Salinigranum_GU951431_Ref_seq | | | | | | | | | | | | ACGGGGCGCAGCAGGCGCGAAACCTTTACACTGCACGCCAGTGCGATAAGGGGACCCCGAGTGCGAGGGCATACTAGCCCTCGCTTTTCACAACCGTAGGGAGGTTGTGGAACAAGAGCTGGGCAAGACCGGTGCCAGCCGCCGCGGTAATACCGGCAGCTCGAGTGATGACCGCTGTTATTGGGCCTAAAGCGTCCGTAGCTGGCCACGCAAGTCCGTCGGGAAATCCACTAGCTCAACTAGTGGGCGTCCGGCGGAAACTGCGTGGCTTGGGACCGGAAGGCTCGAGGGGTACGTCCGGGGTAGGAGTGAAATCCCGTAATCCTGGACGGACCGCCGATGGCGAAAGCACCTCGAGAGAACGGATCCGACAGTGAGGGACGAAAGCTGGGGTCTCGAACCGGATTAGATACCCGGGTAGTCC |

| Phylum_Genus_ASV_number | Presence (1) or Absence (0) in each layer L1 | L2 | L3 | L4 | L5 | L6 | L7 | L8 | L9 | L10 | L11 | Sequence |
|---|---|---|---|---|---|---|---|---|---|---|---|---|
| g__Saliphagus_KY419833_Ref_seq | | | | | | | | | | | | ACGGGGCGCAGCAGGCGCGAAACCTTTACACTGCACGACAGTGCGATAGGGGGACTCCAAGTGCGAGGGCATATAGTCCTCGCTTTTCACTACCGTAAGGTGGTAGTAGAATAAGTGCTGGGCAAGACCGGTGCCAGCCGCCGCGGTAATACCGGCAGCACGAGTGATGACCGCTATTATTGGGCCTAAAGCGTCCGTAGCCGGCCGAACAAGTCCGTCGGGAAATCCGCACGCCTAACGTGCGGGCGTCCGGCGGAAACTGTTCGGCTTGGGACCGGAAGGCTCGAGGGGTACGTCCGGGGTAGGAGTGAAATCCCGTAATCCTGGACGGACCACCGGTGGCGAAAGCGCCTCGAGAGGACGGATCCGACGGTGAGGGACGAAAGCTAGGGTCACGAACCGGATTAGATACCCGGGTAGTCC |
| GCA_000496195.1_A07HB70_Ref_seq | | | | | | | | | | | | ACGGGGCGCAGCAGGCGCGAAACCTTTACACTGCACGCGAGTGCGATAAGGGGACTCCAAGTGCGAGGGCATACAGTCCTCGCTTTTCTGGACCGTAAGGTGGTCCAGGAACAAGTGCTGGGCAAGACCGGTGCCAGCCGCCGCGGTAATACCGGCAGCACGAGTGATGGCCGATCTTATTGGGCCTAAAGCGTCCGTAGCCGGCCGAGCGGGTCCGTCGGGAAATCCACGTGCTCAACGCGTGGGCGTCCGGCGGAAACCACTCGGCTTGGGGCCGGAAGACTCGAGGGGTACGTCTGGGGTAGGAGTGAAATCCCGTAATCCTGGACGGACCGCCGATGGCGAAAGCACCTCGAGAAGACGGACCCGACGGTGAGGGACGAAAGCCAGGGTCTCGAACCGGATTAGATACCCGGGTAGTCC |
| GCA_001399795.1_BA2_Ref_seq | | | | | | | | | | | | ACGGGGCGCAGCAGTCGCGAAAACTCCGCAATGCGCGAAAGCGTGACGGGGCTACCCCGAGTGCCGTCCGCTGAGGATGGCTTTTCCCCGGTGTAATGAGCCTGGGGAATAAGGAGAGGGCAAGCCTGGTGTCAGCCGCCGCGGTAATACCAGCTCTCCGAGTGGTAGGGATGATTATTGGGCTTAAAGCGTCCGTAGCCAGCCCGGCAAGTCTCCCGTTAAATCCAGCGACCTAATCGTTGGGCTGCGGAAGATACTGTTGGGCTAGGGGGCGGGAGAGGCCGACGGTATTCCCGGGGTAGGGGTGAAATCCTATAATCCTGGAGGACCACCAGTGGCGAAGGCTGTCGGCTAGAACGCGCTCGACGGTGAGGGACGAAAGCTGGGGGAGCGAACTGGATTAGATACCCGGGTAGTCC |
| GCA_001443365.1_CSP1-1_Ref_seq | | | | | | | | | | | | ATGGGGCGCAGCAGGCGAGAAACCTTTGCAATGTGCGCAAGCACGACAAGGTTAATCCGAGTGTTTCCTGCTAAAGGAATCTTTTGTTAGTTCTAAAACCACTGATGAATAAGGGGTGGCAAGTTCTGGTGTCAGCCGCCGCGGTAAAACCAGCACCTCAAGTGGTCAGGAGGATTATTGGGCCTAAAGCATCCGTAGCCGGTTCTGTAAGTTTCCGGTTAAATCCACTAGCTCAACTAATGGGCTGCCGGAAATACTGCAGAACTAGGGAGTGGGAGAGGTAGACGGTACTCGGGAGGAAGGGGTAAAATCCTTTGATCTCTTGATGACCACCTGTGGCGAAGGCGGTCTACCAGAACACGTTCGACGGTGAGGGATGAAAGCTGGGGGAGCAAACCGGATTAGATACCCGGGTAGTCC |
| GCA_001595885.1_SG8-5_Ref_seq | | | | | | | | | | | | ACGGGGCGCAGCAGTCGCGAAAACTTGTCAATGCGGGAAACCGCGAACAGGGGAGTCCAAGTGCATGCACAATGTGTATGCTGTTCGGTTGCCTAAAAAGCAGCCGAAGTAAGGGCTGGGTAAGACAGGTGCCAGCCGCCGCGGTAATACCTGCAGCCCGAGTGGTGATCGATATTATTGAGCCTAAAGCGTCCGTAGCCGGTCTGGTAAACTCTTGGGTAAATTGGGCAGCTCAACTGTCCGAATTCCGAGAGGACTGCCAGACTTGGGACCGGGAGAGGTCAGAGGTACTCCAGGGGTAGGGGCTAAATCCTGTAATCCTTGGGGGACCACCAGTGGCGAAGGCGTCTGACTAGAACGGATTCGACGGTAAGGGACGAAACCCTGGGGCGCAAACCGGATTAGATACCCGGGTAGTCC |
| GCA_001871415.1_CG1-02-57-44_Ref_seq | | | | | | | | | | | | ACGGGGTGCAGCAGGCGCGAAACCTTTGCAATGCGTGAAAGCGTGACAAGGGGATTCCAAGTGCCTGCACATTGTGCAGGCTTTTGCCAAGAGTAAAGACCTTGGCGAATAAGTGGTGGGCAAGACCGGTGCCAGCCGCCGCGGTAACACCGGCGCCACGAGTGGGGGCCATGATTATTGGGCCTAAAGCGTCCGTAGCCGGCCTGGTAAATCTCTTGTGAAATCGTACGGCTCAACCGAACGGCGAGCAGGGGACACTGCCAGGCTAGGGACCGGGAGGAGGCAGAGGTATTCGTTGGGGAGCGGTAAAATGCTATAATCCTTCGAAGACCACCTGTGGCGAAGGCGTCTGCCCAGAACGGATCCGACGGTGAGGGACGAAAGCCAGGGGAGCGATCCGGATTAGATACCCGAGTAGTCC |
| GCA_002254415.1_EX4484-52_Ref_seq | | | | | | | | | | | | ACGGGGCGCAGCAGGCGCGAAACCTTCACAATGCACTAATTGTGTGATGGGGAAACTCCAAGTGGTAGCATAGTGCTGCCTTTTGAAAAGTCAAAACATCTTTTTGAATAAGTGGTGGGTAAGACGGTTGCCAGCCGCCGCGGTAATAACCGCGCCACAAGTGGTAATCTCGATTATTGGGTCTAAAGCATCCGTAGCCGGGGATATTAGTTTTCTGTGAAATCCAAATGAAAACATTTGTAATGCAGAAAATACTATATTTCTTGAGACTGGGAGGAGCAAGAGGAACTCCATGGGTAGGGGTAGAATCTTTTAATCCATGGGGGGTCACCAGTGGCGAAGGCGTCTTGCTAGAACAGGTCTGACGGTGAGGGATGAAAGCTGGGGGAGCGAAGAGGATTAGATACCCTTGTAGTCC |
| GCA_002254565.1_EX4484-2_Ref_seq | | | | | | | | | | | | ACGGGGTGCAGCAGGCGCGAAACCTCTGCAATGCACGAAAGTGTGACAGGGGGACCCCAAGTGTCCAAGTACTGCTTGGACTTTTACGGAGTGTAAAAAGCTCCGGGAATAAGGGCTGGGTAAGACCGGTGCCAGCCTCCGCGGTAAAACCGGCAGCTCTAGTGGTAGTCGCTTTTATTGGGCTTAAAGCATCCGTAGCCTGTTTGGAAAGTCCTCTGTGAAATCCTACCGCTTAACGGTAAGACTGGCAAAGGATACTTCCAGGCTTGGGGCCGGAAGAAGCTAGGGGCATTTCTGGGGTAGCGGTGAAATGTTATAATCCCAGAAAGGTCACCAATGGCGAAGGCACCTAGCTAGGACGGGCCCGACGGTGAGGGATGTAAGCTAGGGGAGCGAACGGGATTAGATACCCCGGTAGTCC |
| GCA_002496355.1_SM1-50_Ref_seq | | | | | | | | | | | | ACGGGGCGCAGCAGGCGCGAAAACTTCGCAATGTGCGAAAGCACGACGAGGGAATCCCAAGTGCTGGCTCGTAAGAGTCAGCTGTTTCTCTGTCTAAAAAACAGAGAGAGTAAGAGCTGGGTAAGACGGGTGCCAGCCGCCGCGGTAATACCTGCAGCTCAAGTGGTGGCCGCTATTATTGGGCCTAAAACGTCCGTAGCCGGTTCTATAAATGCCTGGGTAAATCGTACGGCTTAACCGTACGAATTCCGGGTAGACTGTTCAACTTGGGACCGGGAGAGGCTAGAGGTACTCCTGGGGTAGAGGTGAAATTCTGTAATCCTAGGGGGACCACCAGTGGCGAAGGCGTCTAGCTAGAACGGGTCTGACGGTGAGGGACGAAGCCCTGGGGAGCAAACGGGATTAGATACCCCGGTAGTCC |
| GCA_002496385.1_UBA147_Ref_seq | | | | | | | | | | | | ACGGGGCGCAGCAGTCGCGAAAACTTGTCAATGGGGGAAACCCCGAACAGGGGACGCCAAGTGCATGCGTTTTACGTATGCTGTTCAGTTGTCTAAAAAACAGCCGAAGTAAGGGCTGGGTAAGACAGGTGCCAGCCGCCGCGGTAATACCTGCAGCCCAAGTGGTAATCGTTATTATTGAGCCTAAAGCGTCCGTAGCCGGTTTGGTAAATCCTCGGGTAAATCGAGCAGCTTAACTGTTCGAATTCCGGGGGGACTGCCAGACTTGGGACCGGGAGAGGTCAGAGGTACTCCAGGGGTAGGGGCTAAATCCTGTAATCCTTGGAGGACCACCAGTGGCGAAGGCGTCTGACTAGAACGGATCCGACGGTAAGGGACGAAACCCTGGGGCGCAAACCGGATTAGATACCCGGGTAGTCC |
| GCA_002506365.1_UBA583_Ref_seq | | | | | | | | | | | | AAGGGGTGCAGCAGGCGCGAAAACTCTACAATGCACGCAAGTGTGATAGGGCAACCCCTCGTGCTTATCTAACGATAGGCTTTTATTATGTGTAATGTGCATAATGAATAAGTGGTGGGCAAGACCGGTGCCAGCAGCCGCGGTAATACCGGCGCCACGAGTGGAGTTCACTATTATTGGGTCTAAAGTGTTCGTAGCTGGTTTGTTAAGTCTCTTGTGAAATTTCTGTTGCTTAACTCAGAAATTGCAAGAGATACTGGCAAACTTGAGACCGGAAGAGGTTGCAAGTATTGTTTGGGTAGCGGTAAAATGCTATAATCCTTACAAGACTAACTAAGGCGAAGGCATGCAACTTGGACGGATCTGACAGTGAGGAACGAGAGCTGGGGGAGCGAACCGGATTAGATACCCGGGTAGTCC |
| GCA_002778455.1_1-14-0-10-45-29_Ref_seq | | | | | | | | | | | | ACGGGATGCAGCAGGCGCGGAACCTTTGCAATGCGCGAAAGCGTGACAGGGGAACTCCGAGTGGTGAGGGCAAATTGTCCTCATCTTTTAGCCAATTTAAAAAGTTGGCAGAATAAGATCTGGGTAAGACCAGTGCCAGCCGCCATGCGCGTGATTTTCGCACGCACGGCACGGTACGCGGTAATACTGGCAGAGCGAGTGGTACCCACGAATATTGGGTCTAAAGAGTCCGTAGCGGGTCGGTTAAGTCCACTGTGAAATCTGGCAGCTTAACTGTCAGGAGTGCAGTGGATACTGGTCGGCTTGGGAGCGGGGGAGGTCAGGAGTACTTGTGGGGTAGTGGTAAAATACTGTAATCCTACAAGGACTACCAGTGGCGAAGGCGCCTGACCAAAACGCGTCCGACCGTGAGGGACGAAGGCTAGGAGAACGAATCGGATTAGATACCCGAGTAGCCC |
| GCA_002900545.1_SM1-50_Ref_seq | | | | | | | | | | | | ACGGGGCGCAGCAGGCGCGAAATCTTCACAATGTGCGAAAGCACGATGAGGGAATCCCAAGTGCTGGCACGTAAGTGTCAGCTGTTTCTTAGTCTAAAAAACTTAGAGAGTAAGGGCTGGGTAAGACGGGTGCCAGCCGCCGCGGTAATACCTGCAGCTCAAGTGGTGGCCGCTATTATTGGGCCTAAAACGTCCGTAGCCTGTTTAGTAAATGCTTGGGTAAATCGTATAGCTTAACTATACGAATACCGAGTAGACTGCTAGACTTGGGACCGGGAGAGGCTAGAGGTACTCCTGGGGTAGAGGTGAAATTCTGTAATCCTAGGGGGACCACCAGTGGCGAAGGCGTCTAGCTAGAACGGGTCTGACGGTGAGGGACGAAGCCCTGGGGAGCAAACGGGATTAGATACCCCGGTAGTCC |
| GCA_002900545.1_SM1-50_Ref_seq | | | | | | | | | | | | ACGGGGCGCAGCAGGCGCGAAAACTTCGCACTGTGCGAAAGCACGACGAGGGAATCCCAAGTGCTGGCTCGTAAGAGTCAGCTGTTTCTTTGTCTAAAAAACAGAGAGAGTAAGAGCTGGGTAAGACGGGTGCCAGCCGCCGCGGTAATACCTGCAGCTCAAGTGGTGGCCGCTATTATTGGGCCTAAAACGTCCGTAGCCGGTTCATTAAATGCCTGGGTAAATCGTACGGCTTAACCGTACGAATTCCGGGTAGACTGATGGACTTGAGATCGGGAGAAGCTAGAGGTACTCCTGGGGTAGAGGTGAAATTCTGTAATCCTAGGGGGACCACCGGTGGCGAAGGCGTCTAGCTAGAACGAGTCTGACGGTGAGGGACGAAGCCCTGGGGCGCAAACGGGATTAGATACCCCGGTAGTCC |
| GCA_003135935.1_FEN-33_Ref_seq | | | | | | | | | | | | ACGGGGCGCAGCAGTCGCGAAAACTTCACAATGGGAGAAATCCCGATGAGGGAATTCCAAGTGCTAGCACTATGTGTTAGCTGTTCTTTTGTCTAAAAAACAGAAGAAGTAAGGGCCGGGTAAGACGGGTGCCAGCCGCCGCGGTAATACCCGCGGCCCGAGTGGTGGTCGATATTATTGAGCCTAAAACGTCCGTAGCCGGTCTTGTAAATCCTTGGGTAAATCGGCCAGCTTAACTGTCCGAAGTCCGGGGAGACTGCAAGACTTGGGATCGGGAGAGGTCAGAGGTACTTCTGGGGTAGGGGTAAAATCCTGTAATCCTGGAAGGACCACCGGTGGCGAAGGCGTCTGACTAGAACGAATTCGACGGTGAGGGACGAAGCCCTGGGGCGCAAACGGGATTAGATACCCCGGTAGTCC |
| GCA_003551675.1_Natronoplasma_Ref_seq | | | | | | | | | | | | ACGGGGTGCAGCAGTTGCGAAACCTTCACAATGTGGGAAACCACGATGAGGGAACTCCAAGTGCCAACACCCAGTGTTGGCTTTTCACCTGTGTAAAGAGCAGGTAGAATAAGGGCCGGGTAAGACGGGTGCCAGCCGCCGCGGTAACACCCGCGGCCCGAGTGGTGGTCATTTTTATTGAGCTTAAAGCGTTCGTAGCCGGTCTGGTAAATCTCTGGGTAAATCGGACGGCTTAACCGTCCGAATTCCAGGGAGACTGCTAGACTTGGGGCCGGGAAAGGCCAGAGGTACTCTAGGGGTAGAAGTAAAATTCTGTAATCCCTAGAGGACCACCAGTGGCGAAAGCGTCTGGCTTGAACGGACCCGACGGTGAGGGACGAAGCCCTGGGGCGCGAACCGGATTAGATACCCGGGTAGTCC |
| GCA_003561825.1_PWVO01_Ref_seq | | | | | | | | | | | | ACGGGGTGCAGCAGGCGCGAAACCTTTACAATGCACGAAAGTGTGATAAGGGGATCCTAAGTGCCTTGCTCACAGCAAGGCTTTTGCCAAGAGCAAAAATCTTGGAGAATAAGTGGTGGGAAAGACTGGTGCCAGCAGCCGCGGTAATACCAGCGCCACAAGTGGCAATCAGCTTTATTGGGCCTAAAGCGTTCGTAGCCGGTCTGATAAATTTTCTGTGAAATTGTCAGGCTTAACCTGACAGCGCGCAGGAAAGACTATCAGACTAGGAACCGGGAGGAGTCAGAGGTATTCTTGGGGGAGCGGTAAAATGCTATAATCCCGAGAGGACTAACAGTGGCGAAGGCGTCTGACTAGAACGGATTCGACGGTGATTGGACGAAAGCCAGGGGAGCAATCCGGATTAGATACCCGAGTAGTCC |
| GCA_003694805.1_J110_Ref_seq | | | | | | | | | | | | ACGGGGTGCAGCAGGCGCGAAACCTTTCCAATGCACGAAAGTGTGAGAAGGGGATCCCCAGTGCTTGCACATTGTGCAGGCTTTTGCCAAGTGTAAATAGCTTGGCGAATAAGTGGTGGGTAAGACTGGTGCCAGCCGCCGCGGTAACACCAGCGCCACAAGTGGCTATCGCGTTTATTGGGCCTAAAGCGTTCGTAGCCGGCTTGGTAAATCTTTTGTGAAATCGTCTTGCTCAACTTGACGGCGCGCAGAAGACACTGCCAGGCTTGAGACCGGGAGGGGTCAGACGTATTCCGGGGGGACCGGTAAAATGGGATAATCCTCGGAGGACGACCGATGGCGAAGGCATCTGACCAGAACGGATCTGACGGTGAGGAACGAAAGCTAGGGGAGCAAACCGGATTAGATACCCGAGTAGTCC |
| GCA_007117405.1_UBA583_Ref_seq | | | | | | | | | | | | AAGGGGTGCAGCAGGCGCGAAAACTCTACAATGCACGCAAGTGTGATAGGGTGACCTCTCGTGCTTATCGAAAGATAGGCTTTTGTCAAGCTTAGTGCGCTTGACGAATAAGTGGTGGGCAAGACCGGTGCCAGCAGCCGCGGTAATACCGGCGCCACAAGTGGAGATCGCGATTATTGGGTCTAAAGTGTTCGTAGCTGGTTTGTTAAGTCTCTTGTGAAAGTTCTATGGCCTAACCTAGAAATTGCAAGAGATACTGGCAATCTTGAGACCGGAAGAGGTGTGAGGTATTGTTTGGGTAGCGGTAAAATGTTATAATCCTTACAAGACCAACTAAGGCGAAGGCATCACACTTGGACGGATCTGACAGTGAGGAACGAGAGCTGGGGGAGCGAACCGGATTAGATACCCGGGTAGTCC |
| GCA_008974855.1_CSP1-1_Ref_seq | | | | | | | | | | | | ATGGGGCGCAGCAGGCGAGAAACCTTTGCAATGTGCGCAAGCACGACAAGGTTAATCCGAGTGTTTCCTGCTAAAGGAATCTTTTGTTAGTTGTAAAACCACTATCGAATAAGGGGTGGCAAGTTCTGGTGTCAGCCGCCGCGGTAAAACCAGCACCTCAAGTGGTCAGGAGGATTATTGGGCCTAAAGCATCCGTAGCCTGTTCTATAAGTTTCCGGTTAAATCCACTAGCTCAACTAATGGGCTGCCGAAAATACTGTAGAACTAGGGAGTGGGAGAGGTAGACGGTATTCGGTAGGAAGGGGTAAAATCCTTTGATCTATTGAGGACCACCTGTGGCGAAGGCGGTCTACCAGAACACGTTCGACGGTGAGGGATGAAAGCTGGGGGAGCAAACCGGATTAGATACCCGGGTAGTCC |
| GCA_013791785.1_UBA147_Ref_seq | | | | | | | | | | | | ACGGGGCGCAGCAGTCGCGAAAACTTGTCAATGAGGGAAACCTCGAACAGGGGACTCCAAGTGCATGCGTTTTACGTATGCTTTTTGGCTGCCTAAAAAGCAGTCAGAATAAGGGCTGGGTAAGACAGGTGCCAGCCGCCGCGGTAATACCTGCAGCCCAAGTGGTAATCGTTATTATTGAGCCTAAAGCGTCCGTAGCCGGTCCGGTAAACCCTCGGGTAAATCGAGCAGCTCAACTGTTCGAATTCCGGGGGGACTGCCAGACTTGGAACCGGGAGAGGTCAGAGGTACTCCAGGGGTAGGGGCTAAATCCTGTAATCCTTGGAGGACCACCAGTGGCGAAGGCGTCTGACTAGAACGGATTCGACGGTAAGGGACGAAACCCTGGGGCGCAAACCGGATTAGATACCCGGGTAGTCC |
| GCA_013821245.1_Nitrosocosmicus_Ref_seq | | | | | | | | | | | | ATGGGGCGCAGCAGGCGCGAAACCTCTGCAATAGGCGAAAGCTTGACAGGGTTACTCTGAGTGATTTCCGTTAAGGAGATCTTTTGGCACCTCTAAAAATGGTGCAGAATAAGGGGTGGCAAGTCTGGTGTCAGCCGCCGCGGTAATACCAGCACCCCGAGTGGTCGGGACGTTTATTGGGCCTAAAGCATCCGTAGCCGGTTCTACAAGTCTTCCGTTAAATCCACCTGCTTAACAGATGGGCTGCGGAAGATACTATGGATCTAGGAGGCGGGAGAGGCAAGCGGTACTCGATGGTAGGGGTAAAATCCGTTGATCCATTGAAGACCACCAGTGGCGAAGGCGGCTTGCCAGAACGCGCTCGACGGTGAGGGATGAAAGCTGGGGGAGCAAACCGGATTAGATACCCGGGTAGTCC |
| GCA_014730115.1_Bilamarchaeum_Ref_seq | | | | | | | | | | | | ACGGGGTGCAGCAGGCGCGAAACCTCTACAATGCGCGAAAGCGTGATAGGGGGAGTCTGAGTGGCGATGGCAAAGCTATCGTCTTTTGCCAAGTGTAAACAGCTTGGCGAATAAGGGTGTGGGCAATCACACCTTAAGGTATTACTGACAGTCTTTAGGGAGTGGCCTAAGACTGGTGGCAGCCGCCGCGGTAACACCAGCGCCTCGAGTGGTACCCACGGATATTGGGCCTAGAGCGTCCGTAGCCGGACGGTTAAGTCCGCTGTGAAATGTTGGGGCTCAACCCTAACGCTTGCAGTGGATACTGGTCGTCTCGGGAACGGGGGAGGATCGTAGTACTTCCGGGGTAGGGGTAAAATCCGTTGAGCCTGGAAGGACTACCAGTGGCGAAGGCGCCGATCCAAAACGTGTCCGACGGTGAGGGACGAAAGCTGGGGGAGCGAACGGGATTAGATACCCCGGTAGTCC |
| GCA_014894395.1_SM1-50_Ref_seq | | | | | | | | | | | | ACGGGGCGCAGCAGGCGCGAAAACTTCGCACTGTGCGAAAGCACGACGAGGGAATCCCAAGTGCTGGCTCGTAAGAGTCAGCTGTTTCTCTGTCTAAAAAACAGGGAGAGTAAGAGCTGGGTAAGACGGGTGCCAGCCGCCGCGGTAATACCTGCAGCTCAAGTGGTGGCCGCTATTATTGGGCCTAAAACGTCCGTAGCCGGTTCTATAAATGCCTGGGTAAATCGTACGGCTTAACCGTACGAATTCCGGGTAGACTGTTCAACTTGGGACCGGGAGAGGCTAGAGGTACTCCTGGGGTAGAGGTGAAATTCTGTAATCCTAGGGGGACCACCAGTGGCGAAGGCGTCTAGCTAGAACGGGTCTGACGGTGAGGGACGAAGCCCTGGGGAGCAAACGGGATTAGATACCCCGGTAGTCC |
| GCA_016203225.1_DSQK01_Ref_seq | | | | | | | | | | | | ACGGGGTGCAGCAGTCGGGAAAACTTCGCAATGCGCGAAAGCGTGACGAGGGGATTCAGAGTGCTCAGGGATTATCCCTGGGCTTTTGCCAAGGGCAAAAACCTTGGCGAATAAGTGGTGGGCAAGACTGGTGCCAGCCGCCGCGGTAACCCCAGCGCCACAAGTGGGAACCACTTTTATTGGGTCTAAAGCATCCGTAGCCGGCTTTGCAAATGCCTTGTGAAATGGTTGCGCTCAACGTAACCACGTGCAAGGTAGACTACAAAGCTAGGGACCAGAAGAGGTTGGCCGTATTCCTGGGGGAACGGTAAAATGTTATAATCCCAGGAGGACGACCGATGGCGAAGGCAGCCAACCAGTATGGATCCGACGGTGAGGGATGAAAGCTAGGGGAGCGATCCGGATTAGATACCCGAGTAGTCC |
| GCA_016212525.1_Bilamarchaeum_Ref_seq | | | | | | | | | | | | ACGGGATGCAGCAGGCGCGAAACCTCTGCAATGCGCGAAAGCGTGACAGGGAAAGTCTGAGTGGCGGAAACTTAGTTTCCGTCTTTTGCCAAGTGTAAATAGCTTGGAGAATAAGGGTGTGGGCAAGACTGGTGGCAGCCGCCGCGGTAACACCAGCGCCTCGAGTGGTATCCACGAATATTGGGCCTAAAGTGTCCGTAGCCGGACTATTAAGTCCGCTGTGAAATGTTGGGGCTCAACCCTAACGCGTGCAGTGGATACTGATATGTCTTGGGAATGGGGGAGGGTCGGAGTACTTCCAGGGTAGGGGTAAAATCCGTTGAGCCTGGAAGGACTACCAGTGGCGAAGGCGCCGATCCAAAACATGTCCGACGGTGAGGGACGAAAGCTGGGGGAGCGAACGGGATTAGATACCCCGGTAGTCC |

| Phylum_Genus_ASV_number | Presence (1) or Absence (0) in each layer | | | | | | | | | | | |
|---|---|---|---|---|---|---|---|---|---|---|---|---|
| | L1 | L2 | L3 | L4 | L5 | L6 | L7 | L8 | L9 | L10 | L11 | Sequence |
| GCA_016926175.1_Bilamarchaeum_Ref_seq | | | | | | | | | | | | ACGGGGTGCAGCAGGCGCGAAACCTCTACAATGCGCGAAAGCGTGATAGGGGAAGTCTGAGTGGCAGCAGCATTGCTGTTGTCTTTTACCAAGTGTACACAGCTTGGAGAATAAGGGTGTGGGCAAACACATCCTTTATTGGTTGAACAGTCAATAGGGAGTGGCCTATGACTGGTGGCAGCCGCCGCGGTAACACCAGCGCCTCGAGTGGTATCCACGAATACTGGGCCTAAAGCGTCCGTAGCTGGATGGGTAAGTCCACTGTGAAATGTTGGAGCTCAACTCTAACGCTTGCAGTTGATACTGCCTGTCTAGGGAACGGGGGAGGGTCGGAGTACATTCAGGGTAGGGGTAAAATCCGTTGAGCCTGGATGGACTACCAGTGGCGAAGGCGCCGACCCAAAACGTGTCCGACAGTGAGGGACGAAGGCTAGGGGAGCGAACGGGATTAGATACCCCGGTAGTCC |
| GCA_016930415.1_SM23-78_Ref_seq | | | | | | | | | | | | ACGGGGTGCAGCAGGCGCGAAACCTTTTCAATGCGCGAAAGCGTGAGAAGGGAATTCCAAGTGCTCAAGTTTTACTTGGGCTTTTGCCAAGTGTAAAAAGCTTGGCGAATAAGGGGTGGGTAAGACTGGTGCCAGCCGCCGCGGTAACACCAGCGCCTCGAGTGGGGATCGCGATTATTGGGTCTAAAGCGTCCGTAGCAGGCCTTGTAAATGCTTTGTGAAATTGTCCGGCATAACCGGGCAGCGTGCAGAGTAGACTGCAGGGCTCGAGACCGGGAGAGGTTAGAAGTATTCCTGGGGTACCGGTAAAATGGGTTAATCCTGGGAGGACTACCTATGGCGAAGGCATCTGACCAGAACGGTTCTGACTGTGAGGGACGAAAGCTAGGGGAGCAAAACGGATTAGATACCCGAGTAGTCC |
| GCA_018334755.1_Natronoplasma_Ref_seq | | | | | | | | | | | | ACGGGGTGCAGCAGTTGCGAAACCTTCACAATGTGGGAAACCACGATGAGGGGACTCCAAGTGCCAGCACCTAGTGCTGGCTTTTCACCTGTCTAAAGTGCAGGTAGAATAAGGGCCGGGTAAGACGGGTGCCAGCCGCCGCGGTAACACCCGCGGCCCGAGTGGTGGCCATTTTTATTGAGCTTAAAGCGTCCGTAGCCGGTCTGGTAAATCTCTGGGTAAATCGGACGGCTCAACCGTTCGAATTCCAGAGAAACTGCTAGACTTGGGGCCGGGAGAGGTCAGAGGTACTCCTGGGGTAGAAGTAAAATTCTGTAATCCTAGGAGGACCACCAGTGGCGAAAGCGTCTGGCTTGAACGGACCCGACGGTGAGGGACGAAGCCCTGGGGCGCGAACCGGATTAGATACCCGGGTAGTCC |
| GCA_018334835.1_Natronoplasma_Ref_seq | | | | | | | | | | | | ACGGAGTGCAGCAGTTGCGAAACCTTCACAATGTGGGAAACCACGATGAGGGAACTCCAAGTGCCAACACTTAGTGTTGGCTTTTCACCTGTGTAAAGAGCAGGTGGAATAAGGGCCGGGTAAGACGGGTGCCAGCCGCCGCGGTAACACCCGCGGCCCGAGTGGTGACCATTTTTATTGAGCTTAAAGCGTCCGTAGCCGGTCTGGTAAATCTCTGGGTAAATCGTACGGCTTAACCGTTCGAATTCCAGGGAAACTGCCAGACTTGGGGCCGGGAGAAGCTAGAGGTACTCCAGGGGAAGAAGTAAAATTCTGTAATCCTTGGAGGACCACCAGTGGCGAAAGCGTCTAGCTAGAACGGACCCGACGGTGAGGGACGAAGCCCTGGGGCGCGAACCGGATTAGATACCCGGGTAGTCC |
| GCA_018812625.1_Bilamarchaeum_Ref_seq | | | | | | | | | | | | ACGGGGTGCAGCAGGCGCGAAACCTCTACAATGCGCGAAAGCGTGATAGGGGAAGTCTGAGTGGCGATAGCAATGCTATCGTCTTTTACCAAGTCTAAATCGCTTGGAGAATAAGGGTGTGGGCAAACACACCTTTATAGCTAGAACAGTCTATAGGGAGTGGCCTATGACTGGTGGCAGCCGCCGCGGTAACACCAGCGCCTCGAGTGATATCCACGAATACTGGGCCTAAAGCGTCCGTAGCCGGATAAGTAAGTCCACTGTGAAATGTTGGAGCTCAACTCCATCGCTTGCAGTAGATACTGCTAATCTCGGGAACGGGGGAGGGTCGGAGTACGTCCAGGGTAGGGGTAAAATCTGTTGAGCCTGGACGGACTACCAGTGGCGAAGGCGCCGACCCAAAACGTGTTCGACGGTGAGGGACGAAAGCTAGGGGAGCGAACGGGATTAGATACCCCGGTAGTCC |
| GCA_018812715.1_Bilamarchaeum_Ref_seq | | | | | | | | | | | | ACGGGGTGCAGCAGGCGCGAAACCTTTACAATGCACGAAAGTGTGATAAGGGGAGTCTGAGTGGCGATAGCATAGCTATCGTCTTTTGCCAAGTGTAAATAGCTTGGAGAATAAGGGTGTGGGCAATCACACCCTGAACGAACAGTCTTTAGGGAGTGGCCTAAGACTGGTGGCAGCCGCCGCGGTAACACCAGCGCCTCGAGTGGTACCCACGGATATTGGGCCTAGAGCGTCCGTAGCCGGACGATTAAGTCCGCTGTGAAATGTTGGGGCTCAACCCTAACGCTTGCAGTGGATACTGGTTGTCTTGGGAACGGGGGAGGATCGGAGTACTTCCGGGGTAGGGGTAAAAACCCTTGAATGCGTTCACGATTTGCAATCAAGGGATTGCAATCCGTTGAGCCTGGAAGGACTACCAGTGGCGAAGGCGCCGATCCAAAACGTGTCCGACGGTGAGGGACGAAAGCTGGGGGAGCGAACGGATTAGATACCCCGGTAGTCC |
| GCA_018813505.1_Bilamarchaeum_Ref_seq | | | | | | | | | | | | ACGGGATGCAGCAGGCGCGAAACCTCTGCAATGCGCGAAAGCGTGACAGGGGGAATCTGAGTGGCGATAGCATAGCTATCGTCTTTTGCCAAGTGTAAATAACTTGGAGAATAAGGGTGTGGGCAAACACATCTTTAGACAATAAGGAATGGTGTTTTTACAGTCTTAGGAAGTGGCCTATGACTGGTGGCAGCCGCCGCGGTAACACCAGCGCCTCAAGTGGTACCCACGAATATTGGGCCTAAAGCGTCCGTAGCCGGACTGTTAAGTCCGCTGTGAAATGTTGGGGCTCAACCCTAACGCGTGCAGTGGATACTGACAAGTCTTGGGAACGGGGGAGGGTCGGAGTACTCTCGGGGTAGGGGTAAAATAGTTTTCAGTCTCGTGTTTCGGGTTCTGAAAATATTATATTCCTTTGATCCTGGGAGGACTACCAGTGGCGAAGGCGCCGACCCAAAACGTGTCCGACGGTGAGGGACGAAAGCTGGGGGAGCGAACGGGATTAGATACCCCGGTAGTCC |
| GCA_019322375.1_SM23-78_Ref_seq | | | | | | | | | | | | ACGGGGTGCAGCAGGCGCGAAACCTTTACAATGCACGCAAGTGTGATAAGGGGATCCCAAGTGCTTATCATTGATAGGCTTTTGCCAAGTGTAAAAAGCTTGGCGAATAAGGGGTGGGTAAGACTGGTGCCAGCCGCCGCGGTAACACCAGCGCCCCGAGTGGCGATCGCGATTATTGGGCCTAAAGCGTCCGTAGCAGGTCTTGTAAATCCTTTGTGAAATCGGCGAGCTTAACTTGTCGGCGTGCAGAGGACACTGCAGGACTCGAGACCGGGAGAAGTCAGAAGTATTTCCTGGGTACCGGTAAAATGGGTTAATCCAGGGAGGACTACCAATGGCGAAGGCATCTGGCCAGAACGGATTTGACTGTGAGGGACGAAAGCTAGGGGAGCCAAACGGATTAGATACCCGAGTAGTCC |
| GCA_022565615.1_UBA583_Ref_seq | | | | | | | | | | | | AAGGGGTGCAGCAGGCGCGAAACCTTTACAATGCACGAAAGTGTGATAAGGCAACTCCTTGTGCTTATCCAAAAGATAGGCTTTTTTGATACGTAATGTGTATTAAGAATAAGTGGTGGGCAAGACCGGTGCCAGCAGCCGCGGTAATACCGGCGCCACAAGTGGAGATCATGATTACTGGGTCTAAAGTGTTCGTAGCTGGTTATGTAAGTTTCTTGTTAAATATCTATAGCTTAACTTAGATCTGCAAGAAATACTGCATGACTTGAGACCGGAAGGGGTGTGAAGTATGAATTGGGTAGCGGTAAAATGTTATAATCCTTTTCAGACTAACTAAGGCGAAGGCATCACACTAAGACGGATCTGACAGTGAGGAACGAAAGCTGGGGGAGCGAACCGGATTAGATACCCGGGTAGTCC |
| GCA_022573065.1_CSP1-1_Ref_seq | | | | | | | | | | | | ATGGGGCGCAGCAGGCGAGAAACCTTTGCAATGTGCGCAAGCACGACAAGGTTAATCCGAGTGTTTCCTGCTAAAGGAATCTTTTGTTAGTCGTAAAACCACTATCGAATAAGGGGTGGGCAAGTTCTGGTGTCAGCCGCCGCGGTAAAACCAGCACCTCAAGTGGTCAGGAGGATTATTGGGCCTAAAGCATCCGTAGCCTGTTCTATAAGTTTCCGGTTAAATCCATTAGCTCAACTAATGGGCTGCCGAAAATACTGTAGAACTAGGGAGTGGGAGAGGTAGACGGTATTCGGTAGGAAGGGGTAAAATCCTTTGATCTATTGAGGACCACCTGTGGCGAAGGCGGTCTACCAGAACACGTTCGACGGTGAGGGATGAAAGCTGGGGGAGCAAACCGGATTAGATACCCGGGTAGTCC |
| GCA_023137195.1_EX4484-52_Ref_seq | | | | | | | | | | | | ACGGGGTGCAGCAGGCGCGAAACCTTCACAATGCACAATATGTGTGATGGGGAGATTCTAAGTGGCAGCATAACGCTGTCTTTTGAAAAGTTGAAACATCTTTTCGAATAAGTGGTGGGTAAGACGGTTGCCAGCCGCCGCGGTAATAACCGCGCCACAAGTGGTAATCTCGATTATTGGGTCTAAAGCATCCGTAGCCGGTGATATAAGTTTCCTGTGAAATCCAAATGAAAACATTTGGGCATGCAGAAAATACTGTATCACTTGAGACTGGGAGGAGCAAGAGGTACTCGAGGGGAAGGGGTAAAATCTATTAATCCCTCGGGGGTCACCAGTGGCGAAGGCGTCTTGCTAGAACAGGTCTGACGGTGAGGGATGAAAGCCAGGGGAGCGAAGAGGATTAGATACCCTTGTAGTCC |
| GCA_023898345.1_UBA583_Ref_seq | | | | | | | | | | | | ATGGGGTGCAGCAGGCGCGAAACCTCTACAATGCACGCAAGTGTGATAGGGTAACCCCTCGTGCTTATCTTATAGATAGGCTTTTGTCAAGTGTAGTGTACTTGACGAATAAGTGGTGGGCAAGACCGGTGCCAGCAGCCGCGGTAATACCGGCGCCACAAGTGGAGATCGCTATTATTGGGTCTAAAGTGTTCGTAGCTGGTTTGTTAAGTCTCTTGTGAAAGCCCTGCTCTTAATGTAGGAATTGCAAGAGATACTGATAAACTTGGGACCGGGAGAGGTGTGAAGTATTGTTTGGGTAGCGGTAAAATGCTATAATCCTTACAAGACTAACTAAGGCGAAGGCATCACACTAGAACGGATCTGACAGTGAGGAACGAGAGCTGGGGTAGCGAACCGGATTAGATACCCGGGTAGTCC |
| GCA_025457885.1_SM1-50_Ref_seq | | | | | | | | | | | | ACGGGGCGCAGCAGGCGCGAAAACTTCGCACTGTGCGAAAGCACGACGAGGGAATCCCAAGTGCTGGCTCGTAAGAGTCAGCTGTTTCTTTGTCTAAAAAACAGAGAGAGTAAGAGCTGGGTAAGACGGGTGCCAGCCGCCGCGGTAATACCTGCAGCTCAAGTGGTGGCCGCTATTATTGGGCCTAAAACGTCCGTAGCCGGTTCTATAAATGCCTGGGTAAATCGTACGGCTTAACCGTACGAATTCCGGGTAGACTGTTCAACTTGGGACCGGGAGAGGCTAGAGGTACTCCTGGGGTAGAGGTGAAATTCTGTAATCCTAGGGGGACCACCAGTGGCGAAGGCGTCTAGCTAGAACGGGTCTGACGGTGAGGGACGAAGCCCTGGGGAGCAAACGGGATTAGATACCCCGGTAGTCC |
| GCA_027858315.1_UBA583_Ref_seq | | | | | | | | | | | | AAGGAGTGCAGCAGGCGCGAAACCTCTACAATGCACGCAAGTGTGATAGGGCAACCCCTCGTGCTTACCGAAAGGTAGGCTTTTATTATGTGTAATGAGCATAACGAATAAGTGGTGGGCAAGACTGGTGCCAGCAGCCGCGGTAATACCAGCGCCACAAGTGGAGATCACTTTTATTGGGTCTAAAGTGTCCGTAGCTGGTTTGATAAGTCTCTTGTGAAATTTCTTTGGCTTAACCTAGAAACTGCAAGAGAAACTGTTTAACTTGAGACCGGGAGAGGTGTGAAGTATTGTTTGGGTAGCGGTAAAATGTTATAATCCTTACAGGACTAACTAATGCGAAGGCATCACACTAGAACGGATCTGACAGTGAGGGACGAGAGCTAGGGTAGCGAACCGGATTAGATACCCGGGTAGTCC |
| GCA_028703755.1_UBA147_Ref_seq | | | | | | | | | | | | ACGGGGCGCAGCAGTCGCGAAAACTTGTCAATGGGGGAAACCCCGAACAGGGAATGCCAAGTGCATGCGTTTTACGTATGCTGTTCAGTTGTCTAAAAAACAGCCGAAGTAAGGGCTGGGTAAGACAGGTGCCAGCCGCCGCGGTAATACCTGCAGCCCAAGTGGTAATCGTTATTATTGAGCCTAAAGCGTCCGTAGCCGGTTTGGTAAATCCTCGGGTAAATCGAGCAGCTCAACTGTTCGAATACCGGGGAGACTGCCAGACTTGGGACCGGGAGAGGTCAGAGGTACTTCAGGGGTAGGGGCTAAATCCTGTAATCCTTGAAGGACCACCAGTGGCGAAGGCGTCTGACTAGAACGGATCCGACGGTAAGGGACGAAACCCTGGGGCGCAAACCGGATTAGATACCCGGGTAGTCC |
| GCA_030632385.1_EX4484-52_Ref_seq | | | | | | | | | | | | ACGGGGTGCAGCAGGCGCGAAACCTTCACAATGCACAATATGTGTGATGGGGAGAATCTAAGTGGTAGCATAACGCTACCTTTTGAAAAGTGAAAACATCTTTTCGAATAAGTGGCGGGTAAGACGGTTGCCAGCCGCCGCGGTAATAACCGCGCCACGAGTGGTAATCTCGTTTATTTGGTCTAAAGCATTCGTAGCCGGCAATATAAGTTTTCTGTGAAATCTAAATGAAAACATTTAGGCGTGCAGAAAAGACTGTATTGCTTGAGACTGGGAGGAGCAAGAGGTACTCGAAGGGAAGGGGTAAAATCCGTTAATCCTTCGGGGGTCACCAGTGGCGAAGGCGTCTTGCTAGAACAGGTCTGACGGTGAGGGATGAAAGCCAGGGGAGCGAAGAGGATTAGATACCCTTGTAGTCC |
| GCA_030739585.1_GCA-2726865_Ref_seq | | | | | | | | | | | | ACGGGGCGCAGCAGGCGCGAAACCTCCACAATTCACGAAAGTGAGATGGGGTTATCTCGAGTGCCGCCTGATGAAGGCGGCTTTTCCACTGTATAACTAGCCGTGGGAATAAGAGGGGGGCAAGACTGGTGTCAGCCGCCGCGGTAACACCAGCCCCTCGAGTGGTGGGGATGATTATTTGGCCTAAAGCGTCCGTAGCCGGCTTTGTAAATCTCTCGTTAAATCCAGCGTCCTAAGCGTTGGGCTGCGAGGGTAACTGCAGAGCTAGAGGGTGGGAGAGGTCAGCGGTATTCCTGGGGTAGGGGCGAAATCCATTGATCCCAGGAGGACCACCAGTGGCGAAGGCTGCTGACTTGAACACGCCTGACGGTGAGGGACGAAAGCTGGGGTAGCGAATCGGATTAGATACCCGAGTAGTCC |
| GCA_035620615.1_CSP1-1_Ref_seq | | | | | | | | | | | | ATGGGGCGCAGCAGGCGAGAAACCTTTGCAATGTGCGCAAGCACGACAAGGTTAATCCGAGTGTTTCCTGCTAAAGGAATCTTTTGTTAGTTCTAAAACCACTGATGAATAAGGGGTGGGCAAGTTCTGGTGTCAGCCGCCGCGGTAAAACCAGCACCTCAAGTGGTCAGGAGGATTATTGGGCCTAAAGCATCCGTAGCCGGTTCTGTAAGTTCCCGGTTAAATCCACTAGCTCAACTAGTGGGCTGCCGGAAATACTGTAGAACTAGGGAGTGGGAGAGGTAGACGGTACTCGGTAGGAAGGGGTAAAATCCTTTGATCTATTGATGACCACCTGTGGCGAAGGCGGTCTACCAGAACACGTTCGACGGTGAGGGATGAAAGCTGGGGGAGCAAACCGGATTAGATACCCGGGTAGTCC |
| GCA_041488275.1_DSQK01_Ref_seq | | | | | | | | | | | | ACGGGGTGCAGCAGTCGGGAAAACTTCGCAATGCGCGAAAGCGTGACGAGGGGATTCAGAGTGCTTAGGAATTTATTTCCTAGGCTTTTGCCAAGGGCAAAAACCTTGGCGAATAAGTGGTGGGCAAGACTGGTGCCAGCCGCCGCGGTAACCCCAGCGCCACAAGTGGGAACCACTTTTATTGGGTCTAAAGCATCCGTAGCCGGCTCGGCAAATGCCTTGTGAAATGGTCGCGCTCAACGCGACCACGTGCAAGGTAGACTACCGAGCTAGGGACCAGGAGAGGTTGGCAGTATTCCCGGGGAGCGGTAAAATGTTATAATCCCGGGAGGACTACCGATGGCGAAGGCAGCCAACCAGCATGGATCCGACGGTGAGGGATGAAAGCTAGGGGAGCGATCCGGATTAGATACCCGAGTAGTCC |
| GCA_041598975.1_UBA583_Ref_seq | | | | | | | | | | | | AAGGGGTGCAGCAGGCGCGAAACCTCTACAATGCACGCAAGTGTGATAGGGAAACCCCTCGTGCTTACCGAGAGGTAGGCTTTTGTTATGTGTAATGAGCATATCGAATAAGTGATGGGCAAGACTGGTGCCAGCAGCCGCGGTAATACCAGCGTCACAAGTGGAGATCATTTTTATTGGGTCTAAAGAGTCCGTAGCTGGTTTAGTAAGTCTCTTGTGAAATTTCTTTGGCTTAACCAAGAAATTGCAAGAGATACTGCTTGACTTGAGACCGGGGGAGGTAAGAAGTATTGTTTGGGTAGCGGTAAAATGTTATAATCCTTACAAGACTAACTAAGGCGAAGGCATCTTACTAGAACGGATCTGACAGTGAGGGACGAAAGCTGGGGTAGCGAACCGGATTAGATACCCGGGTAGTCC |
| GCA_041631255.1_UBA583_Ref_seq | | | | | | | | | | | | AAGGGGTGCAGCAGGCGCGAAACCTTTACAATGCACGCAAGTGTGATAAGGCGACCCTTCGTGCTTATCGAAAGATAGGCTTTTGTTAAGCTTAGTGCGCTTAACGAATAAGTGGTGGGCAAGACTGGTGCCAGCAGCCGCGGTAATACCAGCGCCACAAGTGGAGATCACGATTACTGGGTCTAAAGTGTTCGTAGCTGGTCGAGTAAGTTTCTTGTGAAAGCCCTCTAGCTTAACTGAGAAATTGCAAGAAATACTGCTCGTCTTGAGACCGGAAGAGATAAGAGGTATTGTTTGGGTAGCGGTAAAATGTTATAATCCTTACAAGACCAACGAAGGCGAAGGCATCTTATTTGGACGGATCTGACAGTGAGGAACGAGAGCCAGGGGAGCGAACCGGATTAGATACCCGGGTAGTCC |
| GCA_041650255.1_1-14-0-10-45-29_Ref_seq | | | | | | | | | | | | ACGGGATGCAGCAGGCGCGAAACCTTTGCAATGCGCGAAAGCGTGACAGGGGAACTCCGAGTGGTGGAGGCATCATGTCTCCACCTTTTGGCCAATTTAAATAGTTGGCAGAATAAGATCTGGGTAAGACTCGTGCCAGCCGCCATGCTATGCAACGCATCAGCATTGGCACGGTACGCGGTAATACGAGCAGAGCAAGTGGTATCCACGAATATTGGGTCTAAAGAGTCCGTAGCGGGTCTGTTAAGTTCACTGTGAAATCTGGCGGCTTAACCGTCAGGCGTGCAGTGGATACTGGCAGGCTTGGGAGTGGGGGAGGCCGGGAGTACTTGCGGGGTAGTGGTAAAATACTGTAATCCTGCAAGGACTACCAGTGGCGAAGGCGCCCGGCCAAAACACGTCCGACCGTGAGGGACGAAGGCTAGGAGAACGAATCGGATTAGATACCCGAGTAGCCC |
| GCA_042217555.1_Nanohalobium_Ref_seq | | | | | | | | | | | | ACGGGGCGCAGCAGGCACGAAACCTTTACAATGCACTAACGTGCGATAAGGGGACTCCGAGTGCGGTCTTCCCACAAGGAAGGCCGCTTTTCTGCACTGTAAGGAGGTGCAGGAACAAGGGGTGGGCAACACTGCTGCCAGCCGCCGCGGTAATAGCAGGACCCCAAGTGGTAGTCGTTATTATTGGGCTTAAAGTGTTCGTAGCCTGCCATGTAAGTCCGCTGTGAAATCCACCTGCTCAACAGGTGGGCGTGCAGCGGATACTGCATTGGCTAGGGACCGGACGGCGTCGAGGCTATTCCCTGGGTAGGAGTGAAATCCTAAAATCCAGGGAGGAGCACCTGTGGCGAAAGCGCTCGGCGAGTACGGATCCGACGGTGAGGAACGAAAGCCAGGGGAGCAAAGCGGATTAGATACCCGCGTAGTCC |
| GCA_043802645.1_1-14-0-10-45-29_Ref_seq | | | | | | | | | | | | ACGGGATGCAGCAGGCGCGAAACCTTTGCAATGCGCGAAAGCGTGACAGGGGAACTCCGAGTGGCAGGGGGCTAATGCTCCCTGTCTTTTGGCCAATCTAAAAAGTTGGCAGAATAAGATCTGGGTAAGACCAGTGCCAGCCGCCGCGGTAATACTGGCAGAGCGAGTGGCATCCACGAATATTGGGGTCTAAAGAGTCCGTAGCGGGCTTGTTAAGTCTACTGTGAAATCCGGCTGCTTAACAGTCGGGCGTGCAGTGGATACTGGCAGGCTTGGGAGCGGGGGAGGTCAGGAGTACTTGCGGGGGAGTGGTAAAATACTGTAATCCTGCAAGGACTACCAGTGGCGAAGGCGCCTGACCAAAACGCGTCCGACCGTGAGGGACGAAGGCTAGGAGAACGAATCGGATTAGATACCCGAGTAGCCC |
| GCA_046485165.1_UBA147_Ref_seq | | | | | | | | | | | | ACGGGGCGCAGCAGTCGCGAAAACTTGTCAATGGGGGAAACCCCGAACAGGGGAGTCCAAGTGCATGCGTTTTACGTATGCTGTTCAGTTGTCTAAAACACAGCCGAAGTAAGGGCTGGGTAAGACAGGTGCCAGCCGCCGCGGTAATACCTGCAGCCCAAGTGGTAATCGTTATTATTGAGCCTAAAGCGTCCGTAGCCGGTTTGGTAAACCCTCGGGTAAATCGAGCAGCTCAACTGTTCGAATTCCGGGGGGACTGCCAGACTTGGGACCGGGAGAGGTCGGAGGTACTTCAGGGGTAGGGGCTAAATCCTGTAATCCTTGAAGGACCACCAGTGGCGAAGGCGTCCGACCAGAACGGATCCGACGGTAAGGGACGAAACCCTGGGGCGCAAACCGGATTAGATACCCGGGTAGTCC |

| | Presence (1) or Absence (0) in each layer | | | | | | | | | | | |
|---|---|---|---|---|---|---|---|---|---|---|---|---|
| Phylum_Genus_ASV_number | L1 | L2 | L3 | L4 | L5 | L6 | L7 | L8 | L9 | L10 | L11 | Sequence |
| GCA_046537845.1_Natronoplasma_Ref_seq | | | | | | | | | | | | ACGGGGTGCAGCAGTTGCGAAACCTTCACAATGTGGGAGACCACGATGAGGGGACTCCAAGTGCCAGCACCCAGTGTTGGCTTTTCACCTGTGTAACGAGCAGGTGGAATAAGGGCCGGGTAAGACGGGTGCCAGCCGCCGCGGTAACACCCGCGGCCCAAGTGGTGGTCATTATTATTGAGCTTAAAGCGTCCGTAGCCGGTCTGGTAAATCTCTGGGTAAATCGGACGGCTCAACCGTTCGAATACCAGGGAGACTGCTAGACTTGGGGCCGGGAGAGGTCAGAGGTACTCCTGGGGTAGAAGTAAAATTCTGTAATCCTAGGAGGACCACCAGTGGCGAAAGCGTCTGGCTAGAACGGACCCGACGGTGAGGGACGAAGCCCTGGGGCGCGAACCGGATTAGATACCCGGGTAGTCC |
| GCA_046540335.1_Natronoplasma_Ref_seq | | | | | | | | | | | | ACGGAGTGCAGCAGTTGCGAAACCTTCACAATGTGGGAAACCACGATGAGGGAACTCCAAGTGCCAGCACCCAGTGCTGGCTTTTCACCTGTGTAAAGAGCAGGTGGAATAAGGGCCGGGTAAGACGGGTGCCAGCCGCCGCGGTAACACCCGCGGCCCGAGTGGTGACCATTTTTATTGAGCTTAAAGCGTCCGTAGCCGGTCTGGTAAATCTCTGGGTAAATCGGACGGCTTAACCGTTCGAATTCCAGAGAAACTGCCAGACTTGGGGCCGGGAGAAGCTAGAGGTACTCCAGGGGTAGAAGTAAAATTCTGTAATCCTTGGAGGACCACCAGTGGCGAAAGCGTCTAGCTAGAACGGACCCGACGGTGAGGGACGAAGCCCTGGGGCGCGAACCGGATTAGATACCCGGGTAGTCT |
| GCA_047602745.1_UBA583_Ref_seq | | | | | | | | | | | | AAGGGGTGCAGCAGGCGCGAAACCTTTACAATGCACGAAAGTGTGATAAGGCAACTCTTTGTGCTTATCTAAAAGATAAGCTTTTTTGATATGTAATGTATATTAAGAATAAGTGGTGGGCAAGACCGGTGCCAGCAGCCGCGGTAATACCGGCGCCACAAGTGGAGATCATGTTTACTGGGTCTAAAGTGTTCGTAGCTGGTTATGTAAGTTTCTTGTTAAATATCTACAGCTTAACTTAGATTTGCAAGAAATACTGTATGACTTGAGACCGGAAGGGGTGTGAAGTATGAATTGGGTAGCGGTAAAATGTTTTAATCCTTTTCAGACTAACTAAGGCGAAAGCATCACACTAAGACGGATCTGACAGTGAGGAACGAAAGCTAGGGGAGCGAACCGGATTAGATACCCGGGTAGTCC |
| GCA_938004075.1_UBA147_Ref_seq | | | | | | | | | | | | ACGGGGCGCAGCAGTCGCGAAAACTTGTCAATGGGGGAAACCCCGAACAGGGAAGTCCAAGTGCATGCGTTTTACGTATGCTGTTCGGTTGTCTAAAACACAGCTGAAGTAAGGGCTGGGTAAGACAGGTGCCAGCCGCCGCGGTAATACCTGCAGCCCAAGTGGTAATCGTTATTATTGAGCCTAAAGCGTCCGTAGCCGGTTTGGTAAACCCTCGGGTAAATCGGGCAGCTTAACTGTCCGAATTCCGGGGGGACTGCCAGACTTGGGACCGGGAGAGGTCAGAGGTACTTCAGGGGTAGGGGCTAAATCCTGTAATCCTTGAAGGACCACCAGTGGCGAAGGCGTCTGACTAGAACGGATCCGACGGTAAGGGACGAAACCCTGGGGCGCAAACCGGATTAGATACCCGGGTAGTCC |
| GCA_963768775.1_FEN-33_Ref_seq | | | | | | | | | | | | ACGGGGCGCAGCAGTCGCGAAAACTTCACAATGGGGGAAACCCCGATGAGGGAATTCCAAGTGCTAGCACTATTGTGTTAGCTGTTCTTTTGTCTAAAAAACAGGAGAAGTAAGGGCCGGGTAAGACGGGTGCCAGCCGCCGCGGTAATACCCGCGGCCCGAGTGGTGGTCGATATTATTGAGCCTAAAACGTCCGTAGCCGGTCTTGTAAATCCTTGGGTAAATCGGCCAGCTTAACTGTCCGAAGTCCGGGGAGACTGCAAGACTTGGGATCGGGAGAGGTCAGAGGTACTTCTGGGGTAGGGGTAAAATCCTGTAATCCTAGAAGGACCACCGGTGGCGAAGGCGTCTGACTAGAACGAATTCGACGGTGAGGGACGAAGCCCTGGGGCGCAAACGGGATTAGATACCCCGGTAGTCC |
| GCA_964550175.1_DSQK01_Ref_seq | | | | | | | | | | | | ACGGGGTGCAGCAGTCGGGAAAACTTCGCAATGCACGAAAGTGTGACGAGGGGATTCAGAGTGCTTAGGATTATGTCCTAGGCTTTTGCCAAGAGCAAATATCTTGGCGAATAAGTGGTGGGCAAGACTGGTGCCAGCCGCCGCGGTAACCCCAGCGCCACAAGTGGGAACCACTTTTATTGGGTCTAAAGCATCCGTAGCCGGTTCGGCAAATGCCTTGTGAAATGGTTGCGCTCAACGTAACCACGTGCAAGGTAGACTACCAAACTAGGGACCAGAAGATGTTAGCAGTATTCCTGGGGAACGGTAAAATGTTATAATCCCAGGAGGACTACCGATGGCGAAGGCAGCTAACAAGTATGGATCCGACGGTGAGGGATGAAAGCTAGGGGAGCGATCCGGATTAGATACCCGAGTAGTCC |

**Supplementary Table 10.- Archaeal genus-level comparison, Poza Roja 2 (dominant genera) versus Archaean Domes (named examples only)**

| Poza Roja 2 genus | Relative abundance in Poza Roja 2 | Reported at Archaean Domes (named in text) |
|---|---|---|
| *SM1-50* | 26.3% | Not reported |
| *Halohasta* | 4.8% | Not reported |
| *Saliphagus* | 2.1% | Not reported |
| *Dethiobacter* | 1.6% (L7) | Not reported |
| *Halorussus* | 1.3% | Not reported |
| *Natrinema* | present | Not reported |
| *Halorubrum* | present | Not reported |
| *Halostagnicola* | present | Not reported |
| *Methanohalobium* | present | Not reported |
| – | – | *Natrialba (named example)* |
| – | – | *Haloferax (named example)* |
| – | – | *Haloterrigena (named example)* |
| – | – | *Methanohalophilus (named example, most abundant OTU)* |
| – | – | *Nanoarchaeum equitans (named example)* |
| – | – | *Thermococcus, Sulfolobus, Ignicoccus, Pyrolobus (named examples)* |
| – | – | *Nitrososphaera, Nitrosopumilus, Cenarchaeum (named examples)* |

Medina-Chávez et al. (2023) report 93 archaeal genera at Archaean Domes overall, but only the ~20 shown here were individually named in the published text; no genus-resolved supplementary table was available for a complete comparison. Absence from this table therefore does not necessarily mean absence from Archaean Domes, only that it was not named in the source publication. Taken together, these comparisons show that Poza Roja 2 shares the same core signature of halophilic and methanogenic Archaea documented at Archaean Domes, but with almost no genus-level overlap and a substantially broader phylum-level footprint, including several DPANN lineages absent from the published Archaean Domes record; notably, Poza Roja 2's single most abundant archaeal genus, the uncultured lineage SM1-50, is not represented among the Archaean Domes taxa in either study.

**Supplementary Table 11. Summary of morphological, physiological, ecological, and metabolic features of representative strains reported in the literature.**

| Strain | Morphology | Growth characteristics | Isolation_source | Metabolic_role | Reference | Paper |
|---|---|---|---|---|---|---|
| *Dethiobacter alkaliphilus (AHT 1)* | Motile spore-forming rods | T: 30 °C; pH: 9.5; Na+: 0.4 M. | Soda lakes, Mongolia | Respiratory anaerobe; facultative autotroph (u $H_2$); utilizes thiosulfate and polysulfide as electron acceptors | Sorokin et al., 2008 | Sorokin, D. Y., Tourova, T. P., Mußmann, M., & Muyzer, G. (2008). Dethiobacter alkaliphilus gen. nov. sp. nov., and Desulfurivibrio alkaliphilus gen. nov. sp. nov.: Two novel representatives of reductive sulfur cycle from soda lakes. Extremophiles, 12(3), 431–439. https://doi.org/10.1007/s00792-008-0148-8 |
| *Halohasta litchfieldiae (tADLT)* | Motile rods | Medium NOM; T: 30 °C; pH: 7.5; NaCl: 18%. | Deep Lake, Antarctica | Aerobic heterotrophic archaeon; uses organic acids (succinate, citrate) and sugars as carbon sources | Mou et al., 2012 | Mou, Y. Z., Qiu, X. X., Zhao, M. L., Cui, H. L., Oh, D., & Dyall-Smith, M. L. (2012). Halohasta litorea gen. nov. sp. nov., and Halohasta litchfieldiae sp. nov., isolated from the Daliang aquaculture farm, China and from Deep Lake, Antarctica, respectively. Extremophiles, 16(6), 895–901. https://doi.org/10.1007/s00792-012-0485-5 |
| *Halohasta litorea (R30T)* | Motile rods | Medium NOM; T: 40 °C; pH: 7.0; NaCl: 15%. | Daliang, China | Aerobic heterotrophic archaeon; utilizes pyruvate and various sugars; produces $H_2S$ from thiosulfate | Mou et al., 2012 | Mou, Y. Z., Qiu, X. X., Zhao, M. L., Cui, H. L., Oh, D., & Dyall-Smith, M. L. (2012). Halohasta litorea gen. nov. sp. nov., and Halohasta litchfieldiae sp. nov., isolated from the Daliang aquaculture farm, China and from Deep Lake, Antarctica, respectively. Extremophiles, 16(6), 895–901. https://doi.org/10.1007/s00792-012-0485-5 |
| *Halorussus amylolyticus (YC93T)* | Motile pleomorphic rods | Medium NHM; T: 37 °C; pH: 7.0; NaCl: 2.0 M. | Yuncheng Salt Lake, China | Aerobic heterotrophic archaeon; notable for its amylolytic, proteolytic, and lipolytic capabilities | Yuan et al., 2015 | Yuan, P. P., Ye, W. T., Pan, J. X., Han, D., Zhang, W. J., & Cui, H. L. (2015). Halorussus amylolyticus sp. Nov., isolated from an inland Salt Lake. International Journal of Systematic and Evolutionary Microbiology, 65(10), 3734–3738. https://doi.org/10.1099/ijsem.0.000487 |
| *Halorussus aquaticus (XZYJ18T)* | Motile rods | Medium NHM; T: 37 °C; pH: 7.0; NaCl: 2.6 M. | Mangkang Saltworks, Tibet | Aerobic heterotrophic archaeon; versatile metabolism; capable of partial denitrification | Zheng et al., 2022 | Zheng, X. W., Wu, Z. P., Sun, Y. P., Wang, B. B., Hou, J., & Cui, H. L. (2022). Halorussus vallis sp. nov., Halorussus aquaticus sp. nov., Halorussus gelatinilyticus sp. nov., Halorussus limi sp. nov., Halorussus salilacus sp. nov., Halorussus salinisoli sp. nov.: six extremely halophilic archaea isolated from solar saltern, salt lake and saline soil. Extremophiles, 26(3). https://doi.org/10.1007/s00792-022-01280-1 |
| *Halorussus caseinilyticus (DT72T)* | Motile rods | Medium NHM; T: 37 °C; pH: 6.5; NaCl: 3.1 M. | Dongtai, China | Aerobic heterotrophic archaeon; hydrolyzes casein and Tween 80; possesses enzymes for trehalose biosynthesis | Hu et al., 2024 | Hu, Y., Ma, X., Tan, S., Li, X. X., Cheng, M., Hou, J., & Cui, H. L. (2024). Genome-based classification of genera Halosegnis and Salella, and description of four novel halophilic archaea isolated from a tidal flat. Antonie van Leeuwenhoek, International Journal of General and Molecular Microbiology, 117(1). https://doi.org/10.1007/s10482-024-01952-2 |
| *Halorussus gelatinilyticus (XZYJT40T)* | Motile pleomorphic rods | Medium NHM; T: 42 °C; pH: 7.0; NaCl: 4.3 M. | Mangkang Saltworks, Tibet | Aerobic heterotrophic archaeon; specialized in gelatin hydrolysis; reduces nitrate | Zheng et al., 2022 | Zheng, X. W., Wu, Z. P., Sun, Y. P., Wang, B. B., Hou, J., & Cui, H. L. (2022). Halorussus vallis sp. nov., Halorussus aquaticus sp. nov., Halorussus gelatinilyticus sp. nov., Halorussus limi sp. nov., Halorussus salilacus sp. nov., Halorussus salinisoli sp. nov.: six extremely halophilic archaea isolated from solar saltern, salt lake and saline soil. Extremophiles, 26(3). https://doi.org/10.1007/s00792-022-01280-1 |
| *Halorussus halobius (HD8-83T)* | Motile pleomorphic rods | Medium NHM; T: 35 °C; pH: 7.5; NaCl: 4.3 M. | Laminaria algae, China | Aerobic heterotrophic archaeon; possesses complete glycolytic and Krebs cycle pathways; reduces nitrate | Han et al., 2022 | Han, D., & Cui, H. L. (2022). Halorussus halobius sp. nov., Halorussus marinus sp. nov. and Halorussus pelagicus sp. nov., isolated from salted brown alga Laminaria. International Journal of Systematic and Evolutionary Microbiology, 72(4). https://doi.org/10.1099/ijsem.0.005313 |
| *Halorussus halophilus (ZS-3T)* | Motile pleomorphic rods | Medium NHM; T: 37 °C; pH: 7.0; NaCl: 2.6 M. | Zhoushan Saltworks, China | Aerobic heterotrophic archaeon; highly proteolytic and amylolytic (hydrolyzes casein, gelatin, and starch) | Ding et al., 2020 | Ding, Y., Han, D., & Cui, H. L. (2020). Halorussus halophilus sp. nov., A Novel Halophilic Archaeon Isolated from a Marine Solar Saltern. Current Microbiology, 77(7), 1321–1327. https://doi.org/10.1007/s00284-020-01921-8 |
| *Halorussus limi (XZYJT49T)* | Motile rods | Medium NHM; T: 37 °C; pH: 7.5; NaCl: 3.9 M. | Mangkang Saltworks, Tibet | Aerobic heterotrophic archaeon; hydrolyzes starch; possesses genes encoding lipases and amylases | Zheng et al., 2022 | Zheng, X. W., Wu, Z. P., Sun, Y. P., Wang, B. B., Hou, J., & Cui, H. L. (2022). Halorussus vallis sp. nov., Halorussus aquaticus sp. nov., Halorussus gelatinilyticus sp. nov., Halorussus limi sp. nov., Halorussus salilacus sp. nov., Halorussus salinisoli sp. nov.: six extremely halophilic archaea isolated from solar saltern, salt lake and saline soil. Extremophiles, 26(3). https://doi.org/10.1007/s00792-022-01280-1 |

| Strain | Morphology | Growth characteristics | Isolation_source | Metabolic_role | Reference | Paper |
|---|---|---|---|---|---|---|
| *Halorussus lipolyticus (DT80T)* | Motile rods | Medium NHM; T: 40 °C; pH: 6.5; NaCl: 2.6 M. | Dongtai, China | Aerobic heterotrophic archaeon; lipolytic (hydrolyzes Tween 80); possesses assimilatory nitrate reductase | Hu et al., 2024 | Hu, Y., Ma, X., Tan, S., Li, X. X., Cheng, M., Hou, J., & Cui, H. L. (2024). Genome-based classification of genera Halosegnis and Salella, and description of four novel halophilic archaea isolated from a tidal flat. Antonie van Leeuwenhoek, International Journal of General and Molecular Microbiology, 117(1). https://doi.org/10.1007/s10482-024-01952-2 |
| *Halorussus litoreus (HD8-51T)* | Motile pleomorphic coccoid cells | Medium NHM; T: 30 °C; pH: 7.0; NaCl: 3.1 M. | Salted Laminaria algae | Aerobic heterotrophic archaeon; capable of anaerobic growth using nitrate or DMSO as electron acceptors | Han et al., 2019 | Han, D., Zhu, L., & Cui, H. L. (2019). Halorussus litoreus sp. Nov., isolated from the salted brown alga laminaria. International Journal of Systematic and Evolutionary Microbiology, 69(3), 767–772. https://doi.org/10.1099/ijsem.0.003233 |
| *Halorussus marinus (LYG-36T)* | Motile pleomorphic rods | Medium NHM; T: 30–40 °C; pH: 7.0–7.5. | Laminaria algae, China | Aerobic heterotrophic archaeon; versatile carbohydrate metabolism; grows anaerobically with DMSO or nitrate | Han et al., 2022 | Han, D., & Cui, H. L. (2022). Halorussus halobius sp. nov., Halorussus marinus sp. nov. and Halorussus pelagicus sp. nov., isolated from salted brown alga Laminaria. International Journal of Systematic and Evolutionary Microbiology, 72(4). https://doi.org/10.1099/ijsem.0.005313 |
| *Halorussus pelagicus (RC-68T)* | Motile pleomorphic rods | Medium NHM; T: 35 °C; pH: 7.0; NaCl: 2.6 M. | Laminaria algae, China | Aerobic heterotrophic archaeon; notable for its ability to hydrolyze gelatin and casein | Han et al., 2022 | Han, D., & Cui, H. L. (2022). Halorussus halobius sp. nov., Halorussus marinus sp. nov. and Halorussus pelagicus sp. nov., isolated from salted brown alga Laminaria. International Journal of Systematic and Evolutionary Microbiology, 72(4). https://doi.org/10.1099/ijsem.0.005313 |
| *Halorussus rarus (TBN4T)* | Motile rods | Medium NOM; T: 37 °C; pH: 7.0; NaCl: 2.1 M. | Taibei Saltworks, China | Aerobic heterotrophic archaeon; utilizes a wide variety of sugars and organic acids; reduces nitrate to nitrite | Cui et al., 2010 | Cui, H. L., Gao, X., Yang, X., & Xu, X. W. (2010). Halorussus rarus gen. nov., sp. nov., a new member of the family Halobacteriaceae isolated from a marine solar saltern. Extremophiles, 14(6), 493–499. https://doi.org/10.1007/s00792-010-0329-0 |
| *Halorussus ruber (YC25T)* | Motile pleomorphic rods | Medium NHM; T: 37 °C; pH: 6.5; NaCl: 1.7 M. | Yuncheng Salt Lake, China | Aerobic heterotrophic archaeon; grows anaerobically with DMSO, nitrate, and arginine; reduces nitrate | Xu et al., 2015 | Xu, W. D., Zhang, W. J., Han, D., Cui, H. L., & Yang, K. (2015). Halorussus ruber sp. nov. isolated from an inland salt lake of China. Archives of Microbiology, 197(1), 91–95. https://doi.org/10.1007/s00203-014-1058-z |
| *Halorussus salilacus (YCN54T)* | Motile rods | Medium NHM; T: 35 °C; pH: 7.0; NaCl: 2.6 M. | Yuncheng Salt Lake, China | Aerobic heterotrophic archaeon; amylolytic; possesses PHA synthase (PhaC) | Zheng et al., 2022 | Zheng, X. W., Wu, Z. P., Sun, Y. P., Wang, B. B., Hou, J., & Cui, H. L. (2022). Halorussus vallis sp. nov., Halorussus aquaticus sp. nov., Halorussus gelatinilyticus sp. nov., Halorussus limi sp. nov., Halorussus salilacus sp. nov., Halorussus salinisoli sp. nov.: six extremely halophilic archaea isolated from solar saltern, salt lake and saline soil. Extremophiles, 26(3). https://doi.org/10.1007/s00792-022-01280-1 |
| *Halorussus salinisoli (LT46T)* | Motile pleomorphic rods | Medium NHM; T: 35 °C; pH: 7.5; NaCl: 3.1 M. | Tarim Basin, China | Aerobic heterotrophic archaeon; amylolytic; utilizes urea and performs partial denitrification | Zheng et al., 2022 | Zheng, X. W., Wu, Z. P., Sun, Y. P., Wang, B. B., Hou, J., & Cui, H. L. (2022). Halorussus vallis sp. nov., Halorussus aquaticus sp. nov., Halorussus gelatinilyticus sp. nov., Halorussus limi sp. nov., Halorussus salilacus sp. nov., Halorussus salinisoli sp. nov.: six extremely halophilic archaea isolated from solar saltern, salt lake and saline soil. Extremophiles, 26(3). https://doi.org/10.1007/s00792-022-01280-1 |
| *Halorussus salinus (YJ-37-HT)* | Motile pleomorphic rods | Medium NHM; T: 37 °C; pH: 7.0; NaCl: 2.6 M. | Yangjiang Saltworks, China | Aerobic heterotrophic archaeon; amylolytic (hydrolyzes starch); produces indole and $H_2S$; reduces nitrate | Xu et al., 2016 | Xu, J. Q., Xu, W. M., Li, Y., Zhou, Y., Lü, Z. Z., Hou, J., Zhu, L., & Cui, H. L. (2016). Halorussus salinus sp. nov., isolated from a marine solar saltern. Archives of Microbiology, 198(10), 957–961. https://doi.org/10.1007/s00203-016-1253-1 |
| *Halorussus vallis (XZYJT10T)* | Motile pleomorphic rods | Medium NHM; T: 37 °C; pH: 7.5; NaCl: 2.0 M. | Mangkang Saltworks, Tibet | Aerobic heterotrophic archaeon; possesses genes encoding amylases and lipases; reduces nitrate; potential for PHA synthesis | Zheng et al., 2022 | Zheng, X. W., Wu, Z. P., Sun, Y. P., Wang, B. B., Hou, J., & Cui, H. L. (2022). Halorussus vallis sp. nov., Halorussus aquaticus sp. nov., Halorussus gelatinilyticus sp. nov., Halorussus limi sp. nov., Halorussus salilacus sp. nov., Halorussus salinisoli sp. nov.: six extremely halophilic archaea isolated from solar saltern, salt lake and saline soil. Extremophiles, 26(3). https://doi.org/10.1007/s00792-022-01280-1 |
| *Heliorestis acidaminivorans (HR10B)* | Motile straight rods | Medium SHC; T: 30–37 °C; pH: 8.0–9.0. | Lake El Hamra, Egypt | Anaerobic anoxygenic phototroph; photoheterotrophic on amino acids and organic acids; capable of $N_2$ fixation | Asao et al., 2012 | Asao, M., Takaichi, S., & Madigan, M. T. (2012). Amino acid-assimilating phototrophic heliobacteria from soda lake environments: Heliorestis acidaminivorans sp. nov. and "Candidatus Heliomonas lunata." Extremophiles, 16(4), 585–595. https://doi.org/10.1007/s00792-012-0458-8 |
| *Heliorestis baculata (OS-H1)* | Motile straight/curved rods | T: 30 °C; pH: 8.5–9.5; Na2CO3: 0.5–1%. | Lake Ostozhe, Russia | Obligate anaerobe; photoheterotrophic; tolerates and oxidizes sulfide to elemental sulfur and polysulfides | Bryantseva et al., 1999 | Bryantseva, I. A., Gorlenko, V. M., Kompantseva, E. I., Tourova, T. P., Kuznetsov, B. B., & Osipov, G. A. (2000). Alkaliphilic heliobacterium Heliorestis baculata sp. nov. and emended description of the genus Heliorestis. Archives of Microbiology, 174(4), 283–291. https://doi.org/10.1007/s002030000204 |

| Strain | Morphology | Growth characteristics | Isolation_source | Metabolic_role | Reference | Paper |
|---|---|---|---|---|---|---|
| *Natranaerovirga hydrolytica (APP2T)* | Non-motile spore-forming rods | T: 45 °C; pH: 10; Na+: 1.0 M. | Soda lakes, Siberia | Anaerobic fermentative saccharolytic organism; hydrolyzes pectin polymers; utilizes glucuronic acid and fructose | Sorokin et al., 2012 | Sorokin, D. Y., Tourova, T. P., Panteleeva, A. N., Kaparullina, E. N., & Muyzer, G. (2012). Anaerobic utilization of pectinous substrates at extremely haloalkaline conditions by Natranaerovirga pectinivora gen. nov., sp. nov., and Natranaerovirga hydrolytica sp. nov., isolated from hypersaline soda lakes. Extremophiles, 16(2), 307–315. https://doi.org/10.1007/s00792-012-0431-6 |
| *Natranaerovirga pectinivora (AP3T)* | Non-motile spore-forming rods | T: 43 °C; pH: 9.7–10; Na+: 0.4–0.6 M. | Soda soils, Siberia | Anaerobic fermentative saccharolytic organism; highly specialized metabolism for pectin and galacturonic acid | Sorokin et al., 2012 | Sorokin, D. Y., Tourova, T. P., Panteleeva, A. N., Kaparullina, E. N., & Muyzer, G. (2012). Anaerobic utilization of pectinous substrates at extremely haloalkaline conditions by Natranaerovirga pectinivora gen. nov., sp. nov., and Natranaerovirga hydrolytica sp. nov., isolated from hypersaline soda lakes. Extremophiles, 16(2), 307–315. https://doi.org/10.1007/s00792-012-0431-6 |
| *Rubribacterium polymorphum* | Motile oval/pleomorphic cells | T: 30 °C; pH: 8.5–9.5; NaCl: 10 g/l. | Soda lake, Siberia | Photoheterotroph and facultative aerobe; grows well in the dark on organic substrates (sugars, acids) | Boldareva et al., 2009 | Boldareva, E. N., Moskalenko, A. A., Makhneva, Z. K., Tourova, T. P., Kolganova, T. v., & Gorlenko, V. M. (2009). Rubribacterium polymorphum gen. nov., sp. nov., a novel alkaliphilic nonsulfur purple bacterium from an Eastern Siberian soda lake. Microbiology, 78(6), 732–740. https://doi.org/10.1134/S0026261709060101 |
| *Salinarimonas chemoclinalis (ML10T)* | Motile ovoid rods | Medium N2; T: 37 °C; pH: 6.5; NaCl: 6%. | Mahoney Lake, Canada | Aerobic anoxygenic phototroph (AAP); synthesizes bacteriochlorophyll a and carotenoids; accumulates PHAs | Messner et al., 2024 | Messner, K., Kyndt, J. A., & Yurkov, V. (2024). Salinarimonas chemoclinalis, an Aerobic Anoxygenic Phototroph Isolated from a Saline, Sulfate-Rich Meromictic Lake. Microorganisms, 12(11). https://doi.org/10.3390/microorganisms12112359 |
| *Salinarimonas ramus (SL014B-41A4T)* | Motile branched rods | Medium ASW; T: 28 °C; pH: 7.0; NaCl: 4%. | Saline soil, China | Chemoorganotroph and facultative aerobe; utilizes various sugars and organic acids; produces PHB | Cai et al., 2011 | Cai, M., Wang, L., Cai, H., Li, Y., Wang, Y. N., Tang, Y. Q., & Wu, X. L. (2011). Salinarimonas ramus sp. nov. and Tessaracoccus oleiagri sp. nov., isolated from a crude oil-contaminated saline soil. International Journal of Systematic and Evolutionary Microbiology, 61(8), 1767–1775. https://doi.org/10.1099/ijs.0.025932-0 |
| *Salinarimonas rosea (YIM YD3T)* | Motile rods | Mediums MA/MY; T: 28–30 °C; pH: 7.0–8.0. | Salt mine, China | Chemoorganotroph and facultative anaerobe; reduces nitrate; degrades esculin, xanthine, and hypoxanthine | Liu et al., 2010 | Liu, J. H., Wang, Y. X., Zhang, X. X., Wang, Z. G., Chen, Y. G., Wen, M. L., Xu, L. H., Peng, Q., & Cui, X. L. (2010). Salinarimonas rosea gen. nov., sp. nov., a new member of the α-2 subgroup of the proteobacteria. International Journal of Systematic and Evolutionary Microbiology, 60(1), 55–60. https://doi.org/10.1099/ijs.0.006981-0 |
| *Salinarimonas soli (BN140002T)* | Motile rods | Medium R2A; T: 25–30 °C; pH: 8.0; NaCl: 0%. | Soil, South Korea | Aerobic chemoorganotroph; possesses pufLM genes (phototrophic potential) and a type I-C CRISPR-Cas system | Jin et al., 2022 | Jin, C. Z., Jin, L., Liu, M. J., Kang, M. K., Park, S. H., Park, D. J., & Kim, C. J. (2022). Salinarimonas soli sp. nov., isolated from soil. International Journal of Systematic and Evolutionary Microbiology, 72(2). https://doi.org/10.1099/ijsem.0.005095 |
| *Saliphagus infecundisoli (YIM 93745T)* | Non-motile cocci | Medium MG; T: 37 °C; pH: 7.0–7.5; NaCl: 15–20%. | Loulan Ancient City | Aerobic heterotrophic archaeon; utilizes a broad range of carbohydrates; reduces nitrate to nitrite | Yin et al., 2017 | Yin, X. Q., Liu, B. B., Chu, X., Salam, N., Li, X., Yang, Z. W., Zhang, Y., Xiao, M., & Li, W. J. (2017). Saliphagus infecundisoli gen. Nov., sp. nov., an extremely halophilic archaeon isolated from a saline soil. International Journal of Systematic and Evolutionary Microbiology, 67(10), 4154–4160. https://doi.org/10.1099/ijsem.0.002270 |
| *Sporosalibacterium faouarense (SOL3f37T)* | Motile spore-forming rods | T: 40 °C; pH: 6.9; NaCl: 40 g/l. | El Faouar, Tunisia | Anaerobic fermentative chemoorganotroph; degrades carbohydrates to acetate, $H_2$, and $CO_2$; reduces fumarate to succinate | Rezgui et al., 2011 | Rezgui, R., Gam, Z. B. A., Hamed, S. ben, Fardeau, M. L., Cayol, J. L., Maaroufi, A., & Labat, M. (2011). Sporosalibacterium faouarense gen. nov., sp. nov., a moderately halophilic bacterium isolated from oil-contaminated soil. International Journal of Systematic and Evolutionary Microbiology, 61(1), 99–104. https://doi.org/10.1099/ijs.0.017715-0 |
| *Sporosalibacterium tautonense (MRo-4T)* | Motile spore-forming rods | T: 42 °C; pH: 7.4–7.9; NaCl: 20 g/l. | TauTona Mine Gold Mine | Anaerobic fermentative chemoorganotroph; degrades complex polysaccharides such as cellulose and chitin | Podos et al., 2017 | Podosokorskaya, O. A., Merkel, A. Y., van Heerden, E., Cason, E. D., Kopitsyn, D. S., Vasilieva, M., Bonch-Osmolovskaya, E. A., & Kublanov, I. v. (2017). Sporosalibacterium tautonense sp. nov., a thermotolerant, halophilic, hydrolytic bacterium isolated from a gold mine, and emended description of the genus Sporosalibacterium. International Journal of Systematic and Evolutionary Microbiology, 67(5), 1457–1461. https://doi.org/10.1099/ijsem.0.001737 |
| *Sulfurimonas aquatica (H1576T)* | Motile rods | Medium basal; T: 13–15 °C; pH: 6.6–6.8. | Lake Harutori, Japan | Aerobic chemolithoautotroph; oxidizes inorganic sulfur compounds; does not utilize $H_2$ or organic matter | Kojima et al., 2022 | Kojima, H., Kato, Y., Watanabe, T., & Fukui, M. (2022). Sulfurimonas aquatica sp. nov., a sulfur-oxidizing bacterium isolated from water of a brackish lake. Archives of Microbiology, 204(9). https://doi.org/10.1007/s00203-022-03167-3 |
| *Sulfurimonas autotrophica (OK10T)* | Motile short rods | Medium MJ; T: 23–26 °C; pH: 6.0–7.5. | Sediment, Okinawa Trough | Aerobic chemolithoautotroph; oxidizes sulfur, sulfide, and thiosulfate to sulfate; fixes $CO_2$ | Inagaki et al., 2003 | Inagaki, F., Takai, K., Kobayashi, H., Nealson, K. H., & Horikoshi, K. (2003). Sulfurimonas autrotrophica gen. nov., sp. nov., a novel sulfur-oxidizing E-proteobacterium isolated from hydrothermal sediments in the Mid-Okinawa Trough. International Journal of Systematic and Evolutionary Microbiology, 53(6), 1801–1805. https://doi.org/10.1099/ijs.0.02682-0 |

| Strain | Morphology | Growth characteristics | Isolation_source | Metabolic_role | Reference | Paper |
|---|---|---|---|---|---|---|
| *Sulfurimonas crateris (SN118T)* | Motile short rods | Medium anóxico; T: 30 °C; pH: 8.0; NaCl: 2–3%. | Mud volcano, Russia | Facultatively anaerobic chemolithoautotroph; performs complete denitrification (reduces nitrate to $N_2$) | Ratnikova et al., 2020 | Ratnikova, N. M., Slobodkin, A. I., Merkel, A. Y., Kopitsyn, D. S., Kevbrin, V. v., Bonch-Osmolovskaya, E. A., & Slobodkina, G. B. (2020). Sulfurimonas crateris sp. nov., a facultative anaerobic sulfur-oxidizing chemolithoautotrophic bacterium isolated from a terrestrial mud volcano. International Journal of Systematic and Evolutionary Microbiology, 70(1), 487–492. https://doi.org/10.1099/ijsem.0.003779 |
| *Sulfurimonas diazotrophicus (HSL3-1T)* | Non-motile rods | Medium MMJHS; T: 37 °C; pH: 7.0; NaCl: 3.0%. | Fujian mangrove, China | Obligate chemolithoautotroph and diazotroph; fixes $N_2$; utilizes $H_2$ and thiosulfate as energy sources | Zhong et al., 2025 | Zhong, Y., Li, Y., Wang, Z., Cui, L., Lv, S., Zhu, H., Yuan, Q., Lai, Q., Wang, S., & Jiang, L. (2025). Sulfurimonas microaerophilic sp. nov. and Sulfurimonas diazotrophicus sp. nov.: Two Novel Nitrogen-Fixing and Hydrogen- and Sulfur-Oxidizing Chemolithoautotrophs Within the Campylobacteria Isolated from Mangrove Sediments. Microorganisms, 13(4). https://doi.org/10.3390/microorganisms13040713 |
| *Sulfurimonas gotlandica (GD1T)* | Motile curved rods | T: 15 °C; pH: 6.7–8.0; requiere NaCl. | Central Baltic Sea | Facultative chemolithoautotroph; oxidizes $H_2$ and sulfur; uses nitrate/nitrite as electron acceptors; mixotrophic with bicarbonate | Labrenz et al., 2013 | Labrenz, M., Grote, J., Mammitzsch, K., Boschker, H. T. S., Laue, M., Jost, G., Glaubitz, S., & Jurgens, K. (2013). Sulfurimonas gotlandica sp. nov., a chemoautotrophic and psychrotolerant epsilonproteobacterium isolated from a pelagic redoxcline, and an emended description of the genus Sulfurimonas. International Journal of Systematic and Evolutionary Microbiology, 63(PART 11), 4141–4148. https://doi.org/10.1099/ijs.0.048827-0 |
| *Sulfurimonas indica (NW8NT)* | Motile short rods | Medium MMJS; T: 33 °C; pH: 5.5; NaCl: 510 mM. | Hydrothermal vent, Indian Ocean | Obligate chemolithoautotroph; oxidizes $H_2$, thiosulfate, and sulfide; fixes $CO_2$ via the rTCA cycle; uses $O_2$ as the electron acceptor | Hu et al., 2021 | Hu, Q., Wang, S., Lai, Q., Shao, Z., & Jiang, L. (2021). Sulfurimonas indica sp. Nov., a hydrogen-and sulfur-oxidizing chemolithoautotroph isolated from a hydrothermal sulfide chimney in the Northwest Indian Ocean. International Journal of Systematic and Evolutionary Microbiology, 71(1), 1–7. https://doi.org/10.1099/ijsem.0.004575 |
| *Sulfurimonas lithotrophica (GYSZ_1T)* | Motile curved rods | Medium MMJHS; T: 33 °C; pH: 6.5; NaCl: 2%. | Coastal sediment, China | Aerobic/anaerobic chemolithoautotroph; prefers $H_2$ as the optimal energy source; oxidizes thiosulfate and sulfide | Wang et al., 2020 | Wang, S., Jiang, L., Liu, X., Yang, S., & Shao, Z. (2020). Sulfurimonas xiamenensis sp. Nov. and Sulfurimonas lithotrophica sp. nov., hydrogen-and sulfur-oxidizing chemolithoautotrophs within the epsilonproteobacteria isolated from coastal sediments, and an emended description of the genus Sulfurimonas. International Journal of Systematic and Evolutionary Microbiology, 70(4), 2657–2663. https://doi.org/10.1099/ijsem.0.004087 |
| *Sulfurimonas marina (B2T)* | Motile rods | Medium MMJS; T: 35 °C; pH: 7.0; NaCl: 3%. | South China Sea | Obligate aerobic chemolithoautotroph; oxidizes sulfur compounds; possesses genes for $N_2$ fixation | Wang et al., 2022 | Wang, Z., Wang, S., Lai, Q., Wei, S., Jiang, L., & Shao, Z. (2022). Sulfurimonas marina sp. nov., an obligately chemolithoautotrophic, sulphur-oxidizing bacterium isolated from a deep-sea sediment sample from the South China Sea. International Journal of Systematic and Evolutionary Microbiology, 72(10). https://doi.org/10.1099/IJSEM.0.005582 |
| *Sulfurimonas microaerophilic (HSL1-7T)* | Motile straight/curved rods | Medium MMJHS; T: 32 °C; pH: 7.0; NaCl: 3.0%. | Fujian mangrove, China | Microaerophilic chemolithoautotroph; oxidizes $H_2$ and thiosulfate; fixes dinitrogen ($N_2$) | Zhong et al., 2025 | Zhong, Y., Li, Y., Wang, Z., Cui, L., Lv, S., Zhu, H., Yuan, Q., Lai, Q., Wang, S., & Jiang, L. (2025). Sulfurimonas microaerophilic sp. nov. and Sulfurimonas diazotrophicus sp. nov.: Two Novel Nitrogen-Fixing and Hydrogen- and Sulfur-Oxidizing Chemolithoautotrophs Within the Campylobacteria Isolated from Mangrove Sediments. Microorganisms, 13(4). https://doi.org/10.3390/microorganisms13040713 |
| *Sulfurimonas paralvinellae (GO25T)* | Motile short rods | Medium MMJHS; T: 30 °C; pH: 6.1; NaCl: 3%. | Okinawa Trough, Japan | Facultatively anaerobic chemolithoautotroph; uses $H_2$ or sulfur as energy sources; reduces nitrate to nitrite, $N_2O$, and $N_2$ | Takai et al., 2006 | Takai, K., Suzuki, M., Nakagawa, S., Miyazaki, M., Suzuki, Y., Inagaki, F., & Horikoshi, K. (2006). Sulfurimonas paralvinellae sp. nov., a novel mesophilic, hydrogen- and sulfur-oxidizing chemolithoautotroph within the Epsilonproteo-bacteria isolated from a deep-sea hydrothermal vent polychaete nest, reclassification of Thiomicrospira denitrificans as Sulfurimonas denitrificans comb. nov. and emended description of the genus Sulfurimonas. International Journal of Systematic and Evolutionary Microbiology, 56(8), 1725–1733. https://doi.org/10.1099/ijs.0.64255-0 |
| *Sulfurimonas xiamenensis (1-1NT)* | Non-motile rods | Medium MMJHS; T: 30 °C; pH: 7.0; NaCl: 5%. | Coastal sediment, China | Aerobic/anaerobic chemolithoautotroph; oxidizes $H_2$, thiosulfate, and tetrathionate; reduces nitrate and sulfur | Wang et al., 2020 | Wang, S., Jiang, L., Liu, X., Yang, S., & Shao, Z. (2020). Sulfurimonas xiamenensis sp. Nov. and Sulfurimonas lithotrophica sp. nov., hydrogen-and sulfur-oxidizing chemolithoautotrophs within the epsilonproteobacteria isolated from coastal sediments, and an emended description of the genus Sulfurimonas. International Journal of Systematic and Evolutionary Microbiology, 70(4), 2657–2663. https://doi.org/10.1099/ijsem.0.004087 |

| Strain | Morphology | Growth characteristics | Isolation_source | Metabolic_role | Reference | Paper |
|---|---|---|---|---|---|---|
| *Tessaracoccus oleiagri (SL014B-20A1T)* | Non-motile oval/rod-shaped cells | Medium ASW; T: 28 °C; pH: 7.0; NaCl: 4%. | Saline soil, China | Chemoorganotroph and facultative aerobe; performs nitrate reduction; utilizes glucose and inositol | Cai et al., 2011 | Cai, M., Wang, L., Cai, H., Li, Y., Wang, Y. N., Tang, Y. Q., & Wu, X. L. (2011). Salinarimonas ramus sp. nov. and Tessaracoccus oleiagri sp. nov., isolated from a crude oil-contaminated saline soil. International Journal of Systematic and Evolutionary Microbiology, 61(8), 1767–1775. https://doi.org/10.1099/ijs.0.025932-0 |
